# Exploring the phylogeny of rosids with a five-locus supermatrix from GenBank

**DOI:** 10.1101/694950

**Authors:** Miao Sun, Ryan A. Folk, Matthew A. Gitzendanner, Stephen A. Smith, Charlotte Germain-Aubrey, Robert P. Guralnick, Pamela S. Soltis, Douglas E. Soltis, Zhiduan Chen

## Abstract

Current advances in sequencing technology have greatly increased the availability of sequence data from public genetic databases. With data from GenBank, we assemble and phylogenetically investigate a 19,740-taxon, five-locus supermatrix (i.e., *atpB, rbcL, matK, matR*, and ITS) for rosids, a large clade containing over 90,000 species, or approximately a quarter of all angiosperms (assuming an estimate of 400,000 angiosperm species). The topology and divergence times of the five-locus tree generally agree with previous estimates of rosid phylogeny, and we recover greater resolution and support in several areas along the rosid backbone, but with a few significant differences (e.g., the placement of the COM clade, as well as Myrtales, Vitales, and Zygophyllales). Our five-locus phylogeny is the most comprehensive DNA data set yet compiled for the rosid clade. Yet, even with 19,740 species, current sampling represents only 16-22% of all rosids, and we also find evidence of strong phylogenetic bias in the accumulation of GenBank data, highlighting continued challenges for species coverage. These limitations also exist in other major angiosperm clades (e.g., asterids, monocots) as well as other large, understudied branches of the Tree of Life, highlighting the need for broader molecular sampling. Nevertheless, the phylogeny presented here improves upon sampling by more than two-fold and will be an important resource for macroevolutionary studies of this pivotal clade.

## I. Introduction

Given their size, rosids (*Rosidae*; Cantino et al., 2007; Wang et al., 2009; APG IV, 2016) have great potential for understanding the evolution and diversification of angiosperms. A clade of 90,000–120,000 species (estimated from the Open Tree of Life and Open Tree Taxonomy database (OTT); Hinchliff et al., 2015), the rosid clade represents more than a quarter of all angiosperms (based on an estimated 400,000 species of angiosperms; Govaerts, 2001). Rosids comprise two large subclades, fabids (i.e., eurosids I, *Fabidae*) and malvids (i.e., eurosids II, *Malvidae*) and are further divided into 17 orders and 135 families (APG IV, 2016). The clade originated in the Early to Late Cretaceous (115 to 93 Million years ago, Myr), followed by rapid diversification yielding the crown groups of fabids (112 to 91 Myr) and malvids (109 to 83 Myr; Wang et al., 2009; Bell et al., 2010; Magallón et al., 2015). The rosid clade diversified rapidly to form the two major lineages in perhaps as little as 4 to 5 million years (Wang et al., 2009; Bell et al., 2010).

Most rosid families have high species diversity in the tropics, but many extend from the tropics to subtropical and temperate areas (e.g., Wang et al., 2009; Soltis et al., 2010). Most ecologically dominant forest trees are found within the clade, as well as diverse aquatics, parasites, arctic, alpine, and desert lineages; the clade also exhibits tremendous diversity in chemistry, reproductive strategy, and life history (Magallón et al., 1999; Wang et al., 2009; Stevens, 2001 onwards). Unique ecological traits are prevalent in the rosids, including nodular association with nitrogen-fixing bacteria (Soltis et al., 1995; Li et al., 2015), chemical defense mechanisms including glucosinolate production in Brassicales (Rodman et al., 1998; Soltis et al., 2005; Edger et al., 2015), and independent origins of parasitism, sometimes associated with rampant horizontal gene transfer (e.g., *Rafflesia*; Davis & Wurdack, 2004; Xi et al., 2012). Many important crops are rosids, including cotton and cacao (Malvaceae), hops (Cannabaceae), legumes (Fabaceae), rubber (Euphorbiaceae), and numerous vegetable and fruit crops (Brassicaceae, Caricaceae, Cucurbitaceae, Rosaceae, Rutaceae, and Vitaceae). Some rosids have been selected as genetic models, including *Arabidopsis thaliana* (*Arabidopsis* Genome Initiative, 2000), *Brassica rapa* (Wang et al., 2011), and various legumes (Sato et al., 2008; Schmutz et al., 2010, 2014; Varshney et al., 2012, 2013; Young et al., 2011).

The “rise of the rosids” yielded most angiosperm-dominated forests present today. Many other lineages of life radiated in the shadow of these rosid-dominated forests (e.g., ants: Moreau et al., 2006; Moreau & Bell, 2013; beetles: Farrell, 1998; Wilf et al., 2000; amphibians: Roelants et al., 2007; mammals: Bininda-Emonds et al., 2007; fungi: Hibbett & Matheny, 2009; liverworts: Feldberg et al., 2014; ferns: Schneider et al., 2004; Watkins & Cardelús, 2012). The initial rise of the rosids and subsequent repeated cycles of radiations within the rosid clade (Soltis & Soltis, 2004; Soltis et al., 2004) have profoundly shaped much of current terrestrial biodiversity (Wang et al., 2009; Boyce et al., 2010).

Although rosids have long been the focus of phylogenetic research (e.g., Wang et al., 2009; Soltis et al., 2011; Zeng et al., 2017; and references therein), the enormous size of this clade has thus far precluded achieving the sampling required for macroevolutionary inferences (Ricklefs, 2007; Smith et al., 2011; Thomas et al., 2013; Folk et al., 2018). Hence, a robust, time-calibrated phylogeny with large-scale species-level sampling is needed for future diversity studies. Additionally, the rosid clade also provides an opportunity to evaluate the implications of taxon and gene sampling. Using data from GenBank, we constructed a 5-locus phylogenetic tree having more than twice the taxon sampling used in earlier studies (e.g., the 4-locus study of Sun et al., 2016)—and compared this tree to all rosid names in the OpenTree of Life (Hinchliff et al., 2015) to quantify bias in DNA sampling across the clade. We hypothesized that (1) taxon sampling remains highly biased across the large rosid clade, and that (2) the use of more genes and increased taxon sampling impacts phylogenetic resolution and divergence time estimation. We tested these hypotheses via a series of comparisons across two trees: a previously published 4-locus, 8,855-taxon supermatrix (Sun et al, 2016) and the more densely sampled 5-locus, 19,740-taxon supermatrix generated here. We then quantified patterns of taxon sampling bias, phylogenetic resolution, and time calibration.

## II. Materials and Methods

### Data mining, alignment, and phylogeny reconstruction

We mined GenBank (Release 214: June 15, 2016) for the chloroplast genes *atpB, matK*, and *rbcL*, the mitochondrial gene *matR*, and the nuclear ribosomal ITS (including ITS-1, 5.8S, and ITS-2 regions) using the PHyLogeny Assembly With Databases pipeline (PHLAWD, version 3.4a, https://github.com/blackrim/phlawd; Smith et al., 2009). These genes represent those most commonly used in phylogenetic studies of plants and therefore the most numerous loci for plants deposited in GenBank; they also represent all three plant genomes.

We employed PHLAWD data mining using three bait sequences of each target locus that represent the phylogenetic diversity of the rosid clade (Hinchliff & Smith, 2014). The quality of sequence data from the five sampled loci was investigated by calculating the best-hit scores from BLAST and plotting the distribution of identity scores ([0, 1]) against coverage scores ([0, 1]). Low-quality and outlier sequences were removed based on these scores (see Results). For all resulting alignments, we (1) validated the species names following The Plant List (TPL; http://www.theplantlist.org/), using the R package Taxonstand v2.0 (Cayuela et al., 2012), and then (2) pruned all taxa with “subsp.”, “var.”, “f.”, “cf.” and “aff.” designations in taxon names. Names for orders and families follow APG IV (2016), and those for major supra-ordinal clades follow Soltis et al. (2011) and Cantino et al. (2007).

We curated the 5-locus data set iteratively by screening individual loci and concatenated matrices (below) for rogue taxon behavior through manual inspection of initial phylogenies for spurious taxon placement and through the RAxML dropsets algorithm (Pattengale et al., 2010a; Sun et al., 2016; Smith & Brown, 2018). The final 5-locus data set contained 19,740 ingroup species (135 families and 17 orders) and 20,294 species, including outgroup taxa (i.e., 554 outgroup species in 17 families and three orders) from Saxifragales, Proteales, and Trochodendrales.

We also compared results from our 5-locus (*atpB, rbcL, matK, matR*, and ITS) data set with those from a previously published 4-locus rosid data set comprising chloroplast and mitochondrial loci (Sun et al., 2016; *atpB, rbcL, matK*, and *matR*). The 4-locus data set contained 8,855 ingroup taxa (9,300 taxa including outgroups; i.e., 445 outgroup species from the same three orders; Sun et al., 2016). Hereafter, all tree sampling statistics will only concern the ingroup.

### Taxon sampling analyses

To evaluate sampling gaps among all taxonomically recorded rosid species and the species included in the 5-locus data set, we mapped the 19,740 validated rosid names in our phylogeny against all rosid species present in OTT v3.0 (https://devtree.opentreeoflife.org/about/taxonomy-version/ott3.0; Hinchliff et al., 2015). Generic names of this complete list were also manually curated via an online tool, Index Nominum Genericorum (Farr & Zijlstra, 1996 onwards), and then invalid, rejected (nom. rej.), illegitimate, and synonymous generic names were all removed, as well as any taxon names with “sp.”, “subsp.”, “var.”, “x”, “cf.” and “aff.”, or other non-species designations (e.g., “spp.”, “clone”, “environmental sample”, “group”).

Additionally, to evaluate whether sampling in rosid DNA data from GenBank is phylogenetically biased, we scored DNA data presence and absence by mapping our phylogeny names to OpenTree (Smith & Brown, 2018) and then executing a *λ* test on this “trait” under an equal rates model (R packge geiger V.2.0.6.2; Pennell et al., 2014). Significance was assessed with a likelihood ratio test.

### Phylogenetics

The edited and pruned alignments of each locus were concatenated into a single supermatrix using FASconCAT v.1.0 (Kück & Meusemann, 2010). We ran maximum likelihood (ML) analyses for each individual locus alignment and for the concatenated matrices using RAxML v.8.2.10 (Stamatakis, 2014) with 100 bootstrap (BS) replicates for topology examination, under an unpartitioned GTRCAT model. For the 5-locus concatenated matrix, the best ML tree was constructed with RAxML using the extended Majority Rule Criterion (autoMRE) as a bootstrap stopping rule (Pattengale et al., 2010b; reached at 352 replicates). We visually examined potential topological conflicts by concatenating different data sets and evaluating strongly supported differences among trees at the family level inferred from the combined supermatrix and of each individual data set (i.e., ITS, *matR*, and the plastid genes; see Results and Discussion). Trees were manipulated for display using Newick utilities (Junier & Zdobnov, 2010), Dendroscope 3 (Huson & Scornavacca, 2012), MEGA (Tamura et al., 2013), and iTOL v3.0 (Letunic & Bork, 2016).

### Divergence Time Estimation

Divergence time estimation was conducted using both the previously published 4-locus (Sun et al., 2016) phylogeny and newly constructed 5-locus phylogeny. In total, 59 calibration points (covering 15/17 rosid orders) were used as time constraints, based on validated fossils frequently used as calibration points in previous molecular dating studies (Davis et al., 2005; Wang et al., 2009; Bell et al., 2010; Sauquet et al., 2012; Magallón et al., 2015; Table S1). The root was constrained to a maximum age of 125 Myr following Wang et al. (2009). For both phylogenies, we used the penalized likelihood program treePL v.1.0 (Smith & O’Meara, 2012) to generate a time-calibrated ultrametric tree. We initially conducted random cross-validation procedures with three options (“randomcv”, “thorough”, and “prime”) to determine the best smoothing value and optimization parameters for both the 4-locus and 5-locus ML trees and then ran 200,000 annealing iterations (default = 5,000) for divergence time estimation.

To estimate variation in the timing of the rosid divergence, we also employed PATHd8 v.1.0 (Britton et al., 2006) using the same 59 calibration points and root constraint as above. Unlike treePL, PATHd8 is a faster heuristic method that sequentially takes averages over path lengths from an internode to all its descending terminals, one pair of sister groups at a time (Ericson et al., 2006; Anderson, 2007).

## III. Results

### Limitations in Taxon and Locus Sampling

For the commonly sequenced locus ITS, PHLAWD initially recovered 42,890 rosid sequences; after removing sequences with non-species designations (cf. Materials and Methods), 39,735 sequences remained. Removal of sequences with low identity and coverage scores (coverage score ≤ 0.1 and identity score ≤ 0.1 were considered low quality) and further duplicate removal and non-ortholog cleaning, 15,100 sequences remained in the single locus matrix; however, only 13,157 sequences were retained in the final combined supermatrix for phylogeny reconstruction, due to two reasons: 1) some ITS sequences still exhibiting characteristics of rogue taxa identified by initial RAxML analyses in primary 5-locus supermatrix (Pattengale et al., 2010; Sun et al., 2016); and 2) in this combined matrix, some species have only one fragment of either ITS1, 5.8S, or ITS2, and the other four genes are not available; therefore, these short single ITS fragments were removed from the combined matrix to avoid introducing large amounts of missing data. Updating the remaining loci (*atpB, rbcL, matK*, and *matR*) with new GenBank data resulted in 1,257, 6,960, 8,489, and 721 sequences, respectively. The alignment lengths for *atpB, rbcL, matK, matR*, and ITS were, respectively, 1,500, 1,401, 1,815, 2,349, and 835 bp, with a concatenated length of 7,900 bp, and 70.55% missing data.

To better understand sampling patterns in our data sets, we matched our recovered phylogenetic tips (species) with those in the Open Tree of Life taxonomic database (OTT v3.0; Hinchliff et al., 2015), which includes rosid clades—e.g., Rutaceae, Francoaceae, and Kirkiaceae—yet to be integrated into the Open Tree topology itself. We sampled 135 families (100% coverage of the rosid families recognized in APG IV, 2016), 3,070 genera (matching 66.34% of OTT), and 19,740 species (matching 16.25% of OTT). The unsampled genera and species mainly reflect absence of DNA data (Fig. 1 and Tables 1, S2), but some are due to taxonomic issues such as synonyms and invalid names. Among these mismatches are names in our 5-gene tree unaccounted for in OTT, comprising 1,134 species (5.74%) and 72 genera (2.35%). Total coverage and taxon representation compared with the rosids present in OTL and OTT are summarized in Table 1, and Table S2, respectively (order and family circumscription adjusted to comply with APG IV [2016]; marked with asterisks in Table S2).

**Fig. 1.**
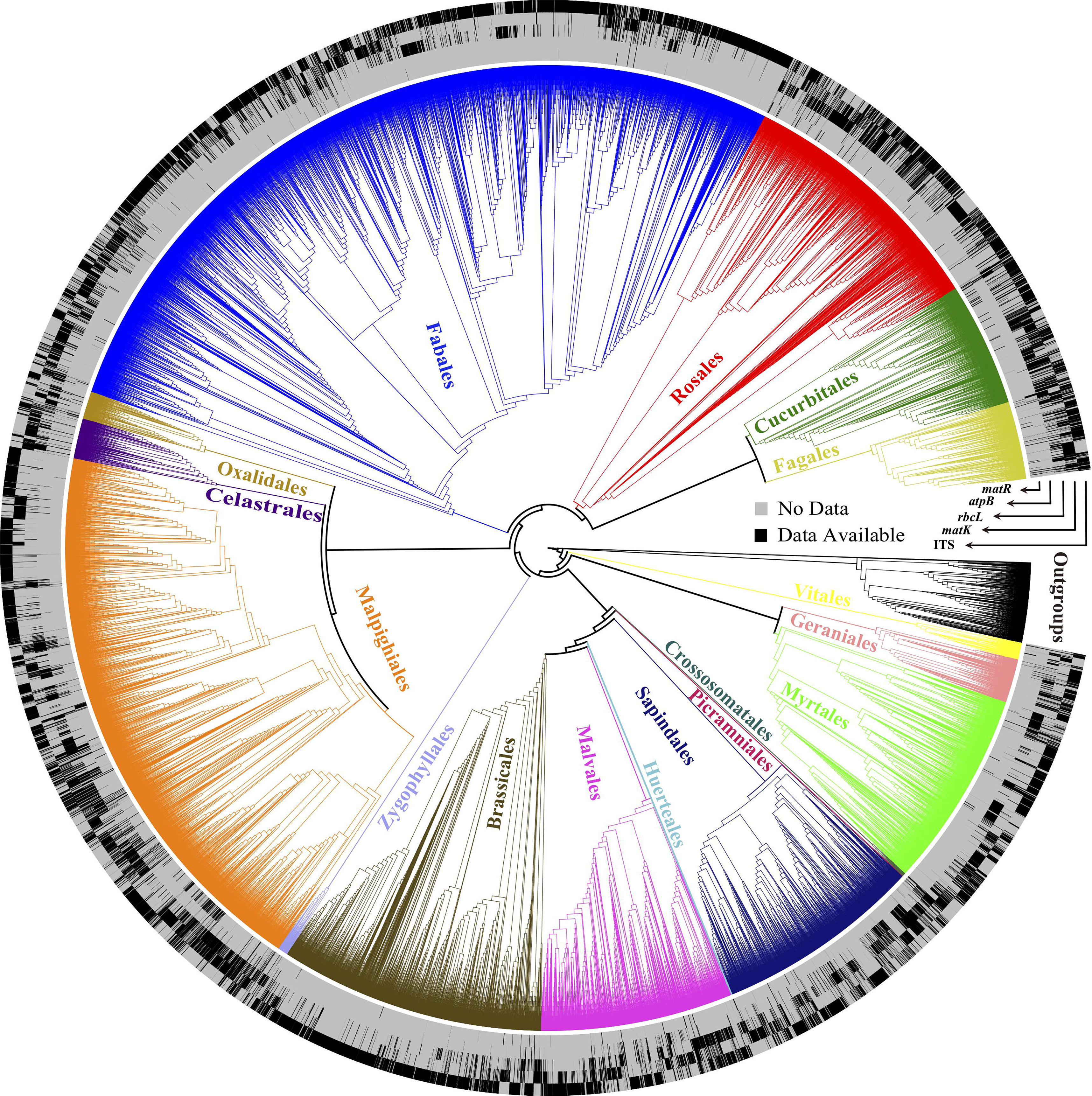
The 5-locus rosid phylogeny showing sampling coverage of sequence data for the 5 loci. This comparison shows substantial phylogenetic bias in each of the 5 loci sampled in the rosid matrix based on presence/absence heatmap layers. The five layers are labeled as *matR, atpB, rbcL, matK*, and ITS from inside toward the outer edge. Species with black tips at each layer mean sequence data are available for a specific locus. Each order is labeled and colored, so phylogenetic bias of DNA data can be viewed by the rough distribution of the black and gray tips within and/or among each order (cf. Tables 1, S2 for further percentage details).

**Table 1.**
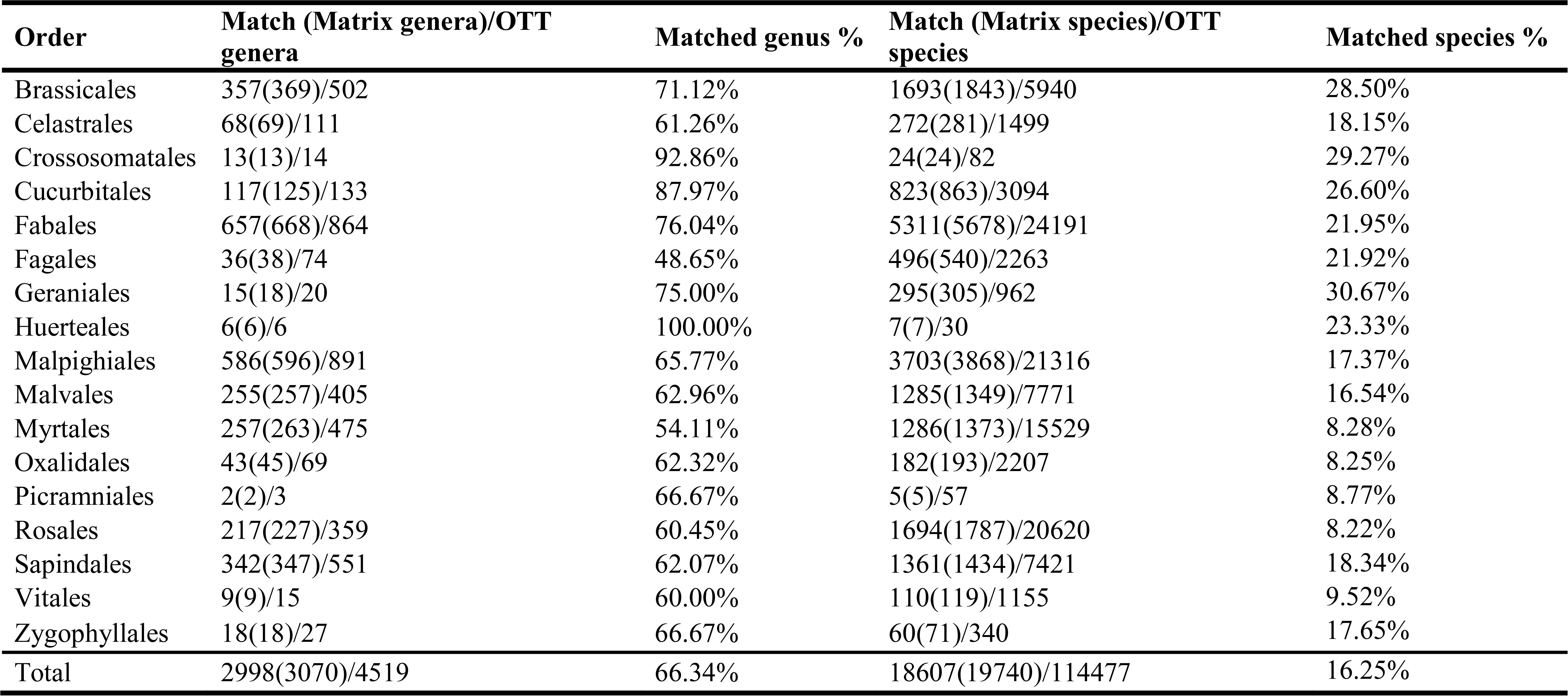
Ordinal-level summary sampling table for the 5-locus rosid supermatrix (“Matrix”) compared to the rosid clade of the Open Tree Taxonomy (“OTT”) database v. 3.0 (https://devtree.opentreeoflife.org/about/taxonomy-version/ott3.0; Hinchliff et al., 2015) and matching taxon names between these data sets. Orders follow APG IV (2016). A summary table at the family level is available in Table S2.

Sampling coverage in our tree shows a strong phylogenetic bias (*p*-value for λ test, *p* ≈ 0; Fig. 1 and Tables 1, S2). Overall, larger orders (> 10,000 species, e.g., Rosales and Myrtales) tend to be more poorly sampled with > 90% of the species unsampled (Table 1). Several large families (> 1,000 species) also have poor coverage; Polygalaceae, Rosaceae, Myrtaceae, Malvaceae, Rutaceae, and Phyllanthaceae have 3.14% to 15.27% sampling (Table S2). Geraniales, Crossosomatales, Brassicales, Cucurbitales, and Huerteales have better sampling, yet no order or family exceeds 50% coverage of known species richness.

### Phylogenetic Analyses

The topology of our 5-locus rosid tree (Fig. S1) generally agrees with that of the 4-locus tree inferred in a previous GenBank mining effort (Sun et al., 2016), but provides greater resolution and support in several areas along the backbone (Fig. S1) as well as greatly improved species-level sampling. The median BS value of the 5-locus rosid tree is lower than that obtained in the 4-locus, 8,855-taxon rosid phylogeny (Fig. 2a), but within methodological expectations (see Discussion).

**Fig. 2.**
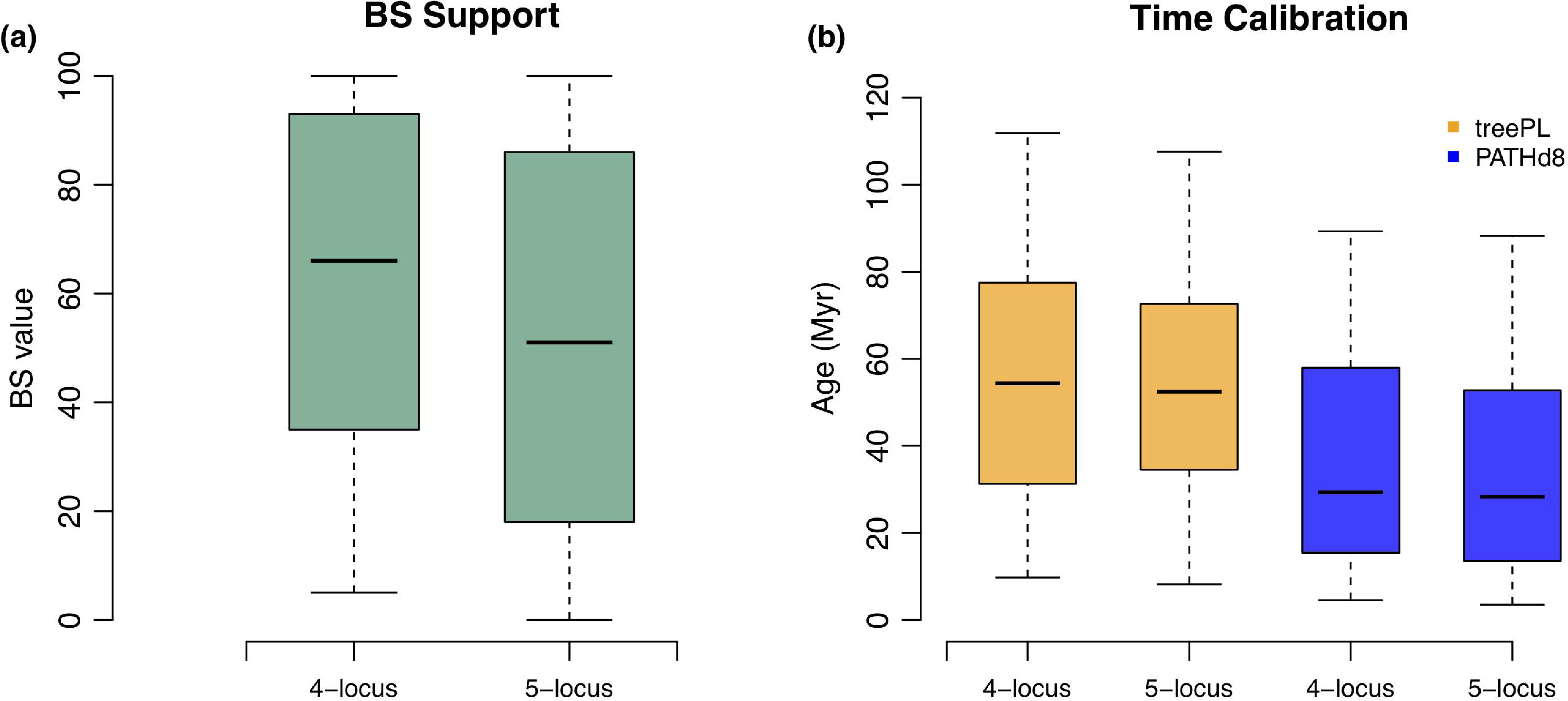
Comparison of phylogenetic resolution (a) and divergence time estimation (b) between the 4-locus, 8,855-taxon rosid phylogeny and the 5-locus, 19,740-taxon rosid phylogeny. BS stands for bootstrap. Orange denotes the treePL method, and blue denotes the PATHd8 method in panel (b).

The topologies inferred from single locus partitions and the concatenated data set are generally consistent, with the exception of the following conflicting phylogenetic placements:

(1) For the COM clade (Celastrales-Oxalidales-Malpighiales), we observed the same conflicting placements when trees from nuclear, plastid, and mitochondrial data are compared as observed in Sun et al. (2015, 2016). Nuclear and mtDNA data favor a placement of the COM clade with malvids, whereas plastid data indicate a placement with fabids. (2) The placement of Myrtales and Geraniales was unstable across locus partitions. In the chloroplast tree, Myrtales and Geraniales were sequential sisters to fabids with strong support, but in the *matR* tree and total evidence tree, Myrtales and Geraniales were sisters to the rest of the rosids (cf. Sun et al., 2015, 2016). The monophyly of Myrtales was not supported in the ITS tree. (3) The placement of Zygophyllales varied among locus partitions. In the *matR* tree, Zygophyllales were resolved as sister to malvids (cf. Sun et al., 2016; Zhao et al., 2016), a different result from the chloroplast and total evidence trees, where the clade was placed within fabids with low to moderate support. (4) The placement of Vitales with respect to the rest of the rosids and Saxifragales was unstable across analyses. In the *matR* and total evidence trees, the three groups were resolved as (rosids + Vitales) + Saxifragales (cf. Sun et al., 2016), a topology that has also been recovered in most studies (Soltis et al., 2007, 2011; Worberg et al., 2007; Zhu et al., 2007; APG, 2009, 2016; Wang et al., 2009; Smith et al., 2010; Barniske et al., 2012; Ruhfel et al., 2014). Our chloroplast data set, by contrast, recovers the relationship among these three clades as rosids + (Saxifragales + Vitales), a result also seen previously (cf. Moore et al., 2010; Ruhfel et al., 2014; Zhang et al., 2012, 2016; Sun et al., 2016). (5) Non-monophyly of some families was occasionally seen within single-gene trees. Two families (Cannabaceae and Euphorbiaceae) are resolved as non-monophyletic in the combined plastid tree, while they are recovered as monophyletic in the total evidence tree as expected; similarly, 12 families are non-monophyletic in the *matR* gene tree and 18 in the ITS tree (see Table S3).

### Dating Analyses

The crown age of rosids is estimated as 117.93 Myr by treePL (89.80 Myr by PATHd8) using the 5-locus tree, and 122.62 Myr by treePL (104.20 Myr by PATHd8) for the 4-locus tree. The ages of other major rosid clades are reported in Table 2. Comparisons of crown ages obtained for all major rosid clades (17 orders and 135 families, *sensu* APG IV) are provided in Figs. 2b and 3, covering both treePL and PATHd8 and both 4-locus and 5-locus trees, as well as previous studies (Wikström et al., 2001; Wang et al., 2009; Bell et al., 2010; Zanne et al., 2014; Magallón et al., 2015; Zeng et al., 2017). The ages of major rosid lineages estimated using the two dating methods and the two trees largely overlap with uncertainty intervals reported in previous studies (Fig 3). However, we did find that some clade ages (e.g., Celastrales, Crossosomatales, and Picramniales; Fig. 3) estimated in our study are younger and outside of the range of uncertainty reported in previous studies (Wikström et al., 2001; Wang et al., 2009; Bell et al., 2010; Zanne et al., 2014; Magallón et al., 2015), even though their placements agree with earlier studies (e.g., Wang et al., 2009; Magallón et al., 2015; APG IV, 2016). This discrepancy is likely due to poor taxon sampling of smaller rosid orders in previous studies; particularly, inclusion of a single species (e.g., Picramniales in the dating analyses of Zanne et al. 2014; Magallón et al. 2015) only allows stem age estimation. The greatly increased rosid sampling in the present study compared to earlier investigations (e.g., Wang et al., 2009; Bell et al., 2010; Magallón et al., 2015) could impact age estimation. Hence, we favor the results from treePL (both 4- and 5-locus; Table 2, Fig 2b). Although differing in overall scaling, estimated divergence times for all nodes estimated from treePL and PATHd8 were highly correlated (R^2^ = 0.762).

**Table 2.** Summary table of ages estimated for rosid major clades by treePL and PATHd8 based on the tree inferred here (“5-locus”) and that inferred in Sun et al. 2016 (“4-locus”). The age unit is million years ago (Myr).

| Clade | 5-locus |  | 4-locus |  |
| --- | --- | --- | --- | --- |
|  | Age treePL | Age PATHd8 | Age treePL | Age PATHd8 |
| **Fabids** | 117.03 | 89.80 | 121.11 | 94.95 |
| **Fagales** | 95.60 | 83.50 | 100.83 | 83.50 |
| Betulaceae | 60.64 | 59.80 | 59.80 | 59.80 |
| Casuarinaceae | 27.57 | 46.63 | 27.06 | 41.62 |
| Fagaceae | 66.27 | 61.12 | 67.04 | 69.46 |
| Juglandaceae | 72.55 | 69.39 | 69.66 | 66.60 |
| Myricaceae | 30.08 | 27.63 | 26.18 | 27.97 |
| Nothofagaceae | 18.53 | 11.42 | 18.26 | 11.30 |
| Ticodendraceae | 66.67 | 59.80 | 64.52 | 63.23 |
| **Cucurbitales** | 111.95 | 65.06 | 113.39 | 53.48 |
| Anisophylleaceae | 48.13 | 35.30 | 49.44 | 39.52 |
| Apodanthaceae | 70.73 | 65.06 | 74.69 | 48.60 |
| Begoniaceae | 52.11 | 35.40 | 23.74 | 20.39 |
| Cucurbitaceae | 34.91 | 28.17 | 42.07 | 29.20 |
| Tetramelaceae | 19.21 | 9.01 | 22.57 | 12.15 |
| Corynocarpaceae | 6.95 | 3.62 | 6.96 | 2.86 |
| Coriariaceae | 14.04 | 7.82 | 22.97 | 10.15 |
| Datisceae | 8.47 | 4.54 | 9.54 | 5.24 |
| **Rosales** | 107.52 | 89.80 | 111.03 | 89.80 |
| Cannabaceae | 67.73 | 65.50 | 67.26 | 65.50 |
| Elaeagnaceae | 27.41 | 20.06 | 28.70 | 27.79 |
| Moraceae | 46.03 | 28.78 | 40.49 | 16.31 |
| Rhamnaceae | 74.55 | 48.60 | 80.11 | 61.70 |
| Rosaceae | 89.80 | 89.80 | 89.80 | 89.80 |
| Ulmaceae | 59.41 | 32.53 | 88.93 | 21.49 |
| Urticaceae | 60.28 | 51.05 | 68.70 | 33.25 |
| Dirachmaceae | 88.51 | 70.60 | 96.98 | 70.60 |
| Barbeyaceae | 84.47 | 53.33 | 90.03 | 70.60 |
| **Fabales** | 96.58 | 59.90 | 105.41 | 75.15 |
| Fabaceae | 83.90 | 55.80 | 93.77 | 65.37 |
| Polygalaceae | 60.89 | 55.80 | 72.51 | 13.46 |
| Surianaceae | 47.39 | 15.50 | 53.52 | 21.96 |
| Quillajaceae | 90.75 | 26.59 | 98.52 | 36.49 |
| **Malpighiales** | 114.67 | 89.30 | 114.47 | 89.30 |
| Achariaceae | 54.72 | 14.37 | 64.21 | 25.88 |
| Bonnetiaceae | 70.08 | 58.52 | 71.27 | 58.67 |
| Calophyllaceae | 55.73 | 21.85 | 47.53 | 17.35 |
| Caryocaraceae | 20.12 | 5.20 | 22.18 | 8.73 |
| Centropalacaceae | 74.12 | 15.13 | 77.14 | 23.34 |
| Chrysobalanaceae | 22.13 | 10.69 | 27.61 | 16.17 |
| Clusiaceae | 52.63 | 27.13 | 52.05 | 22.91 |
| Dichapetalaceae | 17.65 | 5.96 | 22.97 | 10.87 |
| Elatinaceae | 50.37 | 17.24 | 51.62 | 21.99 |
| Erythroxylaceae | 46.31 | 17.55 | 55.76 | 26.26 |
| Euphorbiaceae | 63.30 | 38.14 | 99.38 | 41.95 |
| Humiriaceae | 33.90 | 33.90 | 33.90 | 33.90 |
| Hypericaceae | 81.66 | 63.84 | 81.46 | 63.49 |
| Irvingiaceae | 5.19 | 1.57 | 6.10 | 2.61 |
| Ixonanthaceae | 23.76 | 8.15 | 31.28 | 13.56 |
| Lacistemataceae | 7.82 | 2.19 | 9.54 | 3.73 |
| Linaceae | 58.84 | 25.10 | 60.48 | 37.36 |
| Malpighiaceae | 53.17 | 33.00 | 47.75 | 33.00 |
| Ochnaceae | 51.42 | 15.36 | 54.39 | 25.49 |
| Pandaceae | 38.67 | 8.97 | 42.54 | 15.19 |
| Passifloraceae | 63.15 | 33.74 | 83.25 | 3.24 |
| Peraceae | 53.84 | 13.68 | 68.14 | 22.94 |
| Phyllanthaceae | 68.83 | 28.13 | 101.43 | 50.67 |
| Picrodendraceae | 35.15 | 12.86 | 44.13 | 21.99 |
| Podostemaceae | 80.18 | 89.30 | 82.99 | 89.30 |
| Putranjivaceae | 18.11 | 6.86 | 16.72 | 6.28 |
| Rafflesiaceae | 81.48 | 41.45 | 70.24 | 41.13 |
| Rhizophoraceae | 59.88 | 33.90 | 59.74 | 33.90 |
| Salicaceae | 64.37 | 48.00 | 71.33 | 48.00 |
| Trigoniaceae | 38.07 | 15.32 | 50.40 | 26.28 |
| Violaceae | 57.17 | 27.34 | 68.43 | 47.89 |
| Balanopaceae | 17.87 | 3.61 | 24.51 | 6.52 |
| Ctenolophonaceae | 80.30 | 34.51 | 109.65 | 52.48 |
| Euphroniaceae | 46.86 | 23.35 | 64.26 | 37.35 |
| Goupiaceae | 75.00 | 32.02 | 87.00 | 54.34 |
| Lophopyxidaceae | 60.81 | 19.39 | 77.63 | 31.51 |
| **Celastrales** | 89.44 | 45.93 | 94.99 | 54.88 |
| Lepidobotryaceae | 12.11 | 3.25 | 12.84 | 4.99 |
| Celastraceae | 67.89 | 37.82 | 76.39 | 38.90 |
| **Oxalidales** | 112.85 | 79.20 | 114.28 | 79.20 |
| Connaraceae | 35.38 | 6.59 | 24.76 | 10.10 |
| Cunoniaceae | 43.50 | 79.20 | 32.67 | 65.56 |
| Elaeocarpaceae | 61.70 | 61.70 | 61.70 | 61.70 |
| Huaceae | 14.23 | 4.30 | 16.56 | 6.62 |
| Oxalidaceae | 52.72 | 21.00 | 46.59 | 30.05 |
| Cephalotaceae | 77.57 | 53.13 | 70.61 | 79.20 |
| Brunelliaceae | 2.65 | 3.41 | 3.71 | 3.27 |
| **Zygophyllales** | **96.67** | **42.02** | **96.95** | **74.54** |
| Zygophyllaceae | 91.38 | 41.07 | 90.94 | 69.52 |
| Krameriaceae | 31.36 | 8.25 | 18.16 | 7.87 |
| **Malvids** | **116.49** | **89.30** | **120.33** | **91.47** |
| **Brassicales** | **86.82** | **86.66** | **95.92** | **89.30** |
| Akaniaceae | 4.10 | 1.86 | 3.07 | 1.19 |
| Brassicaceae | 36.20 | 30.70 | 45.27 | 32.55 |
| Capparaceae | 38.94 | 17.68 | 43.31 | 22.21 |
| Caricaceae | 26.58 | 13.58 | 28.74 | 16.33 |
| Cleomaceae | 38.58 | 24.01 | 47.85 | 29.53 |
| Gyrostemonaceae | 6.80 | 3.34 | 9.87 | 5.39 |
| Limnanthaceae | 13.78 | 8.26 | 11.54 | 7.52 |
| Resedaceae | 53.78 | 44.91 | 58.24 | 43.56 |
| Salvadoraceae | 26.97 | 14.59 | 40.37 | 24.73 |
| Tropaeolaceae | 40.20 | 20.73 | 13.06 | 7.64 |
| Bataceae | 32.77 | 20.86 | 36.62 | 24.73 |
| Emblingiaceae | 52.88 | 24.59 | 61.28 | 25.27 |
| Koeberliniaceae | 67.42 | 50.48 | 77.00 | 55.39 |
| Moringaceae | 12.17 | 5.91 | 14.61 | 6.99 |
| Pentadiplandraceae | 63.01 | 62.42 | 70.52 | 70.43 |
| Setchellanthaceae | 78.46 | 79.01 | 87.32 | 89.30 |
| Tovariaceae | 52.88 | 24.59 | 61.28 | 25.27 |
| **Malvales** | **100.59** | **69.12** | **101.97** | **68.91** |
| Bixaceae | 50.29 | 24.36 | 54.37 | 26.98 |
| Cistaceae | 45.68 | 34.47 | 50.40 | 45.35 |
| Cytinaceae | 73.43 | 67.35 | 73.58 | 55.17 |
| Dipterocarpaceae | 35.80 | 17.07 | 36.05 | 19.22 |
| Malvaceae | 50.85 | 36.94 | 44.04 | 25.61 |
| Muntingiaceae | 42.72 | 21.12 | 49.82 | 25.44 |
| Neuradaceae | 10.59 | 5.59 | 11.18 | 5.71 |
| Sarcolaenaceae | 12.94 | 6.04 | 13.37 | 6.31 |
| Sphaerosepalaceae | 11.85 | 4.89 | 13.20 | 5.20 |
| Thymelaeaceae | 48.55 | 40.20 | 47.79 | 12.59 |
| **Picramniales/Picramniaceae** | 36.78 | 9.65 | 36.91 | 14.80 |
| **Huerteales** | 61.57 | 37.20 | 67.93 | 37.20 |
| Dipentodontaceae | 38.06 | 10.16 | 37.79 | 15.23 |
| Tapisciaceae | 37.20 | 37.20 | 37.20 | 37.20 |
| Petenaaceae | 56.88 | 14.16 | 61.53 | 22.34 |
| Gerrardinaceae | 56.88 | 14.16 | 61.53 | 22.34 |
| **Sapindales** | 88.96 | 65.50 | 82.21 | 65.50 |
| Anacardiaceae | 45.59 | 32.35 | 50.69 | 47.00 |
| Burseraceae | 43.87 | 31.43 | 31.32 | 32.09 |
| Meliaceae | 48.60 | 48.60 | 49.04 | 48.60 |
| Nitrariaceae | 62.91 | 17.63 | 32.49 | 16.91 |
| Rutaceae | 65.50 | 65.50 | 65.50 | 65.50 |
| Sapindaceae | 64.76 | 55.80 | 60.52 | 55.80 |
| Simaroubaceae | 46.19 | 12.04 | 47.98 | 21.83 |
| Kirkiaceae | 3.46 | 0.92 | 3.97 | 1.49 |
| Biebersteiniaceae | 19.11 | 5.12 | 17.58 | 7.59 |
| **Crossosomatales** | 88.66 | 28.40 | 90.75 | 34.68 |
| Crossosomataceae | 17.62 | 4.88 | 16.78 | 8.65 |
| Staphyleaceae | 28.40 | 28.40 | 28.40 | 28.40 |
| Strasburgeriaceae | 20.74 | 4.68 | 23.87 | 8.25 |
| Aphloiaceae | 75.92 | 15.79 | 78.94 | 27.26 |
| Geissolomataceae | 46.62 | 9.63 | 56.37 | 17.05 |
| Guamatelaceae | 40.90 | 6.30 | 31.80 | 10.86 |
| Stachyuraceae | 2.84 | 0.63 | 3.67 | 1.01 |
| **Geraniales + Myrtales** | **116.49** | **88.20** | **121.39** | **96.30** |
| **Geraniales** | **107.69** | **47.69** | **117.77** | **81.03** |
| Geraniaceae | 74.66 | 32.08 | 97.14 | 77.94 |
| Francoaceae | 97.34 | 29.59 | 104.62 | 36.60 |
| **Myrtales** | **95.00** | **88.20** | **95.00** | **88.20** |
| Combretaceae | 51.58 | 16.24 | 42.78 | 12.45 |
| Crypteroniaceae | 18.04 | 9.91 | 17.77 | 9.68 |
| Lythraceae | 72.66 | 70.60 | 71.98 | 70.60 |
| Melastomataceae | 49.25 | 51.74 | 47.35 | 60.23 |
| Myrtaceae | 64.10 | 55.80 | 64.27 | 55.80 |
| Onagraceae | 49.56 | 40.09 | 60.65 | 53.07 |
| Penaeaceae | 34.38 | 17.22 | 26.29 | 21.19 |
| Vochysiaceae | 39.18 | 26.95 | 36.65 | 27.24 |
| Alzateaceae | 52.22 | 26.13 | 32.38 | 26.29 |
| **Vitales/Vitaceae** | **75.77** | **66.00** | **78.27** | **66.37** |
| **Rosids** | **117.93** | **89.80** | **122.62** | **104.20** |

**Fig. 3.**
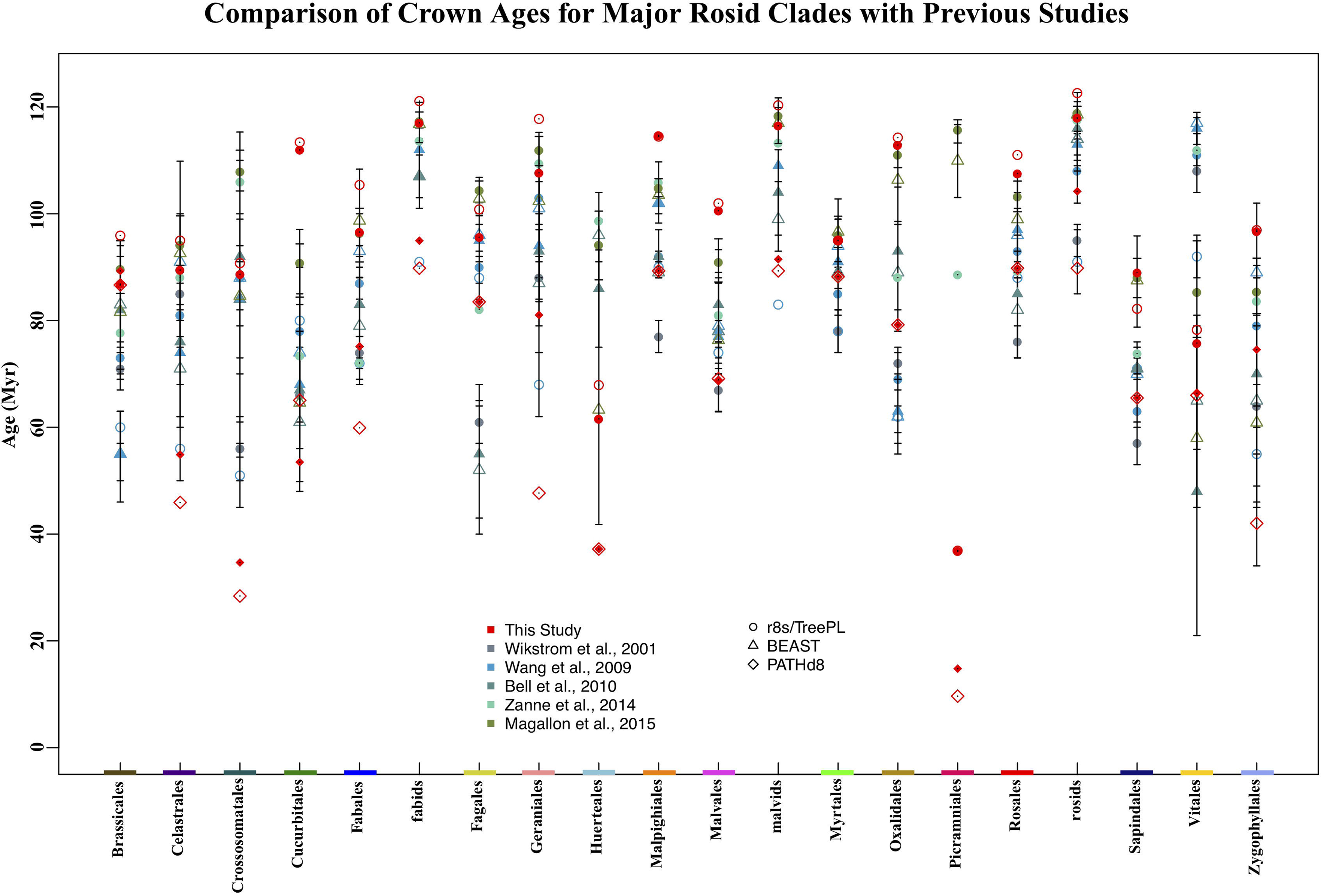
Comparison of crown ages for major rosid clades reported in this study, Wikström et al. (2001), Wang et al. (2009), Bell et al. (2010), Zanne et al. (2014), and Magallón et al. (2015). Error bars represent age ranges reported for the given node. Red hollow circles and diamonds stand for ages estimated from the 4-locus tree; solid symbols are estimated from the 5-locus tree; the color bars at the bottom of the plot correspond to the orders in Fig. 1. In several previous studies, only a single species was sampled for small clades such as Picramniales, preventing the estimation of crown ages; in these cases only the stem age is given here.

## IV. Discussion

Our 5-locus supermatrix represents the most comprehensive DNA data set yet compiled for the rosid clade. However, even with 19,740 ingroup species (out of 114,477 species estimated from OTT), our matrix is far from complete. Only 30,234 species of rosids (ca. 34% estimated from Hinchliff et al., 2015) have any type of DNA data in GenBank (see also Folk et al., 2018), and after a series of filtering steps, our topology represents only 16.25% of all rosid species recorded in OTT (Table 1). This relative sampling level (less than 20%) typifies most major clades of flowering plants (Eiserhardt et al., 2018; Folk et al., 2018).

Taxon sampling within rosids exhibits a strong phylogenetic bias (Fig. 1) in accumulation of molecular data (*p* ≈ 0). Among large families (> 1,000 species), the five with the poorest sampling (Polygalaceae, Myrtaceae, Rosaceae, Phyllanthaceae, and Malvaceae) have only 3.14% to 13.97% of species with at least one of the five loci sampled here in GenBank after matrix assembly and cleaning (cf. Table S2). For the five best-sampled families (Euphorbiaceae, Fabaceae, Passifloraceae, Brassicaceae, Cucurbitaceae), only 21.95% to 40.56% of species have one of the five loci. Hence, no large (> 1,000 species) family of rosids exceeds 45% species coverage, and most are below 30% coverage (Table S2), consistent with molecular sampling patterns across the angiosperms (Eiserhardt et al. 2018, Folk et al., 2018).

Our 5-locus topology is generally in close agreement with previous work (Wang et al., 2009; Soltis et al., 2011; Ruhfel et al., 2014; Sun et al., 2016), but with better overall resolution and without any cases of non-monophyletic families in the total-evidence trees (Fig. S1). Although the overall support across our tree is lower than that obtained in a previously published GenBank mining effort (Sun et al., 2016; Fig 2a), this is not surprising because: (1) studies have shown that BS values tend globally to decrease as the number of taxa increases (e.g., Sanderson & Donoghue, 1996; Sanderson & Wojciechowski, 2000; Soltis & Soltis, 2003; here the 5-locus tree has ∽ 2.2 fold the species sampling of the 4-locus tree); and (2) the standard phylogenetic bootstrap method (e.g., as implemented by RAxML) tends to yield particularly low support for deep branches with large sampling scales (Lemoine et al., 2018).

While our ITS tree yielded low overall resolution for relationships within the rosids, adding ITS data improved the phylogenetic resolution within 14 families compared to that obtained in the 4-locus tree in Sun et al. (2016; Table S3). Additionally, some nodes remain unresolved across data partitions (e.g., placement of Zygophyllales, Myrtales, and Vitales; Table S3), which likely reflect the rapid radiation of the rosid clade (Zhang et al., 2012, 2016; Zhao et al., 2016; Zeng et al., 2017; Green Plant Consortium, submitted). The divergence times here are generally consistent within methods and broadly congruent with the previous literature (Fig. 3). Between the two estimation methods, divergence times from PATHd8 were younger than those from treePL (Figs. 2b and 3), but highly correlated.

Despite the importance of the rosids to terrestrial landscapes, our knowledge of this clade remains limited (Folk et al., 2018), with species sampling gaps and bias that have likewise persisted across flowering plants and in many other major clades of life (Smith & Brown, 2018). Transparently assessing and quantifying data gaps is crucial for studies using large biodiversity data sets (Folk et al., 2018). Our exploration of phylogenetic sampling here highlights the essential role of taxonomic resources like Open Tree for assessing sampling gaps. We anticipate that similar analyses of cladewise sampling and explicit tests of phylogenetic bias will become standard approaches in large-scale studies as these become more numerous, and as discussion continues over their construction and use (e.g., Rabosky, 2015; Beaulieu & O’Meara, 2018; Folk et al., 2018; Donoghue & Edwards, 2019).

While limitations in sampling continue to hinder our understanding of angiosperm evolution, growth of sequence databases continues to be rapid, and our updated supermatrix effort has increased rosid species coverage by more than two-fold while recovering a backbone that is largely robust. Given the importance of deeply sampled global phylogenies for comparative biology (see Folk et al., 2018; Beaulieu & O’Meara, 2018; Smith & Brown, 2018; Allen et al., 2019), the data set we have assembled here will be an important resource for macroevolutionary synthesis across a globally important clade.

## Acknowledgements

This work was supported by the National Science Foundation (DEB-1208809 to D.E.S.), Dimensions of Biodiversity US-China (DEB-1442280 to P.S.S. and D.E.S.), ABI Innovation (DBI-1458640 to P.S.S. and D.E.S.), and National Natural Science Foundation of China (Grant no. 31590822 to Z.D.C.). We thank Dr. Greg Stull for valuable suggestions on the choice of fossil constraints, and Dr. Mark Miller and Dr. Pfeiffer Wayne from Cyber-Infrastructure for Phylogenetic Research (CIPRES) Science Gateway for their extended computation support. We thank the staff at the HiPerGator cluster at the University of Florida and CIPRES for providing us extensive computational resources.

## Author contributions

The authors declare no conflict interests. D.E.S., P.S.S., Z.D.C., and M.S. designed the study; M.S., S.A.S., and M.A.G. conducted GenBank data mining; M.S. and C.G.-A. performed OpenTree mapping; M.S. did the phylogeny and dating analyses; M.S., C.G.-A., and D.E.S. drafted the manuscript; R.A.F., D.E.S., S.A.S., M.A.G., P.S.S., Z.D.C., and R.P.G. revised the manuscript. All authors contributed to and approved the final manuscript.

## Supporting Information

**Fig. S1** The best ML tree obtained from the RAxML analysis.

**Table S1.** Information on the 59 rosid calibration constraints used in this study.

**Table S2.** Family-level summary sampling table for the 5-locus supermatrix (“Matrix”) compared to the rosid clade of the Open Tree Taxonomy (“OTT”) database v.3.0 (https://devtree.opentreeoflife.org/about/taxonomy-version/ott3.0; Hinchliff et al., 2015) and matched taxon names between these data sets.

**Table S3.** List of non-monophyletic families in the *matR* and ITS locus trees.

## Notes

#### Summary of Updates

1. updated one author's academic address 2. update higher resolution figures 3. minor text editing for Supplemental files 4. update the first author as the prior corresponding author for efficient actions upon biorxiv's action

