## Supplementary material for "Exploring the phylogeny of rosids with a five-locus supermatrix from GenBank": Fig. S1

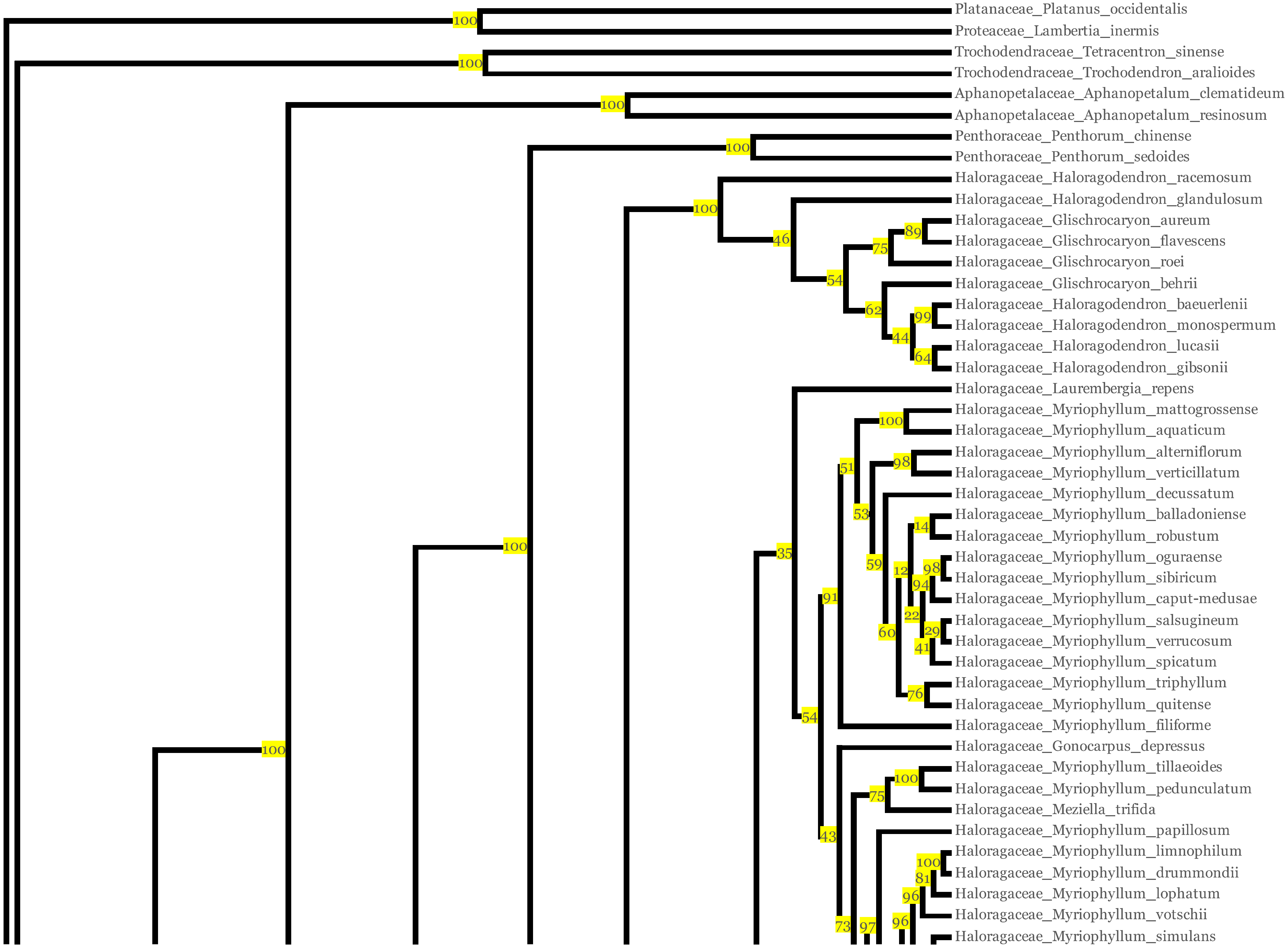
Platanaceae_Platanus_occidentalis
Proteaceae_Lambertia_inermis
Trochodendraceae_Tetracentron_sinense
Trochodendraceae_Trochodendron_aralioides
Aphanopetalaceae_Aphanopetalum_clematideum
Aphanopetalaceae_Aphanopetalum_resinosum
Penthoraceae_Penthorum_chinense
Penthoraceae_Penthorum_sedoides
Haloragaceae_Haloragodendron_racemosum
Haloragaceae_Haloragodendron_glandulosum
Haloragaceae_Glischrocaryon_aureum
Haloragaceae_Glischrocaryon_flavescens
Haloragaceae_Glischrocaryon_roei
Haloragaceae_Glischrocaryon_behrii
Haloragaceae_Haloragodendron_baeuerlenii
Haloragaceae_Haloragodendron_monospermum
Haloragaceae_Haloragodendron_lucasii
Haloragaceae_Haloragodendron_gibsonii
Haloragaceae_Laurembergia_repens
Haloragaceae_Myriophyllum_mattogrossense
Haloragaceae_Myriophyllum_aquaticum
Haloragaceae_Myriophyllum_alterniflorum
Haloragaceae_Myriophyllum_verticillatum
Haloragaceae_Myriophyllum_decussatum
Haloragaceae_Myriophyllum_balladoniense
Haloragaceae_Myriophyllum_robustum
Haloragaceae_Myriophyllum_oguraense
Haloragaceae_Myriophyllum_sibiricum
Haloragaceae_Myriophyllum_caput-medusae
Haloragaceae_Myriophyllum_salsugineum
Haloragaceae_Myriophyllum_verrucosum
Haloragaceae_Myriophyllum_spicatum
Haloragaceae_Myriophyllum_triphyllum
Haloragaceae_Myriophyllum_quitense
Haloragaceae_Myriophyllum_filiforme
Haloragaceae_Gonocarpus_depressus
Haloragaceae_Myriophyllum_tillaeoides
Haloragaceae_Myriophyllum_pedunculatum
Haloragaceae_Meziella_trifida
Haloragaceae_Myriophyllum_papillosum
Haloragaceae_Myriophyllum_limnophilum
Haloragaceae_Myriophyllum_drummondii
Haloragaceae_Myriophyllum_lophatum
Haloragaceae_Myriophyllum_votschii
Haloragaceae_Myriophyllum_simulans

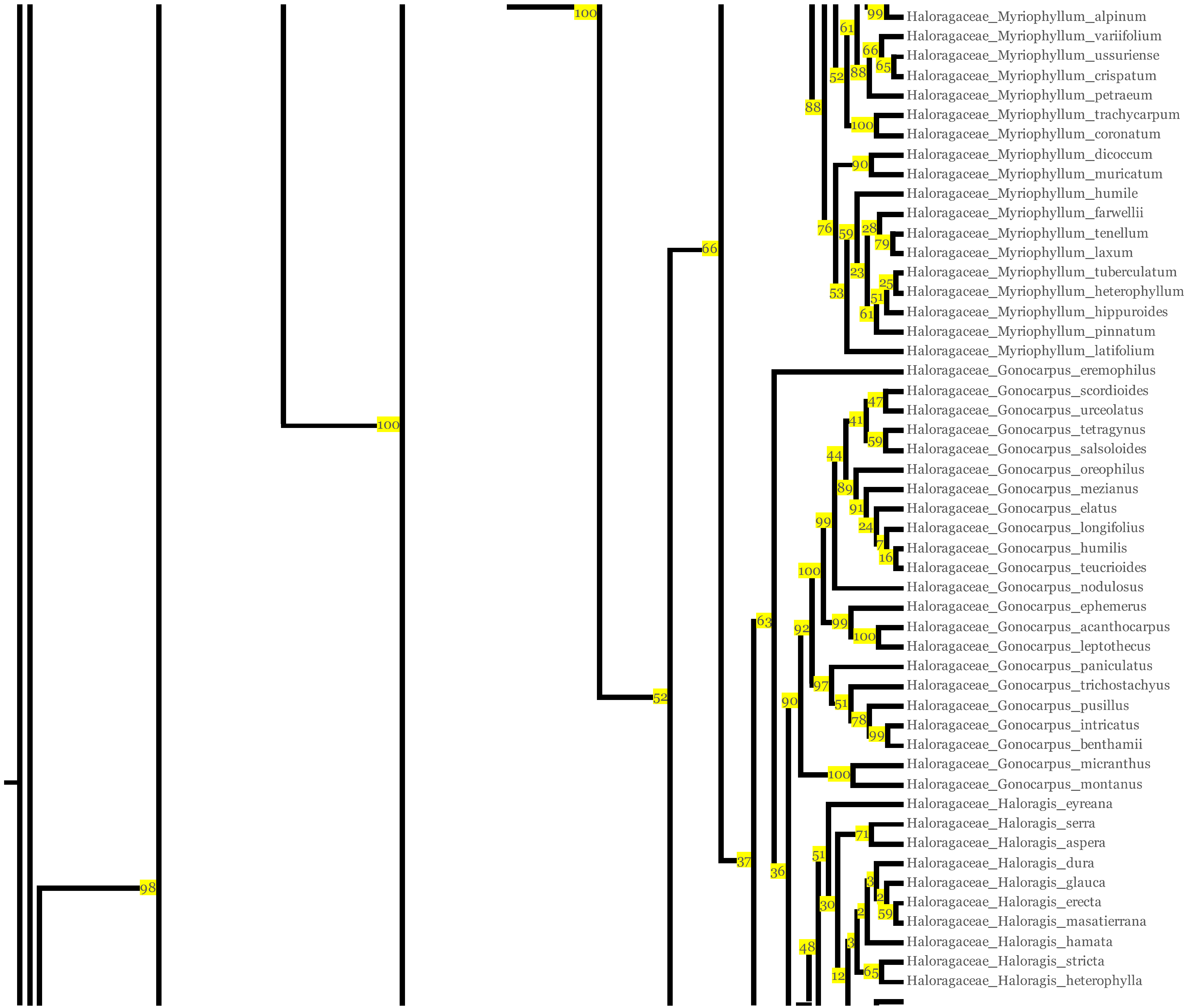
Haloragaceae_Myriophyllum_alpinum
Haloragaceae_Myriophyllum_variifolium
Haloragaceae_Myriophyllum_ussuriense
Haloragaceae_Myriophyllum_crispatum
Haloragaceae_Myriophyllum_petraeum
Haloragaceae_Myriophyllum_trachycarpum
Haloragaceae_Myriophyllum_coronatum
Haloragaceae_Myriophyllum_dicoccum
Haloragaceae_Myriophyllum_muricatum
Haloragaceae_Myriophyllum_humile
Haloragaceae_Myriophyllum_farwellii
Haloragaceae_Myriophyllum_tenellum
Haloragaceae_Myriophyllum_laxum
Haloragaceae_Myriophyllum_tuberculatum
Haloragaceae_Myriophyllum_heterophyllum
Haloragaceae_Myriophyllum_hippuroides
Haloragaceae_Myriophyllum_pinnatum
Haloragaceae_Myriophyllum_latifolium
Haloragaceae_Gonocarpus_eremophilus
Haloragaceae_Gonocarpus_scordioides
Haloragaceae_Gonocarpus_urceolatus
Haloragaceae_Gonocarpus_tetragynus
Haloragaceae_Gonocarpus_salsoloides
Haloragaceae_Gonocarpus_oreophilus
Haloragaceae_Gonocarpus_mezianus
Haloragaceae_Gonocarpus_elatus
Haloragaceae_Gonocarpus_longifolius
Haloragaceae_Gonocarpus_humilis
Haloragaceae_Gonocarpus_teucrioides
Haloragaceae_Gonocarpus_nodulosus
Haloragaceae_Gonocarpus_ephemerus
Haloragaceae_Gonocarpus_acanthocarpus
Haloragaceae_Gonocarpus_leptothecus
Haloragaceae_Gonocarpus_paniculatus
Haloragaceae_Gonocarpus_trichostachyus
Haloragaceae_Gonocarpus_pusillus
Haloragaceae_Gonocarpus_intricatus
Haloragaceae_Gonocarpus_benthamii
Haloragaceae_Gonocarpus_micranthus
Haloragaceae_Gonocarpus_montanus
Haloragaceae_Haloragis_eyreana
Haloragaceae_Haloragis_serra
Haloragaceae_Haloragis_aspera
Haloragaceae_Haloragis_dura
Haloragaceae_Haloragis_glauca
Haloragaceae_Haloragis_erecta
Haloragaceae_Haloragis_masatierrana
Haloragaceae_Haloragis_hamata
Haloragaceae_Haloragis_stricta
Haloragaceae_Haloragis_heterophylla

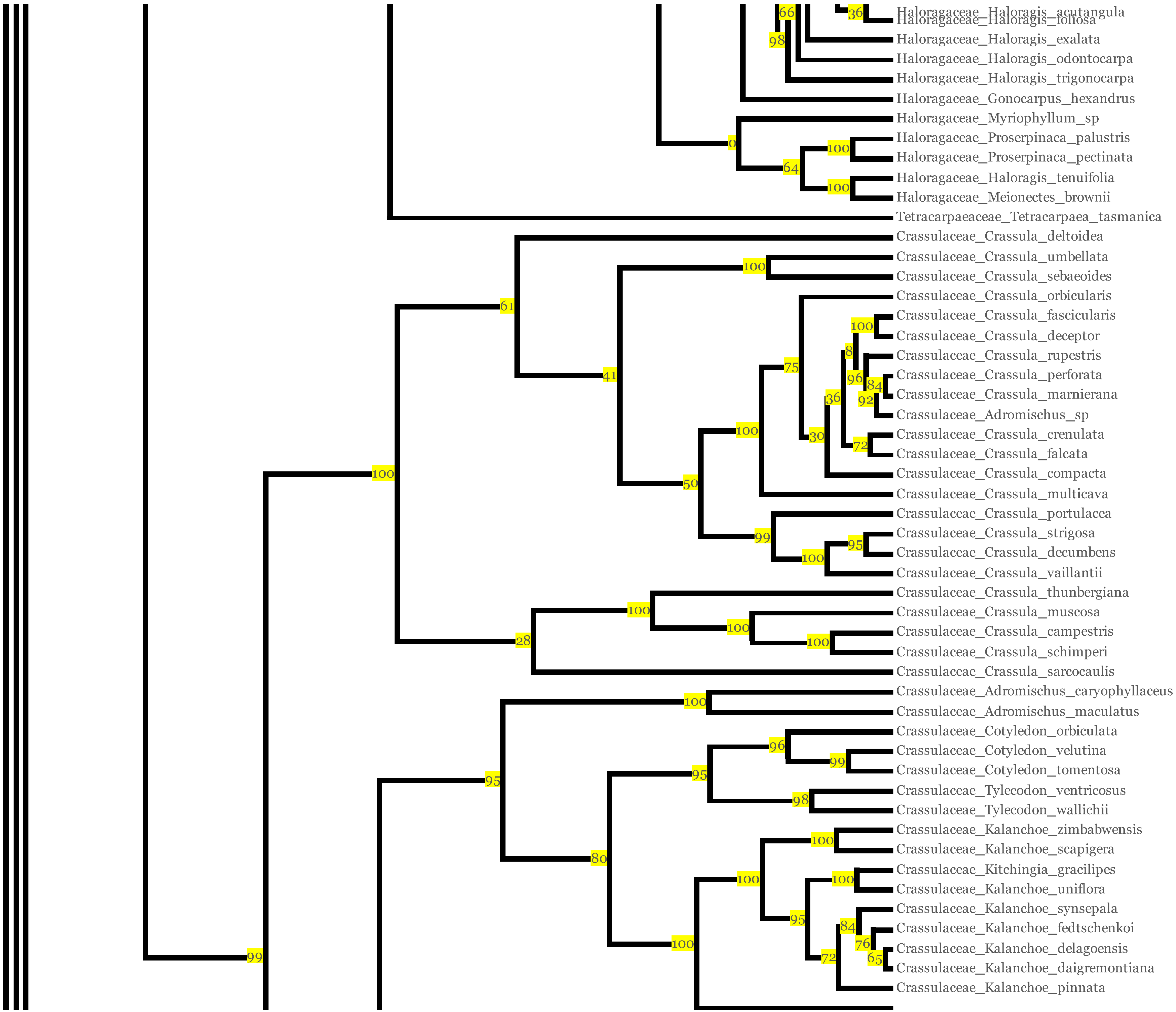
Haloragaceae_Haloragis_acutangula
Haloragaceae_Haloragis_foliosa
Haloragaceae_Haloragis_exalata
Haloragaceae_Haloragis_odontocarpa
Haloragaceae_Haloragis_trigonocarpa
Haloragaceae_Gonocarpus_hexandrus
Haloragaceae_Myriophyllum_sp
Haloragaceae_Proserpinaca_palustris
Haloragaceae_Proserpinaca_pectinata
Haloragaceae_Haloragis_tenuifolia
Haloragaceae_Meionectes_brownii
Tetracarpaeaceae_Tetracarpaea_tasmanica
Crassulaceae_Crassula_deltoidea
Crassulaceae_Crassula_umbellata
Crassulaceae_Crassula_sebaeoides
Crassulaceae_Crassula_orbicularis
Crassulaceae_Crassula_fascicularis
Crassulaceae_Crassula_deceptor
Crassulaceae_Crassula_rupestris
Crassulaceae_Crassula_perforata
Crassulaceae_Crassula_marnierana
Crassulaceae_Adromischus_sp
Crassulaceae_Crassula_crenulata
Crassulaceae_Crassula_falcata
Crassulaceae_Crassula_compacta
Crassulaceae_Crassula_multicava
Crassulaceae_Crassula_portulacea
Crassulaceae_Crassula_strigosa
Crassulaceae_Crassula_decumbens
Crassulaceae_Crassula_vaillantii
Crassulaceae_Crassula_thunbergiana
Crassulaceae_Crassula_muscosa
Crassulaceae_Crassula_campestris
Crassulaceae_Crassula_schimperi
Crassulaceae_Crassula_sarcocaulis
Crassulaceae_Adromischus_caryophyllaceus
Crassulaceae_Adromischus_maculatus
Crassulaceae_Cotyledon_orbiculata
Crassulaceae_Cotyledon_velutina
Crassulaceae_Cotyledon_tomentosa
Crassulaceae_Tylecodon_ventricosus
Crassulaceae_Tylecodon_wallichii
Crassulaceae_Kalanchoe_zimbabwensis
Crassulaceae_Kalanchoe_scapigera
Crassulaceae_Kitchingia_gracilipes
Crassulaceae_Kalanchoe_uniflora
Crassulaceae_Kalanchoe_synsepala
Crassulaceae_Kalanchoe_fedtschenkoi
Crassulaceae_Kalanchoe_delagoensis
Crassulaceae_Kalanchoe_daigremontiana
Crassulaceae_Kalanchoe_pinnata

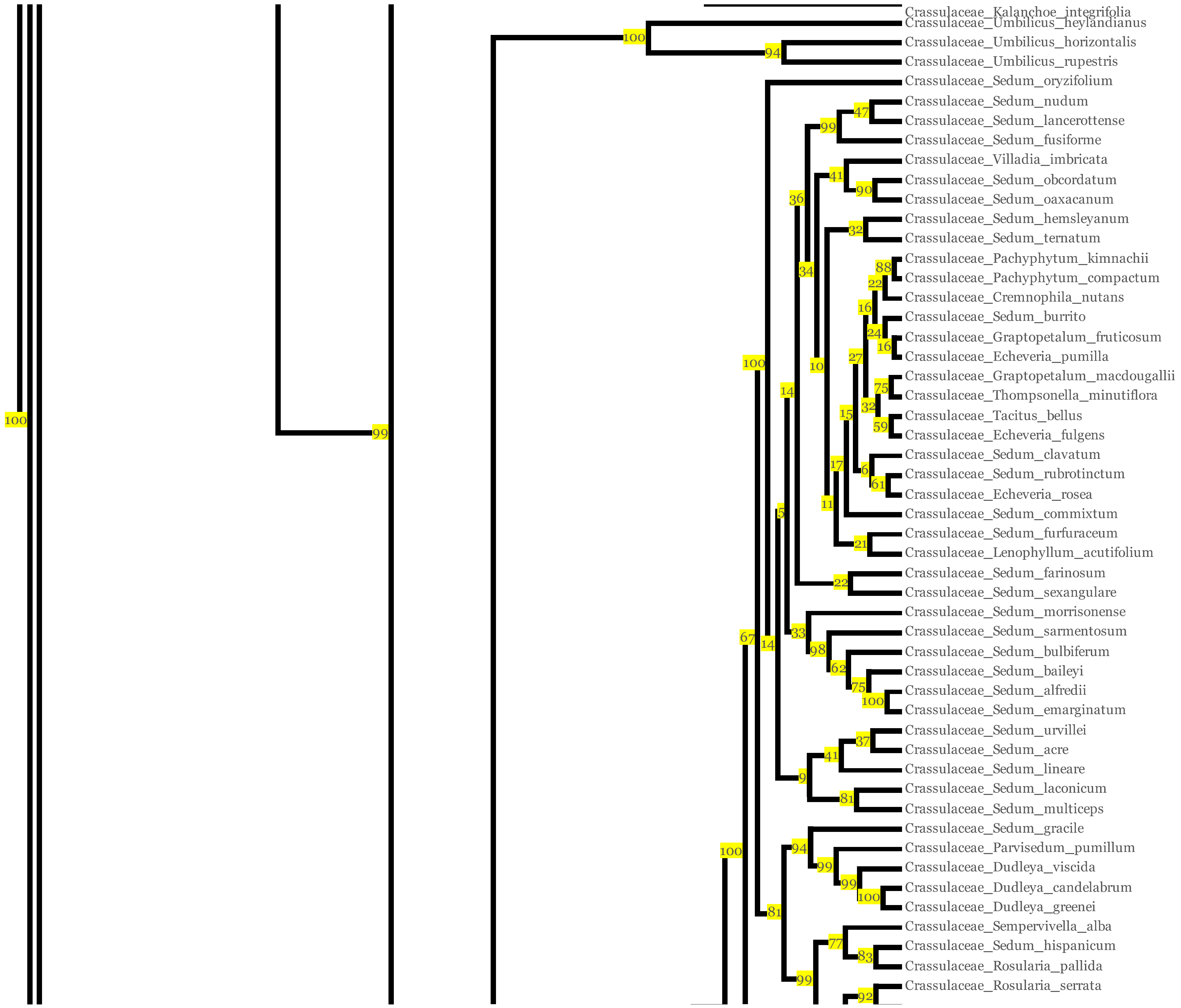
Crassulaceae_Kalanchoe_integrifolia
Crassulaceae_Umbilicus_heylandianus
Crassulaceae_Umbilicus_horizontalis
Crassulaceae_Umbilicus_rupestris
Crassulaceae_Sedum_oryzifolium
Crassulaceae_Sedum_nudum
Crassulaceae_Sedum_lancerottense
Crassulaceae_Sedum_fusiforme
Crassulaceae_Villadia_imbricata
Crassulaceae_Sedum_obcordatum
Crassulaceae_Sedum_oaxacanum
Crassulaceae_Sedum_hemsleyanum
Crassulaceae_Sedum_ternatum
Crassulaceae_Pachyphytum_kimnachii
Crassulaceae_Pachyphytum_compactum
Crassulaceae_Cremnophila_nutans
Crassulaceae_Sedum_burrito
Crassulaceae_Graptopetalum_fruticosum
Crassulaceae_Echeveria_pumilla
Crassulaceae_Graptopetalum_macdougallii
Crassulaceae_Thompsonella_minutiflora
Crassulaceae_Tacitus_bellus
Crassulaceae_Echeveria_fulgens
Crassulaceae_Sedum_clavatum
Crassulaceae_Sedum_rubrotinctum
Crassulaceae_Echeveria_rosea
Crassulaceae_Sedum_commixtum
Crassulaceae_Sedum_furfuraceum
Crassulaceae_Lenophyllum_acutifolium
Crassulaceae_Sedum_farinosum
Crassulaceae_Sedum_sexangulare
Crassulaceae_Sedum_morrisonense
Crassulaceae_Sedum_sarmentosum
Crassulaceae_Sedum_bulbiferum
Crassulaceae_Sedum_baileyi
Crassulaceae_Sedum_alfredii
Crassulaceae_Sedum_emarginatum
Crassulaceae_Sedum_urvillei
Crassulaceae_Sedum_acre
Crassulaceae_Sedum_lineare
Crassulaceae_Sedum_laconicum
Crassulaceae_Sedum_multiceps
Crassulaceae_Sedum_gracile
Crassulaceae_Parvisedum_pumillum
Crassulaceae_Dudleya_viscida
Crassulaceae_Dudleya_candelabrum
Crassulaceae_Dudleya_greenei
Crassulaceae_Sempervivella_alba
Crassulaceae_Sedum_hispanicum
Crassulaceae_Rosularia_pallida
Crassulaceae_Rosularia_serrata

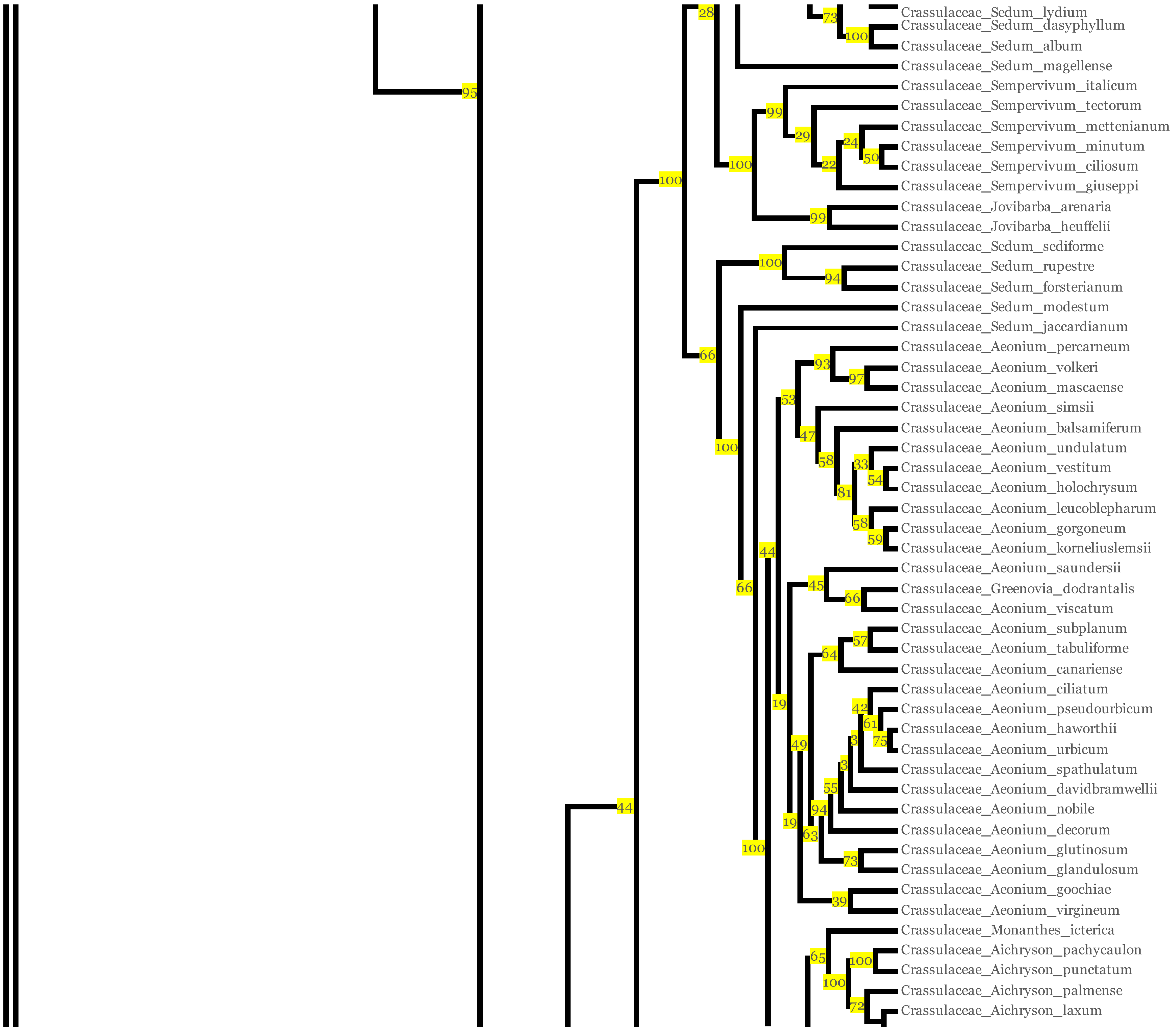
Crassulaceae_Sedum_lydium
Crassulaceae_Sedum_dasyphyllum
Crassulaceae_Sedum_album
Crassulaceae_Sedum_magellense
Crassulaceae_Sempervivum_italicum
Crassulaceae_Sempervivum_tectorum
Crassulaceae_Sempervivum_mettenianum
Crassulaceae_Sempervivum_minutum
Crassulaceae_Sempervivum_ciliosum
Crassulaceae_Sempervivum_giuseppi
Crassulaceae_Jovibarba_arenaria
Crassulaceae_Jovibarba_heuffelii
Crassulaceae_Sedum_sediforme
Crassulaceae_Sedum_rupestre
Crassulaceae_Sedum_forsterianum
Crassulaceae_Sedum_modestum
Crassulaceae_Sedum_jaccardianum
Crassulaceae_Aeonium_percarneum
Crassulaceae_Aeonium_volkeri
Crassulaceae_Aeonium_mascaense
Crassulaceae_Aeonium_simsii
Crassulaceae_Aeonium_balsamiferum
Crassulaceae_Aeonium_undulatum
Crassulaceae_Aeonium_vestitum
Crassulaceae_Aeonium_holochrysum
Crassulaceae_Aeonium_leucoblepharum
Crassulaceae_Aeonium_gorgoneum
Crassulaceae_Aeonium_korneliuslemsii
Crassulaceae_Aeonium_saundersii
Crassulaceae_Greenovia_dodrantalis
Crassulaceae_Aeonium_viscatum
Crassulaceae_Aeonium_subplanum
Crassulaceae_Aeonium_tabuliforme
Crassulaceae_Aeonium_canariense
Crassulaceae_Aeonium_ciliatum
Crassulaceae_Aeonium_pseudourbicum
Crassulaceae_Aeonium_haworthii
Crassulaceae_Aeonium_urbicum
Crassulaceae_Aeonium_spathulatum
Crassulaceae_Aeonium_davidbramwellii
Crassulaceae_Aeonium_nobile
Crassulaceae_Aeonium_decorum
Crassulaceae_Aeonium_glutinosum
Crassulaceae_Aeonium_glandulosum
Crassulaceae_Aeonium_goochiae
Crassulaceae_Aeonium_virgineum
Crassulaceae_Monanthes_icterica
Crassulaceae_Aichryson_pachycaulon
Crassulaceae_Aichryson_punctatum
Crassulaceae_Aichryson_palmense
Crassulaceae_Aichryson_laxum

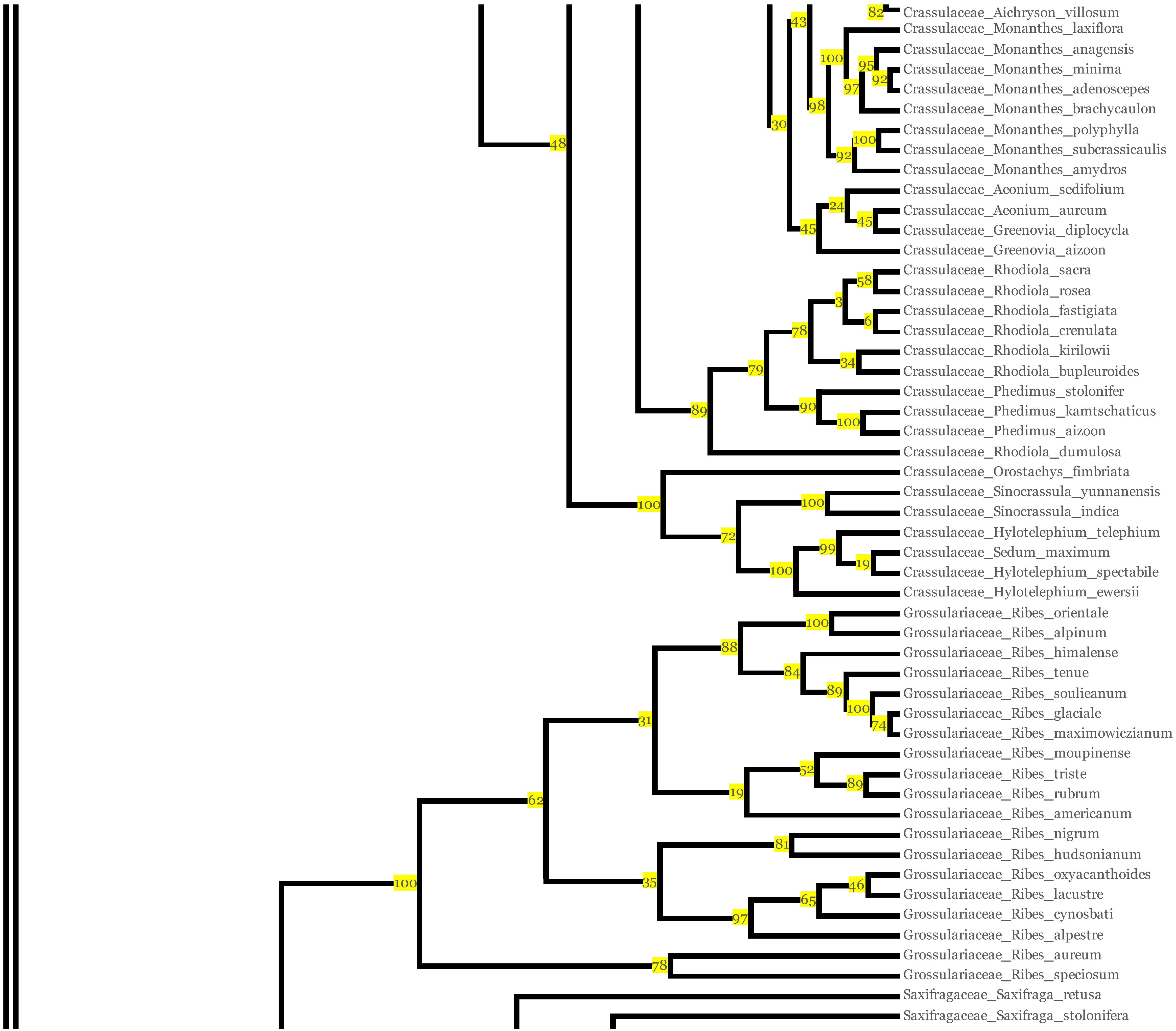
Crassulaceae_Aichryson_villosum
Crassulaceae_Monanthes_laxiflora
Crassulaceae_Monanthes_anagensis
Crassulaceae_Monanthes_minima
Crassulaceae_Monanthes_adenoscepes
Crassulaceae_Monanthes_brachycaulon
Crassulaceae_Monanthes_polyphylla
Crassulaceae_Monanthes_subcrassicaulis
Crassulaceae_Monanthes_amydros
Crassulaceae_Aeonium_sedifolium
Crassulaceae_Aeonium_aureum
Crassulaceae_Greenovia_diplocycla
Crassulaceae_Greenovia_aizoon
Crassulaceae_Rhodiola_sacra
Crassulaceae_Rhodiola_rosea
Crassulaceae_Rhodiola_fastigiata
Crassulaceae_Rhodiola_crenulata
Crassulaceae_Rhodiola_kirilowii
Crassulaceae_Rhodiola_bupleuroides
Crassulaceae_Phedimus_stolonifer
Crassulaceae_Phedimus_kamtschaticus
Crassulaceae_Phedimus_aizoon
Crassulaceae_Rhodiola_dumulosa
Crassulaceae_Orostachys_fimbriata
Crassulaceae_Sinocrassula_yunnanensis
Crassulaceae_Sinocrassula_indica
Crassulaceae_Hylotelephium_telephium
Crassulaceae_Sedum_maximum
Crassulaceae_Hylotelephium_spectabile
Crassulaceae_Hylotelephium_ewersii
Grossulariaceae_Ribes_orientale
Grossulariaceae_Ribes_alpinum
Grossulariaceae_Ribes_himalense
Grossulariaceae_Ribes_tenue
Grossulariaceae_Ribes_soulieanum
Grossulariaceae_Ribes_glaciale
Grossulariaceae_Ribes_maximowiczianum
Grossulariaceae_Ribes_moupinense
Grossulariaceae_Ribes_triste
Grossulariaceae_Ribes_rubrum
Grossulariaceae_Ribes_americanum
Grossulariaceae_Ribes_nigrum
Grossulariaceae_Ribes_hudsonianum
Grossulariaceae_Ribes_oxyacanthoides
Grossulariaceae_Ribes_lacustre
Grossulariaceae_Ribes_cynosbati
Grossulariaceae_Ribes_alpestre
Grossulariaceae_Ribes_aureum
Grossulariaceae_Ribes_speciosum
Saxifragaceae_Saxifraga_retusa
Saxifragaceae_Saxifraga_stolonifera
82
43
100
95
92
97
98
30
48
100
92
24
45
45
58
3
6
78
34
79
90
100
89
100
100
72
99
19
100
100
88
84
89
100
74
31
52
89
19
62
81
46
65
35
97
100
78

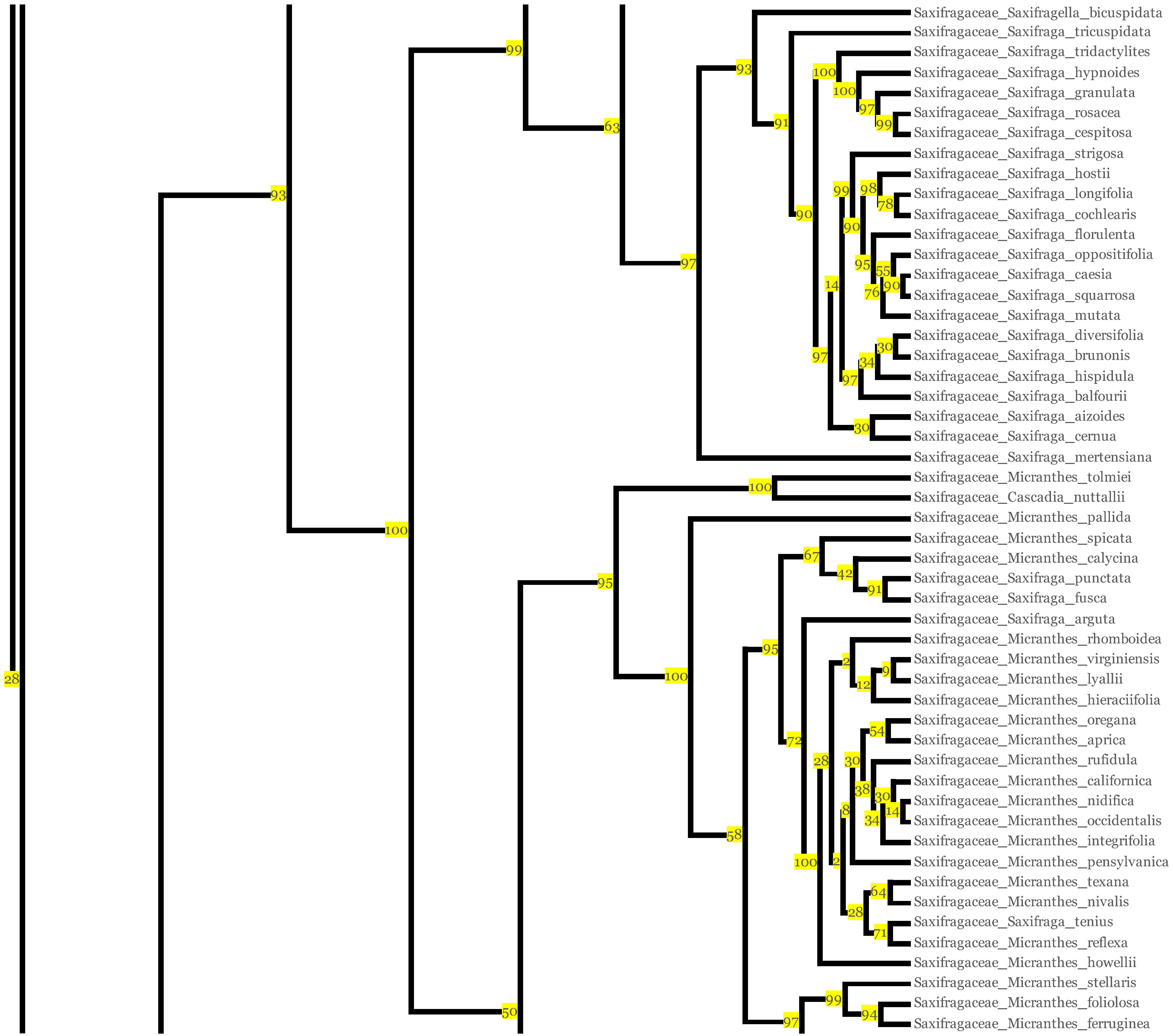
Saxifragaceae_Saxifragella_bicuspidata
Saxifragaceae_Saxifraga_tricuspidata
Saxifragaceae_Saxifraga_tridactylites
Saxifragaceae_Saxifraga_hypnoides
Saxifragaceae_Saxifraga_granulata
Saxifragaceae_Saxifraga_rosacea
Saxifragaceae_Saxifraga_cespitosa
Saxifragaceae_Saxifraga_strigosa
Saxifragaceae_Saxifraga_hostii
Saxifragaceae_Saxifraga_longifolia
Saxifragaceae_Saxifraga_cochlearis
Saxifragaceae_Saxifraga_florulenta
Saxifragaceae_Saxifraga_oppositifolia
Saxifragaceae_Saxifraga_caesia
Saxifragaceae_Saxifraga_squarrosa
Saxifragaceae_Saxifraga_mutata
Saxifragaceae_Saxifraga_diversifolia
Saxifragaceae_Saxifraga_brunonis
Saxifragaceae_Saxifraga_hispidula
Saxifragaceae_Saxifraga_balfourii
Saxifragaceae_Saxifraga_aizoides
Saxifragaceae_Saxifraga_cernua
Saxifragaceae_Saxifraga_mertensiana
Saxifragaceae_Micranthes_tolmiei
Saxifragaceae_Cascadia_nuttallii
Saxifragaceae_Micranthes_pallida
Saxifragaceae_Micranthes_spicata
Saxifragaceae_Micranthes_calycina
Saxifragaceae_Saxifraga_punctata
Saxifragaceae_Saxifraga_fusca
Saxifragaceae_Saxifraga_arguta
Saxifragaceae_Micranthes_rhomboidea
Saxifragaceae_Micranthes_virginiensis
Saxifragaceae_Micranthes_lyallii
Saxifragaceae_Micranthes_hieraciifolia
Saxifragaceae_Micranthes_oregana
Saxifragaceae_Micranthes_aprica
Saxifragaceae_Micranthes_rufidula
Saxifragaceae_Micranthes_californica
Saxifragaceae_Micranthes_nidifica
Saxifragaceae_Micranthes_occidentalis
Saxifragaceae_Micranthes_integrifolia
Saxifragaceae_Micranthes_pensylvanica
Saxifragaceae_Micranthes_texana
Saxifragaceae_Micranthes_nivalis
Saxifragaceae_Saxifraga_tenius
Saxifragaceae_Micranthes_reflexa
Saxifragaceae_Micranthes_howellii
Saxifragaceae_Micranthes_stellaris
Saxifragaceae_Micranthes_foliolosa
Saxifragaceae_Micranthes_ferruginea

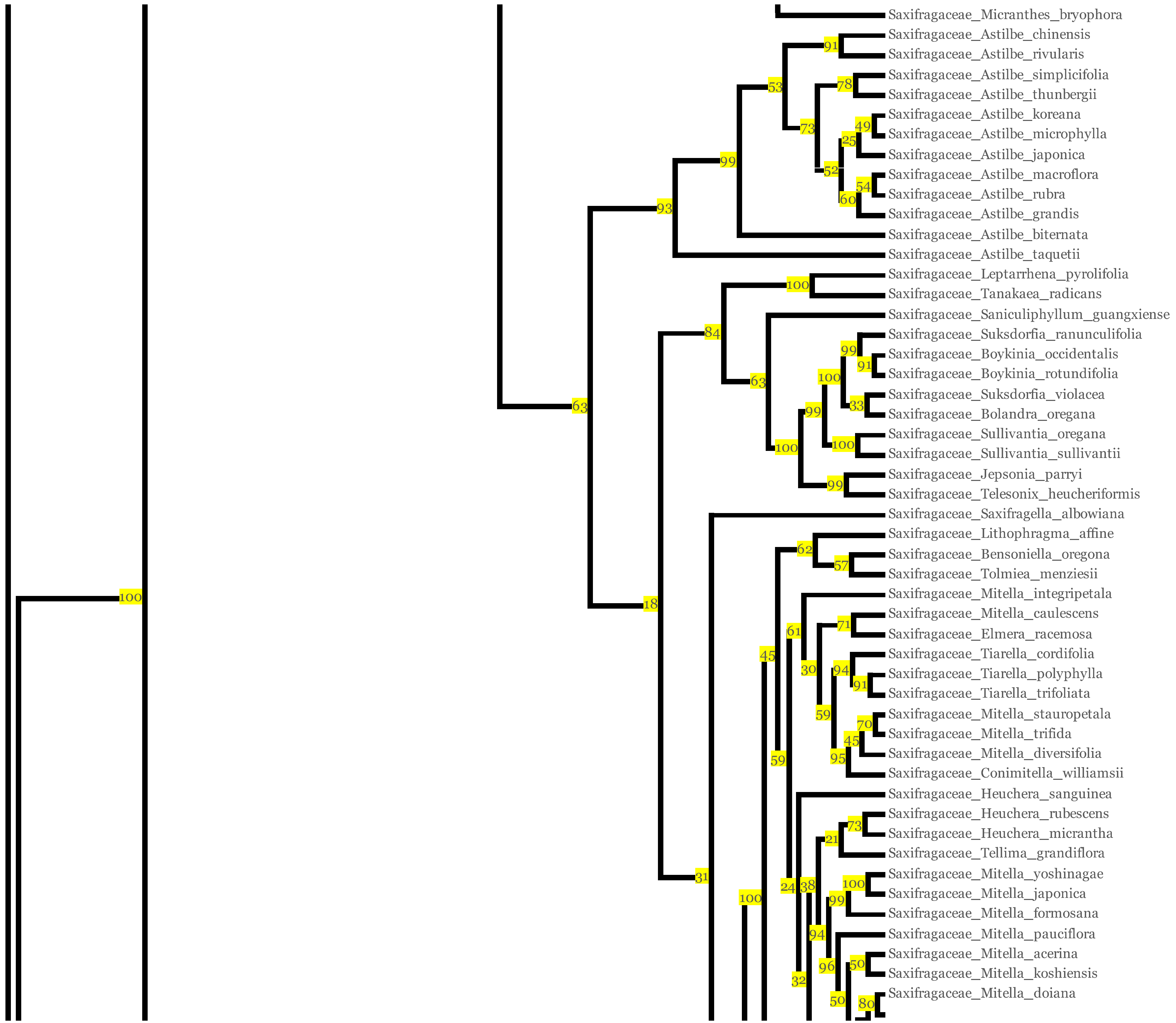
Saxifragaceae_Micranthes_bryophora
Saxifragaceae_Astilbe_chinensis
Saxifragaceae_Astilbe_rivularis
Saxifragaceae_Astilbe_simplicifolia
Saxifragaceae_Astilbe_thunbergii
Saxifragaceae_Astilbe_koreana
Saxifragaceae_Astilbe_microphylla
Saxifragaceae_Astilbe_japonica
Saxifragaceae_Astilbe_macroflora
Saxifragaceae_Astilbe_rubra
Saxifragaceae_Astilbe_grandis
Saxifragaceae_Astilbe_biternata
Saxifragaceae_Astilbe_taquetii
Saxifragaceae_Leptarrhena_pyrolifolia
Saxifragaceae_Tanakaea_radicans
Saxifragaceae_Saniculiphyllum_guangxiense
Saxifragaceae_Suksdorfia_ranunculifolia
Saxifragaceae_Boykinia_occidentalis
Saxifragaceae_Boykinia_rotundifolia
Saxifragaceae_Suksdorfia_violacea
Saxifragaceae_Bolandra_oregana
Saxifragaceae_Sullivantia_oregana
Saxifragaceae_Sullivantia_sullivantii
Saxifragaceae_Jepsonia_parryi
Saxifragaceae_Telesonix_heucheriformis
Saxifragaceae_Saxifragella_albowiana
Saxifragaceae_Lithophragma_affine
Saxifragaceae_Bensoniella_oregona
Saxifragaceae_Tolmiea_menziesii
Saxifragaceae_Mitella_integripetala
Saxifragaceae_Mitella_caulescens
Saxifragaceae_Elmera_racemosa
Saxifragaceae_Tiarella_cordifolia
Saxifragaceae_Tiarella_polyphylla
Saxifragaceae_Tiarella_trifoliata
Saxifragaceae_Mitella_stauropetala
Saxifragaceae_Mitella_trifida
Saxifragaceae_Mitella_diversifolia
Saxifragaceae_Conimitella_williamsii
Saxifragaceae_Heuchera_sanguinea
Saxifragaceae_Heuchera_rubescens
Saxifragaceae_Heuchera_micrantha
Saxifragaceae_Tellima_grandiflora
Saxifragaceae_Mitella_yoshinagae
Saxifragaceae_Mitella_japonica
Saxifragaceae_Mitella_formosana
Saxifragaceae_Mitella_pauciflora
Saxifragaceae_Mitella_acerina
Saxifragaceae_Mitella_koshiensis
Saxifragaceae_Mitella_doiana
91
78
53
73
49
25
52
54
60
99
93
100
84
63
99
91
100
33
99
100
100
99
62
57
63
18
61
71
45
30
94
91
59
70
45
95
59
73
21
100
24
38
99
31
100
94
96
50
32
50
80
100

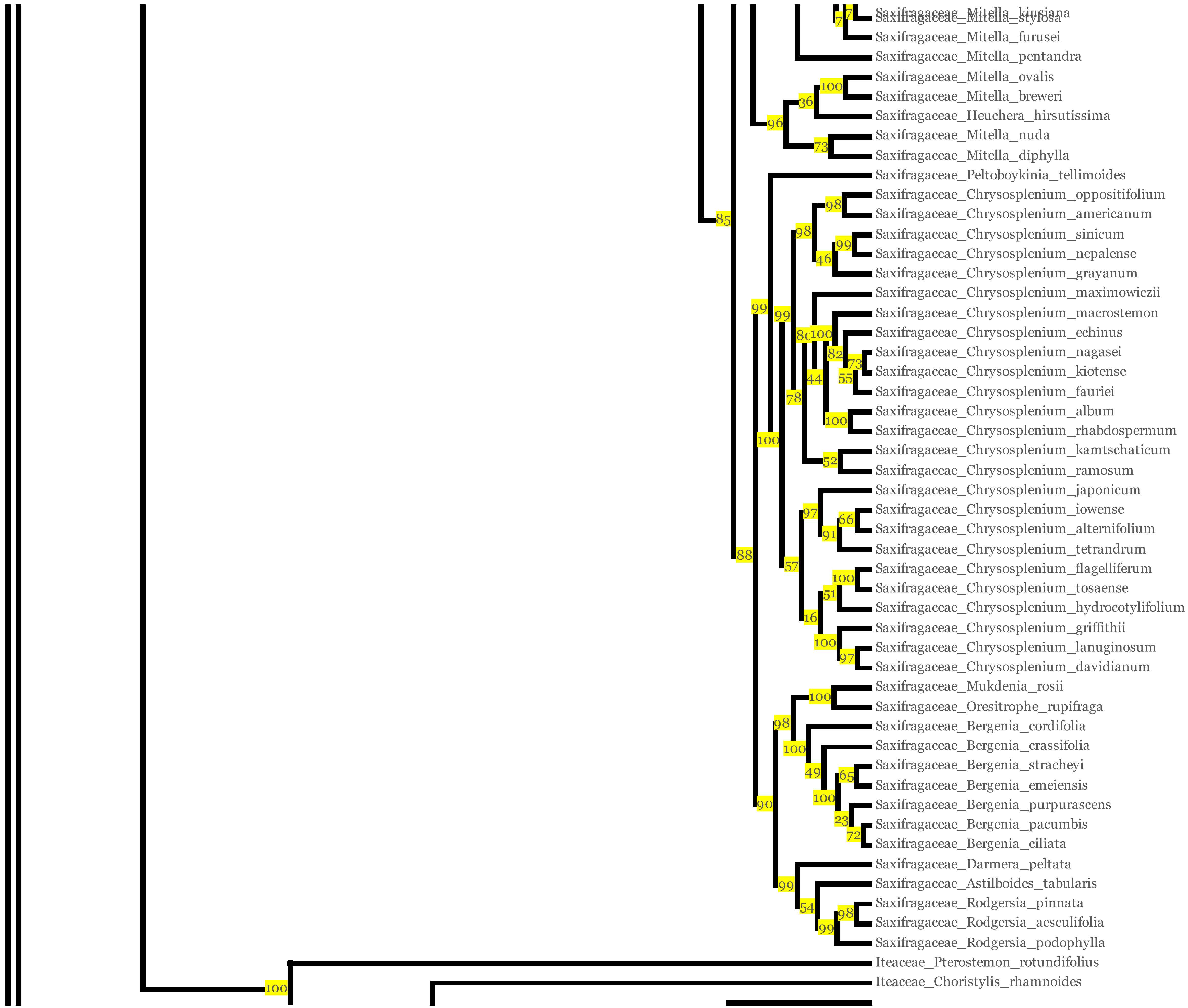
Saxifragaceae_Mitella_kiusiana
Saxifragaceae_Mitella_stylosa
Saxifragaceae_Mitella_furusei
Saxifragaceae_Mitella_pentandra
Saxifragaceae_Mitella_ovalis
Saxifragaceae_Mitella_breweri
Saxifragaceae_Heuchera_hirsutissima
Saxifragaceae_Mitella_nuda
Saxifragaceae_Mitella_diphylla
Saxifragaceae_Peltoboykinia_tellimoides
Saxifragaceae_Chrysosplenium_oppositifolium
Saxifragaceae_Chrysosplenium_americanum
Saxifragaceae_Chrysosplenium_sinicum
Saxifragaceae_Chrysosplenium_nepalense
Saxifragaceae_Chrysosplenium_grayanum
Saxifragaceae_Chrysosplenium_maximowiczii
Saxifragaceae_Chrysosplenium_macrostemon
Saxifragaceae_Chrysosplenium_echinus
Saxifragaceae_Chrysosplenium_nagasei
Saxifragaceae_Chrysosplenium_kiotense
Saxifragaceae_Chrysosplenium_fauriei
Saxifragaceae_Chrysosplenium_album
Saxifragaceae_Chrysosplenium_rhabdospermum
Saxifragaceae_Chrysosplenium_kamtschaticum
Saxifragaceae_Chrysosplenium_ramosum
Saxifragaceae_Chrysosplenium_japonicum
Saxifragaceae_Chrysosplenium_iowense
Saxifragaceae_Chrysosplenium_alternifolium
Saxifragaceae_Chrysosplenium_tetrandrum
Saxifragaceae_Chrysosplenium_flagelliferum
Saxifragaceae_Chrysosplenium_tosaense
Saxifragaceae_Chrysosplenium_hydrocotylifolium
Saxifragaceae_Chrysosplenium_griffithii
Saxifragaceae_Chrysosplenium_lanuginosum
Saxifragaceae_Chrysosplenium_davidianum
Saxifragaceae_Mukdenia_rosii
Saxifragaceae_Oresitrophe_rupifraga
Saxifragaceae_Bergenia_cordifolia
Saxifragaceae_Bergenia_crassifolia
Saxifragaceae_Bergenia_stracheyi
Saxifragaceae_Bergenia_emeiensis
Saxifragaceae_Bergenia_purpurascens
Saxifragaceae_Bergenia_pacumbis
Saxifragaceae_Bergenia_ciliata
Saxifragaceae_Darmera_peltata
Saxifragaceae_Astilboides_tabularis
Saxifragaceae_Rodgersia_pinnata
Saxifragaceae_Rodgersia_aesculifolia
Saxifragaceae_Rodgersia_podophylla
Iteaceae_Pterostemon_rotundifolius
Iteaceae_Choristylis_rhamnoides

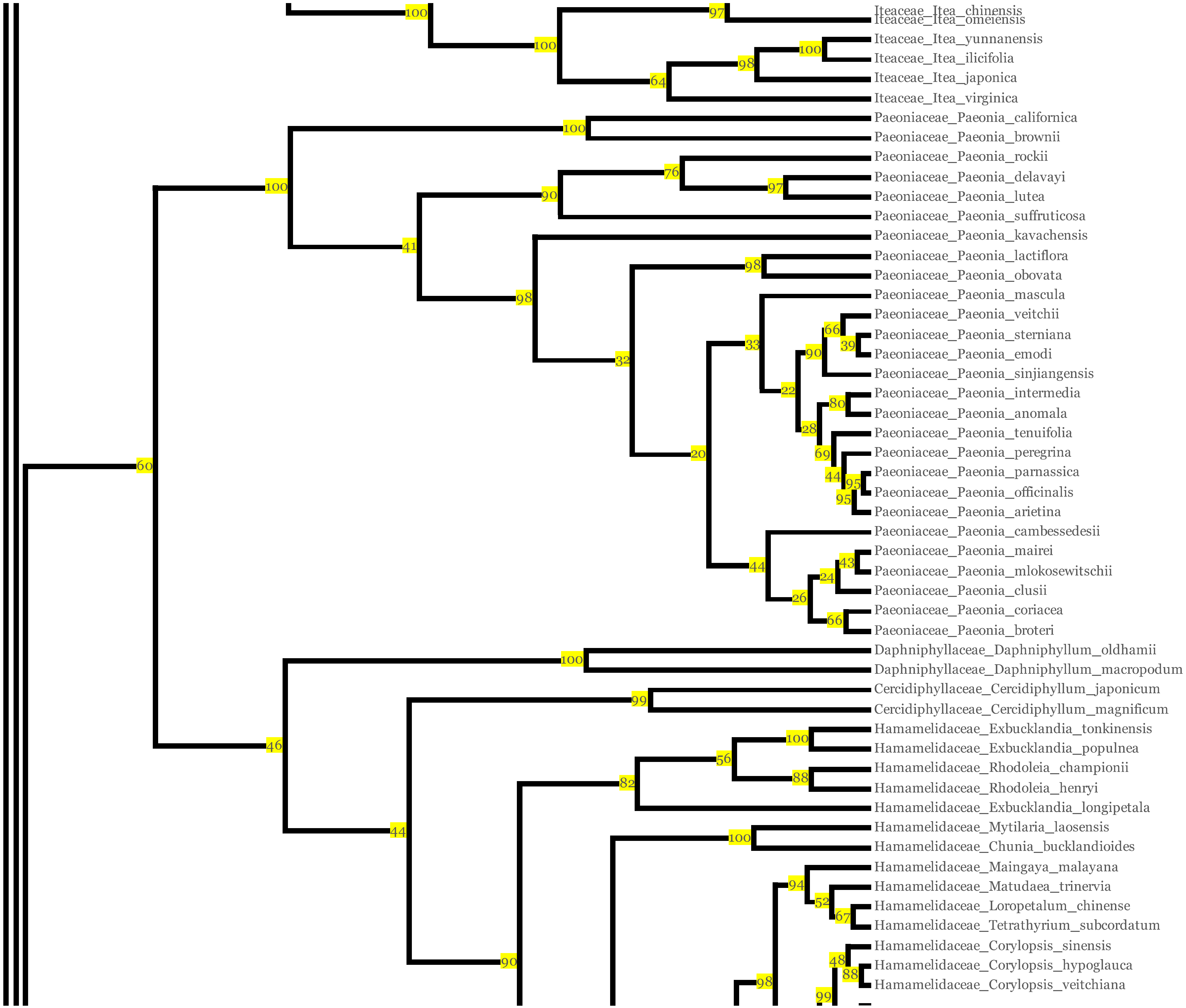
Iteaceae_Itea_chinensis
Iteaceae_Itea_omeiensis
Iteaceae_Itea_yunnanensis
Iteaceae_Itea_ilicifolia
Iteaceae_Itea_japonica
Iteaceae_Itea_virginica
Paeoniaceae_Paeonia_californica
Paeoniaceae_Paeonia_brownii
Paeoniaceae_Paeonia_rockii
Paeoniaceae_Paeonia_delavayi
Paeoniaceae_Paeonia_lutea
Paeoniaceae_Paeonia_suffruticosa
Paeoniaceae_Paeonia_kavachensis
Paeoniaceae_Paeonia_lactiflora
Paeoniaceae_Paeonia_obovata
Paeoniaceae_Paeonia_mascula
Paeoniaceae_Paeonia_veitchii
Paeoniaceae_Paeonia_sterniana
Paeoniaceae_Paeonia_emodi
Paeoniaceae_Paeonia_sinjiangensis
Paeoniaceae_Paeonia_intermedia
Paeoniaceae_Paeonia_anomala
Paeoniaceae_Paeonia_tenuifolia
Paeoniaceae_Paeonia_peregrina
Paeoniaceae_Paeonia_parnassica
Paeoniaceae_Paeonia_officinalis
Paeoniaceae_Paeonia_arietina
Paeoniaceae_Paeonia_cambessedesii
Paeoniaceae_Paeonia_mairei
Paeoniaceae_Paeonia_mlokosewitschii
Paeoniaceae_Paeonia_clusii
Paeoniaceae_Paeonia_coriacea
Paeoniaceae_Paeonia_broteri
Daphniphyllaceae_Daphniphyllum_oldhamii
Daphniphyllaceae_Daphniphyllum_macropodum
Cercidiphyllaceae_Cercidiphyllum_japonicum
Cercidiphyllaceae_Cercidiphyllum_magnificum
Hamamelidaceae_Exbucklandia_tonkinensis
Hamamelidaceae_Exbucklandia_populnea
Hamamelidaceae_Rhodoleia_championii
Hamamelidaceae_Rhodoleia_henryi
Hamamelidaceae_Exbucklandia_longipetala
Hamamelidaceae_Mytilaria_laosensis
Hamamelidaceae_Chunia_bucklandioides
Hamamelidaceae_Maingaya_malayana
Hamamelidaceae_Matudaea_trinervia
Hamamelidaceae_Loropetalum_chinense
Hamamelidaceae_Tetrathyrium_subcordatum
Hamamelidaceae_Corylopsis_sinensis
Hamamelidaceae_Corylopsis_hypoglauca
Hamamelidaceae_Corylopsis_veitchiana

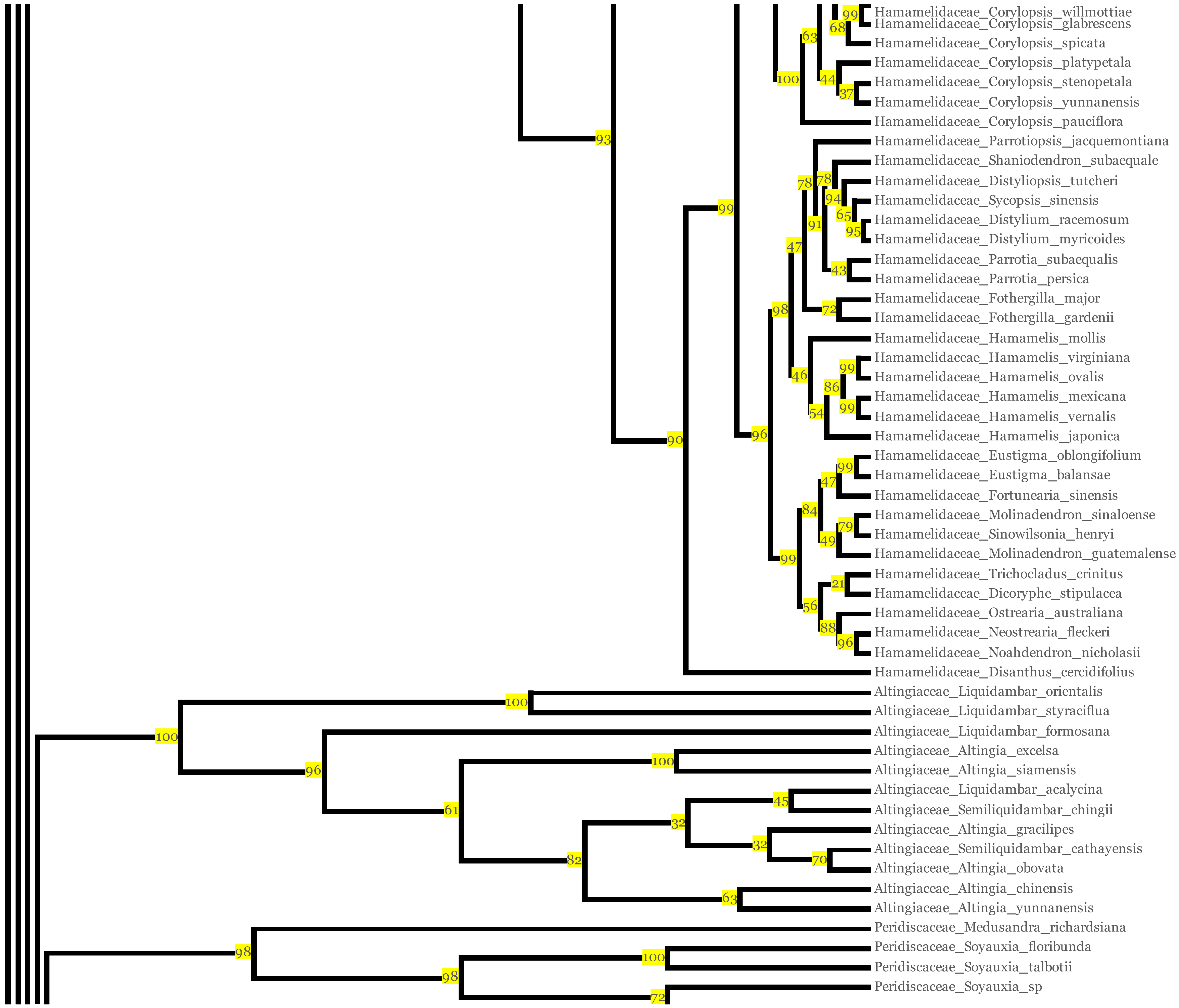
Hamamelidaceae_Corylopsis_willmottiae
Hamamelidaceae_Corylopsis_glabrescens
Hamamelidaceae_Corylopsis_spicata
Hamamelidaceae_Corylopsis_platypetala
Hamamelidaceae_Corylopsis_stenopetala
Hamamelidaceae_Corylopsis_yunnanensis
Hamamelidaceae_Corylopsis_pauciflora
Hamamelidaceae_Parrotiopsis_jacquemontiana
Hamamelidaceae_Shaniodendron_subaequale
Hamamelidaceae_Distyliopsis_tutcheri
Hamamelidaceae_Sycopsis_sinensis
Hamamelidaceae_Distylium_racemosum
Hamamelidaceae_Distylium_myricoides
Hamamelidaceae_Parrotia_subaequalis
Hamamelidaceae_Parrotia_persica
Hamamelidaceae_Fothergilla_major
Hamamelidaceae_Fothergilla_gardenii
Hamamelidaceae_Hamamelis_mollis
Hamamelidaceae_Hamamelis_virginiana
Hamamelidaceae_Hamamelis_ovalis
Hamamelidaceae_Hamamelis_mexicana
Hamamelidaceae_Hamamelis_vernalis
Hamamelidaceae_Hamamelis_japonica
Hamamelidaceae_Eustigma_oblongifolium
Hamamelidaceae_Eustigma_balansae
Hamamelidaceae_Fortunearia_sinensis
Hamamelidaceae_Molinadendron_sinaloense
Hamamelidaceae_Sinowilsonia_henryi
Hamamelidaceae_Molinadendron_guatemalense
Hamamelidaceae_Trichocladus_crinitus
Hamamelidaceae_Dicoryphe_stipulacea
Hamamelidaceae_Ostrearia_australiana
Hamamelidaceae_Neostrearia_fleckeri
Hamamelidaceae_Noahdendron_nicholasii
Hamamelidaceae_Disanthus_cercidifolius
Altingiaceae_Liquidambar_orientalis
Altingiaceae_Liquidambar_styraciflua
Altingiaceae_Liquidambar_formosana
Altingiaceae_Altingia_excelsa
Altingiaceae_Altingia_siamensis
Altingiaceae_Liquidambar_acalycina
Altingiaceae_Semiliquidambar_chingii
Altingiaceae_Altingia_gracilipes
Altingiaceae_Semiliquidambar_cathayensis
Altingiaceae_Altingia_obovata
Altingiaceae_Altingia_chinensis
Altingiaceae_Altingia_yunnanensis
Peridiscaceae_Medusandra_richardsiana
Peridiscaceae_Soyauxia_floribunda
Peridiscaceae_Soyauxia_talbotii
Peridiscaceae_Soyauxia_sp

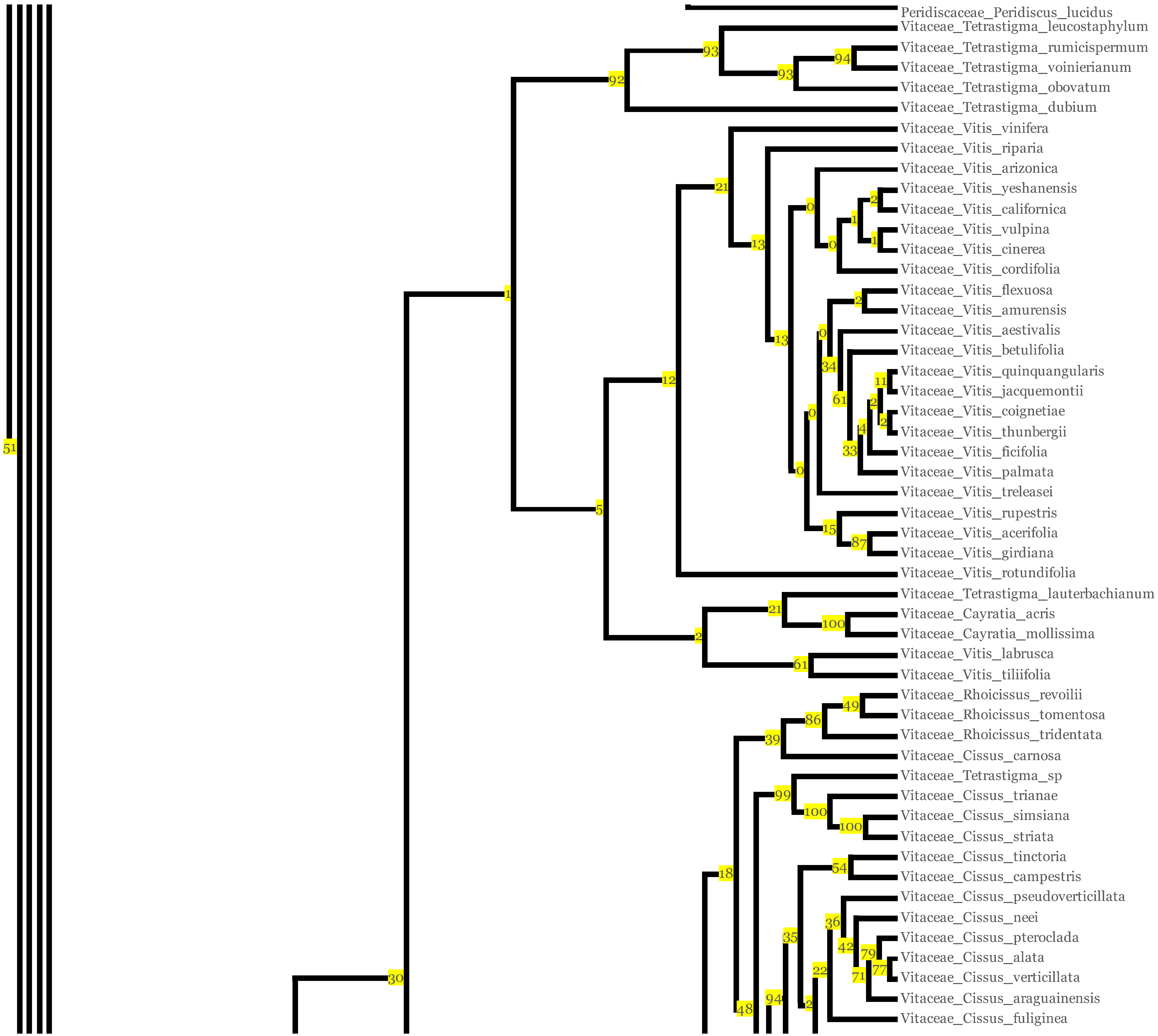
Peridiscaceae_Peridiscus_lucidus
Vitaceae_Tetrastigma_leucostaphylum
Vitaceae_Tetrastigma_rumicispermum
Vitaceae_Tetrastigma_voinierianum
Vitaceae_Tetrastigma_obovatum
Vitaceae_Tetrastigma_dubium
Vitaceae_Vitis_vinifera
Vitaceae_Vitis_riparia
Vitaceae_Vitis_arizonica
Vitaceae_Vitis_yeshanensis
Vitaceae_Vitis_californica
Vitaceae_Vitis_vulpina
Vitaceae_Vitis_cinerea
Vitaceae_Vitis_cordifolia
Vitaceae_Vitis_flexuosa
Vitaceae_Vitis_amurensis
Vitaceae_Vitis_aestivalis
Vitaceae_Vitis_betulifolia
Vitaceae_Vitis_quinquangularis
Vitaceae_Vitis_jacquemontii
Vitaceae_Vitis_coignetiae
Vitaceae_Vitis_thunbergii
Vitaceae_Vitis_ficifolia
Vitaceae_Vitis_palmata
Vitaceae_Vitis_treleasei
Vitaceae_Vitis_rupestris
Vitaceae_Vitis_acerifolia
Vitaceae_Vitis_girdiana
Vitaceae_Vitis_rotundifolia
Vitaceae_Tetrastigma_lauterbachianum
Vitaceae_Cayratia_acris
Vitaceae_Cayratia_mollissima
Vitaceae_Vitis_labrusca
Vitaceae_Vitis_tiliifolia
Vitaceae_Rhoicissus_revoilii
Vitaceae_Rhoicissus_tomentosa
Vitaceae_Rhoicissus_tridentata
Vitaceae_Cissus_carnosa
Vitaceae_Tetrastigma_sp
Vitaceae_Cissus_trianae
Vitaceae_Cissus_simsiana
Vitaceae_Cissus_striata
Vitaceae_Cissus_tinctoria
Vitaceae_Cissus_campestris
Vitaceae_Cissus_pseudoverticillata
Vitaceae_Cissus_neei
Vitaceae_Cissus_pteroclada
Vitaceae_Cissus_alata
Vitaceae_Cissus_verticillata
Vitaceae_Cissus_araguainensis
Vitaceae_Cissus_fuliginea

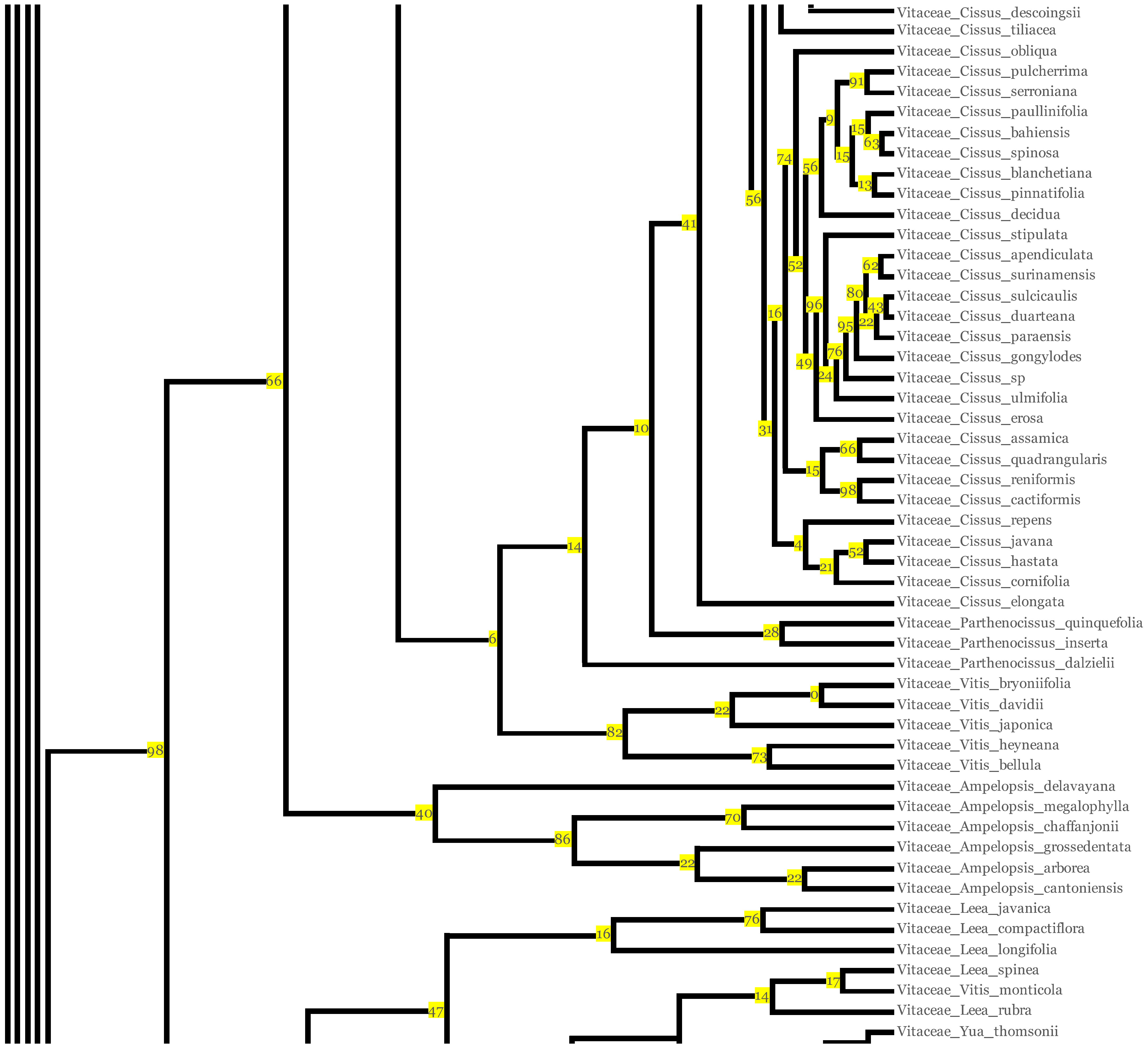
Vitaceae_Cissus_descoingsii
Vitaceae_Cissus_tiliacea
Vitaceae_Cissus_obliqua
Vitaceae_Cissus_pulcherrima
Vitaceae_Cissus_serroniana
Vitaceae_Cissus_paullinifolia
Vitaceae_Cissus_bahiensis
Vitaceae_Cissus_spinosa
Vitaceae_Cissus_blanchetiana
Vitaceae_Cissus_pinnatifolia
Vitaceae_Cissus_decidua
Vitaceae_Cissus_stipulata
Vitaceae_Cissus_apendiculata
Vitaceae_Cissus_surinamensis
Vitaceae_Cissus_sulcicaulis
Vitaceae_Cissus_duarteana
Vitaceae_Cissus_paraensis
Vitaceae_Cissus_gongylodes
Vitaceae_Cissus_sp
Vitaceae_Cissus_ulmifolia
Vitaceae_Cissus_erosa
Vitaceae_Cissus_assamica
Vitaceae_Cissus_quadrangularis
Vitaceae_Cissus_reniformis
Vitaceae_Cissus_cactiformis
Vitaceae_Cissus_repens
Vitaceae_Cissus_javana
Vitaceae_Cissus_hastata
Vitaceae_Cissus_cornifolia
Vitaceae_Cissus_elongata
Vitaceae_Parthenocissus_quinquefolia
Vitaceae_Parthenocissus_inserta
Vitaceae_Parthenocissus_dalzielii
Vitaceae_Vitis_bryoniifolia
Vitaceae_Vitis_davidii
Vitaceae_Vitis_japonica
Vitaceae_Vitis_heyneana
Vitaceae_Vitis_bellula
Vitaceae_Ampelopsis_delavayana
Vitaceae_Ampelopsis_megalophylla
Vitaceae_Ampelopsis_chaffanjonii
Vitaceae_Ampelopsis_grossedentata
Vitaceae_Ampelopsis_arborea
Vitaceae_Ampelopsis_cantoniensis
Vitaceae_Leea_javanica
Vitaceae_Leea_compactiflora
Vitaceae_Leea_longifolia
Vitaceae_Leea_spinea
Vitaceae_Vitis_monticola
Vitaceae_Leea_rubra
Vitaceae_Yua_thomsonii
91
9
15
63
15
74
56
13
56
41
62
52
80
96
43
16
95
22
76
49
24
31
10
66
15
98
4
52
21
14
6
28
0
22
82
73
40
70
86
22
22
76
16
47
17
14
66
98

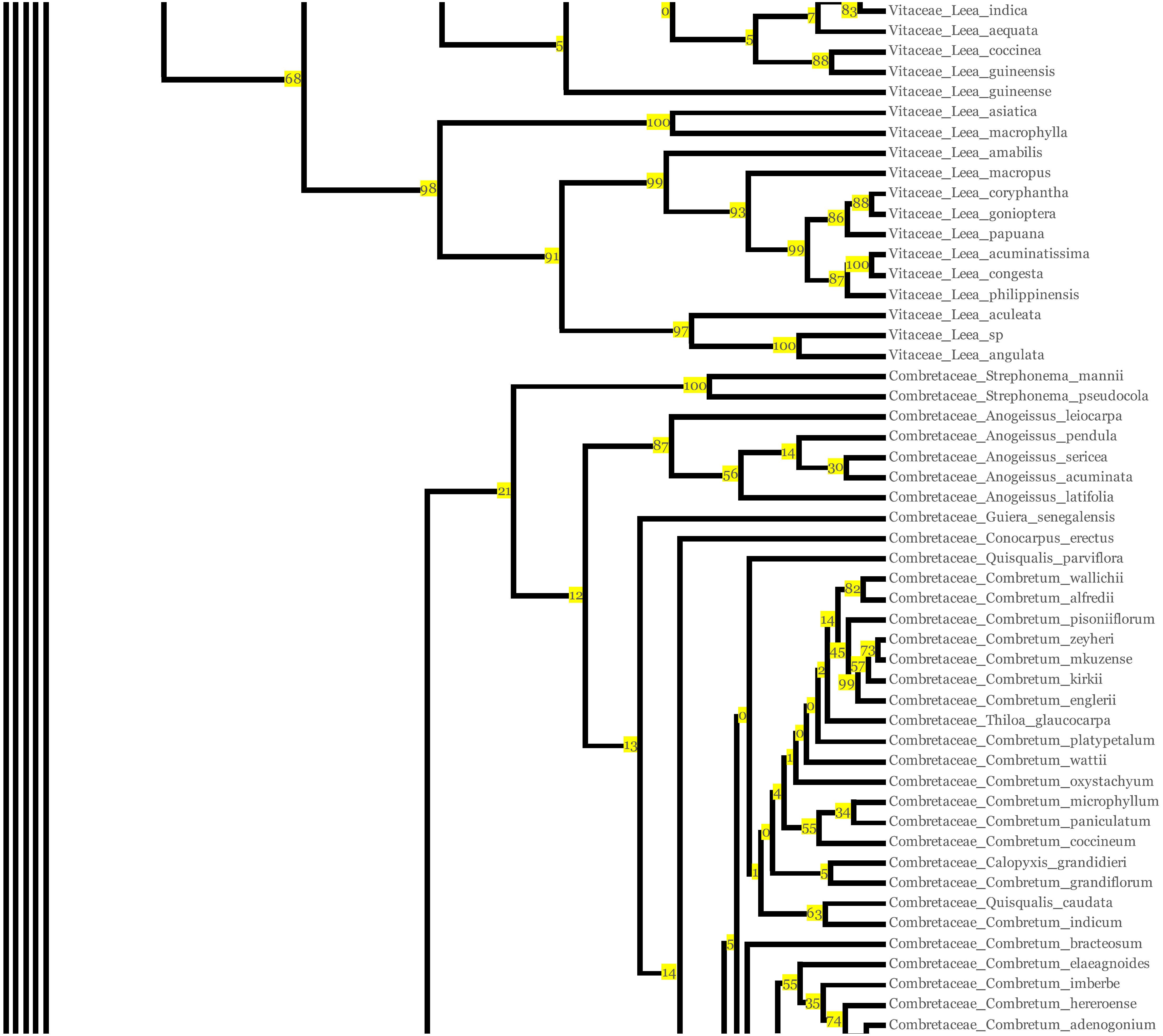
Vitaceae_Leea_indica
Vitaceae_Leea_aequata
Vitaceae_Leea_coccinea
Vitaceae_Leea_guineensis
Vitaceae_Leea_guineense
Vitaceae_Leea_asiatica
Vitaceae_Leea_macrophylla
Vitaceae_Leea_amabilis
Vitaceae_Leea_macropus
Vitaceae_Leea_coryphantha
Vitaceae_Leea_gonioptera
Vitaceae_Leea_papuana
Vitaceae_Leea_acuminatissima
Vitaceae_Leea_congesta
Vitaceae_Leea_philippinensis
Vitaceae_Leea_aculeata
Vitaceae_Leea_sp
Vitaceae_Leea_angulata
Combretaceae_Strephonema_mannii
Combretaceae_Strephonema_pseudocola
Combretaceae_Anogeissus_leiocarpa
Combretaceae_Anogeissus_pendula
Combretaceae_Anogeissus_sericea
Combretaceae_Anogeissus_acuminata
Combretaceae_Anogeissus_latifolia
Combretaceae_Guiera_senegalensis
Combretaceae_Conocarpus_erectus
Combretaceae_Quisqualis_parviflora
Combretaceae_Combretum_wallichii
Combretaceae_Combretum_alfredii
Combretaceae_Combretum_pisoniiflorum
Combretaceae_Combretum_zeyheri
Combretaceae_Combretum_mkuzense
Combretaceae_Combretum_kirkii
Combretaceae_Combretum_englerii
Combretaceae_Thiloa_glaucocarpa
Combretaceae_Combretum_platypetalum
Combretaceae_Combretum_wattii
Combretaceae_Combretum_oxystachyum
Combretaceae_Combretum_microphyllum
Combretaceae_Combretum_paniculatum
Combretaceae_Combretum_coccineum
Combretaceae_Calopyxis_grandidieri
Combretaceae_Combretum_grandiflorum
Combretaceae_Quisqualis_caudata
Combretaceae_Combretum_indicum
Combretaceae_Combretum_bracteosum
Combretaceae_Combretum_elaeagnoides
Combretaceae_Combretum_imberbe
Combretaceae_Combretum_hereroense
Combretaceae_Combretum_adenogonium

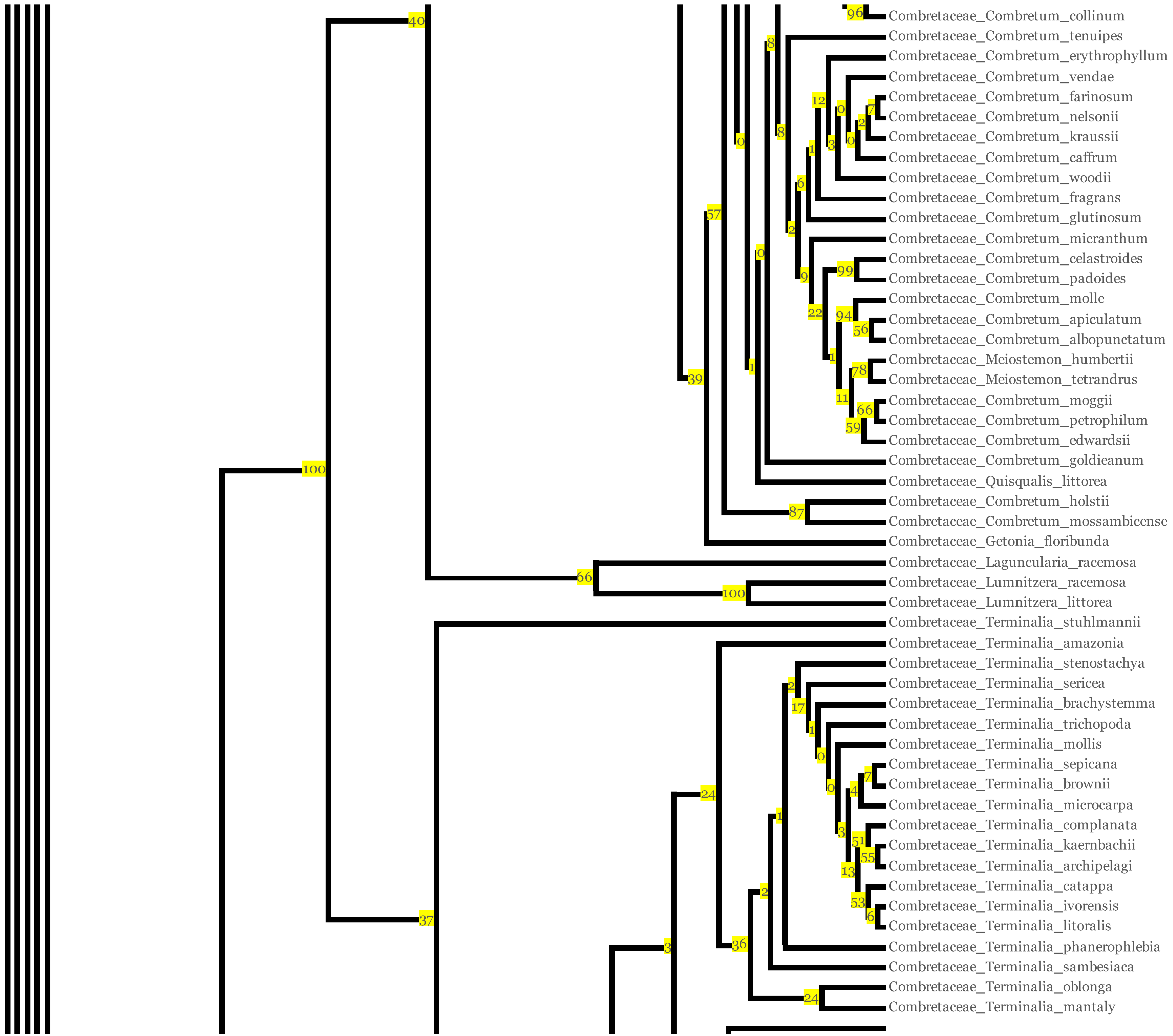
Combretaceae_Combretum_collinum
Combretaceae_Combretum_tenuipes
Combretaceae_Combretum_erythrophyllum
Combretaceae_Combretum_vendae
Combretaceae_Combretum_farinosum
Combretaceae_Combretum_nelsonii
Combretaceae_Combretum_kraussii
Combretaceae_Combretum_caffrum
Combretaceae_Combretum_woodii
Combretaceae_Combretum_fragrans
Combretaceae_Combretum_glutinosum
Combretaceae_Combretum_micranthum
Combretaceae_Combretum_celastroides
Combretaceae_Combretum_padoides
Combretaceae_Combretum_molle
Combretaceae_Combretum_apiculatum
Combretaceae_Combretum_albopunctatum
Combretaceae_Meiostemon_humbertii
Combretaceae_Meiostemon_tetrandrus
Combretaceae_Combretum_moggii
Combretaceae_Combretum_petrophilum
Combretaceae_Combretum_edwardsii
Combretaceae_Combretum_goldieanum
Combretaceae_Quisqualis_littorea
Combretaceae_Combretum_holstii
Combretaceae_Combretum_mossambicense
Combretaceae_Getonia_floribunda
Combretaceae_Laguncularia_racemosa
Combretaceae_Lumnitzera_racemosa
Combretaceae_Lumnitzera_littorea
Combretaceae_Terminalia_stuhlmannii
Combretaceae_Terminalia_amazonia
Combretaceae_Terminalia_stenostachya
Combretaceae_Terminalia_sericea
Combretaceae_Terminalia_brachystemma
Combretaceae_Terminalia_trichopoda
Combretaceae_Terminalia_mollis
Combretaceae_Terminalia_sepicana
Combretaceae_Terminalia_brownii
Combretaceae_Terminalia_microcarpa
Combretaceae_Terminalia_complanata
Combretaceae_Terminalia_kaernbachii
Combretaceae_Terminalia_archipelagi
Combretaceae_Terminalia_catappa
Combretaceae_Terminalia_ivorensis
Combretaceae_Terminalia_litoralis
Combretaceae_Terminalia_phanerophlebia
Combretaceae_Terminalia_sambesiaca
Combretaceae_Terminalia_oblonga
Combretaceae_Terminalia_mantaly
96
8
12
100
40
66
37
24
36
87
57
39
99
94
56
78
66
59
22
51
55
53
13
17

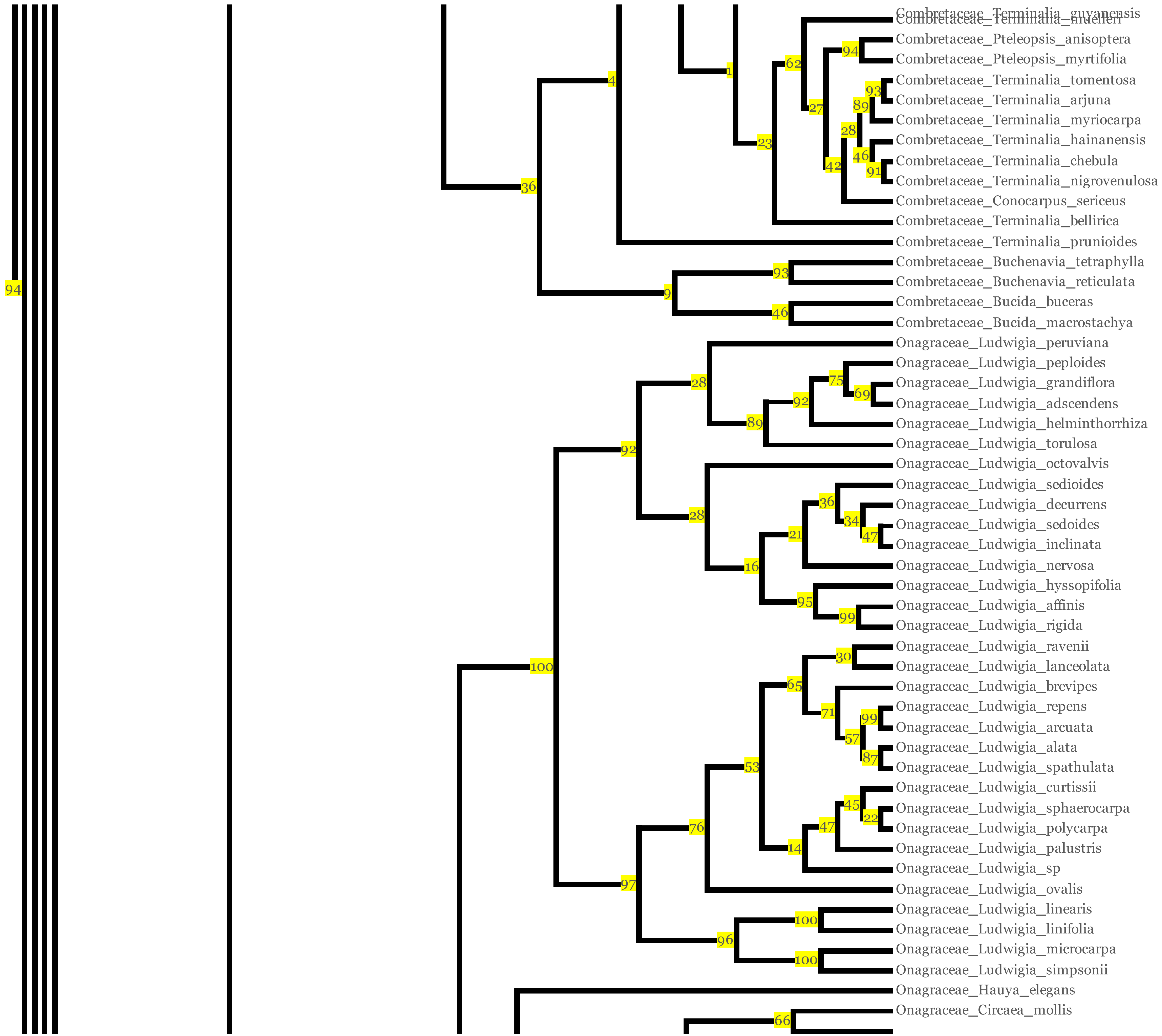
Combretaceae_Terminalia_guyanensis
Combretaceae_Terminalia_muelleri
Combretaceae_Pteleopsis_anisoptera
Combretaceae_Pteleopsis_myrtifolia
Combretaceae_Terminalia_tomentosa
Combretaceae_Terminalia_arjuna
Combretaceae_Terminalia_myriocarpa
Combretaceae_Terminalia_hainanensis
Combretaceae_Terminalia_chebula
Combretaceae_Terminalia_nigrovenulosa
Combretaceae_Conocarpus_sericeus
Combretaceae_Terminalia_bellirica
Combretaceae_Terminalia_prunioides
Combretaceae_Buchenavia_tetraphylla
Combretaceae_Buchenavia_reticulata
Combretaceae_Bucida_buceras
Combretaceae_Bucida_macrostachya
Onagraceae_Ludwigia_peruviana
Onagraceae_Ludwigia_peploides
Onagraceae_Ludwigia_grandiflora
Onagraceae_Ludwigia_adscendens
Onagraceae_Ludwigia_helminthorrhiza
Onagraceae_Ludwigia_torulosa
Onagraceae_Ludwigia_octovalvis
Onagraceae_Ludwigia_sedioides
Onagraceae_Ludwigia_decurrens
Onagraceae_Ludwigia_sedoides
Onagraceae_Ludwigia_inclinata
Onagraceae_Ludwigia_nervosa
Onagraceae_Ludwigia_hyssopifolia
Onagraceae_Ludwigia_affinis
Onagraceae_Ludwigia_rigida
Onagraceae_Ludwigia_ravenii
Onagraceae_Ludwigia_lanceolata
Onagraceae_Ludwigia_brevipes
Onagraceae_Ludwigia_repens
Onagraceae_Ludwigia_arcuata
Onagraceae_Ludwigia_alata
Onagraceae_Ludwigia_spathulata
Onagraceae_Ludwigia_curtissii
Onagraceae_Ludwigia_sphaerocarpa
Onagraceae_Ludwigia_polycarpa
Onagraceae_Ludwigia_palustris
Onagraceae_Ludwigia_sp
Onagraceae_Ludwigia_ovalis
Onagraceae_Ludwigia_linearis
Onagraceae_Ludwigia_linifolia
Onagraceae_Ludwigia_microcarpa
Onagraceae_Ludwigia_simpsonii
Onagraceae_Hauya_elegans
Onagraceae_Circaea_mollis

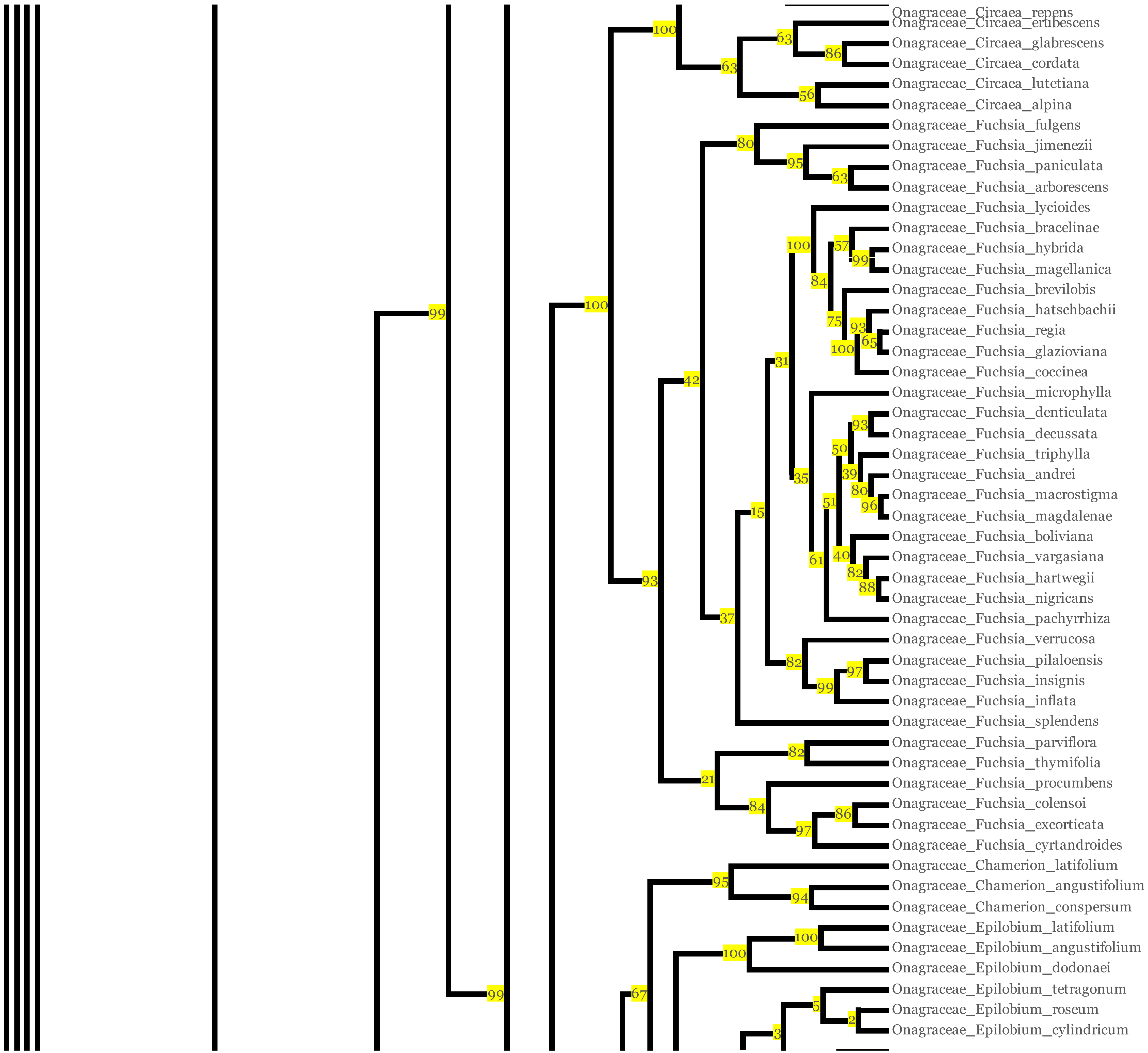
Onagraceae_Circaea_repens
Onagraceae_Circaea_erubescens
Onagraceae_Circaea_glabrescens
Onagraceae_Circaea_cordata
Onagraceae_Circaea_lutetiana
Onagraceae_Circaea_alpina
Onagraceae_Fuchsia_fulgens
Onagraceae_Fuchsia_jimenezii
Onagraceae_Fuchsia_paniculata
Onagraceae_Fuchsia_arborescens
Onagraceae_Fuchsia_lycioides
Onagraceae_Fuchsia_bracelinae
Onagraceae_Fuchsia_hybrida
Onagraceae_Fuchsia_magellanica
Onagraceae_Fuchsia_brevilobis
Onagraceae_Fuchsia_hatschbachii
Onagraceae_Fuchsia_regia
Onagraceae_Fuchsia_glazioviana
Onagraceae_Fuchsia_coccinea
Onagraceae_Fuchsia_microphylla
Onagraceae_Fuchsia_denticulata
Onagraceae_Fuchsia_decussata
Onagraceae_Fuchsia_triphylla
Onagraceae_Fuchsia_andrei
Onagraceae_Fuchsia_macrostigma
Onagraceae_Fuchsia_magdalenae
Onagraceae_Fuchsia_boliviana
Onagraceae_Fuchsia_vargasiana
Onagraceae_Fuchsia_hartwegii
Onagraceae_Fuchsia_nigricans
Onagraceae_Fuchsia_pachyrrhiza
Onagraceae_Fuchsia_verrucosa
Onagraceae_Fuchsia_pilaloensis
Onagraceae_Fuchsia_insignis
Onagraceae_Fuchsia_inflata
Onagraceae_Fuchsia_splendens
Onagraceae_Fuchsia_parviflora
Onagraceae_Fuchsia_thymifolia
Onagraceae_Fuchsia_procumbens
Onagraceae_Fuchsia_colensoi
Onagraceae_Fuchsia_excorticata
Onagraceae_Fuchsia_cyrtandroides
Onagraceae_Chamerion_latifolium
Onagraceae_Chamerion_angustifolium
Onagraceae_Chamerion_conspersum
Onagraceae_Epilobium_latifolium
Onagraceae_Epilobium_angustifolium
Onagraceae_Epilobium_dodonaei
Onagraceae_Epilobium_tetragonum
Onagraceae_Epilobium_roseum
Onagraceae_Epilobium_cylindricum

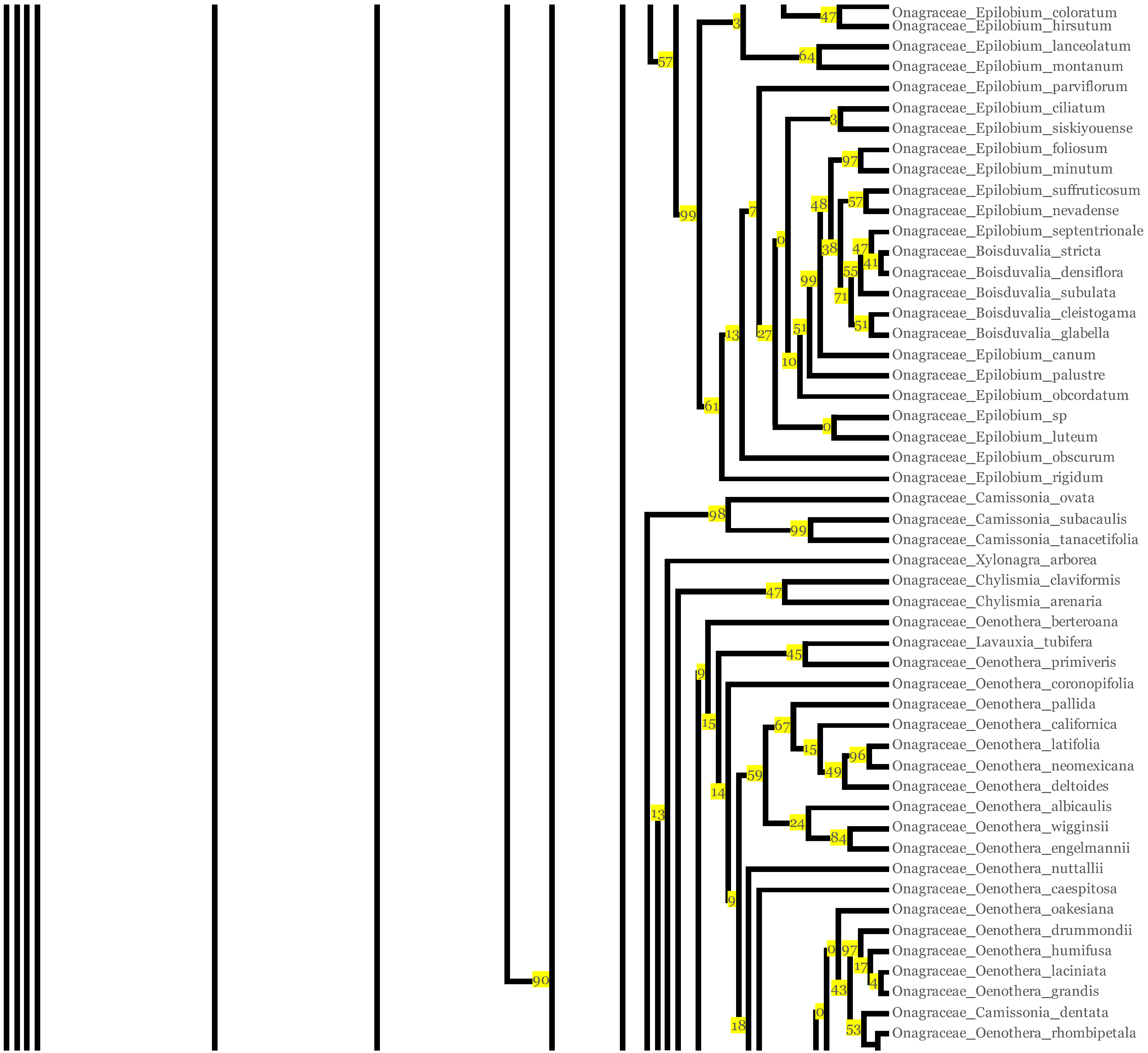
Onagraceae_Epilobium_coloratum
Onagraceae_Epilobium_hirsutum
Onagraceae_Epilobium_lanceolatum
Onagraceae_Epilobium_montanum
Onagraceae_Epilobium_parviflorum
Onagraceae_Epilobium_ciliatum
Onagraceae_Epilobium_siskiyouense
Onagraceae_Epilobium_foliosum
Onagraceae_Epilobium_minutum
Onagraceae_Epilobium_suffruticosum
Onagraceae_Epilobium_nevadense
Onagraceae_Epilobium_septentrionale
Onagraceae_Boisduvalia_stricta
Onagraceae_Boisduvalia_densiflora
Onagraceae_Boisduvalia_subulata
Onagraceae_Boisduvalia_cleistogama
Onagraceae_Boisduvalia_glabella
Onagraceae_Epilobium_canum
Onagraceae_Epilobium_palustre
Onagraceae_Epilobium_obcordatum
Onagraceae_Epilobium_sp
Onagraceae_Epilobium_luteum
Onagraceae_Epilobium_obscurum
Onagraceae_Epilobium_rigidum
Onagraceae_Camissonia_ovata
Onagraceae_Camissonia_subacaulis
Onagraceae_Camissonia_tanacetifolia
Onagraceae_Xylonagra_arborea
Onagraceae_Chylismia_claviformis
Onagraceae_Chylismia_arenaria
Onagraceae_Oenothera_berteroana
Onagraceae_Lavauxia_tubifera
Onagraceae_Oenothera_primiveris
Onagraceae_Oenothera_coronopifolia
Onagraceae_Oenothera_pallida
Onagraceae_Oenothera_californica
Onagraceae_Oenothera_latifolia
Onagraceae_Oenothera_neomexicana
Onagraceae_Oenothera_deltoides
Onagraceae_Oenothera_albicaulis
Onagraceae_Oenothera_wigginsii
Onagraceae_Oenothera_engelmannii
Onagraceae_Oenothera_nuttallii
Onagraceae_Oenothera_caespitosa
Onagraceae_Oenothera_oakesiana
Onagraceae_Oenothera_drummondii
Onagraceae_Oenothera_humifusa
Onagraceae_Oenothera_laciniata
Onagraceae_Oenothera_grandis
Onagraceae_Camissonia_dentata
Onagraceae_Oenothera_rhombipetala

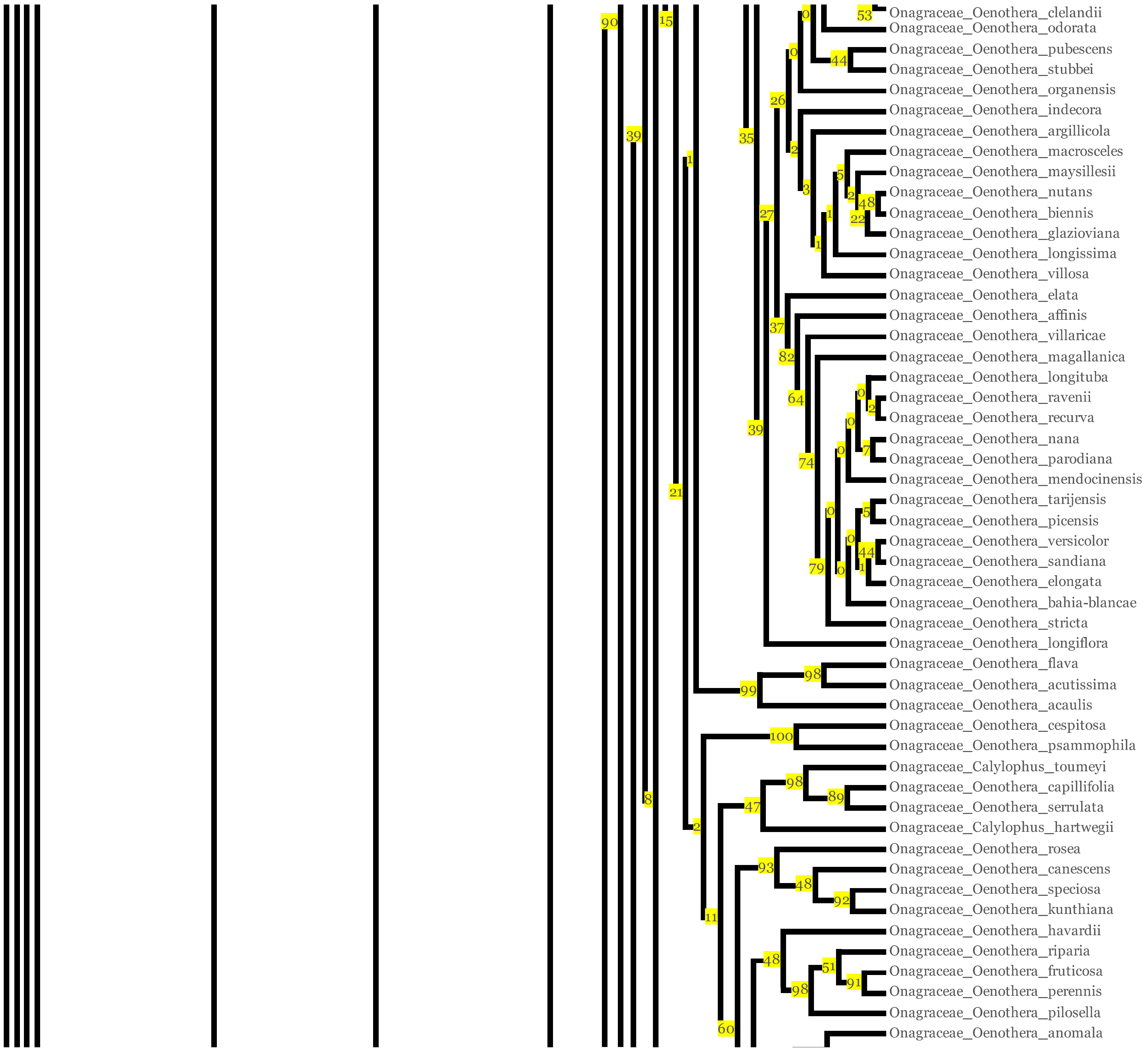
Onagraceae_Oenothera_clelandii
Onagraceae_Oenothera_odorata
Onagraceae_Oenothera_pubescens
Onagraceae_Oenothera_stubbei
Onagraceae_Oenothera_organensis
Onagraceae_Oenothera_indecora
Onagraceae_Oenothera_argillicola
Onagraceae_Oenothera_macrosceles
Onagraceae_Oenothera_maysillesii
Onagraceae_Oenothera_nutans
Onagraceae_Oenothera_biennis
Onagraceae_Oenothera_glazioviana
Onagraceae_Oenothera_longissima
Onagraceae_Oenothera_villosa
Onagraceae_Oenothera_elata
Onagraceae_Oenothera_affinis
Onagraceae_Oenothera_villaricae
Onagraceae_Oenothera_magallanica
Onagraceae_Oenothera_longituba
Onagraceae_Oenothera_ravenii
Onagraceae_Oenothera_recurva
Onagraceae_Oenothera_nana
Onagraceae_Oenothera_parodiana
Onagraceae_Oenothera_mendocinensis
Onagraceae_Oenothera_tarijensis
Onagraceae_Oenothera_picensis
Onagraceae_Oenothera_versicolor
Onagraceae_Oenothera_sandiana
Onagraceae_Oenothera_elongata
Onagraceae_Oenothera_bahia-blancae
Onagraceae_Oenothera_stricta
Onagraceae_Oenothera_longiflora
Onagraceae_Oenothera_flava
Onagraceae_Oenothera_acutissima
Onagraceae_Oenothera_acaulis
Onagraceae_Oenothera_cespitosa
Onagraceae_Oenothera_psammophila
Onagraceae_Calylophus_toumeyi
Onagraceae_Oenothera_capillifolia
Onagraceae_Oenothera_serrulata
Onagraceae_Calylophus_hartwegii
Onagraceae_Oenothera_rosea
Onagraceae_Oenothera_canescens
Onagraceae_Oenothera_speciosa
Onagraceae_Oenothera_kunthiana
Onagraceae_Oenothera_havardii
Onagraceae_Oenothera_riparia
Onagraceae_Oenothera_fruticosa
Onagraceae_Oenothera_perennis
Onagraceae_Oenothera_pilosella
Onagraceae_Oenothera_anomala

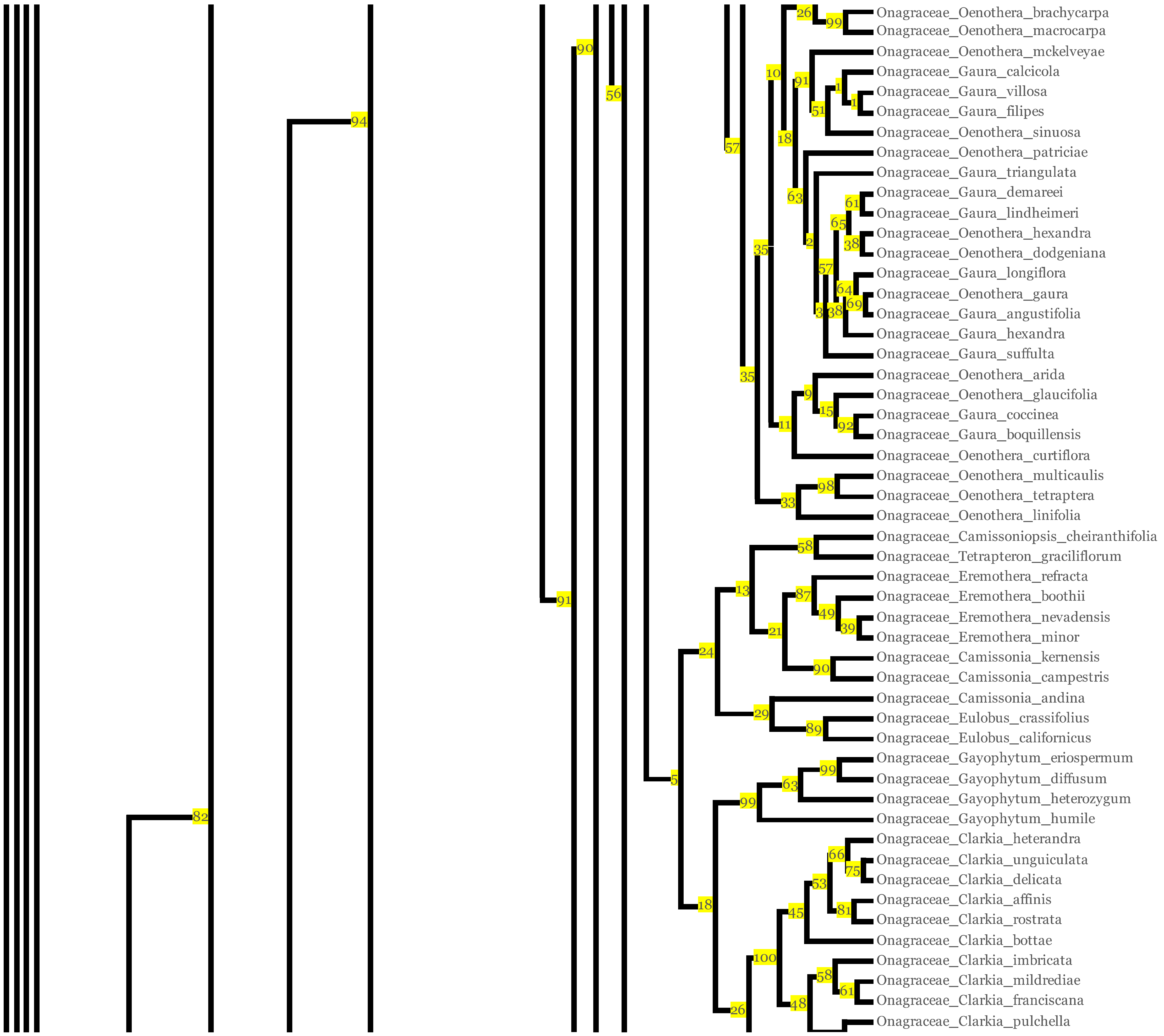
Onagraceae_Oenothera_brachycarpa
Onagraceae_Oenothera_macrocarpa
Onagraceae_Oenothera_mckelveyae
Onagraceae_Gaura_calcicola
Onagraceae_Gaura_villosa
Onagraceae_Gaura_filipes
Onagraceae_Oenothera_sinuosa
Onagraceae_Oenothera_patriciae
Onagraceae_Gaura_triangulata
Onagraceae_Gaura_demareei
Onagraceae_Gaura_lindheimeri
Onagraceae_Oenothera_hexandra
Onagraceae_Oenothera_dodgeniana
Onagraceae_Gaura_longiflora
Onagraceae_Oenothera_gaura
Onagraceae_Gaura_angustifolia
Onagraceae_Gaura_hexandra
Onagraceae_Gaura_suffulta
Onagraceae_Oenothera_arida
Onagraceae_Oenothera_glaucifolia
Onagraceae_Gaura_coccinea
Onagraceae_Gaura_boquillensis
Onagraceae_Oenothera_curtiflora
Onagraceae_Oenothera_multicaulis
Onagraceae_Oenothera_tetraptera
Onagraceae_Oenothera_linifolia
Onagraceae_Camissoniopsis_cheiranthifolia
Onagraceae_Tetrapteron_graciliflorum
Onagraceae_Eremothera_refracta
Onagraceae_Eremothera_boothii
Onagraceae_Eremothera_nevadensis
Onagraceae_Eremothera_minor
Onagraceae_Camissonia_kernensis
Onagraceae_Camissonia_campestris
Onagraceae_Camissonia_andina
Onagraceae_Eulobus_crassifolius
Onagraceae_Eulobus_californicus
Onagraceae_Gayophytum_eriospermum
Onagraceae_Gayophytum_diffusum
Onagraceae_Gayophytum_heterozygum
Onagraceae_Gayophytum_humile
Onagraceae_Clarkia_heterandra
Onagraceae_Clarkia_unguiculata
Onagraceae_Clarkia_delicata
Onagraceae_Clarkia_affinis
Onagraceae_Clarkia_rostrata
Onagraceae_Clarkia_bottae
Onagraceae_Clarkia_imbricata
Onagraceae_Clarkia_mildrediae
Onagraceae_Clarkia_franciscana
Onagraceae_Clarkia_pulchella

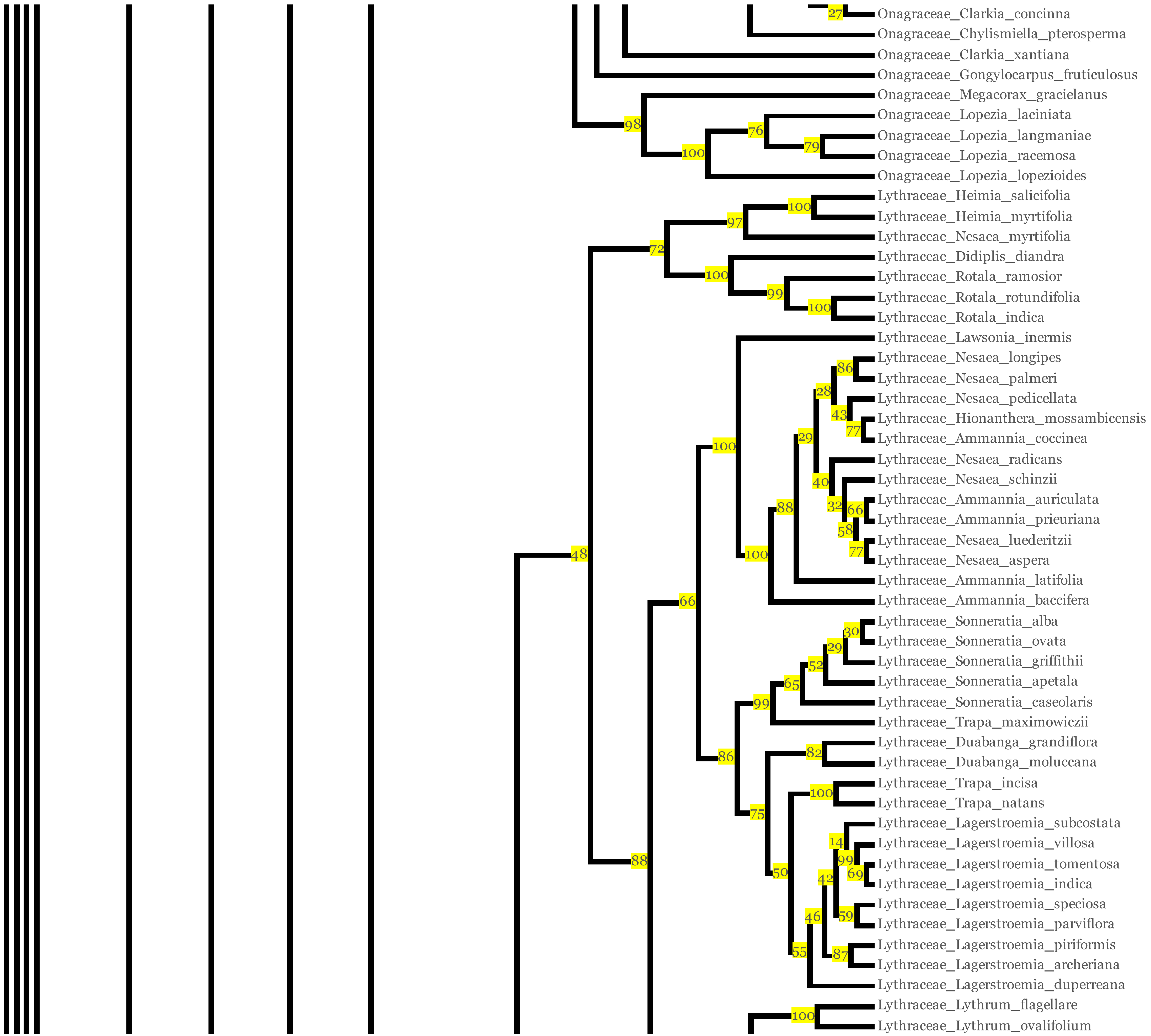

27
Onagraceae_Clarkia_concinna
Onagraceae_Chylismiella_pterosperma
Onagraceae_Clarkia_xantiana
Onagraceae_Gongylocarpus_fruticulosus
Onagraceae_Megacorax_gracielanus
98
76
Onagraceae_Lopezia_laciniata
79
Onagraceae_Lopezia_langmaniae
100
Onagraceae_Lopezia_racemosa
Onagraceae_Lopezia_lopezioides
100
Lythraceae_Heimia_salicifolia
97
Lythraceae_Heimia_myrtifolia
Lythraceae_Nesaea_myrtifolia
72
Lythraceae_Didiplis_diandra
100
Lythraceae_Rotala_ramosior
99
100
Lythraceae_Rotala_rotundifolia
Lythraceae_Rotala_indica
Lythraceae_Lawsonia_inermis
86
Lythraceae_Nesaea_longipes
28
Lythraceae_Nesaea_palmeri
Lythraceae_Nesaea_pedicellata
43
Lythraceae_Hionanthera_mossambicensis
77
29
Lythraceae_Ammannia_coccinea
100
Lythraceae_Nesaea_radicans
40
Lythraceae_Nesaea_schinzii
32
66
Lythraceae_Ammannia_auriculata
88
Lythraceae_Ammannia_prieuriana
58
Lythraceae_Nesaea_luederitzii
77
48
100
Lythraceae_Nesaea_aspera
Lythraceae_Ammannia_latifolia
Lythraceae_Ammannia_baccifera
66
Lythraceae_Sonneratia_alba
30
Lythraceae_Sonneratia_ovata
29
Lythraceae_Sonneratia_griffithii
52
Lythraceae_Sonneratia_apetala
65
Lythraceae_Sonneratia_caseolaris
99
Lythraceae_Trapa_maximowiczii
Lythraceae_Duabanga_grandiflora
82
86
Lythraceae_Duabanga_moluccana
Lythraceae_Trapa_incisa
100
Lythraceae_Trapa_natans
75
Lythraceae_Lagerstroemia_subcostata
14
Lythraceae_Lagerstroemia_villosa
99
Lythraceae_Lagerstroemia_tomentosa
69
50
Lythraceae_Lagerstroemia_indica
42
88
Lythraceae_Lagerstroemia_speciosa
59
Lythraceae_Lagerstroemia_parviflora
46
Lythraceae_Lagerstroemia_piriformis
55
87
Lythraceae_Lagerstroemia_archeriana
Lythraceae_Lagerstroemia_duperreana
Lythraceae_Lythrum_flagellare
100
Lythraceae_Lythrum_ovalifolium

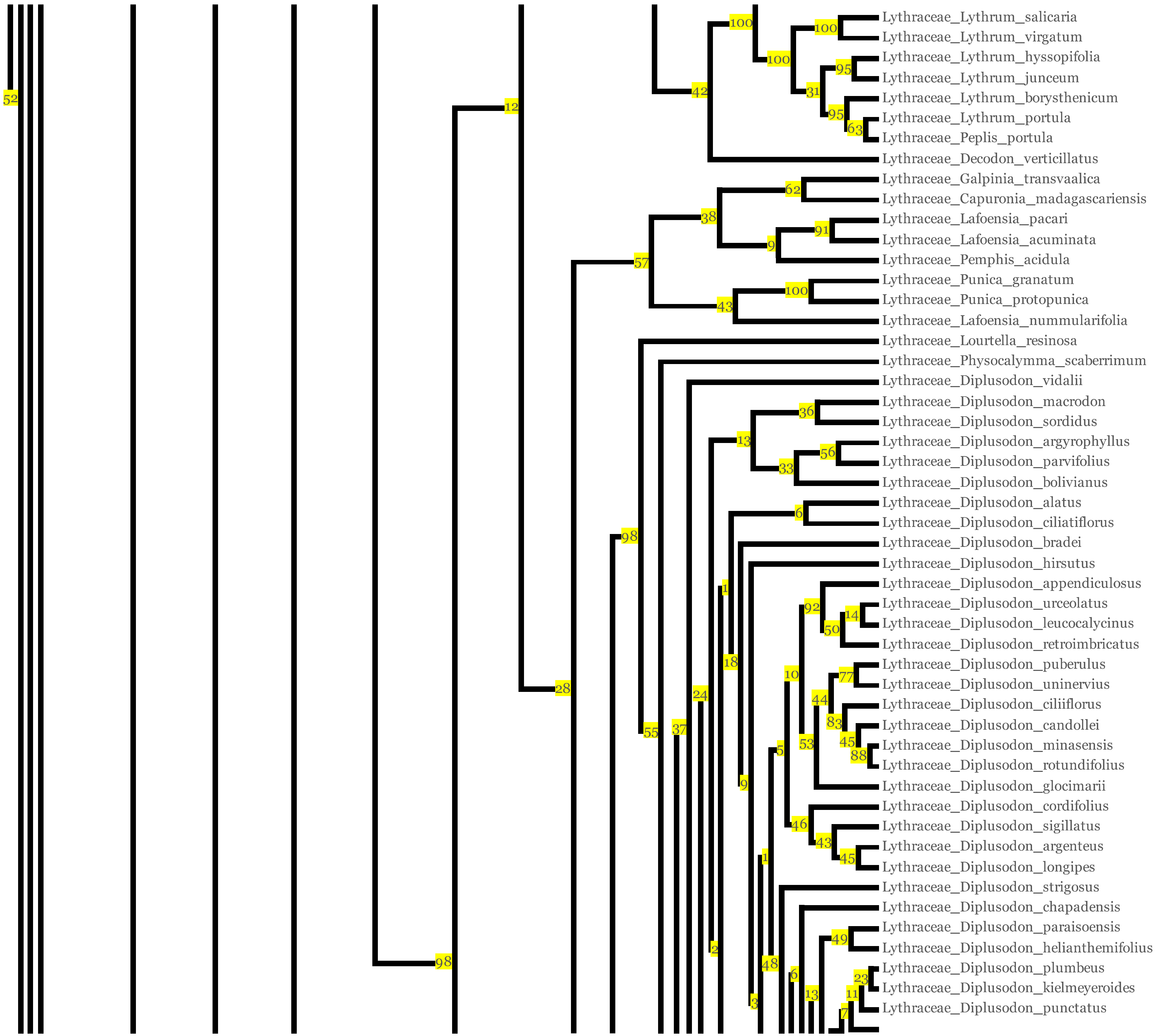
Lythraceae_Lythrum_salicaria
Lythraceae_Lythrum_virgatum
Lythraceae_Lythrum_hyssopifolia
Lythraceae_Lythrum_junceum
Lythraceae_Lythrum_borysthenicum
Lythraceae_Lythrum_portula
Lythraceae_Peplis_portula
Lythraceae_Decodon_verticillatus
Lythraceae_Galpinia_transvaalica
Lythraceae_Capuronia_madagascariensis
Lythraceae_Lafoensia_pacari
Lythraceae_Lafoensia_acuminata
Lythraceae_Pemphis_acidula
Lythraceae_Punica_granatum
Lythraceae_Punica_protopunica
Lythraceae_Lafoensia_nummularifolia
Lythraceae_Lourtella_resinosa
Lythraceae_Physocalymma_scaberrimum
Lythraceae_Diplusodon_vidalii
Lythraceae_Diplusodon_macrodon
Lythraceae_Diplusodon_sordidus
Lythraceae_Diplusodon_argyrophyllus
Lythraceae_Diplusodon_parvifolius
Lythraceae_Diplusodon_bolivianus
Lythraceae_Diplusodon_alatus
Lythraceae_Diplusodon_ciliatiflorus
Lythraceae_Diplusodon_bradei
Lythraceae_Diplusodon_hirsutus
Lythraceae_Diplusodon_appendiculosus
Lythraceae_Diplusodon_urceolatus
Lythraceae_Diplusodon_leucocalycinus
Lythraceae_Diplusodon_retroimbricatus
Lythraceae_Diplusodon_puberulus
Lythraceae_Diplusodon_uninervius
Lythraceae_Diplusodon_ciliiflorus
Lythraceae_Diplusodon_candollei
Lythraceae_Diplusodon_minasensis
Lythraceae_Diplusodon_rotundifolius
Lythraceae_Diplusodon_glocimarii
Lythraceae_Diplusodon_cordifolius
Lythraceae_Diplusodon_sigillatus
Lythraceae_Diplusodon_argenteus
Lythraceae_Diplusodon_longipes
Lythraceae_Diplusodon_strigosus
Lythraceae_Diplusodon_chapadensis
Lythraceae_Diplusodon_paraisoensis
Lythraceae_Diplusodon_helianthemifolius
Lythraceae_Diplusodon_plumbeus
Lythraceae_Diplusodon_kielmeyeroides
Lythraceae_Diplusodon_punctatus
52
12
98
28
98
55
37
24
100
100
100
95
31
95
63
42
62
38
91
9
57
100
43
36
13
56
33
6
1
18
92
14
50
77
10
44
83
45
88
53
5
9
46
43
45
1
2
48
49
6
13
23
11
7
3

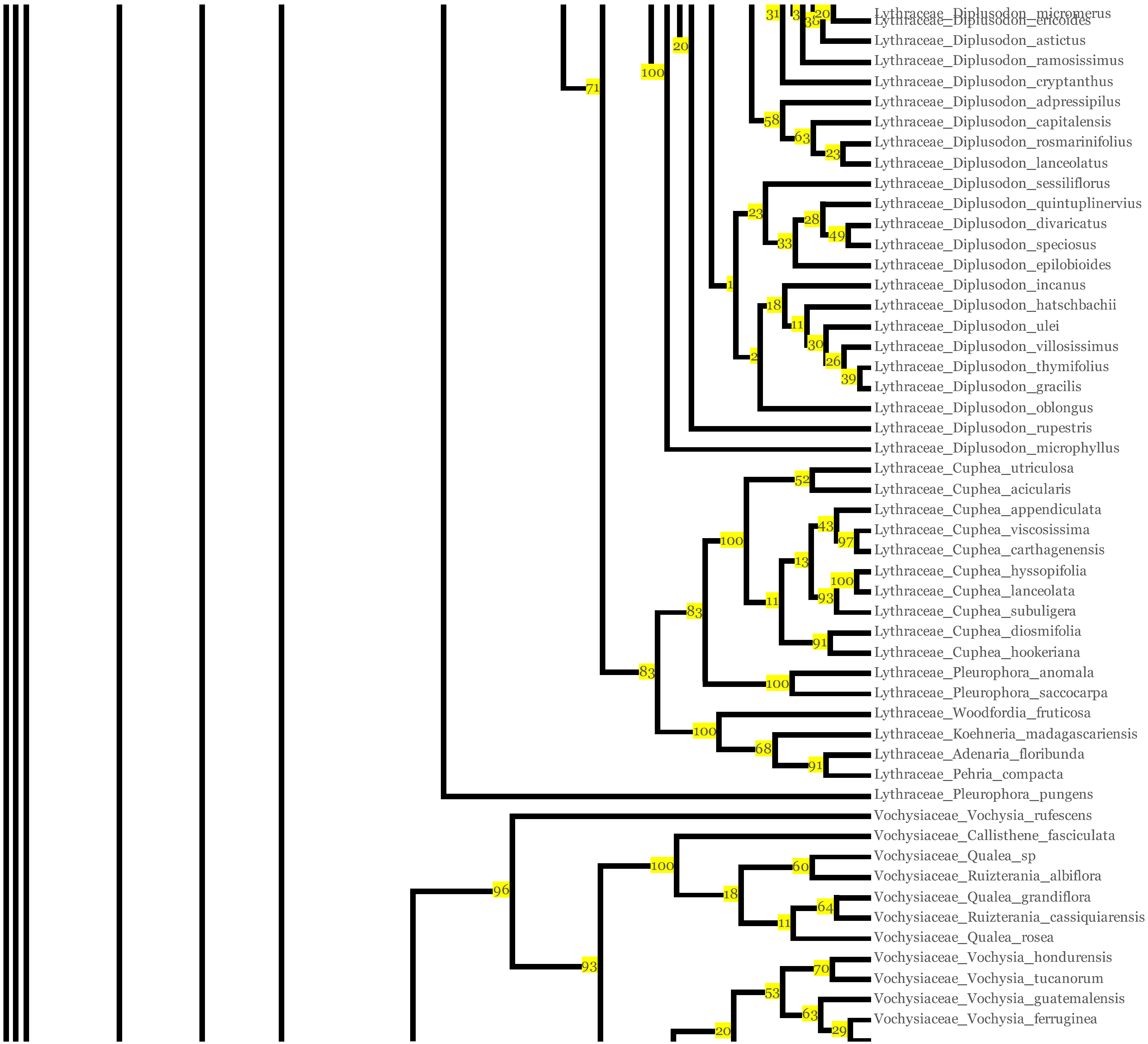

Lythraceae_Diplusodon_micromerus
Lythraceae_Diplusodon_ericoides
Lythraceae_Diplusodon_astictus
Lythraceae_Diplusodon_ramosissimus
Lythraceae_Diplusodon_cryptanthus
Lythraceae_Diplusodon_adpressipilus
Lythraceae_Diplusodon_capitalensis
Lythraceae_Diplusodon_rosmarinifolius
Lythraceae_Diplusodon_lanceolatus
Lythraceae_Diplusodon_sessiliflorus
Lythraceae_Diplusodon_quintuplinervius
Lythraceae_Diplusodon_divaricatus
Lythraceae_Diplusodon_speciosus
Lythraceae_Diplusodon_epilobioides
Lythraceae_Diplusodon_incanus
Lythraceae_Diplusodon_hatschbachii
Lythraceae_Diplusodon_ulei
Lythraceae_Diplusodon_villosissimus
Lythraceae_Diplusodon_thymifolius
Lythraceae_Diplusodon_gracilis
Lythraceae_Diplusodon_oblongus
Lythraceae_Diplusodon_rupestris
Lythraceae_Diplusodon_microphyllus
Lythraceae_Cuphea_utriculosa
Lythraceae_Cuphea_acicularis
Lythraceae_Cuphea_appendiculata
Lythraceae_Cuphea_viscosissima
Lythraceae_Cuphea_carthagenensis
Lythraceae_Cuphea_hyssopifolia
Lythraceae_Cuphea_lanceolata
Lythraceae_Cuphea_subuligera
Lythraceae_Cuphea_diosmifolia
Lythraceae_Cuphea_hookeriana
Lythraceae_Pleurophora_anomala
Lythraceae_Pleurophora_saccocarpa
Lythraceae_Woodfordia_fruticosa
Lythraceae_Koehneria_madagascariensis
Lythraceae_Adenaria_floribunda
Lythraceae_Pehria_compacta
Lythraceae_Pleurophora_pungens
Vochysiaceae_Vochysia_rufescens
Vochysiaceae_Callisthene_fasciculata
Vochysiaceae_Qualea_sp
Vochysiaceae_Ruizterania_albiflora
Vochysiaceae_Qualea_grandiflora
Vochysiaceae_Ruizterania_cassiquiarensis
Vochysiaceae_Qualea_rosea
Vochysiaceae_Vochysia_hondurensis
Vochysiaceae_Vochysia_tucanorum
Vochysiaceae_Vochysia_guatemalensis
Vochysiaceae_Vochysia_ferruginea

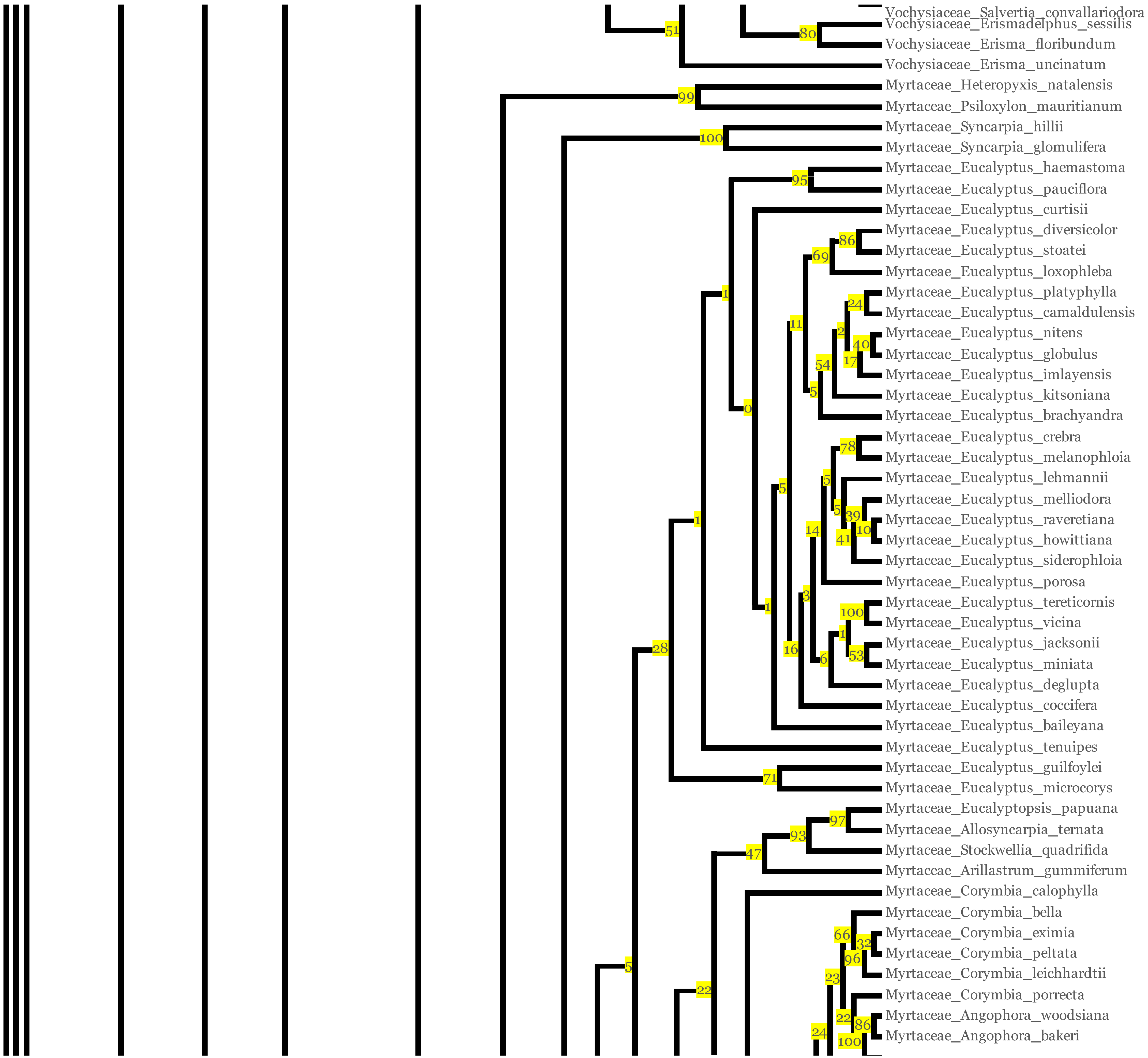

Vochysiaceae_Salvertia_convallariodora
Vochysiaceae_Erismadelphus_sessilis
Vochysiaceae_Erisma_floribundum
Vochysiaceae_Erisma_uncinatum
Myrtaceae_Heteropyxis_natalensis
Myrtaceae_Psiloxylon_mauritianum
Myrtaceae_Syncarpia_hillii
Myrtaceae_Syncarpia_glomulifera
Myrtaceae_Eucalyptus_haemastoma
Myrtaceae_Eucalyptus_pauciflora
Myrtaceae_Eucalyptus_curtisii
Myrtaceae_Eucalyptus_diversicolor
Myrtaceae_Eucalyptus_stoatei
Myrtaceae_Eucalyptus_loxophleba
Myrtaceae_Eucalyptus_platyphylla
Myrtaceae_Eucalyptus_camaldulensis
Myrtaceae_Eucalyptus_nitens
Myrtaceae_Eucalyptus_globulus
Myrtaceae_Eucalyptus_imlayensis
Myrtaceae_Eucalyptus_kitsoniana
Myrtaceae_Eucalyptus_brachyandra
Myrtaceae_Eucalyptus_crebra
Myrtaceae_Eucalyptus_melanophloia
Myrtaceae_Eucalyptus_lehmannii
Myrtaceae_Eucalyptus_melliodora
Myrtaceae_Eucalyptus_raveretiana
Myrtaceae_Eucalyptus_howittiana
Myrtaceae_Eucalyptus_siderophloia
Myrtaceae_Eucalyptus_porosa
Myrtaceae_Eucalyptus_tereticornis
Myrtaceae_Eucalyptus_vicina
Myrtaceae_Eucalyptus_jacksonii
Myrtaceae_Eucalyptus_miniata
Myrtaceae_Eucalyptus_deglupta
Myrtaceae_Eucalyptus_coccifera
Myrtaceae_Eucalyptus_baileyana
Myrtaceae_Eucalyptus_tenuipes
Myrtaceae_Eucalyptus_guilfoylei
Myrtaceae_Eucalyptus_microcorys
Myrtaceae_Eucalyptopsis_papuana
Myrtaceae_Allosyncarpia_ternata
Myrtaceae_Stockwellia_quadrifida
Myrtaceae_Arillastrum_gummiferum
Myrtaceae_Corymbia_calophylla
Myrtaceae_Corymbia_bella
Myrtaceae_Corymbia_eximia
Myrtaceae_Corymbia_peltata
Myrtaceae_Corymbia_leichhardtii
Myrtaceae_Corymbia_porrecta
Myrtaceae_Angophora_woodsiana
Myrtaceae_Angophora_bakeri

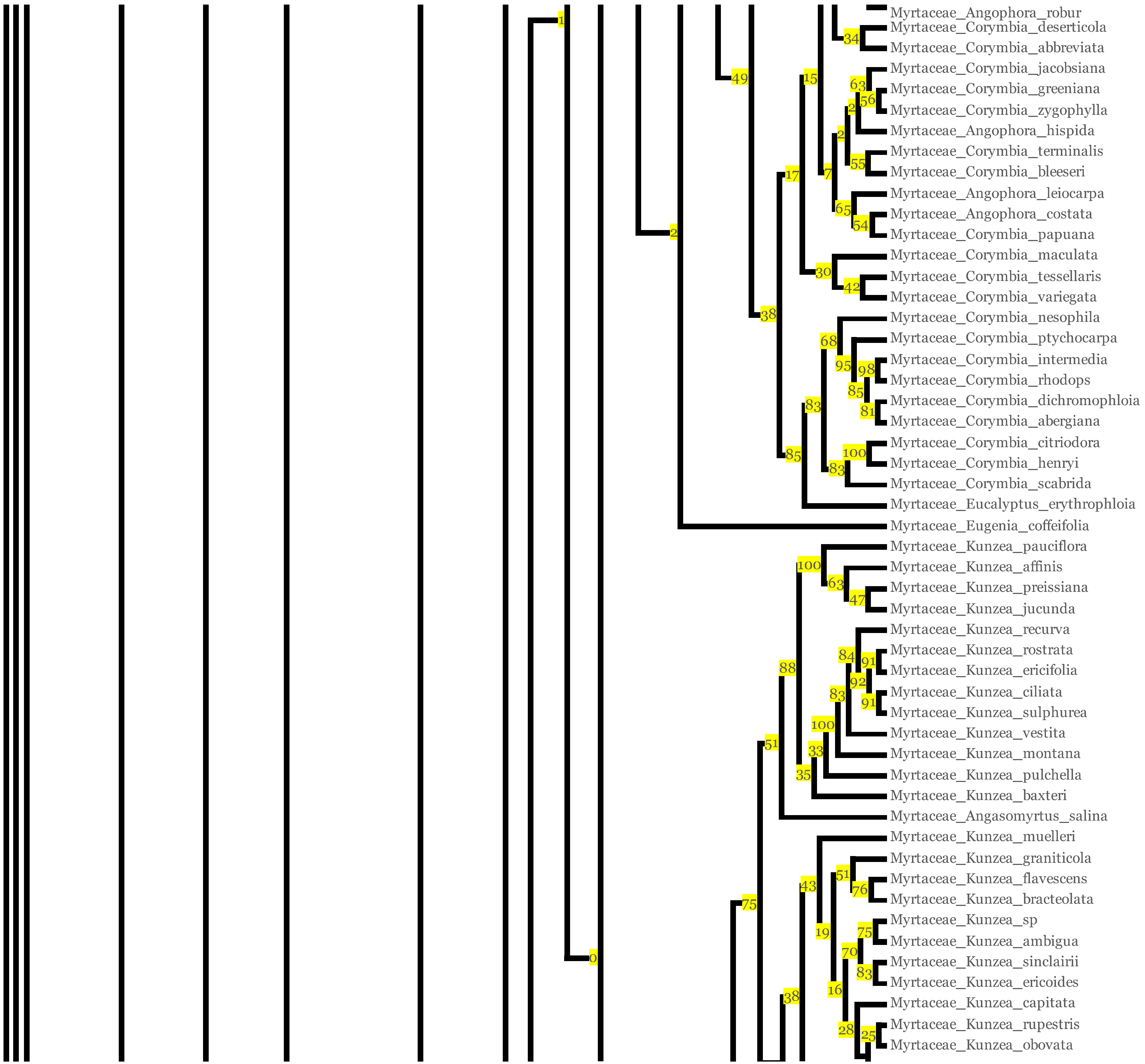
Myrtaceae_Angophora_robur
Myrtaceae_Corymbia_deserticola
Myrtaceae_Corymbia_abbreviata
Myrtaceae_Corymbia_jacobsiana
Myrtaceae_Corymbia_greeniana
Myrtaceae_Corymbia_zygophylla
Myrtaceae_Angophora_hispida
Myrtaceae_Corymbia_terminalis
Myrtaceae_Corymbia_bleeseri
Myrtaceae_Angophora_leiocarpa
Myrtaceae_Angophora_costata
Myrtaceae_Corymbia_papuana
Myrtaceae_Corymbia_maculata
Myrtaceae_Corymbia_tessellaris
Myrtaceae_Corymbia_variegata
Myrtaceae_Corymbia_nesophila
Myrtaceae_Corymbia_ptychocarpa
Myrtaceae_Corymbia_intermedia
Myrtaceae_Corymbia_rhodops
Myrtaceae_Corymbia_dichromophloia
Myrtaceae_Corymbia_abergiana
Myrtaceae_Corymbia_citriodora
Myrtaceae_Corymbia_henryi
Myrtaceae_Corymbia_scabrida
Myrtaceae_Eucalyptus_erythrophloia
Myrtaceae_Eugenia_coffeifolia
Myrtaceae_Kunzea_pauciflora
Myrtaceae_Kunzea_affinis
Myrtaceae_Kunzea_preissiana
Myrtaceae_Kunzea_jucunda
Myrtaceae_Kunzea_recurva
Myrtaceae_Kunzea_rostrata
Myrtaceae_Kunzea_ericifolia
Myrtaceae_Kunzea_ciliata
Myrtaceae_Kunzea_sulphurea
Myrtaceae_Kunzea_vestita
Myrtaceae_Kunzea_montana
Myrtaceae_Kunzea_pulchella
Myrtaceae_Kunzea_baxteri
Myrtaceae_Angasomyrtus_salina
Myrtaceae_Kunzea_muelleri
Myrtaceae_Kunzea_graniticola
Myrtaceae_Kunzea_flavescens
Myrtaceae_Kunzea_bracteolata
Myrtaceae_Kunzea_sp
Myrtaceae_Kunzea_ambigua
Myrtaceae_Kunzea_sinclairii
Myrtaceae_Kunzea_ericoides
Myrtaceae_Kunzea_capitata
Myrtaceae_Kunzea_rupestris
Myrtaceae_Kunzea_obovata

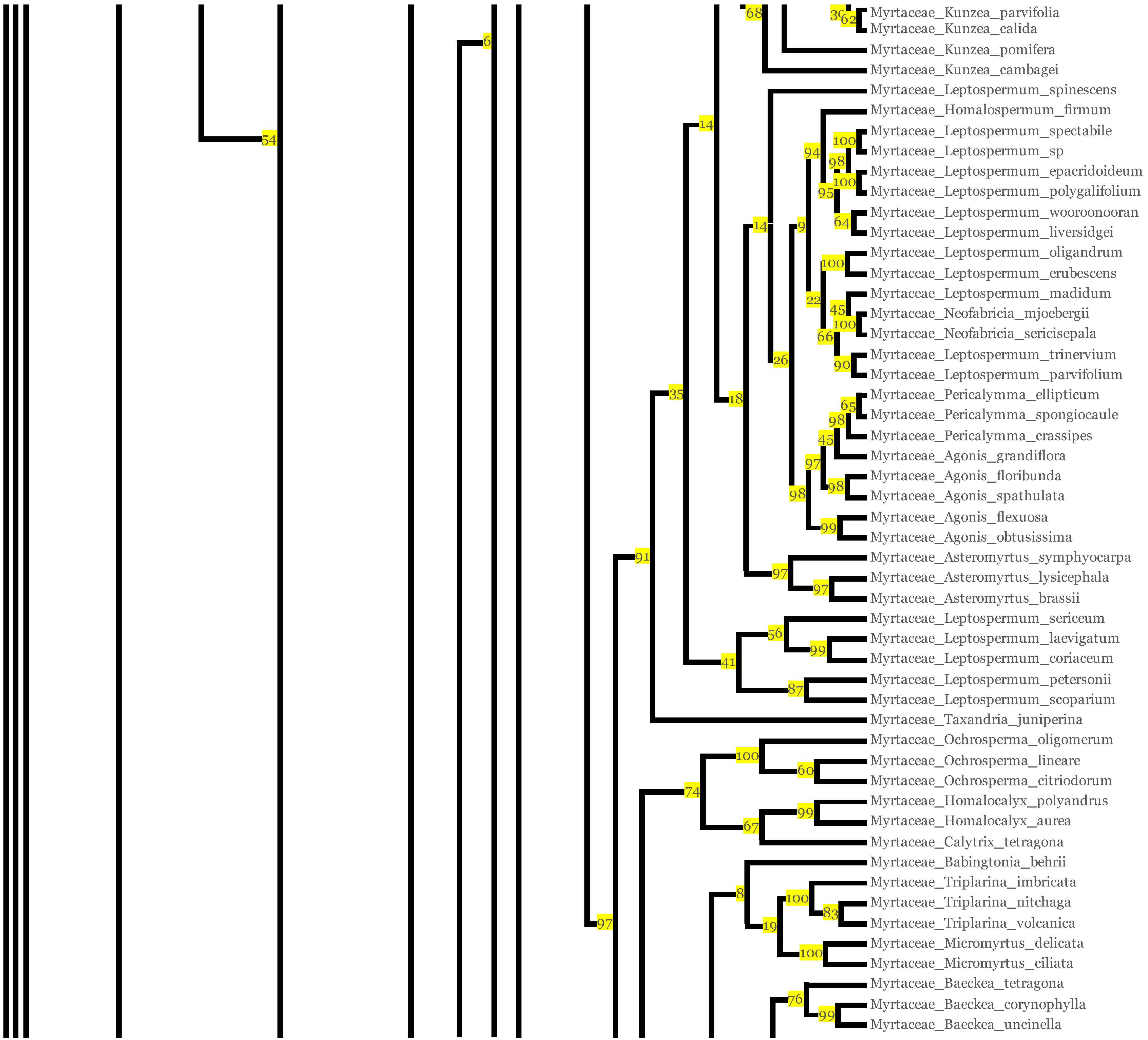
Myrtaceae_Kunzea_parvifolia
Myrtaceae_Kunzea_calida
Myrtaceae_Kunzea_pomifera
Myrtaceae_Kunzea_cambagei
Myrtaceae_Leptospermum_spinescens
Myrtaceae_Homalospermum_firmum
Myrtaceae_Leptospermum_spectabile
Myrtaceae_Leptospermum_sp
Myrtaceae_Leptospermum_epacridoideum
Myrtaceae_Leptospermum_polygalifolium
Myrtaceae_Leptospermum_wooroonooran
Myrtaceae_Leptospermum_liversidgei
Myrtaceae_Leptospermum_oligandrum
Myrtaceae_Leptospermum_erubescens
Myrtaceae_Leptospermum_madidum
Myrtaceae_Neofabricia_mjoebergii
Myrtaceae_Neofabricia_sericisepala
Myrtaceae_Leptospermum_trinervium
Myrtaceae_Leptospermum_parvifolium
Myrtaceae_Pericalymma_ellipticum
Myrtaceae_Pericalymma_spongiocaule
Myrtaceae_Pericalymma_crassipes
Myrtaceae_Agonis_grandiflora
Myrtaceae_Agonis_floribunda
Myrtaceae_Agonis_spathulata
Myrtaceae_Agonis_flexuosa
Myrtaceae_Agonis_obtusissima
Myrtaceae_Asteromyrtus_symphyocarpa
Myrtaceae_Asteromyrtus_lysicephala
Myrtaceae_Asteromyrtus_brassii
Myrtaceae_Leptospermum_sericeum
Myrtaceae_Leptospermum_laevigatum
Myrtaceae_Leptospermum_coriaceum
Myrtaceae_Leptospermum_petersonii
Myrtaceae_Leptospermum_scoparium
Myrtaceae_Taxandria_juniperina
Myrtaceae_Ochrosperma_oligomerum
Myrtaceae_Ochrosperma_lineare
Myrtaceae_Ochrosperma_citriodorum
Myrtaceae_Homalocalyx_polyandrus
Myrtaceae_Homalocalyx_aurea
Myrtaceae_Calytrix_tetragona
Myrtaceae_Babingtonia_behrii
Myrtaceae_Triplarina_imbricata
Myrtaceae_Triplarina_nitchaga
Myrtaceae_Triplarina_volcanica
Myrtaceae_Micromyrtus_delicata
Myrtaceae_Micromyrtus_ciliata
Myrtaceae_Baeckea_tetragona
Myrtaceae_Baeckea_corynophylla
Myrtaceae_Baeckea_uncinella

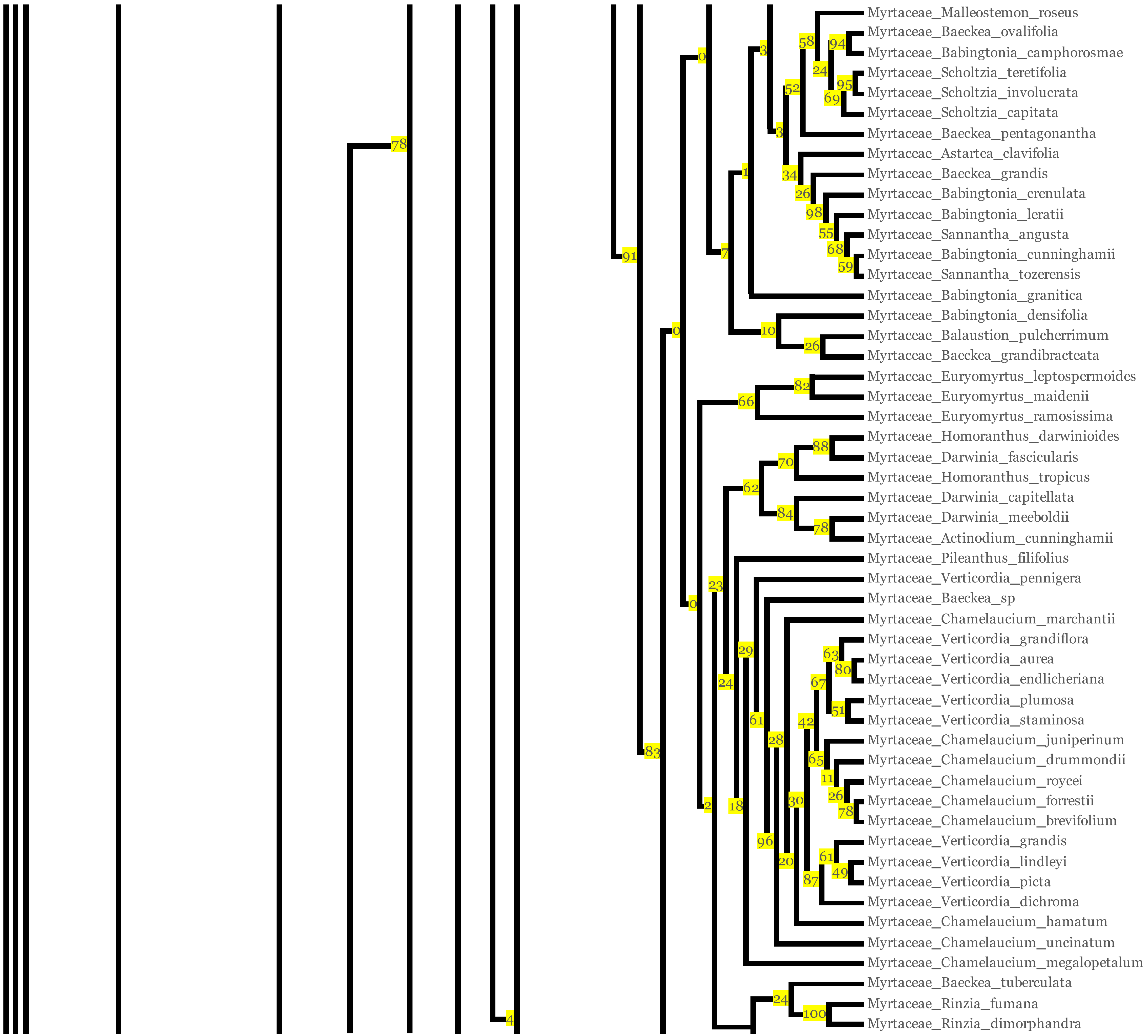
Myrtaceae_Malleostemon_roseus
Myrtaceae_Baeckea_ovalifolia
Myrtaceae_Babingtonia_camphorosmae
Myrtaceae_Scholtzia_teretifolia
Myrtaceae_Scholtzia_involucrata
Myrtaceae_Scholtzia_capitata
Myrtaceae_Baeckea_pentagonantha
Myrtaceae_Astartea_clavifolia
Myrtaceae_Baeckea_grandis
Myrtaceae_Babingtonia_crenulata
Myrtaceae_Babingtonia_leratii
Myrtaceae_Sannantha_angusta
Myrtaceae_Babingtonia_cunninghamii
Myrtaceae_Sannantha_tozerensis
Myrtaceae_Babingtonia_granitica
Myrtaceae_Babingtonia_densifolia
Myrtaceae_Balaustion_pulcherrimum
Myrtaceae_Baeckea_grandibracteata
Myrtaceae_Euryomyrtus_leptospermoides
Myrtaceae_Euryomyrtus_maidenii
Myrtaceae_Euryomyrtus_ramosissima
Myrtaceae_Homoranthus_darwinioides
Myrtaceae_Darwinia_fascicularis
Myrtaceae_Homoranthus_tropicus
Myrtaceae_Darwinia_capitellata
Myrtaceae_Darwinia_meeboldii
Myrtaceae_Actinodium_cunninghamii
Myrtaceae_Pileanthus_filifolius
Myrtaceae_Verticordia_pennigera
Myrtaceae_Baeckea_sp
Myrtaceae_Chamelaucium_marchantii
Myrtaceae_Verticordia_grandiflora
Myrtaceae_Verticordia_aurea
Myrtaceae_Verticordia_endlicheriana
Myrtaceae_Verticordia_plumosa
Myrtaceae_Verticordia_staminosa
Myrtaceae_Chamelaucium_juniperinum
Myrtaceae_Chamelaucium_drummondii
Myrtaceae_Chamelaucium_roycei
Myrtaceae_Chamelaucium_forrestii
Myrtaceae_Chamelaucium_brevifolium
Myrtaceae_Verticordia_grandis
Myrtaceae_Verticordia_lindleyi
Myrtaceae_Verticordia_picta
Myrtaceae_Verticordia_dichroma
Myrtaceae_Chamelaucium_hamatum
Myrtaceae_Chamelaucium_uncinatum
Myrtaceae_Chamelaucium_megalopetalum
Myrtaceae_Baeckea_tuberculata
Myrtaceae_Rinzia_fumana
Myrtaceae_Rinzia_dimorphandra

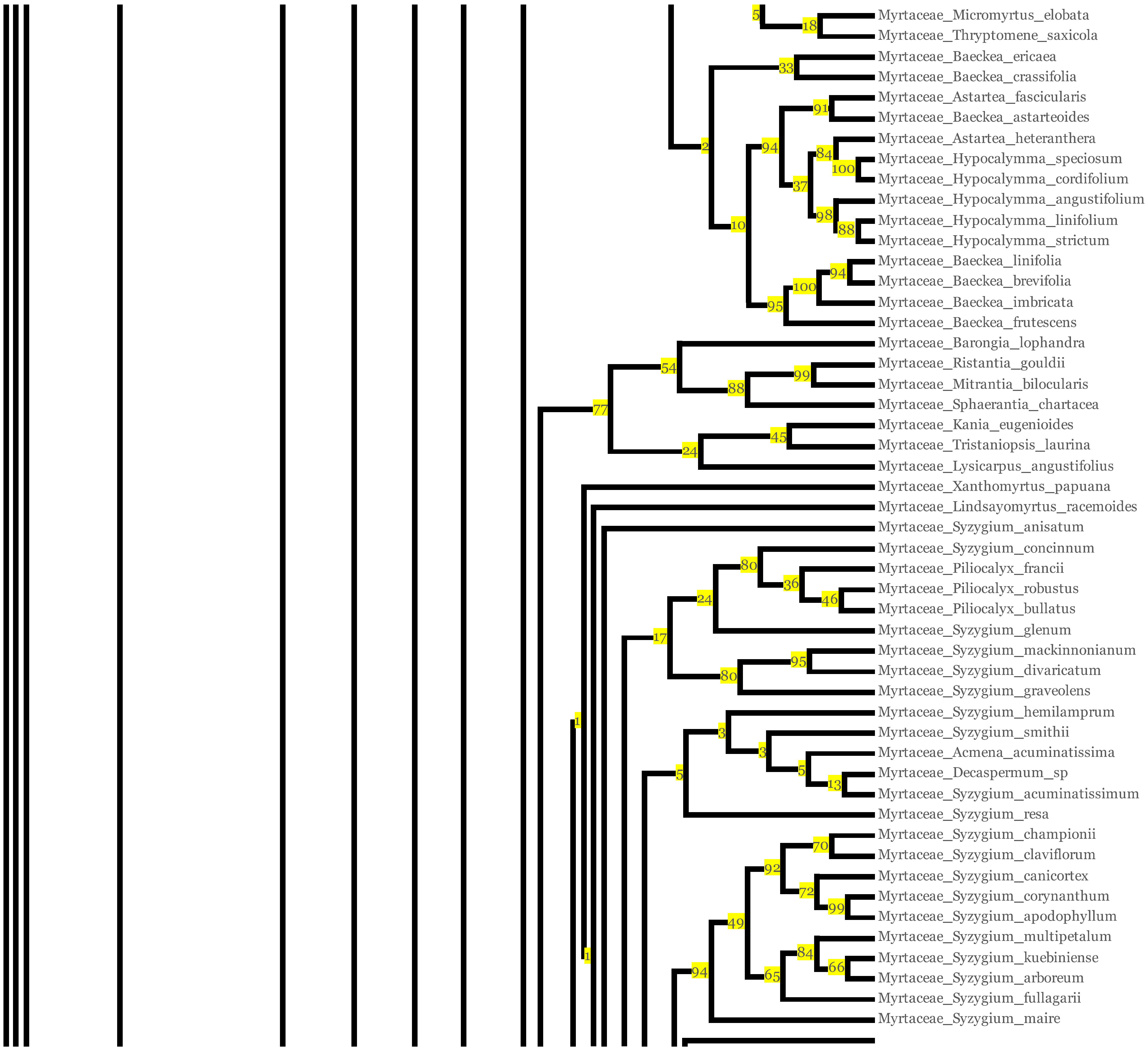
Myrtaceae_Micromyrtus_elobata
Myrtaceae_Thryptomene_saxicola
Myrtaceae_Baeckea_ericaea
Myrtaceae_Baeckea_crassifolia
Myrtaceae_Astartea_fascicularis
Myrtaceae_Baeckea_astarteoides
Myrtaceae_Astartea_heteranthera
Myrtaceae_Hypocalymma_speciosum
Myrtaceae_Hypocalymma_cordifolium
Myrtaceae_Hypocalymma_angustifolium
Myrtaceae_Hypocalymma_linifolium
Myrtaceae_Hypocalymma_strictum
Myrtaceae_Baeckea_linifolia
Myrtaceae_Baeckea_brevifolia
Myrtaceae_Baeckea_imbricata
Myrtaceae_Baeckea_frutescens
Myrtaceae_Barongia_lophandra
Myrtaceae_Ristantia_gouldii
Myrtaceae_Mitrantia_bilocularis
Myrtaceae_Sphaerantia_chartacea
Myrtaceae_Kania_eugenioides
Myrtaceae_Tristaniopsis_laurina
Myrtaceae_Lysicarpus_angustifolius
Myrtaceae_Xanthomyrtus_papuana
Myrtaceae_Lindsayomyrtus_racemoides
Myrtaceae_Syzygium_anisatum
Myrtaceae_Syzygium_concinnum
Myrtaceae_Piliocalyx_francii
Myrtaceae_Piliocalyx_robustus
Myrtaceae_Piliocalyx_bullatus
Myrtaceae_Syzygium_glenum
Myrtaceae_Syzygium_mackinnonianum
Myrtaceae_Syzygium_divaricatum
Myrtaceae_Syzygium_graveolens
Myrtaceae_Syzygium_hemilamprum
Myrtaceae_Syzygium_smithii
Myrtaceae_Acmena_acuminatissima
Myrtaceae_Decaspermum_sp
Myrtaceae_Syzygium_acuminatissimum
Myrtaceae_Syzygium_resa
Myrtaceae_Syzygium_championii
Myrtaceae_Syzygium_claviflorum
Myrtaceae_Syzygium_canicortex
Myrtaceae_Syzygium_corynanthum
Myrtaceae_Syzygium_apodophyllum
Myrtaceae_Syzygium_multipetalum
Myrtaceae_Syzygium_kuebiniense
Myrtaceae_Syzygium_arboreum
Myrtaceae_Syzygium_fullagarii
Myrtaceae_Syzygium_maire

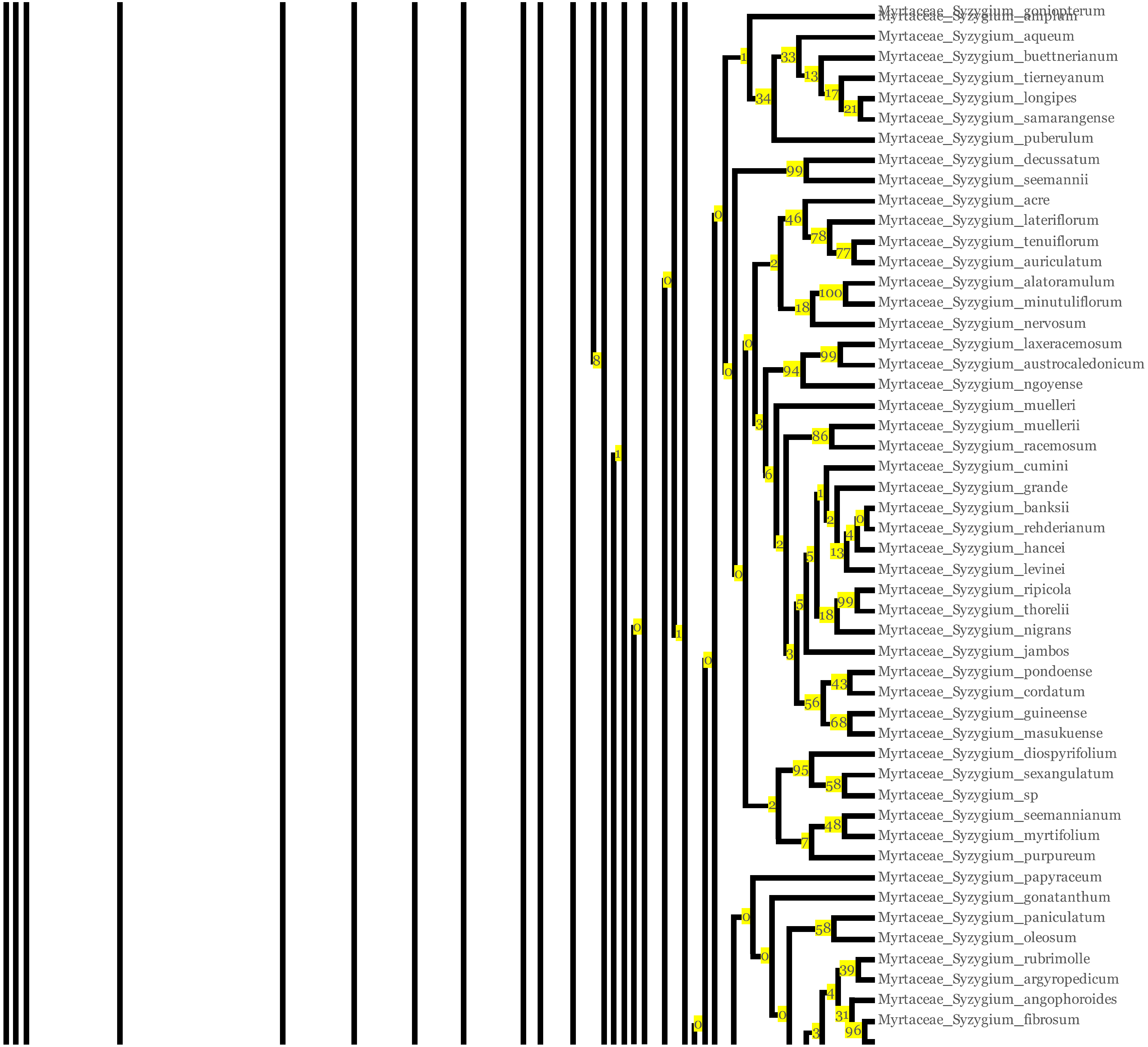
Myrtaceae_Syzygium_goniopterum
Myrtaceae_Syzygium_amplum
Myrtaceae_Syzygium_aqueum
Myrtaceae_Syzygium_buettnerianum
Myrtaceae_Syzygium_tierneyanum
Myrtaceae_Syzygium_longipes
Myrtaceae_Syzygium_samarangense
Myrtaceae_Syzygium_puberulum
Myrtaceae_Syzygium_decussatum
Myrtaceae_Syzygium_seemannii
Myrtaceae_Syzygium_acre
Myrtaceae_Syzygium_lateriflorum
Myrtaceae_Syzygium_tenuiflorum
Myrtaceae_Syzygium_auriculatum
Myrtaceae_Syzygium_alatoramulum
Myrtaceae_Syzygium_minutuliflorum
Myrtaceae_Syzygium_nervosum
Myrtaceae_Syzygium_laxeracemosum
Myrtaceae_Syzygium_austrocaledonicum
Myrtaceae_Syzygium_ngoyense
Myrtaceae_Syzygium_muelleri
Myrtaceae_Syzygium_muellerii
Myrtaceae_Syzygium_racemosum
Myrtaceae_Syzygium_cumini
Myrtaceae_Syzygium_grande
Myrtaceae_Syzygium_banksii
Myrtaceae_Syzygium_rehderianum
Myrtaceae_Syzygium_hancei
Myrtaceae_Syzygium_levinei
Myrtaceae_Syzygium_ripicola
Myrtaceae_Syzygium_thorelii
Myrtaceae_Syzygium_nigrans
Myrtaceae_Syzygium_jambos
Myrtaceae_Syzygium_pondoense
Myrtaceae_Syzygium_cordatum
Myrtaceae_Syzygium_guineense
Myrtaceae_Syzygium_masukuense
Myrtaceae_Syzygium_diospyrifolium
Myrtaceae_Syzygium_sexangulatum
Myrtaceae_Syzygium_sp
Myrtaceae_Syzygium_seemannianum
Myrtaceae_Syzygium_myrtifolium
Myrtaceae_Syzygium_purpureum
Myrtaceae_Syzygium_papyraceum
Myrtaceae_Syzygium_gonatanthum
Myrtaceae_Syzygium_paniculatum
Myrtaceae_Syzygium_oleosum
Myrtaceae_Syzygium_rubrimolle
Myrtaceae_Syzygium_argyropedicum
Myrtaceae_Syzygium_angophoroides
Myrtaceae_Syzygium_fibrosum

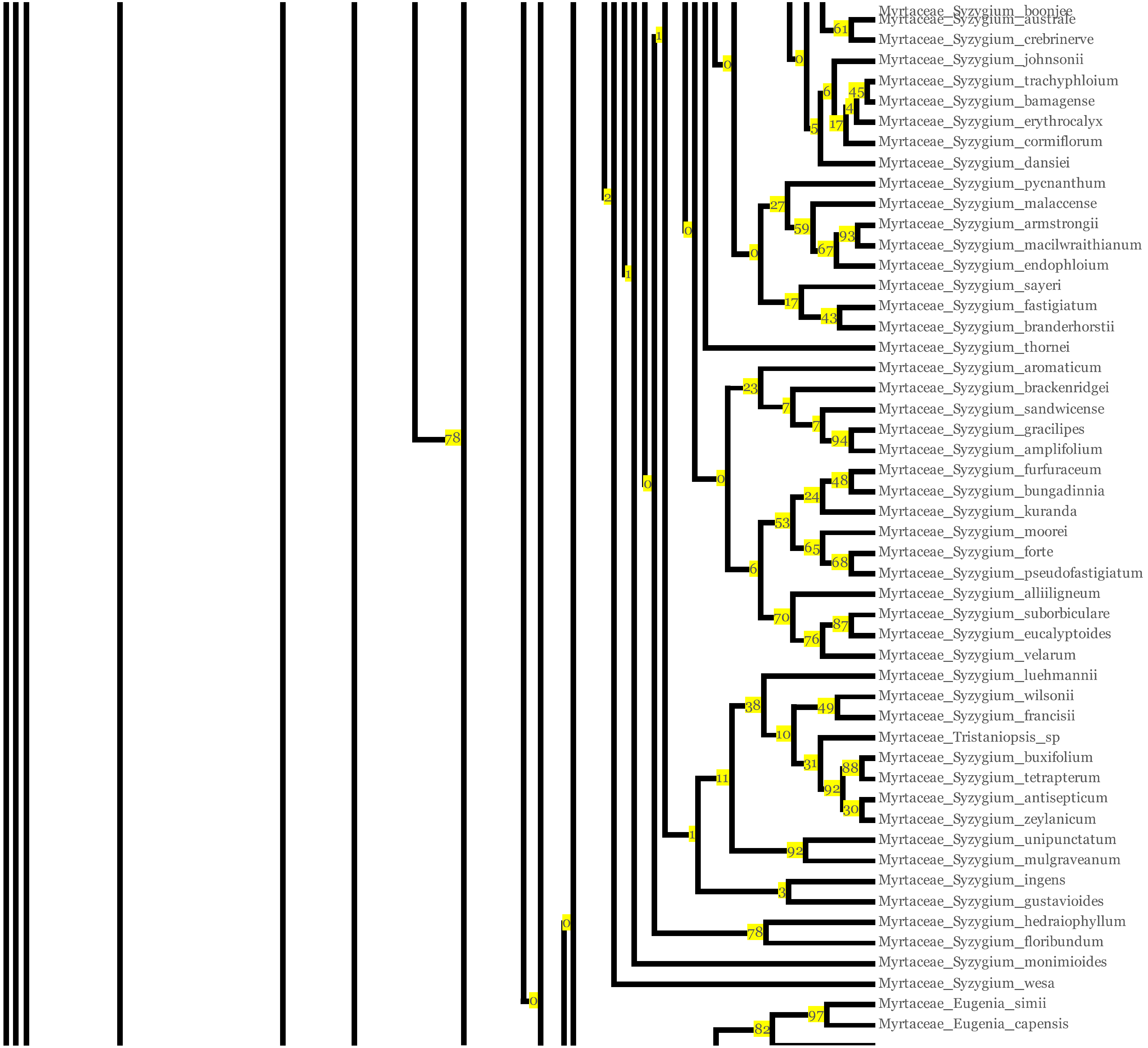
Myrtaceae_Syzygium_boonjee
Myrtaceae_Syzygium_australe
Myrtaceae_Syzygium_crebrinerve
Myrtaceae_Syzygium_johnsonii
Myrtaceae_Syzygium_trachyphloium
Myrtaceae_Syzygium_bamagense
Myrtaceae_Syzygium_erythrocalyx
Myrtaceae_Syzygium_cormiflorum
Myrtaceae_Syzygium_dansiei
Myrtaceae_Syzygium_pycnanthum
Myrtaceae_Syzygium_malaccense
Myrtaceae_Syzygium_armstrongii
Myrtaceae_Syzygium_macilwraithianum
Myrtaceae_Syzygium_endophloium
Myrtaceae_Syzygium_sayeri
Myrtaceae_Syzygium_fastigiatum
Myrtaceae_Syzygium_branderhorstii
Myrtaceae_Syzygium_thornei
Myrtaceae_Syzygium_aromaticum
Myrtaceae_Syzygium_brackenridgei
Myrtaceae_Syzygium_sandwicense
Myrtaceae_Syzygium_gracilipes
Myrtaceae_Syzygium_amplifolium
Myrtaceae_Syzygium_furfuraceum
Myrtaceae_Syzygium_bungadinnia
Myrtaceae_Syzygium_kuranda
Myrtaceae_Syzygium_moorei
Myrtaceae_Syzygium_forte
Myrtaceae_Syzygium_pseudofastigiatum
Myrtaceae_Syzygium_alliiligneum
Myrtaceae_Syzygium_suborbiculare
Myrtaceae_Syzygium_eucalyptoides
Myrtaceae_Syzygium_velarum
Myrtaceae_Syzygium_luehmannii
Myrtaceae_Syzygium_wilsonii
Myrtaceae_Syzygium_francisii
Myrtaceae_Tristaniopsis_sp
Myrtaceae_Syzygium_buxifolium
Myrtaceae_Syzygium_tetrapterum
Myrtaceae_Syzygium_antisepticum
Myrtaceae_Syzygium_zeylanicum
Myrtaceae_Syzygium_unipunctatum
Myrtaceae_Syzygium_mulgraveanum
Myrtaceae_Syzygium_ingens
Myrtaceae_Syzygium_gustavioides
Myrtaceae_Syzygium_hedraiophyllum
Myrtaceae_Syzygium_floribundum
Myrtaceae_Syzygium_monimioides
Myrtaceae_Syzygium_wesa
Myrtaceae_Eugenia_simii
Myrtaceae_Eugenia_capensis

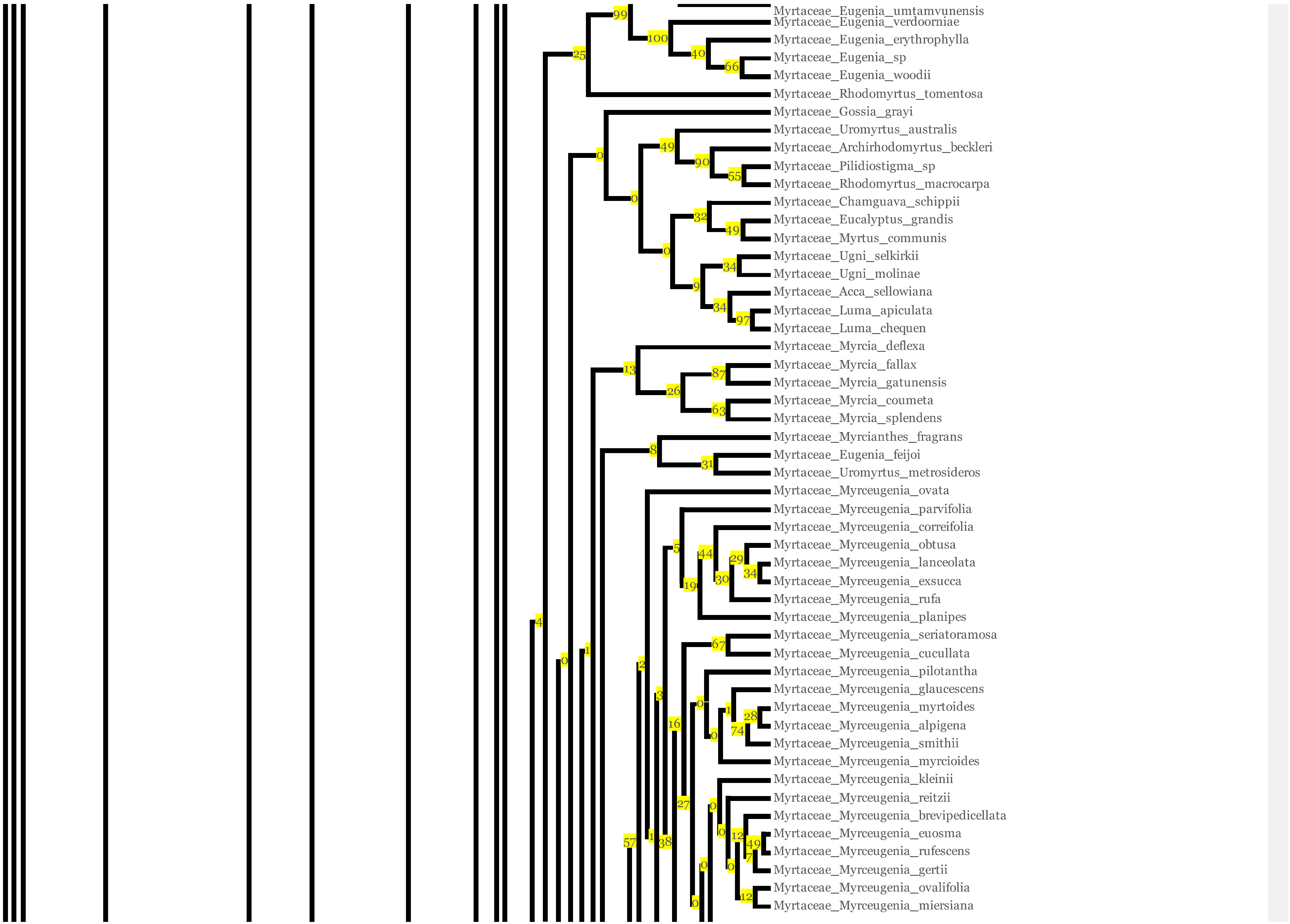
Myrtaceae_Eugenia_umtamvunensis
Myrtaceae_Eugenia_verdoorniae
Myrtaceae_Eugenia_erythrophylla
Myrtaceae_Eugenia_sp
Myrtaceae_Eugenia_woodii
Myrtaceae_Rhodomyrtus_tomentosa
Myrtaceae_Gossia_grayi
Myrtaceae_Uromyrtus_australis
Myrtaceae_Archirhodomyrtus_beckleri
Myrtaceae_Pilidiostigma_sp
Myrtaceae_Rhodomyrtus_macrocarpa
Myrtaceae_Chamguava_schippii
Myrtaceae_Eucalyptus_grandis
Myrtaceae_Myrtus_communis
Myrtaceae_Ugni_selkirkii
Myrtaceae_Ugni_molinae
Myrtaceae_Acca_sellowiana
Myrtaceae_Luma_apiculata
Myrtaceae_Luma_chequen
Myrtaceae_Myrcia_deflexa
Myrtaceae_Myrcia_fallax
Myrtaceae_Myrcia_gatunensis
Myrtaceae_Myrcia_coumeta
Myrtaceae_Myrcia_splendens
Myrtaceae_Myrcianthes_fragrans
Myrtaceae_Eugenia_feijoi
Myrtaceae_Uromyrtus_metrosideros
Myrtaceae_Myrceugenia_ovata
Myrtaceae_Myrceugenia_parvifolia
Myrtaceae_Myrceugenia_correifolia
Myrtaceae_Myrceugenia_obtusa
Myrtaceae_Myrceugenia_lanceolata
Myrtaceae_Myrceugenia_exsucca
Myrtaceae_Myrceugenia_rufa
Myrtaceae_Myrceugenia_planipes
Myrtaceae_Myrceugenia_seriatoramosa
Myrtaceae_Myrceugenia_cucullata
Myrtaceae_Myrceugenia_pilotantha
Myrtaceae_Myrceugenia_glaucescens
Myrtaceae_Myrceugenia_myrtoides
Myrtaceae_Myrceugenia_alpigena
Myrtaceae_Myrceugenia_smithii
Myrtaceae_Myrceugenia_myrcioides
Myrtaceae_Myrceugenia_kleinii
Myrtaceae_Myrceugenia_reitzii
Myrtaceae_Myrceugenia_brevipedicellata
Myrtaceae_Myrceugenia_euosma
Myrtaceae_Myrceugenia_rufescens
Myrtaceae_Myrceugenia_gertii
Myrtaceae_Myrceugenia_ovalifolia
Myrtaceae_Myrceugenia_miersiana

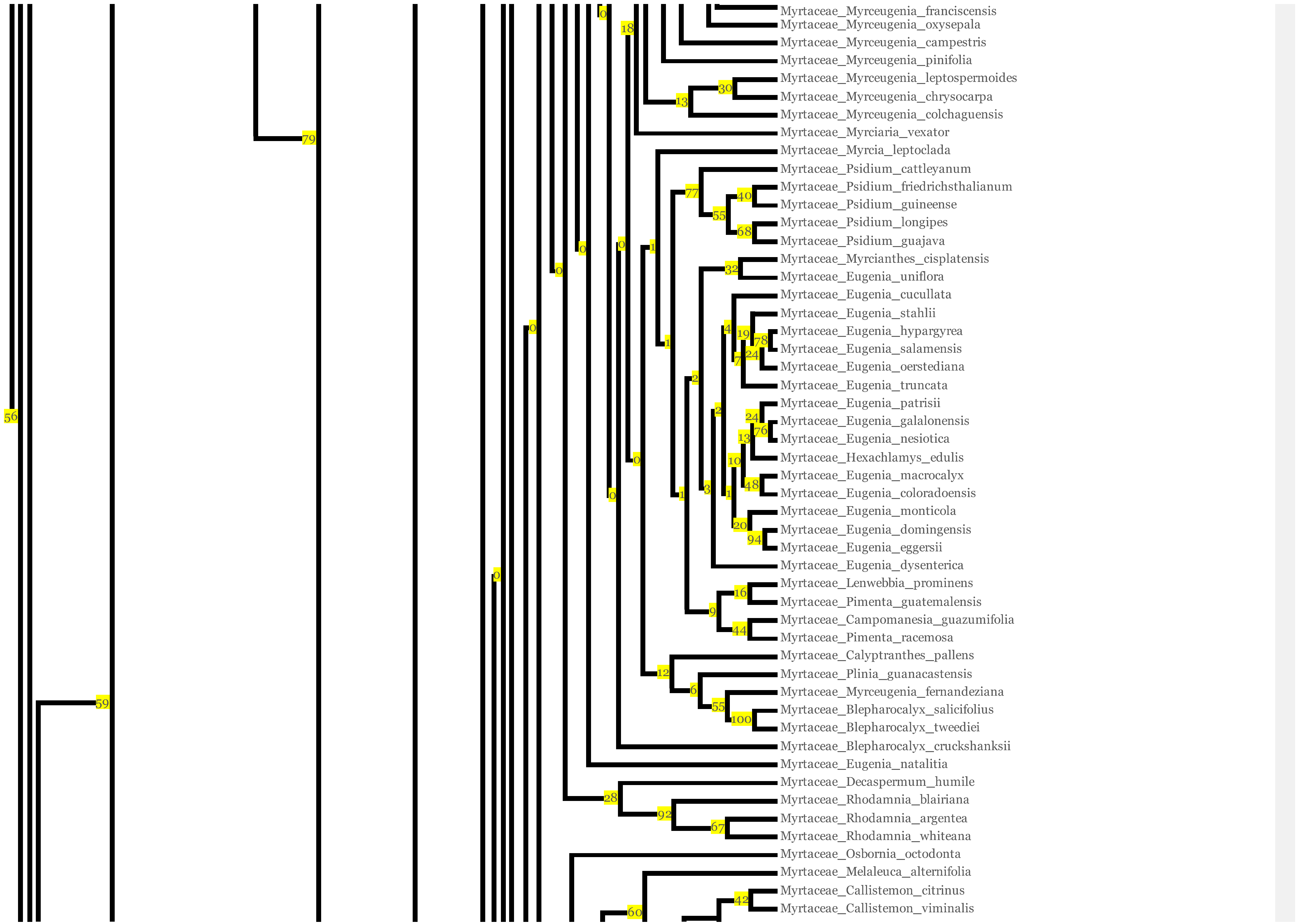
Myrtaceae_Myrceugenia_franciscensis
Myrtaceae_Myrceugenia_oxysepala
Myrtaceae_Myrceugenia_campestris
Myrtaceae_Myrceugenia_pinifolia
Myrtaceae_Myrceugenia_leptospermoides
Myrtaceae_Myrceugenia_chrysocarpa
Myrtaceae_Myrceugenia_colchaguensis
Myrtaceae_Myrciaria_vexator
Myrtaceae_Myrcia_leptoclada
Myrtaceae_Psidium_cattleyanum
Myrtaceae_Psidium_friedrichsthalianum
Myrtaceae_Psidium_guineense
Myrtaceae_Psidium_longipes
Myrtaceae_Psidium_guajava
Myrtaceae_Myrcianthes_cisplatensis
Myrtaceae_Eugenia_uniflora
Myrtaceae_Eugenia_cucullata
Myrtaceae_Eugenia_stahlii
Myrtaceae_Eugenia_hypargyrea
Myrtaceae_Eugenia_salamensis
Myrtaceae_Eugenia_oerstediana
Myrtaceae_Eugenia_truncata
Myrtaceae_Eugenia_patrisii
Myrtaceae_Eugenia_galalonensis
Myrtaceae_Eugenia_nesiotica
Myrtaceae_Hexachlamys_edulis
Myrtaceae_Eugenia_macrocalyx
Myrtaceae_Eugenia_coloradoensis
Myrtaceae_Eugenia_monticola
Myrtaceae_Eugenia_domingensis
Myrtaceae_Eugenia_eggersii
Myrtaceae_Eugenia_dysenterica
Myrtaceae_Lenwebbia_prominens
Myrtaceae_Pimenta_guatemalensis
Myrtaceae_Campomanesia_guazumifolia
Myrtaceae_Pimenta_racemosa
Myrtaceae_Calyptranthes_pallens
Myrtaceae_Plinia_guanacastensis
Myrtaceae_Myrceugenia_fernandeziana
Myrtaceae_Blepharocalyx_salicifolius
Myrtaceae_Blepharocalyx_tweediei
Myrtaceae_Blepharocalyx_cruckshanksii
Myrtaceae_Eugenia_natalitia
Myrtaceae_Decaspermum_humile
Myrtaceae_Rhodamnia_blairiana
Myrtaceae_Rhodamnia_argentea
Myrtaceae_Rhodamnia_whiteana
Myrtaceae_Osbornia_octodonta
Myrtaceae_Melaleuca_alternifolia
Myrtaceae_Callistemon_citrinus
Myrtaceae_Callistemon_viminalis

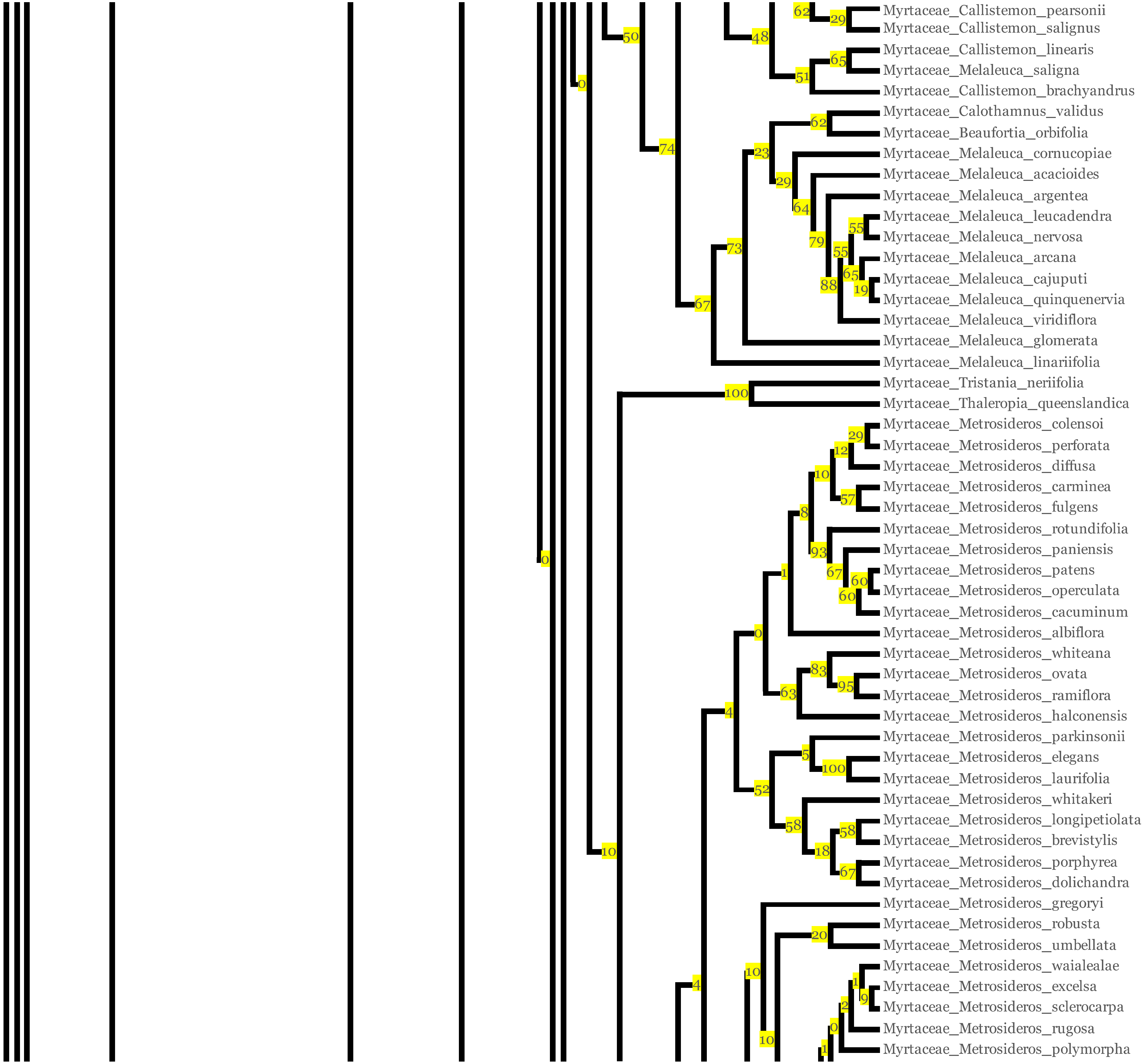
Myrtaceae_Callistemon_pearsonii
Myrtaceae_Callistemon_salignus
Myrtaceae_Callistemon_linearis
Myrtaceae_Melaleuca_saligna
Myrtaceae_Callistemon_brachyandrus
Myrtaceae_Calothamnus_validus
Myrtaceae_Beaufortia_orbifolia
Myrtaceae_Melaleuca_cornucopiae
Myrtaceae_Melaleuca_acacioides
Myrtaceae_Melaleuca_argentea
Myrtaceae_Melaleuca_leucadendra
Myrtaceae_Melaleuca_nervosa
Myrtaceae_Melaleuca_arcana
Myrtaceae_Melaleuca_cajuputi
Myrtaceae_Melaleuca_quinquenervia
Myrtaceae_Melaleuca_viridiflora
Myrtaceae_Melaleuca_glomerata
Myrtaceae_Melaleuca_linariifolia
Myrtaceae_Tristania_neriifolia
Myrtaceae_Thaleropia_queenslandica
Myrtaceae_Metrosideros_colensoi
Myrtaceae_Metrosideros_perforata
Myrtaceae_Metrosideros_diffusa
Myrtaceae_Metrosideros_carminea
Myrtaceae_Metrosideros_fulgens
Myrtaceae_Metrosideros_rotundifolia
Myrtaceae_Metrosideros_paniensis
Myrtaceae_Metrosideros_patens
Myrtaceae_Metrosideros_operculata
Myrtaceae_Metrosideros_cacuminum
Myrtaceae_Metrosideros_albiflora
Myrtaceae_Metrosideros_whiteana
Myrtaceae_Metrosideros_ovata
Myrtaceae_Metrosideros_ramiflora
Myrtaceae_Metrosideros_halconensis
Myrtaceae_Metrosideros_parkinsonii
Myrtaceae_Metrosideros_elegans
Myrtaceae_Metrosideros_laurifolia
Myrtaceae_Metrosideros_whitakeri
Myrtaceae_Metrosideros_longipetiolata
Myrtaceae_Metrosideros_brevistylis
Myrtaceae_Metrosideros_porphyrea
Myrtaceae_Metrosideros_dolichandra
Myrtaceae_Metrosideros_gregoryi
Myrtaceae_Metrosideros_robusta
Myrtaceae_Metrosideros_umbellata
Myrtaceae_Metrosideros_waialealae
Myrtaceae_Metrosideros_excelsa
Myrtaceae_Metrosideros_sclerocarpa
Myrtaceae_Metrosideros_rugosa
Myrtaceae_Metrosideros_polymorpha

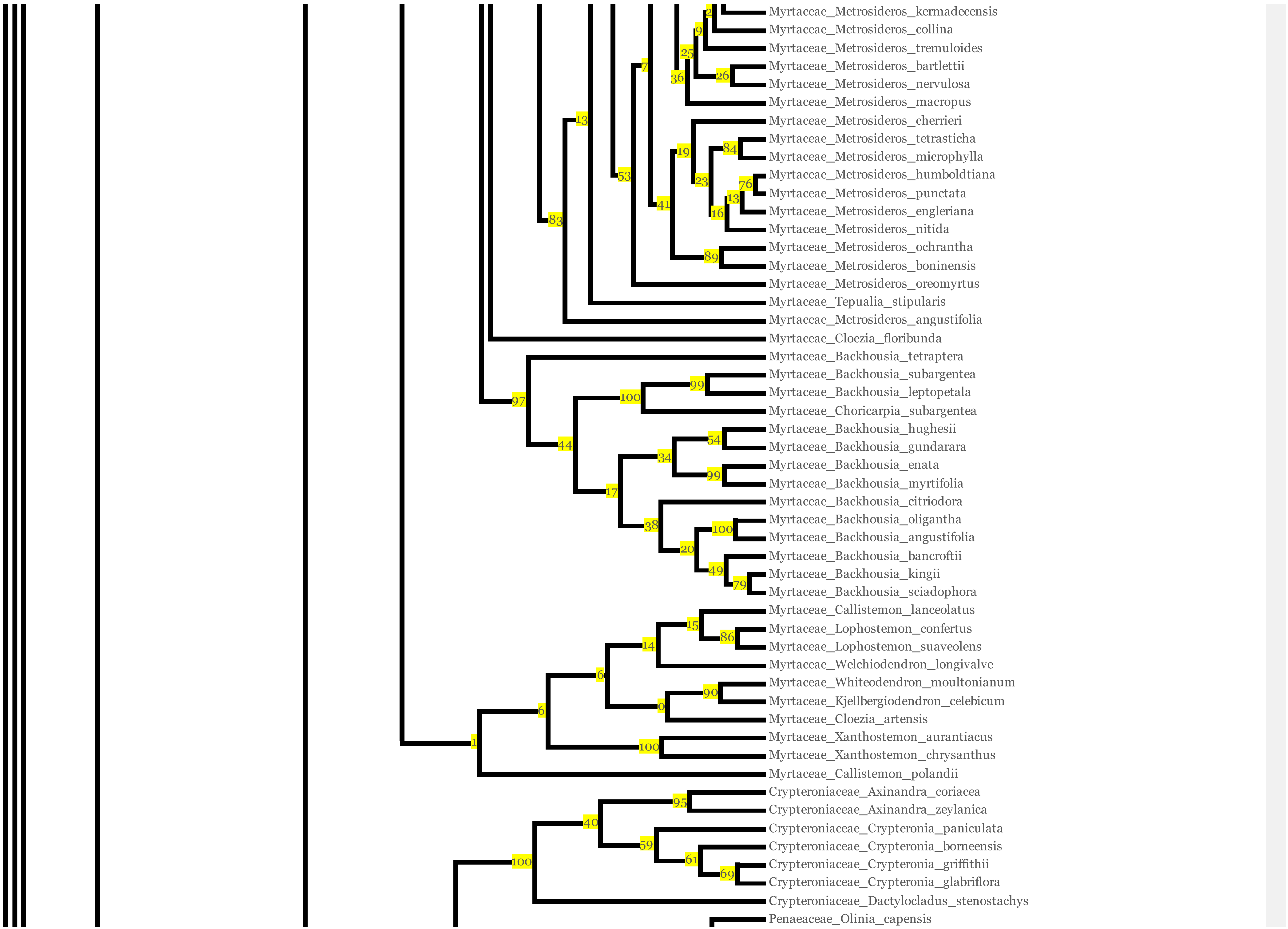
Myrtaceae_Metrosideros_kermadecensis
Myrtaceae_Metrosideros_collina
Myrtaceae_Metrosideros_tremuloides
Myrtaceae_Metrosideros_bartlettii
Myrtaceae_Metrosideros_nervulosa
Myrtaceae_Metrosideros_macropus
Myrtaceae_Metrosideros_cherrieri
Myrtaceae_Metrosideros_tetrasticha
Myrtaceae_Metrosideros_microphylla
Myrtaceae_Metrosideros_humboldtiana
Myrtaceae_Metrosideros_punctata
Myrtaceae_Metrosideros_engleriana
Myrtaceae_Metrosideros_nitida
Myrtaceae_Metrosideros_ochrantha
Myrtaceae_Metrosideros_boninensis
Myrtaceae_Metrosideros_oreomyrtus
Myrtaceae_Tepualia_stipularis
Myrtaceae_Metrosideros_angustifolia
Myrtaceae_Cloezia_floribunda
Myrtaceae_Backhousia_tetraptera
Myrtaceae_Backhousia_subargentea
Myrtaceae_Backhousia_leptopetala
Myrtaceae_Choricarpia_subargentea
Myrtaceae_Backhousia_hughesii
Myrtaceae_Backhousia_gundarara
Myrtaceae_Backhousia_enata
Myrtaceae_Backhousia_myrtifolia
Myrtaceae_Backhousia_citriodora
Myrtaceae_Backhousia_oligantha
Myrtaceae_Backhousia_angustifolia
Myrtaceae_Backhousia_bancroftii
Myrtaceae_Backhousia_kingii
Myrtaceae_Backhousia_sciadophora
Myrtaceae_Callistemon_lanceolatus
Myrtaceae_Lophostemon_confertus
Myrtaceae_Lophostemon_suaveolens
Myrtaceae_Welchiodendron_longivalve
Myrtaceae_Whiteodendron_moultonianum
Myrtaceae_Kjellbergiodendron_celebicum
Myrtaceae_Cloezia_artensis
Myrtaceae_Xanthostemon_aurantiacus
Myrtaceae_Xanthostemon_chrysanthus
Myrtaceae_Callistemon_polandii
Crypteroniaceae_Axinandra_coriacea
Crypteroniaceae_Axinandra_zeylanica
Crypteroniaceae_Crypteronia_paniculata
Crypteroniaceae_Crypteronia_borneensis
Crypteroniaceae_Crypteronia_griffithii
Crypteroniaceae_Crypteronia_glabriflora
Crypteroniaceae_Dactylocladus_stenostachys
Penaeaceae_Olinia_capensis

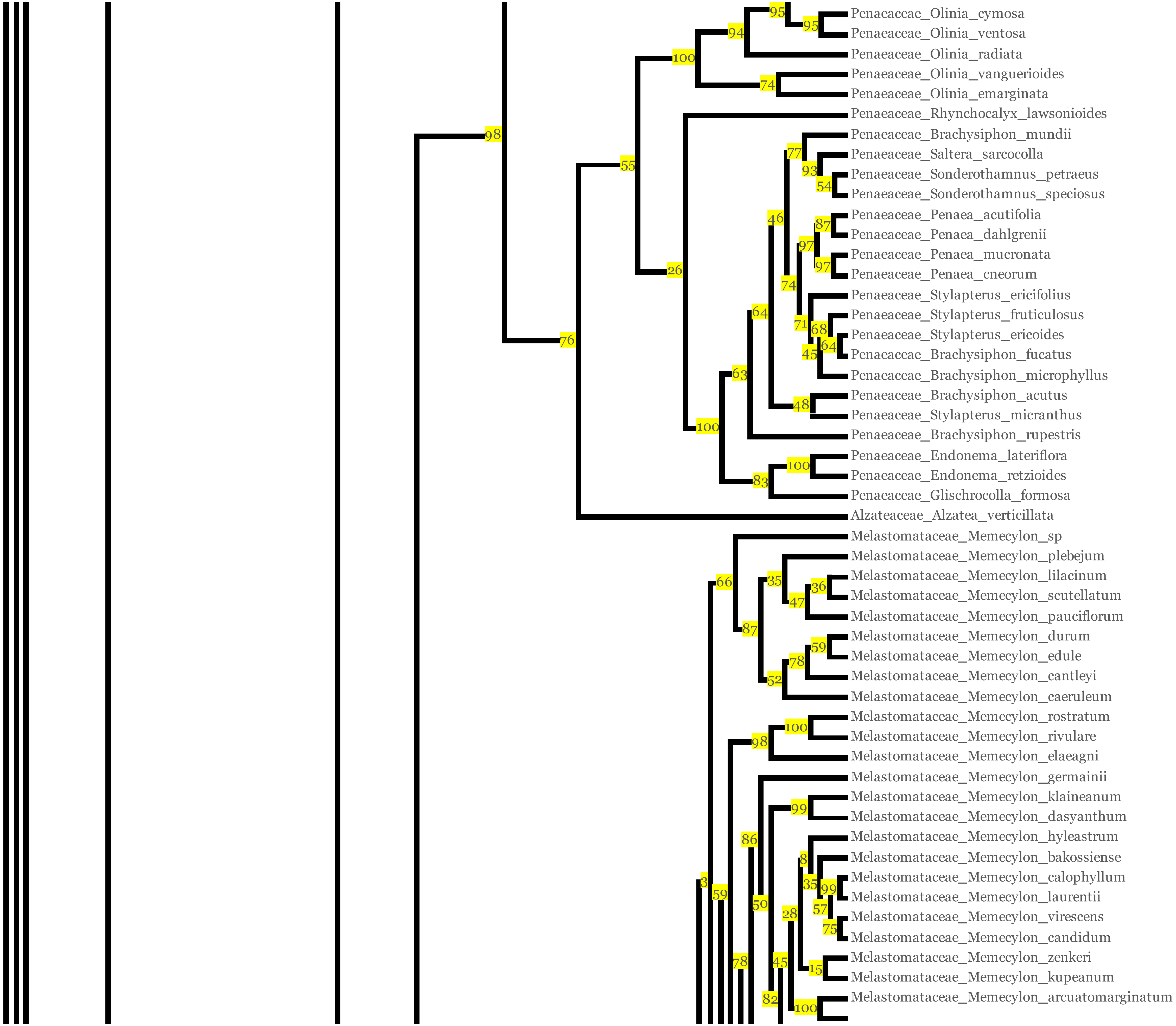
Penaeaceae_Olinia_cymosa
Penaeaceae_Olinia_ventosa
Penaeaceae_Olinia_radiata
Penaeaceae_Olinia_vanguerioides
Penaeaceae_Olinia_emarginata
Penaeaceae_Rhynchocalyx_lawsonioides
Penaeaceae_Brachysiphon_mundii
Penaeaceae_Saltera_sarcocolla
Penaeaceae_Sonderothamnus_petraeus
Penaeaceae_Sonderothamnus_speciosus
Penaeaceae_Penaea_acutifolia
Penaeaceae_Penaea_dahlgrenii
Penaeaceae_Penaea_mucronata
Penaeaceae_Penaea_cneorum
Penaeaceae_Stylapterus_ericifolius
Penaeaceae_Stylapterus_fruticulosus
Penaeaceae_Stylapterus_ericoides
Penaeaceae_Brachysiphon_fucatus
Penaeaceae_Brachysiphon_microphyllus
Penaeaceae_Brachysiphon_acutus
Penaeaceae_Stylapterus_micranthus
Penaeaceae_Brachysiphon_rupestris
Penaeaceae_Endonema_lateriflora
Penaeaceae_Endonema_retzioides
Penaeaceae_Glischrocolla_formosa
Alzateaceae_Alzatea_verticillata
Melastomataceae_Memecylon_sp
Melastomataceae_Memecylon_plebejum
Melastomataceae_Memecylon_lilacinum
Melastomataceae_Memecylon_scutellatum
Melastomataceae_Memecylon_pauciflorum
Melastomataceae_Memecylon_durum
Melastomataceae_Memecylon_edule
Melastomataceae_Memecylon_cantleyi
Melastomataceae_Memecylon_caeruleum
Melastomataceae_Memecylon_rostratum
Melastomataceae_Memecylon_rivulare
Melastomataceae_Memecylon_elaeagni
Melastomataceae_Memecylon_germainii
Melastomataceae_Memecylon_klaineanum
Melastomataceae_Memecylon_dasyanthum
Melastomataceae_Memecylon_hyleastrum
Melastomataceae_Memecylon_bakossiense
Melastomataceae_Memecylon_calophyllum
Melastomataceae_Memecylon_laurentii
Melastomataceae_Memecylon_virescens
Melastomataceae_Memecylon_candidum
Melastomataceae_Memecylon_zenkeri
Melastomataceae_Memecylon_kupeanum
Melastomataceae_Memecylon_arcuatomarginatum

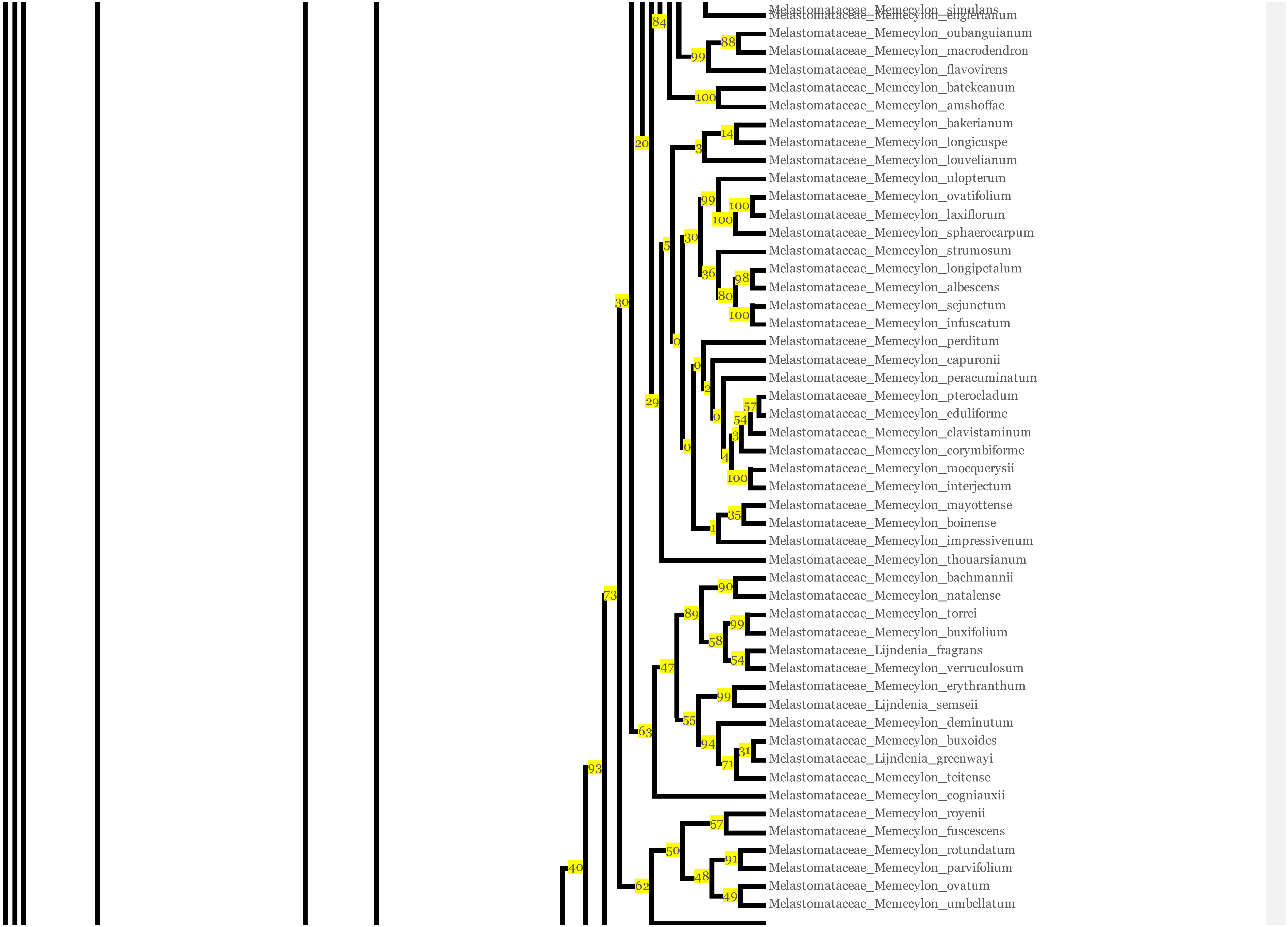
Melastomataceae_Memecylon_simulans
Melastomataceae_Memecylon_englerianum
Melastomataceae_Memecylon_oubanguianum
Melastomataceae_Memecylon_macrodendron
Melastomataceae_Memecylon_flavovirens
Melastomataceae_Memecylon_batekeanum
Melastomataceae_Memecylon_amshoffae
Melastomataceae_Memecylon_bakerianum
Melastomataceae_Memecylon_longicuspe
Melastomataceae_Memecylon_louvelianum
Melastomataceae_Memecylon_ulopterum
Melastomataceae_Memecylon_ovatifolium
Melastomataceae_Memecylon_laxiflorum
Melastomataceae_Memecylon_sphaerocarpum
Melastomataceae_Memecylon_strumosum
Melastomataceae_Memecylon_longipetalum
Melastomataceae_Memecylon_albescens
Melastomataceae_Memecylon_sejunctum
Melastomataceae_Memecylon_infuscatum
Melastomataceae_Memecylon_perditum
Melastomataceae_Memecylon_capuronii
Melastomataceae_Memecylon_peracuminatum
Melastomataceae_Memecylon_pterocladum
Melastomataceae_Memecylon_eduliforme
Melastomataceae_Memecylon_clavistaminum
Melastomataceae_Memecylon_corymbiforme
Melastomataceae_Memecylon_mocquerysii
Melastomataceae_Memecylon_interjectum
Melastomataceae_Memecylon_mayottense
Melastomataceae_Memecylon_boinense
Melastomataceae_Memecylon_impressivenum
Melastomataceae_Memecylon_thouarsianum
Melastomataceae_Memecylon_bachmannii
Melastomataceae_Memecylon_natalense
Melastomataceae_Memecylon_torrei
Melastomataceae_Memecylon_buxifolium
Melastomataceae_Lijndenia_fragrans
Melastomataceae_Memecylon_verruculosum
Melastomataceae_Memecylon_erythranthum
Melastomataceae_Lijndenia_semseii
Melastomataceae_Memecylon_deminutum
Melastomataceae_Memecylon_buxoides
Melastomataceae_Lijndenia_greenwayi
Melastomataceae_Memecylon_teitense
Melastomataceae_Memecylon_cogniauxii
Melastomataceae_Memecylon_royenii
Melastomataceae_Memecylon_fuscescens
Melastomataceae_Memecylon_rotundatum
Melastomataceae_Memecylon_parvifolium
Melastomataceae_Memecylon_ovatum
Melastomataceae_Memecylon_umbellatum

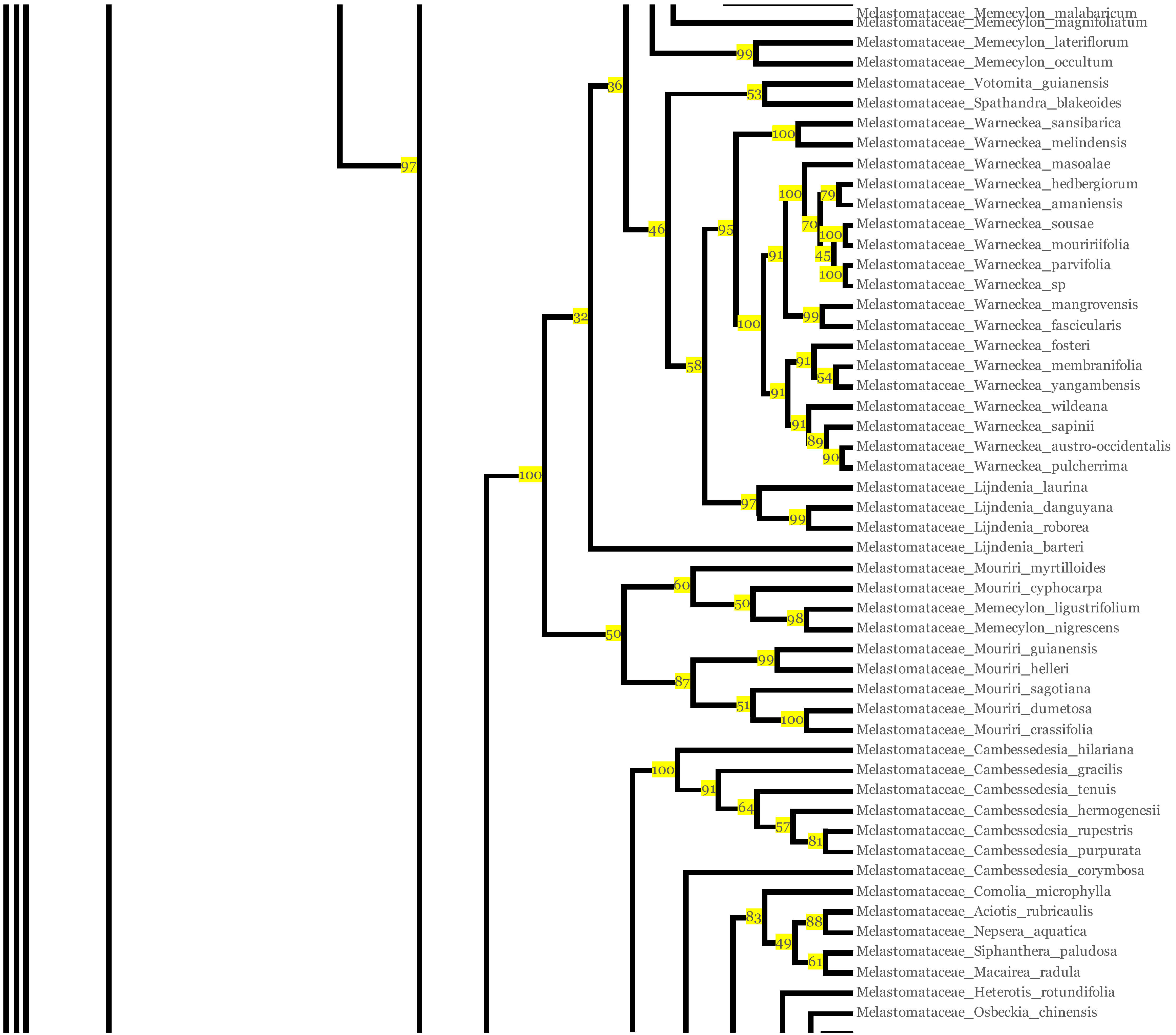
Melastomataceae_Memecylon_malabaricum
Melastomataceae_Memecylon_magnifoliatum
Melastomataceae_Memecylon_lateriflorum
Melastomataceae_Memecylon_occultum
Melastomataceae_Votomita_guianensis
Melastomataceae_Spathandra_blakeoides
Melastomataceae_Warneckea_sansibarica
Melastomataceae_Warneckea_melindensis
Melastomataceae_Warneckea_masoalae
Melastomataceae_Warneckea_hedbergiorum
Melastomataceae_Warneckea_amaniensis
Melastomataceae_Warneckea_sousae
Melastomataceae_Warneckea_mouririifolia
Melastomataceae_Warneckea_parvifolia
Melastomataceae_Warneckea_sp
Melastomataceae_Warneckea_mangrovensis
Melastomataceae_Warneckea_fascicularis
Melastomataceae_Warneckea_fosteri
Melastomataceae_Warneckea_membranifolia
Melastomataceae_Warneckea_yangambensis
Melastomataceae_Warneckea_wildeana
Melastomataceae_Warneckea_sapinii
Melastomataceae_Warneckea_austro-occidentalis
Melastomataceae_Warneckea_pulcherrima
Melastomataceae_Lijndenia_laurina
Melastomataceae_Lijndenia_danguyana
Melastomataceae_Lijndenia_roborea
Melastomataceae_Lijndenia_barteri
Melastomataceae_Mouriri_myrtilloides
Melastomataceae_Mouriri_cyphocarpa
Melastomataceae_Memecylon_ligustrifolium
Melastomataceae_Memecylon_nigrescens
Melastomataceae_Mouriri_guianensis
Melastomataceae_Mouriri_helleri
Melastomataceae_Mouriri_sagotiana
Melastomataceae_Mouriri_dumetosa
Melastomataceae_Mouriri_crassifolia
Melastomataceae_Cambessedesia_hilariana
Melastomataceae_Cambessedesia_gracilis
Melastomataceae_Cambessedesia_tenuis
Melastomataceae_Cambessedesia_hermogenesii
Melastomataceae_Cambessedesia_rupestris
Melastomataceae_Cambessedesia_purpurata
Melastomataceae_Cambessedesia_corymbosa
Melastomataceae_Comolia_microphylla
Melastomataceae_Aciotis_rubricaulis
Melastomataceae_Nepsera_aquatica
Melastomataceae_Siphanthera_paludosa
Melastomataceae_Macairea_radula
Melastomataceae_Heterotis_rotundifolia
Melastomataceae_Osbeckia_chinensis

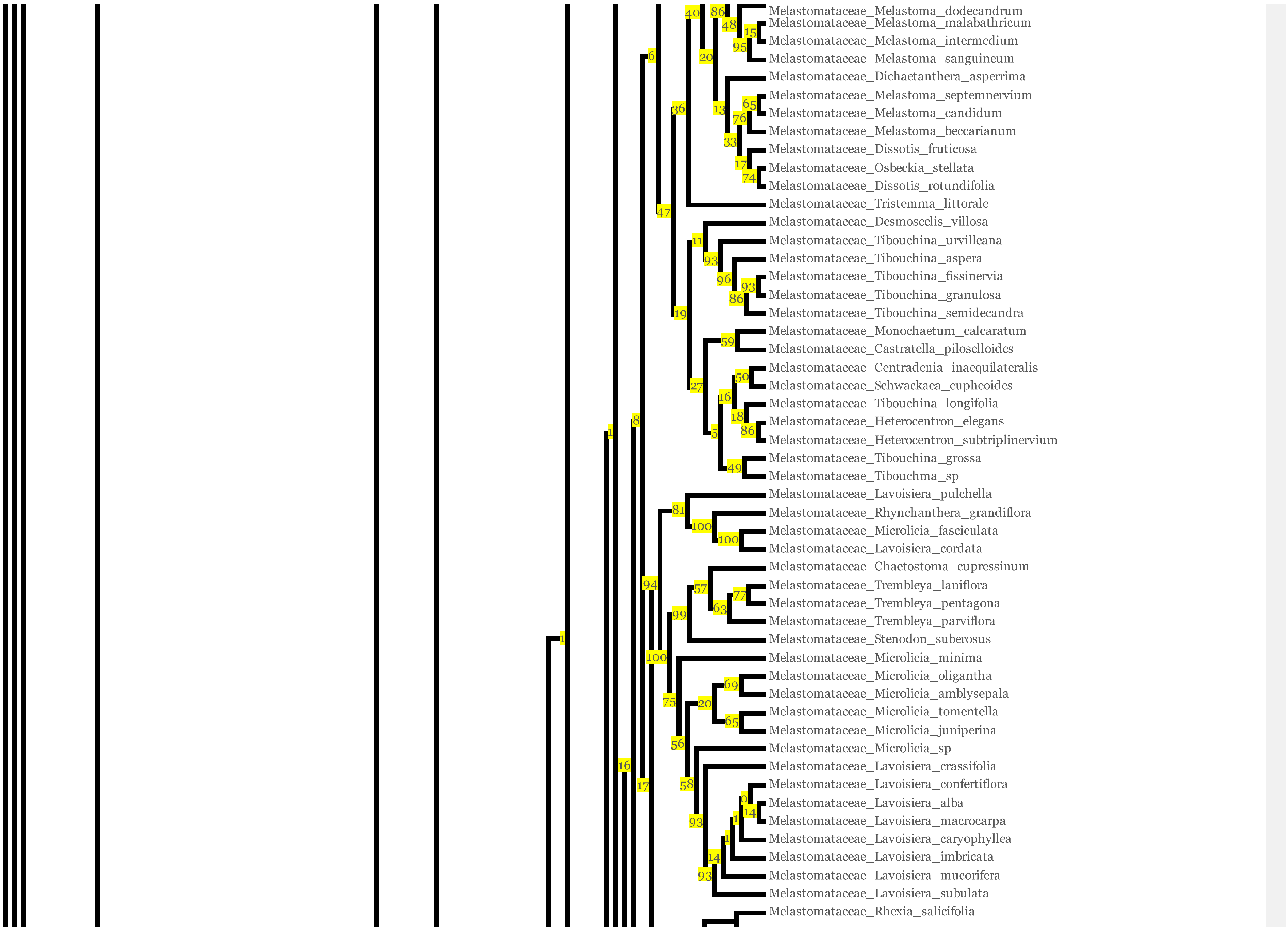
Melastomataceae_Melastoma_dodecandrum
Melastomataceae_Melastoma_malabathricum
Melastomataceae_Melastoma_intermedium
Melastomataceae_Melastoma_sanguineum
Melastomataceae_Dichaetanthera_asperrima
Melastomataceae_Melastoma_septemnervium
Melastomataceae_Melastoma_candidum
Melastomataceae_Melastoma_beccarianum
Melastomataceae_Dissotis_fruticosa
Melastomataceae_Osbeckia_stellata
Melastomataceae_Dissotis_rotundifolia
Melastomataceae_Tristemma_littorale
Melastomataceae_Desmoscelis_villosa
Melastomataceae_Tibouchina_urvilleana
Melastomataceae_Tibouchina_aspera
Melastomataceae_Tibouchina_fissinervia
Melastomataceae_Tibouchina_granulosa
Melastomataceae_Tibouchina_semidecandra
Melastomataceae_Monochaetum_calcaratum
Melastomataceae_Castratella_piloselloides
Melastomataceae_Centradenia_inaequilateralis
Melastomataceae_Schwackaea_cupheoides
Melastomataceae_Tibouchina_longifolia
Melastomataceae_Heterocentron_elegans
Melastomataceae_Heterocentron_subtriplinervium
Melastomataceae_Tibouchina_grossa
Melastomataceae_Tibouchma_sp
Melastomataceae_Lavoisiera_pulchella
Melastomataceae_Rhynchanthera_grandiflora
Melastomataceae_Microlicia_fasciculata
Melastomataceae_Lavoisiera_cordata
Melastomataceae_Chaetostoma_cupressinum
Melastomataceae_Trembleya_laniflora
Melastomataceae_Trembleya_pentagona
Melastomataceae_Trembleya_parviflora
Melastomataceae_Stenodon_suberosus
Melastomataceae_Microlicia_minima
Melastomataceae_Microlicia_oligantha
Melastomataceae_Microlicia_amblysepala
Melastomataceae_Microlicia_tomentella
Melastomataceae_Microlicia_juniperina
Melastomataceae_Microlicia_sp
Melastomataceae_Lavoisiera_crassifolia
Melastomataceae_Lavoisiera_confertiflora
Melastomataceae_Lavoisiera_alba
Melastomataceae_Lavoisiera_macrocarpa
Melastomataceae_Lavoisiera_caryophyllea
Melastomataceae_Lavoisiera_imbricata
Melastomataceae_Lavoisiera_mucorifera
Melastomataceae_Lavoisiera_subulata
Melastomataceae_Rhexia_salicifolia

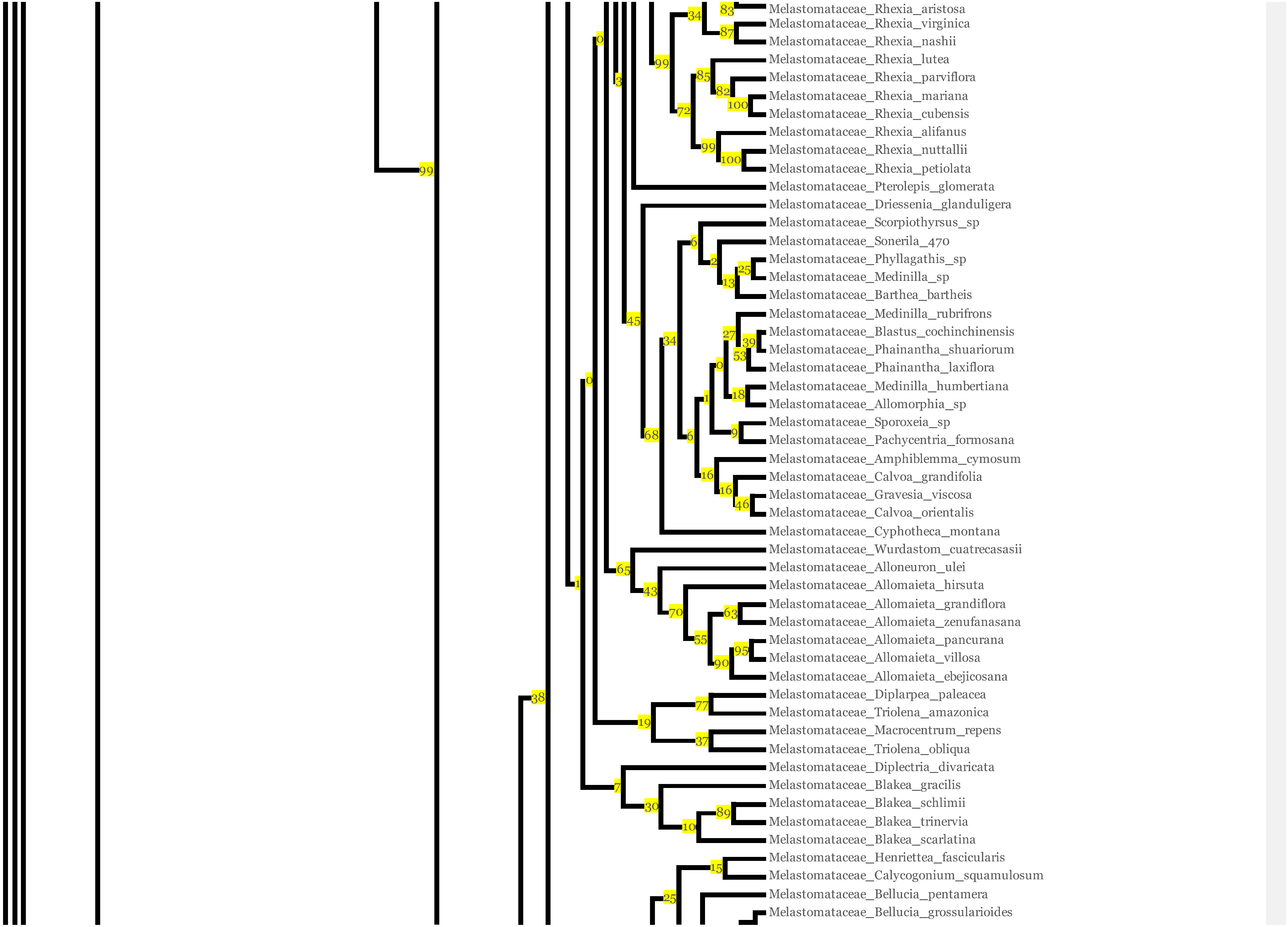
Melastomataceae_Rhexia_aristosa
Melastomataceae_Rhexia_virginica
Melastomataceae_Rhexia_nashii
Melastomataceae_Rhexia_lutea
Melastomataceae_Rhexia_parviflora
Melastomataceae_Rhexia_mariana
Melastomataceae_Rhexia_cubensis
Melastomataceae_Rhexia_alifanus
Melastomataceae_Rhexia_nuttallii
Melastomataceae_Rhexia_petiolata
Melastomataceae_Pterolepis_glomerata
Melastomataceae_Driessenia_glanduligera
Melastomataceae_Scorpiothyrsus_sp
Melastomataceae_Sonerila_470
Melastomataceae_Phyllagathis_sp
Melastomataceae_Medinilla_sp
Melastomataceae_Barthea_bartheis
Melastomataceae_Medinilla_rubrifrons
Melastomataceae_Blastus_cochinchinensis
Melastomataceae_Phainantha_shuariorum
Melastomataceae_Phainantha_laxiflora
Melastomataceae_Medinilla_humbertiana
Melastomataceae_Allomorphia_sp
Melastomataceae_Sporoxeia_sp
Melastomataceae_Pachycentria_formosana
Melastomataceae_Amphiblemma_cymosum
Melastomataceae_Calvoa_grandifolia
Melastomataceae_Gravesia_viscosa
Melastomataceae_Calvoa_orientalis
Melastomataceae_Cyphotheca_montana
Melastomataceae_Wurdastom_cuatrecasasii
Melastomataceae_Alloneuron_ulei
Melastomataceae_Allomaieta_hirsuta
Melastomataceae_Allomaieta_grandiflora
Melastomataceae_Allomaieta_zenufanasana
Melastomataceae_Allomaieta_pancurana
Melastomataceae_Allomaieta_villosa
Melastomataceae_Allomaieta_ebejicosana
Melastomataceae_Diplarpea_paleacea
Melastomataceae_Triolena_amazonica
Melastomataceae_Macrocentrum_repens
Melastomataceae_Triolena_obliqua
Melastomataceae_Diplectria_divaricata
Melastomataceae_Blakea_gracilis
Melastomataceae_Blakea_schlimii
Melastomataceae_Blakea_trinervia
Melastomataceae_Blakea_scarlatina
Melastomataceae_Henriettea_fascicularis
Melastomataceae_Calycogonium_squamulosum
Melastomataceae_Bellucia_pentamera
Melastomataceae_Bellucia_grossularioides

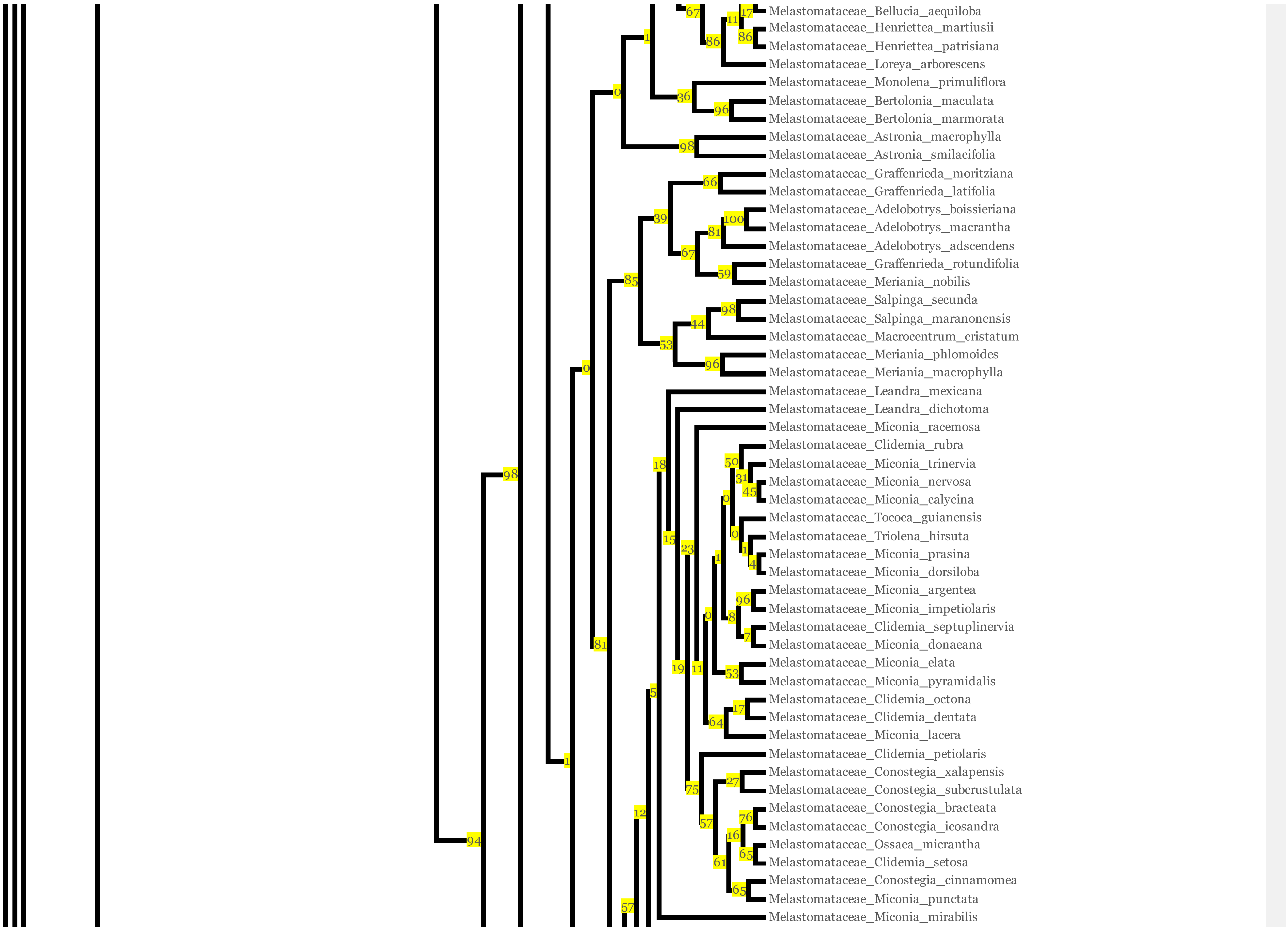
Melastomataceae_Bellucia_aequiloba
Melastomataceae_Henriettea_martiusii
Melastomataceae_Henriettea_patrisiana
Melastomataceae_Loreya_arborescens
Melastomataceae_Monolena_primuliflora
Melastomataceae_Bertolonia_maculata
Melastomataceae_Bertolonia_marmorata
Melastomataceae_Astronia_macrophylla
Melastomataceae_Astronia_smilacifolia
Melastomataceae_Graffenrieda_moritziana
Melastomataceae_Graffenrieda_latifolia
Melastomataceae_Adelobotrys_boissieriana
Melastomataceae_Adelobotrys_macrantha
Melastomataceae_Adelobotrys_adscendens
Melastomataceae_Graffenrieda_rotundifolia
Melastomataceae_Meriania_nobilis
Melastomataceae_Salpinga_secunda
Melastomataceae_Salpinga_maranonensis
Melastomataceae_Macrocentrum_cristatum
Melastomataceae_Meriania_phlomoides
Melastomataceae_Meriania_macrophylla
Melastomataceae_Leandra_mexicana
Melastomataceae_Leandra_dichotoma
Melastomataceae_Miconia_racemosa
Melastomataceae_Clidemia_rubra
Melastomataceae_Miconia_trinervia
Melastomataceae_Miconia_nervosa
Melastomataceae_Miconia_calycina
Melastomataceae_Tococa_guianensis
Melastomataceae_Triolena_hirsuta
Melastomataceae_Miconia_prasina
Melastomataceae_Miconia_dorsiloba
Melastomataceae_Miconia_argentea
Melastomataceae_Miconia_impetiolaris
Melastomataceae_Clidemia_septuplinervia
Melastomataceae_Miconia_donaeana
Melastomataceae_Miconia_elata
Melastomataceae_Miconia_pyramidalis
Melastomataceae_Clidemia_octona
Melastomataceae_Clidemia_dentata
Melastomataceae_Miconia_lacera
Melastomataceae_Clidemia_petiolaris
Melastomataceae_Conostegia_xalapensis
Melastomataceae_Conostegia_subcrustulata
Melastomataceae_Conostegia_bracteata
Melastomataceae_Conostegia_icosandra
Melastomataceae_Ossaea_micrantha
Melastomataceae_Clidemia_setosa
Melastomataceae_Conostegia_cinnamomea
Melastomataceae_Miconia_punctata
Melastomataceae_Miconia_mirabilis

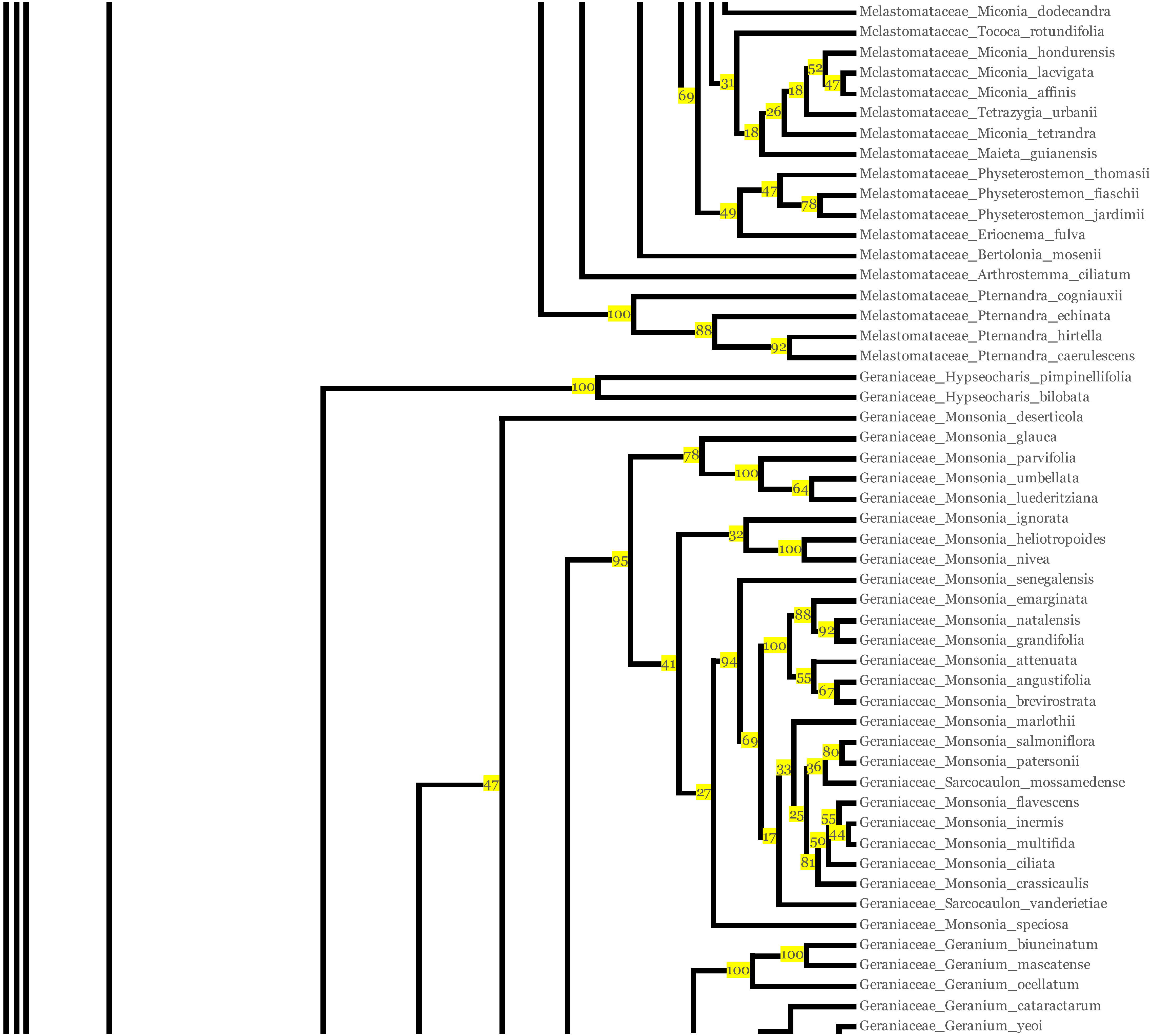
Melastomataceae_Miconia_dodecandra
Melastomataceae_Tococa_rotundifolia
Melastomataceae_Miconia_hondurensis
Melastomataceae_Miconia_laevigata
Melastomataceae_Miconia_affinis
Melastomataceae_Tetrazygia_urbanii
Melastomataceae_Miconia_tetrandra
Melastomataceae_Maieta_guianensis
Melastomataceae_Physeterostemon_thomasii
Melastomataceae_Physeterostemon_fiaschii
Melastomataceae_Physeterostemon_jardimii
Melastomataceae_Eriocnema_fulva
Melastomataceae_Bertolonia_mosenii
Melastomataceae_Arthrostemma_ciliatum
Melastomataceae_Pternandra_cogniauxii
Melastomataceae_Pternandra_echinata
Melastomataceae_Pternandra_hirtella
Melastomataceae_Pternandra_caerulescens
Geraniaceae_Hypseocharis_pimpinellifolia
Geraniaceae_Hypseocharis_bilobata
Geraniaceae_Monsonia_deserticola
Geraniaceae_Monsonia_glauca
Geraniaceae_Monsonia_parvifolia
Geraniaceae_Monsonia_umbellata
Geraniaceae_Monsonia_luederitziana
Geraniaceae_Monsonia_ignorata
Geraniaceae_Monsonia_heliotropoides
Geraniaceae_Monsonia_nivea
Geraniaceae_Monsonia_senegalensis
Geraniaceae_Monsonia_emarginata
Geraniaceae_Monsonia_natalensis
Geraniaceae_Monsonia_grandifolia
Geraniaceae_Monsonia_attenuata
Geraniaceae_Monsonia_angustifolia
Geraniaceae_Monsonia_brevirostrata
Geraniaceae_Monsonia_marlothii
Geraniaceae_Monsonia_salmoniflora
Geraniaceae_Monsonia_patersonii
Geraniaceae_Sarcocaulon_mossamedense
Geraniaceae_Monsonia_flavescens
Geraniaceae_Monsonia_inermis
Geraniaceae_Monsonia_multifida
Geraniaceae_Monsonia_ciliata
Geraniaceae_Monsonia_crassicaulis
Geraniaceae_Sarcocaulon_vanderietiae
Geraniaceae_Monsonia_speciosa
Geraniaceae_Geranium_biuncinatum
Geraniaceae_Geranium_mascatense
Geraniaceae_Geranium_ocellatum
Geraniaceae_Geranium_cataractarum
Geraniaceae_Geranium_yeoi

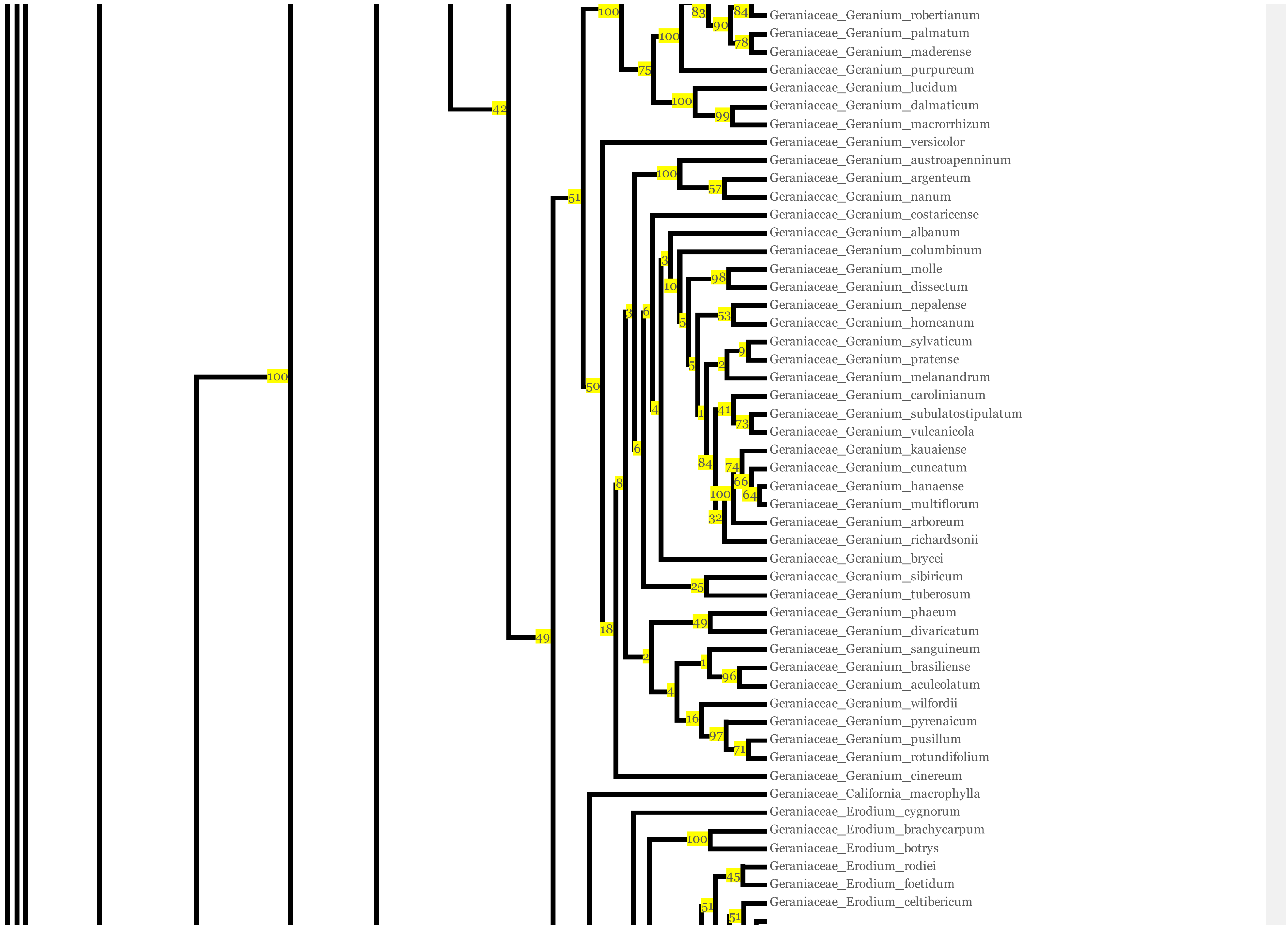
Geraniaceae_Geranium_robertianum
Geraniaceae_Geranium_palmatum
Geraniaceae_Geranium_maderense
Geraniaceae_Geranium_purpureum
Geraniaceae_Geranium_lucidum
Geraniaceae_Geranium_dalmaticum
Geraniaceae_Geranium_macrorrhizum
Geraniaceae_Geranium_versicolor
Geraniaceae_Geranium_austroapenninum
Geraniaceae_Geranium_argenteum
Geraniaceae_Geranium_nanum
Geraniaceae_Geranium_costaricense
Geraniaceae_Geranium_albanum
Geraniaceae_Geranium_columbinum
Geraniaceae_Geranium_molle
Geraniaceae_Geranium_dissectum
Geraniaceae_Geranium_nepalense
Geraniaceae_Geranium_homeanum
Geraniaceae_Geranium_sylvaticum
Geraniaceae_Geranium_pratense
Geraniaceae_Geranium_melanandrum
Geraniaceae_Geranium_carolinianum
Geraniaceae_Geranium_subulatostipulatum
Geraniaceae_Geranium_vulcanicola
Geraniaceae_Geranium_kauaiense
Geraniaceae_Geranium_cuneatum
Geraniaceae_Geranium_hanaense
Geraniaceae_Geranium_multiflorum
Geraniaceae_Geranium_arboreum
Geraniaceae_Geranium_richardsonii
Geraniaceae_Geranium_brycei
Geraniaceae_Geranium_sibiricum
Geraniaceae_Geranium_tuberosum
Geraniaceae_Geranium_phaeum
Geraniaceae_Geranium_divaricatum
Geraniaceae_Geranium_sanguineum
Geraniaceae_Geranium_brasiliense
Geraniaceae_Geranium_aculeolatum
Geraniaceae_Geranium_wilfordii
Geraniaceae_Geranium_pyrenaicum
Geraniaceae_Geranium_pusillum
Geraniaceae_Geranium_rotundifolium
Geraniaceae_Geranium_cinereum
Geraniaceae_California_macrophylla
Geraniaceae_Erodium_cygnorum
Geraniaceae_Erodium_brachycarpum
Geraniaceae_Erodium_botrys
Geraniaceae_Erodium_rodiei
Geraniaceae_Erodium_foetidum
Geraniaceae_Erodium_celtibericum

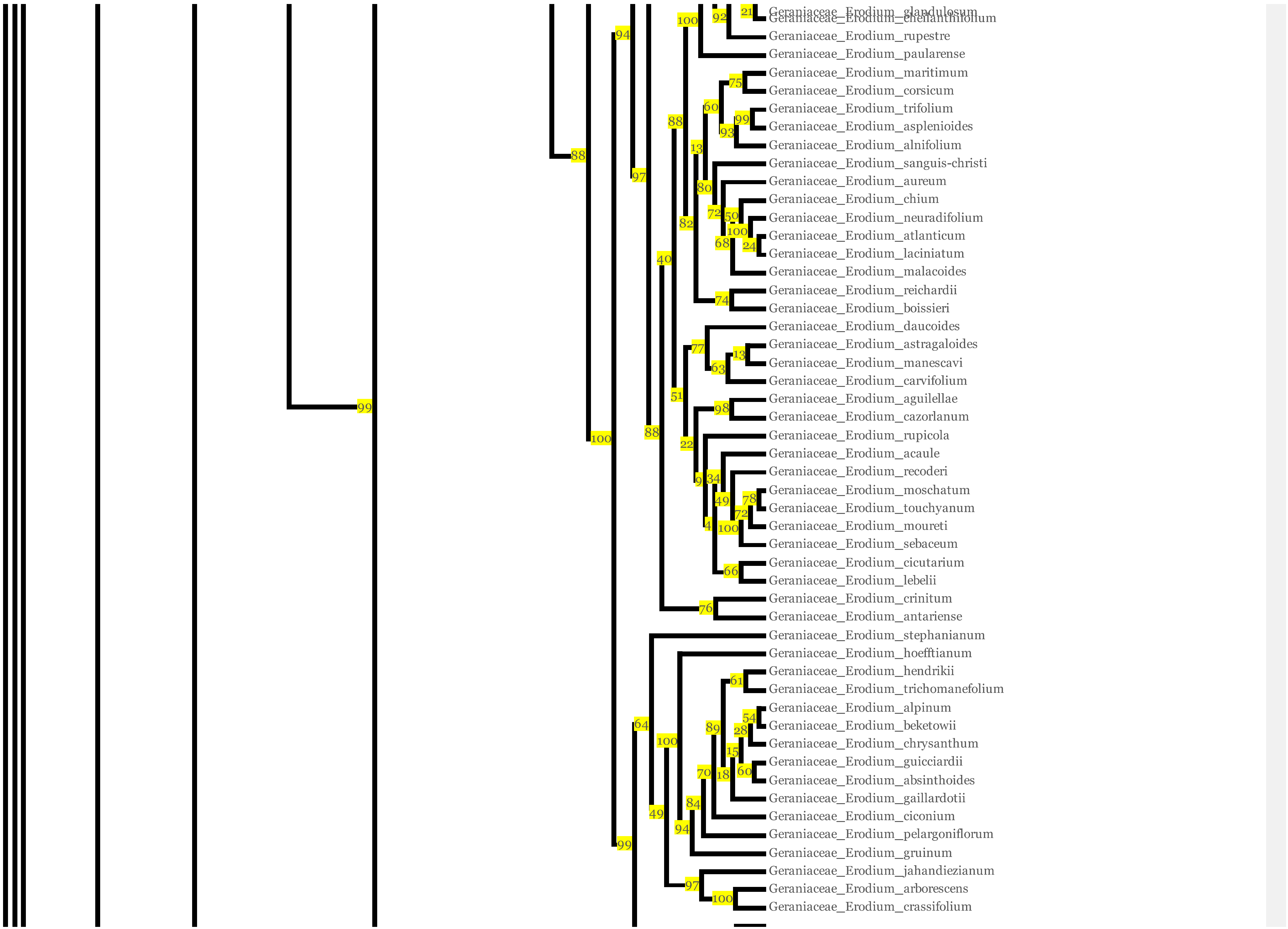

Geraniaceae_Erodium_glandulosum
Geraniaceae_Erodium_cheilanthifolium
Geraniaceae_Erodium_rupestre
Geraniaceae_Erodium_paularense
Geraniaceae_Erodium_maritimum
Geraniaceae_Erodium_corsicum
Geraniaceae_Erodium_trifolium
Geraniaceae_Erodium_asplenioides
Geraniaceae_Erodium_alnifolium
Geraniaceae_Erodium_sanguis-christi
Geraniaceae_Erodium_aureum
Geraniaceae_Erodium_chium
Geraniaceae_Erodium_neuradifolium
Geraniaceae_Erodium_atlanticum
Geraniaceae_Erodium_laciniatum
Geraniaceae_Erodium_malacoides
Geraniaceae_Erodium_reichardii
Geraniaceae_Erodium_boissieri
Geraniaceae_Erodium_daucoides
Geraniaceae_Erodium_astragaloides
Geraniaceae_Erodium_manescavi
Geraniaceae_Erodium_carvifolium
Geraniaceae_Erodium_aguilellae
Geraniaceae_Erodium_cazorlanum
Geraniaceae_Erodium_rupicola
Geraniaceae_Erodium_acaule
Geraniaceae_Erodium_recoderi
Geraniaceae_Erodium_moschatum
Geraniaceae_Erodium_touchyanum
Geraniaceae_Erodium_moureti
Geraniaceae_Erodium_sebaceum
Geraniaceae_Erodium_cicutarium
Geraniaceae_Erodium_lebelii
Geraniaceae_Erodium_crinitum
Geraniaceae_Erodium_antariense
Geraniaceae_Erodium_stephanianum
Geraniaceae_Erodium_hoefftianum
Geraniaceae_Erodium_hendrikii
Geraniaceae_Erodium_trichomanefolium
Geraniaceae_Erodium_alpinum
Geraniaceae_Erodium_beketowii
Geraniaceae_Erodium_chrysanthum
Geraniaceae_Erodium_guicciardii
Geraniaceae_Erodium_absinthoides
Geraniaceae_Erodium_gaillardotii
Geraniaceae_Erodium_ciconium
Geraniaceae_Erodium_pelargoniflorum
Geraniaceae_Erodium_gruinum
Geraniaceae_Erodium_jahandiezianum
Geraniaceae_Erodium_arborescens
Geraniaceae_Erodium_crassifolium

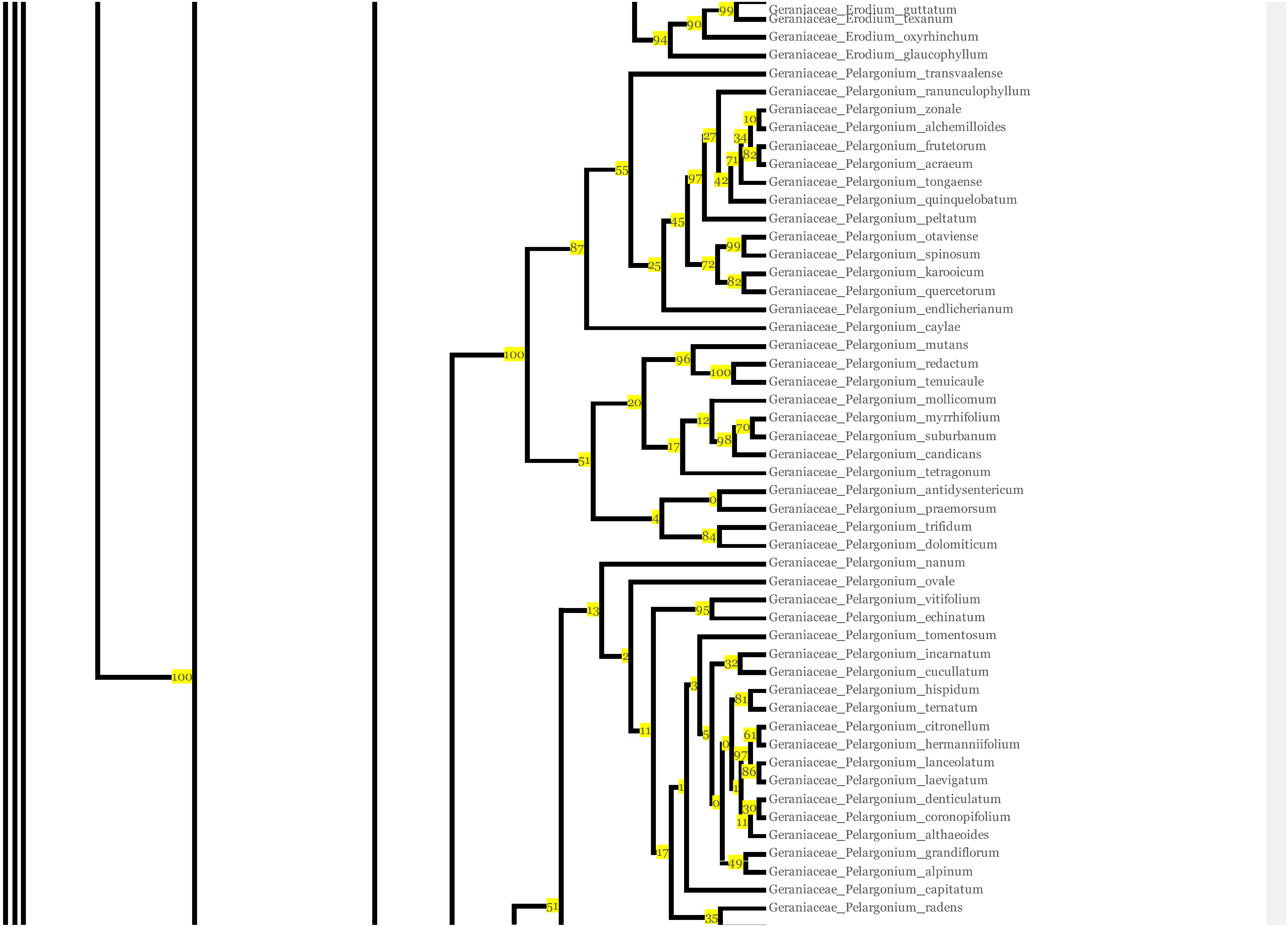

Geraniaceae_Erodium_guttatum
Geraniaceae_Erodium_texanum
Geraniaceae_Erodium_oxyrhinchum
Geraniaceae_Erodium_glaucophyllum
Geraniaceae_Pelargonium_transvaalense
Geraniaceae_Pelargonium_ranunculophyllum
Geraniaceae_Pelargonium_zonale
Geraniaceae_Pelargonium_alchemilloides
Geraniaceae_Pelargonium_frutetorum
Geraniaceae_Pelargonium_acraeum
Geraniaceae_Pelargonium_tongaense
Geraniaceae_Pelargonium_quinquelobatum
Geraniaceae_Pelargonium_peltatum
Geraniaceae_Pelargonium_otaviense
Geraniaceae_Pelargonium_spinosum
Geraniaceae_Pelargonium_karooicum
Geraniaceae_Pelargonium_quercetorum
Geraniaceae_Pelargonium_endlicherianum
Geraniaceae_Pelargonium_caylae
Geraniaceae_Pelargonium_mutans
Geraniaceae_Pelargonium_redactum
Geraniaceae_Pelargonium_tenuicaule
Geraniaceae_Pelargonium_mollicomum
Geraniaceae_Pelargonium_myrrhifolium
Geraniaceae_Pelargonium_suburbanum
Geraniaceae_Pelargonium_candicans
Geraniaceae_Pelargonium_tetragonum
Geraniaceae_Pelargonium_antidysentericum
Geraniaceae_Pelargonium_praemorsum
Geraniaceae_Pelargonium_trifidum
Geraniaceae_Pelargonium_dolomiticum
Geraniaceae_Pelargonium_nanum
Geraniaceae_Pelargonium_ovale
Geraniaceae_Pelargonium_vitifolium
Geraniaceae_Pelargonium_echinatum
Geraniaceae_Pelargonium_tomentosum
Geraniaceae_Pelargonium_incarnatum
Geraniaceae_Pelargonium_cucullatum
Geraniaceae_Pelargonium_hispidum
Geraniaceae_Pelargonium_ternatum
Geraniaceae_Pelargonium_citronellum
Geraniaceae_Pelargonium_hermanniifolium
Geraniaceae_Pelargonium_lanceolatum
Geraniaceae_Pelargonium_laevigatum
Geraniaceae_Pelargonium_denticulatum
Geraniaceae_Pelargonium_coronopifolium
Geraniaceae_Pelargonium_althaeoides
Geraniaceae_Pelargonium_grandiflorum
Geraniaceae_Pelargonium_alpinum
Geraniaceae_Pelargonium_capitatum
Geraniaceae_Pelargonium_radens

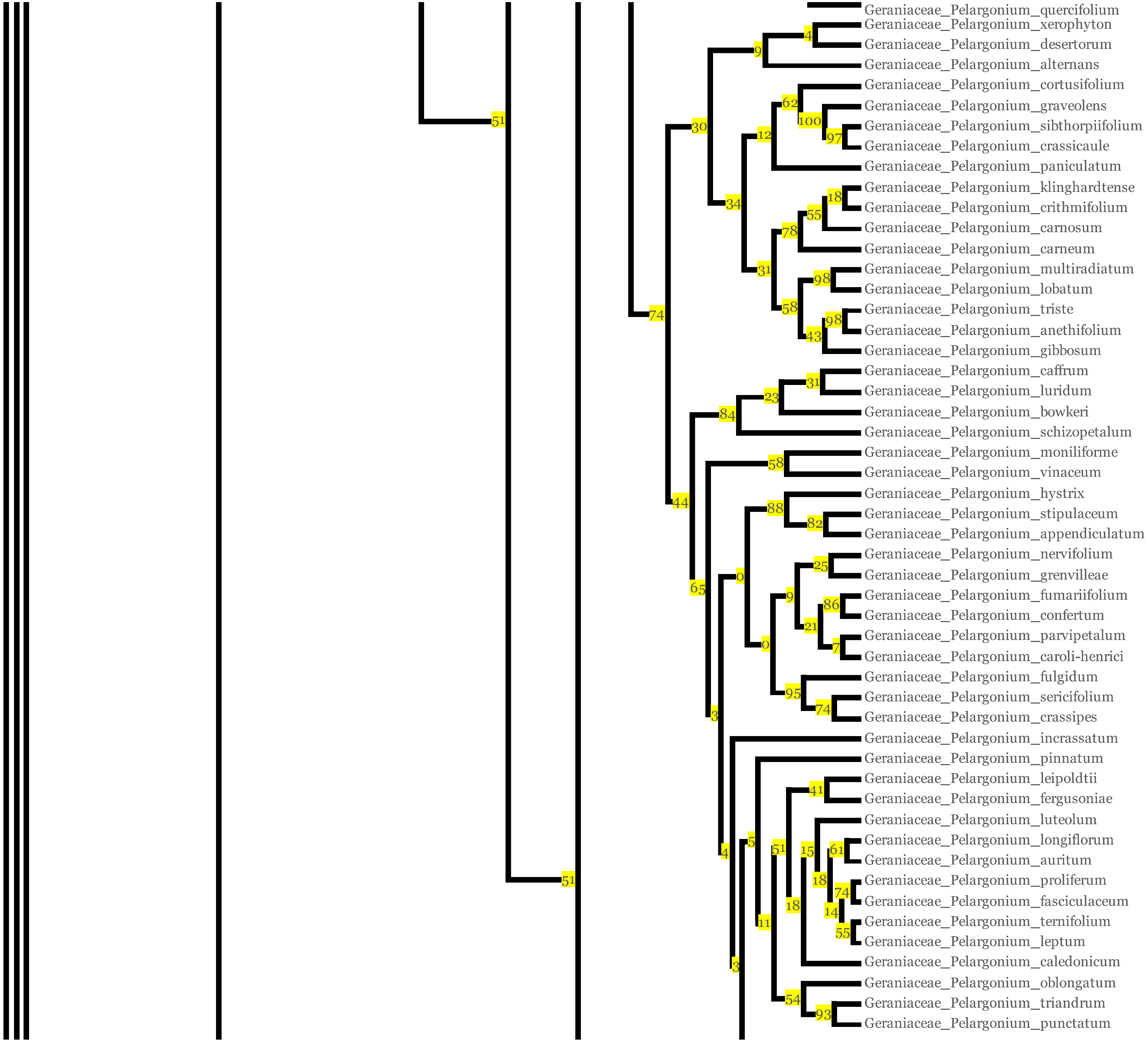
Geraniaceae_Pelargonium_quercifolium
Geraniaceae_Pelargonium_xerophyton
Geraniaceae_Pelargonium_desertorum
Geraniaceae_Pelargonium_alternans
Geraniaceae_Pelargonium_cortusifolium
Geraniaceae_Pelargonium_graveolens
Geraniaceae_Pelargonium_sibthorpiifolium
Geraniaceae_Pelargonium_crassicaule
Geraniaceae_Pelargonium_paniculatum
Geraniaceae_Pelargonium_klinghardtense
Geraniaceae_Pelargonium_crithmifolium
Geraniaceae_Pelargonium_carnosum
Geraniaceae_Pelargonium_carneum
Geraniaceae_Pelargonium_multiradiatum
Geraniaceae_Pelargonium_lobatum
Geraniaceae_Pelargonium_triste
Geraniaceae_Pelargonium_anethifolium
Geraniaceae_Pelargonium_gibbosum
Geraniaceae_Pelargonium_caffrum
Geraniaceae_Pelargonium_luridum
Geraniaceae_Pelargonium_bowkeri
Geraniaceae_Pelargonium_schizopetalum
Geraniaceae_Pelargonium_moniliforme
Geraniaceae_Pelargonium_vinaceum
Geraniaceae_Pelargonium_hystrix
Geraniaceae_Pelargonium_stipulaceum
Geraniaceae_Pelargonium_appendiculatum
Geraniaceae_Pelargonium_nervifolium
Geraniaceae_Pelargonium_grenvilleae
Geraniaceae_Pelargonium_fumariifolium
Geraniaceae_Pelargonium_confertum
Geraniaceae_Pelargonium_parvipetalum
Geraniaceae_Pelargonium_caroli-henrici
Geraniaceae_Pelargonium_fulgidum
Geraniaceae_Pelargonium_sericifolium
Geraniaceae_Pelargonium_crassipes
Geraniaceae_Pelargonium_incrassatum
Geraniaceae_Pelargonium_pinnatum
Geraniaceae_Pelargonium_leipoldtii
Geraniaceae_Pelargonium_fergusoniae
Geraniaceae_Pelargonium_luteolum
Geraniaceae_Pelargonium_longiflorum
Geraniaceae_Pelargonium_auritum
Geraniaceae_Pelargonium_proliferum
Geraniaceae_Pelargonium_fasciculaceum
Geraniaceae_Pelargonium_ternifolium
Geraniaceae_Pelargonium_leptum
Geraniaceae_Pelargonium_caledonicum
Geraniaceae_Pelargonium_oblongatum
Geraniaceae_Pelargonium_triandrum
Geraniaceae_Pelargonium_punctatum

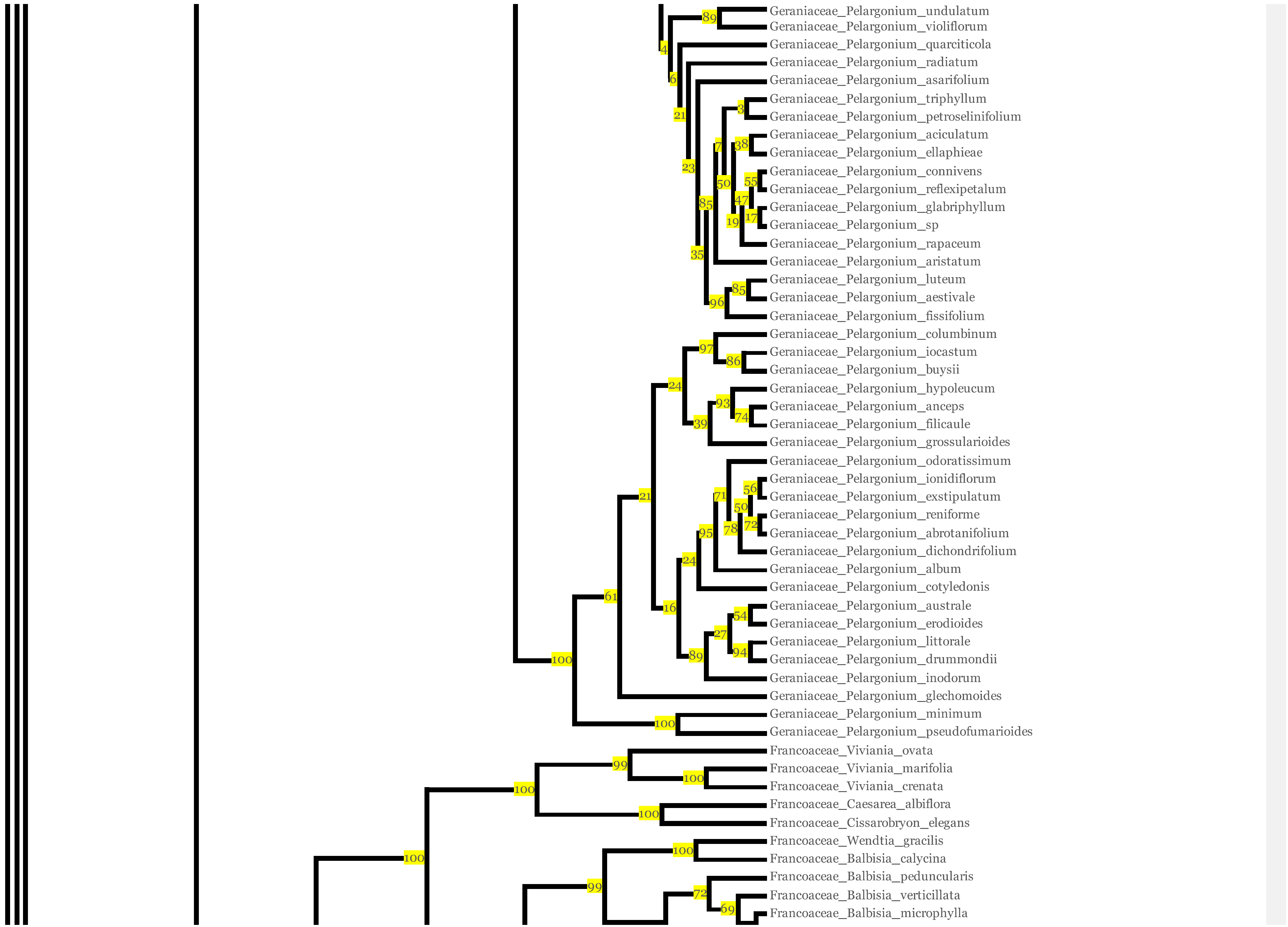

Geraniaceae_Pelargonium_undulatum
Geraniaceae_Pelargonium_violiflorum
Geraniaceae_Pelargonium_quarciticola
Geraniaceae_Pelargonium_radiatum
Geraniaceae_Pelargonium_asarifolium
Geraniaceae_Pelargonium_triphyllum
Geraniaceae_Pelargonium_petroselinifolium
Geraniaceae_Pelargonium_aciculatum
Geraniaceae_Pelargonium_ellaphieae
Geraniaceae_Pelargonium_connivens
Geraniaceae_Pelargonium_reflexipetalum
Geraniaceae_Pelargonium_glabriphyllum
Geraniaceae_Pelargonium_sp
Geraniaceae_Pelargonium_rapaceum
Geraniaceae_Pelargonium_aristatum
Geraniaceae_Pelargonium_luteum
Geraniaceae_Pelargonium_aestivale
Geraniaceae_Pelargonium_fissifolium
Geraniaceae_Pelargonium_columbinum
Geraniaceae_Pelargonium_iocastum
Geraniaceae_Pelargonium_buysii
Geraniaceae_Pelargonium_hypoleucum
Geraniaceae_Pelargonium_anceps
Geraniaceae_Pelargonium_filicaule
Geraniaceae_Pelargonium_grossularioides
Geraniaceae_Pelargonium_odoratissimum
Geraniaceae_Pelargonium_ionidiflorum
Geraniaceae_Pelargonium_exstipulatum
Geraniaceae_Pelargonium_reniforme
Geraniaceae_Pelargonium_abrotanifolium
Geraniaceae_Pelargonium_dichondrifolium
Geraniaceae_Pelargonium_album
Geraniaceae_Pelargonium_cotyledonis
Geraniaceae_Pelargonium_australe
Geraniaceae_Pelargonium_erodioides
Geraniaceae_Pelargonium_littorale
Geraniaceae_Pelargonium_drummondii
Geraniaceae_Pelargonium_inodorum
Geraniaceae_Pelargonium_glechomoides
Geraniaceae_Pelargonium_minimum
Geraniaceae_Pelargonium_pseudofumarioides
Francoaceae_Viviania_ovata
Francoaceae_Viviania_marifolia
Francoaceae_Viviania_crenata
Francoaceae_Caesarea_albiflora
Francoaceae_Cissarobryon_elegans
Francoaceae_Wendtia_gracilis
Francoaceae_Balbisia_calycina
Francoaceae_Balbisia_peduncularis
Francoaceae_Balbisia_verticillata
Francoaceae_Balbisia_microphylla

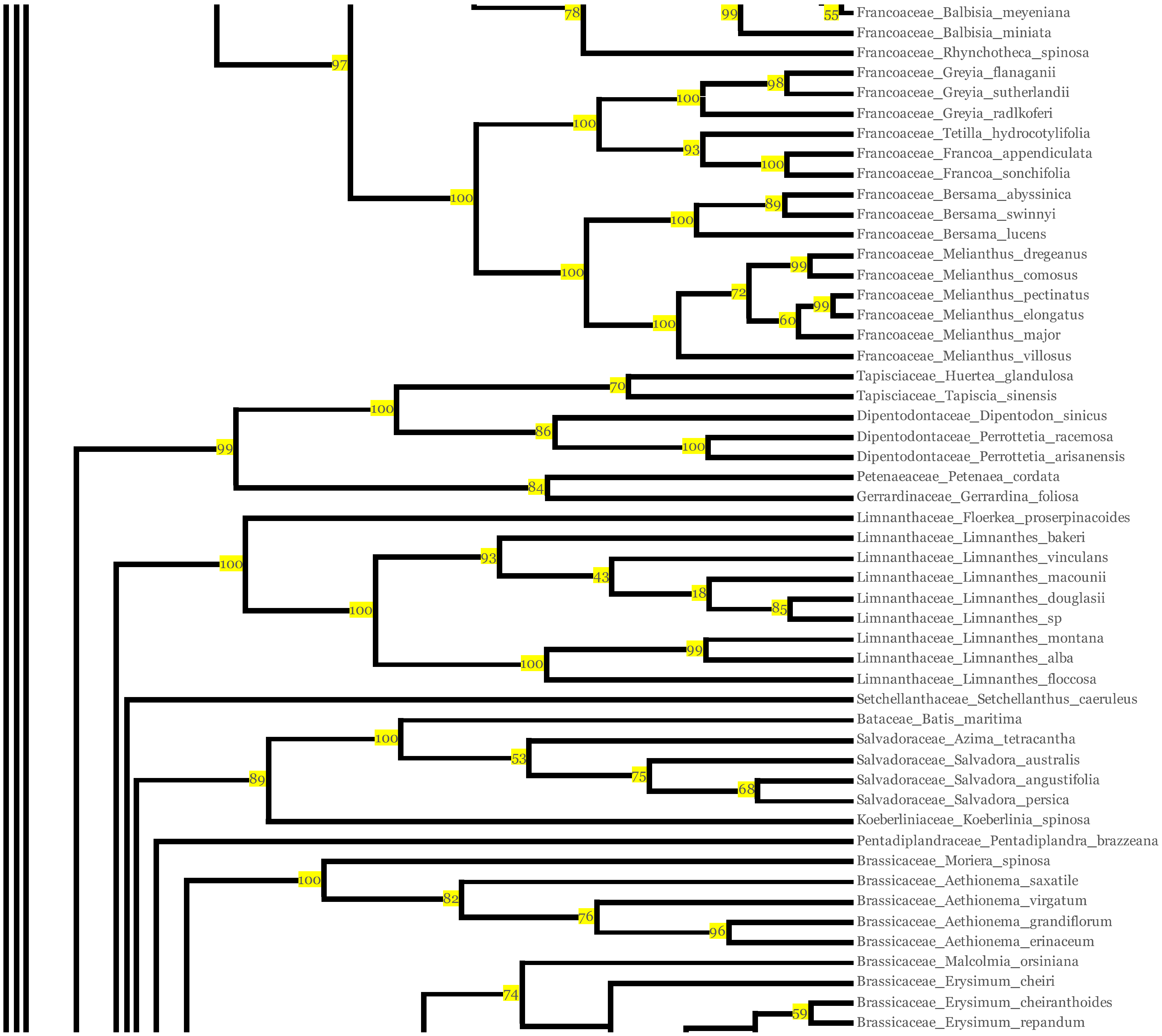
Francoaceae_Balbisia_meyeniana
Francoaceae_Balbisia_miniata
Francoaceae_Rhynchotheca_spinosa
Francoaceae_Greyia_flanaganii
Francoaceae_Greyia_sutherlandii
Francoaceae_Greyia_radlkoferi
Francoaceae_Tetilla_hydrocotylifolia
Francoaceae_Francoa_appendiculata
Francoaceae_Francoa_sonchifolia
Francoaceae_Bersama_abyssinica
Francoaceae_Bersama_swinnyi
Francoaceae_Bersama_lucens
Francoaceae_Melianthus_dregeanus
Francoaceae_Melianthus_comosus
Francoaceae_Melianthus_pectinatus
Francoaceae_Melianthus_elongatus
Francoaceae_Melianthus_major
Francoaceae_Melianthus_villosus
Tapisciaceae_Huertea_glandulosa
Tapisciaceae_Tapiscia_sinensis
Dipentodontaceae_Dipentodon_sinicus
Dipentodontaceae_Perrottetia_racemosa
Dipentodontaceae_Perrottetia_arisanensis
Petenaeaceae_Petenaea_cordata
Gerrardinaceae_Gerrardina_foliosa
Limnanthaceae_Floerkea_proserpinacoides
Limnanthaceae_Limnanthes_bakeri
Limnanthaceae_Limnanthes_vinculans
Limnanthaceae_Limnanthes_macounii
Limnanthaceae_Limnanthes_douglasii
Limnanthaceae_Limnanthes_sp
Limnanthaceae_Limnanthes_montana
Limnanthaceae_Limnanthes_alba
Limnanthaceae_Limnanthes_floccosa
Setchellanthaceae_Setchellanthus_caeruleus
Bataceae_Batis_maritima
Salvadoraceae_Azima_tetracantha
Salvadoraceae_Salvadora_australis
Salvadoraceae_Salvadora_angustifolia
Salvadoraceae_Salvadora_persica
Koeberliniaceae_Koeberlinia_spinosa
Pentadiplandraceae_Pentadiplandra_brazzeana
Brassicaceae_Moriera_spinosa
Brassicaceae_Aethionema_saxatile
Brassicaceae_Aethionema_virgatum
Brassicaceae_Aethionema_grandiflorum
Brassicaceae_Aethionema_erinaceum
Brassicaceae_Malcolmia_orsiniana
Brassicaceae_Erysimum_cheiri
Brassicaceae_Erysimum_cheiranthoides
Brassicaceae_Erysimum_repandum

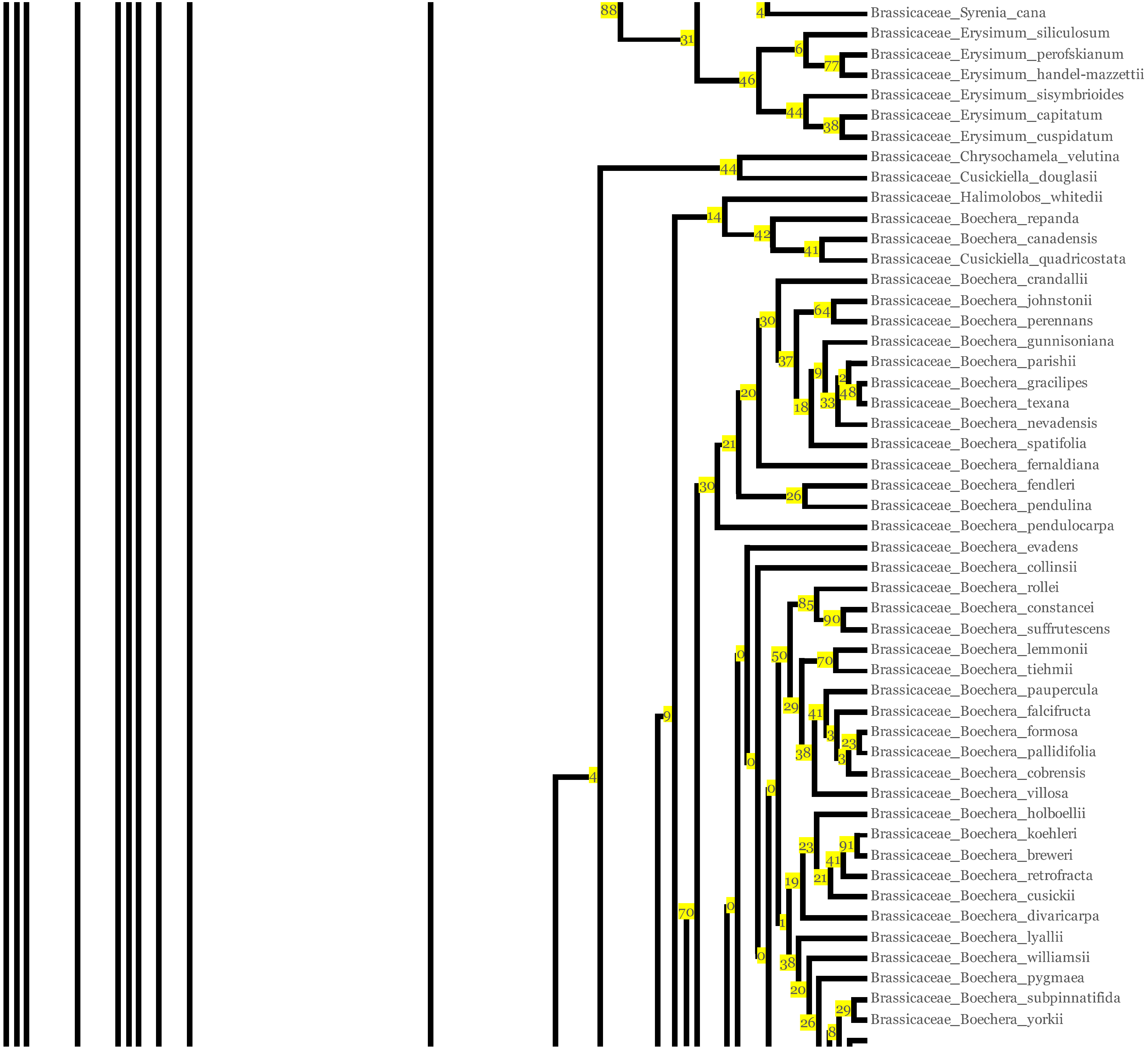
Brassicaceae_Syrenia_cana
Brassicaceae_Erysimum_siliculosum
Brassicaceae_Erysimum_perofskianum
Brassicaceae_Erysimum_handel-mazzettii
Brassicaceae_Erysimum_sisymbrioides
Brassicaceae_Erysimum_capitatum
Brassicaceae_Erysimum_cuspidatum
Brassicaceae_Chrysochamela_velutina
Brassicaceae_Cusickiella_douglasii
Brassicaceae_Halimolobos_whitedii
Brassicaceae_Boechera_repanda
Brassicaceae_Boechera_canadensis
Brassicaceae_Cusickiella_quadricostata
Brassicaceae_Boechera_crandallii
Brassicaceae_Boechera_johnstonii
Brassicaceae_Boechera_perennans
Brassicaceae_Boechera_gunnisoniana
Brassicaceae_Boechera_parishii
Brassicaceae_Boechera_gracilipes
Brassicaceae_Boechera_texana
Brassicaceae_Boechera_nevadensis
Brassicaceae_Boechera_spatifolia
Brassicaceae_Boechera_fernaldiana
Brassicaceae_Boechera_fendleri
Brassicaceae_Boechera_pendulina
Brassicaceae_Boechera_pendulocarpa
Brassicaceae_Boechera_evadens
Brassicaceae_Boechera_collinsii
Brassicaceae_Boechera_rollei
Brassicaceae_Boechera_constancei
Brassicaceae_Boechera_suffrutescens
Brassicaceae_Boechera_lemmonii
Brassicaceae_Boechera_tiehmii
Brassicaceae_Boechera_paupercula
Brassicaceae_Boechera_falcifructa
Brassicaceae_Boechera_formosa
Brassicaceae_Boechera_pallidifolia
Brassicaceae_Boechera_cobrensis
Brassicaceae_Boechera_villosa
Brassicaceae_Boechera_holboellii
Brassicaceae_Boechera_koehleri
Brassicaceae_Boechera_breweri
Brassicaceae_Boechera_retrofracta
Brassicaceae_Boechera_cusickii
Brassicaceae_Boechera_divaricarpa
Brassicaceae_Boechera_lyallii
Brassicaceae_Boechera_williamsii
Brassicaceae_Boechera_pygmaea
Brassicaceae_Boechera_subpinnatifida
Brassicaceae_Boechera_yorkii

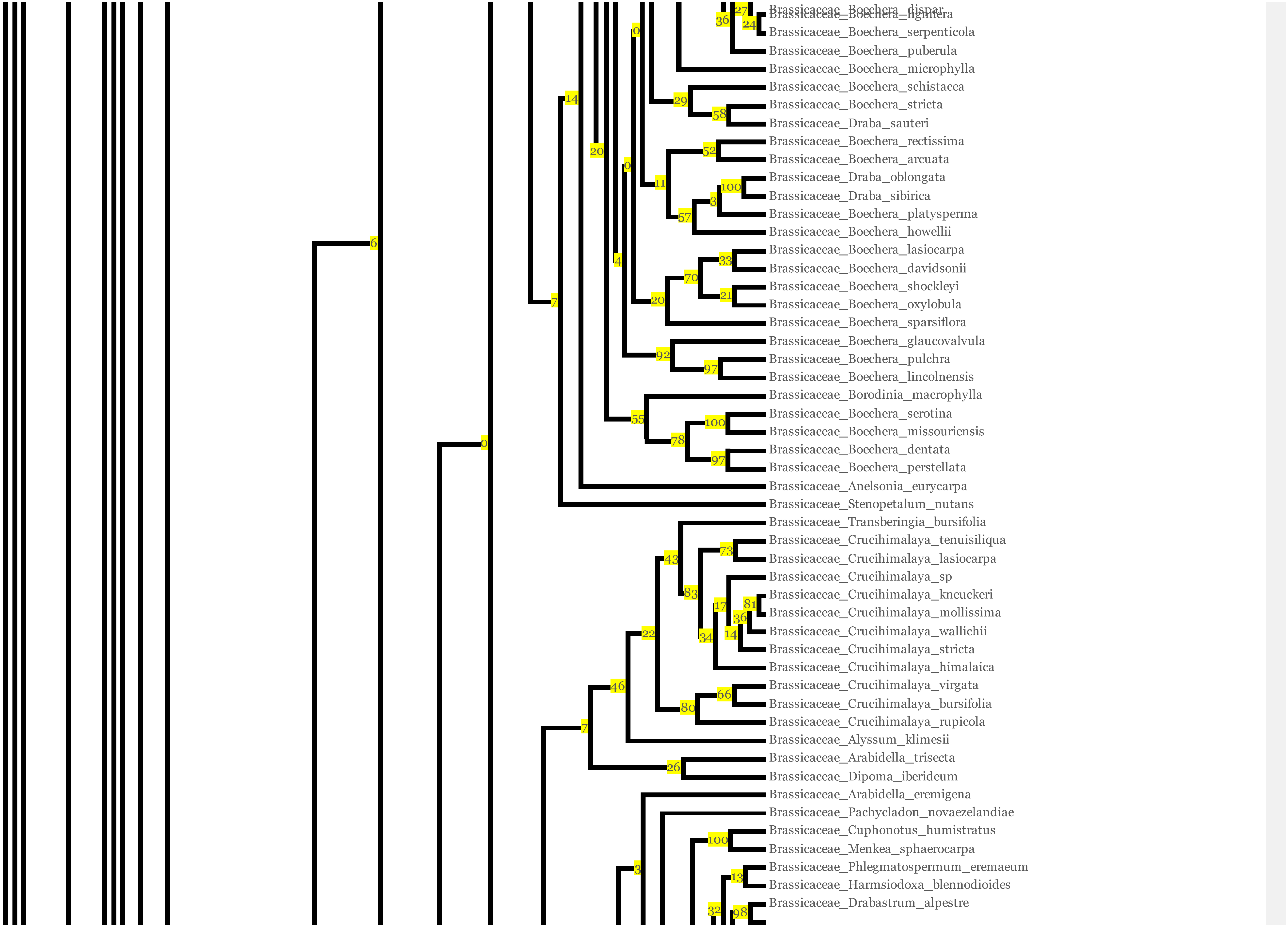

Brassicaceae_Boechera_dispar
Brassicaceae_Boechera_lignifera
Brassicaceae_Boechera_serpenticola
Brassicaceae_Boechera_puberula
Brassicaceae_Boechera_microphylla
Brassicaceae_Boechera_schistacea
Brassicaceae_Boechera_stricta
Brassicaceae_Draba_sauteri
Brassicaceae_Boechera_rectissima
Brassicaceae_Boechera_arcuata
Brassicaceae_Draba_oblongata
Brassicaceae_Draba_sibirica
Brassicaceae_Boechera_platysperma
Brassicaceae_Boechera_howellii
Brassicaceae_Boechera_lasiocarpa
Brassicaceae_Boechera_davidsonii
Brassicaceae_Boechera_shockleyi
Brassicaceae_Boechera_oxylobula
Brassicaceae_Boechera_sparsiflora
Brassicaceae_Boechera_glaucovalvula
Brassicaceae_Boechera_pulchra
Brassicaceae_Boechera_lincolnensis
Brassicaceae_Borodinia_macrophylla
Brassicaceae_Boechera_serotina
Brassicaceae_Boechera_missouriensis
Brassicaceae_Boechera_dentata
Brassicaceae_Boechera_perstellata
Brassicaceae_Anelsonia_eurycarpa
Brassicaceae_Stenopetalum_nutans
Brassicaceae_Transberingia_bursifolia
Brassicaceae_Crucihimalaya_tenuisiliqua
Brassicaceae_Crucihimalaya_lasiocarpa
Brassicaceae_Crucihimalaya_sp
Brassicaceae_Crucihimalaya_kneuckeri
Brassicaceae_Crucihimalaya_mollissima
Brassicaceae_Crucihimalaya_wallichii
Brassicaceae_Crucihimalaya_stricta
Brassicaceae_Crucihimalaya_himalaica
Brassicaceae_Crucihimalaya_virgata
Brassicaceae_Crucihimalaya_bursifolia
Brassicaceae_Crucihimalaya_rupicola
Brassicaceae_Alyssum_klimesii
Brassicaceae_Arabidella_trisecta
Brassicaceae_Dipoma_iberideum
Brassicaceae_Arabidella_eremigena
Brassicaceae_Pachycladon_novaezelandiae
Brassicaceae_Cuphonotus_humistratus
Brassicaceae_Menkea_sphaerocarpa
Brassicaceae_Phlegmatospermum_eremaeum
Brassicaceae_Harmsiodoxa_blennodioides
Brassicaceae_Drabastrum_alpestre

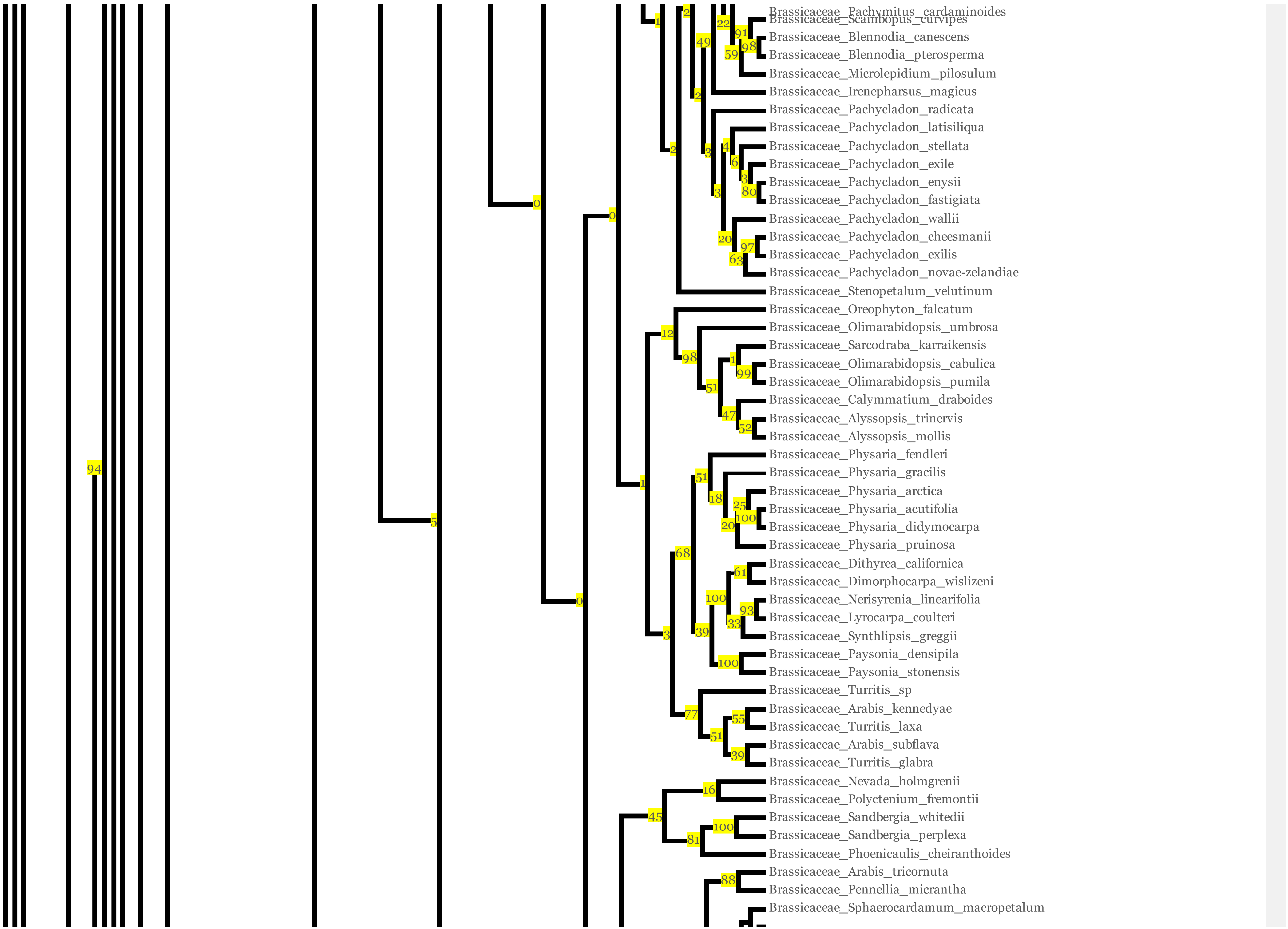
Brassicaceae_Pachymitus_cardaminoides
Brassicaceae_Scambopus_curvipes
Brassicaceae_Blennodia_canescens
Brassicaceae_Blennodia_pterosperma
Brassicaceae_Microlepidium_pilosulum
Brassicaceae_Irenepharsus_magicus
Brassicaceae_Pachycladon_radicata
Brassicaceae_Pachycladon_latisiliqua
Brassicaceae_Pachycladon_stellata
Brassicaceae_Pachycladon_exile
Brassicaceae_Pachycladon_enysii
Brassicaceae_Pachycladon_fastigiata
Brassicaceae_Pachycladon_wallii
Brassicaceae_Pachycladon_cheesmanii
Brassicaceae_Pachycladon_exilis
Brassicaceae_Pachycladon_novae-zelandiae
Brassicaceae_Stenopetalum_velutinum
Brassicaceae_Oreophyton_falcatum
Brassicaceae_Olimarabidopsis_umbrosa
Brassicaceae_Sarcodraba_karraikensis
Brassicaceae_Olimarabidopsis_cabulica
Brassicaceae_Olimarabidopsis_pumila
Brassicaceae_Calymmatium_draboides
Brassicaceae_Alyssopsis_trinervis
Brassicaceae_Alyssopsis_mollis
Brassicaceae_Physaria_fendleri
Brassicaceae_Physaria_gracilis
Brassicaceae_Physaria_arctica
Brassicaceae_Physaria_acutifolia
Brassicaceae_Physaria_didymocarpa
Brassicaceae_Physaria_pruinosa
Brassicaceae_Dithyrea_californica
Brassicaceae_Dimorphocarpa_wislizeni
Brassicaceae_Nerisyrenia_linearifolia
Brassicaceae_Lyrocarpa_coulteri
Brassicaceae_Synthlipsis_greggii
Brassicaceae_Paysonia_densipila
Brassicaceae_Paysonia_stonensis
Brassicaceae_Turritis_sp
Brassicaceae_Arabis_kennedyae
Brassicaceae_Turritis_laxa
Brassicaceae_Arabis_subflava
Brassicaceae_Turritis_glabra
Brassicaceae_Nevada_holmgrenii
Brassicaceae_Polyctenium_fremontii
Brassicaceae_Sandbergia_whitedii
Brassicaceae_Sandbergia_perplexa
Brassicaceae_Phoenicaulis_cheiranthoides
Brassicaceae_Arabis_tricornuta
Brassicaceae_Pennellia_micrantha
Brassicaceae_Sphaerocardamum_macropetalum

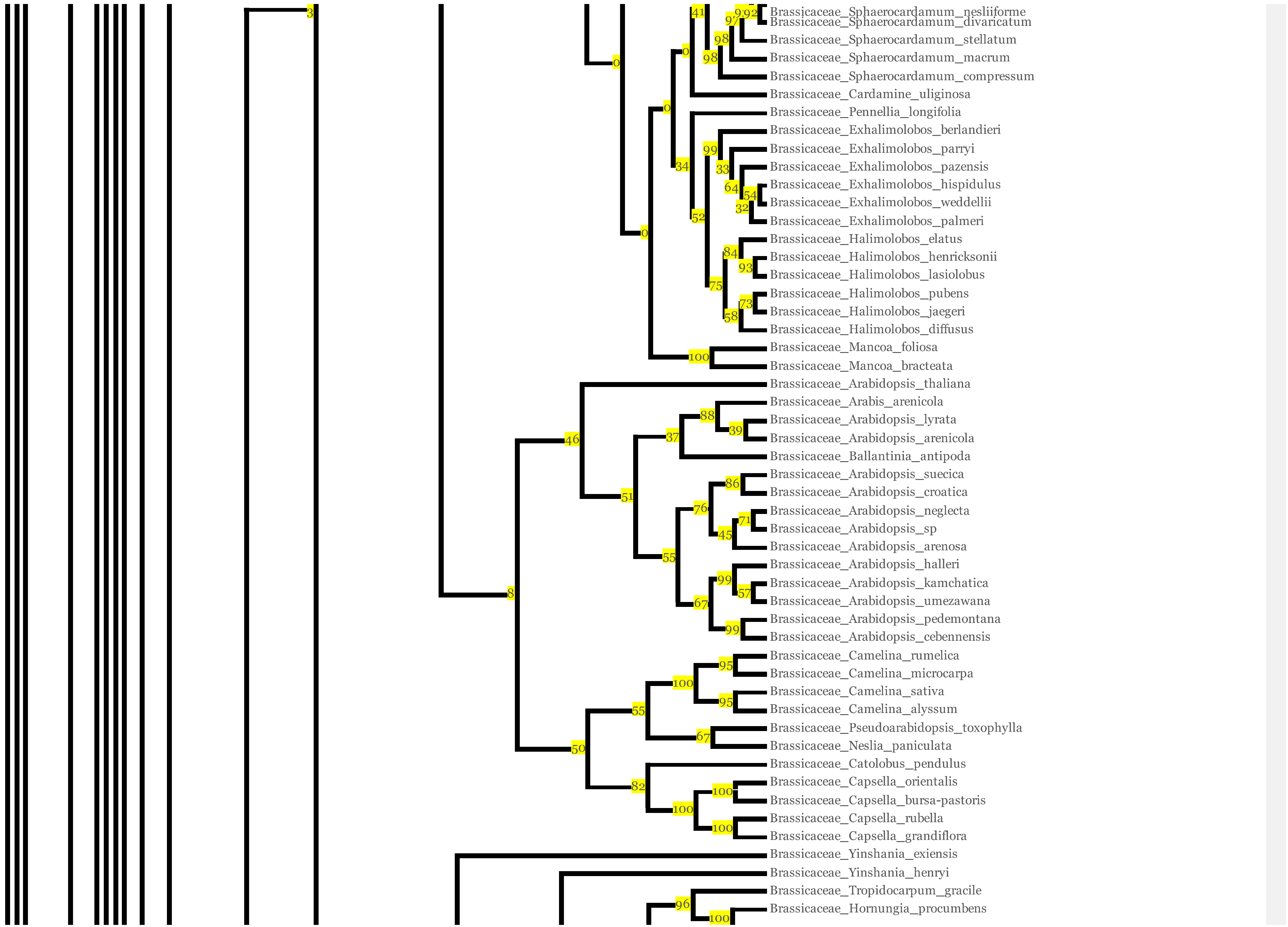
Brassicaceae_Sphaerocardamum_nesliiforme
Brassicaceae_Sphaerocardamum_divaricatum
Brassicaceae_Sphaerocardamum_stellatum
Brassicaceae_Sphaerocardamum_macrum
Brassicaceae_Sphaerocardamum_compressum
Brassicaceae_Cardamine_uliginosa
Brassicaceae_Pennellia_longifolia
Brassicaceae_Exhalimolobos_berlandieri
Brassicaceae_Exhalimolobos_parryi
Brassicaceae_Exhalimolobos_pazensis
Brassicaceae_Exhalimolobos_hispidulus
Brassicaceae_Exhalimolobos_weddellii
Brassicaceae_Exhalimolobos_palmeri
Brassicaceae_Halimolobos_elatus
Brassicaceae_Halimolobos_henricksonii
Brassicaceae_Halimolobos_lasiolobus
Brassicaceae_Halimolobos_pubens
Brassicaceae_Halimolobos_jaegeri
Brassicaceae_Halimolobos_diffusus
Brassicaceae_Mancoa_foliosa
Brassicaceae_Mancoa_bracteata
Brassicaceae_Arabidopsis_thaliana
Brassicaceae_Arabis_arenicola
Brassicaceae_Arabidopsis_lyrata
Brassicaceae_Arabidopsis_arenicola
Brassicaceae_Ballantinia_antipoda
Brassicaceae_Arabidopsis_suecica
Brassicaceae_Arabidopsis_croatica
Brassicaceae_Arabidopsis_neglecta
Brassicaceae_Arabidopsis_sp
Brassicaceae_Arabidopsis_arenosa
Brassicaceae_Arabidopsis_halleri
Brassicaceae_Arabidopsis_kamchatica
Brassicaceae_Arabidopsis_umezawana
Brassicaceae_Arabidopsis_pedemontana
Brassicaceae_Arabidopsis_cebennensis
Brassicaceae_Camelina_rumelica
Brassicaceae_Camelina_microcarpa
Brassicaceae_Camelina_sativa
Brassicaceae_Camelina_alyssum
Brassicaceae_Pseudoarabidopsis_toxophylla
Brassicaceae_Neslia_paniculata
Brassicaceae_Catolobus_pendulus
Brassicaceae_Capsella_orientalis
Brassicaceae_Capsella_bursa-pastoris
Brassicaceae_Capsella_rubella
Brassicaceae_Capsella_grandiflora
Brassicaceae_Yinshania_exiensis
Brassicaceae_Yinshania_henryi
Brassicaceae_Tropidocarpum_gracile
Brassicaceae_Hornungia_procumbens

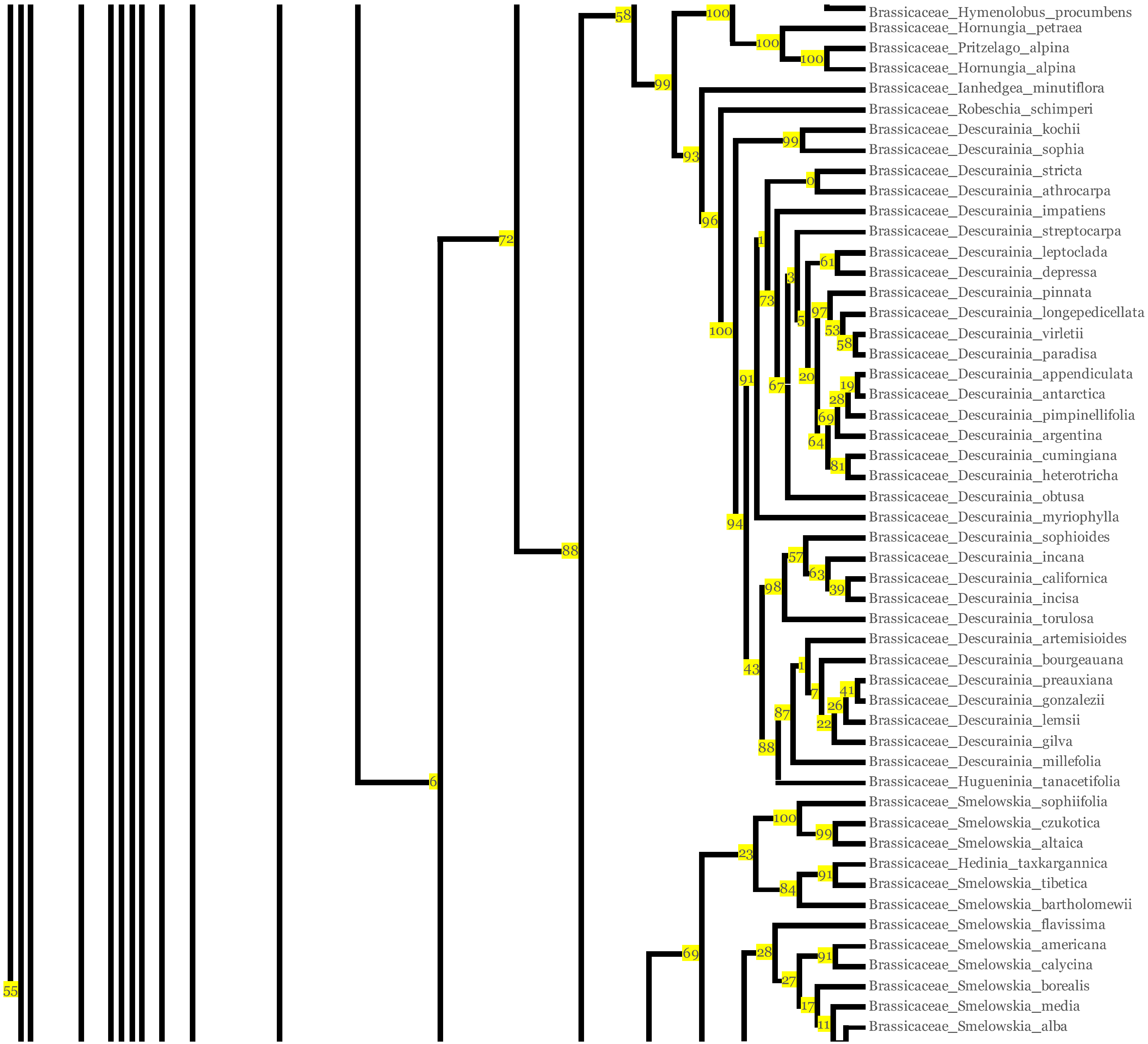
Brassicaceae_Hymenolobus_procumbens
Brassicaceae_Hornungia_petraea
Brassicaceae_Pritzelago_alpina
Brassicaceae_Hornungia_alpina
Brassicaceae_Ianhedgea_minutiflora
Brassicaceae_Robeschia_schimperi
Brassicaceae_Descurainia_kochii
Brassicaceae_Descurainia_sophia
Brassicaceae_Descurainia_stricta
Brassicaceae_Descurainia_athrocarpa
Brassicaceae_Descurainia_impatiens
Brassicaceae_Descurainia_streptocarpa
Brassicaceae_Descurainia_leptoclada
Brassicaceae_Descurainia_depressa
Brassicaceae_Descurainia_pinnata
Brassicaceae_Descurainia_longepedicellata
Brassicaceae_Descurainia_virletii
Brassicaceae_Descurainia_paradisa
Brassicaceae_Descurainia_appendiculata
Brassicaceae_Descurainia_antarctica
Brassicaceae_Descurainia_pimpinellifolia
Brassicaceae_Descurainia_argentina
Brassicaceae_Descurainia_cumingiana
Brassicaceae_Descurainia_heterotricha
Brassicaceae_Descurainia_obtusa
Brassicaceae_Descurainia_myriophylla
Brassicaceae_Descurainia_sophioides
Brassicaceae_Descurainia_incana
Brassicaceae_Descurainia_californica
Brassicaceae_Descurainia_incisa
Brassicaceae_Descurainia_torulosa
Brassicaceae_Descurainia_artemisioides
Brassicaceae_Descurainia_bourgeauana
Brassicaceae_Descurainia_preauxiana
Brassicaceae_Descurainia_gonzalezii
Brassicaceae_Descurainia_lemsii
Brassicaceae_Descurainia_gilva
Brassicaceae_Descurainia_millefolia
Brassicaceae_Huguenina_tanacetifolia
Brassicaceae_Smelowskia_sophiifolia
Brassicaceae_Smelowskia_czukotica
Brassicaceae_Smelowskia_altaica
Brassicaceae_Hedinia_taxkargannica
Brassicaceae_Smelowskia_tibetica
Brassicaceae_Smelowskia_bartholomewii
Brassicaceae_Smelowskia_flavissima
Brassicaceae_Smelowskia_americana
Brassicaceae_Smelowskia_calycina
Brassicaceae_Smelowskia_borealis
Brassicaceae_Smelowskia_media
Brassicaceae_Smelowskia_alba

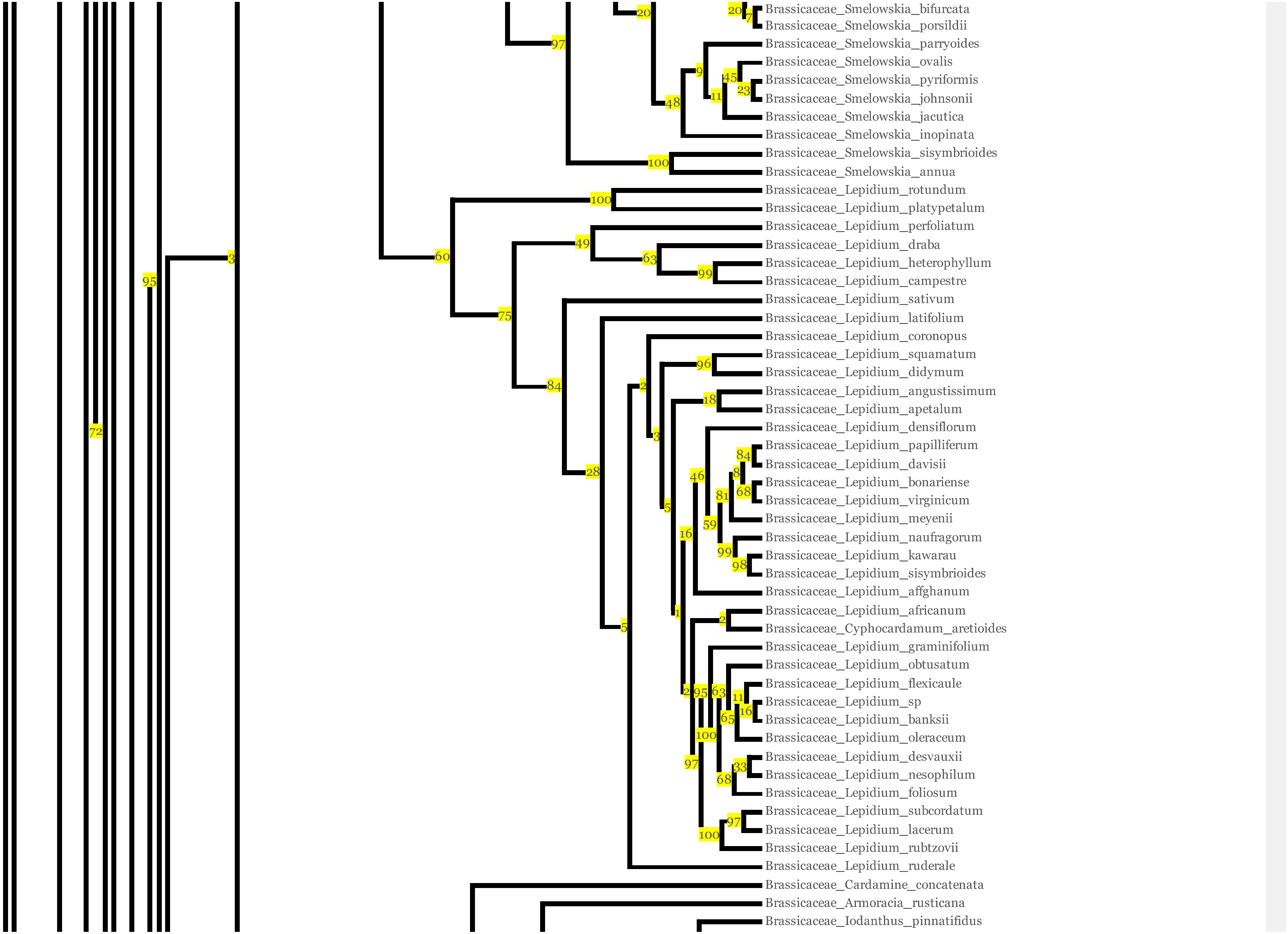
Brassicaceae_Smelowskia_bifurcata
Brassicaceae_Smelowskia_porsildii
Brassicaceae_Smelowskia_parryoides
Brassicaceae_Smelowskia_ovalis
Brassicaceae_Smelowskia_pyriformis
Brassicaceae_Smelowskia_johnsonii
Brassicaceae_Smelowskia_jacutica
Brassicaceae_Smelowskia_inopinata
Brassicaceae_Smelowskia_sisymbrioides
Brassicaceae_Smelowskia_annua
Brassicaceae_Lepidium_rotundum
Brassicaceae_Lepidium_platypetalum
Brassicaceae_Lepidium_perfoliatum
Brassicaceae_Lepidium_draba
Brassicaceae_Lepidium_heterophyllum
Brassicaceae_Lepidium_campestre
Brassicaceae_Lepidium_sativum
Brassicaceae_Lepidium_latifolium
Brassicaceae_Lepidium_coronopus
Brassicaceae_Lepidium_squamatum
Brassicaceae_Lepidium_didymum
Brassicaceae_Lepidium_angustissimum
Brassicaceae_Lepidium_apetalum
Brassicaceae_Lepidium_densiflorum
Brassicaceae_Lepidium_papilliferum
Brassicaceae_Lepidium_davisii
Brassicaceae_Lepidium_bonariense
Brassicaceae_Lepidium_virginicum
Brassicaceae_Lepidium_meyenii
Brassicaceae_Lepidium_naufragorum
Brassicaceae_Lepidium_kawarau
Brassicaceae_Lepidium_sisymbrioides
Brassicaceae_Lepidium_affghanum
Brassicaceae_Lepidium_africanum
Brassicaceae_Cyphocardamum_aretioides
Brassicaceae_Lepidium_graminifolium
Brassicaceae_Lepidium_obtusatum
Brassicaceae_Lepidium_flexicaule
Brassicaceae_Lepidium_sp
Brassicaceae_Lepidium_banksii
Brassicaceae_Lepidium_oleraceum
Brassicaceae_Lepidium_desvauxii
Brassicaceae_Lepidium_nesophilum
Brassicaceae_Lepidium_foliosum
Brassicaceae_Lepidium_subcordatum
Brassicaceae_Lepidium_lacerum
Brassicaceae_Lepidium_rubtzovii
Brassicaceae_Lepidium_ruderale
Brassicaceae_Cardamine_concatenata
Brassicaceae_Armoracia_rusticana
Brassicaceae_Iodanthus_pinnatifidus

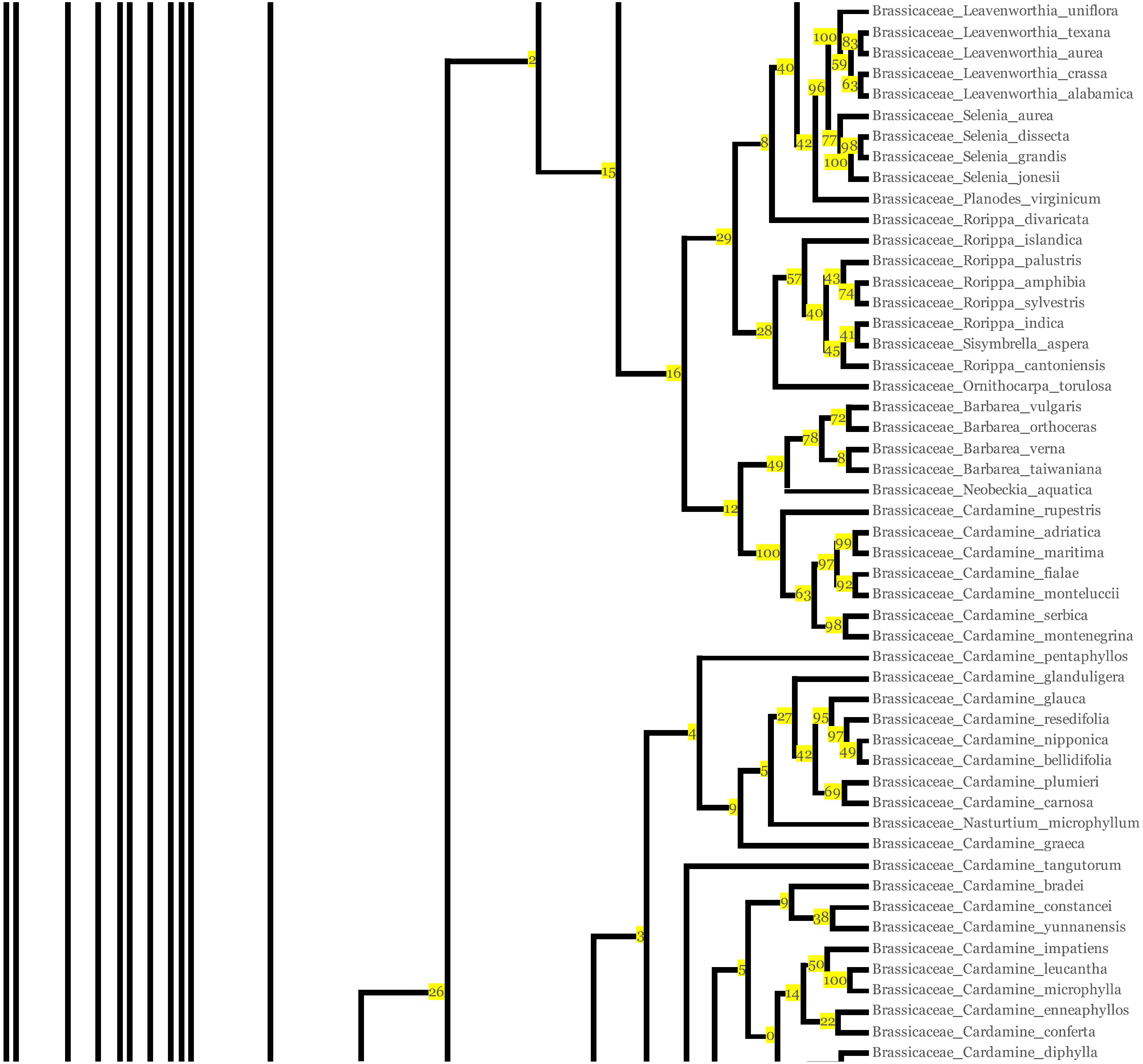
Brassicaceae_Leavenworthia_uniflora
Brassicaceae_Leavenworthia_texana
Brassicaceae_Leavenworthia_aurea
Brassicaceae_Leavenworthia_crassa
Brassicaceae_Leavenworthia_alabamica
Brassicaceae_Selenia_aurea
Brassicaceae_Selenia_dissecta
Brassicaceae_Selenia_grandis
Brassicaceae_Selenia_jonesii
Brassicaceae_Planodes_virginicum
Brassicaceae_Rorippa_divaricata
Brassicaceae_Rorippa_islandica
Brassicaceae_Rorippa_palustris
Brassicaceae_Rorippa_amphibia
Brassicaceae_Rorippa_sylvestris
Brassicaceae_Rorippa_indica
Brassicaceae_Sisymbrella_aspera
Brassicaceae_Rorippa_cantoniensis
Brassicaceae_Ornithocarpa_torulosa
Brassicaceae_Barbarea_vulgaris
Brassicaceae_Barbarea_orthoceras
Brassicaceae_Barbarea_verna
Brassicaceae_Barbarea_taiwaniana
Brassicaceae_Neobeckia_aquatica
Brassicaceae_Cardamine_rupestris
Brassicaceae_Cardamine_adriatica
Brassicaceae_Cardamine_maritima
Brassicaceae_Cardamine_fialae
Brassicaceae_Cardamine_monteluccii
Brassicaceae_Cardamine_serbica
Brassicaceae_Cardamine_montenegrina
Brassicaceae_Cardamine_pentaphyllos
Brassicaceae_Cardamine_glanduligera
Brassicaceae_Cardamine_glauca
Brassicaceae_Cardamine_resedifolia
Brassicaceae_Cardamine_nipponica
Brassicaceae_Cardamine_bellidifolia
Brassicaceae_Cardamine_plumieri
Brassicaceae_Cardamine_carnosa
Brassicaceae_Nasturtium_microphyllum
Brassicaceae_Cardamine_graeca
Brassicaceae_Cardamine_tangutorum
Brassicaceae_Cardamine_bradei
Brassicaceae_Cardamine_constancei
Brassicaceae_Cardamine_yunnanensis
Brassicaceae_Cardamine_impatiens
Brassicaceae_Cardamine_leucantha
Brassicaceae_Cardamine_microphylla
Brassicaceae_Cardamine_enneaphyllos
Brassicaceae_Cardamine_conferta
Brassicaceae_Cardamine_diphylla

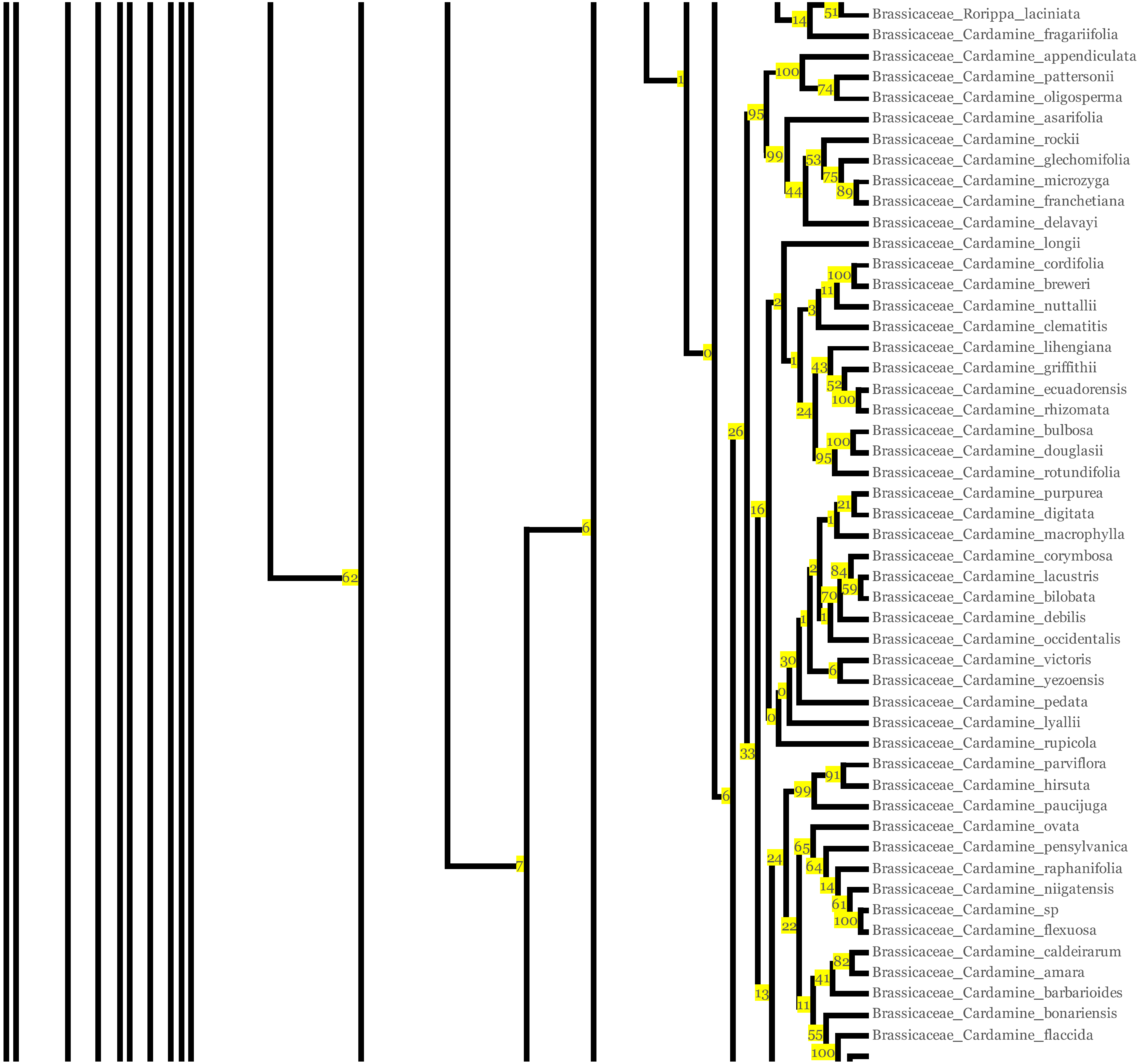
Brassicaceae_Rorippa_laciniata
Brassicaceae_Cardamine_fragariifolia
Brassicaceae_Cardamine_appendiculata
Brassicaceae_Cardamine_pattersonii
Brassicaceae_Cardamine_oligosperma
Brassicaceae_Cardamine_asarifolia
Brassicaceae_Cardamine_rockii
Brassicaceae_Cardamine_glechomifolia
Brassicaceae_Cardamine_microzyga
Brassicaceae_Cardamine_franchetiana
Brassicaceae_Cardamine_delavayi
Brassicaceae_Cardamine_longii
Brassicaceae_Cardamine_cordifolia
Brassicaceae_Cardamine_breweri
Brassicaceae_Cardamine_nuttallii
Brassicaceae_Cardamine_clematitis
Brassicaceae_Cardamine_lihengiana
Brassicaceae_Cardamine_griffithii
Brassicaceae_Cardamine_ecuadorensis
Brassicaceae_Cardamine_rhizomata
Brassicaceae_Cardamine_bulbosa
Brassicaceae_Cardamine_douglasii
Brassicaceae_Cardamine_rotundifolia
Brassicaceae_Cardamine_purpurea
Brassicaceae_Cardamine_digitata
Brassicaceae_Cardamine_macrophylla
Brassicaceae_Cardamine_corymbosa
Brassicaceae_Cardamine_lacustris
Brassicaceae_Cardamine_bilobata
Brassicaceae_Cardamine_debilis
Brassicaceae_Cardamine_occidentalis
Brassicaceae_Cardamine_victoris
Brassicaceae_Cardamine_yezoensis
Brassicaceae_Cardamine_pedata
Brassicaceae_Cardamine_lyallii
Brassicaceae_Cardamine_rupicola
Brassicaceae_Cardamine_parviflora
Brassicaceae_Cardamine_hirsuta
Brassicaceae_Cardamine_paucijuga
Brassicaceae_Cardamine_ovata
Brassicaceae_Cardamine_pensylvanica
Brassicaceae_Cardamine_raphanifolia
Brassicaceae_Cardamine_niigatensis
Brassicaceae_Cardamine_sp
Brassicaceae_Cardamine_flexuosa
Brassicaceae_Cardamine_caldeirarum
Brassicaceae_Cardamine_amara
Brassicaceae_Cardamine_barbarioides
Brassicaceae_Cardamine_bonariensis
Brassicaceae_Cardamine_flaccida

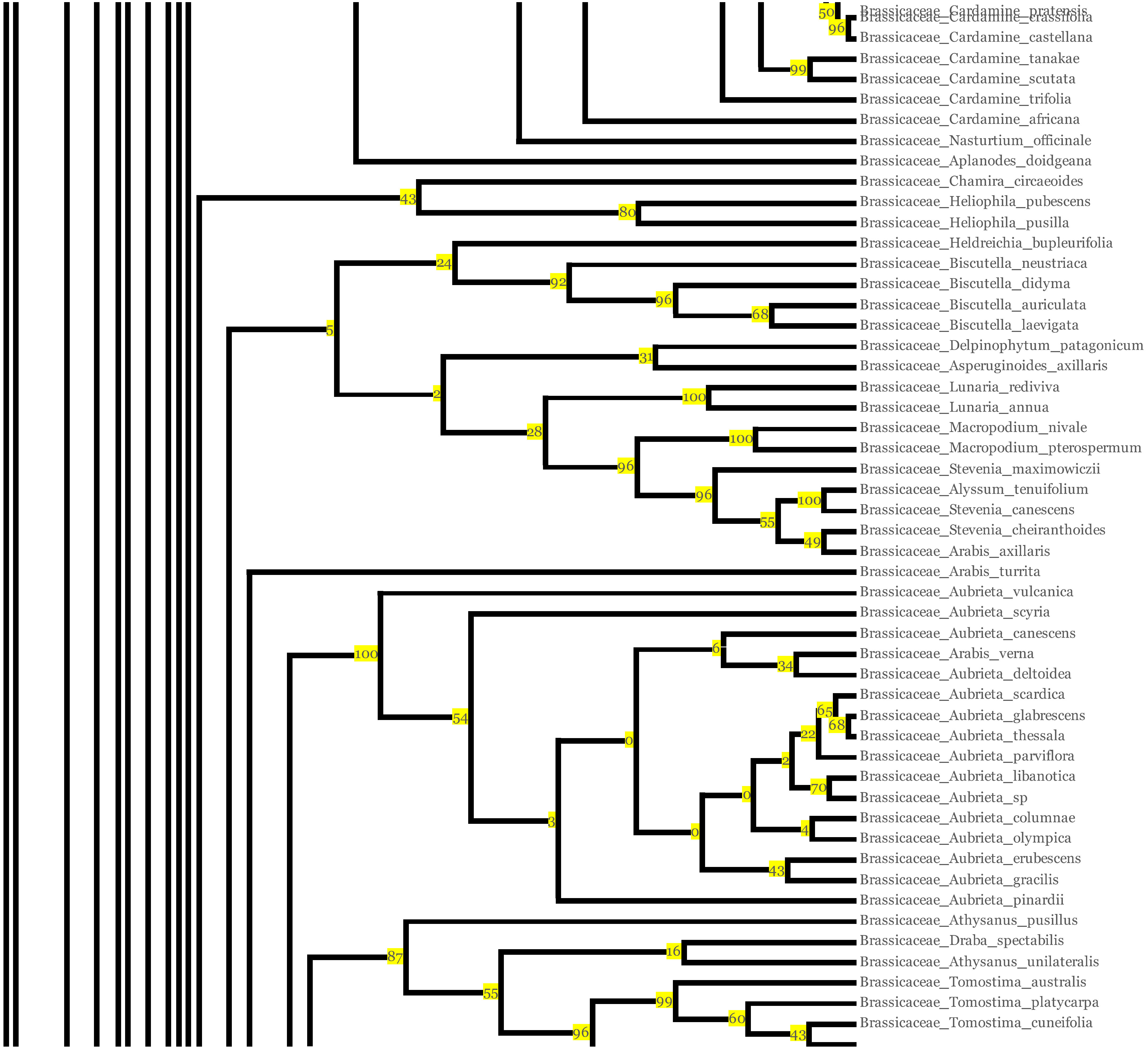
Brassicaceae_Cardamine_pratensis
Brassicaceae_Cardamine_crassifolia
Brassicaceae_Cardamine_castellana
Brassicaceae_Cardamine_tanakae
Brassicaceae_Cardamine_scutata
Brassicaceae_Cardamine_trifolia
Brassicaceae_Cardamine_africana
Brassicaceae_Nasturtium_officinale
Brassicaceae_Aplanodes_doidgeana
Brassicaceae_Chamira_circaeoides
Brassicaceae_Heliophila_pubescens
Brassicaceae_Heliophila_pusilla
Brassicaceae_Heldreichia_bupleurifolia
Brassicaceae_Biscutella_neustriaca
Brassicaceae_Biscutella_didyma
Brassicaceae_Biscutella_auriculata
Brassicaceae_Biscutella_laevigata
Brassicaceae_Delpinophytum_patagonicum
Brassicaceae_Asperuginoides_axillaris
Brassicaceae_Lunaria_rediviva
Brassicaceae_Lunaria_annua
Brassicaceae_Macropodium_nivale
Brassicaceae_Macropodium_pterospermum
Brassicaceae_Stevenia_maximowiczii
Brassicaceae_Alyssum_tenuifolium
Brassicaceae_Stevenia_canescens
Brassicaceae_Stevenia_cheiranthoides
Brassicaceae_Arabis_axillaris
Brassicaceae_Arabis_turrita
Brassicaceae_Aubrieta_vulcanica
Brassicaceae_Aubrieta_scyria
Brassicaceae_Aubrieta_canescens
Brassicaceae_Arabis_verna
Brassicaceae_Aubrieta_deltoidea
Brassicaceae_Aubrieta_scardica
Brassicaceae_Aubrieta_glabrescens
Brassicaceae_Aubrieta_thessala
Brassicaceae_Aubrieta_parviflora
Brassicaceae_Aubrieta_libanotica
Brassicaceae_Aubrieta_sp
Brassicaceae_Aubrieta_columnae
Brassicaceae_Aubrieta_olympica
Brassicaceae_Aubrieta_erubescens
Brassicaceae_Aubrieta_gracilis
Brassicaceae_Aubrieta_pinardii
Brassicaceae_Athysanus_pusillus
Brassicaceae_Draba_spectabilis
Brassicaceae_Athysanus_unilateralis
Brassicaceae_Tomostima_australis
Brassicaceae_Tomostima_platycarpa
Brassicaceae_Tomostima_cuneifolia

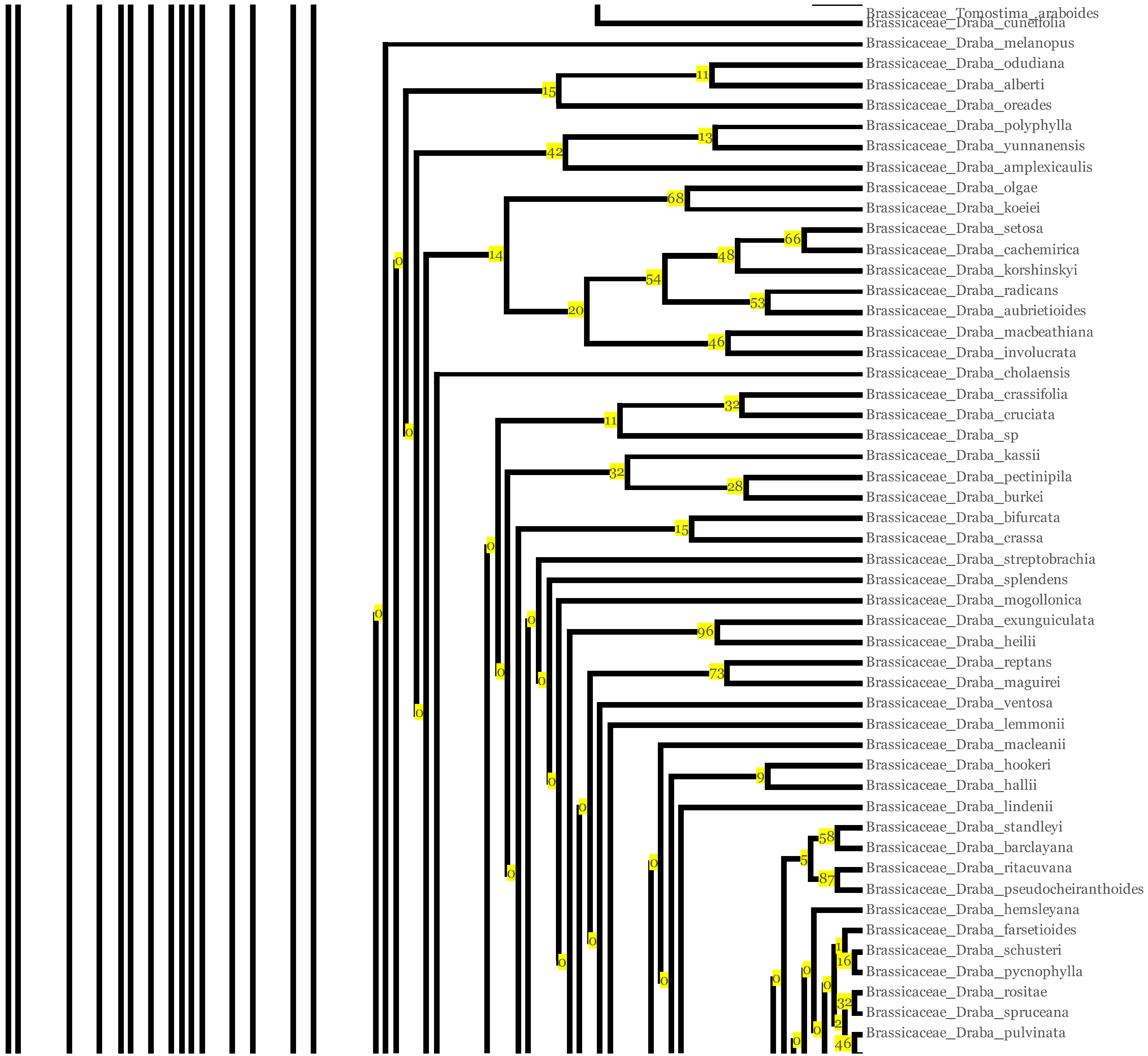
Brassicaceae_Tomostima_araboides
Brassicaceae_Draba_cuneifolia
Brassicaceae_Draba_melanopus
Brassicaceae_Draba_odudiana
Brassicaceae_Draba_alberti
Brassicaceae_Draba_oreades
Brassicaceae_Draba_polyphylla
Brassicaceae_Draba_yunnanensis
Brassicaceae_Draba_amplexicaulis
Brassicaceae_Draba_olgae
Brassicaceae_Draba_koeiei
Brassicaceae_Draba_setosa
Brassicaceae_Draba_cachemirica
Brassicaceae_Draba_korshinskyi
Brassicaceae_Draba_radicans
Brassicaceae_Draba_aubrietioides
Brassicaceae_Draba_macbeathiana
Brassicaceae_Draba_involucrata
Brassicaceae_Draba_cholaensis
Brassicaceae_Draba_crassifolia
Brassicaceae_Draba_cruciata
Brassicaceae_Draba_sp
Brassicaceae_Draba_kassii
Brassicaceae_Draba_pectinipila
Brassicaceae_Draba_burkei
Brassicaceae_Draba_bifurcata
Brassicaceae_Draba_crassa
Brassicaceae_Draba_streptobrachia
Brassicaceae_Draba_splendens
Brassicaceae_Draba_mogollonica
Brassicaceae_Draba_exunguiculata
Brassicaceae_Draba_heilii
Brassicaceae_Draba_reptans
Brassicaceae_Draba_maguirei
Brassicaceae_Draba_ventosa
Brassicaceae_Draba_lemmonii
Brassicaceae_Draba_macleanii
Brassicaceae_Draba_hookeri
Brassicaceae_Draba_hallii
Brassicaceae_Draba_lindenii
Brassicaceae_Draba_standleyi
Brassicaceae_Draba_barclayana
Brassicaceae_Draba_ritacuvana
Brassicaceae_Draba_pseudocheiranthoides
Brassicaceae_Draba_hemsleyana
Brassicaceae_Draba_farsetioides
Brassicaceae_Draba_schusteri
Brassicaceae_Draba_pycnophylla
Brassicaceae_Draba_rositae
Brassicaceae_Draba_spruceana
Brassicaceae_Draba_pulvinata
11
15
13
42
68
66
14
48
54
53
20
46
32
11
32
28
15
96
73
9
58
5
87
16
32
46

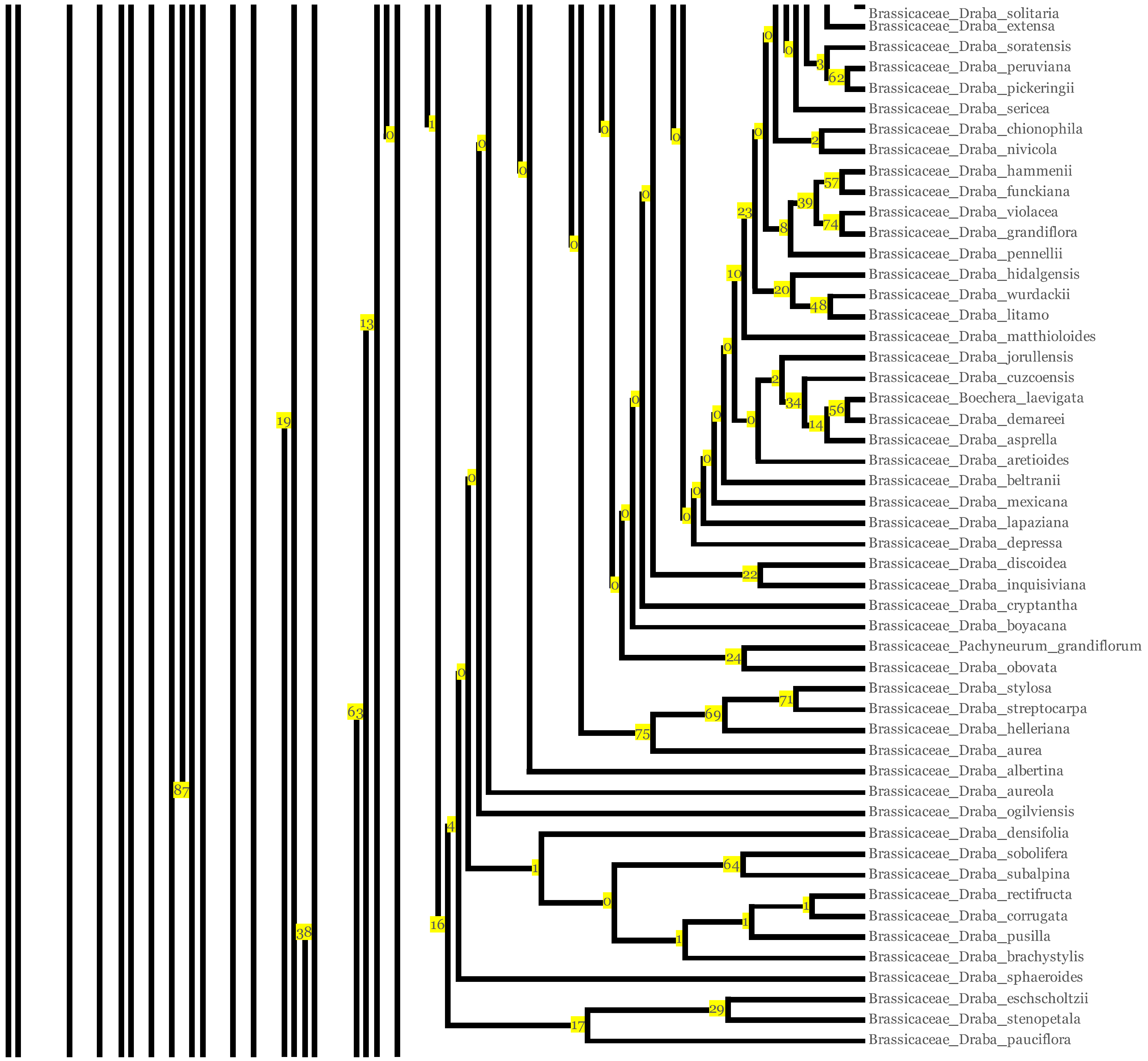

Brassicaceae_Draba_solitaria
Brassicaceae_Draba_extensa
Brassicaceae_Draba_soratensis
Brassicaceae_Draba_peruviana
Brassicaceae_Draba_pickeringii
Brassicaceae_Draba_sericea
Brassicaceae_Draba_chionophila
Brassicaceae_Draba_nivicola
Brassicaceae_Draba_hammenii
Brassicaceae_Draba_funckiana
Brassicaceae_Draba_violacea
Brassicaceae_Draba_grandiflora
Brassicaceae_Draba_pennellii
Brassicaceae_Draba_hidalgensis
Brassicaceae_Draba_wurdackii
Brassicaceae_Draba_litamo
Brassicaceae_Draba_matthioloides
Brassicaceae_Draba_jorullensis
Brassicaceae_Draba_cuzcoensis
Brassicaceae_Boechera_laevigata
Brassicaceae_Draba_demareei
Brassicaceae_Draba_asprella
Brassicaceae_Draba_aretioides
Brassicaceae_Draba_beltranii
Brassicaceae_Draba_mexicana
Brassicaceae_Draba_lapaziana
Brassicaceae_Draba_depressa
Brassicaceae_Draba_discoidea
Brassicaceae_Draba_inquisiviana
Brassicaceae_Draba_cryptantha
Brassicaceae_Draba_boyacana
Brassicaceae_Pachyneurum_grandiflorum
Brassicaceae_Draba_obovata
Brassicaceae_Draba_stylosa
Brassicaceae_Draba_streptocarpa
Brassicaceae_Draba_helleriana
Brassicaceae_Draba_aurea
Brassicaceae_Draba_albertina
Brassicaceae_Draba_aureola
Brassicaceae_Draba_ogilviensis
Brassicaceae_Draba_densifolia
Brassicaceae_Draba_sobolifera
Brassicaceae_Draba_subalpina
Brassicaceae_Draba_rectifructa
Brassicaceae_Draba_corrugata
Brassicaceae_Draba_pusilla
Brassicaceae_Draba_brachystylis
Brassicaceae_Draba_sphaeroides
Brassicaceae_Draba_eschscholtzii
Brassicaceae_Draba_stenopetala
Brassicaceae_Draba_pauciflora

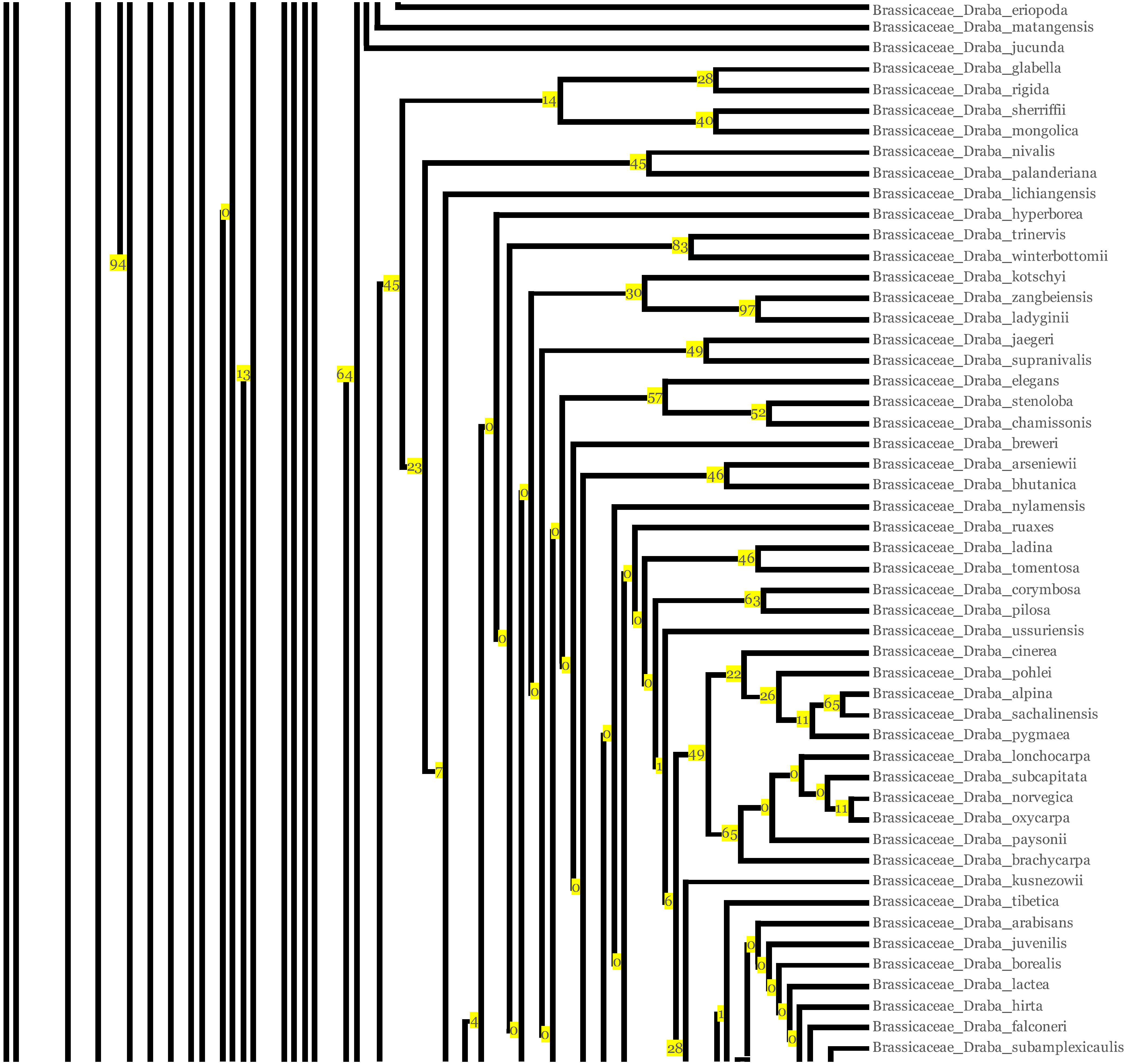

Brassicaceae_Draba_eriopoda
Brassicaceae_Draba_matangensis
Brassicaceae_Draba_jucunda
Brassicaceae_Draba_glabella
Brassicaceae_Draba_rigida
Brassicaceae_Draba_sherriffii
Brassicaceae_Draba_mongolica
Brassicaceae_Draba_nivalis
Brassicaceae_Draba_palanderiana
Brassicaceae_Draba_lichiangensis
Brassicaceae_Draba_hyperborea
Brassicaceae_Draba_trinervis
Brassicaceae_Draba_winterbottomii
Brassicaceae_Draba_kotschyi
Brassicaceae_Draba_zangbeiensis
Brassicaceae_Draba_ladyginii
Brassicaceae_Draba_jaegeri
Brassicaceae_Draba_supranivalis
Brassicaceae_Draba_elegans
Brassicaceae_Draba_stenoloba
Brassicaceae_Draba_chamissonis
Brassicaceae_Draba_breweri
Brassicaceae_Draba_arseniewii
Brassicaceae_Draba_bhutanica
Brassicaceae_Draba_nylamensis
Brassicaceae_Draba_ruaxes
Brassicaceae_Draba_ladina
Brassicaceae_Draba_tomentosa
Brassicaceae_Draba_corymbosa
Brassicaceae_Draba_pilosa
Brassicaceae_Draba_ussuriensis
Brassicaceae_Draba_cinerea
Brassicaceae_Draba_pohlei
Brassicaceae_Draba_alpina
Brassicaceae_Draba_sachalinensis
Brassicaceae_Draba_pygmaea
Brassicaceae_Draba_lonchocarpa
Brassicaceae_Draba_subcapitata
Brassicaceae_Draba_norvegica
Brassicaceae_Draba_oxycarpa
Brassicaceae_Draba_paysonii
Brassicaceae_Draba_brachycarpa
Brassicaceae_Draba_kusnezowii
Brassicaceae_Draba_tibetica
Brassicaceae_Draba_arabisans
Brassicaceae_Draba_juvenilis
Brassicaceae_Draba_borealis
Brassicaceae_Draba_lactea
Brassicaceae_Draba_hirta
Brassicaceae_Draba_falconeri
Brassicaceae_Draba_subamplexicaulis

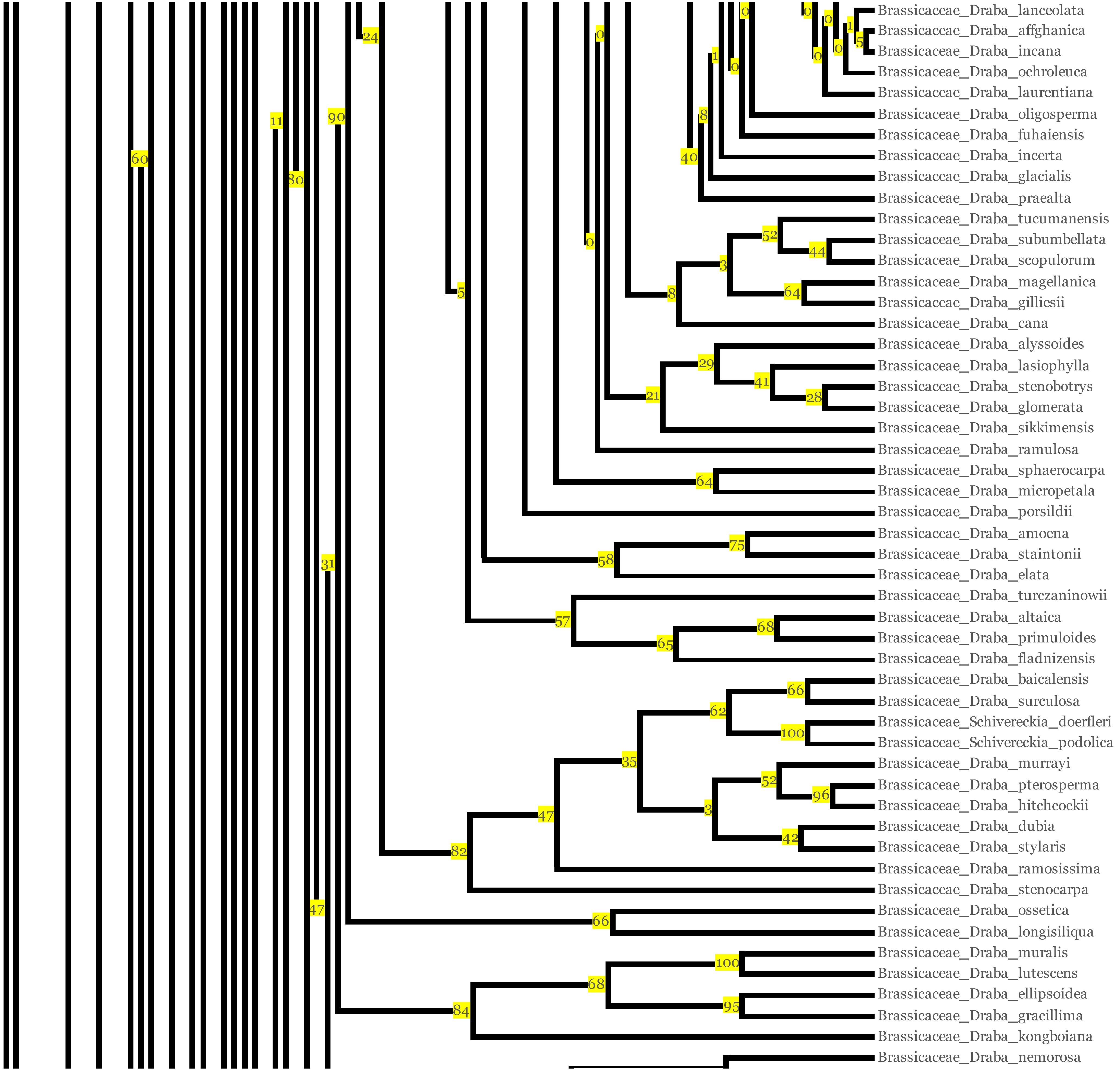
Brassicaceae_Draba_lanceolata
Brassicaceae_Draba_affghanica
Brassicaceae_Draba_incana
Brassicaceae_Draba_ochroleuca
Brassicaceae_Draba_laurentiana
Brassicaceae_Draba_oligosperma
Brassicaceae_Draba_fuhaiensis
Brassicaceae_Draba_incerta
Brassicaceae_Draba_glacialis
Brassicaceae_Draba_praealta
Brassicaceae_Draba_tucumanensis
Brassicaceae_Draba_subumbellata
Brassicaceae_Draba_scopulorum
Brassicaceae_Draba_magellanica
Brassicaceae_Draba_gilliesii
Brassicaceae_Draba_cana
Brassicaceae_Draba_alyssoides
Brassicaceae_Draba_lasiophylla
Brassicaceae_Draba_stenobotrys
Brassicaceae_Draba_glomerata
Brassicaceae_Draba_sikkimensis
Brassicaceae_Draba_ramulosa
Brassicaceae_Draba_sphaerocarpa
Brassicaceae_Draba_micropetala
Brassicaceae_Draba_porsildii
Brassicaceae_Draba_amoena
Brassicaceae_Draba_staintonii
Brassicaceae_Draba_elata
Brassicaceae_Draba_turczaninowii
Brassicaceae_Draba_altaica
Brassicaceae_Draba_primuloides
Brassicaceae_Draba_fladnizensis
Brassicaceae_Draba_baicalensis
Brassicaceae_Draba_surculosa
Brassicaceae_Schivereckia_doerfleri
Brassicaceae_Schivereckia_podolica
Brassicaceae_Draba_murrayi
Brassicaceae_Draba_pterosperma
Brassicaceae_Draba_hitchcockii
Brassicaceae_Draba_dubia
Brassicaceae_Draba_stylaris
Brassicaceae_Draba_ramosissima
Brassicaceae_Draba_stenocarpa
Brassicaceae_Draba_ossetica
Brassicaceae_Draba_longisiliqua
Brassicaceae_Draba_muralis
Brassicaceae_Draba_lutescens
Brassicaceae_Draba_ellipsoidea
Brassicaceae_Draba_gracillima
Brassicaceae_Draba_kongboiana
Brassicaceae_Draba_nemorosa

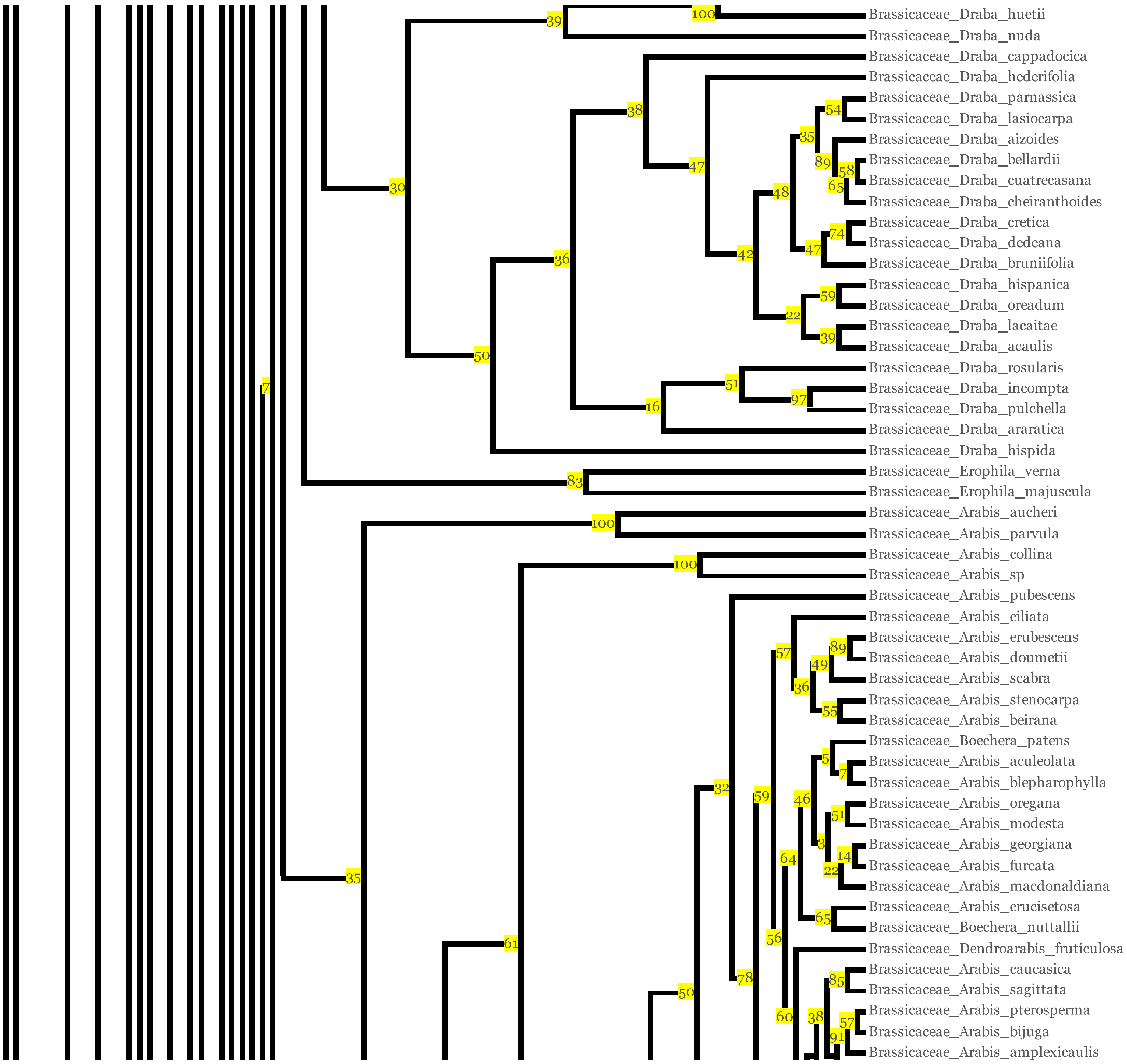
Brassicaceae_Draba_huetii
Brassicaceae_Draba_nuda
Brassicaceae_Draba_cappadocica
Brassicaceae_Draba_hederifolia
Brassicaceae_Draba_parnassica
Brassicaceae_Draba_lasiocarpa
Brassicaceae_Draba_aizoides
Brassicaceae_Draba_bellardii
Brassicaceae_Draba_cuatrecasana
Brassicaceae_Draba_cheiranthoides
Brassicaceae_Draba_cretica
Brassicaceae_Draba_dedeana
Brassicaceae_Draba_bruniifolia
Brassicaceae_Draba_hispanica
Brassicaceae_Draba_oreadum
Brassicaceae_Draba_lacaitae
Brassicaceae_Draba_acaulis
Brassicaceae_Draba_rosularis
Brassicaceae_Draba_incompta
Brassicaceae_Draba_pulchella
Brassicaceae_Draba_araratica
Brassicaceae_Draba_hispida
Brassicaceae_Erophila_verna
Brassicaceae_Erophila_majuscula
Brassicaceae_Arabis_aucheri
Brassicaceae_Arabis_parvula
Brassicaceae_Arabis_collina
Brassicaceae_Arabis_sp
Brassicaceae_Arabis_pubescens
Brassicaceae_Arabis_ciliata
Brassicaceae_Arabis_erubescens
Brassicaceae_Arabis_doumetii
Brassicaceae_Arabis_scabra
Brassicaceae_Arabis_stenocarpa
Brassicaceae_Arabis_beirana
Brassicaceae_Boechera_patens
Brassicaceae_Arabis_aculeolata
Brassicaceae_Arabis_blepharophylla
Brassicaceae_Arabis_oregana
Brassicaceae_Arabis_modesta
Brassicaceae_Arabis_georgiana
Brassicaceae_Arabis_furcata
Brassicaceae_Arabis_macdonaldiana
Brassicaceae_Arabis_crucisetosa
Brassicaceae_Boechera_nuttallii
Brassicaceae_Dendroarabis_fruticulosa
Brassicaceae_Arabis_caucasica
Brassicaceae_Arabis_sagittata
Brassicaceae_Arabis_pterosperma
Brassicaceae_Arabis_bijuga
Brassicaceae_Arabis_amplexicaulis

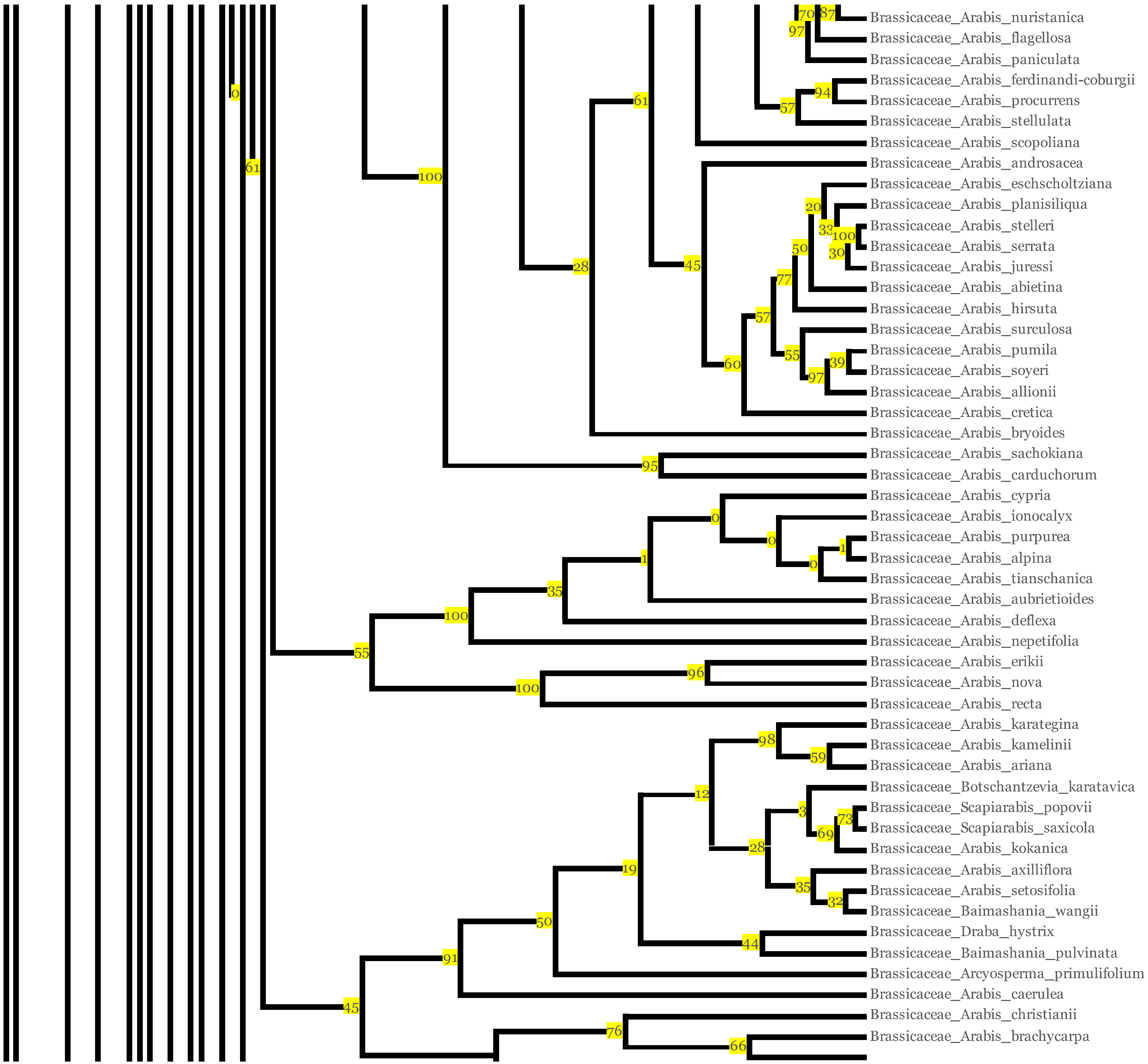
Brassicaceae_Arabis_nuristanica
Brassicaceae_Arabis_flagellosa
Brassicaceae_Arabis_paniculata
Brassicaceae_Arabis_ferdinandi-coburgii
Brassicaceae_Arabis_procurrens
Brassicaceae_Arabis_stellulata
Brassicaceae_Arabis_scopoliana
Brassicaceae_Arabis_androsacea
Brassicaceae_Arabis_eschscholtziana
Brassicaceae_Arabis_planisiliqua
Brassicaceae_Arabis_stelleri
Brassicaceae_Arabis_serrata
Brassicaceae_Arabis_juressi
Brassicaceae_Arabis_abietina
Brassicaceae_Arabis_hirsuta
Brassicaceae_Arabis_surculosa
Brassicaceae_Arabis_pumila
Brassicaceae_Arabis_soyeri
Brassicaceae_Arabis_allionii
Brassicaceae_Arabis_cretica
Brassicaceae_Arabis_bryoides
Brassicaceae_Arabis_sachokiana
Brassicaceae_Arabis_carduchorum
Brassicaceae_Arabis_cypria
Brassicaceae_Arabis_ionocalyx
Brassicaceae_Arabis_purpurea
Brassicaceae_Arabis_alpina
Brassicaceae_Arabis_tianschanica
Brassicaceae_Arabis_aubrietioides
Brassicaceae_Arabis_deflexa
Brassicaceae_Arabis_nepetifolia
Brassicaceae_Arabis_erikii
Brassicaceae_Arabis_nova
Brassicaceae_Arabis_recta
Brassicaceae_Arabis_karategina
Brassicaceae_Arabis_kamelinii
Brassicaceae_Arabis_ariana
Brassicaceae_Botschantzevia_karatavica
Brassicaceae_Scapiarabis_popovii
Brassicaceae_Scapiarabis_saxicola
Brassicaceae_Arabis_kokanica
Brassicaceae_Arabis_axilliflora
Brassicaceae_Arabis_setosifolia
Brassicaceae_Baimashania_wangii
Brassicaceae_Draba_hystrix
Brassicaceae_Baimashania_pulvinata
Brassicaceae_Arcyosperma_primulifolium
Brassicaceae_Arabis_caerulea
Brassicaceae_Arabis_christianii
Brassicaceae_Arabis_brachycarpa

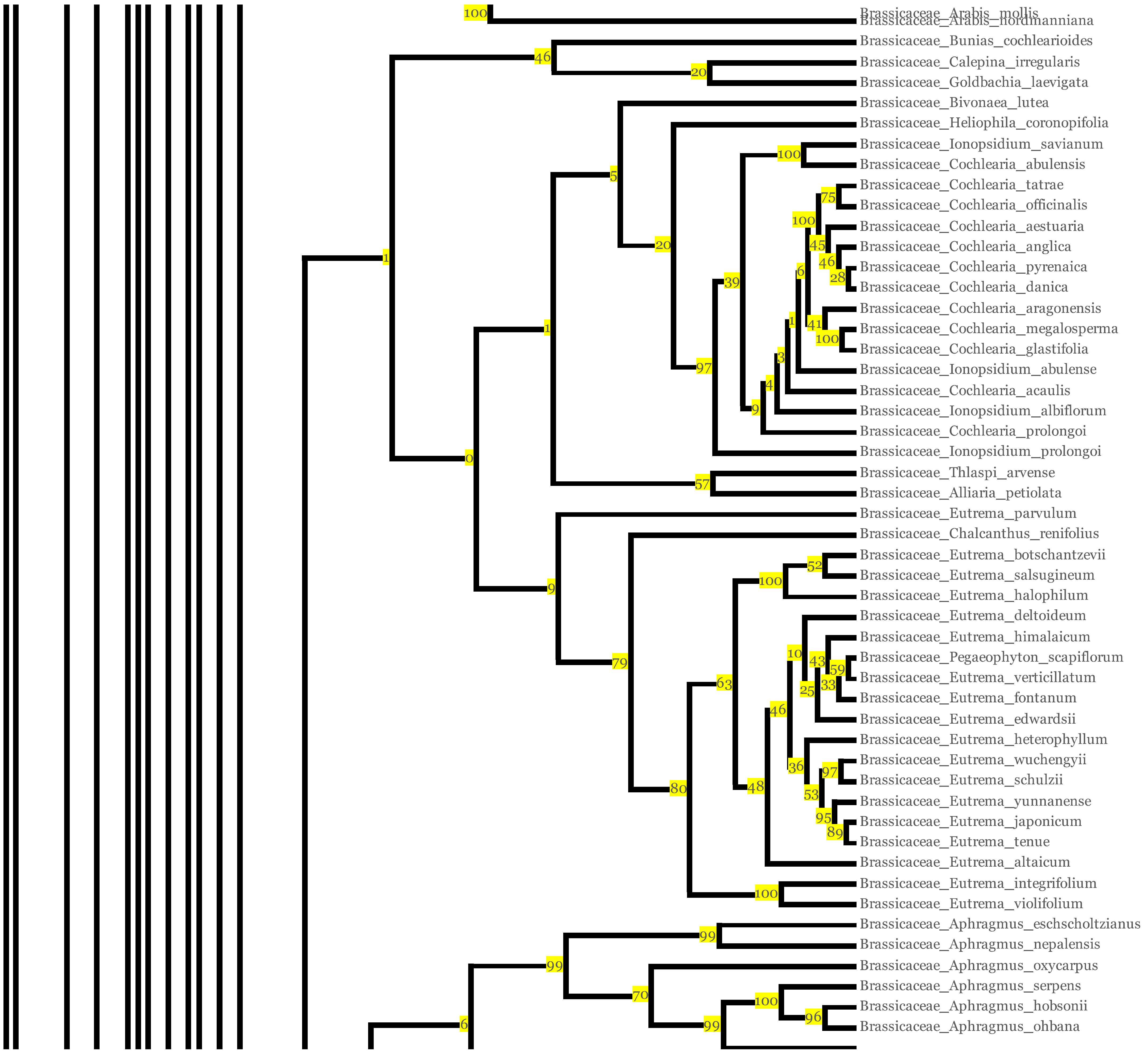
Brassicaceae_Arabis_mollis
Brassicaceae_Arabis_nordmanniana
Brassicaceae_Bunias_cochlearioides
Brassicaceae_Calepina_irregularis
Brassicaceae_Goldbachia_laevigata
Brassicaceae_Bivonaea_lutea
Brassicaceae_Heliophila_coronopifolia
Brassicaceae_Ionopsidium_savianum
Brassicaceae_Cochlearia_abulensis
Brassicaceae_Cochlearia_tatrae
Brassicaceae_Cochlearia_officinalis
Brassicaceae_Cochlearia_aestuaria
Brassicaceae_Cochlearia_anglica
Brassicaceae_Cochlearia_pyrenaica
Brassicaceae_Cochlearia_danica
Brassicaceae_Cochlearia_aragonensis
Brassicaceae_Cochlearia_megalosperma
Brassicaceae_Cochlearia_glastifolia
Brassicaceae_Ionopsidium_abulense
Brassicaceae_Cochlearia_acaulis
Brassicaceae_Ionopsidium_albiflorum
Brassicaceae_Cochlearia_prolongoi
Brassicaceae_Ionopsidium_prolongoi
Brassicaceae_Thlaspi_arvense
Brassicaceae_Alliaria_petiolata
Brassicaceae_Eutrema_parvulum
Brassicaceae_Chalcanthus_renifolius
Brassicaceae_Eutrema_botschantzevii
Brassicaceae_Eutrema_salsugineum
Brassicaceae_Eutrema_halophilum
Brassicaceae_Eutrema_deltoideum
Brassicaceae_Eutrema_himalaicum
Brassicaceae_Pegaeophyton_scapiflorum
Brassicaceae_Eutrema_verticillatum
Brassicaceae_Eutrema_fontanum
Brassicaceae_Eutrema_edwardsii
Brassicaceae_Eutrema_heterophyllum
Brassicaceae_Eutrema_wuchengyii
Brassicaceae_Eutrema_schulzii
Brassicaceae_Eutrema_yunnanense
Brassicaceae_Eutrema_japonicum
Brassicaceae_Eutrema_tenue
Brassicaceae_Eutrema_altaicum
Brassicaceae_Eutrema_integrifolium
Brassicaceae_Eutrema_violifolium
Brassicaceae_Aphragmus_eschscholtzianus
Brassicaceae_Aphragmus_nepalensis
Brassicaceae_Aphragmus_oxycarpus
Brassicaceae_Aphragmus_serpens
Brassicaceae_Aphragmus_hobsonii
Brassicaceae_Aphragmus_ohbana

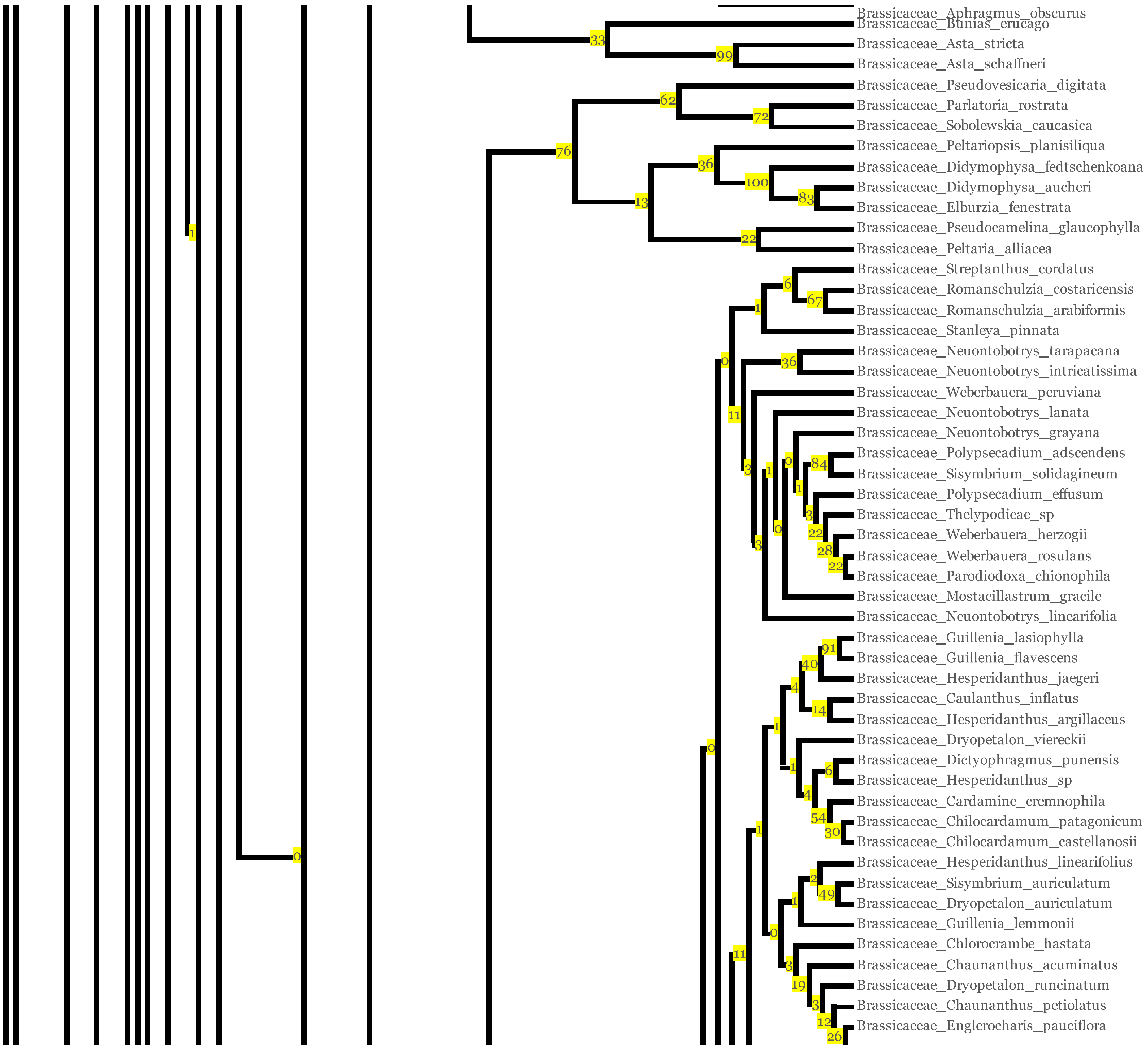
Brassicaceae_Aphragmus_obscurus
Brassicaceae_Bunias_erucago
Brassicaceae_Asta_stricta
Brassicaceae_Asta_schaffneri
Brassicaceae_Pseudovesicaria_digitata
Brassicaceae_Parlatoria_rostrata
Brassicaceae_Sobolewskia_caucasica
Brassicaceae_Peltariopsis_planisiliqua
Brassicaceae_Didymophysa_fedtschenkoana
Brassicaceae_Didymophysa_aucheri
Brassicaceae_Elburzia_fenestrata
Brassicaceae_Pseudocamelina_glaucophylla
Brassicaceae_Peltaria_alliacea
Brassicaceae_Streptanthus_cordatus
Brassicaceae_Romanschulzia_costaricensis
Brassicaceae_Romanschulzia_arabiformis
Brassicaceae_Stanleya_pinnata
Brassicaceae_Neuontobotrys_tarapacana
Brassicaceae_Neuontobotrys_intricatissima
Brassicaceae_Weberbauera_peruviana
Brassicaceae_Neuontobotrys_lanata
Brassicaceae_Neuontobotrys_grayana
Brassicaceae_Polypsecadium_adscendens
Brassicaceae_Sisymbrium_solidagineum
Brassicaceae_Polypsecadium_effusum
Brassicaceae_Thelypodieae_sp
Brassicaceae_Weberbauera_herzogii
Brassicaceae_Weberbauera_rosulans
Brassicaceae_Parodiodoxa_chionophila
Brassicaceae_Mostacillastrum_gracile
Brassicaceae_Neuontobotrys_linearifolia
Brassicaceae_Guillenia_lasiophylla
Brassicaceae_Guillenia_flavescens
Brassicaceae_Hesperidanthus_jaegeri
Brassicaceae_Caulanthus_inflatus
Brassicaceae_Hesperidanthus_argillaceus
Brassicaceae_Dryopetalon_viereckii
Brassicaceae_Dictyophragmus_punensis
Brassicaceae_Hesperidanthus_sp
Brassicaceae_Cardamine_cremnophila
Brassicaceae_Chilocardamum_patagonicum
Brassicaceae_Chilocardamum_castellanosii
Brassicaceae_Hesperidanthus_linearifolius
Brassicaceae_Sisymbrium_auriculatum
Brassicaceae_Dryopetalon_auriculatum
Brassicaceae_Guillenia_lemmonii
Brassicaceae_Chlorocrambe_hastata
Brassicaceae_Chaunanthus_acuminatus
Brassicaceae_Dryopetalon_runcinatum
Brassicaceae_Chaunanthus_petiolatus
Brassicaceae_Englerocharis_pauciflora

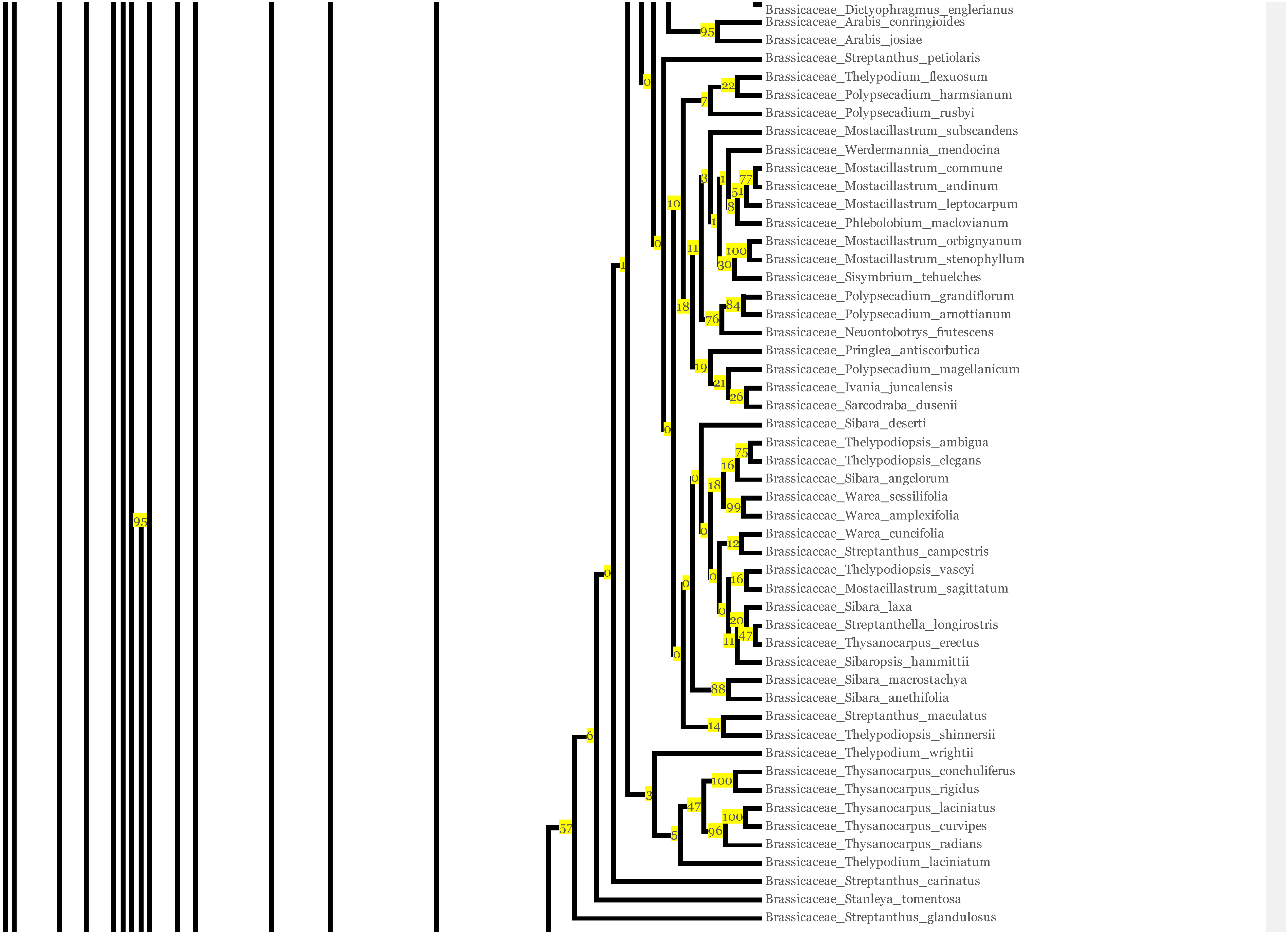
Brassicaceae_Dictyophragmus_englerianus
Brassicaceae_Arabis_conringioides
Brassicaceae_Arabis_josiae
Brassicaceae_Streptanthus_petiolaris
Brassicaceae_Thelypodium_flexuosum
Brassicaceae_Polypsecadium_harmsianum
Brassicaceae_Polypsecadium_rusbyi
Brassicaceae_Mostacillastrum_subscandens
Brassicaceae_Werdermannia_mendocina
Brassicaceae_Mostacillastrum_commune
Brassicaceae_Mostacillastrum_andinum
Brassicaceae_Mostacillastrum_leptocarpum
Brassicaceae_Phlebolobium_maclovianum
Brassicaceae_Mostacillastrum_orbignyanum
Brassicaceae_Mostacillastrum_stenophyllum
Brassicaceae_Sisymbrium_tehuelches
Brassicaceae_Polypsecadium_grandiflorum
Brassicaceae_Polypsecadium_arnottianum
Brassicaceae_Neuontobotrys_frutescens
Brassicaceae_Pringlea_antiscorbutica
Brassicaceae_Polypsecadium_magellanicum
Brassicaceae_Ivania_juncalensis
Brassicaceae_Sarcodraba_dusenii
Brassicaceae_Sibara_deserti
Brassicaceae_Thelypodiopsis_ambigua
Brassicaceae_Thelypodiopsis_elegans
Brassicaceae_Sibara_angelorum
Brassicaceae_Warea_sessilifolia
Brassicaceae_Warea_amplexifolia
Brassicaceae_Warea_cuneifolia
Brassicaceae_Streptanthus_campestris
Brassicaceae_Thelypodiopsis_vaseyi
Brassicaceae_Mostacillastrum_sagittatum
Brassicaceae_Sibara_laxa
Brassicaceae_Streptanthella_longirostris
Brassicaceae_Thysanocarpus_erectus
Brassicaceae_Sibaropsis_hammittii
Brassicaceae_Sibara_macrostachya
Brassicaceae_Sibara_anethifolia
Brassicaceae_Streptanthus_maculatus
Brassicaceae_Thelypodiopsis_shinnersii
Brassicaceae_Thelypodium_wrightii
Brassicaceae_Thysanocarpus_conchuliferus
Brassicaceae_Thysanocarpus_rigidus
Brassicaceae_Thysanocarpus_laciniatus
Brassicaceae_Thysanocarpus_curvipes
Brassicaceae_Thysanocarpus_radians
Brassicaceae_Thelypodium_laciniatum
Brassicaceae_Streptanthus_carinatus
Brassicaceae_Stanleya_tomentosa
Brassicaceae_Streptanthus_glandulosus

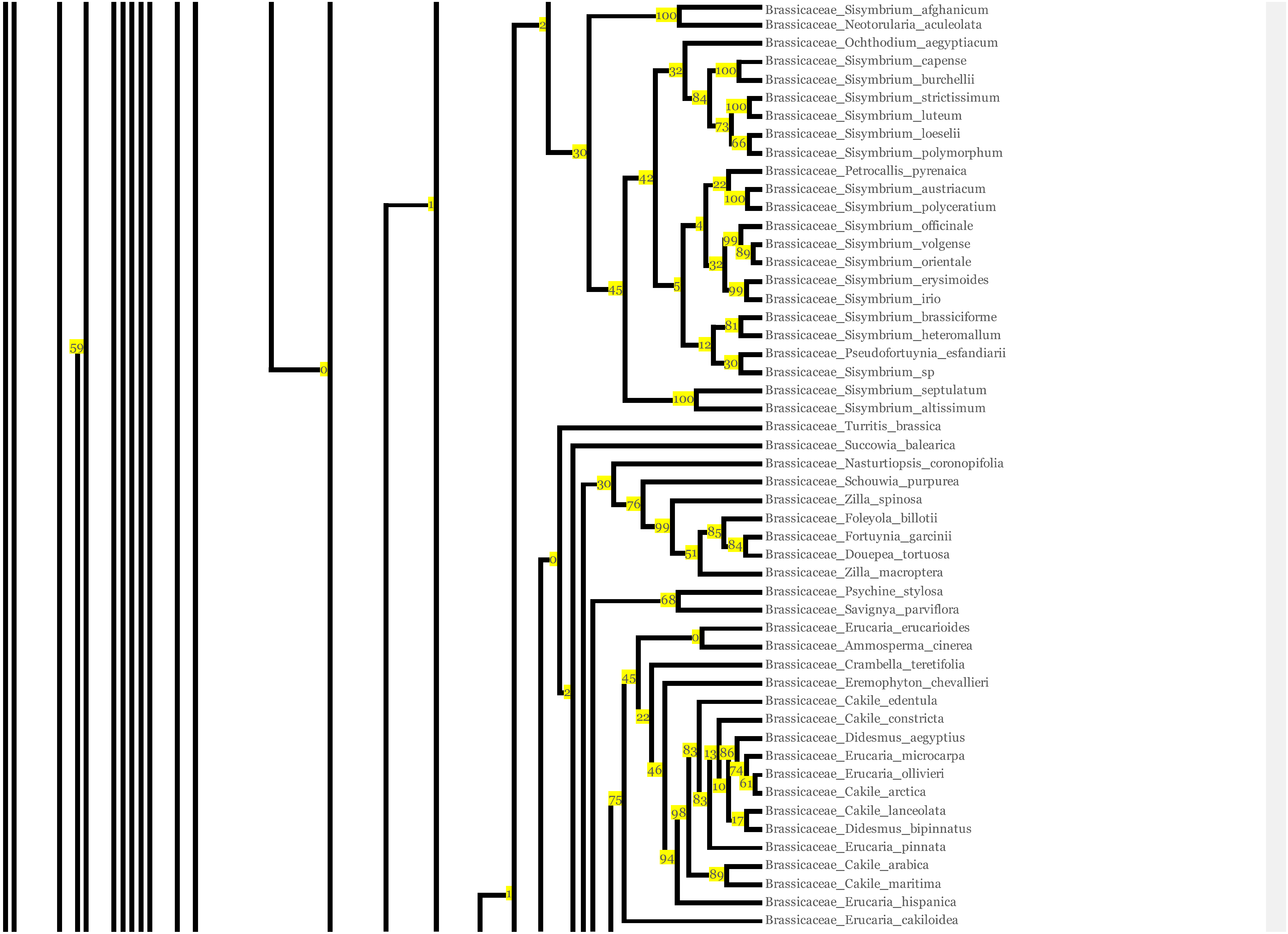
Brassicaceae_Sisymbrium_afghanicum
Brassicaceae_Neotorularia_aculeolata
Brassicaceae_Ochthodium_aegyptiacum
Brassicaceae_Sisymbrium_capense
Brassicaceae_Sisymbrium_burchellii
Brassicaceae_Sisymbrium_strictissimum
Brassicaceae_Sisymbrium_luteum
Brassicaceae_Sisymbrium_loeselii
Brassicaceae_Sisymbrium_polymorphum
Brassicaceae_Petrocallis_pyrenaica
Brassicaceae_Sisymbrium_austriacum
Brassicaceae_Sisymbrium_polyceratium
Brassicaceae_Sisymbrium_officinale
Brassicaceae_Sisymbrium_volgense
Brassicaceae_Sisymbrium_orientale
Brassicaceae_Sisymbrium_erysimoides
Brassicaceae_Sisymbrium_irio
Brassicaceae_Sisymbrium_brassiciforme
Brassicaceae_Sisymbrium_heteromallum
Brassicaceae_Pseudofortuynia_esfandiarii
Brassicaceae_Sisymbrium_sp
Brassicaceae_Sisymbrium_septulatum
Brassicaceae_Sisymbrium_altissimum
Brassicaceae_Turritis_brassica
Brassicaceae_Succowia_balearica
Brassicaceae_Nasturtiopsis_coronopifolia
Brassicaceae_Schouwia_purpurea
Brassicaceae_Zilla_spinosa
Brassicaceae_Foleyola_billotii
Brassicaceae_Fortuynia_garcinii
Brassicaceae_Douepea_tortuosa
Brassicaceae_Zilla_macroptera
Brassicaceae_Psychine_stylosa
Brassicaceae_Savignya_parviflora
Brassicaceae_Erucaria_erucarioides
Brassicaceae_Ammosperma_cinerea
Brassicaceae_Crambella_teretifolia
Brassicaceae_Eremophyton_chevallieri
Brassicaceae_Cakile_edentula
Brassicaceae_Cakile_constricta
Brassicaceae_Didesmus_aegyptius
Brassicaceae_Erucaria_microcarpa
Brassicaceae_Erucaria_ollivieri
Brassicaceae_Cakile_arctica
Brassicaceae_Cakile_lanceolata
Brassicaceae_Didesmus_bipinnatus
Brassicaceae_Erucaria_pinnata
Brassicaceae_Cakile_arabica
Brassicaceae_Cakile_maritima
Brassicaceae_Erucaria_hispanica
Brassicaceae_Erucaria_cakiloidea
100
2
30
32
100
84
100
73
66
42
22
100
4
99
89
32
99
45
5
81
12
30
100
59
0
1
30
76
99
85
84
51
0
68
0
2
45
22
46
83
13
86
74
61
10
83
17
98
94
89
75
1

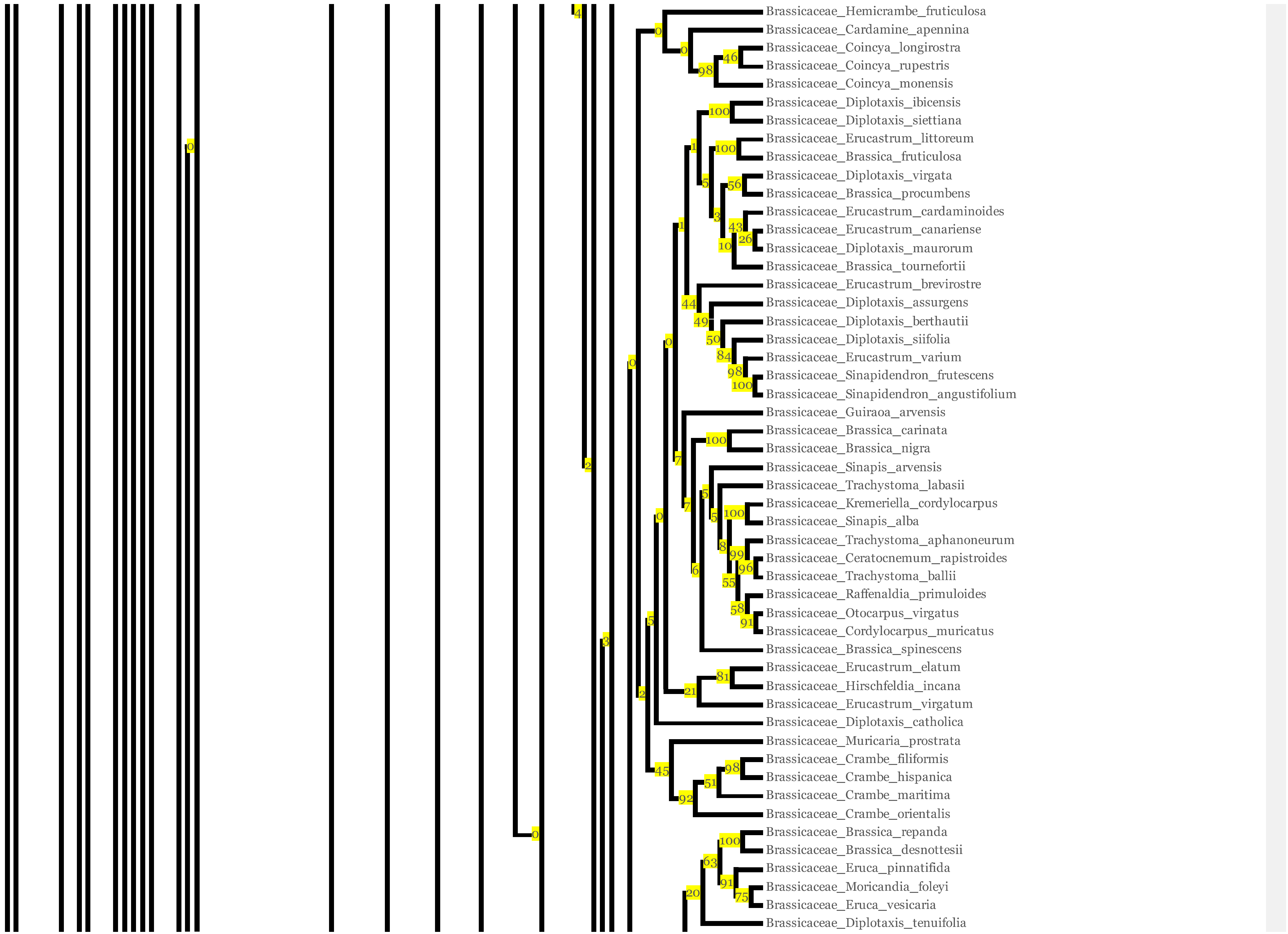
Brassicaceae_Hemicrambe_fruticulosa
Brassicaceae_Cardamine_apennina
Brassicaceae_Coincya_longirostra
Brassicaceae_Coincya_rupestris
Brassicaceae_Coincya_monensis
Brassicaceae_Diplotaxis_ibicensis
Brassicaceae_Diplotaxis_siettiana
Brassicaceae_Erucastrum_littoreum
Brassicaceae_Brassica_fruticulosa
Brassicaceae_Diplotaxis_virgata
Brassicaceae_Brassica_procumbens
Brassicaceae_Erucastrum_cardaminoides
Brassicaceae_Erucastrum_canariense
Brassicaceae_Diplotaxis_maurorum
Brassicaceae_Brassica_tournefortii
Brassicaceae_Erucastrum_brevirostre
Brassicaceae_Diplotaxis_assurgens
Brassicaceae_Diplotaxis_berthautii
Brassicaceae_Diplotaxis_siifolia
Brassicaceae_Erucastrum_varium
Brassicaceae_Sinapidendron_frutescens
Brassicaceae_Sinapidendron_angustifolium
Brassicaceae_Guiraoa_arvensis
Brassicaceae_Brassica_carinata
Brassicaceae_Brassica_nigra
Brassicaceae_Sinapis_arvensis
Brassicaceae_Trachystoma_labasii
Brassicaceae_Kremeriella_cordylocarpus
Brassicaceae_Sinapis_alba
Brassicaceae_Trachystoma_aphanoneurum
Brassicaceae_Ceratocnemum_rapistroides
Brassicaceae_Trachystoma_ballii
Brassicaceae_Raffenaldia_primuloides
Brassicaceae_Otocarpus_virgatus
Brassicaceae_Cordylocarpus_muricatus
Brassicaceae_Brassica_spinescens
Brassicaceae_Erucastrum_elatum
Brassicaceae_Hirschfeldia_incana
Brassicaceae_Erucastrum_virgatum
Brassicaceae_Diplotaxis_catholica
Brassicaceae_Muricaria_prostrata
Brassicaceae_Crambe_filiformis
Brassicaceae_Crambe_hispanica
Brassicaceae_Crambe_maritima
Brassicaceae_Crambe_orientalis
Brassicaceae_Brassica_repanda
Brassicaceae_Brassica_desnottesii
Brassicaceae_Eruca_pinnatifida
Brassicaceae_Moricandia_foleyi
Brassicaceae_Eruca_vesicaria
Brassicaceae_Diplotaxis_tenuifolia
4
0
0
46
98
100
0
1
100
5
56
3
43
26
10
1
44
49
50
0
84
0
98
100
7
100
2
5
7
5
100
0
8
99
6
96
55
58
91
3
5
81
21
2
45
98
51
92
0
100
63
91
20
75

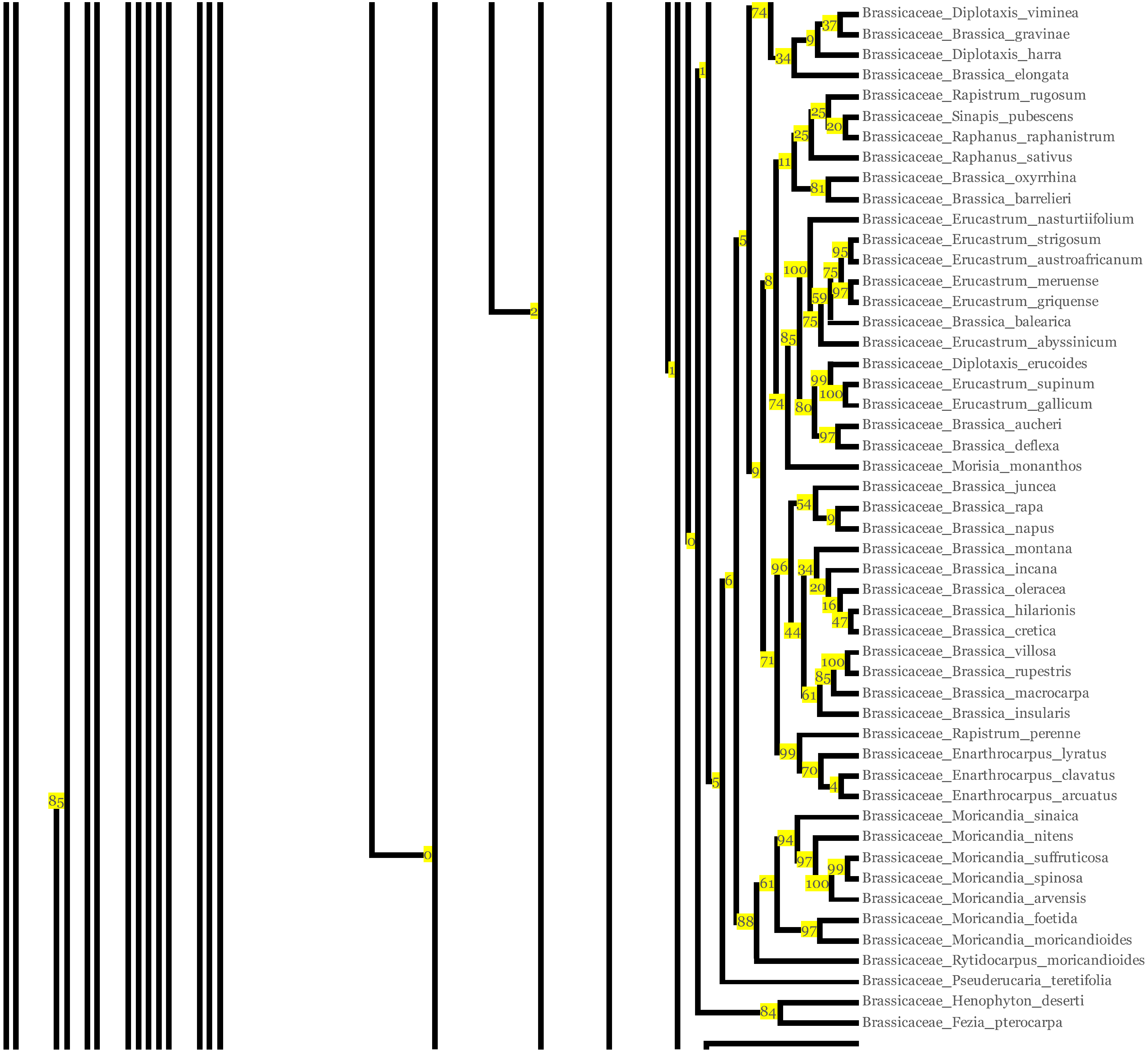
Brassicaceae_Diplotaxis_viminea
Brassicaceae_Brassica_gravinae
Brassicaceae_Diplotaxis_harra
Brassicaceae_Brassica_elongata
Brassicaceae_Rapistrum_rugosum
Brassicaceae_Sinapis_pubescens
Brassicaceae_Raphanus_raphanistrum
Brassicaceae_Raphanus_sativus
Brassicaceae_Brassica_oxyrrhina
Brassicaceae_Brassica_barrelieri
Brassicaceae_Erucastrum_nasturtiifolium
Brassicaceae_Erucastrum_strigosum
Brassicaceae_Erucastrum_austroafricanum
Brassicaceae_Erucastrum_meruense
Brassicaceae_Erucastrum_griquense
Brassicaceae_Brassica_balearica
Brassicaceae_Erucastrum_abyssinicum
Brassicaceae_Diplotaxis_erucoides
Brassicaceae_Erucastrum_supinum
Brassicaceae_Erucastrum_gallicum
Brassicaceae_Brassica_aucheri
Brassicaceae_Brassica_deflexa
Brassicaceae_Morisia_monanthos
Brassicaceae_Brassica_juncea
Brassicaceae_Brassica_rapa
Brassicaceae_Brassica_napus
Brassicaceae_Brassica_montana
Brassicaceae_Brassica_incana
Brassicaceae_Brassica_oleracea
Brassicaceae_Brassica_hilarionis
Brassicaceae_Brassica_cretica
Brassicaceae_Brassica_villosa
Brassicaceae_Brassica_rupestris
Brassicaceae_Brassica_macrocarpa
Brassicaceae_Brassica_insularis
Brassicaceae_Rapistrum_perenne
Brassicaceae_Enarthrocarpus_lyratus
Brassicaceae_Enarthrocarpus_clavatus
Brassicaceae_Enarthrocarpus_arcuatus
Brassicaceae_Moricandia_sinaica
Brassicaceae_Moricandia_nitens
Brassicaceae_Moricandia_suffruticosa
Brassicaceae_Moricandia_spinosa
Brassicaceae_Moricandia_arvensis
Brassicaceae_Moricandia_foetida
Brassicaceae_Moricandia_moricandioides
Brassicaceae_Rytidocarpus_moricandioides
Brassicaceae_Pseuderucaria_teretifolia
Brassicaceae_Henophyton_deserti
Brassicaceae_Fezia_pterocarpa

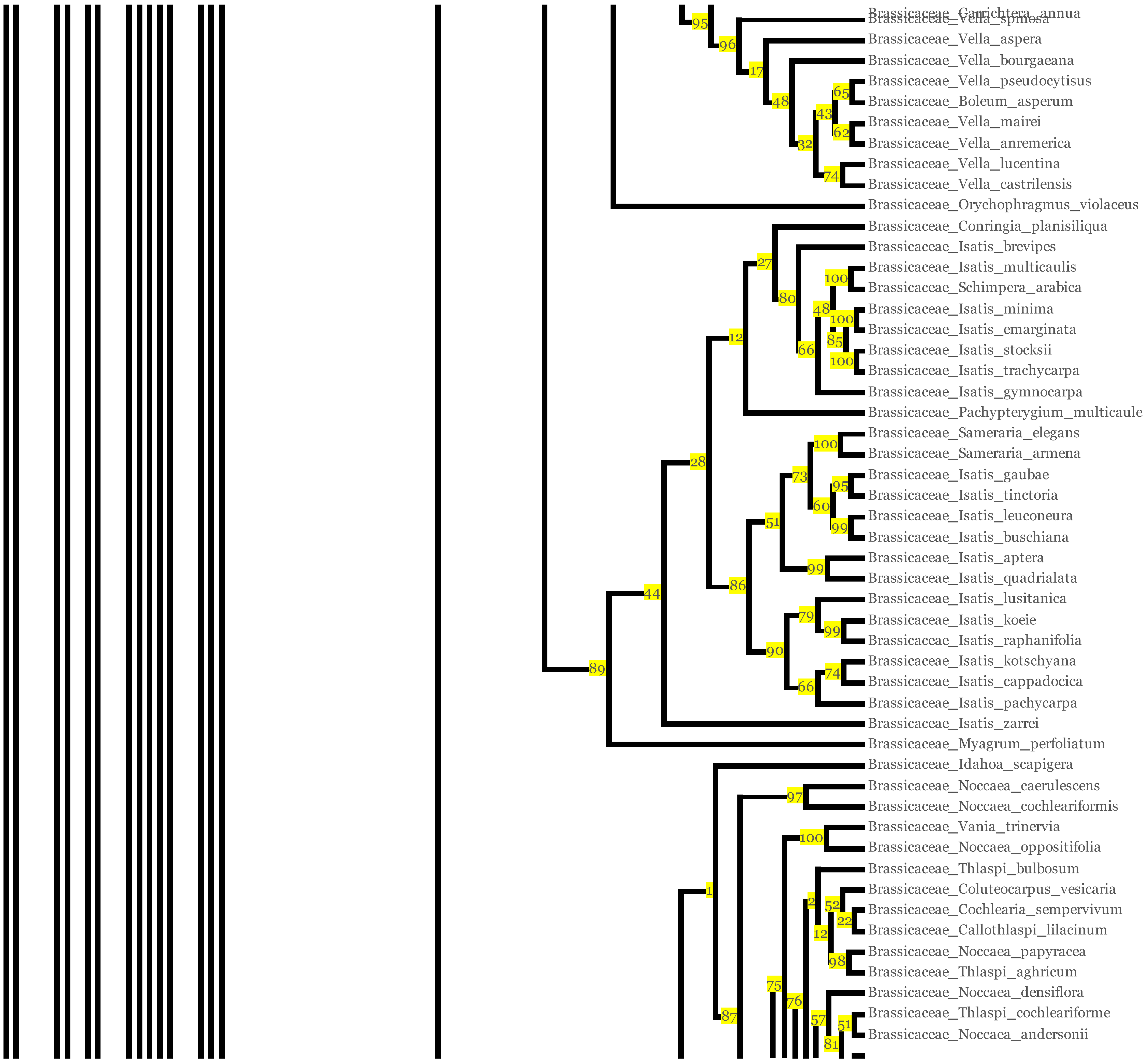
Brassicaceae_Carrichtera_annua
Brassicaceae_Vella_spinosa
Brassicaceae_Vella_aspera
Brassicaceae_Vella_bourgaeana
Brassicaceae_Vella_pseudocytisus
Brassicaceae_Boleum_asperum
Brassicaceae_Vella_mairei
Brassicaceae_Vella_anremerica
Brassicaceae_Vella_lucentina
Brassicaceae_Vella_castrilensis
Brassicaceae_Orychophragmus_violaceus
Brassicaceae_Conringia_planisiliqua
Brassicaceae_Isatis_brevipes
Brassicaceae_Isatis_multicaulis
Brassicaceae_Schimpera_arabica
Brassicaceae_Isatis_minima
Brassicaceae_Isatis_emarginata
Brassicaceae_Isatis_stocksii
Brassicaceae_Isatis_trachycarpa
Brassicaceae_Isatis_gymnocarpa
Brassicaceae_Pachypterygium_multicaule
Brassicaceae_Sameraria_elegans
Brassicaceae_Sameraria_armena
Brassicaceae_Isatis_gaubae
Brassicaceae_Isatis_tinctoria
Brassicaceae_Isatis_leuconeura
Brassicaceae_Isatis_buschiana
Brassicaceae_Isatis_aptera
Brassicaceae_Isatis_quadrialata
Brassicaceae_Isatis_lusitanica
Brassicaceae_Isatis_koeie
Brassicaceae_Isatis_raphanifolia
Brassicaceae_Isatis_kotschyana
Brassicaceae_Isatis_cappadocica
Brassicaceae_Isatis_pachycarpa
Brassicaceae_Isatis_zarrei
Brassicaceae_Myagrum_perfoliatum
Brassicaceae_Idahoa_scapigera
Brassicaceae_Noccaea_caerulescens
Brassicaceae_Noccaea_cochleariformis
Brassicaceae_Vania_trinervia
Brassicaceae_Noccaea_oppositifolia
Brassicaceae_Thlaspi_bulbosum
Brassicaceae_Coluteocarpus_vesicaria
Brassicaceae_Cochlearia_sempervivum
Brassicaceae_Callothlaspi_lilacinum
Brassicaceae_Noccaea_papyracea
Brassicaceae_Thlaspi_aghricum
Brassicaceae_Noccaea_densiflora
Brassicaceae_Thlaspi_cochleariforme
Brassicaceae_Noccaea_andersonii

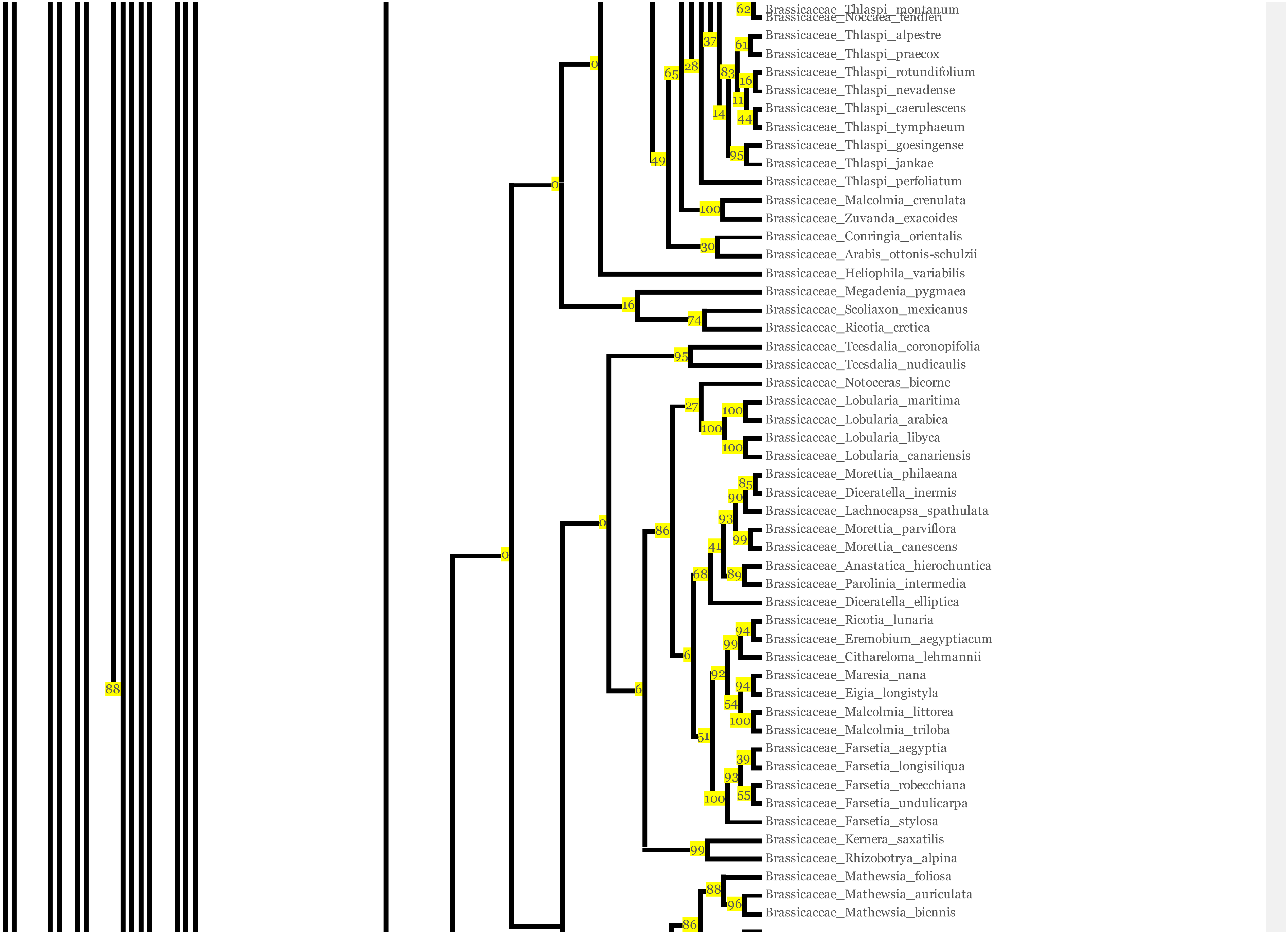
Brassicaceae_Thlaspi_montanum
Brassicaceae_Noccaea_fendleri
Brassicaceae_Thlaspi_alpestre
Brassicaceae_Thlaspi_praecox
Brassicaceae_Thlaspi_rotundifolium
Brassicaceae_Thlaspi_nevadense
Brassicaceae_Thlaspi_caerulescens
Brassicaceae_Thlaspi_tymphaeum
Brassicaceae_Thlaspi_goesingense
Brassicaceae_Thlaspi_jankae
Brassicaceae_Thlaspi_perfoliatum
Brassicaceae_Malcolmia_crenulata
Brassicaceae_Zuvanda_exacoides
Brassicaceae_Conringia_orientalis
Brassicaceae_Arabis_ottonis-schulzii
Brassicaceae_Heliophila_variabilis
Brassicaceae_Megadenia_pygmaea
Brassicaceae_Scoliaxon_mexicanus
Brassicaceae_Ricotia_cretica
Brassicaceae_Teesdalia_coronopifolia
Brassicaceae_Teesdalia_nudicaulis
Brassicaceae_Notoceras_bicorne
Brassicaceae_Lobularia_maritima
Brassicaceae_Lobularia_arabica
Brassicaceae_Lobularia_libyca
Brassicaceae_Lobularia_canariensis
Brassicaceae_Morettia_philaeana
Brassicaceae_Diceratella_inermis
Brassicaceae_Lachnocapsa_spathulata
Brassicaceae_Morettia_parviflora
Brassicaceae_Morettia_canescens
Brassicaceae_Anastatica_hierochuntica
Brassicaceae_Parolinia_intermedia
Brassicaceae_Diceratella_elliptica
Brassicaceae_Ricotia_lunaria
Brassicaceae_Eremobium_aegyptiacum
Brassicaceae_Cithareloma_lehmannii
Brassicaceae_Maresia_nana
Brassicaceae_Eigia_longistyla
Brassicaceae_Malcolmia_littorea
Brassicaceae_Malcolmia_triloba
Brassicaceae_Farsetia_aegyptia
Brassicaceae_Farsetia_longisiliqua
Brassicaceae_Farsetia_robecchiana
Brassicaceae_Farsetia_undulicarpa
Brassicaceae_Farsetia_stylosa
Brassicaceae_Kernera_saxatilis
Brassicaceae_Rhizobotrya_alpina
Brassicaceae_Mathewsia_foliosa
Brassicaceae_Mathewsia_auriculata
Brassicaceae_Mathewsia_biennis

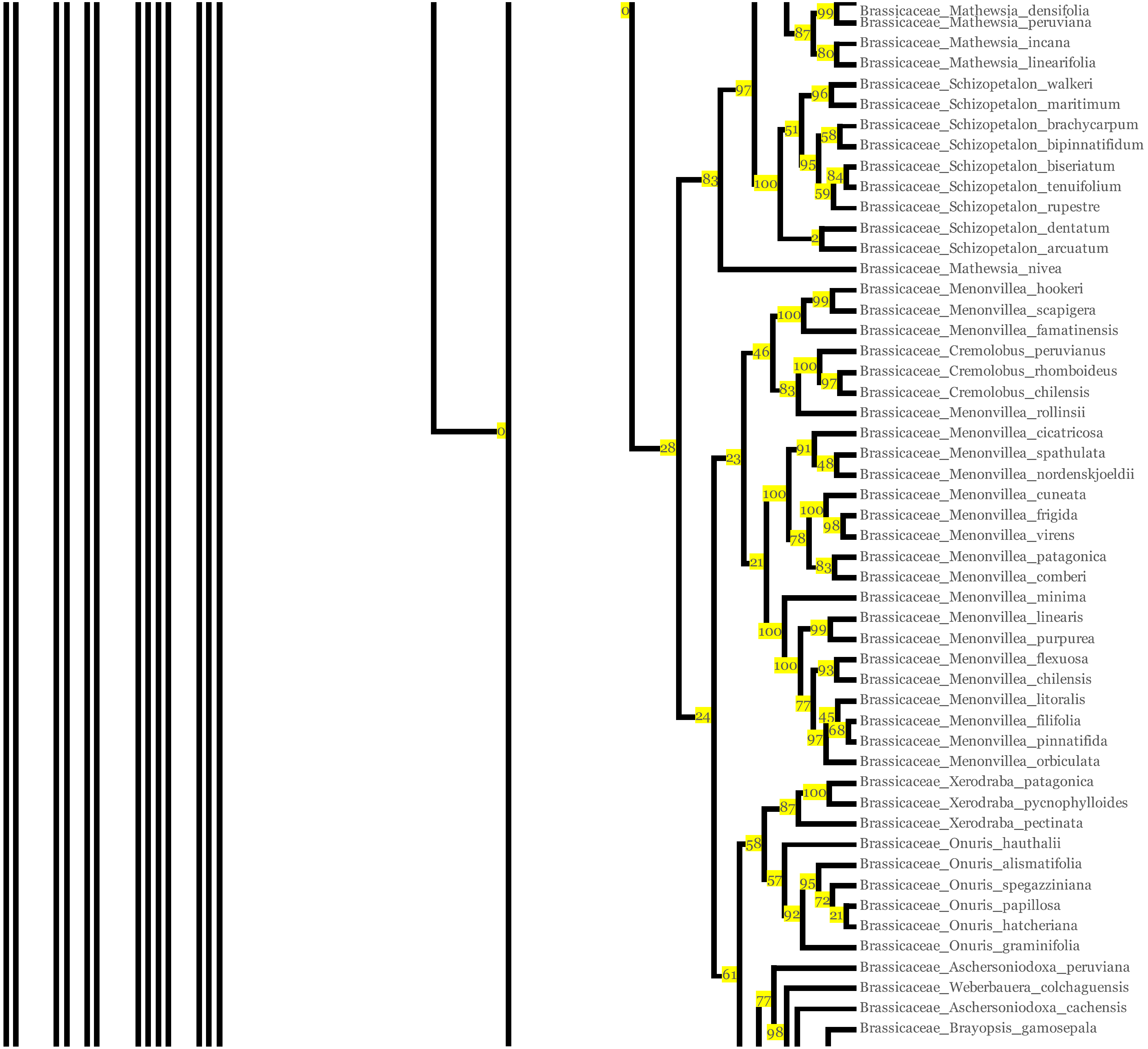
Brassicaceae_Mathewsia_densifolia
Brassicaceae_Mathewsia_peruviana
Brassicaceae_Mathewsia_incana
Brassicaceae_Mathewsia_linearifolia
Brassicaceae_Schizopetalon_walkeri
Brassicaceae_Schizopetalon_maritimum
Brassicaceae_Schizopetalon_brachycarpum
Brassicaceae_Schizopetalon_bipinnatifidum
Brassicaceae_Schizopetalon_biseriatum
Brassicaceae_Schizopetalon_tenuifolium
Brassicaceae_Schizopetalon_rupestre
Brassicaceae_Schizopetalon_dentatum
Brassicaceae_Schizopetalon_arcuatum
Brassicaceae_Mathewsia_nivea
Brassicaceae_Menonvillea_hookeri
Brassicaceae_Menonvillea_scapigera
Brassicaceae_Menonvillea_famatinensis
Brassicaceae_Cremolobus_peruvianus
Brassicaceae_Cremolobus_rhomboideus
Brassicaceae_Cremolobus_chilensis
Brassicaceae_Menonvillea_rollinsii
Brassicaceae_Menonvillea_cicatricosa
Brassicaceae_Menonvillea_spathulata
Brassicaceae_Menonvillea_nordenskjoeldii
Brassicaceae_Menonvillea_cuneata
Brassicaceae_Menonvillea_frigida
Brassicaceae_Menonvillea_virens
Brassicaceae_Menonvillea_patagonica
Brassicaceae_Menonvillea_comberi
Brassicaceae_Menonvillea_minima
Brassicaceae_Menonvillea_linearis
Brassicaceae_Menonvillea_purpurea
Brassicaceae_Menonvillea_flexuosa
Brassicaceae_Menonvillea_chilensis
Brassicaceae_Menonvillea_litoralis
Brassicaceae_Menonvillea_filifolia
Brassicaceae_Menonvillea_pinnatifida
Brassicaceae_Menonvillea_orbiculata
Brassicaceae_Xerodraba_patagonica
Brassicaceae_Xerodraba_pycnophylloides
Brassicaceae_Xerodraba_pectinata
Brassicaceae_Onuris_hauthalii
Brassicaceae_Onuris_alismatifolia
Brassicaceae_Onuris_spegazziniana
Brassicaceae_Onuris_papillosa
Brassicaceae_Onuris_hatcheriana
Brassicaceae_Onuris_graminifolia
Brassicaceae_Aschersoniodoxa_peruviana
Brassicaceae_Weberbauera_colchaguensis
Brassicaceae_Aschersoniodoxa_cachensis
Brassicaceae_Brayopsis_gamosepala

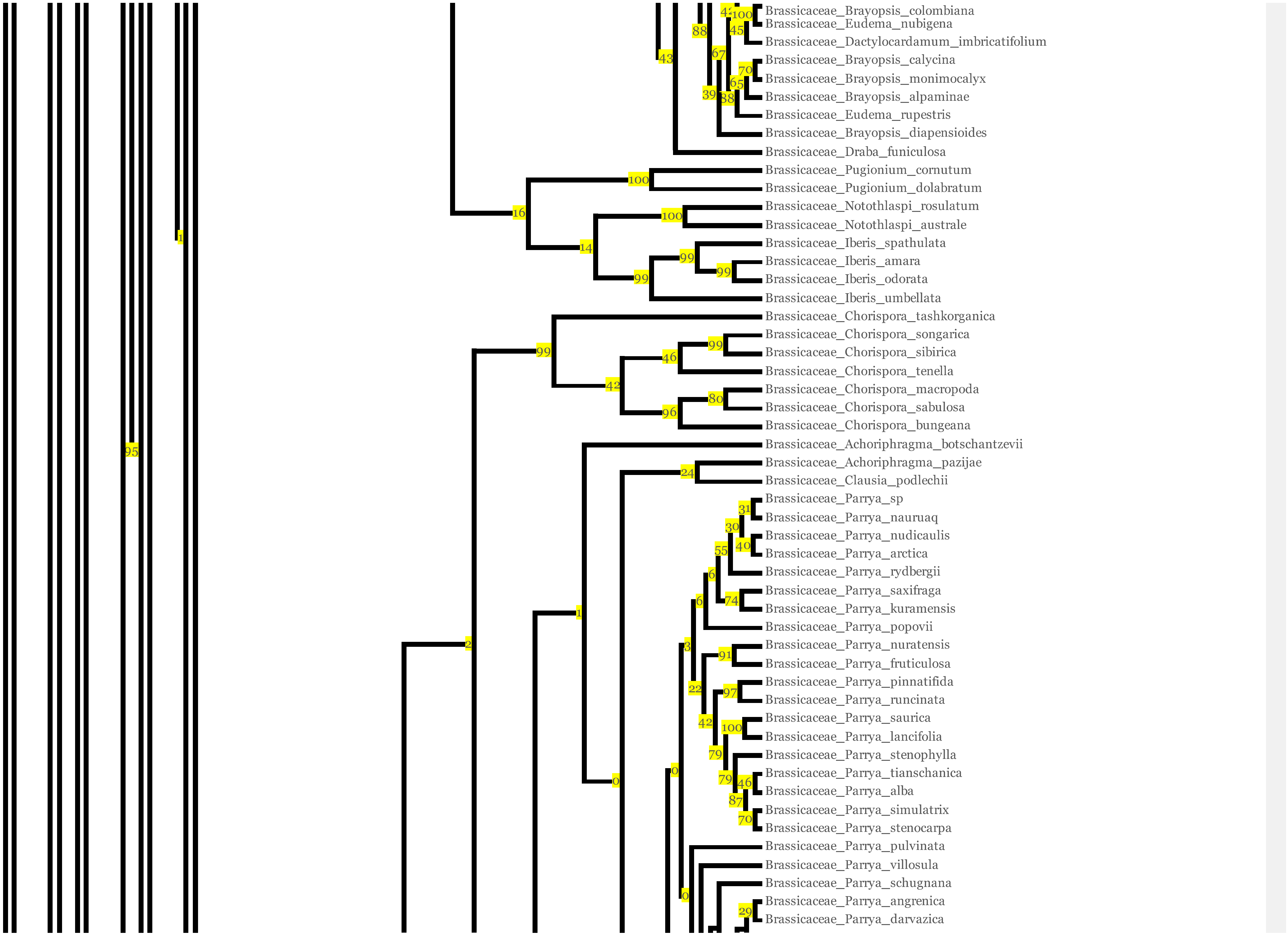
Brassicaceae_Brayopsis_colombiana
Brassicaceae_Eudema_nubigena
Brassicaceae_Dactylocardamum_imbricatifolium
Brassicaceae_Brayopsis_calycina
Brassicaceae_Brayopsis_monimocalyx
Brassicaceae_Brayopsis_alpaminae
Brassicaceae_Eudema_rupestris
Brassicaceae_Brayopsis_diapensioides
Brassicaceae_Draba_funiculosa
Brassicaceae_Pugionium_cornutum
Brassicaceae_Pugionium_dolabratum
Brassicaceae_Notothlaspi_rosulatum
Brassicaceae_Notothlaspi_australe
Brassicaceae_Iberis_spathulata
Brassicaceae_Iberis_amara
Brassicaceae_Iberis_odorata
Brassicaceae_Iberis_umbellata
Brassicaceae_Chorispora_tashkorganica
Brassicaceae_Chorispora_songarica
Brassicaceae_Chorispora_sibirica
Brassicaceae_Chorispora_tenella
Brassicaceae_Chorispora_macropoda
Brassicaceae_Chorispora_sabulosa
Brassicaceae_Chorispora_bungeana
Brassicaceae_Achoriphragma_botschantzevii
Brassicaceae_Achoriphragma_pazijae
Brassicaceae_Clausia_podlechii
Brassicaceae_Parrya_sp
Brassicaceae_Parrya_nauruaq
Brassicaceae_Parrya_nudicaulis
Brassicaceae_Parrya_arctica
Brassicaceae_Parrya_rydbergii
Brassicaceae_Parrya_saxifraga
Brassicaceae_Parrya_kuramensis
Brassicaceae_Parrya_popovii
Brassicaceae_Parrya_nuratensis
Brassicaceae_Parrya_fruticulosa
Brassicaceae_Parrya_pinnatifida
Brassicaceae_Parrya_runcinata
Brassicaceae_Parrya_saurica
Brassicaceae_Parrya_lancifolia
Brassicaceae_Parrya_stenophylla
Brassicaceae_Parrya_tianschanica
Brassicaceae_Parrya_alba
Brassicaceae_Parrya_simulatrix
Brassicaceae_Parrya_stenocarpa
Brassicaceae_Parrya_pulvinata
Brassicaceae_Parrya_villosula
Brassicaceae_Parrya_schugnana
Brassicaceae_Parrya_angrenica
Brassicaceae_Parrya_darvazica

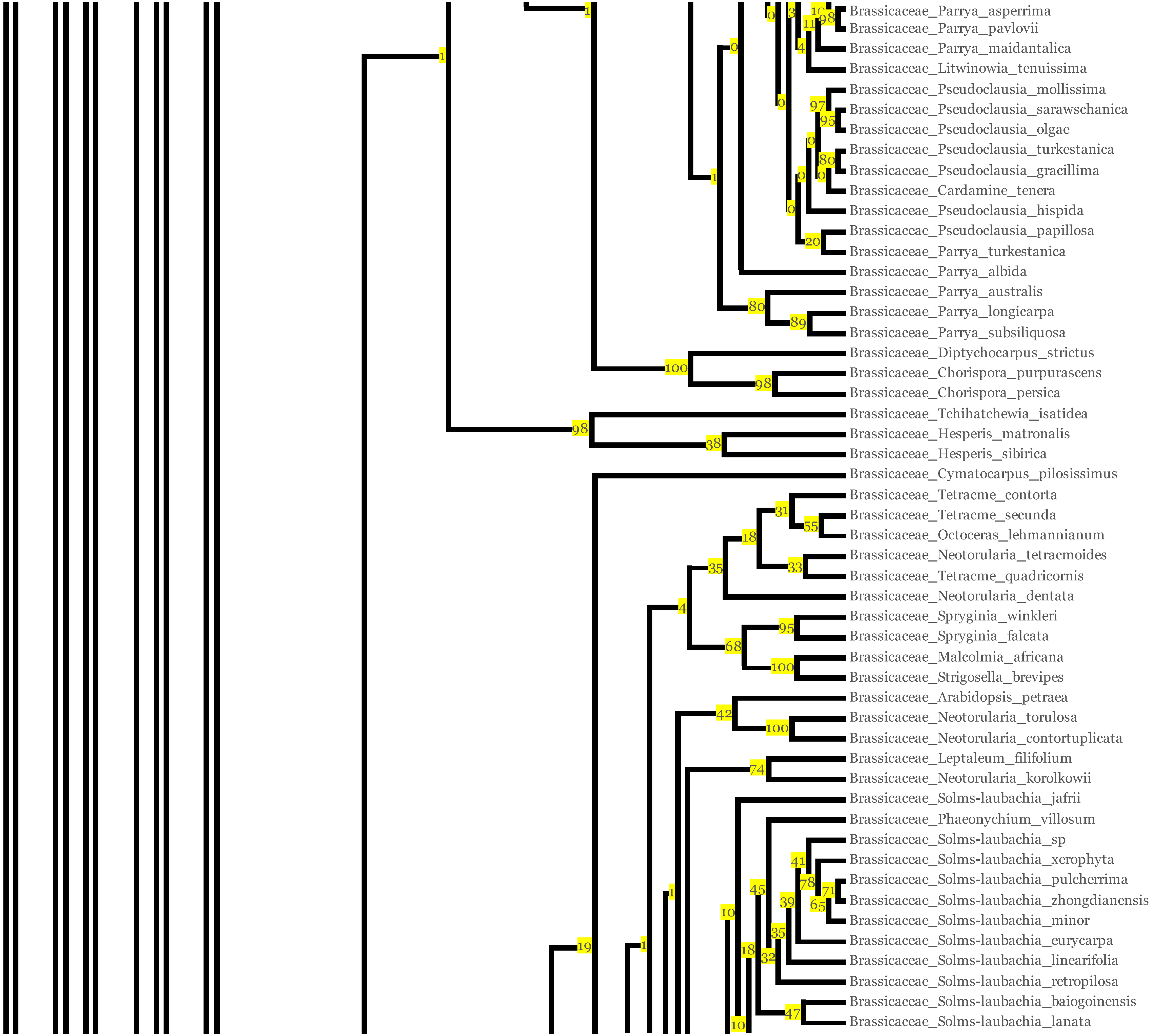
Brassicaceae_Parrya_asperrima
Brassicaceae_Parrya_pavlovii
Brassicaceae_Parrya_maidantalica
Brassicaceae_Litwinowia_tenuissima
Brassicaceae_Pseudoclausia_mollissima
Brassicaceae_Pseudoclausia_sarawschanica
Brassicaceae_Pseudoclausia_olgae
Brassicaceae_Pseudoclausia_turkestanica
Brassicaceae_Pseudoclausia_gracillima
Brassicaceae_Cardamine_tenera
Brassicaceae_Pseudoclausia_hispida
Brassicaceae_Pseudoclausia_papillosa
Brassicaceae_Parrya_turkestanica
Brassicaceae_Parrya_albida
Brassicaceae_Parrya_australis
Brassicaceae_Parrya_longicarpa
Brassicaceae_Parrya_subsiliquosa
Brassicaceae_Diptychocarpus_strictus
Brassicaceae_Chorispora_purpurascens
Brassicaceae_Chorispora_persica
Brassicaceae_Tchihatchewia_isatidea
Brassicaceae_Hesperis_matronalis
Brassicaceae_Hesperis_sibirica
Brassicaceae_Cymatocarpus_pilosissimus
Brassicaceae_Tetracme_contorta
Brassicaceae_Tetracme_secunda
Brassicaceae_Octoceras_lehmannianum
Brassicaceae_Neotorularia_tetracmoides
Brassicaceae_Tetracme_quadricornis
Brassicaceae_Neotorularia_dentata
Brassicaceae_Sprygina_winkleri
Brassicaceae_Sprygina_falcata
Brassicaceae_Malcolmia_africana
Brassicaceae_Strigosella_brevipes
Brassicaceae_Arabidopsis_petraea
Brassicaceae_Neotorularia_torulosa
Brassicaceae_Neotorularia_contortuplicata
Brassicaceae_Leptaleum_filifolium
Brassicaceae_Neotorularia_korolkowii
Brassicaceae_Solms-laubachia_jafrii
Brassicaceae_Phaeonychium_villosum
Brassicaceae_Solms-laubachia_sp
Brassicaceae_Solms-laubachia_xerophyta
Brassicaceae_Solms-laubachia_pulcherrima
Brassicaceae_Solms-laubachia_zhongdianensis
Brassicaceae_Solms-laubachia_minor
Brassicaceae_Solms-laubachia_eurycarpa
Brassicaceae_Solms-laubachia_linearifolia
Brassicaceae_Solms-laubachia_retropilosa
Brassicaceae_Solms-laubachia_baiogoinensis
Brassicaceae_Solms-laubachia_lanata

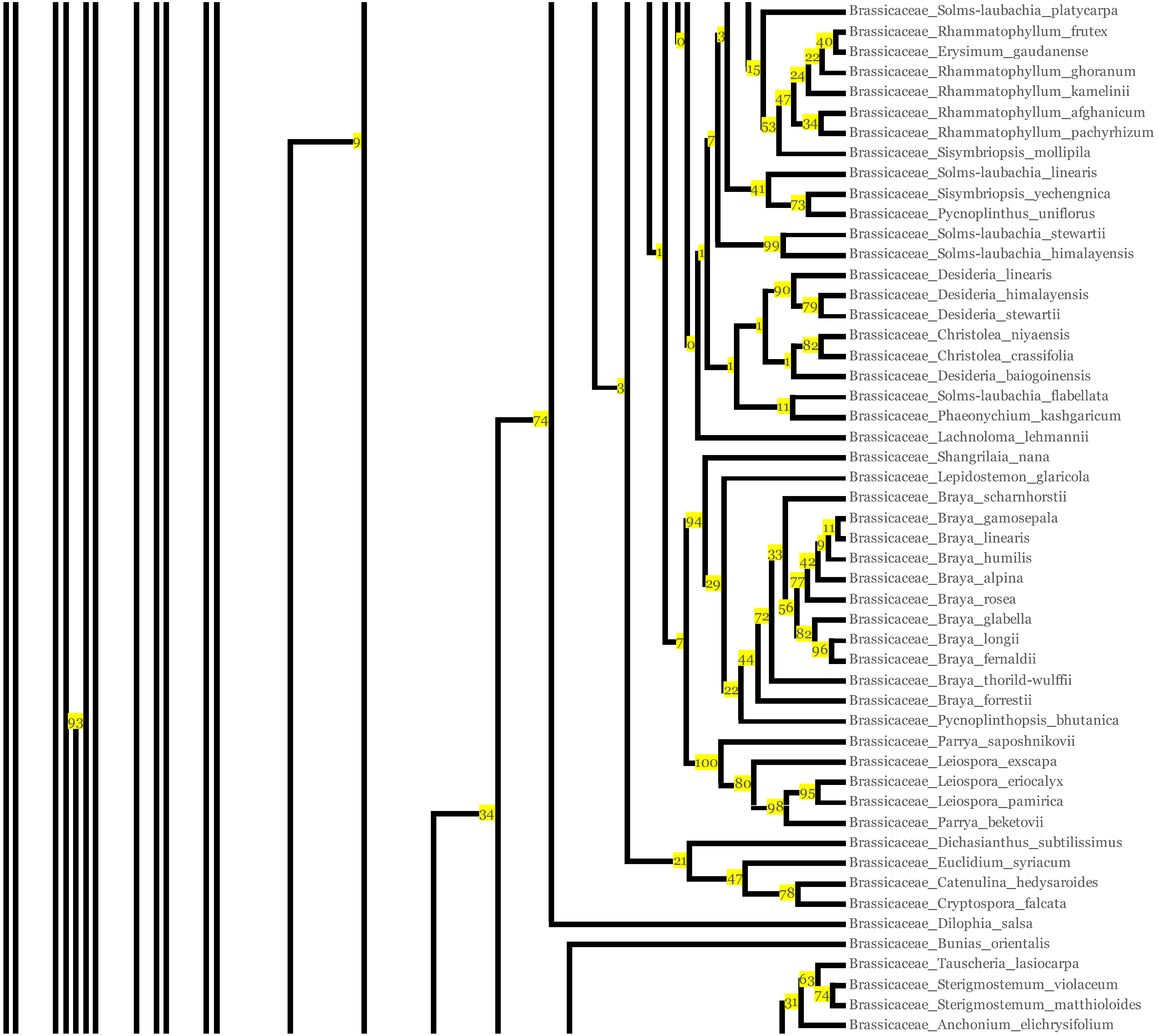
Brassicaceae_Solms-laubachia_platycarpa
Brassicaceae_Rhammatophyllum_frutex
Brassicaceae_Erysimum_gaudanense
Brassicaceae_Rhammatophyllum_ghoranum
Brassicaceae_Rhammatophyllum_kamelinii
Brassicaceae_Rhammatophyllum_afghanicum
Brassicaceae_Rhammatophyllum_pachyrhizum
Brassicaceae_Sisymbriopsis_mollipila
Brassicaceae_Solms-laubachia_linearis
Brassicaceae_Sisymbriopsis_yechengnica
Brassicaceae_Pycnoplinthus_uniflorus
Brassicaceae_Solms-laubachia_stewartii
Brassicaceae_Solms-laubachia_himalayensis
Brassicaceae_Desideria_linearis
Brassicaceae_Desideria_himalayensis
Brassicaceae_Desideria_stewartii
Brassicaceae_Christolea_niyaensis
Brassicaceae_Christolea_crassifolia
Brassicaceae_Desideria_baiogoinensis
Brassicaceae_Solms-laubachia_flabellata
Brassicaceae_Phaeonychium_kashgaricum
Brassicaceae_Lachnoloma_lehmannii
Brassicaceae_Shangrilaia_nana
Brassicaceae_Lepidostemon_glaricola
Brassicaceae_Braya_scharnhorstii
Brassicaceae_Braya_gamosepala
Brassicaceae_Braya_linearis
Brassicaceae_Braya_humilis
Brassicaceae_Braya_alpina
Brassicaceae_Braya_rosea
Brassicaceae_Braya_glabella
Brassicaceae_Braya_longii
Brassicaceae_Braya_fernaldii
Brassicaceae_Braya_thorild-wulffii
Brassicaceae_Braya_forrestii
Brassicaceae_Pycnoplinthopsis_bhutanica
Brassicaceae_Parrya_saposhnikovii
Brassicaceae_Leiospora_exscapa
Brassicaceae_Leiospora_eriocalyx
Brassicaceae_Leiospora_pamirica
Brassicaceae_Parrya_beketovii
Brassicaceae_Dichasianthus_subtilissimus
Brassicaceae_Euclidium_syriacum
Brassicaceae_Catenulina_hedysaroides
Brassicaceae_Cryptospora_falcata
Brassicaceae_Dilophia_salsa
Brassicaceae_Bunias_orientalis
Brassicaceae_Tauscheria_lasiocarpa
Brassicaceae_Sterigmostemum_violaceum
Brassicaceae_Sterigmostemum_matthioloides
Brassicaceae_Anchonium_elichrysifolium

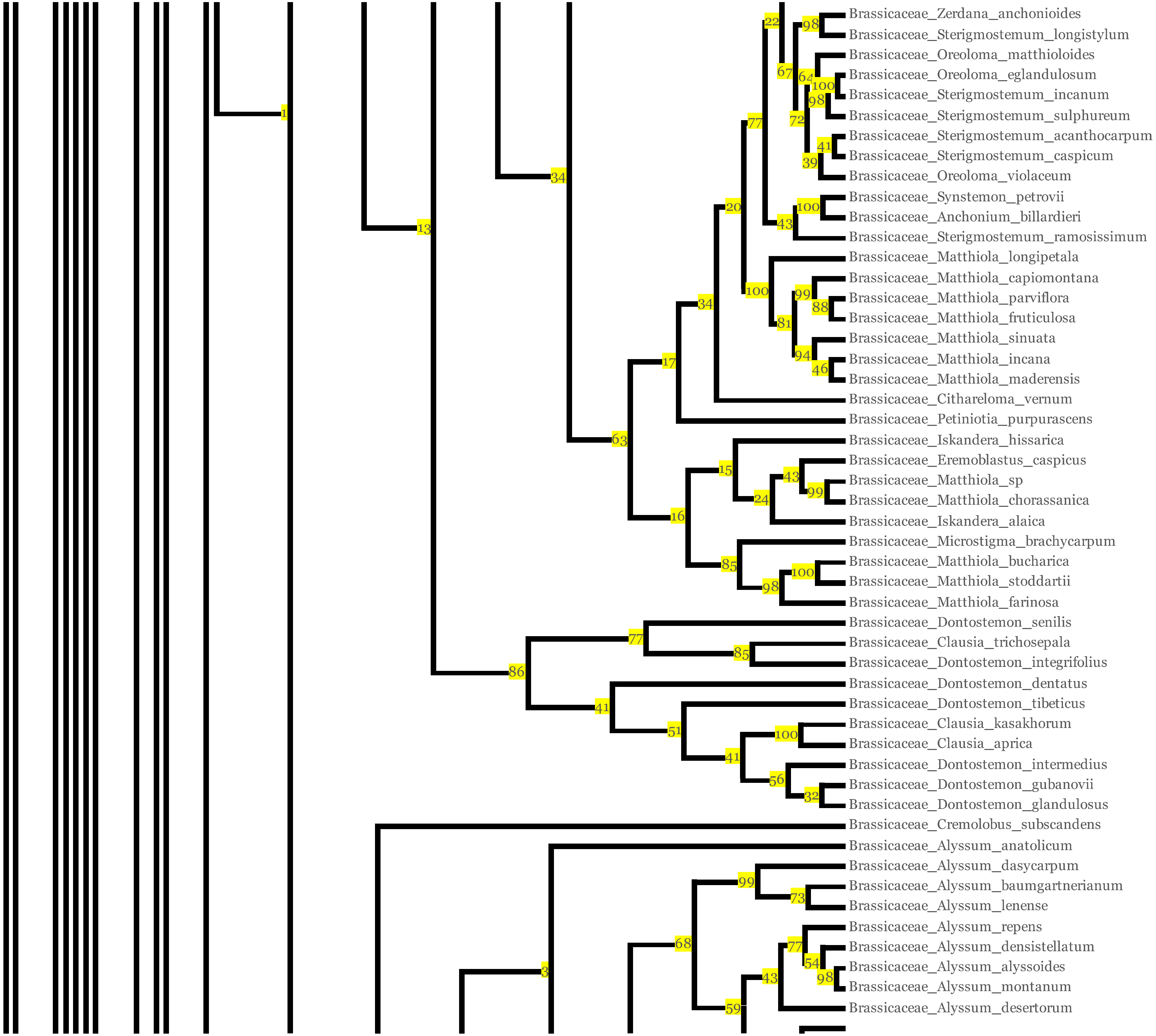
Brassicaceae_Zerdana_anchonioides
Brassicaceae_Sterigmostemum_longistylum
Brassicaceae_Oreoloma_matthioloides
Brassicaceae_Oreoloma_eglandulosum
Brassicaceae_Sterigmostemum_incanum
Brassicaceae_Sterigmostemum_sulphureum
Brassicaceae_Sterigmostemum_acanthocarpum
Brassicaceae_Sterigmostemum_caspicum
Brassicaceae_Oreoloma_violaceum
Brassicaceae_Synstemon_petrovii
Brassicaceae_Anchonium_billardieri
Brassicaceae_Sterigmostemum_ramosissimum
Brassicaceae_Matthiola_longipetala
Brassicaceae_Matthiola_capiomontana
Brassicaceae_Matthiola_parviflora
Brassicaceae_Matthiola_fruticulosa
Brassicaceae_Matthiola_sinuata
Brassicaceae_Matthiola_incana
Brassicaceae_Matthiola_maderensis
Brassicaceae_Cithareloma_vernum
Brassicaceae_Petiniotia_purpurascens
Brassicaceae_Iskandera_hissarica
Brassicaceae_Eremoblastus_caspicus
Brassicaceae_Matthiola_sp
Brassicaceae_Matthiola_chorassanica
Brassicaceae_Iskandera_alaica
Brassicaceae_Microstigma_brachycarpum
Brassicaceae_Matthiola_bucharica
Brassicaceae_Matthiola_stoddartii
Brassicaceae_Matthiola_farinosa
Brassicaceae_Dontostemon_senilis
Brassicaceae_Clausia_trichosepala
Brassicaceae_Dontostemon_integrifolius
Brassicaceae_Dontostemon_dentatus
Brassicaceae_Dontostemon_tibeticus
Brassicaceae_Clausia_kasakhorum
Brassicaceae_Clausia_aprica
Brassicaceae_Dontostemon_intermedius
Brassicaceae_Dontostemon_gubanovii
Brassicaceae_Dontostemon_glandulosus
Brassicaceae_Cremolobus_subscandens
Brassicaceae_Alyssum_anatolicum
Brassicaceae_Alyssum_dasycarpum
Brassicaceae_Alyssum_baumgartnerianum
Brassicaceae_Alyssum_lenense
Brassicaceae_Alyssum_repens
Brassicaceae_Alyssum_densistellatum
Brassicaceae_Alyssum_alyssoides
Brassicaceae_Alyssum_montanum
Brassicaceae_Alyssum_desertorum

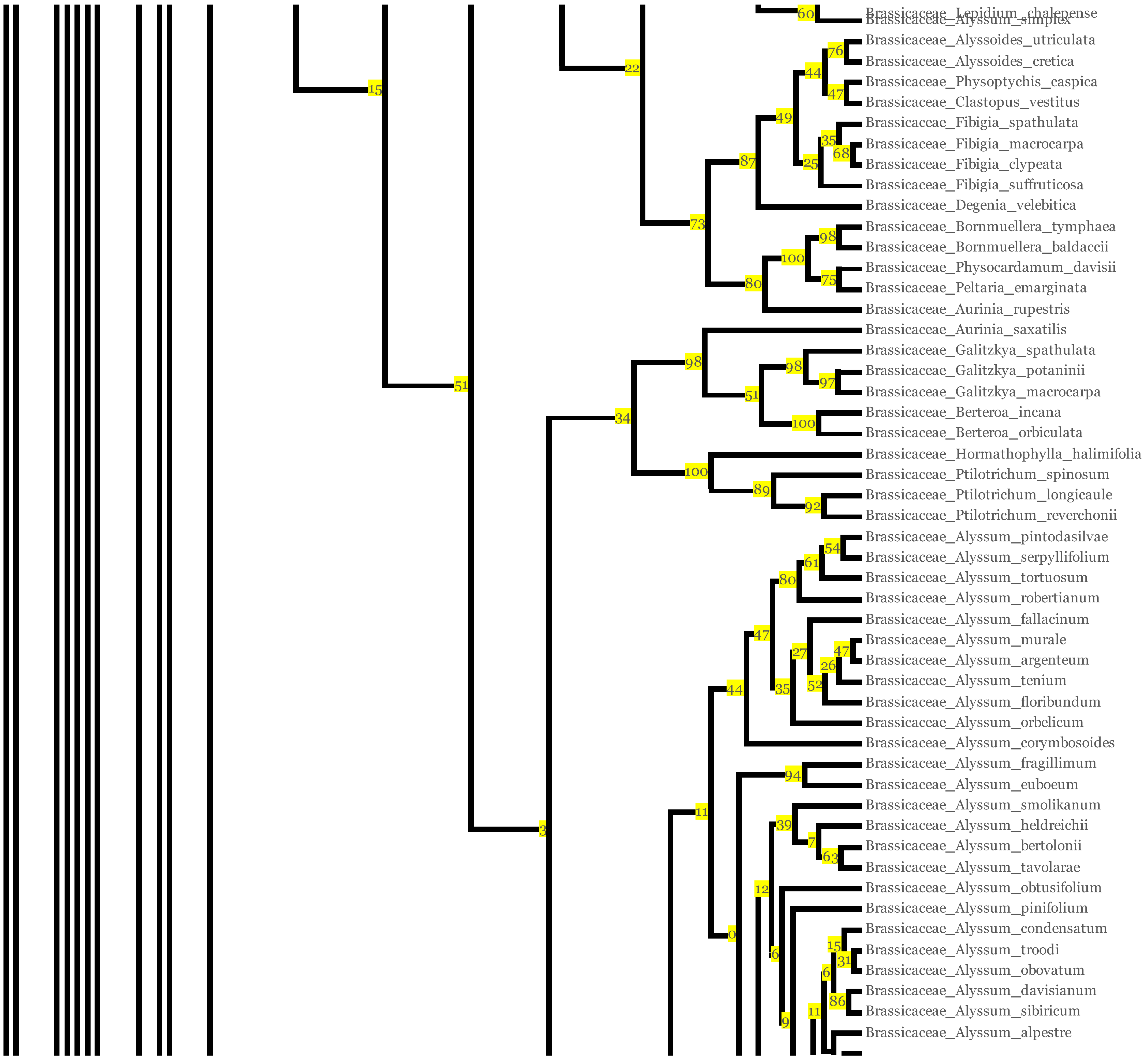
Brassicaceae_Lepidium_chalepense
Brassicaceae_Alyssum_simplex
Brassicaceae_Alyssoides_utriculata
Brassicaceae_Alyssoides_cretica
Brassicaceae_Physoptychis_caspica
Brassicaceae_Clastopus_vestitus
Brassicaceae_Fibigia_spathulata
Brassicaceae_Fibigia_macrocarpa
Brassicaceae_Fibigia_clypeata
Brassicaceae_Fibigia_suffruticosa
Brassicaceae_Degenia_velebitica
Brassicaceae_Bornmuellera_tymphaea
Brassicaceae_Bornmuellera_baldaccii
Brassicaceae_Physocardamum_davisii
Brassicaceae_Peltaria_emarginata
Brassicaceae_Aurinia_rupestris
Brassicaceae_Aurinia_saxatilis
Brassicaceae_Galitzkya_spathulata
Brassicaceae_Galitzkya_potaninii
Brassicaceae_Galitzkya_macrocarpa
Brassicaceae_Berteroa_incana
Brassicaceae_Berteroa_orbiculata
Brassicaceae_Hormathophylla_halimifolia
Brassicaceae_Ptilotrichum_spinosum
Brassicaceae_Ptilotrichum_longicaule
Brassicaceae_Ptilotrichum_reverchonii
Brassicaceae_Alyssum_pintodasilvae
Brassicaceae_Alyssum_serpyllifolium
Brassicaceae_Alyssum_tortuosum
Brassicaceae_Alyssum_robertianum
Brassicaceae_Alyssum_fallacinum
Brassicaceae_Alyssum_murale
Brassicaceae_Alyssum_argenteum
Brassicaceae_Alyssum_tenium
Brassicaceae_Alyssum_floribundum
Brassicaceae_Alyssum_orbelicum
Brassicaceae_Alyssum_corymbosoides
Brassicaceae_Alyssum_fragillimum
Brassicaceae_Alyssum_euboeum
Brassicaceae_Alyssum_smolikanum
Brassicaceae_Alyssum_heldreichii
Brassicaceae_Alyssum_bertolonii
Brassicaceae_Alyssum_tavolarae
Brassicaceae_Alyssum_obtusifolium
Brassicaceae_Alyssum_pinifolium
Brassicaceae_Alyssum_condensatum
Brassicaceae_Alyssum_troodi
Brassicaceae_Alyssum_obovatum
Brassicaceae_Alyssum_davisianum
Brassicaceae_Alyssum_sibiricum
Brassicaceae_Alyssum_alpestre

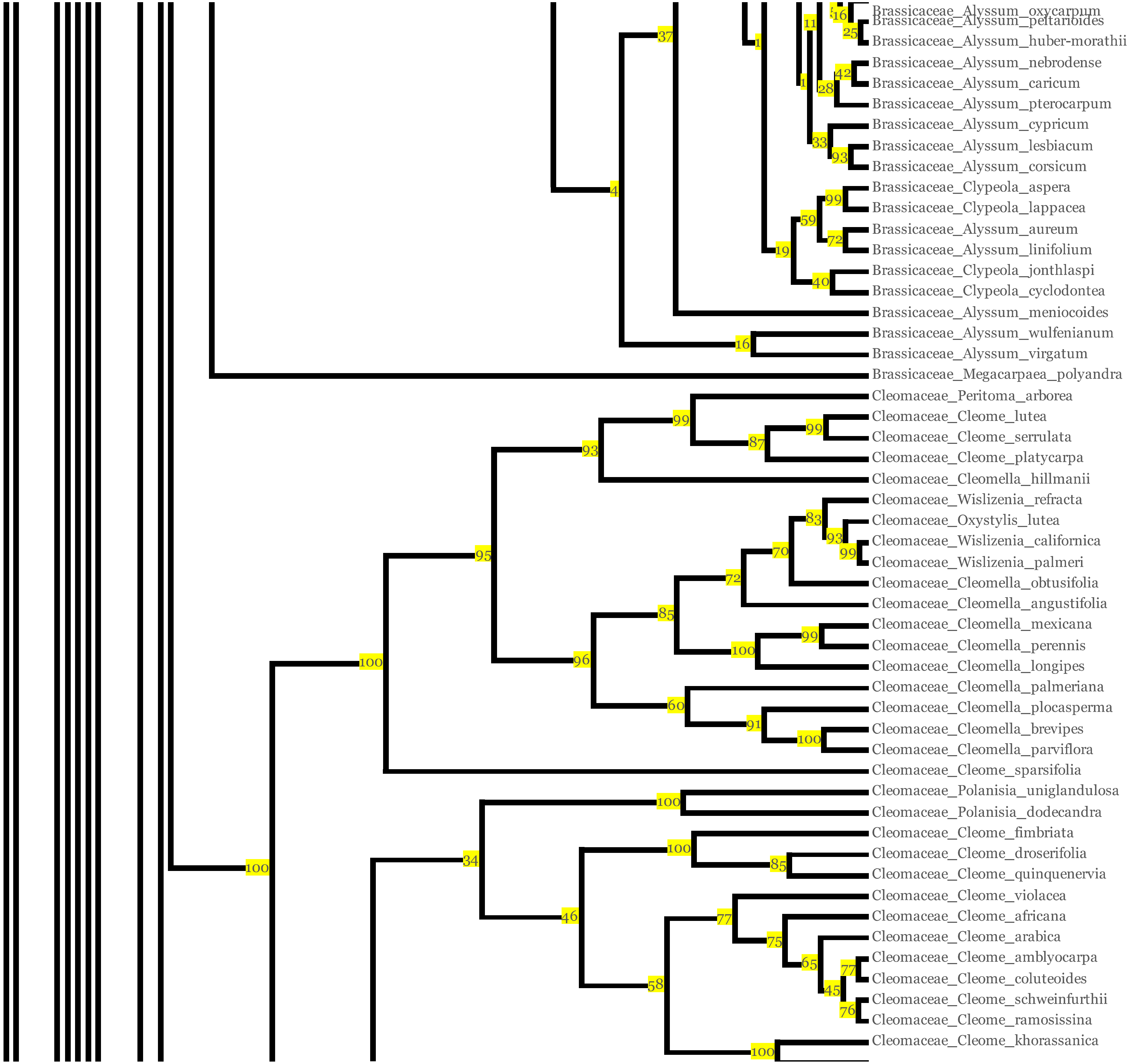
Brassicaceae_Alyssum_oxycarpum
Brassicaceae_Alyssum_peltarioides
Brassicaceae_Alyssum_huber-morathii
Brassicaceae_Alyssum_nebrodense
Brassicaceae_Alyssum_caricum
Brassicaceae_Alyssum_pterocarpum
Brassicaceae_Alyssum_cypricum
Brassicaceae_Alyssum_lesbiacum
Brassicaceae_Alyssum_corsicum
Brassicaceae_Clypeola_aspera
Brassicaceae_Clypeola_lappacea
Brassicaceae_Alyssum_aureum
Brassicaceae_Alyssum_linifolium
Brassicaceae_Clypeola_jonthlaspi
Brassicaceae_Clypeola_cyclodontea
Brassicaceae_Alyssum_meniocoides
Brassicaceae_Alyssum_wulfenianum
Brassicaceae_Alyssum_virgatum
Brassicaceae_Megacarpaea_polyandra
Cleomaceae_Peritoma_arborea
Cleomaceae_Cleome_lutea
Cleomaceae_Cleome_serrulata
Cleomaceae_Cleome_platycarpa
Cleomaceae_Cleomella_hillmanii
Cleomaceae_Wislizenia_refracta
Cleomaceae_Oxystylis_lutea
Cleomaceae_Wislizenia_californica
Cleomaceae_Wislizenia_palmeri
Cleomaceae_Cleomella_obtusifolia
Cleomaceae_Cleomella_angustifolia
Cleomaceae_Cleomella_mexicana
Cleomaceae_Cleomella_perennis
Cleomaceae_Cleomella_longipes
Cleomaceae_Cleomella_palmeriana
Cleomaceae_Cleomella_plocasperma
Cleomaceae_Cleomella_brevipes
Cleomaceae_Cleomella_parviflora
Cleomaceae_Cleome_sparsifolia
Cleomaceae_Polanisia_uniglandulosa
Cleomaceae_Polanisia_dodecandra
Cleomaceae_Cleome_fimbriata
Cleomaceae_Cleome_droserifolia
Cleomaceae_Cleome_quinquenervia
Cleomaceae_Cleome_violacea
Cleomaceae_Cleome_africana
Cleomaceae_Cleome_arabica
Cleomaceae_Cleome_amblyocarpa
Cleomaceae_Cleome_coluteoides
Cleomaceae_Cleome_schweinfurthii
Cleomaceae_Cleome_ramosissina
Cleomaceae_Cleome_khorassanica

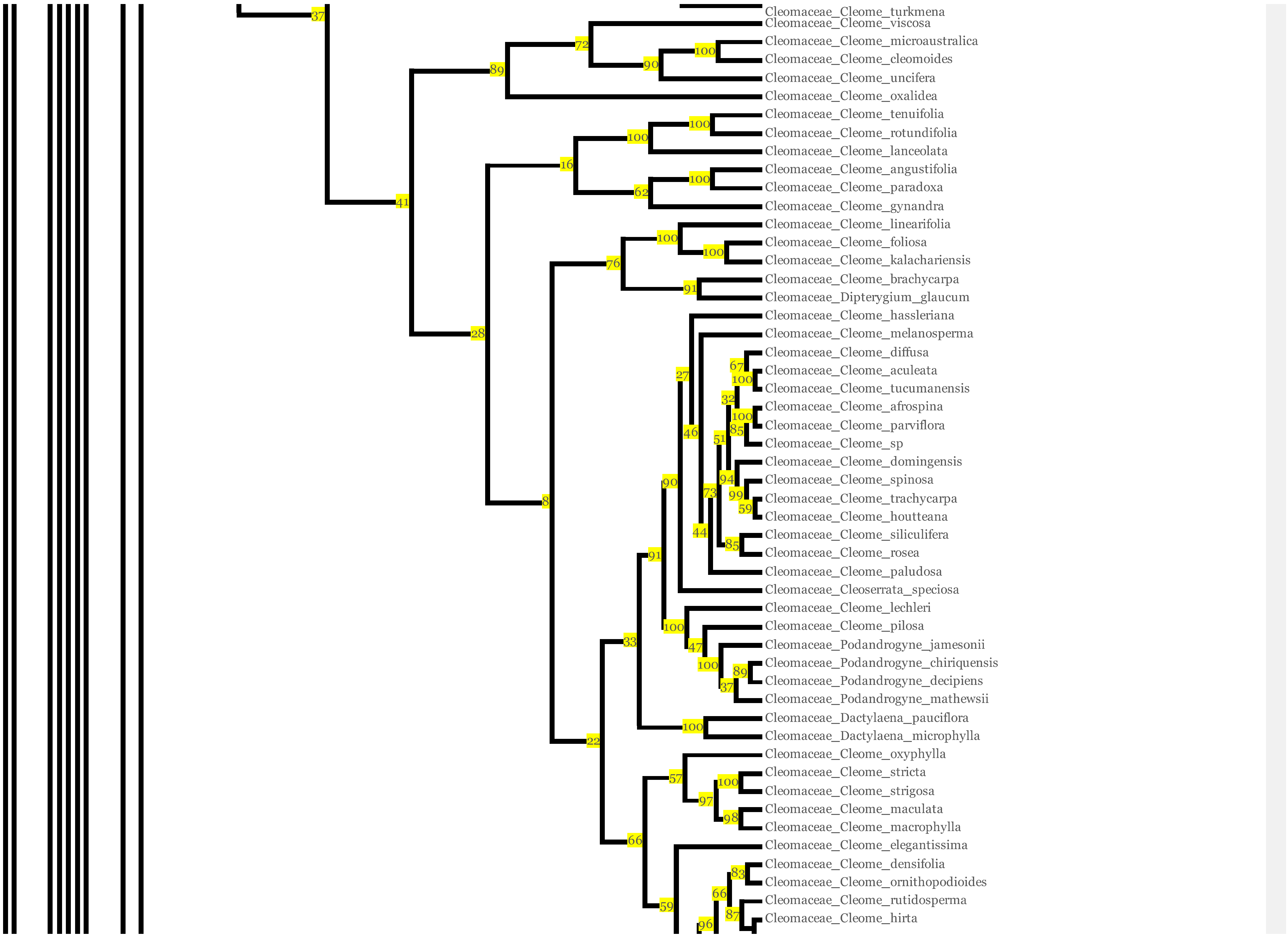
Cleomaceae_Cleome_turkmena
Cleomaceae_Cleome_viscosa
Cleomaceae_Cleome_microaustralica
Cleomaceae_Cleome_cleomoides
Cleomaceae_Cleome_uncifera
Cleomaceae_Cleome_oxalidea
Cleomaceae_Cleome_tenuifolia
Cleomaceae_Cleome_rotundifolia
Cleomaceae_Cleome_lanceolata
Cleomaceae_Cleome_angustifolia
Cleomaceae_Cleome_paradoxa
Cleomaceae_Cleome_gynandra
Cleomaceae_Cleome_linearifolia
Cleomaceae_Cleome_foliosa
Cleomaceae_Cleome_kalachariensis
Cleomaceae_Cleome_brachycarpa
Cleomaceae_Dipterygium_glaucum
Cleomaceae_Cleome_hassleriana
Cleomaceae_Cleome_melanosperma
Cleomaceae_Cleome_diffusa
Cleomaceae_Cleome_aculeata
Cleomaceae_Cleome_tucumanensis
Cleomaceae_Cleome_afrospina
Cleomaceae_Cleome_parviflora
Cleomaceae_Cleome_sp
Cleomaceae_Cleome_domingensis
Cleomaceae_Cleome_spinosa
Cleomaceae_Cleome_trachycarpa
Cleomaceae_Cleome_houtteana
Cleomaceae_Cleome_siliculifera
Cleomaceae_Cleome_rosea
Cleomaceae_Cleome_paludosa
Cleomaceae_Cleoserrata_speciosa
Cleomaceae_Cleome_lechleri
Cleomaceae_Cleome_pilosa
Cleomaceae_Podandrogyne_jamesonii
Cleomaceae_Podandrogyne_chiriquensis
Cleomaceae_Podandrogyne_decipiens
Cleomaceae_Podandrogyne_mathewsii
Cleomaceae_Dactylaena_pauciflora
Cleomaceae_Dactylaena_microphylla
Cleomaceae_Cleome_oxyphylla
Cleomaceae_Cleome_stricta
Cleomaceae_Cleome_strigosa
Cleomaceae_Cleome_maculata
Cleomaceae_Cleome_macrophylla
Cleomaceae_Cleome_elegantissima
Cleomaceae_Cleome_densifolia
Cleomaceae_Cleome_ornithopodioides
Cleomaceae_Cleome_rutidosperma
Cleomaceae_Cleome_hirta

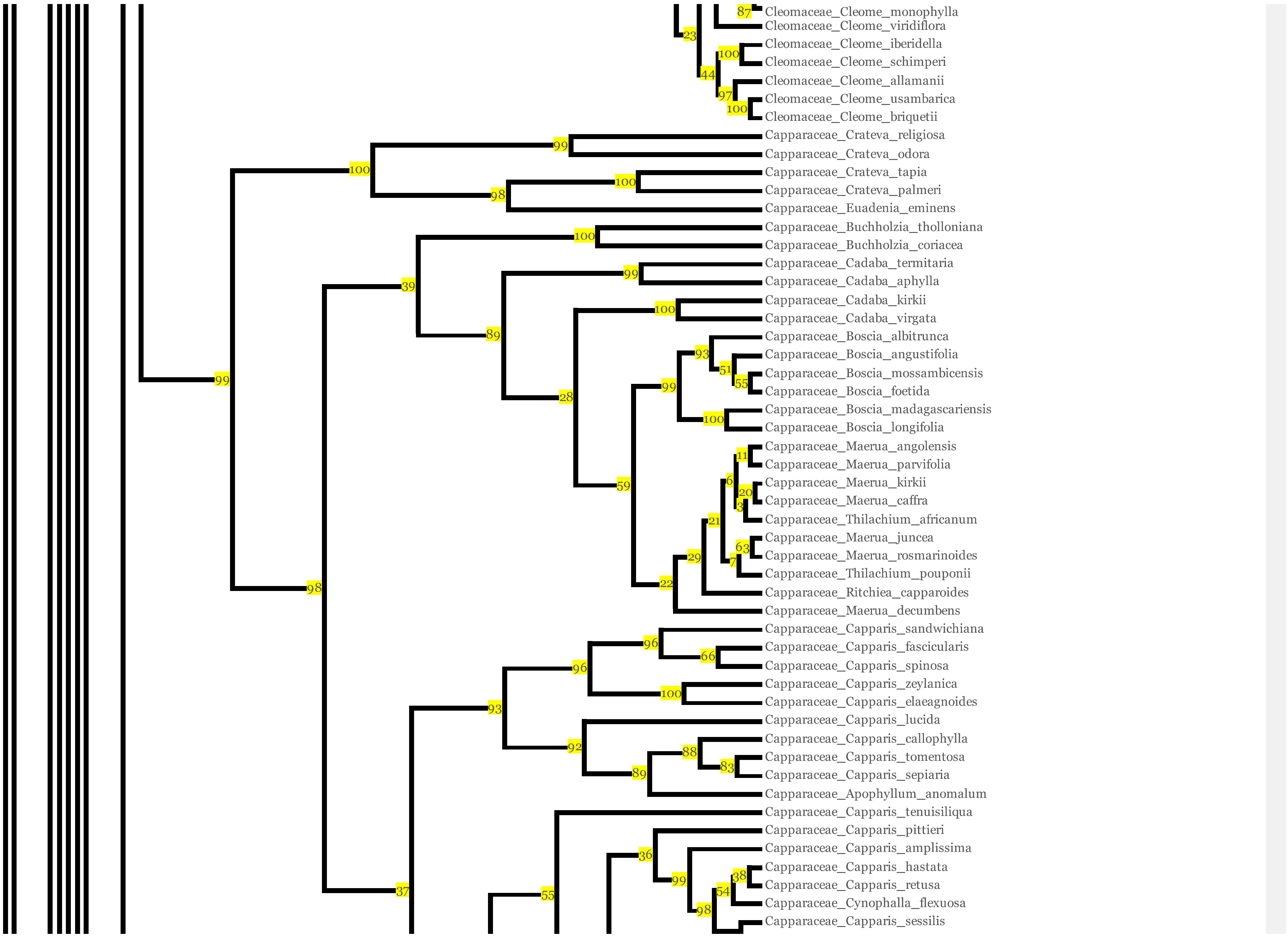
Cleomaceae_Cleome_monophylla
Cleomaceae_Cleome_viridiflora
Cleomaceae_Cleome_iberidella
Cleomaceae_Cleome_schimperi
Cleomaceae_Cleome_allamanii
Cleomaceae_Cleome_usambarica
Cleomaceae_Cleome_briquetii
Capparaceae_Crateva_religiosa
Capparaceae_Crateva_odora
Capparaceae_Crateva_tapia
Capparaceae_Crateva_palmeri
Capparaceae_Euadenia_eminens
Capparaceae_Buchholzia_tholloniana
Capparaceae_Buchholzia_coriacea
Capparaceae_Cadaba_termitaria
Capparaceae_Cadaba_aphylla
Capparaceae_Cadaba_kirkii
Capparaceae_Cadaba_virgata
Capparaceae_Boscia_albitrunca
Capparaceae_Boscia_angustifolia
Capparaceae_Boscia_mossambicensis
Capparaceae_Boscia_foetida
Capparaceae_Boscia_madagascariensis
Capparaceae_Boscia_longifolia
Capparaceae_Maerua_angolensis
Capparaceae_Maerua_parvifolia
Capparaceae_Maerua_kirkii
Capparaceae_Maerua_caffra
Capparaceae_Thilachium_africanum
Capparaceae_Maerua_juncea
Capparaceae_Maerua_rosmarinoides
Capparaceae_Thilachium_pouponii
Capparaceae_Ritchiea_capparoides
Capparaceae_Maerua_decumbens
Capparaceae_Capparis_sandwichiana
Capparaceae_Capparis_fascicularis
Capparaceae_Capparis_spinosa
Capparaceae_Capparis_zeylanica
Capparaceae_Capparis_elaeagnoides
Capparaceae_Capparis_lucida
Capparaceae_Capparis_callophylla
Capparaceae_Capparis_tomentosa
Capparaceae_Capparis_sepiaria
Capparaceae_Apophyllum_anomalum
Capparaceae_Capparis_tenuisiliqua
Capparaceae_Capparis_pittieri
Capparaceae_Capparis_amplissima
Capparaceae_Capparis_hastata
Capparaceae_Capparis_retusa
Capparaceae_Cynophalla_flexuosa
Capparaceae_Capparis_sessilis

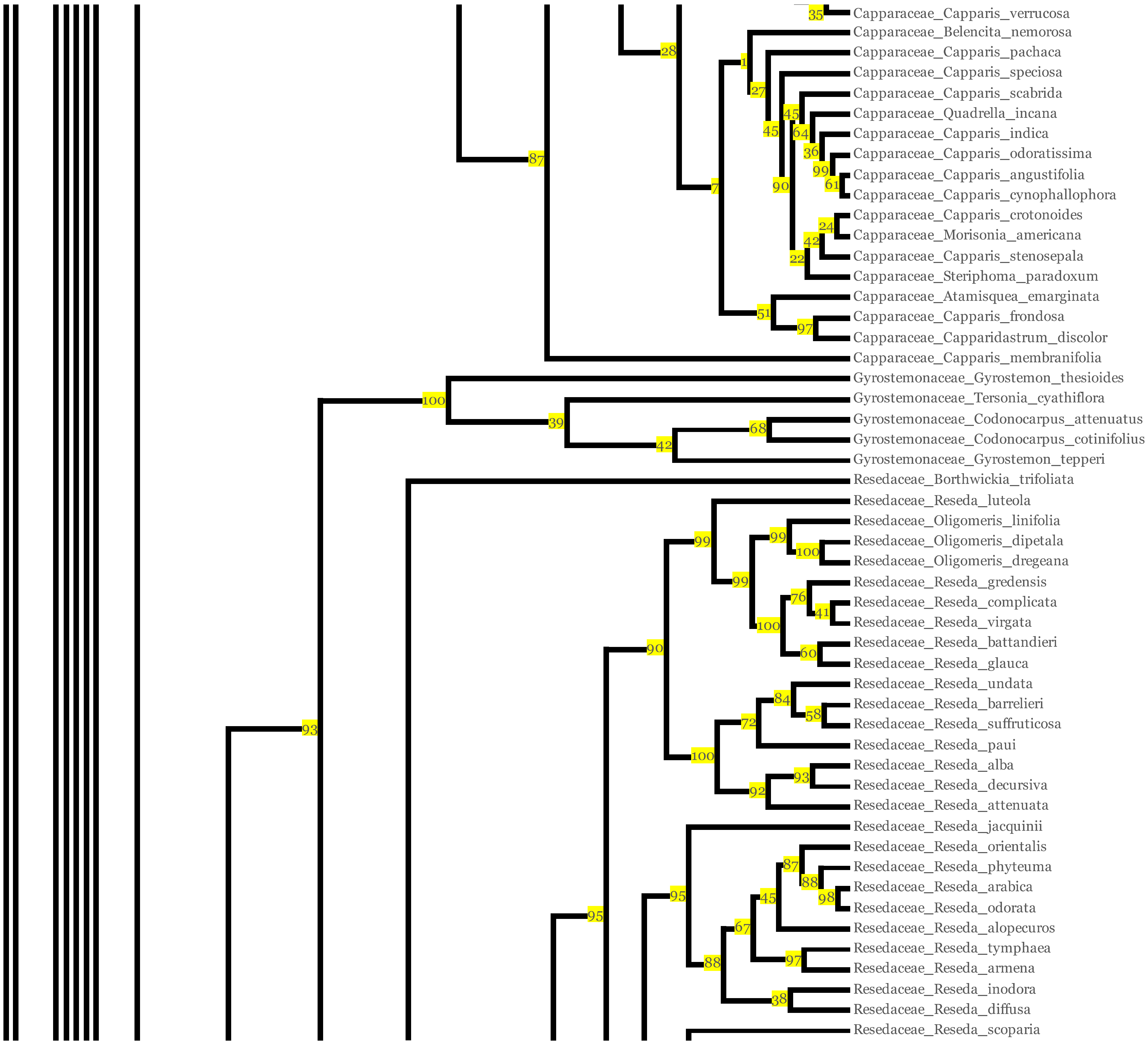
Capparaceae_Capparis_verrucosa
Capparaceae_Belencita_nemorosa
Capparaceae_Capparis_pachaca
Capparaceae_Capparis_speciosa
Capparaceae_Capparis_scabrida
Capparaceae_Quadrella_incana
Capparaceae_Capparis_indica
Capparaceae_Capparis_odoratissima
Capparaceae_Capparis_angustifolia
Capparaceae_Capparis_cynophallophora
Capparaceae_Capparis_crotonoides
Capparaceae_Morisonia_americana
Capparaceae_Capparis_stenosepala
Capparaceae_Steriphoma_paradoxum
Capparaceae_Atamisquea_emarginata
Capparaceae_Capparis_frondosa
Capparaceae_Capparidastrum_discolor
Capparaceae_Capparis_membranifolia
Gyrostemonaceae_Gyrostemon_thesioides
Gyrostemonaceae_Tersonia_cyathiflora
Gyrostemonaceae_Codonocarpus_attenuatus
Gyrostemonaceae_Codonocarpus_cotinifolius
Gyrostemonaceae_Gyrostemon_tepperi
Resedaceae_Borthwickia_trifoliata
Resedaceae_Reseda_luteola
Resedaceae_Oligomeris_linifolia
Resedaceae_Oligomeris_dipetala
Resedaceae_Oligomeris_dregeana
Resedaceae_Reseda_gredensis
Resedaceae_Reseda_complicata
Resedaceae_Reseda_virgata
Resedaceae_Reseda_battandieri
Resedaceae_Reseda_glauca
Resedaceae_Reseda_undata
Resedaceae_Reseda_barrelieri
Resedaceae_Reseda_suffruticosa
Resedaceae_Reseda_paui
Resedaceae_Reseda_alba
Resedaceae_Reseda_decursiva
Resedaceae_Reseda_attenuata
Resedaceae_Reseda_jacquinii
Resedaceae_Reseda_orientalis
Resedaceae_Reseda_phyteuma
Resedaceae_Reseda_arabica
Resedaceae_Reseda_odorata
Resedaceae_Reseda_alopecuros
Resedaceae_Reseda_tymphaea
Resedaceae_Reseda_armena
Resedaceae_Reseda_inodora
Resedaceae_Reseda_diffusa
Resedaceae_Reseda_scoparia

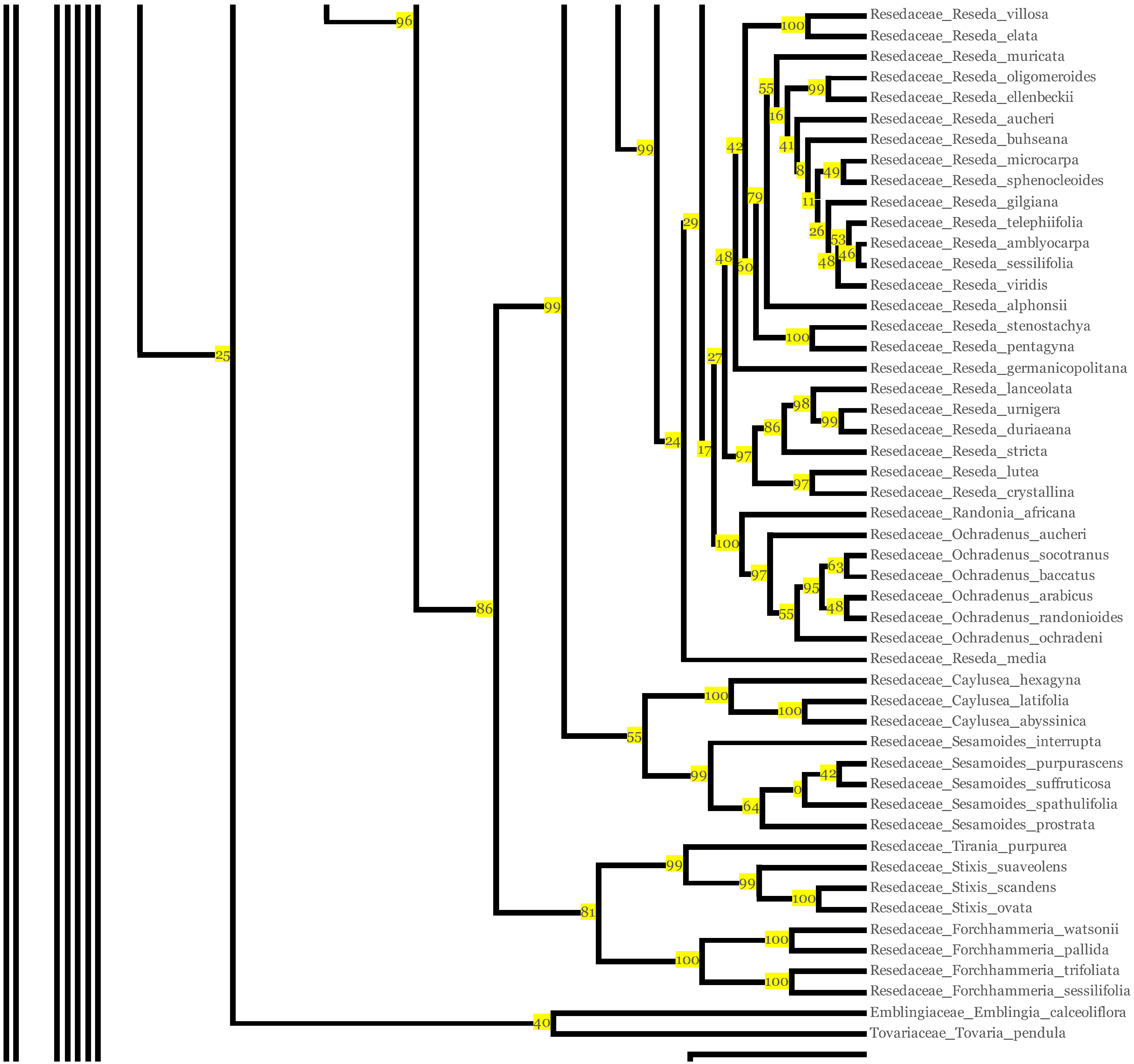
Resedaceae_Reseda_villosa
Resedaceae_Reseda_elata
Resedaceae_Reseda_muricata
Resedaceae_Reseda_oligomeroides
Resedaceae_Reseda_ellenbeckii
Resedaceae_Reseda_aucheri
Resedaceae_Reseda_buhseana
Resedaceae_Reseda_microcarpa
Resedaceae_Reseda_sphenocleoides
Resedaceae_Reseda_gilgiana
Resedaceae_Reseda_telephiifolia
Resedaceae_Reseda_amblyocarpa
Resedaceae_Reseda_sessilifolia
Resedaceae_Reseda_viridis
Resedaceae_Reseda_alphonsii
Resedaceae_Reseda_stenostachya
Resedaceae_Reseda_pentagyna
Resedaceae_Reseda_germanicopolitana
Resedaceae_Reseda_lanceolata
Resedaceae_Reseda_urnigera
Resedaceae_Reseda_duriaeana
Resedaceae_Reseda_stricta
Resedaceae_Reseda_lutea
Resedaceae_Reseda_crystallina
Resedaceae_Randonia_africana
Resedaceae_Ochradenus_aucheri
Resedaceae_Ochradenus_socotranus
Resedaceae_Ochradenus_baccatus
Resedaceae_Ochradenus_arabicus
Resedaceae_Ochradenus_randonioides
Resedaceae_Ochradenus_ochradeni
Resedaceae_Reseda_media
Resedaceae_Caylusea_hexagyna
Resedaceae_Caylusea_latifolia
Resedaceae_Caylusea_abyssinica
Resedaceae_Sesamoides_interrupta
Resedaceae_Sesamoides_purpurascens
Resedaceae_Sesamoides_suffruticosa
Resedaceae_Sesamoides_spathulifolia
Resedaceae_Sesamoides_prostrata
Resedaceae_Tirania_purpurea
Resedaceae_Stixis_suaveolens
Resedaceae_Stixis_scandens
Resedaceae_Stixis_ovata
Resedaceae_Forchhammeria_watsonii
Resedaceae_Forchhammeria_pallida
Resedaceae_Forchhammeria_trifoliata
Resedaceae_Forchhammeria_sessilifolia
Emblingiaceae_Emblingia_calceoliflora
Tovariaceae_Tovaria_pendula

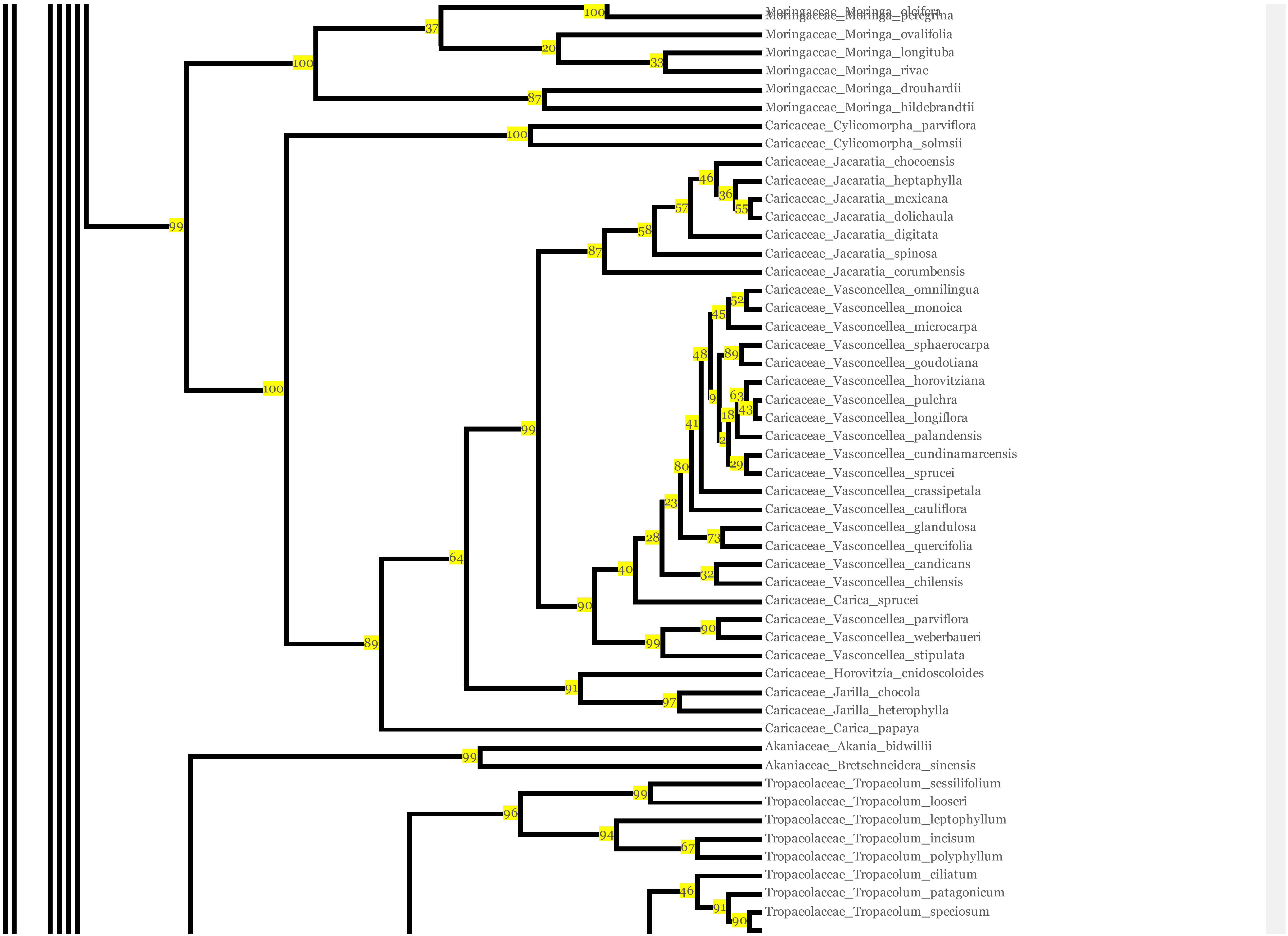
Moringaceae_Moringa_oleifera
Moringaceae_Moringa_peregrina
Moringaceae_Moringa_ovalifolia
Moringaceae_Moringa_longituba
Moringaceae_Moringa_rivae
Moringaceae_Moringa_drouhardii
Moringaceae_Moringa_hildebrandtii
Caricaceae_Cylicomorpha_parviflora
Caricaceae_Cylicomorpha_solmsii
Caricaceae_Jacaratia_chocoensis
Caricaceae_Jacaratia_heptaphylla
Caricaceae_Jacaratia_mexicana
Caricaceae_Jacaratia_dolichaula
Caricaceae_Jacaratia_digitata
Caricaceae_Jacaratia_spinosa
Caricaceae_Jacaratia_corumbensis
Caricaceae_Vasconcellea_omnilingua
Caricaceae_Vasconcellea_monoica
Caricaceae_Vasconcellea_microcarpa
Caricaceae_Vasconcellea_sphaerocarpa
Caricaceae_Vasconcellea_goudotiana
Caricaceae_Vasconcellea_horovitziana
Caricaceae_Vasconcellea_pulchra
Caricaceae_Vasconcellea_longiflora
Caricaceae_Vasconcellea_palandensis
Caricaceae_Vasconcellea_cundinamarcensis
Caricaceae_Vasconcellea_sprucei
Caricaceae_Vasconcellea_crassipetala
Caricaceae_Vasconcellea_cauliflora
Caricaceae_Vasconcellea_glandulosa
Caricaceae_Vasconcellea_quercifolia
Caricaceae_Vasconcellea_candicans
Caricaceae_Vasconcellea_chilensis
Caricaceae_Carica_sprucei
Caricaceae_Vasconcellea_parviflora
Caricaceae_Vasconcellea_weberbaueri
Caricaceae_Vasconcellea_stipulata
Caricaceae_Horovitzia_cnidoscoloides
Caricaceae_Jarilla_chocola
Caricaceae_Jarilla_heterophylla
Caricaceae_Carica_papaya
Akaniaceae_Akania_bidwillii
Akaniaceae_Bretschneidera_sinensis
Tropaeolaceae_Tropaeolum_sessilifolium
Tropaeolaceae_Tropaeolum_looseri
Tropaeolaceae_Tropaeolum_leptophyllum
Tropaeolaceae_Tropaeolum_incisum
Tropaeolaceae_Tropaeolum_polyphyllum
Tropaeolaceae_Tropaeolum_ciliatum
Tropaeolaceae_Tropaeolum_patagonicum
Tropaeolaceae_Tropaeolum_speciosum

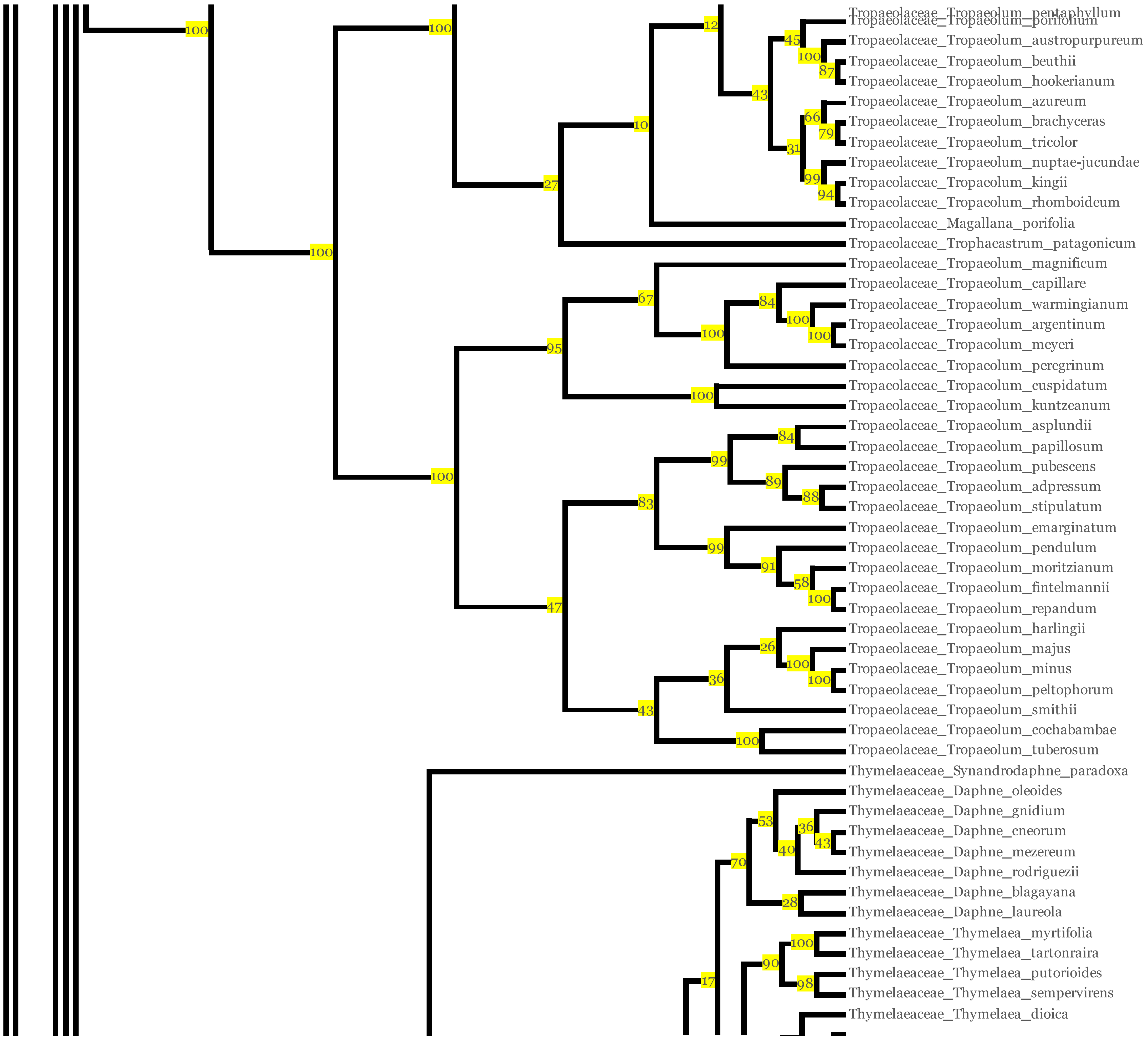

Tropaeolaceae_Tropaeolum_pentaphyllum
Tropaeolaceae_Tropaeolum_porifolium
Tropaeolaceae_Tropaeolum_austropurpureum
Tropaeolaceae_Tropaeolum_beuthii
Tropaeolaceae_Tropaeolum_hookerianum
Tropaeolaceae_Tropaeolum_azureum
Tropaeolaceae_Tropaeolum_brachyceras
Tropaeolaceae_Tropaeolum_tricolor
Tropaeolaceae_Tropaeolum_nuptae-jucundae
Tropaeolaceae_Tropaeolum_kingii
Tropaeolaceae_Tropaeolum_rhomboideum
Tropaeolaceae_Magallana_porifolia
Tropaeolaceae_Trophaeastrum_patagonicum
Tropaeolaceae_Tropaeolum_magnificum
Tropaeolaceae_Tropaeolum_capillare
Tropaeolaceae_Tropaeolum_warmingianum
Tropaeolaceae_Tropaeolum_argentinum
Tropaeolaceae_Tropaeolum_meyeri
Tropaeolaceae_Tropaeolum_peregrinum
Tropaeolaceae_Tropaeolum_cuspidatum
Tropaeolaceae_Tropaeolum_kuntzeanum
Tropaeolaceae_Tropaeolum_asplundii
Tropaeolaceae_Tropaeolum_papillosum
Tropaeolaceae_Tropaeolum_pubescens
Tropaeolaceae_Tropaeolum_adpressum
Tropaeolaceae_Tropaeolum_stipulatum
Tropaeolaceae_Tropaeolum_emarginatum
Tropaeolaceae_Tropaeolum_pendulum
Tropaeolaceae_Tropaeolum_moritzianum
Tropaeolaceae_Tropaeolum_fintelmannii
Tropaeolaceae_Tropaeolum_repandum
Tropaeolaceae_Tropaeolum_harlingii
Tropaeolaceae_Tropaeolum_majus
Tropaeolaceae_Tropaeolum_minus
Tropaeolaceae_Tropaeolum_peltophorum
Tropaeolaceae_Tropaeolum_smithii
Tropaeolaceae_Tropaeolum_cochabambae
Tropaeolaceae_Tropaeolum_tuberosum
Thymelaeaceae_Synandrodaphne_paradoxa
Thymelaeaceae_Daphne_oleoides
Thymelaeaceae_Daphne_gnidium
Thymelaeaceae_Daphne_cneorum
Thymelaeaceae_Daphne_mezereum
Thymelaeaceae_Daphne_rodriguezii
Thymelaeaceae_Daphne_blagayana
Thymelaeaceae_Daphne_laureola
Thymelaeaceae_Thymelaea_myrtifolia
Thymelaeaceae_Thymelaea_tartonraira
Thymelaeaceae_Thymelaea_putorioides
Thymelaeaceae_Thymelaea_sempervirens
Thymelaeaceae_Thymelaea_dioica

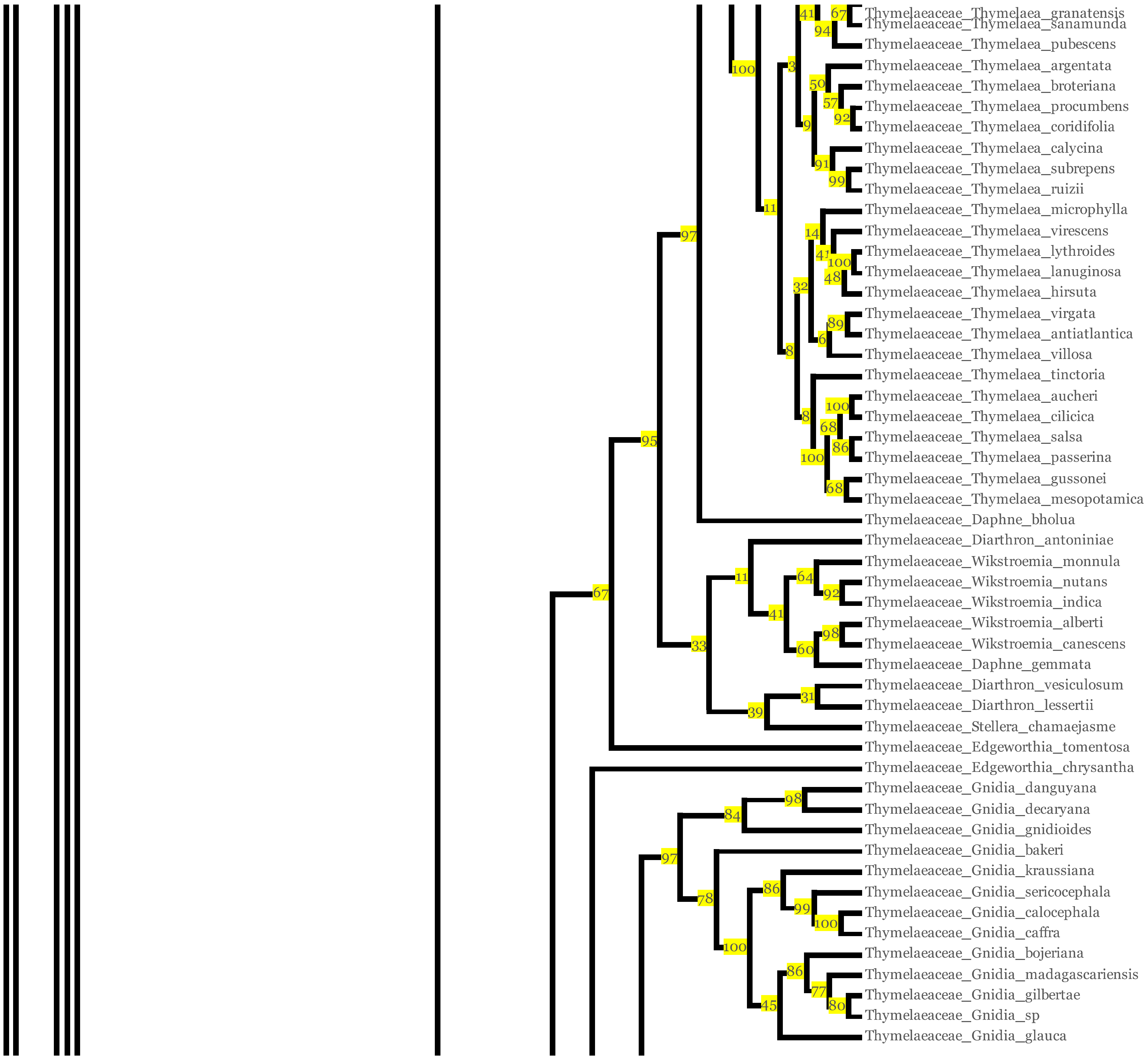
Thymelaeaceae_Thymelaea_granatensis
Thymelaeaceae_Thymelaea_sanamunda
Thymelaeaceae_Thymelaea_pubescens
Thymelaeaceae_Thymelaea_argentata
Thymelaeaceae_Thymelaea_broteriana
Thymelaeaceae_Thymelaea_procumbens
Thymelaeaceae_Thymelaea_coridifolia
Thymelaeaceae_Thymelaea_calycina
Thymelaeaceae_Thymelaea_subrepens
Thymelaeaceae_Thymelaea_ruizii
Thymelaeaceae_Thymelaea_microphylla
Thymelaeaceae_Thymelaea_virescens
Thymelaeaceae_Thymelaea_lythroides
Thymelaeaceae_Thymelaea_lanuginosa
Thymelaeaceae_Thymelaea_hirsuta
Thymelaeaceae_Thymelaea_virgata
Thymelaeaceae_Thymelaea_antiatlantica
Thymelaeaceae_Thymelaea_villosa
Thymelaeaceae_Thymelaea_tinctoria
Thymelaeaceae_Thymelaea_aucheri
Thymelaeaceae_Thymelaea_cilicica
Thymelaeaceae_Thymelaea_salsa
Thymelaeaceae_Thymelaea_passerina
Thymelaeaceae_Thymelaea_gussonei
Thymelaeaceae_Thymelaea_mesopotamica
Thymelaeaceae_Daphne_bholua
Thymelaeaceae_Diarthron_antoniniae
Thymelaeaceae_Wikstroemia_monnula
Thymelaeaceae_Wikstroemia_nutans
Thymelaeaceae_Wikstroemia_indica
Thymelaeaceae_Wikstroemia_alberti
Thymelaeaceae_Wikstroemia_canescens
Thymelaeaceae_Daphne_gemmata
Thymelaeaceae_Diarthron_vesiculosum
Thymelaeaceae_Diarthron_lessertii
Thymelaeaceae_Stellera_chamaejasme
Thymelaeaceae_Edgeworthia_tomentosa
Thymelaeaceae_Edgeworthia_chrysantha
Thymelaeaceae_Gnidia_danguyana
Thymelaeaceae_Gnidia_decaryana
Thymelaeaceae_Gnidia_gnidioides
Thymelaeaceae_Gnidia_bakeri
Thymelaeaceae_Gnidia_kraussiana
Thymelaeaceae_Gnidia_sericocephala
Thymelaeaceae_Gnidia_calocephala
Thymelaeaceae_Gnidia_caffra
Thymelaeaceae_Gnidia_bojeriana
Thymelaeaceae_Gnidia_madagascariensis
Thymelaeaceae_Gnidia_gilbertae
Thymelaeaceae_Gnidia_sp
Thymelaeaceae_Gnidia_glauca

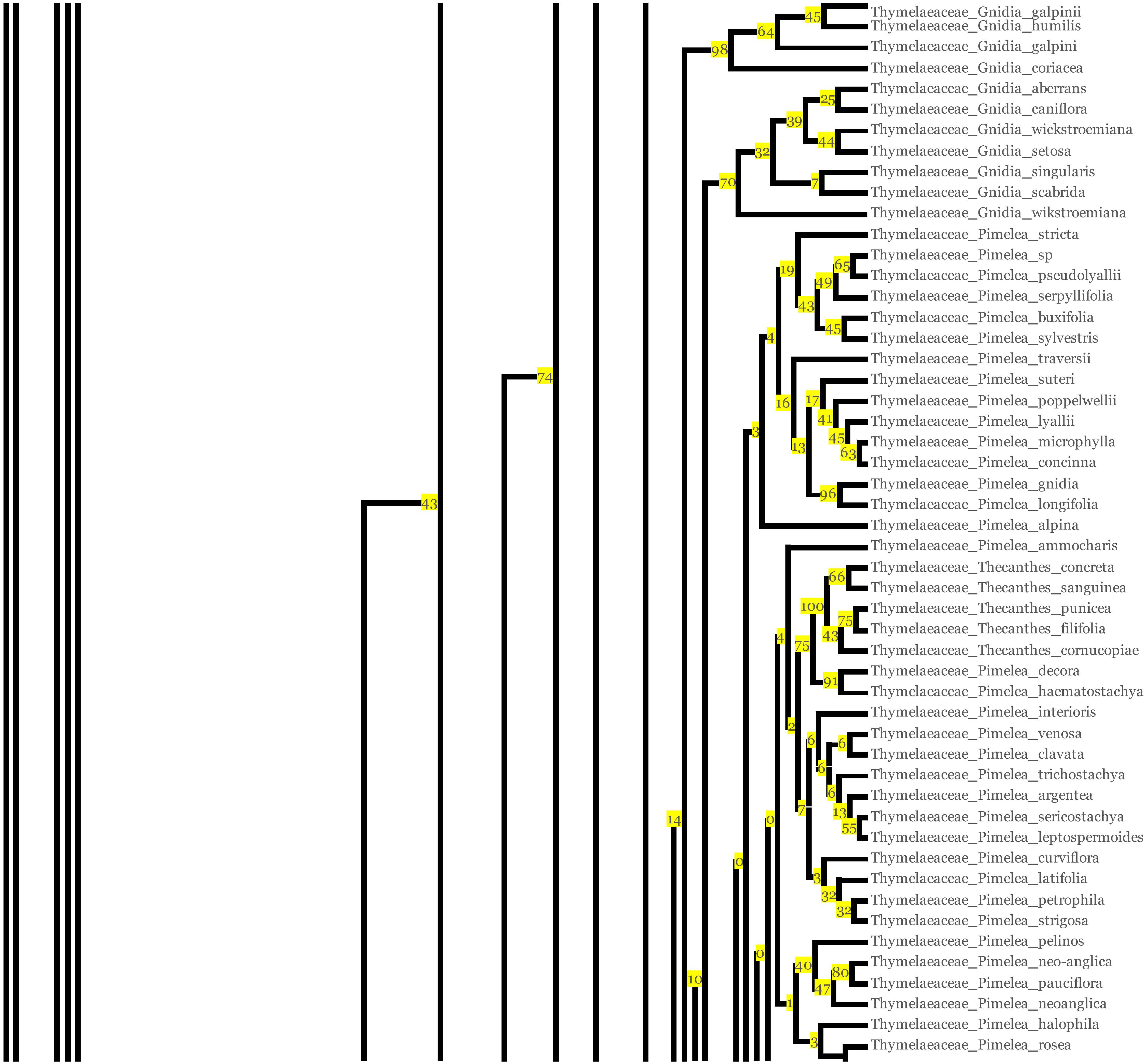
Thymelaeaceae_Gnidia_galpinii
Thymelaeaceae_Gnidia_humilis
Thymelaeaceae_Gnidia_galpini
Thymelaeaceae_Gnidia_coriacea
Thymelaeaceae_Gnidia_aberrans
Thymelaeaceae_Gnidia_caniflora
Thymelaeaceae_Gnidia_wickstroemiana
Thymelaeaceae_Gnidia_setosa
Thymelaeaceae_Gnidia_singularis
Thymelaeaceae_Gnidia_scabrida
Thymelaeaceae_Gnidia_wikstroemiana
Thymelaeaceae_Pimelea_stricta
Thymelaeaceae_Pimelea_sp
Thymelaeaceae_Pimelea_pseudolyallii
Thymelaeaceae_Pimelea_serpyllifolia
Thymelaeaceae_Pimelea_buxifolia
Thymelaeaceae_Pimelea_sylvestris
Thymelaeaceae_Pimelea_traversii
Thymelaeaceae_Pimelea_suteri
Thymelaeaceae_Pimelea_poppelwellii
Thymelaeaceae_Pimelea_lyallii
Thymelaeaceae_Pimelea_microphylla
Thymelaeaceae_Pimelea_concinna
Thymelaeaceae_Pimelea_gnidia
Thymelaeaceae_Pimelea_longifolia
Thymelaeaceae_Pimelea_alpina
Thymelaeaceae_Pimelea_ammocharis
Thymelaeaceae_Thecanthes_concreta
Thymelaeaceae_Thecanthes_sanguinea
Thymelaeaceae_Thecanthes_punicea
Thymelaeaceae_Thecanthes_filifolia
Thymelaeaceae_Thecanthes_cornucopiae
Thymelaeaceae_Pimelea_decora
Thymelaeaceae_Pimelea_haematostachya
Thymelaeaceae_Pimelea_interioris
Thymelaeaceae_Pimelea_venosa
Thymelaeaceae_Pimelea_clavata
Thymelaeaceae_Pimelea_trichostachya
Thymelaeaceae_Pimelea_argentea
Thymelaeaceae_Pimelea_sericostachya
Thymelaeaceae_Pimelea_leptospermoides
Thymelaeaceae_Pimelea_curviflora
Thymelaeaceae_Pimelea_latifolia
Thymelaeaceae_Pimelea_petrophila
Thymelaeaceae_Pimelea_strigosa
Thymelaeaceae_Pimelea_pelinos
Thymelaeaceae_Pimelea_neo-anglica
Thymelaeaceae_Pimelea_pauciflora
Thymelaeaceae_Pimelea_neoanglica
Thymelaeaceae_Pimelea_halophila
Thymelaeaceae_Pimelea_rosea

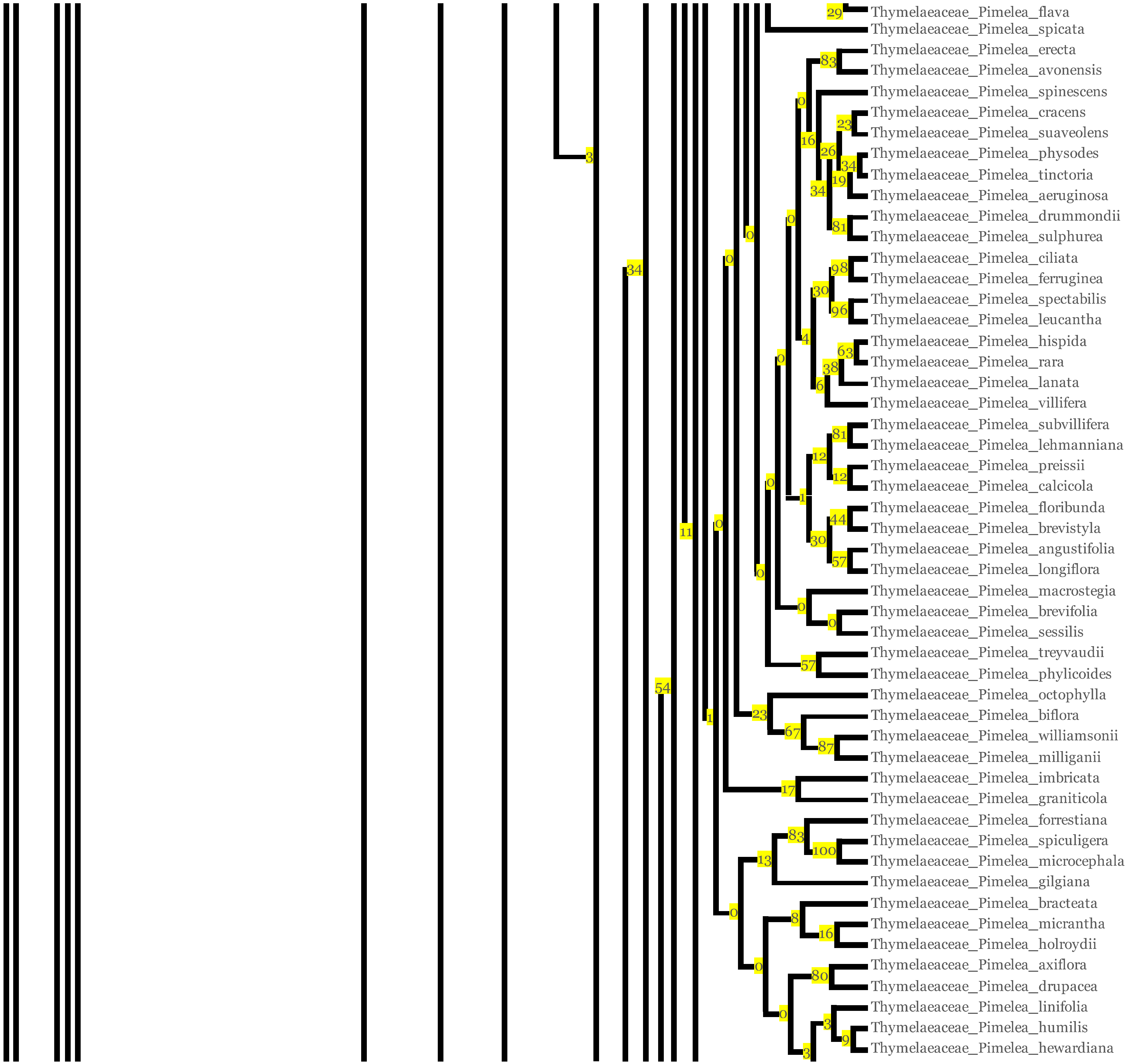
Thymelaeaceae_Pimelea_flava
Thymelaeaceae_Pimelea_spicata
Thymelaeaceae_Pimelea_erecta
Thymelaeaceae_Pimelea_avonensis
Thymelaeaceae_Pimelea_spinescens
Thymelaeaceae_Pimelea_cracens
Thymelaeaceae_Pimelea_suaveolens
Thymelaeaceae_Pimelea_physodes
Thymelaeaceae_Pimelea_tinctoria
Thymelaeaceae_Pimelea_aeruginosa
Thymelaeaceae_Pimelea_drummondii
Thymelaeaceae_Pimelea_sulphurea
Thymelaeaceae_Pimelea_ciliata
Thymelaeaceae_Pimelea_ferruginea
Thymelaeaceae_Pimelea_spectabilis
Thymelaeaceae_Pimelea_leucantha
Thymelaeaceae_Pimelea_hispida
Thymelaeaceae_Pimelea_rara
Thymelaeaceae_Pimelea_lanata
Thymelaeaceae_Pimelea_villifera
Thymelaeaceae_Pimelea_subvillifera
Thymelaeaceae_Pimelea_lehmanniana
Thymelaeaceae_Pimelea_preissii
Thymelaeaceae_Pimelea_calcicola
Thymelaeaceae_Pimelea_floribunda
Thymelaeaceae_Pimelea_brevistyla
Thymelaeaceae_Pimelea_angustifolia
Thymelaeaceae_Pimelea_longiflora
Thymelaeaceae_Pimelea_macrostegia
Thymelaeaceae_Pimelea_brevifolia
Thymelaeaceae_Pimelea_sessilis
Thymelaeaceae_Pimelea_treyvaudii
Thymelaeaceae_Pimelea_phylicoides
Thymelaeaceae_Pimelea_octophylla
Thymelaeaceae_Pimelea_biflora
Thymelaeaceae_Pimelea_williamsonii
Thymelaeaceae_Pimelea_milliganii
Thymelaeaceae_Pimelea_imbricata
Thymelaeaceae_Pimelea_graniticola
Thymelaeaceae_Pimelea_forrestiana
Thymelaeaceae_Pimelea_spiculigera
Thymelaeaceae_Pimelea_microcephala
Thymelaeaceae_Pimelea_gilgiana
Thymelaeaceae_Pimelea_bracteata
Thymelaeaceae_Pimelea_micrantha
Thymelaeaceae_Pimelea_holroydii
Thymelaeaceae_Pimelea_axiflora
Thymelaeaceae_Pimelea_drupacea
Thymelaeaceae_Pimelea_linifolia
Thymelaeaceae_Pimelea_humilis
Thymelaeaceae_Pimelea_hewardiana

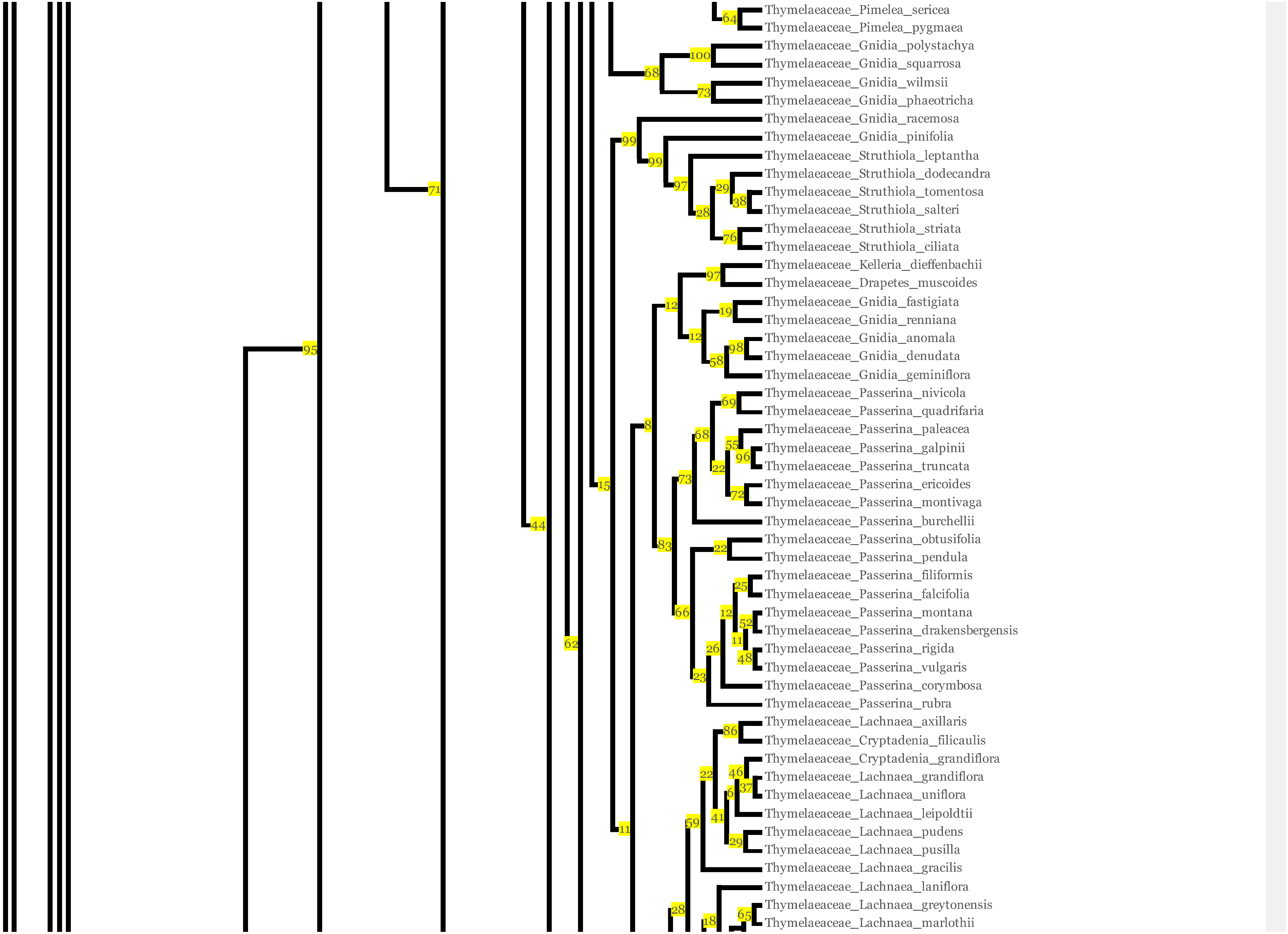
Thymelaeaceae_Pimelea_sericea
Thymelaeaceae_Pimelea_pygmaea
Thymelaeaceae_Gnidia_polystachya
Thymelaeaceae_Gnidia_squarrosa
Thymelaeaceae_Gnidia_wilmsii
Thymelaeaceae_Gnidia_phaeotricha
Thymelaeaceae_Gnidia_racemosa
Thymelaeaceae_Gnidia_pinifolia
Thymelaeaceae_Struthiola_leptantha
Thymelaeaceae_Struthiola_dodecandra
Thymelaeaceae_Struthiola_tomentosa
Thymelaeaceae_Struthiola_salteri
Thymelaeaceae_Struthiola_striata
Thymelaeaceae_Struthiola_ciliata
Thymelaeaceae_Kelleria_dieffenbachii
Thymelaeaceae_Drapetes_muscoides
Thymelaeaceae_Gnidia_fastigiata
Thymelaeaceae_Gnidia_renniana
Thymelaeaceae_Gnidia_anomala
Thymelaeaceae_Gnidia_denudata
Thymelaeaceae_Gnidia_geminiflora
Thymelaeaceae_Passerina_nivicola
Thymelaeaceae_Passerina_quadrifaria
Thymelaeaceae_Passerina_paleacea
Thymelaeaceae_Passerina_galpinii
Thymelaeaceae_Passerina_truncata
Thymelaeaceae_Passerina_ericoides
Thymelaeaceae_Passerina_montivaga
Thymelaeaceae_Passerina_burchellii
Thymelaeaceae_Passerina_obtusifolia
Thymelaeaceae_Passerina_pendula
Thymelaeaceae_Passerina_filiformis
Thymelaeaceae_Passerina_falcifolia
Thymelaeaceae_Passerina_montana
Thymelaeaceae_Passerina_drakensbergensis
Thymelaeaceae_Passerina_rigida
Thymelaeaceae_Passerina_vulgaris
Thymelaeaceae_Passerina_corymbosa
Thymelaeaceae_Passerina_rubra
Thymelaeaceae_Lachnaea_axillaris
Thymelaeaceae_Cryptadenia_filicaulis
Thymelaeaceae_Cryptadenia_grandiflora
Thymelaeaceae_Lachnaea_grandiflora
Thymelaeaceae_Lachnaea_uniflora
Thymelaeaceae_Lachnaea_leipoldtii
Thymelaeaceae_Lachnaea_pudens
Thymelaeaceae_Lachnaea_pusilla
Thymelaeaceae_Lachnaea_gracilis
Thymelaeaceae_Lachnaea_laniflora
Thymelaeaceae_Lachnaea_greytonensis
Thymelaeaceae_Lachnaea_marlothii

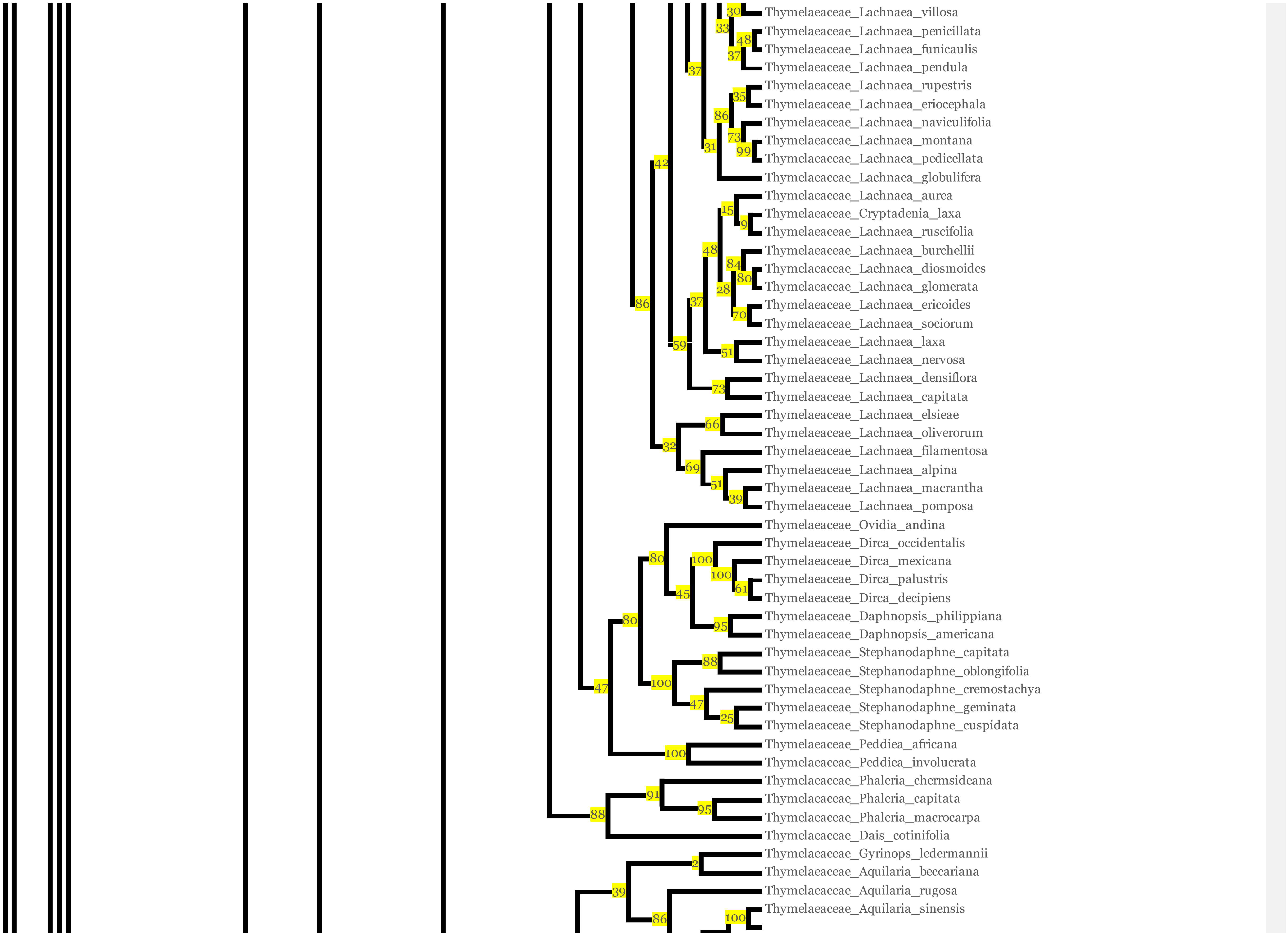
Thymelaeaceae_Lachnaea_villosa
Thymelaeaceae_Lachnaea_penicillata
Thymelaeaceae_Lachnaea_funicaulis
Thymelaeaceae_Lachnaea_pendula
Thymelaeaceae_Lachnaea_rupestris
Thymelaeaceae_Lachnaea_eriocephala
Thymelaeaceae_Lachnaea_naviculifolia
Thymelaeaceae_Lachnaea_montana
Thymelaeaceae_Lachnaea_pedicellata
Thymelaeaceae_Lachnaea_globulifera
Thymelaeaceae_Lachnaea_aurea
Thymelaeaceae_Cryptadenia_laxa
Thymelaeaceae_Lachnaea_ruscifolia
Thymelaeaceae_Lachnaea_burchellii
Thymelaeaceae_Lachnaea_diosmoides
Thymelaeaceae_Lachnaea_glomerata
Thymelaeaceae_Lachnaea_ericoides
Thymelaeaceae_Lachnaea_sociorum
Thymelaeaceae_Lachnaea_laxa
Thymelaeaceae_Lachnaea_nervosa
Thymelaeaceae_Lachnaea_densiflora
Thymelaeaceae_Lachnaea_capitata
Thymelaeaceae_Lachnaea_elsieae
Thymelaeaceae_Lachnaea_oliverorum
Thymelaeaceae_Lachnaea_filamentosa
Thymelaeaceae_Lachnaea_alpina
Thymelaeaceae_Lachnaea_macrantha
Thymelaeaceae_Lachnaea_pomposa
Thymelaeaceae_Ovidia_andina
Thymelaeaceae_Dirca_occidentalis
Thymelaeaceae_Dirca_mexicana
Thymelaeaceae_Dirca_palustris
Thymelaeaceae_Dirca_decipiens
Thymelaeaceae_Daphnopsis_philippiana
Thymelaeaceae_Daphnopsis_americana
Thymelaeaceae_Stephanodaphne_capitata
Thymelaeaceae_Stephanodaphne_oblongifolia
Thymelaeaceae_Stephanodaphne_cremostachya
Thymelaeaceae_Stephanodaphne_geminata
Thymelaeaceae_Stephanodaphne_cuspidata
Thymelaeaceae_Peddiea_africana
Thymelaeaceae_Peddiea_involucrata
Thymelaeaceae_Phaleria_chermsideana
Thymelaeaceae_Phaleria_capitata
Thymelaeaceae_Phaleria_macrocarpa
Thymelaeaceae_Dais_cotinifolia
Thymelaeaceae_Gyrinops_ledermannii
Thymelaeaceae_Aquilaria_beccariana
Thymelaeaceae_Aquilaria_rugosa
Thymelaeaceae_Aquilaria_sinensis

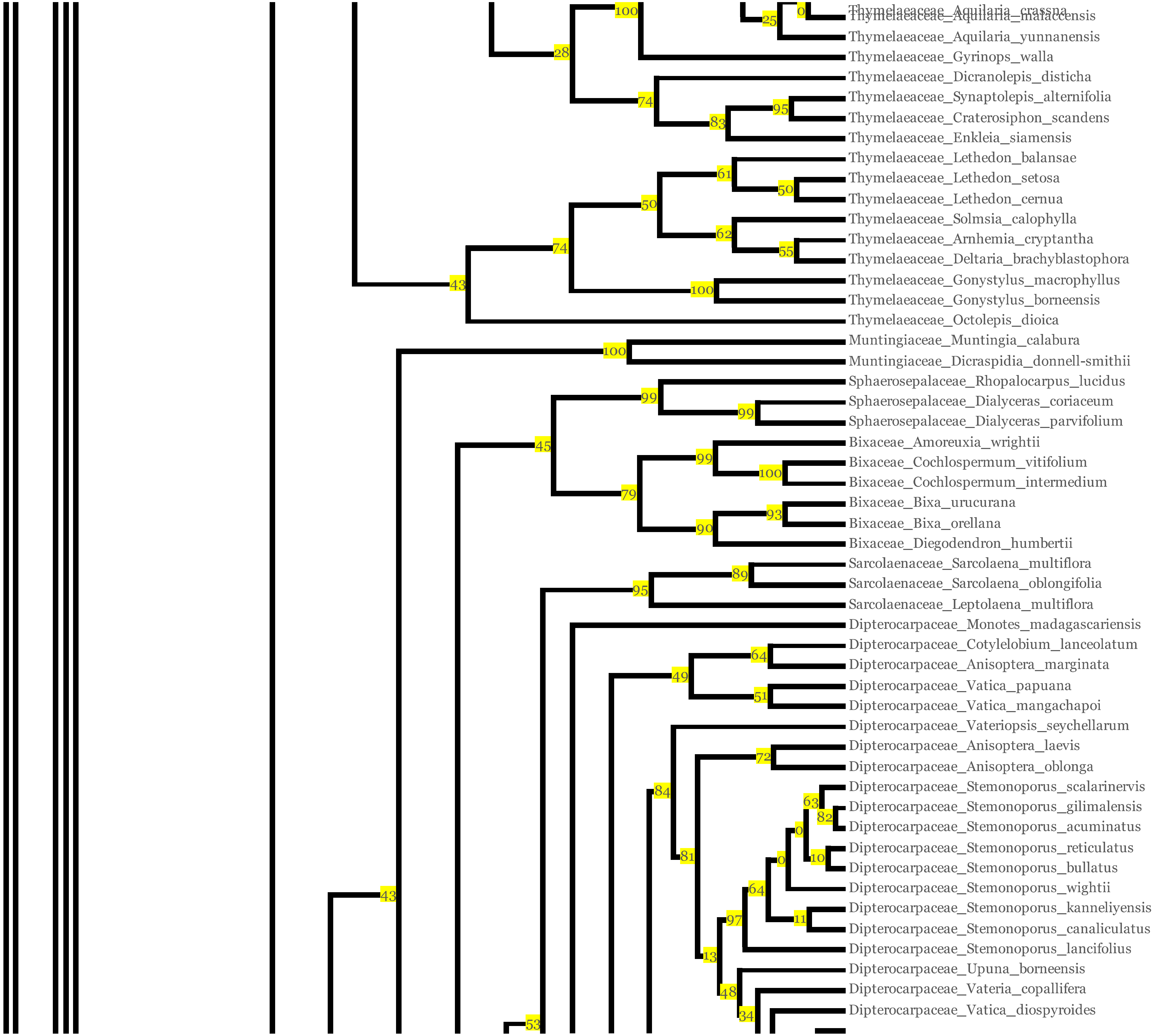
Thymelaeaceae_Aquilaria_crassna
Thymelaeaceae_Aquilaria_malaccensis
Thymelaeaceae_Aquilaria_yunnanensis
Thymelaeaceae_Gyrinops_walla
Thymelaeaceae_Dicranolepis_disticha
Thymelaeaceae_Synaptolepis_alternifolia
Thymelaeaceae_Craterosiphon_scandens
Thymelaeaceae_Enkleia_siamensis
Thymelaeaceae_Lethedon_balansae
Thymelaeaceae_Lethedon_setosa
Thymelaeaceae_Lethedon_cernua
Thymelaeaceae_Solmsia_calophylla
Thymelaeaceae_Arnhemia_cryptantha
Thymelaeaceae_Deltaria_brachyblastophora
Thymelaeaceae_Gonystylus_macrophyllus
Thymelaeaceae_Gonystylus_borneensis
Thymelaeaceae_Octolepis_dioica
Muntingiaceae_Muntingia_calabura
Muntingiaceae_Dicraspidia_donnell-smithii
Sphaerosepalaceae_Rhopalocarpus_lucidus
Sphaerosepalaceae_Dialyceras_coriaceum
Sphaerosepalaceae_Dialyceras_parvifolium
Bixaceae_Amoreuxia_wrightii
Bixaceae_Cochlospermum_vitifolium
Bixaceae_Cochlospermum_intermedium
Bixaceae_Bixa_urucurana
Bixaceae_Bixa_orellana
Bixaceae_Diegodendron_humbertii
Sarcolaenaceae_Sarcolaena_multiflora
Sarcolaenaceae_Sarcolaena_oblongifolia
Sarcolaenaceae_Leptolaena_multiflora
Dipterocarpaceae_Monotes_madagascariensis
Dipterocarpaceae_Cotylelobium_lanceolatum
Dipterocarpaceae_Anisoptera_marginata
Dipterocarpaceae_Vatica_papuana
Dipterocarpaceae_Vatica_mangachapoi
Dipterocarpaceae_Vateriopsis_seychellarum
Dipterocarpaceae_Anisoptera_laevis
Dipterocarpaceae_Anisoptera_oblonga
Dipterocarpaceae_Stemonoporus_scalarinervis
Dipterocarpaceae_Stemonoporus_gilimalensis
Dipterocarpaceae_Stemonoporus_acuminatus
Dipterocarpaceae_Stemonoporus_reticulatus
Dipterocarpaceae_Stemonoporus_bullatus
Dipterocarpaceae_Stemonoporus_wightii
Dipterocarpaceae_Stemonoporus_kanneliyensis
Dipterocarpaceae_Stemonoporus_canaliculatus
Dipterocarpaceae_Stemonoporus_lancifolius
Dipterocarpaceae_Upuna_borneensis
Dipterocarpaceae_Vateria_copallifera
Dipterocarpaceae_Vatica_diospyroides

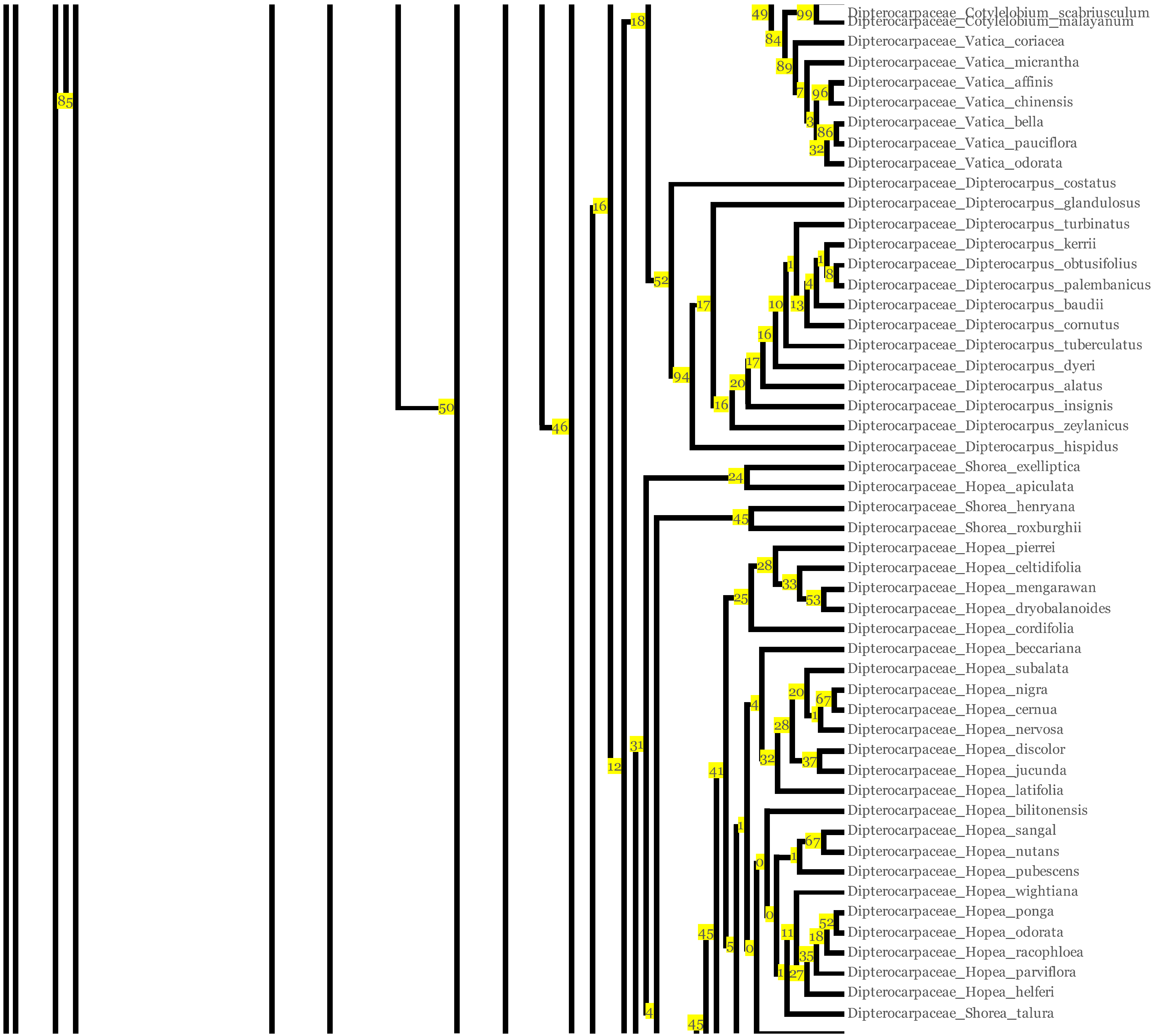
Dipterocarpaceae_Cotylelobium_scabriusculum
Dipterocarpaceae_Cotylelobium_malayanum
Dipterocarpaceae_Vatica_coriacea
Dipterocarpaceae_Vatica_micrantha
Dipterocarpaceae_Vatica_affinis
Dipterocarpaceae_Vatica_chinensis
Dipterocarpaceae_Vatica_bella
Dipterocarpaceae_Vatica_pauciflora
Dipterocarpaceae_Vatica_odorata
Dipterocarpaceae_Dipterocarpus_costatus
Dipterocarpaceae_Dipterocarpus_glandulosus
Dipterocarpaceae_Dipterocarpus_turbinatus
Dipterocarpaceae_Dipterocarpus_kerrii
Dipterocarpaceae_Dipterocarpus_obtusifolius
Dipterocarpaceae_Dipterocarpus_palembanicus
Dipterocarpaceae_Dipterocarpus_baudii
Dipterocarpaceae_Dipterocarpus_cornutus
Dipterocarpaceae_Dipterocarpus_tuberculatus
Dipterocarpaceae_Dipterocarpus_dyeri
Dipterocarpaceae_Dipterocarpus_alatus
Dipterocarpaceae_Dipterocarpus_insignis
Dipterocarpaceae_Dipterocarpus_zeylanicus
Dipterocarpaceae_Dipterocarpus_hispidus
Dipterocarpaceae_Shorea_exelliptica
Dipterocarpaceae_Hopea_apiculata
Dipterocarpaceae_Shorea_henryana
Dipterocarpaceae_Shorea_roxburghii
Dipterocarpaceae_Hopea_pierrei
Dipterocarpaceae_Hopea_celtidifolia
Dipterocarpaceae_Hopea_mengarawan
Dipterocarpaceae_Hopea_dryobalanoides
Dipterocarpaceae_Hopea_cordifolia
Dipterocarpaceae_Hopea_beccariana
Dipterocarpaceae_Hopea_subalata
Dipterocarpaceae_Hopea_nigra
Dipterocarpaceae_Hopea_cernua
Dipterocarpaceae_Hopea_nervosa
Dipterocarpaceae_Hopea_discolor
Dipterocarpaceae_Hopea_jucunda
Dipterocarpaceae_Hopea_latifolia
Dipterocarpaceae_Hopea_bilitonensis
Dipterocarpaceae_Hopea_sangal
Dipterocarpaceae_Hopea_nutans
Dipterocarpaceae_Hopea_pubescens
Dipterocarpaceae_Hopea_wightiana
Dipterocarpaceae_Hopea_ponga
Dipterocarpaceae_Hopea_odorata
Dipterocarpaceae_Hopea_racophloea
Dipterocarpaceae_Hopea_parviflora
Dipterocarpaceae_Hopea_helferi
Dipterocarpaceae_Shorea_talura

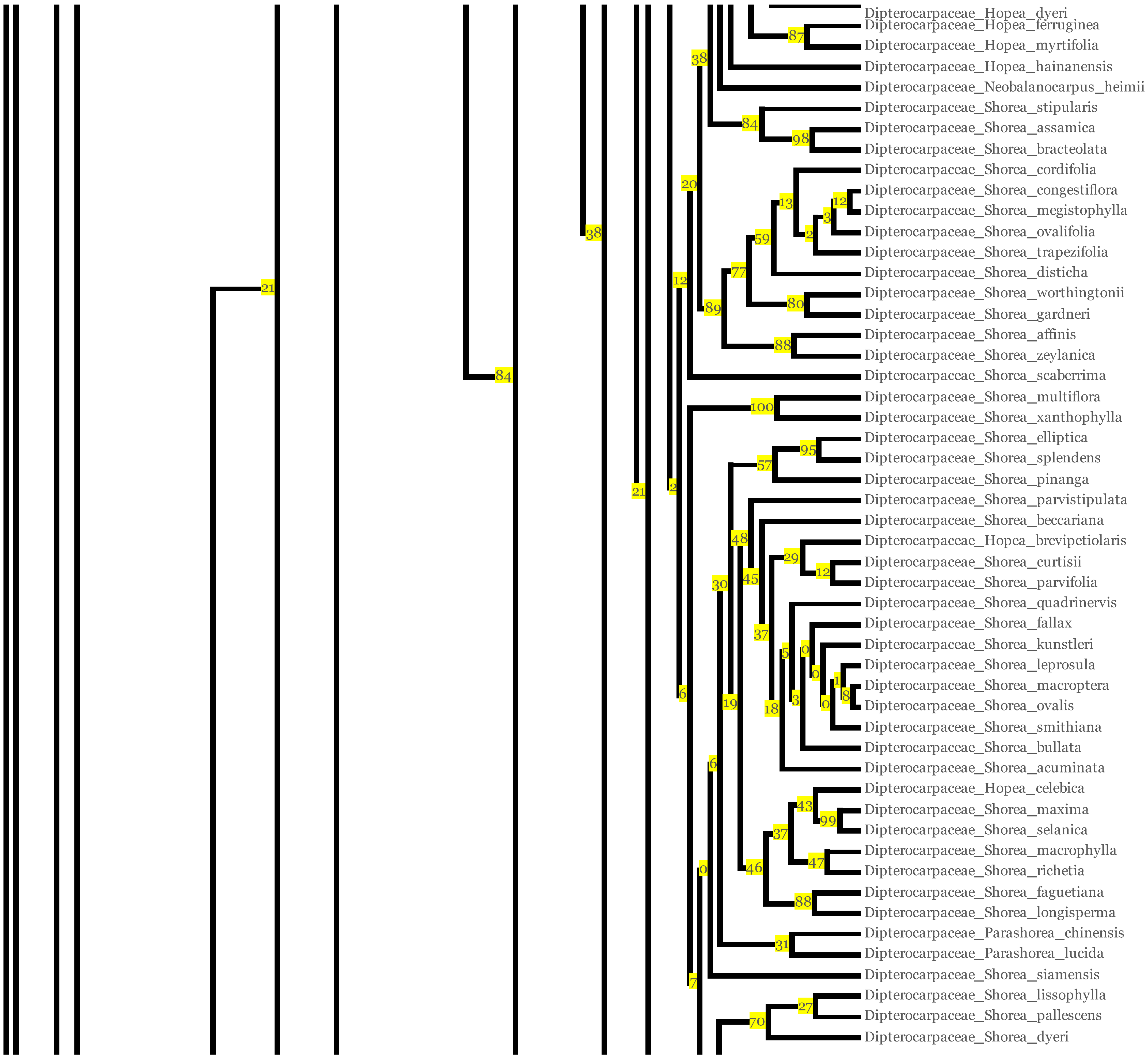
Dipterocarpaceae_Hopea_dyeri
Dipterocarpaceae_Hopea_ferruginea
Dipterocarpaceae_Hopea_myrtifolia
Dipterocarpaceae_Hopea_hainanensis
Dipterocarpaceae_Neobalanocarpus_heimii
Dipterocarpaceae_Shorea_stipularis
Dipterocarpaceae_Shorea_assamica
Dipterocarpaceae_Shorea_bracteolata
Dipterocarpaceae_Shorea_cordifolia
Dipterocarpaceae_Shorea_congestiflora
Dipterocarpaceae_Shorea_megistophylla
Dipterocarpaceae_Shorea_ovalifolia
Dipterocarpaceae_Shorea_trapezifolia
Dipterocarpaceae_Shorea_disticha
Dipterocarpaceae_Shorea_worthingtonii
Dipterocarpaceae_Shorea_gardneri
Dipterocarpaceae_Shorea_affinis
Dipterocarpaceae_Shorea_zeylanica
Dipterocarpaceae_Shorea_scaberrima
Dipterocarpaceae_Shorea_multiflora
Dipterocarpaceae_Shorea_xanthophylla
Dipterocarpaceae_Shorea_elliptica
Dipterocarpaceae_Shorea_splendens
Dipterocarpaceae_Shorea_pinanga
Dipterocarpaceae_Shorea_parvistipulata
Dipterocarpaceae_Shorea_beccariana
Dipterocarpaceae_Hopea_brevipetiolaris
Dipterocarpaceae_Shorea_curtisii
Dipterocarpaceae_Shorea_parvifolia
Dipterocarpaceae_Shorea_quadrinervis
Dipterocarpaceae_Shorea_fallax
Dipterocarpaceae_Shorea_kunstleri
Dipterocarpaceae_Shorea_leprosula
Dipterocarpaceae_Shorea_macroptera
Dipterocarpaceae_Shorea_ovalis
Dipterocarpaceae_Shorea_smithiana
Dipterocarpaceae_Shorea_bullata
Dipterocarpaceae_Shorea_acuminata
Dipterocarpaceae_Hopea_celebica
Dipterocarpaceae_Shorea_maxima
Dipterocarpaceae_Shorea_selanica
Dipterocarpaceae_Shorea_macrophylla
Dipterocarpaceae_Shorea_richetia
Dipterocarpaceae_Shorea_faguetiana
Dipterocarpaceae_Shorea_longisperma
Dipterocarpaceae_Parashorea_chinensis
Dipterocarpaceae_Parashorea_lucida
Dipterocarpaceae_Shorea_siamensis
Dipterocarpaceae_Shorea_lissophylla
Dipterocarpaceae_Shorea_pallescens
Dipterocarpaceae_Shorea_dyeri

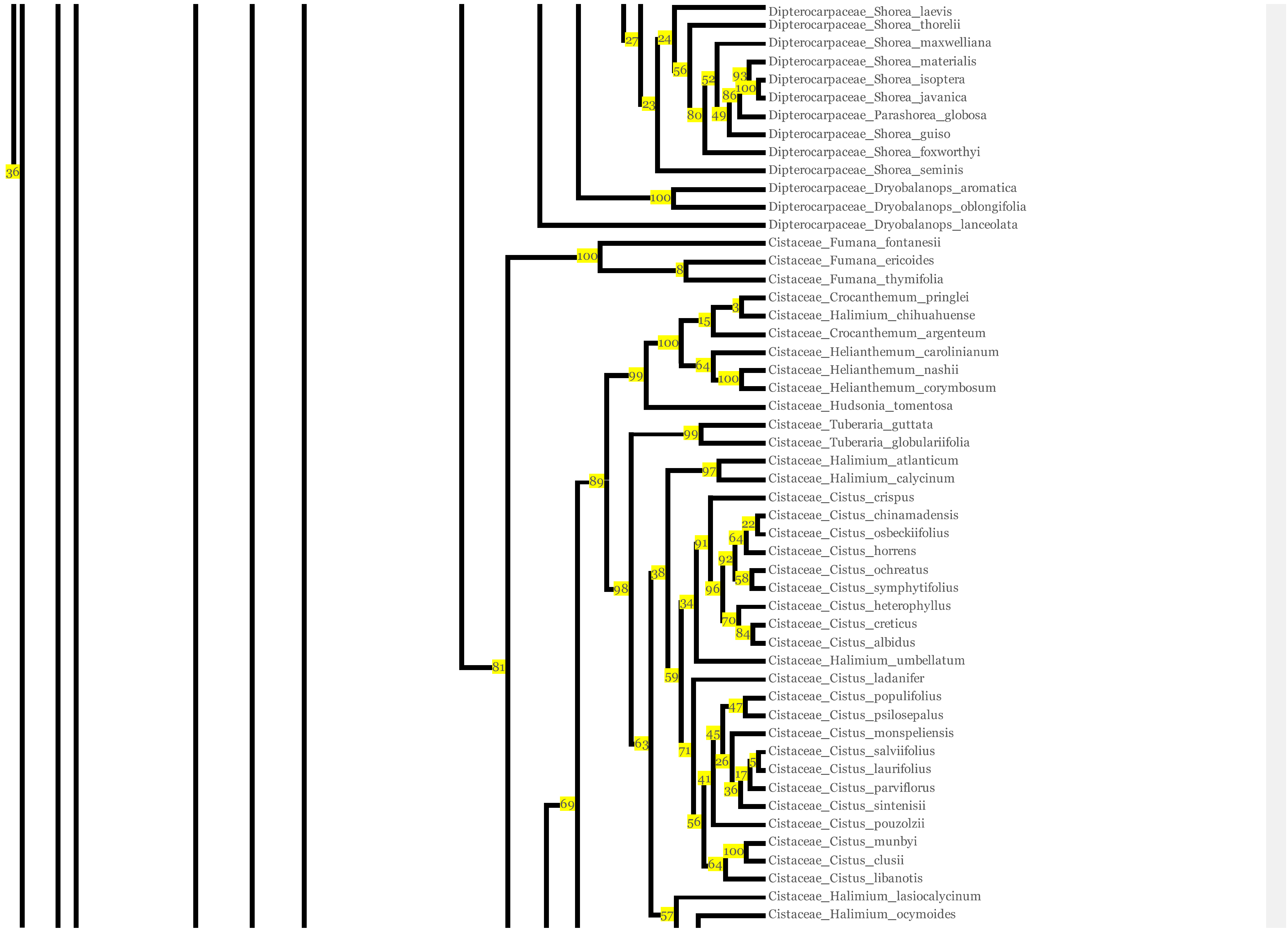
Dipterocarpaceae_Shorea_laevis
Dipterocarpaceae_Shorea_thorelii
Dipterocarpaceae_Shorea_maxwelliana
Dipterocarpaceae_Shorea_materialis
Dipterocarpaceae_Shorea_isoptera
Dipterocarpaceae_Shorea_javanica
Dipterocarpaceae_Parashorea_globosa
Dipterocarpaceae_Shorea_guiso
Dipterocarpaceae_Shorea_foxworthyi
Dipterocarpaceae_Shorea_seminis
Dipterocarpaceae_Dryobalanops_aromatica
Dipterocarpaceae_Dryobalanops_oblongifolia
Dipterocarpaceae_Dryobalanops_lanceolata
Cistaceae_Fumana_fontanesii
Cistaceae_Fumana_ericoides
Cistaceae_Fumana_thymifolia
Cistaceae_Crocanthemum_pringlei
Cistaceae_Halimium_chihuahuense
Cistaceae_Crocanthemum_argenteum
Cistaceae_Helianthemum_carolinianum
Cistaceae_Helianthemum_nashii
Cistaceae_Helianthemum_corymbosum
Cistaceae_Hudsonia_tomentosa
Cistaceae_Tuberaria_guttata
Cistaceae_Tuberaria_globulariifolia
Cistaceae_Halimium_atlanticum
Cistaceae_Halimium_calycinum
Cistaceae_Cistus_crispus
Cistaceae_Cistus_chinamadensis
Cistaceae_Cistus_osbeckiifolius
Cistaceae_Cistus_horrens
Cistaceae_Cistus_ochreatus
Cistaceae_Cistus_symphytifolius
Cistaceae_Cistus_heterophyllus
Cistaceae_Cistus_creticus
Cistaceae_Cistus_albidus
Cistaceae_Halimium_umbellatum
Cistaceae_Cistus_ladanifer
Cistaceae_Cistus_populifolius
Cistaceae_Cistus_psilosepalus
Cistaceae_Cistus_monspeliensis
Cistaceae_Cistus_salviifolius
Cistaceae_Cistus_laurifolius
Cistaceae_Cistus_parviflorus
Cistaceae_Cistus_sintenisii
Cistaceae_Cistus_pouzolzii
Cistaceae_Cistus_munbyi
Cistaceae_Cistus_clusii
Cistaceae_Cistus_libanotis
Cistaceae_Halimium_lasiocalycinum
Cistaceae_Halimium_ocymoides

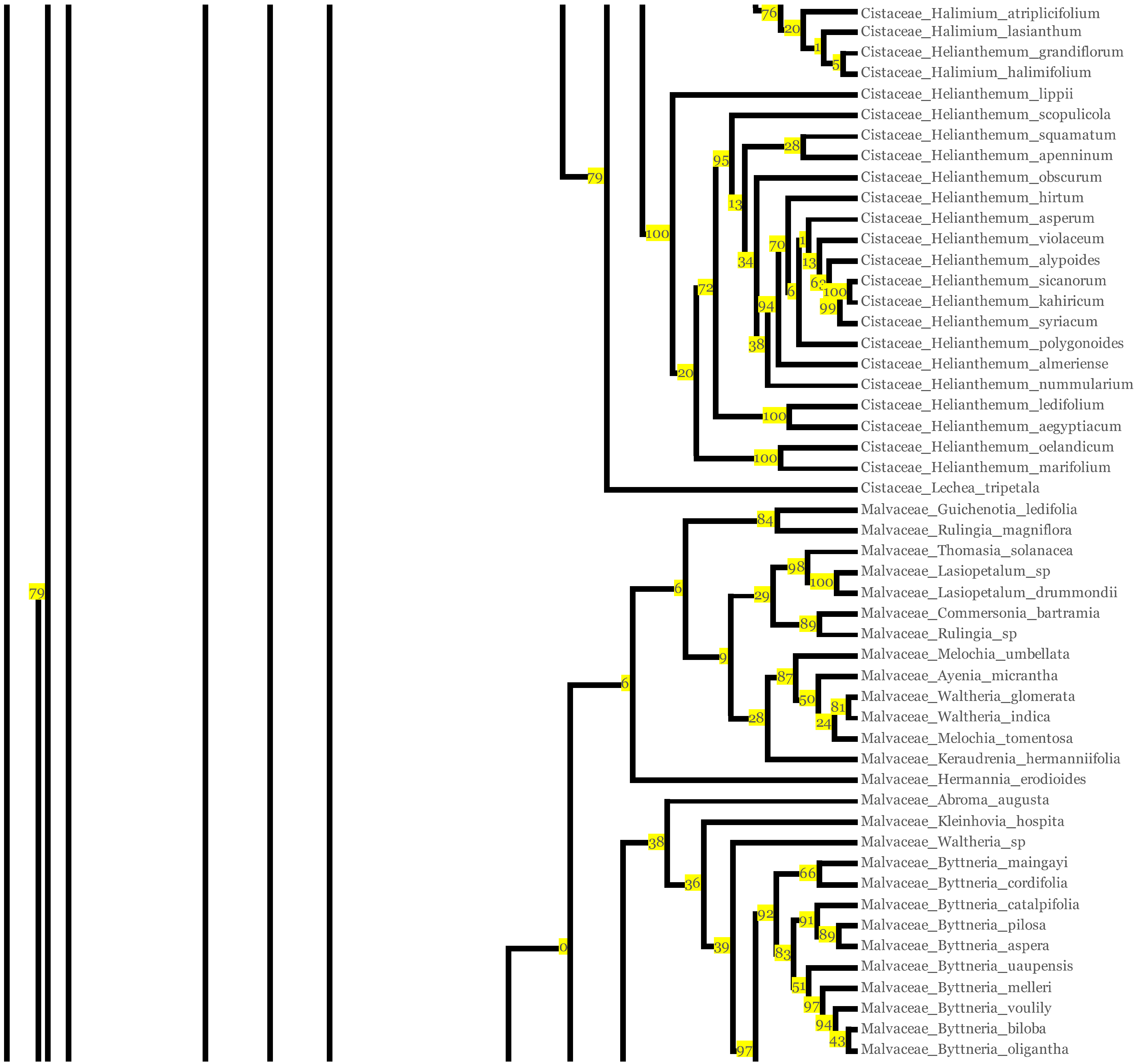
Cistaceae_Halimium_atriplicifolium
Cistaceae_Halimium_lasianthum
Cistaceae_Helianthemum_grandiflorum
Cistaceae_Halimium_halimifolium
Cistaceae_Helianthemum_lippii
Cistaceae_Helianthemum_scopulicola
Cistaceae_Helianthemum_squamatum
Cistaceae_Helianthemum_apenninum
Cistaceae_Helianthemum_obscurum
Cistaceae_Helianthemum_hirtum
Cistaceae_Helianthemum_asperum
Cistaceae_Helianthemum_violaceum
Cistaceae_Helianthemum_alypoides
Cistaceae_Helianthemum_sicanorum
Cistaceae_Helianthemum_kahiricum
Cistaceae_Helianthemum_syriacum
Cistaceae_Helianthemum_polygonoides
Cistaceae_Helianthemum_almeriense
Cistaceae_Helianthemum_nummularium
Cistaceae_Helianthemum_ledifolium
Cistaceae_Helianthemum_aegyptiacum
Cistaceae_Helianthemum_oelandicum
Cistaceae_Helianthemum_marifolium
Cistaceae_Lechea_tripetala
Malvaceae_Guichenotia_ledifolia
Malvaceae_Rulingia_magniflora
Malvaceae_Thomasia_solanacea
Malvaceae_Lasiopetalum_sp
Malvaceae_Lasiopetalum_drummondii
Malvaceae_Commersonia_bartramia
Malvaceae_Rulingia_sp
Malvaceae_Melochia_umbellata
Malvaceae_Ayenia_micrantha
Malvaceae_Waltheria_glomerata
Malvaceae_Waltheria_indica
Malvaceae_Melochia_tomentosa
Malvaceae_Keraudrenia_hermanniifolia
Malvaceae_Hermannia_erodioides
Malvaceae_Abroma_augusta
Malvaceae_Kleinhovia_hospita
Malvaceae_Waltheria_sp
Malvaceae_Byttneria_maingayi
Malvaceae_Byttneria_cordifolia
Malvaceae_Byttneria_catalpifolia
Malvaceae_Byttneria_pilosa
Malvaceae_Byttneria_aspera
Malvaceae_Byttneria_uaupensis
Malvaceae_Byttneria_melleri
Malvaceae_Byttneria_voulily
Malvaceae_Byttneria_biloba
Malvaceae_Byttneria_oligantha

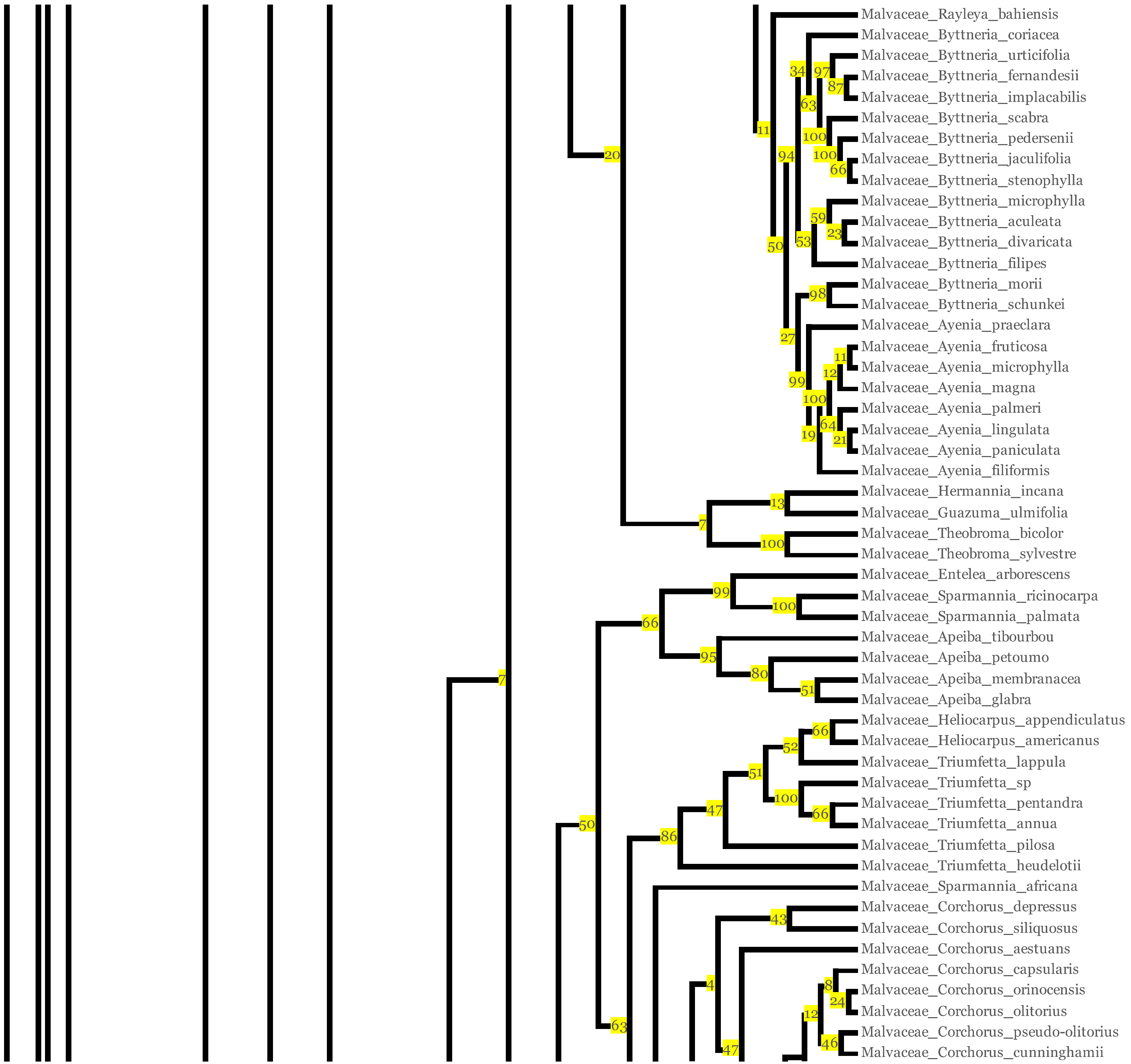
Malvaceae_Rayleya_bahiensis
Malvaceae_Byttneria_coriacea
Malvaceae_Byttneria_urticifolia
Malvaceae_Byttneria_fernandesii
Malvaceae_Byttneria_implacabilis
Malvaceae_Byttneria_scabra
Malvaceae_Byttneria_pedersenii
Malvaceae_Byttneria_jaculifolia
Malvaceae_Byttneria_stenophylla
Malvaceae_Byttneria_microphylla
Malvaceae_Byttneria_aculeata
Malvaceae_Byttneria_divaricata
Malvaceae_Byttneria_filipes
Malvaceae_Byttneria_morii
Malvaceae_Byttneria_schunkei
Malvaceae_Ayenia_praeclara
Malvaceae_Ayenia_fruticosa
Malvaceae_Ayenia_microphylla
Malvaceae_Ayenia_magna
Malvaceae_Ayenia_palmeri
Malvaceae_Ayenia_lingulata
Malvaceae_Ayenia_paniculata
Malvaceae_Ayenia_filiformis
Malvaceae_Hermannia_incana
Malvaceae_Guazuma_ulmifolia
Malvaceae_Theobroma_bicolor
Malvaceae_Theobroma_sylvestre
Malvaceae_Entelea_arborescens
Malvaceae_Sparmannia_ricinocarpa
Malvaceae_Sparmannia_palmata
Malvaceae_Apeiba_tibourbou
Malvaceae_Apeiba_petoumo
Malvaceae_Apeiba_membranacea
Malvaceae_Apeiba_glabra
Malvaceae_Heliocarpus_appendiculatus
Malvaceae_Heliocarpus_americanus
Malvaceae_Triumfetta_lappula
Malvaceae_Triumfetta_sp
Malvaceae_Triumfetta_pentandra
Malvaceae_Triumfetta_annua
Malvaceae_Triumfetta_pilosa
Malvaceae_Triumfetta_heudelotii
Malvaceae_Sparmannia_africana
Malvaceae_Corchorus_depressus
Malvaceae_Corchorus_siliquosus
Malvaceae_Corchorus_aestuans
Malvaceae_Corchorus_capsularis
Malvaceae_Corchorus_orinocensis
Malvaceae_Corchorus_olitorius
Malvaceae_Corchorus_pseudo-olitorius
Malvaceae_Corchorus_cunninghamii

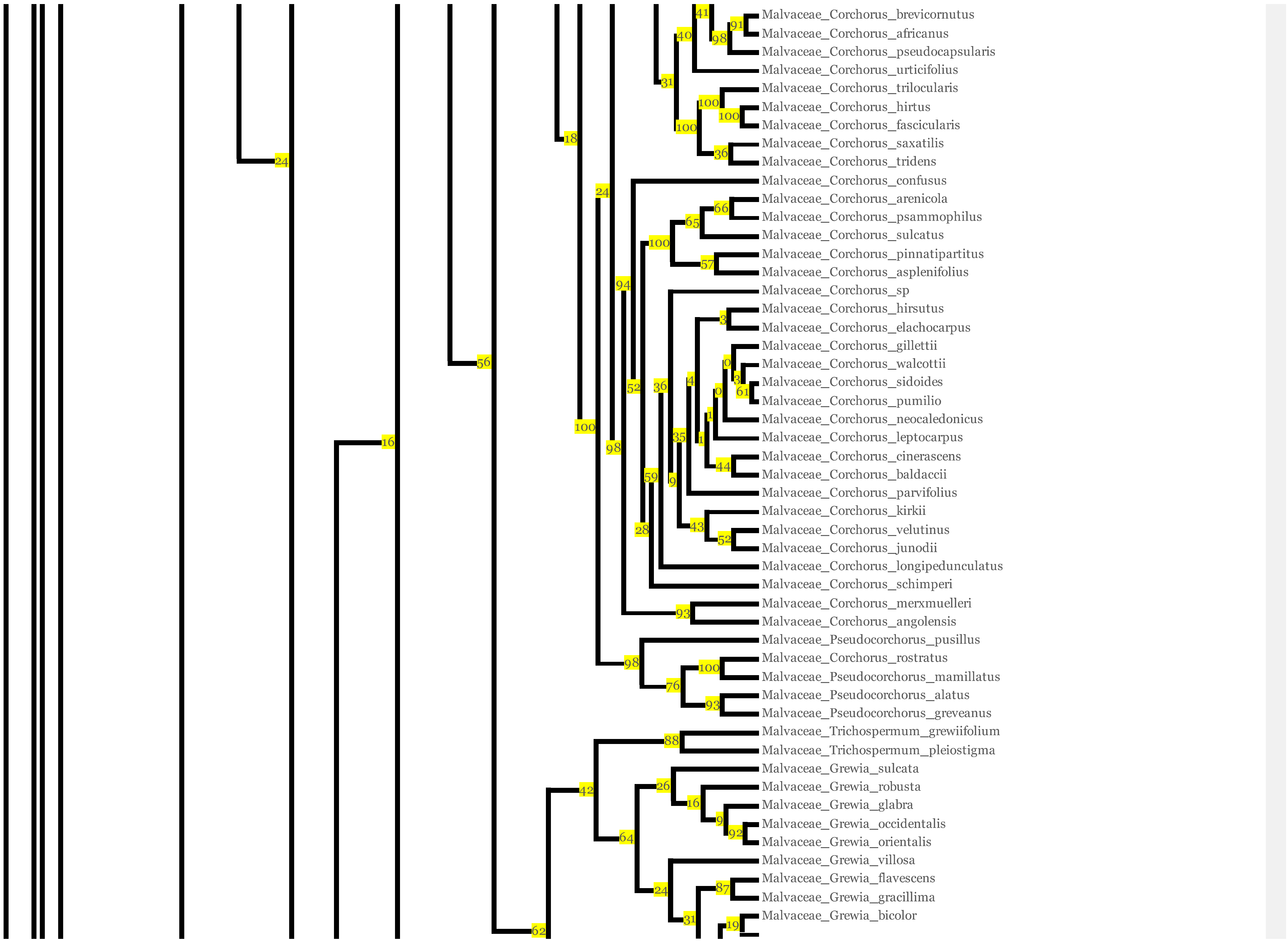
Malvaceae_Corchorus_brevicornutus
Malvaceae_Corchorus_africanus
Malvaceae_Corchorus_pseudocapsularis
Malvaceae_Corchorus_urticifolius
Malvaceae_Corchorus_trilocularis
Malvaceae_Corchorus_hirtus
Malvaceae_Corchorus_fascicularis
Malvaceae_Corchorus_saxatilis
Malvaceae_Corchorus_tridens
Malvaceae_Corchorus_confusus
Malvaceae_Corchorus_arenicola
Malvaceae_Corchorus_psammophilus
Malvaceae_Corchorus_sulcatus
Malvaceae_Corchorus_pinnatipartitus
Malvaceae_Corchorus_asplenifolius
Malvaceae_Corchorus_sp
Malvaceae_Corchorus_hirsutus
Malvaceae_Corchorus_elachocarpus
Malvaceae_Corchorus_gillettii
Malvaceae_Corchorus_walcottii
Malvaceae_Corchorus_sidoides
Malvaceae_Corchorus_pumilio
Malvaceae_Corchorus_neocaledonicus
Malvaceae_Corchorus_leptocarpus
Malvaceae_Corchorus_cinerascens
Malvaceae_Corchorus_baldaccii
Malvaceae_Corchorus_parvifolius
Malvaceae_Corchorus_kirkii
Malvaceae_Corchorus_velutinus
Malvaceae_Corchorus_junodii
Malvaceae_Corchorus_longipedunculatus
Malvaceae_Corchorus_schimperi
Malvaceae_Corchorus_merxmuelleri
Malvaceae_Corchorus_angolensis
Malvaceae_Pseudocorchorus_pusillus
Malvaceae_Corchorus_rostratus
Malvaceae_Pseudocorchorus_mamillatus
Malvaceae_Pseudocorchorus_alatus
Malvaceae_Pseudocorchorus_greveanus
Malvaceae_Trichospermum_grewiifolium
Malvaceae_Trichospermum_pleiostigma
Malvaceae_Grewia_sulcata
Malvaceae_Grewia_robusta
Malvaceae_Grewia_glabra
Malvaceae_Grewia_occidentalis
Malvaceae_Grewia_orientalis
Malvaceae_Grewia_villosa
Malvaceae_Grewia_flavescens
Malvaceae_Grewia_gracillima
Malvaceae_Grewia_bicolor

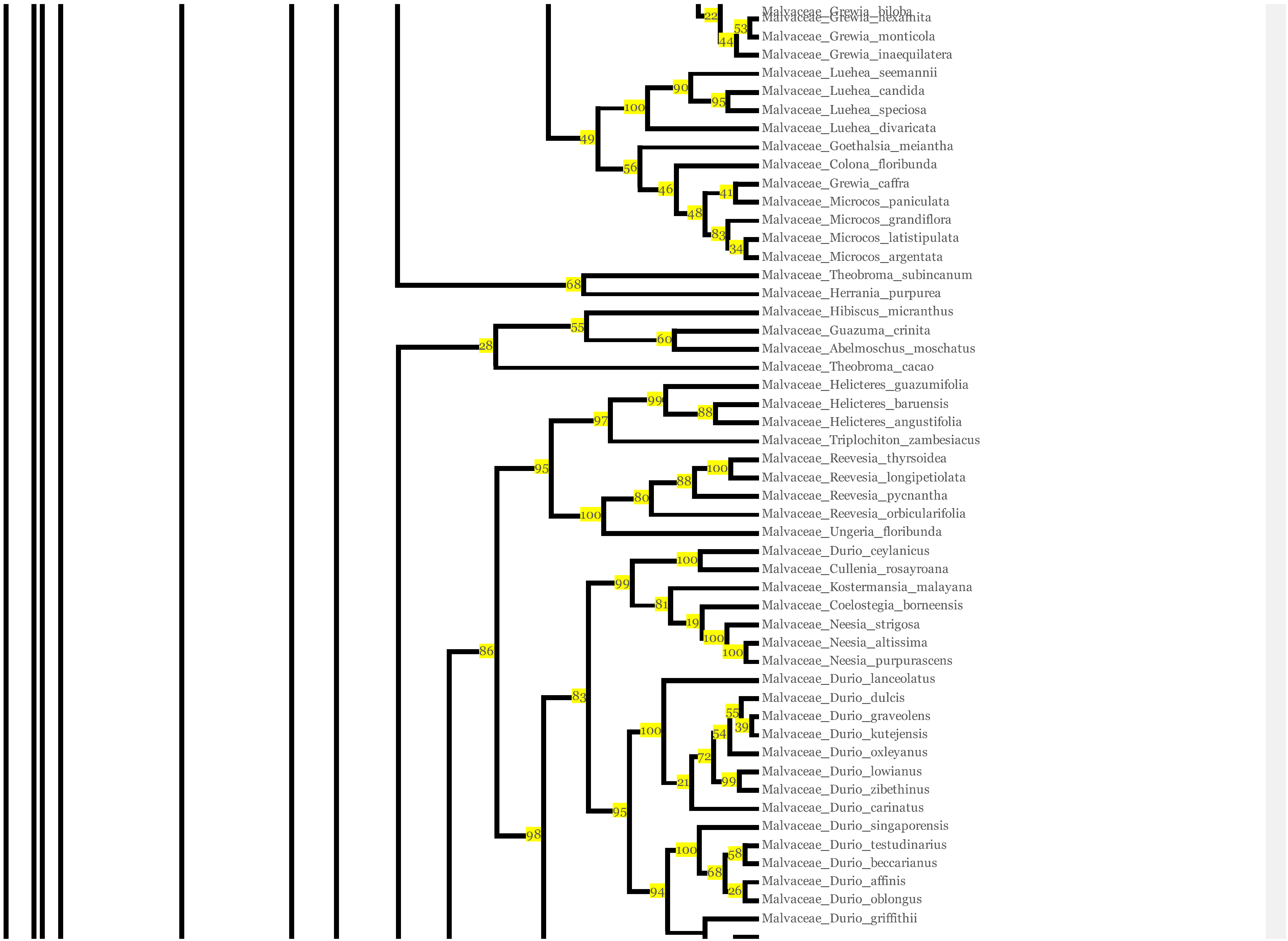
Malvaceae_Grewia_biloba
Malvaceae_Grewia_hexamita
Malvaceae_Grewia_monticola
Malvaceae_Grewia_inaequilatera
Malvaceae_Luehea_seemannii
Malvaceae_Luehea_candida
Malvaceae_Luehea_speciosa
Malvaceae_Luehea_divaricata
Malvaceae_Goethalsia_meiantha
Malvaceae_Colona_floribunda
Malvaceae_Grewia_caffra
Malvaceae_Microcos_paniculata
Malvaceae_Microcos_grandiflora
Malvaceae_Microcos_latistipulata
Malvaceae_Microcos_argentata
Malvaceae_Theobroma_subincanum
Malvaceae_Herrania_purpurea
Malvaceae_Hibiscus_micranthus
Malvaceae_Guazuma_crinita
Malvaceae_Abelmoschus_moschatus
Malvaceae_Theobroma_cacao
Malvaceae_Helicteres_guazumifolia
Malvaceae_Helicteres_baruensis
Malvaceae_Helicteres_angustifolia
Malvaceae_Triplochiton_zambesiacus
Malvaceae_Reevesia_thyrsoidea
Malvaceae_Reevesia_longipetiolata
Malvaceae_Reevesia_pycnantha
Malvaceae_Reevesia_orbicularifolia
Malvaceae_Ungeria_floribunda
Malvaceae_Durio_ceylanicus
Malvaceae_Cullenia_rosayroana
Malvaceae_Kostermansia_malayana
Malvaceae_Coelostegia_borneensis
Malvaceae_Neesia_strigosa
Malvaceae_Neesia_altissima
Malvaceae_Neesia_purpurascens
Malvaceae_Durio_lanceolatus
Malvaceae_Durio_dulcis
Malvaceae_Durio_graveolens
Malvaceae_Durio_kutejensis
Malvaceae_Durio_oxleyanus
Malvaceae_Durio_lowianus
Malvaceae_Durio_zibethinus
Malvaceae_Durio_carinatus
Malvaceae_Durio_singaporensis
Malvaceae_Durio_testudinarius
Malvaceae_Durio_beccarianus
Malvaceae_Durio_affinis
Malvaceae_Durio_oblongus
Malvaceae_Durio_griffithii

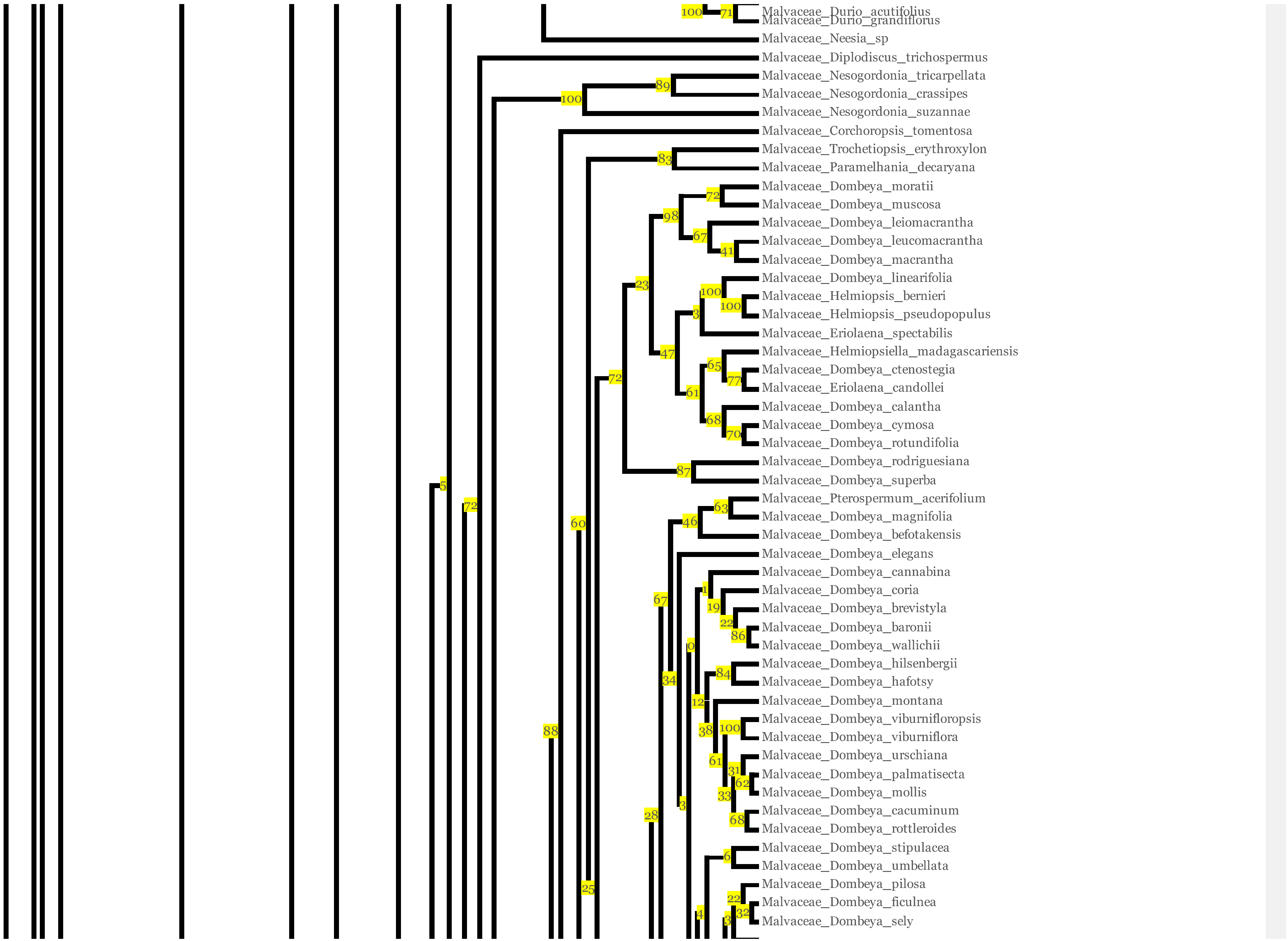
Malvaceae_Durio_acutifolius
Malvaceae_Durio_grandiflorus
Malvaceae_Neesia_sp
Malvaceae_Diplodiscus_trichospermus
Malvaceae_Nesogordonia_tricarpellata
Malvaceae_Nesogordonia_crassipes
Malvaceae_Nesogordonia_suzannae
Malvaceae_Corchoropsis_tomentosa
Malvaceae_Trochetiopsis_erythroxylon
Malvaceae_Paramelhania_decaryana
Malvaceae_Dombeya_moratii
Malvaceae_Dombeya_muscosa
Malvaceae_Dombeya_leiomacrantha
Malvaceae_Dombeya_leucomacrantha
Malvaceae_Dombeya_macrantha
Malvaceae_Dombeya_linearifolia
Malvaceae_Helmiopsis_bernieri
Malvaceae_Helmiopsis_pseudopopulus
Malvaceae_Eriolaena_spectabilis
Malvaceae_Helmiopsiella_madagascariensis
Malvaceae_Dombeya_ctenostegia
Malvaceae_Eriolaena_candollei
Malvaceae_Dombeya_calantha
Malvaceae_Dombeya_cymosa
Malvaceae_Dombeya_rotundifolia
Malvaceae_Dombeya_rodriguesiana
Malvaceae_Dombeya_superba
Malvaceae_Pterospermum_acerifolium
Malvaceae_Dombeya_magnifolia
Malvaceae_Dombeya_befotakensis
Malvaceae_Dombeya_elegans
Malvaceae_Dombeya_cannabina
Malvaceae_Dombeya_coria
Malvaceae_Dombeya_brevistyla
Malvaceae_Dombeya_baronii
Malvaceae_Dombeya_wallichii
Malvaceae_Dombeya_hilsenbergii
Malvaceae_Dombeya_hafotsy
Malvaceae_Dombeya_montana
Malvaceae_Dombeya_viburnifloropsis
Malvaceae_Dombeya_viburniflora
Malvaceae_Dombeya_urschiana
Malvaceae_Dombeya_palmatisecta
Malvaceae_Dombeya_mollis
Malvaceae_Dombeya_cacuminum
Malvaceae_Dombeya_rottleroides
Malvaceae_Dombeya_stipulacea
Malvaceae_Dombeya_umbellata
Malvaceae_Dombeya_pilosa
Malvaceae_Dombeya_ficulnea
Malvaceae_Dombeya_sely

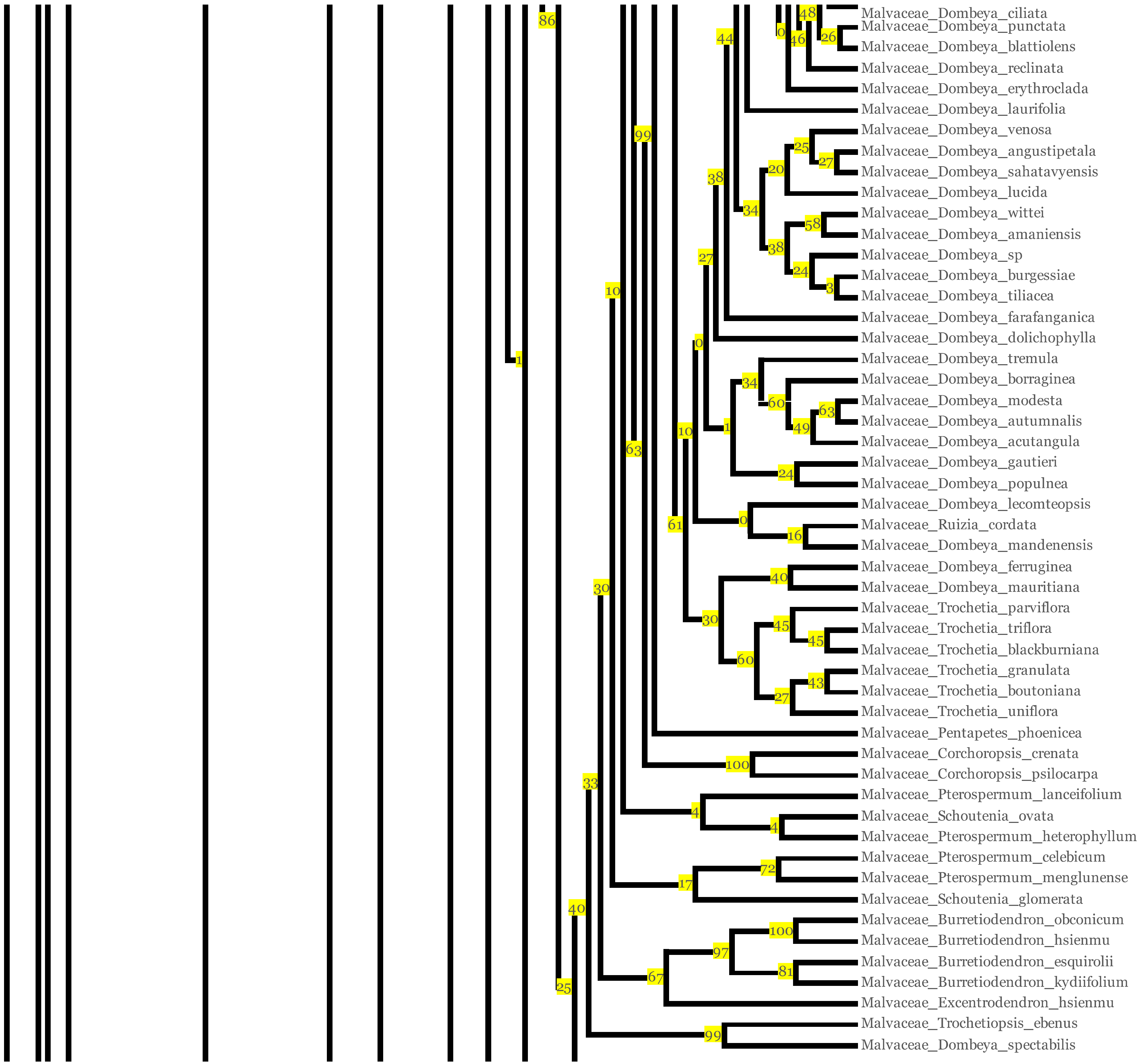
Malvaceae_Dombeya_ciliata
Malvaceae_Dombeya_punctata
Malvaceae_Dombeya_blattiolens
Malvaceae_Dombeya_reclinata
Malvaceae_Dombeya_erythroclada
Malvaceae_Dombeya_laurifolia
Malvaceae_Dombeya_venosa
Malvaceae_Dombeya_angustipetala
Malvaceae_Dombeya_sahatavyensis
Malvaceae_Dombeya_lucida
Malvaceae_Dombeya_wittei
Malvaceae_Dombeya_amaniensis
Malvaceae_Dombeya_sp
Malvaceae_Dombeya_burgessiae
Malvaceae_Dombeya_tiliacea
Malvaceae_Dombeya_farafanganica
Malvaceae_Dombeya_dolichophylla
Malvaceae_Dombeya_tremula
Malvaceae_Dombeya_borraginea
Malvaceae_Dombeya_modesta
Malvaceae_Dombeya_autumnalis
Malvaceae_Dombeya_acutangula
Malvaceae_Dombeya_gautieri
Malvaceae_Dombeya_populnea
Malvaceae_Dombeya_lecomteopsis
Malvaceae_Ruizia_cordata
Malvaceae_Dombeya_mandenensis
Malvaceae_Dombeya_ferruginea
Malvaceae_Dombeya_mauritiana
Malvaceae_Trochetia_parviflora
Malvaceae_Trochetia_triflora
Malvaceae_Trochetia_blackburniana
Malvaceae_Trochetia_granulata
Malvaceae_Trochetia_boutoniana
Malvaceae_Trochetia_uniflora
Malvaceae_Pentapetes_phoenicea
Malvaceae_Corchoropsis_crenata
Malvaceae_Corchoropsis_psilocarpa
Malvaceae_Pterospermum_lanceifolium
Malvaceae_Schoutenia_ovata
Malvaceae_Pterospermum_heterophyllum
Malvaceae_Pterospermum_celebicum
Malvaceae_Pterospermum_menglunense
Malvaceae_Schoutenia_glomerata
Malvaceae_Burretiodendron_obconicum
Malvaceae_Burretiodendron_hsienmu
Malvaceae_Burretiodendron_esquirolii
Malvaceae_Burretiodendron_kydiifolium
Malvaceae_Excentrodendron_hsienmu
Malvaceae_Trochetiopsis_ebenus
Malvaceae_Dombeya_spectabilis

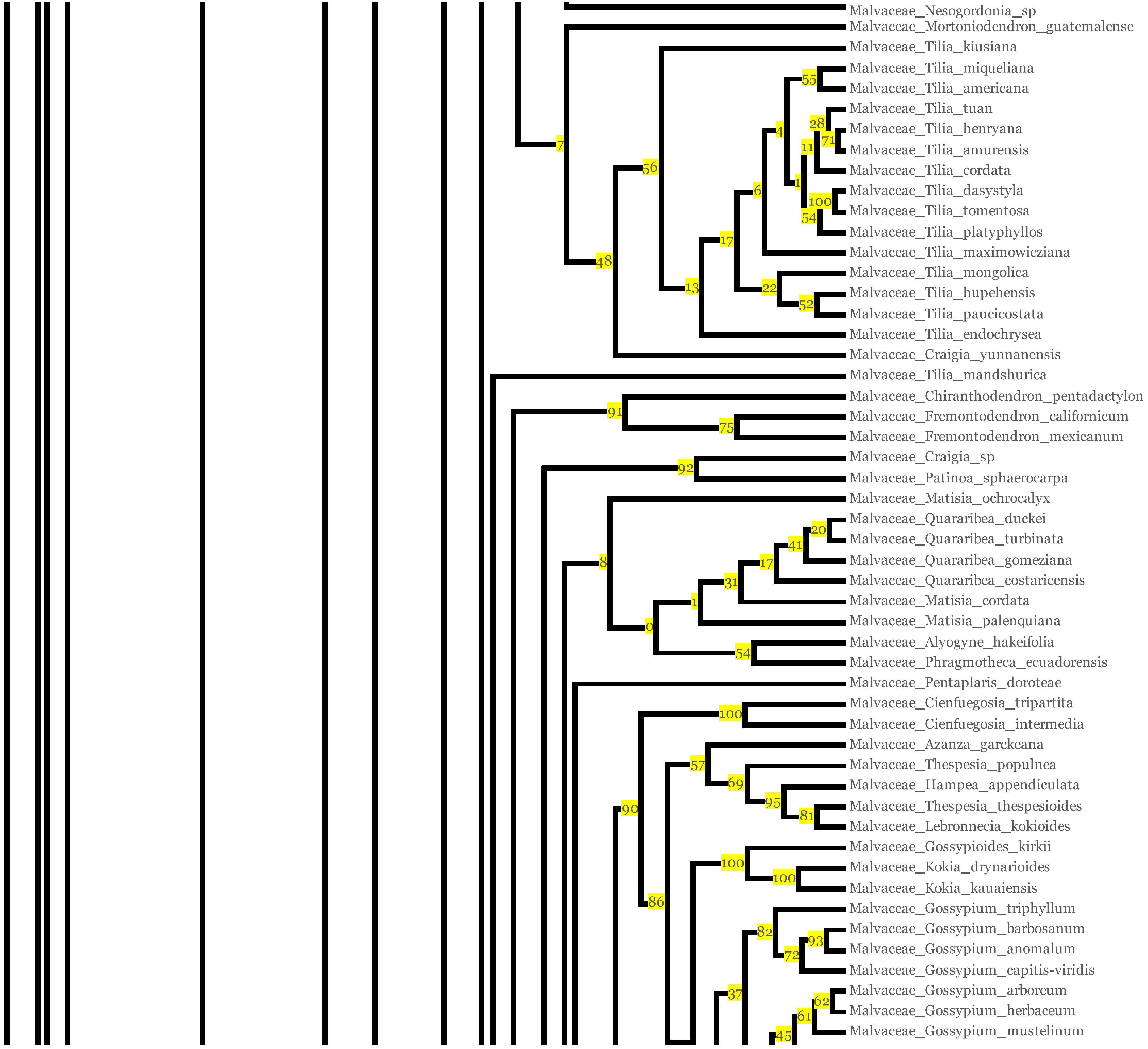
Malvaceae_Nesogordonia_sp
Malvaceae_Mortoniodendron_guatemalense
Malvaceae_Tilia_kiusiana
Malvaceae_Tilia_miqueliana
Malvaceae_Tilia_americana
Malvaceae_Tilia_tuan
Malvaceae_Tilia_henryana
Malvaceae_Tilia_amurensis
Malvaceae_Tilia_cordata
Malvaceae_Tilia_dasystyla
Malvaceae_Tilia_tomentosa
Malvaceae_Tilia_platyphyllos
Malvaceae_Tilia_maximowicziana
Malvaceae_Tilia_mongolica
Malvaceae_Tilia_hupehensis
Malvaceae_Tilia_paucicostata
Malvaceae_Tilia_endochrysea
Malvaceae_Craigia_yunnanensis
Malvaceae_Tilia_mandshurica
Malvaceae_Chiranthodendron_pentadactylon
Malvaceae_Fremontodendron_californicum
Malvaceae_Fremontodendron_mexicanum
Malvaceae_Craigia_sp
Malvaceae_Patinoa_sphaerocarpa
Malvaceae_Matisia_ochrocalyx
Malvaceae_Quararibea_duckei
Malvaceae_Quararibea_turbinata
Malvaceae_Quararibea_gomeziana
Malvaceae_Quararibea_costaricensis
Malvaceae_Matisia_cordata
Malvaceae_Matisia_palenquiana
Malvaceae_Alyogyne_hakeifolia
Malvaceae_Phragmotheca_ecuadorensis
Malvaceae_Pentaplaris_doroteae
Malvaceae_Cienfuegosia_tripartita
Malvaceae_Cienfuegosia_intermedia
Malvaceae_Azanza_garckeana
Malvaceae_Thespesia_populnea
Malvaceae_Hampea_appendiculata
Malvaceae_Thespesia_thespesioides
Malvaceae_Lebronnecia_kokioides
Malvaceae_Gossypioides_kirkii
Malvaceae_Kokia_drynarioides
Malvaceae_Kokia_kauaiensis
Malvaceae_Gossypium_triphyllum
Malvaceae_Gossypium_barbosanum
Malvaceae_Gossypium_anomalum
Malvaceae_Gossypium_capitis-viridis
Malvaceae_Gossypium_arboreum
Malvaceae_Gossypium_herbaceum
Malvaceae_Gossypium_mustelinum

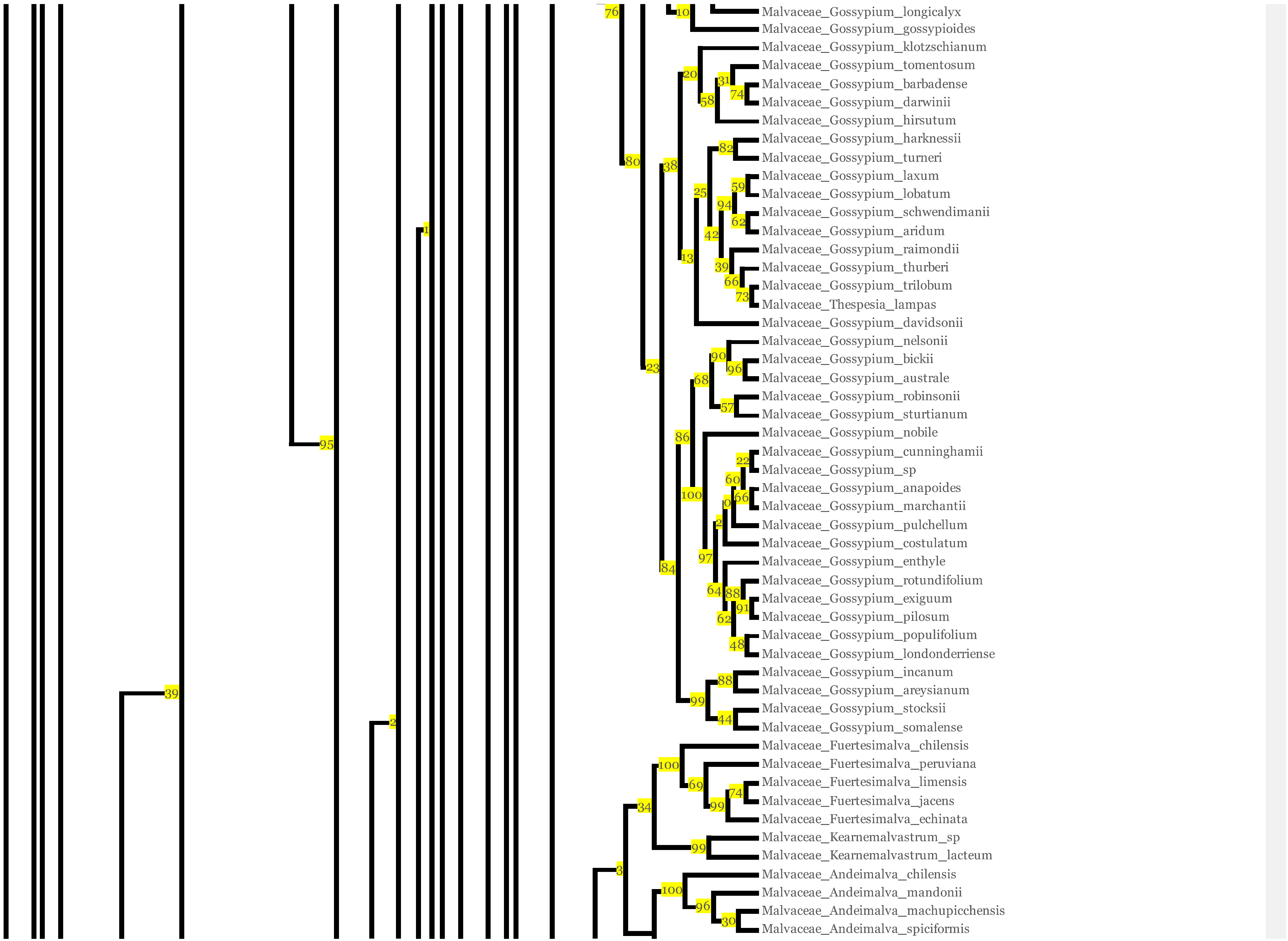
Malvaceae_Gossypium_longicalyx
Malvaceae_Gossypium_gossypioides
Malvaceae_Gossypium_klotzschianum
Malvaceae_Gossypium_tomentosum
Malvaceae_Gossypium_barbadense
Malvaceae_Gossypium_darwinii
Malvaceae_Gossypium_hirsutum
Malvaceae_Gossypium_harknessii
Malvaceae_Gossypium_turneri
Malvaceae_Gossypium_laxum
Malvaceae_Gossypium_lobatum
Malvaceae_Gossypium_schwendimanii
Malvaceae_Gossypium_aridum
Malvaceae_Gossypium_raimondii
Malvaceae_Gossypium_thurberi
Malvaceae_Gossypium_trilobum
Malvaceae_Thespesia_lampas
Malvaceae_Gossypium_davidsonii
Malvaceae_Gossypium_nelsonii
Malvaceae_Gossypium_bickii
Malvaceae_Gossypium_australe
Malvaceae_Gossypium_robinsonii
Malvaceae_Gossypium_sturtianum
Malvaceae_Gossypium_nobile
Malvaceae_Gossypium_cunninghamii
Malvaceae_Gossypium_sp
Malvaceae_Gossypium_anapoides
Malvaceae_Gossypium_marchantii
Malvaceae_Gossypium_pulchellum
Malvaceae_Gossypium_costulatum
Malvaceae_Gossypium_enthyle
Malvaceae_Gossypium_rotundifolium
Malvaceae_Gossypium_exiguum
Malvaceae_Gossypium_pilosum
Malvaceae_Gossypium_populifolium
Malvaceae_Gossypium_londonderriense
Malvaceae_Gossypium_incanum
Malvaceae_Gossypium_areysianum
Malvaceae_Gossypium_stocksii
Malvaceae_Gossypium_somalense
Malvaceae_Fuertesimalva_chilensis
Malvaceae_Fuertesimalva_peruviana
Malvaceae_Fuertesimalva_limensis
Malvaceae_Fuertesimalva_jacens
Malvaceae_Fuertesimalva_echinata
Malvaceae_Kearnemalvastrum_sp
Malvaceae_Kearnemalvastrum_lacteum
Malvaceae_Andeimalva_chilensis
Malvaceae_Andeimalva_mandonii
Malvaceae_Andeimalva_machupicchensis
Malvaceae_Andeimalva_spiciformis

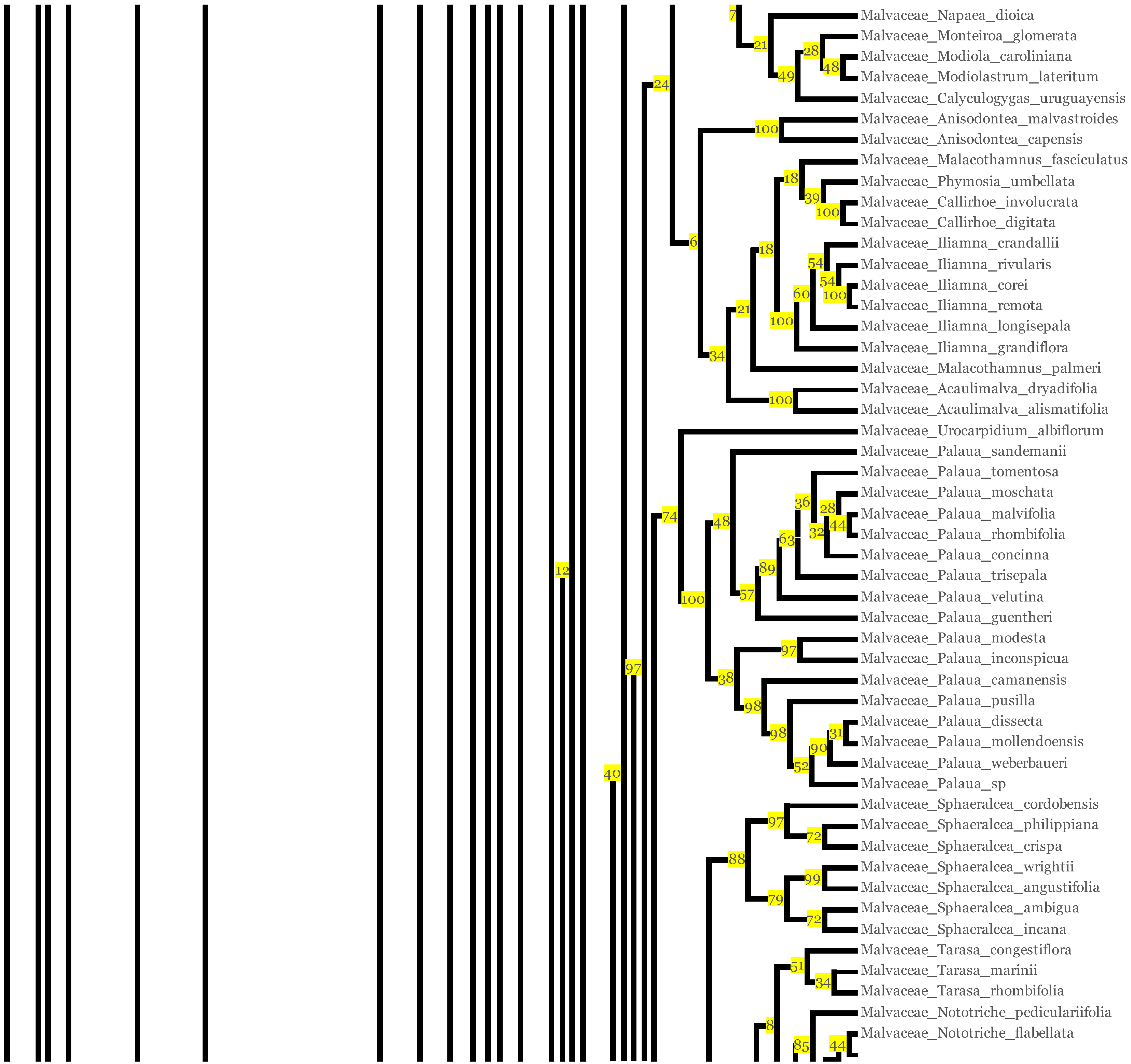
Malvaceae_Napaea_dioica
Malvaceae_Monteiroa_glomerata
Malvaceae_Modiola_caroliniana
Malvaceae_Modiolastrum_lateritum
Malvaceae_Calyculogygas_uruguayensis
Malvaceae_Anisodontea_malvastroides
Malvaceae_Anisodontea_capensis
Malvaceae_Malacothamnus_fasciculatus
Malvaceae_Phymosia_umbellata
Malvaceae_Callirhoe_involucrata
Malvaceae_Callirhoe_digitata
Malvaceae_Iliamna_crandallii
Malvaceae_Iliamna_rivularis
Malvaceae_Iliamna_corei
Malvaceae_Iliamna_remota
Malvaceae_Iliamna_longisepala
Malvaceae_Iliamna_grandiflora
Malvaceae_Malacothamnus_palmeri
Malvaceae_Acaulimalva_dryadifolia
Malvaceae_Acaulimalva_alismatifolia
Malvaceae_Urocarpidium_albiflorum
Malvaceae_Palaua_sandemanii
Malvaceae_Palaua_tomentosa
Malvaceae_Palaua_moschata
Malvaceae_Palaua_malvifolia
Malvaceae_Palaua_rhombifolia
Malvaceae_Palaua_concinna
Malvaceae_Palaua_trisepala
Malvaceae_Palaua_velutina
Malvaceae_Palaua_guentheri
Malvaceae_Palaua_modesta
Malvaceae_Palaua_inconspicua
Malvaceae_Palaua_camanensis
Malvaceae_Palaua_pusilla
Malvaceae_Palaua_dissecta
Malvaceae_Palaua_mollendoensis
Malvaceae_Palaua_weberbaueri
Malvaceae_Palaua_sp
Malvaceae_Sphaeralcea_cordobensis
Malvaceae_Sphaeralcea_philippiana
Malvaceae_Sphaeralcea_crispa
Malvaceae_Sphaeralcea_wrightii
Malvaceae_Sphaeralcea_angustifolia
Malvaceae_Sphaeralcea_ambigua
Malvaceae_Sphaeralcea_incana
Malvaceae_Tarasa_congestiflora
Malvaceae_Tarasa_marinii
Malvaceae_Tarasa_rhombifolia
Malvaceae_Nototriche_pedicularifolia
Malvaceae_Nototriche_flabellata

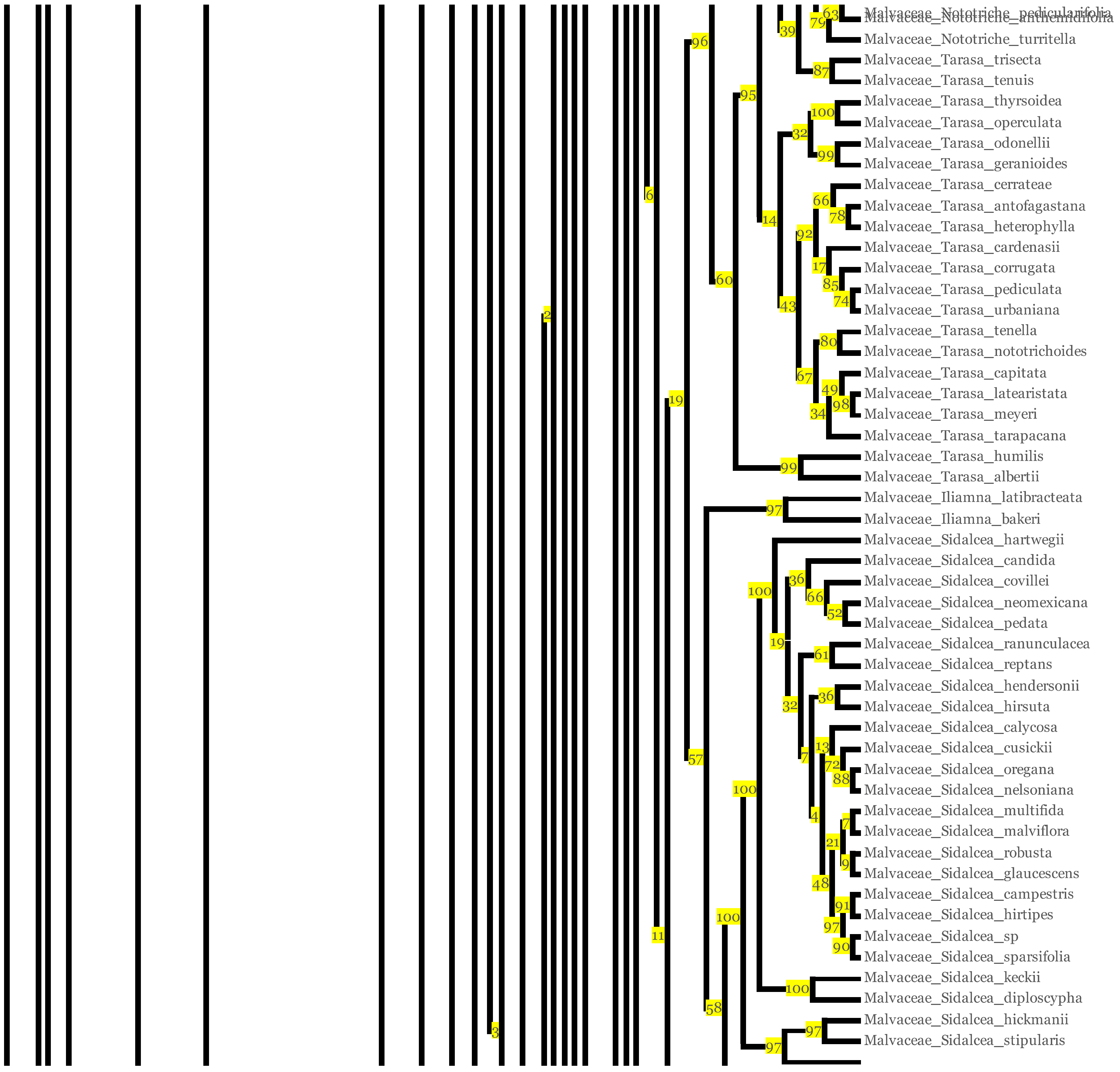
Malvaceae_Nototriche_pedicularifolia
Malvaceae_Nototriche_anthemidifolia
Malvaceae_Nototriche_turritella
Malvaceae_Tarasa_trisecta
Malvaceae_Tarasa_tenuis
Malvaceae_Tarasa_thyrsoidea
Malvaceae_Tarasa_operculata
Malvaceae_Tarasa_odonellii
Malvaceae_Tarasa_geranioides
Malvaceae_Tarasa_cerrateae
Malvaceae_Tarasa_antofagastana
Malvaceae_Tarasa_heterophylla
Malvaceae_Tarasa_cardenasii
Malvaceae_Tarasa_corrugata
Malvaceae_Tarasa_pediculata
Malvaceae_Tarasa_urbaniana
Malvaceae_Tarasa_tenella
Malvaceae_Tarasa_nototrichoides
Malvaceae_Tarasa_capitata
Malvaceae_Tarasa_latearistata
Malvaceae_Tarasa_meyeri
Malvaceae_Tarasa_tarapacana
Malvaceae_Tarasa_humilis
Malvaceae_Tarasa_albertii
Malvaceae_Iliamna_latibracteata
Malvaceae_Iliamna_bakeri
Malvaceae_Sidalcea_hartwegii
Malvaceae_Sidalcea_candida
Malvaceae_Sidalcea_covillei
Malvaceae_Sidalcea_neomexicana
Malvaceae_Sidalcea_pedata
Malvaceae_Sidalcea_ranunculacea
Malvaceae_Sidalcea_reptans
Malvaceae_Sidalcea_hendersonii
Malvaceae_Sidalcea_hirsuta
Malvaceae_Sidalcea_calycosa
Malvaceae_Sidalcea_cusickii
Malvaceae_Sidalcea_oregana
Malvaceae_Sidalcea_nelsoniana
Malvaceae_Sidalcea_multifida
Malvaceae_Sidalcea_malviflora
Malvaceae_Sidalcea_robusta
Malvaceae_Sidalcea_glaucescens
Malvaceae_Sidalcea_campestris
Malvaceae_Sidalcea_hirtipes
Malvaceae_Sidalcea_sp
Malvaceae_Sidalcea_sparsifolia
Malvaceae_Sidalcea_keckii
Malvaceae_Sidalcea_diploscypha
Malvaceae_Sidalcea_hickmanii
Malvaceae_Sidalcea_stipularis

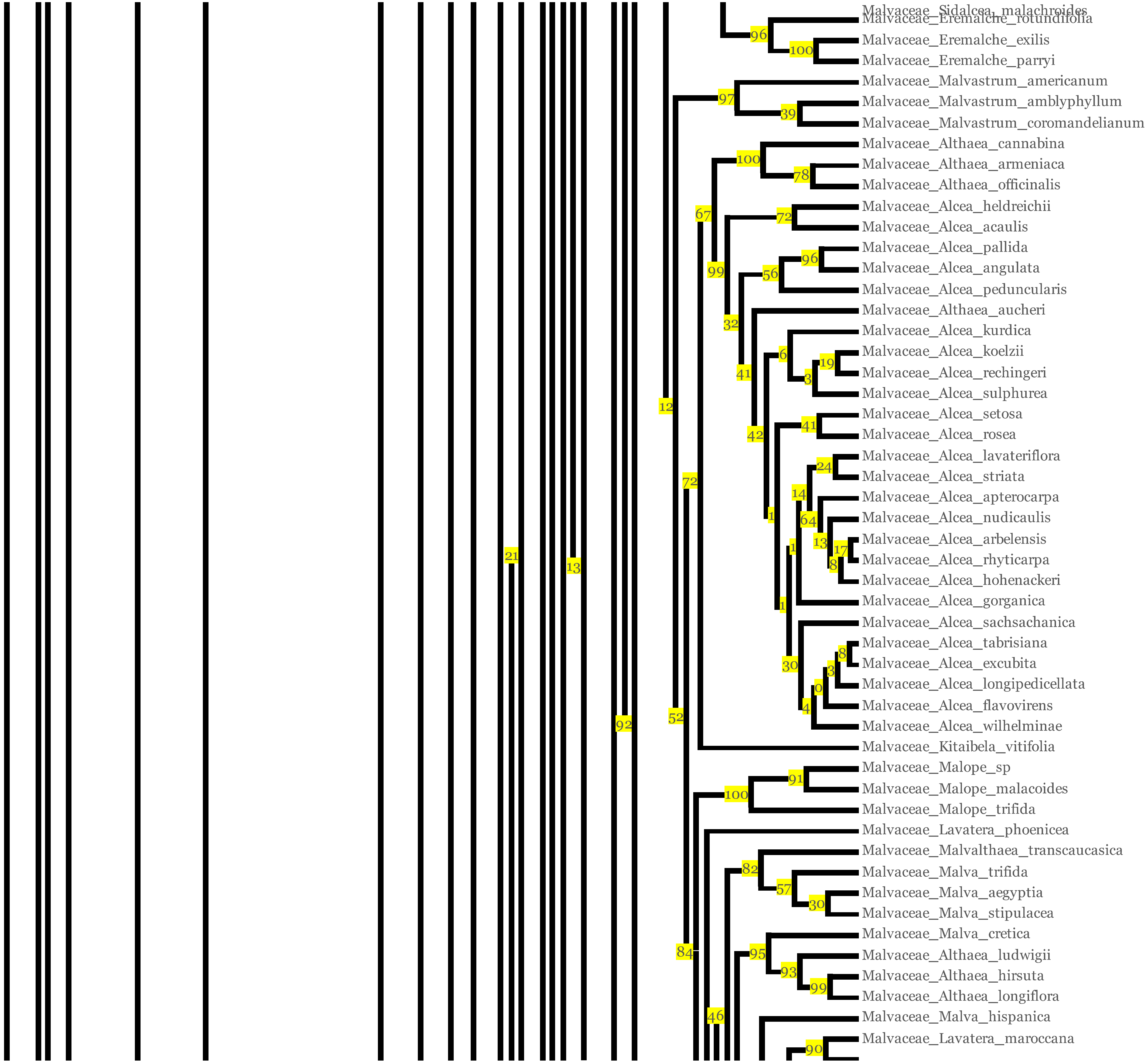
Malvaceae_Sidalcea_malachroides
Malvaceae_Eremalche_rotundifolia
Malvaceae_Eremalche_exilis
Malvaceae_Eremalche_parryi
Malvaceae_Malvastrum_americanum
Malvaceae_Malvastrum_amblyphyllum
Malvaceae_Malvastrum_coromandelianum
Malvaceae_Althaea_cannabina
Malvaceae_Althaea_armeniaca
Malvaceae_Althaea_officinalis
Malvaceae_Alcea_heldreichii
Malvaceae_Alcea_acaulis
Malvaceae_Alcea_pallida
Malvaceae_Alcea_angulata
Malvaceae_Alcea_peduncularis
Malvaceae_Althaea_aucheri
Malvaceae_Alcea_kurdica
Malvaceae_Alcea_koelzii
Malvaceae_Alcea_rechingeri
Malvaceae_Alcea_sulphurea
Malvaceae_Alcea_setosa
Malvaceae_Alcea_rosea
Malvaceae_Alcea_lavateriflora
Malvaceae_Alcea_striata
Malvaceae_Alcea_apterocarpa
Malvaceae_Alcea_nudicaulis
Malvaceae_Alcea_arbelensis
Malvaceae_Alcea_rhyticarpa
Malvaceae_Alcea_hohenackeri
Malvaceae_Alcea_gorganica
Malvaceae_Alcea_sachsachanica
Malvaceae_Alcea_tabrisiana
Malvaceae_Alcea_excubita
Malvaceae_Alcea_longipedicellata
Malvaceae_Alcea_flavovirens
Malvaceae_Alcea_wilhelminae
Malvaceae_Kitaibela_vitifolia
Malvaceae_Malope_sp
Malvaceae_Malope_malacoides
Malvaceae_Malope_trifida
Malvaceae_Lavatera_phoenicea
Malvaceae_Malvalthaea_transcaucasica
Malvaceae_Malva_trifida
Malvaceae_Malva_aegyptia
Malvaceae_Malva_stipulacea
Malvaceae_Malva_cretica
Malvaceae_Althaea_ludwigii
Malvaceae_Althaea_hirsuta
Malvaceae_Althaea_longiflora
Malvaceae_Malva_hispanica
Malvaceae_Lavatera_maroccana

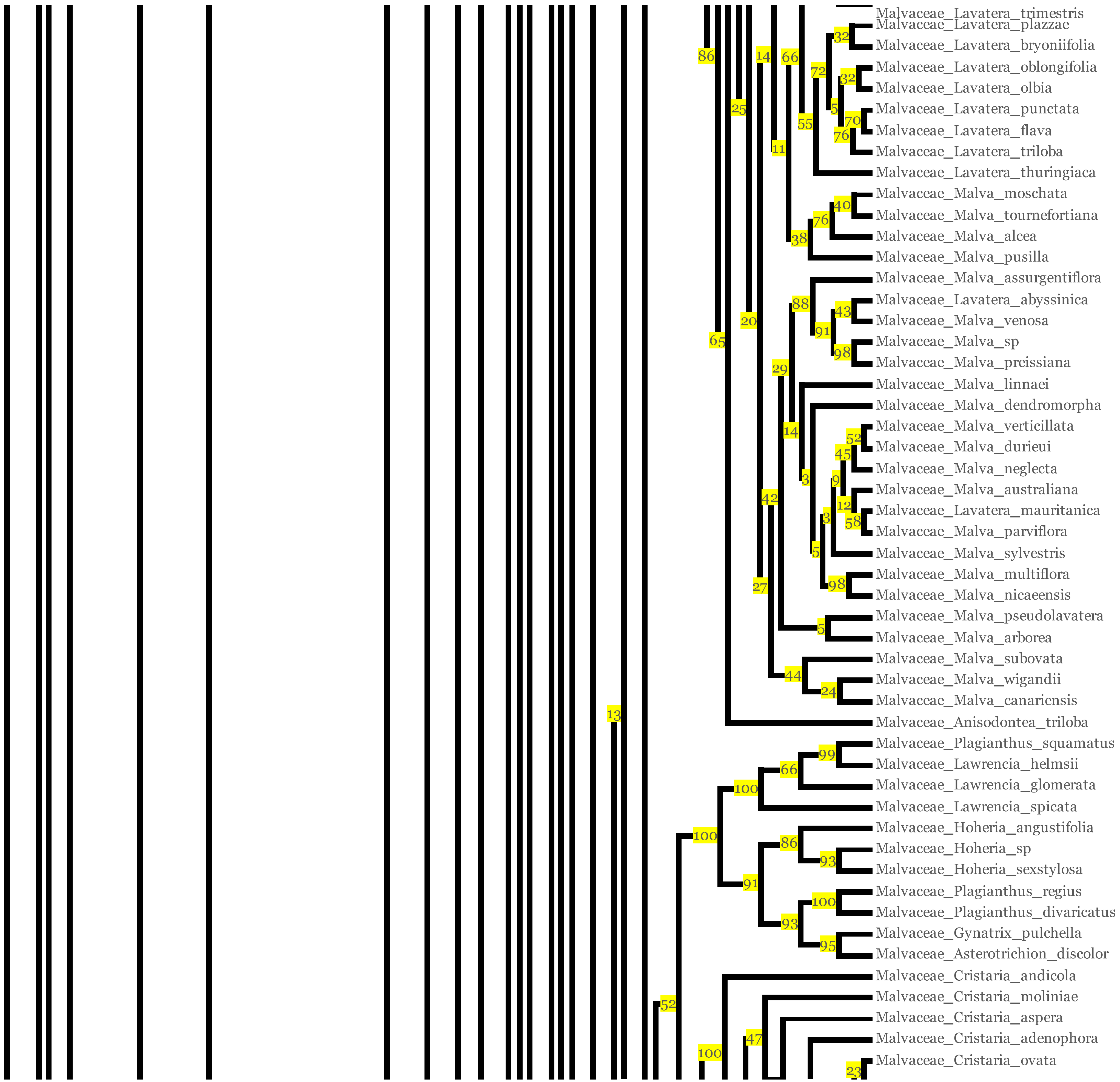
Malvaceae_Lavatera_trimestris
Malvaceae_Lavatera_plazzae
Malvaceae_Lavatera_bryoniifolia
Malvaceae_Lavatera_oblongifolia
Malvaceae_Lavatera_olbia
Malvaceae_Lavatera_punctata
Malvaceae_Lavatera_flava
Malvaceae_Lavatera_triloba
Malvaceae_Lavatera_thuringiaca
Malvaceae_Malva_moschata
Malvaceae_Malva_tournefortiana
Malvaceae_Malva_alcea
Malvaceae_Malva_pusilla
Malvaceae_Malva_assurgentiflora
Malvaceae_Lavatera_abyssinica
Malvaceae_Malva_venosa
Malvaceae_Malva_sp
Malvaceae_Malva_preissiana
Malvaceae_Malva_linnaei
Malvaceae_Malva_dendromorpha
Malvaceae_Malva_verticillata
Malvaceae_Malva_durieui
Malvaceae_Malva_neglecta
Malvaceae_Malva_australiana
Malvaceae_Lavatera_mauritanica
Malvaceae_Malva_parviflora
Malvaceae_Malva_sylvestris
Malvaceae_Malva_multiflora
Malvaceae_Malva_nicaeensis
Malvaceae_Malva_pseudolavatera
Malvaceae_Malva_arborea
Malvaceae_Malva_subovata
Malvaceae_Malva_wigandii
Malvaceae_Malva_canariensis
Malvaceae_Anisodontea_triloba
Malvaceae_Plagianthus_squamatus
Malvaceae_Lawrencia_helmsii
Malvaceae_Lawrencia_glomerata
Malvaceae_Lawrencia_spicata
Malvaceae_Hoheria_angustifolia
Malvaceae_Hoheria_sp
Malvaceae_Hoheria_sexstylosa
Malvaceae_Plagianthus_regius
Malvaceae_Plagianthus_divaricatus
Malvaceae_Gynatrix_pulchella
Malvaceae_Asterotrichion_discolor
Malvaceae_Cristaria_andicola
Malvaceae_Cristaria_moliniae
Malvaceae_Cristaria_aspera
Malvaceae_Cristaria_adenophora
Malvaceae_Cristaria_ovata

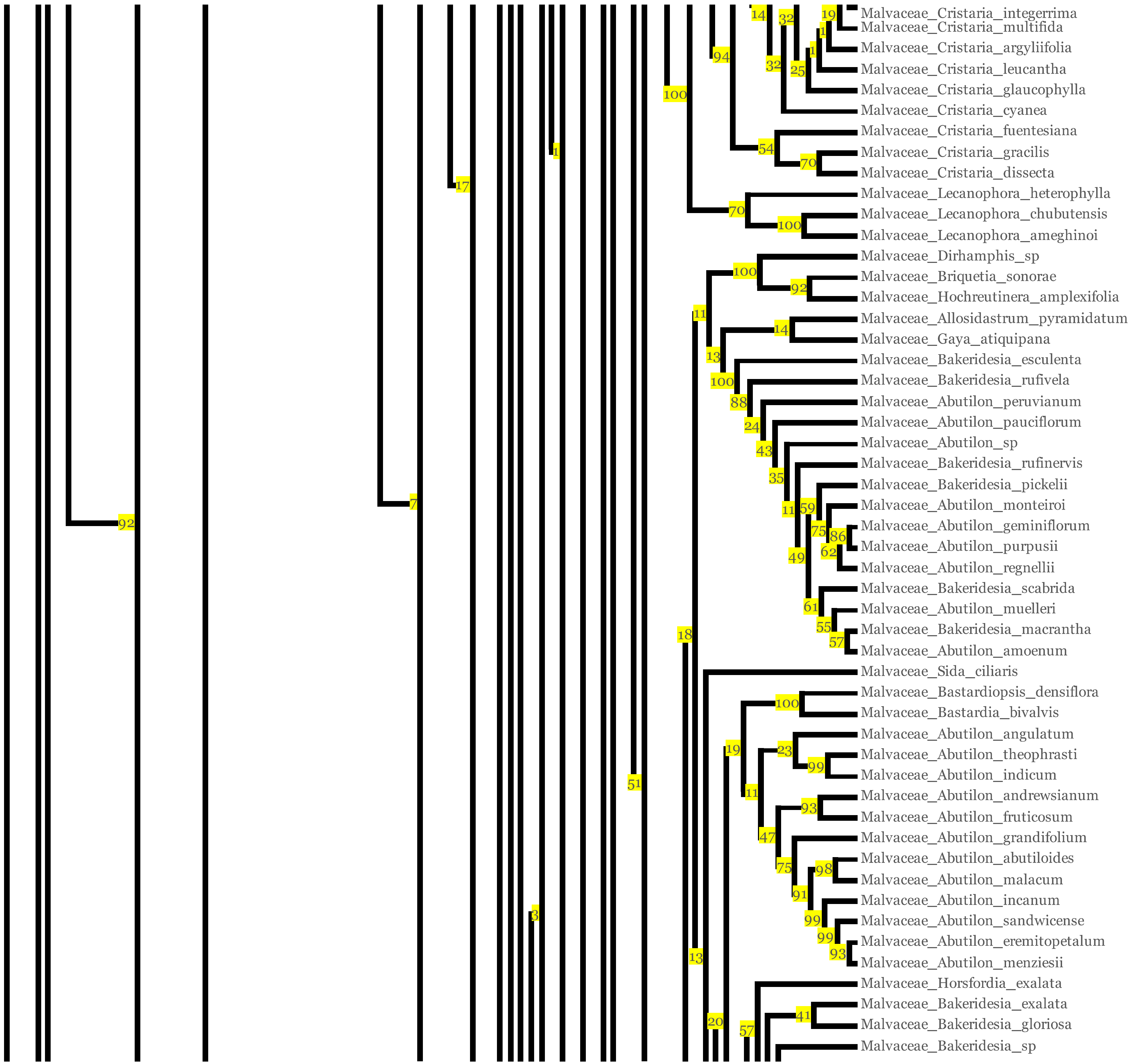
Malvaceae_Cristaria_integerrima
Malvaceae_Cristaria_multifida
Malvaceae_Cristaria_argyliifolia
Malvaceae_Cristaria_leucantha
Malvaceae_Cristaria_glaucophylla
Malvaceae_Cristaria_cyanea
Malvaceae_Cristaria_fuentesiana
Malvaceae_Cristaria_gracilis
Malvaceae_Cristaria_dissecta
Malvaceae_Lecanophora_heterophylla
Malvaceae_Lecanophora_chubutensis
Malvaceae_Lecanophora_ameghinoi
Malvaceae_Dirhamphis_sp
Malvaceae_Briquetia_sonorae
Malvaceae_Hochreutinera_amplexifolia
Malvaceae_Allosidastrum_pyramidatum
Malvaceae_Gaya_atiquipana
Malvaceae_Bakeridesia_esculenta
Malvaceae_Bakeridesia_rufivela
Malvaceae_Abutilon_peruvianum
Malvaceae_Abutilon_pauciflorum
Malvaceae_Abutilon_sp
Malvaceae_Bakeridesia_rufinervis
Malvaceae_Bakeridesia_pickelii
Malvaceae_Abutilon_monteiroi
Malvaceae_Abutilon_geminiflorum
Malvaceae_Abutilon_purpusii
Malvaceae_Abutilon_regnellii
Malvaceae_Bakeridesia_scabrida
Malvaceae_Abutilon_muelleri
Malvaceae_Bakeridesia_macrantha
Malvaceae_Abutilon_amoenum
Malvaceae_Sida_ciliaris
Malvaceae_Bastardiopsis_densiflora
Malvaceae_Bastardia_bivalvis
Malvaceae_Abutilon_angulatum
Malvaceae_Abutilon_theophrasti
Malvaceae_Abutilon_indicum
Malvaceae_Abutilon_andrewsianum
Malvaceae_Abutilon_fruticosum
Malvaceae_Abutilon_grandifolium
Malvaceae_Abutilon_abutiloides
Malvaceae_Abutilon_malacum
Malvaceae_Abutilon_incanum
Malvaceae_Abutilon_sandwicense
Malvaceae_Abutilon_eremitopetalum
Malvaceae_Abutilon_menziesii
Malvaceae_Horsfordia_exalata
Malvaceae_Bakeridesia_exalata
Malvaceae_Bakeridesia_gloriosa
Malvaceae_Bakeridesia_sp

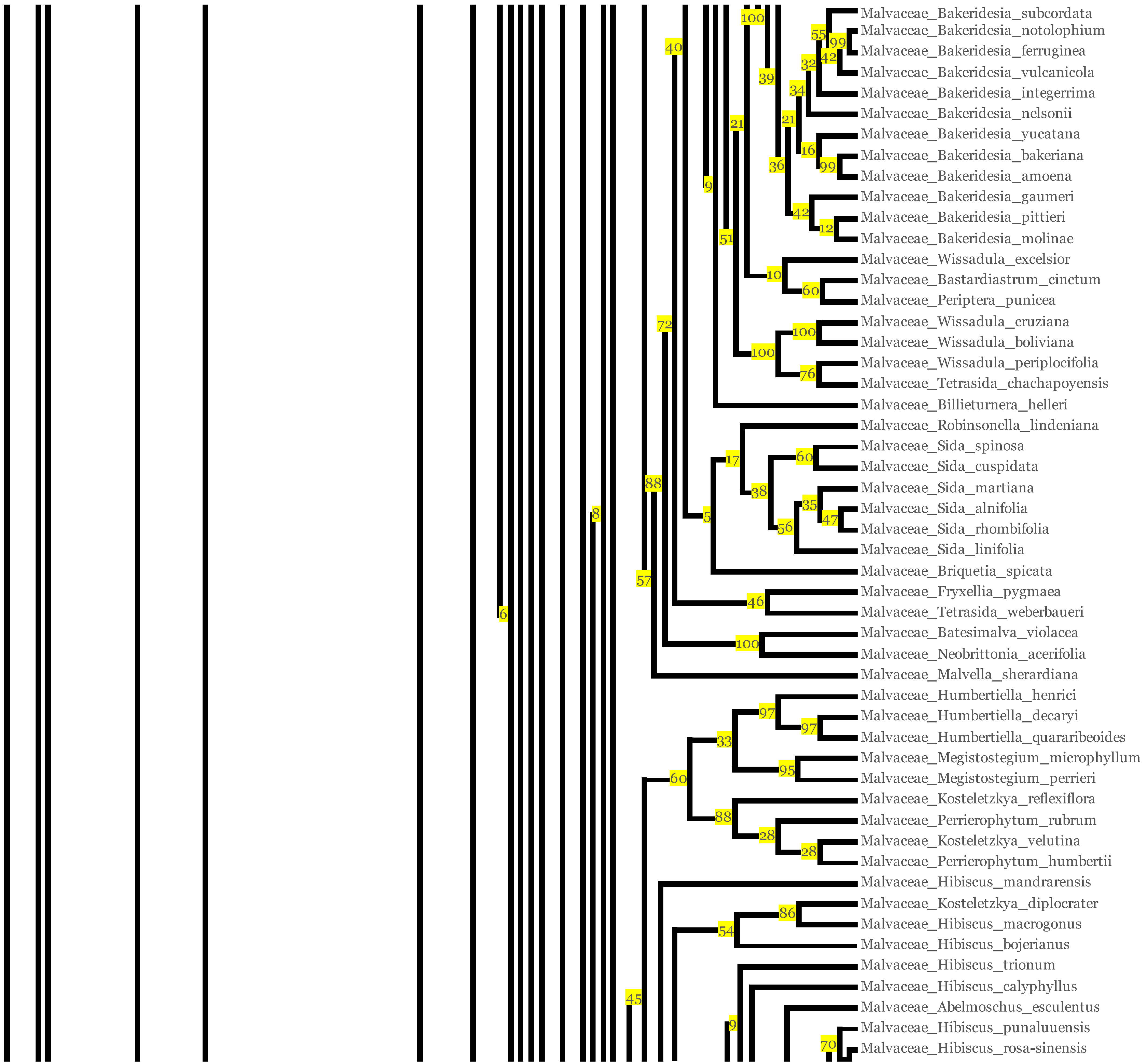
Malvaceae_Bakeridesia_subcordata
Malvaceae_Bakeridesia_notolophium
Malvaceae_Bakeridesia_ferruginea
Malvaceae_Bakeridesia_vulcanicola
Malvaceae_Bakeridesia_integerrima
Malvaceae_Bakeridesia_nelsonii
Malvaceae_Bakeridesia_yucatana
Malvaceae_Bakeridesia_bakeriana
Malvaceae_Bakeridesia_amoena
Malvaceae_Bakeridesia_gaumeri
Malvaceae_Bakeridesia_pittieri
Malvaceae_Bakeridesia_molinae
Malvaceae_Wissadula_excelsior
Malvaceae_Bastardiastrum_cinctum
Malvaceae_Periptera_punicea
Malvaceae_Wissadula_cruziana
Malvaceae_Wissadula_boliviana
Malvaceae_Wissadula_periplocifolia
Malvaceae_Tetrasida_chachapoyensis
Malvaceae_Billieturnera_helleri
Malvaceae_Robinsonella_lindeniana
Malvaceae_Sida_spinosa
Malvaceae_Sida_cuspidata
Malvaceae_Sida_martiana
Malvaceae_Sida_alnifolia
Malvaceae_Sida_rhombifolia
Malvaceae_Sida_linifolia
Malvaceae_Briquetia_spicata
Malvaceae_Fryxellia_pygmaea
Malvaceae_Tetrasida_weberbaueri
Malvaceae_Batesimalva_violacea
Malvaceae_Neobrittonia_acerifolia
Malvaceae_Malvella_sherardiana
Malvaceae_Humbertiella_henrici
Malvaceae_Humbertiella_decaryi
Malvaceae_Humbertiella_quararibeoides
Malvaceae_Megistostegium_microphyllum
Malvaceae_Megistostegium_perrieri
Malvaceae_Kosteletzkya_reflexiflora
Malvaceae_Perrierophytum_rubrum
Malvaceae_Kosteletzkya_velutina
Malvaceae_Perrierophytum_humbertii
Malvaceae_Hibiscus_mandrarensis
Malvaceae_Kosteletzkya_diplocrater
Malvaceae_Hibiscus_macrogonus
Malvaceae_Hibiscus_bojerianus
Malvaceae_Hibiscus_trionum
Malvaceae_Hibiscus_calyphyllus
Malvaceae_Abelmoschus_esculentus
Malvaceae_Hibiscus_punaluuensis
Malvaceae_Hibiscus_rosa-sinensis

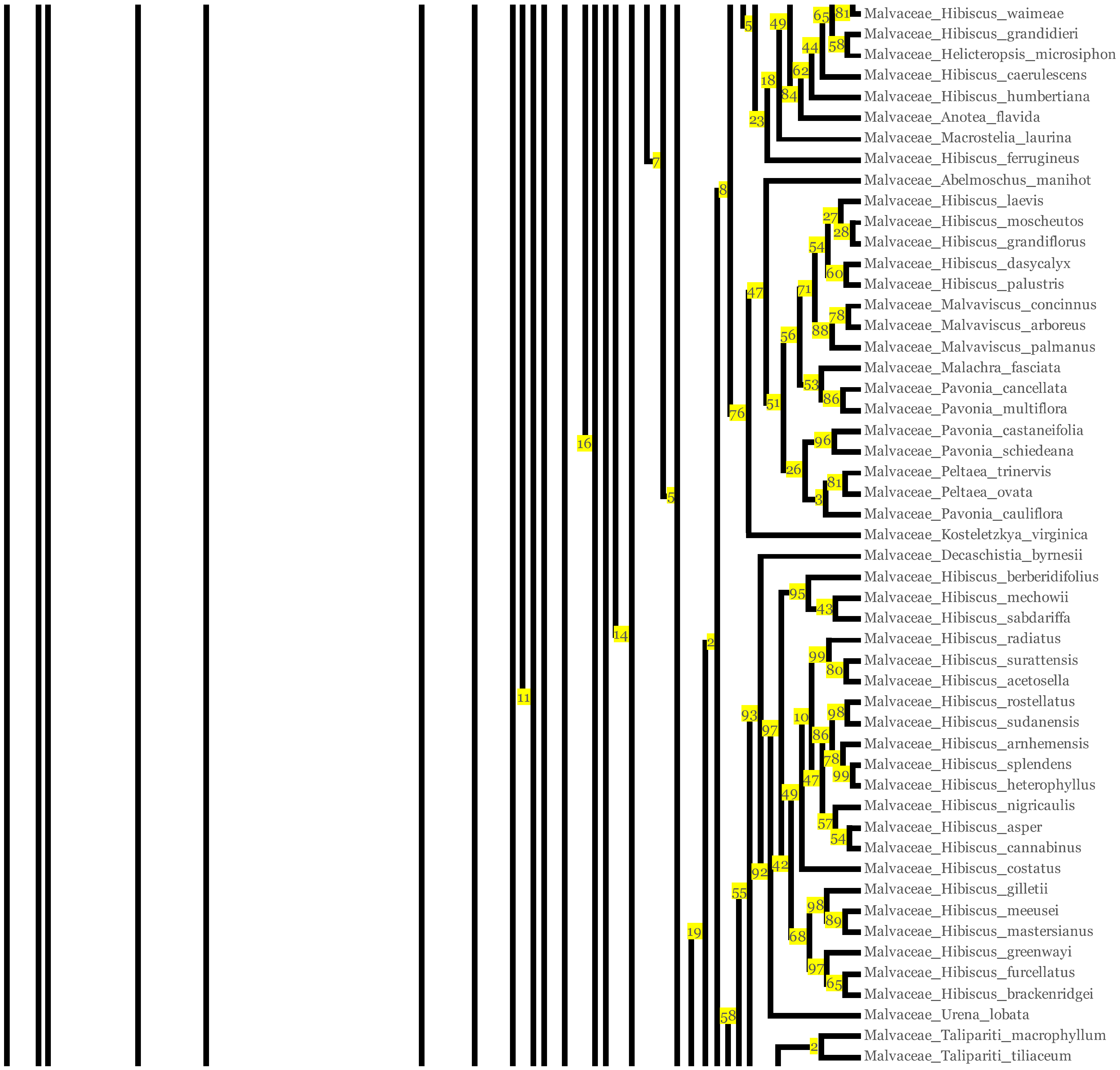
Malvaceae_Hibiscus_waimeae
Malvaceae_Hibiscus_grandidieri
Malvaceae_Helicteropsis_microsiphon
Malvaceae_Hibiscus_caerulescens
Malvaceae_Hibiscus_humbertiana
Malvaceae_Anotea_flavida
Malvaceae_Macrostelia_laurina
Malvaceae_Hibiscus_ferrugineus
Malvaceae_Abelmoschus_manihot
Malvaceae_Hibiscus_laevis
Malvaceae_Hibiscus_moscheutos
Malvaceae_Hibiscus_grandiflorus
Malvaceae_Hibiscus_dasycalyx
Malvaceae_Hibiscus_palustris
Malvaceae_Malvaviscus_concinnus
Malvaceae_Malvaviscus_arboreus
Malvaceae_Malvaviscus_palmanus
Malvaceae_Malachra_fasciata
Malvaceae_Pavonia_cancellata
Malvaceae_Pavonia_multiflora
Malvaceae_Pavonia_castaneifolia
Malvaceae_Pavonia_schiedeana
Malvaceae_Peltaea_trinervis
Malvaceae_Peltaea_ovata
Malvaceae_Pavonia_cauliflora
Malvaceae_Kosteletzkya_virginica
Malvaceae_Decaschistia_byrnesii
Malvaceae_Hibiscus_berberidifolius
Malvaceae_Hibiscus_mechowii
Malvaceae_Hibiscus_sabdariffa
Malvaceae_Hibiscus_radiatus
Malvaceae_Hibiscus_surattensis
Malvaceae_Hibiscus_acetosella
Malvaceae_Hibiscus_rostellatus
Malvaceae_Hibiscus_sudanensis
Malvaceae_Hibiscus_arnhemensis
Malvaceae_Hibiscus_splendens
Malvaceae_Hibiscus_heterophyllus
Malvaceae_Hibiscus_nigricaulis
Malvaceae_Hibiscus_asper
Malvaceae_Hibiscus_cannabinus
Malvaceae_Hibiscus_costatus
Malvaceae_Hibiscus_gilletii
Malvaceae_Hibiscus_meeusei
Malvaceae_Hibiscus_mastersianus
Malvaceae_Hibiscus_greenwayi
Malvaceae_Hibiscus_furcellatus
Malvaceae_Hibiscus_brackenridgei
Malvaceae_Urena_lobata
Malvaceae_Talipariti_macrophyllum
Malvaceae_Talipariti_tiliaceum

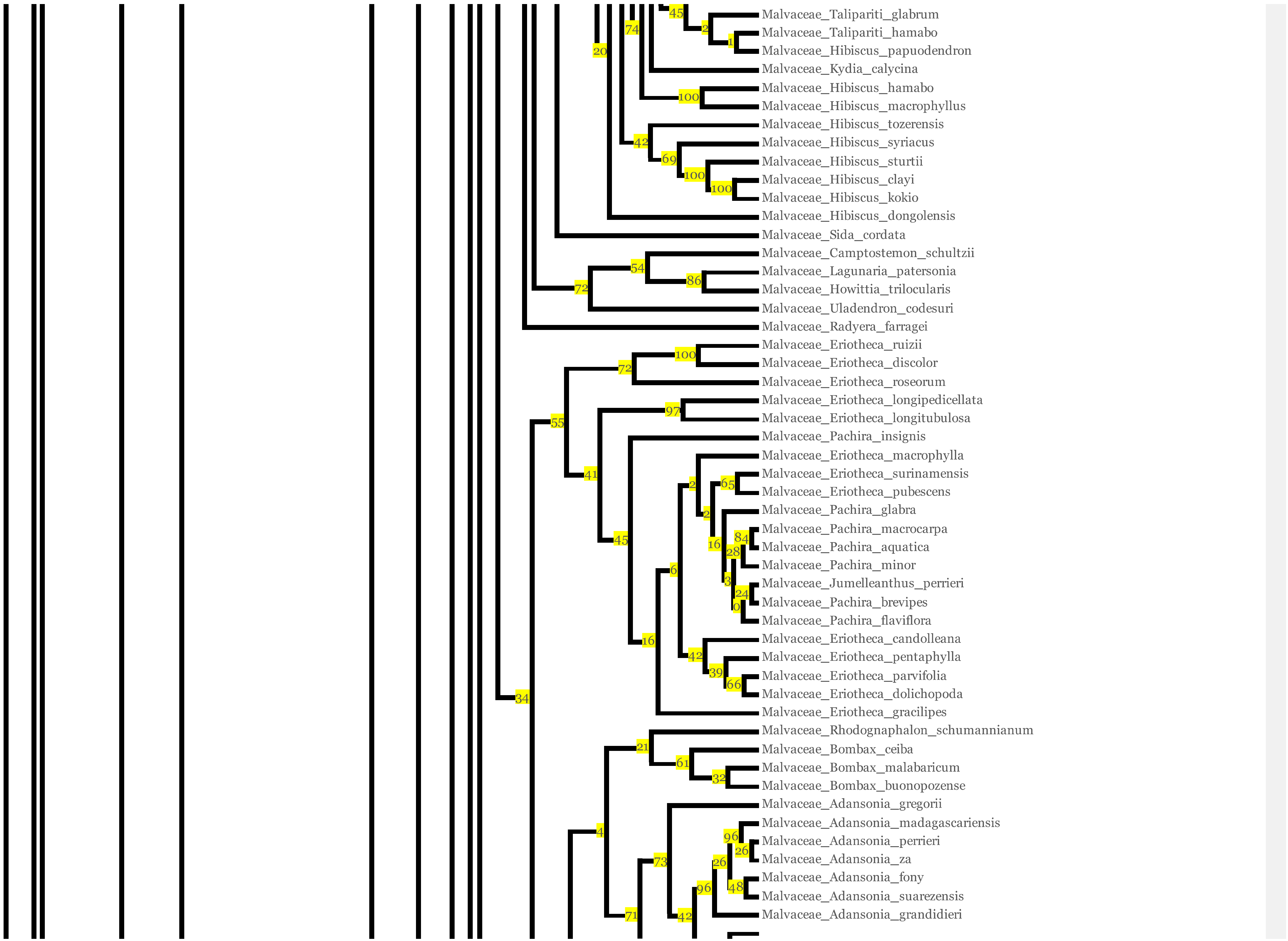
Malvaceae_Talipariti_glabrum
Malvaceae_Talipariti_hamabo
Malvaceae_Hibiscus_papuodendron
Malvaceae_Kydia_calycina
Malvaceae_Hibiscus_hamabo
Malvaceae_Hibiscus_macrophyllus
Malvaceae_Hibiscus_tozerensis
Malvaceae_Hibiscus_syriacus
Malvaceae_Hibiscus_sturtii
Malvaceae_Hibiscus_clayi
Malvaceae_Hibiscus_kokio
Malvaceae_Hibiscus_dongolensis
Malvaceae_Sida_cordata
Malvaceae_Camptostemon_schultzii
Malvaceae_Lagunaria_patersonia
Malvaceae_Howittia_trilocularis
Malvaceae_Uladendron_codesuri
Malvaceae_Radyera_farragei
Malvaceae_Eriotheca_ruizii
Malvaceae_Eriotheca_discolor
Malvaceae_Eriotheca_roseorum
Malvaceae_Eriotheca_longipedicellata
Malvaceae_Eriotheca_longitubulosa
Malvaceae_Pachira_insignis
Malvaceae_Eriotheca_macrophylla
Malvaceae_Eriotheca_surinamensis
Malvaceae_Eriotheca_pubescens
Malvaceae_Pachira_glabra
Malvaceae_Pachira_macrocarpa
Malvaceae_Pachira_aquatica
Malvaceae_Pachira_minor
Malvaceae_Jumelleanthus_perrieri
Malvaceae_Pachira_brevipes
Malvaceae_Pachira_flaviflora
Malvaceae_Eriotheca_candolleana
Malvaceae_Eriotheca_pentaphylla
Malvaceae_Eriotheca_parvifolia
Malvaceae_Eriotheca_dolichopoda
Malvaceae_Eriotheca_gracilipes
Malvaceae_Rhodognaphalon_schumannianum
Malvaceae_Bombax_ceiba
Malvaceae_Bombax_malabaricum
Malvaceae_Bombax_buonopozense
Malvaceae_Adansonia_gregorii
Malvaceae_Adansonia_madagascariensis
Malvaceae_Adansonia_perrieri
Malvaceae_Adansonia_za
Malvaceae_Adansonia_fony
Malvaceae_Adansonia_suarezensis
Malvaceae_Adansonia_grandidieri

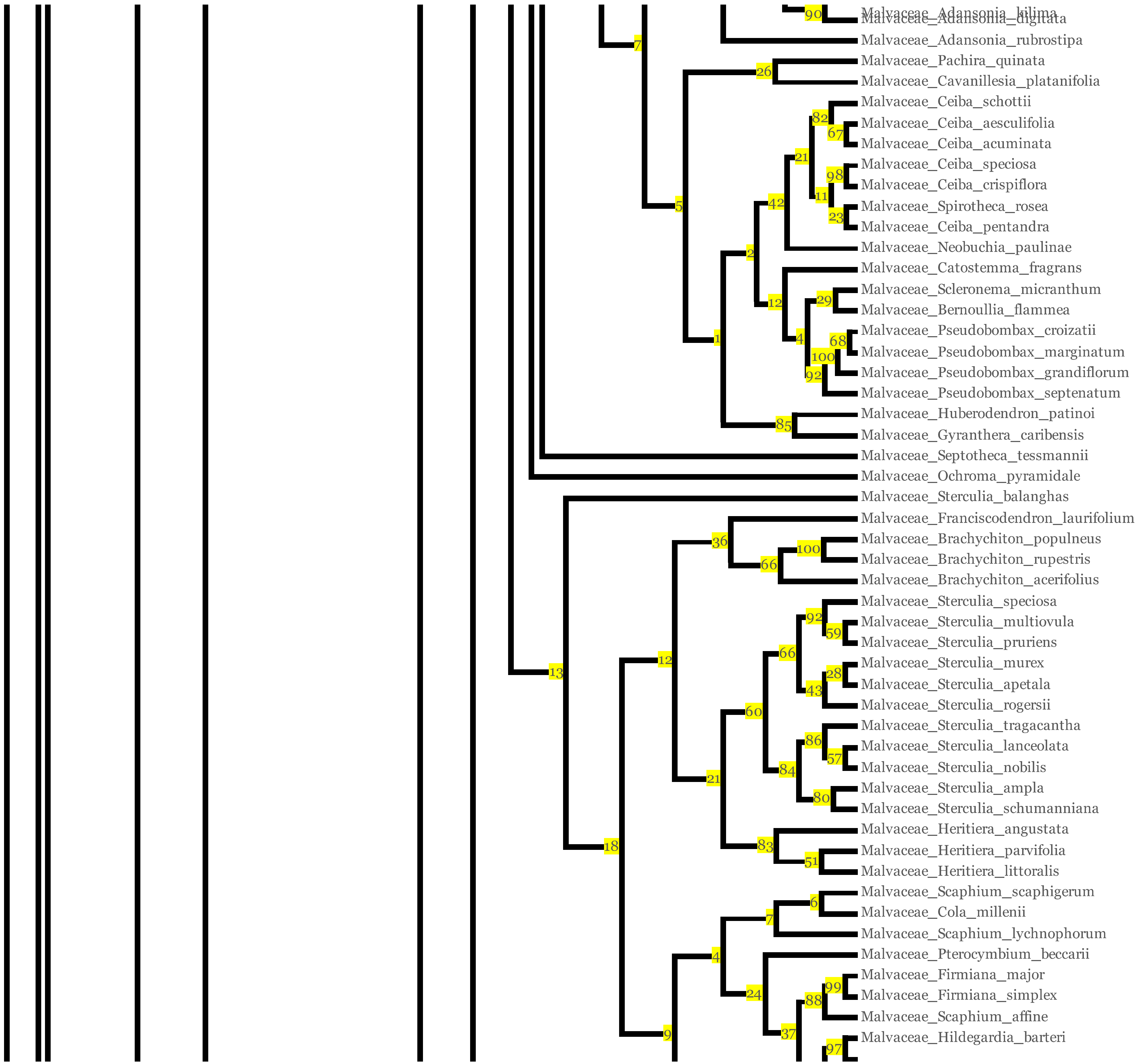
Malvaceae_Adansonia_kilima
Malvaceae_Adansonia_digitata
Malvaceae_Adansonia_rubrostipa
Malvaceae_Pachira_quinata
Malvaceae_Cavanillesia_platanifolia
Malvaceae_Ceiba_schottii
Malvaceae_Ceiba_aesculifolia
Malvaceae_Ceiba_acuminata
Malvaceae_Ceiba_speciosa
Malvaceae_Ceiba_crispiflora
Malvaceae_Spirotheca_rosea
Malvaceae_Ceiba_pentandra
Malvaceae_Neobuchia_paulinae
Malvaceae_Catostemma_fragrans
Malvaceae_Scleronema_micranthum
Malvaceae_Bernoullia_flammea
Malvaceae_Pseudobombax_croizatii
Malvaceae_Pseudobombax_marginatum
Malvaceae_Pseudobombax_grandiflorum
Malvaceae_Pseudobombax_septenatum
Malvaceae_Huberodendron_patinoi
Malvaceae_Gyranthera_caribensis
Malvaceae_Septotheca_tessmannii
Malvaceae_Ochroma_pyramidale
Malvaceae_Sterculia_balanghas
Malvaceae_Franciscodendron_laurifolium
Malvaceae_Brachychiton_populneus
Malvaceae_Brachychiton_rupestris
Malvaceae_Brachychiton_acerifolius
Malvaceae_Sterculia_speciosa
Malvaceae_Sterculia_multiovula
Malvaceae_Sterculia_pruriens
Malvaceae_Sterculia_murex
Malvaceae_Sterculia_apetala
Malvaceae_Sterculia_rogersii
Malvaceae_Sterculia_tragacantha
Malvaceae_Sterculia_lanceolata
Malvaceae_Sterculia_nobilis
Malvaceae_Sterculia_ampla
Malvaceae_Sterculia_schumanniana
Malvaceae_Heritiera_angustata
Malvaceae_Heritiera_parvifolia
Malvaceae_Heritiera_littoralis
Malvaceae_Scaphium_scaphigerum
Malvaceae_Cola_millenii
Malvaceae_Scaphium_lychnophorum
Malvaceae_Pterocymbium_beccarii
Malvaceae_Firmiana_major
Malvaceae_Firmiana_simplex
Malvaceae_Scaphium_affine
Malvaceae_Hildegardia_barteri

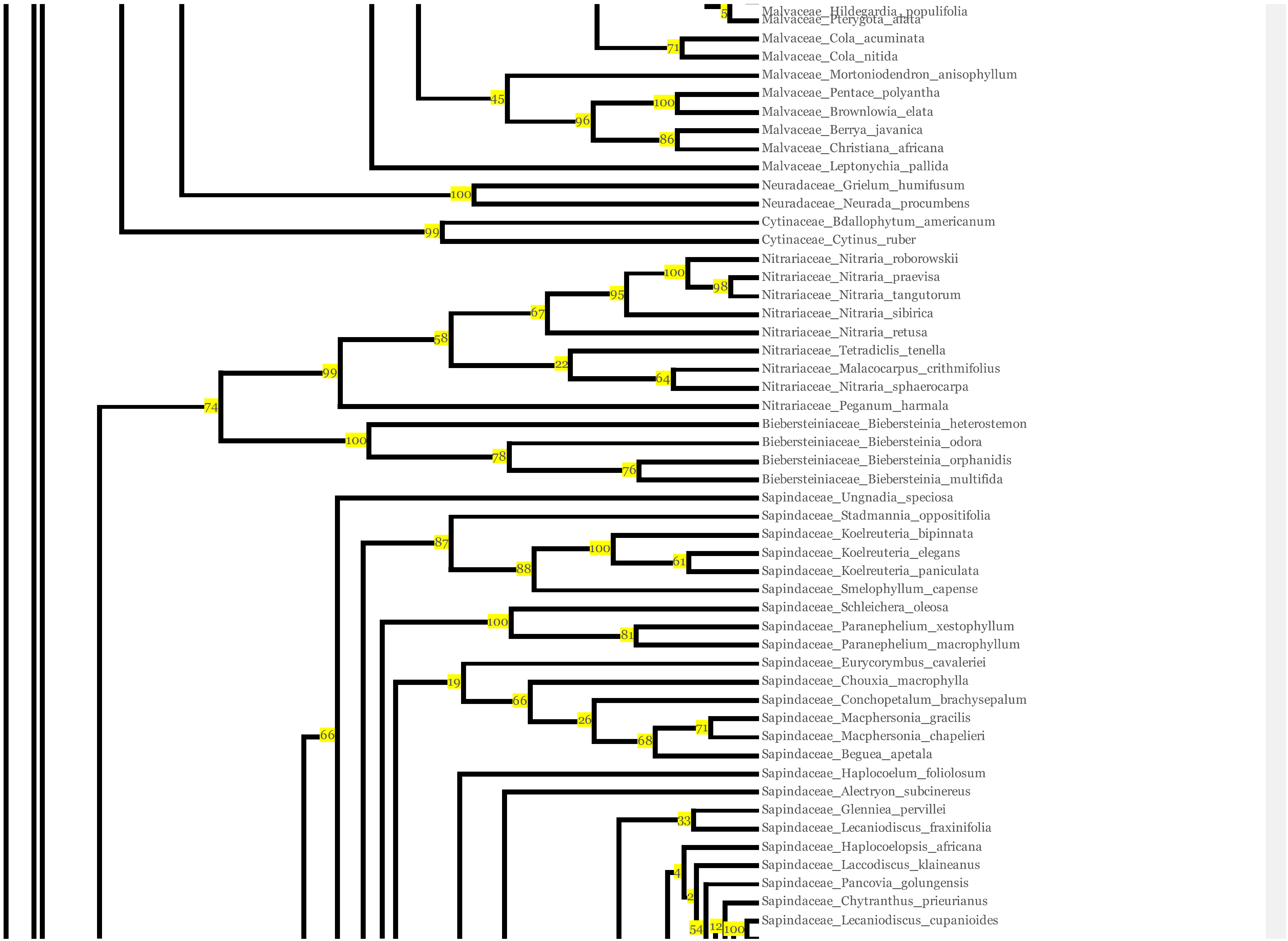
Malvaceae_Hildegardia_populifolia
Malvaceae_Pterygota_alata
Malvaceae_Cola_acuminata
Malvaceae_Cola_nitida
Malvaceae_Mortoniodendron_anisophyllum
Malvaceae_Pentace_polyantha
Malvaceae_Brownlowia_elata
Malvaceae_Berrya_javanica
Malvaceae_Christiana_africana
Malvaceae_Leptonychia_pallida
Neuradaceae_Grielum_humifusum
Neuradaceae_Neurada_procumbens
Cytinaceae_Bdallophytum_americanum
Cytinaceae_Cytinus_ruber
Nitrariaceae_Nitraria_roborowskii
Nitrariaceae_Nitraria_praevisa
Nitrariaceae_Nitraria_tangutorum
Nitrariaceae_Nitraria_sibirica
Nitrariaceae_Nitraria_retusa
Nitrariaceae_Tetradiclis_tenella
Nitrariaceae_Malacocarpus_crithmifolius
Nitrariaceae_Nitraria_sphaerocarpa
Nitrariaceae_Peganum_harmala
Biebersteiniaceae_Biebersteinia_heterostemon
Biebersteiniaceae_Biebersteinia_odora
Biebersteiniaceae_Biebersteinia_orphanidis
Biebersteiniaceae_Biebersteinia_multifida
Sapindaceae_Ungnadia_speciosa
Sapindaceae_Stadmannia_oppositifolia
Sapindaceae_Koelreuteria_bipinnata
Sapindaceae_Koelreuteria_elegans
Sapindaceae_Koelreuteria_paniculata
Sapindaceae_Smelophyllum_capense
Sapindaceae_Schleichera_oleosa
Sapindaceae_Paranephelium_xestophyllum
Sapindaceae_Paranephelium_macrophyllum
Sapindaceae_Eurycorymbus_cavaleriei
Sapindaceae_Chouxia_macrophylla
Sapindaceae_Conchopetalum_brachysepalum
Sapindaceae_Macphersonia_gracilis
Sapindaceae_Macphersonia_chapelieri
Sapindaceae_Beguea_apetala
Sapindaceae_Haplocoelum_foliolosum
Sapindaceae_Alectryon_subcinereus
Sapindaceae_Glenniea_pervillei
Sapindaceae_Lecaniodiscus_fraxinifolia
Sapindaceae_Haplocoelopsis_africana
Sapindaceae_Laccodiscus_klaineanus
Sapindaceae_Pancovia_golungensis
Sapindaceae_Chytranthus_prieurianus
Sapindaceae_Lecaniodiscus_cupanioides
71
45
100
96
86
100
99
100
98
95
67
58
22
64
99
74
100
78
76
87
100
61
88
100
81
19
66
26
71
68
66
33
54
12
100

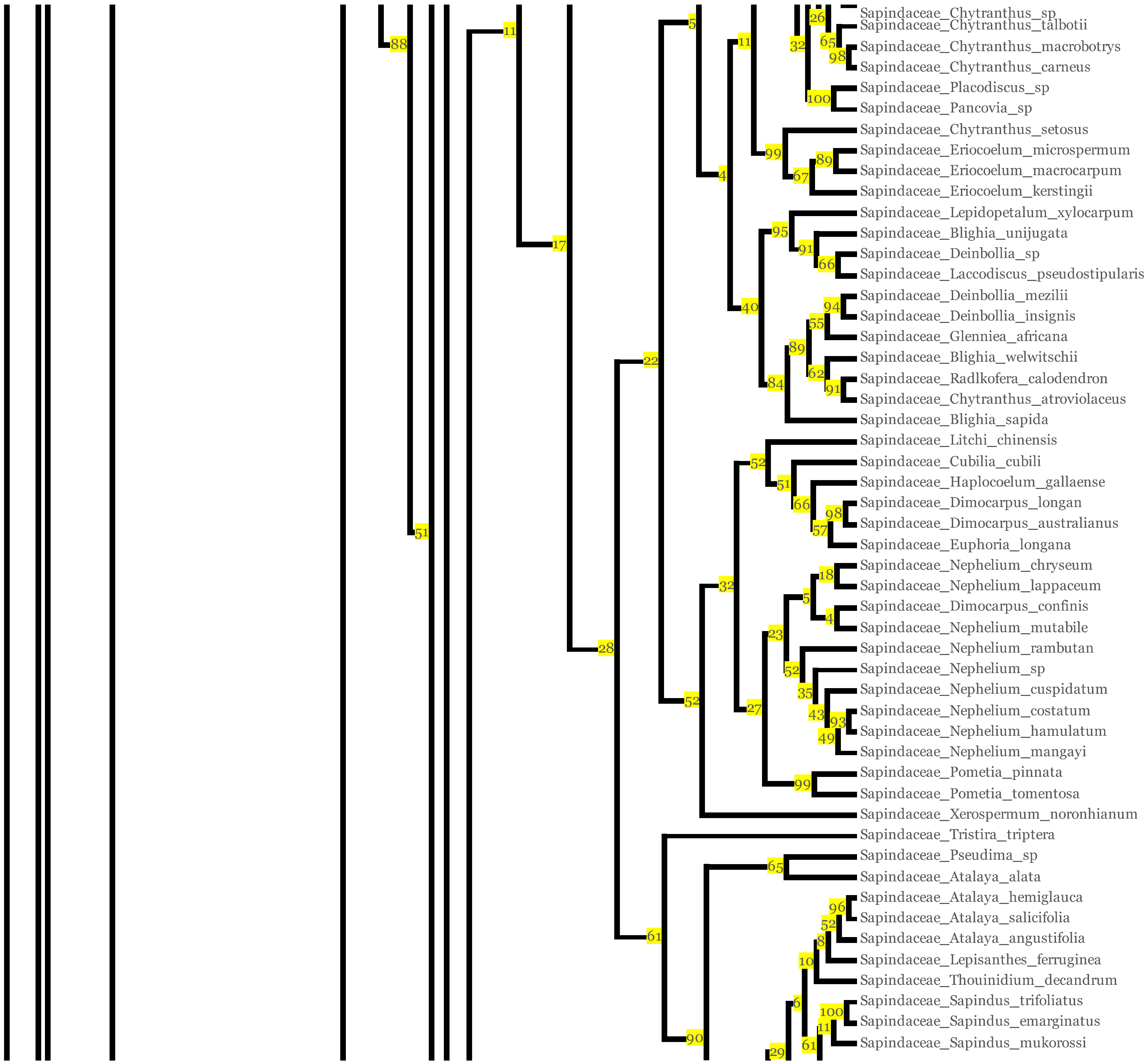
Sapindaceae_Chytranthus_sp
Sapindaceae_Chytranthus_talbotii
Sapindaceae_Chytranthus_macrobotrys
Sapindaceae_Chytranthus_carneus
Sapindaceae_Placodiscus_sp
Sapindaceae_Pancovia_sp
Sapindaceae_Chytranthus_setosus
Sapindaceae_Eriocoelum_microspermum
Sapindaceae_Eriocoelum_macrocarpum
Sapindaceae_Eriocoelum_kerstingii
Sapindaceae_Lepidopetalum_xylocarpum
Sapindaceae_Blighia_unijugata
Sapindaceae_Deinbollia_sp
Sapindaceae_Laccodiscus_pseudostipularis
Sapindaceae_Deinbollia_mezilii
Sapindaceae_Deinbollia_insignis
Sapindaceae_Glenniea_africana
Sapindaceae_Blighia_welwitschii
Sapindaceae_Radlkofera_calodendron
Sapindaceae_Chytranthus_atroviolaceus
Sapindaceae_Blighia_sapida
Sapindaceae_Litchi_chinensis
Sapindaceae_Cubilia_cubili
Sapindaceae_Haplocoelum_gallaense
Sapindaceae_Dimocarpus_longan
Sapindaceae_Dimocarpus_australianus
Sapindaceae_Euphoria_longana
Sapindaceae_Nephelium_chryseum
Sapindaceae_Nephelium_lappaceum
Sapindaceae_Dimocarpus_confinis
Sapindaceae_Nephelium_mutabile
Sapindaceae_Nephelium_rambutan
Sapindaceae_Nephelium_sp
Sapindaceae_Nephelium_cuspidatum
Sapindaceae_Nephelium_costatum
Sapindaceae_Nephelium_hamulatum
Sapindaceae_Nephelium_mangayi
Sapindaceae_Pometia_pinnata
Sapindaceae_Pometia_tomentosa
Sapindaceae_Xerospermum_noronhianum
Sapindaceae_Tristira_triptera
Sapindaceae_Pseudima_sp
Sapindaceae_Atalaya_alata
Sapindaceae_Atalaya_hemiglauca
Sapindaceae_Atalaya_salicifolia
Sapindaceae_Atalaya_angustifolia
Sapindaceae_Lepisanthes_ferruginea
Sapindaceae_Thouinidium_decandrum
Sapindaceae_Sapindus_trifoliatus
Sapindaceae_Sapindus_emarginatus
Sapindaceae_Sapindus_mukorossi

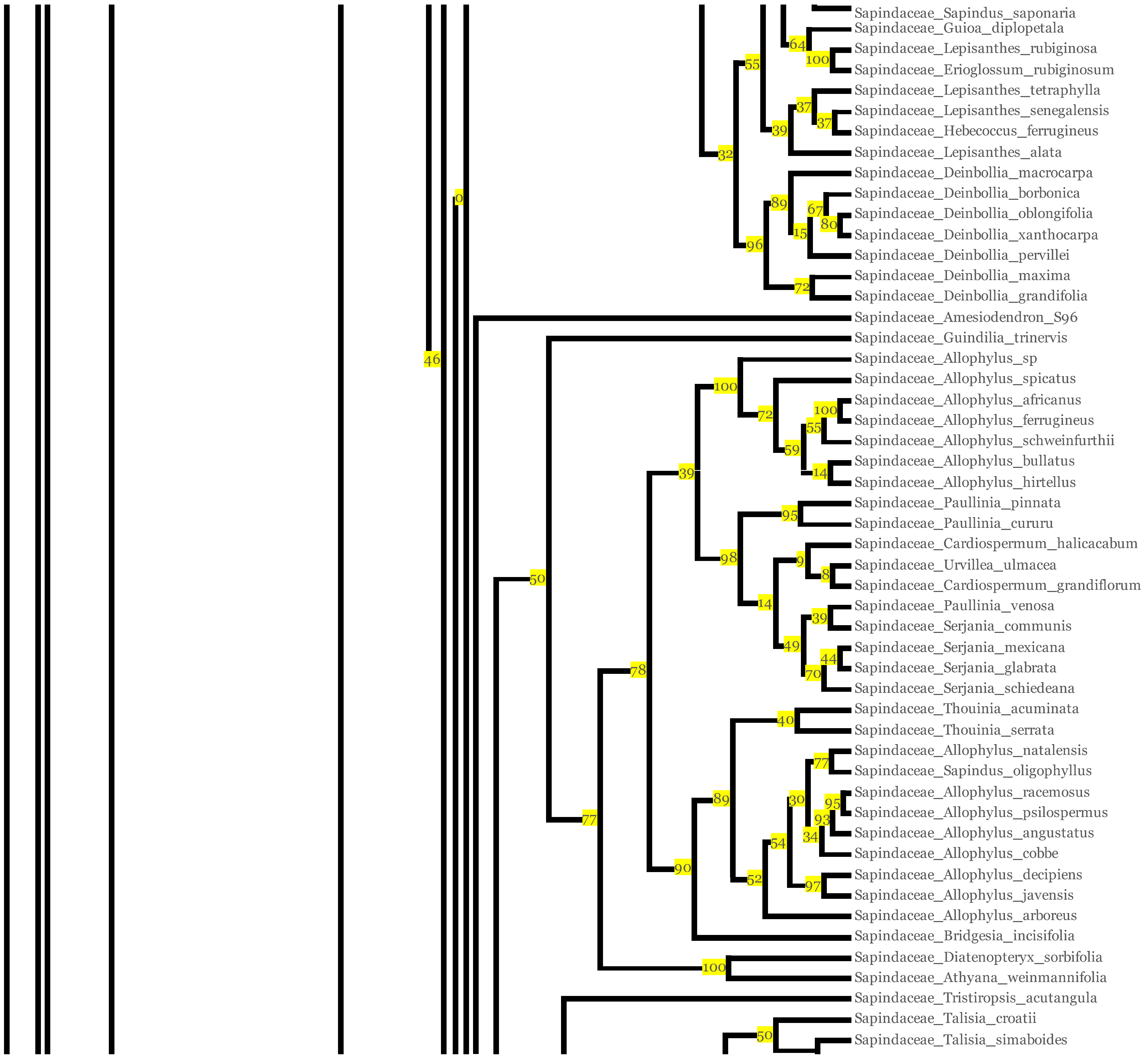
Sapindaceae_Sapindus_saponaria
Sapindaceae_Guioa_diplopetala
Sapindaceae_Lepisanthes_rubiginosa
Sapindaceae_Erioglossum_rubiginosum
Sapindaceae_Lepisanthes_tetraphylla
Sapindaceae_Lepisanthes_senegalensis
Sapindaceae_Hebecoccus_ferrugineus
Sapindaceae_Lepisanthes_alata
Sapindaceae_Deinbollia_macrocarpa
Sapindaceae_Deinbollia_borbonica
Sapindaceae_Deinbollia_oblongifolia
Sapindaceae_Deinbollia_xanthocarpa
Sapindaceae_Deinbollia_pervillei
Sapindaceae_Deinbollia_maxima
Sapindaceae_Deinbollia_grandifolia
Sapindaceae_Amesiodendron_S96
Sapindaceae_Guindilia_trinervis
Sapindaceae_Allophylus_sp
Sapindaceae_Allophylus_spicatus
Sapindaceae_Allophylus_africanus
Sapindaceae_Allophylus_ferrugineus
Sapindaceae_Allophylus_schweinfurthii
Sapindaceae_Allophylus_bullatus
Sapindaceae_Allophylus_hirtellus
Sapindaceae_Paullinia_pinnata
Sapindaceae_Paullinia_cururu
Sapindaceae_Cardiospermum_halicacabum
Sapindaceae_Urvillea_ulmacea
Sapindaceae_Cardiospermum_grandiflorum
Sapindaceae_Paullinia_venosa
Sapindaceae_Serjania_communis
Sapindaceae_Serjania_mexicana
Sapindaceae_Serjania_glabrata
Sapindaceae_Serjania_schiedeana
Sapindaceae_Thouinia_acuminata
Sapindaceae_Thouinia_serrata
Sapindaceae_Allophylus_natalensis
Sapindaceae_Sapindus_oligophyllus
Sapindaceae_Allophylus_racemosus
Sapindaceae_Allophylus_psilospermus
Sapindaceae_Allophylus_angustatus
Sapindaceae_Allophylus_cobbe
Sapindaceae_Allophylus_decipiens
Sapindaceae_Allophylus_javensis
Sapindaceae_Allophylus_arboreus
Sapindaceae_Bridgesia_incisifolia
Sapindaceae_Diatenopteryx_sorbifolia
Sapindaceae_Athyana_weinmannifolia
Sapindaceae_Tristiropsis_acutangula
Sapindaceae_Talisia_croatii
Sapindaceae_Talisia_simaboides

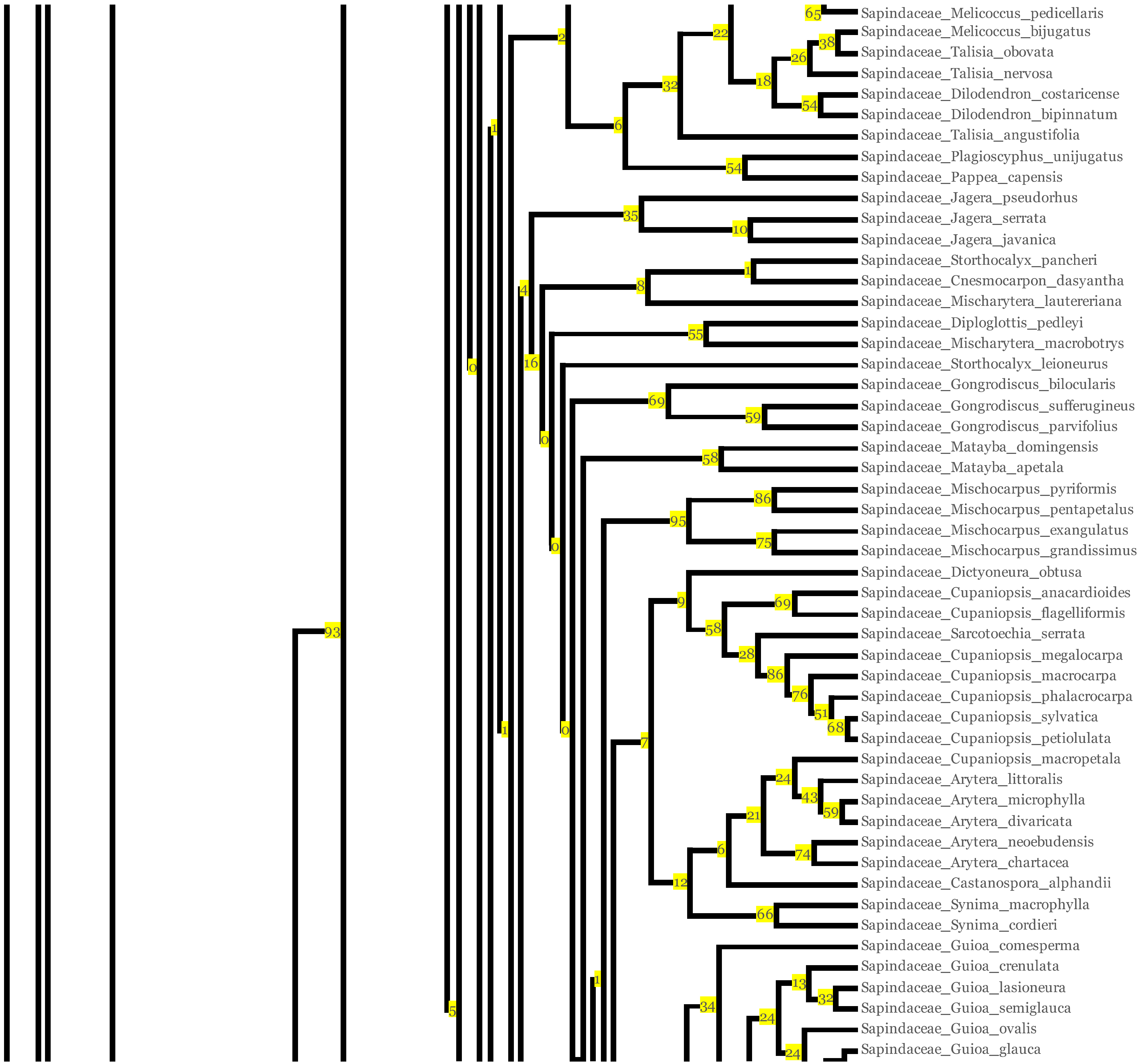
Sapindaceae_Melicoccus_pedicellaris
Sapindaceae_Melicoccus_bijugatus
Sapindaceae_Talisia_obovata
Sapindaceae_Talisia_nervosa
Sapindaceae_Dilodendron_costaricense
Sapindaceae_Dilodendron_bipinnatum
Sapindaceae_Talisia_angustifolia
Sapindaceae_Plagioscyphus_unijugatus
Sapindaceae_Pappea_capensis
Sapindaceae_Jagera_pseudorhus
Sapindaceae_Jagera_serrata
Sapindaceae_Jagera_javanica
Sapindaceae_Storthocalyx_pancheri
Sapindaceae_Cnesmocarpon_dasyantha
Sapindaceae_Mischarytera_lautereriana
Sapindaceae_Diploglottis_pedleyi
Sapindaceae_Mischarytera_macrobotrys
Sapindaceae_Storthocalyx_leioneurus
Sapindaceae_Gongrodiscus_bilocularis
Sapindaceae_Gongrodiscus_sufferugineus
Sapindaceae_Gongrodiscus_parvifolius
Sapindaceae_Matayba_domingensis
Sapindaceae_Matayba_apetala
Sapindaceae_Mischocarpus_pyriformis
Sapindaceae_Mischocarpus_pentapetalus
Sapindaceae_Mischocarpus_exangulatus
Sapindaceae_Mischocarpus_grandissimus
Sapindaceae_Dictyoneura_obtusa
Sapindaceae_Cupaniopsis_anacardioides
Sapindaceae_Cupaniopsis_flagelliformis
Sapindaceae_Sarcotoechia_serrata
Sapindaceae_Cupaniopsis_megalocarpa
Sapindaceae_Cupaniopsis_macrocarpa
Sapindaceae_Cupaniopsis_phalacrocarpa
Sapindaceae_Cupaniopsis_sylvatica
Sapindaceae_Cupaniopsis_petiolulata
Sapindaceae_Cupaniopsis_macropetala
Sapindaceae_Arytera_littoralis
Sapindaceae_Arytera_microphylla
Sapindaceae_Arytera_divaricata
Sapindaceae_Arytera_neoebudensis
Sapindaceae_Arytera_chartacea
Sapindaceae_Castanospora_alphandii
Sapindaceae_Synima_macrophylla
Sapindaceae_Synima_cordieri
Sapindaceae_Guioa_comesperma
Sapindaceae_Guioa_crenulata
Sapindaceae_Guioa_lasioneura
Sapindaceae_Guioa_semiglauca
Sapindaceae_Guioa_ovalis
Sapindaceae_Guioa_glauca

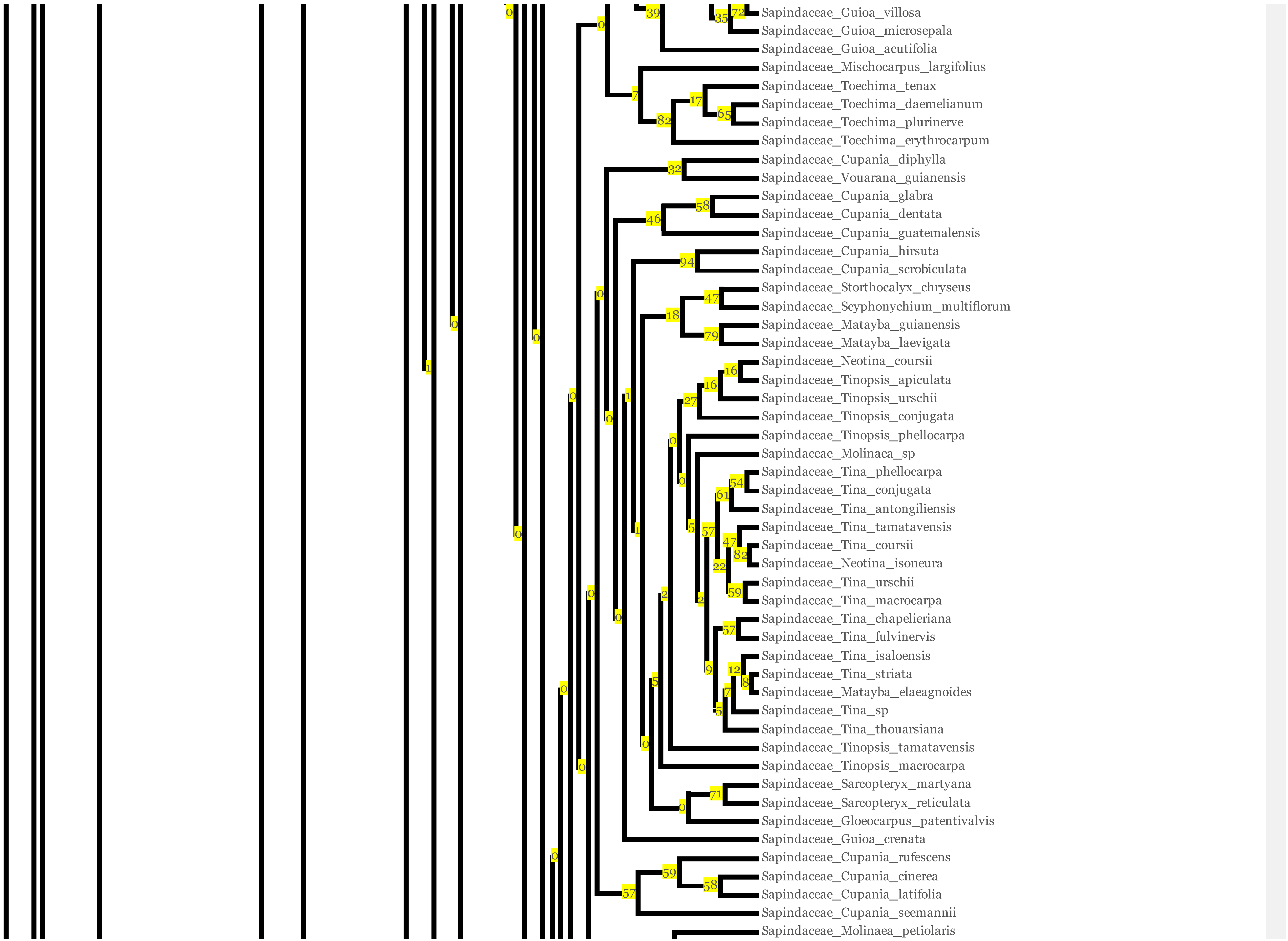
Sapindaceae_Guioa_villosa
Sapindaceae_Guioa_microsepala
Sapindaceae_Guioa_acutifolia
Sapindaceae_Mischocarpus_largifolius
Sapindaceae_Toechima_tenax
Sapindaceae_Toechima_daemelianum
Sapindaceae_Toechima_plurinerve
Sapindaceae_Toechima_erythrocarpum
Sapindaceae_Cupania_diphylla
Sapindaceae_Vouarana_guianensis
Sapindaceae_Cupania_glabra
Sapindaceae_Cupania_dentata
Sapindaceae_Cupania_guatemalensis
Sapindaceae_Cupania_hirsuta
Sapindaceae_Cupania_scrobiculata
Sapindaceae_Storthocalyx_chryseus
Sapindaceae_Scyphonychium_multiflorum
Sapindaceae_Matayba_guianensis
Sapindaceae_Matayba_laevigata
Sapindaceae_Neotina_coursii
Sapindaceae_Tinopsis_apiculata
Sapindaceae_Tinopsis_urschii
Sapindaceae_Tinopsis_conjugata
Sapindaceae_Tinopsis_phellocarpa
Sapindaceae_Molinaea_sp
Sapindaceae_Tina_phellocarpa
Sapindaceae_Tina_conjugata
Sapindaceae_Tina_antongiliensis
Sapindaceae_Tina_tamatavensis
Sapindaceae_Tina_coursii
Sapindaceae_Neotina_isoneura
Sapindaceae_Tina_urschii
Sapindaceae_Tina_macrocarpa
Sapindaceae_Tina_chapelieriana
Sapindaceae_Tina_fulvinervis
Sapindaceae_Tina_isaloensis
Sapindaceae_Tina_striata
Sapindaceae_Matayba_elaeagnoides
Sapindaceae_Tina_sp
Sapindaceae_Tina_thouarsiana
Sapindaceae_Tinopsis_tamatavensis
Sapindaceae_Tinopsis_macrocarpa
Sapindaceae_Sarcopteryx_martyana
Sapindaceae_Sarcopteryx_reticulata
Sapindaceae_Gloeocarpus_patentivalvis
Sapindaceae_Guioa_crenata
Sapindaceae_Cupania_rufescens
Sapindaceae_Cupania_cinerea
Sapindaceae_Cupania_latifolia
Sapindaceae_Cupania_seemannii
Sapindaceae_Molinaea_petiolaris

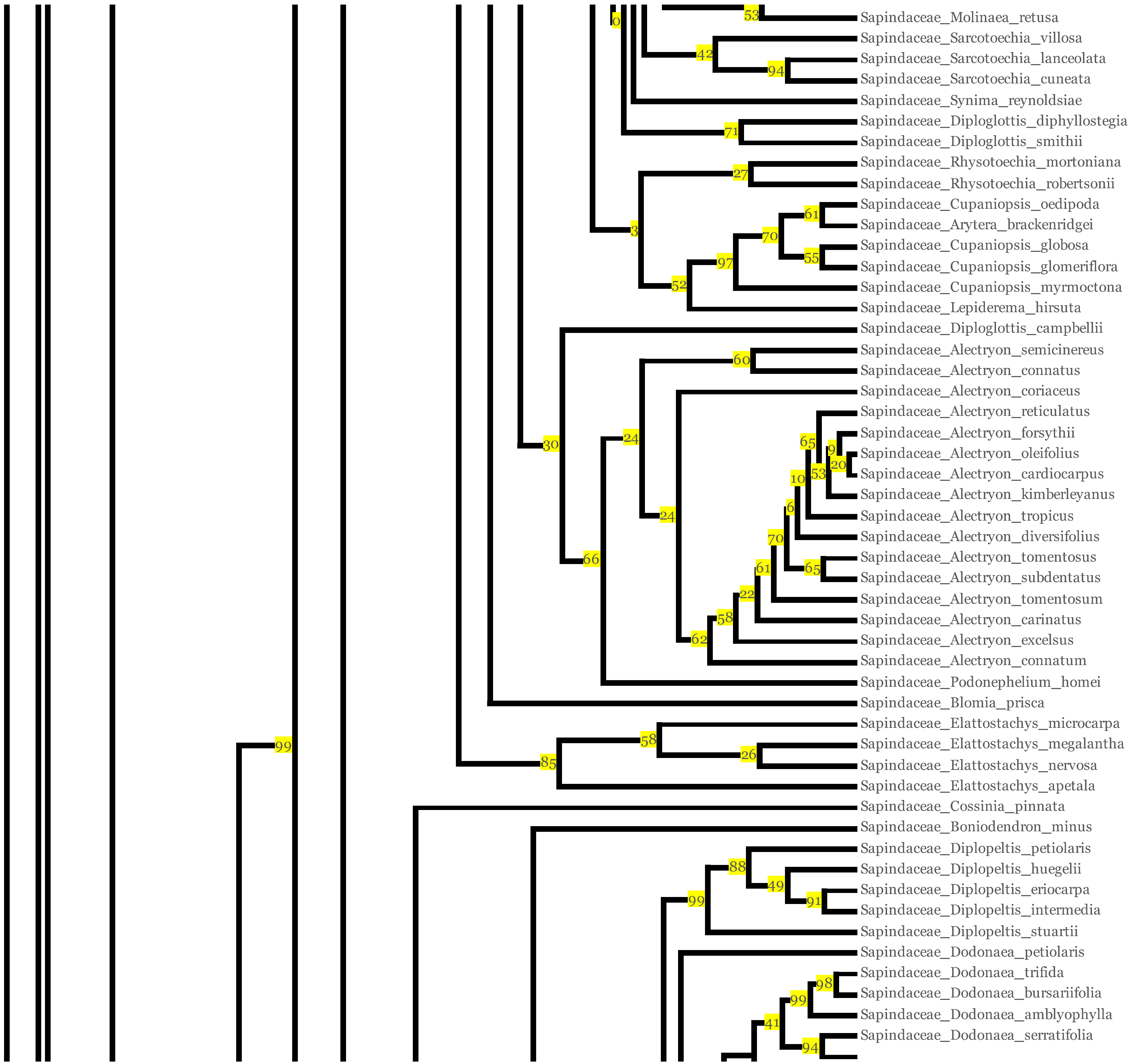
Sapindaceae_Molinaea_retusa
Sapindaceae_Sarcotoechia_villosa
Sapindaceae_Sarcotoechia_lanceolata
Sapindaceae_Sarcotoechia_cuneata
Sapindaceae_Synima_reynoldsiae
Sapindaceae_Diploglottis_diphyllostegia
Sapindaceae_Diploglottis_smithii
Sapindaceae_Rhysotoechia_mortoniana
Sapindaceae_Rhysotoechia_robertsonii
Sapindaceae_Cupaniopsis_oedipoda
Sapindaceae_Arytera_brackenridgei
Sapindaceae_Cupaniopsis_globosa
Sapindaceae_Cupaniopsis_glomeriflora
Sapindaceae_Cupaniopsis_myrmoctona
Sapindaceae_Lepiderema_hirsuta
Sapindaceae_Diploglottis_campbellii
Sapindaceae_Alectryon_semicinereus
Sapindaceae_Alectryon_connatus
Sapindaceae_Alectryon_coriaceus
Sapindaceae_Alectryon_reticulatus
Sapindaceae_Alectryon_forsythii
Sapindaceae_Alectryon_oleifolius
Sapindaceae_Alectryon_cardiocarpus
Sapindaceae_Alectryon_kimberleyanus
Sapindaceae_Alectryon_tropicus
Sapindaceae_Alectryon_diversifolius
Sapindaceae_Alectryon_tomentosus
Sapindaceae_Alectryon_subdentatus
Sapindaceae_Alectryon_tomentosum
Sapindaceae_Alectryon_carinatus
Sapindaceae_Alectryon_excelsus
Sapindaceae_Alectryon_connatum
Sapindaceae_Podonephelium_homei
Sapindaceae_Blomia_prisca
Sapindaceae_Elattostachys_microcarpa
Sapindaceae_Elattostachys_megalantha
Sapindaceae_Elattostachys_nervosa
Sapindaceae_Elattostachys_apetala
Sapindaceae_Cossinia_pinnata
Sapindaceae_Boniodendron_minus
Sapindaceae_Diplopeltis_petiolaris
Sapindaceae_Diplopeltis_huegelii
Sapindaceae_Diplopeltis_eriocarpa
Sapindaceae_Diplopeltis_intermedia
Sapindaceae_Diplopeltis_stuartii
Sapindaceae_Dodonaea_petiolaris
Sapindaceae_Dodonaea_trifida
Sapindaceae_Dodonaea_bursariifolia
Sapindaceae_Dodonaea_amblyophylla
Sapindaceae_Dodonaea_serratifolia

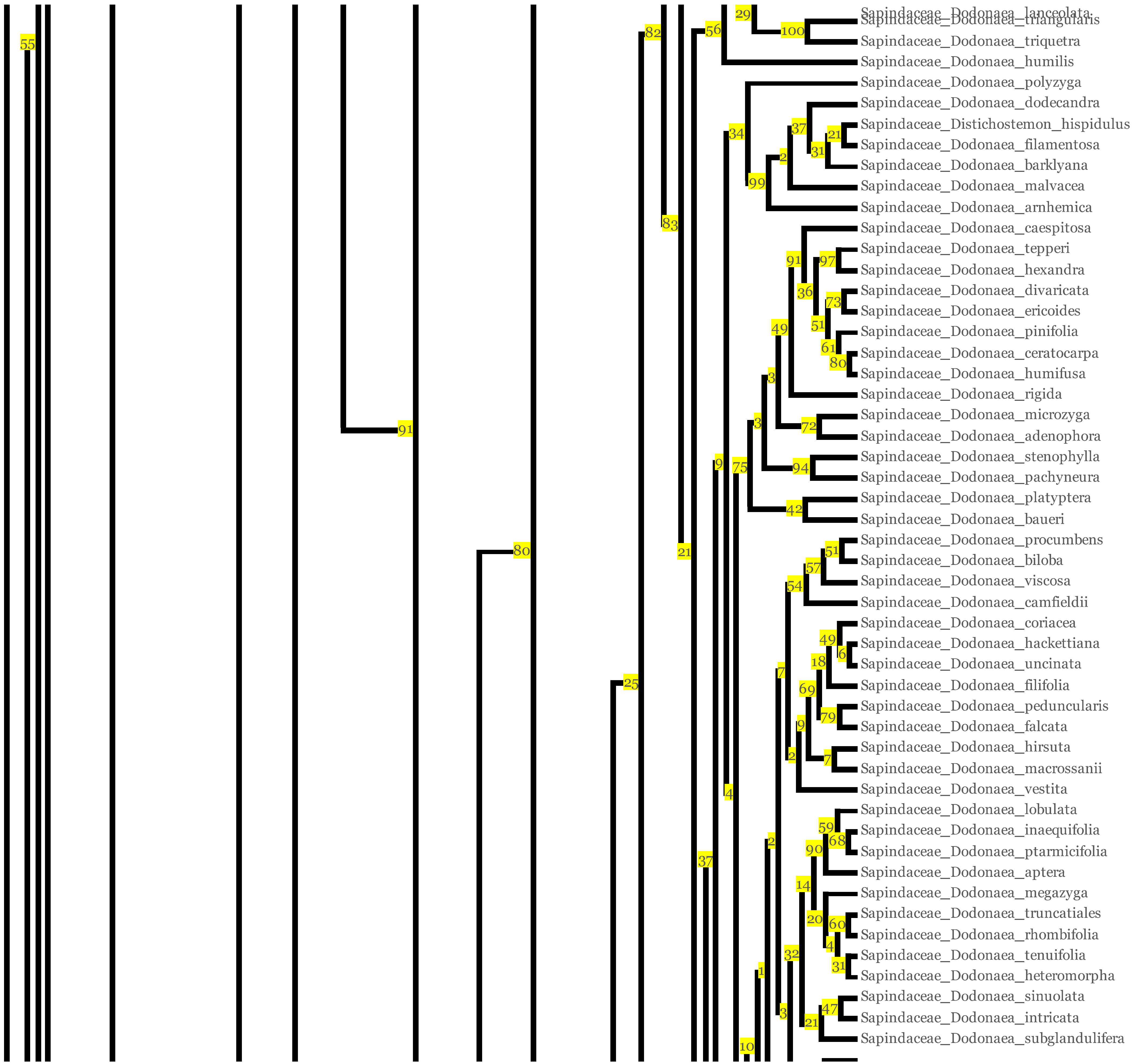
Sapindaceae_Dodonaea_lanceolata
Sapindaceae_Dodonaea_triangularis
Sapindaceae_Dodonaea_triquetra
Sapindaceae_Dodonaea_humilis
Sapindaceae_Dodonaea_polyzyga
Sapindaceae_Dodonaea_dodecandra
Sapindaceae_Distichostemon_hispidulus
Sapindaceae_Dodonaea_filamentosa
Sapindaceae_Dodonaea_barklyana
Sapindaceae_Dodonaea_malvacea
Sapindaceae_Dodonaea_arnhemica
Sapindaceae_Dodonaea_caespitosa
Sapindaceae_Dodonaea_tepperi
Sapindaceae_Dodonaea_hexandra
Sapindaceae_Dodonaea_divaricata
Sapindaceae_Dodonaea_ericoides
Sapindaceae_Dodonaea_pinifolia
Sapindaceae_Dodonaea_ceratocarpa
Sapindaceae_Dodonaea_humifusa
Sapindaceae_Dodonaea_rigida
Sapindaceae_Dodonaea_microzyga
Sapindaceae_Dodonaea_adenophora
Sapindaceae_Dodonaea_stenophylla
Sapindaceae_Dodonaea_pachyneura
Sapindaceae_Dodonaea_platyptera
Sapindaceae_Dodonaea_baueri
Sapindaceae_Dodonaea_procumbens
Sapindaceae_Dodonaea_biloba
Sapindaceae_Dodonaea_viscosa
Sapindaceae_Dodonaea_camfieldii
Sapindaceae_Dodonaea_coriacea
Sapindaceae_Dodonaea_hackettiana
Sapindaceae_Dodonaea_uncinata
Sapindaceae_Dodonaea_filifolia
Sapindaceae_Dodonaea_peduncularis
Sapindaceae_Dodonaea_falcata
Sapindaceae_Dodonaea_hirsuta
Sapindaceae_Dodonaea_macrossanii
Sapindaceae_Dodonaea_vestita
Sapindaceae_Dodonaea_lobulata
Sapindaceae_Dodonaea_inaequifolia
Sapindaceae_Dodonaea_ptarmicifolia
Sapindaceae_Dodonaea_aptera
Sapindaceae_Dodonaea_megazyga
Sapindaceae_Dodonaea_truncatiales
Sapindaceae_Dodonaea_rhombifolia
Sapindaceae_Dodonaea_tenuifolia
Sapindaceae_Dodonaea_heteromorpha
Sapindaceae_Dodonaea_sinuolata
Sapindaceae_Dodonaea_intricata
Sapindaceae_Dodonaea_subglandulifera

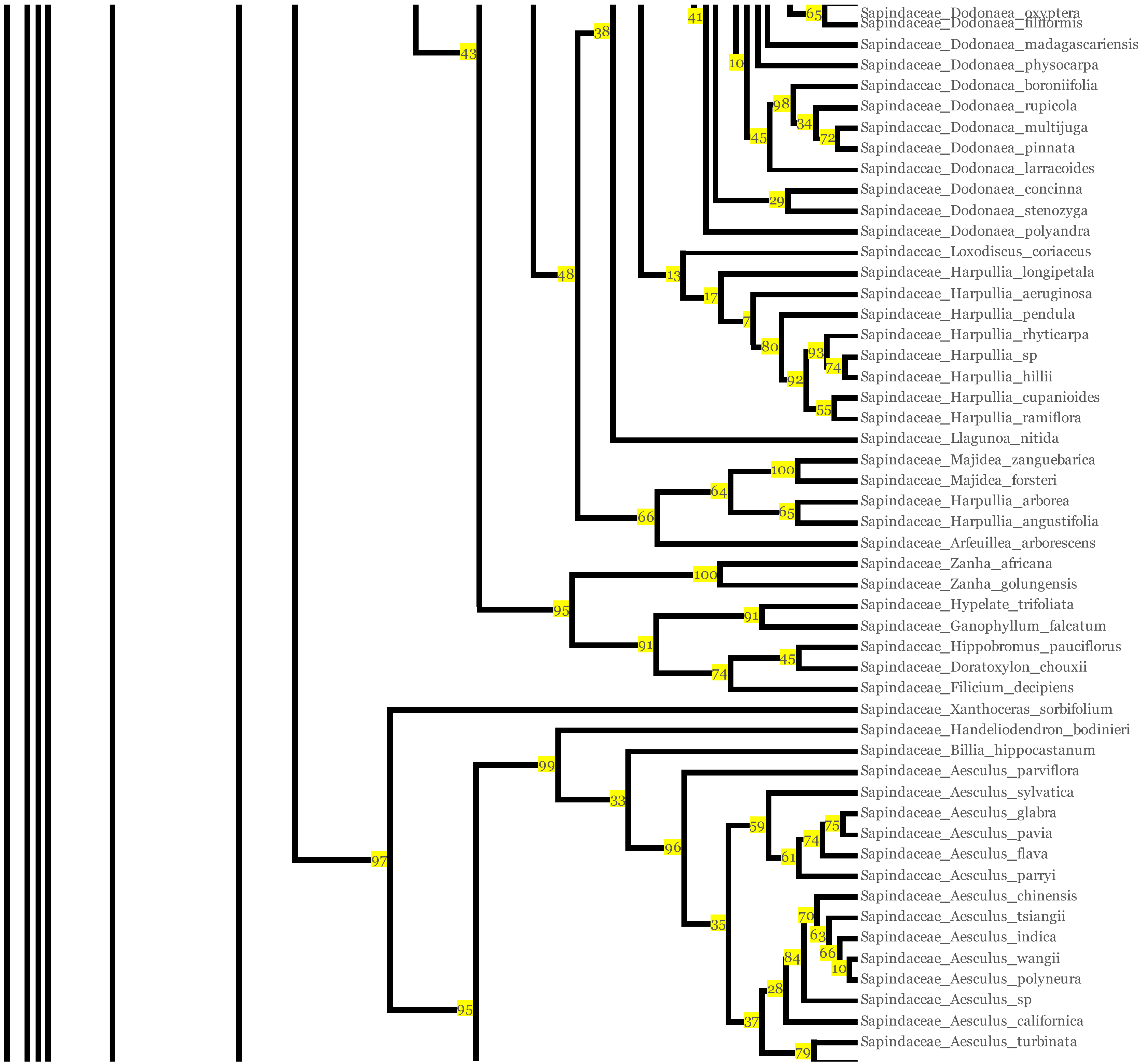
Sapindaceae_Dodonaea_oxyptera
Sapindaceae_Dodonaea_filiformis
Sapindaceae_Dodonaea_madagascariensis
Sapindaceae_Dodonaea_physocarpa
Sapindaceae_Dodonaea_boroniifolia
Sapindaceae_Dodonaea_rupicola
Sapindaceae_Dodonaea_multijuga
Sapindaceae_Dodonaea_pinnata
Sapindaceae_Dodonaea_larraeoides
Sapindaceae_Dodonaea_concinna
Sapindaceae_Dodonaea_stenozyga
Sapindaceae_Dodonaea_polyandra
Sapindaceae_Loxodiscus_coriaceus
Sapindaceae_Harpullia_longipetala
Sapindaceae_Harpullia_aeruginosa
Sapindaceae_Harpullia_pendula
Sapindaceae_Harpullia_rhyticarpa
Sapindaceae_Harpullia_sp
Sapindaceae_Harpullia_hillii
Sapindaceae_Harpullia_cupanioides
Sapindaceae_Harpullia_ramiflora
Sapindaceae_Llagunoa_nitida
Sapindaceae_Majidea_zanguebarica
Sapindaceae_Majidea_forsteri
Sapindaceae_Harpullia_arborea
Sapindaceae_Harpullia_angustifolia
Sapindaceae_Arfeuillea_arborescens
Sapindaceae_Zanha_africana
Sapindaceae_Zanha_golungensis
Sapindaceae_Hypelate_trifoliata
Sapindaceae_Ganophyllum_falcatum
Sapindaceae_Hippobromus_pauciflorus
Sapindaceae_Doratoxylon_chouxii
Sapindaceae_Filicium_decipiens
Sapindaceae_Xanthoceras_sorbifolium
Sapindaceae_Handeliodendron_bodinieri
Sapindaceae_Billia_hippocastanum
Sapindaceae_Aesculus_parviflora
Sapindaceae_Aesculus_sylvatica
Sapindaceae_Aesculus_glabra
Sapindaceae_Aesculus_pavia
Sapindaceae_Aesculus_flava
Sapindaceae_Aesculus_parryi
Sapindaceae_Aesculus_chinensis
Sapindaceae_Aesculus_tsiangii
Sapindaceae_Aesculus_indica
Sapindaceae_Aesculus_wangii
Sapindaceae_Aesculus_polyneura
Sapindaceae_Aesculus_sp
Sapindaceae_Aesculus_californica
Sapindaceae_Aesculus_turbinata

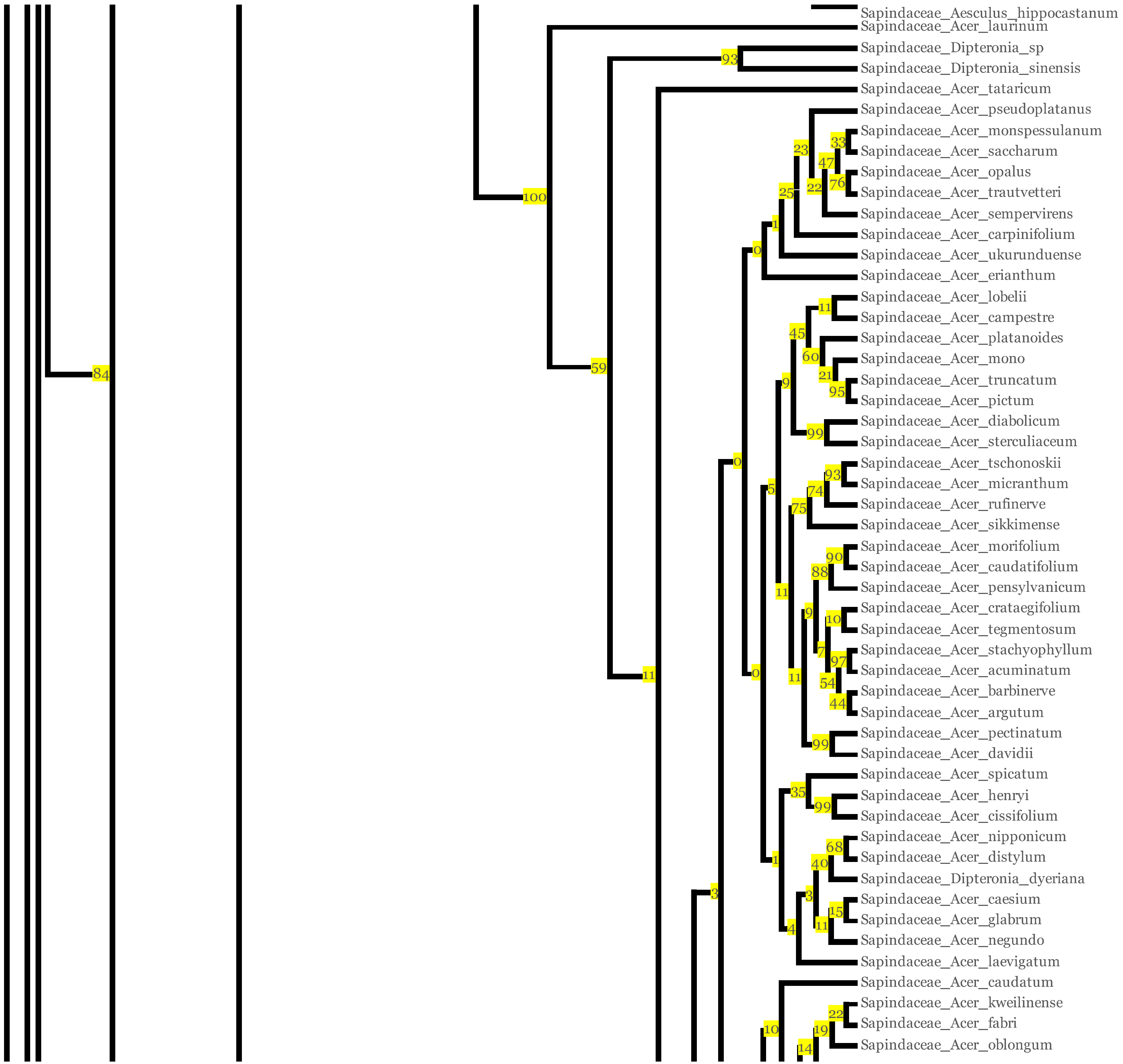
Sapindaceae_Aesculus_hippocastanum
Sapindaceae_Acer_laurinum
Sapindaceae_Dipteronia_sp
Sapindaceae_Dipteronia_sinensis
Sapindaceae_Acer_tataricum
Sapindaceae_Acer_pseudoplatanus
Sapindaceae_Acer_monspessulanum
Sapindaceae_Acer_saccharum
Sapindaceae_Acer_opalus
Sapindaceae_Acer_trautvetteri
Sapindaceae_Acer_sempervirens
Sapindaceae_Acer_carpinifolium
Sapindaceae_Acer_ukurunduense
Sapindaceae_Acer_erianthum
Sapindaceae_Acer_lobelii
Sapindaceae_Acer_campestre
Sapindaceae_Acer_platanoides
Sapindaceae_Acer_mono
Sapindaceae_Acer_truncatum
Sapindaceae_Acer_pictum
Sapindaceae_Acer_diabolicum
Sapindaceae_Acer_sterculiaceum
Sapindaceae_Acer_tschonoskii
Sapindaceae_Acer_micranthum
Sapindaceae_Acer_rufinerve
Sapindaceae_Acer_sikkimense
Sapindaceae_Acer_morifolium
Sapindaceae_Acer_caudatifolium
Sapindaceae_Acer_pensylvanicum
Sapindaceae_Acer_crataegifolium
Sapindaceae_Acer_tegmentosum
Sapindaceae_Acer_stachyophyllum
Sapindaceae_Acer_acuminatum
Sapindaceae_Acer_barbinerve
Sapindaceae_Acer_argutum
Sapindaceae_Acer_pectinatum
Sapindaceae_Acer_davidii
Sapindaceae_Acer_spicatum
Sapindaceae_Acer_henryi
Sapindaceae_Acer_cissifolium
Sapindaceae_Acer_nipponicum
Sapindaceae_Acer_distylum
Sapindaceae_Dipteronia_dyeriana
Sapindaceae_Acer_caesium
Sapindaceae_Acer_glabrum
Sapindaceae_Acer_negundo
Sapindaceae_Acer_laevigatum
Sapindaceae_Acer_caudatum
Sapindaceae_Acer_kweilinense
Sapindaceae_Acer_fabri
Sapindaceae_Acer_oblongum

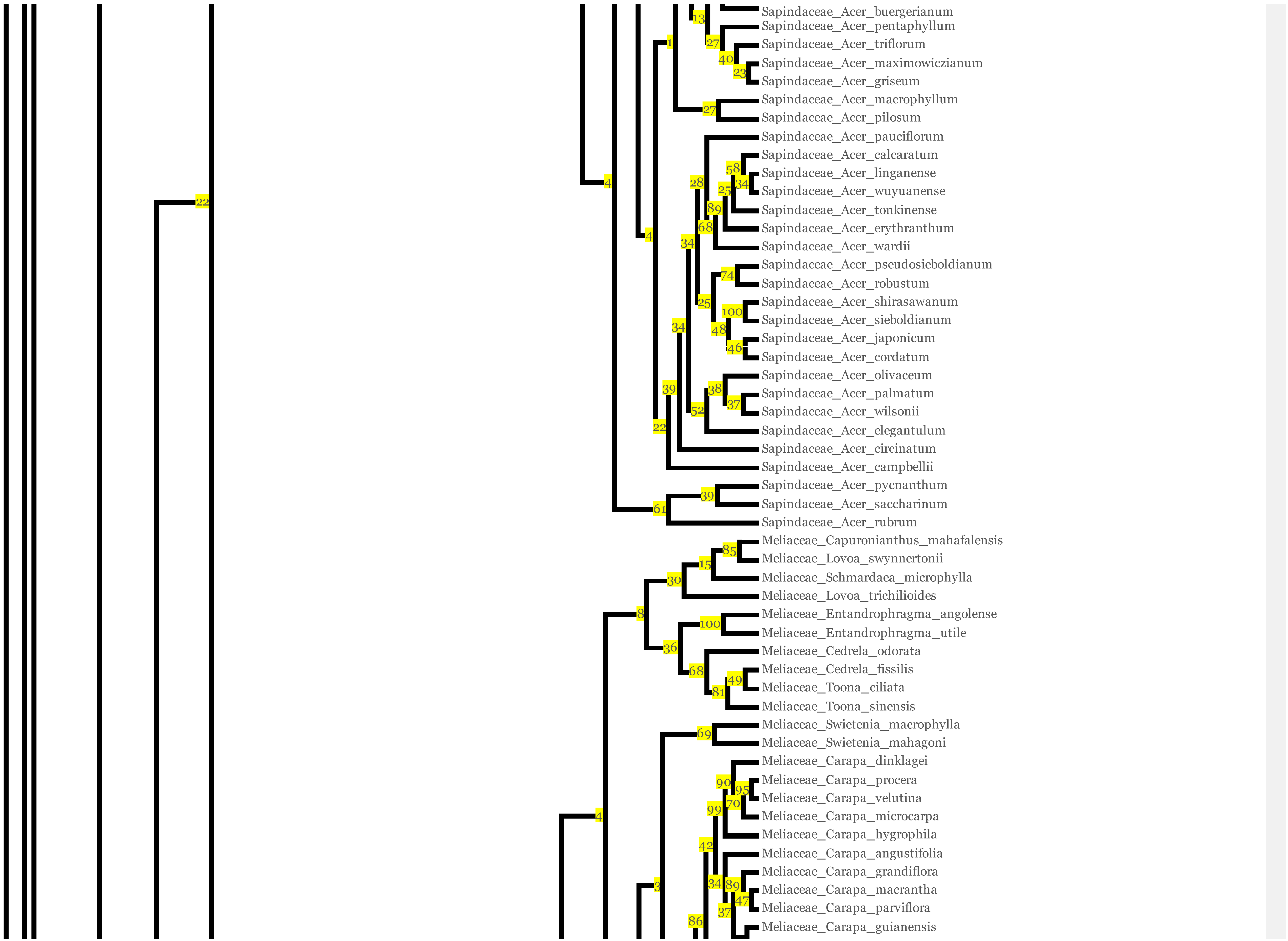
Sapindaceae_Acer_buergerianum
Sapindaceae_Acer_pentaphyllum
Sapindaceae_Acer_triflorum
Sapindaceae_Acer_maximowiczianum
Sapindaceae_Acer_griseum
Sapindaceae_Acer_macrophyllum
Sapindaceae_Acer_pilosum
Sapindaceae_Acer_pauciflorum
Sapindaceae_Acer_calcaratum
Sapindaceae_Acer_linganense
Sapindaceae_Acer_wuyuanense
Sapindaceae_Acer_tonkinense
Sapindaceae_Acer_erythranthum
Sapindaceae_Acer_wardii
Sapindaceae_Acer_pseudosieboldianum
Sapindaceae_Acer_robustum
Sapindaceae_Acer_shirasawanum
Sapindaceae_Acer_sieboldianum
Sapindaceae_Acer_japonicum
Sapindaceae_Acer_cordatum
Sapindaceae_Acer_olivaceum
Sapindaceae_Acer_palmatum
Sapindaceae_Acer_wilsonii
Sapindaceae_Acer_elegantulum
Sapindaceae_Acer_circinatum
Sapindaceae_Acer_campbellii
Sapindaceae_Acer_pycnanthum
Sapindaceae_Acer_saccharinum
Sapindaceae_Acer_rubrum
Meliaceae_Capuronianthus_mahafalensis
Meliaceae_Lovoa_swynnertonii
Meliaceae_Schmardaea_microphylla
Meliaceae_Lovoa_trichilioides
Meliaceae_Entandrophragma_angolense
Meliaceae_Entandrophragma_utile
Meliaceae_Cedrela_odorata
Meliaceae_Cedrela_fissilis
Meliaceae_Toona_ciliata
Meliaceae_Toona_sinensis
Meliaceae_Swietenia_macrophylla
Meliaceae_Swietenia_mahagoni
Meliaceae_Carapa_dinklagei
Meliaceae_Carapa_procera
Meliaceae_Carapa_velutina
Meliaceae_Carapa_microcarpa
Meliaceae_Carapa_hygrophila
Meliaceae_Carapa_angustifolia
Meliaceae_Carapa_grandiflora
Meliaceae_Carapa_macrantha
Meliaceae_Carapa_parviflora
Meliaceae_Carapa_guianensis
22
4
13
27
1
40
23
27
58
28
34
25
89
4
68
34
74
25
100
34
48
46
38
39
37
52
22
39
61
85
15
30
8
100
36
68
49
81
69
90
95
70
99
4
42
3
34
89
47
37
86

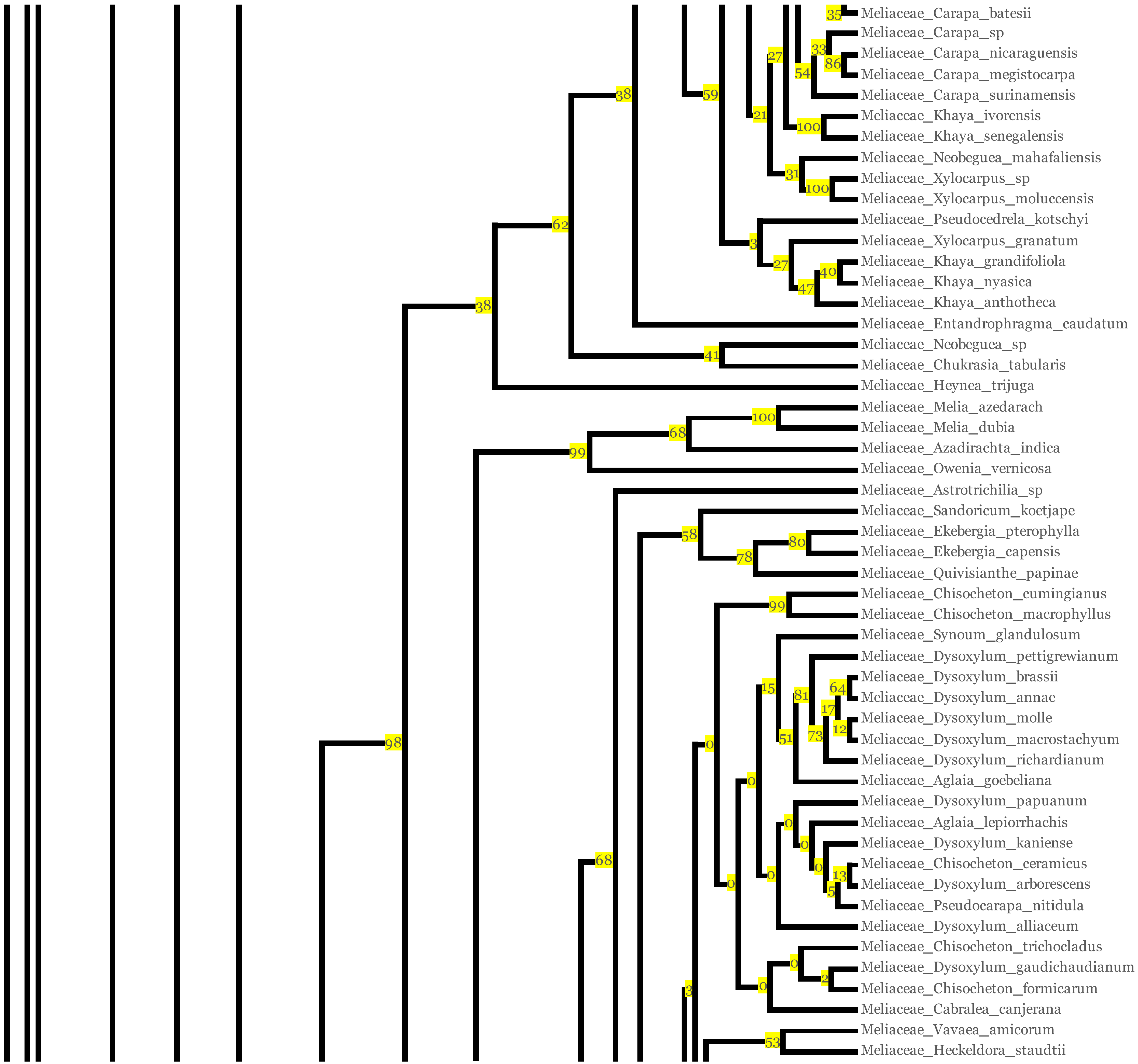
Meliaceae_Carapa_batesii
Meliaceae_Carapa_sp
Meliaceae_Carapa_nicaraguensis
Meliaceae_Carapa_megistocarpa
Meliaceae_Carapa_surinamensis
Meliaceae_Khaya_ivorensis
Meliaceae_Khaya_senegalensis
Meliaceae_Neobeguea_mahafaliensis
Meliaceae_Xylocarpus_sp
Meliaceae_Xylocarpus_moluccensis
Meliaceae_Pseudocedrela_kotschyi
Meliaceae_Xylocarpus_granatum
Meliaceae_Khaya_grandifoliola
Meliaceae_Khaya_nyasica
Meliaceae_Khaya_anthotheca
Meliaceae_Entandrophragma_caudatum
Meliaceae_Neobeguea_sp
Meliaceae_Chukrasia_tabularis
Meliaceae_Heynea_trijuga
Meliaceae_Melia_azedarach
Meliaceae_Melia_dubia
Meliaceae_Azadirachta_indica
Meliaceae_Owenia_vernicosa
Meliaceae_Astrotrichilia_sp
Meliaceae_Sandoricum_koetjape
Meliaceae_Ekebergia_pterophylla
Meliaceae_Ekebergia_capensis
Meliaceae_Quivisianthe_papinae
Meliaceae_Chisocheton_cumingianus
Meliaceae_Chisocheton_macrophyllus
Meliaceae_Synoum_glandulosum
Meliaceae_Dysoxylum_pettigrewianum
Meliaceae_Dysoxylum_brassii
Meliaceae_Dysoxylum_annae
Meliaceae_Dysoxylum_molle
Meliaceae_Dysoxylum_macrostachyum
Meliaceae_Dysoxylum_richardianum
Meliaceae_Aglaia_goebeliana
Meliaceae_Dysoxylum_papuanum
Meliaceae_Aglaia_lepiorrhachis
Meliaceae_Dysoxylum_kaniense
Meliaceae_Chisocheton_ceramicus
Meliaceae_Dysoxylum_arborescens
Meliaceae_Pseudocarapa_nitidula
Meliaceae_Dysoxylum_alliaceum
Meliaceae_Chisocheton_trichocladus
Meliaceae_Dysoxylum_gaudichaudianum
Meliaceae_Chisocheton_formicarum
Meliaceae_Cabralea_canjerana
Meliaceae_Vavaea_amicorum
Meliaceae_Heckeldora_staudtii

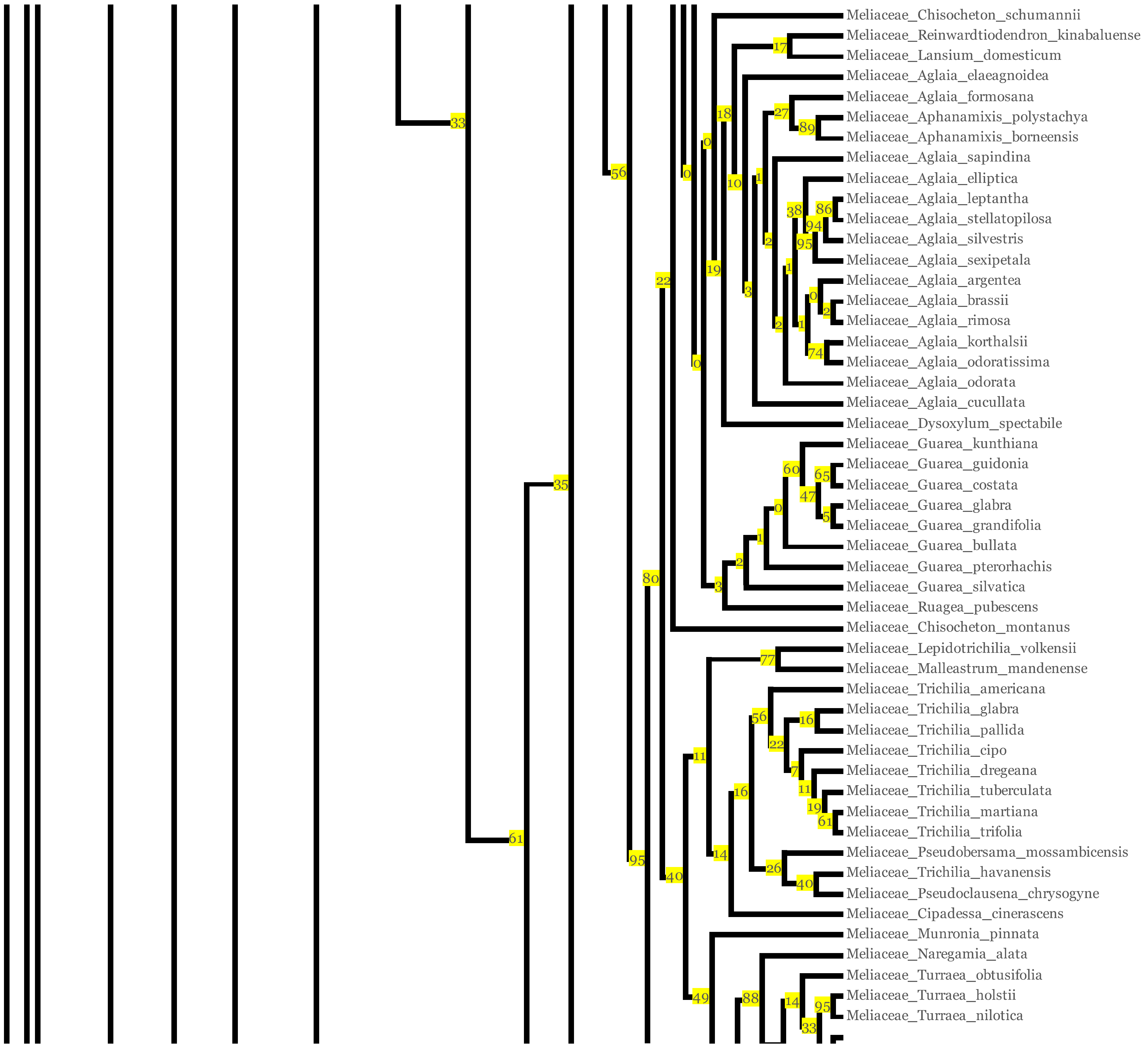
Meliaceae_Chisocheton_schumannii
Meliaceae_Reinwardtiodendron_kinabaluense
Meliaceae_Lansium_domesticum
Meliaceae_Aglaia_elaeagnoidea
Meliaceae_Aglaia_formosana
Meliaceae_Aphanamixis_polystachya
Meliaceae_Aphanamixis_borneensis
Meliaceae_Aglaia_sapindina
Meliaceae_Aglaia_elliptica
Meliaceae_Aglaia_leptantha
Meliaceae_Aglaia_stellatopilosa
Meliaceae_Aglaia_silvestris
Meliaceae_Aglaia_sexipetala
Meliaceae_Aglaia_argentea
Meliaceae_Aglaia_brassii
Meliaceae_Aglaia_rimosa
Meliaceae_Aglaia_korthalsii
Meliaceae_Aglaia_odoratissima
Meliaceae_Aglaia_odorata
Meliaceae_Aglaia_cucullata
Meliaceae_Dysoxylum_spectabile
Meliaceae_Guarea_kunthiana
Meliaceae_Guarea_guidonia
Meliaceae_Guarea_costata
Meliaceae_Guarea_glabra
Meliaceae_Guarea_grandifolia
Meliaceae_Guarea_bullata
Meliaceae_Guarea_pterorhachis
Meliaceae_Guarea_silvatica
Meliaceae_Ruagea_pubescens
Meliaceae_Chisocheton_montanus
Meliaceae_Lepidotrichilia_volkensii
Meliaceae_Malleastrum_mandenense
Meliaceae_Trichilia_americana
Meliaceae_Trichilia_glabra
Meliaceae_Trichilia_pallida
Meliaceae_Trichilia_cipo
Meliaceae_Trichilia_dregeana
Meliaceae_Trichilia_tuberculata
Meliaceae_Trichilia_martiana
Meliaceae_Trichilia_trifolia
Meliaceae_Pseudobersama_mossambicensis
Meliaceae_Trichilia_havanensis
Meliaceae_Pseudoclausena_chrysogyne
Meliaceae_Cipadessa_cinerascens
Meliaceae_Munronia_pinnata
Meliaceae_Naregamia_alata
Meliaceae_Turraea_obtusifolia
Meliaceae_Turraea_holstii
Meliaceae_Turraea_nilotica

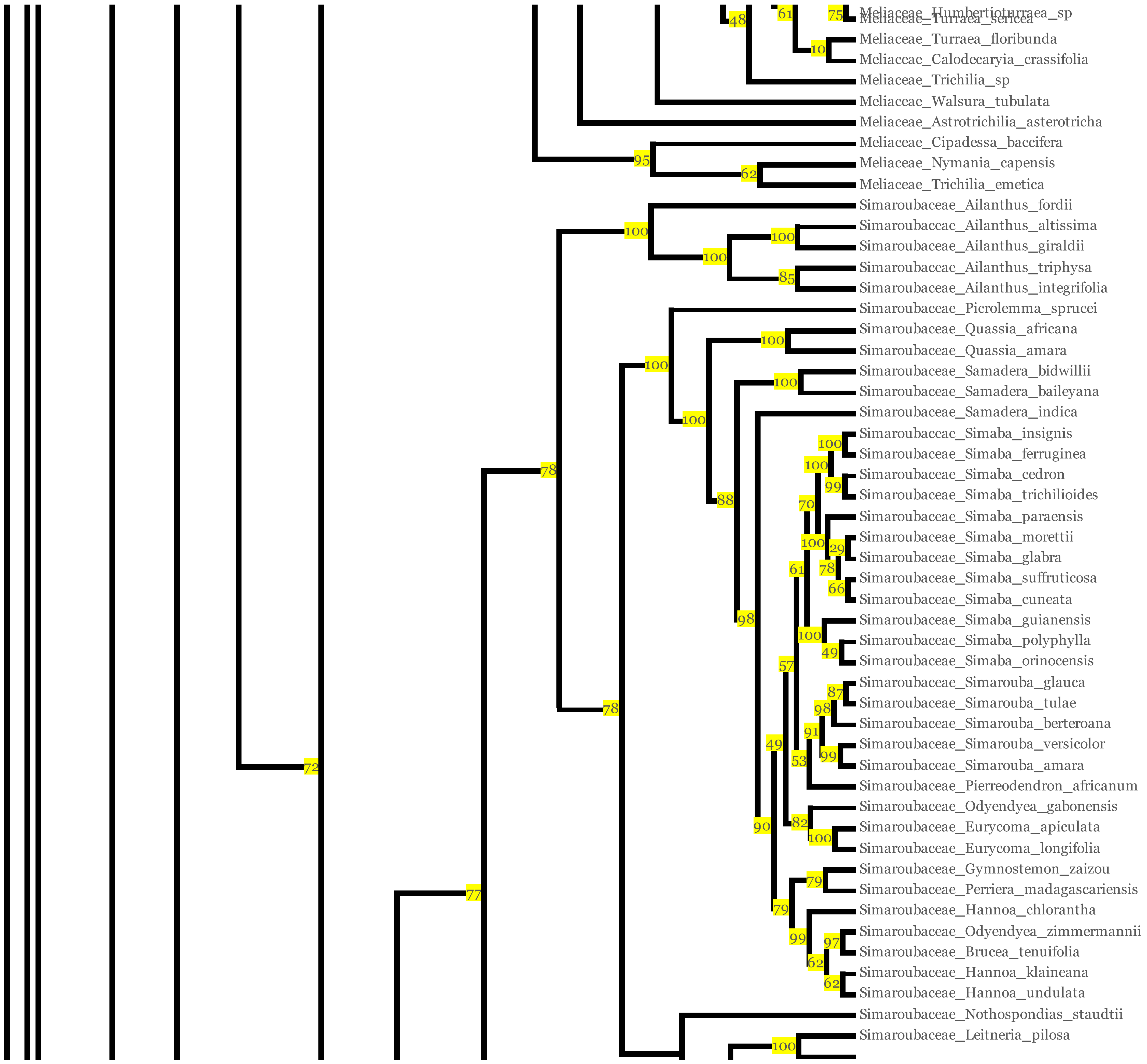
Meliaceae_Humbertioturraea_sp
Meliaceae_Turraea_sericea
Meliaceae_Turraea_floribunda
Meliaceae_Calodecaryia_crassifolia
Meliaceae_Trichilia_sp
Meliaceae_Walsura_tubulata
Meliaceae_Astrotrichilia_asterotricha
Meliaceae_Cipadessa_baccifera
Meliaceae_Nymania_capensis
Meliaceae_Trichilia_emetica
Simaroubaceae_Ailanthus_fordii
Simaroubaceae_Ailanthus_altissima
Simaroubaceae_Ailanthus_giraldii
Simaroubaceae_Ailanthus_triphysa
Simaroubaceae_Ailanthus_integrifolia
Simaroubaceae_Picrolemma_sprucei
Simaroubaceae_Quassia_africana
Simaroubaceae_Quassia_amara
Simaroubaceae_Samadera_bidwillii
Simaroubaceae_Samadera_baileyana
Simaroubaceae_Samadera_indica
Simaroubaceae_Simaba_insignis
Simaroubaceae_Simaba_ferruginea
Simaroubaceae_Simaba_cedron
Simaroubaceae_Simaba_trichilioides
Simaroubaceae_Simaba_paraensis
Simaroubaceae_Simaba_morettii
Simaroubaceae_Simaba_glabra
Simaroubaceae_Simaba_suffruticosa
Simaroubaceae_Simaba_cuneata
Simaroubaceae_Simaba_guianensis
Simaroubaceae_Simaba_polyphylla
Simaroubaceae_Simaba_orinocensis
Simaroubaceae_Simarouba_glauca
Simaroubaceae_Simarouba_tulae
Simaroubaceae_Simarouba_berteroana
Simaroubaceae_Simarouba_versicolor
Simaroubaceae_Simarouba_amara
Simaroubaceae_Pierreodendron_africanum
Simaroubaceae_Odyendyea_gabonensis
Simaroubaceae_Eurycoma_apiculata
Simaroubaceae_Eurycoma_longifolia
Simaroubaceae_Gymnostemon_zaizou
Simaroubaceae_Perriera_madagascariensis
Simaroubaceae_Hannoa_chlorantha
Simaroubaceae_Odyendyea_zimmermannii
Simaroubaceae_Brucea_tenuifolia
Simaroubaceae_Hannoa_klaineana
Simaroubaceae_Hannoa_undulata
Simaroubaceae_Nothospondias_staudtii
Simaroubaceae_Leitneria_pilosa

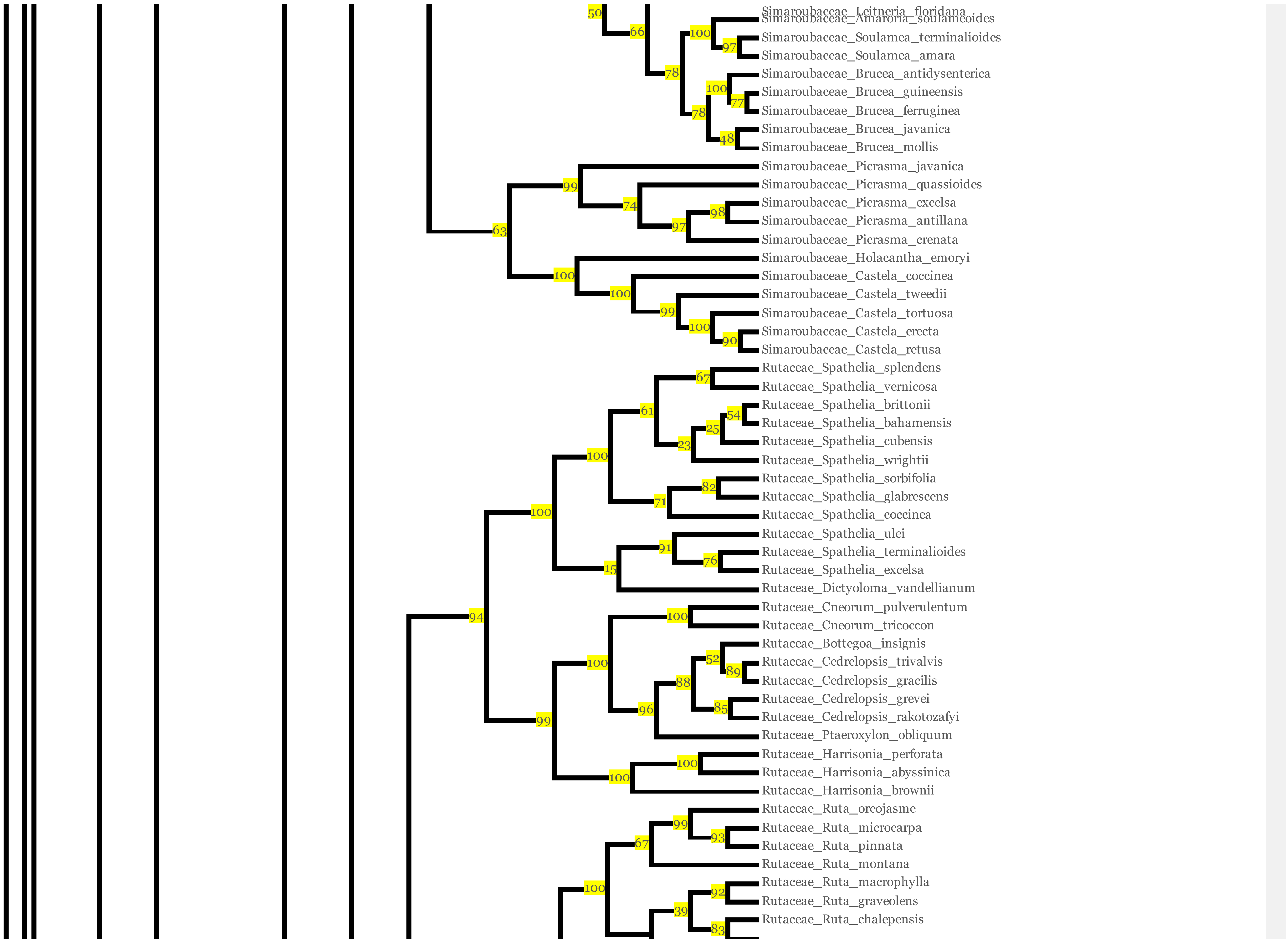
Simaroubaceae_Leitneria_floridana
Simaroubaceae_Amaroria_soulameoides
Simaroubaceae_Soulamea_terminalioides
Simaroubaceae_Soulamea_amara
Simaroubaceae_Brucea_antidysenterica
Simaroubaceae_Brucea_guineensis
Simaroubaceae_Brucea_ferruginea
Simaroubaceae_Brucea_javanica
Simaroubaceae_Brucea_mollis
Simaroubaceae_Picrasma_javanica
Simaroubaceae_Picrasma_quassioides
Simaroubaceae_Picrasma_excelsa
Simaroubaceae_Picrasma_antillana
Simaroubaceae_Picrasma_crenata
Simaroubaceae_Holacantha_emoryi
Simaroubaceae_Castela_coccinea
Simaroubaceae_Castela_tweedii
Simaroubaceae_Castela_tortuosa
Simaroubaceae_Castela_erecta
Simaroubaceae_Castela_retusa
Rutaceae_Spathelia_splendens
Rutaceae_Spathelia_vernicosa
Rutaceae_Spathelia_brittonii
Rutaceae_Spathelia_bahamensis
Rutaceae_Spathelia_cubensis
Rutaceae_Spathelia_wrightii
Rutaceae_Spathelia_sorbifolia
Rutaceae_Spathelia_glabrescens
Rutaceae_Spathelia_coccinea
Rutaceae_Spathelia_ulei
Rutaceae_Spathelia_terminalioides
Rutaceae_Spathelia_excelsa
Rutaceae_Dictyoloma_vandellianum
Rutaceae_Cneorum_pulverulentum
Rutaceae_Cneorum_tricoccon
Rutaceae_Bottegoa_insignis
Rutaceae_Cedrelopsis_trivalvis
Rutaceae_Cedrelopsis_gracilis
Rutaceae_Cedrelopsis_grevei
Rutaceae_Cedrelopsis_rakotozafyi
Rutaceae_Ptaeroxylon_obliquum
Rutaceae_Harrisonia_perforata
Rutaceae_Harrisonia_abyssinica
Rutaceae_Harrisonia_brownii
Rutaceae_Ruta_oreojasme
Rutaceae_Ruta_microcarpa
Rutaceae_Ruta_pinnata
Rutaceae_Ruta_montana
Rutaceae_Ruta_macrophylla
Rutaceae_Ruta_graveolens
Rutaceae_Ruta_chalepensis

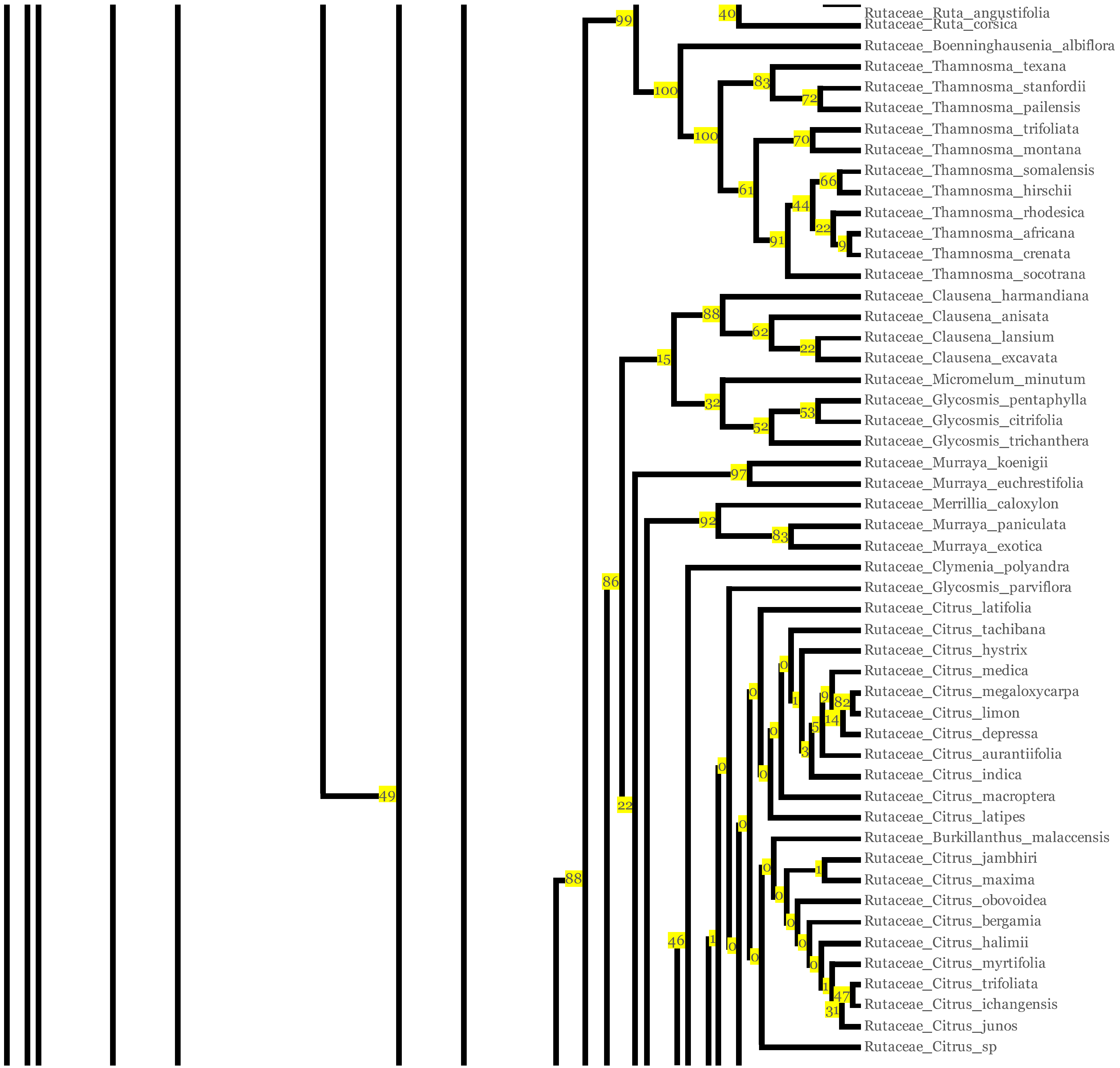
Rutaceae_Ruta_angustifolia
Rutaceae_Ruta_corsica
Rutaceae_Boenninghausenia_albiflora
Rutaceae_Thamnosma_texana
Rutaceae_Thamnosma_stanfordii
Rutaceae_Thamnosma_pailensis
Rutaceae_Thamnosma_trifoliata
Rutaceae_Thamnosma_montana
Rutaceae_Thamnosma_somalensis
Rutaceae_Thamnosma_hirschii
Rutaceae_Thamnosma_rhodesica
Rutaceae_Thamnosma_africana
Rutaceae_Thamnosma_crenata
Rutaceae_Thamnosma_socotrana
Rutaceae_Clausena_harmandiana
Rutaceae_Clausena_anisata
Rutaceae_Clausena_lansium
Rutaceae_Clausena_excavata
Rutaceae_Micromelum_minutum
Rutaceae_Glycosmis_pentaphylla
Rutaceae_Glycosmis_citrifolia
Rutaceae_Glycosmis_trichanthera
Rutaceae_Murraya_koenigii
Rutaceae_Murraya_euchrestifolia
Rutaceae_Merrillia_caloxylon
Rutaceae_Murraya_paniculata
Rutaceae_Murraya_exotica
Rutaceae_Clymenia_polyandra
Rutaceae_Glycosmis_parviflora
Rutaceae_Citrus_latifolia
Rutaceae_Citrus_tachibana
Rutaceae_Citrus_hystrix
Rutaceae_Citrus_medica
Rutaceae_Citrus_megaloxycarpa
Rutaceae_Citrus_limon
Rutaceae_Citrus_depressa
Rutaceae_Citrus_aurantiifolia
Rutaceae_Citrus_indica
Rutaceae_Citrus_macroptera
Rutaceae_Citrus_latipes
Rutaceae_Burkillanthus_malaccensis
Rutaceae_Citrus_jambhiri
Rutaceae_Citrus_maxima
Rutaceae_Citrus_obovoidea
Rutaceae_Citrus_bergamia
Rutaceae_Citrus_halimii
Rutaceae_Citrus_myrtifolia
Rutaceae_Citrus_trifoliata
Rutaceae_Citrus_ichangensis
Rutaceae_Citrus_junos
Rutaceae_Citrus_sp

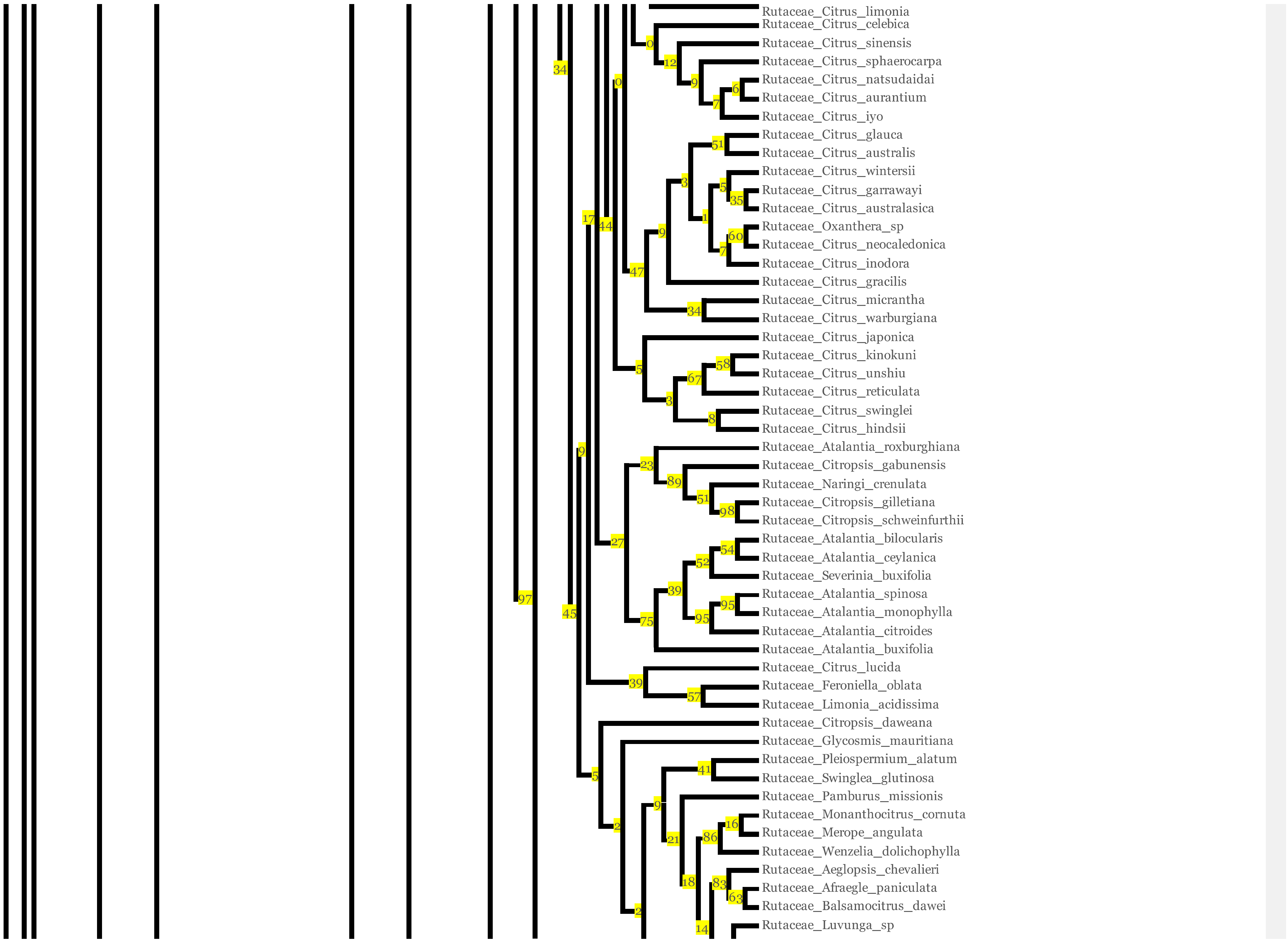
Rutaceae_Citrus_limonia
Rutaceae_Citrus_celebica
Rutaceae_Citrus_sinensis
Rutaceae_Citrus_sphaerocarpa
Rutaceae_Citrus_natsudaidai
Rutaceae_Citrus_aurantium
Rutaceae_Citrus_iyo
Rutaceae_Citrus_glauca
Rutaceae_Citrus_australis
Rutaceae_Citrus_wintersii
Rutaceae_Citrus_garrawayi
Rutaceae_Citrus_australasica
Rutaceae_Oxanthera_sp
Rutaceae_Citrus_neocaledonica
Rutaceae_Citrus_inodora
Rutaceae_Citrus_gracilis
Rutaceae_Citrus_micrantha
Rutaceae_Citrus_warburgiana
Rutaceae_Citrus_japonica
Rutaceae_Citrus_kinokuni
Rutaceae_Citrus_unshiu
Rutaceae_Citrus_reticulata
Rutaceae_Citrus_swinglei
Rutaceae_Citrus_hindsii
Rutaceae_Atalantia_roxburghiana
Rutaceae_Citropsis_gabunensis
Rutaceae_Naringi_crenulata
Rutaceae_Citropsis_gilletiana
Rutaceae_Citropsis_schweinfurthii
Rutaceae_Atalantia_bilocularis
Rutaceae_Atalantia_ceylanica
Rutaceae_Severinia_buxifolia
Rutaceae_Atalantia_spinosa
Rutaceae_Atalantia_monophylla
Rutaceae_Atalantia_citroides
Rutaceae_Atalantia_buxifolia
Rutaceae_Citrus_lucida
Rutaceae_Feroniella_oblata
Rutaceae_Limonia_acidissima
Rutaceae_Citropsis_daweana
Rutaceae_Glycosmis_mauritiana
Rutaceae_Pleiospermium_alatum
Rutaceae_Swinglea_glutinosa
Rutaceae_Pamburus_missionis
Rutaceae_Monanthocitrus_cornuta
Rutaceae_Merope_angulata
Rutaceae_Wenzelia_dolichophylla
Rutaceae_Aeglopsis_chevalieri
Rutaceae_Afraegle_paniculata
Rutaceae_Balsamocitrus_dawei
Rutaceae_Luvunga_sp

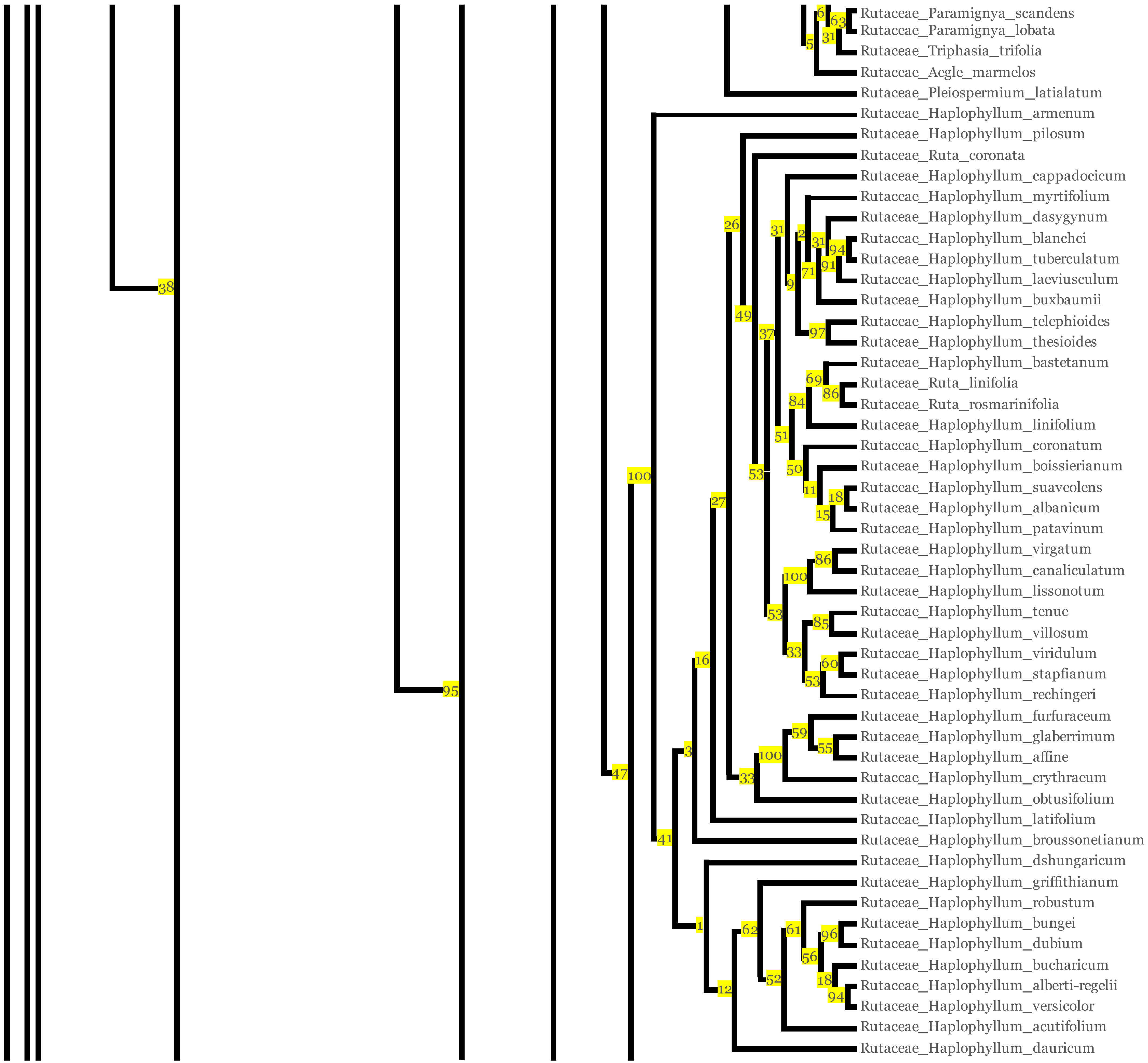
Rutaceae_Paramignya_scandens
Rutaceae_Paramignya_lobata
Rutaceae_Triphasia_trifolia
Rutaceae_Aegle_marmelos
Rutaceae_Pleiospermium_latialatum
Rutaceae_Haplophyllum_armenum
Rutaceae_Haplophyllum_pilosum
Rutaceae_Ruta_coronata
Rutaceae_Haplophyllum_cappadocicum
Rutaceae_Haplophyllum_myrtifolium
Rutaceae_Haplophyllum_dasygynum
Rutaceae_Haplophyllum_blanchei
Rutaceae_Haplophyllum_tuberculatum
Rutaceae_Haplophyllum_laeviusculum
Rutaceae_Haplophyllum_buxbaumii
Rutaceae_Haplophyllum_telephioides
Rutaceae_Haplophyllum_thesioides
Rutaceae_Haplophyllum_bastetanum
Rutaceae_Ruta_linifolia
Rutaceae_Ruta_rosmarinifolia
Rutaceae_Haplophyllum_linifolium
Rutaceae_Haplophyllum_coronatum
Rutaceae_Haplophyllum_boissierianum
Rutaceae_Haplophyllum_suaveolens
Rutaceae_Haplophyllum_albanicum
Rutaceae_Haplophyllum_patavinum
Rutaceae_Haplophyllum_virgatum
Rutaceae_Haplophyllum_canaliculatum
Rutaceae_Haplophyllum_lissonotum
Rutaceae_Haplophyllum_tenue
Rutaceae_Haplophyllum_villosum
Rutaceae_Haplophyllum_viridulum
Rutaceae_Haplophyllum_stapfianum
Rutaceae_Haplophyllum_rechingeri
Rutaceae_Haplophyllum_furfuraceum
Rutaceae_Haplophyllum_glaberrimum
Rutaceae_Haplophyllum_affine
Rutaceae_Haplophyllum_erythraeum
Rutaceae_Haplophyllum_obtusifolium
Rutaceae_Haplophyllum_latifolium
Rutaceae_Haplophyllum_broussonetianum
Rutaceae_Haplophyllum_dshungaricum
Rutaceae_Haplophyllum_griffithianum
Rutaceae_Haplophyllum_robustum
Rutaceae_Haplophyllum_bungei
Rutaceae_Haplophyllum_dubium
Rutaceae_Haplophyllum_bucharicum
Rutaceae_Haplophyllum_alberti-regelii
Rutaceae_Haplophyllum_versicolor
Rutaceae_Haplophyllum_acutifolium
Rutaceae_Haplophyllum_dauricum

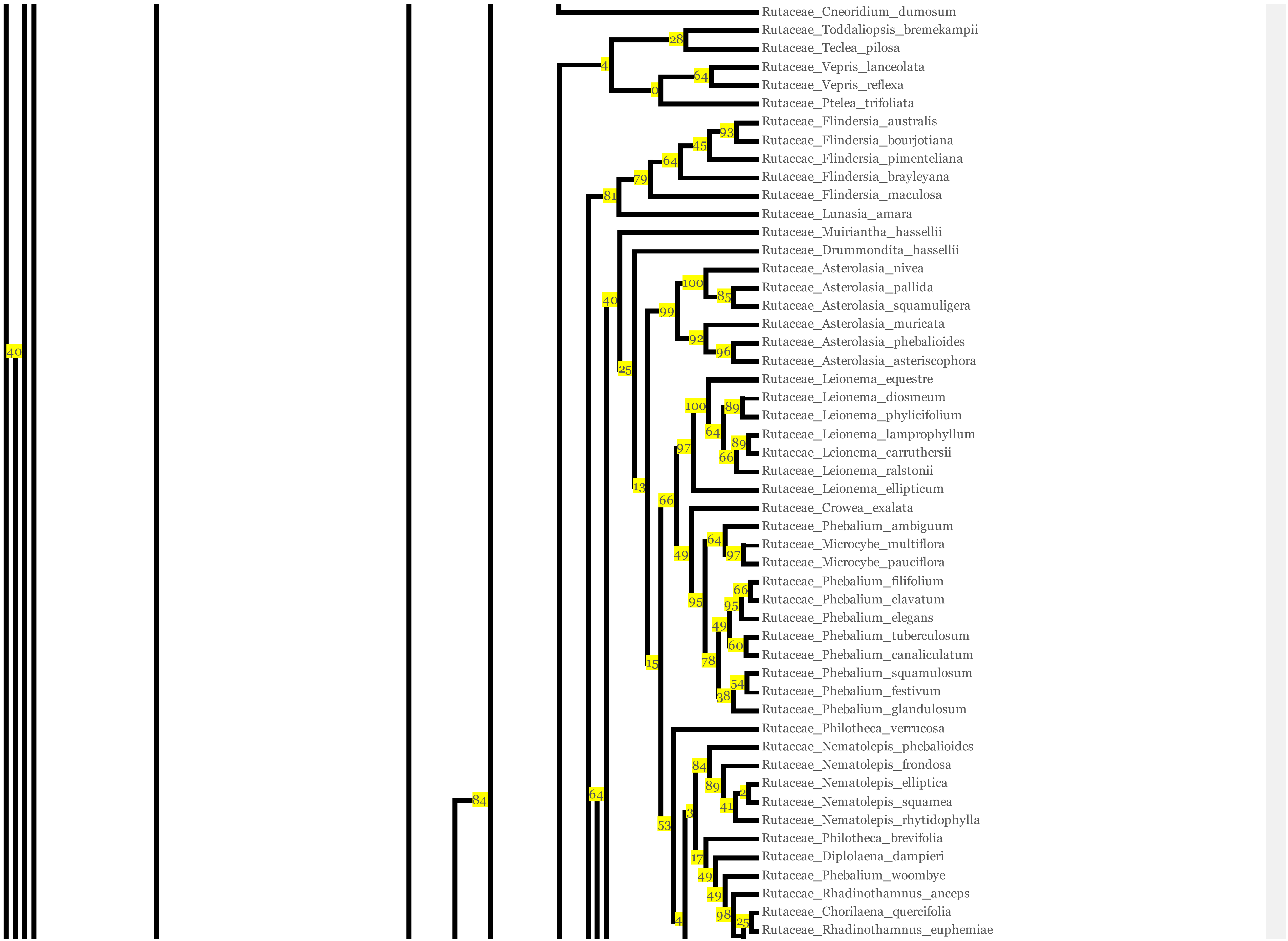

Rutaceae_Cneoridium_dumosum
Rutaceae_Toddaliopsis_bremekampii
Rutaceae_Teclea_pilosa
Rutaceae_Vepris_lanceolata
Rutaceae_Vepris_reflexa
Rutaceae_Ptelea_trifoliata
Rutaceae_Flindersia_australis
Rutaceae_Flindersia_bourjotiana
Rutaceae_Flindersia_pimenteliana
Rutaceae_Flindersia_brayleyana
Rutaceae_Flindersia_maculosa
Rutaceae_Lunasia_amara
Rutaceae_Muiriantha_hassellii
Rutaceae_Drummondita_hassellii
Rutaceae_Asterolasia_nivea
Rutaceae_Asterolasia_pallida
Rutaceae_Asterolasia_squamuligera
Rutaceae_Asterolasia_muricata
Rutaceae_Asterolasia_phebalioides
Rutaceae_Asterolasia_asteriscophora
Rutaceae_Leionema_equestre
Rutaceae_Leionema_diosmeum
Rutaceae_Leionema_phylicifolium
Rutaceae_Leionema_lamprophyllum
Rutaceae_Leionema_carruthersii
Rutaceae_Leionema_ralstonii
Rutaceae_Leionema_ellipticum
Rutaceae_Crowea_exalata
Rutaceae_Phebalium_ambiguum
Rutaceae_Microcybe_multiflora
Rutaceae_Microcybe_pauciflora
Rutaceae_Phebalium_filifolium
Rutaceae_Phebalium_clavatum
Rutaceae_Phebalium_elegans
Rutaceae_Phebalium_tuberculosum
Rutaceae_Phebalium_canaliculatum
Rutaceae_Phebalium_squamulosum
Rutaceae_Phebalium_festivum
Rutaceae_Phebalium_glandulosum
Rutaceae_Philotheca_verrucosa
Rutaceae_Nematolepis_phebalioides
Rutaceae_Nematolepis_frondosa
Rutaceae_Nematolepis_elliptica
Rutaceae_Nematolepis_squamea
Rutaceae_Nematolepis_rhytidophylla
Rutaceae_Philotheca_brevifolia
Rutaceae_Diplolaena_dampieri
Rutaceae_Phebalium_woombye
Rutaceae_Rhadinothamnus_anceps
Rutaceae_Chorilaena_quercifolia
Rutaceae_Rhadinothamnus_euphemiae

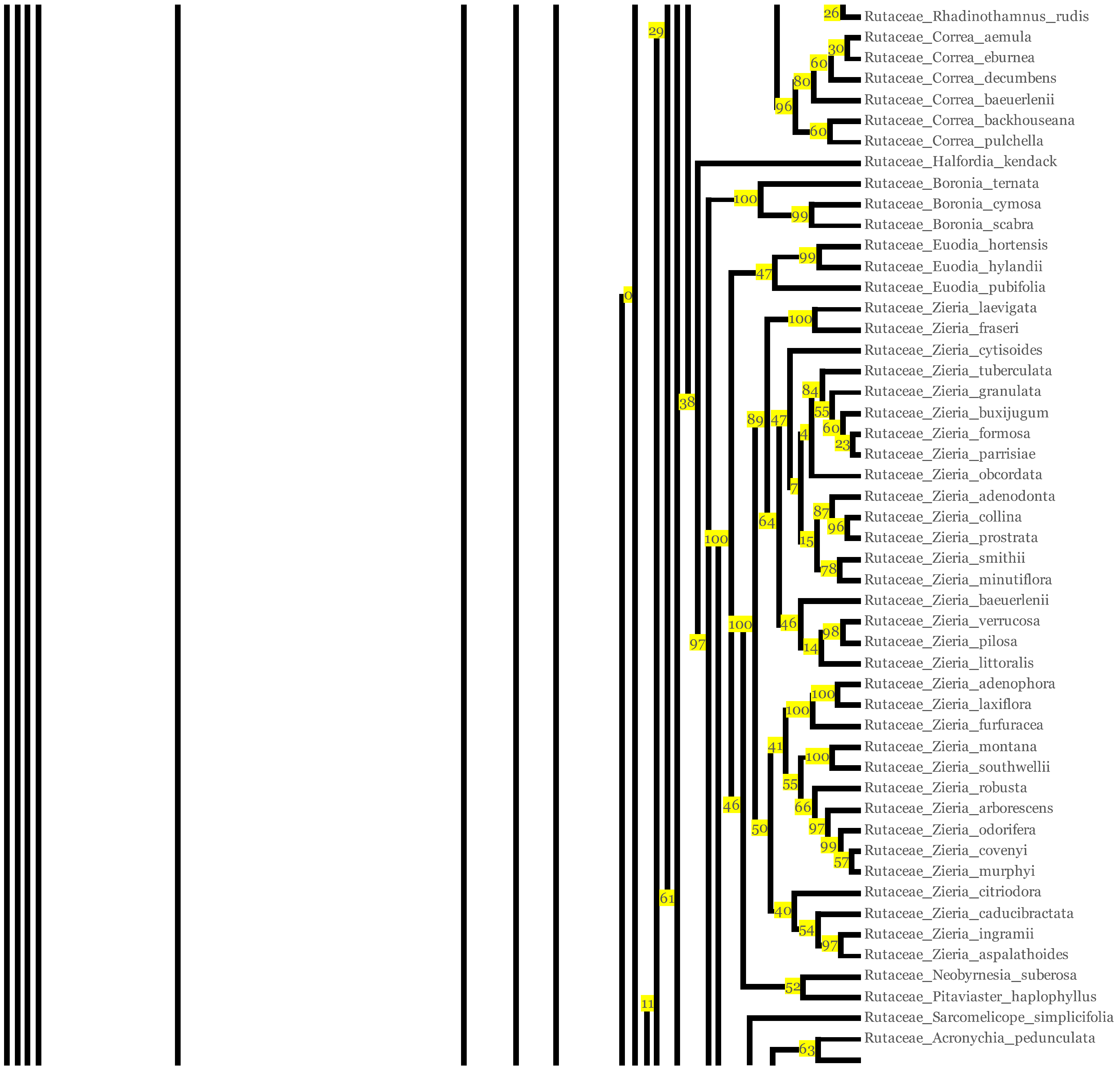
Rutaceae_Rhadinothamnus_rudis
Rutaceae_Correa_aemula
Rutaceae_Correa_eburnea
Rutaceae_Correa_decumbens
Rutaceae_Correa_baeuerlenii
Rutaceae_Correa_backhouseana
Rutaceae_Correa_pulchella
Rutaceae_Halfordia_kendack
Rutaceae_Boronia_ternata
Rutaceae_Boronia_cymosa
Rutaceae_Boronia_scabra
Rutaceae_Euodia_hortensis
Rutaceae_Euodia_hylandii
Rutaceae_Euodia_pubifolia
Rutaceae_Zieria_laevigata
Rutaceae_Zieria_fraseri
Rutaceae_Zieria_cytisoides
Rutaceae_Zieria_tuberculata
Rutaceae_Zieria_granulata
Rutaceae_Zieria_buxijugum
Rutaceae_Zieria_formosa
Rutaceae_Zieria_parrisiae
Rutaceae_Zieria_obcordata
Rutaceae_Zieria_adenodonta
Rutaceae_Zieria_collina
Rutaceae_Zieria_prostrata
Rutaceae_Zieria_smithii
Rutaceae_Zieria_minutiflora
Rutaceae_Zieria_baeuerlenii
Rutaceae_Zieria_verrucosa
Rutaceae_Zieria_pilosa
Rutaceae_Zieria_littoralis
Rutaceae_Zieria_adenophora
Rutaceae_Zieria_laxiflora
Rutaceae_Zieria_furfuracea
Rutaceae_Zieria_montana
Rutaceae_Zieria_southwellii
Rutaceae_Zieria_robusta
Rutaceae_Zieria_arborescens
Rutaceae_Zieria_odorifera
Rutaceae_Zieria_covenyi
Rutaceae_Zieria_murphyi
Rutaceae_Zieria_citriodora
Rutaceae_Zieria_caducibractata
Rutaceae_Zieria_ingramii
Rutaceae_Zieria_aspalathoides
Rutaceae_Neobyrnesia_suberosa
Rutaceae_Pitaviaster_haplophyllus
Rutaceae_Sarcomelicope_simplicifolia
Rutaceae_Acronychia_pedunculata

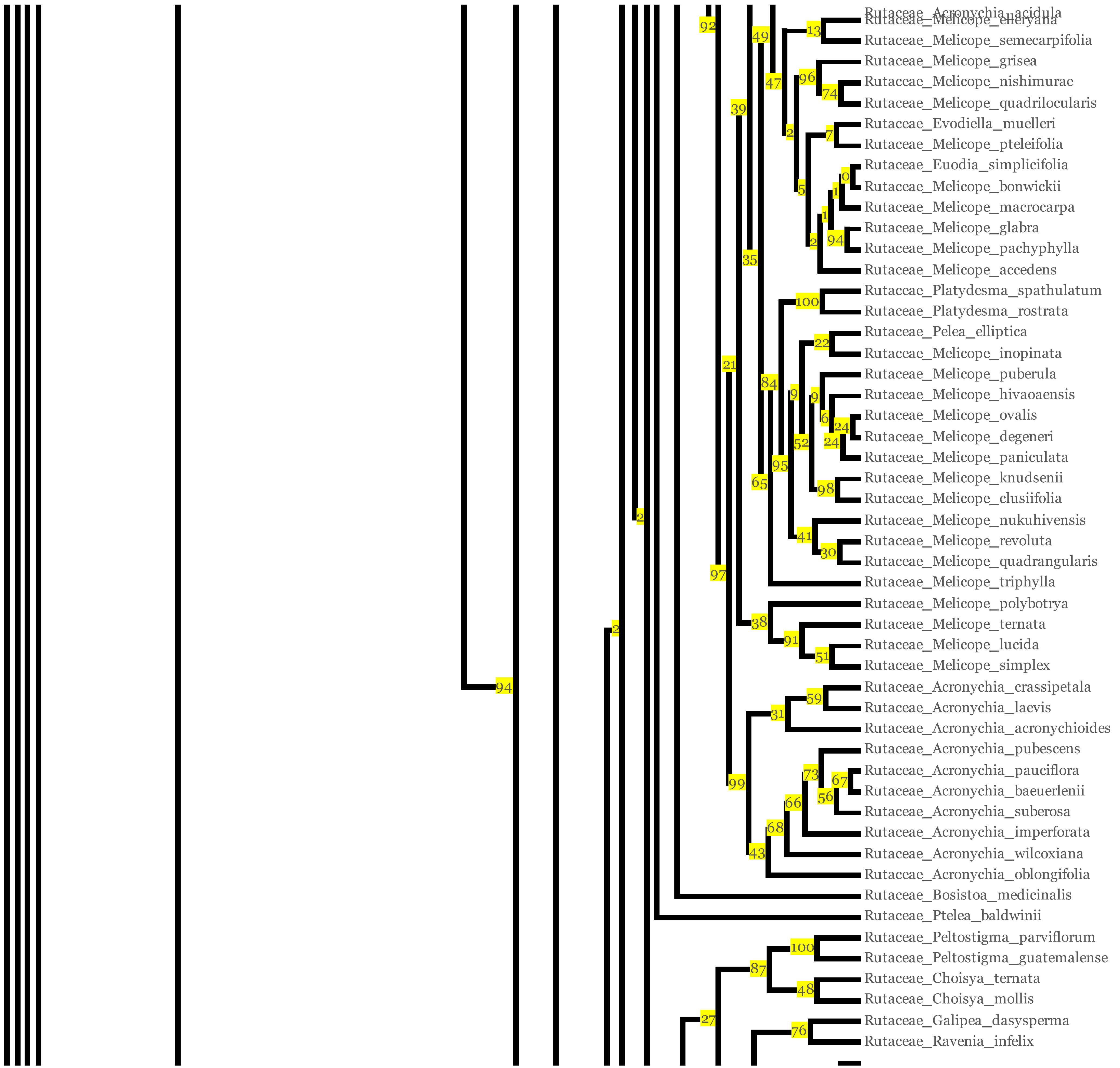

Rutaceae_Acronychia_acidula
Rutaceae_Melicope_elleryana
Rutaceae_Melicope_semecarpifolia
Rutaceae_Melicope_grisea
Rutaceae_Melicope_nishimurae
Rutaceae_Melicope_quadrilocularis
Rutaceae_Evodiella_muelleri
Rutaceae_Melicope_pteleifolia
Rutaceae_Euodia_simplicifolia
Rutaceae_Melicope_bonwickii
Rutaceae_Melicope_macrocarpa
Rutaceae_Melicope_glabra
Rutaceae_Melicope_pachyphylla
Rutaceae_Melicope_accedens
Rutaceae_Platydesma_spathulatum
Rutaceae_Platydesma_rostrata
Rutaceae_Pelea_elliptica
Rutaceae_Melicope_inopinata
Rutaceae_Melicope_puberula
Rutaceae_Melicope_hivaoaensis
Rutaceae_Melicope_ovalis
Rutaceae_Melicope_degeneri
Rutaceae_Melicope_paniculata
Rutaceae_Melicope_knudsenii
Rutaceae_Melicope_clusiifolia
Rutaceae_Melicope_nukuhivensis
Rutaceae_Melicope_revoluta
Rutaceae_Melicope_quadrangularis
Rutaceae_Melicope_triphylla
Rutaceae_Melicope_polybotrya
Rutaceae_Melicope_ternata
Rutaceae_Melicope_lucida
Rutaceae_Melicope_simplex
Rutaceae_Acronychia_crassipetala
Rutaceae_Acronychia_laevis
Rutaceae_Acronychia_acronychioides
Rutaceae_Acronychia_pubescens
Rutaceae_Acronychia_pauciflora
Rutaceae_Acronychia_baeuerlenii
Rutaceae_Acronychia_suberosa
Rutaceae_Acronychia_imperforata
Rutaceae_Acronychia_wilcoxiana
Rutaceae_Acronychia_oblongifolia
Rutaceae_Bosistoa_medicinalis
Rutaceae_Ptelea_baldwinii
Rutaceae_Peltostigma_parviflorum
Rutaceae_Peltostigma_guatemalense
Rutaceae_Choisya_ternata
Rutaceae_Choisya_mollis
Rutaceae_Galipea_dasysperma
Rutaceae_Ravenia_infelix

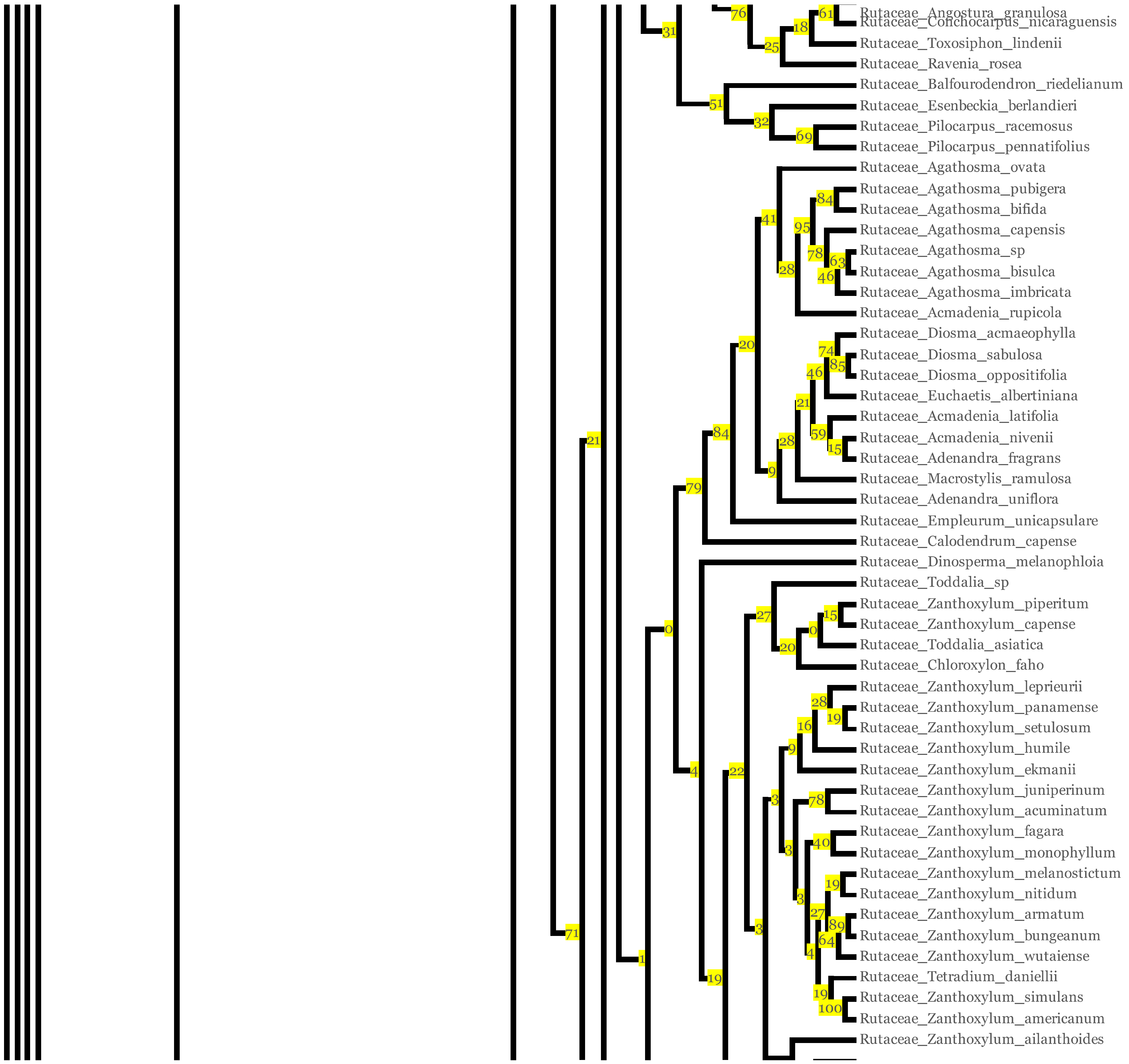
Rutaceae_Angostura_granulosa
Rutaceae_Conchocarpus_nicaraguensis
Rutaceae_Toxosiphon_lindenii
Rutaceae_Ravenia_rosea
Rutaceae_Balfourodendron_riedelianum
Rutaceae_Esenbeckia_berlandieri
Rutaceae_Pilocarpus_racemosus
Rutaceae_Pilocarpus_pennatifolius
Rutaceae_Agathosma_ovata
Rutaceae_Agathosma_pubigera
Rutaceae_Agathosma_bifida
Rutaceae_Agathosma_capensis
Rutaceae_Agathosma_sp
Rutaceae_Agathosma_bisulca
Rutaceae_Agathosma_imbricata
Rutaceae_Acmadenia_rupicola
Rutaceae_Diosma_acmaeophylla
Rutaceae_Diosma_sabulosa
Rutaceae_Diosma_oppositifolia
Rutaceae_Euchaetis_albertiniana
Rutaceae_Acmadenia_latifolia
Rutaceae_Acmadenia_nivenii
Rutaceae_Adenandra_fragrans
Rutaceae_Macrostylis_ramulosa
Rutaceae_Adenandra_uniflora
Rutaceae_Empleurum_unicapsulare
Rutaceae_Calodendrum_capense
Rutaceae_Dinosperma_melanophloia
Rutaceae_Toddalia_sp
Rutaceae_Zanthoxylum_piperitum
Rutaceae_Zanthoxylum_capense
Rutaceae_Toddalia_asiatica
Rutaceae_Chloroxylon_faho
Rutaceae_Zanthoxylum_leprieurii
Rutaceae_Zanthoxylum_panamense
Rutaceae_Zanthoxylum_setulosum
Rutaceae_Zanthoxylum_humile
Rutaceae_Zanthoxylum_ekmanii
Rutaceae_Zanthoxylum_juniperinum
Rutaceae_Zanthoxylum_acuminatum
Rutaceae_Zanthoxylum_fagara
Rutaceae_Zanthoxylum_monophyllum
Rutaceae_Zanthoxylum_melanostictum
Rutaceae_Zanthoxylum_nitidum
Rutaceae_Zanthoxylum_armatum
Rutaceae_Zanthoxylum_bungeanum
Rutaceae_Zanthoxylum_wutaiense
Rutaceae_Tetradium_daniellii
Rutaceae_Zanthoxylum_simulans
Rutaceae_Zanthoxylum_americanum
Rutaceae_Zanthoxylum_ailanthoides

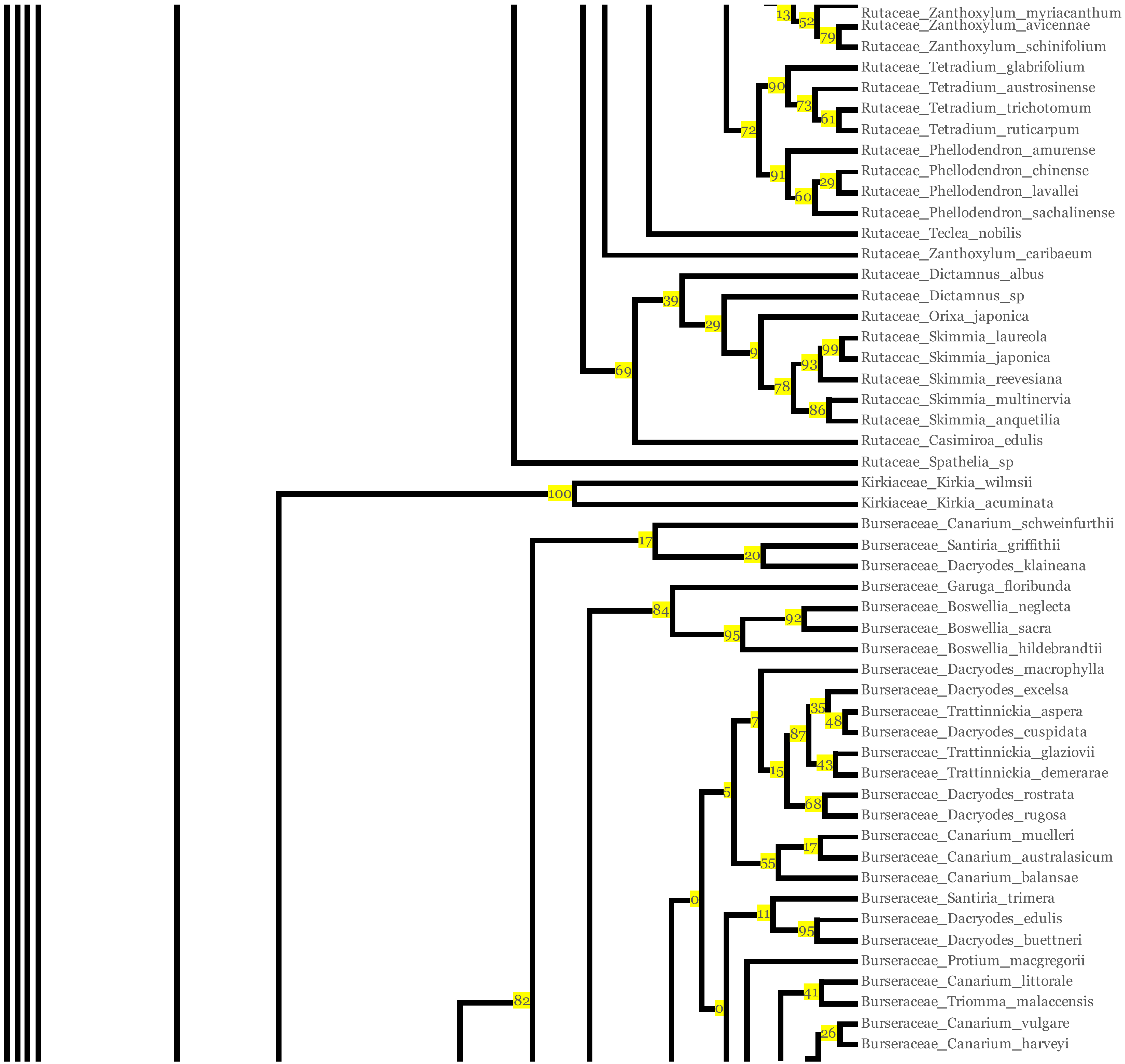
Rutaceae_Zanthoxylum_myriacanthum
Rutaceae_Zanthoxylum_avicennae
Rutaceae_Zanthoxylum_schinifolium
Rutaceae_Tetradium_glabrifolium
Rutaceae_Tetradium_austrosinense
Rutaceae_Tetradium_trichotomum
Rutaceae_Tetradium_ruticarpum
Rutaceae_Phellodendron_amurense
Rutaceae_Phellodendron_chinense
Rutaceae_Phellodendron_lavallei
Rutaceae_Phellodendron_sachalinense
Rutaceae_Teclea_nobilis
Rutaceae_Zanthoxylum_caribaeum
Rutaceae_Dictamnus_albus
Rutaceae_Dictamnus_sp
Rutaceae_Orixa_japonica
Rutaceae_Skimmia_laureola
Rutaceae_Skimmia_japonica
Rutaceae_Skimmia_reevesiana
Rutaceae_Skimmia_multinervia
Rutaceae_Skimmia_anquetilia
Rutaceae_Casimiroa_edulis
Rutaceae_Spathelia_sp
Kirkiaceae_Kirkia_wilmsii
Kirkiaceae_Kirkia_acuminata
Burseraceae_Canarium_schweinfurthii
Burseraceae_Santiria_griffithii
Burseraceae_Dacryodes_klaineana
Burseraceae_Garuga_floribunda
Burseraceae_Boswellia_neglecta
Burseraceae_Boswellia_sacra
Burseraceae_Boswellia_hildebrandtii
Burseraceae_Dacryodes_macrophylla
Burseraceae_Dacryodes_excelsa
Burseraceae_Trattinnickia_aspera
Burseraceae_Dacryodes_cuspidata
Burseraceae_Trattinnickia_glaziovii
Burseraceae_Trattinnickia_demerarae
Burseraceae_Dacryodes_rostrata
Burseraceae_Dacryodes_rugosa
Burseraceae_Canarium_muelleri
Burseraceae_Canarium_australasicum
Burseraceae_Canarium_balansae
Burseraceae_Santiria_trimera
Burseraceae_Dacryodes_edulis
Burseraceae_Dacryodes_buettneri
Burseraceae_Protium_macgregorii
Burseraceae_Canarium_littorale
Burseraceae_Triomma_malaccensis
Burseraceae_Canarium_vulgare
Burseraceae_Canarium_harveyi

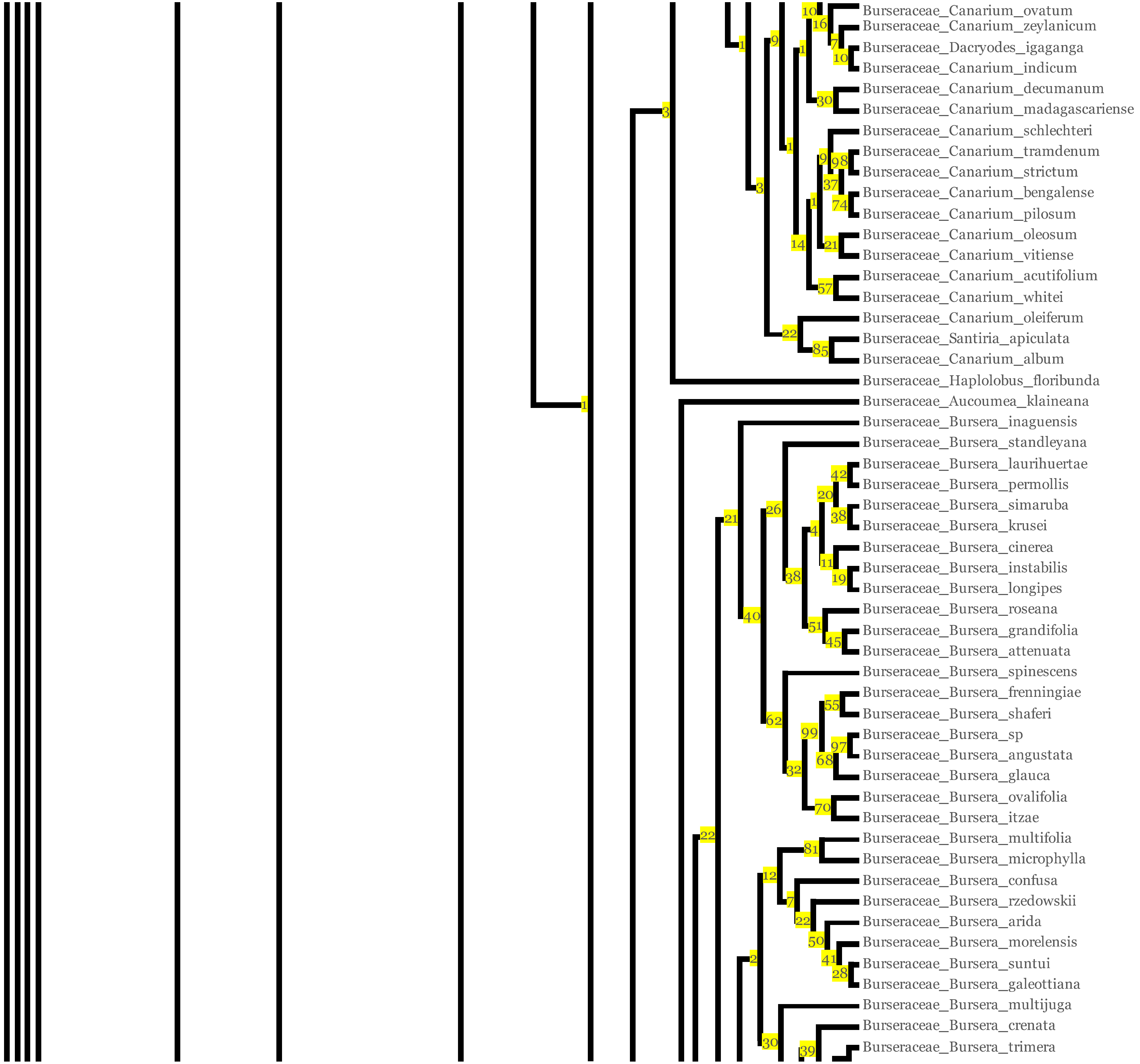
Burseraceae_Canarium_ovatum
Burseraceae_Canarium_zeylanicum
Burseraceae_Dacryodes_igaganga
Burseraceae_Canarium_indicum
Burseraceae_Canarium_decumanum
Burseraceae_Canarium_madagascariense
Burseraceae_Canarium_schlechteri
Burseraceae_Canarium_tramdenum
Burseraceae_Canarium_strictum
Burseraceae_Canarium_bengalense
Burseraceae_Canarium_pilosum
Burseraceae_Canarium_oleosum
Burseraceae_Canarium_vitiense
Burseraceae_Canarium_acutifolium
Burseraceae_Canarium_whitei
Burseraceae_Canarium_oleiferum
Burseraceae_Santiria_apiculata
Burseraceae_Canarium_album
Burseraceae_Haplolobus_floribunda
Burseraceae_Aucoumea_klaineana
Burseraceae_Bursera_inaguensis
Burseraceae_Bursera_standleyana
Burseraceae_Bursera_laurihuertae
Burseraceae_Bursera_permollis
Burseraceae_Bursera_simaruba
Burseraceae_Bursera_krusei
Burseraceae_Bursera_cinerea
Burseraceae_Bursera_instabilis
Burseraceae_Bursera_longipes
Burseraceae_Bursera_roseana
Burseraceae_Bursera_grandifolia
Burseraceae_Bursera_attenuata
Burseraceae_Bursera_spinescens
Burseraceae_Bursera_frenningiae
Burseraceae_Bursera_shaferi
Burseraceae_Bursera_sp
Burseraceae_Bursera_angustata
Burseraceae_Bursera_glauca
Burseraceae_Bursera_ovalifolia
Burseraceae_Bursera_itzae
Burseraceae_Bursera_multifolia
Burseraceae_Bursera_microphylla
Burseraceae_Bursera_confusa
Burseraceae_Bursera_rzedowskii
Burseraceae_Bursera_arida
Burseraceae_Bursera_morelensis
Burseraceae_Bursera_suntui
Burseraceae_Bursera_galeottiana
Burseraceae_Bursera_multijuga
Burseraceae_Bursera_crenata
Burseraceae_Bursera_trimera

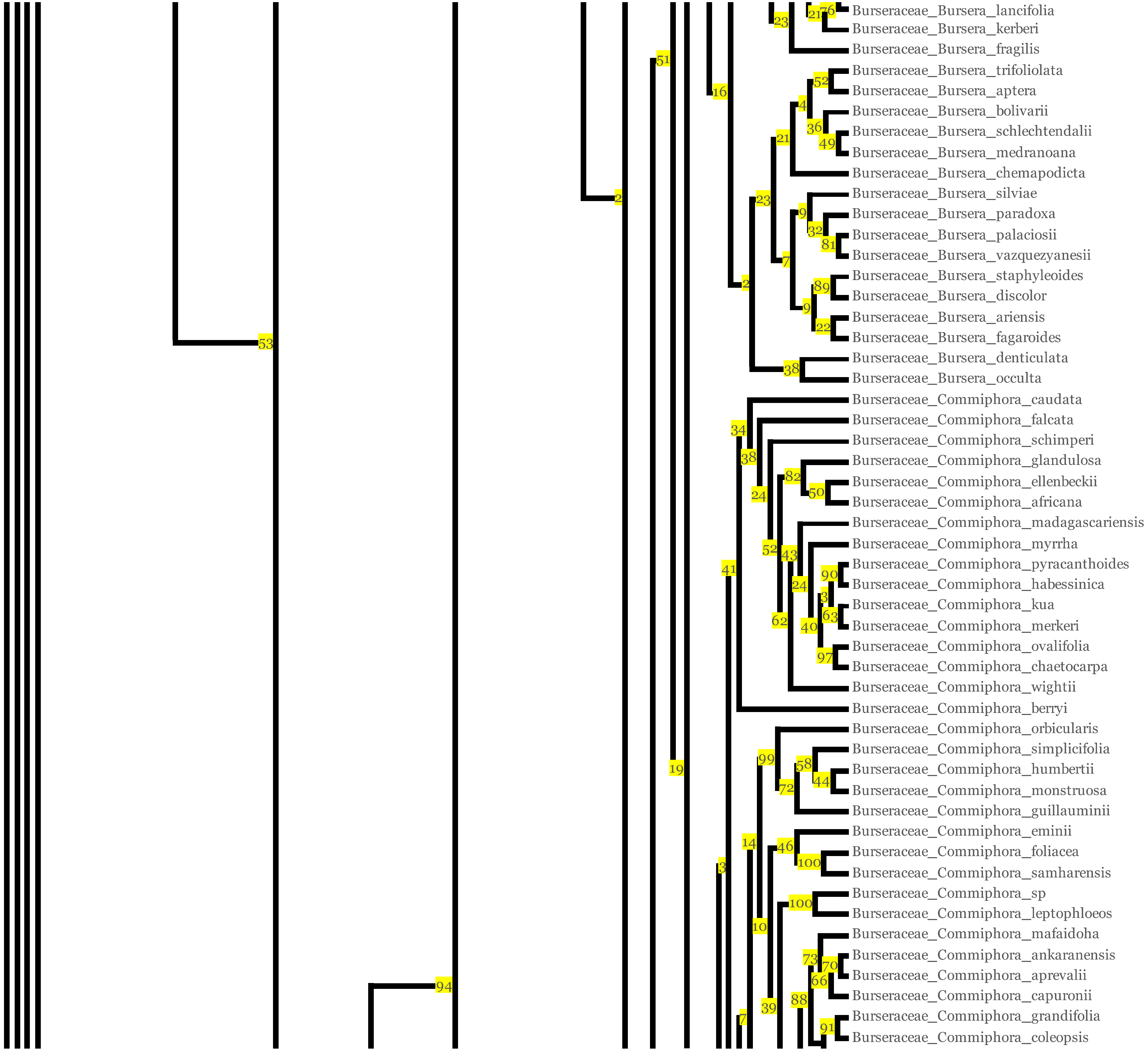

Burseraceae_Bursera_lancifolia
Burseraceae_Bursera_kerberi
Burseraceae_Bursera_fragilis
Burseraceae_Bursera_trifoliolata
Burseraceae_Bursera_aptera
Burseraceae_Bursera_bolivarii
Burseraceae_Bursera_schlechtendalii
Burseraceae_Bursera_medranoana
Burseraceae_Bursera_chemapodicta
Burseraceae_Bursera_silviae
Burseraceae_Bursera_paradoxa
Burseraceae_Bursera_palaciosii
Burseraceae_Bursera_vazquezyanesii
Burseraceae_Bursera_staphyleoides
Burseraceae_Bursera_discolor
Burseraceae_Bursera_ariensis
Burseraceae_Bursera_fagaroides
Burseraceae_Bursera_denticulata
Burseraceae_Bursera_occulta
Burseraceae_Commiphora_caudata
Burseraceae_Commiphora_falcata
Burseraceae_Commiphora_schimperi
Burseraceae_Commiphora_glandulosa
Burseraceae_Commiphora_ellenbeckii
Burseraceae_Commiphora_africana
Burseraceae_Commiphora_madagascariensis
Burseraceae_Commiphora_myrrha
Burseraceae_Commiphora_pyracanthoides
Burseraceae_Commiphora_habessinica
Burseraceae_Commiphora_kua
Burseraceae_Commiphora_merkeri
Burseraceae_Commiphora_ovalifolia
Burseraceae_Commiphora_chaetocarpa
Burseraceae_Commiphora_wightii
Burseraceae_Commiphora_berryi
Burseraceae_Commiphora_orbicularis
Burseraceae_Commiphora_simplicifolia
Burseraceae_Commiphora_humbertii
Burseraceae_Commiphora_monstruosa
Burseraceae_Commiphora_guillauminii
Burseraceae_Commiphora_eminii
Burseraceae_Commiphora_foliacea
Burseraceae_Commiphora_samharensis
Burseraceae_Commiphora_sp
Burseraceae_Commiphora_leptophloeos
Burseraceae_Commiphora_mafaidoha
Burseraceae_Commiphora_ankaranensis
Burseraceae_Commiphora_aprevalii
Burseraceae_Commiphora_capuronii
Burseraceae_Commiphora_grandifolia
Burseraceae_Commiphora_coleopsis

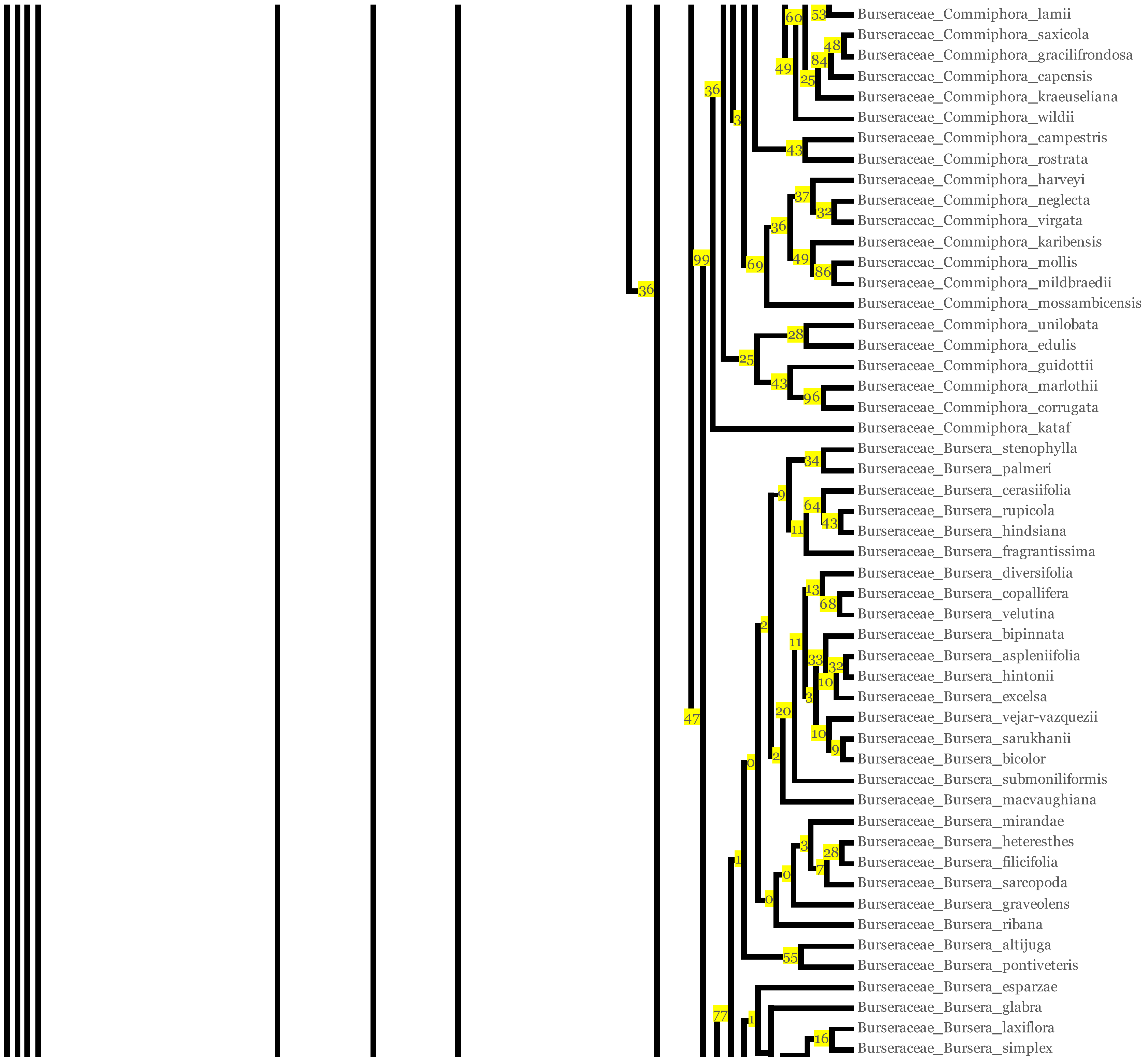
Burseraceae_Commiphora_lamii
Burseraceae_Commiphora_saxicola
Burseraceae_Commiphora_gracilifrondosa
Burseraceae_Commiphora_capensis
Burseraceae_Commiphora_kraeuseliana
Burseraceae_Commiphora_wildii
Burseraceae_Commiphora_campestris
Burseraceae_Commiphora_rostrata
Burseraceae_Commiphora_harveyi
Burseraceae_Commiphora_neglecta
Burseraceae_Commiphora_virgata
Burseraceae_Commiphora_karibensis
Burseraceae_Commiphora_mollis
Burseraceae_Commiphora_mildbraedii
Burseraceae_Commiphora_mossambicensis
Burseraceae_Commiphora_unilobata
Burseraceae_Commiphora_edulis
Burseraceae_Commiphora_guidottii
Burseraceae_Commiphora_marlothii
Burseraceae_Commiphora_corrugata
Burseraceae_Commiphora_kataf
Burseraceae_Bursera_stenophylla
Burseraceae_Bursera_palmeri
Burseraceae_Bursera_cerasiifolia
Burseraceae_Bursera_rupicola
Burseraceae_Bursera_hindsiana
Burseraceae_Bursera_fragrantissima
Burseraceae_Bursera_diversifolia
Burseraceae_Bursera_copallifera
Burseraceae_Bursera_velutina
Burseraceae_Bursera_bipinnata
Burseraceae_Bursera_aspleniifolia
Burseraceae_Bursera_hintonii
Burseraceae_Bursera_excelsa
Burseraceae_Bursera_vejar-vazquezii
Burseraceae_Bursera_sarukhanii
Burseraceae_Bursera_bicolor
Burseraceae_Bursera_submoniliformis
Burseraceae_Bursera_macvaughiana
Burseraceae_Bursera_mirandae
Burseraceae_Bursera_heteresthes
Burseraceae_Bursera_filicifolia
Burseraceae_Bursera_sarcopoda
Burseraceae_Bursera_graveolens
Burseraceae_Bursera_ribana
Burseraceae_Bursera_altijuga
Burseraceae_Bursera_pontiveteris
Burseraceae_Bursera_esparzae
Burseraceae_Bursera_glabra
Burseraceae_Bursera_laxiflora
Burseraceae_Bursera_simplex
60 53 48 84 49 25 36 3 43 37 32 36 49 99 69 86 36 28 25 43 96 34 9 64 43 11 13 68 2 11 33 32 10 3 47 20 10 9 0 2 3 28 1 7 0 0 55 77 1 16

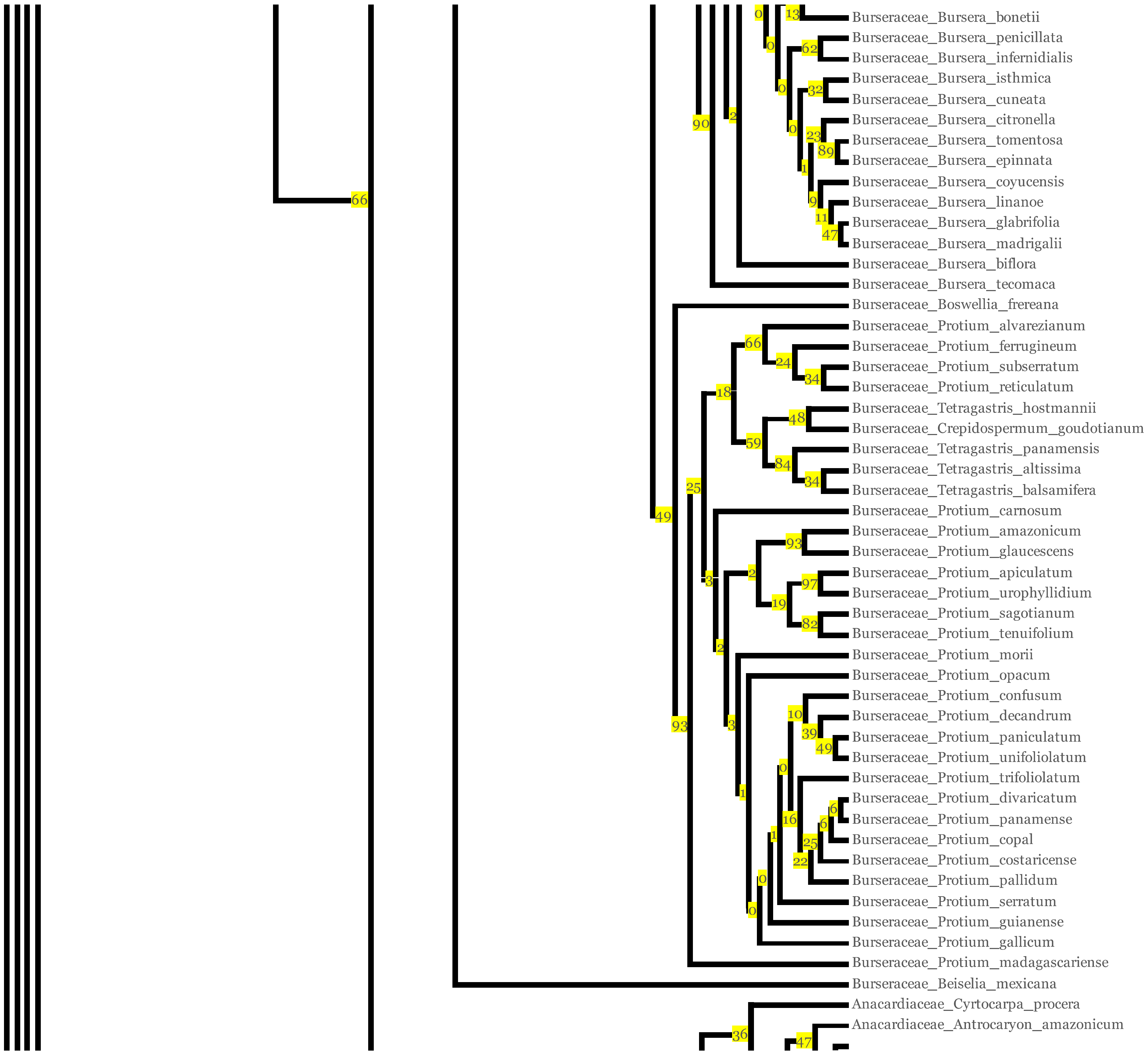

Burseraceae_Bursera_bonetii
Burseraceae_Bursera_penicillata
Burseraceae_Bursera_infernidialis
Burseraceae_Bursera_isthmica
Burseraceae_Bursera_cuneata
Burseraceae_Bursera_citronella
Burseraceae_Bursera_tomentosa
Burseraceae_Bursera_epinnata
Burseraceae_Bursera_coyucensis
Burseraceae_Bursera_linanoe
Burseraceae_Bursera_glabrifolia
Burseraceae_Bursera_madrigalii
Burseraceae_Bursera_biflora
Burseraceae_Bursera_tecomaca
Burseraceae_Boswellia_frereana
Burseraceae_Protium_alvarezianum
Burseraceae_Protium_ferrugineum
Burseraceae_Protium_subserratum
Burseraceae_Protium_reticulatum
Burseraceae_Tetragastris_hostmannii
Burseraceae_Crepidospermum_goudotianum
Burseraceae_Tetragastris_panamensis
Burseraceae_Tetragastris_altissima
Burseraceae_Tetragastris_balsamifera
Burseraceae_Protium_carnosum
Burseraceae_Protium_amazonicum
Burseraceae_Protium_glaucescens
Burseraceae_Protium_apiculatum
Burseraceae_Protium_urophyllidium
Burseraceae_Protium_sagotianum
Burseraceae_Protium_tenuifolium
Burseraceae_Protium_morii
Burseraceae_Protium_opacum
Burseraceae_Protium_confusum
Burseraceae_Protium_decandrum
Burseraceae_Protium_paniculatum
Burseraceae_Protium_unifoliolatum
Burseraceae_Protium_trifoliolatum
Burseraceae_Protium_divaricatum
Burseraceae_Protium_panamense
Burseraceae_Protium_copal
Burseraceae_Protium_costaricense
Burseraceae_Protium_pallidum
Burseraceae_Protium_serratum
Burseraceae_Protium_guianense
Burseraceae_Protium_gallicum
Burseraceae_Protium_madagascariense
Burseraceae_Beiselia_mexicana
Anacardiaceae_Cyrtocarpa_procera
Anacardiaceae_Antrocaryon_amazonicum

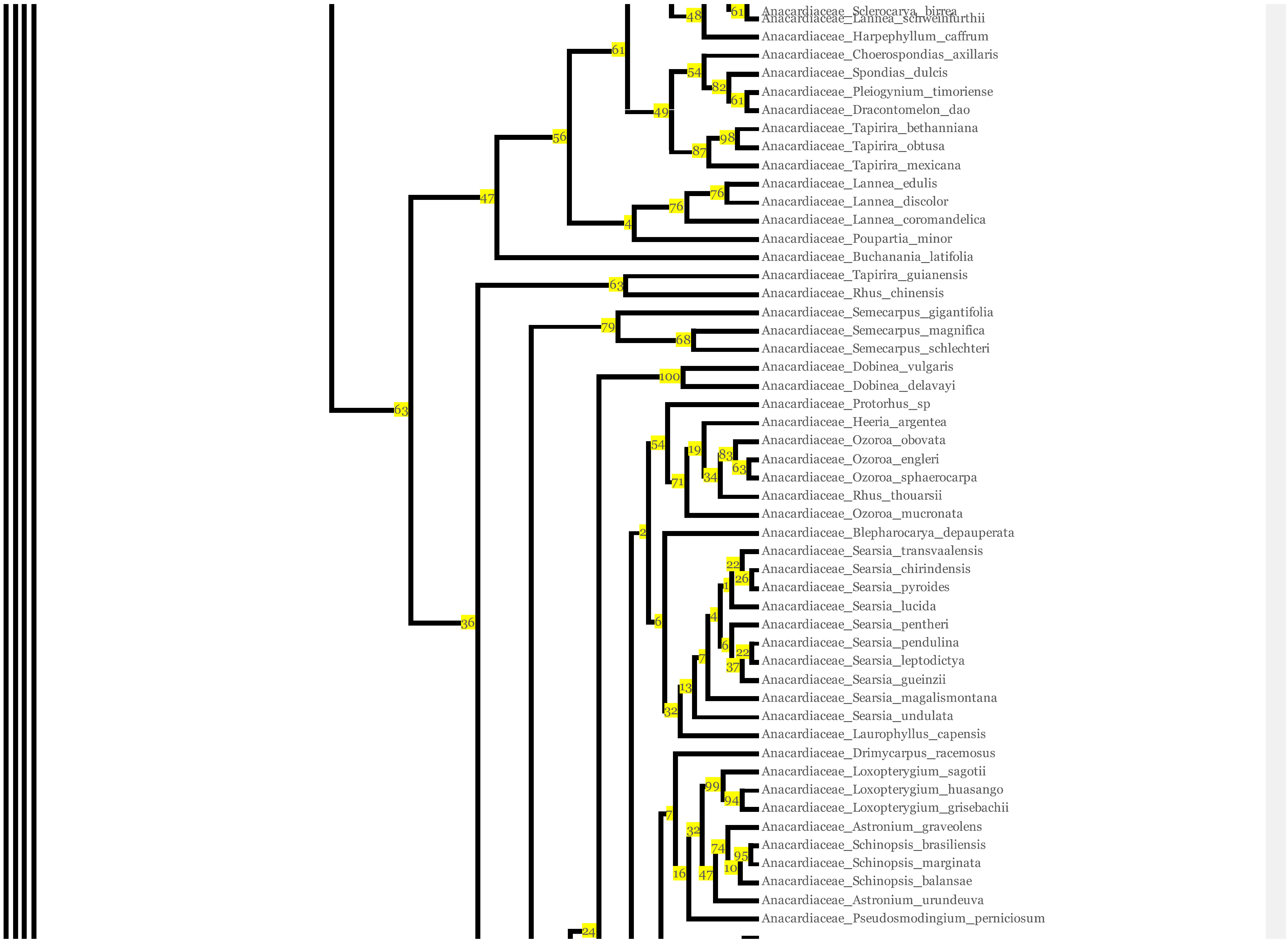
Anacardiaceae_Sclerocarya_birrea
Anacardiaceae_Lannea_schweinfurthii
Anacardiaceae_Harpephyllum_caffrum
Anacardiaceae_Choerospondias_axillaris
Anacardiaceae_Spondias_dulcis
Anacardiaceae_Pleiogynium_timoriense
Anacardiaceae_Dracontomelon_dao
Anacardiaceae_Tapirira_bethanniana
Anacardiaceae_Tapirira_obtusa
Anacardiaceae_Tapirira_mexicana
Anacardiaceae_Lannea_edulis
Anacardiaceae_Lannea_discolor
Anacardiaceae_Lannea_coromandelica
Anacardiaceae_Poupartia_minor
Anacardiaceae_Buchanania_latifolia
Anacardiaceae_Tapirira_guianensis
Anacardiaceae_Rhus_chinensis
Anacardiaceae_Semecarpus_gigantifolia
Anacardiaceae_Semecarpus_magnifica
Anacardiaceae_Semecarpus_schlechteri
Anacardiaceae_Dobinea_vulgaris
Anacardiaceae_Dobinea_delavayi
Anacardiaceae_Protorhus_sp
Anacardiaceae_Heeria_argentea
Anacardiaceae_Ozoroa_obovata
Anacardiaceae_Ozoroa_engleri
Anacardiaceae_Ozoroa_sphaerocarpa
Anacardiaceae_Rhus_thouarsii
Anacardiaceae_Ozoroa_mucronata
Anacardiaceae_Blepharocarya_depauperata
Anacardiaceae_Searsia_transvaalensis
Anacardiaceae_Searsia_chirindensis
Anacardiaceae_Searsia_pyroides
Anacardiaceae_Searsia_lucida
Anacardiaceae_Searsia_pentheri
Anacardiaceae_Searsia_pendulina
Anacardiaceae_Searsia_leptodictya
Anacardiaceae_Searsia_gueinzii
Anacardiaceae_Searsia_magalismontana
Anacardiaceae_Searsia_undulata
Anacardiaceae_Laurophyllus_capensis
Anacardiaceae_Drimycarpus_racemosus
Anacardiaceae_Loxopterygium_sagotii
Anacardiaceae_Loxopterygium_huasango
Anacardiaceae_Loxopterygium_grisebachii
Anacardiaceae_Astronium_graveolens
Anacardiaceae_Schinopsis_brasiliensis
Anacardiaceae_Schinopsis_marginata
Anacardiaceae_Schinopsis_balansae
Anacardiaceae_Astronium_urundeuva
Anacardiaceae_Pseudosmodingium_perniciosum

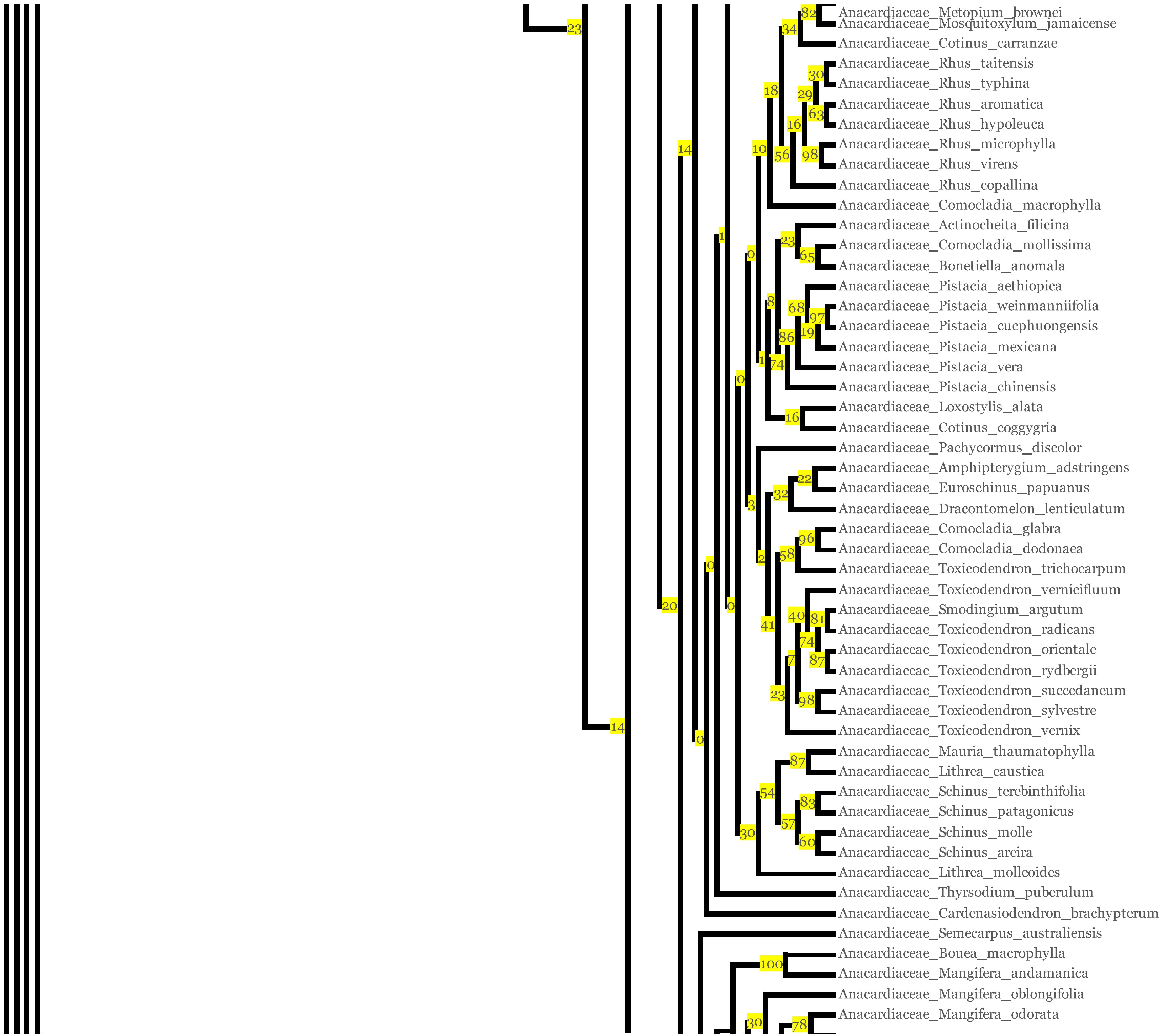

Anacardiaceae_Metopium_brownei
Anacardiaceae_Mosquitoxylum_jamaicense
Anacardiaceae_Cotinus_carranzae
Anacardiaceae_Rhus_taitensis
Anacardiaceae_Rhus_typhina
Anacardiaceae_Rhus_aromatica
Anacardiaceae_Rhus_hypoleuca
Anacardiaceae_Rhus_microphylla
Anacardiaceae_Rhus_virens
Anacardiaceae_Rhus_copallina
Anacardiaceae_Comocladia_macrophylla
Anacardiaceae_Actinocheita_filicina
Anacardiaceae_Comocladia_mollissima
Anacardiaceae_Bonetiella_anomala
Anacardiaceae_Pistacia_aethiopica
Anacardiaceae_Pistacia_weinmanniifolia
Anacardiaceae_Pistacia_cucphuongensis
Anacardiaceae_Pistacia_mexicana
Anacardiaceae_Pistacia_vera
Anacardiaceae_Pistacia_chinensis
Anacardiaceae_Loxostylis_alata
Anacardiaceae_Cotinus_coggygria
Anacardiaceae_Pachycormus_discolor
Anacardiaceae_Amphipterygium_adstringens
Anacardiaceae_Euroschinus_papuanus
Anacardiaceae_Dracontomelon_lenticulatum
Anacardiaceae_Comocladia_glabra
Anacardiaceae_Comocladia_dodonaea
Anacardiaceae_Toxicodendron_trichocarpum
Anacardiaceae_Toxicodendron_vernicifluum
Anacardiaceae_Smodingium_argutum
Anacardiaceae_Toxicodendron_radicans
Anacardiaceae_Toxicodendron_orientale
Anacardiaceae_Toxicodendron_rydbergii
Anacardiaceae_Toxicodendron_succedaneum
Anacardiaceae_Toxicodendron_sylvestre
Anacardiaceae_Toxicodendron_vernix
Anacardiaceae_Mauria_thaumatophylla
Anacardiaceae_Lithrea_caustica
Anacardiaceae_Schinus_terebinthifolia
Anacardiaceae_Schinus_patagonicus
Anacardiaceae_Schinus_molle
Anacardiaceae_Schinus_areira
Anacardiaceae_Lithrea_molleoides
Anacardiaceae_Thyrsodium_puberulum
Anacardiaceae_Cardenasiodendron_brachypterum
Anacardiaceae_Semecarpus_australiensis
Anacardiaceae_Bouea_macrophylla
Anacardiaceae_Mangifera_andamanica
Anacardiaceae_Mangifera_oblongifolia
Anacardiaceae_Mangifera_odorata

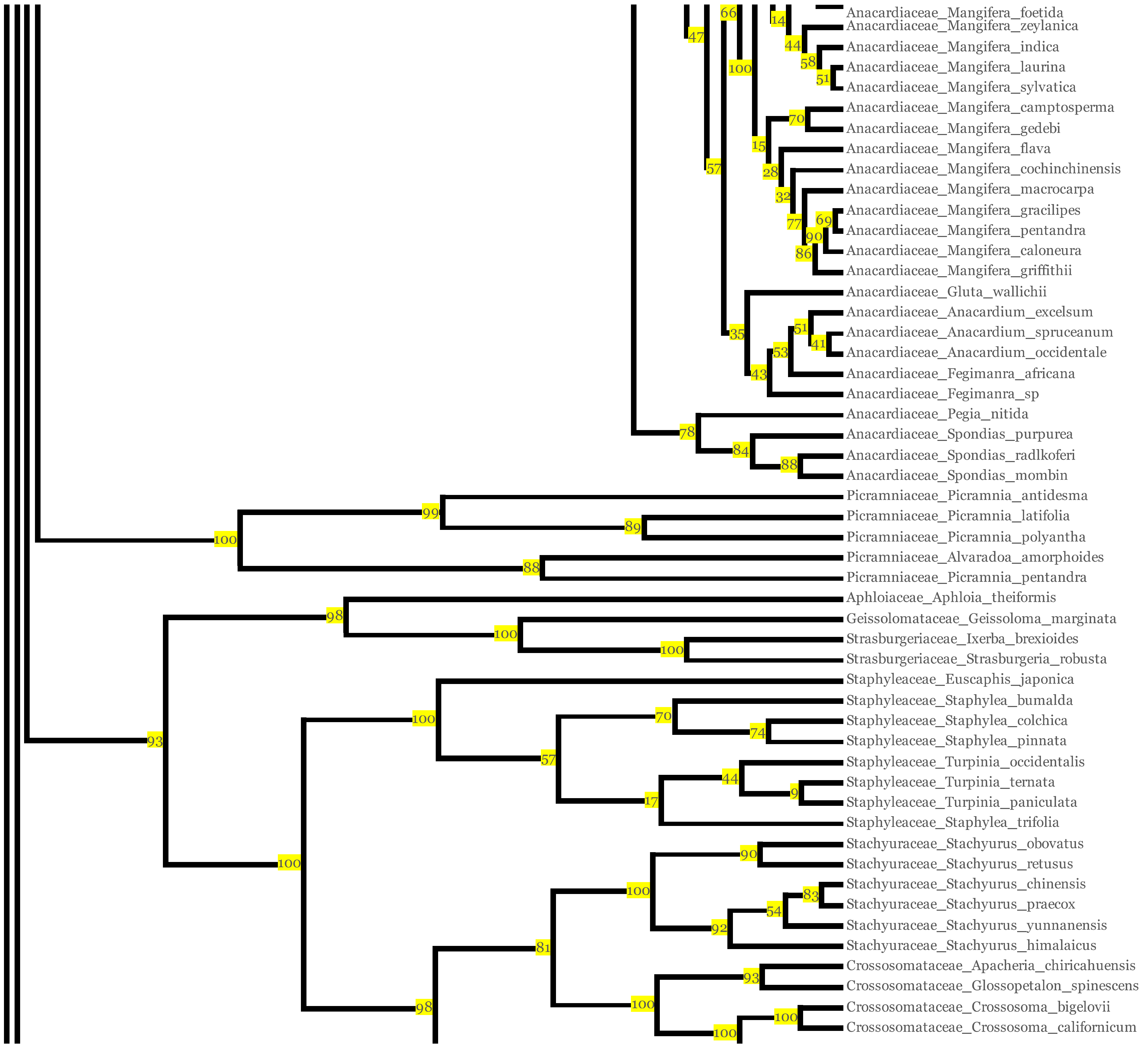
Anacardiaceae_Mangifera_foetida
Anacardiaceae_Mangifera_zeylanica
Anacardiaceae_Mangifera_indica
Anacardiaceae_Mangifera_laurina
Anacardiaceae_Mangifera_sylvatica
Anacardiaceae_Mangifera_camptosperma
Anacardiaceae_Mangifera_gedebi
Anacardiaceae_Mangifera_flava
Anacardiaceae_Mangifera_cochinchinensis
Anacardiaceae_Mangifera_macrocarpa
Anacardiaceae_Mangifera_gracilipes
Anacardiaceae_Mangifera_pentandra
Anacardiaceae_Mangifera_caloneura
Anacardiaceae_Mangifera_griffithii
Anacardiaceae_Gluta_wallichii
Anacardiaceae_Anacardium_excelsum
Anacardiaceae_Anacardium_spruceanum
Anacardiaceae_Anacardium_occidentale
Anacardiaceae_Fegimanra_africana
Anacardiaceae_Fegimanra_sp
Anacardiaceae_Pegia_nitida
Anacardiaceae_Spondias_purpurea
Anacardiaceae_Spondias_radlkoferi
Anacardiaceae_Spondias_mombin
Picramniaceae_Picramnia_antidesma
Picramniaceae_Picramnia_latifolia
Picramniaceae_Picramnia_polyantha
Picramniaceae_Alvaradoa_amorphoides
Picramniaceae_Picramnia_pentandra
Aphloiaceae_Aphloia_theiformis
Geissolomataceae_Geissoloma_marginata
Strasburgeriaceae_Ixerba_brexioides
Strasburgeriaceae_Strasburgeria_robusta
Staphyleaceae_Euscaphis_japonica
Staphyleaceae_Staphylea_bumalda
Staphyleaceae_Staphylea_colchica
Staphyleaceae_Staphylea_pinnata
Staphyleaceae_Turpinia_occidentalis
Staphyleaceae_Turpinia_ternata
Staphyleaceae_Turpinia_paniculata
Staphyleaceae_Staphylea_trifolia
Stachyuraceae_Stachyurus_obovatus
Stachyuraceae_Stachyurus_retusus
Stachyuraceae_Stachyurus_chinensis
Stachyuraceae_Stachyurus_praecox
Stachyuraceae_Stachyurus_yunnanensis
Stachyuraceae_Stachyurus_himalaicus
Crossosomataceae_Apacheria_chiricahuensis
Crossosomataceae_Glossopetalon_spinescens
Crossosomataceae_Crossosoma_bigelovii
Crossosomataceae_Crossosoma_californicum

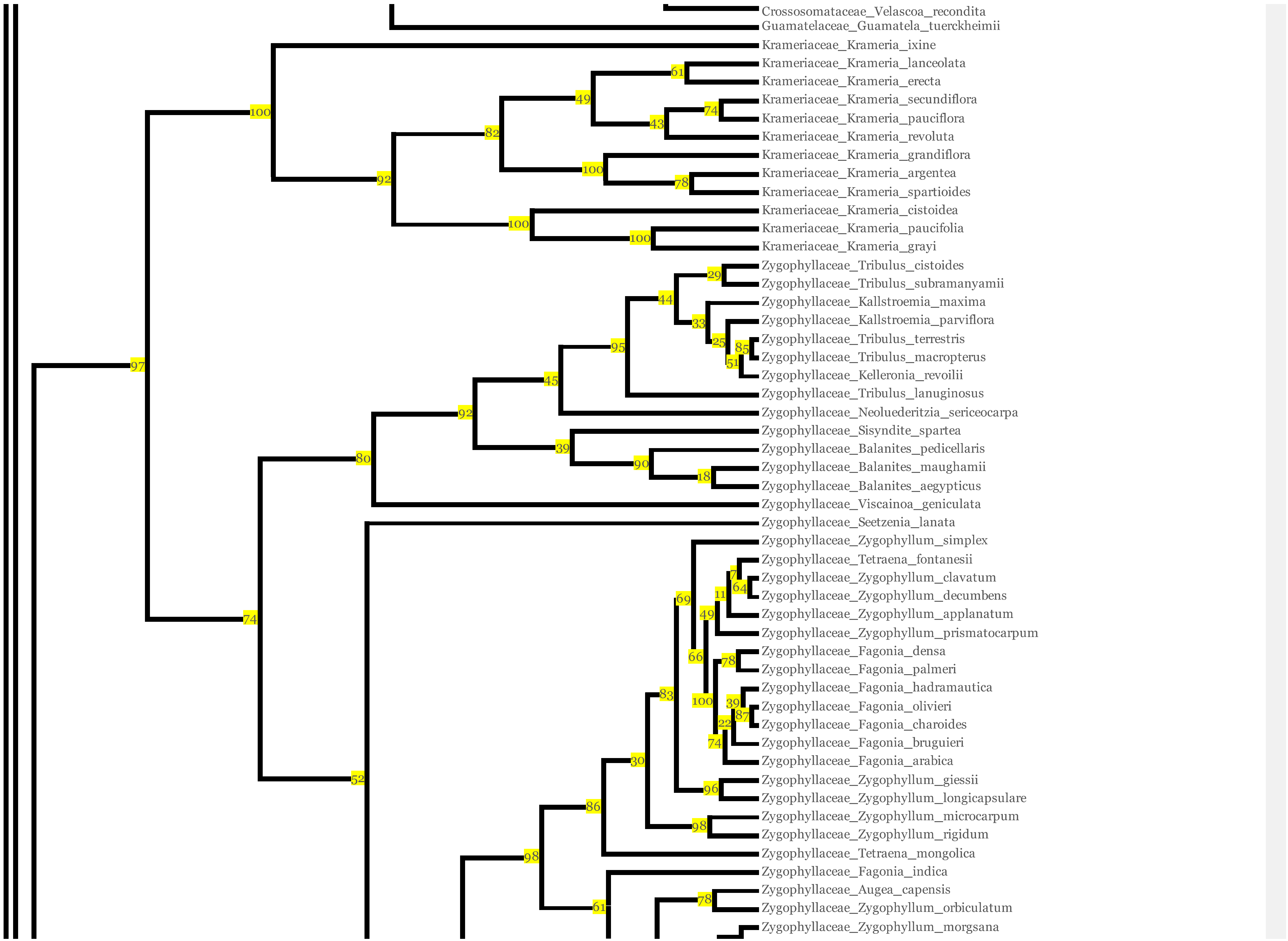
Crossosomataceae_Velascoa_recondita
Guamatelaceae_Guamatela_tuerckheimii
Krameriaceae_Krameria_ixine
Krameriaceae_Krameria_lanceolata
Krameriaceae_Krameria_erecta
Krameriaceae_Krameria_secundiflora
Krameriaceae_Krameria_pauciflora
Krameriaceae_Krameria_revoluta
Krameriaceae_Krameria_grandiflora
Krameriaceae_Krameria_argentea
Krameriaceae_Krameria_spartioides
Krameriaceae_Krameria_cistoidea
Krameriaceae_Krameria_paucifolia
Krameriaceae_Krameria_grayi
Zygophyllaceae_Tribulus_cistoides
Zygophyllaceae_Tribulus_subramanyamii
Zygophyllaceae_Kallstroemia_maxima
Zygophyllaceae_Kallstroemia_parviflora
Zygophyllaceae_Tribulus_terrestris
Zygophyllaceae_Tribulus_macropterus
Zygophyllaceae_Kelleronia_revoilii
Zygophyllaceae_Tribulus_lanuginosus
Zygophyllaceae_Neoluederitzia_sericeocarpa
Zygophyllaceae_Sisyndite_spartea
Zygophyllaceae_Balanites_pedicellaris
Zygophyllaceae_Balanites_maughamii
Zygophyllaceae_Balanites_aegypticus
Zygophyllaceae_Viscainoa_geniculata
Zygophyllaceae_Seetzenia_lanata
Zygophyllaceae_Zygophyllum_simplex
Zygophyllaceae_Tetraena_fontanesii
Zygophyllaceae_Zygophyllum_clavatum
Zygophyllaceae_Zygophyllum_decumbens
Zygophyllaceae_Zygophyllum_applanatum
Zygophyllaceae_Zygophyllum_prismatocarpum
Zygophyllaceae_Fagonia_densa
Zygophyllaceae_Fagonia_palmeri
Zygophyllaceae_Fagonia_hadramautica
Zygophyllaceae_Fagonia_olivieri
Zygophyllaceae_Fagonia_charoides
Zygophyllaceae_Fagonia_bruguieri
Zygophyllaceae_Fagonia_arabica
Zygophyllaceae_Zygophyllum_giessii
Zygophyllaceae_Zygophyllum_longicapsulare
Zygophyllaceae_Zygophyllum_microcarpum
Zygophyllaceae_Zygophyllum_rigidum
Zygophyllaceae_Tetraena_mongolica
Zygophyllaceae_Fagonia_indica
Zygophyllaceae_Augea_capensis
Zygophyllaceae_Zygophyllum_orbiculatum
Zygophyllaceae_Zygophyllum_morgsana

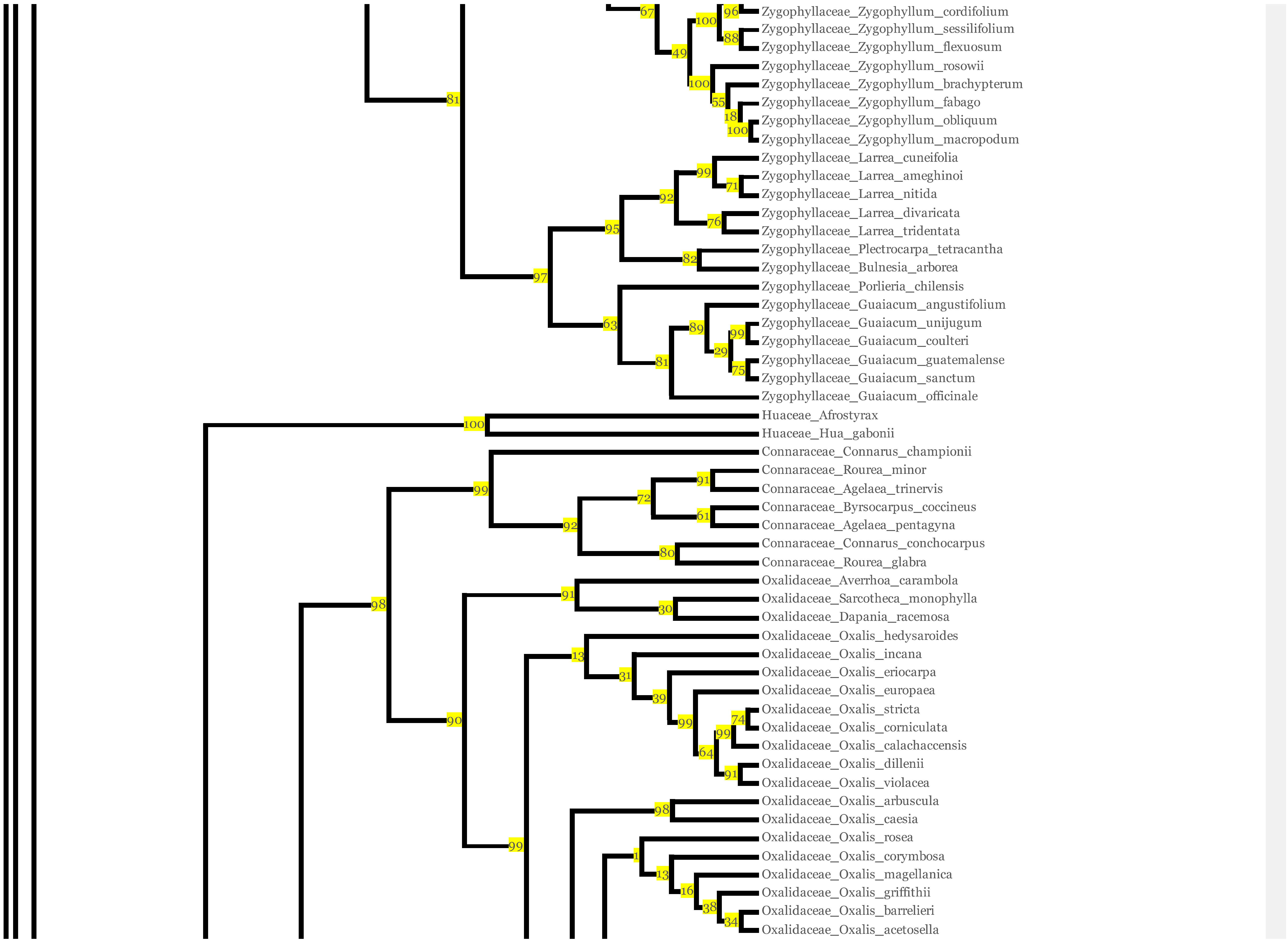
Zygophyllaceae_Zygophyllum_cordifolium
Zygophyllaceae_Zygophyllum_sessilifolium
Zygophyllaceae_Zygophyllum_flexuosum
Zygophyllaceae_Zygophyllum_rosowii
Zygophyllaceae_Zygophyllum_brachypterum
Zygophyllaceae_Zygophyllum_fabago
Zygophyllaceae_Zygophyllum_obliquum
Zygophyllaceae_Zygophyllum_macropodum
Zygophyllaceae_Larrea_cuneifolia
Zygophyllaceae_Larrea_ameghinoi
Zygophyllaceae_Larrea_nitida
Zygophyllaceae_Larrea_divaricata
Zygophyllaceae_Larrea_tridentata
Zygophyllaceae_Plectrocarpa_tetracantha
Zygophyllaceae_Bulnesia_arborea
Zygophyllaceae_Porlieria_chilensis
Zygophyllaceae_Guaiacum_angustifolium
Zygophyllaceae_Guaiacum_unijugum
Zygophyllaceae_Guaiacum_coulteri
Zygophyllaceae_Guaiacum_guatemalense
Zygophyllaceae_Guaiacum_sanctum
Zygophyllaceae_Guaiacum_officinale
Huaceae_Afrostyrax
Huaceae_Hua_gabonii
Connaraceae_Connarus_championii
Connaraceae_Rourea_minor
Connaraceae_Agelaea_trinervis
Connaraceae_Byrsocarpus_coccineus
Connaraceae_Agelaea_pentagyna
Connaraceae_Connarus_conchocarpus
Connaraceae_Rourea_glabra
Oxalidaceae_Averrhoa_carambola
Oxalidaceae_Sarcotheca_monophylla
Oxalidaceae_Dapania_racemosa
Oxalidaceae_Oxalis_hedysaroides
Oxalidaceae_Oxalis_incana
Oxalidaceae_Oxalis_eriocarpa
Oxalidaceae_Oxalis_europaea
Oxalidaceae_Oxalis_stricta
Oxalidaceae_Oxalis_corniculata
Oxalidaceae_Oxalis_calachaccensis
Oxalidaceae_Oxalis_dillenii
Oxalidaceae_Oxalis_violacea
Oxalidaceae_Oxalis_arbuscula
Oxalidaceae_Oxalis_caesia
Oxalidaceae_Oxalis_rosea
Oxalidaceae_Oxalis_corymbosa
Oxalidaceae_Oxalis_magellanica
Oxalidaceae_Oxalis_griffithii
Oxalidaceae_Oxalis_barrelieri
Oxalidaceae_Oxalis_acetosella

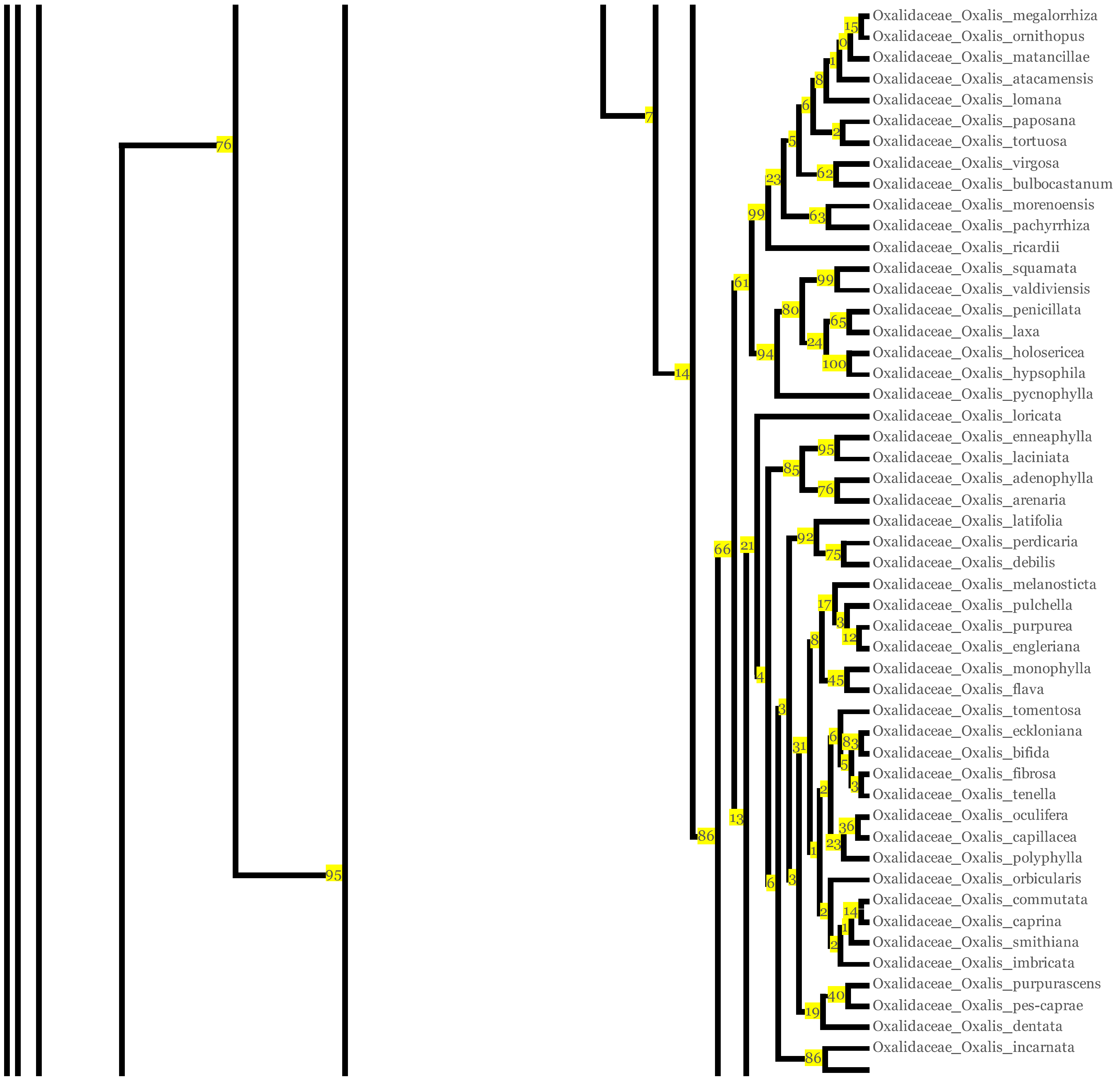
Oxalidaceae_Oxalis_megalorrhiza
Oxalidaceae_Oxalis_ornithopus
Oxalidaceae_Oxalis_matancillae
Oxalidaceae_Oxalis_atacamensis
Oxalidaceae_Oxalis_lomana
Oxalidaceae_Oxalis_paposana
Oxalidaceae_Oxalis_tortuosa
Oxalidaceae_Oxalis_virgosa
Oxalidaceae_Oxalis_bulbocastanum
Oxalidaceae_Oxalis_morenoensis
Oxalidaceae_Oxalis_pachyrrhiza
Oxalidaceae_Oxalis_ricardii
Oxalidaceae_Oxalis_squamata
Oxalidaceae_Oxalis_valdiviensis
Oxalidaceae_Oxalis_penicillata
Oxalidaceae_Oxalis_laxa
Oxalidaceae_Oxalis_holosericea
Oxalidaceae_Oxalis_hypsophila
Oxalidaceae_Oxalis_pycnophylla
Oxalidaceae_Oxalis_loricata
Oxalidaceae_Oxalis_enneaphylla
Oxalidaceae_Oxalis_laciniata
Oxalidaceae_Oxalis_adenophylla
Oxalidaceae_Oxalis_arenaria
Oxalidaceae_Oxalis_latifolia
Oxalidaceae_Oxalis_perdicaria
Oxalidaceae_Oxalis_debilis
Oxalidaceae_Oxalis_melanosticta
Oxalidaceae_Oxalis_pulchella
Oxalidaceae_Oxalis_purpurea
Oxalidaceae_Oxalis_engleriana
Oxalidaceae_Oxalis_monophylla
Oxalidaceae_Oxalis_flava
Oxalidaceae_Oxalis_tomentosa
Oxalidaceae_Oxalis_eckloniana
Oxalidaceae_Oxalis_bifida
Oxalidaceae_Oxalis_fibrosa
Oxalidaceae_Oxalis_tenella
Oxalidaceae_Oxalis_oculifera
Oxalidaceae_Oxalis_capillacea
Oxalidaceae_Oxalis_polyphylla
Oxalidaceae_Oxalis_orbicularis
Oxalidaceae_Oxalis_commutata
Oxalidaceae_Oxalis_caprina
Oxalidaceae_Oxalis_smithiana
Oxalidaceae_Oxalis_imbricata
Oxalidaceae_Oxalis_purpurascens
Oxalidaceae_Oxalis_pes-caprae
Oxalidaceae_Oxalis_dentata
Oxalidaceae_Oxalis_incarnata

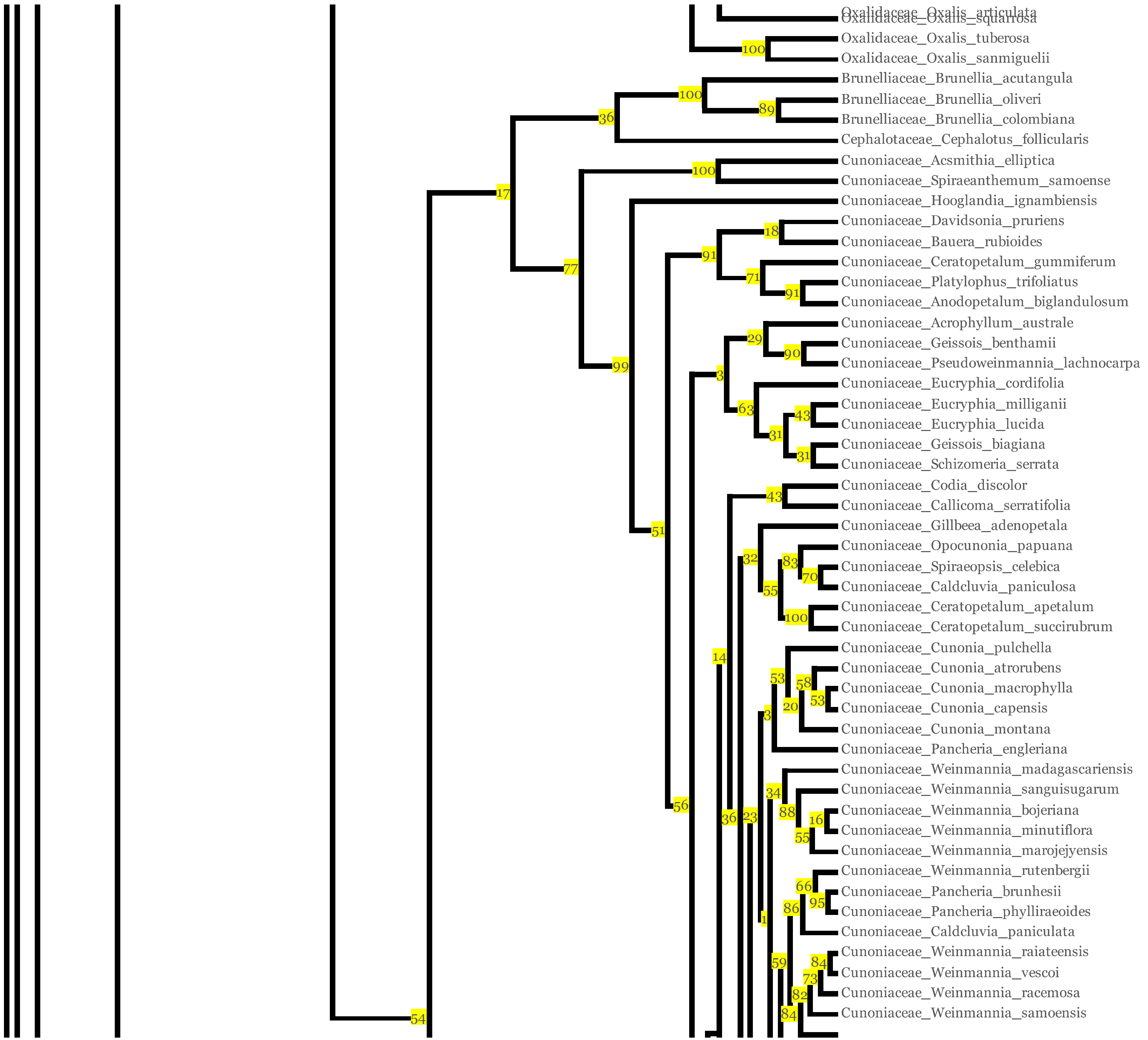
Oxalidaceae_Oxalis_articulata
Oxalidaceae_Oxalis_squarrosa
Oxalidaceae_Oxalis_tuberosa
Oxalidaceae_Oxalis_sanmiguelii
Brunelliaceae_Brunellia_acutangula
Brunelliaceae_Brunellia_oliveri
Brunelliaceae_Brunellia_colombiana
Cephalotaceae_Cephalotus_follicularis
Cunoniaceae_Acsmithia_elliptica
Cunoniaceae_Spiraeanthemum_samoense
Cunoniaceae_Hooglandia_ignambiensis
Cunoniaceae_Davidsonia_pruriens
Cunoniaceae_Bauera_rubioides
Cunoniaceae_Ceratopetalum_gummiferum
Cunoniaceae_Platylophus_trifoliatus
Cunoniaceae_Anodopetalum_biglandulosum
Cunoniaceae_Acrophyllum_australe
Cunoniaceae_Geissois_benthamii
Cunoniaceae_Pseudoweinmannia_lachnocarpa
Cunoniaceae_Eucryphia_cordifolia
Cunoniaceae_Eucryphia_milliganii
Cunoniaceae_Eucryphia_lucida
Cunoniaceae_Geissois_biagiana
Cunoniaceae_Schizomeria_serrata
Cunoniaceae_Codia_discolor
Cunoniaceae_Callicoma_serratifolia
Cunoniaceae_Gillbeea_adenopetala
Cunoniaceae_Opocunonia_papuana
Cunoniaceae_Spiraeopsis_celebica
Cunoniaceae_Caldcluvia_paniculosa
Cunoniaceae_Ceratopetalum_apetalum
Cunoniaceae_Ceratopetalum_succirubrum
Cunoniaceae_Cunonia_pulchella
Cunoniaceae_Cunonia_atrorubens
Cunoniaceae_Cunonia_macrophylla
Cunoniaceae_Cunonia_capensis
Cunoniaceae_Cunonia_montana
Cunoniaceae_Pancheria_engleriana
Cunoniaceae_Weinmannia_madagascariensis
Cunoniaceae_Weinmannia_sanguisugarum
Cunoniaceae_Weinmannia_bojeriana
Cunoniaceae_Weinmannia_minutiflora
Cunoniaceae_Weinmannia_marojejyensis
Cunoniaceae_Weinmannia_rutenbergii
Cunoniaceae_Pancheria_brunhesii
Cunoniaceae_Pancheria_phylliraeoides
Cunoniaceae_Caldcluvia_paniculata
Cunoniaceae_Weinmannia_raiateensis
Cunoniaceae_Weinmannia_vescoi
Cunoniaceae_Weinmannia_racemosa
Cunoniaceae_Weinmannia_samoensis

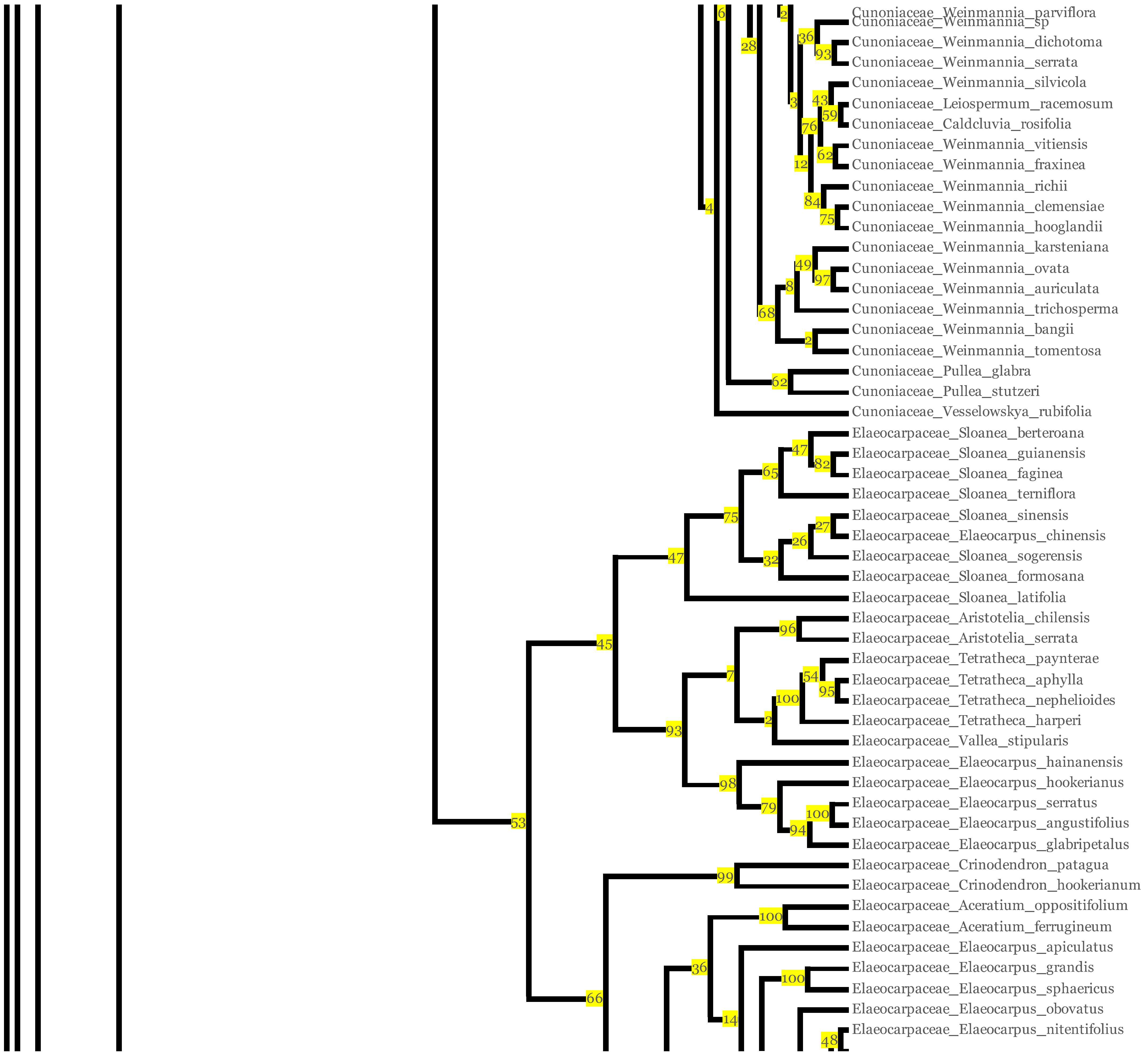
Cunoniaceae_Weinmannia_parviflora
Cunoniaceae_Weinmannia_sp
Cunoniaceae_Weinmannia_dichotoma
Cunoniaceae_Weinmannia_serrata
Cunoniaceae_Weinmannia_silvicola
Cunoniaceae_Leiospermum_racemosum
Cunoniaceae_Caldcluvia_rosifolia
Cunoniaceae_Weinmannia_vitiensis
Cunoniaceae_Weinmannia_fraxinea
Cunoniaceae_Weinmannia_richii
Cunoniaceae_Weinmannia_clemensiae
Cunoniaceae_Weinmannia_hooglandii
Cunoniaceae_Weinmannia_karsteniana
Cunoniaceae_Weinmannia_ovata
Cunoniaceae_Weinmannia_auriculata
Cunoniaceae_Weinmannia_trichosperma
Cunoniaceae_Weinmannia_bangii
Cunoniaceae_Weinmannia_tomentosa
Cunoniaceae_Pullea_glabra
Cunoniaceae_Pullea_stutzeri
Cunoniaceae_Vesselowskya_rubifolia
Elaeocarpaceae_Sloanea_berteroana
Elaeocarpaceae_Sloanea_guianensis
Elaeocarpaceae_Sloanea_faginea
Elaeocarpaceae_Sloanea_terniflora
Elaeocarpaceae_Sloanea_sinensis
Elaeocarpaceae_Elaeocarpus_chinensis
Elaeocarpaceae_Sloanea_sogerensis
Elaeocarpaceae_Sloanea_formosana
Elaeocarpaceae_Sloanea_latifolia
Elaeocarpaceae_Aristotelia_chilensis
Elaeocarpaceae_Aristotelia_serrata
Elaeocarpaceae_Tetratheca_paynterae
Elaeocarpaceae_Tetratheca_aphylla
Elaeocarpaceae_Tetratheca_nephelioides
Elaeocarpaceae_Tetratheca_harperi
Elaeocarpaceae_Vallea_stipularis
Elaeocarpaceae_Elaeocarpus_hainanensis
Elaeocarpaceae_Elaeocarpus_hookerianus
Elaeocarpaceae_Elaeocarpus_serratus
Elaeocarpaceae_Elaeocarpus_angustifolius
Elaeocarpaceae_Elaeocarpus_glabripetalus
Elaeocarpaceae_Crinodendron_patagua
Elaeocarpaceae_Crinodendron_hookerianum
Elaeocarpaceae_Aceratium_oppositifolium
Elaeocarpaceae_Aceratium_ferrugineum
Elaeocarpaceae_Elaeocarpus_apiculatus
Elaeocarpaceae_Elaeocarpus_grandis
Elaeocarpaceae_Elaeocarpus_sphaericus
Elaeocarpaceae_Elaeocarpus_obovatus
Elaeocarpaceae_Elaeocarpus_nitentifolius

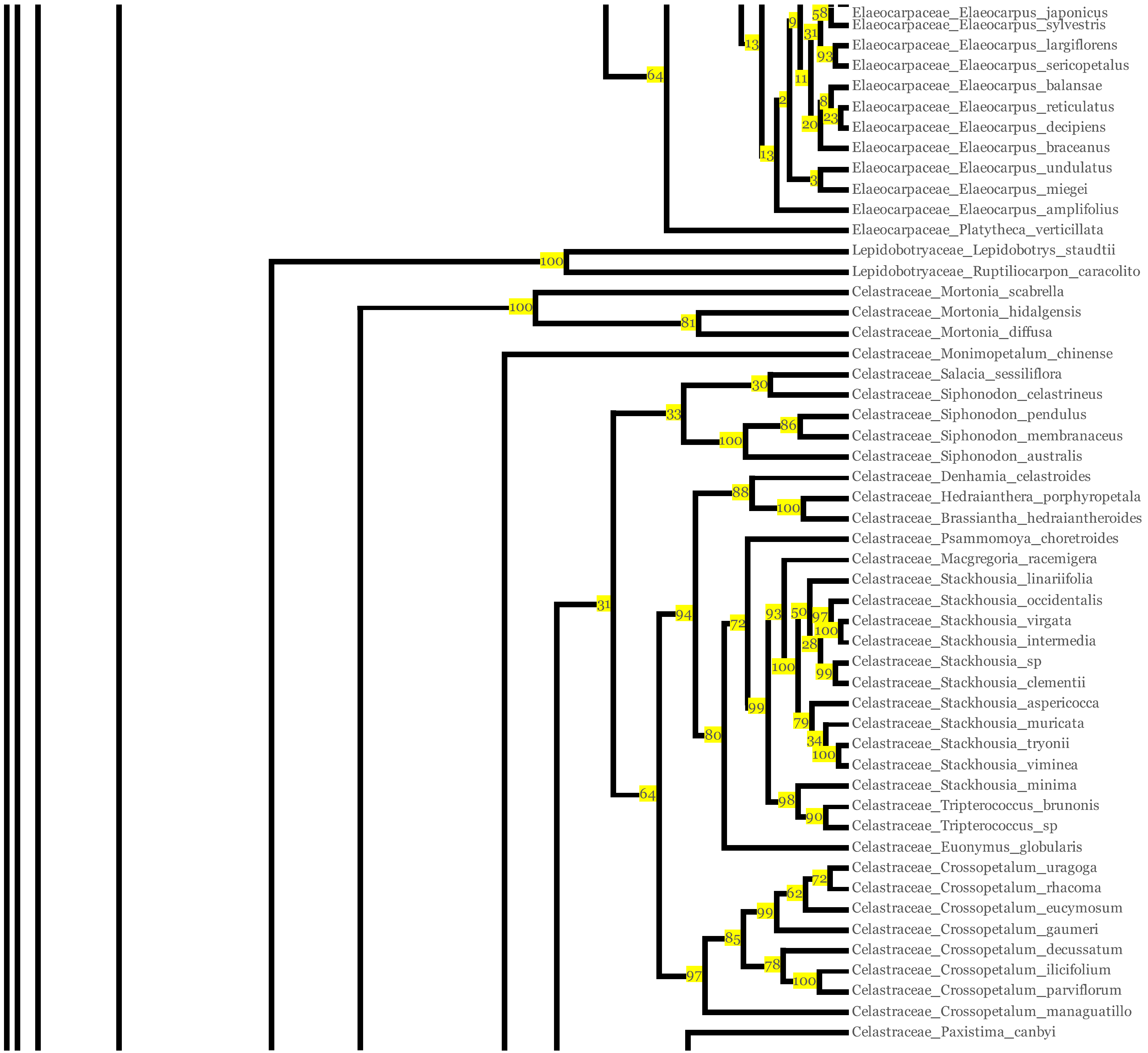
Elaeocarpaceae_Elaeocarpus_japonicus
Elaeocarpaceae_Elaeocarpus_sylvestris
Elaeocarpaceae_Elaeocarpus_largiflorens
Elaeocarpaceae_Elaeocarpus_sericopetalus
Elaeocarpaceae_Elaeocarpus_balansae
Elaeocarpaceae_Elaeocarpus_reticulatus
Elaeocarpaceae_Elaeocarpus_decipiens
Elaeocarpaceae_Elaeocarpus_braceanus
Elaeocarpaceae_Elaeocarpus_undulatus
Elaeocarpaceae_Elaeocarpus_miegei
Elaeocarpaceae_Elaeocarpus_amplifolius
Elaeocarpaceae_Platytheca_verticillata
Lepidobotryaceae_Lepidobotrys_staudtii
Lepidobotryaceae_Ruptiliocarpon_caracolito
Celastraceae_Mortonia_scabrella
Celastraceae_Mortonia_hidalgensis
Celastraceae_Mortonia_diffusa
Celastraceae_Monimopetalum_chinense
Celastraceae_Salacia_sessiliflora
Celastraceae_Siphonodon_celastrineus
Celastraceae_Siphonodon_pendulus
Celastraceae_Siphonodon_membranaceus
Celastraceae_Siphonodon_australis
Celastraceae_Denhamia_celastroides
Celastraceae_Hedraianthera_porphyropetala
Celastraceae_Brassiantha_hedraiantheroides
Celastraceae_Psammomoya_choretroides
Celastraceae_Macgregoria_racemigera
Celastraceae_Stackhousia_linariifolia
Celastraceae_Stackhousia_occidentalis
Celastraceae_Stackhousia_virgata
Celastraceae_Stackhousia_intermedia
Celastraceae_Stackhousia_sp
Celastraceae_Stackhousia_clementii
Celastraceae_Stackhousia_aspericocca
Celastraceae_Stackhousia_muricata
Celastraceae_Stackhousia_tryonii
Celastraceae_Stackhousia_viminea
Celastraceae_Stackhousia_minima
Celastraceae_Tripterococcus_brunonis
Celastraceae_Tripterococcus_sp
Celastraceae_Euonymus_globularis
Celastraceae_Crossopetalum_uragoga
Celastraceae_Crossopetalum_rhacoma
Celastraceae_Crossopetalum_eucymosum
Celastraceae_Crossopetalum_gaumeri
Celastraceae_Crossopetalum_decussatum
Celastraceae_Crossopetalum_ilicifolium
Celastraceae_Crossopetalum_parviflorum
Celastraceae_Crossopetalum_managuatillo
Celastraceae_Paxistima_canbyi

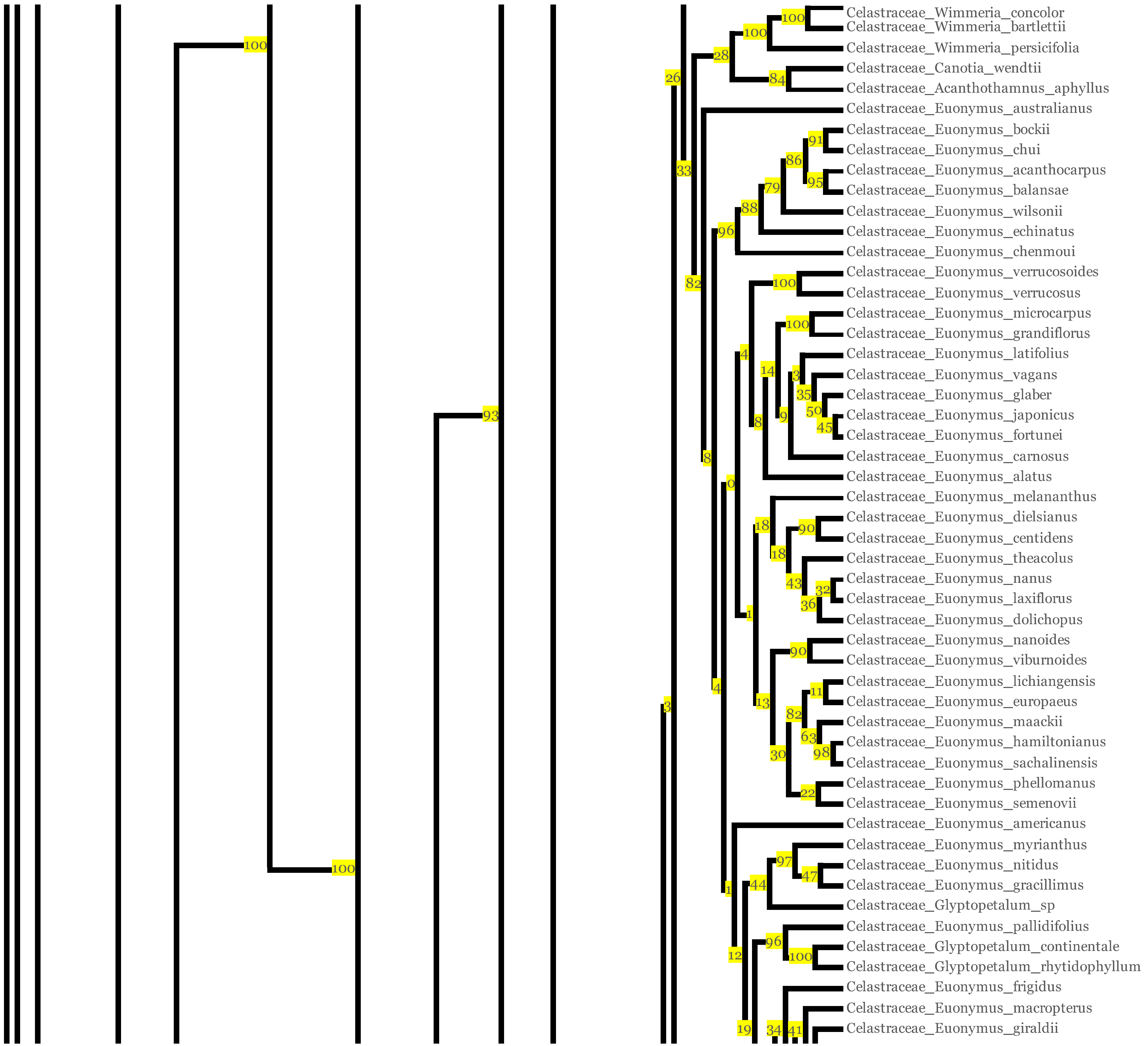
Celastraceae_Wimmeria_concolor
Celastraceae_Wimmeria_bartlettii
Celastraceae_Wimmeria_persicifolia
Celastraceae_Canotia_wendtii
Celastraceae_Acanthothamnus_aphyllus
Celastraceae_Euonymus_australianus
Celastraceae_Euonymus_bockii
Celastraceae_Euonymus_chui
Celastraceae_Euonymus_acanthocarpus
Celastraceae_Euonymus_balansae
Celastraceae_Euonymus_wilsonii
Celastraceae_Euonymus_echinatus
Celastraceae_Euonymus_chenmoui
Celastraceae_Euonymus_verrucosoides
Celastraceae_Euonymus_verrucosus
Celastraceae_Euonymus_microcarpus
Celastraceae_Euonymus_grandiflorus
Celastraceae_Euonymus_latifolius
Celastraceae_Euonymus_vagans
Celastraceae_Euonymus_glaber
Celastraceae_Euonymus_japonicus
Celastraceae_Euonymus_fortunei
Celastraceae_Euonymus_carnosus
Celastraceae_Euonymus_alatus
Celastraceae_Euonymus_melananthus
Celastraceae_Euonymus_dielsianus
Celastraceae_Euonymus_centidens
Celastraceae_Euonymus_theacolus
Celastraceae_Euonymus_nanus
Celastraceae_Euonymus_laxiflorus
Celastraceae_Euonymus_dolichopus
Celastraceae_Euonymus_nanoides
Celastraceae_Euonymus_viburnoides
Celastraceae_Euonymus_lichiangensis
Celastraceae_Euonymus_europaeus
Celastraceae_Euonymus_maackii
Celastraceae_Euonymus_hamiltonianus
Celastraceae_Euonymus_sachalinensis
Celastraceae_Euonymus_phellomanus
Celastraceae_Euonymus_semenovii
Celastraceae_Euonymus_americanus
Celastraceae_Euonymus_myrianthus
Celastraceae_Euonymus_nitidus
Celastraceae_Euonymus_gracillimus
Celastraceae_Glyptopetalum_sp
Celastraceae_Euonymus_pallidifolius
Celastraceae_Glyptopetalum_continentale
Celastraceae_Glyptopetalum_rhytidophyllum
Celastraceae_Euonymus_frigidus
Celastraceae_Euonymus_macropterus
Celastraceae_Euonymus_giraldii

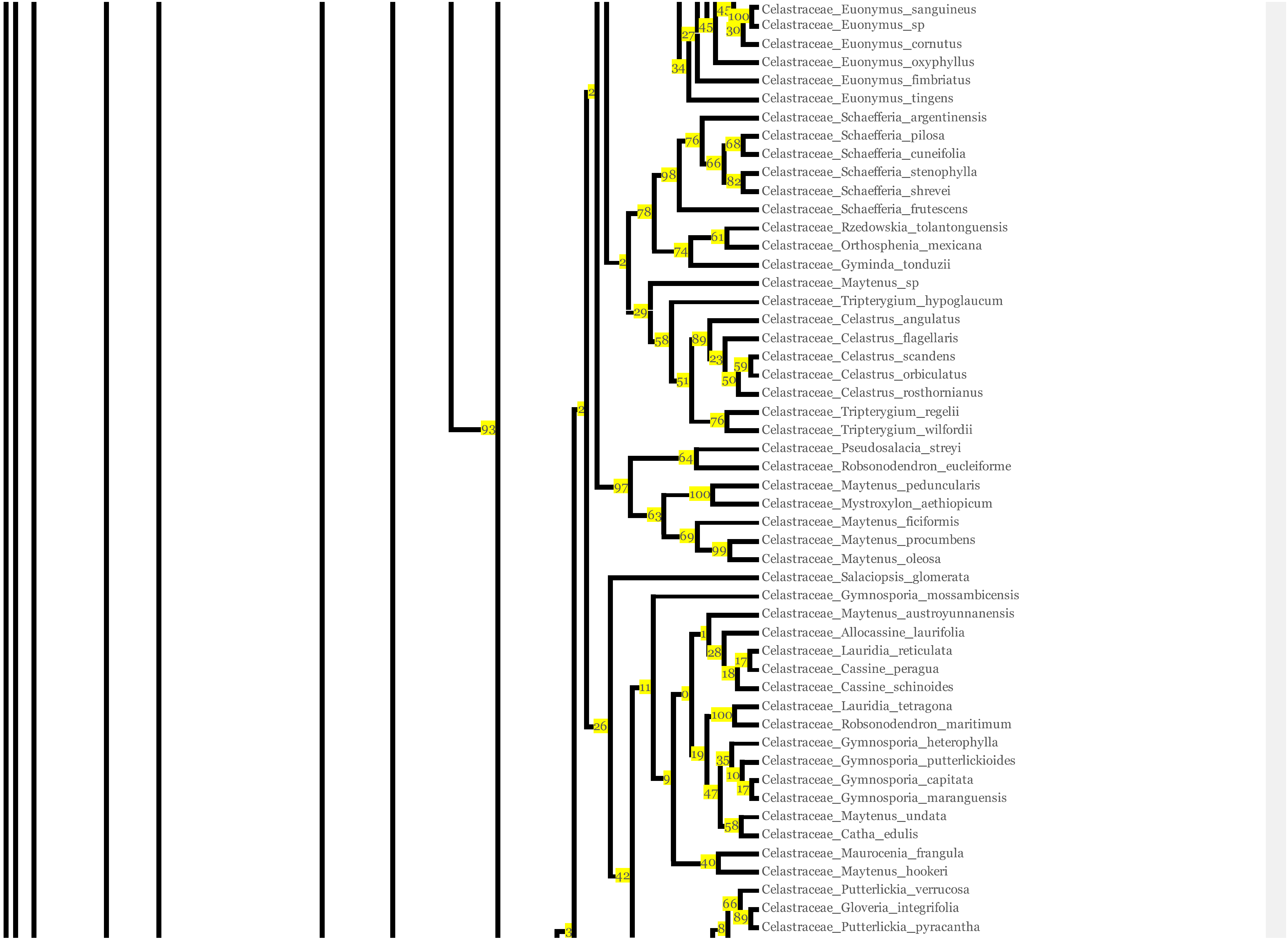
Celastraceae_Euonymus_sanguineus
Celastraceae_Euonymus_sp
Celastraceae_Euonymus_cornutus
Celastraceae_Euonymus_oxyphyllus
Celastraceae_Euonymus_fimbriatus
Celastraceae_Euonymus_tingens
Celastraceae_Schaefferia_argentinensis
Celastraceae_Schaefferia_pilosa
Celastraceae_Schaefferia_cuneifolia
Celastraceae_Schaefferia_stenophylla
Celastraceae_Schaefferia_shrevei
Celastraceae_Schaefferia_frutescens
Celastraceae_Rzedowskia_tolantonguensis
Celastraceae_Orthosphenia_mexicana
Celastraceae_Gyminda_tonduzii
Celastraceae_Maytenus_sp
Celastraceae_Tripterygium_hypoglaucum
Celastraceae_Celastrus_angulatus
Celastraceae_Celastrus_flagellaris
Celastraceae_Celastrus_scandens
Celastraceae_Celastrus_orbiculatus
Celastraceae_Celastrus_rosthornianus
Celastraceae_Tripterygium_regelii
Celastraceae_Tripterygium_wilfordii
Celastraceae_Pseudosalacia_streyi
Celastraceae_Robsonodendron_eucleiforme
Celastraceae_Maytenus_peduncularis
Celastraceae_Mystroxylon_aethiopicum
Celastraceae_Maytenus_ficiformis
Celastraceae_Maytenus_procumbens
Celastraceae_Maytenus_oleosa
Celastraceae_Salaciopsis_glomerata
Celastraceae_Gymnosporia_mossambicensis
Celastraceae_Maytenus_austroyunnanensis
Celastraceae_Allocassine_laurifolia
Celastraceae_Lauridia_reticulata
Celastraceae_Cassine_peragua
Celastraceae_Cassine_schinoides
Celastraceae_Lauridia_tetragona
Celastraceae_Robsonodendron_maritimum
Celastraceae_Gymnosporia_heterophylla
Celastraceae_Gymnosporia_putterlickioides
Celastraceae_Gymnosporia_capitata
Celastraceae_Gymnosporia_maranguensis
Celastraceae_Maytenus_undata
Celastraceae_Catha_edulis
Celastraceae_Maurocenia_frangula
Celastraceae_Maytenus_hookeri
Celastraceae_Putterlickia_verrucosa
Celastraceae_Gloveria_integrifolia
Celastraceae_Putterlickia_pyracantha

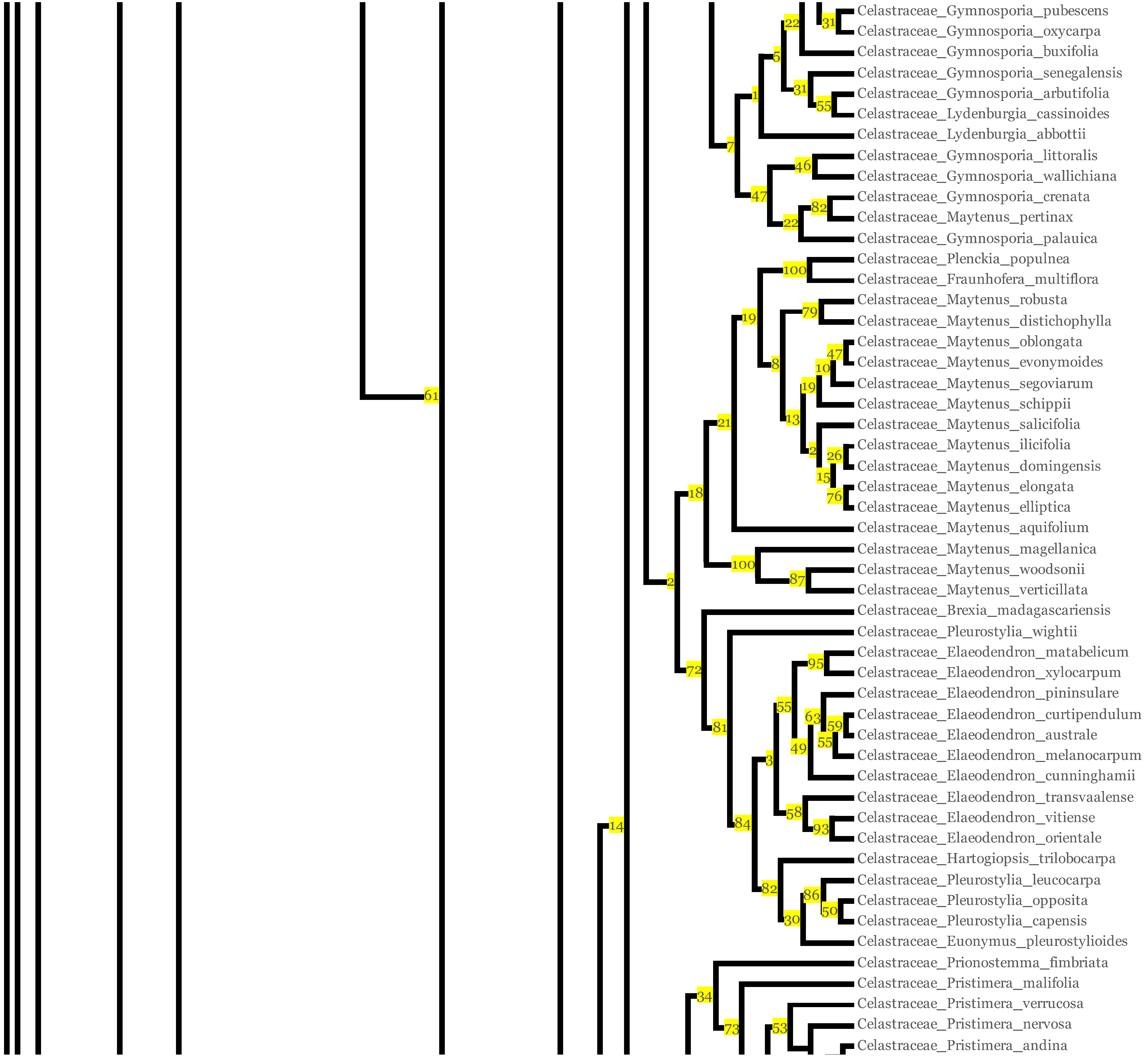

Celastraceae_Gymnosporia_pubescens
Celastraceae_Gymnosporia_oxycarpa
Celastraceae_Gymnosporia_buxifolia
Celastraceae_Gymnosporia_senegalensis
Celastraceae_Gymnosporia_arbutifolia
Celastraceae_Lydenburgia_cassinoides
Celastraceae_Lydenburgia_abbottii
Celastraceae_Gymnosporia_littoralis
Celastraceae_Gymnosporia_wallichiana
Celastraceae_Gymnosporia_crenata
Celastraceae_Maytenus_pertinax
Celastraceae_Gymnosporia_palauica
Celastraceae_Plenckia_populnea
Celastraceae_Fraunhofera_multiflora
Celastraceae_Maytenus_robusta
Celastraceae_Maytenus_distichophylla
Celastraceae_Maytenus_oblongata
Celastraceae_Maytenus_evonymoides
Celastraceae_Maytenus_segoviarum
Celastraceae_Maytenus_schippii
Celastraceae_Maytenus_salicifolia
Celastraceae_Maytenus_ilicifolia
Celastraceae_Maytenus_domingensis
Celastraceae_Maytenus_elongata
Celastraceae_Maytenus_elliptica
Celastraceae_Maytenus_aquifolium
Celastraceae_Maytenus_magellanica
Celastraceae_Maytenus_woodsonii
Celastraceae_Maytenus_verticillata
Celastraceae_Brexia_madagascariensis
Celastraceae_Pleurostylia_wightii
Celastraceae_Elaeodendron_matabelicum
Celastraceae_Elaeodendron_xylocarpum
Celastraceae_Elaeodendron_pininsulare
Celastraceae_Elaeodendron_curtipendulum
Celastraceae_Elaeodendron_australe
Celastraceae_Elaeodendron_melanocarpum
Celastraceae_Elaeodendron_cunninghamii
Celastraceae_Elaeodendron_transvaalense
Celastraceae_Elaeodendron_vitiense
Celastraceae_Elaeodendron_orientale
Celastraceae_Hartogiopsis_trilobocarpa
Celastraceae_Pleurostylia_leucocarpa
Celastraceae_Pleurostylia_opposita
Celastraceae_Pleurostylia_capensis
Celastraceae_Euonymus_pleurostylioides
Celastraceae_Prionostemma_fimbriata
Celastraceae_Pristimera_malifolia
Celastraceae_Pristimera_verrucosa
Celastraceae_Pristimera_nervosa
Celastraceae_Pristimera_andina

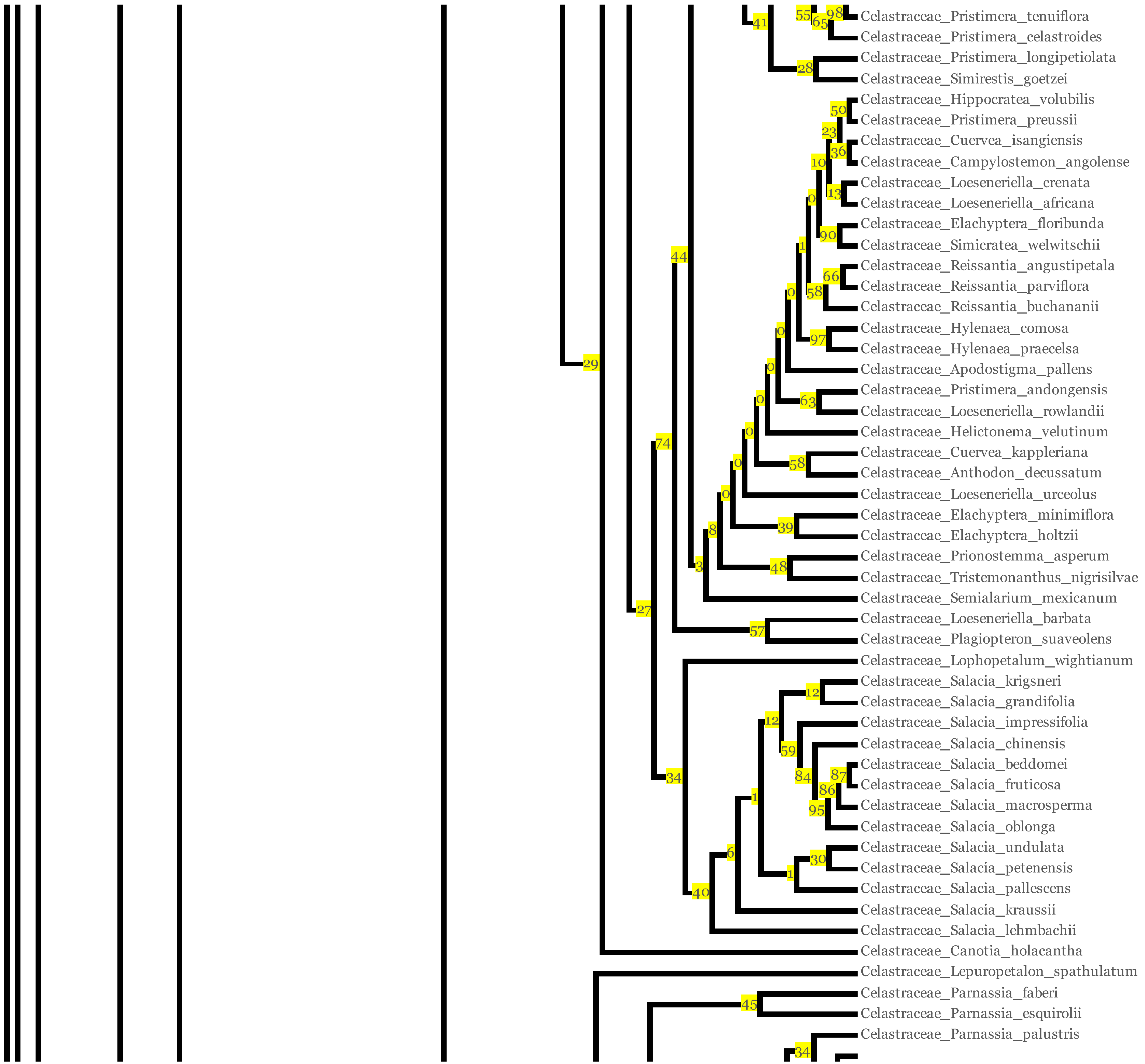
Celastraceae_Pristimera_tenuiflora
Celastraceae_Pristimera_celastroides
Celastraceae_Pristimera_longipetiolata
Celastraceae_Simirestis_goetzei
Celastraceae_Hippocratea_volubilis
Celastraceae_Pristimera_preussii
Celastraceae_Cuervea_isangiensis
Celastraceae_Campylostemon_angolense
Celastraceae_Loeseneriella_crenata
Celastraceae_Loeseneriella_africana
Celastraceae_Elachyptera_floribunda
Celastraceae_Simicratea_welwitschii
Celastraceae_Reissantia_angustipetala
Celastraceae_Reissantia_parviflora
Celastraceae_Reissantia_buchananii
Celastraceae_Hylenaea_comosa
Celastraceae_Hylenaea_praecelsa
Celastraceae_Apodostigma_pallens
Celastraceae_Pristimera_andongensis
Celastraceae_Loeseneriella_rowlandii
Celastraceae_Helictonema_velutinum
Celastraceae_Cuervea_kappleriana
Celastraceae_Anthodon_decussatum
Celastraceae_Loeseneriella_urceolus
Celastraceae_Elachyptera_minimiflora
Celastraceae_Elachyptera_holtzii
Celastraceae_Prionostemma_asperum
Celastraceae_Tristemonanthus_nigrisilvae
Celastraceae_Semialarium_mexicanum
Celastraceae_Loeseneriella_barbata
Celastraceae_Plagiopteron_suaveolens
Celastraceae_Lophopetalum_wightianum
Celastraceae_Salacia_krigsneri
Celastraceae_Salacia_grandifolia
Celastraceae_Salacia_impressifolia
Celastraceae_Salacia_chinensis
Celastraceae_Salacia_beddomei
Celastraceae_Salacia_fruticosa
Celastraceae_Salacia_macrosperma
Celastraceae_Salacia_oblonga
Celastraceae_Salacia_undulata
Celastraceae_Salacia_petenensis
Celastraceae_Salacia_pallescens
Celastraceae_Salacia_kraussii
Celastraceae_Salacia_lehmbachii
Celastraceae_Canotia_holacantha
Celastraceae_Lepuropetalon_spathulatum
Celastraceae_Parnassia_faberi
Celastraceae_Parnassia_esquirolii
Celastraceae_Parnassia_palustris

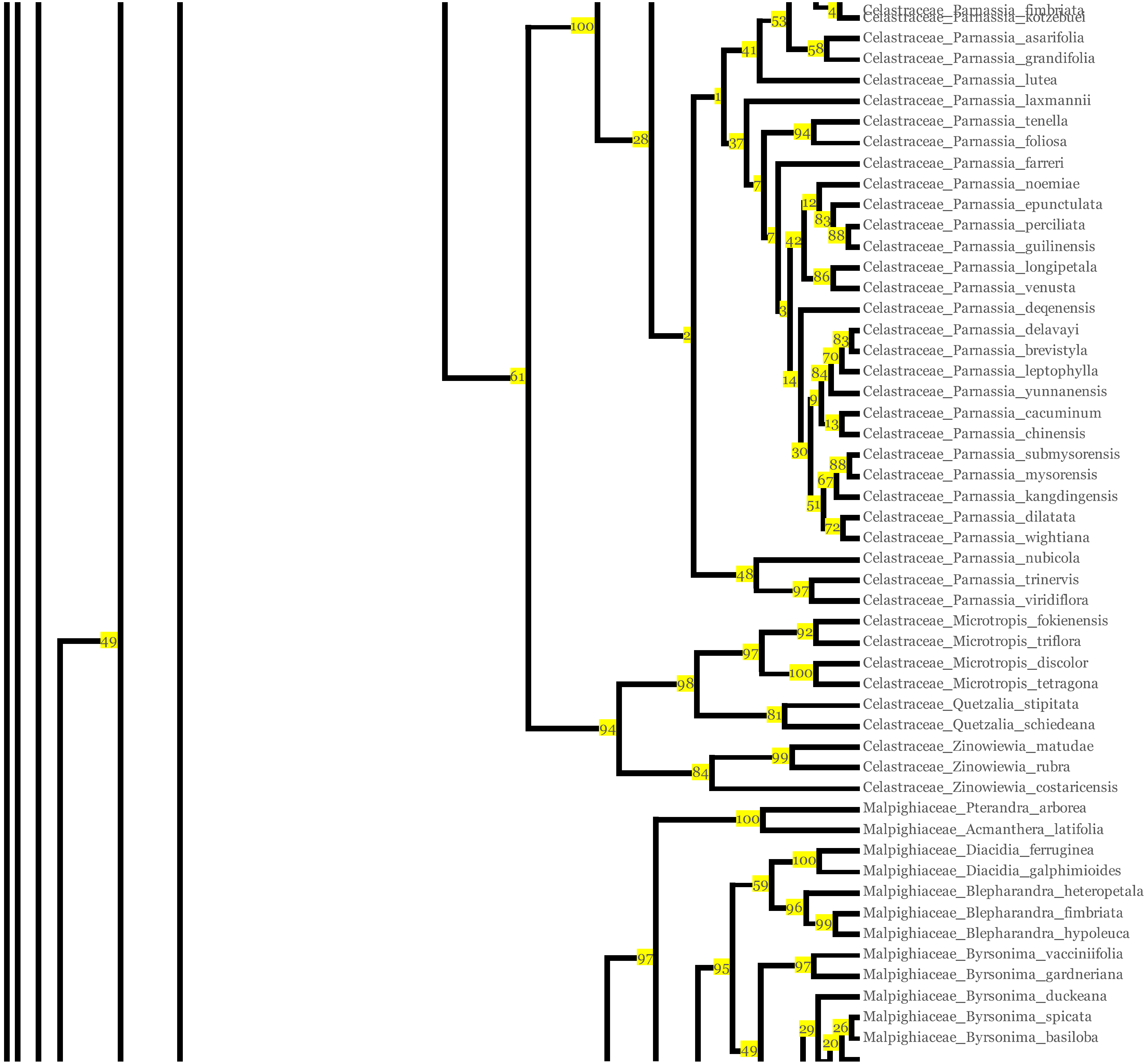
Celastraceae_Parnassia_fimbriata
Celastraceae_Parnassia_kotzebuei
Celastraceae_Parnassia_asarifolia
Celastraceae_Parnassia_grandifolia
Celastraceae_Parnassia_lutea
Celastraceae_Parnassia_laxmannii
Celastraceae_Parnassia_tenella
Celastraceae_Parnassia_foliosa
Celastraceae_Parnassia_farreri
Celastraceae_Parnassia_noemiae
Celastraceae_Parnassia_epunctulata
Celastraceae_Parnassia_perciliata
Celastraceae_Parnassia_guilinensis
Celastraceae_Parnassia_longipetala
Celastraceae_Parnassia_venusta
Celastraceae_Parnassia_deqenensis
Celastraceae_Parnassia_delavayi
Celastraceae_Parnassia_brevistyla
Celastraceae_Parnassia_leptophylla
Celastraceae_Parnassia_yunnanensis
Celastraceae_Parnassia_cacuminum
Celastraceae_Parnassia_chinensis
Celastraceae_Parnassia_submysorensis
Celastraceae_Parnassia_mysorensis
Celastraceae_Parnassia_kangdingensis
Celastraceae_Parnassia_dilatata
Celastraceae_Parnassia_wightiana
Celastraceae_Parnassia_nubicola
Celastraceae_Parnassia_trinervis
Celastraceae_Parnassia_viridiflora
Celastraceae_Microtropis_fokienensis
Celastraceae_Microtropis_triflora
Celastraceae_Microtropis_discolor
Celastraceae_Microtropis_tetragona
Celastraceae_Quetzalia_stipitata
Celastraceae_Quetzalia_schiedeana
Celastraceae_Zinowiewia_matudae
Celastraceae_Zinowiewia_rubra
Celastraceae_Zinowiewia_costaricensis
Malpighiaceae_Pterandra_arborea
Malpighiaceae_Acmanthera_latifolia
Malpighiaceae_Diacidia_ferruginea
Malpighiaceae_Diacidia_galphimioides
Malpighiaceae_Blepharandra_heteropetala
Malpighiaceae_Blepharandra_fimbriata
Malpighiaceae_Blepharandra_hypoleuca
Malpighiaceae_Byrsonima_vacciniifolia
Malpighiaceae_Byrsonima_gardneriana
Malpighiaceae_Byrsonima_duckeana
Malpighiaceae_Byrsonima_spicata
Malpighiaceae_Byrsonima_basiloba

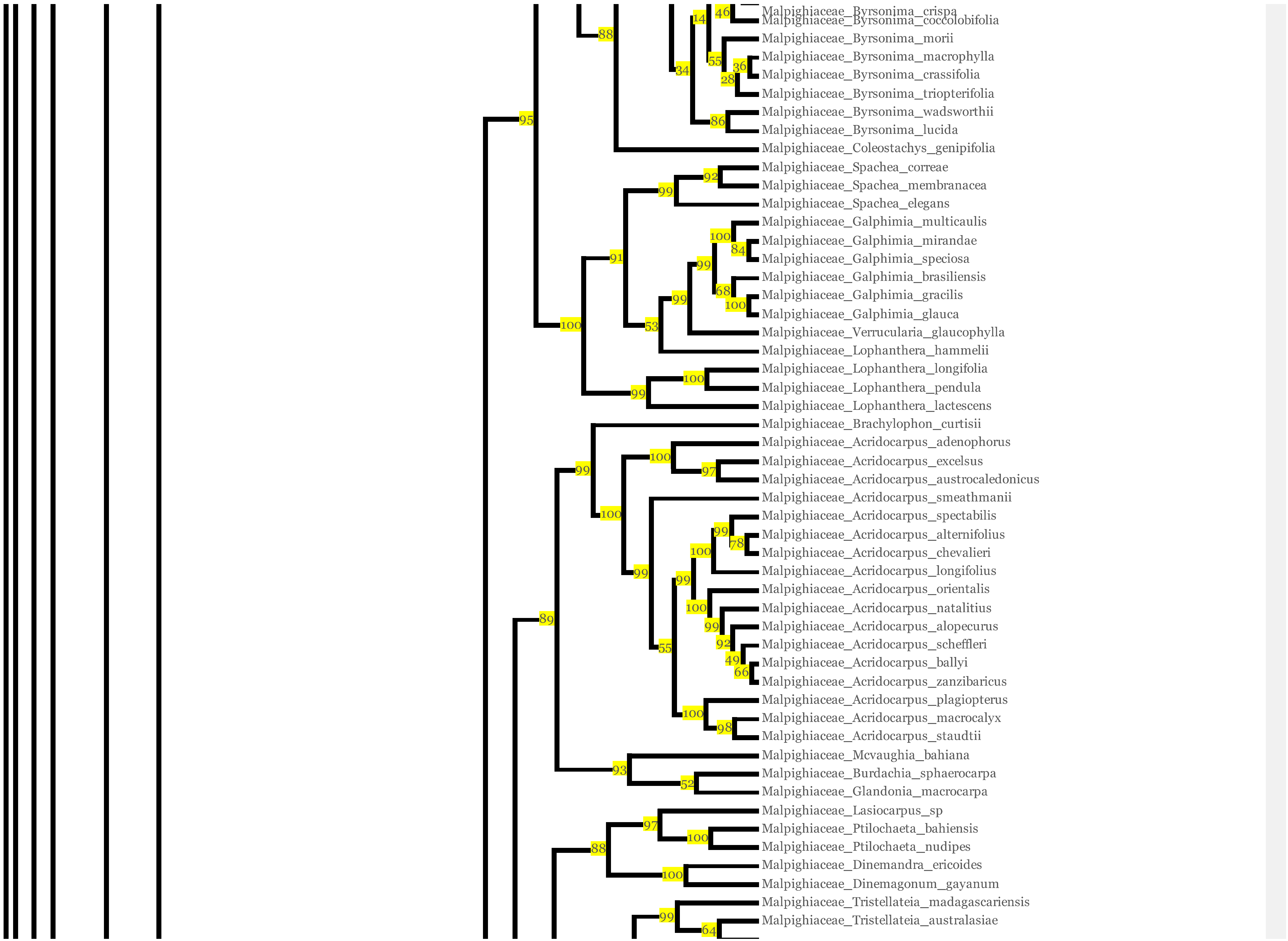

Malpighiaceae_Byrsonima_crispa
Malpighiaceae_Byrsonima_coccolobifolia
Malpighiaceae_Byrsonima_morii
Malpighiaceae_Byrsonima_macrophylla
Malpighiaceae_Byrsonima_crassifolia
Malpighiaceae_Byrsonima_triopterifolia
Malpighiaceae_Byrsonima_wadsworthii
Malpighiaceae_Byrsonima_lucida
Malpighiaceae_Coleostachys_genipifolia
Malpighiaceae_Spachea_correae
Malpighiaceae_Spachea_membranacea
Malpighiaceae_Spachea_elegans
Malpighiaceae_Galphimia_multicaulis
Malpighiaceae_Galphimia_mirandae
Malpighiaceae_Galphimia_speciosa
Malpighiaceae_Galphimia_brasiliensis
Malpighiaceae_Galphimia_gracilis
Malpighiaceae_Galphimia_glauca
Malpighiaceae_Verrucularia_glaucophylla
Malpighiaceae_Lophanthera_hammelii
Malpighiaceae_Lophanthera_longifolia
Malpighiaceae_Lophanthera_pendula
Malpighiaceae_Lophanthera_lactescens
Malpighiaceae_Brachylophon_curtisii
Malpighiaceae_Acridocarpus_adenophorus
Malpighiaceae_Acridocarpus_excelsus
Malpighiaceae_Acridocarpus_austrocaledonicus
Malpighiaceae_Acridocarpus_smeathmanii
Malpighiaceae_Acridocarpus_spectabilis
Malpighiaceae_Acridocarpus_alternifolius
Malpighiaceae_Acridocarpus_chevalieri
Malpighiaceae_Acridocarpus_longifolius
Malpighiaceae_Acridocarpus_orientalis
Malpighiaceae_Acridocarpus_natalitius
Malpighiaceae_Acridocarpus_alopecurus
Malpighiaceae_Acridocarpus_scheffleri
Malpighiaceae_Acridocarpus_ballyi
Malpighiaceae_Acridocarpus_zanzibaricus
Malpighiaceae_Acridocarpus_plagiopterus
Malpighiaceae_Acridocarpus_macrocalyx
Malpighiaceae_Acridocarpus_staudtii
Malpighiaceae_Mcvaughia_bahiana
Malpighiaceae_Burdachia_sphaerocarpa
Malpighiaceae_Glandonia_macrocarpa
Malpighiaceae_Lasiocarpus_sp
Malpighiaceae_Ptilochaeta_bahiensis
Malpighiaceae_Ptilochaeta_nudipes
Malpighiaceae_Dinemandra_ericoides
Malpighiaceae_Dinemagonum_gayanum
Malpighiaceae_Tristellateia_madagascariensis
Malpighiaceae_Tristellateia_australasiae

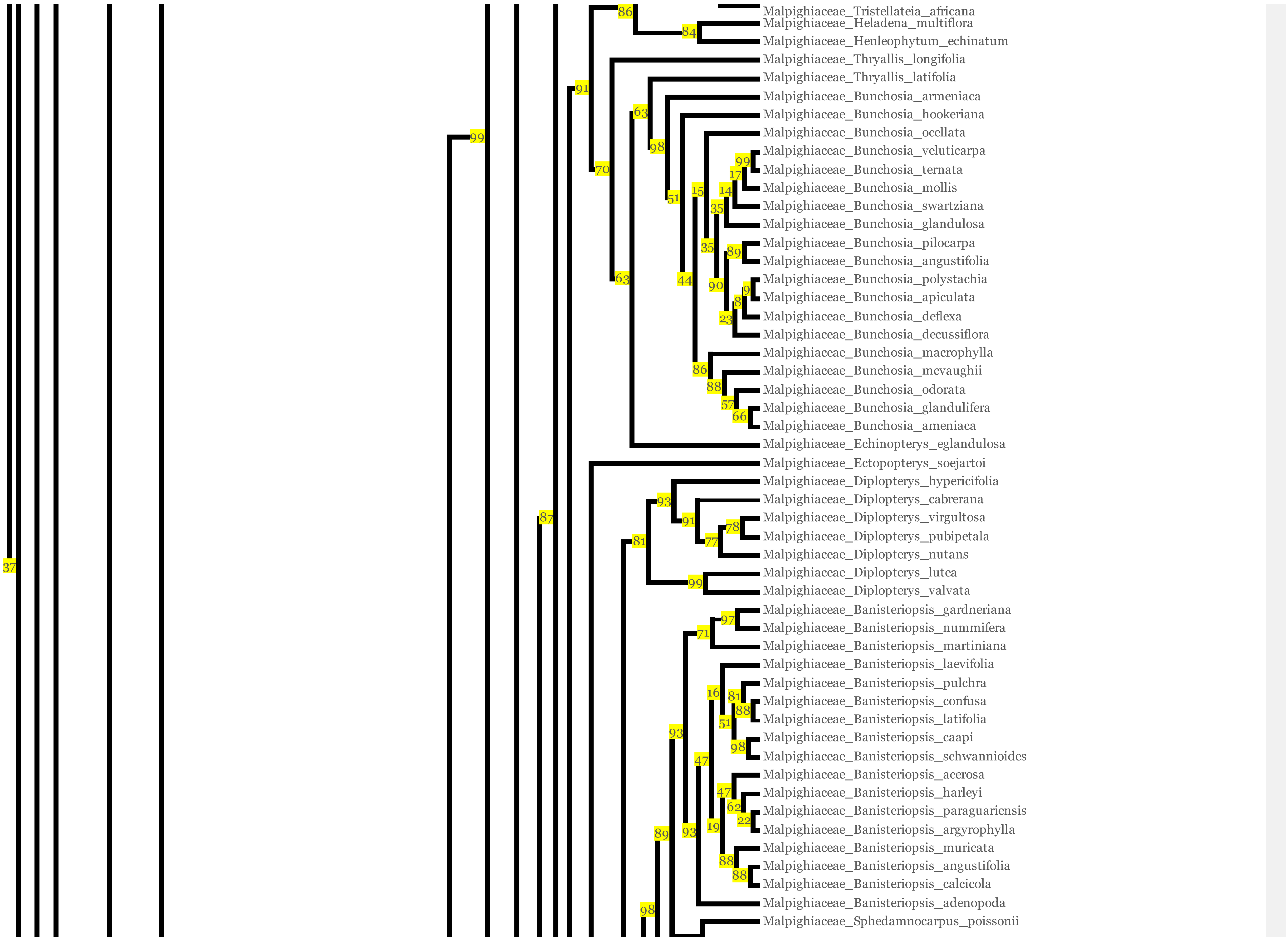
Malpighiaceae_Tristellateia_africana
Malpighiaceae_Heladena_multiflora
Malpighiaceae_Henleophytum_echinatum
Malpighiaceae_Thryallis_longifolia
Malpighiaceae_Thryallis_latifolia
Malpighiaceae_Bunchosia_armeniaca
Malpighiaceae_Bunchosia_hookeriana
Malpighiaceae_Bunchosia_ocellata
Malpighiaceae_Bunchosia_veluticarpa
Malpighiaceae_Bunchosia_ternata
Malpighiaceae_Bunchosia_mollis
Malpighiaceae_Bunchosia_swartziana
Malpighiaceae_Bunchosia_glandulosa
Malpighiaceae_Bunchosia_pilocarpa
Malpighiaceae_Bunchosia_angustifolia
Malpighiaceae_Bunchosia_polystachia
Malpighiaceae_Bunchosia_apiculata
Malpighiaceae_Bunchosia_deflexa
Malpighiaceae_Bunchosia_decussiflora
Malpighiaceae_Bunchosia_macrophylla
Malpighiaceae_Bunchosia_mcvaughii
Malpighiaceae_Bunchosia_odorata
Malpighiaceae_Bunchosia_glandulifera
Malpighiaceae_Bunchosia_ameniaca
Malpighiaceae_Echinopterys_eglandulosa
Malpighiaceae_Ectopopterys_soejartoi
Malpighiaceae_Diplopterys_hypericifolia
Malpighiaceae_Diplopterys_cabrerana
Malpighiaceae_Diplopterys_virgultosa
Malpighiaceae_Diplopterys_pubipetala
Malpighiaceae_Diplopterys_nutans
Malpighiaceae_Diplopterys_lutea
Malpighiaceae_Diplopterys_valvata
Malpighiaceae_Banisteriopsis_gardneriana
Malpighiaceae_Banisteriopsis_nummifera
Malpighiaceae_Banisteriopsis_martiniana
Malpighiaceae_Banisteriopsis_laevifolia
Malpighiaceae_Banisteriopsis_pulchra
Malpighiaceae_Banisteriopsis_confusa
Malpighiaceae_Banisteriopsis_latifolia
Malpighiaceae_Banisteriopsis_caapi
Malpighiaceae_Banisteriopsis_schwannioides
Malpighiaceae_Banisteriopsis_acerosa
Malpighiaceae_Banisteriopsis_harleyi
Malpighiaceae_Banisteriopsis_paraguariensis
Malpighiaceae_Banisteriopsis_argyrophylla
Malpighiaceae_Banisteriopsis_muricata
Malpighiaceae_Banisteriopsis_angustifolia
Malpighiaceae_Banisteriopsis_calcicola
Malpighiaceae_Banisteriopsis_adenopoda
Malpighiaceae_Sphedamnocarpus_poissonii

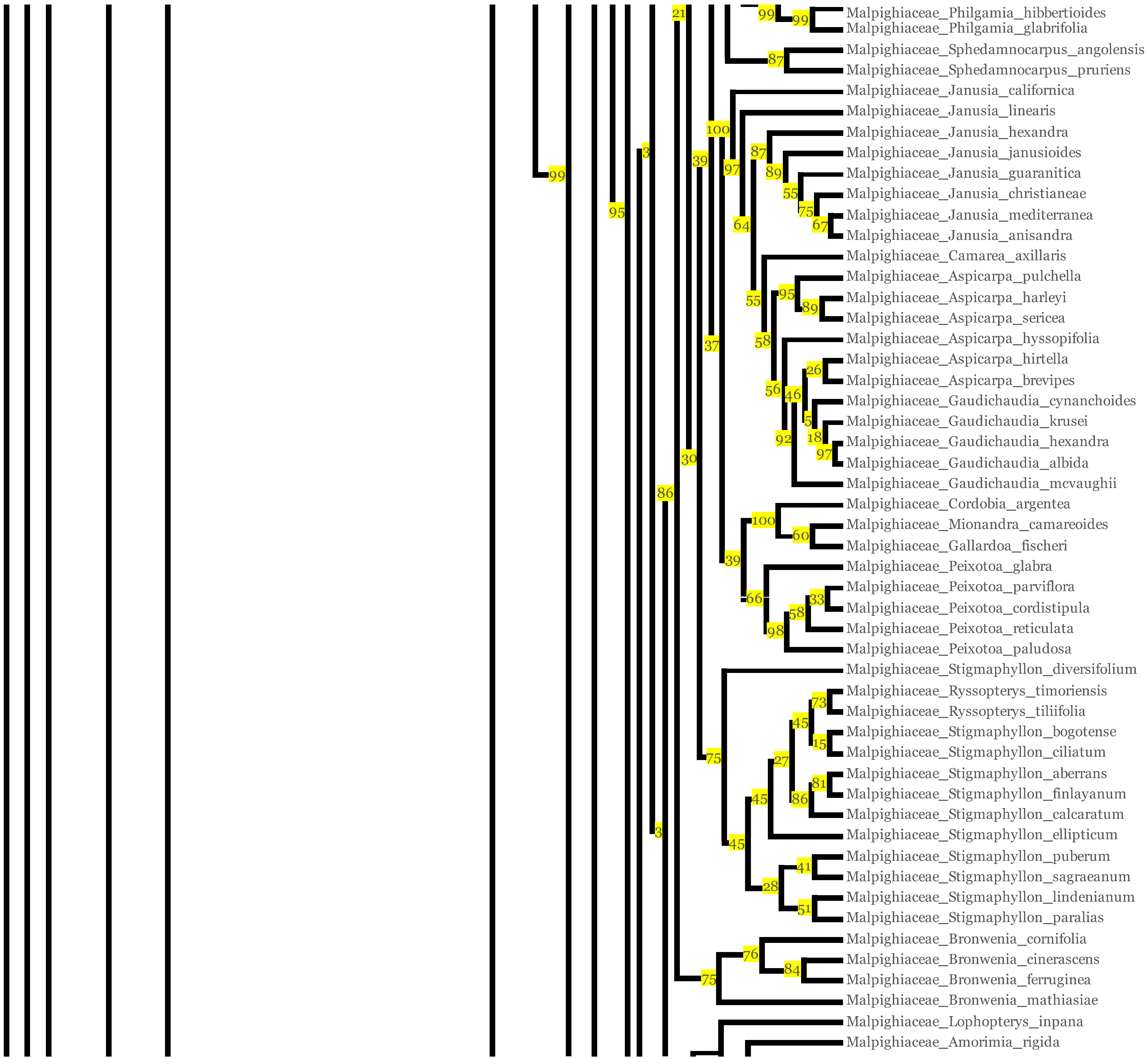

Malpighiaceae_Philgamia_hibbertioides
Malpighiaceae_Philgamia_glabrifolia
Malpighiaceae_Sphedamnocarpus_angolensis
Malpighiaceae_Sphedamnocarpus_pruriens
Malpighiaceae_Janusia_californica
Malpighiaceae_Janusia_linearis
Malpighiaceae_Janusia_hexandra
Malpighiaceae_Janusia_janusioides
Malpighiaceae_Janusia_guaranitica
Malpighiaceae_Janusia_christianeae
Malpighiaceae_Janusia_mediterranea
Malpighiaceae_Janusia_anisandra
Malpighiaceae_Camarea_axillaris
Malpighiaceae_Aspicarpa_pulchella
Malpighiaceae_Aspicarpa_harleyi
Malpighiaceae_Aspicarpa_sericea
Malpighiaceae_Aspicarpa_hyssopifolia
Malpighiaceae_Aspicarpa_hirtella
Malpighiaceae_Aspicarpa_brevipes
Malpighiaceae_Gaudichaudia_cynanchoides
Malpighiaceae_Gaudichaudia_krusei
Malpighiaceae_Gaudichaudia_hexandra
Malpighiaceae_Gaudichaudia_albida
Malpighiaceae_Gaudichaudia_mcvaughii
Malpighiaceae_Cordobia_argentea
Malpighiaceae_Mionandra_camareoides
Malpighiaceae_Gallardoa_fischeri
Malpighiaceae_Peixotoa_glabra
Malpighiaceae_Peixotoa_parviflora
Malpighiaceae_Peixotoa_cordistipula
Malpighiaceae_Peixotoa_reticulata
Malpighiaceae_Peixotoa_paludosa
Malpighiaceae_Stigmaphyllon_diversifolium
Malpighiaceae_Ryssopterys_timoriensis
Malpighiaceae_Ryssopterys_tiliifolia
Malpighiaceae_Stigmaphyllon_bogotense
Malpighiaceae_Stigmaphyllon_ciliatum
Malpighiaceae_Stigmaphyllon_aberrans
Malpighiaceae_Stigmaphyllon_finlayanum
Malpighiaceae_Stigmaphyllon_calcaratum
Malpighiaceae_Stigmaphyllon_ellipticum
Malpighiaceae_Stigmaphyllon_puberum
Malpighiaceae_Stigmaphyllon_sagraeanum
Malpighiaceae_Stigmaphyllon_lindenianum
Malpighiaceae_Stigmaphyllon_paralias
Malpighiaceae_Bronwenia_cornifolia
Malpighiaceae_Bronwenia_cinerascens
Malpighiaceae_Bronwenia_ferruginea
Malpighiaceae_Bronwenia_mathiasiae
Malpighiaceae_Lophopterys_inpana
Malpighiaceae_Amorimia_rigida

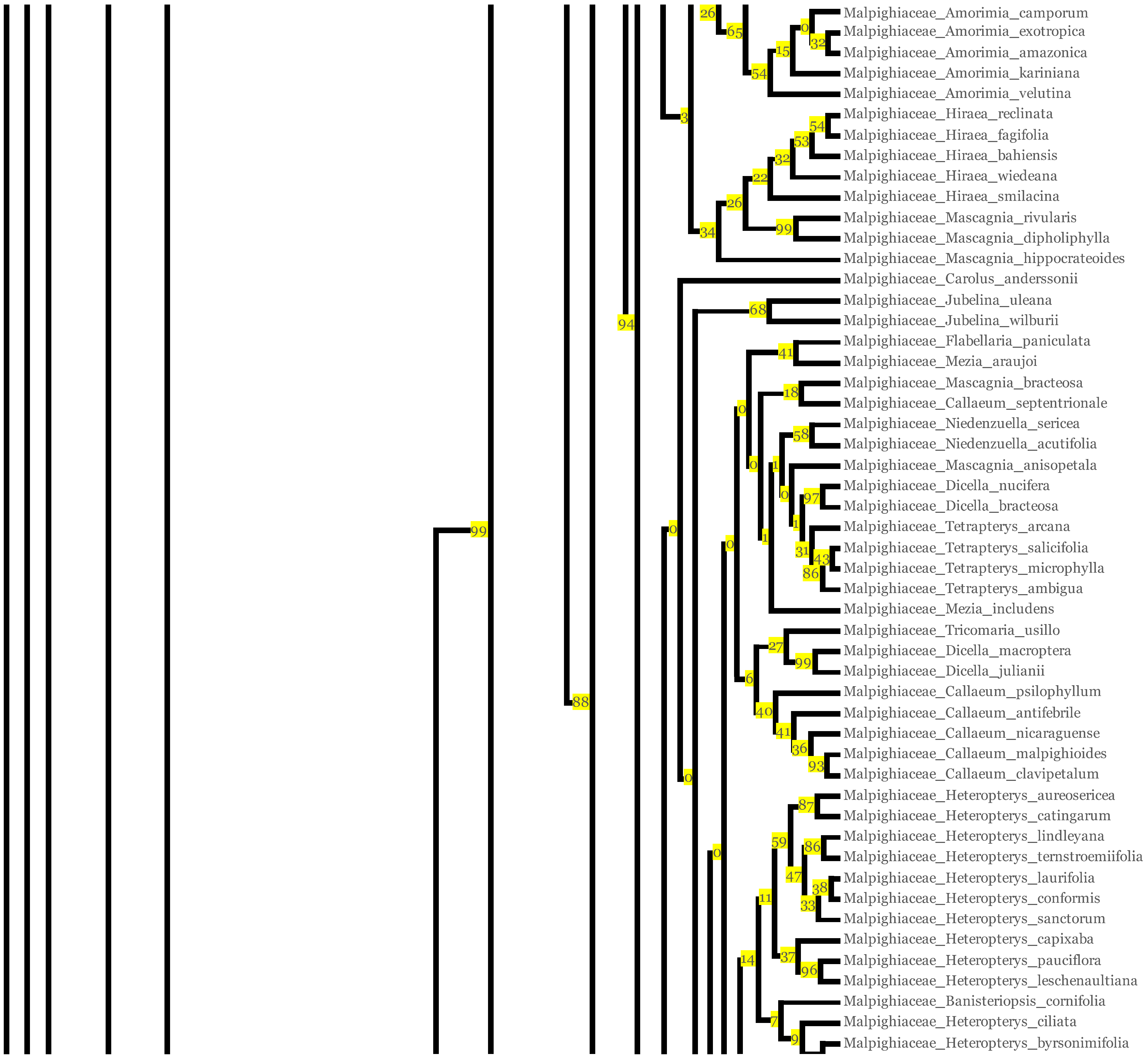
Malpighiaceae_Amorimia_camporum
Malpighiaceae_Amorimia_exotropica
Malpighiaceae_Amorimia_amazonica
Malpighiaceae_Amorimia_kariniana
Malpighiaceae_Amorimia_velutina
Malpighiaceae_Hiraea_reclinata
Malpighiaceae_Hiraea_fagifolia
Malpighiaceae_Hiraea_bahiensis
Malpighiaceae_Hiraea_wiedeana
Malpighiaceae_Hiraea_smilacina
Malpighiaceae_Mascagnia_rivularis
Malpighiaceae_Mascagnia_dipholiphylla
Malpighiaceae_Mascagnia_hippocrateoides
Malpighiaceae_Carolus_anderssonii
Malpighiaceae_Jubelina_uleana
Malpighiaceae_Jubelina_wilburii
Malpighiaceae_Flabellaria_paniculata
Malpighiaceae_Mezia_araujoi
Malpighiaceae_Mascagnia_bracteosa
Malpighiaceae_Callaeum_septentrionale
Malpighiaceae_Niedenzuella_sericea
Malpighiaceae_Niedenzuella_acutifolia
Malpighiaceae_Mascagnia_anisopetala
Malpighiaceae_Dicella_nucifera
Malpighiaceae_Dicella_bracteosa
Malpighiaceae_Tetrapterys_arcana
Malpighiaceae_Tetrapterys_salicifolia
Malpighiaceae_Tetrapterys_microphylla
Malpighiaceae_Tetrapterys_ambigua
Malpighiaceae_Mezia_includens
Malpighiaceae_Tricomaria_usillo
Malpighiaceae_Dicella_macroptera
Malpighiaceae_Dicella_julianii
Malpighiaceae_Callaeum_psilophyllum
Malpighiaceae_Callaeum_antifebrile
Malpighiaceae_Callaeum_nicaraguense
Malpighiaceae_Callaeum_malpighioides
Malpighiaceae_Callaeum_clavipetalum
Malpighiaceae_Heteropterys_aureosericea
Malpighiaceae_Heteropterys_catingarum
Malpighiaceae_Heteropterys_lindleyana
Malpighiaceae_Heteropterys_ternstroemiifolia
Malpighiaceae_Heteropterys_laurifolia
Malpighiaceae_Heteropterys_conformis
Malpighiaceae_Heteropterys_sanctorum
Malpighiaceae_Heteropterys_capixaba
Malpighiaceae_Heteropterys_pauciflora
Malpighiaceae_Heteropterys_leschenaultiana
Malpighiaceae_Banisteriopsis_cornifolia
Malpighiaceae_Heteropterys_ciliata
Malpighiaceae_Heteropterys_byrsonimifolia

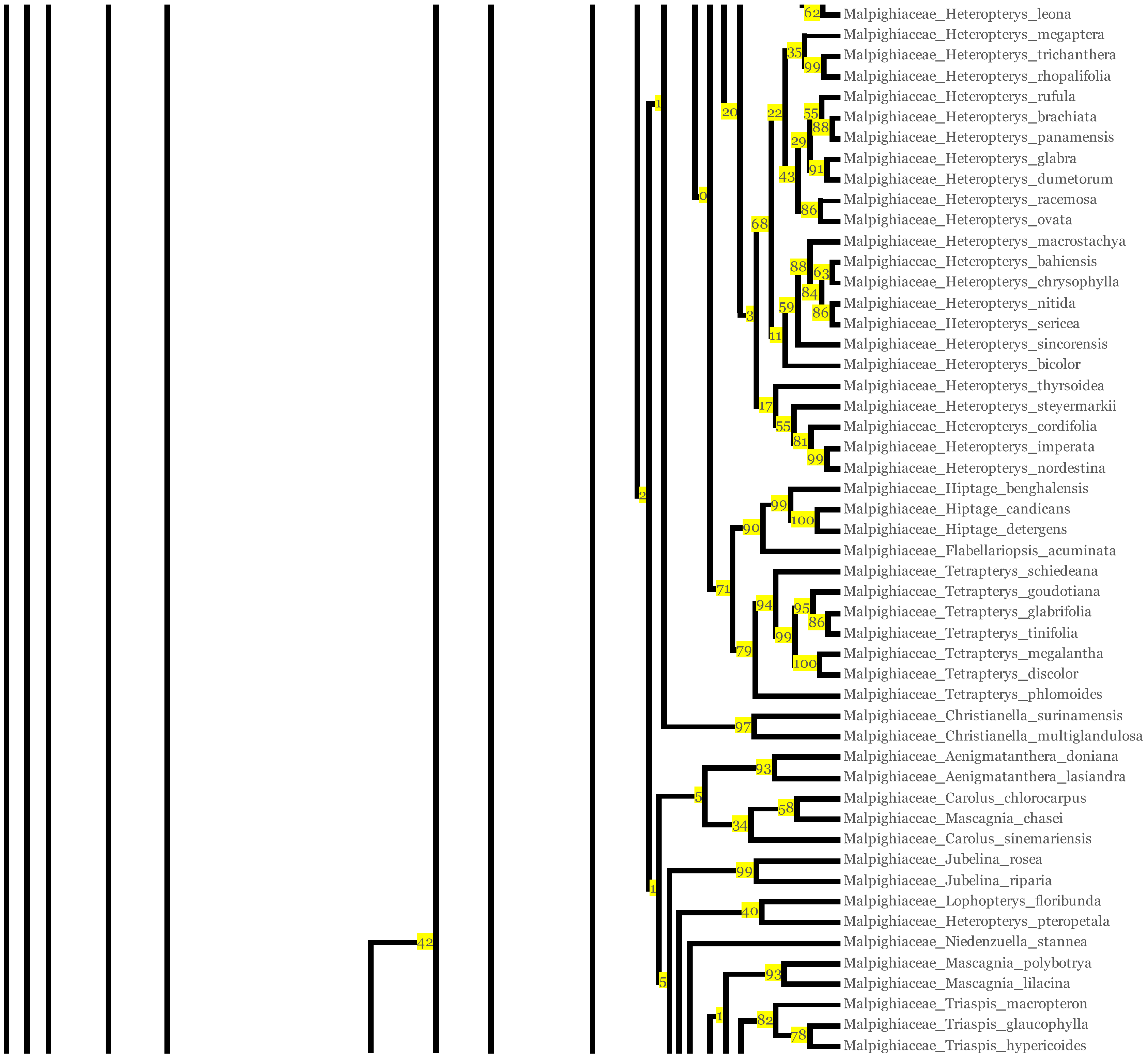
Malpighiaceae_Heteropterys_leona
Malpighiaceae_Heteropterys_megaptera
Malpighiaceae_Heteropterys_trichanthera
Malpighiaceae_Heteropterys_rhopalifolia
Malpighiaceae_Heteropterys_rufula
Malpighiaceae_Heteropterys_brachiata
Malpighiaceae_Heteropterys_panamensis
Malpighiaceae_Heteropterys_glabra
Malpighiaceae_Heteropterys_dumetorum
Malpighiaceae_Heteropterys_racemosa
Malpighiaceae_Heteropterys_ovata
Malpighiaceae_Heteropterys_macrostachya
Malpighiaceae_Heteropterys_bahiensis
Malpighiaceae_Heteropterys_chrysophylla
Malpighiaceae_Heteropterys_nitida
Malpighiaceae_Heteropterys_sericea
Malpighiaceae_Heteropterys_sincorensis
Malpighiaceae_Heteropterys_bicolor
Malpighiaceae_Heteropterys_thyrsoidea
Malpighiaceae_Heteropterys_steyermarkii
Malpighiaceae_Heteropterys_cordifolia
Malpighiaceae_Heteropterys_imperata
Malpighiaceae_Heteropterys_nordestina
Malpighiaceae_Hiptage_benghalensis
Malpighiaceae_Hiptage_candicans
Malpighiaceae_Hiptage_detergens
Malpighiaceae_Flabellariopsis_acuminata
Malpighiaceae_Tetrapterys_schiedeana
Malpighiaceae_Tetrapterys_goudotiana
Malpighiaceae_Tetrapterys_glabrifolia
Malpighiaceae_Tetrapterys_tinifolia
Malpighiaceae_Tetrapterys_megalantha
Malpighiaceae_Tetrapterys_discolor
Malpighiaceae_Tetrapterys_phlomoides
Malpighiaceae_Christianella_surinamensis
Malpighiaceae_Christianella_multiglandulosa
Malpighiaceae_Aenigmatanthera_doniana
Malpighiaceae_Aenigmatanthera_lasiandra
Malpighiaceae_Carolus_chlorocarpus
Malpighiaceae_Mascagnia_chasei
Malpighiaceae_Carolus_sinemariensis
Malpighiaceae_Jubelina_rosea
Malpighiaceae_Jubelina_riparia
Malpighiaceae_Lophopterys_floribunda
Malpighiaceae_Heteropterys_pteropetala
Malpighiaceae_Niedenzuella_stannea
Malpighiaceae_Mascagnia_polybotrya
Malpighiaceae_Mascagnia_lilacina
Malpighiaceae_Triaspis_macropteron
Malpighiaceae_Triaspis_glaucophylla
Malpighiaceae_Triaspis_hypericoides

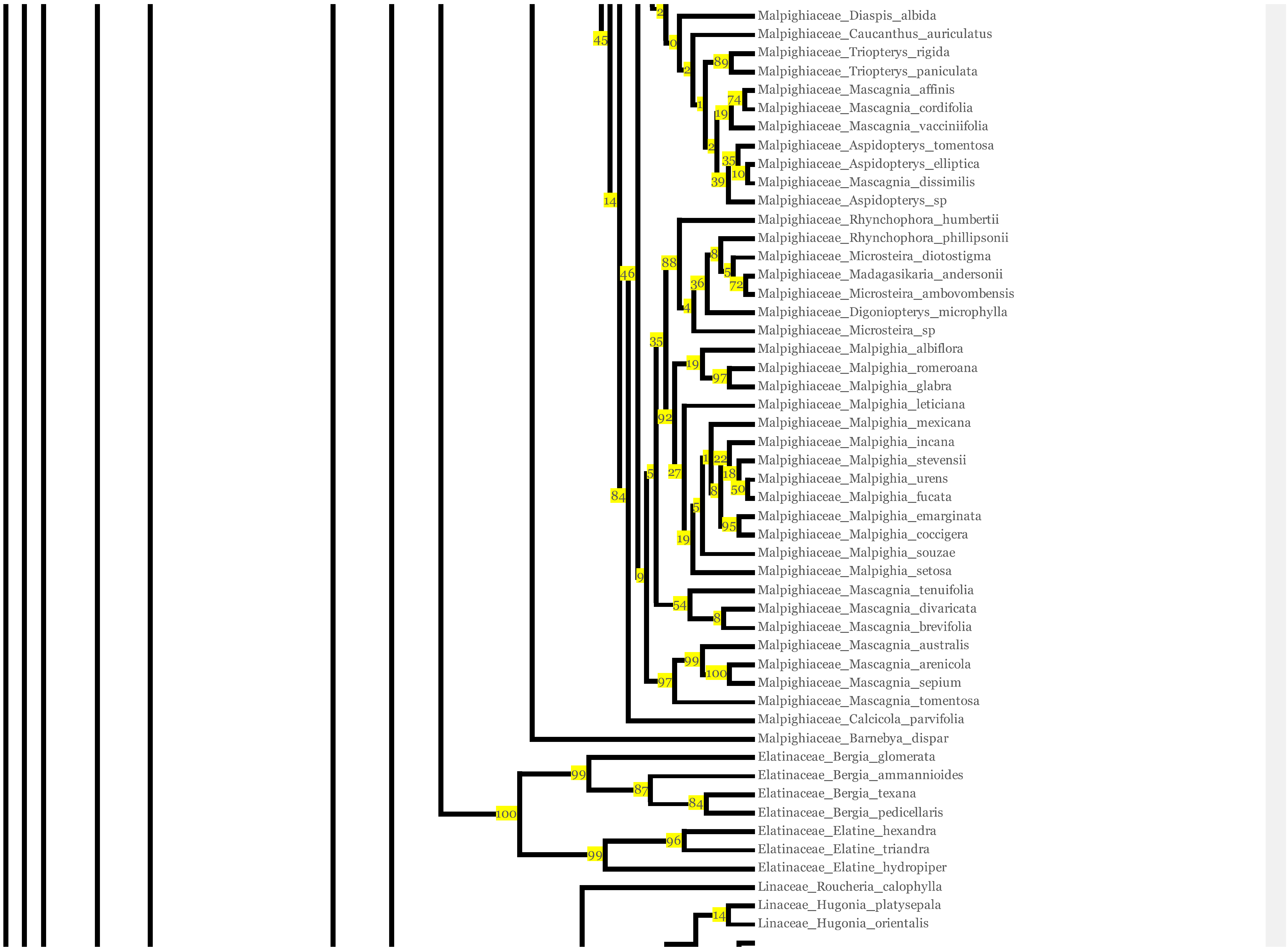
Malpighiaceae_Diaspis_albida
Malpighiaceae_Caucanthus_auriculatus
Malpighiaceae_Triopterys_rigida
Malpighiaceae_Triopterys_paniculata
Malpighiaceae_Mascagnia_affinis
Malpighiaceae_Mascagnia_cordifolia
Malpighiaceae_Mascagnia_vacciniifolia
Malpighiaceae_Aspidopterys_tomentosa
Malpighiaceae_Aspidopterys_elliptica
Malpighiaceae_Mascagnia_dissimilis
Malpighiaceae_Aspidopterys_sp
Malpighiaceae_Rhynchophora_humbertii
Malpighiaceae_Rhynchophora_phillipsonii
Malpighiaceae_Microsteira_diotostigma
Malpighiaceae_Madagasikaria_andersonii
Malpighiaceae_Microsteira_ambovombensis
Malpighiaceae_Digoniopterys_microphylla
Malpighiaceae_Microsteira_sp
Malpighiaceae_Malpighia_albiflora
Malpighiaceae_Malpighia_romeroana
Malpighiaceae_Malpighia_glabra
Malpighiaceae_Malpighia_leticiana
Malpighiaceae_Malpighia_mexicana
Malpighiaceae_Malpighia_incana
Malpighiaceae_Malpighia_stevensii
Malpighiaceae_Malpighia_urens
Malpighiaceae_Malpighia_fucata
Malpighiaceae_Malpighia_emarginata
Malpighiaceae_Malpighia_coccigera
Malpighiaceae_Malpighia_souzae
Malpighiaceae_Malpighia_setosa
Malpighiaceae_Mascagnia_tenuifolia
Malpighiaceae_Mascagnia_divaricata
Malpighiaceae_Mascagnia_brevifolia
Malpighiaceae_Mascagnia_australis
Malpighiaceae_Mascagnia_arenicola
Malpighiaceae_Mascagnia_sepium
Malpighiaceae_Mascagnia_tomentosa
Malpighiaceae_Calcicola_parvifolia
Malpighiaceae_Barnebya_dispar
Elatinaceae_Bergia_glomerata
Elatinaceae_Bergia_ammannioides
Elatinaceae_Bergia_texana
Elatinaceae_Bergia_pedicellaris
Elatinaceae_Elatine_hexandra
Elatinaceae_Elatine_triandra
Elatinaceae_Elatine_hydropiper
Linaceae_Roucheria_calophylla
Linaceae_Hugonia_platysepala
Linaceae_Hugonia_orientalis

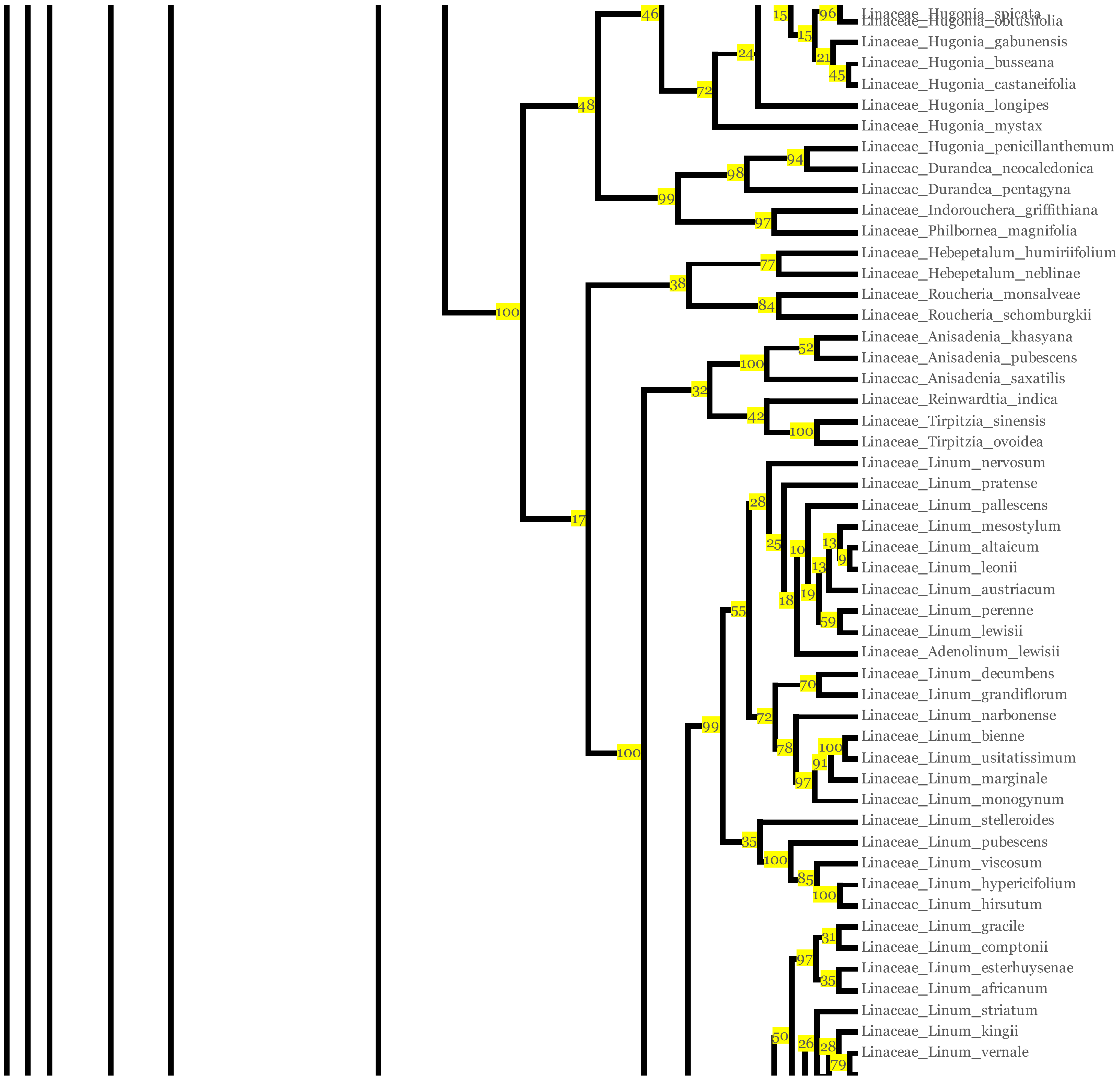

Linaceae_Hugonia_spicata
Linaceae_Hugonia_obtusifolia
Linaceae_Hugonia_gabunensis
Linaceae_Hugonia_busseana
Linaceae_Hugonia_castaneifolia
Linaceae_Hugonia_longipes
Linaceae_Hugonia_mystax
Linaceae_Hugonia_penicillanthemum
Linaceae_Durandea_neocaledonica
Linaceae_Durandea_pentagyna
Linaceae_Indorouchera_griffithiana
Linaceae_Philbornea_magnifolia
Linaceae_Hebepetalum_humiriifolium
Linaceae_Hebepetalum_neblinae
Linaceae_Roucheria_monsalveae
Linaceae_Roucheria_schomburgkii
Linaceae_Anisadenia_khasyana
Linaceae_Anisadenia_pubescens
Linaceae_Anisadenia_saxatilis
Linaceae_Reinwardtia_indica
Linaceae_Tirpitzia_sinensis
Linaceae_Tirpitzia_ovoidea
Linaceae_Linum_nervosum
Linaceae_Linum_pratense
Linaceae_Linum_pallescens
Linaceae_Linum_mesostylum
Linaceae_Linum_altaicum
Linaceae_Linum_leonii
Linaceae_Linum_austriacum
Linaceae_Linum_perenne
Linaceae_Linum_lewisii
Linaceae_Adenolinum_lewisii
Linaceae_Linum_decumbens
Linaceae_Linum_grandiflorum
Linaceae_Linum_narbonense
Linaceae_Linum_bienne
Linaceae_Linum_usitatissimum
Linaceae_Linum_marginale
Linaceae_Linum_monogynum
Linaceae_Linum_stelleroides
Linaceae_Linum_pubescens
Linaceae_Linum_viscosum
Linaceae_Linum_hypericifolium
Linaceae_Linum_hirsutum
Linaceae_Linum_gracile
Linaceae_Linum_comptonii
Linaceae_Linum_esterhuysenae
Linaceae_Linum_africanum
Linaceae_Linum_striatum
Linaceae_Linum_kingii
Linaceae_Linum_vernale

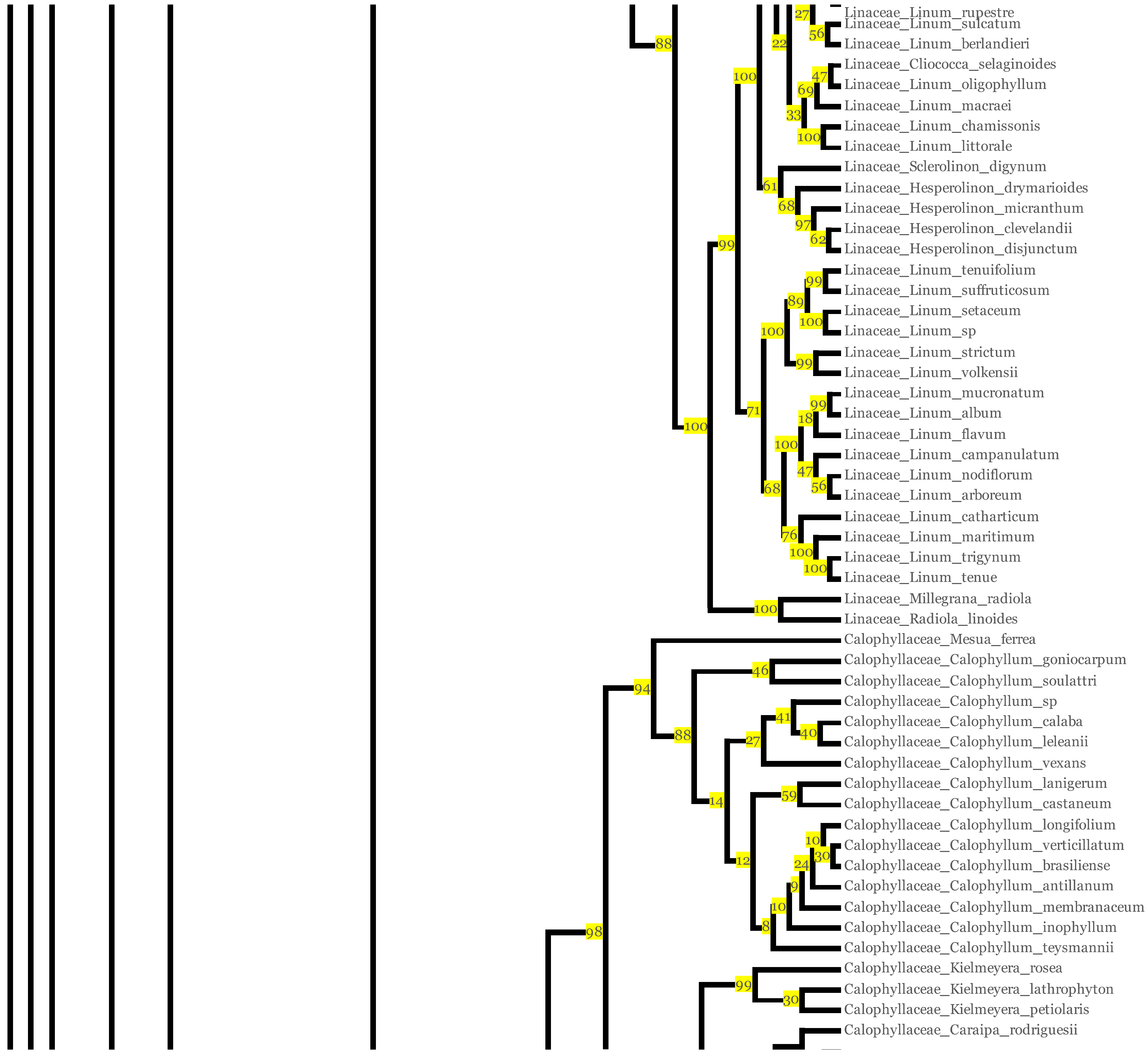

Linaceae_Linum_rupestre
Linaceae_Linum_sulcatum
Linaceae_Linum_berlandieri
Linaceae_Cliococca_selaginoides
Linaceae_Linum_oligophyllum
Linaceae_Linum_macraei
Linaceae_Linum_chamissonis
Linaceae_Linum_littorale
Linaceae_Sclerolinon_digynum
Linaceae_Hesperolinon_drymarioides
Linaceae_Hesperolinon_micranthum
Linaceae_Hesperolinon_clevelandii
Linaceae_Hesperolinon_disjunctum
Linaceae_Linum_tenuifolium
Linaceae_Linum_suffruticosum
Linaceae_Linum_setaceum
Linaceae_Linum_sp
Linaceae_Linum_strictum
Linaceae_Linum_volkensii
Linaceae_Linum_mucronatum
Linaceae_Linum_album
Linaceae_Linum_flavum
Linaceae_Linum_campanulatum
Linaceae_Linum_nodiflorum
Linaceae_Linum_arboreum
Linaceae_Linum_catharticum
Linaceae_Linum_maritimum
Linaceae_Linum_trigynum
Linaceae_Linum_tenue
Linaceae_Millegrana_radiola
Linaceae_Radiola_linoides
Calophyllaceae_Mesua_ferrea
Calophyllaceae_Calophyllum_goniocarpum
Calophyllaceae_Calophyllum_soulattri
Calophyllaceae_Calophyllum_sp
Calophyllaceae_Calophyllum_calaba
Calophyllaceae_Calophyllum_leleanii
Calophyllaceae_Calophyllum_vexans
Calophyllaceae_Calophyllum_lanigerum
Calophyllaceae_Calophyllum_castaneum
Calophyllaceae_Calophyllum_longifolium
Calophyllaceae_Calophyllum_verticillatum
Calophyllaceae_Calophyllum_brasiliense
Calophyllaceae_Calophyllum_antillanum
Calophyllaceae_Calophyllum_membranaceum
Calophyllaceae_Calophyllum_inophyllum
Calophyllaceae_Calophyllum_teysmannii
Calophyllaceae_Kielmeyera_rosea
Calophyllaceae_Kielmeyera_lathrophyton
Calophyllaceae_Kielmeyera_petiolaris
Calophyllaceae_Caraipa_rodriguesii

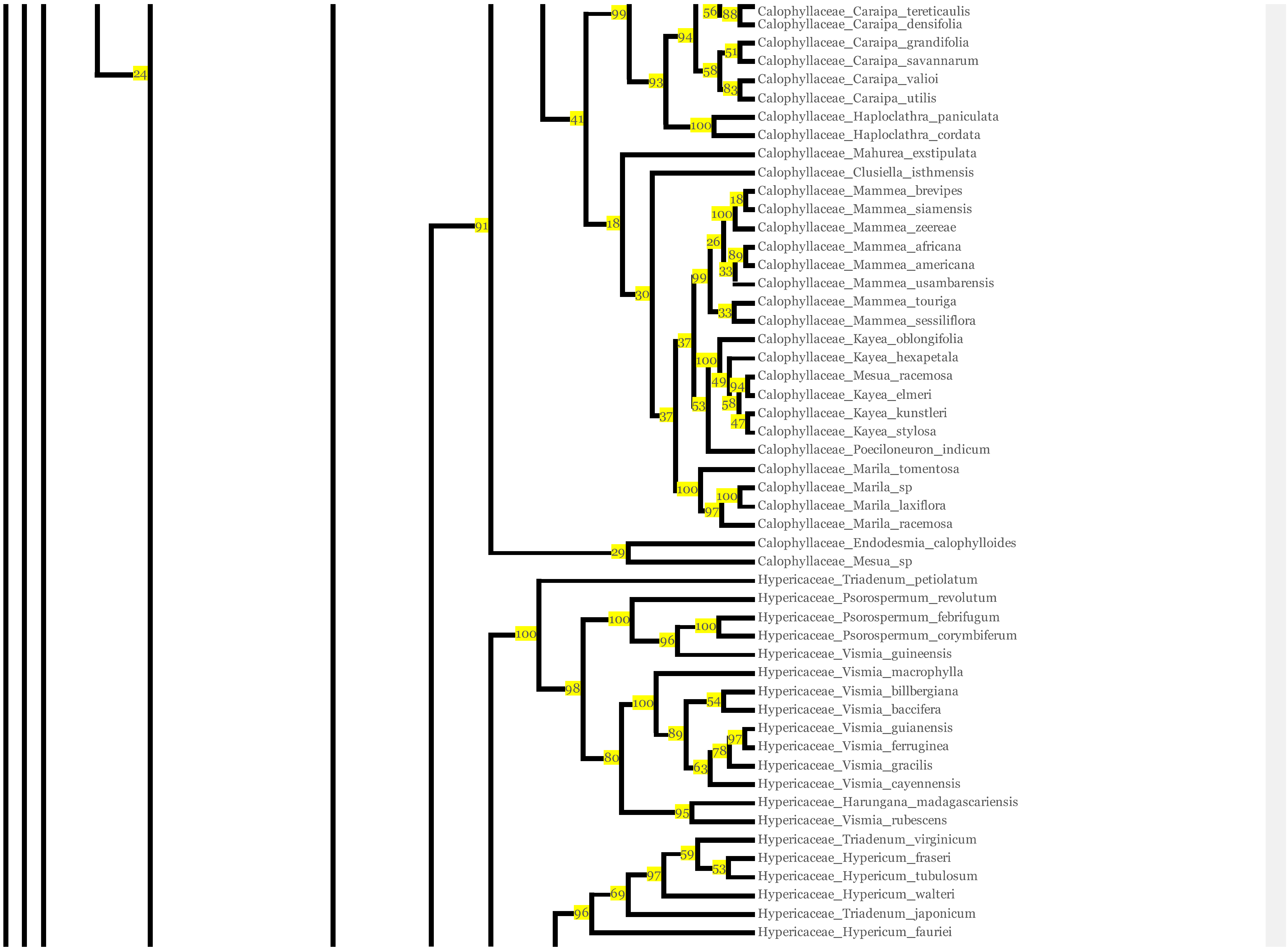
Calophyllaceae_Caraipa_tereticaulis
Calophyllaceae_Caraipa_densifolia
Calophyllaceae_Caraipa_grandifolia
Calophyllaceae_Caraipa_savannarum
Calophyllaceae_Caraipa_valioi
Calophyllaceae_Caraipa_utilis
Calophyllaceae_Haploclathra_paniculata
Calophyllaceae_Haploclathra_cordata
Calophyllaceae_Mahurea_exstipulata
Calophyllaceae_Clusiella_isthmensis
Calophyllaceae_Mammea_brevipes
Calophyllaceae_Mammea_siamensis
Calophyllaceae_Mammea_zeereae
Calophyllaceae_Mammea_africana
Calophyllaceae_Mammea_americana
Calophyllaceae_Mammea_usambarensis
Calophyllaceae_Mammea_touriga
Calophyllaceae_Mammea_sessiliflora
Calophyllaceae_Kayea_oblongifolia
Calophyllaceae_Kayea_hexapetala
Calophyllaceae_Mesua_racemosa
Calophyllaceae_Kayea_elmeri
Calophyllaceae_Kayea_kunstleri
Calophyllaceae_Kayea_stylosa
Calophyllaceae_Poeciloneuron_indicum
Calophyllaceae_Marila_tomentosa
Calophyllaceae_Marila_sp
Calophyllaceae_Marila_laxiflora
Calophyllaceae_Marila_racemosa
Calophyllaceae_Endodesmia_calophylloides
Calophyllaceae_Mesua_sp
Hypericaceae_Triadenum_petiolatum
Hypericaceae_Psorospermum_revolutum
Hypericaceae_Psorospermum_febrifugum
Hypericaceae_Psorospermum_corymbiferum
Hypericaceae_Vismia_guineensis
Hypericaceae_Vismia_macrophylla
Hypericaceae_Vismia_billbergiana
Hypericaceae_Vismia_baccifera
Hypericaceae_Vismia_guianensis
Hypericaceae_Vismia_ferruginea
Hypericaceae_Vismia_gracilis
Hypericaceae_Vismia_cayennensis
Hypericaceae_Harungana_madagascariensis
Hypericaceae_Vismia_rubescens
Hypericaceae_Triadenum_virginicum
Hypericaceae_Hypericum_fraseri
Hypericaceae_Hypericum_tubulosum
Hypericaceae_Hypericum_walteri
Hypericaceae_Triadenum_japonicum
Hypericaceae_Hypericum_fauriei

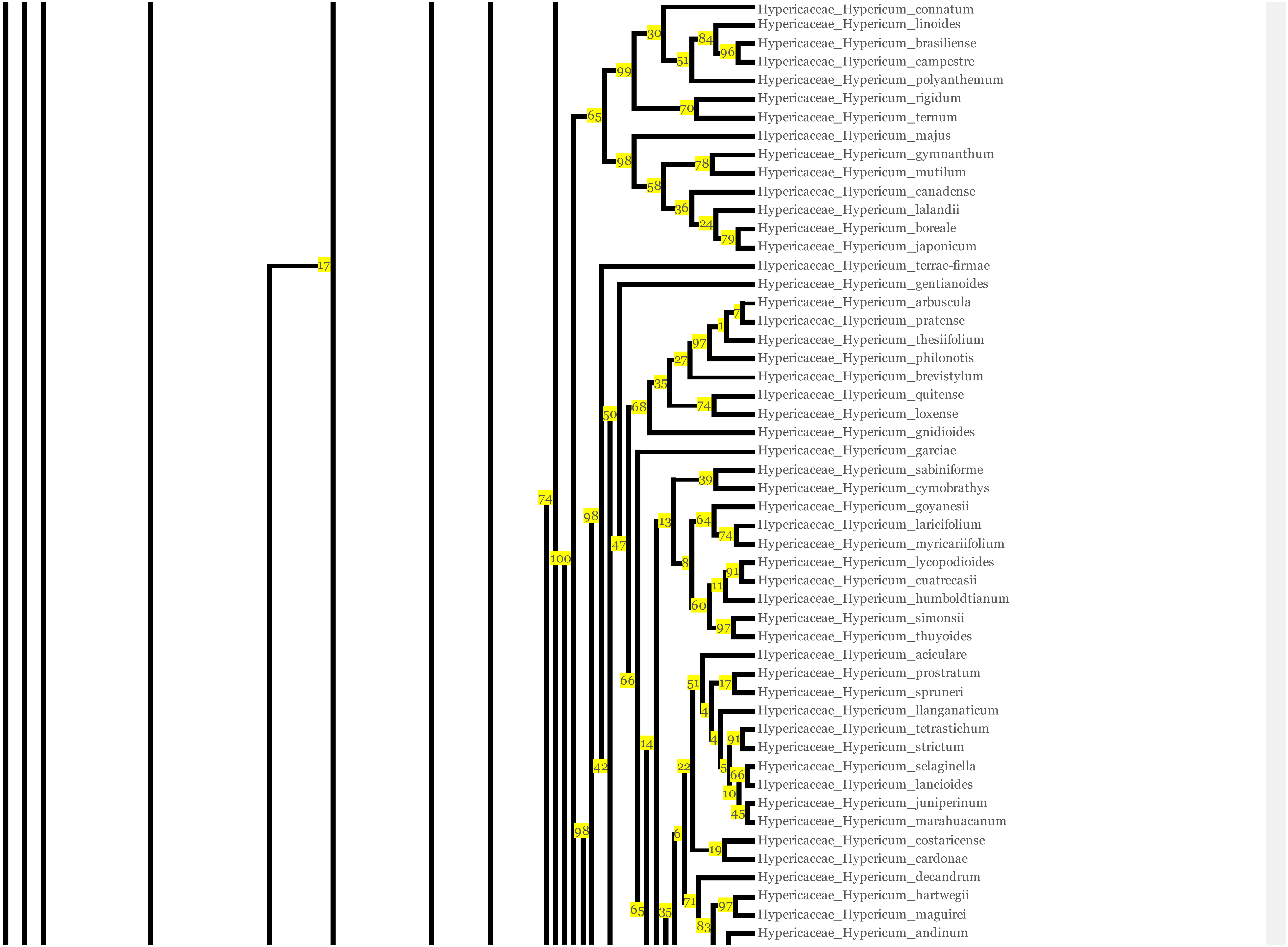
Hypericaceae_Hypericum_connatum
Hypericaceae_Hypericum_linoides
Hypericaceae_Hypericum_brasiliense
Hypericaceae_Hypericum_campestre
Hypericaceae_Hypericum_polyanthemum
Hypericaceae_Hypericum_rigidum
Hypericaceae_Hypericum_ternum
Hypericaceae_Hypericum_majus
Hypericaceae_Hypericum_gymnanthum
Hypericaceae_Hypericum_mutilum
Hypericaceae_Hypericum_canadense
Hypericaceae_Hypericum_lalandii
Hypericaceae_Hypericum_boreale
Hypericaceae_Hypericum_japonicum
Hypericaceae_Hypericum_terrae-firmae
Hypericaceae_Hypericum_gentianoides
Hypericaceae_Hypericum_arbuscula
Hypericaceae_Hypericum_pratense
Hypericaceae_Hypericum_thesiifolium
Hypericaceae_Hypericum_philonotis
Hypericaceae_Hypericum_brevistylum
Hypericaceae_Hypericum_quitense
Hypericaceae_Hypericum_loxense
Hypericaceae_Hypericum_gnidioides
Hypericaceae_Hypericum_garciae
Hypericaceae_Hypericum_sabiniforme
Hypericaceae_Hypericum_cymobrathys
Hypericaceae_Hypericum_goyanesii
Hypericaceae_Hypericum_laricifolium
Hypericaceae_Hypericum_myricariifolium
Hypericaceae_Hypericum_lycopodioides
Hypericaceae_Hypericum_cuatrecasii
Hypericaceae_Hypericum_humboldtianum
Hypericaceae_Hypericum_simonsii
Hypericaceae_Hypericum_thuyoides
Hypericaceae_Hypericum_aciculare
Hypericaceae_Hypericum_prostratum
Hypericaceae_Hypericum_spruneri
Hypericaceae_Hypericum_llanganaticum
Hypericaceae_Hypericum_tetrastichum
Hypericaceae_Hypericum_strictum
Hypericaceae_Hypericum_selaginella
Hypericaceae_Hypericum_lancioides
Hypericaceae_Hypericum_juniperinum
Hypericaceae_Hypericum_marahuacanum
Hypericaceae_Hypericum_costaricense
Hypericaceae_Hypericum_cardonae
Hypericaceae_Hypericum_decandrum
Hypericaceae_Hypericum_hartwegii
Hypericaceae_Hypericum_maguirei
Hypericaceae_Hypericum_andinum

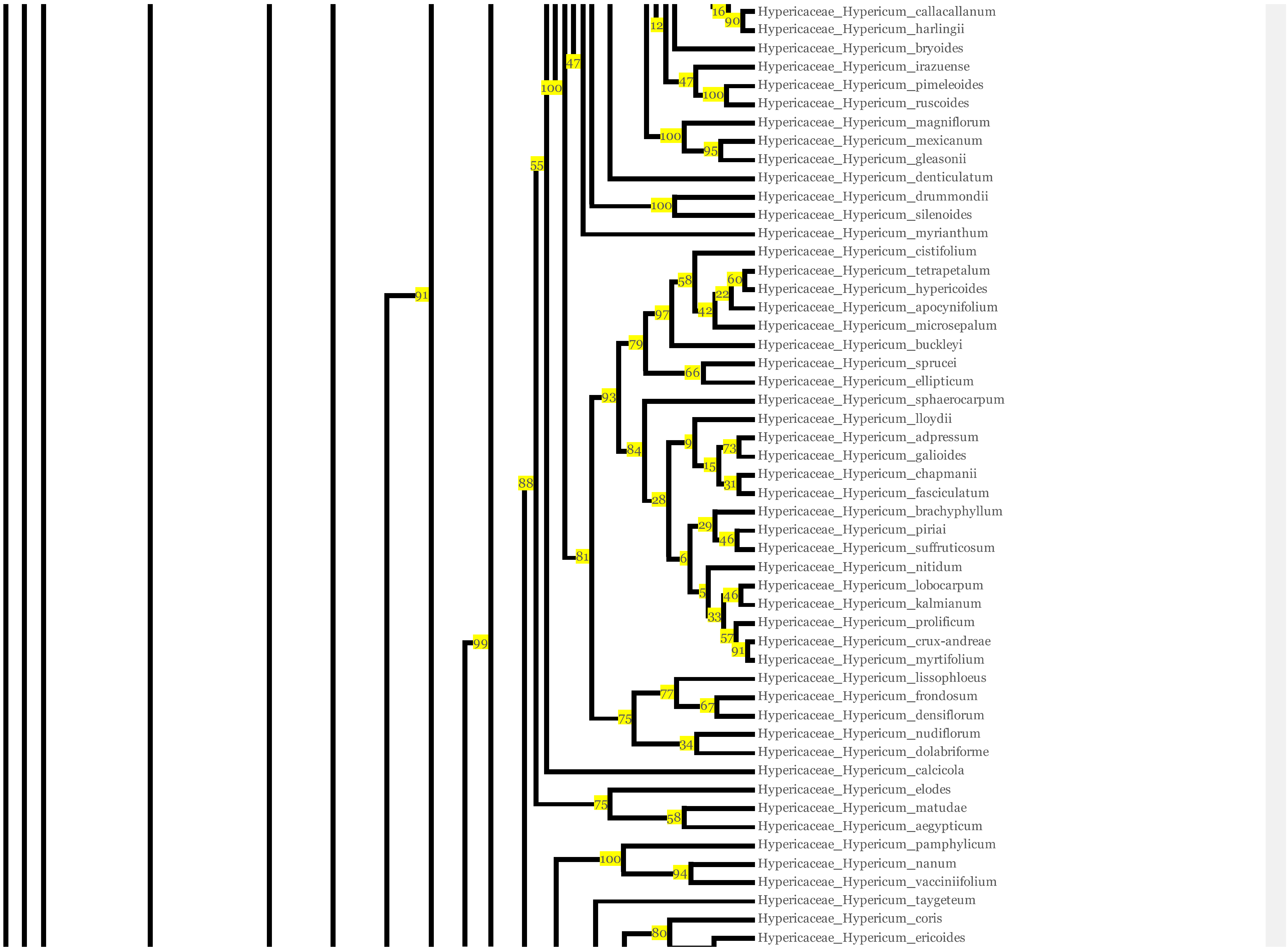
Hypericaceae_Hypericum_callacallanum
Hypericaceae_Hypericum_harlingii
Hypericaceae_Hypericum_bryoides
Hypericaceae_Hypericum_irazuense
Hypericaceae_Hypericum_pimeleoides
Hypericaceae_Hypericum_ruscoides
Hypericaceae_Hypericum_magniflorum
Hypericaceae_Hypericum_mexicanum
Hypericaceae_Hypericum_gleasonii
Hypericaceae_Hypericum_denticulatum
Hypericaceae_Hypericum_drummondii
Hypericaceae_Hypericum_silenoides
Hypericaceae_Hypericum_myrianthum
Hypericaceae_Hypericum_cistifolium
Hypericaceae_Hypericum_tetrapetalum
Hypericaceae_Hypericum_hypericoides
Hypericaceae_Hypericum_apocynifolium
Hypericaceae_Hypericum_microsepalum
Hypericaceae_Hypericum_buckleyi
Hypericaceae_Hypericum_sprucei
Hypericaceae_Hypericum_ellipticum
Hypericaceae_Hypericum_sphaerocarpum
Hypericaceae_Hypericum_lloydii
Hypericaceae_Hypericum_adpressum
Hypericaceae_Hypericum_galioides
Hypericaceae_Hypericum_chapmanii
Hypericaceae_Hypericum_fasciculatum
Hypericaceae_Hypericum_brachyphyllum
Hypericaceae_Hypericum_piriai
Hypericaceae_Hypericum_suffruticosum
Hypericaceae_Hypericum_nitidum
Hypericaceae_Hypericum_lobocarpum
Hypericaceae_Hypericum_kalmianum
Hypericaceae_Hypericum_prolificum
Hypericaceae_Hypericum_crux-andreae
Hypericaceae_Hypericum_myrtifolium
Hypericaceae_Hypericum_lissophloeus
Hypericaceae_Hypericum_frondosum
Hypericaceae_Hypericum_densiflorum
Hypericaceae_Hypericum_nudiflorum
Hypericaceae_Hypericum_dolabriforme
Hypericaceae_Hypericum_calcicola
Hypericaceae_Hypericum_elodes
Hypericaceae_Hypericum_matudae
Hypericaceae_Hypericum_aegypticum
Hypericaceae_Hypericum_pamphylicum
Hypericaceae_Hypericum_nanum
Hypericaceae_Hypericum_vacciniifolium
Hypericaceae_Hypericum_taygeteum
Hypericaceae_Hypericum_coris
Hypericaceae_Hypericum_ericoides

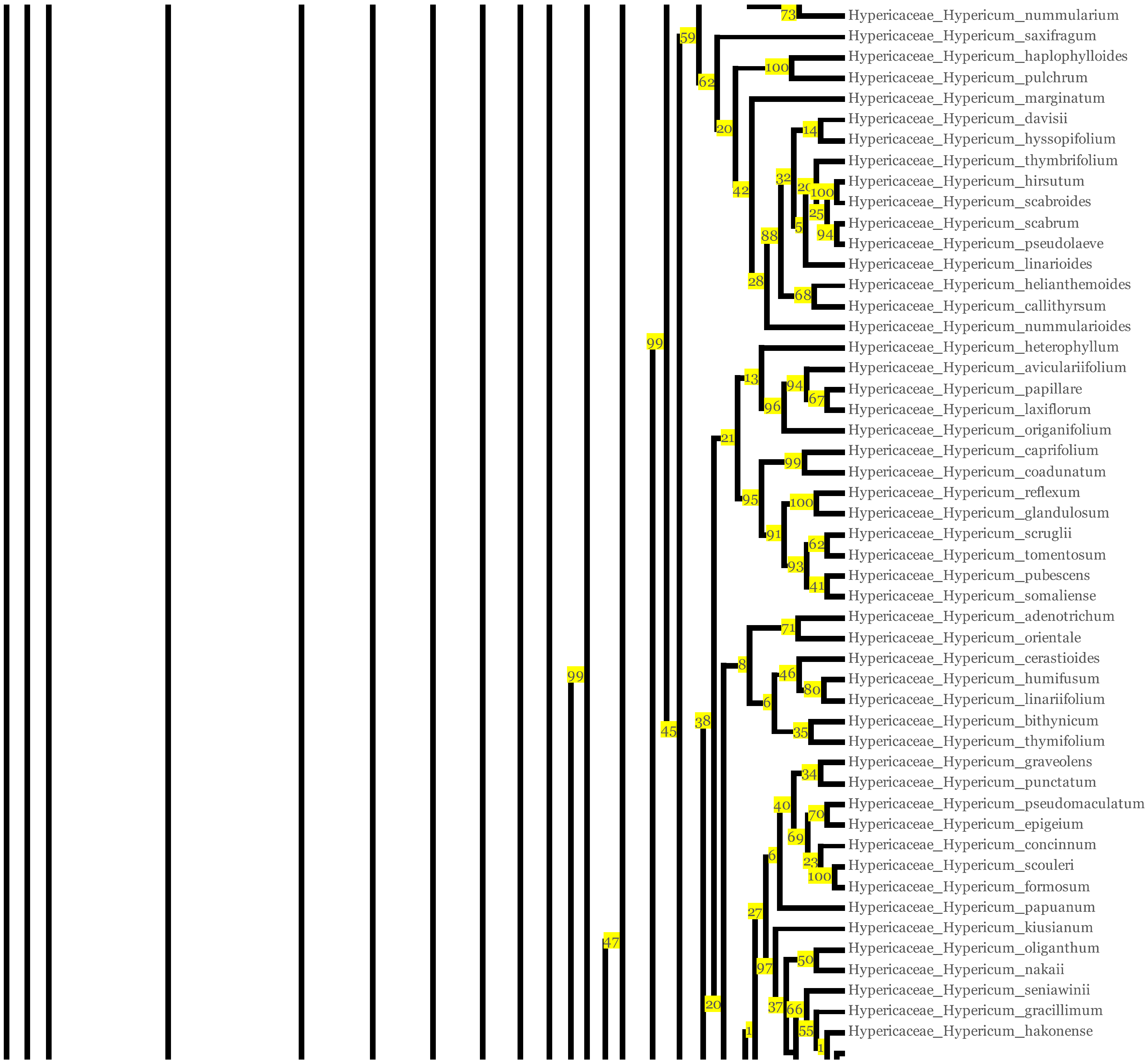

Hypericaceae_Hypericum_nummularium
Hypericaceae_Hypericum_saxifragum
Hypericaceae_Hypericum_haplophylloides
Hypericaceae_Hypericum_pulchrum
Hypericaceae_Hypericum_marginatum
Hypericaceae_Hypericum_davisii
Hypericaceae_Hypericum_hyssopifolium
Hypericaceae_Hypericum_thymbrifolium
Hypericaceae_Hypericum_hirsutum
Hypericaceae_Hypericum_scabroides
Hypericaceae_Hypericum_scabrum
Hypericaceae_Hypericum_pseudolaeve
Hypericaceae_Hypericum_linarioides
Hypericaceae_Hypericum_helianthemoides
Hypericaceae_Hypericum_callithyrsum
Hypericaceae_Hypericum_nummularioides
Hypericaceae_Hypericum_heterophyllum
Hypericaceae_Hypericum_aviculariifolium
Hypericaceae_Hypericum_papillare
Hypericaceae_Hypericum_laxiflorum
Hypericaceae_Hypericum_origanifolium
Hypericaceae_Hypericum_caprifolium
Hypericaceae_Hypericum_coadunatum
Hypericaceae_Hypericum_reflexum
Hypericaceae_Hypericum_glandulosum
Hypericaceae_Hypericum_scruglii
Hypericaceae_Hypericum_tomentosum
Hypericaceae_Hypericum_pubescens
Hypericaceae_Hypericum_somaliense
Hypericaceae_Hypericum_adenotrichum
Hypericaceae_Hypericum_orientale
Hypericaceae_Hypericum_cerastioides
Hypericaceae_Hypericum_humifusum
Hypericaceae_Hypericum_linariifolium
Hypericaceae_Hypericum_bithynicum
Hypericaceae_Hypericum_thymifolium
Hypericaceae_Hypericum_graveolens
Hypericaceae_Hypericum_punctatum
Hypericaceae_Hypericum_pseudomaculatum
Hypericaceae_Hypericum_epigeium
Hypericaceae_Hypericum_concinnum
Hypericaceae_Hypericum_scouleri
Hypericaceae_Hypericum_formosum
Hypericaceae_Hypericum_papuanum
Hypericaceae_Hypericum_kiusianum
Hypericaceae_Hypericum_oliganthum
Hypericaceae_Hypericum_nakaii
Hypericaceae_Hypericum_seniawinii
Hypericaceae_Hypericum_gracillimum
Hypericaceae_Hypericum_hakonense
73
59
100
62
20
14
42
32
100
25
94
88
28
68
99
13
94
67
96
21
99
100
95
91
62
93
41
71
8
46
80
99
45
38
35
34
40
70
69
23
100
27
47
97
50
37
66
20
55

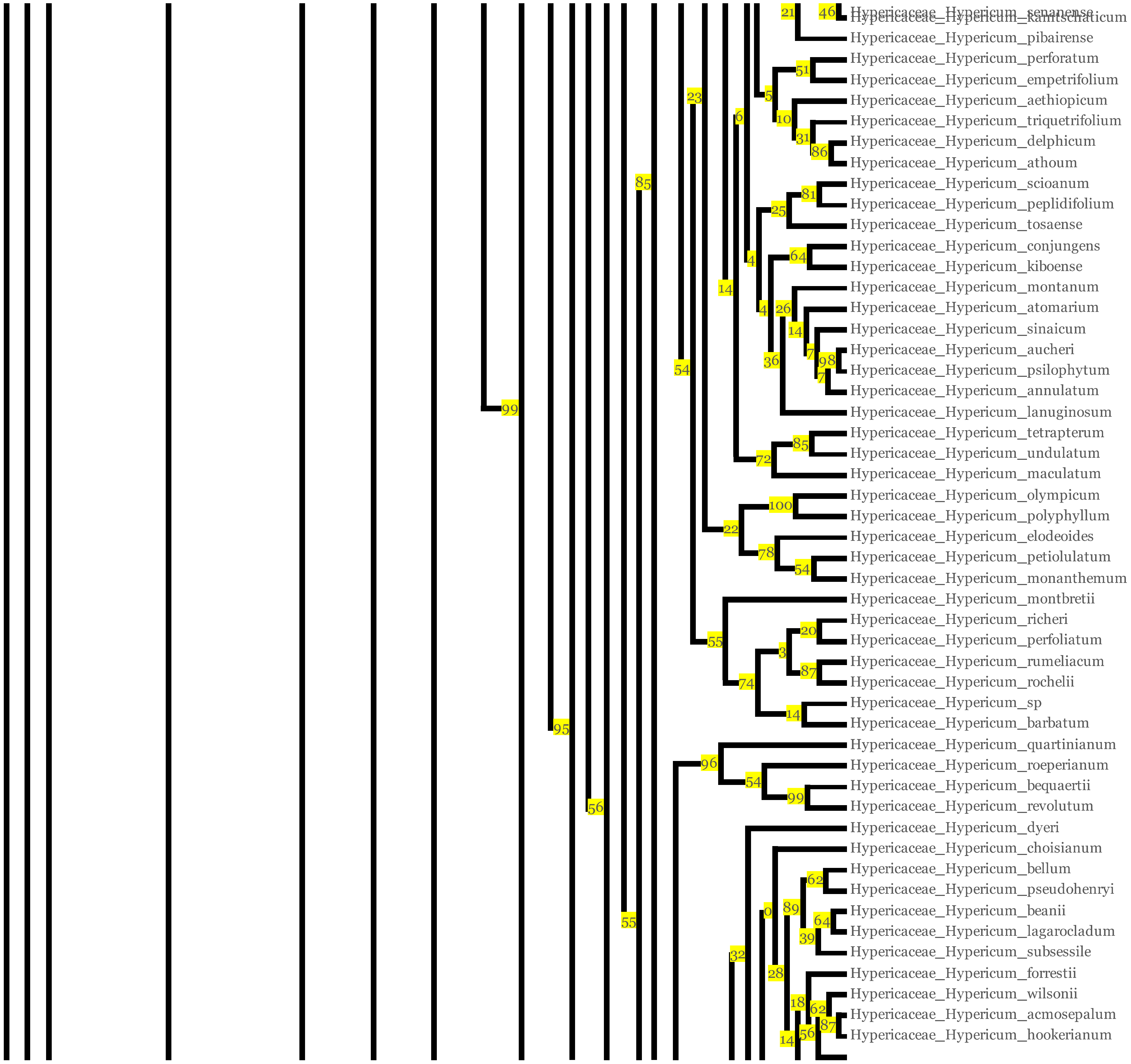
Hypericaceae_Hypericum_senanense
Hypericaceae_Hypericum_kamtschaticum
Hypericaceae_Hypericum_pibairense
Hypericaceae_Hypericum_perforatum
Hypericaceae_Hypericum_empetrifolium
Hypericaceae_Hypericum_aethiopicum
Hypericaceae_Hypericum_triquetrifolium
Hypericaceae_Hypericum_delphicum
Hypericaceae_Hypericum_athoum
Hypericaceae_Hypericum_scioanum
Hypericaceae_Hypericum_peplidifolium
Hypericaceae_Hypericum_tosaense
Hypericaceae_Hypericum_conjungens
Hypericaceae_Hypericum_kiboense
Hypericaceae_Hypericum_montanum
Hypericaceae_Hypericum_atomarium
Hypericaceae_Hypericum_sinaicum
Hypericaceae_Hypericum_aucheri
Hypericaceae_Hypericum_psilophytum
Hypericaceae_Hypericum_annulatum
Hypericaceae_Hypericum_lanuginosum
Hypericaceae_Hypericum_tetrapterum
Hypericaceae_Hypericum_undulatum
Hypericaceae_Hypericum_maculatum
Hypericaceae_Hypericum_olympicum
Hypericaceae_Hypericum_polyphyllum
Hypericaceae_Hypericum_elodeoides
Hypericaceae_Hypericum_petiolulatum
Hypericaceae_Hypericum_monanthemum
Hypericaceae_Hypericum_montbretii
Hypericaceae_Hypericum_richeri
Hypericaceae_Hypericum_perfoliatum
Hypericaceae_Hypericum_rumeliacum
Hypericaceae_Hypericum_rochelii
Hypericaceae_Hypericum_sp
Hypericaceae_Hypericum_barbatum
Hypericaceae_Hypericum_quartinianum
Hypericaceae_Hypericum_roeperianum
Hypericaceae_Hypericum_bequaertii
Hypericaceae_Hypericum_revolutum
Hypericaceae_Hypericum_dyeri
Hypericaceae_Hypericum_choisianum
Hypericaceae_Hypericum_bellum
Hypericaceae_Hypericum_pseudohenryi
Hypericaceae_Hypericum_beanii
Hypericaceae_Hypericum_lagarocladum
Hypericaceae_Hypericum_subsessile
Hypericaceae_Hypericum_forrestii
Hypericaceae_Hypericum_wilsonii
Hypericaceae_Hypericum_acmosepalum
Hypericaceae_Hypericum_hookerianum

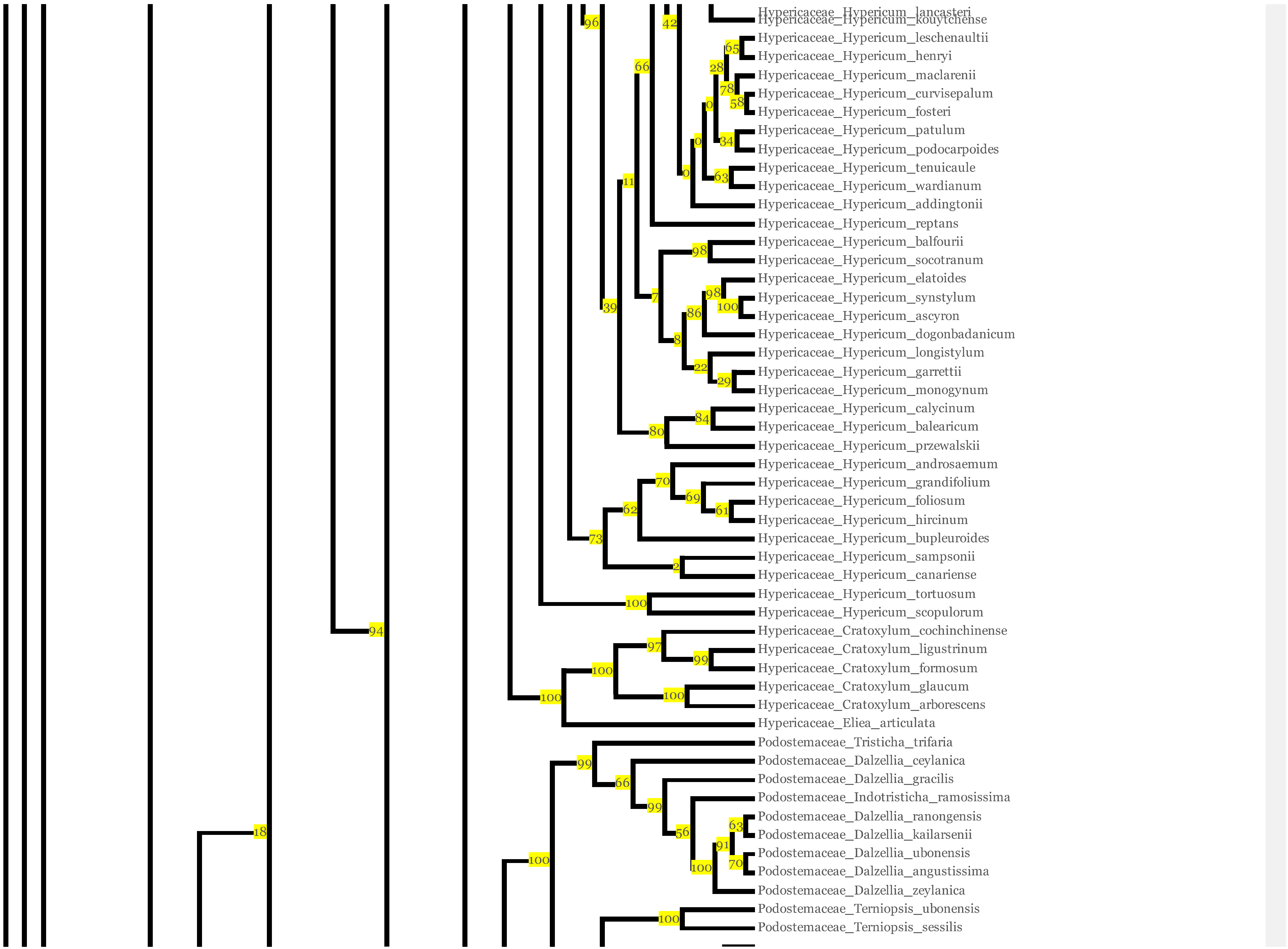
Hypericaceae_Hypericum_lancasteri
Hypericaceae_Hypericum_kouytchense
Hypericaceae_Hypericum_leschenaultii
Hypericaceae_Hypericum_henryi
Hypericaceae_Hypericum_maclarenii
Hypericaceae_Hypericum_curvisepalum
Hypericaceae_Hypericum_fosteri
Hypericaceae_Hypericum_patulum
Hypericaceae_Hypericum_podocarpoides
Hypericaceae_Hypericum_tenuicaule
Hypericaceae_Hypericum_wardianum
Hypericaceae_Hypericum_addingtonii
Hypericaceae_Hypericum_reptans
Hypericaceae_Hypericum_balfourii
Hypericaceae_Hypericum_socotranum
Hypericaceae_Hypericum_elatoides
Hypericaceae_Hypericum_synstylum
Hypericaceae_Hypericum_ascyron
Hypericaceae_Hypericum_dogonbadanicum
Hypericaceae_Hypericum_longistylum
Hypericaceae_Hypericum_garrettii
Hypericaceae_Hypericum_monogynum
Hypericaceae_Hypericum_calycinum
Hypericaceae_Hypericum_balearicum
Hypericaceae_Hypericum_przewalskii
Hypericaceae_Hypericum_androsaemum
Hypericaceae_Hypericum_grandifolium
Hypericaceae_Hypericum_foliosum
Hypericaceae_Hypericum_hircinum
Hypericaceae_Hypericum_bupleuroides
Hypericaceae_Hypericum_sampsonii
Hypericaceae_Hypericum_canariense
Hypericaceae_Hypericum_tortuosum
Hypericaceae_Hypericum_scopulorum
Hypericaceae_Cratoxylum_cochinchinense
Hypericaceae_Cratoxylum_ligustrinum
Hypericaceae_Cratoxylum_formosum
Hypericaceae_Cratoxylum_glaucum
Hypericaceae_Cratoxylum_arborescens
Hypericaceae_Eliea_articulata
Podostemaceae_Tristicha_trifaria
Podostemaceae_Dalzellia_ceylanica
Podostemaceae_Dalzellia_gracilis
Podostemaceae_Indotristicha_ramosissima
Podostemaceae_Dalzellia_ranongensis
Podostemaceae_Dalzellia_kailarsenii
Podostemaceae_Dalzellia_ubonensis
Podostemaceae_Dalzellia_angustissima
Podostemaceae_Dalzellia_zeylanica
Podostemaceae_Terniopsis_ubonensis
Podostemaceae_Terniopsis_sessilis

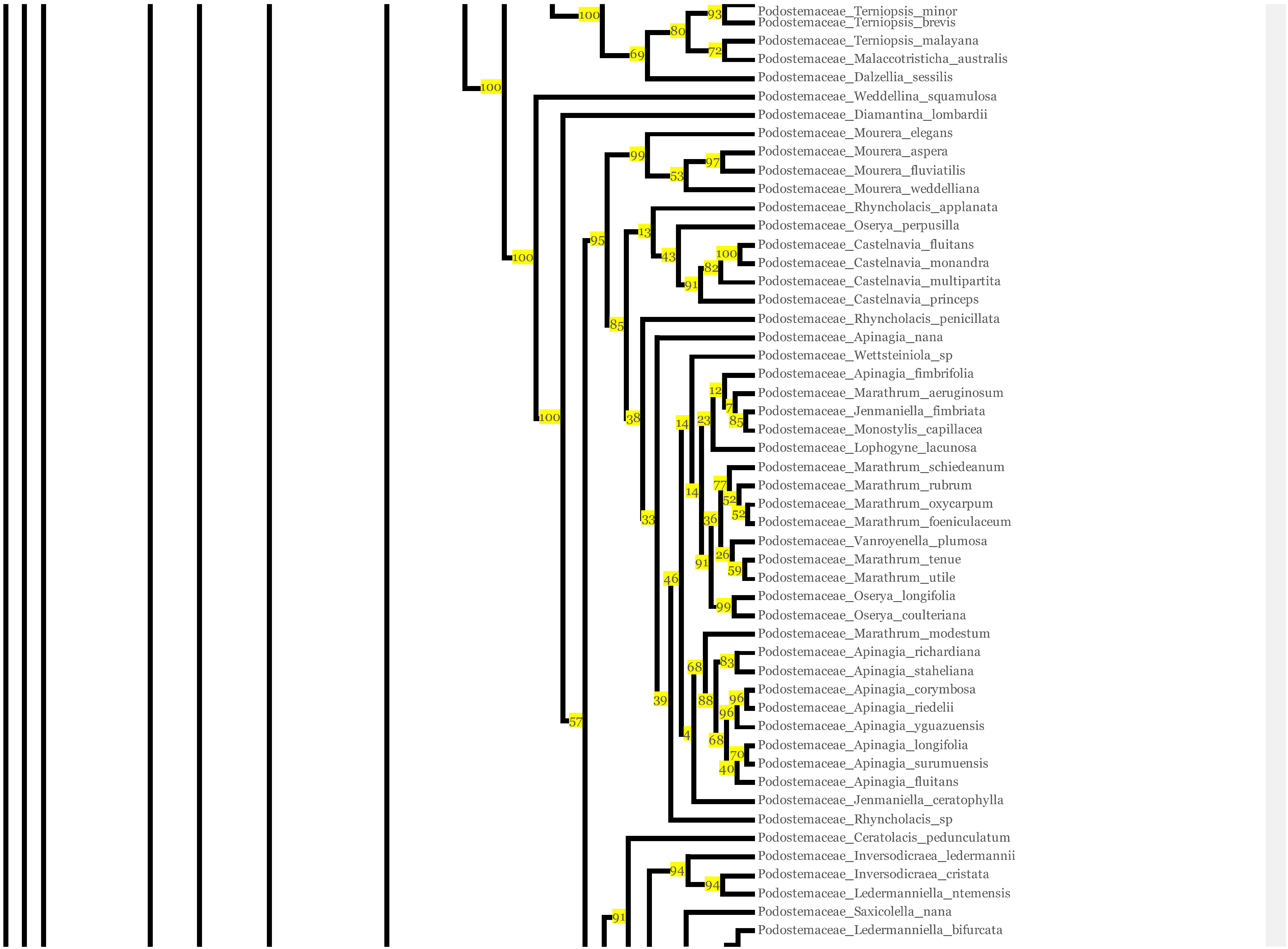
Podostemaceae_Terniopsis_minor
Podostemaceae_Terniopsis_brevis
Podostemaceae_Terniopsis_malayana
Podostemaceae_Malaccotristicha_australis
Podostemaceae_Dalzellia_sessilis
Podostemaceae_Weddellina_squamulosa
Podostemaceae_Diamantina_lombardii
Podostemaceae_Mourera_elegans
Podostemaceae_Mourera_aspera
Podostemaceae_Mourera_fluviatilis
Podostemaceae_Mourera_weddelliana
Podostemaceae_Rhyncholacis_applanata
Podostemaceae_Oserya_perpusilla
Podostemaceae_Castelnavia_fluitans
Podostemaceae_Castelnavia_monandra
Podostemaceae_Castelnavia_multipartita
Podostemaceae_Castelnavia_princeps
Podostemaceae_Rhyncholacis_penicillata
Podostemaceae_Apinagia_nana
Podostemaceae_Wettsteiniola_sp
Podostemaceae_Apinagia_fimbrifolia
Podostemaceae_Marathrum_aeruginosum
Podostemaceae_Jenmaniella_fimbriata
Podostemaceae_Monostylis_capillacea
Podostemaceae_Lophogyne_lacunosa
Podostemaceae_Marathrum_schiedeanum
Podostemaceae_Marathrum_rubrum
Podostemaceae_Marathrum_oxycarpum
Podostemaceae_Marathrum_foeniculaceum
Podostemaceae_Vanroyenella_plumosa
Podostemaceae_Marathrum_tenue
Podostemaceae_Marathrum_utile
Podostemaceae_Oserya_longifolia
Podostemaceae_Oserya_coulteriana
Podostemaceae_Marathrum_modestum
Podostemaceae_Apinagia_richardiana
Podostemaceae_Apinagia_staheliana
Podostemaceae_Apinagia_corymbosa
Podostemaceae_Apinagia_riedelii
Podostemaceae_Apinagia_yguazuensis
Podostemaceae_Apinagia_longifolia
Podostemaceae_Apinagia_surumuensis
Podostemaceae_Apinagia_fluitans
Podostemaceae_Jenmaniella_ceratophylla
Podostemaceae_Rhyncholacis_sp
Podostemaceae_Ceratolacis_pedunculatum
Podostemaceae_Inversodicraea_ledermannii
Podostemaceae_Inversodicraea_cristata
Podostemaceae_Ledermanniella_ntemensis
Podostemaceae_Saxicolella_nana
Podostemaceae_Ledermanniella_bifurcata

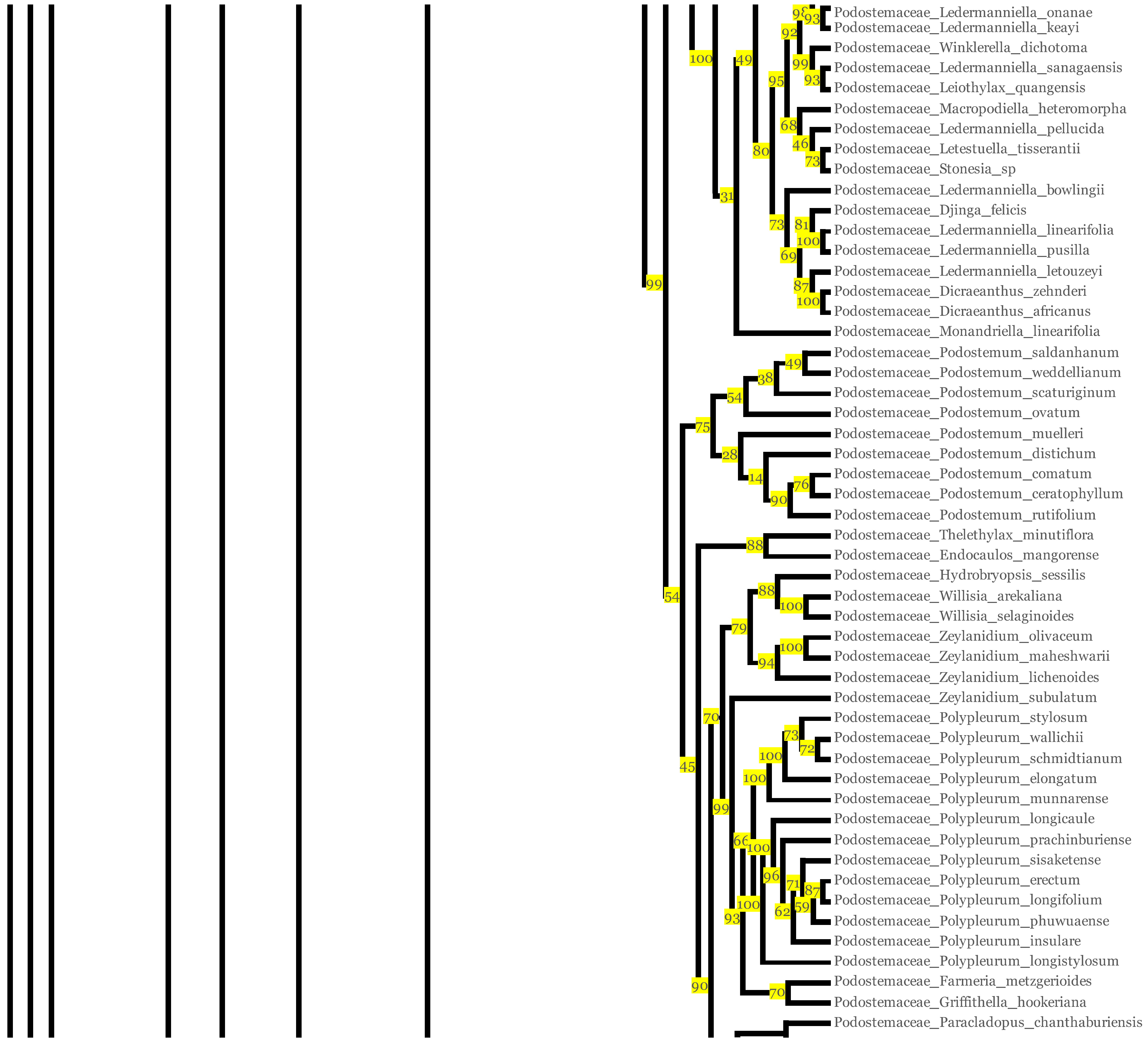
Podostemaceae_Ledermanniella_onanae
Podostemaceae_Ledermanniella_keayi
Podostemaceae_Winklerella_dichotoma
Podostemaceae_Ledermanniella_sanagaensis
Podostemaceae_Leiothylax_quangensis
Podostemaceae_Macropodiella_heteromorpha
Podostemaceae_Ledermanniella_pellucida
Podostemaceae_Letestuella_tisserantii
Podostemaceae_Stonesia_sp
Podostemaceae_Ledermanniella_bowlingii
Podostemaceae_Djinga_felicis
Podostemaceae_Ledermanniella_linearifolia
Podostemaceae_Ledermanniella_pusilla
Podostemaceae_Ledermanniella_letouzeyi
Podostemaceae_Dicraeanthus_zehnderi
Podostemaceae_Dicraeanthus_africanus
Podostemaceae_Monandriella_linearifolia
Podostemaceae_Podostemum_saldanhanum
Podostemaceae_Podostemum_weddellianum
Podostemaceae_Podostemum_scaturiginum
Podostemaceae_Podostemum_ovatum
Podostemaceae_Podostemum_muelleri
Podostemaceae_Podostemum_distichum
Podostemaceae_Podostemum_comatum
Podostemaceae_Podostemum_ceratophyllum
Podostemaceae_Podostemum_rutifolium
Podostemaceae_Thelethylax_minutiflora
Podostemaceae_Endocaulos_mangorense
Podostemaceae_Hydrobryopsis_sessilis
Podostemaceae_Willisia_arekaliana
Podostemaceae_Willisia_selaginoides
Podostemaceae_Zeylanidium_olivaceum
Podostemaceae_Zeylanidium_maheshwarii
Podostemaceae_Zeylanidium_lichenoides
Podostemaceae_Zeylanidium_subulatum
Podostemaceae_Polypleurum_stylosum
Podostemaceae_Polypleurum_wallichii
Podostemaceae_Polypleurum_schmidtianum
Podostemaceae_Polypleurum_elongatum
Podostemaceae_Polypleurum_munnarense
Podostemaceae_Polypleurum_longicaule
Podostemaceae_Polypleurum_prachinburiense
Podostemaceae_Polypleurum_sisaketense
Podostemaceae_Polypleurum_erectum
Podostemaceae_Polypleurum_longifolium
Podostemaceae_Polypleurum_phuwuaense
Podostemaceae_Polypleurum_insulare
Podostemaceae_Polypleurum_longistylosum
Podostemaceae_Farmeria_metzgerioides
Podostemaceae_Griffithella_hookeriana
Podostemaceae_Paracladopus_chanthaburiensis

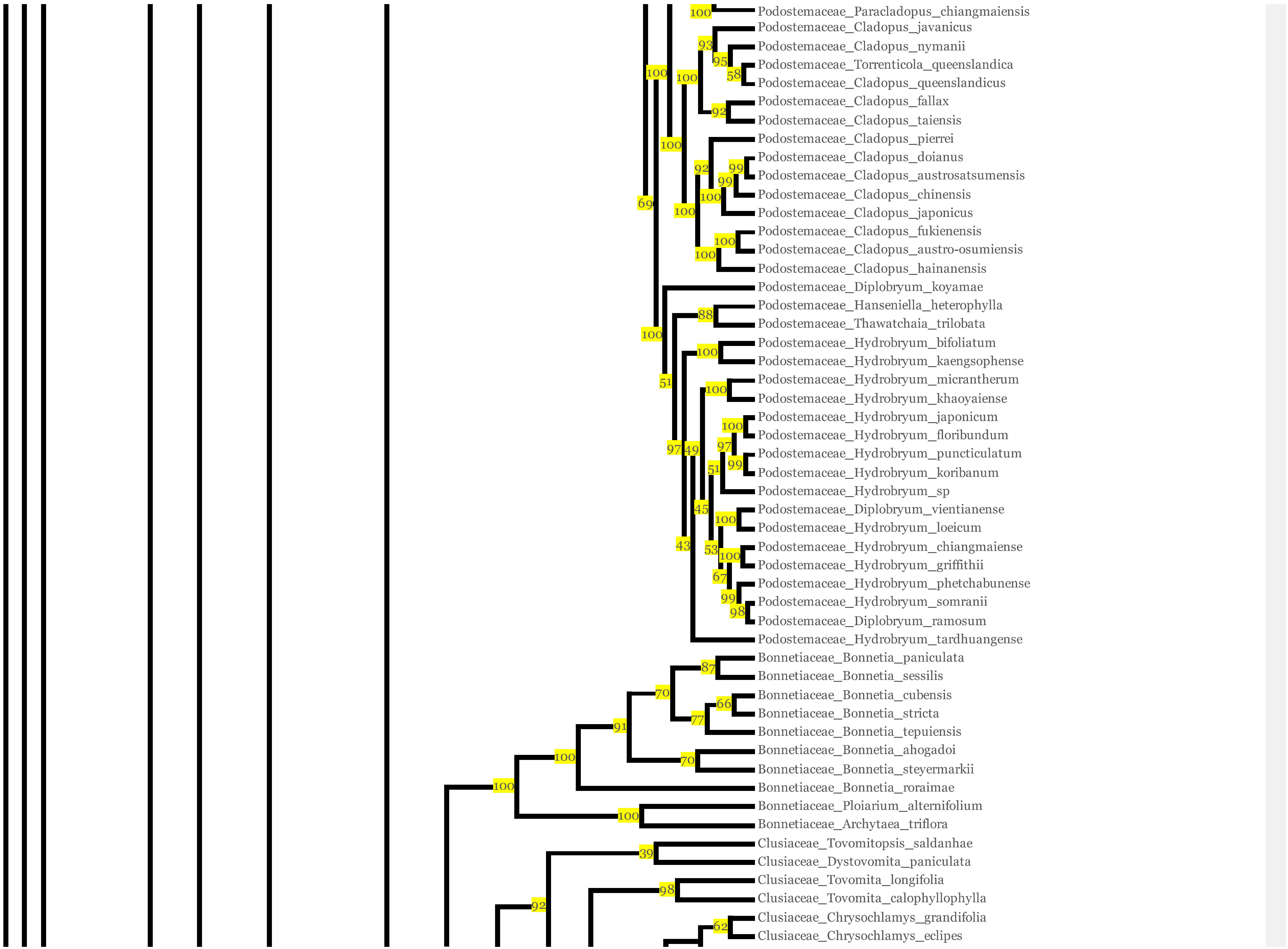
Podostemaceae_Paracladopus_chiangmaiensis
Podostemaceae_Cladopus_javanicus
Podostemaceae_Cladopus_nymanii
Podostemaceae_Torrenticola_queenslandica
Podostemaceae_Cladopus_queenslandicus
Podostemaceae_Cladopus_fallax
Podostemaceae_Cladopus_taiensis
Podostemaceae_Cladopus_pierrei
Podostemaceae_Cladopus_doianus
Podostemaceae_Cladopus_austrosatsumensis
Podostemaceae_Cladopus_chinensis
Podostemaceae_Cladopus_japonicus
Podostemaceae_Cladopus_fukienensis
Podostemaceae_Cladopus_austro-osumiensis
Podostemaceae_Cladopus_hainanensis
Podostemaceae_Diplobryum_koyamae
Podostemaceae_Hanseniella_heterophylla
Podostemaceae_Thawatchaia_trilobata
Podostemaceae_Hydrobryum_bifoliatum
Podostemaceae_Hydrobryum_kaengsophense
Podostemaceae_Hydrobryum_micrantherum
Podostemaceae_Hydrobryum_khaoyaiense
Podostemaceae_Hydrobryum_japonicum
Podostemaceae_Hydrobryum_floribundum
Podostemaceae_Hydrobryum_puncticulatum
Podostemaceae_Hydrobryum_koribanum
Podostemaceae_Hydrobryum_sp
Podostemaceae_Diplobryum_vientianense
Podostemaceae_Hydrobryum_loeicum
Podostemaceae_Hydrobryum_chiangmaiense
Podostemaceae_Hydrobryum_griffithii
Podostemaceae_Hydrobryum_phetchabunense
Podostemaceae_Hydrobryum_somranii
Podostemaceae_Diplobryum_ramosum
Podostemaceae_Hydrobryum_tardhuangense
Bonnetiaceae_Bonnetia_paniculata
Bonnetiaceae_Bonnetia_sessilis
Bonnetiaceae_Bonnetia_cubensis
Bonnetiaceae_Bonnetia_stricta
Bonnetiaceae_Bonnetia_tepuiensis
Bonnetiaceae_Bonnetia_ahogadoi
Bonnetiaceae_Bonnetia_steyermarkii
Bonnetiaceae_Bonnetia_roraimae
Bonnetiaceae_Ploiarium_alternifolium
Bonnetiaceae_Archytaea_triflora
Clusiaceae_Tovomitopsis_saldanhae
Clusiaceae_Dystovomita_paniculata
Clusiaceae_Tovomita_longifolia
Clusiaceae_Tovomita_calophyllophylla
Clusiaceae_Chrysochlamys_grandifolia
Clusiaceae_Chrysochlamys_eclipes

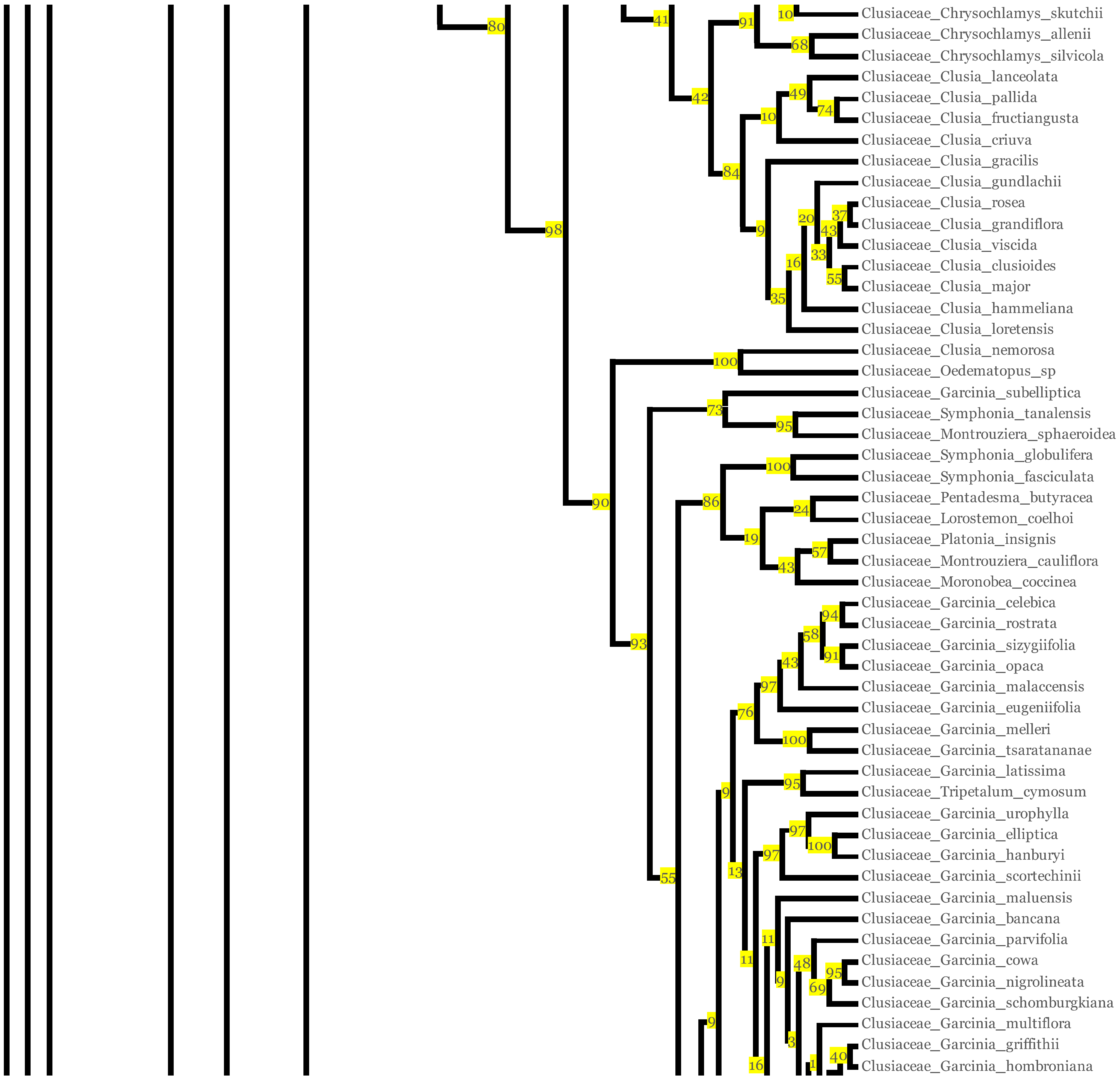
Clusiaceae_Chrysochlamys_skutchii
Clusiaceae_Chrysochlamys_allenii
Clusiaceae_Chrysochlamys_silvicola
Clusiaceae_Clusia_lanceolata
Clusiaceae_Clusia_pallida
Clusiaceae_Clusia_fructiangusta
Clusiaceae_Clusia_criuva
Clusiaceae_Clusia_gracilis
Clusiaceae_Clusia_gundlachii
Clusiaceae_Clusia_rosea
Clusiaceae_Clusia_grandiflora
Clusiaceae_Clusia_viscida
Clusiaceae_Clusia_clusioides
Clusiaceae_Clusia_major
Clusiaceae_Clusia_hammeliana
Clusiaceae_Clusia_loretensis
Clusiaceae_Clusia_nemorosa
Clusiaceae_Oedematopus_sp
Clusiaceae_Garcinia_subelliptica
Clusiaceae_Symphonia_tanalensis
Clusiaceae_Montrouziera_sphaeroidea
Clusiaceae_Symphonia_globulifera
Clusiaceae_Symphonia_fasciculata
Clusiaceae_Pentadesma_butyracea
Clusiaceae_Lorostemon_coelhoi
Clusiaceae_Platonia_insignis
Clusiaceae_Montrouziera_cauliflora
Clusiaceae_Moronobea_coccinea
Clusiaceae_Garcinia_celebica
Clusiaceae_Garcinia_rostrata
Clusiaceae_Garcinia_sizygiifolia
Clusiaceae_Garcinia_opaca
Clusiaceae_Garcinia_malaccensis
Clusiaceae_Garcinia_eugeniifolia
Clusiaceae_Garcinia_melleri
Clusiaceae_Garcinia_tsaratananae
Clusiaceae_Garcinia_latissima
Clusiaceae_Tripetalum_cymosum
Clusiaceae_Garcinia_urophylla
Clusiaceae_Garcinia_elliptica
Clusiaceae_Garcinia_hanburyi
Clusiaceae_Garcinia_scortechinii
Clusiaceae_Garcinia_maluensis
Clusiaceae_Garcinia_bancana
Clusiaceae_Garcinia_parvifolia
Clusiaceae_Garcinia_cowa
Clusiaceae_Garcinia_nigrolineata
Clusiaceae_Garcinia_schomburgkiana
Clusiaceae_Garcinia_multiflora
Clusiaceae_Garcinia_griffithii
Clusiaceae_Garcinia_hombroniana

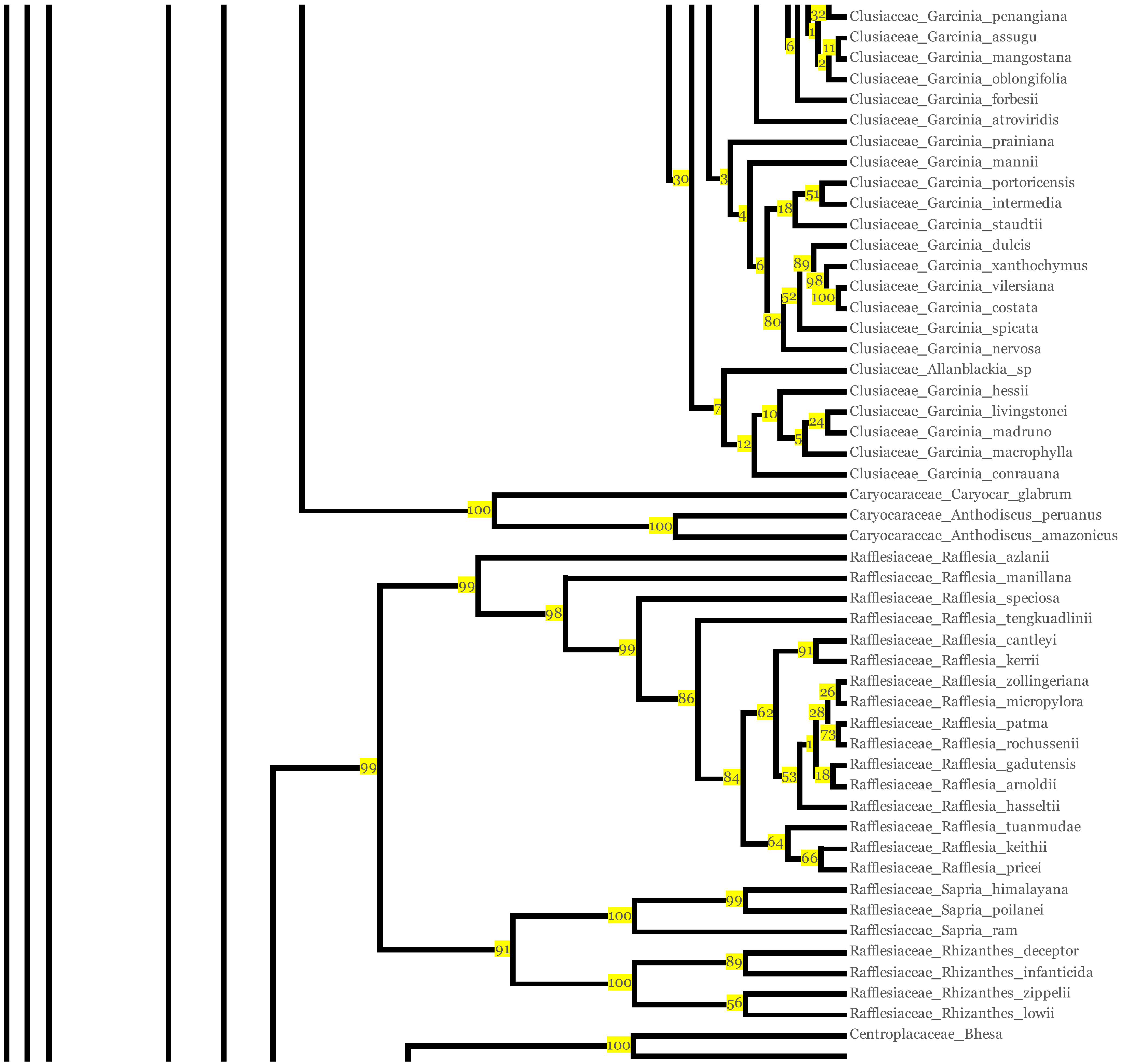
Clusiaceae_Garcinia_penangiana
Clusiaceae_Garcinia_assugu
Clusiaceae_Garcinia_mangostana
Clusiaceae_Garcinia_oblongifolia
Clusiaceae_Garcinia_forbesii
Clusiaceae_Garcinia_atroviridis
Clusiaceae_Garcinia_prainiana
Clusiaceae_Garcinia_mannii
Clusiaceae_Garcinia_portoricensis
Clusiaceae_Garcinia_intermedia
Clusiaceae_Garcinia_staudtii
Clusiaceae_Garcinia_dulcis
Clusiaceae_Garcinia_xanthochymus
Clusiaceae_Garcinia_vilersiana
Clusiaceae_Garcinia_costata
Clusiaceae_Garcinia_spicata
Clusiaceae_Garcinia_nervosa
Clusiaceae_Allanblackia_sp
Clusiaceae_Garcinia_hessii
Clusiaceae_Garcinia_livingstonei
Clusiaceae_Garcinia_madruno
Clusiaceae_Garcinia_macrophylla
Clusiaceae_Garcinia_conrauana
Caryocaraceae_Caryocar_glabrum
Caryocaraceae_Anthodiscus_peruanus
Caryocaraceae_Anthodiscus_amazonicus
Rafflesiaceae_Rafflesia_azlanii
Rafflesiaceae_Rafflesia_manillana
Rafflesiaceae_Rafflesia_speciosa
Rafflesiaceae_Rafflesia_tengkuadlinii
Rafflesiaceae_Rafflesia_cantleyi
Rafflesiaceae_Rafflesia_kerrii
Rafflesiaceae_Rafflesia_zollingeriana
Rafflesiaceae_Rafflesia_micropylora
Rafflesiaceae_Rafflesia_patma
Rafflesiaceae_Rafflesia_rochussenii
Rafflesiaceae_Rafflesia_gadutensis
Rafflesiaceae_Rafflesia_arnoldii
Rafflesiaceae_Rafflesia_hasseltii
Rafflesiaceae_Rafflesia_tuanmudae
Rafflesiaceae_Rafflesia_keithii
Rafflesiaceae_Rafflesia_pricei
Rafflesiaceae_Sapria_himalayana
Rafflesiaceae_Sapria_poilanei
Rafflesiaceae_Sapria_ram
Rafflesiaceae_Rhizanthes_deceptor
Rafflesiaceae_Rhizanthes_infanticida
Rafflesiaceae_Rhizanthes_zippelii
Rafflesiaceae_Rhizanthes_lowii
Centroplacaceae_Bhesa

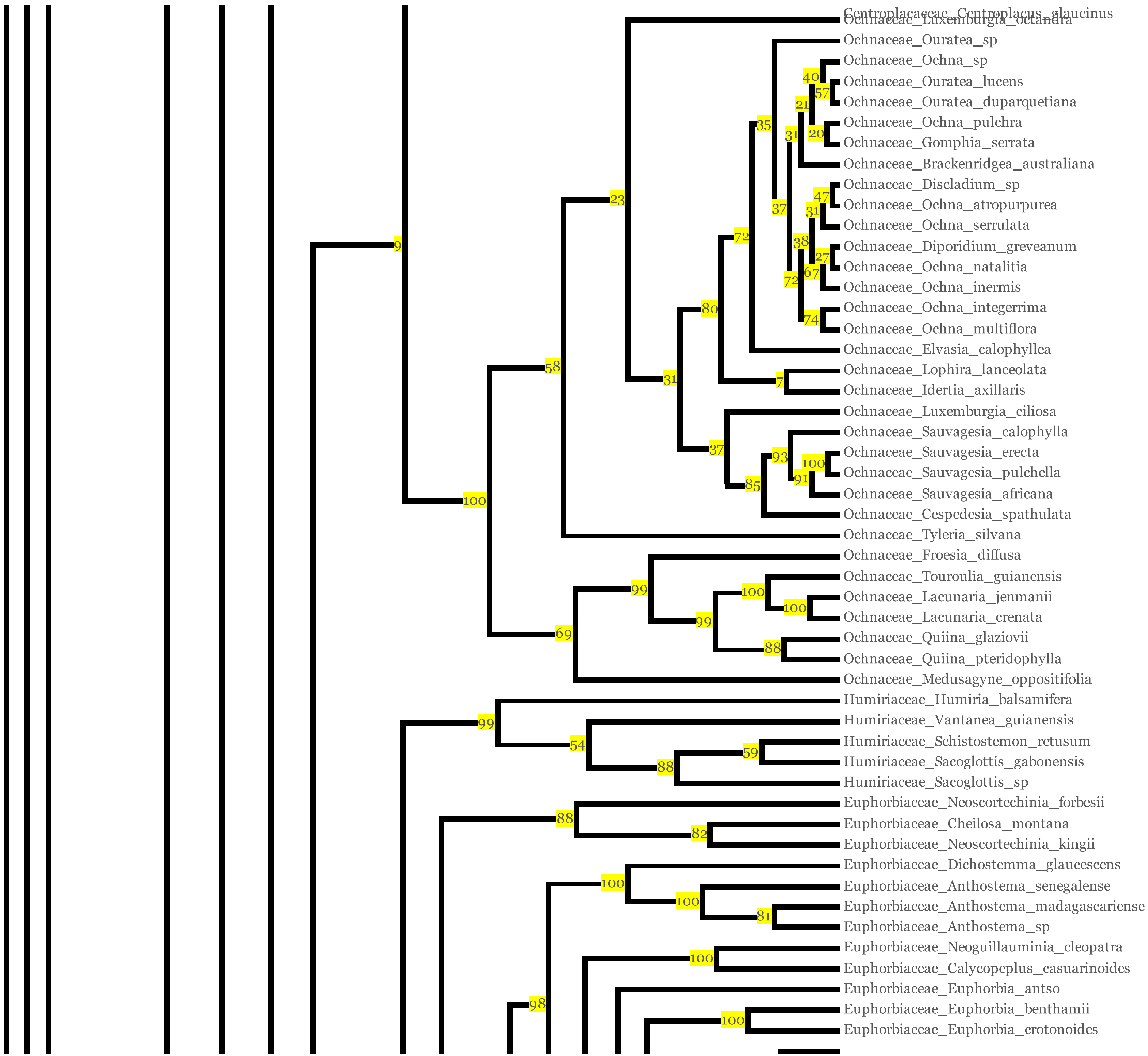
Centroplacaceae_Centroplacus_glaucinus
Ochnaceae_Luxemburgia_octandra
Ochnaceae_Ouratea_sp
Ochnaceae_Ochna_sp
Ochnaceae_Ouratea_lucens
Ochnaceae_Ouratea_duparquetiana
Ochnaceae_Ochna_pulchra
Ochnaceae_Gomphia_serrata
Ochnaceae_Brackenridgea_australiana
Ochnaceae_Discladium_sp
Ochnaceae_Ochna_atropurpurea
Ochnaceae_Ochna_serrulata
Ochnaceae_Diporidium_greveanum
Ochnaceae_Ochna_natalitia
Ochnaceae_Ochna_inermis
Ochnaceae_Ochna_integerrima
Ochnaceae_Ochna_multiflora
Ochnaceae_Elvasia_calophyllea
Ochnaceae_Lophira_lanceolata
Ochnaceae_Idertia_axillaris
Ochnaceae_Luxemburgia_ciliosa
Ochnaceae_Sauvagesia_calophylla
Ochnaceae_Sauvagesia_erecta
Ochnaceae_Sauvagesia_pulchella
Ochnaceae_Sauvagesia_africana
Ochnaceae_Cespedesia_spathulata
Ochnaceae_Tyleria_silvana
Ochnaceae_Froesia_diffusa
Ochnaceae_Touroulia_guianensis
Ochnaceae_Lacunaria_jenmanii
Ochnaceae_Lacunaria_crenata
Ochnaceae_Quiina_glaziovii
Ochnaceae_Quiina_pteridophylla
Ochnaceae_Medusagyne_oppositifolia
Humiriaceae_Humiria_balsamifera
Humiriaceae_Vantanea_guianensis
Humiriaceae_Schistostemon_retusum
Humiriaceae_Sacoglottis_gabonensis
Humiriaceae_Sacoglottis_sp
Euphorbiaceae_Neoscortechinia_forbesii
Euphorbiaceae_Cheilosa_montana
Euphorbiaceae_Neoscortechinia_kingii
Euphorbiaceae_Dichostemma_glaucescens
Euphorbiaceae_Anthostema_senegalense
Euphorbiaceae_Anthostema_madagascariense
Euphorbiaceae_Anthostema_sp
Euphorbiaceae_Neoguillauminia_cleopatra
Euphorbiaceae_Calycopeplus_casuarinoides
Euphorbiaceae_Euphorbia_antso
Euphorbiaceae_Euphorbia_benthamii
Euphorbiaceae_Euphorbia_crotonoides

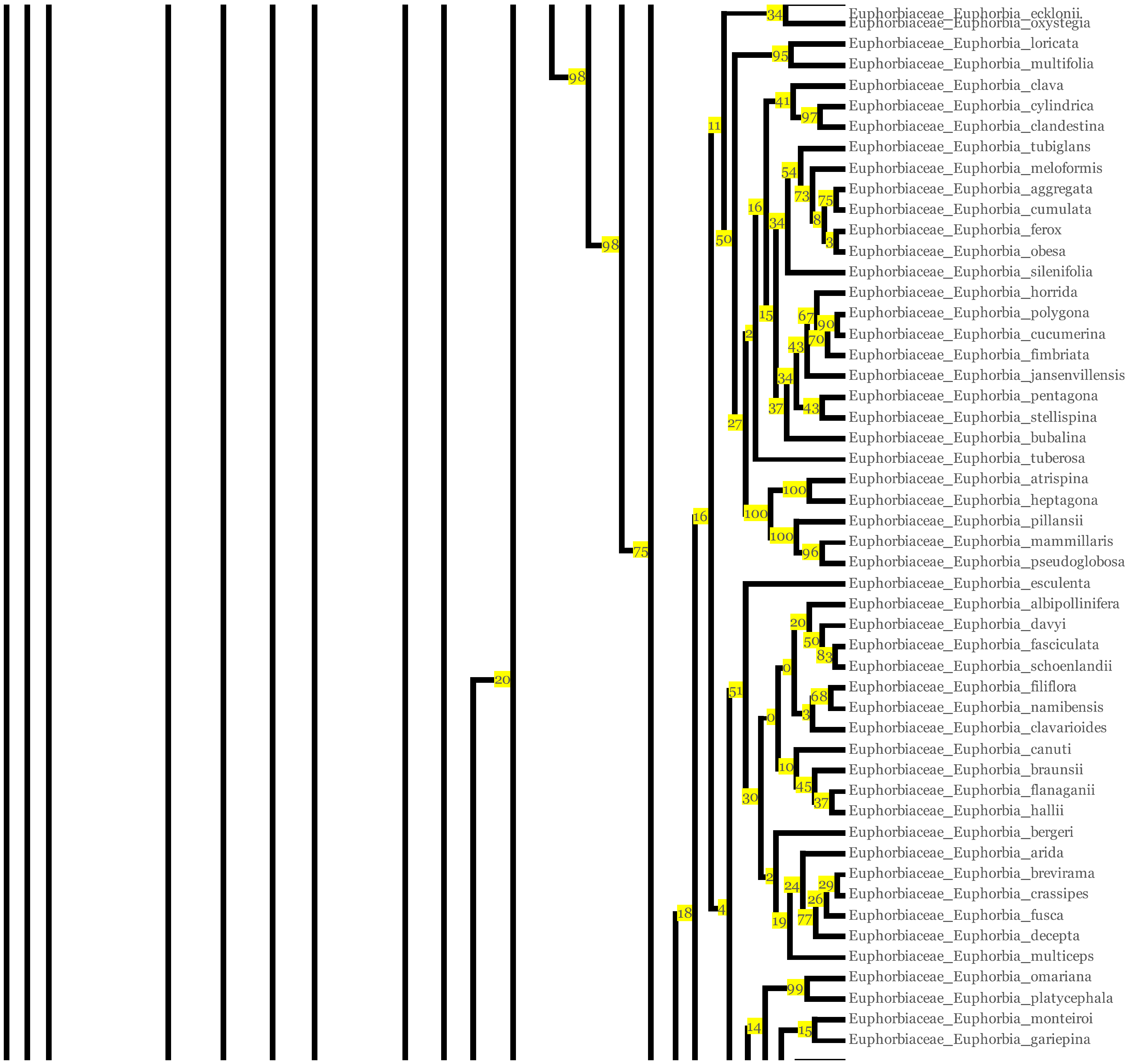
Euphorbiaceae_Euphorbia_ecklonii
Euphorbiaceae_Euphorbia_oxystegia
Euphorbiaceae_Euphorbia_loricata
Euphorbiaceae_Euphorbia_multifolia
Euphorbiaceae_Euphorbia_clava
Euphorbiaceae_Euphorbia_cylindrica
Euphorbiaceae_Euphorbia_clandestina
Euphorbiaceae_Euphorbia_tubiglans
Euphorbiaceae_Euphorbia_meloformis
Euphorbiaceae_Euphorbia_aggregata
Euphorbiaceae_Euphorbia_cumulata
Euphorbiaceae_Euphorbia_ferox
Euphorbiaceae_Euphorbia_obesa
Euphorbiaceae_Euphorbia_silenifolia
Euphorbiaceae_Euphorbia_horrida
Euphorbiaceae_Euphorbia_polygona
Euphorbiaceae_Euphorbia_cucumerina
Euphorbiaceae_Euphorbia_fimbriata
Euphorbiaceae_Euphorbia_jansenvillensis
Euphorbiaceae_Euphorbia_pentagona
Euphorbiaceae_Euphorbia_stellispina
Euphorbiaceae_Euphorbia_bubalina
Euphorbiaceae_Euphorbia_tuberosa
Euphorbiaceae_Euphorbia_atrispina
Euphorbiaceae_Euphorbia_heptagona
Euphorbiaceae_Euphorbia_pillansii
Euphorbiaceae_Euphorbia_mammillaris
Euphorbiaceae_Euphorbia_pseudoglobosa
Euphorbiaceae_Euphorbia_esculenta
Euphorbiaceae_Euphorbia_albipollinifera
Euphorbiaceae_Euphorbia_davyi
Euphorbiaceae_Euphorbia_fasciculata
Euphorbiaceae_Euphorbia_schoenlandii
Euphorbiaceae_Euphorbia_filiflora
Euphorbiaceae_Euphorbia_namibensis
Euphorbiaceae_Euphorbia_clavarioides
Euphorbiaceae_Euphorbia_canuti
Euphorbiaceae_Euphorbia_braunsii
Euphorbiaceae_Euphorbia_flanaganii
Euphorbiaceae_Euphorbia_hallii
Euphorbiaceae_Euphorbia_bergeri
Euphorbiaceae_Euphorbia_arida
Euphorbiaceae_Euphorbia_brevirama
Euphorbiaceae_Euphorbia_crassipes
Euphorbiaceae_Euphorbia_fusca
Euphorbiaceae_Euphorbia_decepta
Euphorbiaceae_Euphorbia_multiceps
Euphorbiaceae_Euphorbia_omariana
Euphorbiaceae_Euphorbia_platycephala
Euphorbiaceae_Euphorbia_monteiroi
Euphorbiaceae_Euphorbia_gariepina

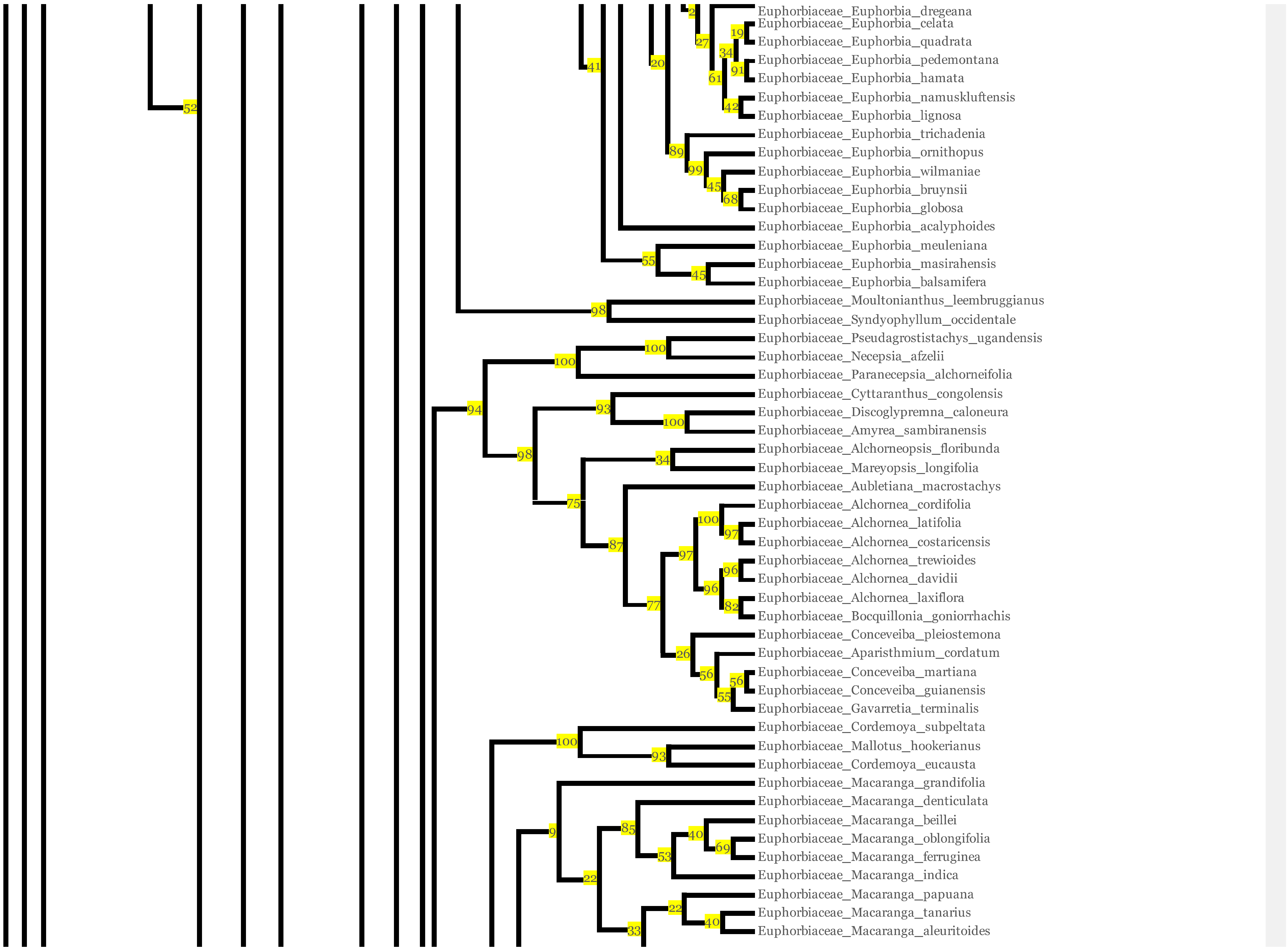
Euphorbiaceae_Euphorbia_dregeana
Euphorbiaceae_Euphorbia_celata
Euphorbiaceae_Euphorbia_quadrata
Euphorbiaceae_Euphorbia_pedemontana
Euphorbiaceae_Euphorbia_hamata
Euphorbiaceae_Euphorbia_namuskluftensis
Euphorbiaceae_Euphorbia_lignosa
Euphorbiaceae_Euphorbia_trichadenia
Euphorbiaceae_Euphorbia_ornithopus
Euphorbiaceae_Euphorbia_wilmaniae
Euphorbiaceae_Euphorbia_bruynsii
Euphorbiaceae_Euphorbia_globosa
Euphorbiaceae_Euphorbia_acalyphoides
Euphorbiaceae_Euphorbia_meuleniana
Euphorbiaceae_Euphorbia_masirahensis
Euphorbiaceae_Euphorbia_balsamifera
Euphorbiaceae_Moultonianthus_leembruggianus
Euphorbiaceae_Syndyophyllum_occidentale
Euphorbiaceae_Pseudagrostistachys_ugandensis
Euphorbiaceae_Necepsia_afzelii
Euphorbiaceae_Paranecepsia_alchorneifolia
Euphorbiaceae_Cyttaranthus_congolensis
Euphorbiaceae_Discoglypremna_caloneura
Euphorbiaceae_Amyrea_sambiranensis
Euphorbiaceae_Alchorneopsis_floribunda
Euphorbiaceae_Mareyopsis_longifolia
Euphorbiaceae_Aubletiana_macrostachys
Euphorbiaceae_Alchornea_cordifolia
Euphorbiaceae_Alchornea_latifolia
Euphorbiaceae_Alchornea_costaricensis
Euphorbiaceae_Alchornea_trewioides
Euphorbiaceae_Alchornea_davidii
Euphorbiaceae_Alchornea_laxiflora
Euphorbiaceae_Bocquillonia_goniorrhachis
Euphorbiaceae_Conceveiba_pleiostemona
Euphorbiaceae_Aparisthmium_cordatum
Euphorbiaceae_Conceveiba_martiana
Euphorbiaceae_Conceveiba_guianensis
Euphorbiaceae_Gavarretia_terminalis
Euphorbiaceae_Cordemoya_subpeltata
Euphorbiaceae_Mallotus_hookerianus
Euphorbiaceae_Cordemoya_eucausta
Euphorbiaceae_Macaranga_grandifolia
Euphorbiaceae_Macaranga_denticulata
Euphorbiaceae_Macaranga_beillei
Euphorbiaceae_Macaranga_oblongifolia
Euphorbiaceae_Macaranga_ferruginea
Euphorbiaceae_Macaranga_indica
Euphorbiaceae_Macaranga_papuana
Euphorbiaceae_Macaranga_tanarius
Euphorbiaceae_Macaranga_aleuritoides

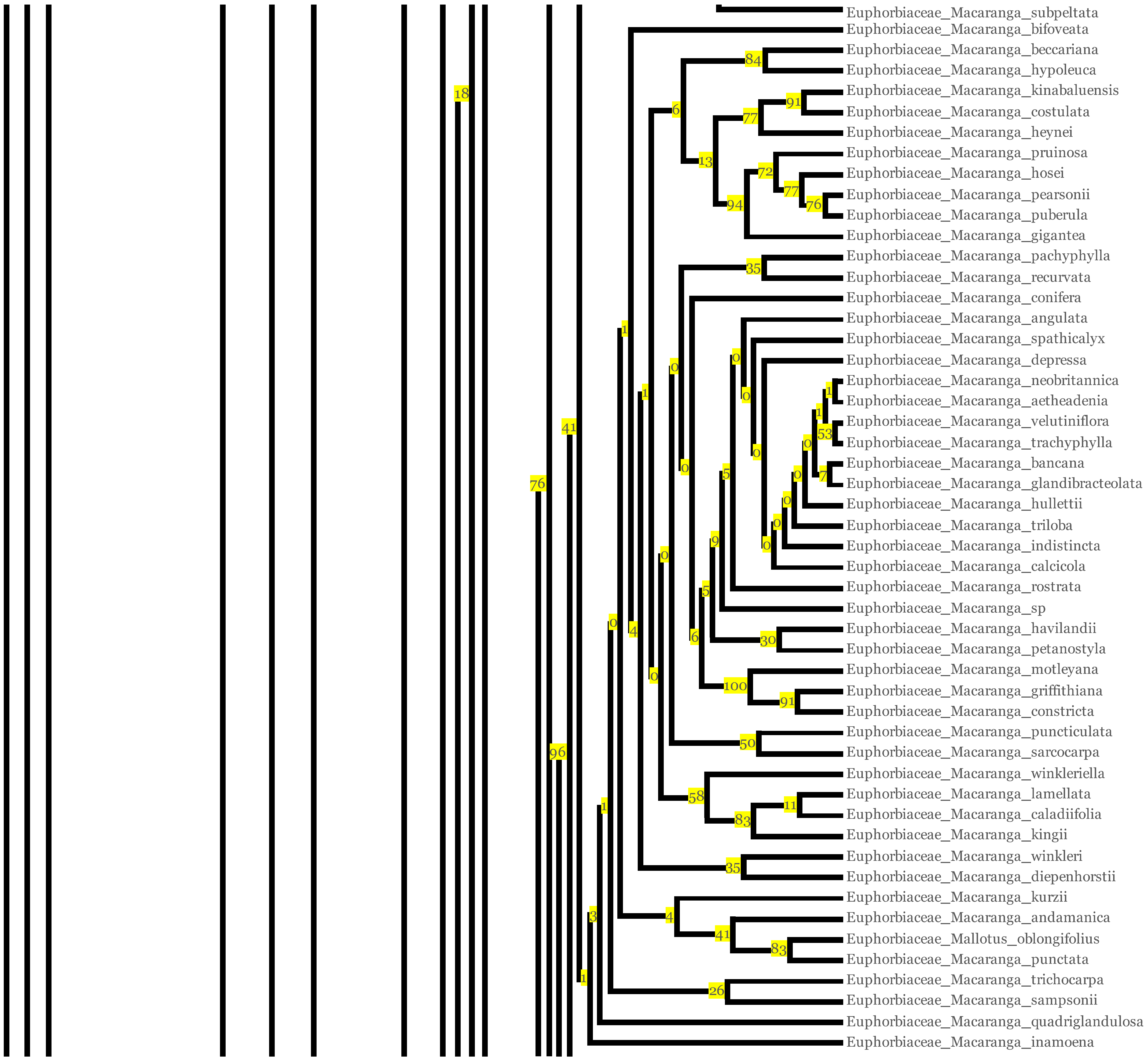
Euphorbiaceae_Macaranga_subpeltata
Euphorbiaceae_Macaranga_bifoveata
Euphorbiaceae_Macaranga_beccariana
Euphorbiaceae_Macaranga_hypoleuca
Euphorbiaceae_Macaranga_kinabaluensis
Euphorbiaceae_Macaranga_costulata
Euphorbiaceae_Macaranga_heynei
Euphorbiaceae_Macaranga_pruinosa
Euphorbiaceae_Macaranga_hosei
Euphorbiaceae_Macaranga_pearsonii
Euphorbiaceae_Macaranga_puberula
Euphorbiaceae_Macaranga_gigantea
Euphorbiaceae_Macaranga_pachyphylla
Euphorbiaceae_Macaranga_recurvata
Euphorbiaceae_Macaranga_conifera
Euphorbiaceae_Macaranga_angulata
Euphorbiaceae_Macaranga_spathicalyx
Euphorbiaceae_Macaranga_depressa
Euphorbiaceae_Macaranga_neobritannica
Euphorbiaceae_Macaranga_aetheadenia
Euphorbiaceae_Macaranga_velutiniflora
Euphorbiaceae_Macaranga_trachyphylla
Euphorbiaceae_Macaranga_bancana
Euphorbiaceae_Macaranga_glandibracteolata
Euphorbiaceae_Macaranga_hullettii
Euphorbiaceae_Macaranga_triloba
Euphorbiaceae_Macaranga_indistincta
Euphorbiaceae_Macaranga_calcicola
Euphorbiaceae_Macaranga_rostrata
Euphorbiaceae_Macaranga_sp
Euphorbiaceae_Macaranga_havilandii
Euphorbiaceae_Macaranga_petanostyla
Euphorbiaceae_Macaranga_motleyana
Euphorbiaceae_Macaranga_griffithiana
Euphorbiaceae_Macaranga_constricta
Euphorbiaceae_Macaranga_puncticulata
Euphorbiaceae_Macaranga_sarcocarpa
Euphorbiaceae_Macaranga_winkleriella
Euphorbiaceae_Macaranga_lamellata
Euphorbiaceae_Macaranga_caladiifolia
Euphorbiaceae_Macaranga_kingii
Euphorbiaceae_Macaranga_winkleri
Euphorbiaceae_Macaranga_diepenhorstii
Euphorbiaceae_Macaranga_kurzii
Euphorbiaceae_Macaranga_andamanica
Euphorbiaceae_Mallotus_oblongifolius
Euphorbiaceae_Macaranga_punctata
Euphorbiaceae_Macaranga_trichocarpa
Euphorbiaceae_Macaranga_sampsonii
Euphorbiaceae_Macaranga_quadriglandulosa
Euphorbiaceae_Macaranga_inamoena

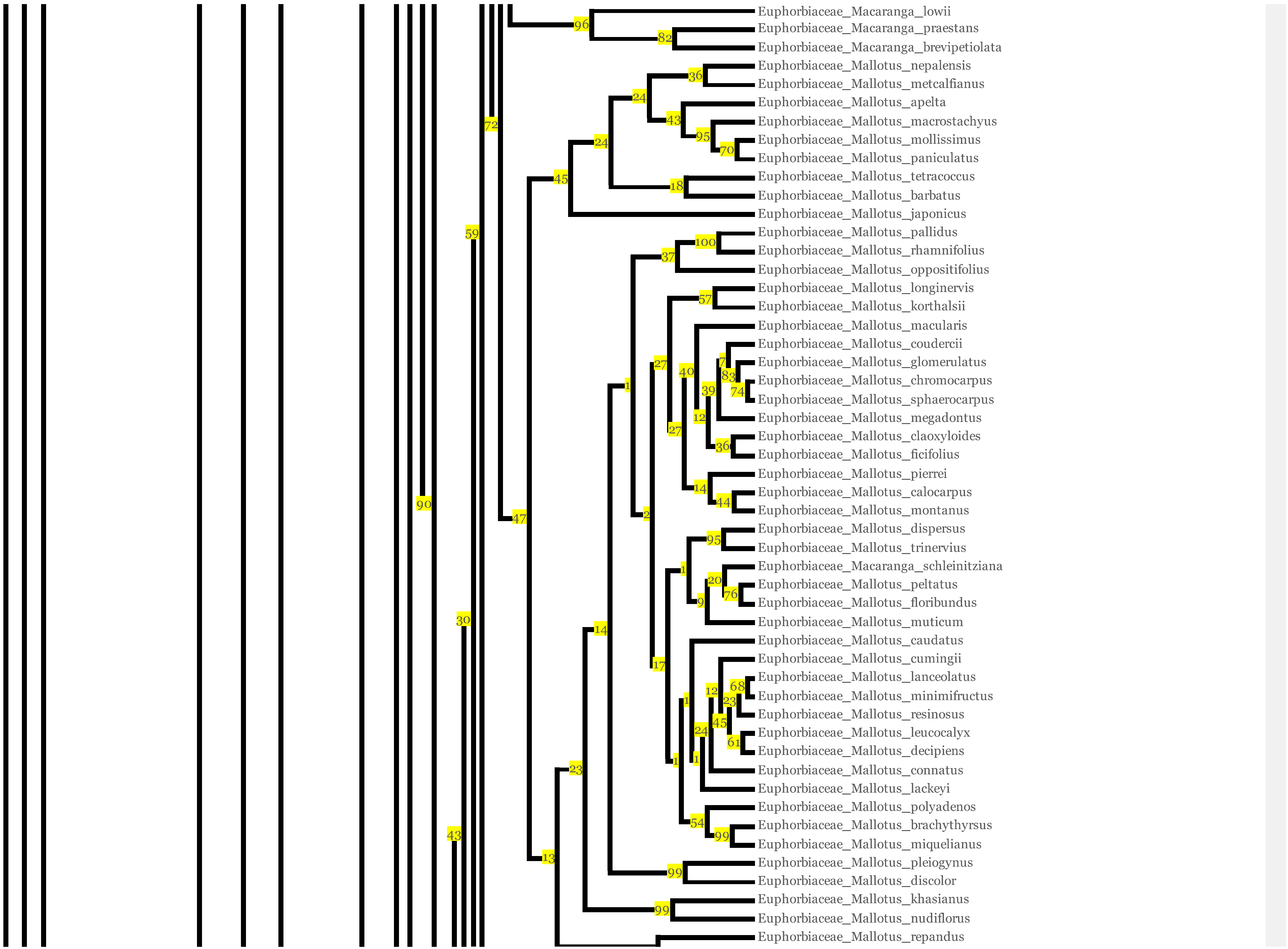
Euphorbiaceae_Macaranga_lowii
Euphorbiaceae_Macaranga_praestans
Euphorbiaceae_Macaranga_brevipetiolata
Euphorbiaceae_Mallotus_nepalensis
Euphorbiaceae_Mallotus_metcalfianus
Euphorbiaceae_Mallotus_apelta
Euphorbiaceae_Mallotus_macrostachyus
Euphorbiaceae_Mallotus_mollissimus
Euphorbiaceae_Mallotus_paniculatus
Euphorbiaceae_Mallotus_tetracoccus
Euphorbiaceae_Mallotus_barbatus
Euphorbiaceae_Mallotus_japonicus
Euphorbiaceae_Mallotus_pallidus
Euphorbiaceae_Mallotus_rhamnifolius
Euphorbiaceae_Mallotus_oppositifolius
Euphorbiaceae_Mallotus_longinervis
Euphorbiaceae_Mallotus_korthalsii
Euphorbiaceae_Mallotus_macularis
Euphorbiaceae_Mallotus_coudercii
Euphorbiaceae_Mallotus_glomerulatus
Euphorbiaceae_Mallotus_chromocarpus
Euphorbiaceae_Mallotus_sphaerocarpus
Euphorbiaceae_Mallotus_megadontus
Euphorbiaceae_Mallotus_claoxyloides
Euphorbiaceae_Mallotus_ficifolius
Euphorbiaceae_Mallotus_pierrei
Euphorbiaceae_Mallotus_calocarpus
Euphorbiaceae_Mallotus_montanus
Euphorbiaceae_Mallotus_dispersus
Euphorbiaceae_Mallotus_trinervius
Euphorbiaceae_Macaranga_schleinitziana
Euphorbiaceae_Mallotus_peltatus
Euphorbiaceae_Mallotus_floribundus
Euphorbiaceae_Mallotus_muticum
Euphorbiaceae_Mallotus_caudatus
Euphorbiaceae_Mallotus_cumingii
Euphorbiaceae_Mallotus_lanceolatus
Euphorbiaceae_Mallotus_minimifructus
Euphorbiaceae_Mallotus_resinosus
Euphorbiaceae_Mallotus_leucocalyx
Euphorbiaceae_Mallotus_decipiens
Euphorbiaceae_Mallotus_connatus
Euphorbiaceae_Mallotus_lackeyi
Euphorbiaceae_Mallotus_polyadenos
Euphorbiaceae_Mallotus_brachythyrsus
Euphorbiaceae_Mallotus_miquelianus
Euphorbiaceae_Mallotus_pleiogynus
Euphorbiaceae_Mallotus_discolor
Euphorbiaceae_Mallotus_khasianus
Euphorbiaceae_Mallotus_nudiflorus
Euphorbiaceae_Mallotus_repandus
96
82
36
24
43
95
70
72
24
18
45
59
100
37
57
27
40
7
83
39
74
12
1
27
36
14
44
90
47
2
95
1
20
76
9
30
14
17
12
68
23
1
45
24
61
1
1
23
54
99
43
13
99
99

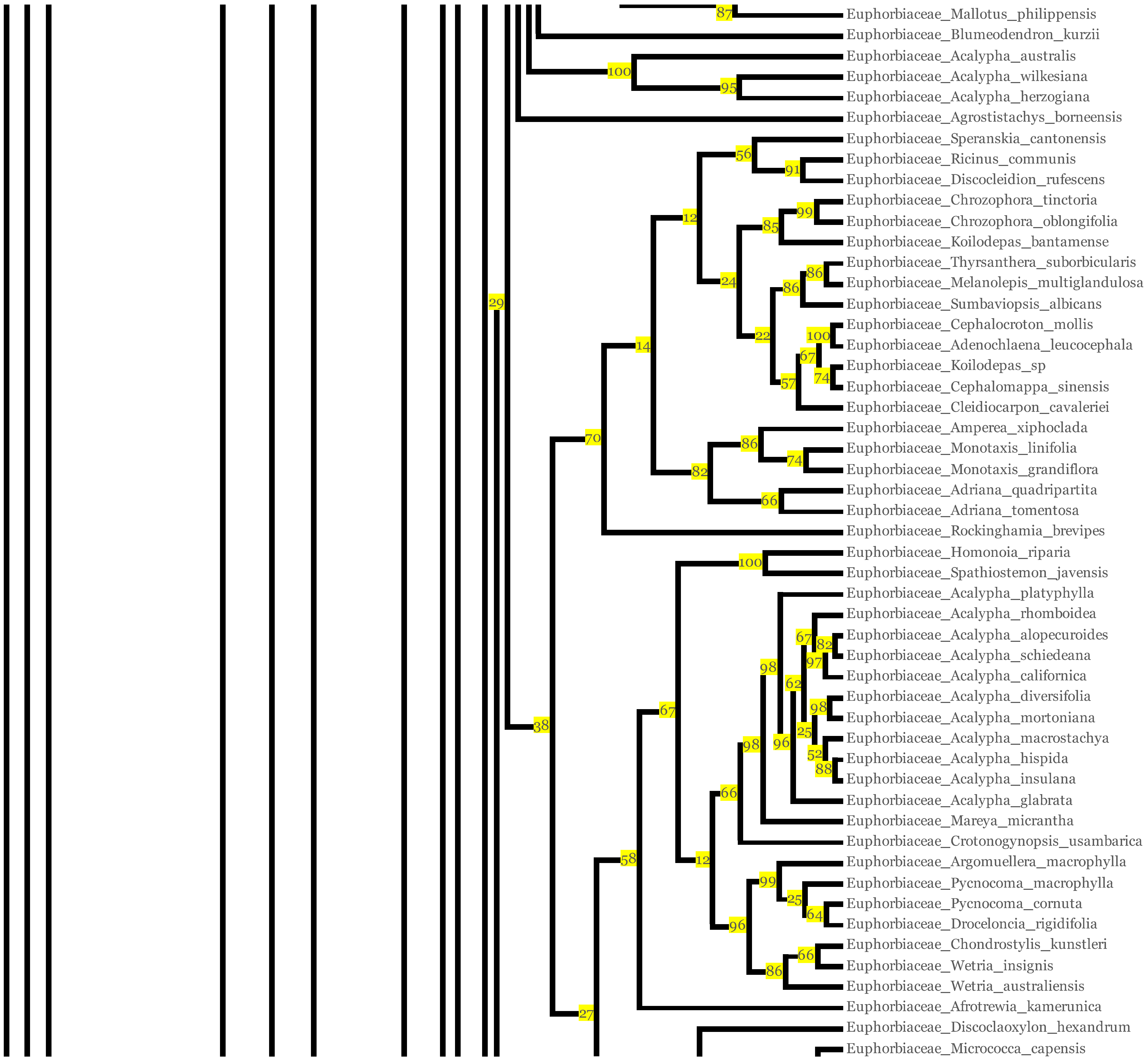
Euphorbiaceae_Mallotus_philippensis
Euphorbiaceae_Blumeodendron_kurzii
Euphorbiaceae_Acalypha_australis
Euphorbiaceae_Acalypha_wilkesiana
Euphorbiaceae_Acalypha_herzogiana
Euphorbiaceae_Agrostistachys_borneensis
Euphorbiaceae_Speranskia_cantonensis
Euphorbiaceae_Ricinus_communis
Euphorbiaceae_Discocleidion_rufescens
Euphorbiaceae_Chrozophora_tinctoria
Euphorbiaceae_Chrozophora_oblongifolia
Euphorbiaceae_Koilodepas_bantamense
Euphorbiaceae_Thyrsanthera_suborbicularis
Euphorbiaceae_Melanolepis_multiglandulosa
Euphorbiaceae_Sumbaviopsis_albicans
Euphorbiaceae_Cephalocroton_mollis
Euphorbiaceae_Adenochlaena_leucocephala
Euphorbiaceae_Koilodepas_sp
Euphorbiaceae_Cephalomappa_sinensis
Euphorbiaceae_Cleidiocarpon_cavaleriei
Euphorbiaceae_Amperea_xiphoclada
Euphorbiaceae_Monotaxis_linifolia
Euphorbiaceae_Monotaxis_grandiflora
Euphorbiaceae_Adriana_quadripartita
Euphorbiaceae_Adriana_tomentosa
Euphorbiaceae_Rockinghamia_brevipes
Euphorbiaceae_Homonoia_riparia
Euphorbiaceae_Spathiostemon_javensis
Euphorbiaceae_Acalypha_platyphylla
Euphorbiaceae_Acalypha_rhomboidea
Euphorbiaceae_Acalypha_alopecuroides
Euphorbiaceae_Acalypha_schiedeana
Euphorbiaceae_Acalypha_californica
Euphorbiaceae_Acalypha_diversifolia
Euphorbiaceae_Acalypha_mortoniana
Euphorbiaceae_Acalypha_macrostachya
Euphorbiaceae_Acalypha_hispida
Euphorbiaceae_Acalypha_insulana
Euphorbiaceae_Acalypha_glabrata
Euphorbiaceae_Mareya_micrantha
Euphorbiaceae_Crotonogynopsis_usambarica
Euphorbiaceae_Argomuellera_macrophylla
Euphorbiaceae_Pycnocoma_macrophylla
Euphorbiaceae_Pycnocoma_cornuta
Euphorbiaceae_Droceloncia_rigidifolia
Euphorbiaceae_Chondrostylis_kunstleri
Euphorbiaceae_Wetria_insignis
Euphorbiaceae_Wetria_australiensis
Euphorbiaceae_Afrotrewia_kamerunica
Euphorbiaceae_Discoclaoxylon_hexandrum
Euphorbiaceae_Micrococca_capensis

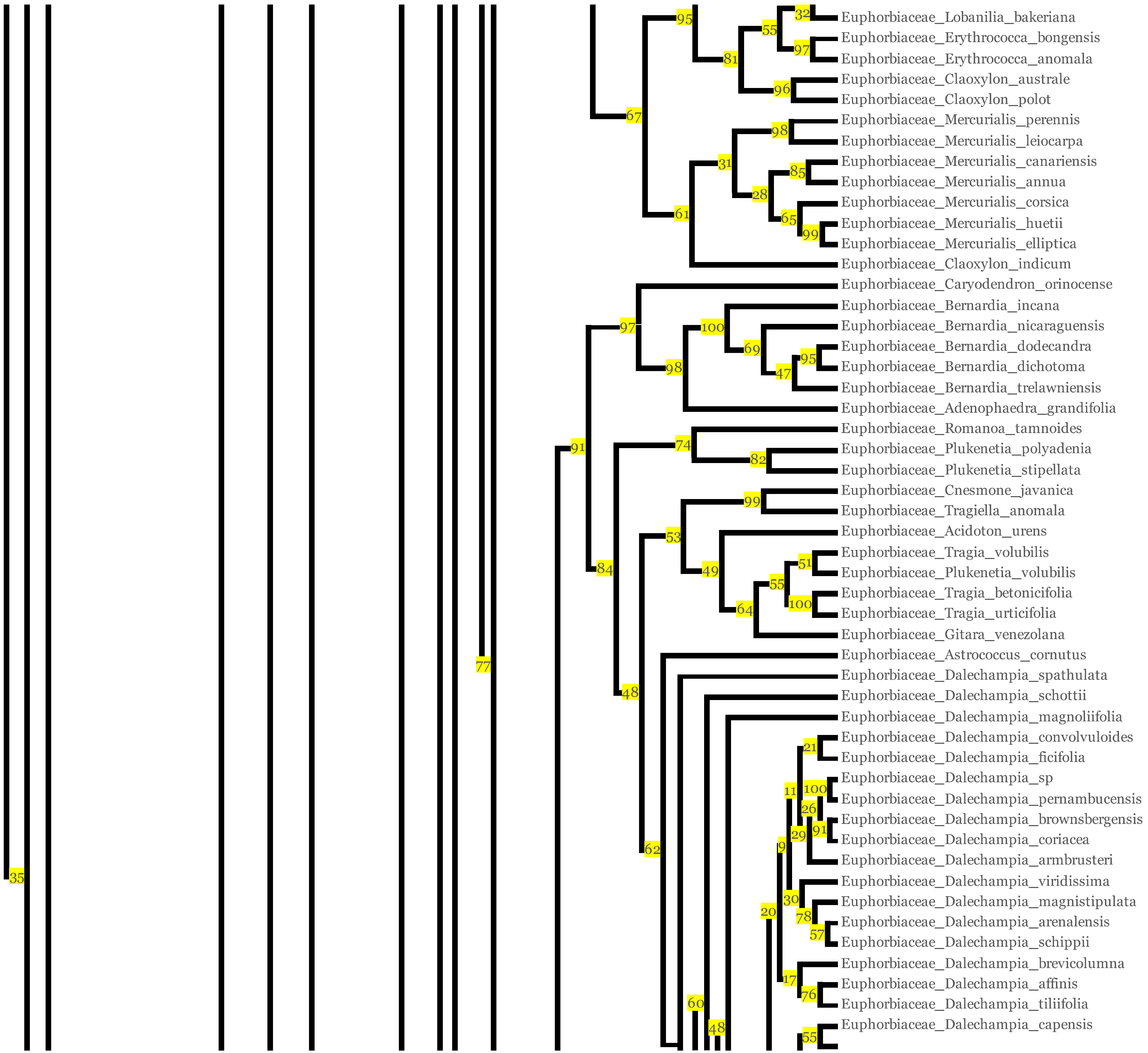
Euphorbiaceae_Lobanilia_bakeriana
Euphorbiaceae_Erythrococca_bongensis
Euphorbiaceae_Erythrococca_anomala
Euphorbiaceae_Claoxylon_australe
Euphorbiaceae_Claoxylon_polot
Euphorbiaceae_Mercurialis_perennis
Euphorbiaceae_Mercurialis_leiocarpa
Euphorbiaceae_Mercurialis_canariensis
Euphorbiaceae_Mercurialis_annua
Euphorbiaceae_Mercurialis_corsica
Euphorbiaceae_Mercurialis_huetii
Euphorbiaceae_Mercurialis_elliptica
Euphorbiaceae_Claoxylon_indicum
Euphorbiaceae_Caryodendron_orinocense
Euphorbiaceae_Bernardia_incana
Euphorbiaceae_Bernardia_nicaraguensis
Euphorbiaceae_Bernardia_dodecandra
Euphorbiaceae_Bernardia_dichotoma
Euphorbiaceae_Bernardia_trelawniensis
Euphorbiaceae_Adenophaedra_grandifolia
Euphorbiaceae_Romanoa_tamnoides
Euphorbiaceae_Plukenetia_polyadenia
Euphorbiaceae_Plukenetia_stipellata
Euphorbiaceae_Cnesmone_javanica
Euphorbiaceae_Tragiella_anomala
Euphorbiaceae_Acidoton_urens
Euphorbiaceae_Tragia_volubilis
Euphorbiaceae_Plukenetia_volubilis
Euphorbiaceae_Tragia_betonicifolia
Euphorbiaceae_Tragia_urticifolia
Euphorbiaceae_Gitara_venezolana
Euphorbiaceae_Astrococcus_cornutus
Euphorbiaceae_Dalechampia_spathulata
Euphorbiaceae_Dalechampia_schottii
Euphorbiaceae_Dalechampia_magnoliifolia
Euphorbiaceae_Dalechampia_convolvuloides
Euphorbiaceae_Dalechampia_ficifolia
Euphorbiaceae_Dalechampia_sp
Euphorbiaceae_Dalechampia_pernambucensis
Euphorbiaceae_Dalechampia_brownsbergensis
Euphorbiaceae_Dalechampia_coriacea
Euphorbiaceae_Dalechampia_armbrusteri
Euphorbiaceae_Dalechampia_viridissima
Euphorbiaceae_Dalechampia_magnistipulata
Euphorbiaceae_Dalechampia_arenalensis
Euphorbiaceae_Dalechampia_schippii
Euphorbiaceae_Dalechampia_brevicolumna
Euphorbiaceae_Dalechampia_affinis
Euphorbiaceae_Dalechampia_tiliifolia
Euphorbiaceae_Dalechampia_capensis

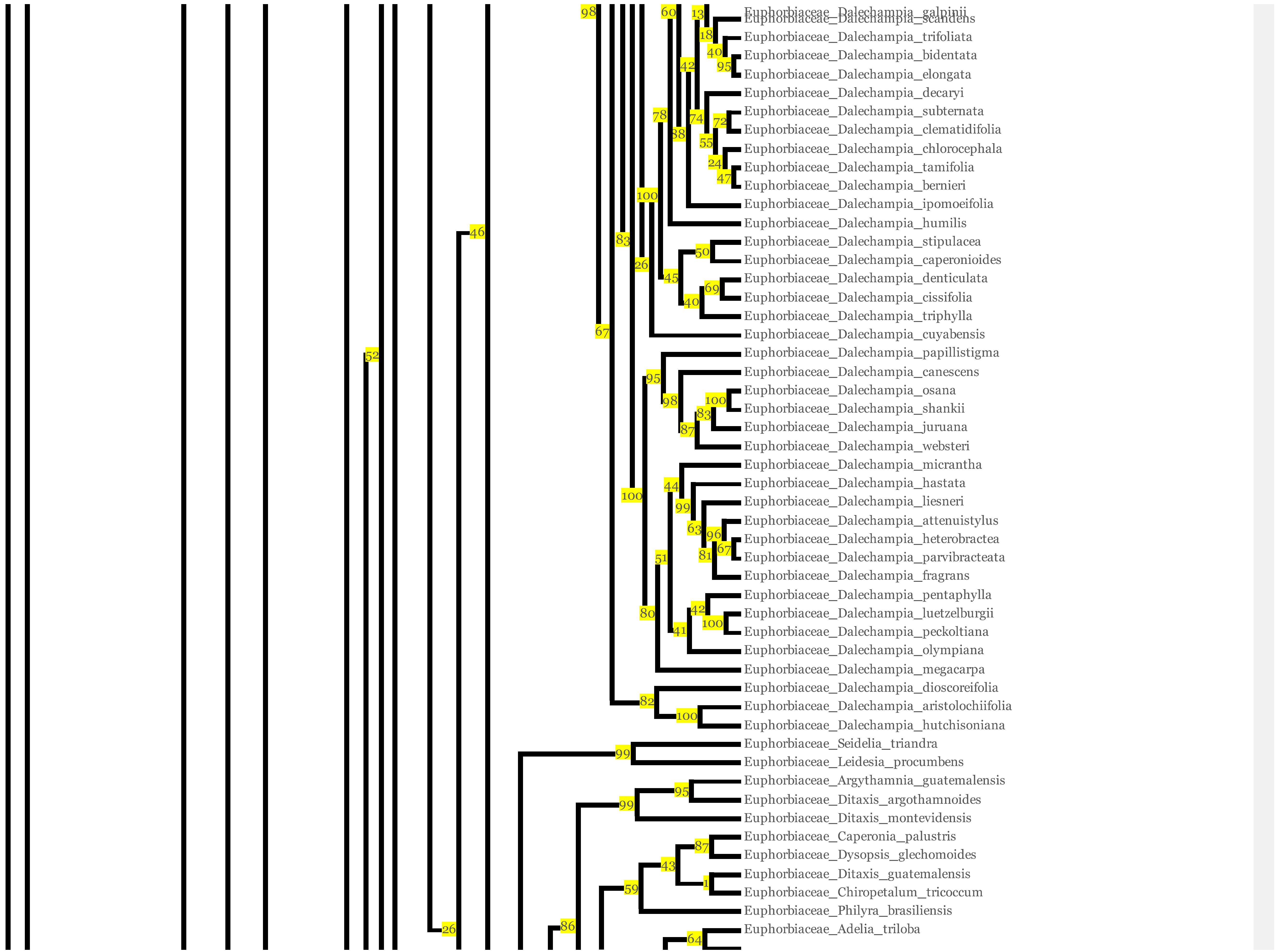

Euphorbiaceae_Dalechampia_galpinii
Euphorbiaceae_Dalechampia_scandens
Euphorbiaceae_Dalechampia_trifoliata
Euphorbiaceae_Dalechampia_bidentata
Euphorbiaceae_Dalechampia_elongata
Euphorbiaceae_Dalechampia_decaryi
Euphorbiaceae_Dalechampia_subternata
Euphorbiaceae_Dalechampia_clematidifolia
Euphorbiaceae_Dalechampia_chlorocephala
Euphorbiaceae_Dalechampia_tamifolia
Euphorbiaceae_Dalechampia_bernieri
Euphorbiaceae_Dalechampia_ipomoeifolia
Euphorbiaceae_Dalechampia_humilis
Euphorbiaceae_Dalechampia_stipulacea
Euphorbiaceae_Dalechampia_caperonioides
Euphorbiaceae_Dalechampia_denticulata
Euphorbiaceae_Dalechampia_cissifolia
Euphorbiaceae_Dalechampia_triphylla
Euphorbiaceae_Dalechampia_cuyabensis
Euphorbiaceae_Dalechampia_papillistigma
Euphorbiaceae_Dalechampia_canescens
Euphorbiaceae_Dalechampia_osana
Euphorbiaceae_Dalechampia_shankii
Euphorbiaceae_Dalechampia_juruana
Euphorbiaceae_Dalechampia_websteri
Euphorbiaceae_Dalechampia_micrantha
Euphorbiaceae_Dalechampia_hastata
Euphorbiaceae_Dalechampia_liesneri
Euphorbiaceae_Dalechampia_attenuistylus
Euphorbiaceae_Dalechampia_heterobractea
Euphorbiaceae_Dalechampia_parvibracteata
Euphorbiaceae_Dalechampia_fragrans
Euphorbiaceae_Dalechampia_pentaphylla
Euphorbiaceae_Dalechampia_luetzelburgii
Euphorbiaceae_Dalechampia_peckoltiana
Euphorbiaceae_Dalechampia_olympiana
Euphorbiaceae_Dalechampia_megacarpa
Euphorbiaceae_Dalechampia_dioscoreifolia
Euphorbiaceae_Dalechampia_aristolochiifolia
Euphorbiaceae_Dalechampia_hutchisoniana
Euphorbiaceae_Seidelia_triandra
Euphorbiaceae_Leidesia_procumbens
Euphorbiaceae_Argythamnia_guatemalensis
Euphorbiaceae_Ditaxis_argothamnoides
Euphorbiaceae_Ditaxis_montevidensis
Euphorbiaceae_Caperonia_palustris
Euphorbiaceae_Dysopsis_glechomoides
Euphorbiaceae_Ditaxis_guatemalensis
Euphorbiaceae_Chiropetalum_tricoccum
Euphorbiaceae_Philyra_brasiliensis
Euphorbiaceae_Adelia_triloba

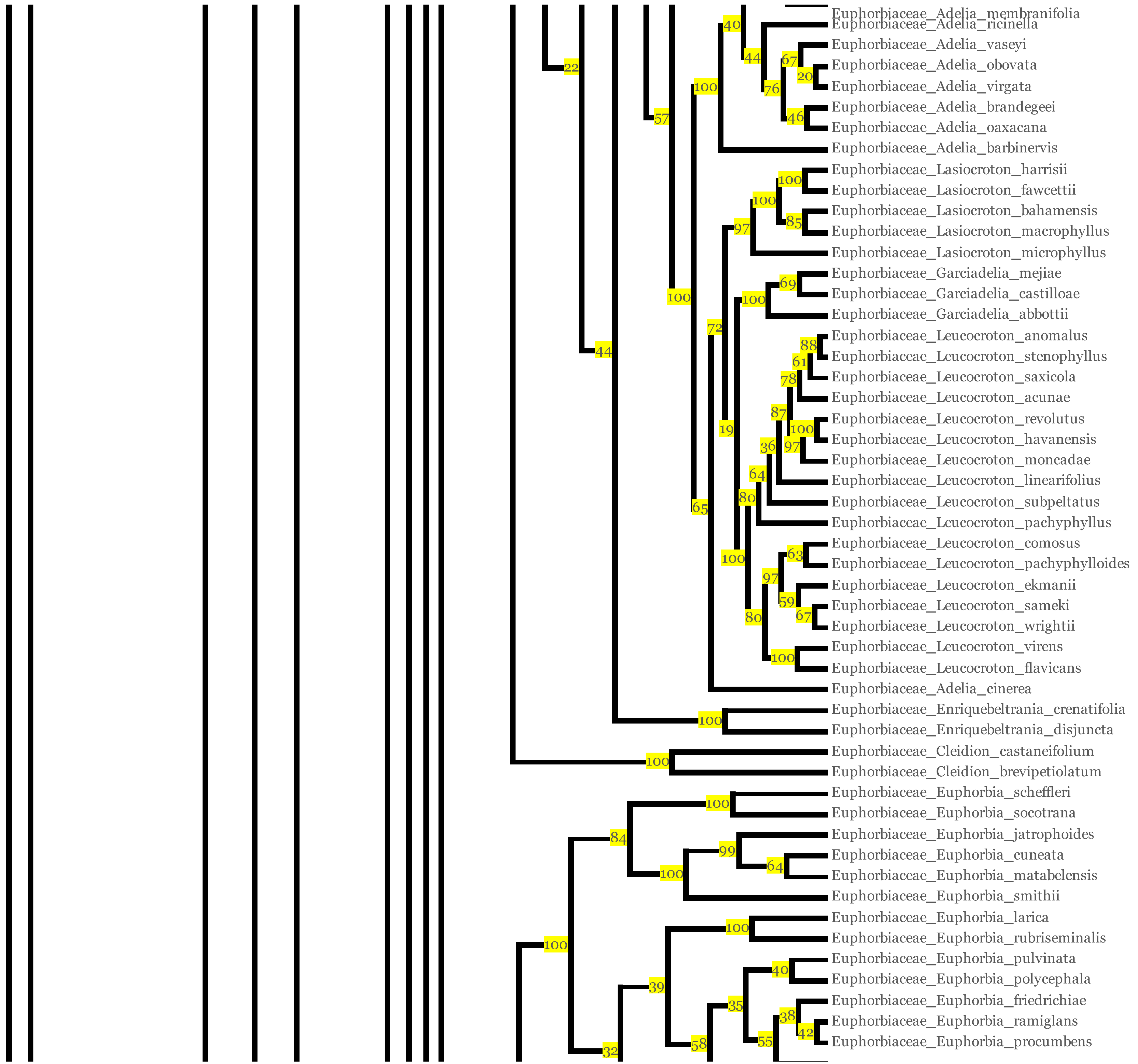
Euphorbiaceae_Adelia_membranifolia
Euphorbiaceae_Adelia_ricinella
Euphorbiaceae_Adelia_vaseyi
Euphorbiaceae_Adelia_obovata
Euphorbiaceae_Adelia_virgata
Euphorbiaceae_Adelia_brandegeei
Euphorbiaceae_Adelia_oaxacana
Euphorbiaceae_Adelia_barbinervis
Euphorbiaceae_Lasiocroton_harrisii
Euphorbiaceae_Lasiocroton_fawcettii
Euphorbiaceae_Lasiocroton_bahamensis
Euphorbiaceae_Lasiocroton_macrophyllus
Euphorbiaceae_Lasiocroton_microphyllus
Euphorbiaceae_Garciadelia_mejiae
Euphorbiaceae_Garciadelia_castilloae
Euphorbiaceae_Garciadelia_abbottii
Euphorbiaceae_Leucocroton_anomalus
Euphorbiaceae_Leucocroton_stenophyllus
Euphorbiaceae_Leucocroton_saxicola
Euphorbiaceae_Leucocroton_acunae
Euphorbiaceae_Leucocroton_revolutus
Euphorbiaceae_Leucocroton_havanensis
Euphorbiaceae_Leucocroton_moncadae
Euphorbiaceae_Leucocroton_linearifolius
Euphorbiaceae_Leucocroton_subpeltatus
Euphorbiaceae_Leucocroton_pachyphyllus
Euphorbiaceae_Leucocroton_comosus
Euphorbiaceae_Leucocroton_pachyphylloides
Euphorbiaceae_Leucocroton_ekmanii
Euphorbiaceae_Leucocroton_sameki
Euphorbiaceae_Leucocroton_wrightii
Euphorbiaceae_Leucocroton_virens
Euphorbiaceae_Leucocroton_flavicans
Euphorbiaceae_Adelia_cinerea
Euphorbiaceae_Enriquebeltrania_crenatifolia
Euphorbiaceae_Enriquebeltrania_disjuncta
Euphorbiaceae_Cleidion_castaneifolium
Euphorbiaceae_Cleidion_brevipetiolatum
Euphorbiaceae_Euphorbia_scheffleri
Euphorbiaceae_Euphorbia_socotrana
Euphorbiaceae_Euphorbia_jatrophoides
Euphorbiaceae_Euphorbia_cuneata
Euphorbiaceae_Euphorbia_matabelensis
Euphorbiaceae_Euphorbia_smithii
Euphorbiaceae_Euphorbia_larica
Euphorbiaceae_Euphorbia_rubriseminalis
Euphorbiaceae_Euphorbia_pulvinata
Euphorbiaceae_Euphorbia_polycephala
Euphorbiaceae_Euphorbia_friedrichiae
Euphorbiaceae_Euphorbia_ramiglans
Euphorbiaceae_Euphorbia_procumbens

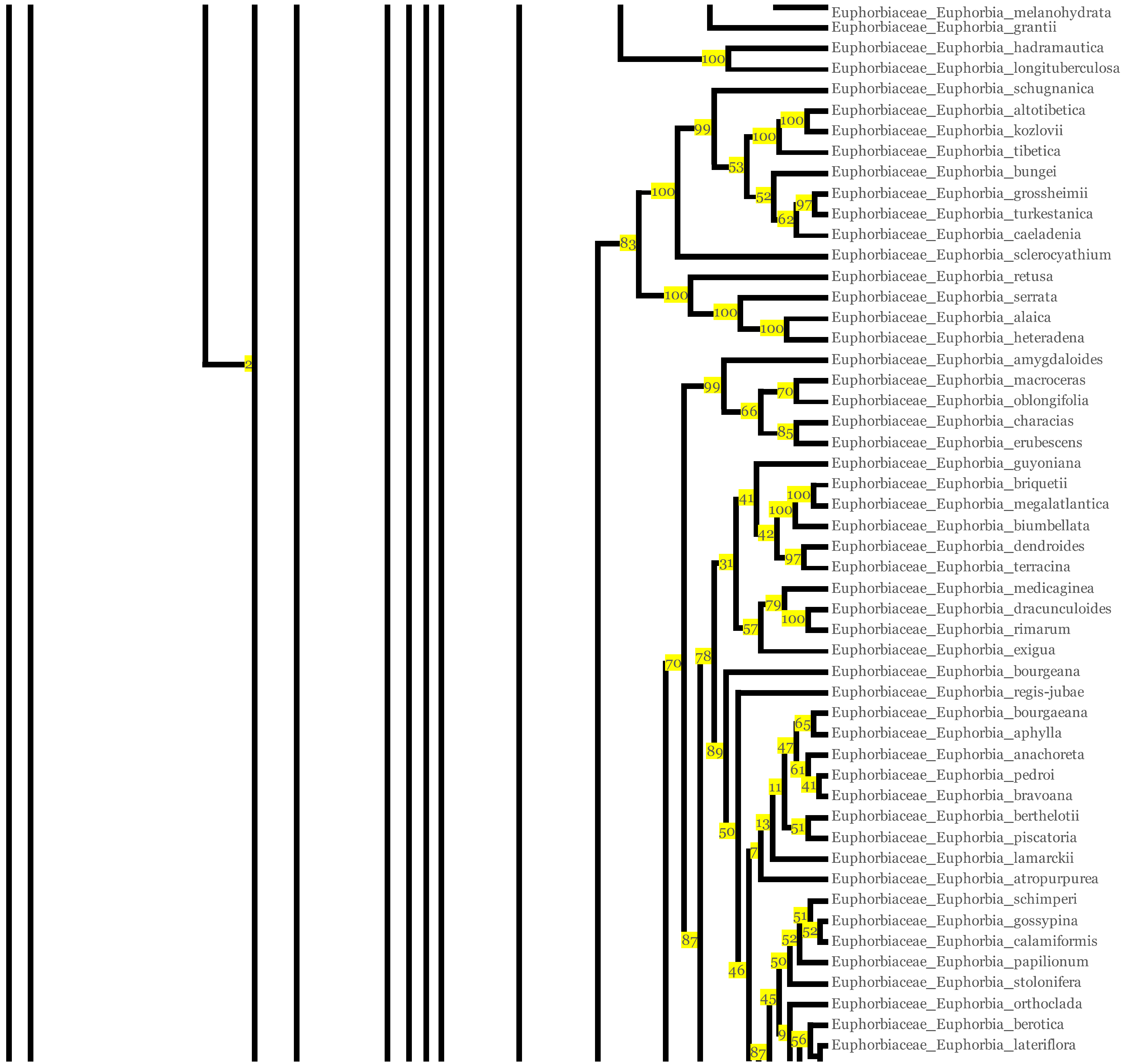
Euphorbiaceae_Euphorbia_melanohydrata
Euphorbiaceae_Euphorbia_grantii
Euphorbiaceae_Euphorbia_hadramautica
Euphorbiaceae_Euphorbia_longituberculosa
Euphorbiaceae_Euphorbia_schugnanica
Euphorbiaceae_Euphorbia_altotibetica
Euphorbiaceae_Euphorbia_kozlovii
Euphorbiaceae_Euphorbia_tibetica
Euphorbiaceae_Euphorbia_bungei
Euphorbiaceae_Euphorbia_grossheimii
Euphorbiaceae_Euphorbia_turkestanica
Euphorbiaceae_Euphorbia_caeladenia
Euphorbiaceae_Euphorbia_sclerocyathium
Euphorbiaceae_Euphorbia_retusa
Euphorbiaceae_Euphorbia_serrata
Euphorbiaceae_Euphorbia_alaica
Euphorbiaceae_Euphorbia_heteradena
Euphorbiaceae_Euphorbia_amygdaloides
Euphorbiaceae_Euphorbia_macroceras
Euphorbiaceae_Euphorbia_oblongifolia
Euphorbiaceae_Euphorbia_characias
Euphorbiaceae_Euphorbia_erubescens
Euphorbiaceae_Euphorbia_guyoniana
Euphorbiaceae_Euphorbia_briquetii
Euphorbiaceae_Euphorbia_megalatlantica
Euphorbiaceae_Euphorbia_biumbellata
Euphorbiaceae_Euphorbia_dendroides
Euphorbiaceae_Euphorbia_terracina
Euphorbiaceae_Euphorbia_medicaginea
Euphorbiaceae_Euphorbia_dracunculoides
Euphorbiaceae_Euphorbia_rimarum
Euphorbiaceae_Euphorbia_exigua
Euphorbiaceae_Euphorbia_bourgeana
Euphorbiaceae_Euphorbia_regis-jubae
Euphorbiaceae_Euphorbia_bourgaeana
Euphorbiaceae_Euphorbia_aphylla
Euphorbiaceae_Euphorbia_anachoreta
Euphorbiaceae_Euphorbia_pedroi
Euphorbiaceae_Euphorbia_bravoana
Euphorbiaceae_Euphorbia_berthelotii
Euphorbiaceae_Euphorbia_piscatoria
Euphorbiaceae_Euphorbia_lamarckii
Euphorbiaceae_Euphorbia_atropurpurea
Euphorbiaceae_Euphorbia_schimperi
Euphorbiaceae_Euphorbia_gossypina
Euphorbiaceae_Euphorbia_calamiformis
Euphorbiaceae_Euphorbia_papilionum
Euphorbiaceae_Euphorbia_stolonifera
Euphorbiaceae_Euphorbia_orthoclada
Euphorbiaceae_Euphorbia_berotica
Euphorbiaceae_Euphorbia_lateriflora

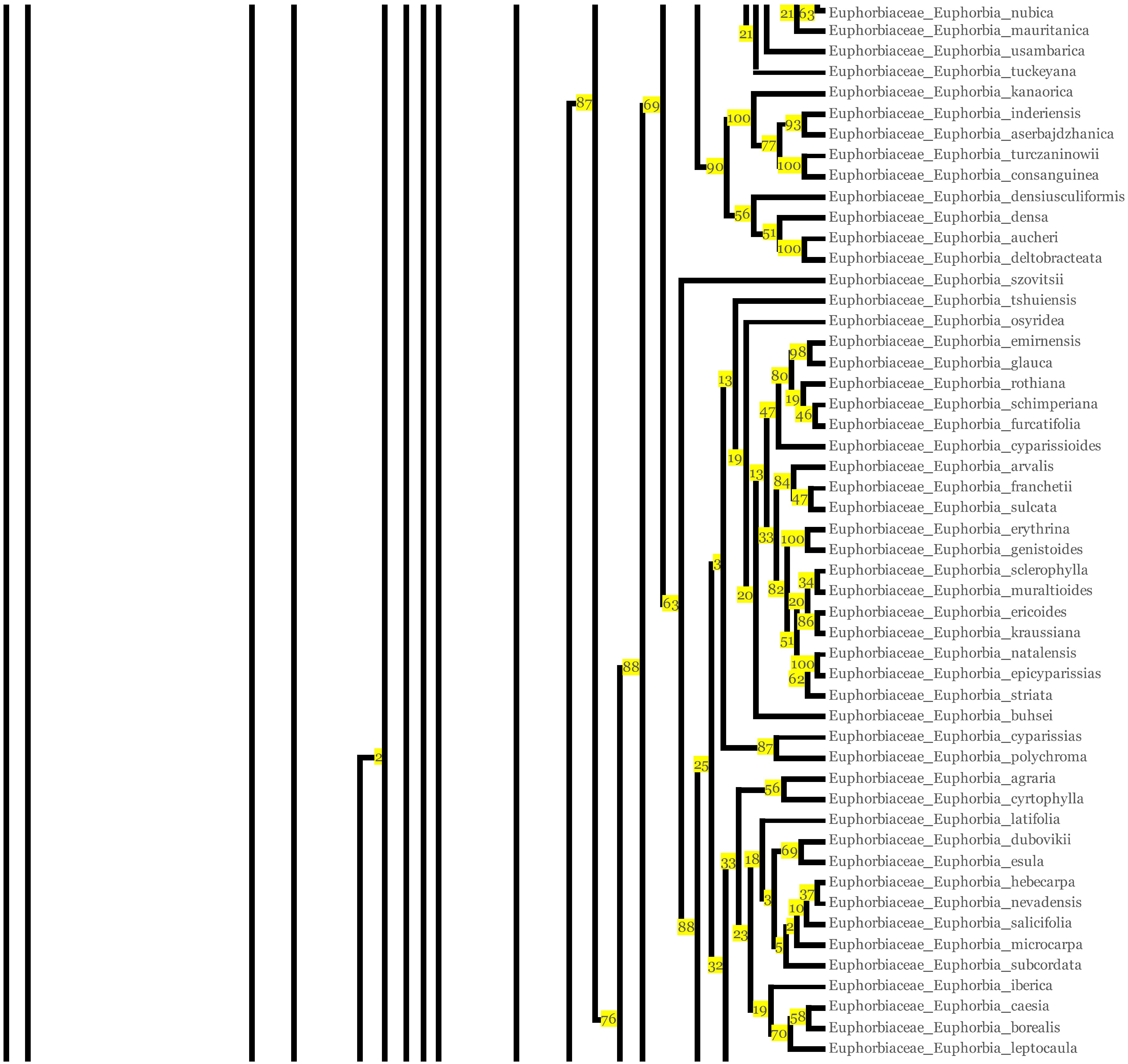
Euphorbiaceae_Euphorbia_nubica
Euphorbiaceae_Euphorbia_mauritanica
Euphorbiaceae_Euphorbia_usambarica
Euphorbiaceae_Euphorbia_tuckeyana
Euphorbiaceae_Euphorbia_kanaorica
Euphorbiaceae_Euphorbia_inderiensis
Euphorbiaceae_Euphorbia_aserbajdzhanica
Euphorbiaceae_Euphorbia_turczaninowii
Euphorbiaceae_Euphorbia_consanguinea
Euphorbiaceae_Euphorbia_densiusculiformis
Euphorbiaceae_Euphorbia_densa
Euphorbiaceae_Euphorbia_aucheri
Euphorbiaceae_Euphorbia_deltobracteata
Euphorbiaceae_Euphorbia_szovitsii
Euphorbiaceae_Euphorbia_tshuiensis
Euphorbiaceae_Euphorbia_osyridea
Euphorbiaceae_Euphorbia_emirnensis
Euphorbiaceae_Euphorbia_glauca
Euphorbiaceae_Euphorbia_rothiana
Euphorbiaceae_Euphorbia_schimperiana
Euphorbiaceae_Euphorbia_furcatifolia
Euphorbiaceae_Euphorbia_cyparissioides
Euphorbiaceae_Euphorbia_arvalis
Euphorbiaceae_Euphorbia_franchetii
Euphorbiaceae_Euphorbia_sulcata
Euphorbiaceae_Euphorbia_erythrina
Euphorbiaceae_Euphorbia_genistoides
Euphorbiaceae_Euphorbia_sclerophylla
Euphorbiaceae_Euphorbia_muraltioides
Euphorbiaceae_Euphorbia_ericoides
Euphorbiaceae_Euphorbia_kraussiana
Euphorbiaceae_Euphorbia_natalensis
Euphorbiaceae_Euphorbia_epicyparissias
Euphorbiaceae_Euphorbia_striata
Euphorbiaceae_Euphorbia_buhsei
Euphorbiaceae_Euphorbia_cyparissias
Euphorbiaceae_Euphorbia_polychroma
Euphorbiaceae_Euphorbia_agraria
Euphorbiaceae_Euphorbia_cyrtophylla
Euphorbiaceae_Euphorbia_latifolia
Euphorbiaceae_Euphorbia_dubovikii
Euphorbiaceae_Euphorbia_esula
Euphorbiaceae_Euphorbia_hebecarpa
Euphorbiaceae_Euphorbia_nevadensis
Euphorbiaceae_Euphorbia_salicifolia
Euphorbiaceae_Euphorbia_microcarpa
Euphorbiaceae_Euphorbia_subcordata
Euphorbiaceae_Euphorbia_iberica
Euphorbiaceae_Euphorbia_caesia
Euphorbiaceae_Euphorbia_borealis
Euphorbiaceae_Euphorbia_leptocaula

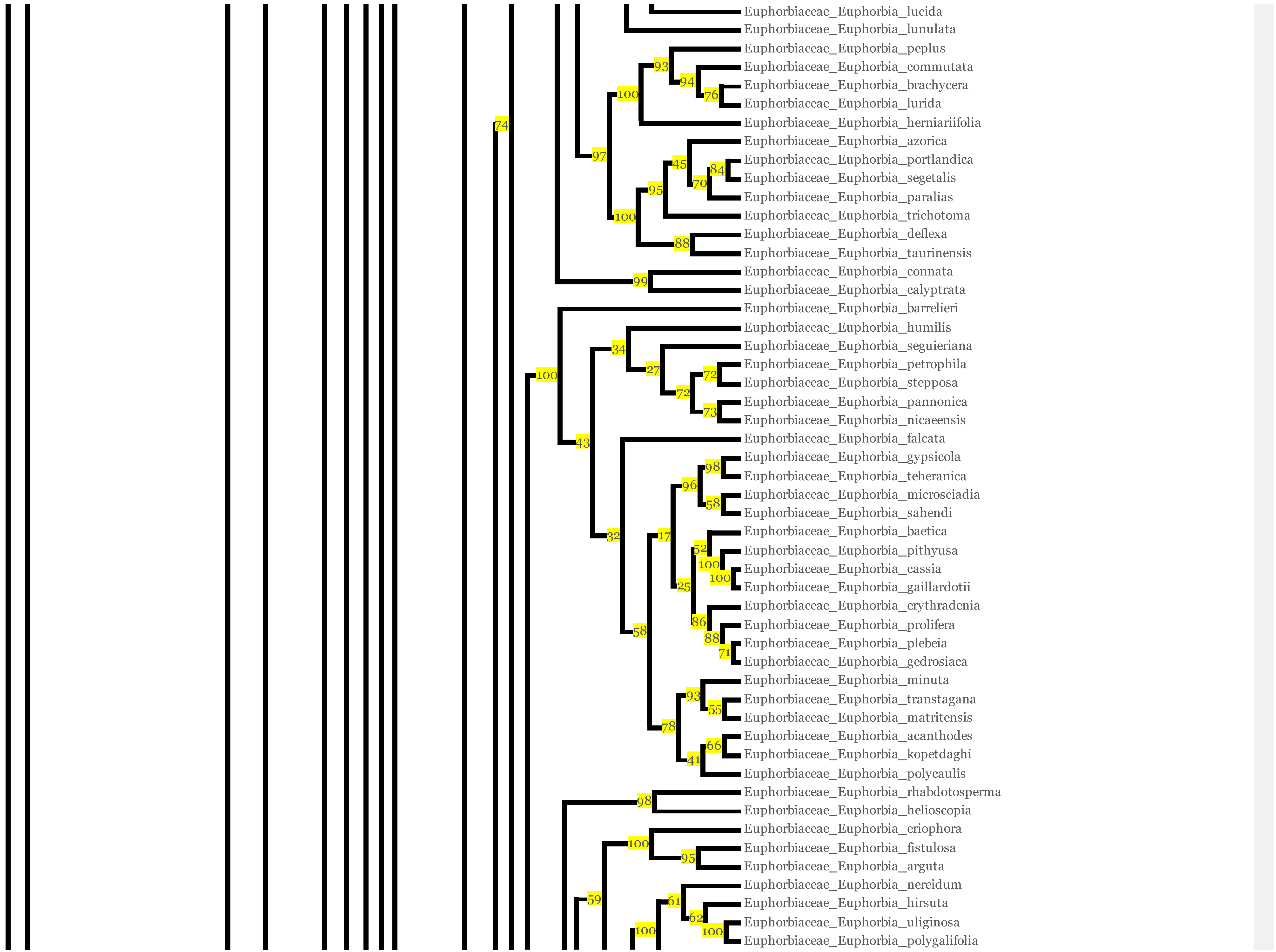
Euphorbiaceae_Euphorbia_lucida
Euphorbiaceae_Euphorbia_lunulata
Euphorbiaceae_Euphorbia_peplus
Euphorbiaceae_Euphorbia_commutata
Euphorbiaceae_Euphorbia_brachycera
Euphorbiaceae_Euphorbia_lurida
Euphorbiaceae_Euphorbia_herniariifolia
Euphorbiaceae_Euphorbia_azorica
Euphorbiaceae_Euphorbia_portlandica
Euphorbiaceae_Euphorbia_segetalis
Euphorbiaceae_Euphorbia_paralias
Euphorbiaceae_Euphorbia_trichotoma
Euphorbiaceae_Euphorbia_deflexa
Euphorbiaceae_Euphorbia_taurinensis
Euphorbiaceae_Euphorbia_connata
Euphorbiaceae_Euphorbia_calyptrata
Euphorbiaceae_Euphorbia_barrelieri
Euphorbiaceae_Euphorbia_humilis
Euphorbiaceae_Euphorbia_seguieriana
Euphorbiaceae_Euphorbia_petrophila
Euphorbiaceae_Euphorbia_stepposa
Euphorbiaceae_Euphorbia_pannonica
Euphorbiaceae_Euphorbia_nicaeensis
Euphorbiaceae_Euphorbia_falcata
Euphorbiaceae_Euphorbia_gypsicola
Euphorbiaceae_Euphorbia_teheranica
Euphorbiaceae_Euphorbia_microsciadia
Euphorbiaceae_Euphorbia_sahendi
Euphorbiaceae_Euphorbia_baetica
Euphorbiaceae_Euphorbia_pithyusa
Euphorbiaceae_Euphorbia_cassia
Euphorbiaceae_Euphorbia_gaillardotii
Euphorbiaceae_Euphorbia_erythradenia
Euphorbiaceae_Euphorbia_prolifera
Euphorbiaceae_Euphorbia_plebeia
Euphorbiaceae_Euphorbia_gedrosiaca
Euphorbiaceae_Euphorbia_minuta
Euphorbiaceae_Euphorbia_transtagana
Euphorbiaceae_Euphorbia_matritensis
Euphorbiaceae_Euphorbia_acanthodes
Euphorbiaceae_Euphorbia_kopetdaghi
Euphorbiaceae_Euphorbia_polycaulis
Euphorbiaceae_Euphorbia_rhabdotosperma
Euphorbiaceae_Euphorbia_helioscopia
Euphorbiaceae_Euphorbia_eriophora
Euphorbiaceae_Euphorbia_fistulosa
Euphorbiaceae_Euphorbia_arguta
Euphorbiaceae_Euphorbia_nereidum
Euphorbiaceae_Euphorbia_hirsuta
Euphorbiaceae_Euphorbia_uliginosa
Euphorbiaceae_Euphorbia_polygalifolia

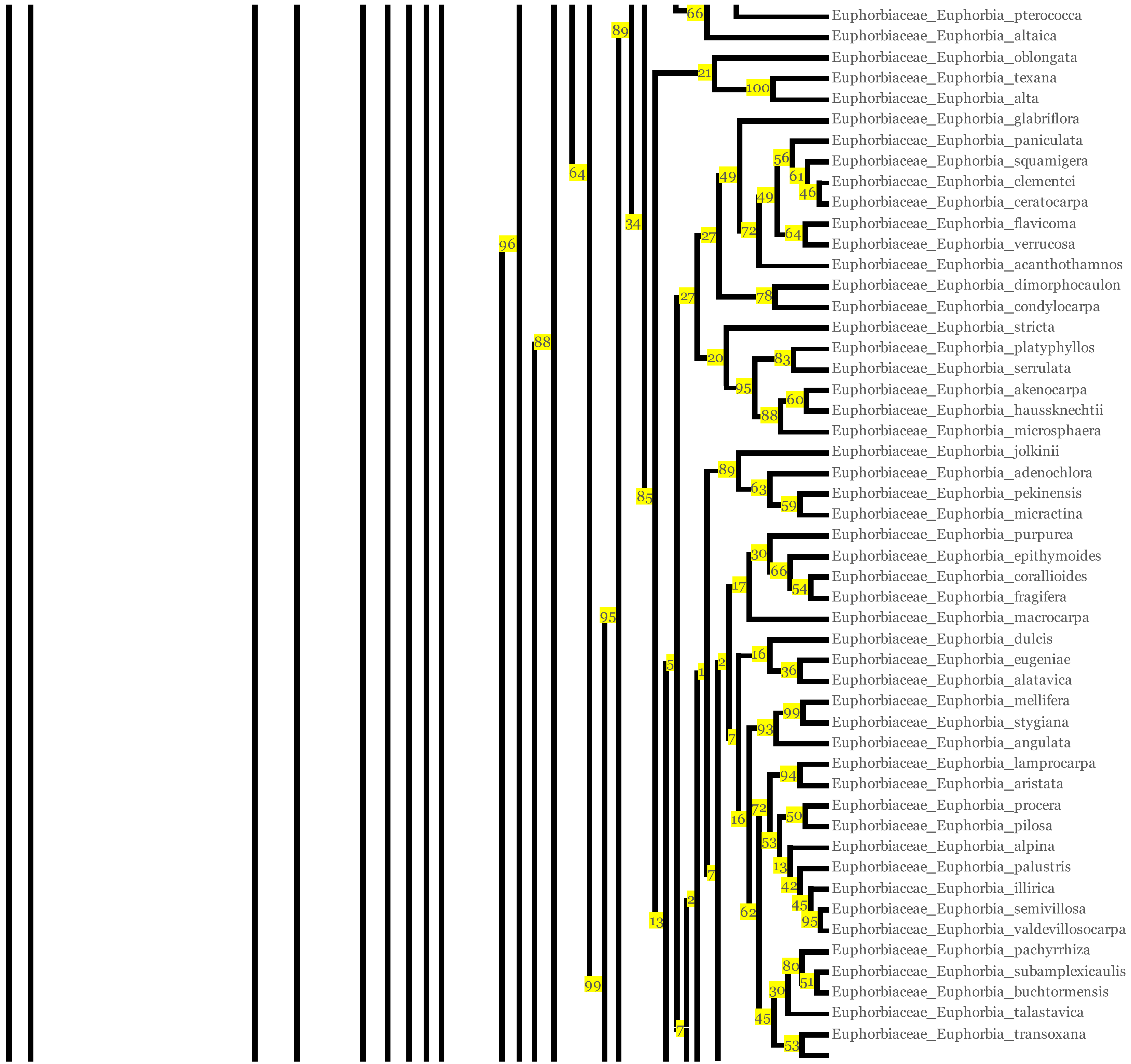
Euphorbiaceae_Euphorbia_pterococca
Euphorbiaceae_Euphorbia_altaica
Euphorbiaceae_Euphorbia_oblongata
Euphorbiaceae_Euphorbia_texana
Euphorbiaceae_Euphorbia_alta
Euphorbiaceae_Euphorbia_glabriflora
Euphorbiaceae_Euphorbia_paniculata
Euphorbiaceae_Euphorbia_squamigera
Euphorbiaceae_Euphorbia_clementei
Euphorbiaceae_Euphorbia_ceratocarpa
Euphorbiaceae_Euphorbia_flavicoma
Euphorbiaceae_Euphorbia_verrucosa
Euphorbiaceae_Euphorbia_acanthothamnos
Euphorbiaceae_Euphorbia_dimorphocaulon
Euphorbiaceae_Euphorbia_condylocarpa
Euphorbiaceae_Euphorbia_stricta
Euphorbiaceae_Euphorbia_platyphyllos
Euphorbiaceae_Euphorbia_serrulata
Euphorbiaceae_Euphorbia_akenocarpa
Euphorbiaceae_Euphorbia_haussknechtii
Euphorbiaceae_Euphorbia_microsphaera
Euphorbiaceae_Euphorbia_jolkinii
Euphorbiaceae_Euphorbia_adenochlora
Euphorbiaceae_Euphorbia_pekinensis
Euphorbiaceae_Euphorbia_micractina
Euphorbiaceae_Euphorbia_purpurea
Euphorbiaceae_Euphorbia_epithymoides
Euphorbiaceae_Euphorbia_corallioides
Euphorbiaceae_Euphorbia_fragifera
Euphorbiaceae_Euphorbia_macrocarpa
Euphorbiaceae_Euphorbia_dulcis
Euphorbiaceae_Euphorbia_eugeniae
Euphorbiaceae_Euphorbia_alatavica
Euphorbiaceae_Euphorbia_mellifera
Euphorbiaceae_Euphorbia_stygiana
Euphorbiaceae_Euphorbia_angulata
Euphorbiaceae_Euphorbia_lamprocarpa
Euphorbiaceae_Euphorbia_aristata
Euphorbiaceae_Euphorbia_procera
Euphorbiaceae_Euphorbia_pilosa
Euphorbiaceae_Euphorbia_alpina
Euphorbiaceae_Euphorbia_palustris
Euphorbiaceae_Euphorbia_illirica
Euphorbiaceae_Euphorbia_semivillosa
Euphorbiaceae_Euphorbia_valdevillosocarpa
Euphorbiaceae_Euphorbia_pachyrrhiza
Euphorbiaceae_Euphorbia_subamplexicaulis
Euphorbiaceae_Euphorbia_buchtormensis
Euphorbiaceae_Euphorbia_talastavica
Euphorbiaceae_Euphorbia_transoxana

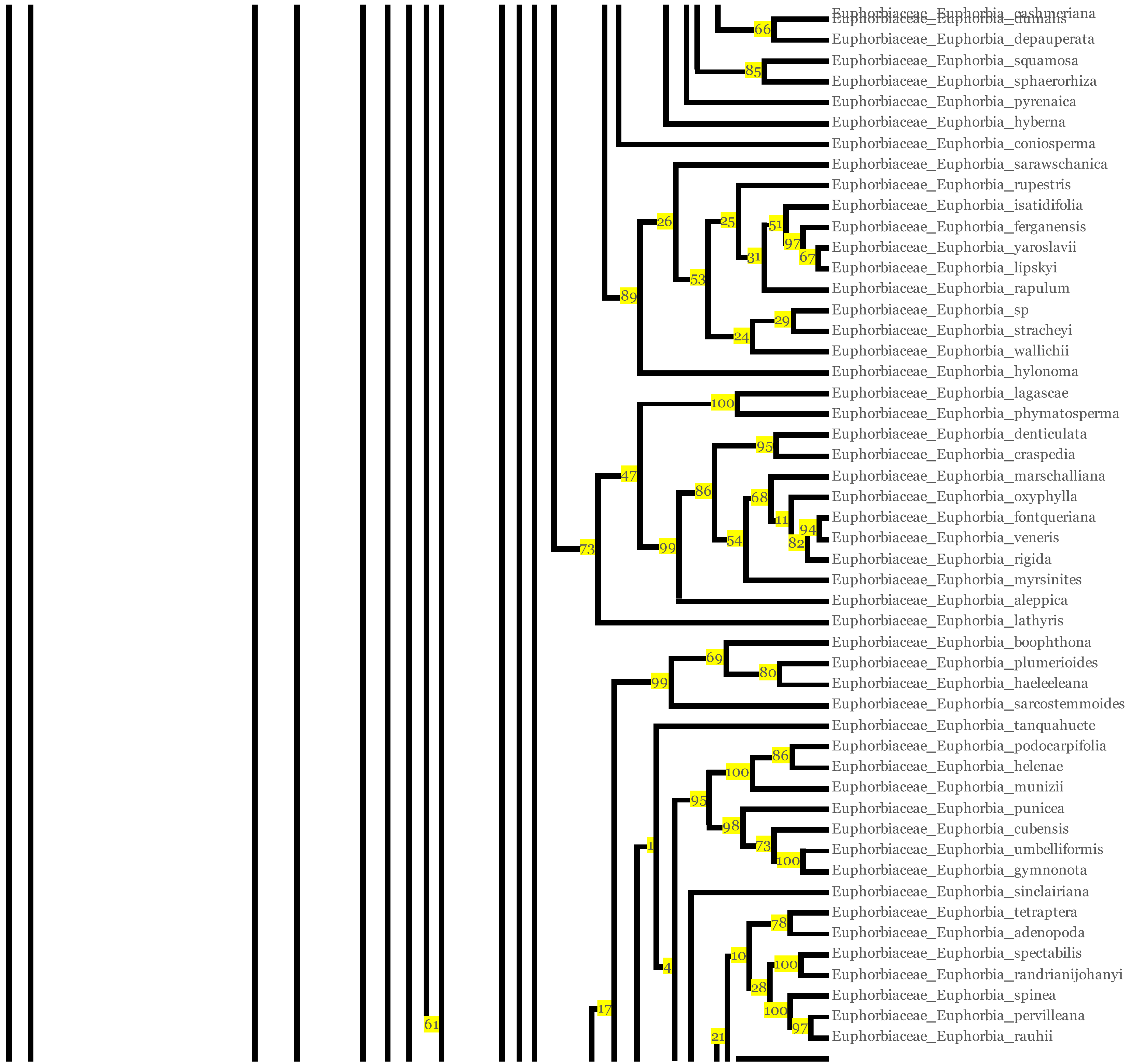
Euphorbiaceae_Euphorbia_cashmeriana
Euphorbiaceae_Euphorbia_depauperata
Euphorbiaceae_Euphorbia_squamosa
Euphorbiaceae_Euphorbia_sphaerorhiza
Euphorbiaceae_Euphorbia_pyrenaica
Euphorbiaceae_Euphorbia_hyberna
Euphorbiaceae_Euphorbia_coniosperma
Euphorbiaceae_Euphorbia_sarawschanica
Euphorbiaceae_Euphorbia_rupestris
Euphorbiaceae_Euphorbia_isatidifolia
Euphorbiaceae_Euphorbia_ferganensis
Euphorbiaceae_Euphorbia_yaroslavii
Euphorbiaceae_Euphorbia_lipskyi
Euphorbiaceae_Euphorbia_rapulum
Euphorbiaceae_Euphorbia_sp
Euphorbiaceae_Euphorbia_stracheyi
Euphorbiaceae_Euphorbia_wallichii
Euphorbiaceae_Euphorbia_hylonoma
Euphorbiaceae_Euphorbia_lagascae
Euphorbiaceae_Euphorbia_phymatosperma
Euphorbiaceae_Euphorbia_denticulata
Euphorbiaceae_Euphorbia_craspedia
Euphorbiaceae_Euphorbia_marschalliana
Euphorbiaceae_Euphorbia_oxyphylla
Euphorbiaceae_Euphorbia_fontqueriana
Euphorbiaceae_Euphorbia_veneris
Euphorbiaceae_Euphorbia_rigida
Euphorbiaceae_Euphorbia_myrsinites
Euphorbiaceae_Euphorbia_aleppica
Euphorbiaceae_Euphorbia_lathyris
Euphorbiaceae_Euphorbia_boophthona
Euphorbiaceae_Euphorbia_plumerioides
Euphorbiaceae_Euphorbia_haeleeleana
Euphorbiaceae_Euphorbia_sarcostemmoides
Euphorbiaceae_Euphorbia_tanquahuete
Euphorbiaceae_Euphorbia_podocarpifolia
Euphorbiaceae_Euphorbia_helenae
Euphorbiaceae_Euphorbia_munizii
Euphorbiaceae_Euphorbia_punicea
Euphorbiaceae_Euphorbia_cubensis
Euphorbiaceae_Euphorbia_umbelliformis
Euphorbiaceae_Euphorbia_gymnonota
Euphorbiaceae_Euphorbia_sinclairiana
Euphorbiaceae_Euphorbia_tetraptera
Euphorbiaceae_Euphorbia_adenopoda
Euphorbiaceae_Euphorbia_spectabilis
Euphorbiaceae_Euphorbia_randrianjohanyi
Euphorbiaceae_Euphorbia_spinea
Euphorbiaceae_Euphorbia_pervilleana
Euphorbiaceae_Euphorbia_rauhii

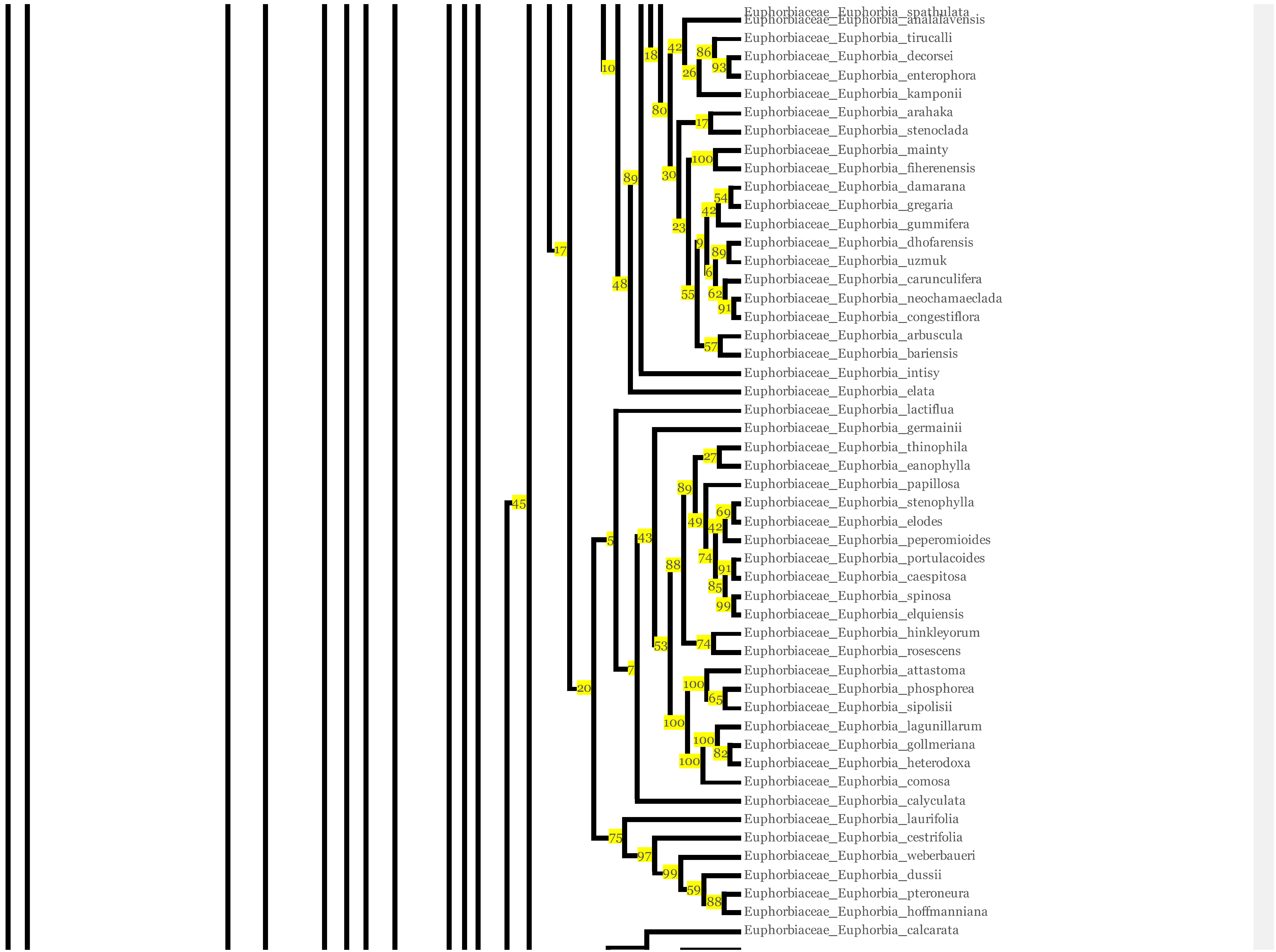
Euphorbiaceae_Euphorbia_spathulata
Euphorbiaceae_Euphorbia_analalavensis
Euphorbiaceae_Euphorbia_tirucalli
Euphorbiaceae_Euphorbia_decorsei
Euphorbiaceae_Euphorbia_enterophora
Euphorbiaceae_Euphorbia_kamponii
Euphorbiaceae_Euphorbia_arahaka
Euphorbiaceae_Euphorbia_stenoclada
Euphorbiaceae_Euphorbia_mainty
Euphorbiaceae_Euphorbia_fiherenensis
Euphorbiaceae_Euphorbia_damarana
Euphorbiaceae_Euphorbia_gregaria
Euphorbiaceae_Euphorbia_gummifera
Euphorbiaceae_Euphorbia_dhofarensis
Euphorbiaceae_Euphorbia_uzmuk
Euphorbiaceae_Euphorbia_carunculifera
Euphorbiaceae_Euphorbia_neochamaeclada
Euphorbiaceae_Euphorbia_congestiflora
Euphorbiaceae_Euphorbia_arbuscula
Euphorbiaceae_Euphorbia_bariensis
Euphorbiaceae_Euphorbia_intisy
Euphorbiaceae_Euphorbia_elata
Euphorbiaceae_Euphorbia_lactiflua
Euphorbiaceae_Euphorbia_germainii
Euphorbiaceae_Euphorbia_thinophila
Euphorbiaceae_Euphorbia_eanophylla
Euphorbiaceae_Euphorbia_papillosa
Euphorbiaceae_Euphorbia_stenophylla
Euphorbiaceae_Euphorbia_elodes
Euphorbiaceae_Euphorbia_peperomioides
Euphorbiaceae_Euphorbia_portulacoides
Euphorbiaceae_Euphorbia_caespitosa
Euphorbiaceae_Euphorbia_spinosa
Euphorbiaceae_Euphorbia_elquiensis
Euphorbiaceae_Euphorbia_hinkleyorum
Euphorbiaceae_Euphorbia_rosescens
Euphorbiaceae_Euphorbia_attastoma
Euphorbiaceae_Euphorbia_phosphorea
Euphorbiaceae_Euphorbia_sipolisii
Euphorbiaceae_Euphorbia_lagunillarum
Euphorbiaceae_Euphorbia_gollmeriana
Euphorbiaceae_Euphorbia_heterodoxa
Euphorbiaceae_Euphorbia_comosa
Euphorbiaceae_Euphorbia_calyculata
Euphorbiaceae_Euphorbia_laurifolia
Euphorbiaceae_Euphorbia_cestrifolia
Euphorbiaceae_Euphorbia_weberbaueri
Euphorbiaceae_Euphorbia_dussii
Euphorbiaceae_Euphorbia_pteroneura
Euphorbiaceae_Euphorbia_hoffmanniana
Euphorbiaceae_Euphorbia_calcarata

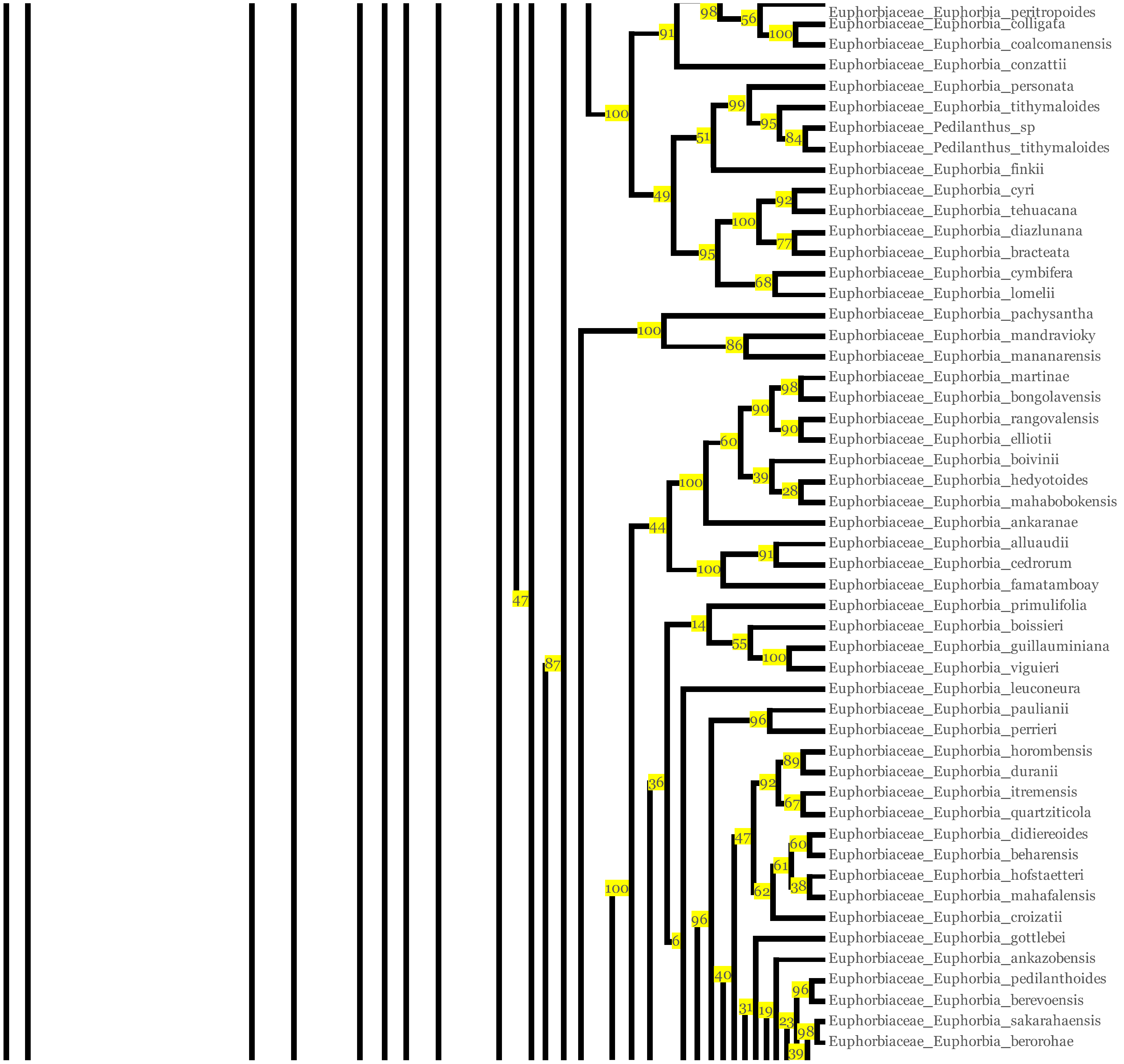
Euphorbiaceae_Euphorbia_peritropoides
Euphorbiaceae_Euphorbia_colligata
Euphorbiaceae_Euphorbia_coalcomanensis
Euphorbiaceae_Euphorbia_conzattii
Euphorbiaceae_Euphorbia_personata
Euphorbiaceae_Euphorbia_tithymaloides
Euphorbiaceae_Pedilanthus_sp
Euphorbiaceae_Pedilanthus_tithymaloides
Euphorbiaceae_Euphorbia_finkii
Euphorbiaceae_Euphorbia_cyri
Euphorbiaceae_Euphorbia_tehuacana
Euphorbiaceae_Euphorbia_diazlunana
Euphorbiaceae_Euphorbia_bracteata
Euphorbiaceae_Euphorbia_cymbifera
Euphorbiaceae_Euphorbia_lomelii
Euphorbiaceae_Euphorbia_pachysantha
Euphorbiaceae_Euphorbia_mandravioky
Euphorbiaceae_Euphorbia_mananarensis
Euphorbiaceae_Euphorbia_martinae
Euphorbiaceae_Euphorbia_bongolavensis
Euphorbiaceae_Euphorbia_rangovalensis
Euphorbiaceae_Euphorbia_elliotii
Euphorbiaceae_Euphorbia_boivinii
Euphorbiaceae_Euphorbia_hedyotoides
Euphorbiaceae_Euphorbia_mahabobokensis
Euphorbiaceae_Euphorbia_ankaranae
Euphorbiaceae_Euphorbia_alluaudii
Euphorbiaceae_Euphorbia_cedrorum
Euphorbiaceae_Euphorbia_famatamboay
Euphorbiaceae_Euphorbia_primulifolia
Euphorbiaceae_Euphorbia_boissieri
Euphorbiaceae_Euphorbia_guillauminiana
Euphorbiaceae_Euphorbia_viguieri
Euphorbiaceae_Euphorbia_leuconeura
Euphorbiaceae_Euphorbia_paulianii
Euphorbiaceae_Euphorbia_perrieri
Euphorbiaceae_Euphorbia_horombensis
Euphorbiaceae_Euphorbia_duranii
Euphorbiaceae_Euphorbia_itremensis
Euphorbiaceae_Euphorbia_quartziticola
Euphorbiaceae_Euphorbia_didiereoides
Euphorbiaceae_Euphorbia_beharensis
Euphorbiaceae_Euphorbia_hofstaetteri
Euphorbiaceae_Euphorbia_mahafalensis
Euphorbiaceae_Euphorbia_croizatii
Euphorbiaceae_Euphorbia_gottlebei
Euphorbiaceae_Euphorbia_ankazobensis
Euphorbiaceae_Euphorbia_pedilanthoides
Euphorbiaceae_Euphorbia_berevoensis
Euphorbiaceae_Euphorbia_sakarahaensis
Euphorbiaceae_Euphorbia_berorohae

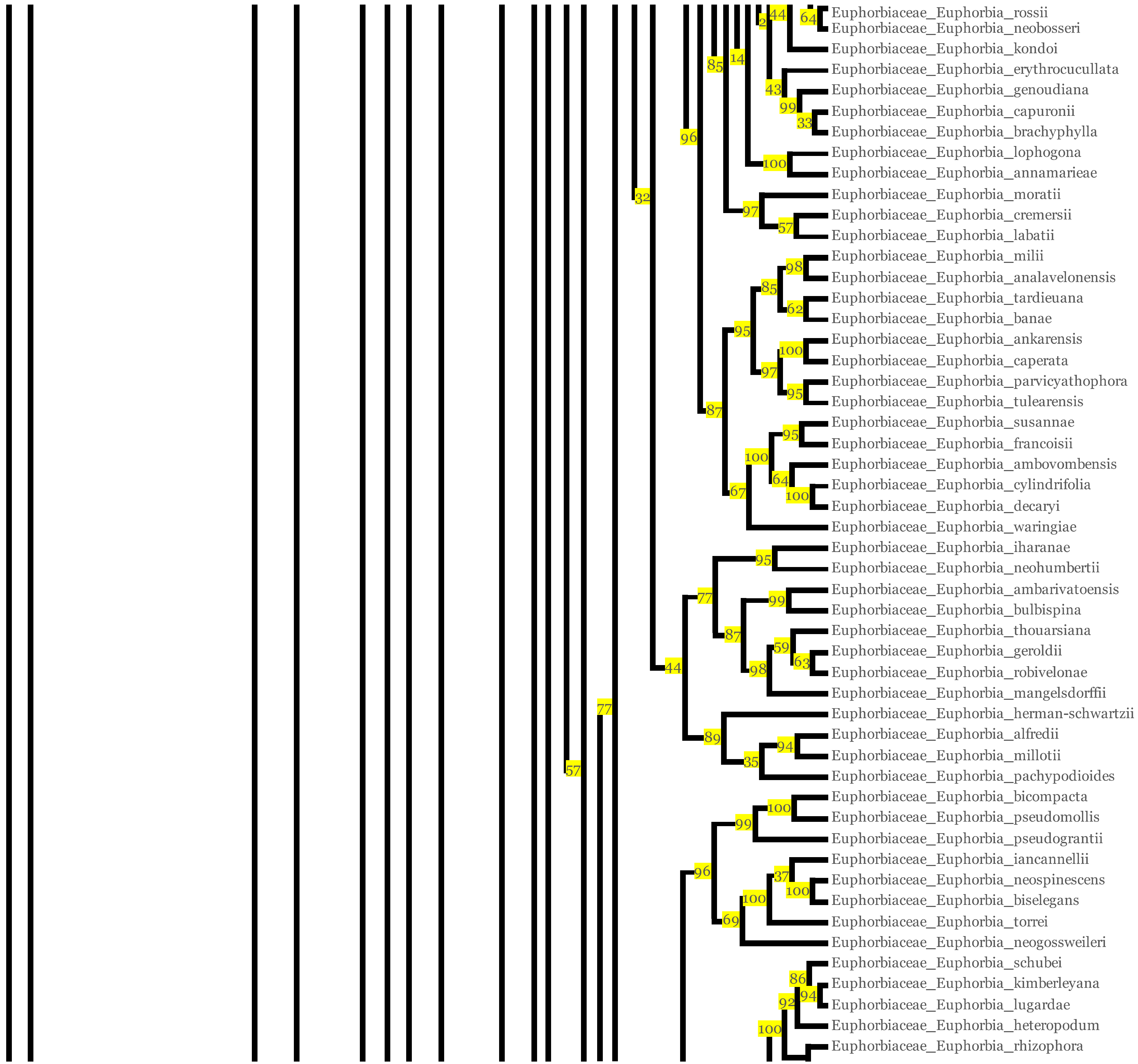
Euphorbiaceae_Euphorbia_rossii
Euphorbiaceae_Euphorbia_neobosseri
Euphorbiaceae_Euphorbia_kondoi
Euphorbiaceae_Euphorbia_erythrocucullata
Euphorbiaceae_Euphorbia_genoudiana
Euphorbiaceae_Euphorbia_capuronii
Euphorbiaceae_Euphorbia_brachyphylla
Euphorbiaceae_Euphorbia_lophogona
Euphorbiaceae_Euphorbia_annamarieae
Euphorbiaceae_Euphorbia_moratii
Euphorbiaceae_Euphorbia_cremersii
Euphorbiaceae_Euphorbia_labatii
Euphorbiaceae_Euphorbia_milii
Euphorbiaceae_Euphorbia_analavelonensis
Euphorbiaceae_Euphorbia_tardieuana
Euphorbiaceae_Euphorbia_banae
Euphorbiaceae_Euphorbia_ankarensis
Euphorbiaceae_Euphorbia_caperata
Euphorbiaceae_Euphorbia_parvicyathophora
Euphorbiaceae_Euphorbia_tulearensis
Euphorbiaceae_Euphorbia_susannae
Euphorbiaceae_Euphorbia_francoisii
Euphorbiaceae_Euphorbia_ambovombensis
Euphorbiaceae_Euphorbia_cylindrifolia
Euphorbiaceae_Euphorbia_decaryi
Euphorbiaceae_Euphorbia_waringiae
Euphorbiaceae_Euphorbia_iharanae
Euphorbiaceae_Euphorbia_neohumbertii
Euphorbiaceae_Euphorbia_ambarivatoensis
Euphorbiaceae_Euphorbia_bulbispina
Euphorbiaceae_Euphorbia_thouarsiana
Euphorbiaceae_Euphorbia_geroldii
Euphorbiaceae_Euphorbia_robivelonae
Euphorbiaceae_Euphorbia_mangelsdorffii
Euphorbiaceae_Euphorbia_herman-schwartzii
Euphorbiaceae_Euphorbia_alfredii
Euphorbiaceae_Euphorbia_millotii
Euphorbiaceae_Euphorbia_pachypodioides
Euphorbiaceae_Euphorbia_bicompacta
Euphorbiaceae_Euphorbia_pseudomollis
Euphorbiaceae_Euphorbia_pseudograntii
Euphorbiaceae_Euphorbia_iancannellii
Euphorbiaceae_Euphorbia_neospinescens
Euphorbiaceae_Euphorbia_biselegans
Euphorbiaceae_Euphorbia_torrei
Euphorbiaceae_Euphorbia_neogossweileri
Euphorbiaceae_Euphorbia_schubei
Euphorbiaceae_Euphorbia_kimberleyana
Euphorbiaceae_Euphorbia_lugardae
Euphorbiaceae_Euphorbia_heteropodum
Euphorbiaceae_Euphorbia_rhizophora

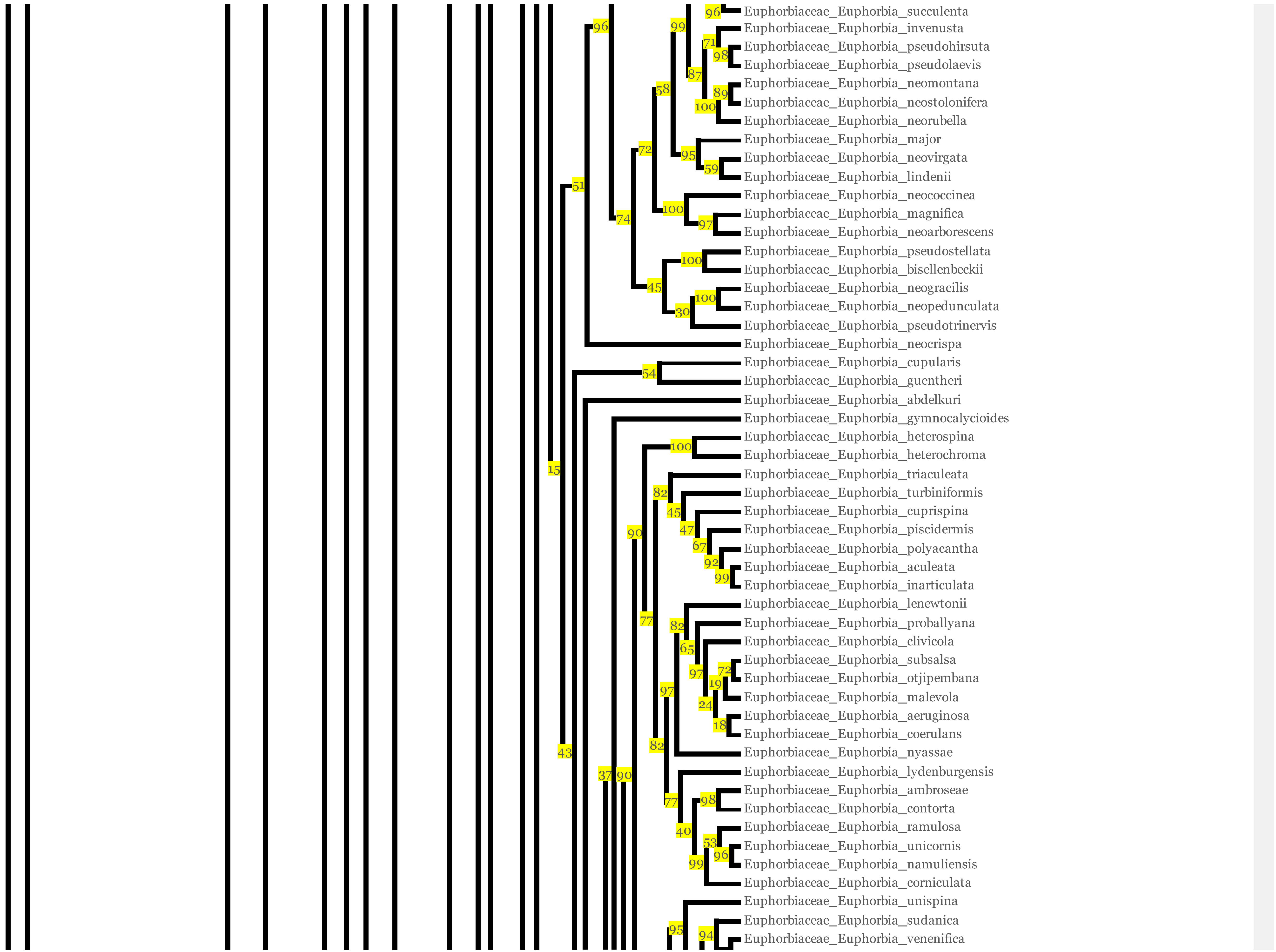
Euphorbiaceae_Euphorbia_succulenta
Euphorbiaceae_Euphorbia_invenusta
Euphorbiaceae_Euphorbia_pseudohirsuta
Euphorbiaceae_Euphorbia_pseudolaevis
Euphorbiaceae_Euphorbia_neomontana
Euphorbiaceae_Euphorbia_neostolonifera
Euphorbiaceae_Euphorbia_neorubella
Euphorbiaceae_Euphorbia_major
Euphorbiaceae_Euphorbia_neovirgata
Euphorbiaceae_Euphorbia_lindenii
Euphorbiaceae_Euphorbia_neococcinea
Euphorbiaceae_Euphorbia_magnifica
Euphorbiaceae_Euphorbia_neoarborescens
Euphorbiaceae_Euphorbia_pseudostellata
Euphorbiaceae_Euphorbia_bisellenbeckii
Euphorbiaceae_Euphorbia_neogracilis
Euphorbiaceae_Euphorbia_neopedunculata
Euphorbiaceae_Euphorbia_pseudotrinervis
Euphorbiaceae_Euphorbia_neocrispa
Euphorbiaceae_Euphorbia_cupularis
Euphorbiaceae_Euphorbia_guentheri
Euphorbiaceae_Euphorbia_abdelkuri
Euphorbiaceae_Euphorbia_gymnocalycioides
Euphorbiaceae_Euphorbia_heterospina
Euphorbiaceae_Euphorbia_heterochroma
Euphorbiaceae_Euphorbia_triaculeata
Euphorbiaceae_Euphorbia_turbiniformis
Euphorbiaceae_Euphorbia_cuprispina
Euphorbiaceae_Euphorbia_piscidermis
Euphorbiaceae_Euphorbia_polyacantha
Euphorbiaceae_Euphorbia_aculeata
Euphorbiaceae_Euphorbia_inarticulata
Euphorbiaceae_Euphorbia_lenewtonii
Euphorbiaceae_Euphorbia_proballyana
Euphorbiaceae_Euphorbia_clivicola
Euphorbiaceae_Euphorbia_subsalsa
Euphorbiaceae_Euphorbia_otjipembana
Euphorbiaceae_Euphorbia_malevola
Euphorbiaceae_Euphorbia_aeruginosa
Euphorbiaceae_Euphorbia_coerulans
Euphorbiaceae_Euphorbia_nyassae
Euphorbiaceae_Euphorbia_lydenburgensis
Euphorbiaceae_Euphorbia_ambroseae
Euphorbiaceae_Euphorbia_contorta
Euphorbiaceae_Euphorbia_ramulosa
Euphorbiaceae_Euphorbia_unicornis
Euphorbiaceae_Euphorbia_namuliensis
Euphorbiaceae_Euphorbia_corniculata
Euphorbiaceae_Euphorbia_unispina
Euphorbiaceae_Euphorbia_sudanica
Euphorbiaceae_Euphorbia_venenifica

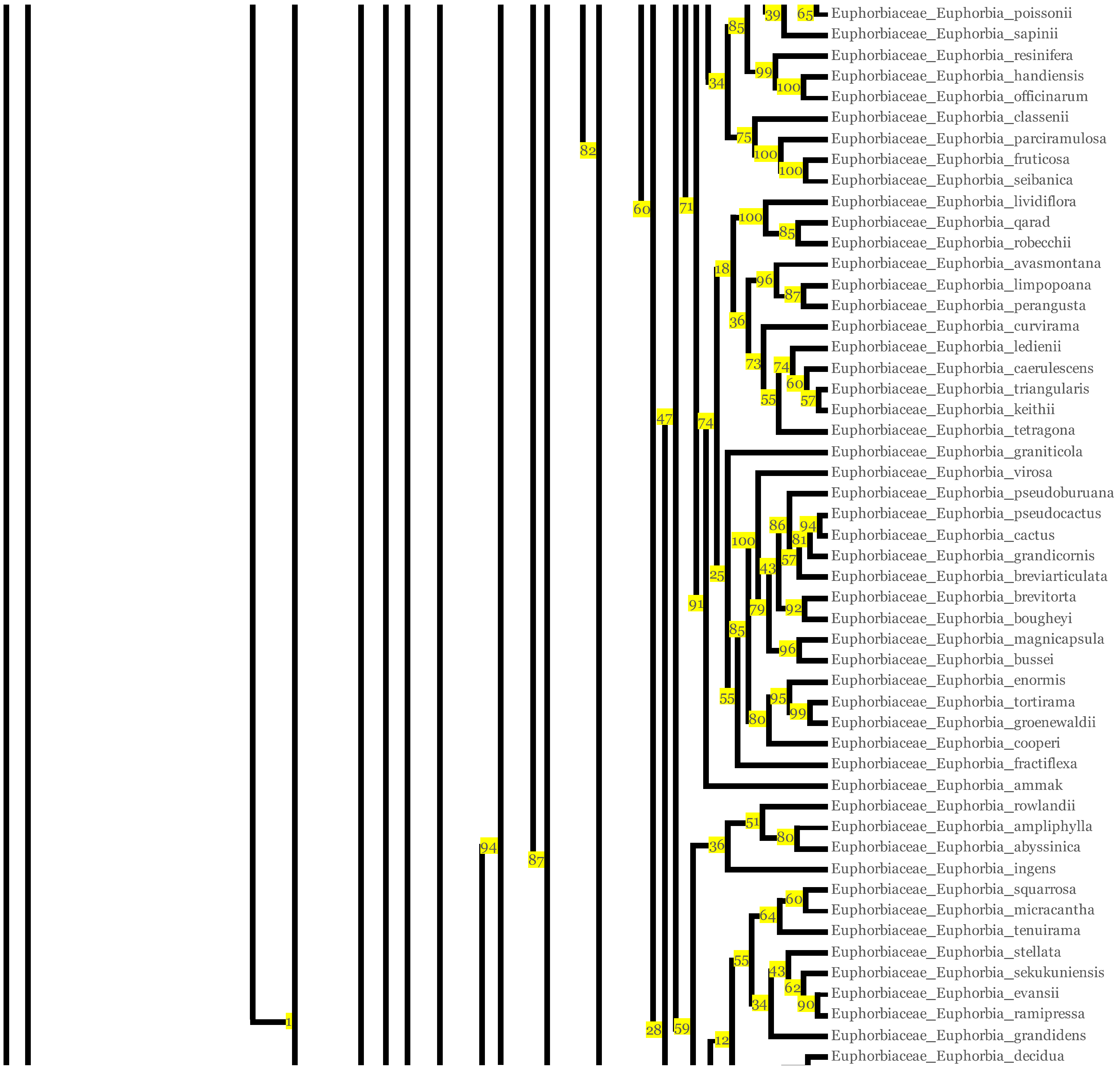

Euphorbiaceae_Euphorbia_poissonii
Euphorbiaceae_Euphorbia_sapinii
Euphorbiaceae_Euphorbia_resinifera
Euphorbiaceae_Euphorbia_handiensis
Euphorbiaceae_Euphorbia_officinarum
Euphorbiaceae_Euphorbia_classenii
Euphorbiaceae_Euphorbia_parciramulosa
Euphorbiaceae_Euphorbia_fruticosa
Euphorbiaceae_Euphorbia_seibanica
Euphorbiaceae_Euphorbia_lividiflora
Euphorbiaceae_Euphorbia_qarad
Euphorbiaceae_Euphorbia_robecchii
Euphorbiaceae_Euphorbia_avasmontana
Euphorbiaceae_Euphorbia_limpopoana
Euphorbiaceae_Euphorbia_perangusta
Euphorbiaceae_Euphorbia_curvirama
Euphorbiaceae_Euphorbia_ledienii
Euphorbiaceae_Euphorbia_caerulescens
Euphorbiaceae_Euphorbia_triangularis
Euphorbiaceae_Euphorbia_keithii
Euphorbiaceae_Euphorbia_tetragona
Euphorbiaceae_Euphorbia_graniticola
Euphorbiaceae_Euphorbia_virosa
Euphorbiaceae_Euphorbia_pseudoburuana
Euphorbiaceae_Euphorbia_pseudocactus
Euphorbiaceae_Euphorbia_cactus
Euphorbiaceae_Euphorbia_grandicornis
Euphorbiaceae_Euphorbia_breviarticulata
Euphorbiaceae_Euphorbia_brevitorta
Euphorbiaceae_Euphorbia_bougheyi
Euphorbiaceae_Euphorbia_magnicapsula
Euphorbiaceae_Euphorbia_bussei
Euphorbiaceae_Euphorbia_enormis
Euphorbiaceae_Euphorbia_tortirama
Euphorbiaceae_Euphorbia_groenewaldii
Euphorbiaceae_Euphorbia_cooperi
Euphorbiaceae_Euphorbia_fractiflexa
Euphorbiaceae_Euphorbia_ammak
Euphorbiaceae_Euphorbia_rowlandii
Euphorbiaceae_Euphorbia_ampliphylla
Euphorbiaceae_Euphorbia_abyssinica
Euphorbiaceae_Euphorbia_ingens
Euphorbiaceae_Euphorbia_squarrosa
Euphorbiaceae_Euphorbia_micracantha
Euphorbiaceae_Euphorbia_tenuirama
Euphorbiaceae_Euphorbia_stellata
Euphorbiaceae_Euphorbia_sekukuniensis
Euphorbiaceae_Euphorbia_evansii
Euphorbiaceae_Euphorbia_ramipressa
Euphorbiaceae_Euphorbia_grandidens
Euphorbiaceae_Euphorbia_decidua

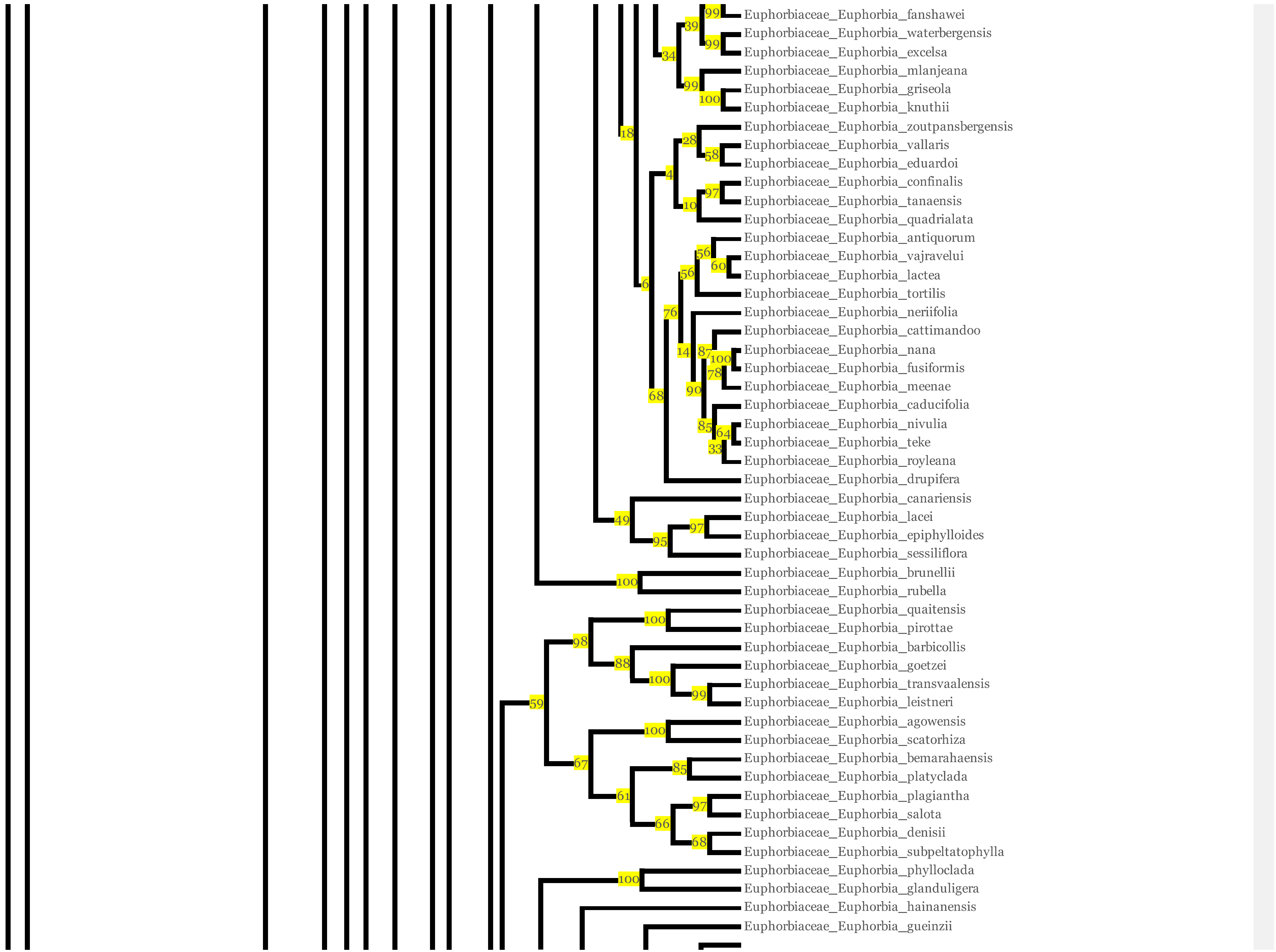
Euphorbiaceae_Euphorbia_fanshawei
Euphorbiaceae_Euphorbia_waterbergensis
Euphorbiaceae_Euphorbia_excelsa
Euphorbiaceae_Euphorbia_mlanjeana
Euphorbiaceae_Euphorbia_griseola
Euphorbiaceae_Euphorbia_knuthii
Euphorbiaceae_Euphorbia_zoutpansbergensis
Euphorbiaceae_Euphorbia_vallaris
Euphorbiaceae_Euphorbia_eduardoi
Euphorbiaceae_Euphorbia_confinalis
Euphorbiaceae_Euphorbia_tanaensis
Euphorbiaceae_Euphorbia_quadrialata
Euphorbiaceae_Euphorbia_antiquorum
Euphorbiaceae_Euphorbia_vajravelui
Euphorbiaceae_Euphorbia_lactea
Euphorbiaceae_Euphorbia_tortilis
Euphorbiaceae_Euphorbia_neriifolia
Euphorbiaceae_Euphorbia_cattimandoo
Euphorbiaceae_Euphorbia_nana
Euphorbiaceae_Euphorbia_fusiformis
Euphorbiaceae_Euphorbia_meenae
Euphorbiaceae_Euphorbia_caducifolia
Euphorbiaceae_Euphorbia_nivulia
Euphorbiaceae_Euphorbia_teke
Euphorbiaceae_Euphorbia_royleana
Euphorbiaceae_Euphorbia_drupifera
Euphorbiaceae_Euphorbia_canariensis
Euphorbiaceae_Euphorbia_lacei
Euphorbiaceae_Euphorbia_epiphylloides
Euphorbiaceae_Euphorbia_sessiliflora
Euphorbiaceae_Euphorbia_brunellii
Euphorbiaceae_Euphorbia_rubella
Euphorbiaceae_Euphorbia_quaitensis
Euphorbiaceae_Euphorbia_pirottae
Euphorbiaceae_Euphorbia_barbicollis
Euphorbiaceae_Euphorbia_goetzei
Euphorbiaceae_Euphorbia_transvaalensis
Euphorbiaceae_Euphorbia_leistneri
Euphorbiaceae_Euphorbia_agowensis
Euphorbiaceae_Euphorbia_scatorhiza
Euphorbiaceae_Euphorbia_bemarahaensis
Euphorbiaceae_Euphorbia_platyclada
Euphorbiaceae_Euphorbia_plagiantha
Euphorbiaceae_Euphorbia_salota
Euphorbiaceae_Euphorbia_denisii
Euphorbiaceae_Euphorbia_subpeltatophylla
Euphorbiaceae_Euphorbia_phylloclada
Euphorbiaceae_Euphorbia_glanduligera
Euphorbiaceae_Euphorbia_hainanensis
Euphorbiaceae_Euphorbia_gueinzii
99
39
99
34
99
100
18
28
58
4
97
10
56
60
56
6
76
14
87
100
78
90
68
85
64
33
49
97
95
100
100
98
88
100
99
59
100
67
85
61
97
66
68
100

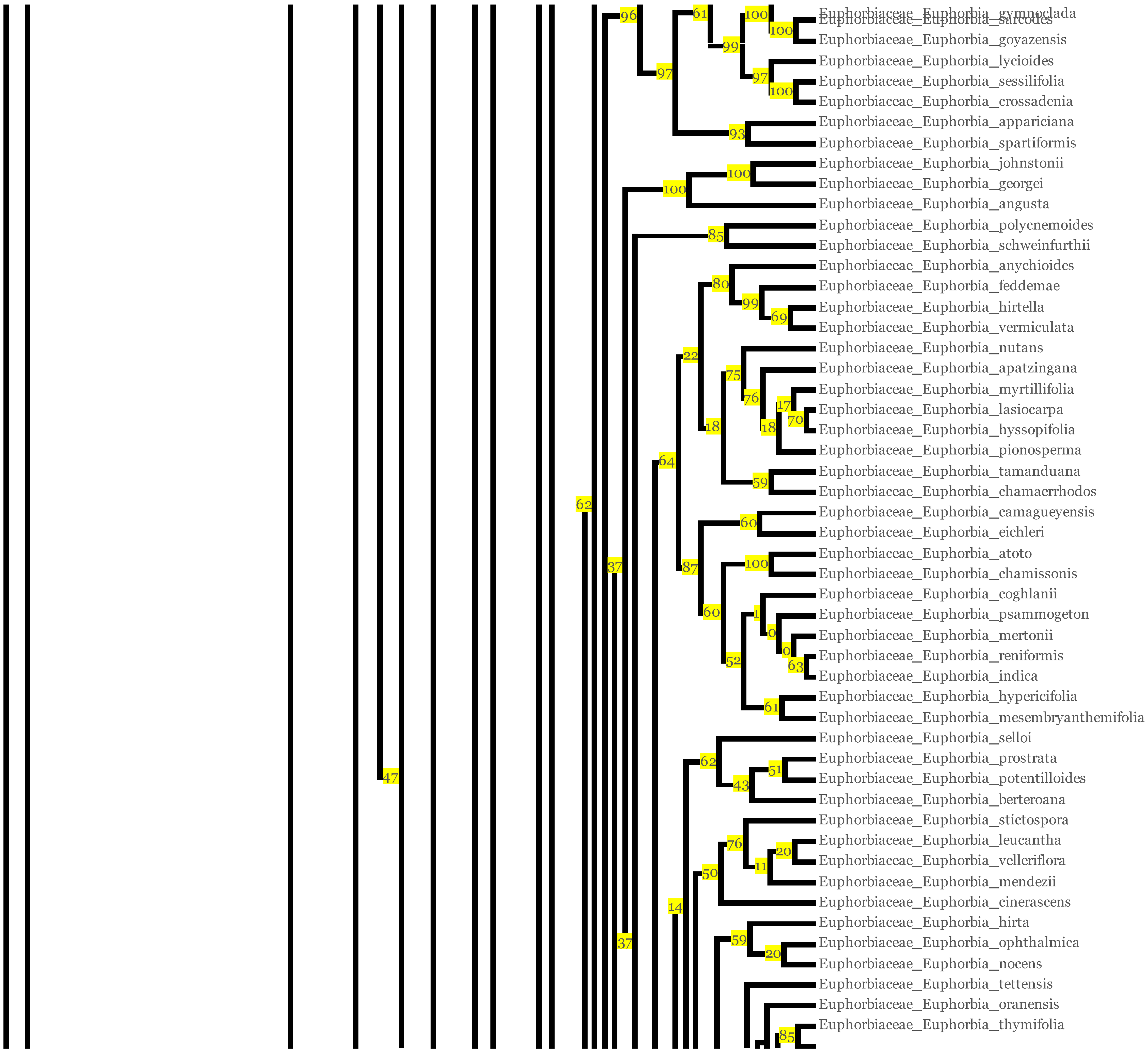
Euphorbiaceae_Euphorbia_gymnoclada
Euphorbiaceae_Euphorbia_sarcodes
Euphorbiaceae_Euphorbia_goyazensis
Euphorbiaceae_Euphorbia_lycioides
Euphorbiaceae_Euphorbia_sessilifolia
Euphorbiaceae_Euphorbia_crossadenia
Euphorbiaceae_Euphorbia_appariciana
Euphorbiaceae_Euphorbia_spartiformis
Euphorbiaceae_Euphorbia_johnstonii
Euphorbiaceae_Euphorbia_georgei
Euphorbiaceae_Euphorbia_angusta
Euphorbiaceae_Euphorbia_polycnemoides
Euphorbiaceae_Euphorbia_schweinfurthii
Euphorbiaceae_Euphorbia_anychioides
Euphorbiaceae_Euphorbia_feddemae
Euphorbiaceae_Euphorbia_hirtella
Euphorbiaceae_Euphorbia_vermiculata
Euphorbiaceae_Euphorbia_nutans
Euphorbiaceae_Euphorbia_apatzingana
Euphorbiaceae_Euphorbia_myrtillifolia
Euphorbiaceae_Euphorbia_lasiocarpa
Euphorbiaceae_Euphorbia_hyssopifolia
Euphorbiaceae_Euphorbia_pionosperma
Euphorbiaceae_Euphorbia_tamanduana
Euphorbiaceae_Euphorbia_chamaerrhodos
Euphorbiaceae_Euphorbia_camagueyensis
Euphorbiaceae_Euphorbia_eichleri
Euphorbiaceae_Euphorbia_atoto
Euphorbiaceae_Euphorbia_chamissonis
Euphorbiaceae_Euphorbia_coghlanii
Euphorbiaceae_Euphorbia_psammogeton
Euphorbiaceae_Euphorbia_mertonii
Euphorbiaceae_Euphorbia_reniformis
Euphorbiaceae_Euphorbia_indica
Euphorbiaceae_Euphorbia_hypericifolia
Euphorbiaceae_Euphorbia_mesembryanthemifolia
Euphorbiaceae_Euphorbia_selloi
Euphorbiaceae_Euphorbia_prostrata
Euphorbiaceae_Euphorbia_potentilloides
Euphorbiaceae_Euphorbia_berteroana
Euphorbiaceae_Euphorbia_stictospora
Euphorbiaceae_Euphorbia_leucantha
Euphorbiaceae_Euphorbia_velleriflora
Euphorbiaceae_Euphorbia_mendezii
Euphorbiaceae_Euphorbia_cinerascens
Euphorbiaceae_Euphorbia_hirta
Euphorbiaceae_Euphorbia_ophthalmica
Euphorbiaceae_Euphorbia_nocens
Euphorbiaceae_Euphorbia_tettensis
Euphorbiaceae_Euphorbia_oranensis
Euphorbiaceae_Euphorbia_thymifolia

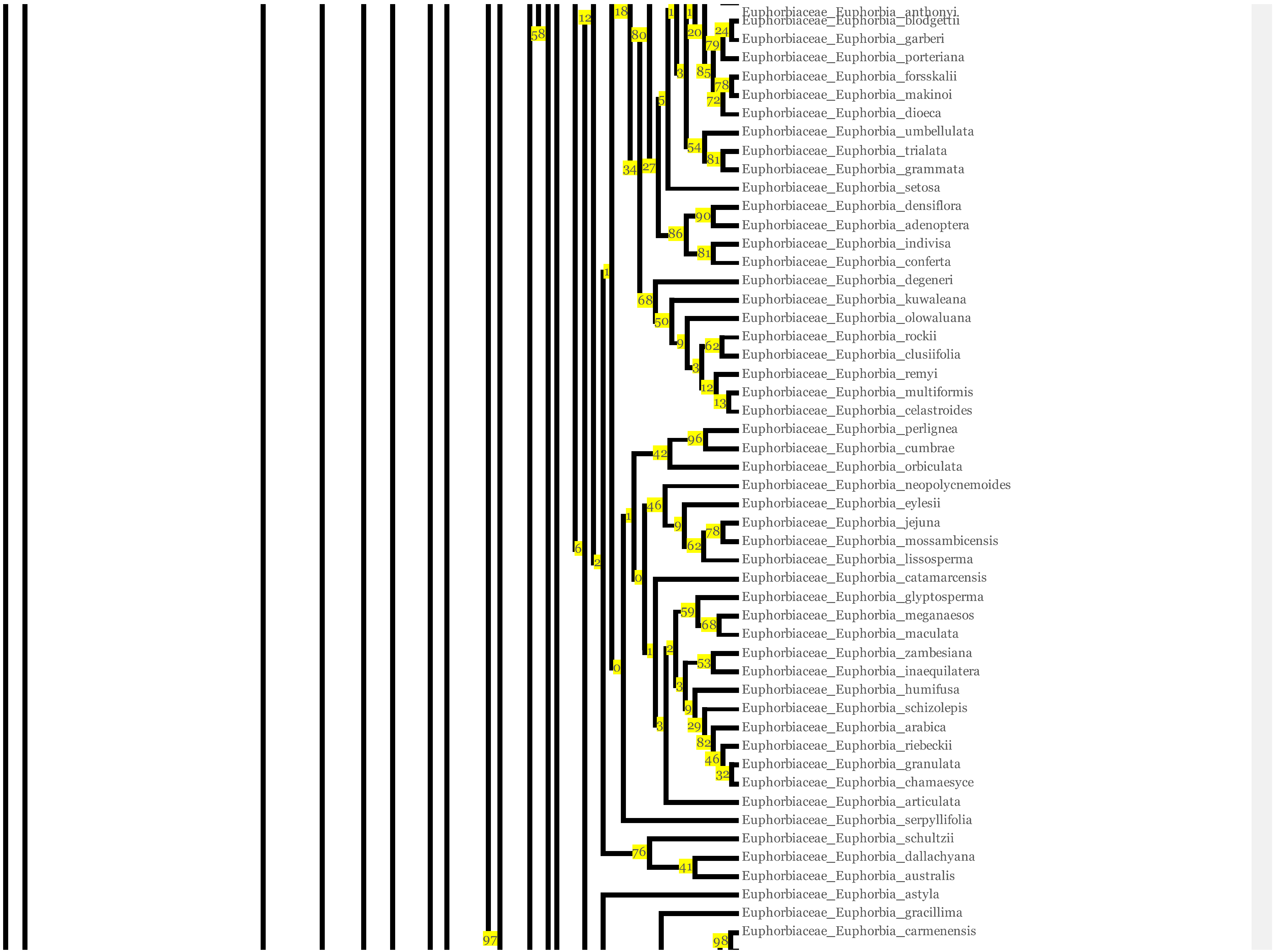
Euphorbiaceae_Euphorbia_anthonyi
Euphorbiaceae_Euphorbia_blodgettii
Euphorbiaceae_Euphorbia_garberi
Euphorbiaceae_Euphorbia_porteriana
Euphorbiaceae_Euphorbia_forsskalii
Euphorbiaceae_Euphorbia_makinoi
Euphorbiaceae_Euphorbia_dioeca
Euphorbiaceae_Euphorbia_umbellulata
Euphorbiaceae_Euphorbia_trialata
Euphorbiaceae_Euphorbia_grammata
Euphorbiaceae_Euphorbia_setosa
Euphorbiaceae_Euphorbia_densiflora
Euphorbiaceae_Euphorbia_adenoptera
Euphorbiaceae_Euphorbia_indivisa
Euphorbiaceae_Euphorbia_conferta
Euphorbiaceae_Euphorbia_degeneri
Euphorbiaceae_Euphorbia_kuwaleana
Euphorbiaceae_Euphorbia_olowaluana
Euphorbiaceae_Euphorbia_rockii
Euphorbiaceae_Euphorbia_clusiifolia
Euphorbiaceae_Euphorbia_remyi
Euphorbiaceae_Euphorbia_multiformis
Euphorbiaceae_Euphorbia_celastroides
Euphorbiaceae_Euphorbia_perlignea
Euphorbiaceae_Euphorbia_cumbrae
Euphorbiaceae_Euphorbia_orbiculata
Euphorbiaceae_Euphorbia_neopolycnemoides
Euphorbiaceae_Euphorbia_eylesii
Euphorbiaceae_Euphorbia_jejuna
Euphorbiaceae_Euphorbia_mossambicensis
Euphorbiaceae_Euphorbia_lissosperma
Euphorbiaceae_Euphorbia_catamarcensis
Euphorbiaceae_Euphorbia_glyptosperma
Euphorbiaceae_Euphorbia_meganaesos
Euphorbiaceae_Euphorbia_maculata
Euphorbiaceae_Euphorbia_zambesiana
Euphorbiaceae_Euphorbia_inaequilatera
Euphorbiaceae_Euphorbia_humifusa
Euphorbiaceae_Euphorbia_schizolepis
Euphorbiaceae_Euphorbia_arabica
Euphorbiaceae_Euphorbia_riebeckii
Euphorbiaceae_Euphorbia_granulata
Euphorbiaceae_Euphorbia_chamaesyce
Euphorbiaceae_Euphorbia_articulata
Euphorbiaceae_Euphorbia_serpyllifolia
Euphorbiaceae_Euphorbia_schultzii
Euphorbiaceae_Euphorbia_dallachyana
Euphorbiaceae_Euphorbia_australis
Euphorbiaceae_Euphorbia_astyla
Euphorbiaceae_Euphorbia_gracillima
Euphorbiaceae_Euphorbia_carmenensis

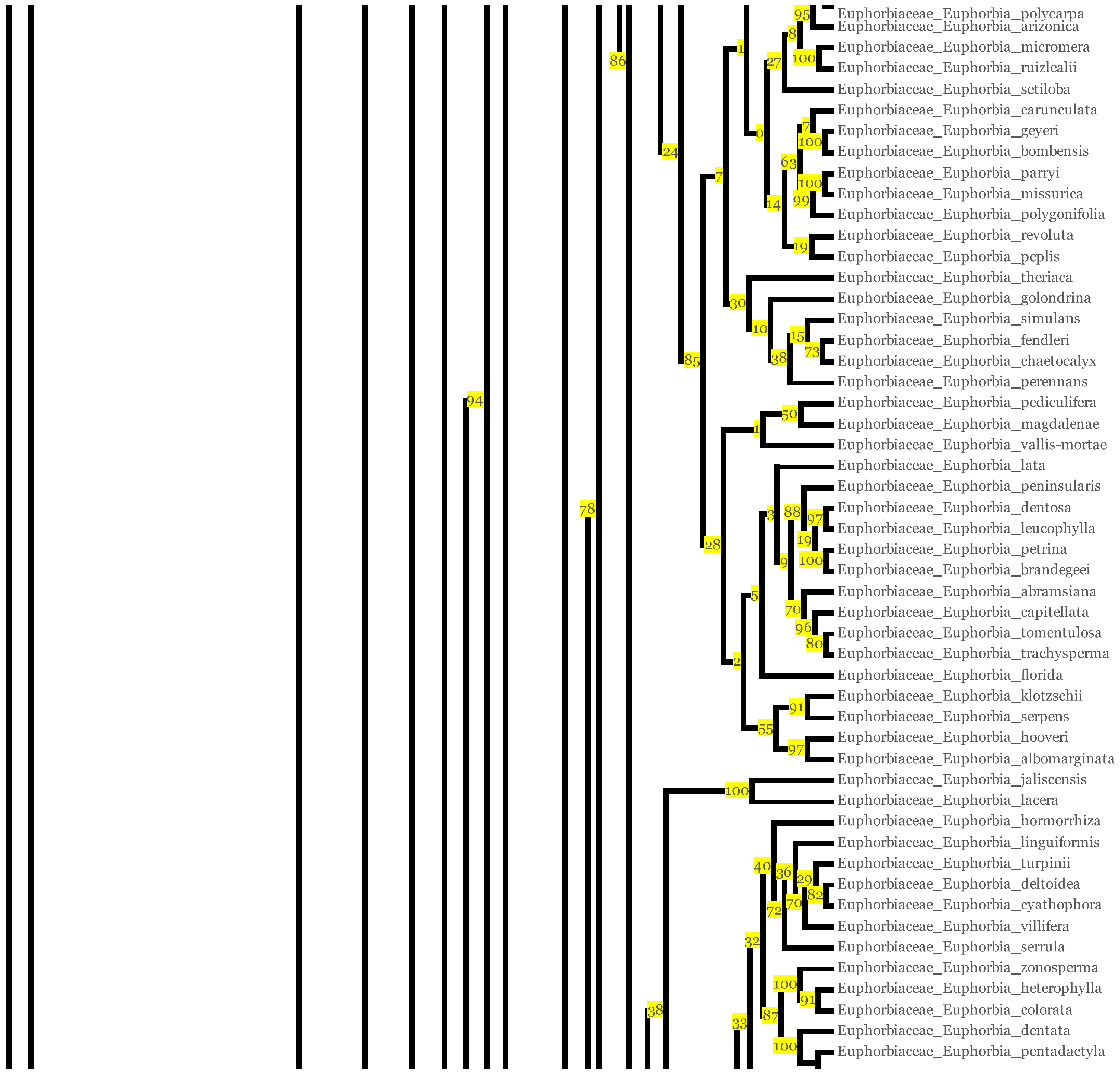
Euphorbiaceae_Euphorbia_polycarpa
Euphorbiaceae_Euphorbia_arizonica
Euphorbiaceae_Euphorbia_micromera
Euphorbiaceae_Euphorbia_ruizlealii
Euphorbiaceae_Euphorbia_setiloba
Euphorbiaceae_Euphorbia_carunculata
Euphorbiaceae_Euphorbia_geyeri
Euphorbiaceae_Euphorbia_bombensis
Euphorbiaceae_Euphorbia_parryi
Euphorbiaceae_Euphorbia_missurica
Euphorbiaceae_Euphorbia_polygonifolia
Euphorbiaceae_Euphorbia_revoluta
Euphorbiaceae_Euphorbia_peplis
Euphorbiaceae_Euphorbia_theriaca
Euphorbiaceae_Euphorbia_golondrina
Euphorbiaceae_Euphorbia_simulans
Euphorbiaceae_Euphorbia_fendleri
Euphorbiaceae_Euphorbia_chaetocalyx
Euphorbiaceae_Euphorbia_perennans
Euphorbiaceae_Euphorbia_pediculifera
Euphorbiaceae_Euphorbia_magdalenae
Euphorbiaceae_Euphorbia_vallis-mortae
Euphorbiaceae_Euphorbia_lata
Euphorbiaceae_Euphorbia_peninsularis
Euphorbiaceae_Euphorbia_dentosa
Euphorbiaceae_Euphorbia_leucophylla
Euphorbiaceae_Euphorbia_petrina
Euphorbiaceae_Euphorbia_brandegeei
Euphorbiaceae_Euphorbia_abramsiana
Euphorbiaceae_Euphorbia_capitellata
Euphorbiaceae_Euphorbia_tomentulosa
Euphorbiaceae_Euphorbia_trachysperma
Euphorbiaceae_Euphorbia_florida
Euphorbiaceae_Euphorbia_klotzschii
Euphorbiaceae_Euphorbia_serpens
Euphorbiaceae_Euphorbia_hooveri
Euphorbiaceae_Euphorbia_albomarginata
Euphorbiaceae_Euphorbia_jaliscensis
Euphorbiaceae_Euphorbia_lacera
Euphorbiaceae_Euphorbia_hormorrhiza
Euphorbiaceae_Euphorbia_linguiformis
Euphorbiaceae_Euphorbia_turpinii
Euphorbiaceae_Euphorbia_deltoidea
Euphorbiaceae_Euphorbia_cyathophora
Euphorbiaceae_Euphorbia_villifera
Euphorbiaceae_Euphorbia_serrula
Euphorbiaceae_Euphorbia_zonosperma
Euphorbiaceae_Euphorbia_heterophylla
Euphorbiaceae_Euphorbia_colorata
Euphorbiaceae_Euphorbia_dentata
Euphorbiaceae_Euphorbia_pentadactyla

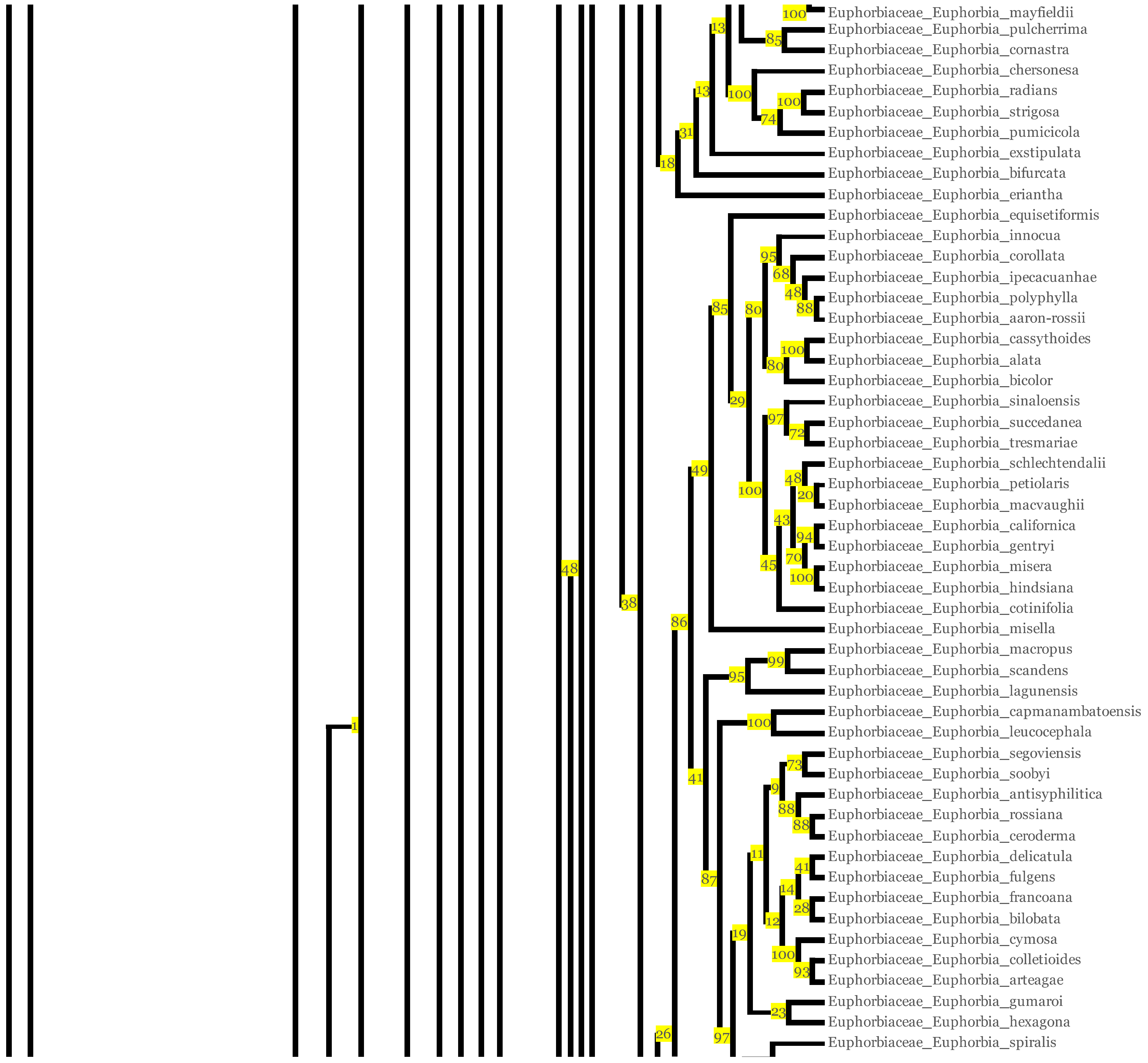
Euphorbiaceae_Euphorbia_mayfieldii
Euphorbiaceae_Euphorbia_pulcherrima
Euphorbiaceae_Euphorbia_cornastra
Euphorbiaceae_Euphorbia_chersonesa
Euphorbiaceae_Euphorbia_radians
Euphorbiaceae_Euphorbia_strigosa
Euphorbiaceae_Euphorbia_pumicicola
Euphorbiaceae_Euphorbia_exstipulata
Euphorbiaceae_Euphorbia_bifurcata
Euphorbiaceae_Euphorbia_eriantha
Euphorbiaceae_Euphorbia_equisetiformis
Euphorbiaceae_Euphorbia_innocua
Euphorbiaceae_Euphorbia_corollata
Euphorbiaceae_Euphorbia_ipecacuanhae
Euphorbiaceae_Euphorbia_polyphylla
Euphorbiaceae_Euphorbia_aaron-rossii
Euphorbiaceae_Euphorbia_cassythoides
Euphorbiaceae_Euphorbia_alata
Euphorbiaceae_Euphorbia_bicolor
Euphorbiaceae_Euphorbia_sinaloensis
Euphorbiaceae_Euphorbia_succedanea
Euphorbiaceae_Euphorbia_tresmariae
Euphorbiaceae_Euphorbia_schlechtendalii
Euphorbiaceae_Euphorbia_petiolaris
Euphorbiaceae_Euphorbia_macvaughii
Euphorbiaceae_Euphorbia_californica
Euphorbiaceae_Euphorbia_gentryi
Euphorbiaceae_Euphorbia_misera
Euphorbiaceae_Euphorbia_hindsiana
Euphorbiaceae_Euphorbia_cotinifolia
Euphorbiaceae_Euphorbia_misella
Euphorbiaceae_Euphorbia_macropus
Euphorbiaceae_Euphorbia_scandens
Euphorbiaceae_Euphorbia_lagunensis
Euphorbiaceae_Euphorbia_capmanambatoensis
Euphorbiaceae_Euphorbia_leucocephala
Euphorbiaceae_Euphorbia_segoviensis
Euphorbiaceae_Euphorbia_soobyi
Euphorbiaceae_Euphorbia_antisyphilitica
Euphorbiaceae_Euphorbia_rossiana
Euphorbiaceae_Euphorbia_ceroderma
Euphorbiaceae_Euphorbia_delicatula
Euphorbiaceae_Euphorbia_fulgens
Euphorbiaceae_Euphorbia_francoana
Euphorbiaceae_Euphorbia_bilobata
Euphorbiaceae_Euphorbia_cymosa
Euphorbiaceae_Euphorbia_colletioides
Euphorbiaceae_Euphorbia_arteagae
Euphorbiaceae_Euphorbia_gumaroi
Euphorbiaceae_Euphorbia_hexagona
Euphorbiaceae_Euphorbia_spiralis

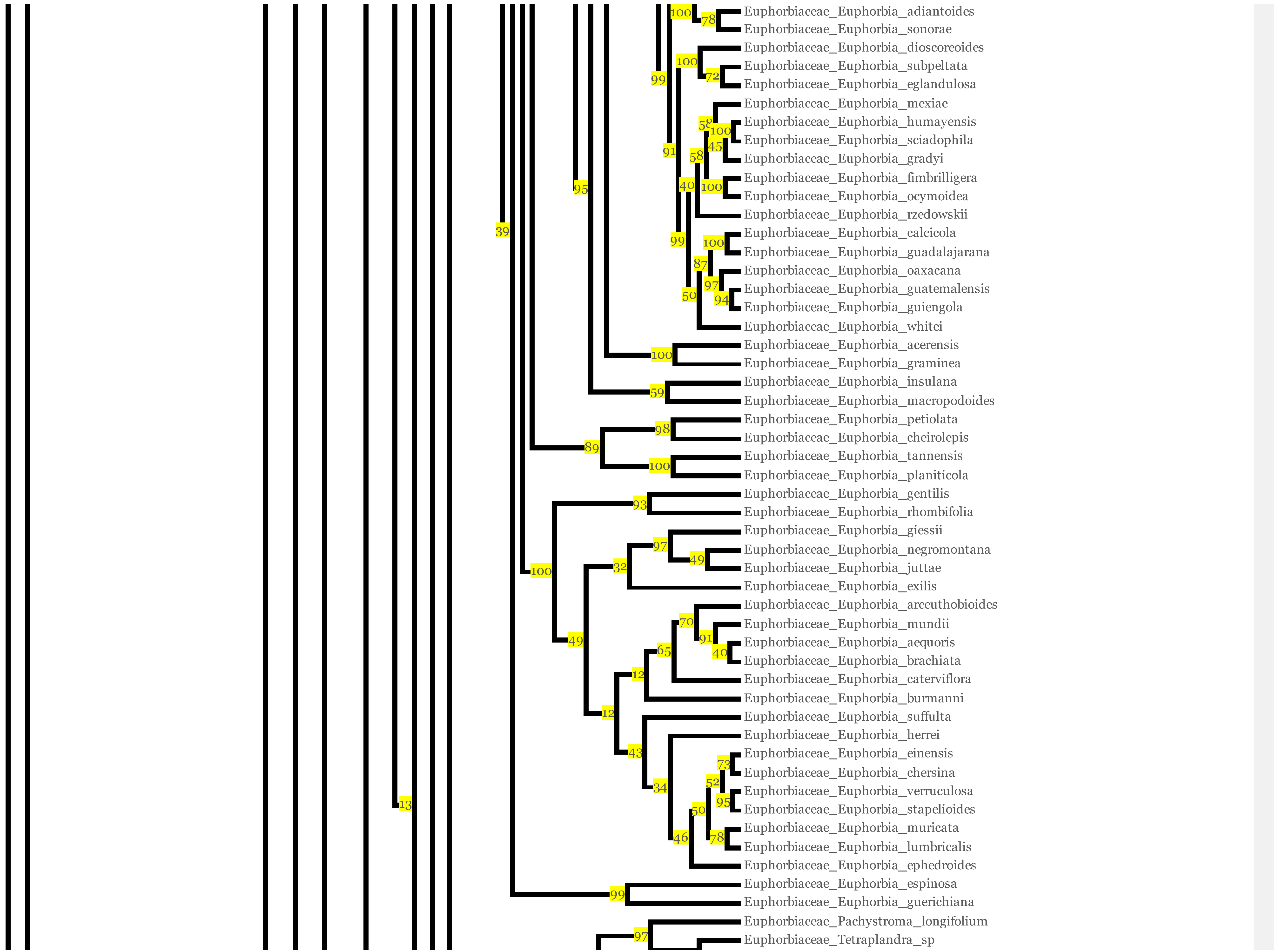

Euphorbiaceae_Euphorbia_adiantoides
Euphorbiaceae_Euphorbia_sonorae
Euphorbiaceae_Euphorbia_dioscoreoides
Euphorbiaceae_Euphorbia_subpeltata
Euphorbiaceae_Euphorbia_eglandulosa
Euphorbiaceae_Euphorbia_mexiae
Euphorbiaceae_Euphorbia_humayensis
Euphorbiaceae_Euphorbia_sciadophila
Euphorbiaceae_Euphorbia_gradyi
Euphorbiaceae_Euphorbia_fimbrilligera
Euphorbiaceae_Euphorbia_ocymoidea
Euphorbiaceae_Euphorbia_rzedowskii
Euphorbiaceae_Euphorbia_calcicola
Euphorbiaceae_Euphorbia_guadalajarana
Euphorbiaceae_Euphorbia_oaxacana
Euphorbiaceae_Euphorbia_guatemalensis
Euphorbiaceae_Euphorbia_guiengola
Euphorbiaceae_Euphorbia_whitei
Euphorbiaceae_Euphorbia_acerensis
Euphorbiaceae_Euphorbia_graminea
Euphorbiaceae_Euphorbia_insulana
Euphorbiaceae_Euphorbia_macropodoides
Euphorbiaceae_Euphorbia_petiolata
Euphorbiaceae_Euphorbia_cheirolepis
Euphorbiaceae_Euphorbia_tannensis
Euphorbiaceae_Euphorbia_planiticola
Euphorbiaceae_Euphorbia_gentilis
Euphorbiaceae_Euphorbia_rhombifolia
Euphorbiaceae_Euphorbia_giessii
Euphorbiaceae_Euphorbia_negromontana
Euphorbiaceae_Euphorbia_juttae
Euphorbiaceae_Euphorbia_exilis
Euphorbiaceae_Euphorbia_arceuthobioides
Euphorbiaceae_Euphorbia_mundii
Euphorbiaceae_Euphorbia_aequoris
Euphorbiaceae_Euphorbia_brachiata
Euphorbiaceae_Euphorbia_caterviflora
Euphorbiaceae_Euphorbia_burmanni
Euphorbiaceae_Euphorbia_suffulta
Euphorbiaceae_Euphorbia_herrei
Euphorbiaceae_Euphorbia_einensis
Euphorbiaceae_Euphorbia_chersina
Euphorbiaceae_Euphorbia_verruculosa
Euphorbiaceae_Euphorbia_stapelioides
Euphorbiaceae_Euphorbia_muricata
Euphorbiaceae_Euphorbia_lumbricalis
Euphorbiaceae_Euphorbia_ephedroides
Euphorbiaceae_Euphorbia_espinosa
Euphorbiaceae_Euphorbia_guerichiana
Euphorbiaceae_Pachystroma_longifolium
Euphorbiaceae_Tetraplandra_sp
100
78
100
72
99
58
100
45
91
58
100
40
95
39
99
100
87
97
50
94
100
59
98
89
100
93
97
49
32
100
70
91
40
65
49
12
12
43
73
52
95
34
50
78
46
13
99
97

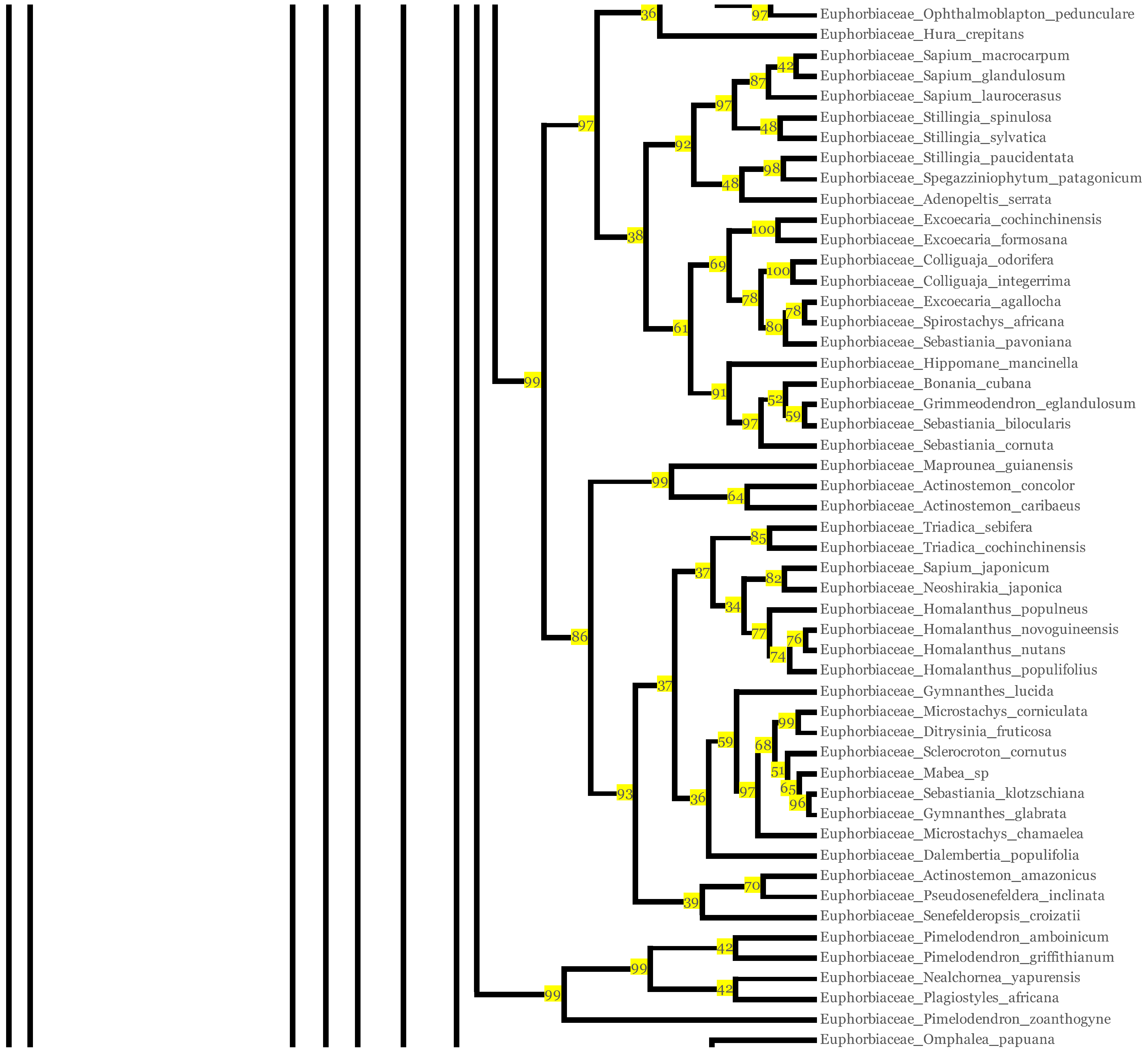
Euphorbiaceae_Ophthalmoblapton_pedunculare
Euphorbiaceae_Hura_crepitans
Euphorbiaceae_Sapium_macrocarpum
Euphorbiaceae_Sapium_glandulosum
Euphorbiaceae_Sapium_laurocerasus
Euphorbiaceae_Stillingia_spinulosa
Euphorbiaceae_Stillingia_sylvatica
Euphorbiaceae_Stillingia_paucidentata
Euphorbiaceae_Spegazziniophytum_patagonicum
Euphorbiaceae_Adenopeltis_serrata
Euphorbiaceae_Excoecaria_cochinchinensis
Euphorbiaceae_Excoecaria_formosana
Euphorbiaceae_Colliguaja_odorifera
Euphorbiaceae_Colliguaja_integerrima
Euphorbiaceae_Excoecaria_agallocha
Euphorbiaceae_Spirostachys_africana
Euphorbiaceae_Sebastiania_pavoniana
Euphorbiaceae_Hippomane_mancinella
Euphorbiaceae_Bonania_cubana
Euphorbiaceae_Grimmeodendron_eglandulosum
Euphorbiaceae_Sebastiania_bilocularis
Euphorbiaceae_Sebastiania_cornuta
Euphorbiaceae_Maprounea_guianensis
Euphorbiaceae_Actinostemon_concolor
Euphorbiaceae_Actinostemon_caribaeus
Euphorbiaceae_Triadica_sebifera
Euphorbiaceae_Triadica_cochinchinensis
Euphorbiaceae_Sapium_japonicum
Euphorbiaceae_Neoshirakia_japonica
Euphorbiaceae_Homalanthus_populneus
Euphorbiaceae_Homalanthus_novoguineensis
Euphorbiaceae_Homalanthus_nutans
Euphorbiaceae_Homalanthus_populifolius
Euphorbiaceae_Gymnanthes_lucida
Euphorbiaceae_Microstachys_corniculata
Euphorbiaceae_Ditrysinia_fruticosa
Euphorbiaceae_Sclerocroton_cornutus
Euphorbiaceae_Mabea_sp
Euphorbiaceae_Sebastiania_klotzschiana
Euphorbiaceae_Gymnanthes_glabrata
Euphorbiaceae_Microstachys_chamaelea
Euphorbiaceae_Dalembertia_populifolia
Euphorbiaceae_Actinostemon_amazonicus
Euphorbiaceae_Pseudosenefeldera_inclinata
Euphorbiaceae_Senefelderopsis_croizatii
Euphorbiaceae_Pimelodendron_amboinicum
Euphorbiaceae_Pimelodendron_griffithianum
Euphorbiaceae_Nealchornea_yapurensis
Euphorbiaceae_Plagiostyles_africana
Euphorbiaceae_Pimelodendron_zoanthogyne
Euphorbiaceae_Omphalea_papuana

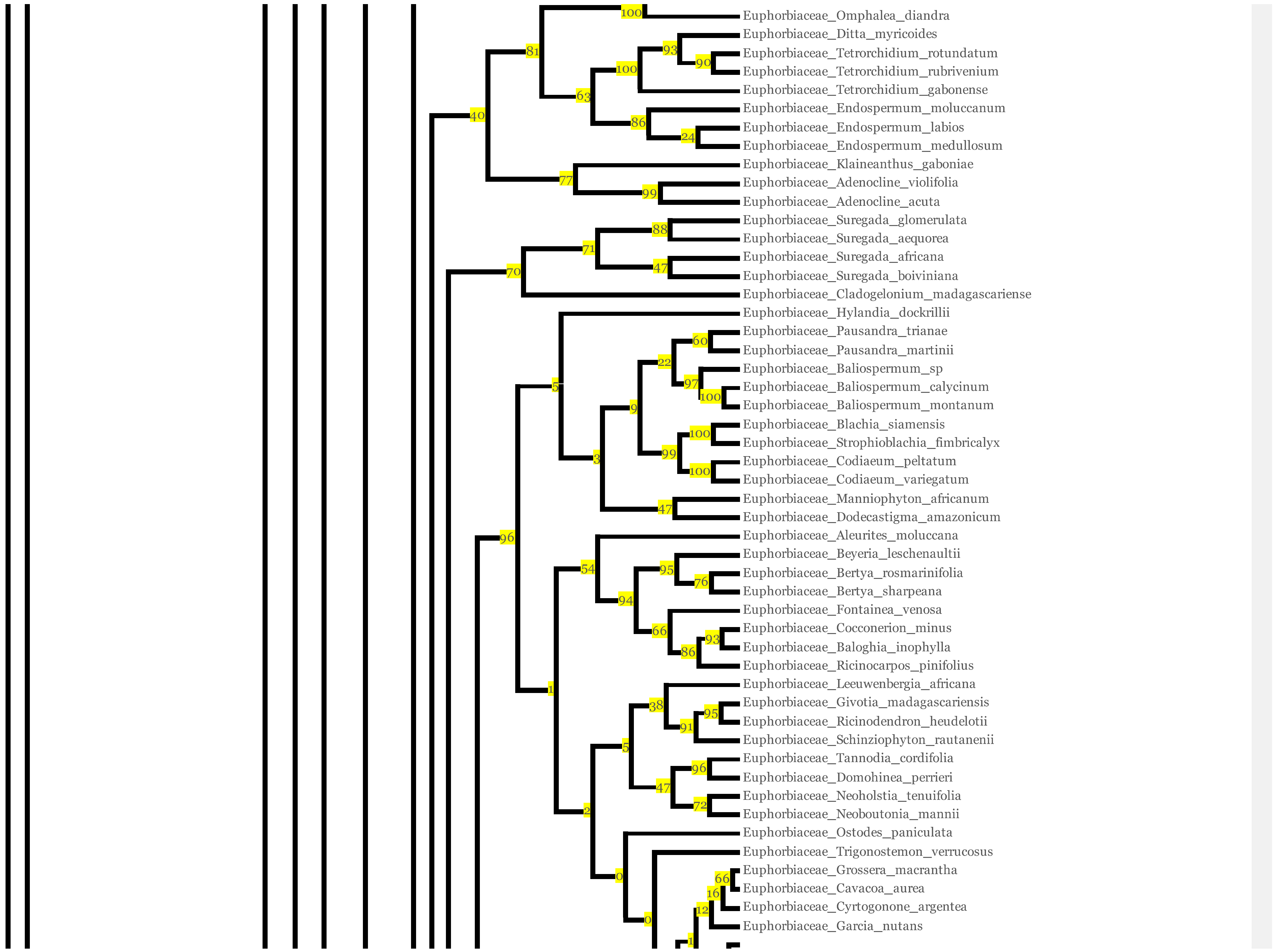
Euphorbiaceae_Omphalea_diandra
Euphorbiaceae_Ditta_myricoides
Euphorbiaceae_Tetrorchidium_rotundatum
Euphorbiaceae_Tetrorchidium_rubrivenium
Euphorbiaceae_Tetrorchidium_gabonense
Euphorbiaceae_Endospermum_moluccanum
Euphorbiaceae_Endospermum_labios
Euphorbiaceae_Endospermum_medullosum
Euphorbiaceae_Klaineanthus_gaboniae
Euphorbiaceae_Adenocline_violifolia
Euphorbiaceae_Adenocline_acuta
Euphorbiaceae_Suregada_glomerulata
Euphorbiaceae_Suregada_aequorea
Euphorbiaceae_Suregada_africana
Euphorbiaceae_Suregada_boiviniana
Euphorbiaceae_Cladogelonium_madagascariense
Euphorbiaceae_Hylandia_dockrillii
Euphorbiaceae_Pausandra_trianae
Euphorbiaceae_Pausandra_martinii
Euphorbiaceae_Baliospermum_sp
Euphorbiaceae_Baliospermum_calycinum
Euphorbiaceae_Baliospermum_montanum
Euphorbiaceae_Blachia_siamensis
Euphorbiaceae_Strophioblachia_fimbricalyx
Euphorbiaceae_Codiaeum_peltatum
Euphorbiaceae_Codiaeum_variegatum
Euphorbiaceae_Manniophyton_africanum
Euphorbiaceae_Dodecastigma_amazonicum
Euphorbiaceae_Aleurites_moluccana
Euphorbiaceae_Beyeria_leschenaultii
Euphorbiaceae_Bertya_rosmarinifolia
Euphorbiaceae_Bertya_sharpeana
Euphorbiaceae_Fontainea_venosa
Euphorbiaceae_Cocconerion_minus
Euphorbiaceae_Baloghia_inophylla
Euphorbiaceae_Ricinocarpos_pinifolius
Euphorbiaceae_Leeuwenbergia_africana
Euphorbiaceae_Givotia_madagascariensis
Euphorbiaceae_Ricinodendron_heudelotii
Euphorbiaceae_Schinziophyton_rautanenii
Euphorbiaceae_Tannodia_cordifolia
Euphorbiaceae_Domohinea_perrieri
Euphorbiaceae_Neoholstia_tenuifolia
Euphorbiaceae_Neoboutonia_mannii
Euphorbiaceae_Ostodes_paniculata
Euphorbiaceae_Trigonostemon_verrucosus
Euphorbiaceae_Grossera_macrantha
Euphorbiaceae_Cavacoa_aurea
Euphorbiaceae_Cyrtogonone_argentea
Euphorbiaceae_Garcia_nutans

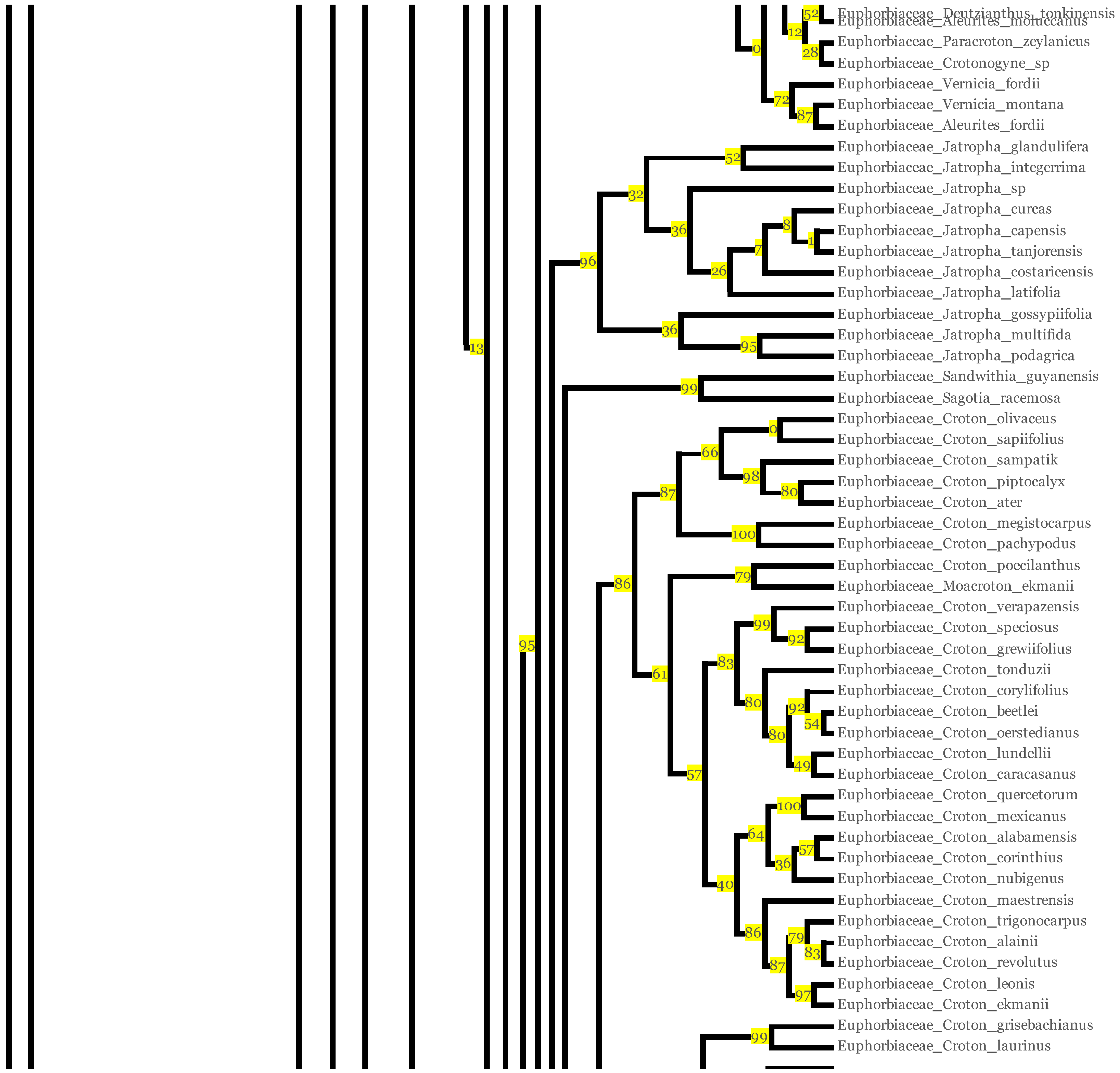
Euphorbiaceae_Deutzianthus_tonkinensis
Euphorbiaceae_Aleurites_moluccanus
Euphorbiaceae_Paracroton_zeylanicus
Euphorbiaceae_Crotonogyne_sp
Euphorbiaceae_Vernicia_fordii
Euphorbiaceae_Vernicia_montana
Euphorbiaceae_Aleurites_fordii
Euphorbiaceae_Jatropha_glandulifera
Euphorbiaceae_Jatropha_integerrima
Euphorbiaceae_Jatropha_sp
Euphorbiaceae_Jatropha_curcas
Euphorbiaceae_Jatropha_capensis
Euphorbiaceae_Jatropha_tanjorensis
Euphorbiaceae_Jatropha_costaricensis
Euphorbiaceae_Jatropha_latifolia
Euphorbiaceae_Jatropha_gossypiifolia
Euphorbiaceae_Jatropha_multifida
Euphorbiaceae_Jatropha_podagrica
Euphorbiaceae_Sandwithia_guyanensis
Euphorbiaceae_Sagotia_racemosa
Euphorbiaceae_Croton_olivaceus
Euphorbiaceae_Croton_sapiifolius
Euphorbiaceae_Croton_sampatik
Euphorbiaceae_Croton_piptocalyx
Euphorbiaceae_Croton_ater
Euphorbiaceae_Croton_megistocarpus
Euphorbiaceae_Croton_pachypodus
Euphorbiaceae_Croton_poecilanthus
Euphorbiaceae_Moacroton_ekmanii
Euphorbiaceae_Croton_verapazensis
Euphorbiaceae_Croton_speciosus
Euphorbiaceae_Croton_grewiifolius
Euphorbiaceae_Croton_tonduzii
Euphorbiaceae_Croton_corylifolius
Euphorbiaceae_Croton_beetlei
Euphorbiaceae_Croton_oerstedianus
Euphorbiaceae_Croton_lundellii
Euphorbiaceae_Croton_caracasanus
Euphorbiaceae_Croton_quercetorum
Euphorbiaceae_Croton_mexicanus
Euphorbiaceae_Croton_alabamensis
Euphorbiaceae_Croton_corinthius
Euphorbiaceae_Croton_nubigenus
Euphorbiaceae_Croton_maestrensis
Euphorbiaceae_Croton_trigonocarpus
Euphorbiaceae_Croton_alainii
Euphorbiaceae_Croton_revolutus
Euphorbiaceae_Croton_leonis
Euphorbiaceae_Croton_ekmanii
Euphorbiaceae_Croton_grisebachianus
Euphorbiaceae_Croton_laurinus

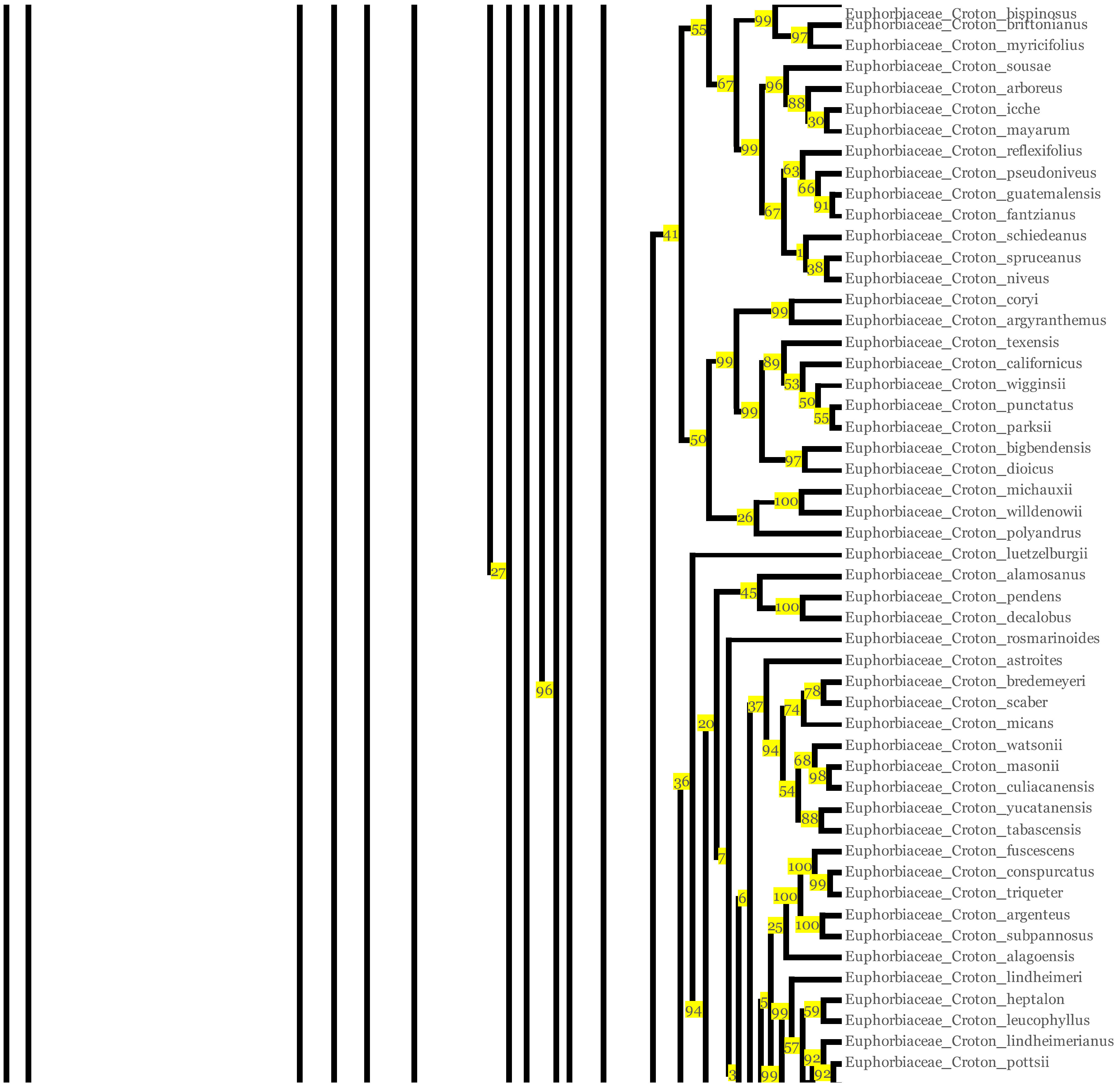
Euphorbiaceae_Croton_bispinosus
Euphorbiaceae_Croton_brittonianus
Euphorbiaceae_Croton_myricifolius
Euphorbiaceae_Croton_sousae
Euphorbiaceae_Croton_arboreus
Euphorbiaceae_Croton_icche
Euphorbiaceae_Croton_mayarum
Euphorbiaceae_Croton_reflexifolius
Euphorbiaceae_Croton_pseudoniveus
Euphorbiaceae_Croton_guatemalensis
Euphorbiaceae_Croton_fantzianus
Euphorbiaceae_Croton_schiedeanus
Euphorbiaceae_Croton_spruceanus
Euphorbiaceae_Croton_niveus
Euphorbiaceae_Croton_coryi
Euphorbiaceae_Croton_argyranthemus
Euphorbiaceae_Croton_texensis
Euphorbiaceae_Croton_californicus
Euphorbiaceae_Croton_wigginsii
Euphorbiaceae_Croton_punctatus
Euphorbiaceae_Croton_parksii
Euphorbiaceae_Croton_bigbendensis
Euphorbiaceae_Croton_dioicus
Euphorbiaceae_Croton_michauxii
Euphorbiaceae_Croton_willdenowii
Euphorbiaceae_Croton_polyandrus
Euphorbiaceae_Croton_luetzelburgii
Euphorbiaceae_Croton_alamosanus
Euphorbiaceae_Croton_pendens
Euphorbiaceae_Croton_decalobus
Euphorbiaceae_Croton_rosmarinoides
Euphorbiaceae_Croton_astroites
Euphorbiaceae_Croton_bredemeyeri
Euphorbiaceae_Croton_scaber
Euphorbiaceae_Croton_micans
Euphorbiaceae_Croton_watsonii
Euphorbiaceae_Croton_masonii
Euphorbiaceae_Croton_culiacanensis
Euphorbiaceae_Croton_yucatanensis
Euphorbiaceae_Croton_tabascensis
Euphorbiaceae_Croton_fuscescens
Euphorbiaceae_Croton_conspurcatus
Euphorbiaceae_Croton_triqueter
Euphorbiaceae_Croton_argenteus
Euphorbiaceae_Croton_subpannosus
Euphorbiaceae_Croton_alagoensis
Euphorbiaceae_Croton_lindheimeri
Euphorbiaceae_Croton_heptalon
Euphorbiaceae_Croton_leucophyllus
Euphorbiaceae_Croton_lindheimerianus
Euphorbiaceae_Croton_pottsii

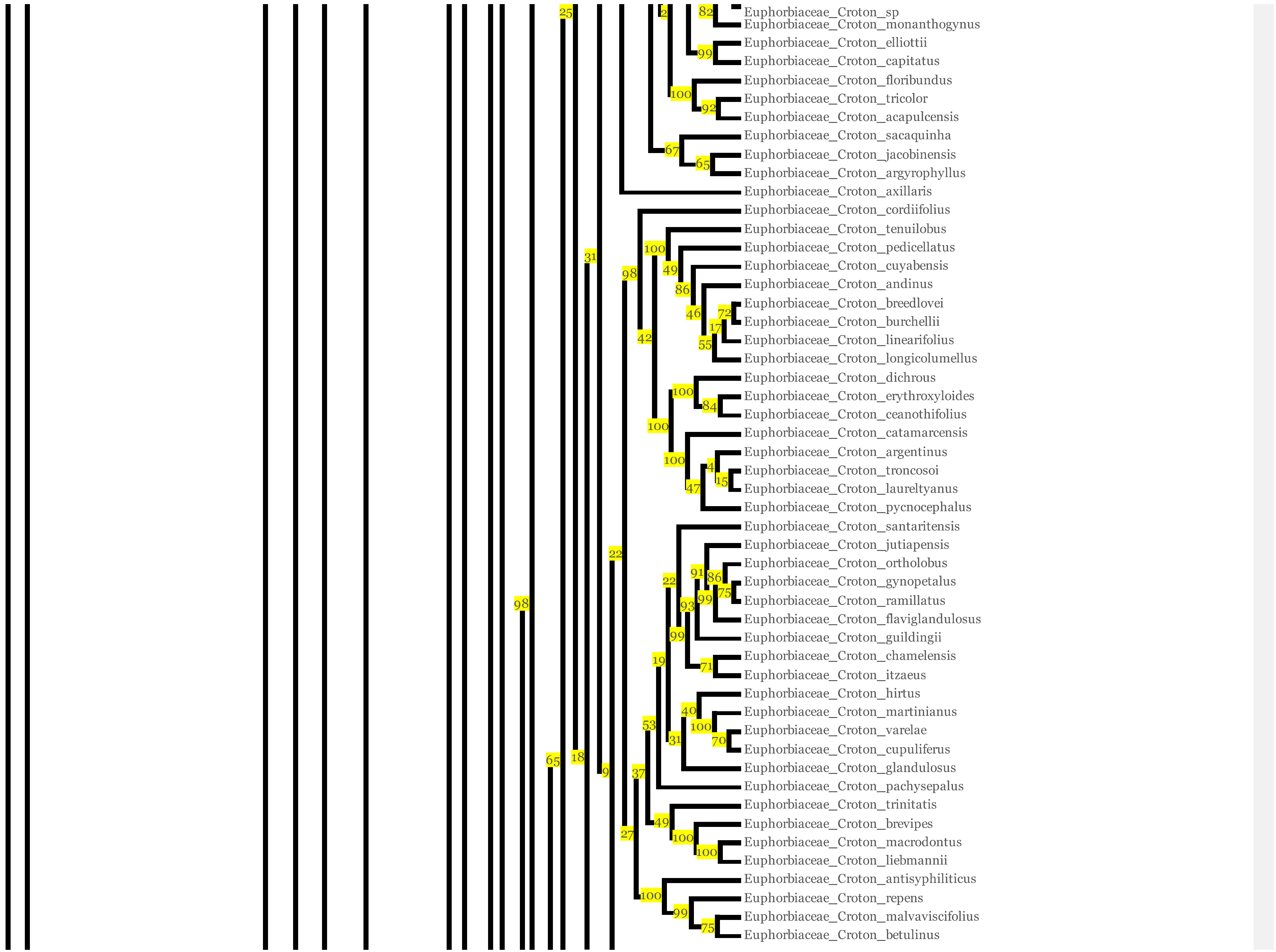
Euphorbiaceae_Croton_sp
Euphorbiaceae_Croton_monanthogynus
Euphorbiaceae_Croton_elliottii
Euphorbiaceae_Croton_capitatus
Euphorbiaceae_Croton_floribundus
Euphorbiaceae_Croton_tricolor
Euphorbiaceae_Croton_acapulcensis
Euphorbiaceae_Croton_sacaquinha
Euphorbiaceae_Croton_jacobinensis
Euphorbiaceae_Croton_argyrophyllus
Euphorbiaceae_Croton_axillaris
Euphorbiaceae_Croton_cordiifolius
Euphorbiaceae_Croton_tenuilobus
Euphorbiaceae_Croton_pedicellatus
Euphorbiaceae_Croton_cuyabensis
Euphorbiaceae_Croton_andinus
Euphorbiaceae_Croton_breedlovei
Euphorbiaceae_Croton_burchellii
Euphorbiaceae_Croton_linearifolius
Euphorbiaceae_Croton_longicolumellus
Euphorbiaceae_Croton_dichrous
Euphorbiaceae_Croton_erythroxyloides
Euphorbiaceae_Croton_ceanothifolius
Euphorbiaceae_Croton_catamarcensis
Euphorbiaceae_Croton_argentinus
Euphorbiaceae_Croton_troncosoi
Euphorbiaceae_Croton_laureltyanus
Euphorbiaceae_Croton_pycnocephalus
Euphorbiaceae_Croton_santaritensis
Euphorbiaceae_Croton_jutiapensis
Euphorbiaceae_Croton_ortholobus
Euphorbiaceae_Croton_gynopetalus
Euphorbiaceae_Croton_ramillatus
Euphorbiaceae_Croton_flaviglandulosus
Euphorbiaceae_Croton_guildingii
Euphorbiaceae_Croton_chamelensis
Euphorbiaceae_Croton_itzaeus
Euphorbiaceae_Croton_hirtus
Euphorbiaceae_Croton_martinianus
Euphorbiaceae_Croton_varelae
Euphorbiaceae_Croton_cupuliferus
Euphorbiaceae_Croton_glandulosus
Euphorbiaceae_Croton_pachysepalus
Euphorbiaceae_Croton_trinitatis
Euphorbiaceae_Croton_brevipes
Euphorbiaceae_Croton_macrodontus
Euphorbiaceae_Croton_liebmannii
Euphorbiaceae_Croton_antisyphiliticus
Euphorbiaceae_Croton_repens
Euphorbiaceae_Croton_malvaviscifolius
Euphorbiaceae_Croton_betulinus

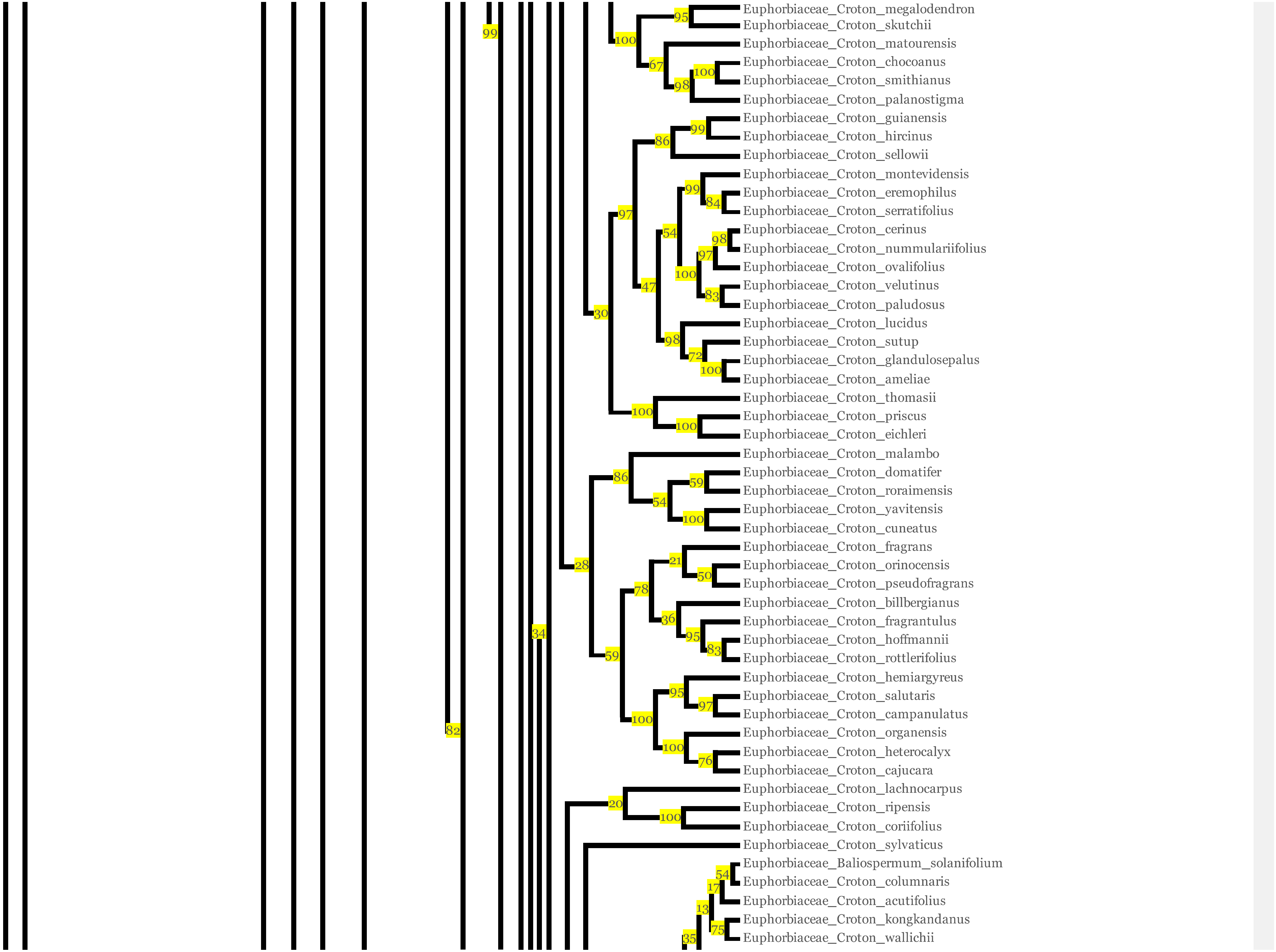
Euphorbiaceae_Croton_megalodendron
Euphorbiaceae_Croton_skutchii
Euphorbiaceae_Croton_matourensis
Euphorbiaceae_Croton_chocoanus
Euphorbiaceae_Croton_smithianus
Euphorbiaceae_Croton_palanostigma
Euphorbiaceae_Croton_guianensis
Euphorbiaceae_Croton_hircinus
Euphorbiaceae_Croton_sellowii
Euphorbiaceae_Croton_montevidensis
Euphorbiaceae_Croton_eremophilus
Euphorbiaceae_Croton_serratifolius
Euphorbiaceae_Croton_cerinus
Euphorbiaceae_Croton_nummulariifolius
Euphorbiaceae_Croton_ovalifolius
Euphorbiaceae_Croton_velutinus
Euphorbiaceae_Croton_paludosus
Euphorbiaceae_Croton_lucidus
Euphorbiaceae_Croton_sutup
Euphorbiaceae_Croton_glandulosepalus
Euphorbiaceae_Croton_ameliae
Euphorbiaceae_Croton_thomasii
Euphorbiaceae_Croton_priscus
Euphorbiaceae_Croton_eichleri
Euphorbiaceae_Croton_malambo
Euphorbiaceae_Croton_domatifer
Euphorbiaceae_Croton_roraimensis
Euphorbiaceae_Croton_yavitensis
Euphorbiaceae_Croton_cuneatus
Euphorbiaceae_Croton_fragrans
Euphorbiaceae_Croton_orinocensis
Euphorbiaceae_Croton_pseudofragrans
Euphorbiaceae_Croton_billbergianus
Euphorbiaceae_Croton_fragrantulus
Euphorbiaceae_Croton_hoffmannii
Euphorbiaceae_Croton_rottlerifolius
Euphorbiaceae_Croton_hemiargyreus
Euphorbiaceae_Croton_salutaris
Euphorbiaceae_Croton_campanulatus
Euphorbiaceae_Croton_organensis
Euphorbiaceae_Croton_heterocalyx
Euphorbiaceae_Croton_cajucara
Euphorbiaceae_Croton_lachnocarpus
Euphorbiaceae_Croton_ripensis
Euphorbiaceae_Croton_coriifolius
Euphorbiaceae_Croton_sylvaticus
Euphorbiaceae_Baliospermum_solanifolium
Euphorbiaceae_Croton_columnaris
Euphorbiaceae_Croton_acutifolius
Euphorbiaceae_Croton_kongkandanus
Euphorbiaceae_Croton_wallichii

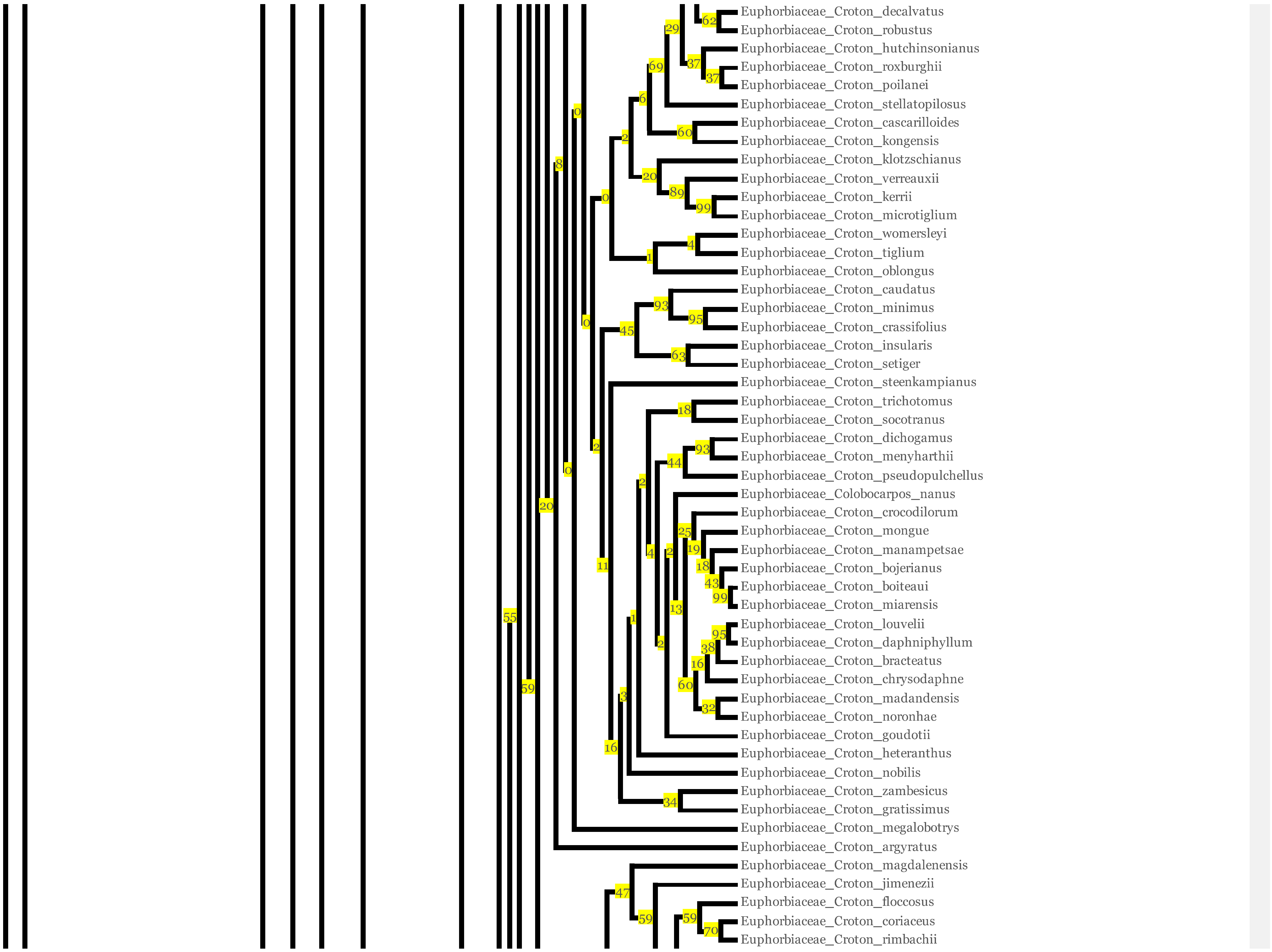
Euphorbiaceae_Croton_decalvatus
Euphorbiaceae_Croton_robustus
Euphorbiaceae_Croton_hutchinsonianus
Euphorbiaceae_Croton_roxburghii
Euphorbiaceae_Croton_poilanei
Euphorbiaceae_Croton_stellatopilosus
Euphorbiaceae_Croton_cascarilloides
Euphorbiaceae_Croton_kongensis
Euphorbiaceae_Croton_klotzschianus
Euphorbiaceae_Croton_verreauxii
Euphorbiaceae_Croton_kerrii
Euphorbiaceae_Croton_microtiglium
Euphorbiaceae_Croton_womersleyi
Euphorbiaceae_Croton_tiglium
Euphorbiaceae_Croton_oblongus
Euphorbiaceae_Croton_caudatus
Euphorbiaceae_Croton_minimus
Euphorbiaceae_Croton_crassifolius
Euphorbiaceae_Croton_insularis
Euphorbiaceae_Croton_setiger
Euphorbiaceae_Croton_steenkampianus
Euphorbiaceae_Croton_trichotomus
Euphorbiaceae_Croton_socotranus
Euphorbiaceae_Croton_dichogamus
Euphorbiaceae_Croton_menyharthii
Euphorbiaceae_Croton_pseudopulchellus
Euphorbiaceae_Colobocarpos_nanus
Euphorbiaceae_Croton_crocodilorum
Euphorbiaceae_Croton_mongue
Euphorbiaceae_Croton_manampetsae
Euphorbiaceae_Croton_bojerianus
Euphorbiaceae_Croton_boiteaui
Euphorbiaceae_Croton_miarensis
Euphorbiaceae_Croton_louvelii
Euphorbiaceae_Croton_daphniphyllum
Euphorbiaceae_Croton_bracteatus
Euphorbiaceae_Croton_chrysodaphne
Euphorbiaceae_Croton_madandensis
Euphorbiaceae_Croton_noronhae
Euphorbiaceae_Croton_goudotii
Euphorbiaceae_Croton_heteranthus
Euphorbiaceae_Croton_nobilis
Euphorbiaceae_Croton_zambesicus
Euphorbiaceae_Croton_gratissimus
Euphorbiaceae_Croton_megalobotrys
Euphorbiaceae_Croton_argyratus
Euphorbiaceae_Croton_magdalenensis
Euphorbiaceae_Croton_jimenezii
Euphorbiaceae_Croton_floccosus
Euphorbiaceae_Croton_coriaceus
Euphorbiaceae_Croton_rimbachii

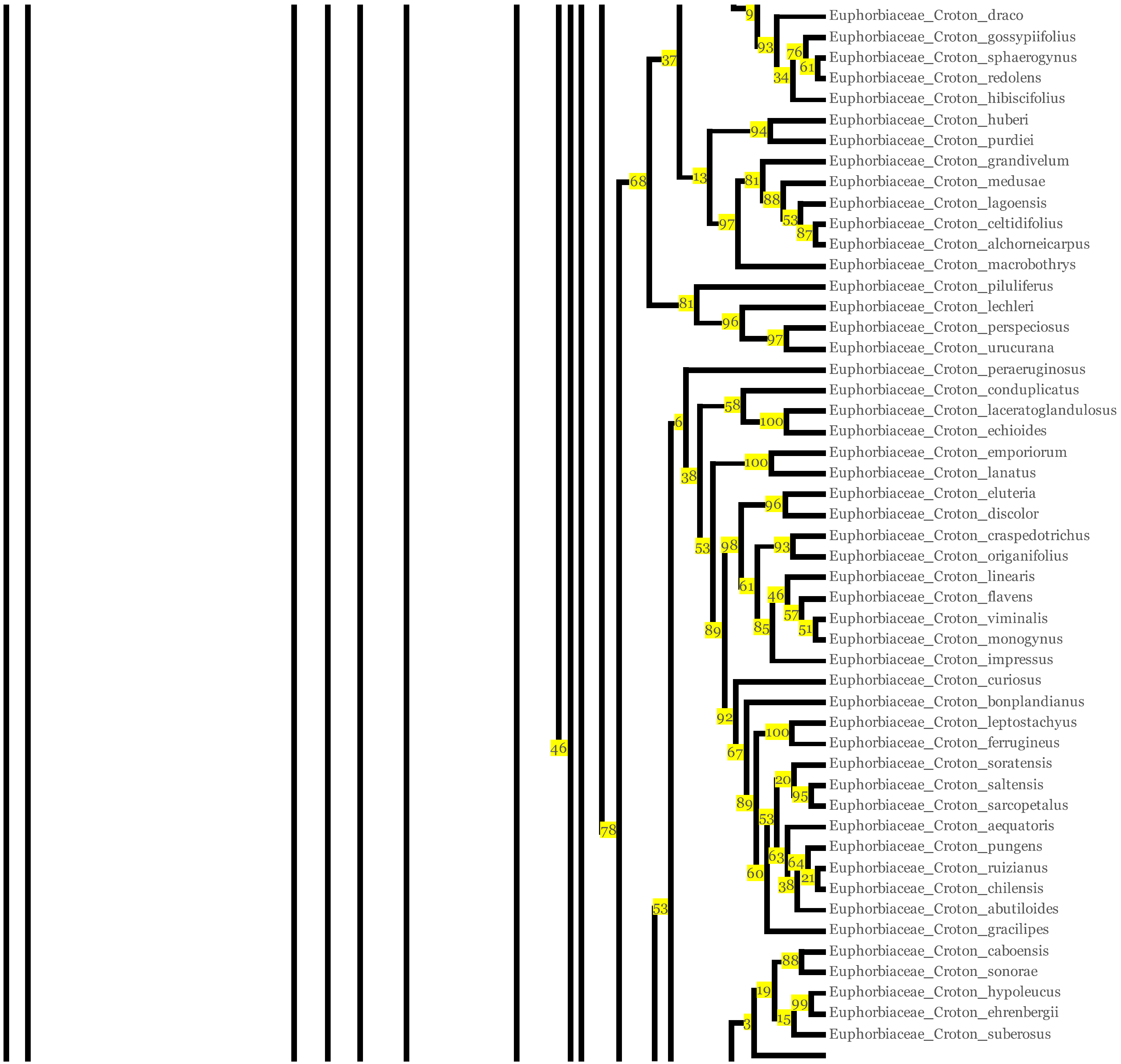
Euphorbiaceae_Croton_draco
Euphorbiaceae_Croton_gossypiifolius
Euphorbiaceae_Croton_sphaerogynus
Euphorbiaceae_Croton_redolens
Euphorbiaceae_Croton_hibiscifolius
Euphorbiaceae_Croton_huberi
Euphorbiaceae_Croton_purdiei
Euphorbiaceae_Croton_grandivelum
Euphorbiaceae_Croton_medusae
Euphorbiaceae_Croton_lagoensis
Euphorbiaceae_Croton_celtidifolius
Euphorbiaceae_Croton_alchorneicarpus
Euphorbiaceae_Croton_macrobothrys
Euphorbiaceae_Croton_piluliferus
Euphorbiaceae_Croton_lechleri
Euphorbiaceae_Croton_perspeciosus
Euphorbiaceae_Croton_urucurana
Euphorbiaceae_Croton_peraeruginosus
Euphorbiaceae_Croton_conduplicatus
Euphorbiaceae_Croton_laceratoglandulosus
Euphorbiaceae_Croton_echioides
Euphorbiaceae_Croton_emporiorum
Euphorbiaceae_Croton_lanatus
Euphorbiaceae_Croton_eluteria
Euphorbiaceae_Croton_discolor
Euphorbiaceae_Croton_craspedotrichus
Euphorbiaceae_Croton_origanifolius
Euphorbiaceae_Croton_linearis
Euphorbiaceae_Croton_flavens
Euphorbiaceae_Croton_viminalis
Euphorbiaceae_Croton_monogynus
Euphorbiaceae_Croton_impressus
Euphorbiaceae_Croton_curiosus
Euphorbiaceae_Croton_bonplandianus
Euphorbiaceae_Croton_leptostachyus
Euphorbiaceae_Croton_ferrugineus
Euphorbiaceae_Croton_soratensis
Euphorbiaceae_Croton_saltensis
Euphorbiaceae_Croton_sarcopetalus
Euphorbiaceae_Croton_aequatoris
Euphorbiaceae_Croton_pungens
Euphorbiaceae_Croton_ruizianus
Euphorbiaceae_Croton_chilensis
Euphorbiaceae_Croton_abutiloides
Euphorbiaceae_Croton_gracilipes
Euphorbiaceae_Croton_caboensis
Euphorbiaceae_Croton_sonorae
Euphorbiaceae_Croton_hypoleucus
Euphorbiaceae_Croton_ehrenbergii
Euphorbiaceae_Croton_suberosus
9
93
76
61
34
37
94
13
81
88
53
87
97
68
81
96
97
6
58
100
100
38
96
53
98
93
61
46
57
51
85
89
92
100
67
20
95
89
53
63
64
21
38
60
46
78
53
88
19
99
3
15

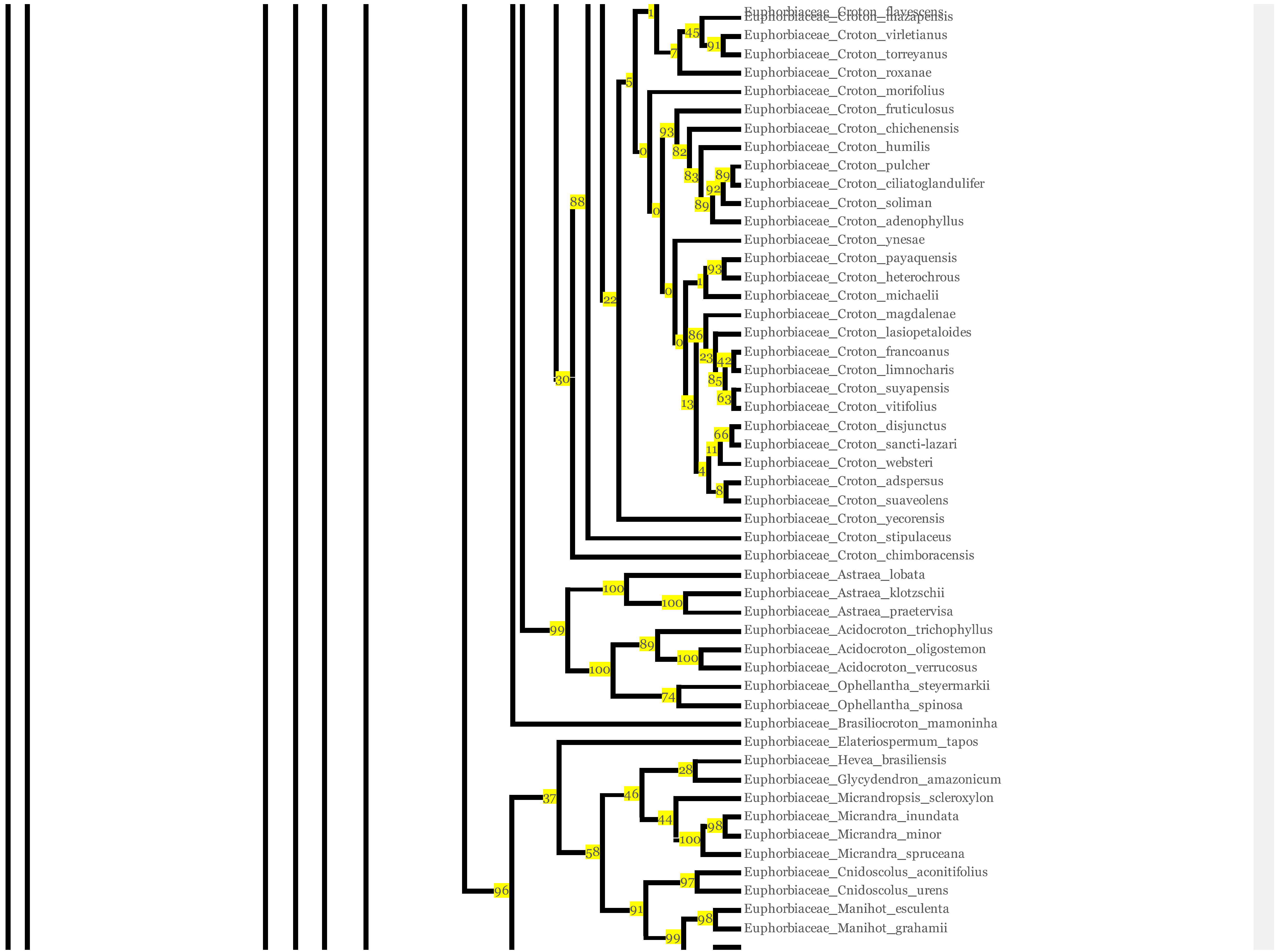

Euphorbiaceae_Croton_flavescens
Euphorbiaceae_Croton_mazapensis
Euphorbiaceae_Croton_virletianus
Euphorbiaceae_Croton_torreyanus
Euphorbiaceae_Croton_roxanae
Euphorbiaceae_Croton_morifolius
Euphorbiaceae_Croton_fruticulosus
Euphorbiaceae_Croton_chichenensis
Euphorbiaceae_Croton_humilis
Euphorbiaceae_Croton_pulcher
Euphorbiaceae_Croton_ciliatoglandulifer
Euphorbiaceae_Croton_soliman
Euphorbiaceae_Croton_adenophyllus
Euphorbiaceae_Croton_ynesae
Euphorbiaceae_Croton_payaquensis
Euphorbiaceae_Croton_heterochrous
Euphorbiaceae_Croton_michaelii
Euphorbiaceae_Croton_magdalenae
Euphorbiaceae_Croton_lasiopetaloides
Euphorbiaceae_Croton_francoanus
Euphorbiaceae_Croton_limnocharis
Euphorbiaceae_Croton_suyapensis
Euphorbiaceae_Croton_vitifolius
Euphorbiaceae_Croton_disjunctus
Euphorbiaceae_Croton_sancti-lazari
Euphorbiaceae_Croton_websteri
Euphorbiaceae_Croton_adspersus
Euphorbiaceae_Croton_suaveolens
Euphorbiaceae_Croton_yecorensis
Euphorbiaceae_Croton_stipulaceus
Euphorbiaceae_Croton_chimboracensis
Euphorbiaceae_Astraea_lobata
Euphorbiaceae_Astraea_klotzschii
Euphorbiaceae_Astraea_praetervisa
Euphorbiaceae_Acidocroton_trichophyllus
Euphorbiaceae_Acidocroton_oligostemon
Euphorbiaceae_Acidocroton_verrucosus
Euphorbiaceae_Ophellantha_steyermarkii
Euphorbiaceae_Ophellantha_spinosa
Euphorbiaceae_Brasiliocroton_mamoninha
Euphorbiaceae_Elateriospermum_tapos
Euphorbiaceae_Hevea_brasiliensis
Euphorbiaceae_Glycydendron_amazonicum
Euphorbiaceae_Micrandropsis_scleroxylon
Euphorbiaceae_Micrandra_inundata
Euphorbiaceae_Micrandra_minor
Euphorbiaceae_Micrandra_spruceana
Euphorbiaceae_Cnidoscolus_aconitifolius
Euphorbiaceae_Cnidoscolus_urens
Euphorbiaceae_Manihot_esculenta
Euphorbiaceae_Manihot_grahamii

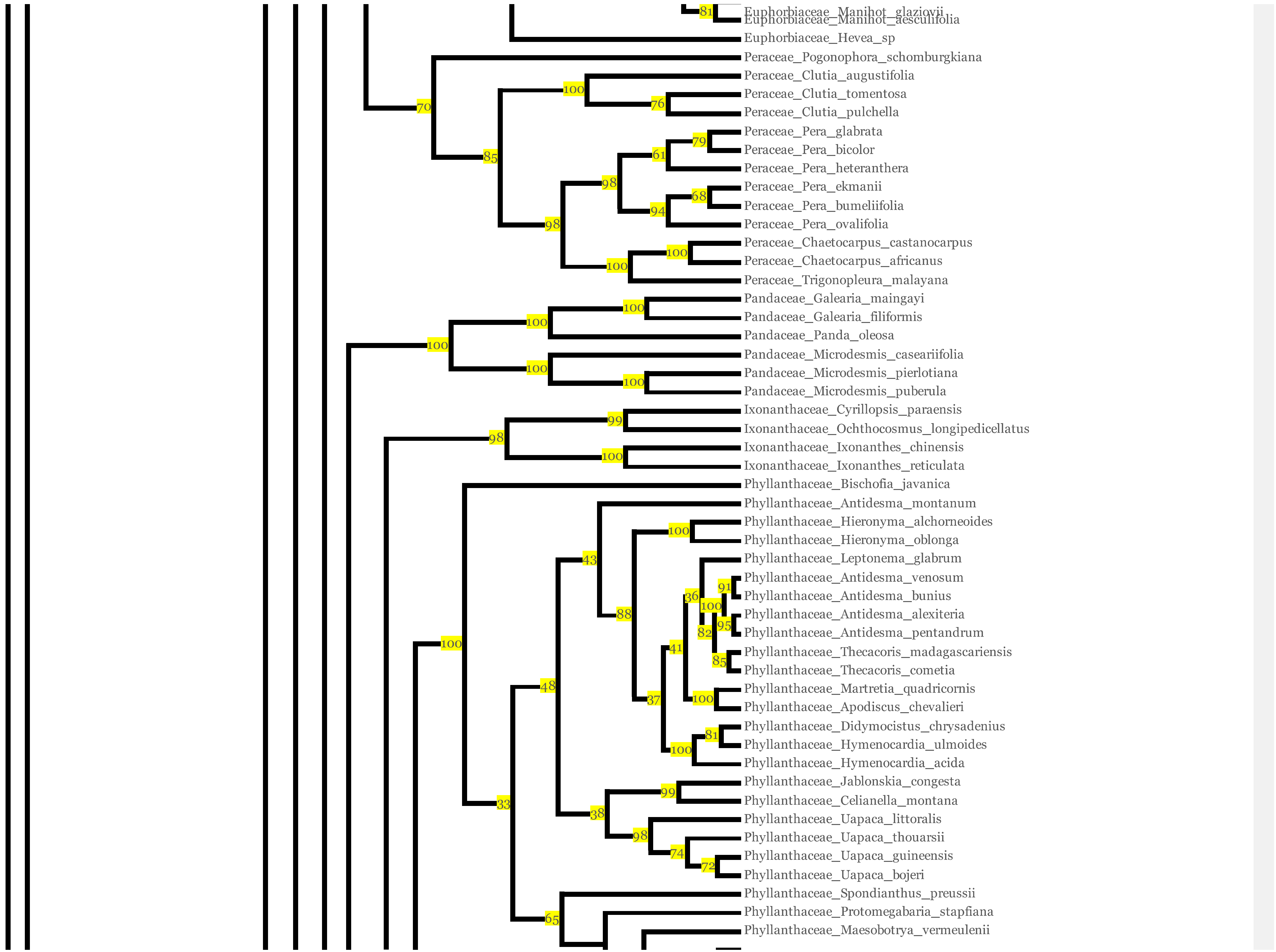
Euphorbiaceae_Manihot_glaziovii
Euphorbiaceae_Manihot_aesculifolia
Euphorbiaceae_Hevea_sp
Peraceae_Pogonophora_schomburgkiana
Peraceae_Clutia_augustifolia
Peraceae_Clutia_tomentosa
Peraceae_Clutia_pulchella
Peraceae_Pera_glabrata
Peraceae_Pera_bicolor
Peraceae_Pera_heteranthera
Peraceae_Pera_ekmanii
Peraceae_Pera_bumeliifolia
Peraceae_Pera_ovalifolia
Peraceae_Chaetocarpus_castanocarpus
Peraceae_Chaetocarpus_africanus
Peraceae_Trigonopleura_malayana
Pandaceae_Galearia_maingayi
Pandaceae_Galearia_filiformis
Pandaceae_Panda_oleosa
Pandaceae_Microdesmis_caseariifolia
Pandaceae_Microdesmis_pierlotiana
Pandaceae_Microdesmis_puberula
Ixonanthaceae_Cyrillopsis_paraensis
Ixonanthaceae_Ochthocosmus_longipedicellatus
Ixonanthaceae_Ixonanthes_chinensis
Ixonanthaceae_Ixonanthes_reticulata
Phyllanthaceae_Bischofia_javanica
Phyllanthaceae_Antidesma_montanum
Phyllanthaceae_Hieronyma_alchorneoides
Phyllanthaceae_Hieronyma_oblonga
Phyllanthaceae_Leptonema_glabrum
Phyllanthaceae_Antidesma_venosum
Phyllanthaceae_Antidesma_bunius
Phyllanthaceae_Antidesma_alexiteria
Phyllanthaceae_Antidesma_pentandrum
Phyllanthaceae_Thecacoris_madagascariensis
Phyllanthaceae_Thecacoris_cometia
Phyllanthaceae_Martretia_quadricornis
Phyllanthaceae_Apodiscus_chevalieri
Phyllanthaceae_Didymocistus_chrysadenius
Phyllanthaceae_Hymenocardia_ulmoides
Phyllanthaceae_Hymenocardia_acida
Phyllanthaceae_Jablonskia_congesta
Phyllanthaceae_Celianella_montana
Phyllanthaceae_Uapaca_littoralis
Phyllanthaceae_Uapaca_thouarsii
Phyllanthaceae_Uapaca_guineensis
Phyllanthaceae_Uapaca_bojeri
Phyllanthaceae_Spondianthus_preussii
Phyllanthaceae_Protomegabaria_stapfiana
Phyllanthaceae_Maesobotrya_vermeulenii

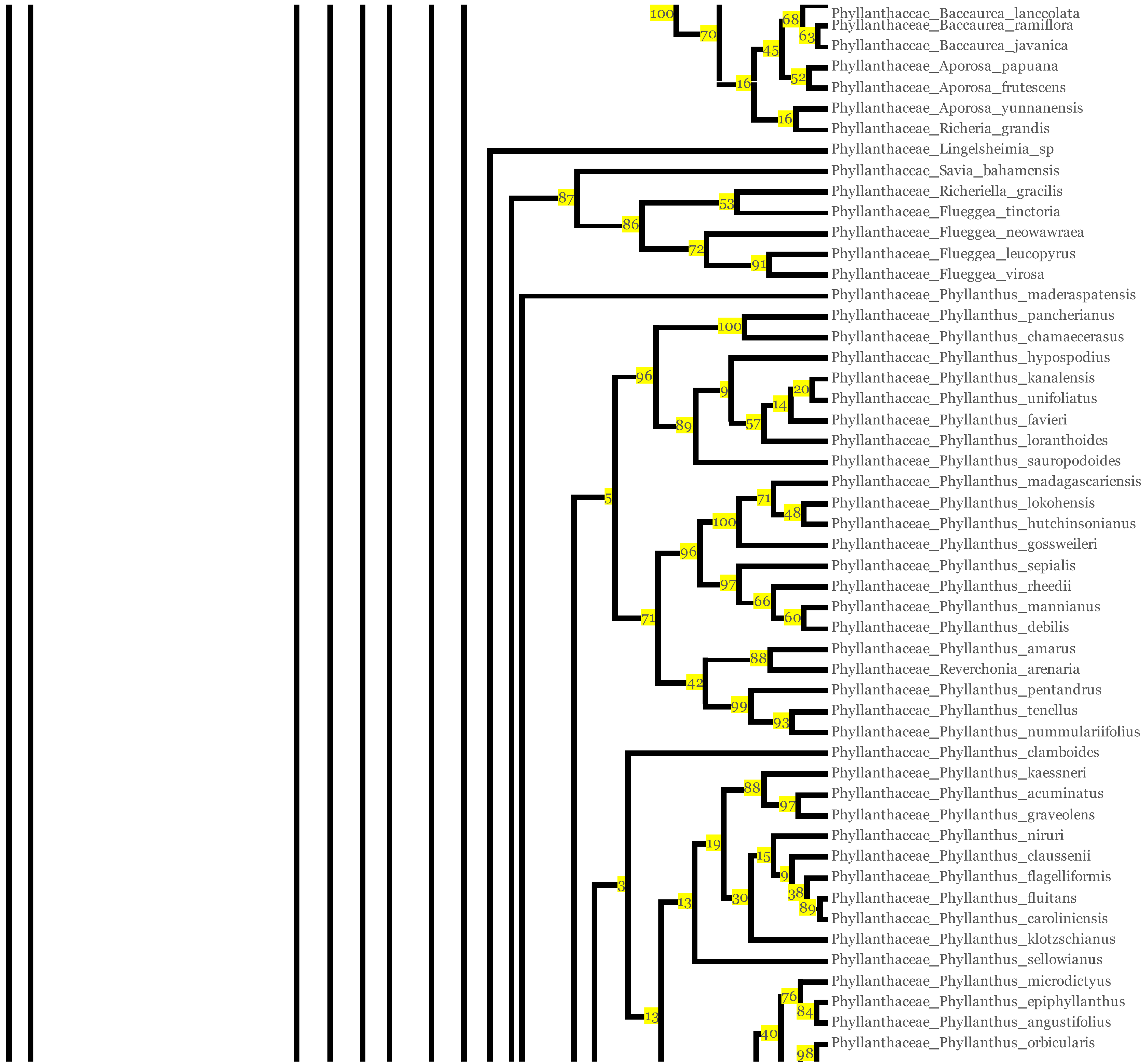
Phyllanthaceae_Baccaurea_lanceolata
Phyllanthaceae_Baccaurea_ramiflora
Phyllanthaceae_Baccaurea_javanica
Phyllanthaceae_Aporosa_papuana
Phyllanthaceae_Aporosa_frutescens
Phyllanthaceae_Aporosa_yunnanensis
Phyllanthaceae_Richeria_grandis
Phyllanthaceae_Lingelsheimia_sp
Phyllanthaceae_Savia_bahamensis
Phyllanthaceae_Richeriella_gracilis
Phyllanthaceae_Flueggea_tinctoria
Phyllanthaceae_Flueggea_neowawraea
Phyllanthaceae_Flueggea_leucopyrus
Phyllanthaceae_Flueggea_virosa
Phyllanthaceae_Phyllanthus_maderaspatensis
Phyllanthaceae_Phyllanthus_pancherianus
Phyllanthaceae_Phyllanthus_chamaecerasus
Phyllanthaceae_Phyllanthus_hypospodius
Phyllanthaceae_Phyllanthus_kanalensis
Phyllanthaceae_Phyllanthus_unifoliatus
Phyllanthaceae_Phyllanthus_favieri
Phyllanthaceae_Phyllanthus_loranthoides
Phyllanthaceae_Phyllanthus_sauropodoides
Phyllanthaceae_Phyllanthus_madagascariensis
Phyllanthaceae_Phyllanthus_lokohensis
Phyllanthaceae_Phyllanthus_hutchinsonianus
Phyllanthaceae_Phyllanthus_gossweileri
Phyllanthaceae_Phyllanthus_sepialis
Phyllanthaceae_Phyllanthus_rheedii
Phyllanthaceae_Phyllanthus_mannianus
Phyllanthaceae_Phyllanthus_debilis
Phyllanthaceae_Phyllanthus_amarus
Phyllanthaceae_Reverchonia_arenaria
Phyllanthaceae_Phyllanthus_pentandrus
Phyllanthaceae_Phyllanthus_tenellus
Phyllanthaceae_Phyllanthus_nummulariifolius
Phyllanthaceae_Phyllanthus_clamboides
Phyllanthaceae_Phyllanthus_kaessneri
Phyllanthaceae_Phyllanthus_acuminatus
Phyllanthaceae_Phyllanthus_graveolens
Phyllanthaceae_Phyllanthus_niruri
Phyllanthaceae_Phyllanthus_claussenii
Phyllanthaceae_Phyllanthus_flagelliformis
Phyllanthaceae_Phyllanthus_fluitans
Phyllanthaceae_Phyllanthus_caroliniensis
Phyllanthaceae_Phyllanthus_klotzschianus
Phyllanthaceae_Phyllanthus_sellowianus
Phyllanthaceae_Phyllanthus_microdictyus
Phyllanthaceae_Phyllanthus_epiphyllanthus
Phyllanthaceae_Phyllanthus_angustifolius
Phyllanthaceae_Phyllanthus_orbicularis

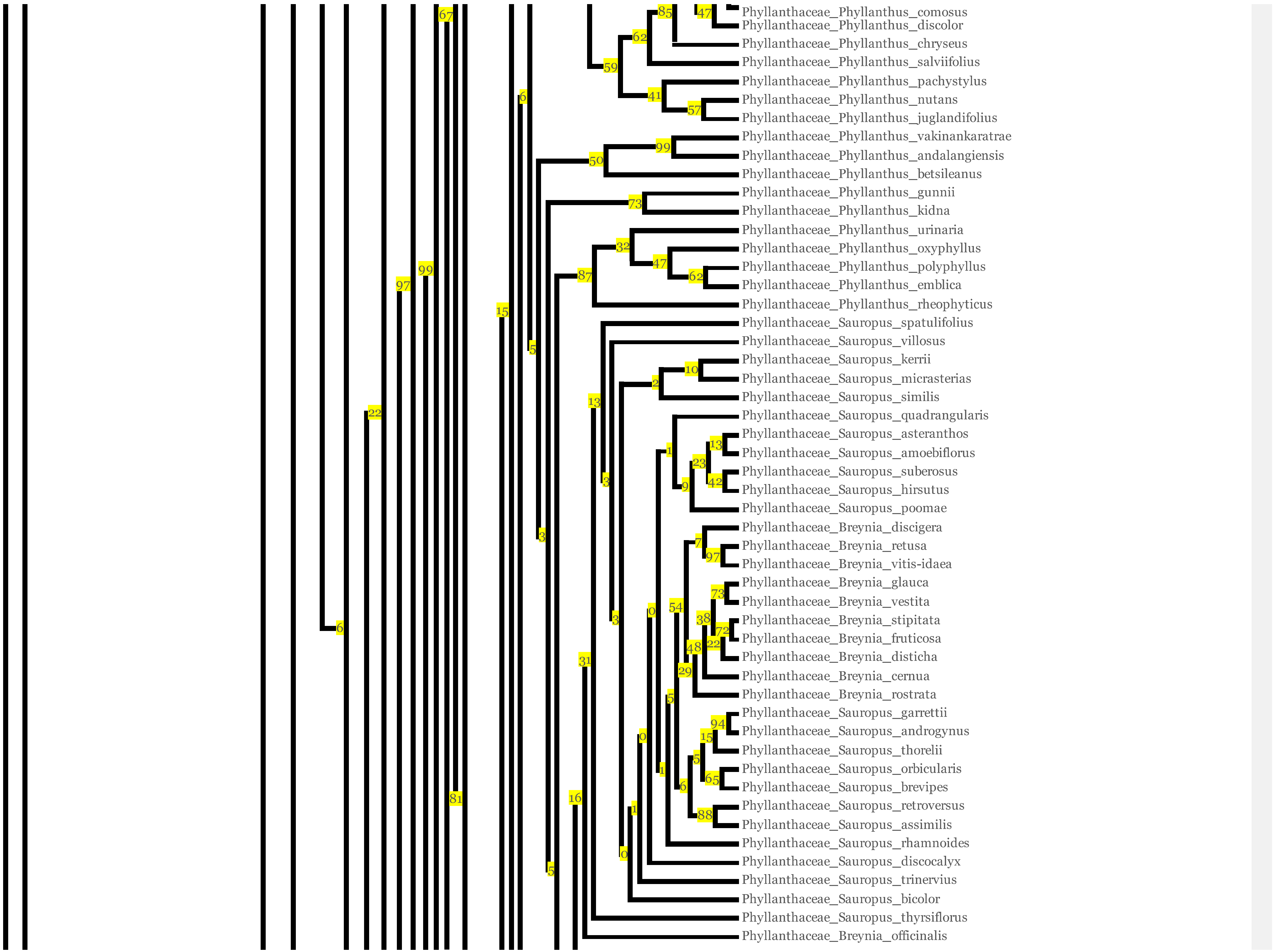
Phyllanthaceae_Phyllanthus_comosus
Phyllanthaceae_Phyllanthus_discolor
Phyllanthaceae_Phyllanthus_chryseus
Phyllanthaceae_Phyllanthus_salviifolius
Phyllanthaceae_Phyllanthus_pachystylus
Phyllanthaceae_Phyllanthus_nutans
Phyllanthaceae_Phyllanthus_juglandifolius
Phyllanthaceae_Phyllanthus_vakinankaratrae
Phyllanthaceae_Phyllanthus_andalangiensis
Phyllanthaceae_Phyllanthus_betsileanus
Phyllanthaceae_Phyllanthus_gunnii
Phyllanthaceae_Phyllanthus_kidna
Phyllanthaceae_Phyllanthus_urinaria
Phyllanthaceae_Phyllanthus_oxyphyllus
Phyllanthaceae_Phyllanthus_polyphyllus
Phyllanthaceae_Phyllanthus_emblica
Phyllanthaceae_Phyllanthus_rheophyticus
Phyllanthaceae_Sauropus_spatulifolius
Phyllanthaceae_Sauropus_villosus
Phyllanthaceae_Sauropus_kerrii
Phyllanthaceae_Sauropus_micrasterias
Phyllanthaceae_Sauropus_similis
Phyllanthaceae_Sauropus_quadrangularis
Phyllanthaceae_Sauropus_asteranthos
Phyllanthaceae_Sauropus_amoebiflorus
Phyllanthaceae_Sauropus_suberosus
Phyllanthaceae_Sauropus_hirsutus
Phyllanthaceae_Sauropus_poomae
Phyllanthaceae_Breynia_discigera
Phyllanthaceae_Breynia_retusa
Phyllanthaceae_Breynia_vitis-idaea
Phyllanthaceae_Breynia_glauca
Phyllanthaceae_Breynia_vestita
Phyllanthaceae_Breynia_stipitata
Phyllanthaceae_Breynia_fruticosa
Phyllanthaceae_Breynia_disticha
Phyllanthaceae_Breynia_cernua
Phyllanthaceae_Breynia_rostrata
Phyllanthaceae_Sauropus_garrettii
Phyllanthaceae_Sauropus_androgynus
Phyllanthaceae_Sauropus_thorelii
Phyllanthaceae_Sauropus_orbicularis
Phyllanthaceae_Sauropus_brevipes
Phyllanthaceae_Sauropus_retroversus
Phyllanthaceae_Sauropus_assimilis
Phyllanthaceae_Sauropus_rhamnoides
Phyllanthaceae_Sauropus_discocalyx
Phyllanthaceae_Sauropus_trinervius
Phyllanthaceae_Sauropus_bicolor
Phyllanthaceae_Sauropus_thyrsiflorus
Phyllanthaceae_Breynia_officinalis

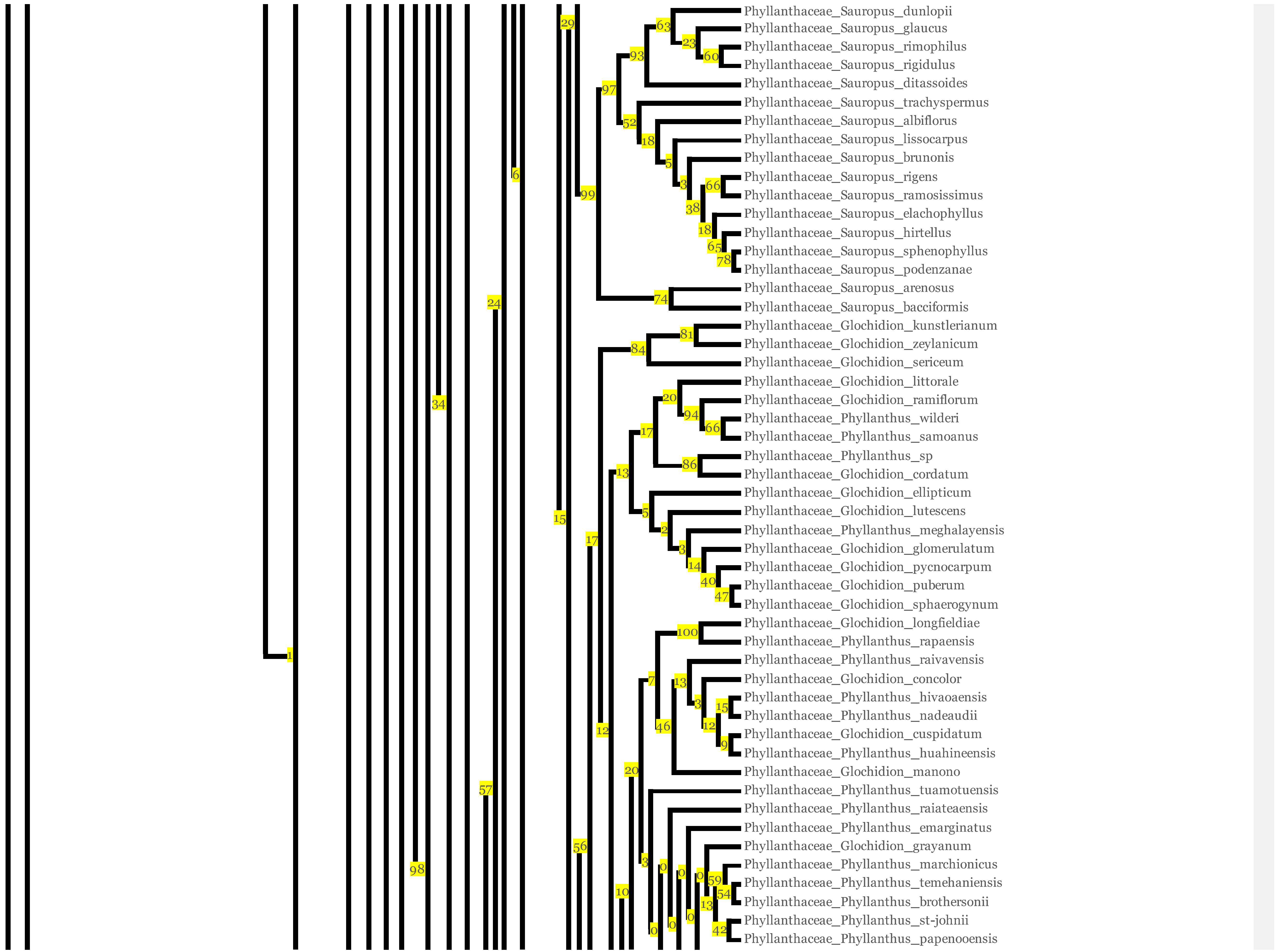
Phyllanthaceae_Sauropus_dunlopii
Phyllanthaceae_Sauropus_glaucus
Phyllanthaceae_Sauropus_rimophilus
Phyllanthaceae_Sauropus_rigidulus
Phyllanthaceae_Sauropus_ditassoides
Phyllanthaceae_Sauropus_trachyspermus
Phyllanthaceae_Sauropus_albiflorus
Phyllanthaceae_Sauropus_lissocarpus
Phyllanthaceae_Sauropus_brunonis
Phyllanthaceae_Sauropus_rigens
Phyllanthaceae_Sauropus_ramosissimus
Phyllanthaceae_Sauropus_elachophyllus
Phyllanthaceae_Sauropus_hirtellus
Phyllanthaceae_Sauropus_sphenophyllus
Phyllanthaceae_Sauropus_podenzanae
Phyllanthaceae_Sauropus_arenosus
Phyllanthaceae_Sauropus_bacciformis
Phyllanthaceae_Glochidion_kunstlerianum
Phyllanthaceae_Glochidion_zeylanicum
Phyllanthaceae_Glochidion_sericeum
Phyllanthaceae_Glochidion_littorale
Phyllanthaceae_Glochidion_ramiflorum
Phyllanthaceae_Phyllanthus_wilderi
Phyllanthaceae_Phyllanthus_samoanus
Phyllanthaceae_Phyllanthus_sp
Phyllanthaceae_Glochidion_cordatum
Phyllanthaceae_Glochidion_ellipticum
Phyllanthaceae_Glochidion_lutescens
Phyllanthaceae_Phyllanthus_meghalayensis
Phyllanthaceae_Glochidion_glomerulatum
Phyllanthaceae_Glochidion_pycnocarpum
Phyllanthaceae_Glochidion_puberum
Phyllanthaceae_Glochidion_sphaerogynum
Phyllanthaceae_Glochidion_longfieldiae
Phyllanthaceae_Phyllanthus_rapaensis
Phyllanthaceae_Phyllanthus_raivavensis
Phyllanthaceae_Glochidion_concolor
Phyllanthaceae_Phyllanthus_hivaoaensis
Phyllanthaceae_Phyllanthus_nadeaudii
Phyllanthaceae_Glochidion_cuspidatum
Phyllanthaceae_Phyllanthus_huahineensis
Phyllanthaceae_Glochidion_manono
Phyllanthaceae_Phyllanthus_tuamotuensis
Phyllanthaceae_Phyllanthus_raiateaensis
Phyllanthaceae_Phyllanthus_emarginatus
Phyllanthaceae_Glochidion_grayanum
Phyllanthaceae_Phyllanthus_marchionicus
Phyllanthaceae_Phyllanthus_temehaniensis
Phyllanthaceae_Phyllanthus_brothersonii
Phyllanthaceae_Phyllanthus_st-johnii
Phyllanthaceae_Phyllanthus_papenooensis

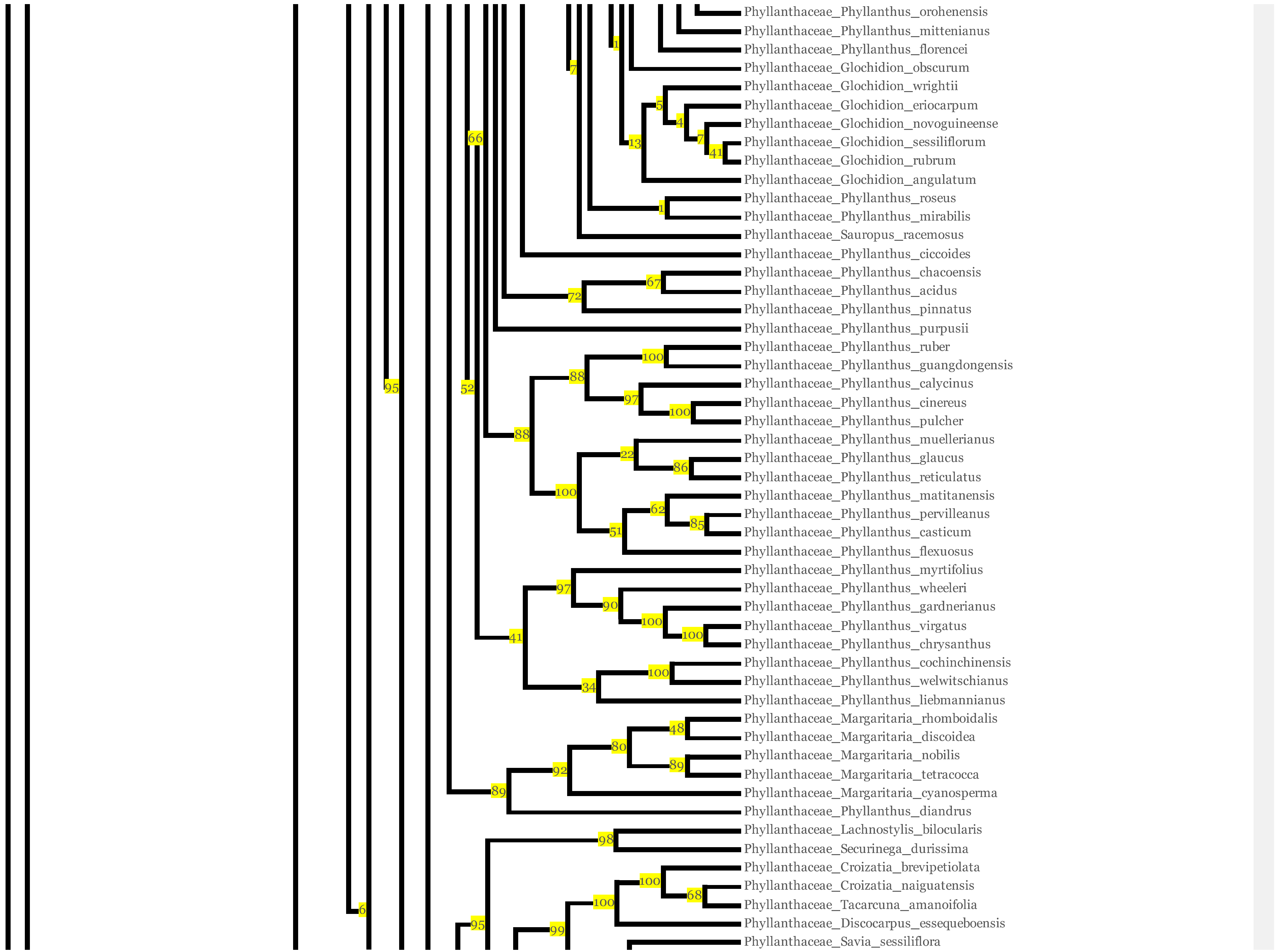
Phyllanthaceae_Phyllanthus_orohenensis
Phyllanthaceae_Phyllanthus_mittenianus
Phyllanthaceae_Phyllanthus_florencei
Phyllanthaceae_Glochidion_obscurum
Phyllanthaceae_Glochidion_wrightii
Phyllanthaceae_Glochidion_eriocarpum
Phyllanthaceae_Glochidion_novoguineense
Phyllanthaceae_Glochidion_sessiliflorum
Phyllanthaceae_Glochidion_rubrum
Phyllanthaceae_Glochidion_angulatum
Phyllanthaceae_Phyllanthus_roseus
Phyllanthaceae_Phyllanthus_mirabilis
Phyllanthaceae_Sauropus_racemosus
Phyllanthaceae_Phyllanthus_ciccoides
Phyllanthaceae_Phyllanthus_chacoensis
Phyllanthaceae_Phyllanthus_acidus
Phyllanthaceae_Phyllanthus_pinnatus
Phyllanthaceae_Phyllanthus_purpusii
Phyllanthaceae_Phyllanthus_ruber
Phyllanthaceae_Phyllanthus_guangdongensis
Phyllanthaceae_Phyllanthus_calycinus
Phyllanthaceae_Phyllanthus_cinereus
Phyllanthaceae_Phyllanthus_pulcher
Phyllanthaceae_Phyllanthus_muellerianus
Phyllanthaceae_Phyllanthus_glaucus
Phyllanthaceae_Phyllanthus_reticulatus
Phyllanthaceae_Phyllanthus_matitanensis
Phyllanthaceae_Phyllanthus_pervilleanus
Phyllanthaceae_Phyllanthus_casticum
Phyllanthaceae_Phyllanthus_flexuosus
Phyllanthaceae_Phyllanthus_myrtifolius
Phyllanthaceae_Phyllanthus_wheeleri
Phyllanthaceae_Phyllanthus_gardnerianus
Phyllanthaceae_Phyllanthus_virgatus
Phyllanthaceae_Phyllanthus_chrysanthus
Phyllanthaceae_Phyllanthus_cochinchinensis
Phyllanthaceae_Phyllanthus_welwitschianus
Phyllanthaceae_Phyllanthus_liebmannianus
Phyllanthaceae_Margaritaria_rhomboidalis
Phyllanthaceae_Margaritaria_discoidea
Phyllanthaceae_Margaritaria_nobilis
Phyllanthaceae_Margaritaria_tetracocca
Phyllanthaceae_Margaritaria_cyanosperma
Phyllanthaceae_Phyllanthus_diandrus
Phyllanthaceae_Lachnostylis_bilocularis
Phyllanthaceae_Securinega_durissima
Phyllanthaceae_Croizatia_brevipetiolata
Phyllanthaceae_Croizatia_naiguatensis
Phyllanthaceae_Tacarcuna_amanoifolia
Phyllanthaceae_Discocarpus_essequeboensis
Phyllanthaceae_Savia_sessiliflora

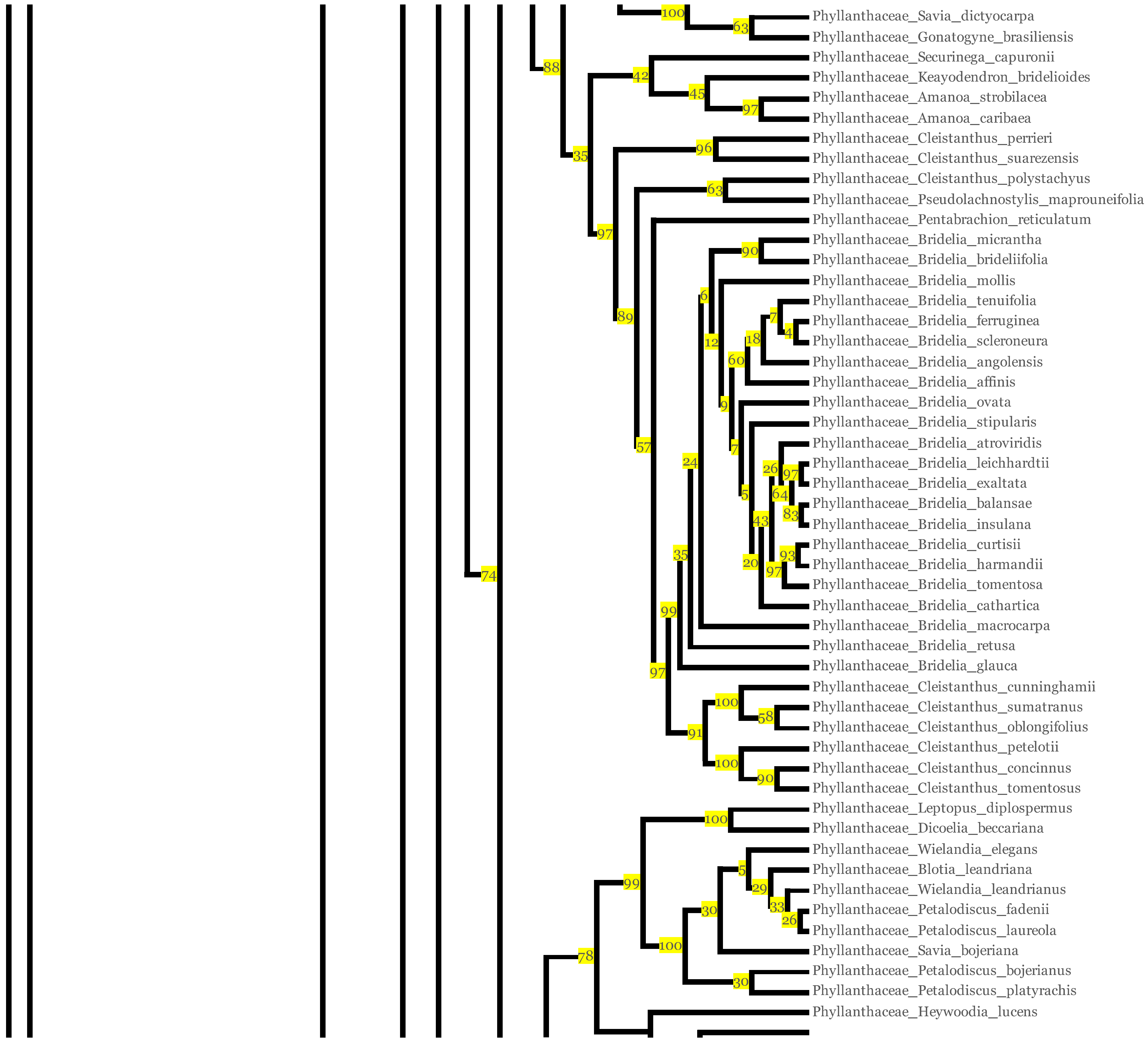
Phyllanthaceae_Savia_dictyocarpa
Phyllanthaceae_Gonatogyne_brasiliensis
Phyllanthaceae_Securinega_capuronii
Phyllanthaceae_Keayodendron_bridelioides
Phyllanthaceae_Amanoa_strobilacea
Phyllanthaceae_Amanoa_caribaea
Phyllanthaceae_Cleistanthus_perrieri
Phyllanthaceae_Cleistanthus_suarezensis
Phyllanthaceae_Cleistanthus_polystachyus
Phyllanthaceae_Pseudolachnostylis_maprouneifolia
Phyllanthaceae_Pentabrachion_reticulatum
Phyllanthaceae_Bridelia_micrantha
Phyllanthaceae_Bridelia_brideliifolia
Phyllanthaceae_Bridelia_mollis
Phyllanthaceae_Bridelia_tenuifolia
Phyllanthaceae_Bridelia_ferruginea
Phyllanthaceae_Bridelia_scleroneura
Phyllanthaceae_Bridelia_angolensis
Phyllanthaceae_Bridelia_affinis
Phyllanthaceae_Bridelia_ovata
Phyllanthaceae_Bridelia_stipularis
Phyllanthaceae_Bridelia_atroviridis
Phyllanthaceae_Bridelia_leichhardtii
Phyllanthaceae_Bridelia_exaltata
Phyllanthaceae_Bridelia_balansae
Phyllanthaceae_Bridelia_insulana
Phyllanthaceae_Bridelia_curtisii
Phyllanthaceae_Bridelia_harmandii
Phyllanthaceae_Bridelia_tomentosa
Phyllanthaceae_Bridelia_cathartica
Phyllanthaceae_Bridelia_macrocarpa
Phyllanthaceae_Bridelia_retusa
Phyllanthaceae_Bridelia_glauca
Phyllanthaceae_Cleistanthus_cunninghamii
Phyllanthaceae_Cleistanthus_sumatranus
Phyllanthaceae_Cleistanthus_oblongifolius
Phyllanthaceae_Cleistanthus_petelotii
Phyllanthaceae_Cleistanthus_concinnus
Phyllanthaceae_Cleistanthus_tomentosus
Phyllanthaceae_Leptopus_diplospermus
Phyllanthaceae_Dicoelia_beccariana
Phyllanthaceae_Wielandia_elegans
Phyllanthaceae_Blotia_leandriana
Phyllanthaceae_Wielandia_leandrianus
Phyllanthaceae_Petalodiscus_fadenii
Phyllanthaceae_Petalodiscus_laureola
Phyllanthaceae_Savia_bojeriana
Phyllanthaceae_Petalodiscus_bojerianus
Phyllanthaceae_Petalodiscus_platyrachis
Phyllanthaceae_Heywoodia_lucens

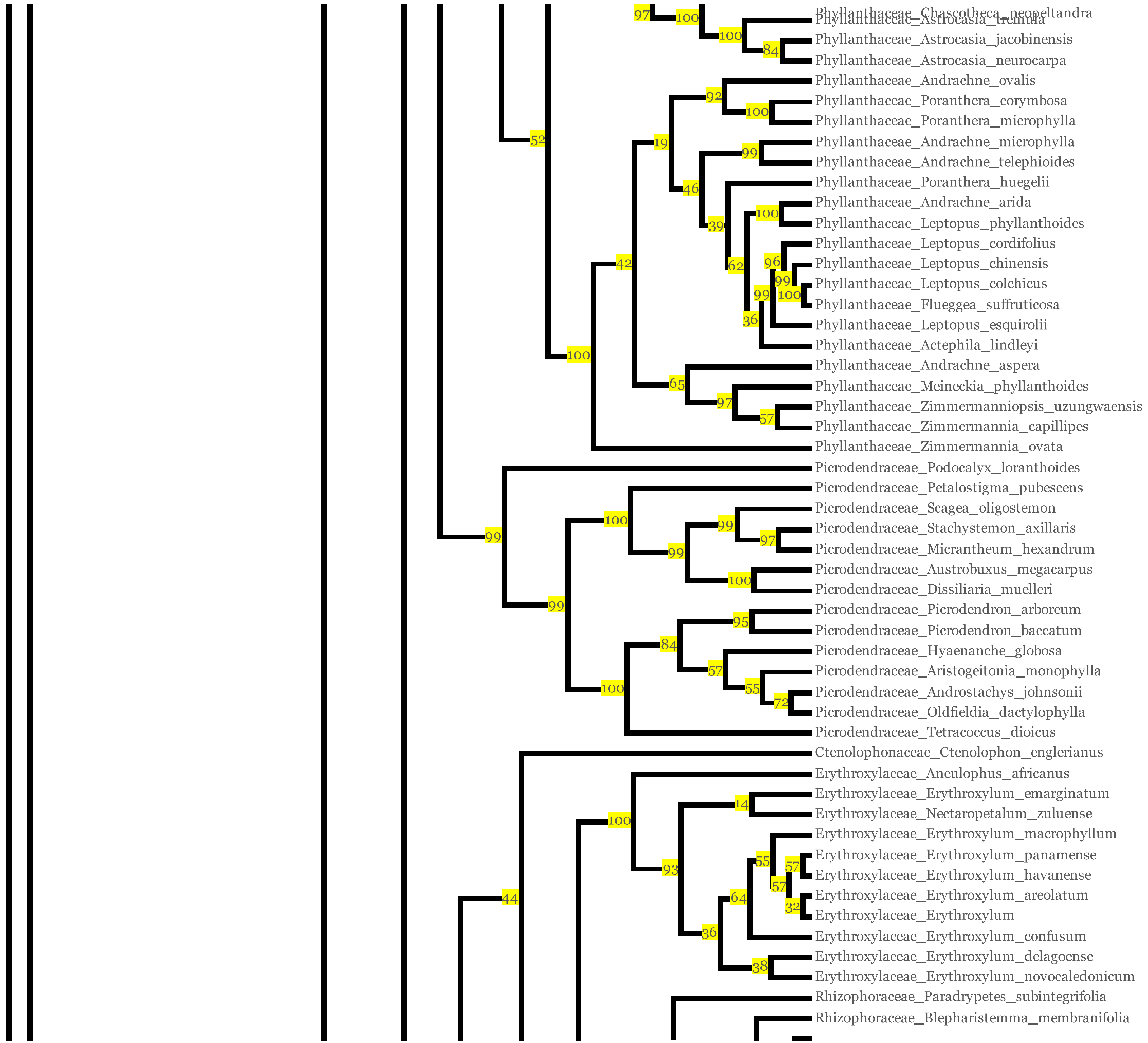
Phyllanthaceae_Chascotheca_neopeltandra
Phyllanthaceae_Astrocasia_tremula
Phyllanthaceae_Astrocasia_jacobinensis
Phyllanthaceae_Astrocasia_neurocarpa
Phyllanthaceae_Andrachne_ovalis
Phyllanthaceae_Poranthera_corymbosa
Phyllanthaceae_Poranthera_microphylla
Phyllanthaceae_Andrachne_microphylla
Phyllanthaceae_Andrachne_telephioides
Phyllanthaceae_Poranthera_huegelii
Phyllanthaceae_Andrachne_arida
Phyllanthaceae_Leptopus_phyllanthoides
Phyllanthaceae_Leptopus_cordifolius
Phyllanthaceae_Leptopus_chinensis
Phyllanthaceae_Leptopus_colchicus
Phyllanthaceae_Flueggea_suffruticosa
Phyllanthaceae_Leptopus_esquirolii
Phyllanthaceae_Actephila_lindleyi
Phyllanthaceae_Andrachne_aspera
Phyllanthaceae_Meineckia_phyllanthoides
Phyllanthaceae_Zimmermanniopsis_uzungwaensis
Phyllanthaceae_Zimmermannia_capillipes
Phyllanthaceae_Zimmermannia_ovata
Picrodendraceae_Podocalyx_loranthoides
Picrodendraceae_Petalostigma_pubescens
Picrodendraceae_Scagea_oligostemon
Picrodendraceae_Stachystemon_axillaris
Picrodendraceae_Micrantheum_hexandrum
Picrodendraceae_Austrobuxus_megacarpus
Picrodendraceae_Dissiliaria_muelleri
Picrodendraceae_Picrodendron_arboreum
Picrodendraceae_Picrodendron_baccatum
Picrodendraceae_Hyaenanche_globosa
Picrodendraceae_Aristogeitonia_monophylla
Picrodendraceae_Androstachys_johnsonii
Picrodendraceae_Oldfieldia_dactylophylla
Picrodendraceae_Tetracoccus_dioicus
Ctenolophonaceae_Ctenolophon_englerianus
Erythroxylaceae_Aneulophus_africanus
Erythroxylaceae_Erythroxylum_emarginatum
Erythroxylaceae_Nectaropetalum_zuluense
Erythroxylaceae_Erythroxylum_macrophyllum
Erythroxylaceae_Erythroxylum_panamense
Erythroxylaceae_Erythroxylum_havanense
Erythroxylaceae_Erythroxylum_areolatum
Erythroxylaceae_Erythroxylum
Erythroxylaceae_Erythroxylum_confusum
Erythroxylaceae_Erythroxylum_delagoense
Erythroxylaceae_Erythroxylum_novocaledonicum
Rhizophoraceae_Paradrypetes_subintegrifolia
Rhizophoraceae_Blepharistemma_membranifolia

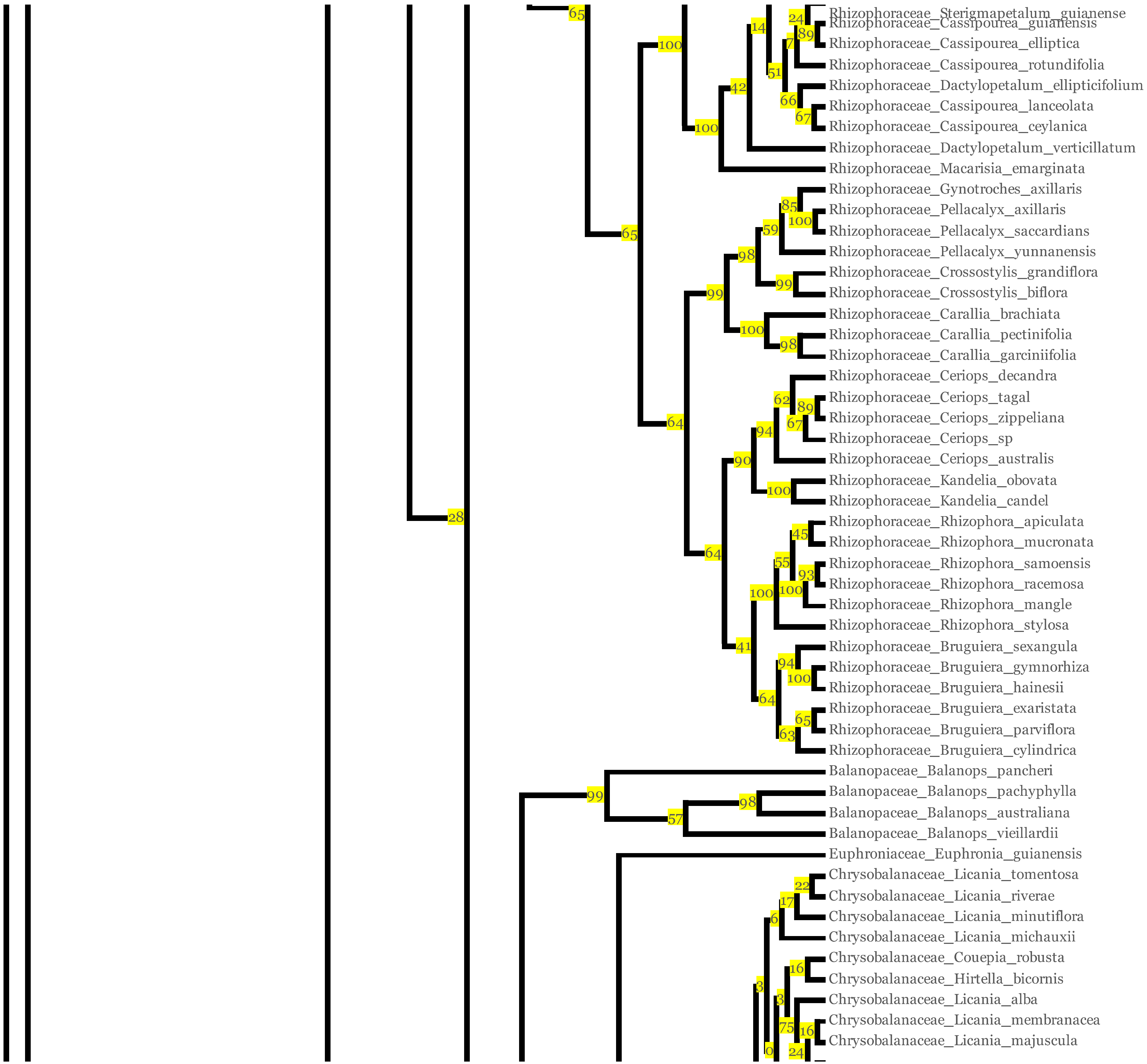
Rhizophoraceae_Sterigmapetalum_guianense
Rhizophoraceae_Cassipourea_guianensis
Rhizophoraceae_Cassipourea_elliptica
Rhizophoraceae_Cassipourea_rotundifolia
Rhizophoraceae_Dactylopetalum_ellipticifolium
Rhizophoraceae_Cassipourea_lanceolata
Rhizophoraceae_Cassipourea_ceylanica
Rhizophoraceae_Dactylopetalum_verticillatum
Rhizophoraceae_Macarisia_emarginata
Rhizophoraceae_Gynotroches_axillaris
Rhizophoraceae_Pellacalyx_axillaris
Rhizophoraceae_Pellacalyx_saccardians
Rhizophoraceae_Pellacalyx_yunnanensis
Rhizophoraceae_Crossostylis_grandiflora
Rhizophoraceae_Crossostylis_biflora
Rhizophoraceae_Carallia_brachiata
Rhizophoraceae_Carallia_pectinifolia
Rhizophoraceae_Carallia_garciniifolia
Rhizophoraceae_Ceriops_decandra
Rhizophoraceae_Ceriops_tagal
Rhizophoraceae_Ceriops_zippeliana
Rhizophoraceae_Ceriops_sp
Rhizophoraceae_Ceriops_australis
Rhizophoraceae_Kandelia_obovata
Rhizophoraceae_Kandelia_candel
Rhizophoraceae_Rhizophora_apiculata
Rhizophoraceae_Rhizophora_mucronata
Rhizophoraceae_Rhizophora_samoensis
Rhizophoraceae_Rhizophora_racemosa
Rhizophoraceae_Rhizophora_mangle
Rhizophoraceae_Rhizophora_stylosa
Rhizophoraceae_Bruguiera_sexangula
Rhizophoraceae_Bruguiera_gymnorhiza
Rhizophoraceae_Bruguiera_hainesii
Rhizophoraceae_Bruguiera_exaristata
Rhizophoraceae_Bruguiera_parviflora
Rhizophoraceae_Bruguiera_cylindrica
Balanopaceae_Balanops_pancheri
Balanopaceae_Balanops_pachyphylla
Balanopaceae_Balanops_australiana
Balanopaceae_Balanops_vieillardii
Euphroniaceae_Euphronia_guianensis
Chrysobalanaceae_Licania_tomentosa
Chrysobalanaceae_Licania_riverae
Chrysobalanaceae_Licania_minutiflora
Chrysobalanaceae_Licania_michauxii
Chrysobalanaceae_Couepia_robusta
Chrysobalanaceae_Hirtella_bicornis
Chrysobalanaceae_Licania_alba
Chrysobalanaceae_Licania_membranacea
Chrysobalanaceae_Licania_majuscula

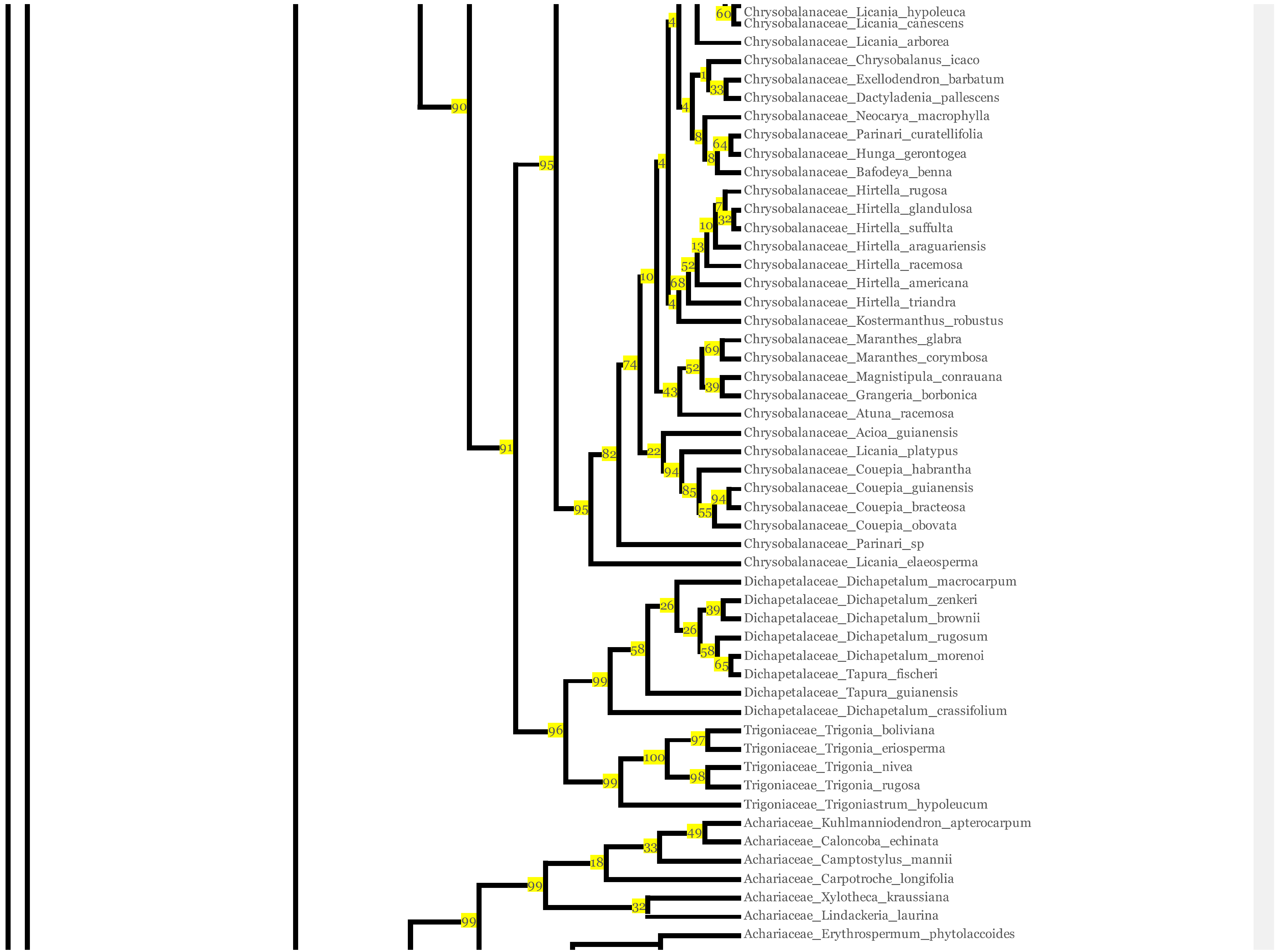
Chrysobalanaceae_Licania_hypoleuca
Chrysobalanaceae_Licania_canescens
Chrysobalanaceae_Licania_arborea
Chrysobalanaceae_Chrysobalanus_icaco
Chrysobalanaceae_Exellodendron_barbatum
Chrysobalanaceae_Dactyladenia_pallescens
Chrysobalanaceae_Neocarya_macrophylla
Chrysobalanaceae_Parinari_curatellifolia
Chrysobalanaceae_Hunga_gerontogea
Chrysobalanaceae_Bafodeya_benna
Chrysobalanaceae_Hirtella_rugosa
Chrysobalanaceae_Hirtella_glandulosa
Chrysobalanaceae_Hirtella_suffulta
Chrysobalanaceae_Hirtella_araguariensis
Chrysobalanaceae_Hirtella_racemosa
Chrysobalanaceae_Hirtella_americana
Chrysobalanaceae_Hirtella_triandra
Chrysobalanaceae_Kostermanthus_robustus
Chrysobalanaceae_Maranthes_glabra
Chrysobalanaceae_Maranthes_corymbosa
Chrysobalanaceae_Magnistipula_conrauana
Chrysobalanaceae_Grangeria_borbonica
Chrysobalanaceae_Atuna_racemosa
Chrysobalanaceae_Acioa_guianensis
Chrysobalanaceae_Licania_platypus
Chrysobalanaceae_Couepia_habrantha
Chrysobalanaceae_Couepia_guianensis
Chrysobalanaceae_Couepia_bracteosa
Chrysobalanaceae_Couepia_obovata
Chrysobalanaceae_Parinari_sp
Chrysobalanaceae_Licania_elaeosperma
Dichapetalaceae_Dichapetalum_macrocarpum
Dichapetalaceae_Dichapetalum_zenkeri
Dichapetalaceae_Dichapetalum_brownii
Dichapetalaceae_Dichapetalum_rugosum
Dichapetalaceae_Dichapetalum_morenoi
Dichapetalaceae_Tapura_fischeri
Dichapetalaceae_Tapura_guianensis
Dichapetalaceae_Dichapetalum_crassifolium
Trigoniaceae_Trigonia_boliviana
Trigoniaceae_Trigonia_eriosperma
Trigoniaceae_Trigonia_nivea
Trigoniaceae_Trigonia_rugosa
Trigoniaceae_Trigoniastrum_hypoleucum
Achariaceae_Kuhlmanniodendron_apterocarpum
Achariaceae_Caloncoba_echinata
Achariaceae_Camptostylus_mannii
Achariaceae_Carpotroche_longifolia
Achariaceae_Xylotheca_kraussiana
Achariaceae_Lindackeria_laurina
Achariaceae_Erythrospermum_phytolaccoides

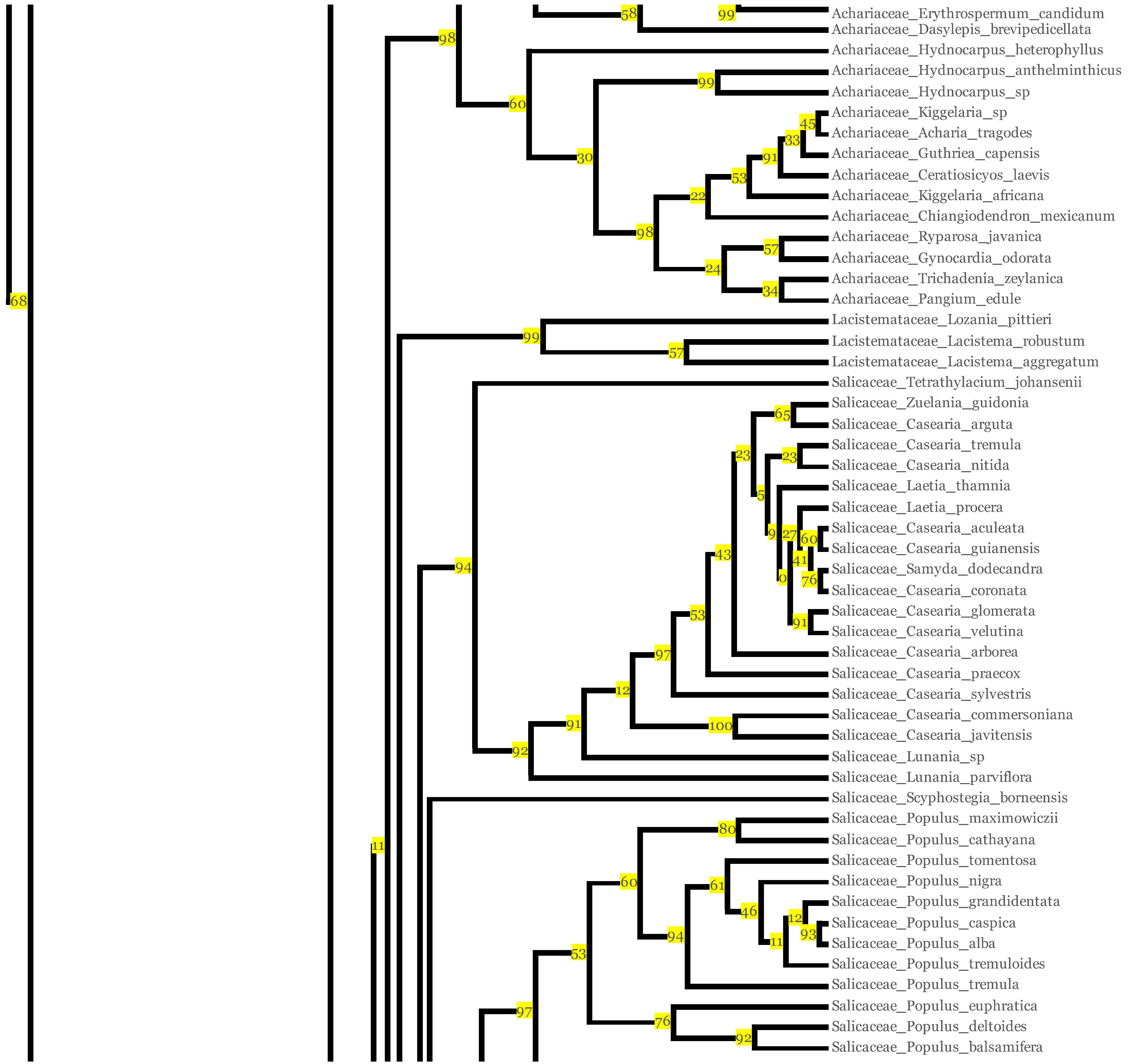
Achariaceae_Erythrospermum_candidum
Achariaceae_Dasylepis_brevipedicellata
Achariaceae_Hydnocarpus_heterophyllus
Achariaceae_Hydnocarpus_anthelminthicus
Achariaceae_Hydnocarpus_sp
Achariaceae_Kiggelaria_sp
Achariaceae_Acharia_tragodes
Achariaceae_Guthriea_capensis
Achariaceae_Ceratiosicyos_laevis
Achariaceae_Kiggelaria_africana
Achariaceae_Chiangiodendron_mexicanum
Achariaceae_Ryparosa_javanica
Achariaceae_Gynocardia_odorata
Achariaceae_Trichadenia_zeylanica
Achariaceae_Pangium_edule
Lacistemataceae_Lozania_pittieri
Lacistemataceae_Lacistema_robustum
Lacistemataceae_Lacistema_aggregatum
Salicaceae_Tetrathylacium_johansenii
Salicaceae_Zuelania_guidonia
Salicaceae_Casearia_arguta
Salicaceae_Casearia_tremula
Salicaceae_Casearia_nitida
Salicaceae_Laetia_thamnia
Salicaceae_Laetia_procera
Salicaceae_Casearia_aculeata
Salicaceae_Casearia_guianensis
Salicaceae_Samyda_dodecandra
Salicaceae_Casearia_coronata
Salicaceae_Casearia_glomerata
Salicaceae_Casearia_velutina
Salicaceae_Casearia_arborea
Salicaceae_Casearia_praecox
Salicaceae_Casearia_sylvestris
Salicaceae_Casearia_commersoniana
Salicaceae_Casearia_javitensis
Salicaceae_Lunania_sp
Salicaceae_Lunania_parviflora
Salicaceae_Scyphostegia_borneensis
Salicaceae_Populus_maximowiczii
Salicaceae_Populus_cathayana
Salicaceae_Populus_tomentosa
Salicaceae_Populus_nigra
Salicaceae_Populus_grandidentata
Salicaceae_Populus_caspica
Salicaceae_Populus_alba
Salicaceae_Populus_tremuloides
Salicaceae_Populus_tremula
Salicaceae_Populus_euphratica
Salicaceae_Populus_deltoides
Salicaceae_Populus_balsamifera

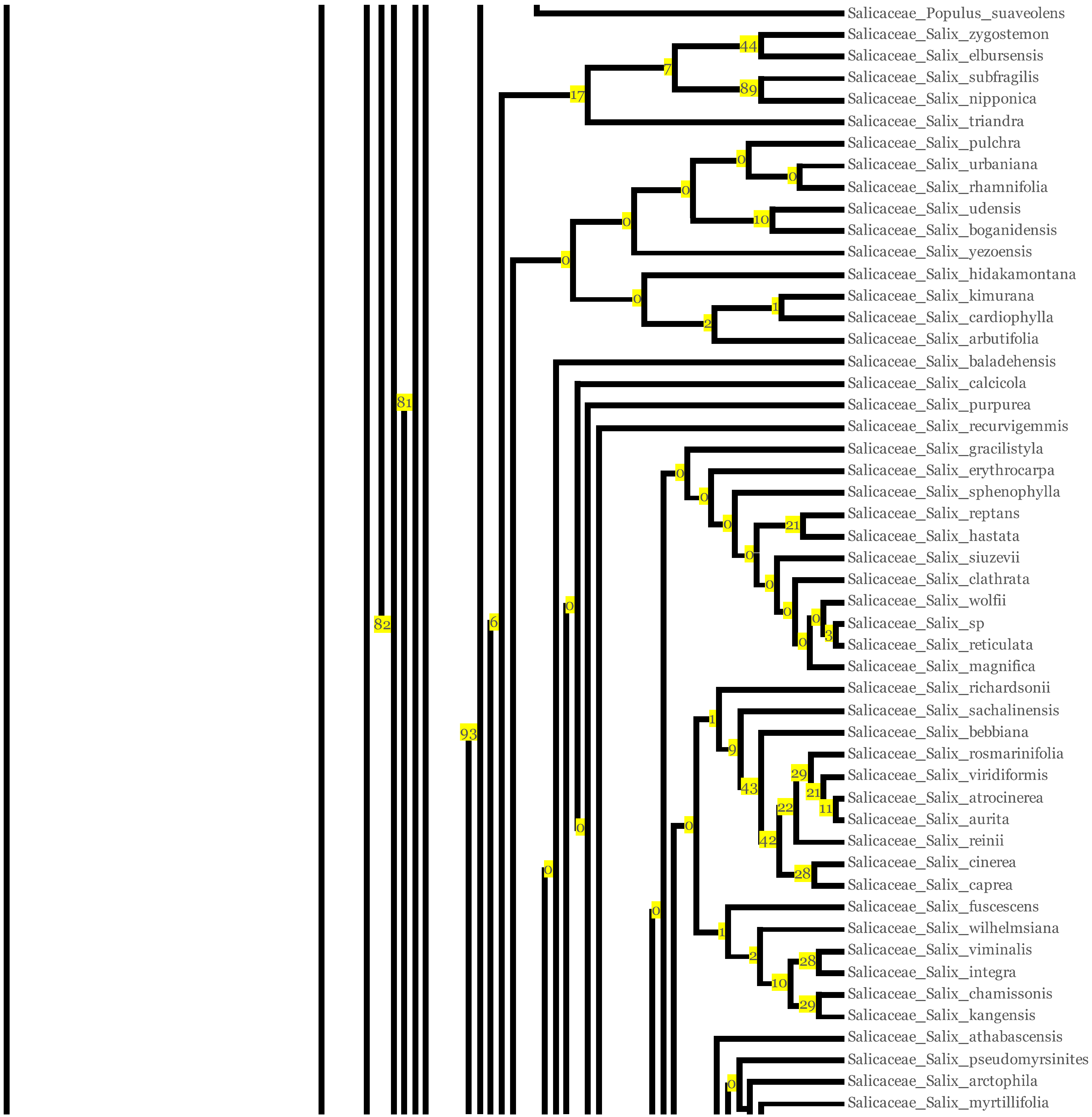
Salicaceae_Populus_suaveolens
Salicaceae_Salix_zygostemon
Salicaceae_Salix_elbursensis
Salicaceae_Salix_subfragilis
Salicaceae_Salix_nipponica
Salicaceae_Salix_triandra
Salicaceae_Salix_pulchra
Salicaceae_Salix_urbaniana
Salicaceae_Salix_rhamnifolia
Salicaceae_Salix_udensis
Salicaceae_Salix_boganidensis
Salicaceae_Salix_yezoensis
Salicaceae_Salix_hidakamontana
Salicaceae_Salix_kimurana
Salicaceae_Salix_cardiophylla
Salicaceae_Salix_arbutifolia
Salicaceae_Salix_baladehensis
Salicaceae_Salix_calcicola
Salicaceae_Salix_purpurea
Salicaceae_Salix_recurvigemmis
Salicaceae_Salix_gracilistyla
Salicaceae_Salix_erythrocarpa
Salicaceae_Salix_sphenophylla
Salicaceae_Salix_reptans
Salicaceae_Salix_hastata
Salicaceae_Salix_siuzevii
Salicaceae_Salix_clathrata
Salicaceae_Salix_wolfii
Salicaceae_Salix_sp
Salicaceae_Salix_reticulata
Salicaceae_Salix_magnifica
Salicaceae_Salix_richardsonii
Salicaceae_Salix_sachalinensis
Salicaceae_Salix_bebbiana
Salicaceae_Salix_rosmarinifolia
Salicaceae_Salix_viridiformis
Salicaceae_Salix_atrocinerea
Salicaceae_Salix_aurita
Salicaceae_Salix_reinii
Salicaceae_Salix_cinerea
Salicaceae_Salix_caprea
Salicaceae_Salix_fuscescens
Salicaceae_Salix_wilhelmsiana
Salicaceae_Salix_viminalis
Salicaceae_Salix_integra
Salicaceae_Salix_chamissonis
Salicaceae_Salix_kangensis
Salicaceae_Salix_athabascensis
Salicaceae_Salix_pseudomyrsinites
Salicaceae_Salix_arctophila
Salicaceae_Salix_myrtillifolia

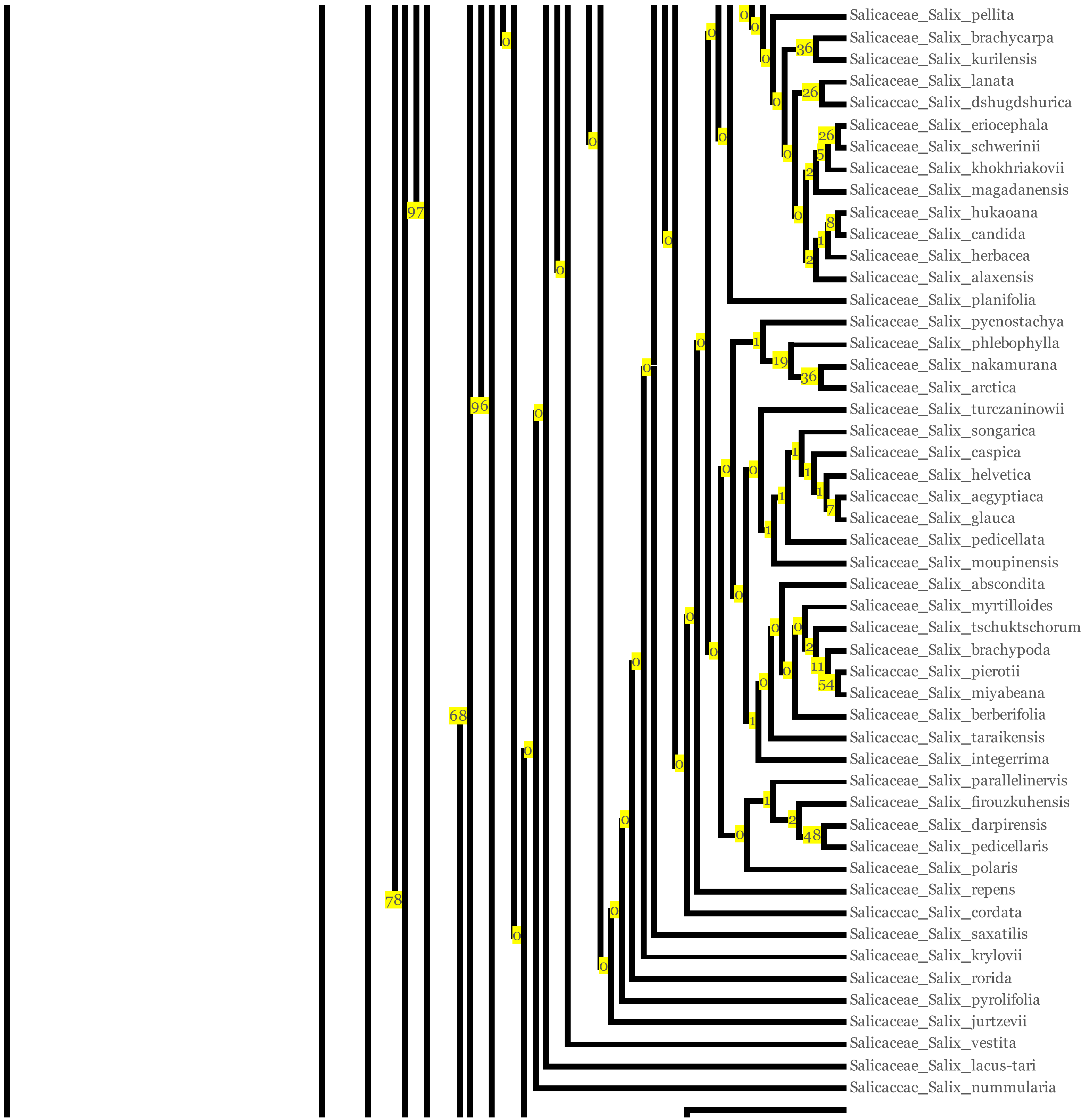
Salicaceae_Salix_pellita
Salicaceae_Salix_brachycarpa
Salicaceae_Salix_kurilensis
Salicaceae_Salix_lanata
Salicaceae_Salix_dshugdshurica
Salicaceae_Salix_eriocephala
Salicaceae_Salix_schwerinii
Salicaceae_Salix_khokhriakovii
Salicaceae_Salix_magadanensis
Salicaceae_Salix_hukaoana
Salicaceae_Salix_candida
Salicaceae_Salix_herbacea
Salicaceae_Salix_alaxensis
Salicaceae_Salix_planifolia
Salicaceae_Salix_pycnostachya
Salicaceae_Salix_phlebophylla
Salicaceae_Salix_nakamurana
Salicaceae_Salix_arctica
Salicaceae_Salix_turczaninowii
Salicaceae_Salix_songarica
Salicaceae_Salix_caspica
Salicaceae_Salix_helvetica
Salicaceae_Salix_aegyptiaca
Salicaceae_Salix_glauca
Salicaceae_Salix_pedicellata
Salicaceae_Salix_moupinensis
Salicaceae_Salix_abscondita
Salicaceae_Salix_myrtilloides
Salicaceae_Salix_tschuktschorum
Salicaceae_Salix_brachypoda
Salicaceae_Salix_pierotii
Salicaceae_Salix_miyabeana
Salicaceae_Salix_berberifolia
Salicaceae_Salix_taraikensis
Salicaceae_Salix_integerrima
Salicaceae_Salix_parallelinervis
Salicaceae_Salix_firouzkuhensis
Salicaceae_Salix_darpirensis
Salicaceae_Salix_pedicellaris
Salicaceae_Salix_polaris
Salicaceae_Salix_repens
Salicaceae_Salix_cordata
Salicaceae_Salix_saxatilis
Salicaceae_Salix_krylovii
Salicaceae_Salix_rorida
Salicaceae_Salix_pyrolifolia
Salicaceae_Salix_jurtzevii
Salicaceae_Salix_vestita
Salicaceae_Salix_lacus-tari
Salicaceae_Salix_nummularia

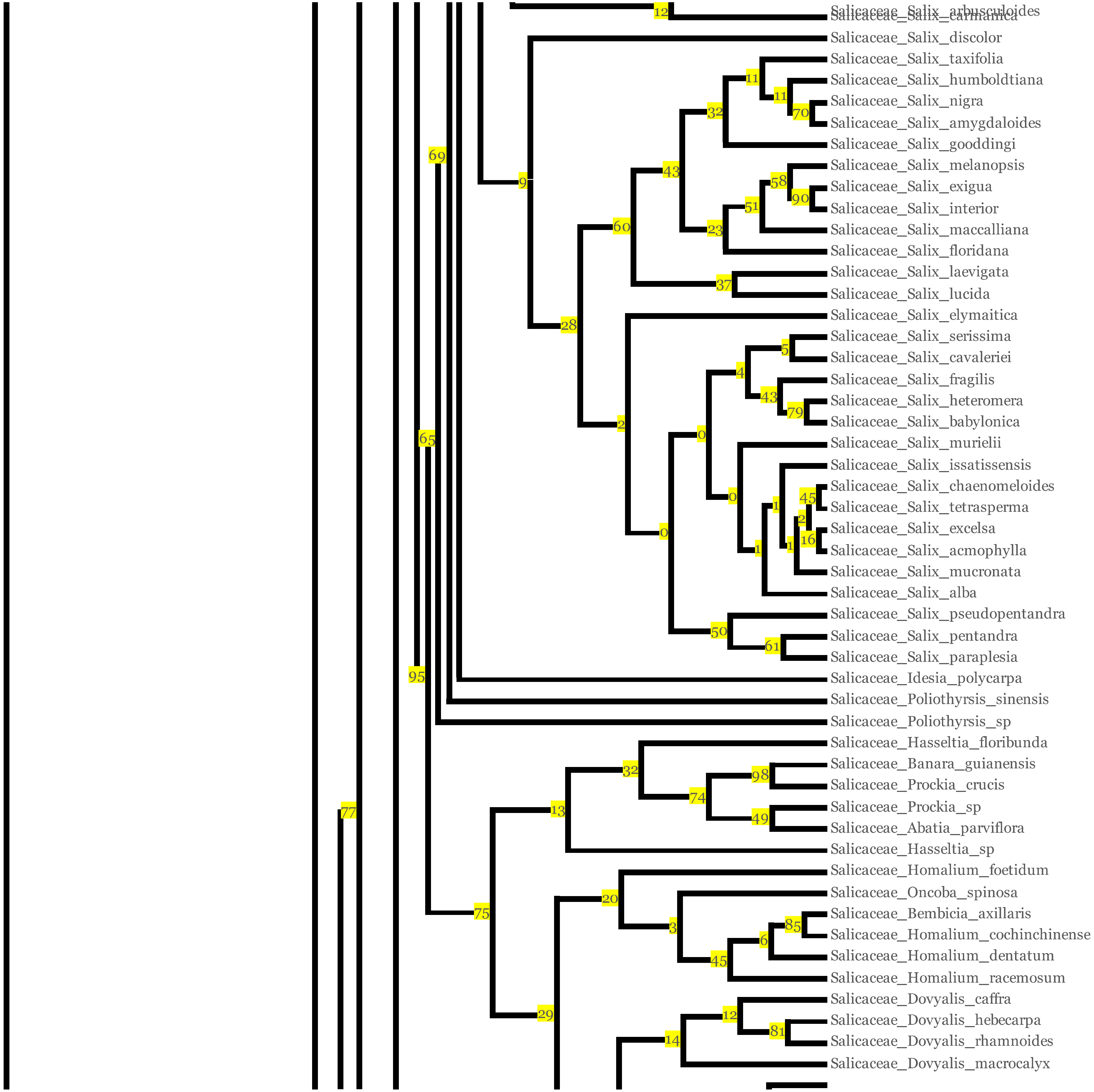
Salicaceae_Salix_arbusculoides
Salicaceae_Salix_carmanica
Salicaceae_Salix_discolor
Salicaceae_Salix_taxifolia
Salicaceae_Salix_humboldtiana
Salicaceae_Salix_nigra
Salicaceae_Salix_amygdaloides
Salicaceae_Salix_gooddingi
Salicaceae_Salix_melanopsis
Salicaceae_Salix_exigua
Salicaceae_Salix_interior
Salicaceae_Salix_maccalliana
Salicaceae_Salix_floridana
Salicaceae_Salix_laevigata
Salicaceae_Salix_lucida
Salicaceae_Salix_elymaitica
Salicaceae_Salix_serissima
Salicaceae_Salix_cavaleriei
Salicaceae_Salix_fragilis
Salicaceae_Salix_heteromera
Salicaceae_Salix_babylonica
Salicaceae_Salix_murielii
Salicaceae_Salix_issatissensis
Salicaceae_Salix_chaenomeloides
Salicaceae_Salix_tetrasperma
Salicaceae_Salix_excelsa
Salicaceae_Salix_acmophylla
Salicaceae_Salix_mucronata
Salicaceae_Salix_alba
Salicaceae_Salix_pseudopentandra
Salicaceae_Salix_pentandra
Salicaceae_Salix_paraplesia
Salicaceae_Idesia_polycarpa
Salicaceae_Poliothyrsis_sinensis
Salicaceae_Poliothyrsis_sp
Salicaceae_Hasseltia_floribunda
Salicaceae_Banara_guianensis
Salicaceae_Prockia_crucis
Salicaceae_Prockia_sp
Salicaceae_Abatia_parviflora
Salicaceae_Hasseltia_sp
Salicaceae_Homalium_foetidum
Salicaceae_Oncoba_spinosa
Salicaceae_Bembicia_axillaris
Salicaceae_Homalium_cochinchinense
Salicaceae_Homalium_dentatum
Salicaceae_Homalium_racemosum
Salicaceae_Dovyalis_caffra
Salicaceae_Dovyalis_hebecarpa
Salicaceae_Dovyalis_rhamnoides
Salicaceae_Dovyalis_macrocalyx

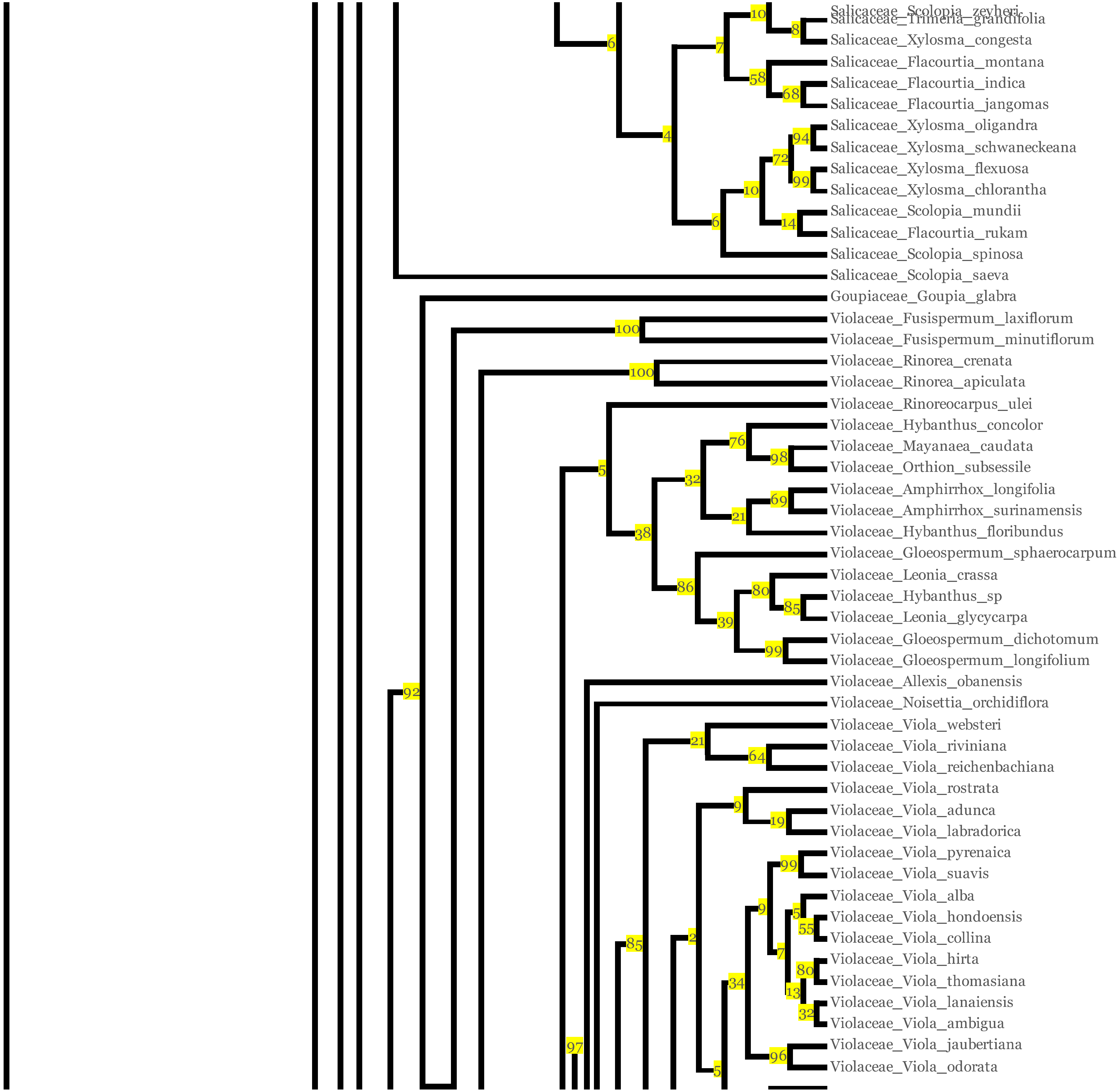
Salicaceae_Scolopia_zeyheri
Salicaceae_Trimeria_grandifolia
Salicaceae_Xylosma_congesta
Salicaceae_Flacourtia_montana
Salicaceae_Flacourtia_indica
Salicaceae_Flacourtia_jangomas
Salicaceae_Xylosma_oligandra
Salicaceae_Xylosma_schwaneckeana
Salicaceae_Xylosma_flexuosa
Salicaceae_Xylosma_chlorantha
Salicaceae_Scolopia_mundii
Salicaceae_Flacourtia_rukam
Salicaceae_Scolopia_spinosa
Salicaceae_Scolopia_saeva
Goupiaceae_Goupia_glabra
Violaceae_Fusispermum_laxiflorum
Violaceae_Fusispermum_minutiflorum
Violaceae_Rinorea_crenata
Violaceae_Rinorea_apiculata
Violaceae_Rinoreocarpus_ulei
Violaceae_Hybanthus_concolor
Violaceae_Mayanaea_caudata
Violaceae_Orthion_subsessile
Violaceae_Amphirrhox_longifolia
Violaceae_Amphirrhox_surinamensis
Violaceae_Hybanthus_floribundus
Violaceae_Gloeospermum_sphaerocarpum
Violaceae_Leonia_crassa
Violaceae_Hybanthus_sp
Violaceae_Leonia_glycycarpa
Violaceae_Gloeospermum_dichotomum
Violaceae_Gloeospermum_longifolium
Violaceae_Allexis_obanensis
Violaceae_Noisettia_orchidiflora
Violaceae_Viola_websteri
Violaceae_Viola_riviniana
Violaceae_Viola_reichenbachiana
Violaceae_Viola_rostrata
Violaceae_Viola_adunca
Violaceae_Viola_labradorica
Violaceae_Viola_pyrenaica
Violaceae_Viola_suavis
Violaceae_Viola_alba
Violaceae_Viola_hondoensis
Violaceae_Viola_collina
Violaceae_Viola_hirta
Violaceae_Viola_thomasiana
Violaceae_Viola_lanaiensis
Violaceae_Viola_ambigua
Violaceae_Viola_jaubertiana
Violaceae_Viola_odorata

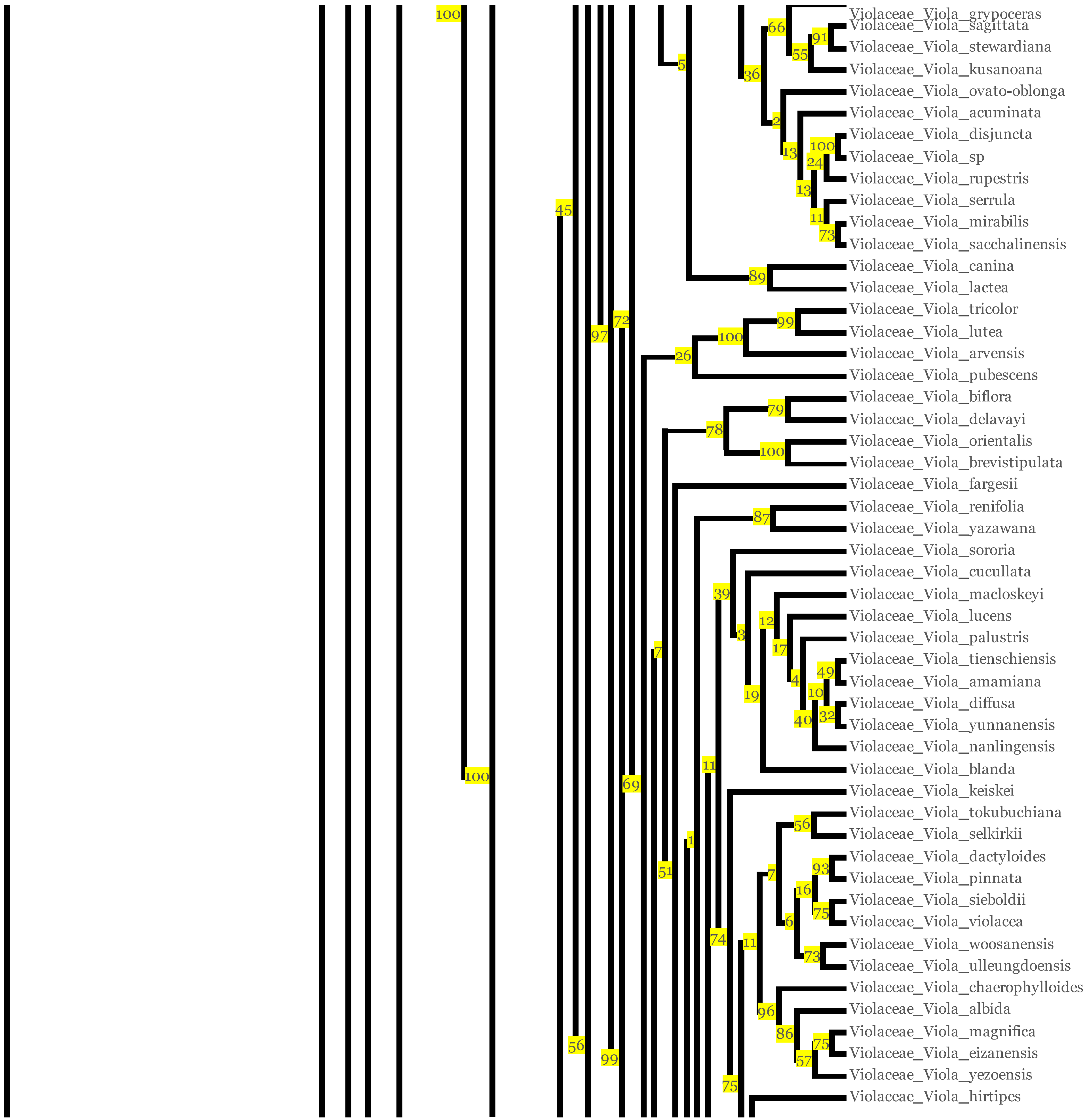
Violaceae_Viola_grypoceras
Violaceae_Viola_sagittata
Violaceae_Viola_stewardiana
Violaceae_Viola_kusanoana
Violaceae_Viola_ovato-oblonga
Violaceae_Viola_acuminata
Violaceae_Viola_disjuncta
Violaceae_Viola_sp
Violaceae_Viola_rupestris
Violaceae_Viola_serrula
Violaceae_Viola_mirabilis
Violaceae_Viola_sacchalinensis
Violaceae_Viola_canina
Violaceae_Viola_lactea
Violaceae_Viola_tricolor
Violaceae_Viola_lutea
Violaceae_Viola_arvensis
Violaceae_Viola_pubescens
Violaceae_Viola_biflora
Violaceae_Viola_delavayi
Violaceae_Viola_orientalis
Violaceae_Viola_brevistipulata
Violaceae_Viola_fargesii
Violaceae_Viola_renifolia
Violaceae_Viola_yazawana
Violaceae_Viola_sororia
Violaceae_Viola_cucullata
Violaceae_Viola_macloskeyi
Violaceae_Viola_lucens
Violaceae_Viola_palustris
Violaceae_Viola_tienschiensis
Violaceae_Viola_amamiana
Violaceae_Viola_diffusa
Violaceae_Viola_yunnanensis
Violaceae_Viola_nanlingensis
Violaceae_Viola_blanda
Violaceae_Viola_keiskei
Violaceae_Viola_tokubuchiana
Violaceae_Viola_selkirkii
Violaceae_Viola_dactyloides
Violaceae_Viola_pinnata
Violaceae_Viola_sieboldii
Violaceae_Viola_violacea
Violaceae_Viola_woosanensis
Violaceae_Viola_ulleungdoensis
Violaceae_Viola_chaerophylloides
Violaceae_Viola_albida
Violaceae_Viola_magnifica
Violaceae_Viola_eizanensis
Violaceae_Viola_yezoensis
Violaceae_Viola_hirtipes

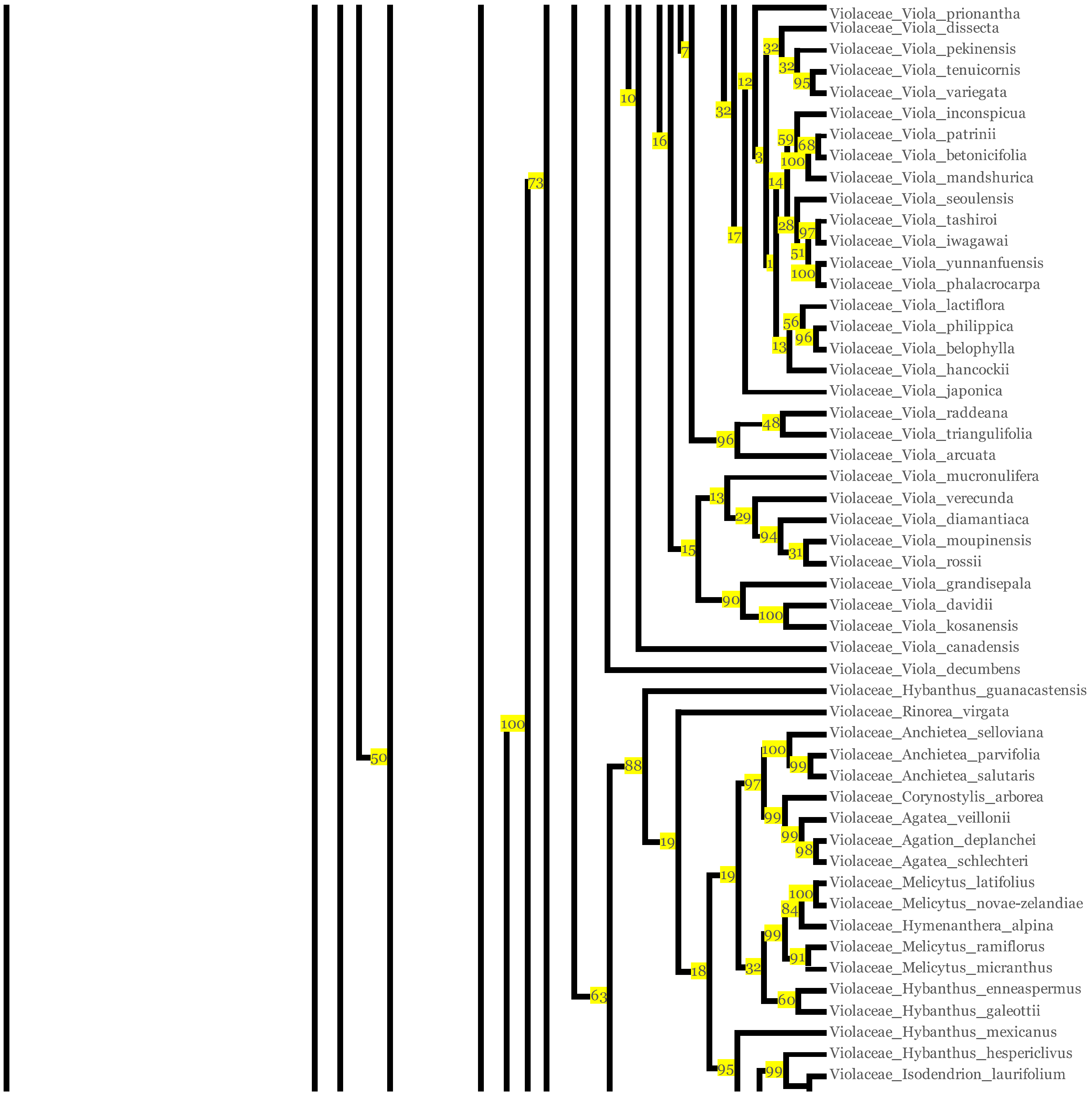
Violaceae_Viola_prionantha
Violaceae_Viola_dissecta
Violaceae_Viola_pekinensis
Violaceae_Viola_tenuicornis
Violaceae_Viola_variegata
Violaceae_Viola_inconspicua
Violaceae_Viola_patrinii
Violaceae_Viola_betonicifolia
Violaceae_Viola_mandshurica
Violaceae_Viola_seoulensis
Violaceae_Viola_tashiroi
Violaceae_Viola_iwagawai
Violaceae_Viola_yunnanfuensis
Violaceae_Viola_phalacrocarpa
Violaceae_Viola_lactiflora
Violaceae_Viola_philippica
Violaceae_Viola_belophylla
Violaceae_Viola_hancockii
Violaceae_Viola_japonica
Violaceae_Viola_raddeana
Violaceae_Viola_triangulifolia
Violaceae_Viola_arcuata
Violaceae_Viola_mucronulifera
Violaceae_Viola_verecunda
Violaceae_Viola_diamantiaca
Violaceae_Viola_moupinensis
Violaceae_Viola_rossii
Violaceae_Viola_grandisepala
Violaceae_Viola_davidii
Violaceae_Viola_kosanensis
Violaceae_Viola_canadensis
Violaceae_Viola_decumbens
Violaceae_Hybanthus_guanacastensis
Violaceae_Rinorea_virgata
Violaceae_Anchietea_selloviana
Violaceae_Anchietea_parvifolia
Violaceae_Anchietea_salutaris
Violaceae_Corynostylis_arborea
Violaceae_Agatea_veillonii
Violaceae_Agation_deplanchei
Violaceae_Agatea_schlechteri
Violaceae_Melicytus_latifolius
Violaceae_Melicytus_novae-zelandiae
Violaceae_Hymenanthera_alpina
Violaceae_Melicytus_ramiflorus
Violaceae_Melicytus_micranthus
Violaceae_Hybanthus_enneaspermus
Violaceae_Hybanthus_galeottii
Violaceae_Hybanthus_mexicanus
Violaceae_Hybanthus_hespericlivus
Violaceae_Isodendrion_laurifolium

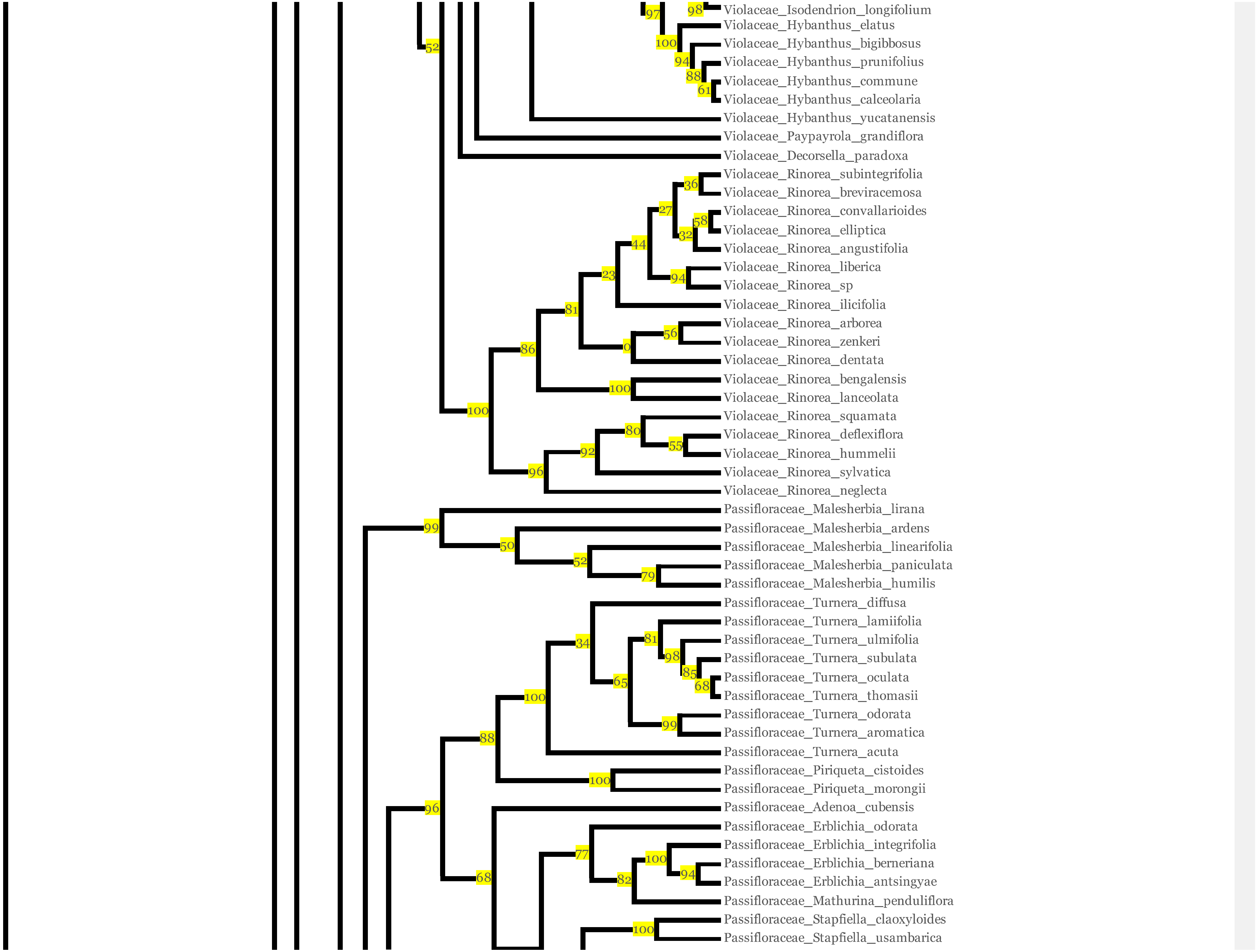
Violaceae_Isodendrion_longifolium
Violaceae_Hybanthus_elatus
Violaceae_Hybanthus_bigibbosus
Violaceae_Hybanthus_prunifolius
Violaceae_Hybanthus_commune
Violaceae_Hybanthus_calceolaria
Violaceae_Hybanthus_yucatanensis
Violaceae_Paypayrola_grandiflora
Violaceae_Decorsella_paradoxa
Violaceae_Rinorea_subintegrifolia
Violaceae_Rinorea_breviracemosa
Violaceae_Rinorea_convallarioides
Violaceae_Rinorea_elliptica
Violaceae_Rinorea_angustifolia
Violaceae_Rinorea_liberica
Violaceae_Rinorea_sp
Violaceae_Rinorea_ilicifolia
Violaceae_Rinorea_arborea
Violaceae_Rinorea_zenkeri
Violaceae_Rinorea_dentata
Violaceae_Rinorea_bengalensis
Violaceae_Rinorea_lanceolata
Violaceae_Rinorea_squamata
Violaceae_Rinorea_deflexiflora
Violaceae_Rinorea_hummelii
Violaceae_Rinorea_sylvatica
Violaceae_Rinorea_neglecta
Passifloraceae_Malesherbia_lirana
Passifloraceae_Malesherbia_ardens
Passifloraceae_Malesherbia_linearifolia
Passifloraceae_Malesherbia_paniculata
Passifloraceae_Malesherbia_humilis
Passifloraceae_Turnera_diffusa
Passifloraceae_Turnera_lamiifolia
Passifloraceae_Turnera_ulmifolia
Passifloraceae_Turnera_subulata
Passifloraceae_Turnera_oculata
Passifloraceae_Turnera_thomasii
Passifloraceae_Turnera_odorata
Passifloraceae_Turnera_aromatica
Passifloraceae_Turnera_acuta
Passifloraceae_Piriqueta_cistoides
Passifloraceae_Piriqueta_morongii
Passifloraceae_Adenoa_cubensis
Passifloraceae_Erblichia_odorata
Passifloraceae_Erblichia_integrifolia
Passifloraceae_Erblichia_berneriana
Passifloraceae_Erblichia_antsingyae
Passifloraceae_Mathurina_penduliflora
Passifloraceae_Stapfiella_claoxyloides
Passifloraceae_Stapfiella_usambarica

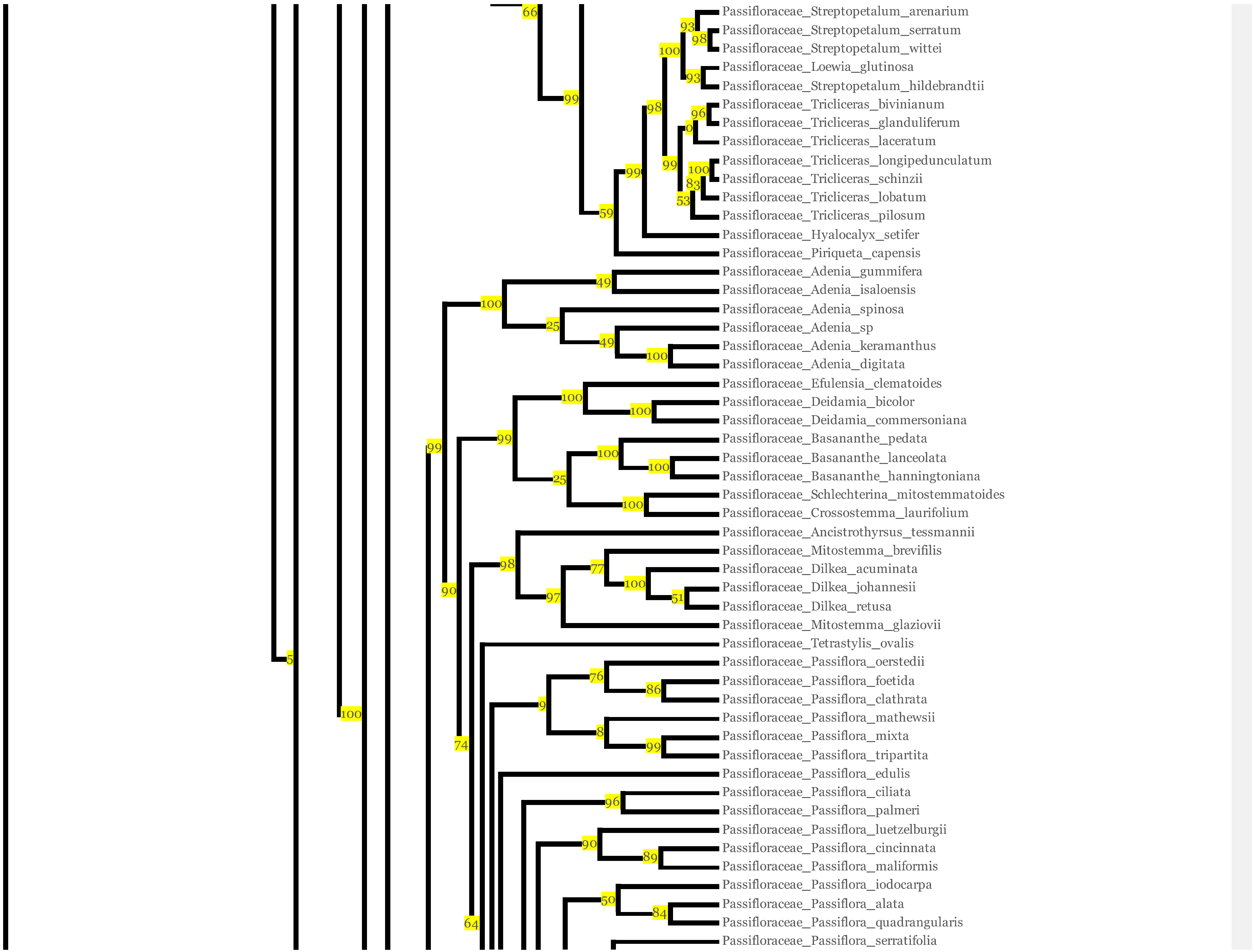
Passifloraceae_Streptopetalum_arenarium
Passifloraceae_Streptopetalum_serratum
Passifloraceae_Streptopetalum_wittei
Passifloraceae_Loewia_glutinosa
Passifloraceae_Streptopetalum_hildebrandtii
Passifloraceae_Tricliceras_bivinianum
Passifloraceae_Tricliceras_glanduliferum
Passifloraceae_Tricliceras_laceratum
Passifloraceae_Tricliceras_longipedunculatum
Passifloraceae_Tricliceras_schinzii
Passifloraceae_Tricliceras_lobatum
Passifloraceae_Tricliceras_pilosum
Passifloraceae_Hyalocalyx_setifer
Passifloraceae_Piriqueta_capensis
Passifloraceae_Adenia_gummifera
Passifloraceae_Adenia_isaloensis
Passifloraceae_Adenia_spinosa
Passifloraceae_Adenia_sp
Passifloraceae_Adenia_keramanthus
Passifloraceae_Adenia_digitata
Passifloraceae_Efulensia_clematoides
Passifloraceae_Deidamia_bicolor
Passifloraceae_Deidamia_commersoniana
Passifloraceae_Basananthe_pedata
Passifloraceae_Basananthe_lanceolata
Passifloraceae_Basananthe_hanningtoniana
Passifloraceae_Schlechterina_mitostemmatoides
Passifloraceae_Crossostemma_laurifolium
Passifloraceae_Ancistrothyrsus_tessmannii
Passifloraceae_Mitostemma_brevifilis
Passifloraceae_Dilkea_acuminata
Passifloraceae_Dilkea_johannesii
Passifloraceae_Dilkea_retusa
Passifloraceae_Mitostemma_glaziovii
Passifloraceae_Tetrastylis_ovalis
Passifloraceae_Passiflora_oerstedii
Passifloraceae_Passiflora_foetida
Passifloraceae_Passiflora_clathrata
Passifloraceae_Passiflora_mathewsii
Passifloraceae_Passiflora_mixta
Passifloraceae_Passiflora_tripartita
Passifloraceae_Passiflora_edulis
Passifloraceae_Passiflora_ciliata
Passifloraceae_Passiflora_palmeri
Passifloraceae_Passiflora_luetzelburgii
Passifloraceae_Passiflora_cincinnata
Passifloraceae_Passiflora_maliformis
Passifloraceae_Passiflora_iodocarpa
Passifloraceae_Passiflora_alata
Passifloraceae_Passiflora_quadrangularis
Passifloraceae_Passiflora_serratifolia

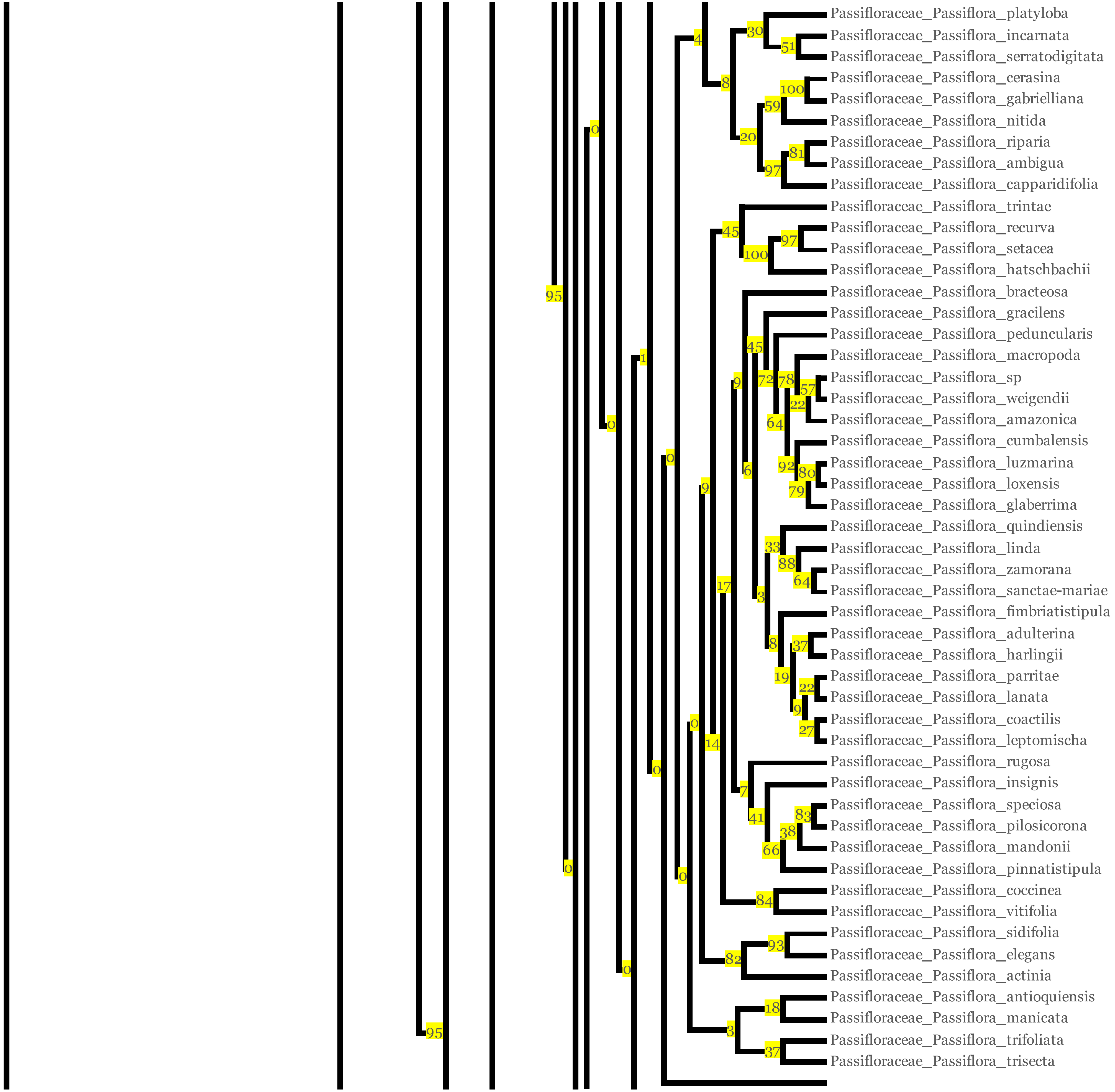
Passifloraceae_Passiflora_platyloba
Passifloraceae_Passiflora_incarnata
Passifloraceae_Passiflora_serratodigitata
Passifloraceae_Passiflora_cerasina
Passifloraceae_Passiflora_gabrielliana
Passifloraceae_Passiflora_nitida
Passifloraceae_Passiflora_riparia
Passifloraceae_Passiflora_ambigua
Passifloraceae_Passiflora_capparidifolia
Passifloraceae_Passiflora_trintae
Passifloraceae_Passiflora_recurva
Passifloraceae_Passiflora_setacea
Passifloraceae_Passiflora_hatschbachii
Passifloraceae_Passiflora_bracteosa
Passifloraceae_Passiflora_gracilens
Passifloraceae_Passiflora_peduncularis
Passifloraceae_Passiflora_macropoda
Passifloraceae_Passiflora_sp
Passifloraceae_Passiflora_weigendii
Passifloraceae_Passiflora_amazonica
Passifloraceae_Passiflora_cumbalensis
Passifloraceae_Passiflora_luzmarina
Passifloraceae_Passiflora_loxensis
Passifloraceae_Passiflora_glaberrima
Passifloraceae_Passiflora_quindiensis
Passifloraceae_Passiflora_linda
Passifloraceae_Passiflora_zamorana
Passifloraceae_Passiflora_sanctae-mariae
Passifloraceae_Passiflora_fimbriatistipula
Passifloraceae_Passiflora_adulterina
Passifloraceae_Passiflora_harlingii
Passifloraceae_Passiflora_parritae
Passifloraceae_Passiflora_lanata
Passifloraceae_Passiflora_coactilis
Passifloraceae_Passiflora_leptomischa
Passifloraceae_Passiflora_rugosa
Passifloraceae_Passiflora_insignis
Passifloraceae_Passiflora_speciosa
Passifloraceae_Passiflora_pilosicorona
Passifloraceae_Passiflora_mandonii
Passifloraceae_Passiflora_pinnatistipula
Passifloraceae_Passiflora_coccinea
Passifloraceae_Passiflora_vitifolia
Passifloraceae_Passiflora_sidifolia
Passifloraceae_Passiflora_elegans
Passifloraceae_Passiflora_actinia
Passifloraceae_Passiflora_antioquiensis
Passifloraceae_Passiflora_manicata
Passifloraceae_Passiflora_trifoliata
Passifloraceae_Passiflora_trisecta

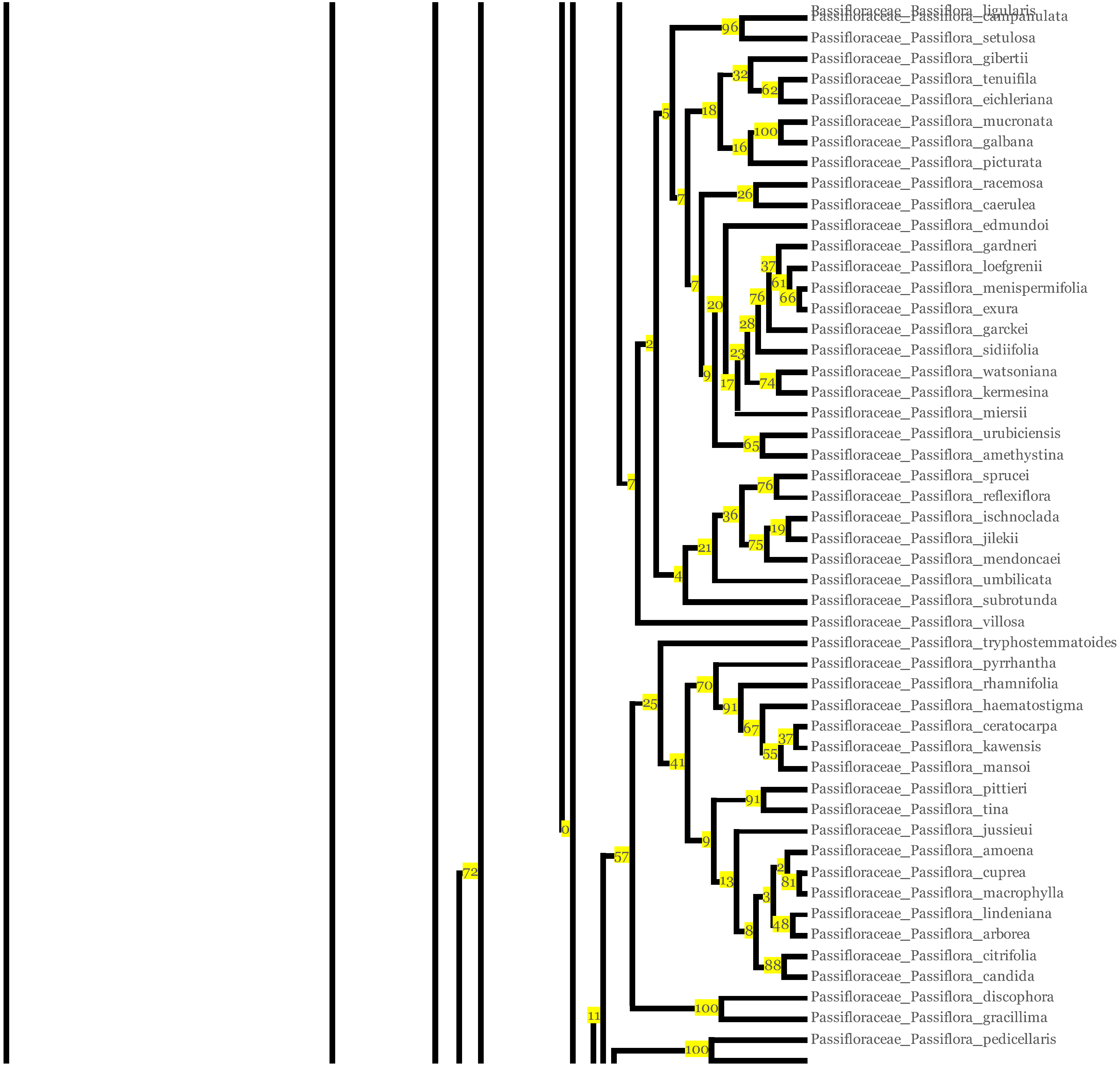
Passifloraceae_Passiflora_ligularis
Passifloraceae_Passiflora_campanulata
Passifloraceae_Passiflora_setulosa
Passifloraceae_Passiflora_gibertii
Passifloraceae_Passiflora_tenuifila
Passifloraceae_Passiflora_eichleriana
Passifloraceae_Passiflora_mucronata
Passifloraceae_Passiflora_galbana
Passifloraceae_Passiflora_picturata
Passifloraceae_Passiflora_racemosa
Passifloraceae_Passiflora_caerulea
Passifloraceae_Passiflora_edmundoi
Passifloraceae_Passiflora_gardneri
Passifloraceae_Passiflora_loefgrenii
Passifloraceae_Passiflora_menispermifolia
Passifloraceae_Passiflora_exura
Passifloraceae_Passiflora_garckei
Passifloraceae_Passiflora_sidiifolia
Passifloraceae_Passiflora_watsoniana
Passifloraceae_Passiflora_kermesina
Passifloraceae_Passiflora_miersii
Passifloraceae_Passiflora_urubiciensis
Passifloraceae_Passiflora_amethystina
Passifloraceae_Passiflora_sprucei
Passifloraceae_Passiflora_reflexiflora
Passifloraceae_Passiflora_ischnoclada
Passifloraceae_Passiflora_jilekii
Passifloraceae_Passiflora_mendoncaei
Passifloraceae_Passiflora_umbilicata
Passifloraceae_Passiflora_subrotunda
Passifloraceae_Passiflora_villosa
Passifloraceae_Passiflora_tryphostemmatoides
Passifloraceae_Passiflora_pyrrhantha
Passifloraceae_Passiflora_rhamnifolia
Passifloraceae_Passiflora_haematostigma
Passifloraceae_Passiflora_ceratocarpa
Passifloraceae_Passiflora_kawensis
Passifloraceae_Passiflora_mansoi
Passifloraceae_Passiflora_pittieri
Passifloraceae_Passiflora_tina
Passifloraceae_Passiflora_jussieui
Passifloraceae_Passiflora_amoena
Passifloraceae_Passiflora_cuprea
Passifloraceae_Passiflora_macrophylla
Passifloraceae_Passiflora_lindeniana
Passifloraceae_Passiflora_arborea
Passifloraceae_Passiflora_citrifolia
Passifloraceae_Passiflora_candida
Passifloraceae_Passiflora_discophora
Passifloraceae_Passiflora_gracillima
Passifloraceae_Passiflora_pedicellaris

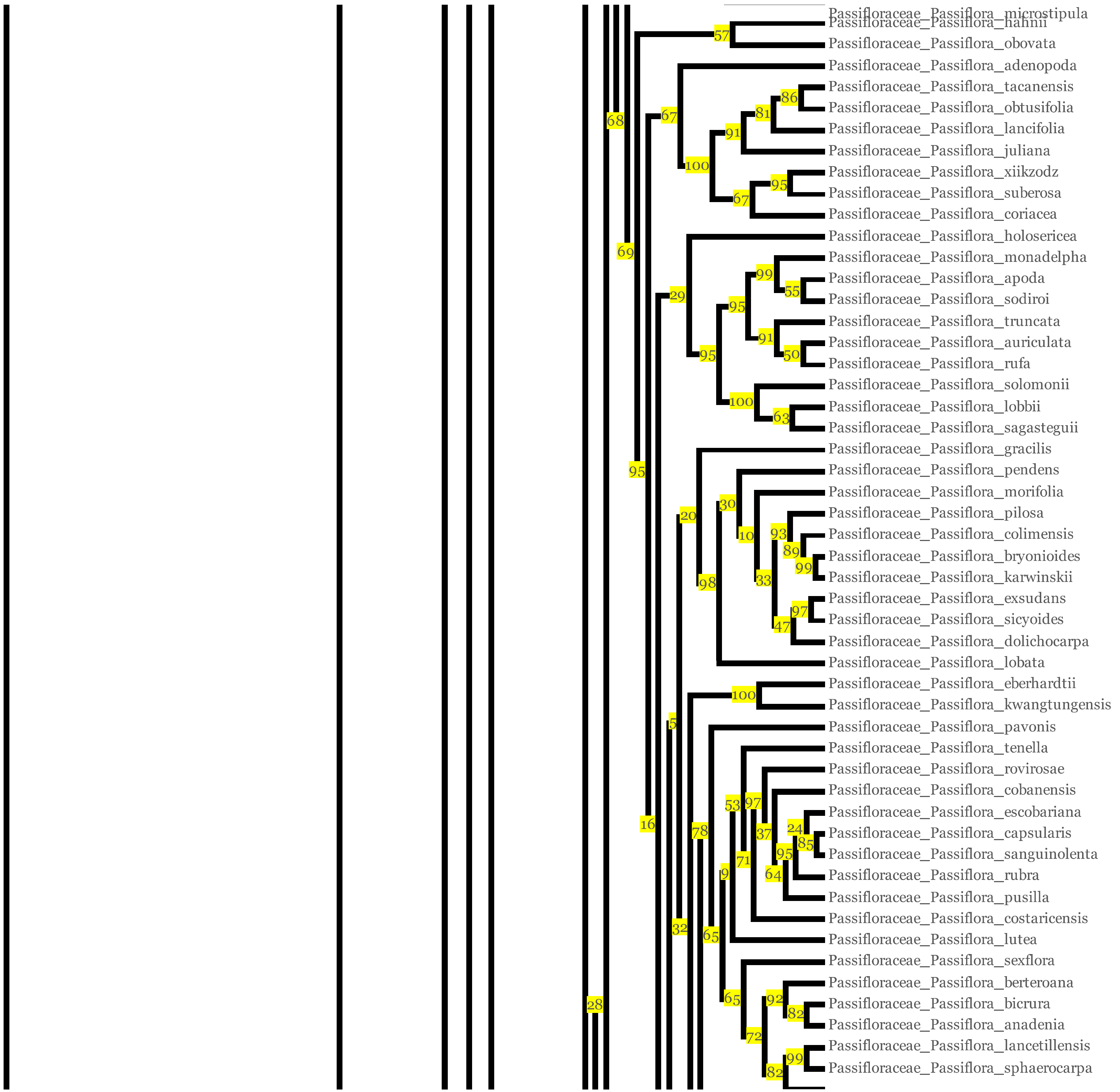
Passifloraceae_Passiflora_microstipula
Passifloraceae_Passiflora_hahnii
Passifloraceae_Passiflora_obovata
Passifloraceae_Passiflora_adenopoda
Passifloraceae_Passiflora_tacanensis
Passifloraceae_Passiflora_obtusifolia
Passifloraceae_Passiflora_lancifolia
Passifloraceae_Passiflora_juliana
Passifloraceae_Passiflora_xiikzodz
Passifloraceae_Passiflora_suberosa
Passifloraceae_Passiflora_coriacea
Passifloraceae_Passiflora_holosericea
Passifloraceae_Passiflora_monadelpha
Passifloraceae_Passiflora_apoda
Passifloraceae_Passiflora_sodiroi
Passifloraceae_Passiflora_truncata
Passifloraceae_Passiflora_auriculata
Passifloraceae_Passiflora_rufa
Passifloraceae_Passiflora_solomonii
Passifloraceae_Passiflora_lobbii
Passifloraceae_Passiflora_sagasteguii
Passifloraceae_Passiflora_gracilis
Passifloraceae_Passiflora_pendens
Passifloraceae_Passiflora_morifolia
Passifloraceae_Passiflora_pilosa
Passifloraceae_Passiflora_colimensis
Passifloraceae_Passiflora_bryonioides
Passifloraceae_Passiflora_karwinskii
Passifloraceae_Passiflora_exsudans
Passifloraceae_Passiflora_sicyoides
Passifloraceae_Passiflora_dolichocarpa
Passifloraceae_Passiflora_lobata
Passifloraceae_Passiflora_eberhardtii
Passifloraceae_Passiflora_kwangtungensis
Passifloraceae_Passiflora_pavonis
Passifloraceae_Passiflora_tenella
Passifloraceae_Passiflora_rovirosae
Passifloraceae_Passiflora_cobanensis
Passifloraceae_Passiflora_escobariana
Passifloraceae_Passiflora_capsularis
Passifloraceae_Passiflora_sanguinolenta
Passifloraceae_Passiflora_rubra
Passifloraceae_Passiflora_pusilla
Passifloraceae_Passiflora_costaricensis
Passifloraceae_Passiflora_lutea
Passifloraceae_Passiflora_sexflora
Passifloraceae_Passiflora_berteroana
Passifloraceae_Passiflora_bicrura
Passifloraceae_Passiflora_anadenia
Passifloraceae_Passiflora_lancetillensis
Passifloraceae_Passiflora_sphaerocarpa

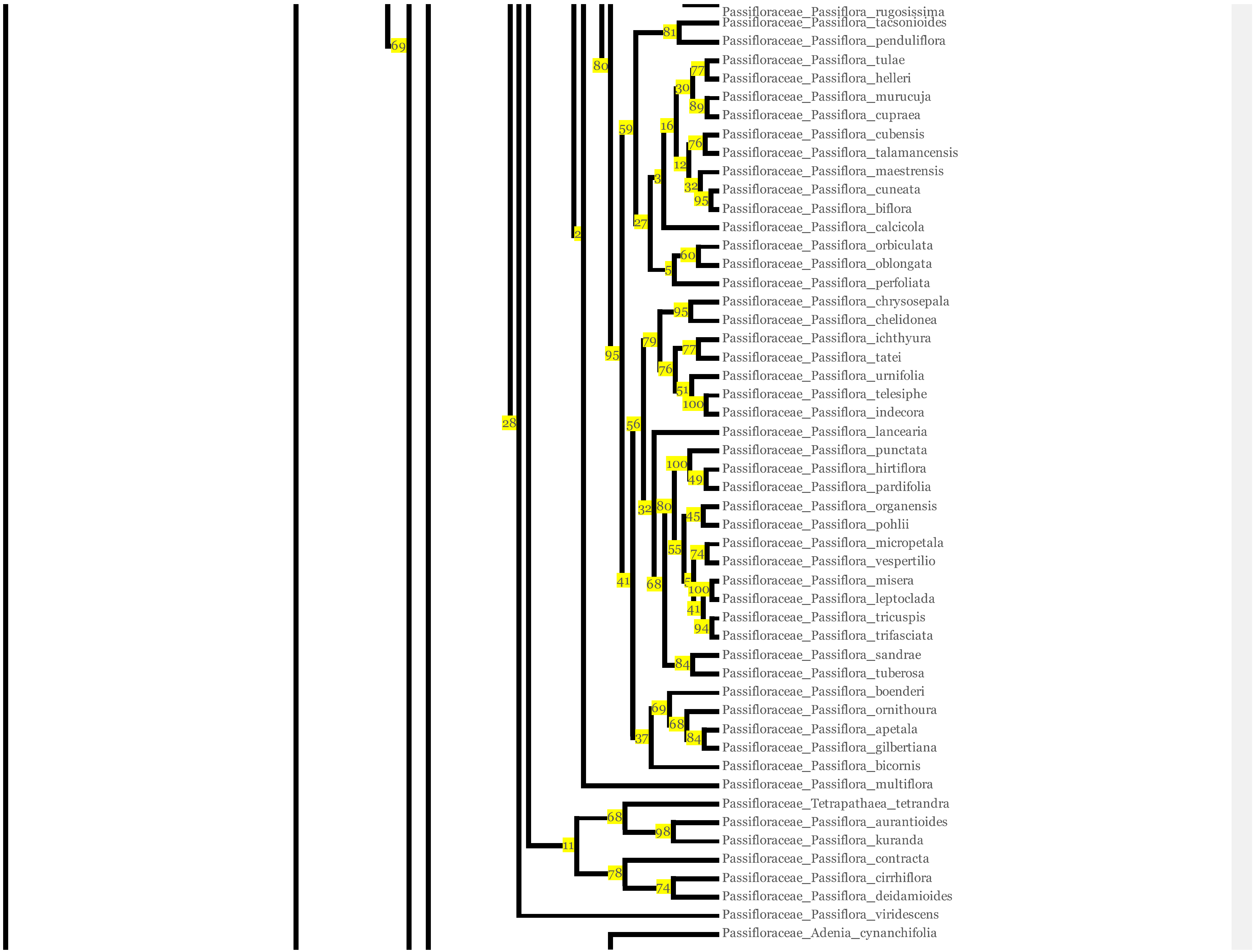

Passifloraceae_Passiflora_rugosissima
Passifloraceae_Passiflora_tacsonioides
Passifloraceae_Passiflora_penduliflora
Passifloraceae_Passiflora_tulae
Passifloraceae_Passiflora_helleri
Passifloraceae_Passiflora_murucuja
Passifloraceae_Passiflora_cupraea
Passifloraceae_Passiflora_cubensis
Passifloraceae_Passiflora_talamancensis
Passifloraceae_Passiflora_maestrensis
Passifloraceae_Passiflora_cuneata
Passifloraceae_Passiflora_biflora
Passifloraceae_Passiflora_calcicola
Passifloraceae_Passiflora_orbiculata
Passifloraceae_Passiflora_oblongata
Passifloraceae_Passiflora_perfoliata
Passifloraceae_Passiflora_chrysosepala
Passifloraceae_Passiflora_chelidonea
Passifloraceae_Passiflora_ichthyura
Passifloraceae_Passiflora_tatei
Passifloraceae_Passiflora_urnifolia
Passifloraceae_Passiflora_telesiphe
Passifloraceae_Passiflora_indecora
Passifloraceae_Passiflora_lancearia
Passifloraceae_Passiflora_punctata
Passifloraceae_Passiflora_hirtiflora
Passifloraceae_Passiflora_pardifolia
Passifloraceae_Passiflora_organensis
Passifloraceae_Passiflora_pohlii
Passifloraceae_Passiflora_micropetala
Passifloraceae_Passiflora_vespertilio
Passifloraceae_Passiflora_misera
Passifloraceae_Passiflora_leptoclada
Passifloraceae_Passiflora_tricuspis
Passifloraceae_Passiflora_trifasciata
Passifloraceae_Passiflora_sandrae
Passifloraceae_Passiflora_tuberosa
Passifloraceae_Passiflora_boenderi
Passifloraceae_Passiflora_ornithoura
Passifloraceae_Passiflora_apetala
Passifloraceae_Passiflora_gilbertiana
Passifloraceae_Passiflora_bicornis
Passifloraceae_Passiflora_multiflora
Passifloraceae_Tetrapathaea_tetrandra
Passifloraceae_Passiflora_aurantioides
Passifloraceae_Passiflora_kuranda
Passifloraceae_Passiflora_contracta
Passifloraceae_Passiflora_cirrhiflora
Passifloraceae_Passiflora_deidamioides
Passifloraceae_Passiflora_viridescens
Passifloraceae_Adenia_cynanchifolia

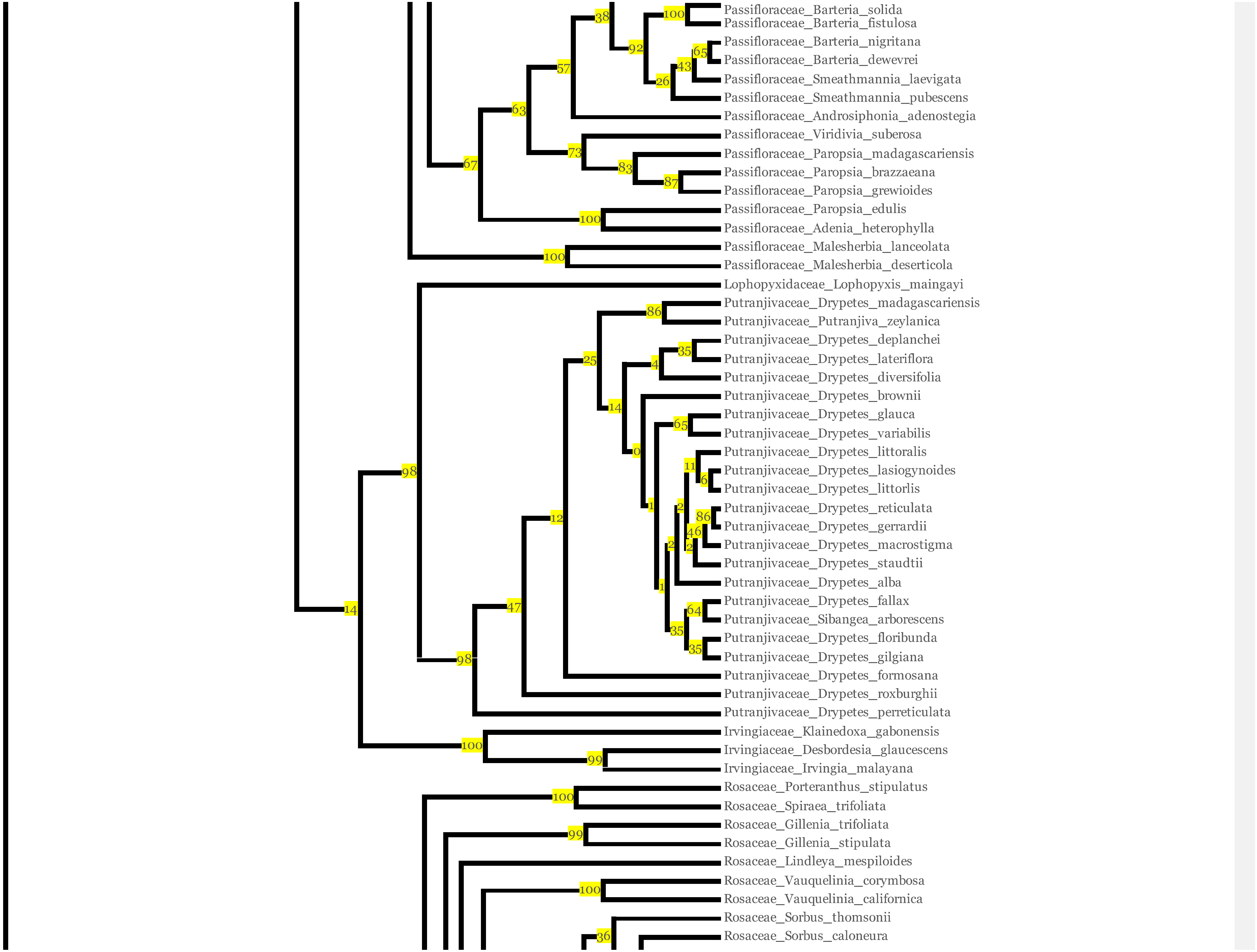
Passifloraceae_Barteria_solida
Passifloraceae_Barteria_fistulosa
Passifloraceae_Barteria_nigritana
Passifloraceae_Barteria_dewevrei
Passifloraceae_Smeathmannia_laevigata
Passifloraceae_Smeathmannia_pubescens
Passifloraceae_Androsiphonia_adenostegia
Passifloraceae_Viridivia_suberosa
Passifloraceae_Paropsia_madagascariensis
Passifloraceae_Paropsia_brazzaeana
Passifloraceae_Paropsia_grewioides
Passifloraceae_Paropsia_edulis
Passifloraceae_Adenia_heterophylla
Passifloraceae_Malesherbia_lanceolata
Passifloraceae_Malesherbia_deserticola
Lophopyxidaceae_Lophopyxis_maingayi
Putranjivaceae_Drypetes_madagascariensis
Putranjivaceae_Putranjiva_zeylanica
Putranjivaceae_Drypetes_deplanchei
Putranjivaceae_Drypetes_lateriflora
Putranjivaceae_Drypetes_diversifolia
Putranjivaceae_Drypetes_brownii
Putranjivaceae_Drypetes_glauca
Putranjivaceae_Drypetes_variabilis
Putranjivaceae_Drypetes_littoralis
Putranjivaceae_Drypetes_lasiogynoides
Putranjivaceae_Drypetes_littorlis
Putranjivaceae_Drypetes_reticulata
Putranjivaceae_Drypetes_gerrardii
Putranjivaceae_Drypetes_macrostigma
Putranjivaceae_Drypetes_staudtii
Putranjivaceae_Drypetes_alba
Putranjivaceae_Drypetes_fallax
Putranjivaceae_Sibangea_arborescens
Putranjivaceae_Drypetes_floribunda
Putranjivaceae_Drypetes_gilgiana
Putranjivaceae_Drypetes_formosana
Putranjivaceae_Drypetes_roxburghii
Putranjivaceae_Drypetes_perreticulata
Irvingiaceae_Klainedoxa_gabonensis
Irvingiaceae_Desbordesia_glaucescens
Irvingiaceae_Irvingia_malayana
Rosaceae_Porteranthus_stipulatus
Rosaceae_Spiraea_trifoliata
Rosaceae_Gillenia_trifoliata
Rosaceae_Gillenia_stipulata
Rosaceae_Lindleya_mespiloides
Rosaceae_Vauquelinia_corymbosa
Rosaceae_Vauquelinia_californica
Rosaceae_Sorbus_thomsonii
Rosaceae_Sorbus_caloneura

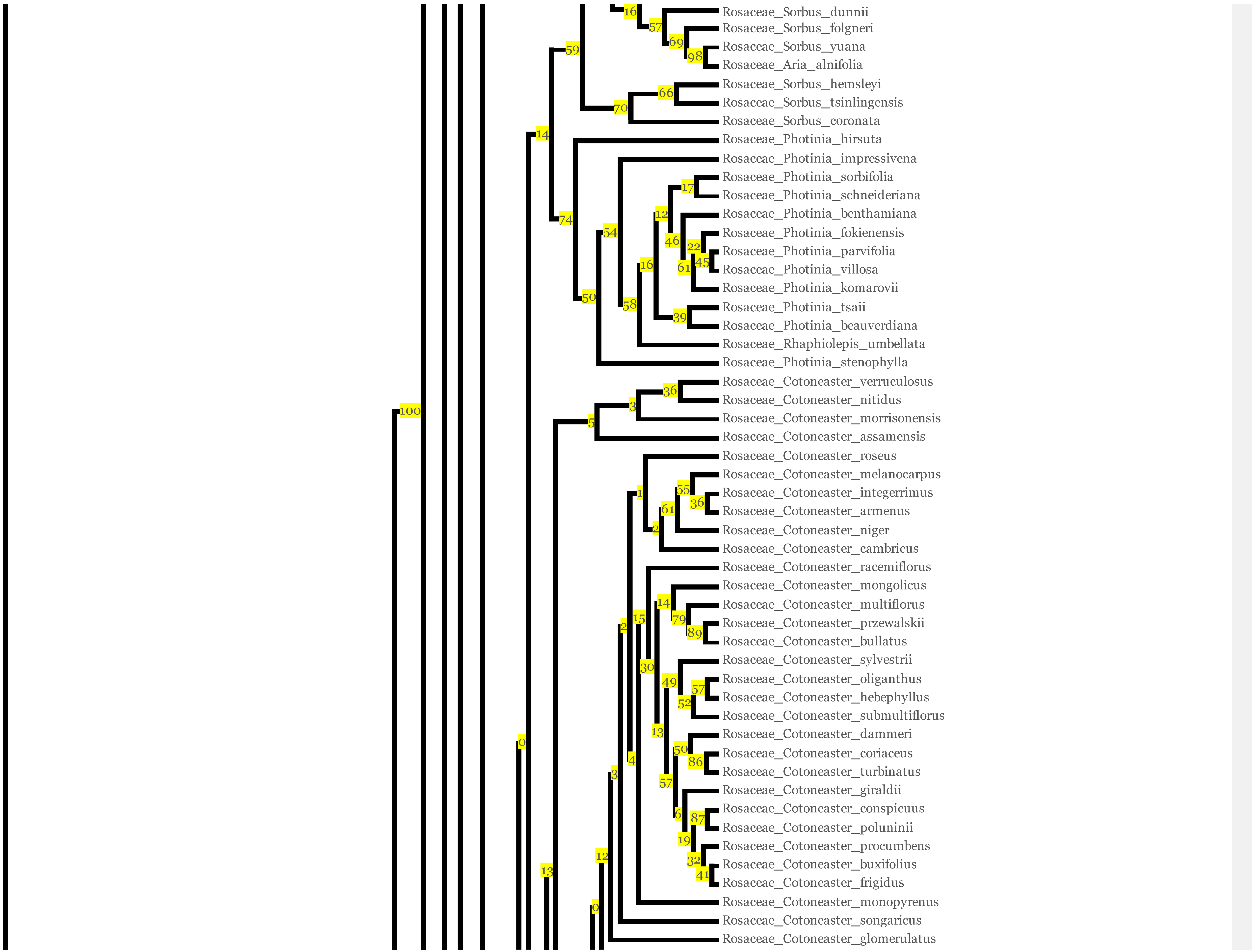
Rosaceae_Sorbus_dunnii
Rosaceae_Sorbus_folgneri
Rosaceae_Sorbus_yuana
Rosaceae_Aria_alnifolia
Rosaceae_Sorbus_hemsleyi
Rosaceae_Sorbus_tsinlingensis
Rosaceae_Sorbus_coronata
Rosaceae_Photinia_hirsuta
Rosaceae_Photinia_impressivena
Rosaceae_Photinia_sorbifolia
Rosaceae_Photinia_schneideriana
Rosaceae_Photinia_benthamiana
Rosaceae_Photinia_fokienensis
Rosaceae_Photinia_parvifolia
Rosaceae_Photinia_villosa
Rosaceae_Photinia_komarovii
Rosaceae_Photinia_tsaii
Rosaceae_Photinia_beauverdiana
Rosaceae_Rhaphiolepis_umbellata
Rosaceae_Photinia_stenophylla
Rosaceae_Cotoneaster_verruculosus
Rosaceae_Cotoneaster_nitidus
Rosaceae_Cotoneaster_morrisonensis
Rosaceae_Cotoneaster_assamensis
Rosaceae_Cotoneaster_roseus
Rosaceae_Cotoneaster_melanocarpus
Rosaceae_Cotoneaster_integerrimus
Rosaceae_Cotoneaster_armenus
Rosaceae_Cotoneaster_niger
Rosaceae_Cotoneaster_cambricus
Rosaceae_Cotoneaster_racemiflorus
Rosaceae_Cotoneaster_mongolicus
Rosaceae_Cotoneaster_multiflorus
Rosaceae_Cotoneaster_przewalskii
Rosaceae_Cotoneaster_bullatus
Rosaceae_Cotoneaster_sylvestrii
Rosaceae_Cotoneaster_oliganthus
Rosaceae_Cotoneaster_hebephyllus
Rosaceae_Cotoneaster_submultiflorus
Rosaceae_Cotoneaster_dammeri
Rosaceae_Cotoneaster_coriaceus
Rosaceae_Cotoneaster_turbinatus
Rosaceae_Cotoneaster_giraldii
Rosaceae_Cotoneaster_conspicuus
Rosaceae_Cotoneaster_poluninii
Rosaceae_Cotoneaster_procumbens
Rosaceae_Cotoneaster_buxifolius
Rosaceae_Cotoneaster_frigidus
Rosaceae_Cotoneaster_monopyrenus
Rosaceae_Cotoneaster_songaricus
Rosaceae_Cotoneaster_glomerulatus

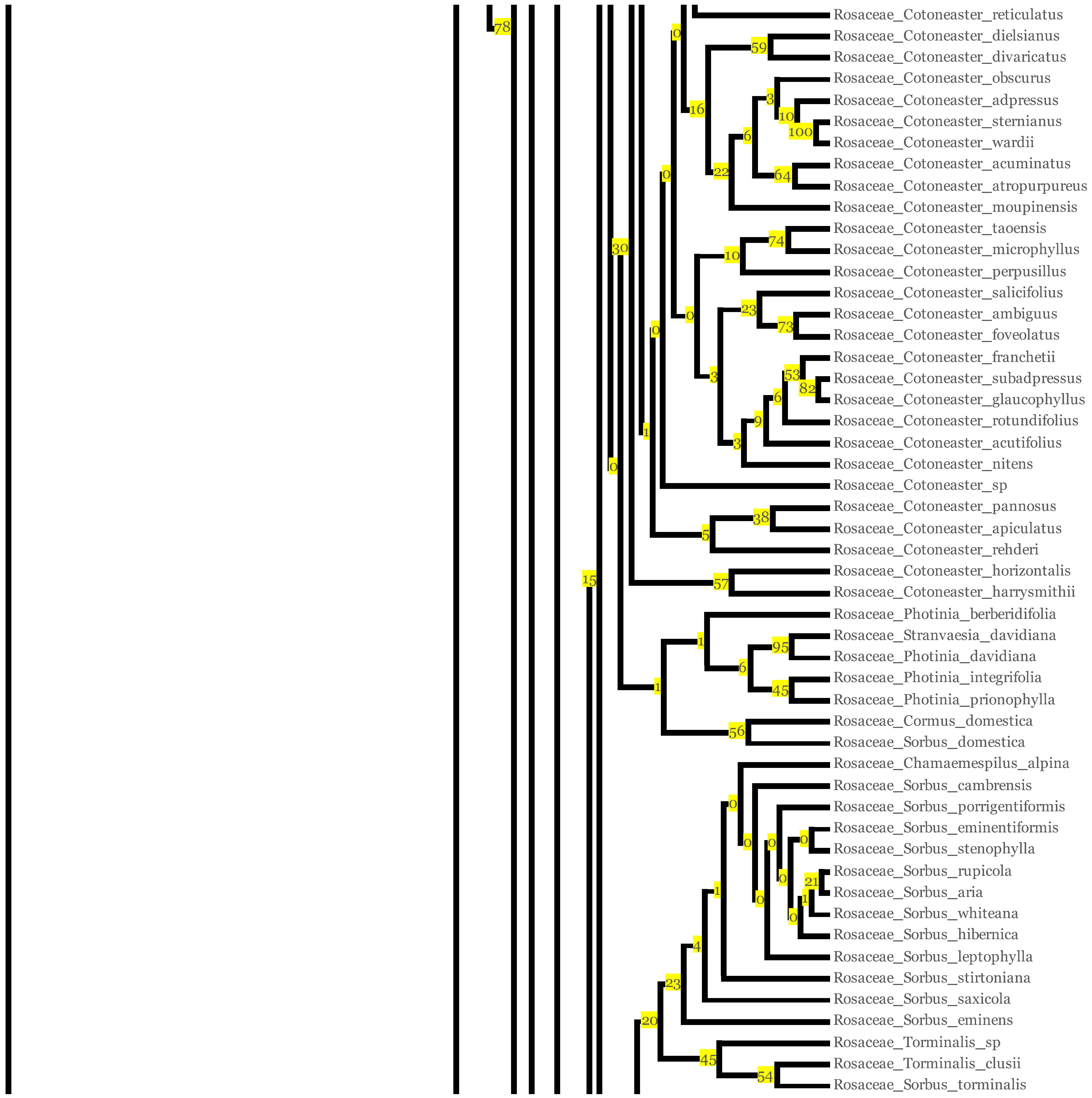
Rosaceae_Cotoneaster_reticulatus
Rosaceae_Cotoneaster_dielsianus
Rosaceae_Cotoneaster_divaricatus
Rosaceae_Cotoneaster_obscurus
Rosaceae_Cotoneaster_adpressus
Rosaceae_Cotoneaster_sternianus
Rosaceae_Cotoneaster_wardii
Rosaceae_Cotoneaster_acuminatus
Rosaceae_Cotoneaster_atropurpureus
Rosaceae_Cotoneaster_moupinensis
Rosaceae_Cotoneaster_taoensis
Rosaceae_Cotoneaster_microphyllus
Rosaceae_Cotoneaster_perpusillus
Rosaceae_Cotoneaster_salicifolius
Rosaceae_Cotoneaster_ambiguus
Rosaceae_Cotoneaster_foveolatus
Rosaceae_Cotoneaster_franchetii
Rosaceae_Cotoneaster_subadpressus
Rosaceae_Cotoneaster_glaucophyllus
Rosaceae_Cotoneaster_rotundifolius
Rosaceae_Cotoneaster_acutifolius
Rosaceae_Cotoneaster_nitens
Rosaceae_Cotoneaster_sp
Rosaceae_Cotoneaster_pannosus
Rosaceae_Cotoneaster_apiculatus
Rosaceae_Cotoneaster_rehderi
Rosaceae_Cotoneaster_horizontalis
Rosaceae_Cotoneaster_harrysmithii
Rosaceae_Photinia_berberidifolia
Rosaceae_Stranvaesia_davidiana
Rosaceae_Photinia_davidiana
Rosaceae_Photinia_integrifolia
Rosaceae_Photinia_prionophylla
Rosaceae_Cormus_domestica
Rosaceae_Sorbus_domestica
Rosaceae_Chamaemespilus_alpina
Rosaceae_Sorbus_cambrensis
Rosaceae_Sorbus_porrigentiformis
Rosaceae_Sorbus_eminentiformis
Rosaceae_Sorbus_stenophylla
Rosaceae_Sorbus_rupicola
Rosaceae_Sorbus_aria
Rosaceae_Sorbus_whiteana
Rosaceae_Sorbus_hibernica
Rosaceae_Sorbus_leptophylla
Rosaceae_Sorbus_stirtoniana
Rosaceae_Sorbus_saxicola
Rosaceae_Sorbus_eminens
Rosaceae_Torminalis_sp
Rosaceae_Torminalis_clusii
Rosaceae_Sorbus_torminalis

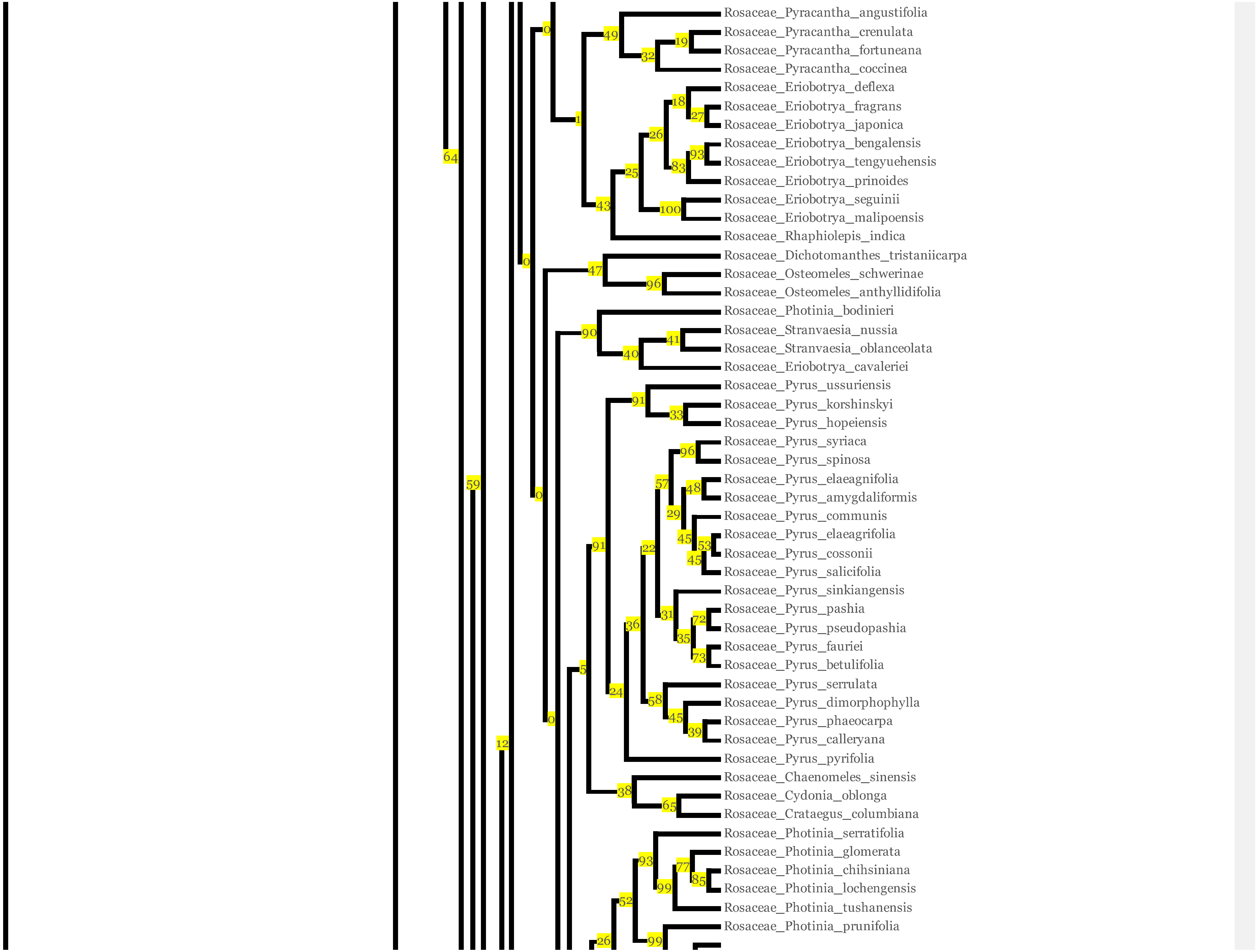
Rosaceae_Pyracantha_angustifolia
Rosaceae_Pyracantha_crenulata
Rosaceae_Pyracantha_fortuneana
Rosaceae_Pyracantha_coccinea
Rosaceae_Eriobotrya_deflexa
Rosaceae_Eriobotrya_fragrans
Rosaceae_Eriobotrya_japonica
Rosaceae_Eriobotrya_bengalensis
Rosaceae_Eriobotrya_tengyuehensis
Rosaceae_Eriobotrya_prinoides
Rosaceae_Eriobotrya_seguinii
Rosaceae_Eriobotrya_malipoensis
Rosaceae_Rhaphiolepis_indica
Rosaceae_Dichotomanthes_tristaniicarpa
Rosaceae_Osteomeles_schwerinae
Rosaceae_Osteomeles_anthyllidifolia
Rosaceae_Photinia_bodinieri
Rosaceae_Stranvaesia_nussia
Rosaceae_Stranvaesia_oblanceolata
Rosaceae_Eriobotrya_cavaleriei
Rosaceae_Pyrus_ussuriensis
Rosaceae_Pyrus_korshinskyi
Rosaceae_Pyrus_hopeiensis
Rosaceae_Pyrus_syriaca
Rosaceae_Pyrus_spinosa
Rosaceae_Pyrus_elaeagnifolia
Rosaceae_Pyrus_amygdaliformis
Rosaceae_Pyrus_communis
Rosaceae_Pyrus_elaeagrifolia
Rosaceae_Pyrus_cossonii
Rosaceae_Pyrus_salicifolia
Rosaceae_Pyrus_sinkiangensis
Rosaceae_Pyrus_pashia
Rosaceae_Pyrus_pseudopashia
Rosaceae_Pyrus_fauriei
Rosaceae_Pyrus_betulifolia
Rosaceae_Pyrus_serrulata
Rosaceae_Pyrus_dimorphophylla
Rosaceae_Pyrus_phaeocarpa
Rosaceae_Pyrus_calleryana
Rosaceae_Pyrus_pyrifolia
Rosaceae_Chaenomeles_sinensis
Rosaceae_Cydonia_oblonga
Rosaceae_Crataegus_columbiana
Rosaceae_Photinia_serratifolia
Rosaceae_Photinia_glomerata
Rosaceae_Photinia_chihsiniana
Rosaceae_Photinia_lochengensis
Rosaceae_Photinia_tushanensis
Rosaceae_Photinia_prunifolia

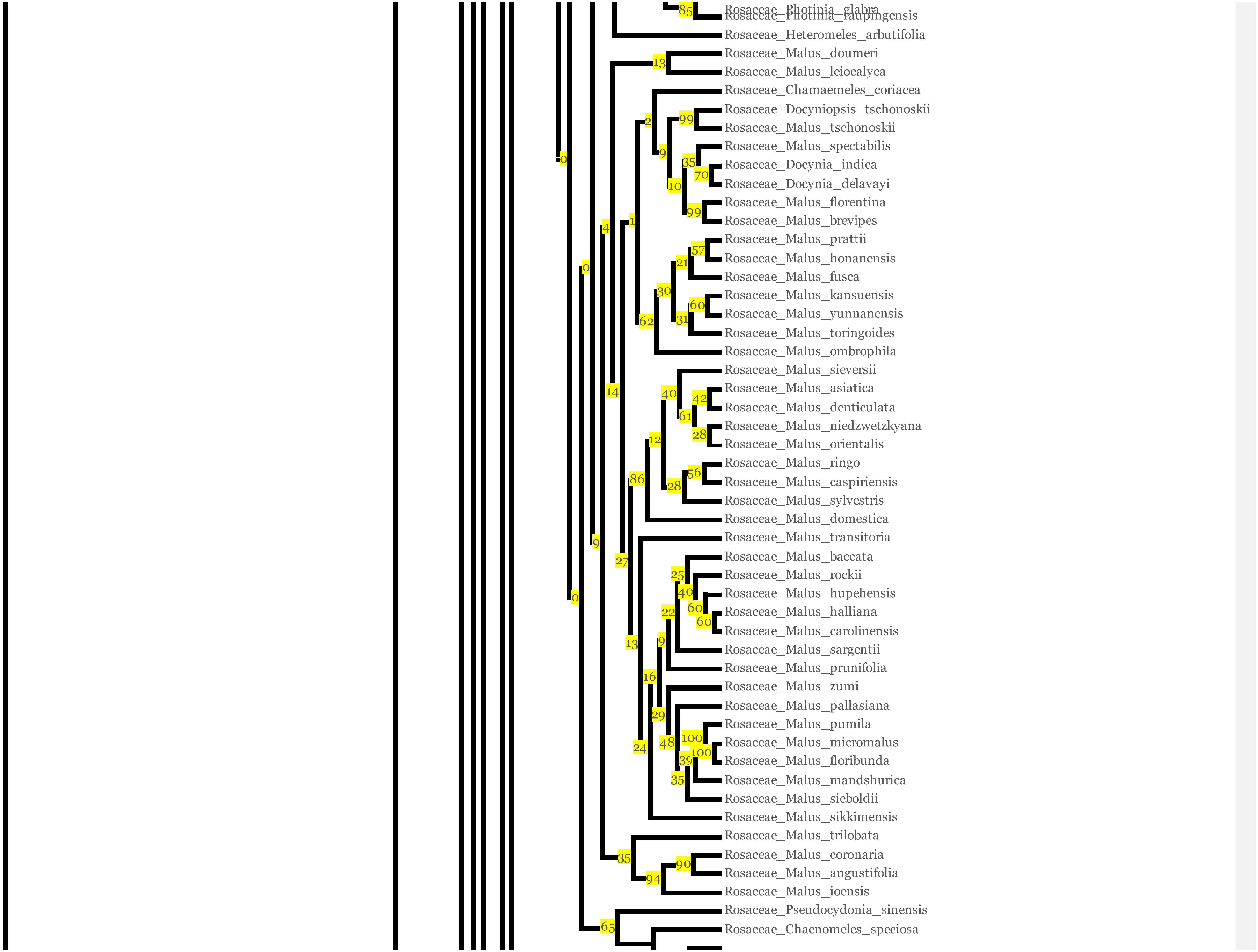
Rosaceae_Photinia_glabra
Rosaceae_Photinia_raupingensis
Rosaceae_Heteromeles_arbutifolia
Rosaceae_Malus_doumeri
Rosaceae_Malus_leiocalyca
Rosaceae_Chamaemeles_coriacea
Rosaceae_Docyniopsis_tschonoskii
Rosaceae_Malus_tschonoskii
Rosaceae_Malus_spectabilis
Rosaceae_Docynia_indica
Rosaceae_Docynia_delavayi
Rosaceae_Malus_florentina
Rosaceae_Malus_brevipes
Rosaceae_Malus_prattii
Rosaceae_Malus_honanensis
Rosaceae_Malus_fusca
Rosaceae_Malus_kansuensis
Rosaceae_Malus_yunnanensis
Rosaceae_Malus_toringoides
Rosaceae_Malus_ombrophila
Rosaceae_Malus_sieversii
Rosaceae_Malus_asiatica
Rosaceae_Malus_denticulata
Rosaceae_Malus_niedzwetzkyana
Rosaceae_Malus_orientalis
Rosaceae_Malus_ringo
Rosaceae_Malus_caspiriensis
Rosaceae_Malus_sylvestris
Rosaceae_Malus_domestica
Rosaceae_Malus_transitoria
Rosaceae_Malus_baccata
Rosaceae_Malus_rockii
Rosaceae_Malus_hupehensis
Rosaceae_Malus_halliana
Rosaceae_Malus_carolinensis
Rosaceae_Malus_sargentii
Rosaceae_Malus_prunifolia
Rosaceae_Malus_zumi
Rosaceae_Malus_pallasiana
Rosaceae_Malus_pumila
Rosaceae_Malus_micromalus
Rosaceae_Malus_floribunda
Rosaceae_Malus_mandshurica
Rosaceae_Malus_sieboldii
Rosaceae_Malus_sikkimensis
Rosaceae_Malus_trilobata
Rosaceae_Malus_coronaria
Rosaceae_Malus_angustifolia
Rosaceae_Malus_ioensis
Rosaceae_Pseudocydonia_sinensis
Rosaceae_Chaenomeles_speciosa
85
13
2
99
9
35
70
10
99
0
1
4
0
57
21
30
60
31
62
14
40
42
61
28
12
86
56
28
9
27
25
40
60
60
22
0
9
13
16
29
100
48
100
39
35
24
35
90
94
65

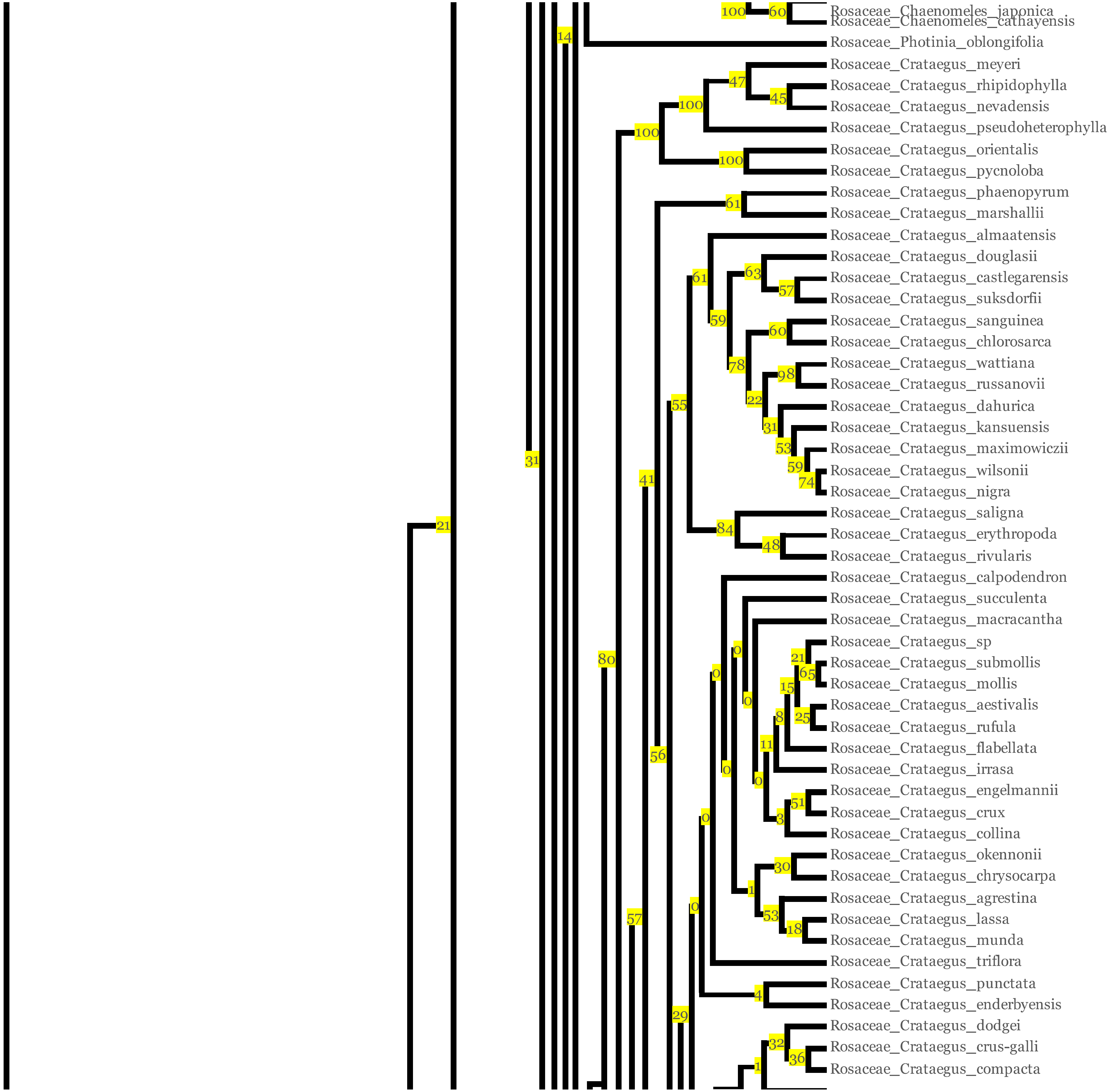
Rosaceae_Chaenomeles_japonica
Rosaceae_Chaenomeles_cathayensis
Rosaceae_Photinia_oblongifolia
Rosaceae_Crataegus_meyeri
Rosaceae_Crataegus_rhipidophylla
Rosaceae_Crataegus_nevadensis
Rosaceae_Crataegus_pseudoheterophylla
Rosaceae_Crataegus_orientalis
Rosaceae_Crataegus_pycnoloba
Rosaceae_Crataegus_phaenopyrum
Rosaceae_Crataegus_marshallii
Rosaceae_Crataegus_almaatensis
Rosaceae_Crataegus_douglasii
Rosaceae_Crataegus_castlegarensis
Rosaceae_Crataegus_suksdorfii
Rosaceae_Crataegus_sanguinea
Rosaceae_Crataegus_chlorosarca
Rosaceae_Crataegus_wattiana
Rosaceae_Crataegus_russanovii
Rosaceae_Crataegus_dahurica
Rosaceae_Crataegus_kansuensis
Rosaceae_Crataegus_maximowiczii
Rosaceae_Crataegus_wilsonii
Rosaceae_Crataegus_nigra
Rosaceae_Crataegus_saligna
Rosaceae_Crataegus_erythropoda
Rosaceae_Crataegus_rivularis
Rosaceae_Crataegus_calpodendron
Rosaceae_Crataegus_succulenta
Rosaceae_Crataegus_macracantha
Rosaceae_Crataegus_sp
Rosaceae_Crataegus_submollis
Rosaceae_Crataegus_mollis
Rosaceae_Crataegus_aestivalis
Rosaceae_Crataegus_rufula
Rosaceae_Crataegus_flabellata
Rosaceae_Crataegus_irrasa
Rosaceae_Crataegus_engelmannii
Rosaceae_Crataegus_crux
Rosaceae_Crataegus_collina
Rosaceae_Crataegus_okennonii
Rosaceae_Crataegus_chrysocarpa
Rosaceae_Crataegus_agrestina
Rosaceae_Crataegus_lassa
Rosaceae_Crataegus_munda
Rosaceae_Crataegus_triflora
Rosaceae_Crataegus_punctata
Rosaceae_Crataegus_enderbyensis
Rosaceae_Crataegus_dodgei
Rosaceae_Crataegus_crus-galli
Rosaceae_Crataegus_compacta

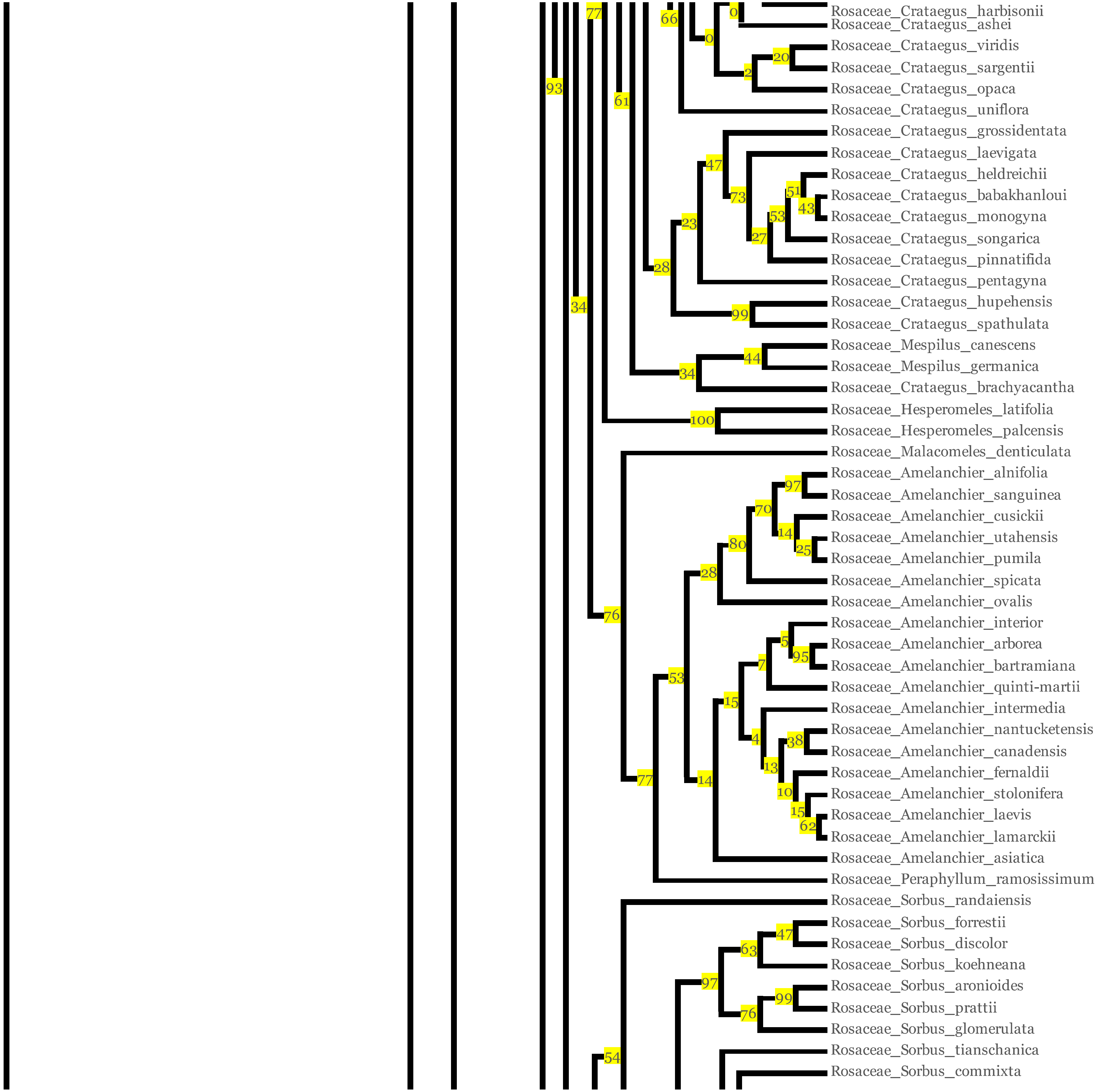
Rosaceae_Crataegus_harbisonii
Rosaceae_Crataegus_ashei
Rosaceae_Crataegus_viridis
Rosaceae_Crataegus_sargentii
Rosaceae_Crataegus_opaca
Rosaceae_Crataegus_uniflora
Rosaceae_Crataegus_grossidentata
Rosaceae_Crataegus_laevigata
Rosaceae_Crataegus_heldreichii
Rosaceae_Crataegus_babakhanloui
Rosaceae_Crataegus_monogyna
Rosaceae_Crataegus_songarica
Rosaceae_Crataegus_pinnatifida
Rosaceae_Crataegus_pentagyna
Rosaceae_Crataegus_hupehensis
Rosaceae_Crataegus_spathulata
Rosaceae_Mespilus_canescens
Rosaceae_Mespilus_germanica
Rosaceae_Crataegus_brachyacantha
Rosaceae_Hesperomeles_latifolia
Rosaceae_Hesperomeles_palcensis
Rosaceae_Malacomeles_denticulata
Rosaceae_Amelanchier_alnifolia
Rosaceae_Amelanchier_sanguinea
Rosaceae_Amelanchier_cusickii
Rosaceae_Amelanchier_utahensis
Rosaceae_Amelanchier_pumila
Rosaceae_Amelanchier_spicata
Rosaceae_Amelanchier_ovalis
Rosaceae_Amelanchier_interior
Rosaceae_Amelanchier_arborea
Rosaceae_Amelanchier_bartramiana
Rosaceae_Amelanchier_quinti-martii
Rosaceae_Amelanchier_intermedia
Rosaceae_Amelanchier_nantucketensis
Rosaceae_Amelanchier_canadensis
Rosaceae_Amelanchier_fernaldii
Rosaceae_Amelanchier_stolonifera
Rosaceae_Amelanchier_laevis
Rosaceae_Amelanchier_lamarckii
Rosaceae_Amelanchier_asiatica
Rosaceae_Peraphyllum_ramosissimum
Rosaceae_Sorbus_randaiensis
Rosaceae_Sorbus_forrestii
Rosaceae_Sorbus_discolor
Rosaceae_Sorbus_koehneana
Rosaceae_Sorbus_aronioides
Rosaceae_Sorbus_prattii
Rosaceae_Sorbus_glomerulata
Rosaceae_Sorbus_tianschanica
Rosaceae_Sorbus_commixta

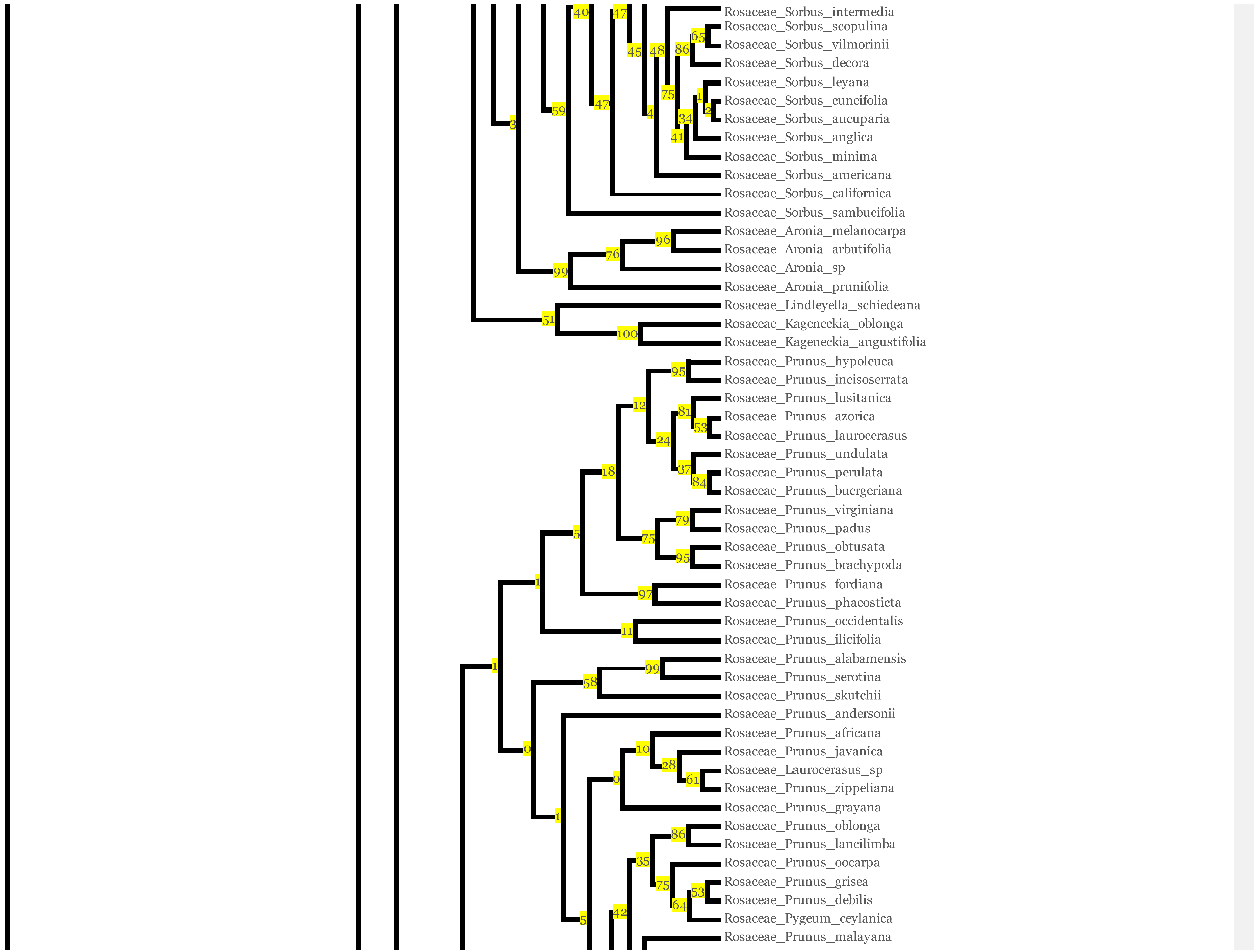
Rosaceae_Sorbus_intermedia
Rosaceae_Sorbus_scopulina
Rosaceae_Sorbus_vilmorinii
Rosaceae_Sorbus_decora
Rosaceae_Sorbus_leyana
Rosaceae_Sorbus_cuneifolia
Rosaceae_Sorbus_aucuparia
Rosaceae_Sorbus_anglica
Rosaceae_Sorbus_minima
Rosaceae_Sorbus_americana
Rosaceae_Sorbus_californica
Rosaceae_Sorbus_sambucifolia
Rosaceae_Aronia_melanocarpa
Rosaceae_Aronia_arbutifolia
Rosaceae_Aronia_sp
Rosaceae_Aronia_prunifolia
Rosaceae_Lindleyella_schiedeana
Rosaceae_Kageneckia_oblonga
Rosaceae_Kageneckia_angustifolia
Rosaceae_Prunus_hypoleuca
Rosaceae_Prunus_incisoserrata
Rosaceae_Prunus_lusitanica
Rosaceae_Prunus_azorica
Rosaceae_Prunus_laurocerasus
Rosaceae_Prunus_undulata
Rosaceae_Prunus_perulata
Rosaceae_Prunus_buergeriana
Rosaceae_Prunus_virginiana
Rosaceae_Prunus_padus
Rosaceae_Prunus_obtusata
Rosaceae_Prunus_brachypoda
Rosaceae_Prunus_fordiana
Rosaceae_Prunus_phaeosticta
Rosaceae_Prunus_occidentalis
Rosaceae_Prunus_ilicifolia
Rosaceae_Prunus_alabamensis
Rosaceae_Prunus_serotina
Rosaceae_Prunus_skutchii
Rosaceae_Prunus_andersonii
Rosaceae_Prunus_africana
Rosaceae_Prunus_javanica
Rosaceae_Laurocerasus_sp
Rosaceae_Prunus_zippeliana
Rosaceae_Prunus_grayana
Rosaceae_Prunus_oblonga
Rosaceae_Prunus_lancilimba
Rosaceae_Prunus_oocarpa
Rosaceae_Prunus_grisea
Rosaceae_Prunus_debilis
Rosaceae_Pygeum_ceylanica
Rosaceae_Prunus_malayana

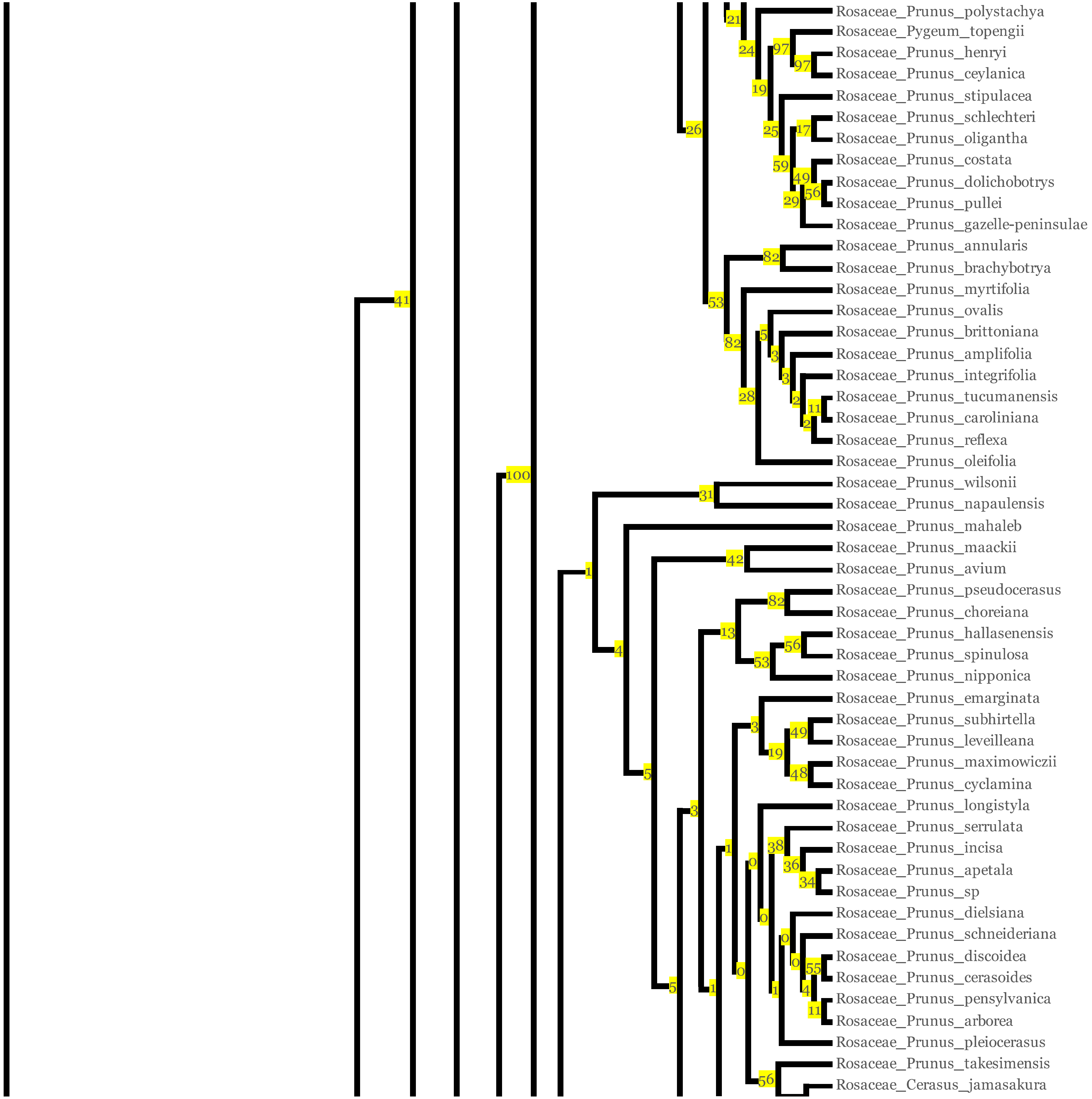
Rosaceae_Prunus_polystachya
Rosaceae_Pygeum_topengii
Rosaceae_Prunus_henryi
Rosaceae_Prunus_ceylanica
Rosaceae_Prunus_stipulacea
Rosaceae_Prunus_schlechteri
Rosaceae_Prunus_oligantha
Rosaceae_Prunus_costata
Rosaceae_Prunus_dolichobotrys
Rosaceae_Prunus_pullei
Rosaceae_Prunus_gazelle-peninsulae
Rosaceae_Prunus_annularis
Rosaceae_Prunus_brachybotrya
Rosaceae_Prunus_myrtifolia
Rosaceae_Prunus_ovalis
Rosaceae_Prunus_brittoniana
Rosaceae_Prunus_amplifolia
Rosaceae_Prunus_integrifolia
Rosaceae_Prunus_tucumanensis
Rosaceae_Prunus_caroliniana
Rosaceae_Prunus_reflexa
Rosaceae_Prunus_oleifolia
Rosaceae_Prunus_wilsonii
Rosaceae_Prunus_napaulensis
Rosaceae_Prunus_mahaleb
Rosaceae_Prunus_maackii
Rosaceae_Prunus_avium
Rosaceae_Prunus_pseudocerasus
Rosaceae_Prunus_choreiana
Rosaceae_Prunus_hallasenensis
Rosaceae_Prunus_spinulosa
Rosaceae_Prunus_nipponica
Rosaceae_Prunus_emarginata
Rosaceae_Prunus_subhirtella
Rosaceae_Prunus_leveilleana
Rosaceae_Prunus_maximowiczii
Rosaceae_Prunus_cyclamina
Rosaceae_Prunus_longistyla
Rosaceae_Prunus_serrulata
Rosaceae_Prunus_incisa
Rosaceae_Prunus_apetala
Rosaceae_Prunus_sp
Rosaceae_Prunus_dielsiana
Rosaceae_Prunus_schneideriana
Rosaceae_Prunus_discoidea
Rosaceae_Prunus_cerasoides
Rosaceae_Prunus_pensylvanica
Rosaceae_Prunus_arborea
Rosaceae_Prunus_pleiocerasus
Rosaceae_Prunus_takesimensis
Rosaceae_Cerasus_jamasakura

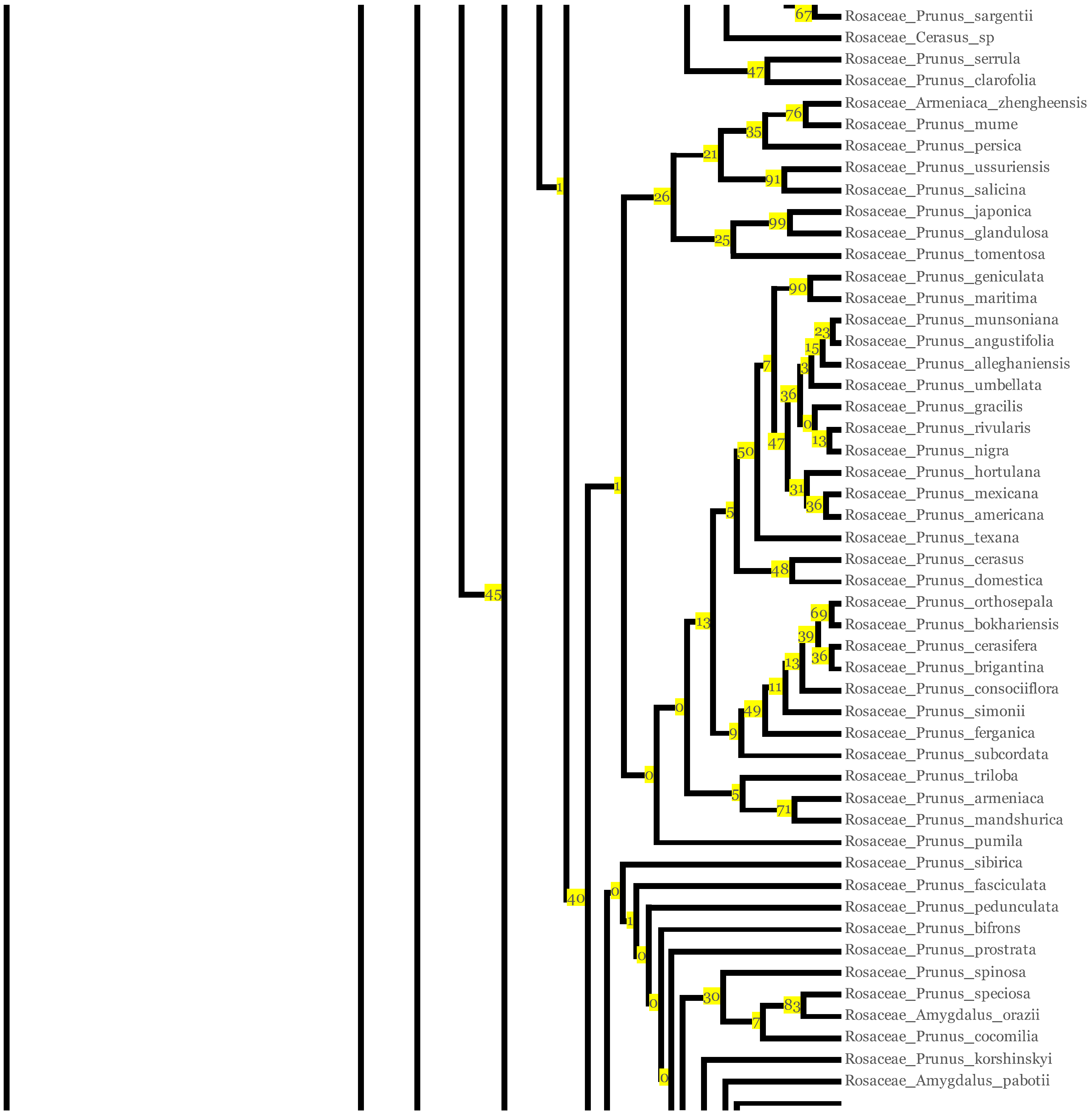
Rosaceae_Prunus_sargentii
Rosaceae_Cerasus_sp
Rosaceae_Prunus_serrula
Rosaceae_Prunus_clarofolia
Rosaceae_Armeniaca_zhengheensis
Rosaceae_Prunus_mume
Rosaceae_Prunus_persica
Rosaceae_Prunus_ussuriensis
Rosaceae_Prunus_salicina
Rosaceae_Prunus_japonica
Rosaceae_Prunus_glandulosa
Rosaceae_Prunus_tomentosa
Rosaceae_Prunus_geniculata
Rosaceae_Prunus_maritima
Rosaceae_Prunus_munsoniana
Rosaceae_Prunus_angustifolia
Rosaceae_Prunus_alleghaniensis
Rosaceae_Prunus_umbellata
Rosaceae_Prunus_gracilis
Rosaceae_Prunus_rivularis
Rosaceae_Prunus_nigra
Rosaceae_Prunus_hortulana
Rosaceae_Prunus_mexicana
Rosaceae_Prunus_americana
Rosaceae_Prunus_texana
Rosaceae_Prunus_cerasus
Rosaceae_Prunus_domestica
Rosaceae_Prunus_orthosepala
Rosaceae_Prunus_bokhariensis
Rosaceae_Prunus_cerasifera
Rosaceae_Prunus_brigantina
Rosaceae_Prunus_consociiflora
Rosaceae_Prunus_simonii
Rosaceae_Prunus_ferganica
Rosaceae_Prunus_subcordata
Rosaceae_Prunus_triloba
Rosaceae_Prunus_armeniaca
Rosaceae_Prunus_mandshurica
Rosaceae_Prunus_pumila
Rosaceae_Prunus_sibirica
Rosaceae_Prunus_fasciculata
Rosaceae_Prunus_pedunculata
Rosaceae_Prunus_bifrons
Rosaceae_Prunus_prostrata
Rosaceae_Prunus_spinosa
Rosaceae_Prunus_speciosa
Rosaceae_Amygdalus_orazii
Rosaceae_Prunus_cocomilia
Rosaceae_Prunus_korshinskyi
Rosaceae_Amygdalus_pabotii

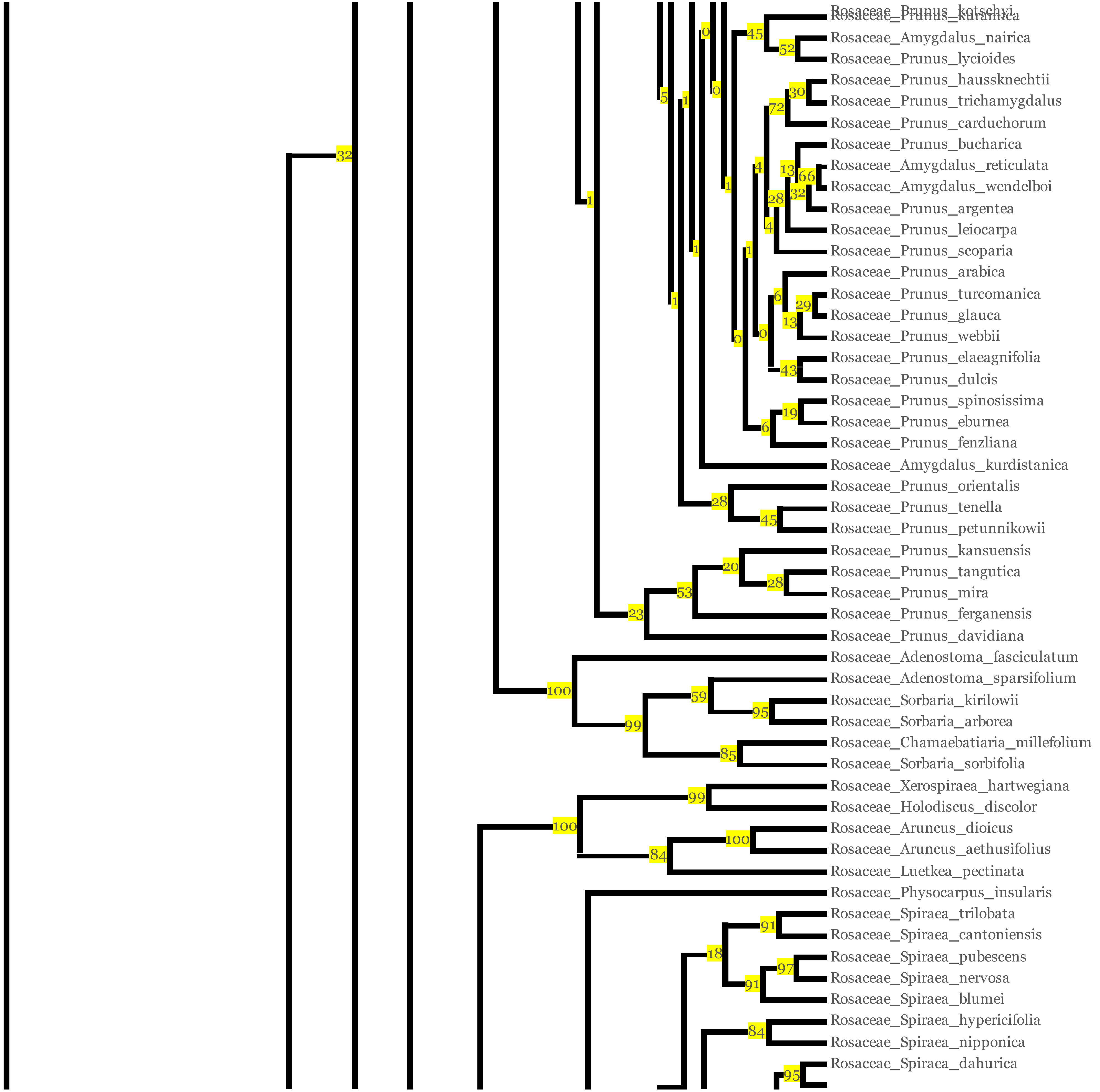
Rosaceae_Prunus_kotschyi
Rosaceae_Prunus_kuramica
Rosaceae_Amygdalus_nairica
Rosaceae_Prunus_lycioides
Rosaceae_Prunus_haussknechtii
Rosaceae_Prunus_trichamygdalus
Rosaceae_Prunus_carduchorum
Rosaceae_Prunus_bucharica
Rosaceae_Amygdalus_reticulata
Rosaceae_Amygdalus_wendelboi
Rosaceae_Prunus_argentea
Rosaceae_Prunus_leiocarpa
Rosaceae_Prunus_scoparia
Rosaceae_Prunus_arabica
Rosaceae_Prunus_turcomanica
Rosaceae_Prunus_glauca
Rosaceae_Prunus_webbii
Rosaceae_Prunus_elaeagnifolia
Rosaceae_Prunus_dulcis
Rosaceae_Prunus_spinosissima
Rosaceae_Prunus_eburnea
Rosaceae_Prunus_fenzliana
Rosaceae_Amygdalus_kurdistanica
Rosaceae_Prunus_orientalis
Rosaceae_Prunus_tenella
Rosaceae_Prunus_petunnikowii
Rosaceae_Prunus_kansuensis
Rosaceae_Prunus_tangutica
Rosaceae_Prunus_mira
Rosaceae_Prunus_ferganensis
Rosaceae_Prunus_davidiana
Rosaceae_Adenostoma_fasciculatum
Rosaceae_Adenostoma_sparsifolium
Rosaceae_Sorbaria_kirilowii
Rosaceae_Sorbaria_arborea
Rosaceae_Chamaebatiaria_millefolium
Rosaceae_Sorbaria_sorbifolia
Rosaceae_Xerospiraea_hartwegiana
Rosaceae_Holodiscus_discolor
Rosaceae_Aruncus_dioicus
Rosaceae_Aruncus_aethusifolius
Rosaceae_Luetkea_pectinata
Rosaceae_Physocarpus_insularis
Rosaceae_Spiraea_trilobata
Rosaceae_Spiraea_cantoniensis
Rosaceae_Spiraea_pubescens
Rosaceae_Spiraea_nervosa
Rosaceae_Spiraea_blumei
Rosaceae_Spiraea_hypericifolia
Rosaceae_Spiraea_nipponica
Rosaceae_Spiraea_dahurica

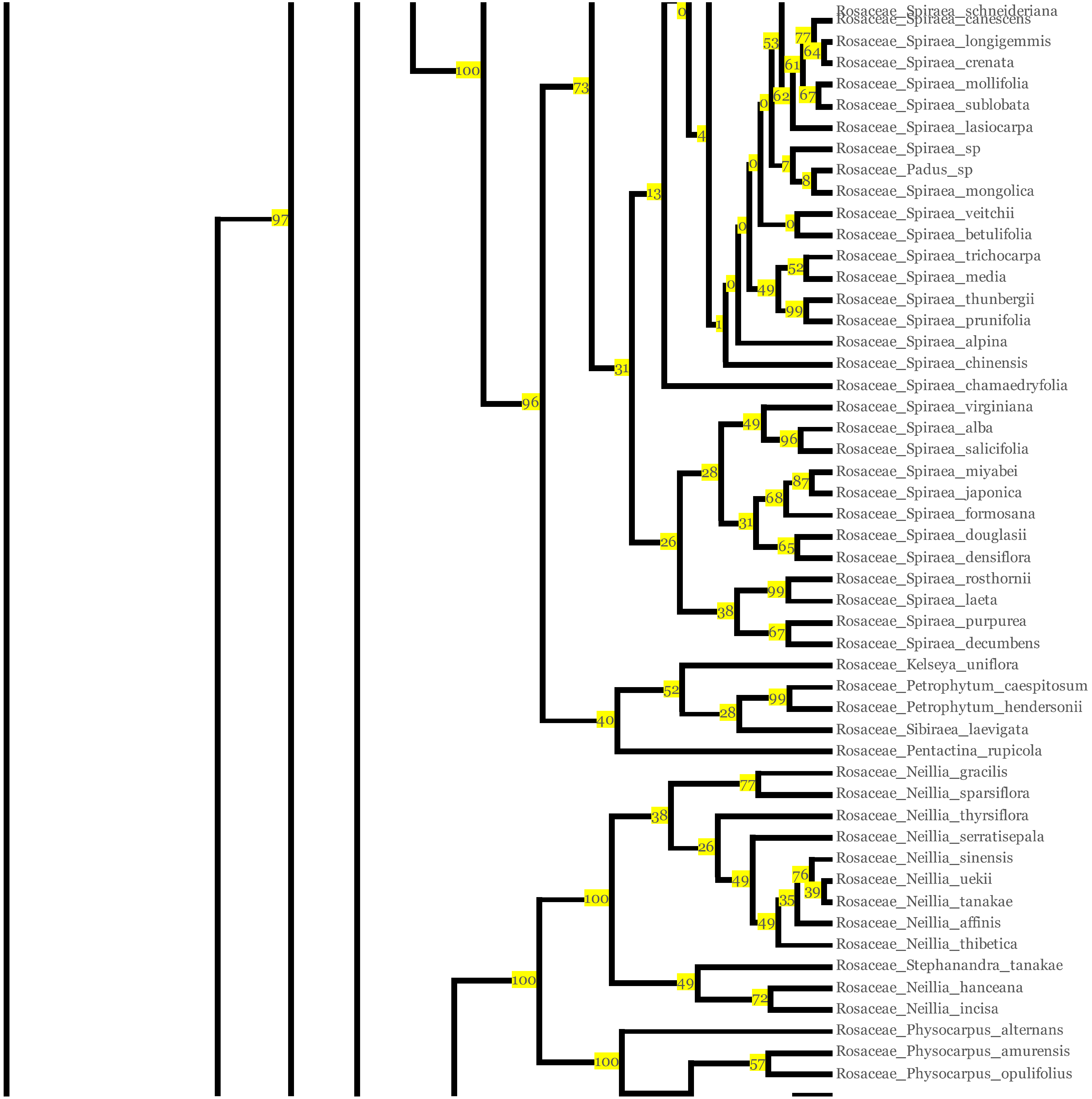
Rosaceae_Spiraea_schneideriana
Rosaceae_Spiraea_canescens
Rosaceae_Spiraea_longigemmis
Rosaceae_Spiraea_crenata
Rosaceae_Spiraea_mollifolia
Rosaceae_Spiraea_sublobata
Rosaceae_Spiraea_lasiocarpa
Rosaceae_Spiraea_sp
Rosaceae_Padus_sp
Rosaceae_Spiraea_mongolica
Rosaceae_Spiraea_veitchii
Rosaceae_Spiraea_betulifolia
Rosaceae_Spiraea_trichocarpa
Rosaceae_Spiraea_media
Rosaceae_Spiraea_thunbergii
Rosaceae_Spiraea_prunifolia
Rosaceae_Spiraea_alpina
Rosaceae_Spiraea_chinensis
Rosaceae_Spiraea_chamaedryfolia
Rosaceae_Spiraea_virginiana
Rosaceae_Spiraea_alba
Rosaceae_Spiraea_salicifolia
Rosaceae_Spiraea_miyabei
Rosaceae_Spiraea_japonica
Rosaceae_Spiraea_formosana
Rosaceae_Spiraea_douglasii
Rosaceae_Spiraea_densiflora
Rosaceae_Spiraea_rosthornii
Rosaceae_Spiraea_laeta
Rosaceae_Spiraea_purpurea
Rosaceae_Spiraea_decumbens
Rosaceae_Kelseya_uniflora
Rosaceae_Petrophytum_caespitosum
Rosaceae_Petrophytum_hendersonii
Rosaceae_Sibiraea_laevigata
Rosaceae_Pentactina_rupicola
Rosaceae_Neillia_gracilis
Rosaceae_Neillia_sparsiflora
Rosaceae_Neillia_thyrsiflora
Rosaceae_Neillia_serratisepala
Rosaceae_Neillia_sinensis
Rosaceae_Neillia_uekii
Rosaceae_Neillia_tanakae
Rosaceae_Neillia_affinis
Rosaceae_Neillia_thibetica
Rosaceae_Stephanandra_tanakae
Rosaceae_Neillia_hanceana
Rosaceae_Neillia_incisa
Rosaceae_Physocarpus_alternans
Rosaceae_Physocarpus_amurensis
Rosaceae_Physocarpus_opulifolius

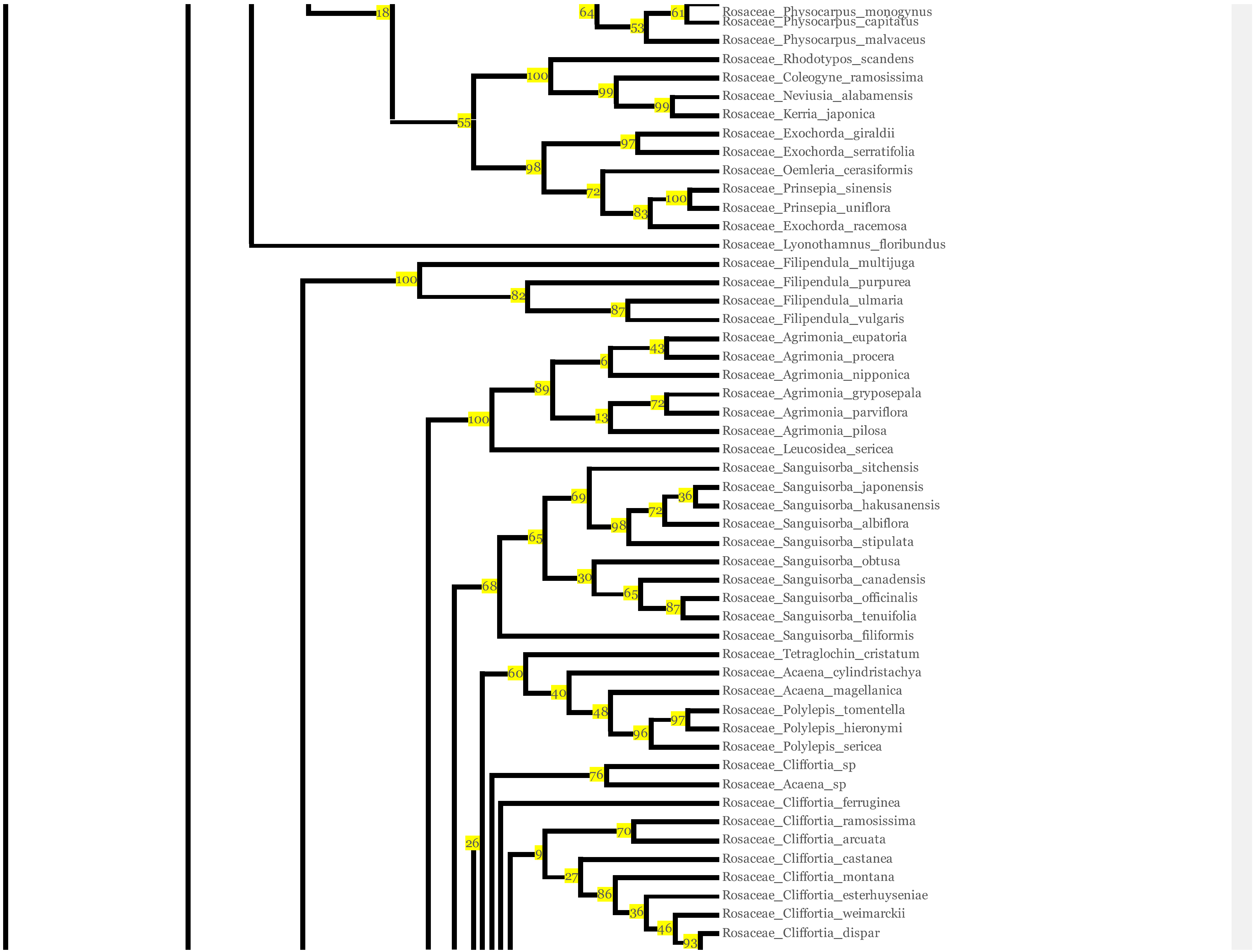
Rosaceae_Physocarpus_monogynus
Rosaceae_Physocarpus_capitatus
Rosaceae_Physocarpus_malvaceus
Rosaceae_Rhodotypos_scandens
Rosaceae_Coleogyne_ramosissima
Rosaceae_Neviusia_alabamensis
Rosaceae_Kerria_japonica
Rosaceae_Exochorda_giraldii
Rosaceae_Exochorda_serratifolia
Rosaceae_Oemleria_cerasiformis
Rosaceae_Prinsepia_sinensis
Rosaceae_Prinsepia_uniflora
Rosaceae_Exochorda_racemosa
Rosaceae_Lyonothamnus_floribundus
Rosaceae_Filipendula_multijuga
Rosaceae_Filipendula_purpurea
Rosaceae_Filipendula_ulmaria
Rosaceae_Filipendula_vulgaris
Rosaceae_Agrimonia_eupatoria
Rosaceae_Agrimonia_procera
Rosaceae_Agrimonia_nipponica
Rosaceae_Agrimonia_gryposepala
Rosaceae_Agrimonia_parviflora
Rosaceae_Agrimonia_pilosa
Rosaceae_Leucosidea_sericea
Rosaceae_Sanguisorba_sitchensis
Rosaceae_Sanguisorba_japonensis
Rosaceae_Sanguisorba_hakusanensis
Rosaceae_Sanguisorba_albiflora
Rosaceae_Sanguisorba_stipulata
Rosaceae_Sanguisorba_obtusa
Rosaceae_Sanguisorba_canadensis
Rosaceae_Sanguisorba_officinalis
Rosaceae_Sanguisorba_tenuifolia
Rosaceae_Sanguisorba_filiformis
Rosaceae_Tetraglochin_cristatum
Rosaceae_Acaena_cylindristachya
Rosaceae_Acaena_magellanica
Rosaceae_Polylepis_tomentella
Rosaceae_Polylepis_hieronymi
Rosaceae_Polylepis_sericea
Rosaceae_Cliffortia_sp
Rosaceae_Acaena_sp
Rosaceae_Cliffortia_ferruginea
Rosaceae_Cliffortia_ramosissima
Rosaceae_Cliffortia_arcuata
Rosaceae_Cliffortia_castanea
Rosaceae_Cliffortia_montana
Rosaceae_Cliffortia_esterhuyseniae
Rosaceae_Cliffortia_weimarckii
Rosaceae_Cliffortia_dispar

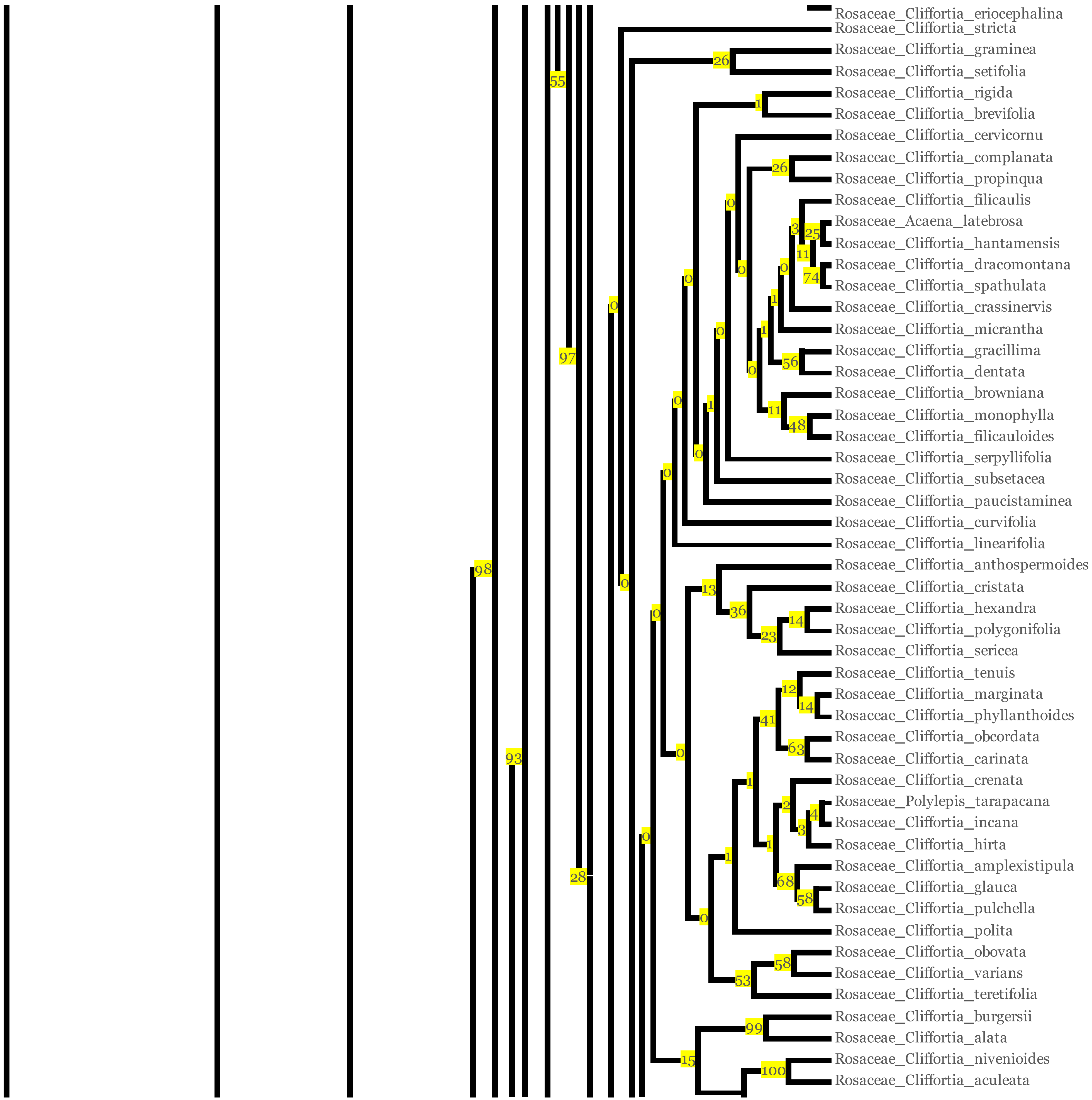

Rosaceae_Cliffortia_eriocephalina
Rosaceae_Cliffortia_stricta
Rosaceae_Cliffortia_graminea
Rosaceae_Cliffortia_setifolia
Rosaceae_Cliffortia_rigida
Rosaceae_Cliffortia_brevifolia
Rosaceae_Cliffortia_cervicornu
Rosaceae_Cliffortia_complanata
Rosaceae_Cliffortia_propinqua
Rosaceae_Cliffortia_filicaulis
Rosaceae_Acaena_latebrosa
Rosaceae_Cliffortia_hantamensis
Rosaceae_Cliffortia_dracomontana
Rosaceae_Cliffortia_spathulata
Rosaceae_Cliffortia_crassinervis
Rosaceae_Cliffortia_micrantha
Rosaceae_Cliffortia_gracillima
Rosaceae_Cliffortia_dentata
Rosaceae_Cliffortia_browniana
Rosaceae_Cliffortia_monophylla
Rosaceae_Cliffortia_filicauloides
Rosaceae_Cliffortia_serpyllifolia
Rosaceae_Cliffortia_subsetacea
Rosaceae_Cliffortia_paucistaminea
Rosaceae_Cliffortia_curvifolia
Rosaceae_Cliffortia_linearifolia
Rosaceae_Cliffortia_anthospermoides
Rosaceae_Cliffortia_cristata
Rosaceae_Cliffortia_hexandra
Rosaceae_Cliffortia_polygonifolia
Rosaceae_Cliffortia_sericea
Rosaceae_Cliffortia_tenuis
Rosaceae_Cliffortia_marginata
Rosaceae_Cliffortia_phyllanthoides
Rosaceae_Cliffortia_obcordata
Rosaceae_Cliffortia_carinata
Rosaceae_Cliffortia_crenata
Rosaceae_Polylepis_tarapacana
Rosaceae_Cliffortia_incana
Rosaceae_Cliffortia_hirta
Rosaceae_Cliffortia_amplexistipula
Rosaceae_Cliffortia_glauca
Rosaceae_Cliffortia_pulchella
Rosaceae_Cliffortia_polita
Rosaceae_Cliffortia_obovata
Rosaceae_Cliffortia_varians
Rosaceae_Cliffortia_teretifolia
Rosaceae_Cliffortia_burgersii
Rosaceae_Cliffortia_alata
Rosaceae_Cliffortia_nivenioides
Rosaceae_Cliffortia_aculeata

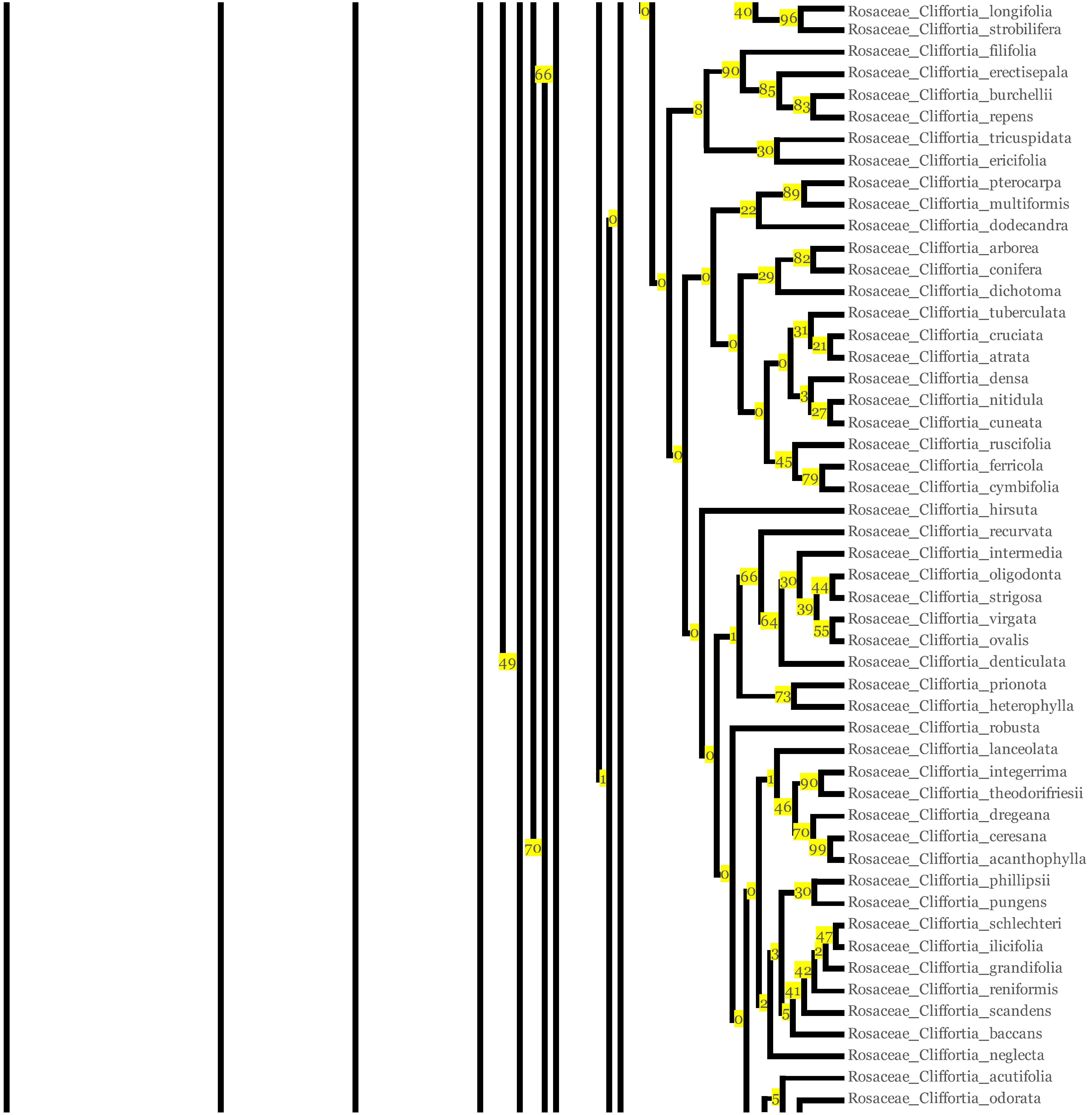
Rosaceae_Cliffortia_longifolia
Rosaceae_Cliffortia_strobilifera
Rosaceae_Cliffortia_filifolia
Rosaceae_Cliffortia_erectisepala
Rosaceae_Cliffortia_burchellii
Rosaceae_Cliffortia_repens
Rosaceae_Cliffortia_tricuspidata
Rosaceae_Cliffortia_ericifolia
Rosaceae_Cliffortia_pterocarpa
Rosaceae_Cliffortia_multiformis
Rosaceae_Cliffortia_dodecandra
Rosaceae_Cliffortia_arborea
Rosaceae_Cliffortia_conifera
Rosaceae_Cliffortia_dichotoma
Rosaceae_Cliffortia_tuberculata
Rosaceae_Cliffortia_cruciata
Rosaceae_Cliffortia_atrata
Rosaceae_Cliffortia_densa
Rosaceae_Cliffortia_nitidula
Rosaceae_Cliffortia_cuneata
Rosaceae_Cliffortia_ruscifolia
Rosaceae_Cliffortia_ferricola
Rosaceae_Cliffortia_cymbifolia
Rosaceae_Cliffortia_hirsuta
Rosaceae_Cliffortia_recurvata
Rosaceae_Cliffortia_intermedia
Rosaceae_Cliffortia_oligodonta
Rosaceae_Cliffortia_strigosa
Rosaceae_Cliffortia_virgata
Rosaceae_Cliffortia_ovalis
Rosaceae_Cliffortia_denticulata
Rosaceae_Cliffortia_prionota
Rosaceae_Cliffortia_heterophylla
Rosaceae_Cliffortia_robusta
Rosaceae_Cliffortia_lanceolata
Rosaceae_Cliffortia_integerrima
Rosaceae_Cliffortia_theodorifriesii
Rosaceae_Cliffortia_dregeana
Rosaceae_Cliffortia_ceresana
Rosaceae_Cliffortia_acanthophylla
Rosaceae_Cliffortia_phillipsii
Rosaceae_Cliffortia_pungens
Rosaceae_Cliffortia_schlechteri
Rosaceae_Cliffortia_ilicifolia
Rosaceae_Cliffortia_grandifolia
Rosaceae_Cliffortia_reniformis
Rosaceae_Cliffortia_scandens
Rosaceae_Cliffortia_baccans
Rosaceae_Cliffortia_neglecta
Rosaceae_Cliffortia_acutifolia
Rosaceae_Cliffortia_odorata

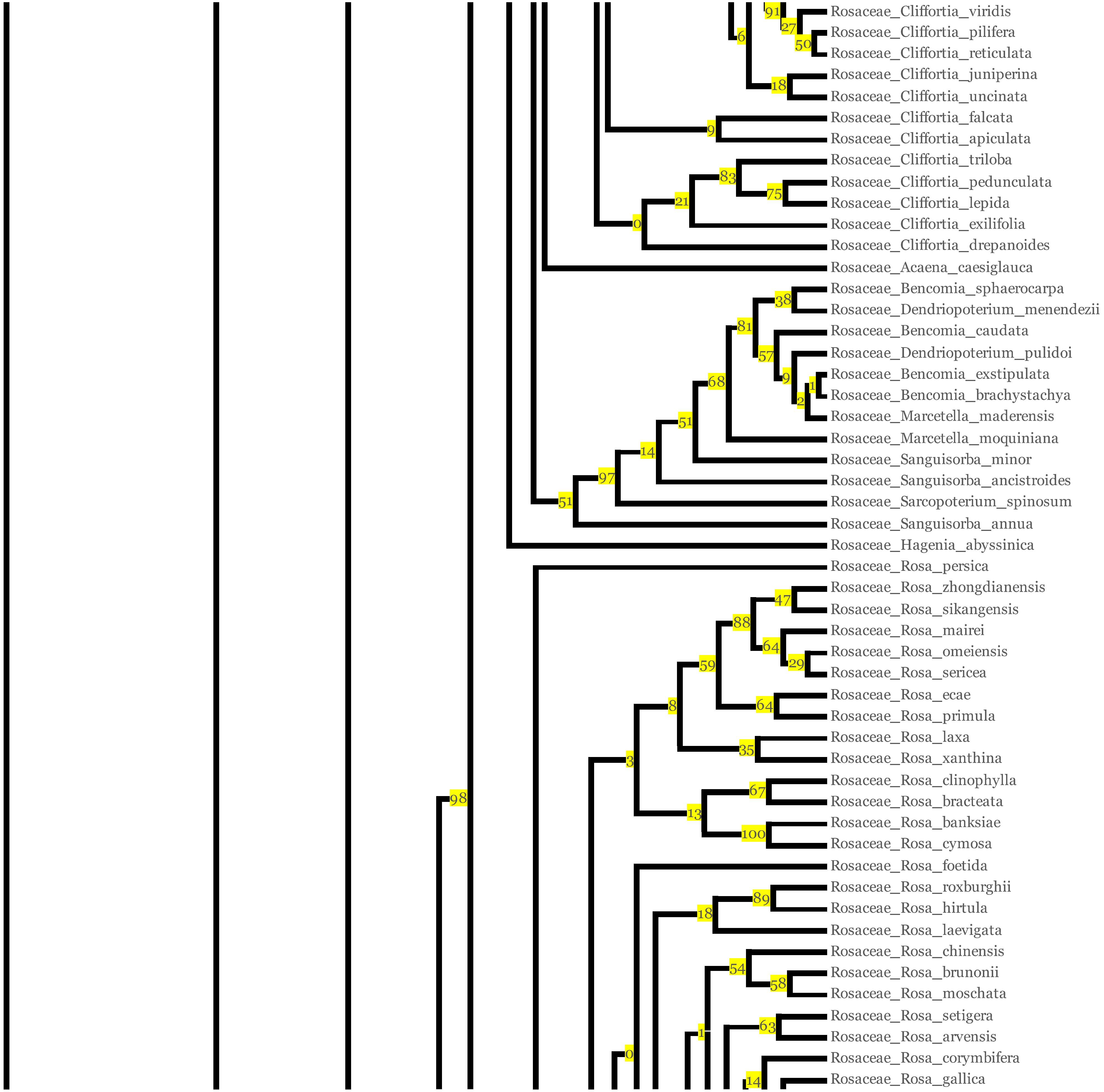
Rosaceae_Cliffortia_viridis
Rosaceae_Cliffortia_pilifera
Rosaceae_Cliffortia_reticulata
Rosaceae_Cliffortia_juniperina
Rosaceae_Cliffortia_uncinata
Rosaceae_Cliffortia_falcata
Rosaceae_Cliffortia_apiculata
Rosaceae_Cliffortia_triloba
Rosaceae_Cliffortia_pedunculata
Rosaceae_Cliffortia_lepida
Rosaceae_Cliffortia_exilifolia
Rosaceae_Cliffortia_drepanoides
Rosaceae_Acaena_caesiglauca
Rosaceae_Bencomia_sphaerocarpa
Rosaceae_Dendriopoterium_menendezii
Rosaceae_Bencomia_caudata
Rosaceae_Dendriopoterium_pulidoi
Rosaceae_Bencomia_exstipulata
Rosaceae_Bencomia_brachystachya
Rosaceae_Marcetella_maderensis
Rosaceae_Marcetella_moquiniana
Rosaceae_Sanguisorba_minor
Rosaceae_Sanguisorba_ancistroides
Rosaceae_Sarcopoterium_spinosum
Rosaceae_Sanguisorba_annua
Rosaceae_Hagenia_abyssinica
Rosaceae_Rosa_persica
Rosaceae_Rosa_zhongdianensis
Rosaceae_Rosa_sikangensis
Rosaceae_Rosa_mairei
Rosaceae_Rosa_omeiensis
Rosaceae_Rosa_sericea
Rosaceae_Rosa_ecae
Rosaceae_Rosa_primula
Rosaceae_Rosa_laxa
Rosaceae_Rosa_xanthina
Rosaceae_Rosa_clinophylla
Rosaceae_Rosa_bracteata
Rosaceae_Rosa_banksiae
Rosaceae_Rosa_cymosa
Rosaceae_Rosa_foetida
Rosaceae_Rosa_roxburghii
Rosaceae_Rosa_hirtula
Rosaceae_Rosa_laevigata
Rosaceae_Rosa_chinensis
Rosaceae_Rosa_brunonii
Rosaceae_Rosa_moschata
Rosaceae_Rosa_setigera
Rosaceae_Rosa_arvensis
Rosaceae_Rosa_corymbifera
Rosaceae_Rosa_gallica

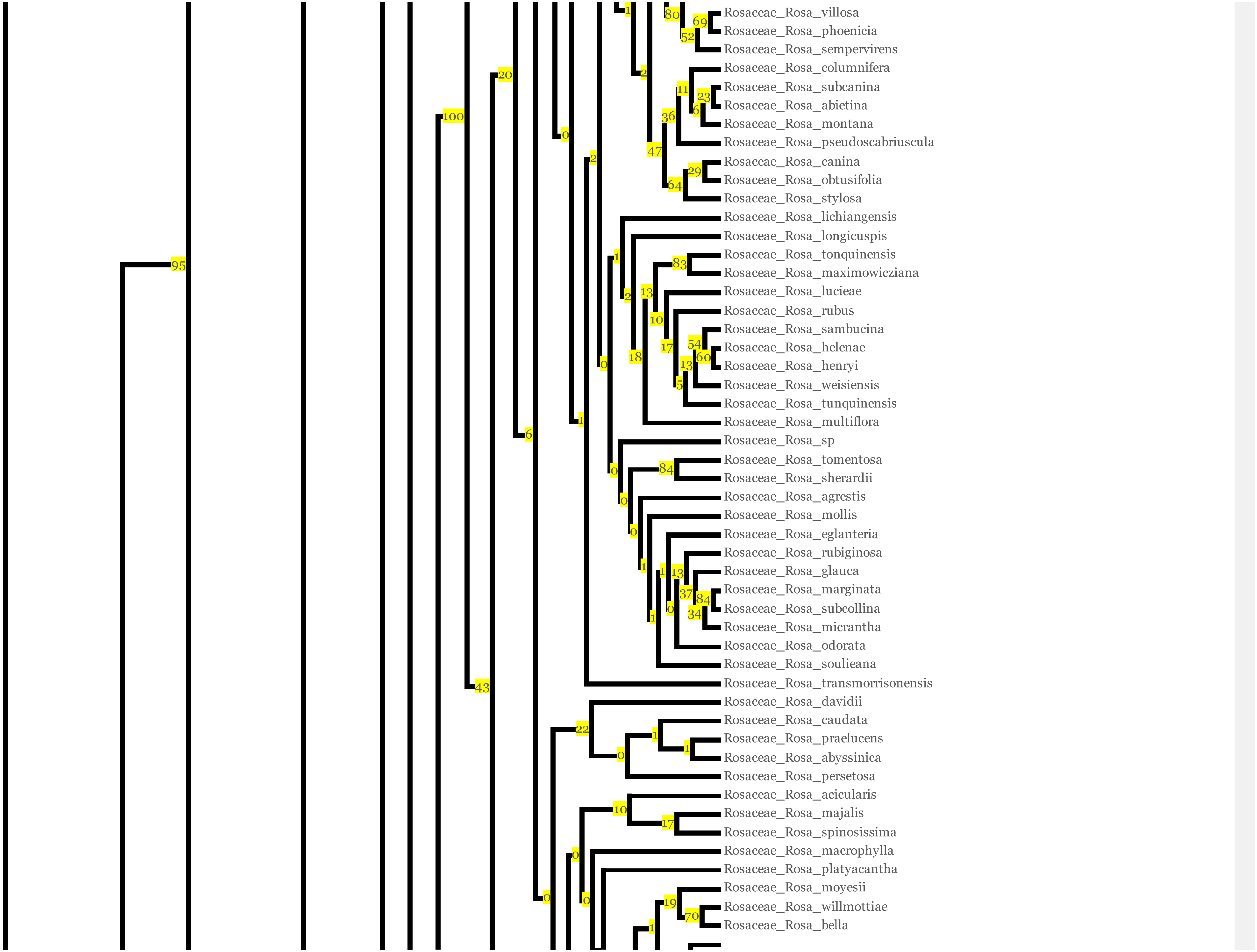
Rosaceae_Rosa_villosa
Rosaceae_Rosa_phoenicia
Rosaceae_Rosa_sempervirens
Rosaceae_Rosa_columnifera
Rosaceae_Rosa_subcanina
Rosaceae_Rosa_abietina
Rosaceae_Rosa_montana
Rosaceae_Rosa_pseudoscabriuscula
Rosaceae_Rosa_canina
Rosaceae_Rosa_obtusifolia
Rosaceae_Rosa_stylosa
Rosaceae_Rosa_lichiangensis
Rosaceae_Rosa_longicuspis
Rosaceae_Rosa_tonquinensis
Rosaceae_Rosa_maximowicziana
Rosaceae_Rosa_lucieae
Rosaceae_Rosa_rubus
Rosaceae_Rosa_sambucina
Rosaceae_Rosa_helenae
Rosaceae_Rosa_henryi
Rosaceae_Rosa_weisiensis
Rosaceae_Rosa_tunquinensis
Rosaceae_Rosa_multiflora
Rosaceae_Rosa_sp
Rosaceae_Rosa_tomentosa
Rosaceae_Rosa_sherardii
Rosaceae_Rosa_agrestis
Rosaceae_Rosa_mollis
Rosaceae_Rosa_eglanteria
Rosaceae_Rosa_rubiginosa
Rosaceae_Rosa_glauca
Rosaceae_Rosa_marginata
Rosaceae_Rosa_subcollina
Rosaceae_Rosa_micrantha
Rosaceae_Rosa_odorata
Rosaceae_Rosa_soulieana
Rosaceae_Rosa_transmorrisonensis
Rosaceae_Rosa_davidii
Rosaceae_Rosa_caudata
Rosaceae_Rosa_praelucens
Rosaceae_Rosa_abyssinica
Rosaceae_Rosa_persetosa
Rosaceae_Rosa_acicularis
Rosaceae_Rosa_majalis
Rosaceae_Rosa_spinosissima
Rosaceae_Rosa_macrophylla
Rosaceae_Rosa_platyacantha
Rosaceae_Rosa_moyesii
Rosaceae_Rosa_willmottiae
Rosaceae_Rosa_bella

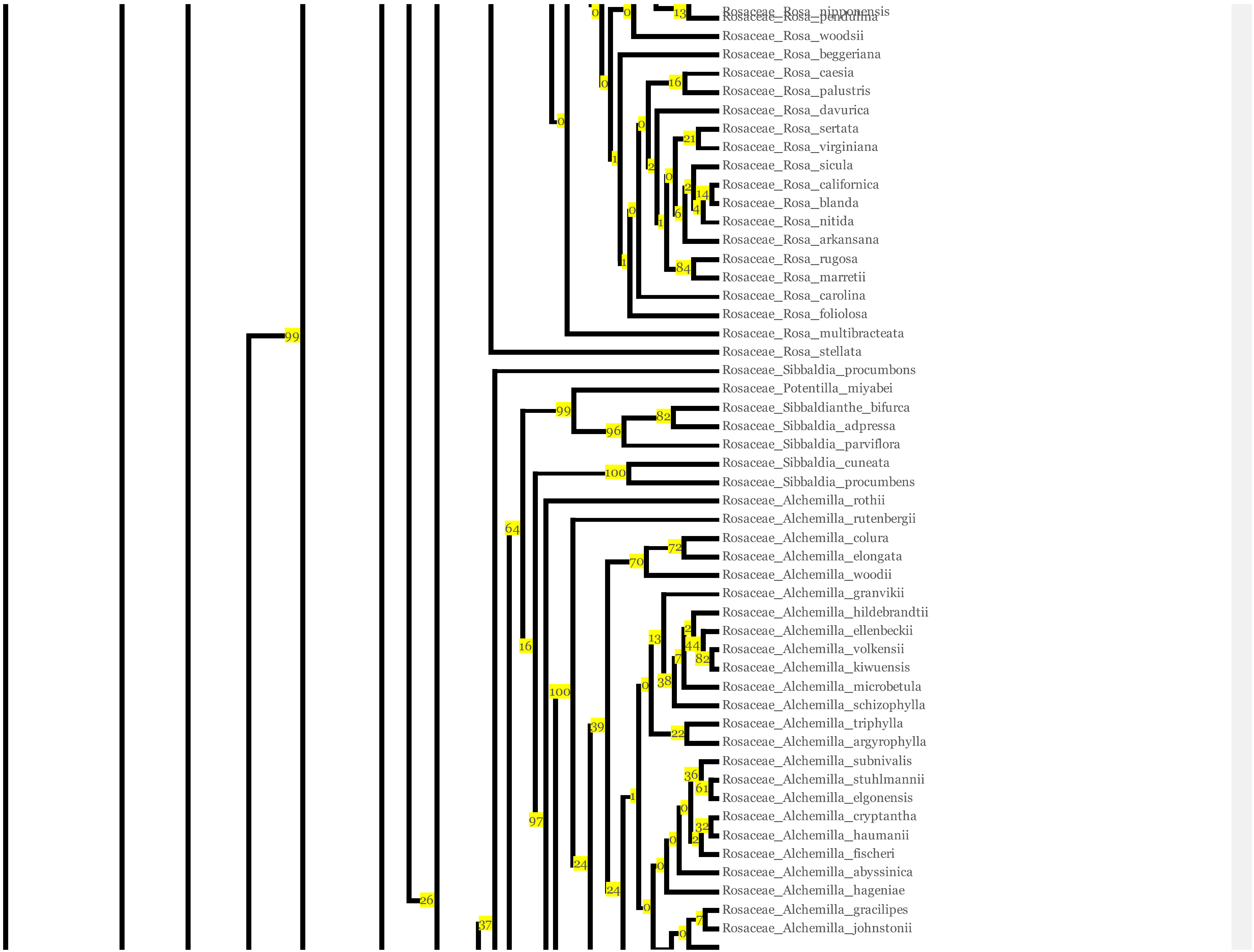
Rosaceae_Rosa_nipponensis
Rosaceae_Rosa_pendulina
Rosaceae_Rosa_woodsii
Rosaceae_Rosa_beggeriana
Rosaceae_Rosa_caesia
Rosaceae_Rosa_palustris
Rosaceae_Rosa_davurica
Rosaceae_Rosa_sertata
Rosaceae_Rosa_virginiana
Rosaceae_Rosa_sicula
Rosaceae_Rosa_californica
Rosaceae_Rosa_blanda
Rosaceae_Rosa_nitida
Rosaceae_Rosa_arkansana
Rosaceae_Rosa_rugosa
Rosaceae_Rosa_marretii
Rosaceae_Rosa_carolina
Rosaceae_Rosa_foliolosa
Rosaceae_Rosa_multibracteata
Rosaceae_Rosa_stellata
Rosaceae_Sibbaldia_procumbons
Rosaceae_Potentilla_miyabei
Rosaceae_Sibbaldianthe_bifurca
Rosaceae_Sibbaldia_adpressa
Rosaceae_Sibbaldia_parviflora
Rosaceae_Sibbaldia_cuneata
Rosaceae_Sibbaldia_procumbens
Rosaceae_Alchemilla_rothii
Rosaceae_Alchemilla_rutenbergii
Rosaceae_Alchemilla_colura
Rosaceae_Alchemilla_elongata
Rosaceae_Alchemilla_woodii
Rosaceae_Alchemilla_granvikii
Rosaceae_Alchemilla_hildebrandtii
Rosaceae_Alchemilla_ellenbeckii
Rosaceae_Alchemilla_volkensii
Rosaceae_Alchemilla_kiwuensis
Rosaceae_Alchemilla_microbetula
Rosaceae_Alchemilla_schizophylla
Rosaceae_Alchemilla_triphylla
Rosaceae_Alchemilla_argyrophylla
Rosaceae_Alchemilla_subnivalis
Rosaceae_Alchemilla_stuhlmannii
Rosaceae_Alchemilla_elgonensis
Rosaceae_Alchemilla_cryptantha
Rosaceae_Alchemilla_haumanii
Rosaceae_Alchemilla_fischeri
Rosaceae_Alchemilla_abyssinica
Rosaceae_Alchemilla_hageniae
Rosaceae_Alchemilla_gracilipes
Rosaceae_Alchemilla_johnstonii

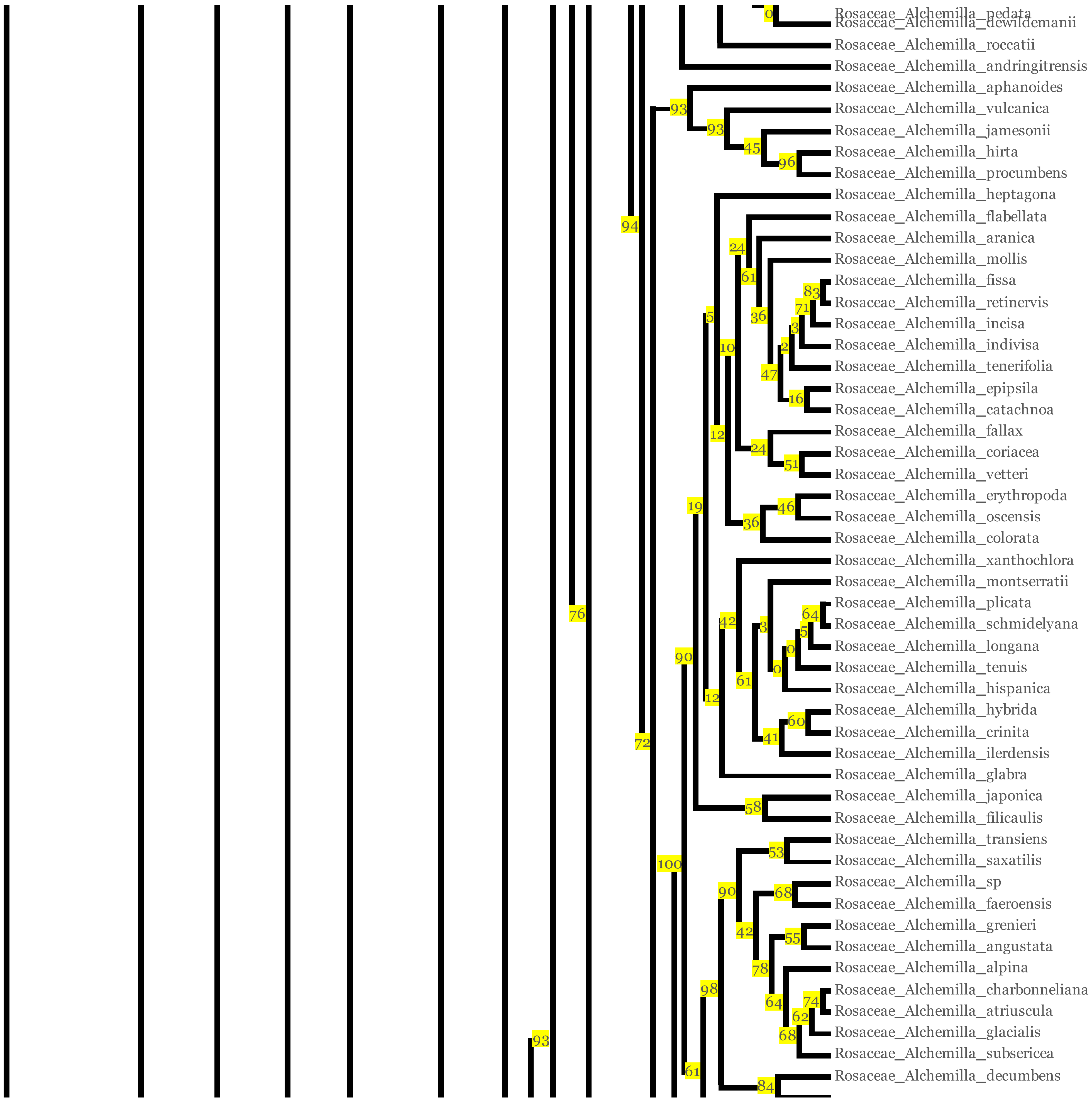
Rosaceae_Alchemilla_pedata
Rosaceae_Alchemilla_dewildemanii
Rosaceae_Alchemilla_roccatii
Rosaceae_Alchemilla_andringitrensis
Rosaceae_Alchemilla_aphanoides
Rosaceae_Alchemilla_vulcanica
Rosaceae_Alchemilla_jamesonii
Rosaceae_Alchemilla_hirta
Rosaceae_Alchemilla_procumbens
Rosaceae_Alchemilla_heptagona
Rosaceae_Alchemilla_flabellata
Rosaceae_Alchemilla_aranica
Rosaceae_Alchemilla_mollis
Rosaceae_Alchemilla_fissa
Rosaceae_Alchemilla_retinervis
Rosaceae_Alchemilla_incisa
Rosaceae_Alchemilla_indivisa
Rosaceae_Alchemilla_tenerifolia
Rosaceae_Alchemilla_epipsila
Rosaceae_Alchemilla_catachnoa
Rosaceae_Alchemilla_fallax
Rosaceae_Alchemilla_coriacea
Rosaceae_Alchemilla_vetteri
Rosaceae_Alchemilla_erythropoda
Rosaceae_Alchemilla_oscensis
Rosaceae_Alchemilla_colorata
Rosaceae_Alchemilla_xanthochlora
Rosaceae_Alchemilla_montserratii
Rosaceae_Alchemilla_plicata
Rosaceae_Alchemilla_schmidelyana
Rosaceae_Alchemilla_longana
Rosaceae_Alchemilla_tenuis
Rosaceae_Alchemilla_hispanica
Rosaceae_Alchemilla_hybrida
Rosaceae_Alchemilla_crinita
Rosaceae_Alchemilla_ilerdensis
Rosaceae_Alchemilla_glabra
Rosaceae_Alchemilla_japonica
Rosaceae_Alchemilla_filicaulis
Rosaceae_Alchemilla_transiens
Rosaceae_Alchemilla_saxatilis
Rosaceae_Alchemilla_sp
Rosaceae_Alchemilla_faeroensis
Rosaceae_Alchemilla_grenieri
Rosaceae_Alchemilla_angustata
Rosaceae_Alchemilla_alpina
Rosaceae_Alchemilla_charbonneliana
Rosaceae_Alchemilla_atriuscula
Rosaceae_Alchemilla_glacialis
Rosaceae_Alchemilla_subsericea
Rosaceae_Alchemilla_decumbens

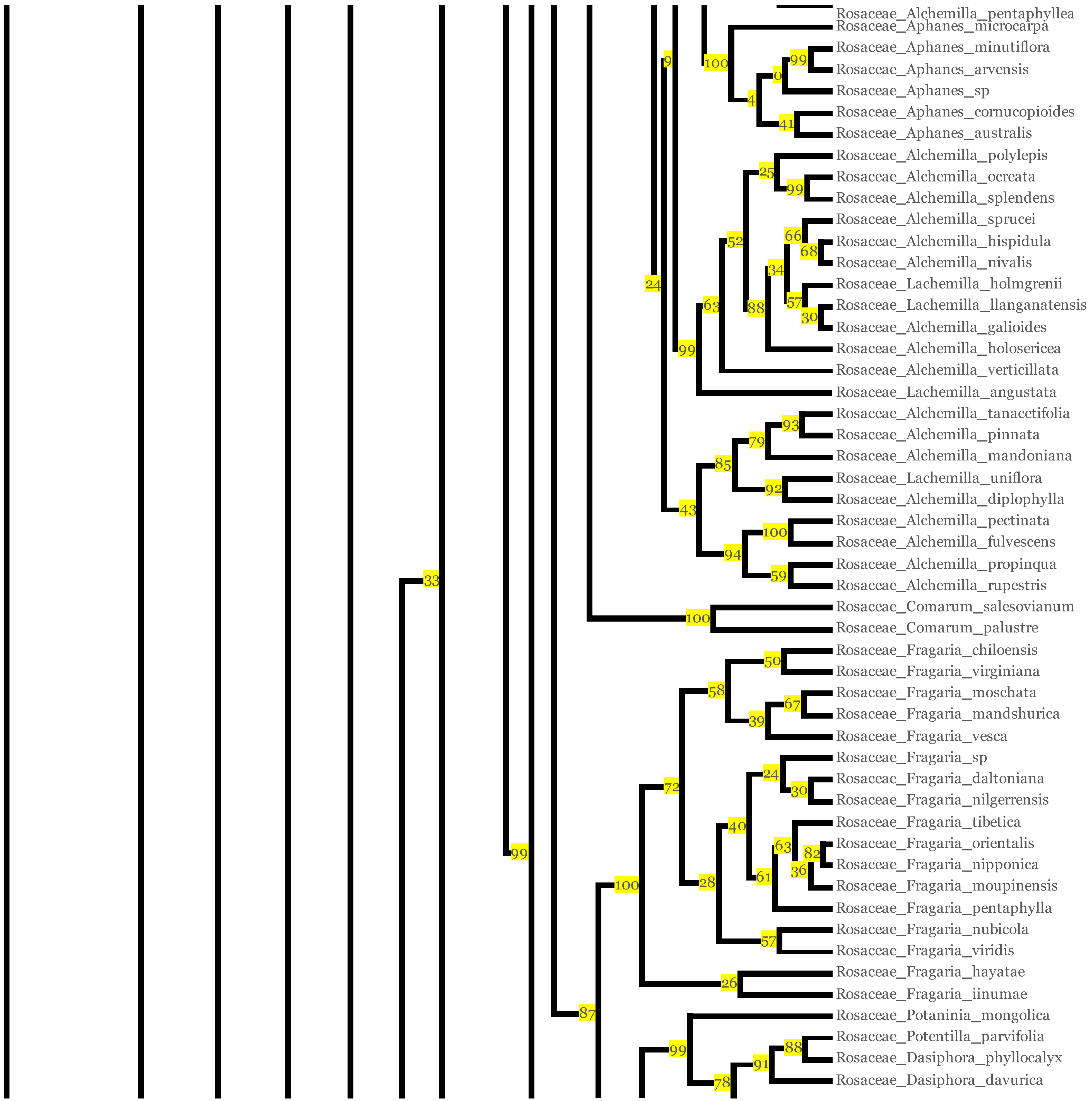
Rosaceae_Alchemilla_pentaphyllea
Rosaceae_Aphanes_microcarpa
Rosaceae_Aphanes_minutiflora
Rosaceae_Aphanes_arvensis
Rosaceae_Aphanes_sp
Rosaceae_Aphanes_cornucopioides
Rosaceae_Aphanes_australis
Rosaceae_Alchemilla_polylepis
Rosaceae_Alchemilla_ocreata
Rosaceae_Alchemilla_splendens
Rosaceae_Alchemilla_sprucei
Rosaceae_Alchemilla_hispidula
Rosaceae_Alchemilla_nivalis
Rosaceae_Lachemilla_holmgrenii
Rosaceae_Lachemilla_llanganatensis
Rosaceae_Alchemilla_galioides
Rosaceae_Alchemilla_holosericea
Rosaceae_Alchemilla_verticillata
Rosaceae_Lachemilla_angustata
Rosaceae_Alchemilla_tanacetifolia
Rosaceae_Alchemilla_pinnata
Rosaceae_Alchemilla_mandoniana
Rosaceae_Lachemilla_uniflora
Rosaceae_Alchemilla_diplophylla
Rosaceae_Alchemilla_pectinata
Rosaceae_Alchemilla_fulvescens
Rosaceae_Alchemilla_propinqua
Rosaceae_Alchemilla_rupestris
Rosaceae_Comarum_salesovianum
Rosaceae_Comarum_palustre
Rosaceae_Fragaria_chiloensis
Rosaceae_Fragaria_virginiana
Rosaceae_Fragaria_moschata
Rosaceae_Fragaria_mandshurica
Rosaceae_Fragaria_vesca
Rosaceae_Fragaria_sp
Rosaceae_Fragaria_daltoniana
Rosaceae_Fragaria_nilgerrensis
Rosaceae_Fragaria_tibetica
Rosaceae_Fragaria_orientalis
Rosaceae_Fragaria_nipponica
Rosaceae_Fragaria_moupinensis
Rosaceae_Fragaria_pentaphylla
Rosaceae_Fragaria_nubicola
Rosaceae_Fragaria_viridis
Rosaceae_Fragaria_hayatae
Rosaceae_Fragaria_iinumae
Rosaceae_Potaninia_mongolica
Rosaceae_Potentilla_parvifolia
Rosaceae_Dasiphora_phyllocalyx
Rosaceae_Dasiphora_davurica

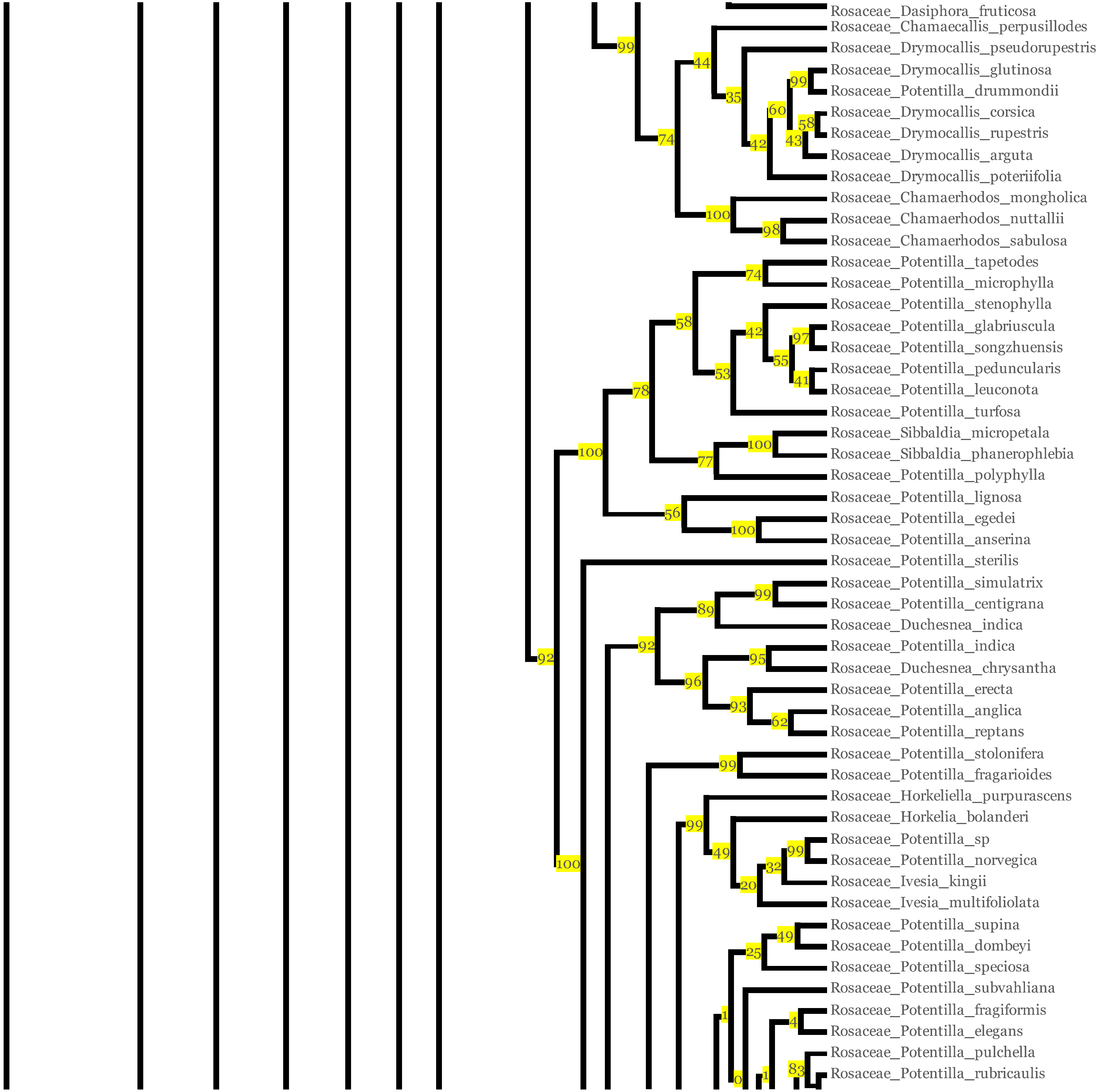
Rosaceae_Dasiphora_fruticosa
Rosaceae_Chamaecallis_perpusillodes
Rosaceae_Drymocallis_pseudorupestris
Rosaceae_Drymocallis_glutinosa
Rosaceae_Potentilla_drummondii
Rosaceae_Drymocallis_corsica
Rosaceae_Drymocallis_rupestris
Rosaceae_Drymocallis_arguta
Rosaceae_Drymocallis_poteriifolia
Rosaceae_Chamaerhodos_mongholica
Rosaceae_Chamaerhodos_nuttallii
Rosaceae_Chamaerhodos_sabulosa
Rosaceae_Potentilla_tapetodes
Rosaceae_Potentilla_microphylla
Rosaceae_Potentilla_stenophylla
Rosaceae_Potentilla_glabriuscula
Rosaceae_Potentilla_songzhuensis
Rosaceae_Potentilla_peduncularis
Rosaceae_Potentilla_leuconota
Rosaceae_Potentilla_turfosa
Rosaceae_Sibbaldia_micropetala
Rosaceae_Sibbaldia_phanerophlebia
Rosaceae_Potentilla_polyphylla
Rosaceae_Potentilla_lignosa
Rosaceae_Potentilla_egedei
Rosaceae_Potentilla_anserina
Rosaceae_Potentilla_sterilis
Rosaceae_Potentilla_simulatrix
Rosaceae_Potentilla_centigrana
Rosaceae_Duchesnea_indica
Rosaceae_Potentilla_indica
Rosaceae_Duchesnea_chrysantha
Rosaceae_Potentilla_erecta
Rosaceae_Potentilla_anglica
Rosaceae_Potentilla_reptans
Rosaceae_Potentilla_stolonifera
Rosaceae_Potentilla_fragarioides
Rosaceae_Horkeliella_purpurascens
Rosaceae_Horkelia_bolanderi
Rosaceae_Potentilla_sp
Rosaceae_Potentilla_norvegica
Rosaceae_Ivesia_kingii
Rosaceae_Ivesia_multifoliolata
Rosaceae_Potentilla_supina
Rosaceae_Potentilla_dombeyi
Rosaceae_Potentilla_speciosa
Rosaceae_Potentilla_subvahliana
Rosaceae_Potentilla_fragiformis
Rosaceae_Potentilla_elegans
Rosaceae_Potentilla_pulchella
Rosaceae_Potentilla_rubricaulis

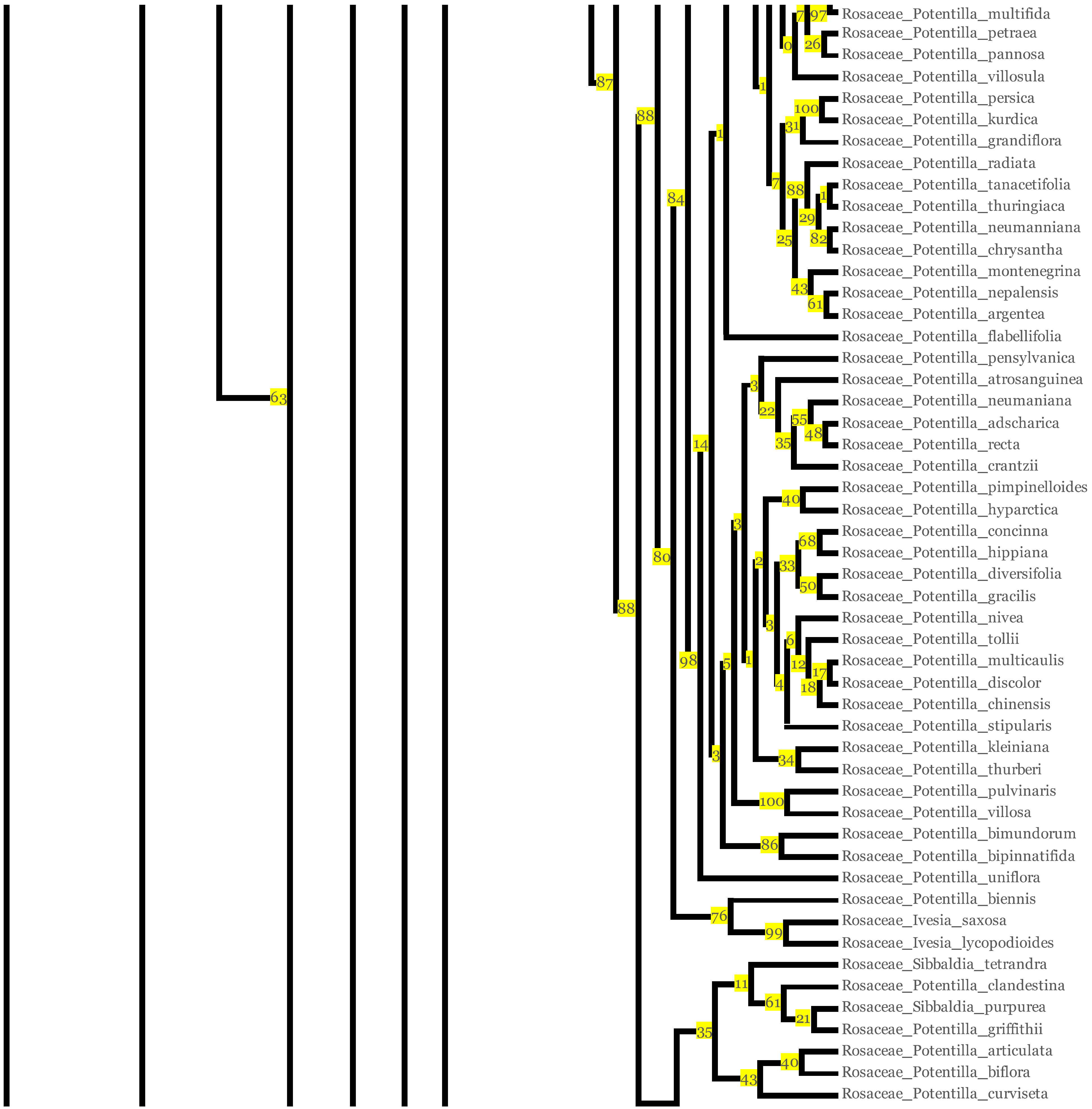
Rosaceae_Potentilla_multifida
Rosaceae_Potentilla_petraea
Rosaceae_Potentilla_pannosa
Rosaceae_Potentilla_villosula
Rosaceae_Potentilla_persica
Rosaceae_Potentilla_kurdica
Rosaceae_Potentilla_grandiflora
Rosaceae_Potentilla_radiata
Rosaceae_Potentilla_tanacetifolia
Rosaceae_Potentilla_thuringiaca
Rosaceae_Potentilla_neumanniana
Rosaceae_Potentilla_chrysantha
Rosaceae_Potentilla_montenegrina
Rosaceae_Potentilla_nepalensis
Rosaceae_Potentilla_argentea
Rosaceae_Potentilla_flabellifolia
Rosaceae_Potentilla_pensylvanica
Rosaceae_Potentilla_atrosanguinea
Rosaceae_Potentilla_neumaniana
Rosaceae_Potentilla_adscharica
Rosaceae_Potentilla_recta
Rosaceae_Potentilla_crantzii
Rosaceae_Potentilla_pimpinelloides
Rosaceae_Potentilla_hyparctica
Rosaceae_Potentilla_concinna
Rosaceae_Potentilla_hippiana
Rosaceae_Potentilla_diversifolia
Rosaceae_Potentilla_gracilis
Rosaceae_Potentilla_nivea
Rosaceae_Potentilla_tollii
Rosaceae_Potentilla_multicaulis
Rosaceae_Potentilla_discolor
Rosaceae_Potentilla_chinensis
Rosaceae_Potentilla_stipularis
Rosaceae_Potentilla_kleiniana
Rosaceae_Potentilla_thurberi
Rosaceae_Potentilla_pulvinaris
Rosaceae_Potentilla_villosa
Rosaceae_Potentilla_bimundorum
Rosaceae_Potentilla_bipinnatifida
Rosaceae_Potentilla_uniflora
Rosaceae_Potentilla_biennis
Rosaceae_Ivesia_saxosa
Rosaceae_Ivesia_lycopodioides
Rosaceae_Sibbaldia_tetrandra
Rosaceae_Potentilla_clandestina
Rosaceae_Sibbaldia_purpurea
Rosaceae_Potentilla_griffithii
Rosaceae_Potentilla_articulata
Rosaceae_Potentilla_biflora
Rosaceae_Potentilla_curviseta

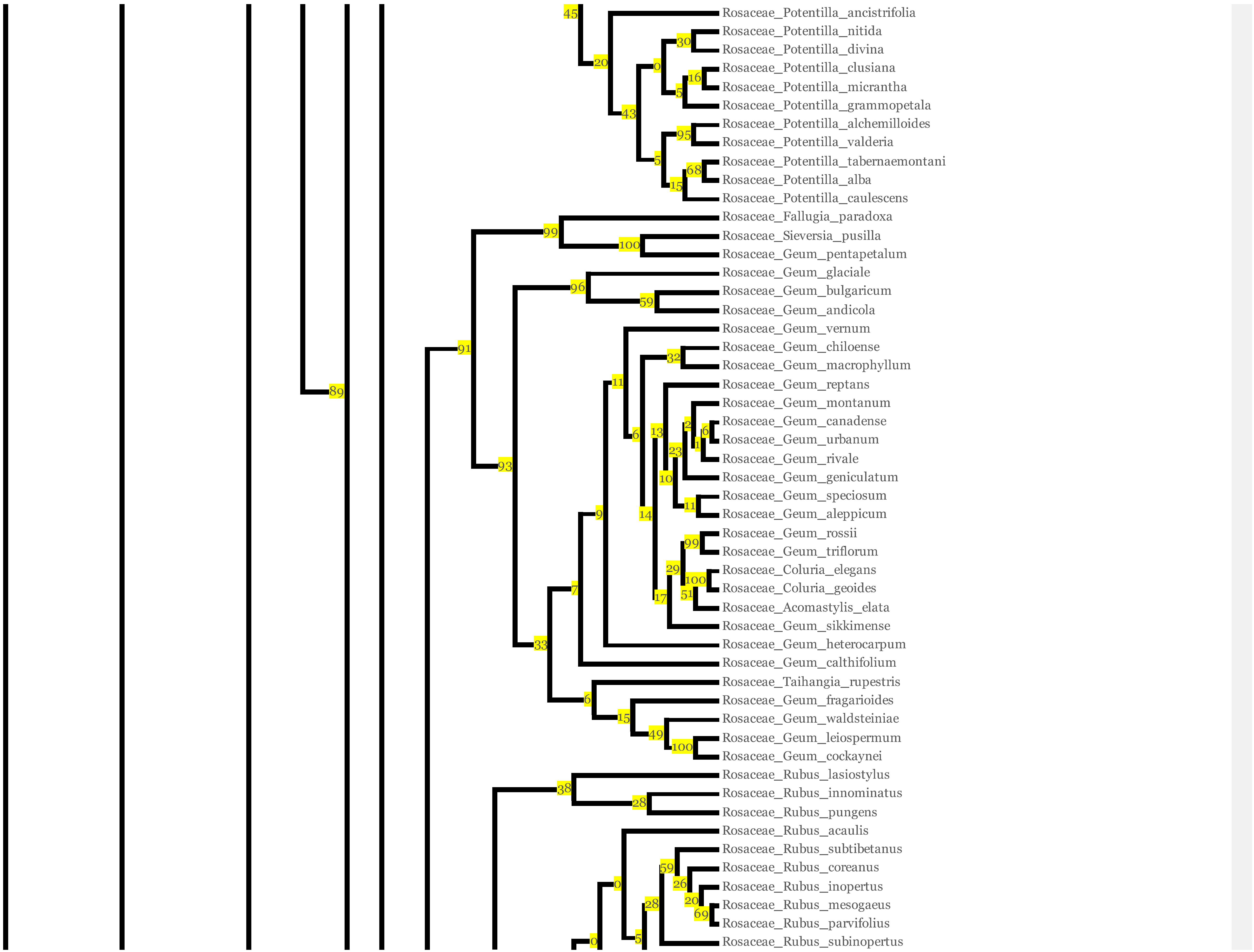
Rosaceae_Potentilla_ancistrifolia
Rosaceae_Potentilla_nitida
Rosaceae_Potentilla_divina
Rosaceae_Potentilla_clusiana
Rosaceae_Potentilla_micrantha
Rosaceae_Potentilla_grammopetala
Rosaceae_Potentilla_alchemilloides
Rosaceae_Potentilla_valderia
Rosaceae_Potentilla_tabernaemontani
Rosaceae_Potentilla_alba
Rosaceae_Potentilla_caulescens
Rosaceae_Fallugia_paradoxa
Rosaceae_Sieversia_pusilla
Rosaceae_Geum_pentapetalum
Rosaceae_Geum_glaciale
Rosaceae_Geum_bulgaricum
Rosaceae_Geum_andicola
Rosaceae_Geum_vernum
Rosaceae_Geum_chiloense
Rosaceae_Geum_macrophyllum
Rosaceae_Geum_reptans
Rosaceae_Geum_montanum
Rosaceae_Geum_canadense
Rosaceae_Geum_urbanum
Rosaceae_Geum_rivale
Rosaceae_Geum_geniculatum
Rosaceae_Geum_speciosum
Rosaceae_Geum_aleppicum
Rosaceae_Geum_rossii
Rosaceae_Geum_triflorum
Rosaceae_Coluria_elegans
Rosaceae_Coluria_geoides
Rosaceae_Acomastylis_elata
Rosaceae_Geum_sikkimense
Rosaceae_Geum_heterocarpum
Rosaceae_Geum_calthifolium
Rosaceae_Taihangia_rupestris
Rosaceae_Geum_fragarioides
Rosaceae_Geum_waldsteiniae
Rosaceae_Geum_leiospermum
Rosaceae_Geum_cockaynei
Rosaceae_Rubus_lasiostylus
Rosaceae_Rubus_innominatus
Rosaceae_Rubus_pungens
Rosaceae_Rubus_acaulis
Rosaceae_Rubus_subtibetanus
Rosaceae_Rubus_coreanus
Rosaceae_Rubus_inopertus
Rosaceae_Rubus_mesogaeus
Rosaceae_Rubus_parvifolius
Rosaceae_Rubus_subinopertus

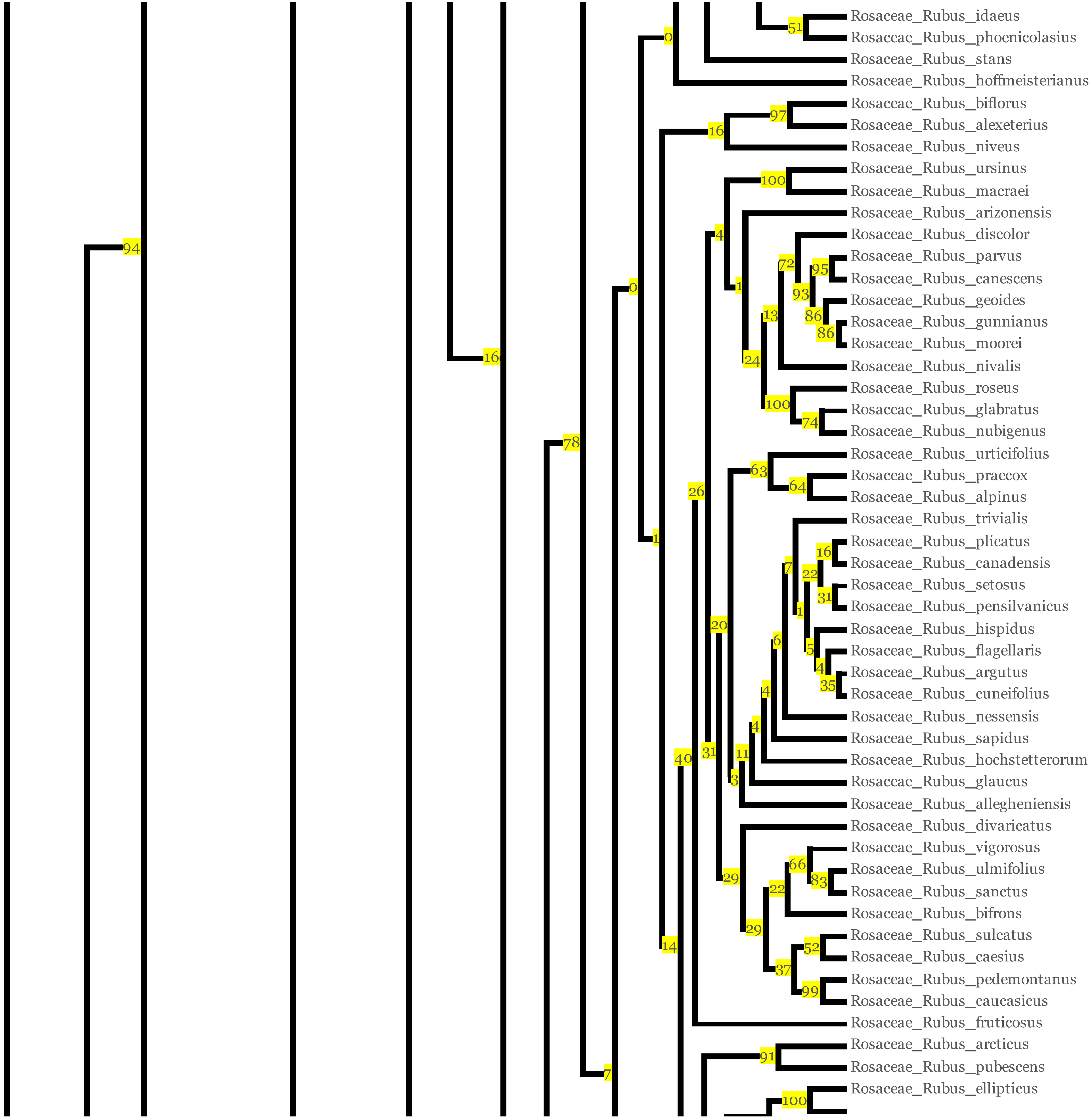
Rosaceae_Rubus_idaeus
Rosaceae_Rubus_phoenicolasius
Rosaceae_Rubus_stans
Rosaceae_Rubus_hoffmeisterianus
Rosaceae_Rubus_biflorus
Rosaceae_Rubus_alexeterius
Rosaceae_Rubus_niveus
Rosaceae_Rubus_ursinus
Rosaceae_Rubus_macraei
Rosaceae_Rubus_arizonensis
Rosaceae_Rubus_discolor
Rosaceae_Rubus_parvus
Rosaceae_Rubus_canescens
Rosaceae_Rubus_geoides
Rosaceae_Rubus_gunnianus
Rosaceae_Rubus_moorei
Rosaceae_Rubus_nivalis
Rosaceae_Rubus_roseus
Rosaceae_Rubus_glabratus
Rosaceae_Rubus_nubigenus
Rosaceae_Rubus_urticifolius
Rosaceae_Rubus_praecox
Rosaceae_Rubus_alpinus
Rosaceae_Rubus_trivialis
Rosaceae_Rubus_plicatus
Rosaceae_Rubus_canadensis
Rosaceae_Rubus_setosus
Rosaceae_Rubus_pensilvanicus
Rosaceae_Rubus_hispidus
Rosaceae_Rubus_flagellaris
Rosaceae_Rubus_argutus
Rosaceae_Rubus_cuneifolius
Rosaceae_Rubus_nessensis
Rosaceae_Rubus_sapidus
Rosaceae_Rubus_hochstetterorum
Rosaceae_Rubus_glaucus
Rosaceae_Rubus_allegheniensis
Rosaceae_Rubus_divaricatus
Rosaceae_Rubus_vigorosus
Rosaceae_Rubus_ulmifolius
Rosaceae_Rubus_sanctus
Rosaceae_Rubus_bifrons
Rosaceae_Rubus_sulcatus
Rosaceae_Rubus_caesius
Rosaceae_Rubus_pedemontanus
Rosaceae_Rubus_caucasicus
Rosaceae_Rubus_fruticosus
Rosaceae_Rubus_arcticus
Rosaceae_Rubus_pubescens
Rosaceae_Rubus_ellipticus

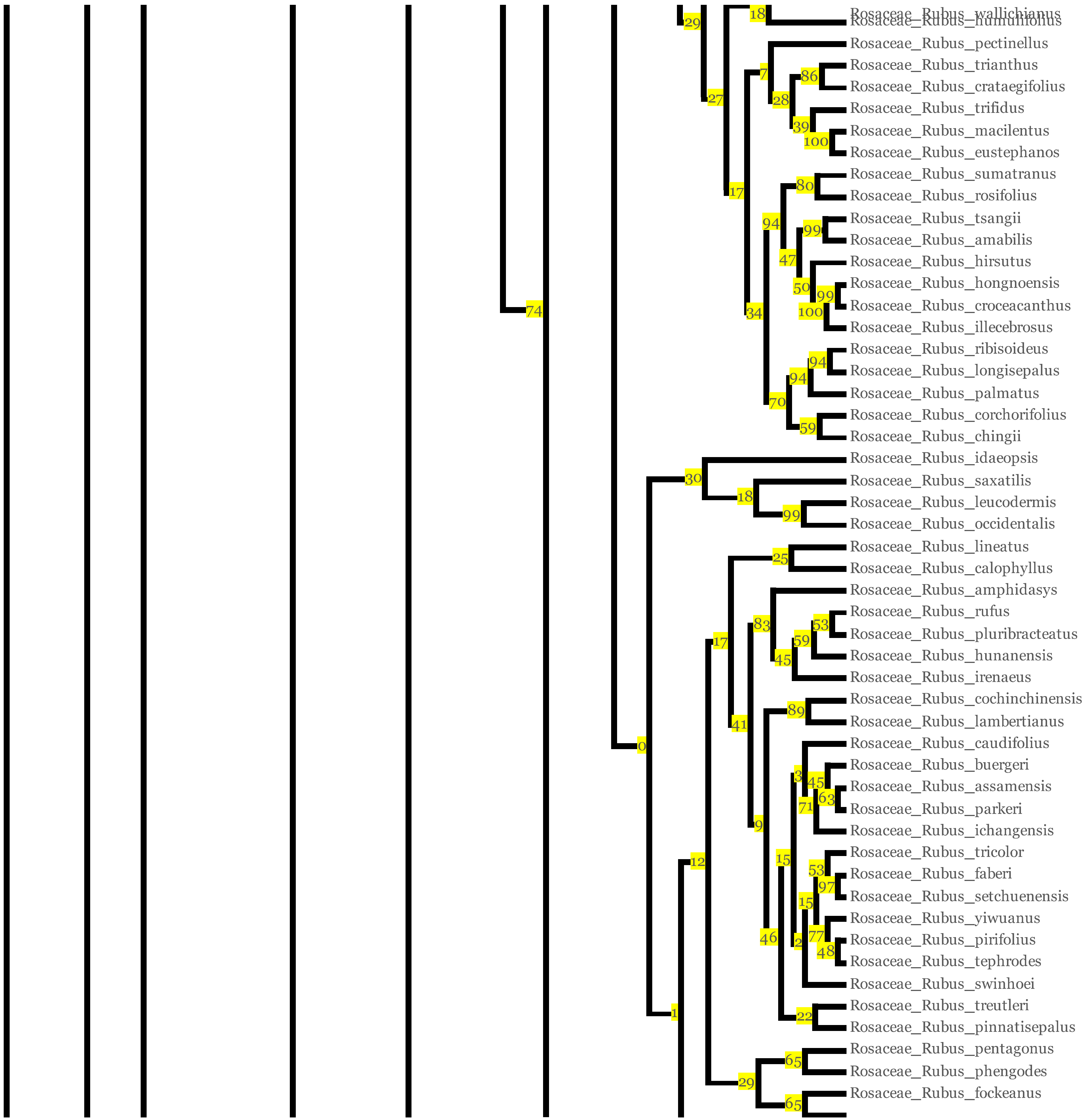
Rosaceae_Rubus_wallichianus
Rosaceae_Rubus_humulifolius
Rosaceae_Rubus_pectinellus
Rosaceae_Rubus_trianthus
Rosaceae_Rubus_crataegifolius
Rosaceae_Rubus_trifidus
Rosaceae_Rubus_macilentus
Rosaceae_Rubus_eustephanos
Rosaceae_Rubus_sumatranus
Rosaceae_Rubus_rosifolius
Rosaceae_Rubus_tsangii
Rosaceae_Rubus_amabilis
Rosaceae_Rubus_hirsutus
Rosaceae_Rubus_hongnoensis
Rosaceae_Rubus_croceacanthus
Rosaceae_Rubus_illecebrosus
Rosaceae_Rubus_ribisoideus
Rosaceae_Rubus_longisepalus
Rosaceae_Rubus_palmatus
Rosaceae_Rubus_corchorifolius
Rosaceae_Rubus_chingii
Rosaceae_Rubus_idaeopsis
Rosaceae_Rubus_saxatilis
Rosaceae_Rubus_leucodermis
Rosaceae_Rubus_occidentalis
Rosaceae_Rubus_lineatus
Rosaceae_Rubus_calophyllus
Rosaceae_Rubus_amphidasys
Rosaceae_Rubus_rufus
Rosaceae_Rubus_pluribracteatus
Rosaceae_Rubus_hunanensis
Rosaceae_Rubus_irenaeus
Rosaceae_Rubus_cochinchinensis
Rosaceae_Rubus_lambertianus
Rosaceae_Rubus_caudifolius
Rosaceae_Rubus_buergeri
Rosaceae_Rubus_assamensis
Rosaceae_Rubus_parkeri
Rosaceae_Rubus_ichangensis
Rosaceae_Rubus_tricolor
Rosaceae_Rubus_faberi
Rosaceae_Rubus_setchuenensis
Rosaceae_Rubus_yiwuanus
Rosaceae_Rubus_pirifolius
Rosaceae_Rubus_tephrodes
Rosaceae_Rubus_swinhoei
Rosaceae_Rubus_treutleri
Rosaceae_Rubus_pinnatisepalus
Rosaceae_Rubus_pentagonus
Rosaceae_Rubus_phengodes
Rosaceae_Rubus_fockeanus

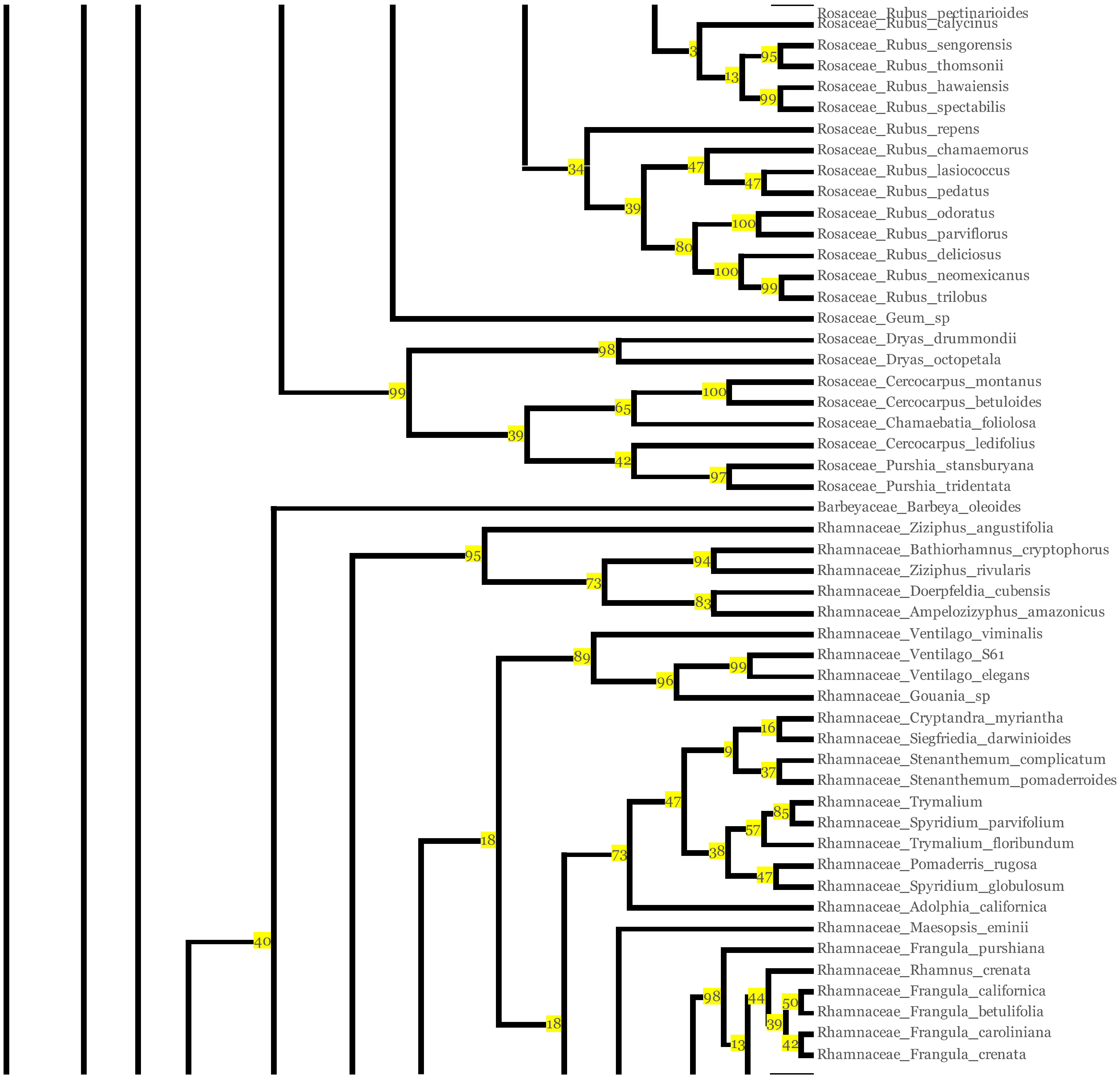
Rosaceae_Rubus_pectinarioides
Rosaceae_Rubus_calycinus
Rosaceae_Rubus_sengorensis
Rosaceae_Rubus_thomsonii
Rosaceae_Rubus_hawaiensis
Rosaceae_Rubus_spectabilis
Rosaceae_Rubus_repens
Rosaceae_Rubus_chamaemorus
Rosaceae_Rubus_lasiococcus
Rosaceae_Rubus_pedatus
Rosaceae_Rubus_odoratus
Rosaceae_Rubus_parviflorus
Rosaceae_Rubus_deliciosus
Rosaceae_Rubus_neomexicanus
Rosaceae_Rubus_trilobus
Rosaceae_Geum_sp
Rosaceae_Dryas_drummondii
Rosaceae_Dryas_octopetala
Rosaceae_Cercocarpus_montanus
Rosaceae_Cercocarpus_betuloides
Rosaceae_Chamaebatia_foliolosa
Rosaceae_Cercocarpus_ledifolius
Rosaceae_Purshia_stansburyana
Rosaceae_Purshia_tridentata
Barbeyaceae_Barbeya_oleoides
Rhamnaceae_Ziziphus_angustifolia
Rhamnaceae_Bathiorhamnus_cryptophorus
Rhamnaceae_Ziziphus_rivularis
Rhamnaceae_Doerpfeldia_cubensis
Rhamnaceae_Ampelozizyphus_amazonicus
Rhamnaceae_Ventilago_viminalis
Rhamnaceae_Ventilago_S61
Rhamnaceae_Ventilago_elegans
Rhamnaceae_Gouania_sp
Rhamnaceae_Cryptandra_myriantha
Rhamnaceae_Siegfriedia_darwinioides
Rhamnaceae_Stenanthemum_complicatum
Rhamnaceae_Stenanthemum_pomaderroides
Rhamnaceae_Trymalium
Rhamnaceae_Spyridium_parvifolium
Rhamnaceae_Trymalium_floribundum
Rhamnaceae_Pomaderris_rugosa
Rhamnaceae_Spyridium_globulosum
Rhamnaceae_Adolphia_californica
Rhamnaceae_Maesopsis_eminii
Rhamnaceae_Frangula_purshiana
Rhamnaceae_Rhamnus_crenata
Rhamnaceae_Frangula_californica
Rhamnaceae_Frangula_betulifolia
Rhamnaceae_Frangula_caroliniana
Rhamnaceae_Frangula_crenata

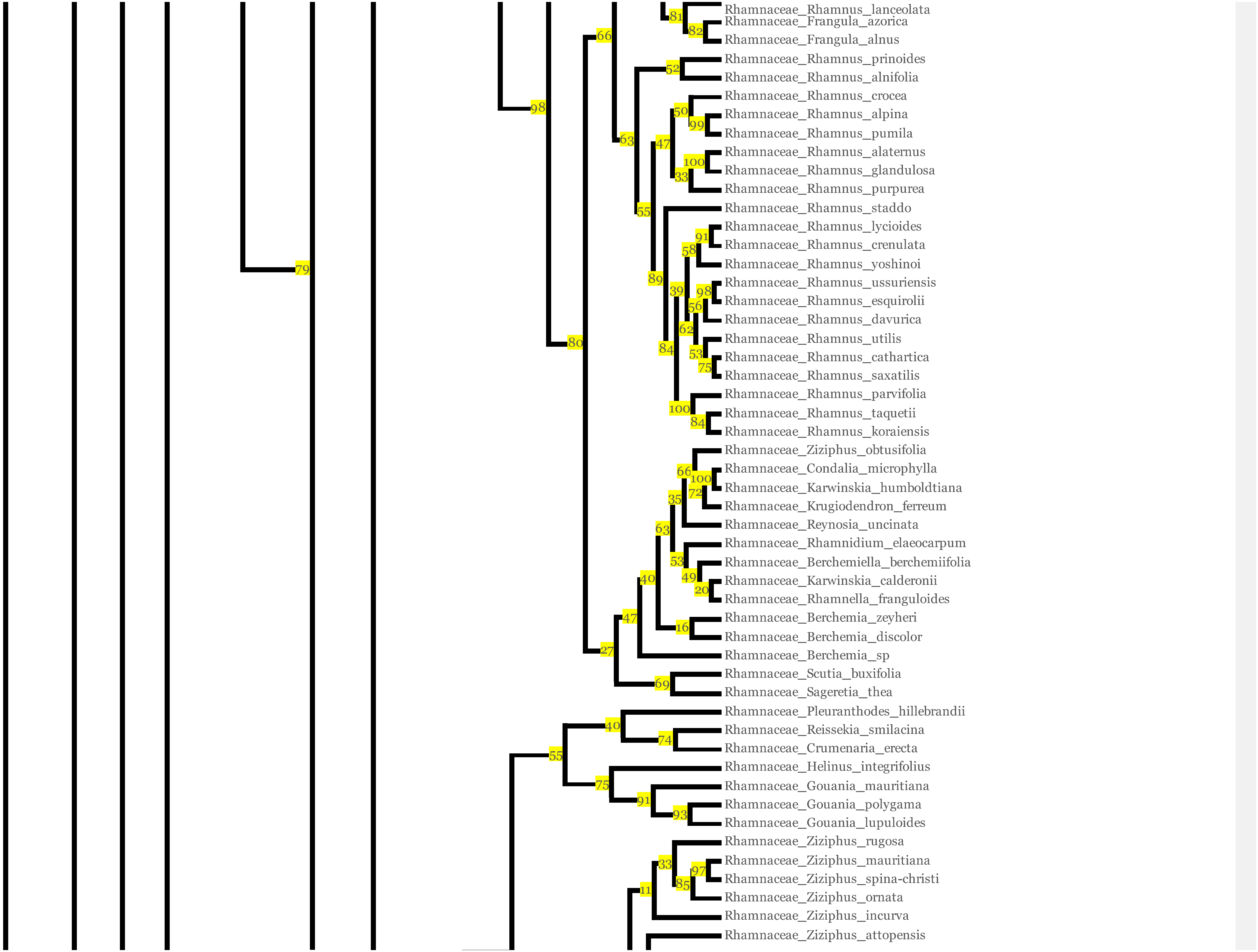
Rhamnaceae_Rhamnus_lanceolata
Rhamnaceae_Frangula_azorica
Rhamnaceae_Frangula_alnus
Rhamnaceae_Rhamnus_prinoides
Rhamnaceae_Rhamnus_alnifolia
Rhamnaceae_Rhamnus_crocea
Rhamnaceae_Rhamnus_alpina
Rhamnaceae_Rhamnus_pumila
Rhamnaceae_Rhamnus_alaternus
Rhamnaceae_Rhamnus_glandulosa
Rhamnaceae_Rhamnus_purpurea
Rhamnaceae_Rhamnus_staddo
Rhamnaceae_Rhamnus_lycioides
Rhamnaceae_Rhamnus_crenulata
Rhamnaceae_Rhamnus_yoshinoi
Rhamnaceae_Rhamnus_ussuriensis
Rhamnaceae_Rhamnus_esquirolii
Rhamnaceae_Rhamnus_davurica
Rhamnaceae_Rhamnus_utilis
Rhamnaceae_Rhamnus_cathartica
Rhamnaceae_Rhamnus_saxatilis
Rhamnaceae_Rhamnus_parvifolia
Rhamnaceae_Rhamnus_taquetii
Rhamnaceae_Rhamnus_koraiensis
Rhamnaceae_Ziziphus_obtusifolia
Rhamnaceae_Condalia_microphylla
Rhamnaceae_Karwinskia_humboldtiana
Rhamnaceae_Krugiodendron_ferreum
Rhamnaceae_Reynosia_uncinata
Rhamnaceae_Rhamnidium_elaeocarpum
Rhamnaceae_Berchemiella_berchemiifolia
Rhamnaceae_Karwinskia_calderonii
Rhamnaceae_Rhamnella_franguloides
Rhamnaceae_Berchemia_zeyheri
Rhamnaceae_Berchemia_discolor
Rhamnaceae_Berchemia_sp
Rhamnaceae_Scutia_buxifolia
Rhamnaceae_Sageretia_thea
Rhamnaceae_Pleuranthodes_hillebrandii
Rhamnaceae_Reissekia_smilacina
Rhamnaceae_Crumenaria_erecta
Rhamnaceae_Helinus_integrifolius
Rhamnaceae_Gouania_mauritiana
Rhamnaceae_Gouania_polygama
Rhamnaceae_Gouania_lupuloides
Rhamnaceae_Ziziphus_rugosa
Rhamnaceae_Ziziphus_mauritiana
Rhamnaceae_Ziziphus_spina-christi
Rhamnaceae_Ziziphus_ornata
Rhamnaceae_Ziziphus_incurva
Rhamnaceae_Ziziphus_attopensis

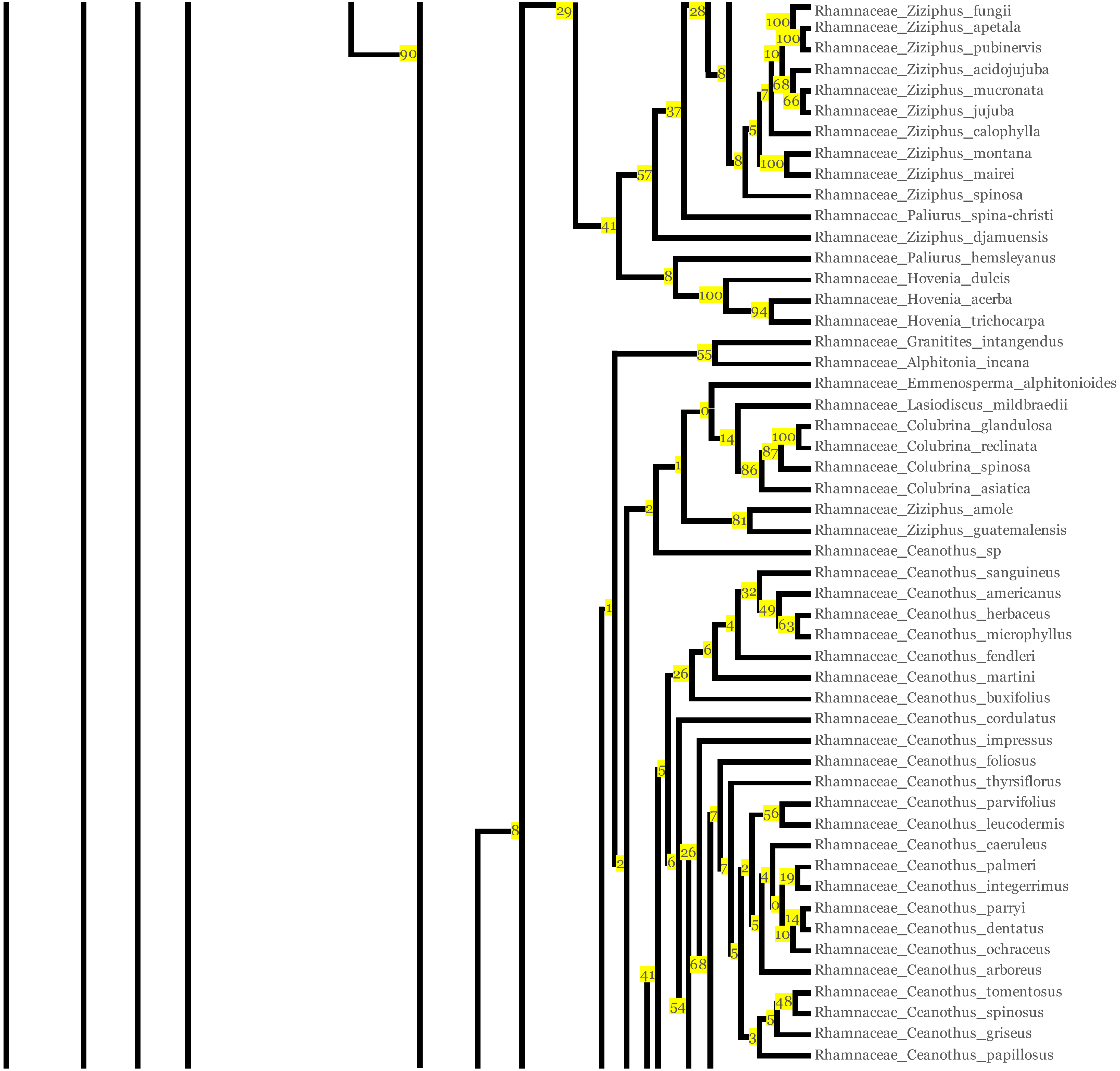
Rhamnaceae_Ziziphus_fungii
Rhamnaceae_Ziziphus_apetala
Rhamnaceae_Ziziphus_pubinervis
Rhamnaceae_Ziziphus_acidojujuba
Rhamnaceae_Ziziphus_mucronata
Rhamnaceae_Ziziphus_jujuba
Rhamnaceae_Ziziphus_calophylla
Rhamnaceae_Ziziphus_montana
Rhamnaceae_Ziziphus_mairei
Rhamnaceae_Ziziphus_spinosa
Rhamnaceae_Paliurus_spina-christi
Rhamnaceae_Ziziphus_djamuensis
Rhamnaceae_Paliurus_hemsleyanus
Rhamnaceae_Hovenia_dulcis
Rhamnaceae_Hovenia_acerba
Rhamnaceae_Hovenia_trichocarpa
Rhamnaceae_Granitites_intangendus
Rhamnaceae_Alphitonia_incana
Rhamnaceae_Emmenosperma_alphitonioides
Rhamnaceae_Lasiodiscus_mildbraedii
Rhamnaceae_Colubrina_glandulosa
Rhamnaceae_Colubrina_reclinata
Rhamnaceae_Colubrina_spinosa
Rhamnaceae_Colubrina_asiatica
Rhamnaceae_Ziziphus_amole
Rhamnaceae_Ziziphus_guatemalensis
Rhamnaceae_Ceanothus_sp
Rhamnaceae_Ceanothus_sanguineus
Rhamnaceae_Ceanothus_americanus
Rhamnaceae_Ceanothus_herbaceus
Rhamnaceae_Ceanothus_microphyllus
Rhamnaceae_Ceanothus_fendleri
Rhamnaceae_Ceanothus_martini
Rhamnaceae_Ceanothus_buxifolius
Rhamnaceae_Ceanothus_cordulatus
Rhamnaceae_Ceanothus_impressus
Rhamnaceae_Ceanothus_foliosus
Rhamnaceae_Ceanothus_thyrsiflorus
Rhamnaceae_Ceanothus_parvifolius
Rhamnaceae_Ceanothus_leucodermis
Rhamnaceae_Ceanothus_caeruleus
Rhamnaceae_Ceanothus_palmeri
Rhamnaceae_Ceanothus_integerrimus
Rhamnaceae_Ceanothus_parryi
Rhamnaceae_Ceanothus_dentatus
Rhamnaceae_Ceanothus_ochraceus
Rhamnaceae_Ceanothus_arboreus
Rhamnaceae_Ceanothus_tomentosus
Rhamnaceae_Ceanothus_spinosus
Rhamnaceae_Ceanothus_griseus
Rhamnaceae_Ceanothus_papillosus

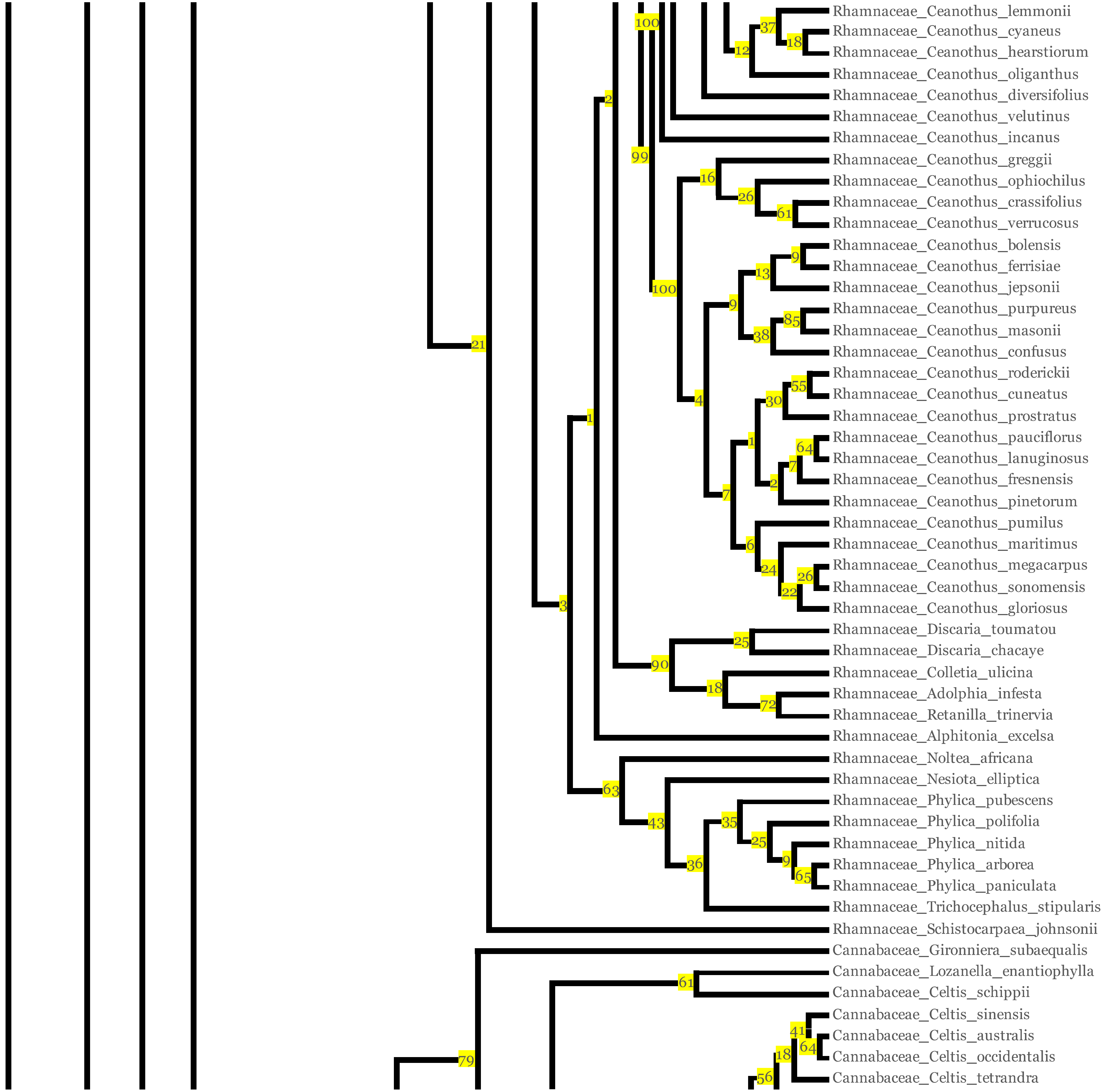
Rhamnaceae_Ceanothus_lemmonii
Rhamnaceae_Ceanothus_cyaneus
Rhamnaceae_Ceanothus_hearstiorum
Rhamnaceae_Ceanothus_oliganthus
Rhamnaceae_Ceanothus_diversifolius
Rhamnaceae_Ceanothus_velutinus
Rhamnaceae_Ceanothus_incanus
Rhamnaceae_Ceanothus_greggii
Rhamnaceae_Ceanothus_ophiochilus
Rhamnaceae_Ceanothus_crassifolius
Rhamnaceae_Ceanothus_verrucosus
Rhamnaceae_Ceanothus_bolensis
Rhamnaceae_Ceanothus_ferrisiae
Rhamnaceae_Ceanothus_jepsonii
Rhamnaceae_Ceanothus_purpureus
Rhamnaceae_Ceanothus_masonii
Rhamnaceae_Ceanothus_confusus
Rhamnaceae_Ceanothus_roderickii
Rhamnaceae_Ceanothus_cuneatus
Rhamnaceae_Ceanothus_prostratus
Rhamnaceae_Ceanothus_pauciflorus
Rhamnaceae_Ceanothus_lanuginosus
Rhamnaceae_Ceanothus_fresnensis
Rhamnaceae_Ceanothus_pinetorum
Rhamnaceae_Ceanothus_pumilus
Rhamnaceae_Ceanothus_maritimus
Rhamnaceae_Ceanothus_megacarpus
Rhamnaceae_Ceanothus_sonomensis
Rhamnaceae_Ceanothus_gloriosus
Rhamnaceae_Discaria_toumatou
Rhamnaceae_Discaria_chacaye
Rhamnaceae_Colletia_ulicina
Rhamnaceae_Adolphia_infesta
Rhamnaceae_Retanilla_trinervia
Rhamnaceae_Alphitonia_excelsa
Rhamnaceae_Noltea_africana
Rhamnaceae_Nesiota_elliptica
Rhamnaceae_Phylica_pubescens
Rhamnaceae_Phylica_polifolia
Rhamnaceae_Phylica_nitida
Rhamnaceae_Phylica_arborea
Rhamnaceae_Phylica_paniculata
Rhamnaceae_Trichocephalus_stipularis
Rhamnaceae_Schistocarpaea_johnsonii
Cannabaceae_Gironniera_subaequalis
Cannabaceae_Lozanella_enantiophylla
Cannabaceae_Celtis_schippii
Cannabaceae_Celtis_sinensis
Cannabaceae_Celtis_australis
Cannabaceae_Celtis_occidentalis
Cannabaceae_Celtis_tetrandra

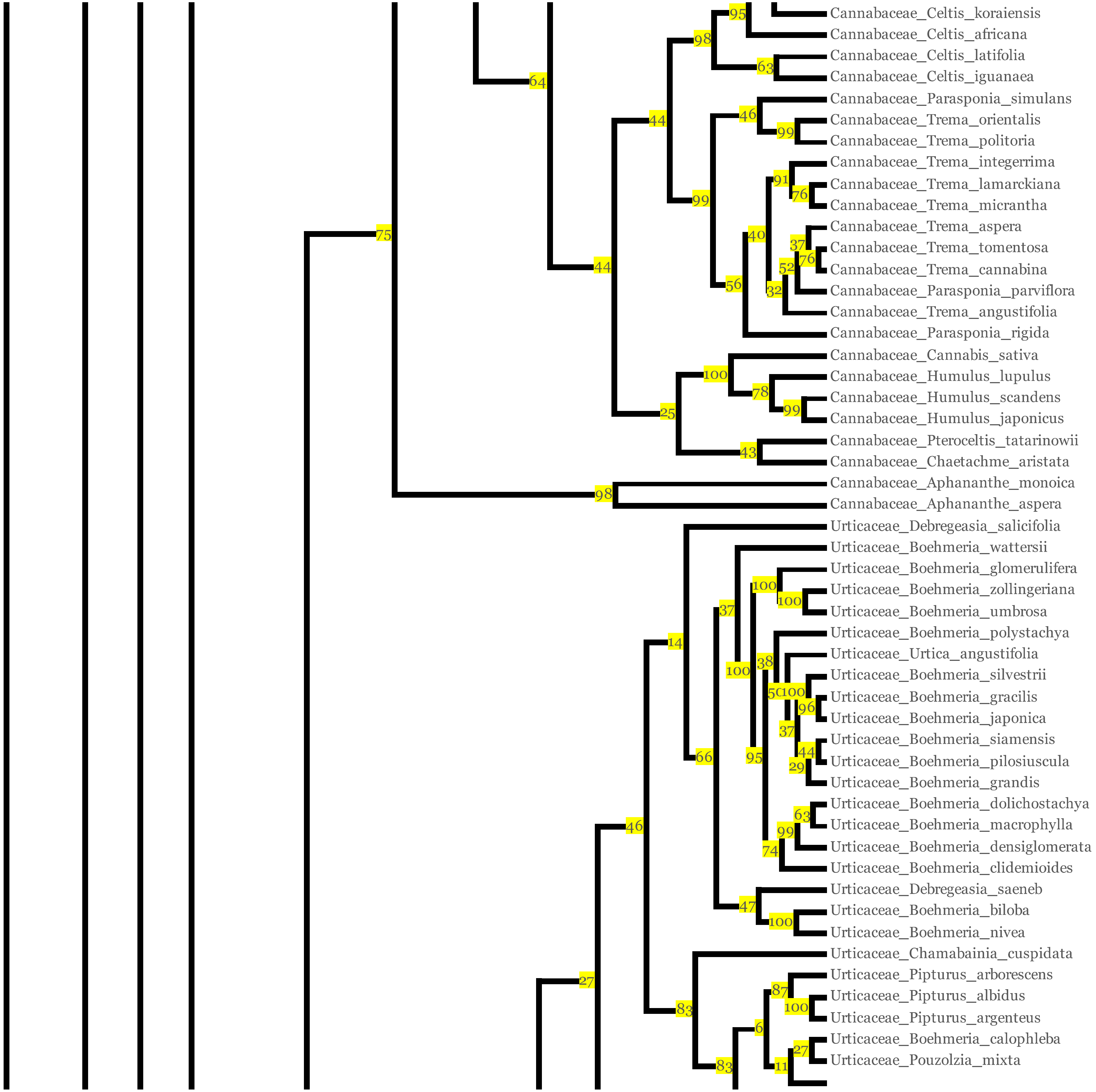
Cannabaceae_Celtis_koraiensis
Cannabaceae_Celtis_africana
Cannabaceae_Celtis_latifolia
Cannabaceae_Celtis_iguanaea
Cannabaceae_Parasponia_simulans
Cannabaceae_Trema_orientalis
Cannabaceae_Trema_politoria
Cannabaceae_Trema_integerrima
Cannabaceae_Trema_lamarckiana
Cannabaceae_Trema_micrantha
Cannabaceae_Trema_aspera
Cannabaceae_Trema_tomentosa
Cannabaceae_Trema_cannabina
Cannabaceae_Parasponia_parviflora
Cannabaceae_Trema_angustifolia
Cannabaceae_Parasponia_rigida
Cannabaceae_Cannabis_sativa
Cannabaceae_Humulus_lupulus
Cannabaceae_Humulus_scandens
Cannabaceae_Humulus_japonicus
Cannabaceae_Pteroceltis_tatarinowii
Cannabaceae_Chaetachme_aristata
Cannabaceae_Aphananthe_monoica
Cannabaceae_Aphananthe_aspera
Urticaceae_Debregeasia_salicifolia
Urticaceae_Boehmeria_wattersii
Urticaceae_Boehmeria_glomerulifera
Urticaceae_Boehmeria_zollingeriana
Urticaceae_Boehmeria_umbrosa
Urticaceae_Boehmeria_polystachya
Urticaceae_Urtica_angustifolia
Urticaceae_Boehmeria_silvestrii
Urticaceae_Boehmeria_gracilis
Urticaceae_Boehmeria_japonica
Urticaceae_Boehmeria_siamensis
Urticaceae_Boehmeria_pilosiuscula
Urticaceae_Boehmeria_grandis
Urticaceae_Boehmeria_dolichostachya
Urticaceae_Boehmeria_macrophylla
Urticaceae_Boehmeria_densiglomerata
Urticaceae_Boehmeria_clidemioides
Urticaceae_Debregeasia_saeneb
Urticaceae_Boehmeria_biloba
Urticaceae_Boehmeria_nivea
Urticaceae_Chamabainia_cuspidata
Urticaceae_Pipturus_arborescens
Urticaceae_Pipturus_albidus
Urticaceae_Pipturus_argenteus
Urticaceae_Boehmeria_calophleba
Urticaceae_Pouzolzia_mixta

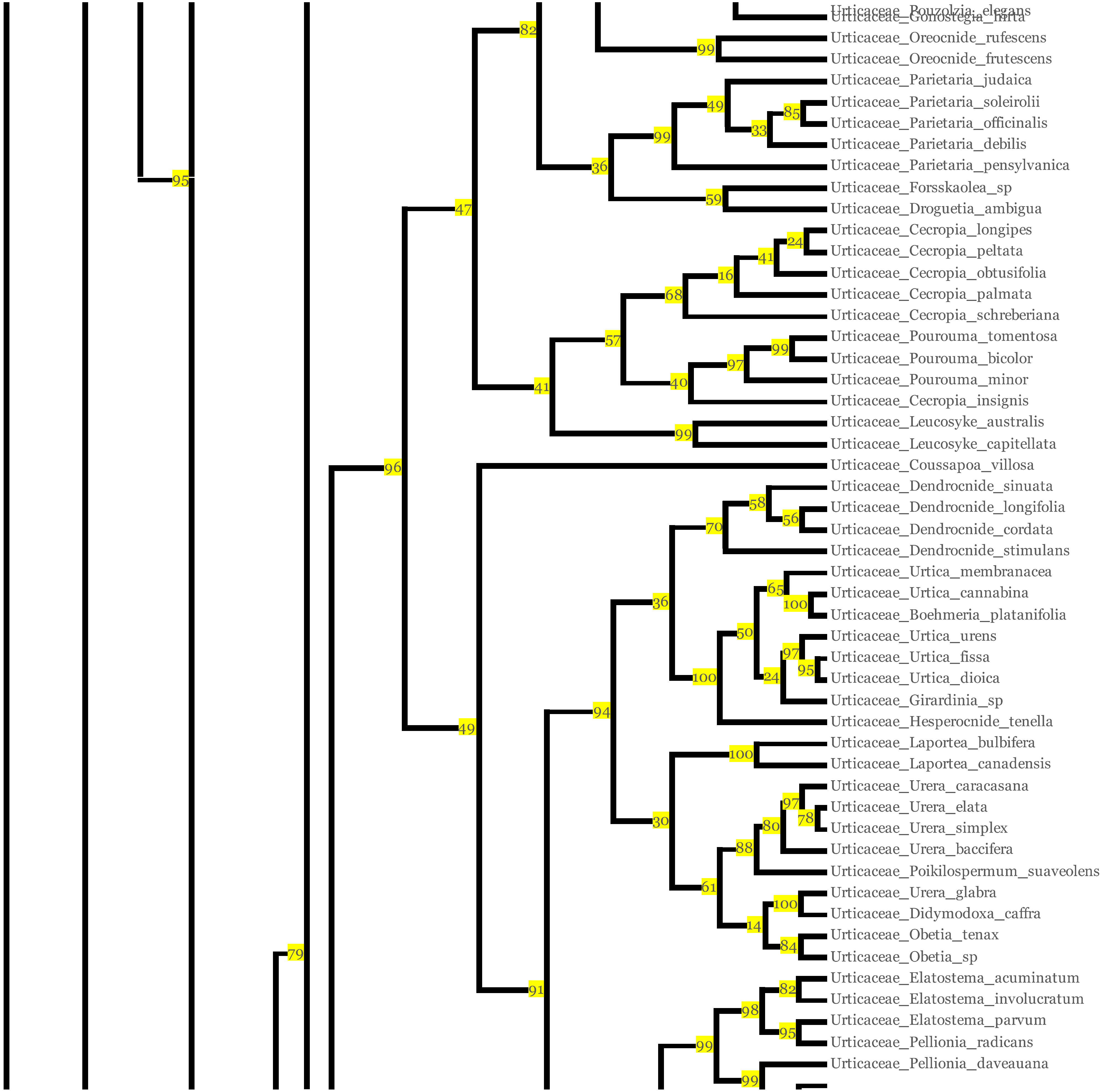
Urticaceae_Pouzolzia_elegans
Urticaceae_Gonostegia_hirta
Urticaceae_Oreocnide_rufescens
Urticaceae_Oreocnide_frutescens
Urticaceae_Parietaria_judaica
Urticaceae_Parietaria_soleirolii
Urticaceae_Parietaria_officinalis
Urticaceae_Parietaria_debilis
Urticaceae_Parietaria_pensylvanica
Urticaceae_Forsskaolea_sp
Urticaceae_Droguetia_ambigua
Urticaceae_Cecropia_longipes
Urticaceae_Cecropia_peltata
Urticaceae_Cecropia_obtusifolia
Urticaceae_Cecropia_palmata
Urticaceae_Cecropia_schreberiana
Urticaceae_Pourouma_tomentosa
Urticaceae_Pourouma_bicolor
Urticaceae_Pourouma_minor
Urticaceae_Cecropia_insignis
Urticaceae_Leucosyke_australis
Urticaceae_Leucosyke_capitellata
Urticaceae_Coussapoa_villosa
Urticaceae_Dendrocnide_sinuata
Urticaceae_Dendrocnide_longifolia
Urticaceae_Dendrocnide_cordata
Urticaceae_Dendrocnide_stimulans
Urticaceae_Urtica_membranacea
Urticaceae_Urtica_cannabina
Urticaceae_Boehmeria_platanifolia
Urticaceae_Urtica_urens
Urticaceae_Urtica_fissa
Urticaceae_Urtica_dioica
Urticaceae_Girardinia_sp
Urticaceae_Hesperocnide_tenella
Urticaceae_Laportea_bulbifera
Urticaceae_Laportea_canadensis
Urticaceae_Urera_caracasana
Urticaceae_Urera_elata
Urticaceae_Urera_simplex
Urticaceae_Urera_baccifera
Urticaceae_Poikilospermum_suaveolens
Urticaceae_Urera_glabra
Urticaceae_Didymodoxa_caffra
Urticaceae_Obetia_tenax
Urticaceae_Obetia_sp
Urticaceae_Elatostema_acuminatum
Urticaceae_Elatostema_involucratum
Urticaceae_Elatostema_parvum
Urticaceae_Pellionia_radicans
Urticaceae_Pellionia_daveauana

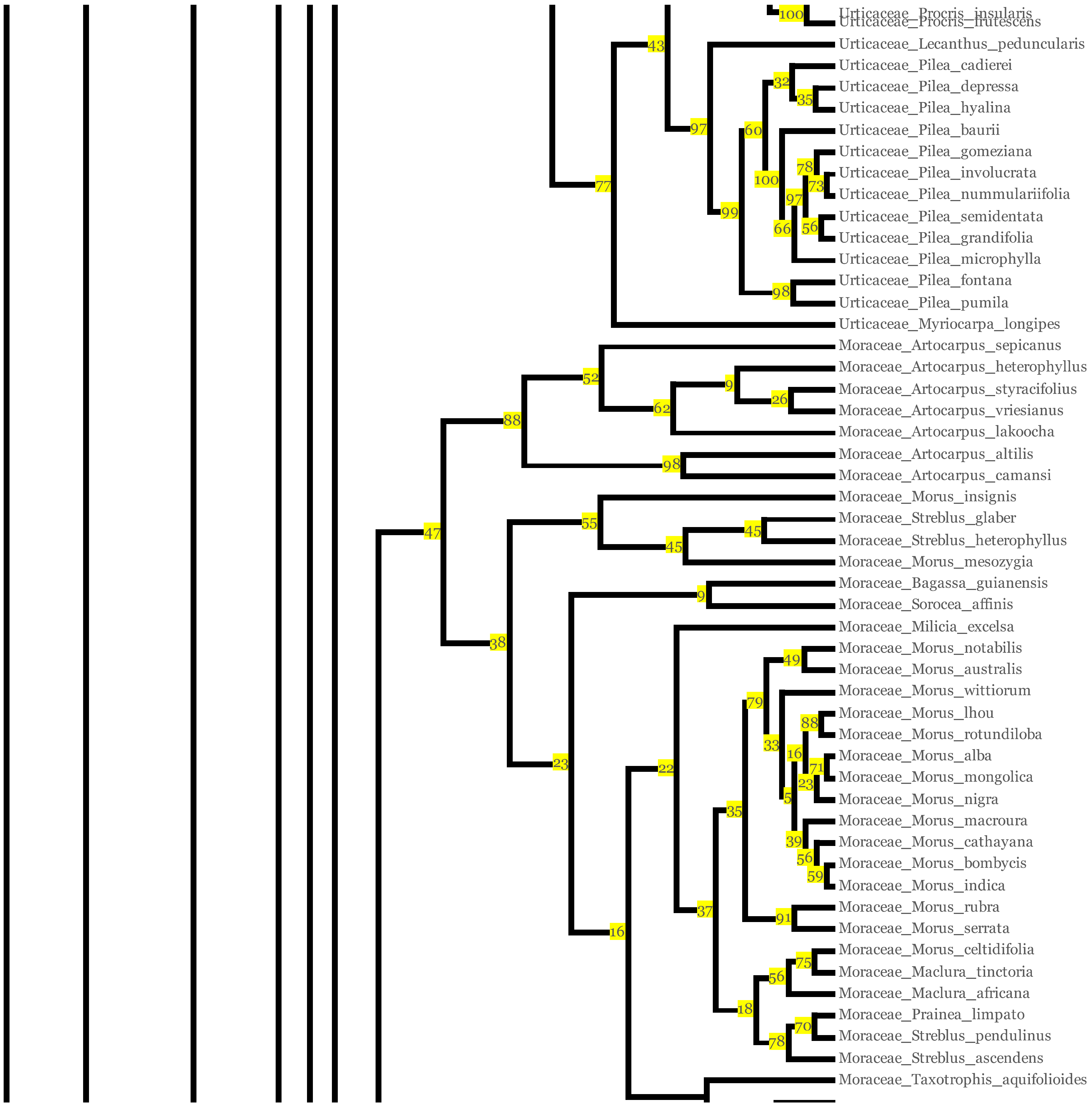
Urticaceae_Procris_insularis
Urticaceae_Procris_frutescens
Urticaceae_Lecanthus_peduncularis
Urticaceae_Pilea_cadierei
Urticaceae_Pilea_depressa
Urticaceae_Pilea_hyalina
Urticaceae_Pilea_baurii
Urticaceae_Pilea_gomeziana
Urticaceae_Pilea_involucrata
Urticaceae_Pilea_nummulariifolia
Urticaceae_Pilea_semidentata
Urticaceae_Pilea_grandifolia
Urticaceae_Pilea_microphylla
Urticaceae_Pilea_fontana
Urticaceae_Pilea_pumila
Urticaceae_Myriocarpa_longipes
Moraceae_Artocarpus_sepicanus
Moraceae_Artocarpus_heterophyllus
Moraceae_Artocarpus_styracifolius
Moraceae_Artocarpus_vriesianus
Moraceae_Artocarpus_lakoocha
Moraceae_Artocarpus_altilis
Moraceae_Artocarpus_camansi
Moraceae_Morus_insignis
Moraceae_Streblus_glaber
Moraceae_Streblus_heterophyllus
Moraceae_Morus_mesozygia
Moraceae_Bagassa_guianensis
Moraceae_Sorocea_affinis
Moraceae_Milicia_excelsa
Moraceae_Morus_notabilis
Moraceae_Morus_australis
Moraceae_Morus_wittiorum
Moraceae_Morus_lhou
Moraceae_Morus_rotundiloba
Moraceae_Morus_alba
Moraceae_Morus_mongolica
Moraceae_Morus_nigra
Moraceae_Morus_macroura
Moraceae_Morus_cathayana
Moraceae_Morus_bombycis
Moraceae_Morus_indica
Moraceae_Morus_rubra
Moraceae_Morus_serrata
Moraceae_Morus_celtidifolia
Moraceae_Maclura_tinctoria
Moraceae_Maclura_africana
Moraceae_Prainea_limpato
Moraceae_Streblus_pendulinus
Moraceae_Streblus_ascendens
Moraceae_Taxotrophis_aquifolioides

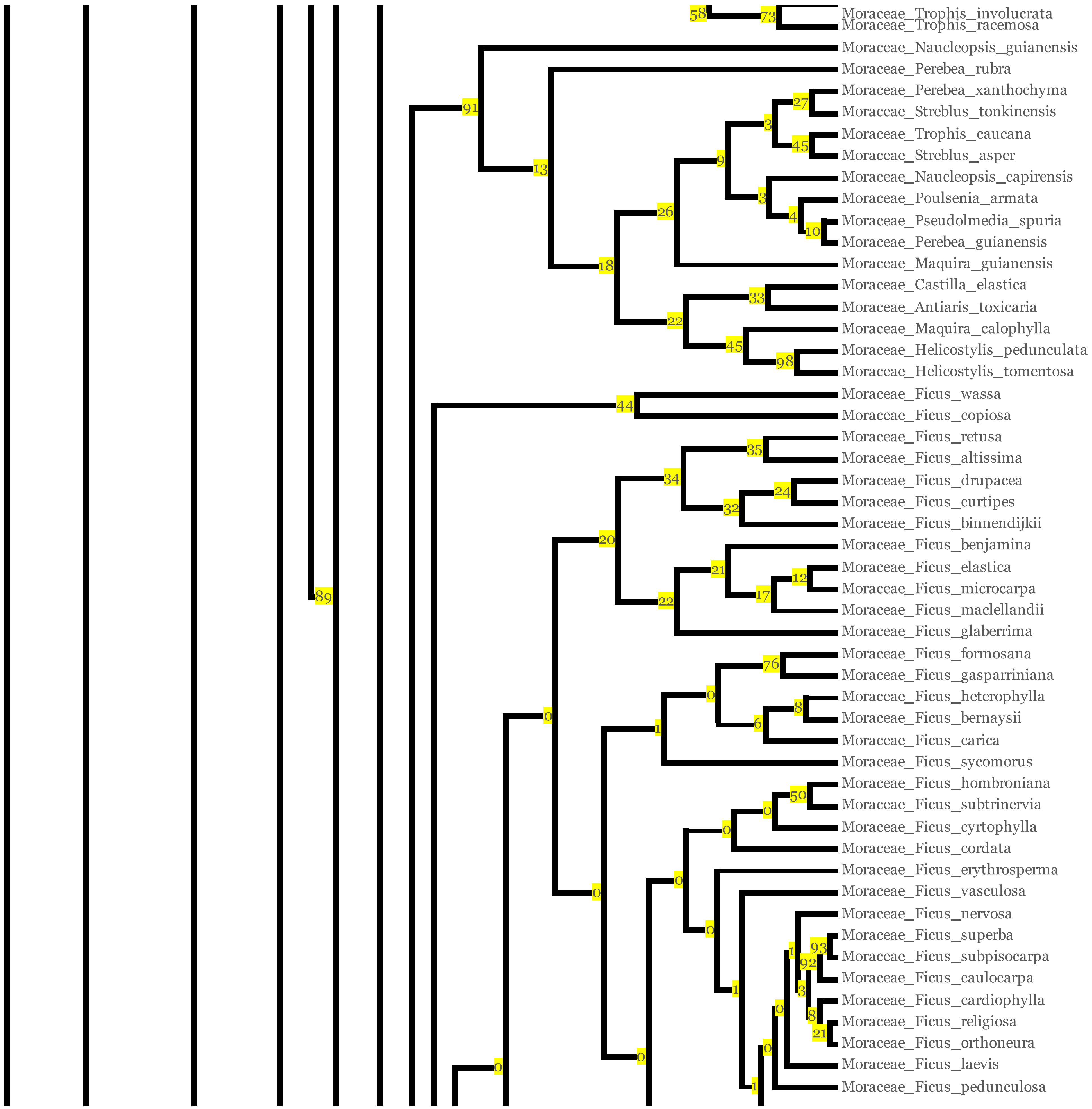

Moraceae_Trophis_involucrata
Moraceae_Trophis_racemosa
Moraceae_Naucleopsis_guianensis
Moraceae_Perebea_rubra
Moraceae_Perebea_xanthochyma
Moraceae_Streblus_tonkinensis
Moraceae_Trophis_caucana
Moraceae_Streblus_asper
Moraceae_Naucleopsis_capirensis
Moraceae_Poulsenia_armata
Moraceae_Pseudolmedia_spuria
Moraceae_Perebea_guianensis
Moraceae_Maquira_guianensis
Moraceae_Castilla_elastica
Moraceae_Antiaris_toxicaria
Moraceae_Maquira_calophylla
Moraceae_Helicostylis_pedunculata
Moraceae_Helicostylis_tomentosa
Moraceae_Ficus_wassa
Moraceae_Ficus_copiosa
Moraceae_Ficus_retusa
Moraceae_Ficus_altissima
Moraceae_Ficus_drupacea
Moraceae_Ficus_curtipes
Moraceae_Ficus_binnendijkii
Moraceae_Ficus_benjamina
Moraceae_Ficus_elastica
Moraceae_Ficus_microcarpa
Moraceae_Ficus_maclellandii
Moraceae_Ficus_glaberrima
Moraceae_Ficus_formosana
Moraceae_Ficus_gasparriniana
Moraceae_Ficus_heterophylla
Moraceae_Ficus_bernaysii
Moraceae_Ficus_carica
Moraceae_Ficus_sycomorus
Moraceae_Ficus_hombroniana
Moraceae_Ficus_subtrinervia
Moraceae_Ficus_cyrtophylla
Moraceae_Ficus_cordata
Moraceae_Ficus_erythrosperma
Moraceae_Ficus_vasculosa
Moraceae_Ficus_nervosa
Moraceae_Ficus_superba
Moraceae_Ficus_subpisocarpa
Moraceae_Ficus_caulocarpa
Moraceae_Ficus_cardiophylla
Moraceae_Ficus_religiosa
Moraceae_Ficus_orthoneura
Moraceae_Ficus_laevis
Moraceae_Ficus_pedunculosa

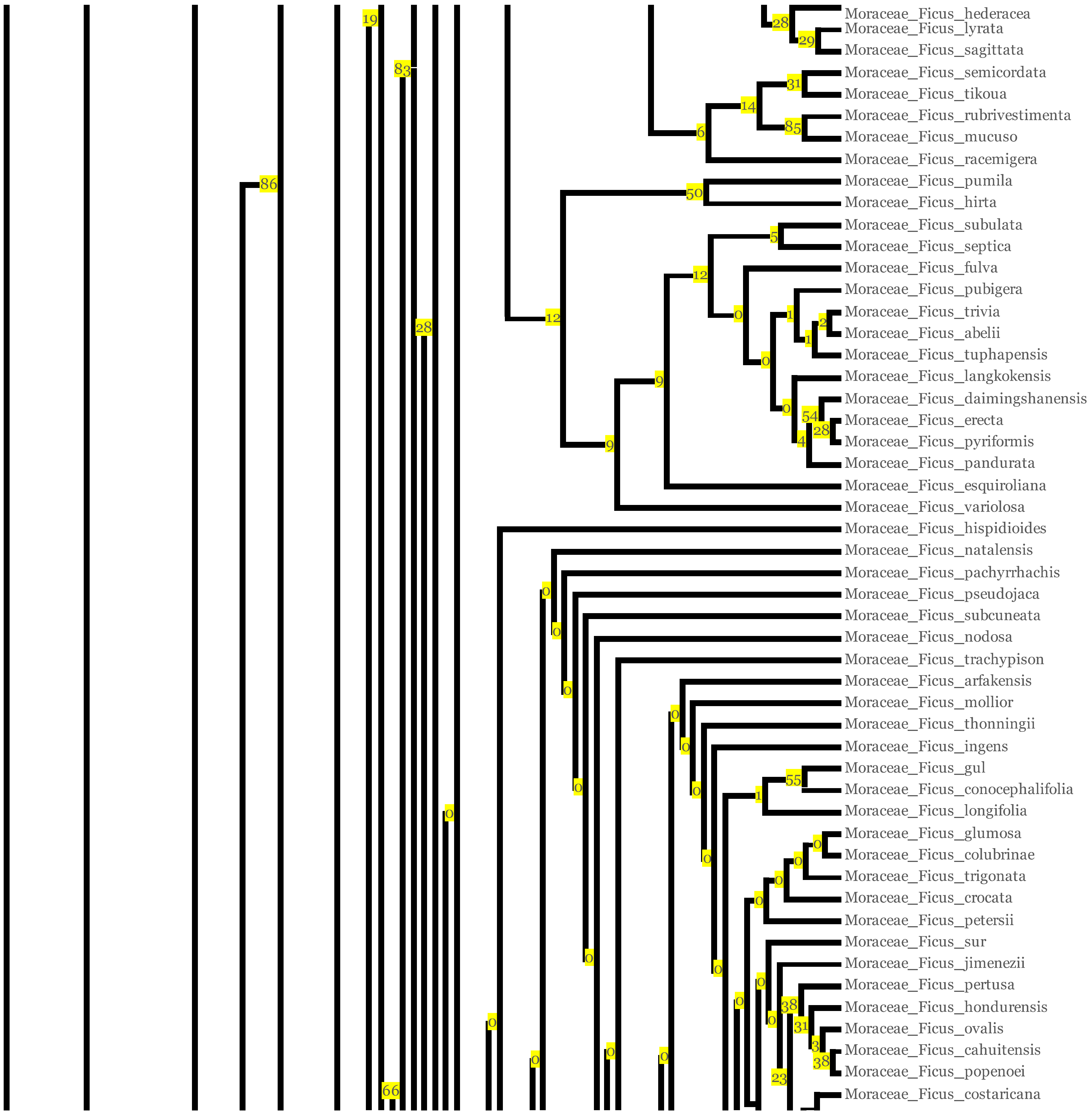
Moraceae_Ficus_hederacea
Moraceae_Ficus_lyrata
Moraceae_Ficus_sagittata
Moraceae_Ficus_semicordata
Moraceae_Ficus_tikoua
Moraceae_Ficus_rubrivestimenta
Moraceae_Ficus_mucuso
Moraceae_Ficus_racemigera
Moraceae_Ficus_pumila
Moraceae_Ficus_hirta
Moraceae_Ficus_subulata
Moraceae_Ficus_septica
Moraceae_Ficus_fulva
Moraceae_Ficus_pubigera
Moraceae_Ficus_trivia
Moraceae_Ficus_abelii
Moraceae_Ficus_tuphapensis
Moraceae_Ficus_langkokensis
Moraceae_Ficus_daimingshanensis
Moraceae_Ficus_erecta
Moraceae_Ficus_pyriformis
Moraceae_Ficus_pandurata
Moraceae_Ficus_esquiroliana
Moraceae_Ficus_variolosa
Moraceae_Ficus_hispidioides
Moraceae_Ficus_natalensis
Moraceae_Ficus_pachyrrhachis
Moraceae_Ficus_pseudojaca
Moraceae_Ficus_subcuneata
Moraceae_Ficus_nodosa
Moraceae_Ficus_trachypison
Moraceae_Ficus_arfakensis
Moraceae_Ficus_mollior
Moraceae_Ficus_thonningii
Moraceae_Ficus_ingens
Moraceae_Ficus_gul
Moraceae_Ficus_conocephalifolia
Moraceae_Ficus_longifolia
Moraceae_Ficus_glumosa
Moraceae_Ficus_colubrinae
Moraceae_Ficus_trigonata
Moraceae_Ficus_crocata
Moraceae_Ficus_petersii
Moraceae_Ficus_sur
Moraceae_Ficus_jimenezii
Moraceae_Ficus_pertusa
Moraceae_Ficus_hondurensis
Moraceae_Ficus_ovalis
Moraceae_Ficus_cahuitensis
Moraceae_Ficus_popenoei
Moraceae_Ficus_costaricana

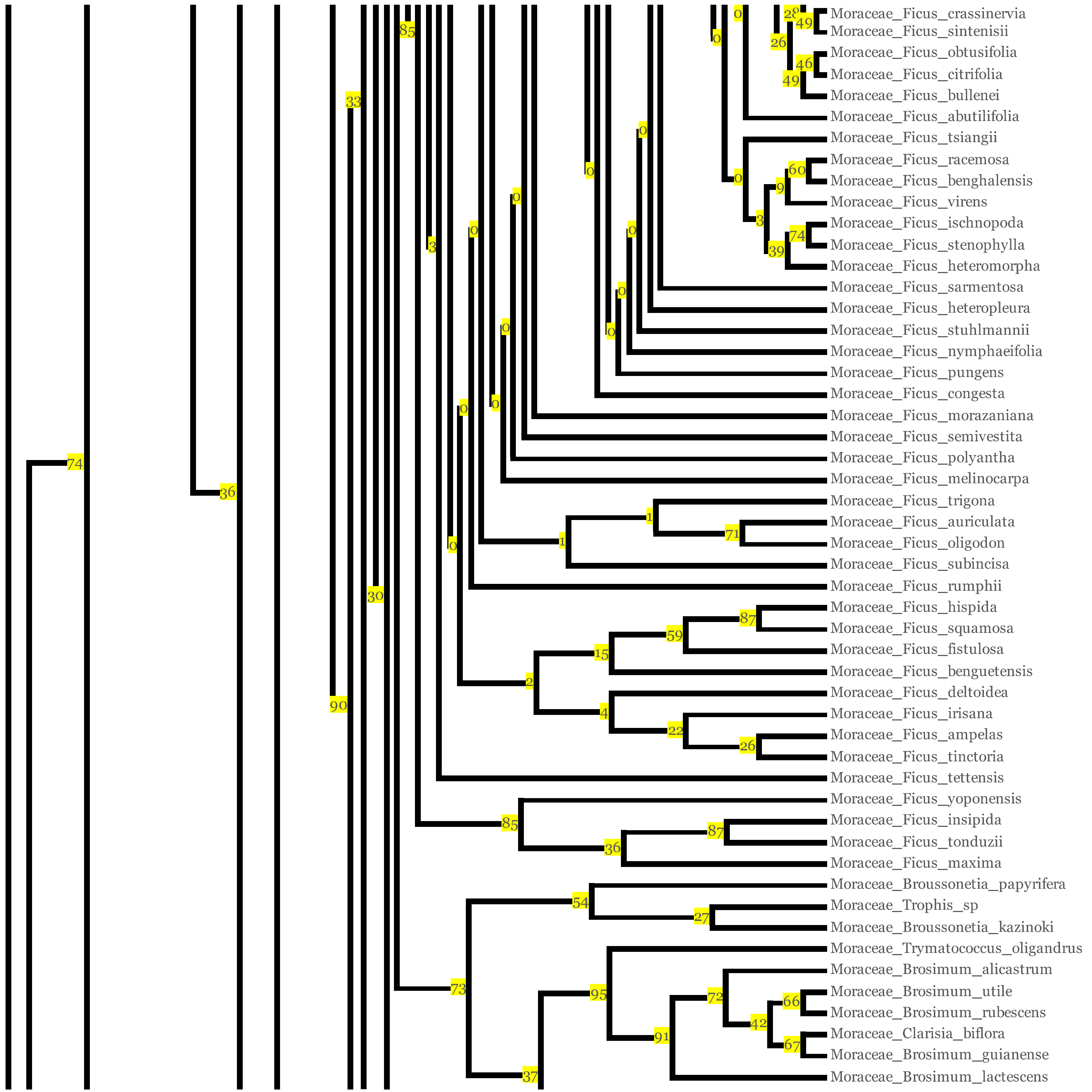

Moraceae_Ficus_crassinervia
Moraceae_Ficus_sintenisii
Moraceae_Ficus_obtusifolia
Moraceae_Ficus_citrifolia
Moraceae_Ficus_bullenei
Moraceae_Ficus_abutilifolia
Moraceae_Ficus_tsiangii
Moraceae_Ficus_racemosa
Moraceae_Ficus_benghalensis
Moraceae_Ficus_virens
Moraceae_Ficus_ischnopoda
Moraceae_Ficus_stenophylla
Moraceae_Ficus_heteromorpha
Moraceae_Ficus_sarmentosa
Moraceae_Ficus_heteropleura
Moraceae_Ficus_stuhlmannii
Moraceae_Ficus_nymphaeifolia
Moraceae_Ficus_pungens
Moraceae_Ficus_congesta
Moraceae_Ficus_morazaniana
Moraceae_Ficus_semivestita
Moraceae_Ficus_polyantha
Moraceae_Ficus_melinocarpa
Moraceae_Ficus_trigona
Moraceae_Ficus_auriculata
Moraceae_Ficus_oligodon
Moraceae_Ficus_subincisa
Moraceae_Ficus_rumphii
Moraceae_Ficus_hispida
Moraceae_Ficus_squamosa
Moraceae_Ficus_fistulosa
Moraceae_Ficus_benguetensis
Moraceae_Ficus_deltoidea
Moraceae_Ficus_irisana
Moraceae_Ficus_ampelas
Moraceae_Ficus_tinctoria
Moraceae_Ficus_tettensis
Moraceae_Ficus_yoponensis
Moraceae_Ficus_insipida
Moraceae_Ficus_tonduzii
Moraceae_Ficus_maxima
Moraceae_Broussonetia_papyrifera
Moraceae_Trophis_sp
Moraceae_Broussonetia_kazinoki
Moraceae_Trymatococcus_oligandrus
Moraceae_Brosimum_alicastrum
Moraceae_Brosimum_utile
Moraceae_Brosimum_rubescens
Moraceae_Clarisia_biflora
Moraceae_Brosimum_guianense
Moraceae_Brosimum_lactescens

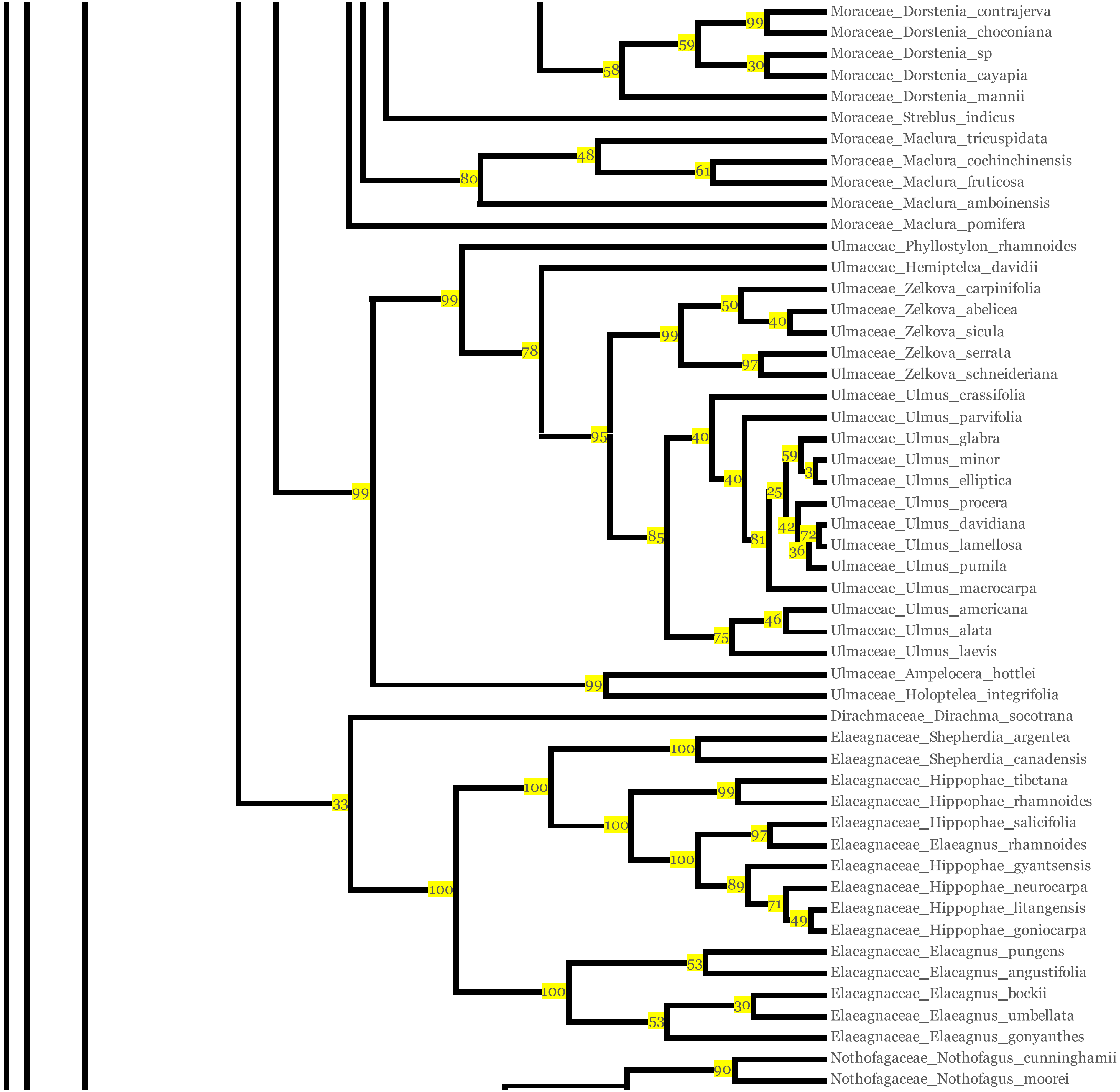
Moraceae_Dorstenia_contrajerva
Moraceae_Dorstenia_choconiana
Moraceae_Dorstenia_sp
Moraceae_Dorstenia_cayapia
Moraceae_Dorstenia_mannii
Moraceae_Streblus_indicus
Moraceae_Maclura_tricuspidata
Moraceae_Maclura_cochinchinensis
Moraceae_Maclura_fruticosa
Moraceae_Maclura_amboinensis
Moraceae_Maclura_pomifera
Ulmaceae_Phyllostylon_rhamnoides
Ulmaceae_Hemiptelea_davidii
Ulmaceae_Zelkova_carpinifolia
Ulmaceae_Zelkova_abelicea
Ulmaceae_Zelkova_sicula
Ulmaceae_Zelkova_serrata
Ulmaceae_Zelkova_schneideriana
Ulmaceae_Ulmus_crassifolia
Ulmaceae_Ulmus_parvifolia
Ulmaceae_Ulmus_glabra
Ulmaceae_Ulmus_minor
Ulmaceae_Ulmus_elliptica
Ulmaceae_Ulmus_procera
Ulmaceae_Ulmus_davidiana
Ulmaceae_Ulmus_lamellosa
Ulmaceae_Ulmus_pumila
Ulmaceae_Ulmus_macrocarpa
Ulmaceae_Ulmus_americana
Ulmaceae_Ulmus_alata
Ulmaceae_Ulmus_laevis
Ulmaceae_Ampelocera_hottlei
Ulmaceae_Holoptelea_integrifolia
Dirachmaceae_Dirachma_socotrana
Elaeagnaceae_Shepherdia_argentea
Elaeagnaceae_Shepherdia_canadensis
Elaeagnaceae_Hippophae_tibetana
Elaeagnaceae_Hippophae_rhamnoides
Elaeagnaceae_Hippophae_salicifolia
Elaeagnaceae_Elaeagnus_rhamnoides
Elaeagnaceae_Hippophae_gyantsensis
Elaeagnaceae_Hippophae_neurocarpa
Elaeagnaceae_Hippophae_litangensis
Elaeagnaceae_Hippophae_goniocarpa
Elaeagnaceae_Elaeagnus_pungens
Elaeagnaceae_Elaeagnus_angustifolia
Elaeagnaceae_Elaeagnus_bockii
Elaeagnaceae_Elaeagnus_umbellata
Elaeagnaceae_Elaeagnus_gonyanthes
Nothofagaceae_Nothofagus_cunninghamii
Nothofagaceae_Nothofagus_moorei

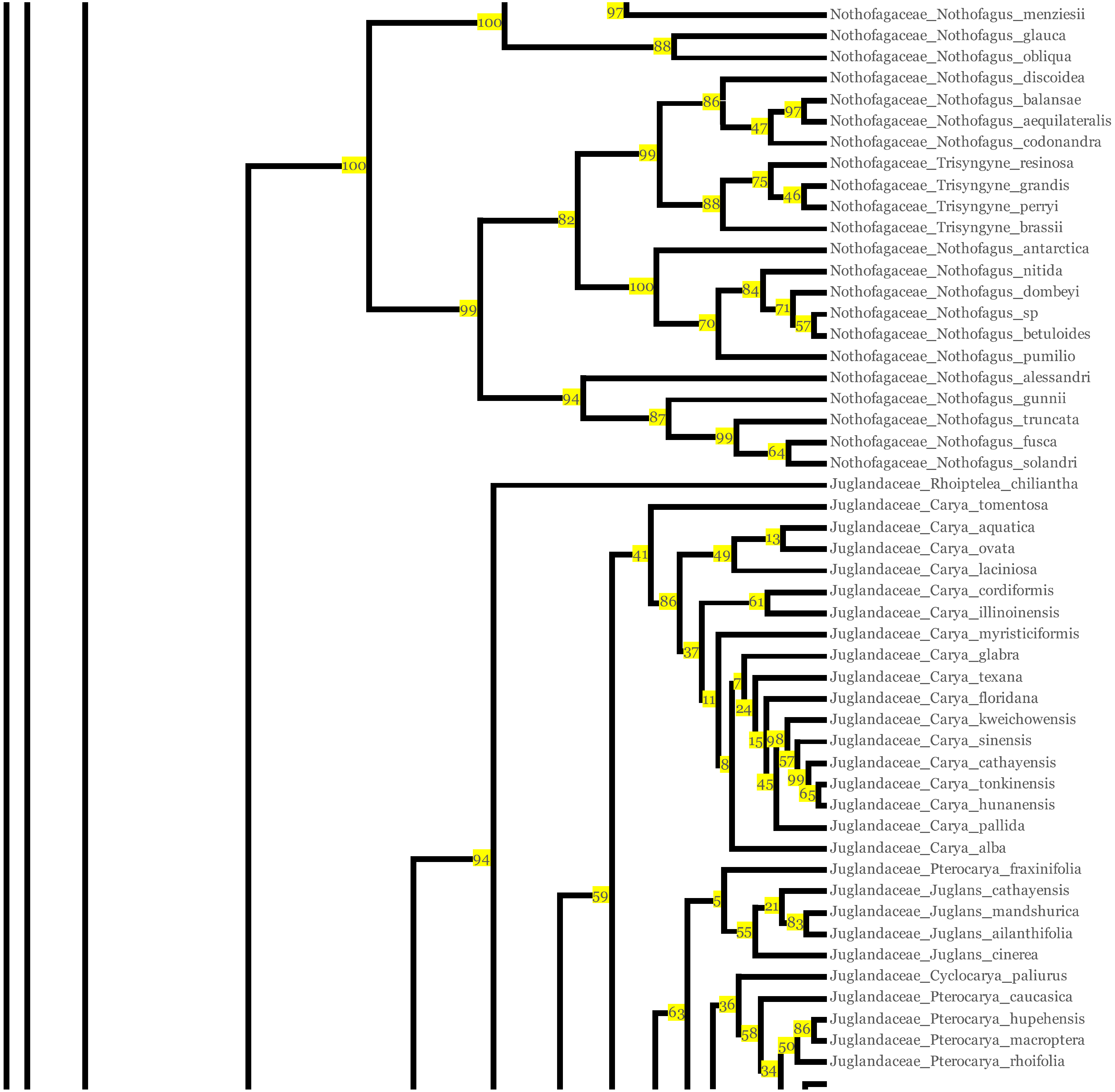
Nothofagaceae_Nothofagus_menziesii
Nothofagaceae_Nothofagus_glauca
Nothofagaceae_Nothofagus_obliqua
Nothofagaceae_Nothofagus_discoidea
Nothofagaceae_Nothofagus_balansae
Nothofagaceae_Nothofagus_aequilateralis
Nothofagaceae_Nothofagus_codonandra
Nothofagaceae_Trisyngyne_resinosa
Nothofagaceae_Trisyngyne_grandis
Nothofagaceae_Trisyngyne_perryi
Nothofagaceae_Trisyngyne_brassii
Nothofagaceae_Nothofagus_antarctica
Nothofagaceae_Nothofagus_nitida
Nothofagaceae_Nothofagus_dombeyi
Nothofagaceae_Nothofagus_sp
Nothofagaceae_Nothofagus_betuloides
Nothofagaceae_Nothofagus_pumilio
Nothofagaceae_Nothofagus_alessandri
Nothofagaceae_Nothofagus_gunnii
Nothofagaceae_Nothofagus_truncata
Nothofagaceae_Nothofagus_fusca
Nothofagaceae_Nothofagus_solandri
Juglandaceae_Rhoiptelea_chiliantha
Juglandaceae_Carya_tomentosa
Juglandaceae_Carya_aquatica
Juglandaceae_Carya_ovata
Juglandaceae_Carya_laciniosa
Juglandaceae_Carya_cordiformis
Juglandaceae_Carya_illinoinensis
Juglandaceae_Carya_myristiciformis
Juglandaceae_Carya_glabra
Juglandaceae_Carya_texana
Juglandaceae_Carya_floridana
Juglandaceae_Carya_kweichowensis
Juglandaceae_Carya_sinensis
Juglandaceae_Carya_cathayensis
Juglandaceae_Carya_tonkinensis
Juglandaceae_Carya_hunanensis
Juglandaceae_Carya_pallida
Juglandaceae_Carya_alba
Juglandaceae_Pterocarya_fraxinifolia
Juglandaceae_Juglans_cathayensis
Juglandaceae_Juglans_mandshurica
Juglandaceae_Juglans_ailanthifolia
Juglandaceae_Juglans_cinerea
Juglandaceae_Cyclocarya_paliurus
Juglandaceae_Pterocarya_caucasica
Juglandaceae_Pterocarya_hupehensis
Juglandaceae_Pterocarya_macroptera
Juglandaceae_Pterocarya_rhoifolia

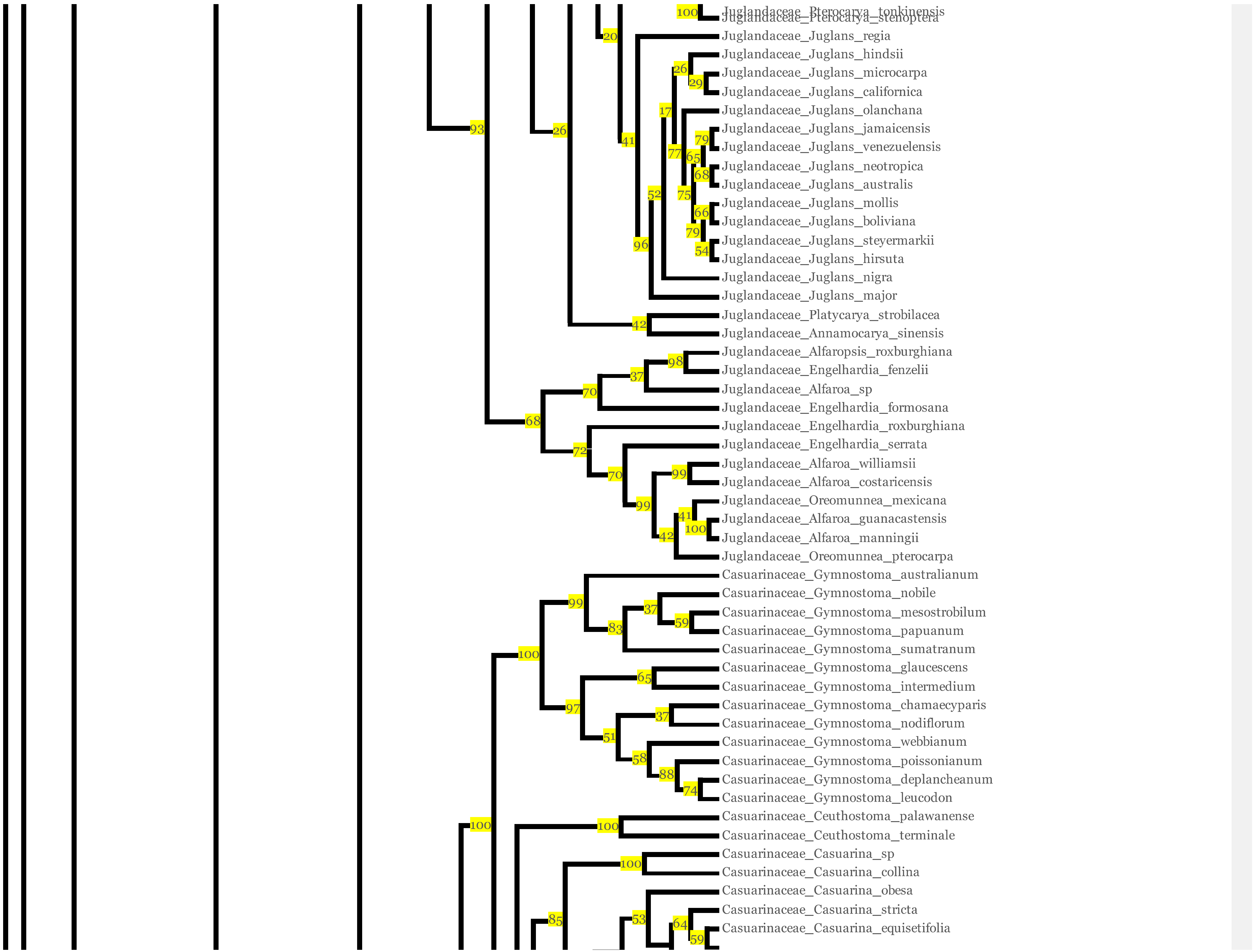
Juglandaceae_Pterocarya_tonkinensis
Juglandaceae_Pterocarya_stenoptera
Juglandaceae_Juglans_regia
Juglandaceae_Juglans_hindsii
Juglandaceae_Juglans_microcarpa
Juglandaceae_Juglans_californica
Juglandaceae_Juglans_olanchana
Juglandaceae_Juglans_jamaicensis
Juglandaceae_Juglans_venezuelensis
Juglandaceae_Juglans_neotropica
Juglandaceae_Juglans_australis
Juglandaceae_Juglans_mollis
Juglandaceae_Juglans_boliviana
Juglandaceae_Juglans_steyermarkii
Juglandaceae_Juglans_hirsuta
Juglandaceae_Juglans_nigra
Juglandaceae_Juglans_major
Juglandaceae_Platycarya_strobilacea
Juglandaceae_Annamocarya_sinensis
Juglandaceae_Alfaropsis_roxburghiana
Juglandaceae_Engelhardia_fenzelii
Juglandaceae_Alfaroa_sp
Juglandaceae_Engelhardia_formosana
Juglandaceae_Engelhardia_roxburghiana
Juglandaceae_Engelhardia_serrata
Juglandaceae_Alfaroa_williamsii
Juglandaceae_Alfaroa_costaricensis
Juglandaceae_Oreomunnea_mexicana
Juglandaceae_Alfaroa_guanacastensis
Juglandaceae_Alfaroa_manningii
Juglandaceae_Oreomunnea_pterocarpa
Casuarinaceae_Gymnostoma_australianum
Casuarinaceae_Gymnostoma_nobile
Casuarinaceae_Gymnostoma_mesostrobilum
Casuarinaceae_Gymnostoma_papuanum
Casuarinaceae_Gymnostoma_sumatranum
Casuarinaceae_Gymnostoma_glaucescens
Casuarinaceae_Gymnostoma_intermedium
Casuarinaceae_Gymnostoma_chamaecyparis
Casuarinaceae_Gymnostoma_nodiflorum
Casuarinaceae_Gymnostoma_webbianum
Casuarinaceae_Gymnostoma_poissonianum
Casuarinaceae_Gymnostoma_deplancheanum
Casuarinaceae_Gymnostoma_leucodon
Casuarinaceae_Ceuthostoma_palawanense
Casuarinaceae_Ceuthostoma_terminale
Casuarinaceae_Casuarina_sp
Casuarinaceae_Casuarina_collina
Casuarinaceae_Casuarina_obesa
Casuarinaceae_Casuarina_stricta
Casuarinaceae_Casuarina_equisetifolia

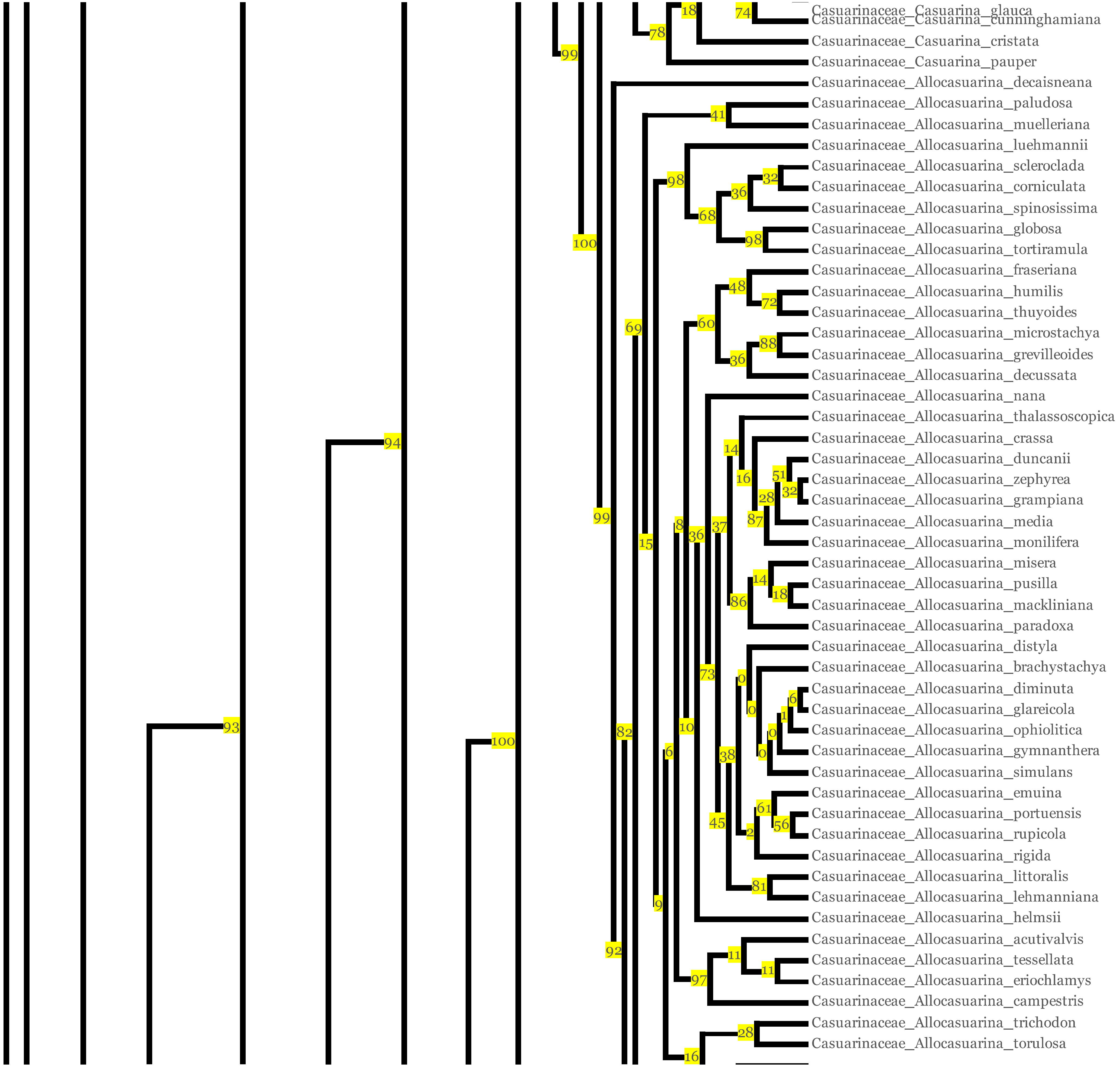
Casuarinaceae_Casuarina_glauca
Casuarinaceae_Casuarina_cunninghamiana
Casuarinaceae_Casuarina_cristata
Casuarinaceae_Casuarina_pauper
Casuarinaceae_Allocasuarina_decaisneana
Casuarinaceae_Allocasuarina_paludosa
Casuarinaceae_Allocasuarina_muelleriana
Casuarinaceae_Allocasuarina_luehmannii
Casuarinaceae_Allocasuarina_scleroclada
Casuarinaceae_Allocasuarina_corniculata
Casuarinaceae_Allocasuarina_spinosissima
Casuarinaceae_Allocasuarina_globosa
Casuarinaceae_Allocasuarina_tortiramula
Casuarinaceae_Allocasuarina_fraseriana
Casuarinaceae_Allocasuarina_humilis
Casuarinaceae_Allocasuarina_thuyoides
Casuarinaceae_Allocasuarina_microstachya
Casuarinaceae_Allocasuarina_grevilleoides
Casuarinaceae_Allocasuarina_decussata
Casuarinaceae_Allocasuarina_nana
Casuarinaceae_Allocasuarina_thalassoscopica
Casuarinaceae_Allocasuarina_crassa
Casuarinaceae_Allocasuarina_duncanii
Casuarinaceae_Allocasuarina_zephyrea
Casuarinaceae_Allocasuarina_grampiana
Casuarinaceae_Allocasuarina_media
Casuarinaceae_Allocasuarina_monilifera
Casuarinaceae_Allocasuarina_misera
Casuarinaceae_Allocasuarina_pusilla
Casuarinaceae_Allocasuarina_mackliniana
Casuarinaceae_Allocasuarina_paradoxa
Casuarinaceae_Allocasuarina_distyla
Casuarinaceae_Allocasuarina_brachystachya
Casuarinaceae_Allocasuarina_diminuta
Casuarinaceae_Allocasuarina_glareicola
Casuarinaceae_Allocasuarina_ophiolitica
Casuarinaceae_Allocasuarina_gymnanthera
Casuarinaceae_Allocasuarina_simulans
Casuarinaceae_Allocasuarina_emuina
Casuarinaceae_Allocasuarina_portuensis
Casuarinaceae_Allocasuarina_rupicola
Casuarinaceae_Allocasuarina_rigida
Casuarinaceae_Allocasuarina_littoralis
Casuarinaceae_Allocasuarina_lehmanniana
Casuarinaceae_Allocasuarina_helmsii
Casuarinaceae_Allocasuarina_acutivalvis
Casuarinaceae_Allocasuarina_tessellata
Casuarinaceae_Allocasuarina_eriochlamys
Casuarinaceae_Allocasuarina_campestris
Casuarinaceae_Allocasuarina_trichodon
Casuarinaceae_Allocasuarina_torulosa

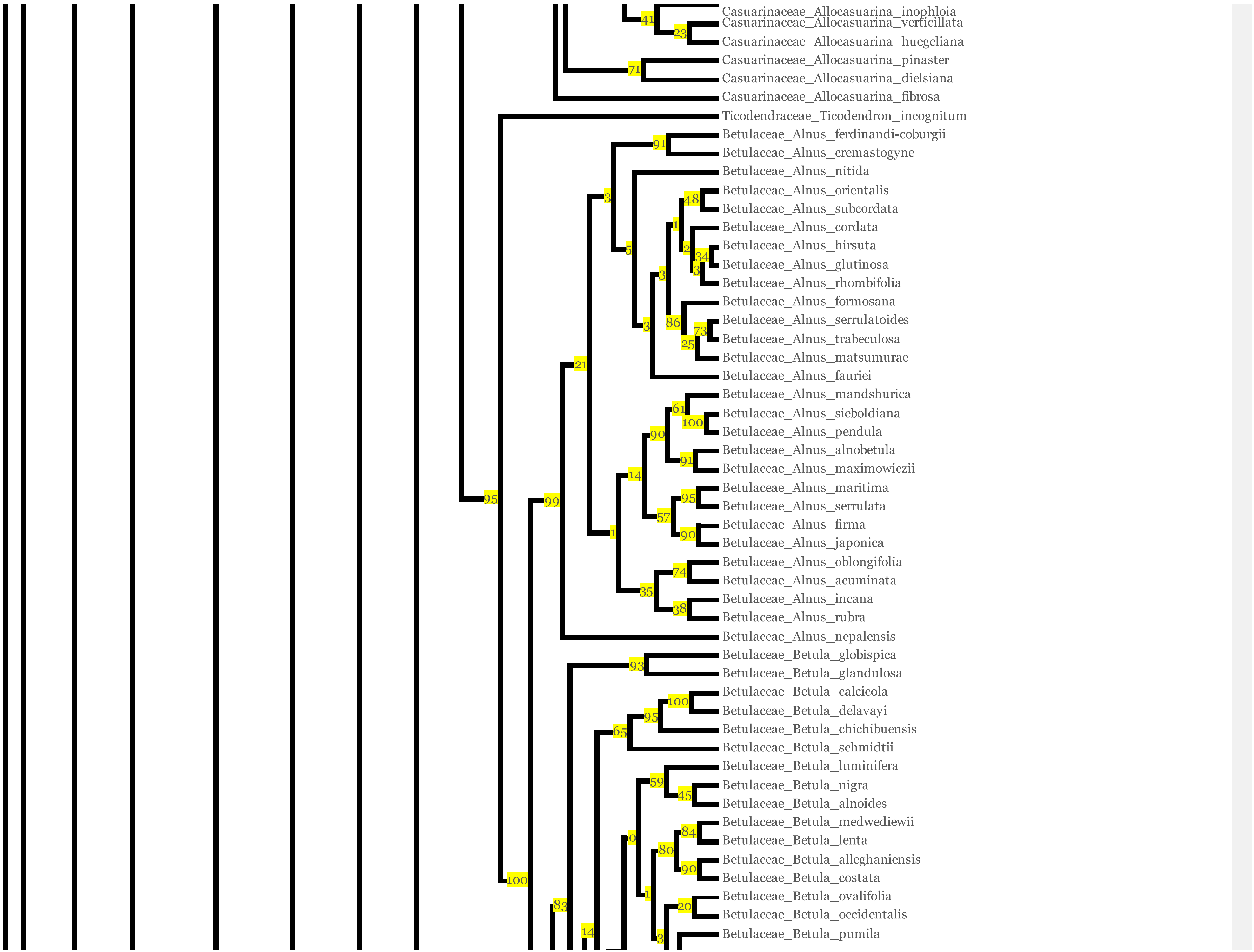
Casuarinaceae_Allocasuarina_inophloia
Casuarinaceae_Allocasuarina_verticillata
Casuarinaceae_Allocasuarina_huegeliana
Casuarinaceae_Allocasuarina_pinaster
Casuarinaceae_Allocasuarina_dielsiana
Casuarinaceae_Allocasuarina_fibrosa
Ticodendraceae_Ticodendron_incognitum
Betulaceae_Alnus_ferdinandi-coburgii
Betulaceae_Alnus_cremastogyne
Betulaceae_Alnus_nitida
Betulaceae_Alnus_orientalis
Betulaceae_Alnus_subcordata
Betulaceae_Alnus_cordata
Betulaceae_Alnus_hirsuta
Betulaceae_Alnus_glutinosa
Betulaceae_Alnus_rhombifolia
Betulaceae_Alnus_formosana
Betulaceae_Alnus_serrulatoides
Betulaceae_Alnus_trabeculosa
Betulaceae_Alnus_matsumurae
Betulaceae_Alnus_fauriei
Betulaceae_Alnus_mandshurica
Betulaceae_Alnus_sieboldiana
Betulaceae_Alnus_pendula
Betulaceae_Alnus_alnobetula
Betulaceae_Alnus_maximowiczii
Betulaceae_Alnus_maritima
Betulaceae_Alnus_serrulata
Betulaceae_Alnus_firma
Betulaceae_Alnus_japonica
Betulaceae_Alnus_oblongifolia
Betulaceae_Alnus_acuminata
Betulaceae_Alnus_incana
Betulaceae_Alnus_rubra
Betulaceae_Alnus_nepalensis
Betulaceae_Betula_globispica
Betulaceae_Betula_glandulosa
Betulaceae_Betula_calcicola
Betulaceae_Betula_delavayi
Betulaceae_Betula_chichibuensis
Betulaceae_Betula_schmidtii
Betulaceae_Betula_luminifera
Betulaceae_Betula_nigra
Betulaceae_Betula_alnoides
Betulaceae_Betula_medwediewii
Betulaceae_Betula_lenta
Betulaceae_Betula_alleghaniensis
Betulaceae_Betula_costata
Betulaceae_Betula_ovalifolia
Betulaceae_Betula_occidentalis
Betulaceae_Betula_pumila
41
23
71
91
3
48
1
5
2
34
3
3
86
73
3
25
21
61
100
90
91
14
95
57
95
99
1
90
74
35
38
93
100
95
65
59
45
0
84
80
90
100
83
1
20
14
3

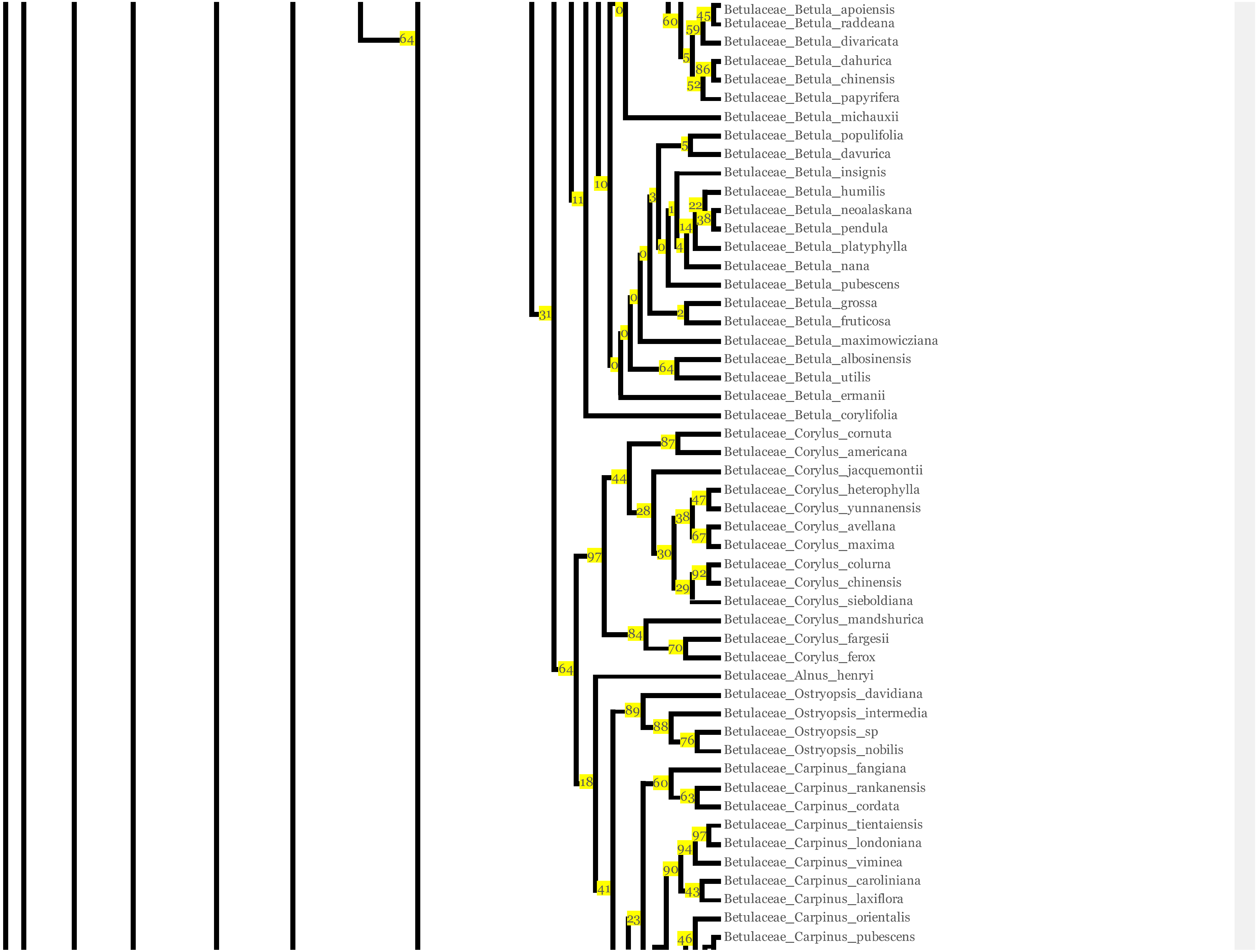
Betulaceae_Betula_apoiensis
Betulaceae_Betula_raddeana
Betulaceae_Betula_divaricata
Betulaceae_Betula_dahurica
Betulaceae_Betula_chinensis
Betulaceae_Betula_papyrifera
Betulaceae_Betula_michauxii
Betulaceae_Betula_populifolia
Betulaceae_Betula_davurica
Betulaceae_Betula_insignis
Betulaceae_Betula_humilis
Betulaceae_Betula_neoalaskana
Betulaceae_Betula_pendula
Betulaceae_Betula_platyphylla
Betulaceae_Betula_nana
Betulaceae_Betula_pubescens
Betulaceae_Betula_grossa
Betulaceae_Betula_fruticosa
Betulaceae_Betula_maximowicziana
Betulaceae_Betula_albosinensis
Betulaceae_Betula_utilis
Betulaceae_Betula_ermanii
Betulaceae_Betula_corylifolia
Betulaceae_Corylus_cornuta
Betulaceae_Corylus_americana
Betulaceae_Corylus_jacquemontii
Betulaceae_Corylus_heterophylla
Betulaceae_Corylus_yunnanensis
Betulaceae_Corylus_avellana
Betulaceae_Corylus_maxima
Betulaceae_Corylus_colurna
Betulaceae_Corylus_chinensis
Betulaceae_Corylus_sieboldiana
Betulaceae_Corylus_mandshurica
Betulaceae_Corylus_fargesii
Betulaceae_Corylus_ferox
Betulaceae_Alnus_henryi
Betulaceae_Ostryopsis_davidiana
Betulaceae_Ostryopsis_intermedia
Betulaceae_Ostryopsis_sp
Betulaceae_Ostryopsis_nobilis
Betulaceae_Carpinus_fangiana
Betulaceae_Carpinus_rankanensis
Betulaceae_Carpinus_cordata
Betulaceae_Carpinus_tientaiensis
Betulaceae_Carpinus_londoniana
Betulaceae_Carpinus_viminea
Betulaceae_Carpinus_caroliniana
Betulaceae_Carpinus_laxiflora
Betulaceae_Carpinus_orientalis
Betulaceae_Carpinus_pubescens

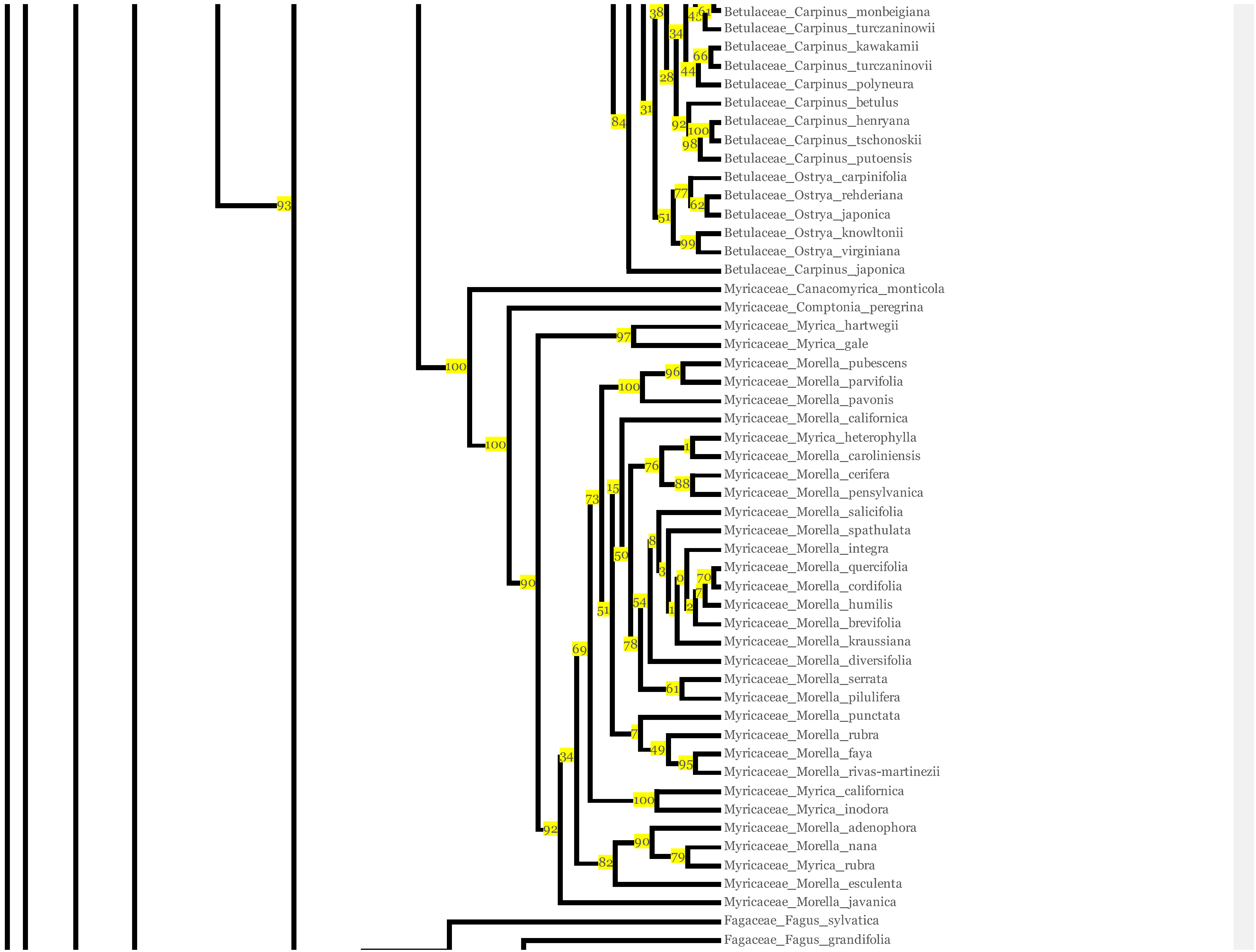
Betulaceae_Carpinus_monbeigiana
Betulaceae_Carpinus_turczaninowii
Betulaceae_Carpinus_kawakamii
Betulaceae_Carpinus_turczaninovii
Betulaceae_Carpinus_polyneura
Betulaceae_Carpinus_betulus
Betulaceae_Carpinus_henryana
Betulaceae_Carpinus_tschonoskii
Betulaceae_Carpinus_putoensis
Betulaceae_Ostrya_carpinifolia
Betulaceae_Ostrya_rehderiana
Betulaceae_Ostrya_japonica
Betulaceae_Ostrya_knowltonii
Betulaceae_Ostrya_virginiana
Betulaceae_Carpinus_japonica
Myricaceae_Canacomyrica_monticola
Myricaceae_Comptonia_peregrina
Myricaceae_Myrica_hartwegii
Myricaceae_Myrica_gale
Myricaceae_Morella_pubescens
Myricaceae_Morella_parvifolia
Myricaceae_Morella_pavonis
Myricaceae_Morella_californica
Myricaceae_Myrica_heterophylla
Myricaceae_Morella_caroliniensis
Myricaceae_Morella_cerifera
Myricaceae_Morella_pensylvanica
Myricaceae_Morella_salicifolia
Myricaceae_Morella_spathulata
Myricaceae_Morella_integra
Myricaceae_Morella_quercifolia
Myricaceae_Morella_cordifolia
Myricaceae_Morella_humilis
Myricaceae_Morella_brevifolia
Myricaceae_Morella_kraussiana
Myricaceae_Morella_diversifolia
Myricaceae_Morella_serrata
Myricaceae_Morella_pilulifera
Myricaceae_Morella_punctata
Myricaceae_Morella_rubra
Myricaceae_Morella_faya
Myricaceae_Morella_rivas-martinezii
Myricaceae_Myrica_californica
Myricaceae_Myrica_inodora
Myricaceae_Morella_adenophora
Myricaceae_Morella_nana
Myricaceae_Myrica_rubra
Myricaceae_Morella_esculenta
Myricaceae_Morella_javanica
Fagaceae_Fagus_sylvatica
Fagaceae_Fagus_grandifolia

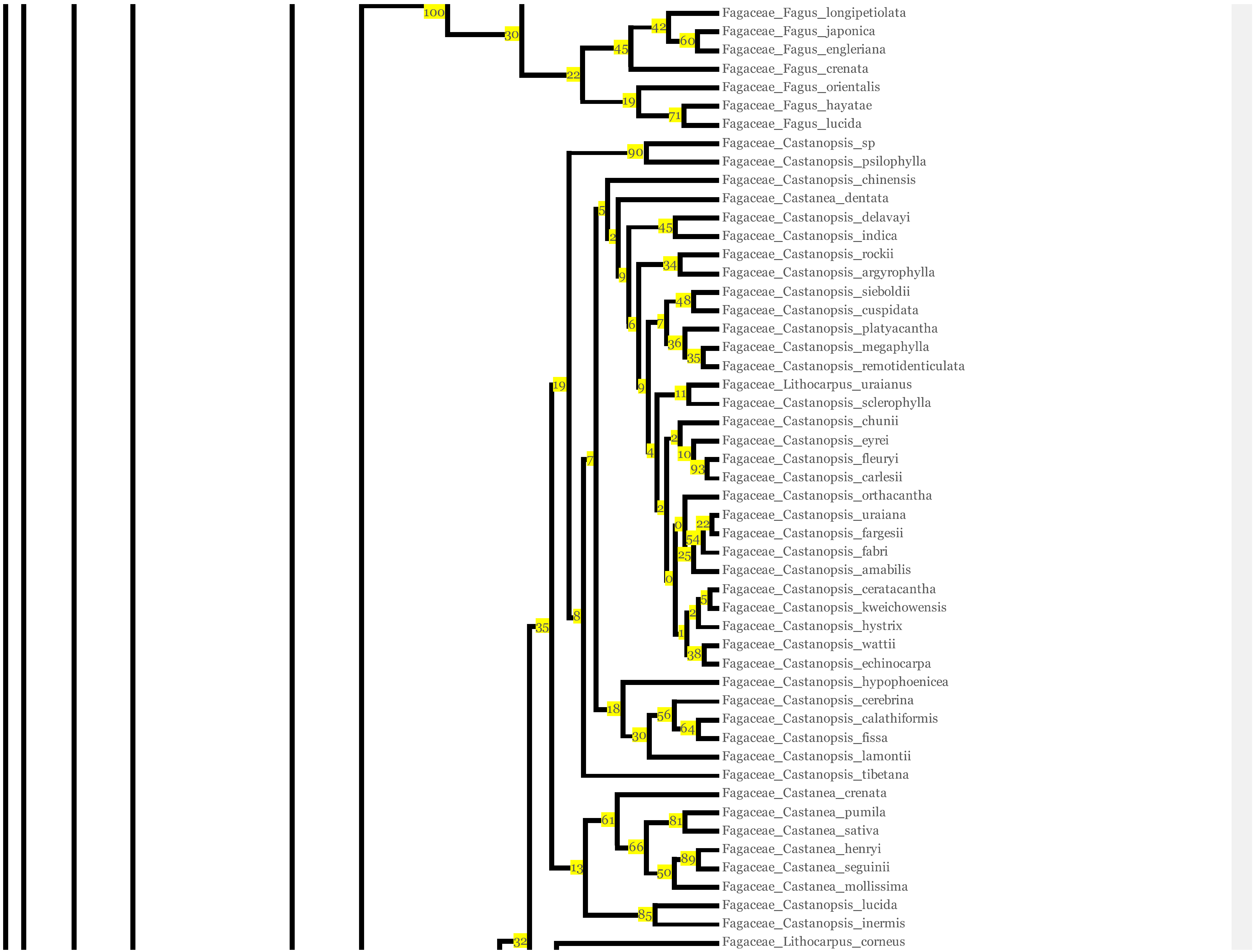
Fagaceae_Fagus_longipetiolata
Fagaceae_Fagus_japonica
Fagaceae_Fagus_engleriana
Fagaceae_Fagus_crenata
Fagaceae_Fagus_orientalis
Fagaceae_Fagus_hayatae
Fagaceae_Fagus_lucida
Fagaceae_Castanopsis_sp
Fagaceae_Castanopsis_psilophylla
Fagaceae_Castanopsis_chinensis
Fagaceae_Castanea_dentata
Fagaceae_Castanopsis_delavayi
Fagaceae_Castanopsis_indica
Fagaceae_Castanopsis_rockii
Fagaceae_Castanopsis_argyrophylla
Fagaceae_Castanopsis_sieboldii
Fagaceae_Castanopsis_cuspidata
Fagaceae_Castanopsis_platyacantha
Fagaceae_Castanopsis_megaphylla
Fagaceae_Castanopsis_remotidenticulata
Fagaceae_Lithocarpus_uraianus
Fagaceae_Castanopsis_sclerophylla
Fagaceae_Castanopsis_chunii
Fagaceae_Castanopsis_eyrei
Fagaceae_Castanopsis_fleuryi
Fagaceae_Castanopsis_carlesii
Fagaceae_Castanopsis_orthacantha
Fagaceae_Castanopsis_uraiana
Fagaceae_Castanopsis_fargesii
Fagaceae_Castanopsis_fabri
Fagaceae_Castanopsis_amabilis
Fagaceae_Castanopsis_ceratacantha
Fagaceae_Castanopsis_kweichowensis
Fagaceae_Castanopsis_hystrix
Fagaceae_Castanopsis_wattii
Fagaceae_Castanopsis_echinocarpa
Fagaceae_Castanopsis_hypophoenicea
Fagaceae_Castanopsis_cerebrina
Fagaceae_Castanopsis_calathiformis
Fagaceae_Castanopsis_fissa
Fagaceae_Castanopsis_lamontii
Fagaceae_Castanopsis_tibetana
Fagaceae_Castanea_crenata
Fagaceae_Castanea_pumila
Fagaceae_Castanea_sativa
Fagaceae_Castanea_henryi
Fagaceae_Castanea_seguinii
Fagaceae_Castanea_mollissima
Fagaceae_Castanopsis_lucida
Fagaceae_Castanopsis_inermis
Fagaceae_Lithocarpus_corneus

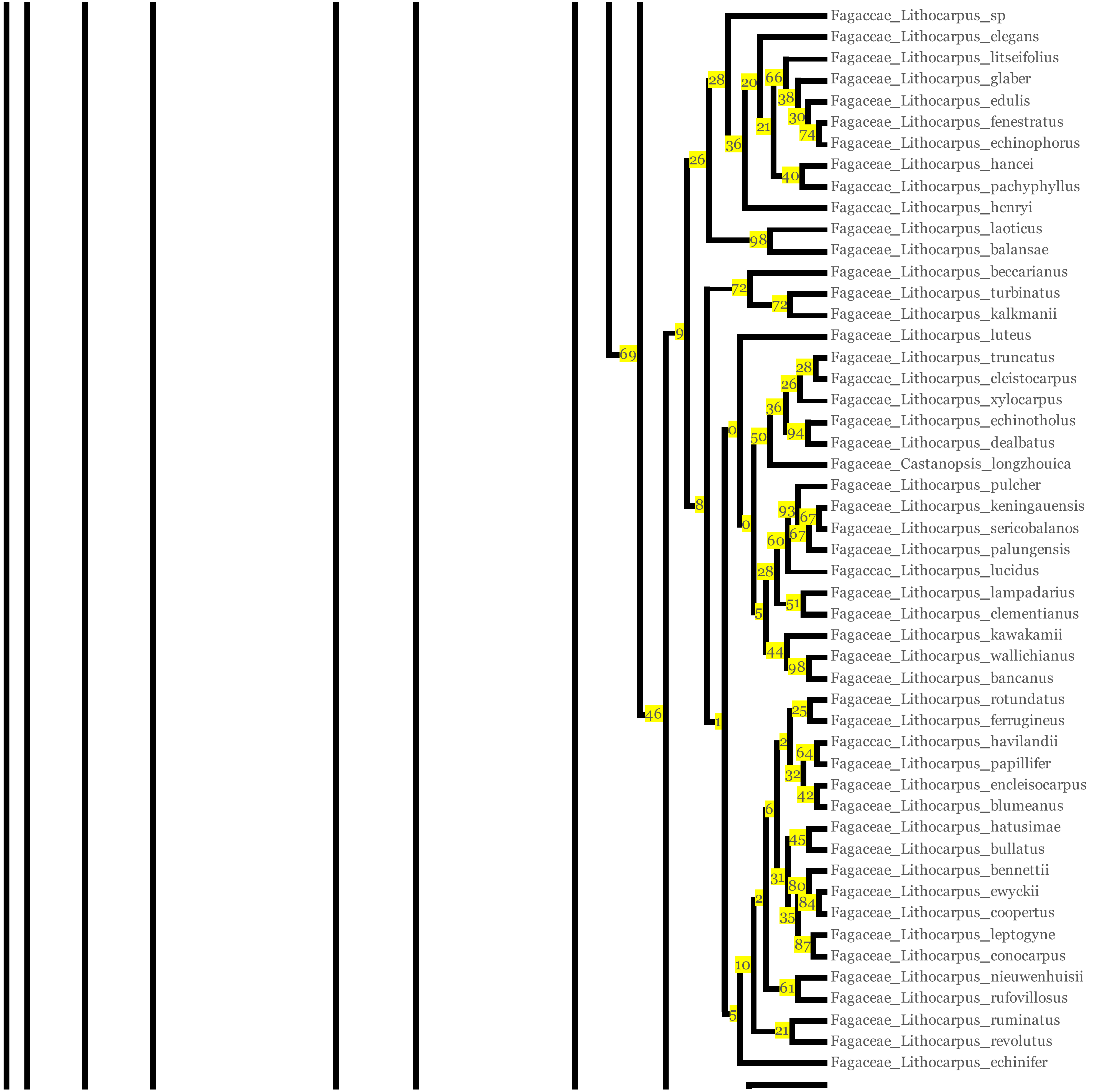
Fagaceae_Lithocarpus_sp
Fagaceae_Lithocarpus_elegans
Fagaceae_Lithocarpus_litseifolius
Fagaceae_Lithocarpus_glaber
Fagaceae_Lithocarpus_edulis
Fagaceae_Lithocarpus_fenestratus
Fagaceae_Lithocarpus_echinophorus
Fagaceae_Lithocarpus_hancei
Fagaceae_Lithocarpus_pachyphyllus
Fagaceae_Lithocarpus_henryi
Fagaceae_Lithocarpus_laoticus
Fagaceae_Lithocarpus_balansae
Fagaceae_Lithocarpus_beccarianus
Fagaceae_Lithocarpus_turbinatus
Fagaceae_Lithocarpus_kalkmanii
Fagaceae_Lithocarpus_luteus
Fagaceae_Lithocarpus_truncatus
Fagaceae_Lithocarpus_cleistocarpus
Fagaceae_Lithocarpus_xylocarpus
Fagaceae_Lithocarpus_echinotholus
Fagaceae_Lithocarpus_dealbatus
Fagaceae_Castanopsis_longzhouica
Fagaceae_Lithocarpus_pulcher
Fagaceae_Lithocarpus_keningauensis
Fagaceae_Lithocarpus_sericobalanos
Fagaceae_Lithocarpus_palungensis
Fagaceae_Lithocarpus_lucidus
Fagaceae_Lithocarpus_lampadarius
Fagaceae_Lithocarpus_clementianus
Fagaceae_Lithocarpus_kawakamii
Fagaceae_Lithocarpus_wallichianus
Fagaceae_Lithocarpus_bancanus
Fagaceae_Lithocarpus_rotundatus
Fagaceae_Lithocarpus_ferrugineus
Fagaceae_Lithocarpus_havilandii
Fagaceae_Lithocarpus_papillifer
Fagaceae_Lithocarpus_encleisocarpus
Fagaceae_Lithocarpus_blumeanus
Fagaceae_Lithocarpus_hatusimae
Fagaceae_Lithocarpus_bullatus
Fagaceae_Lithocarpus_bennettii
Fagaceae_Lithocarpus_ewyckii
Fagaceae_Lithocarpus_coopertus
Fagaceae_Lithocarpus_leptogyne
Fagaceae_Lithocarpus_conocarpus
Fagaceae_Lithocarpus_nieuwenhuisii
Fagaceae_Lithocarpus_rufovillosus
Fagaceae_Lithocarpus_ruminatus
Fagaceae_Lithocarpus_revolutus
Fagaceae_Lithocarpus_echinifer

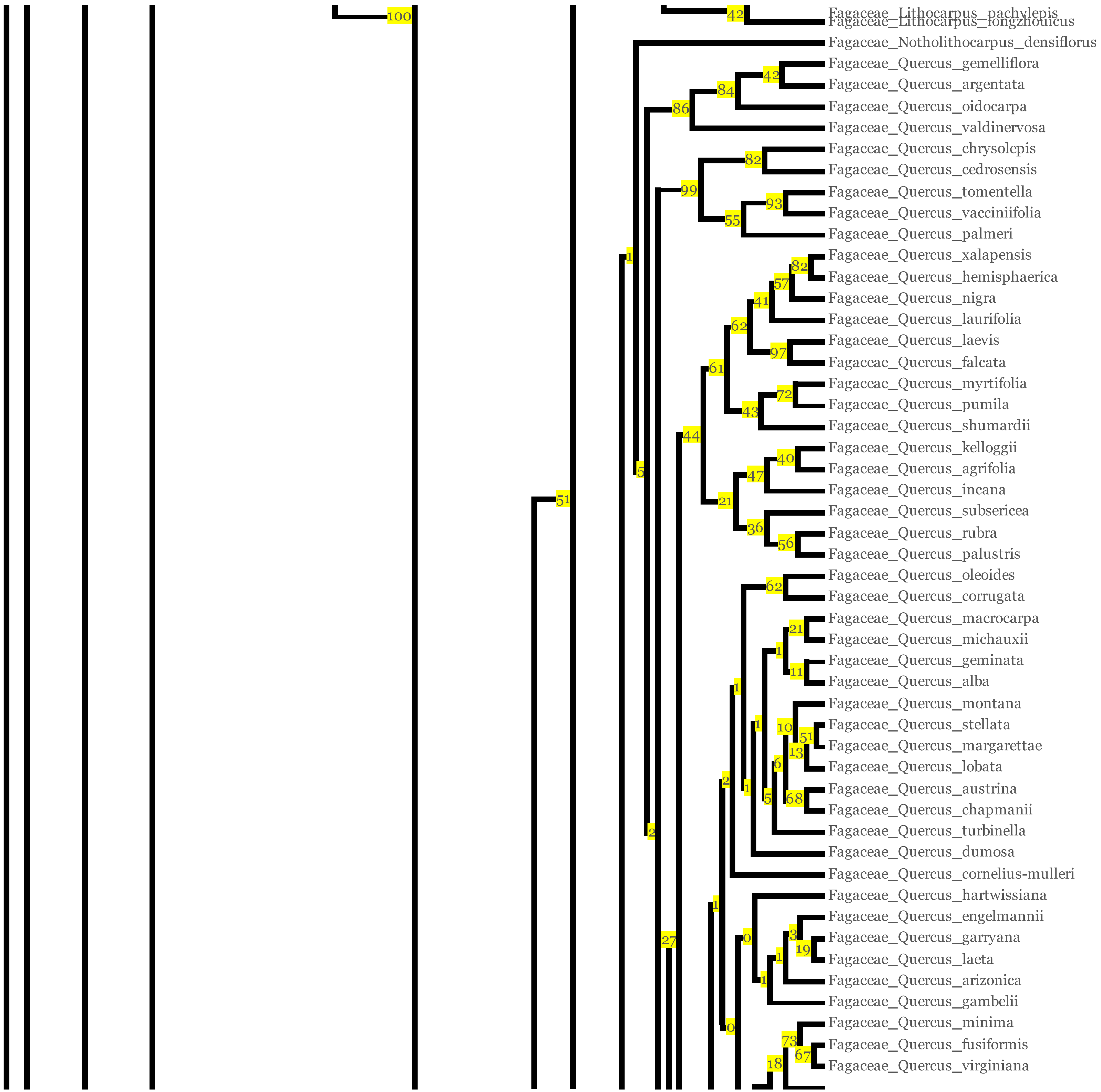
Fagaceae_Lithocarpus_pachylepis
Fagaceae_Lithocarpus_longzhouicus
Fagaceae_Notholithocarpus_densiflorus
Fagaceae_Quercus_gemelliflora
Fagaceae_Quercus_argentata
Fagaceae_Quercus_oidocarpa
Fagaceae_Quercus_valdinervosa
Fagaceae_Quercus_chrysolepis
Fagaceae_Quercus_cedrosensis
Fagaceae_Quercus_tomentella
Fagaceae_Quercus_vacciniifolia
Fagaceae_Quercus_palmeri
Fagaceae_Quercus_xalapensis
Fagaceae_Quercus_hemisphaerica
Fagaceae_Quercus_nigra
Fagaceae_Quercus_laurifolia
Fagaceae_Quercus_laevis
Fagaceae_Quercus_falcata
Fagaceae_Quercus_myrtifolia
Fagaceae_Quercus_pumila
Fagaceae_Quercus_shumardii
Fagaceae_Quercus_kelloggii
Fagaceae_Quercus_agrifolia
Fagaceae_Quercus_incana
Fagaceae_Quercus_subsericea
Fagaceae_Quercus_rubra
Fagaceae_Quercus_palustris
Fagaceae_Quercus_oleoides
Fagaceae_Quercus_corrugata
Fagaceae_Quercus_macrocarpa
Fagaceae_Quercus_michauxii
Fagaceae_Quercus_geminata
Fagaceae_Quercus_alba
Fagaceae_Quercus_montana
Fagaceae_Quercus_stellata
Fagaceae_Quercus_margarettae
Fagaceae_Quercus_lobata
Fagaceae_Quercus_austrina
Fagaceae_Quercus_chapmanii
Fagaceae_Quercus_turbinella
Fagaceae_Quercus_dumosa
Fagaceae_Quercus_cornelius-mulleri
Fagaceae_Quercus_hartwissiana
Fagaceae_Quercus_engelmannii
Fagaceae_Quercus_garryana
Fagaceae_Quercus_laeta
Fagaceae_Quercus_arizonica
Fagaceae_Quercus_gambelii
Fagaceae_Quercus_minima
Fagaceae_Quercus_fusiformis
Fagaceae_Quercus_virginiana

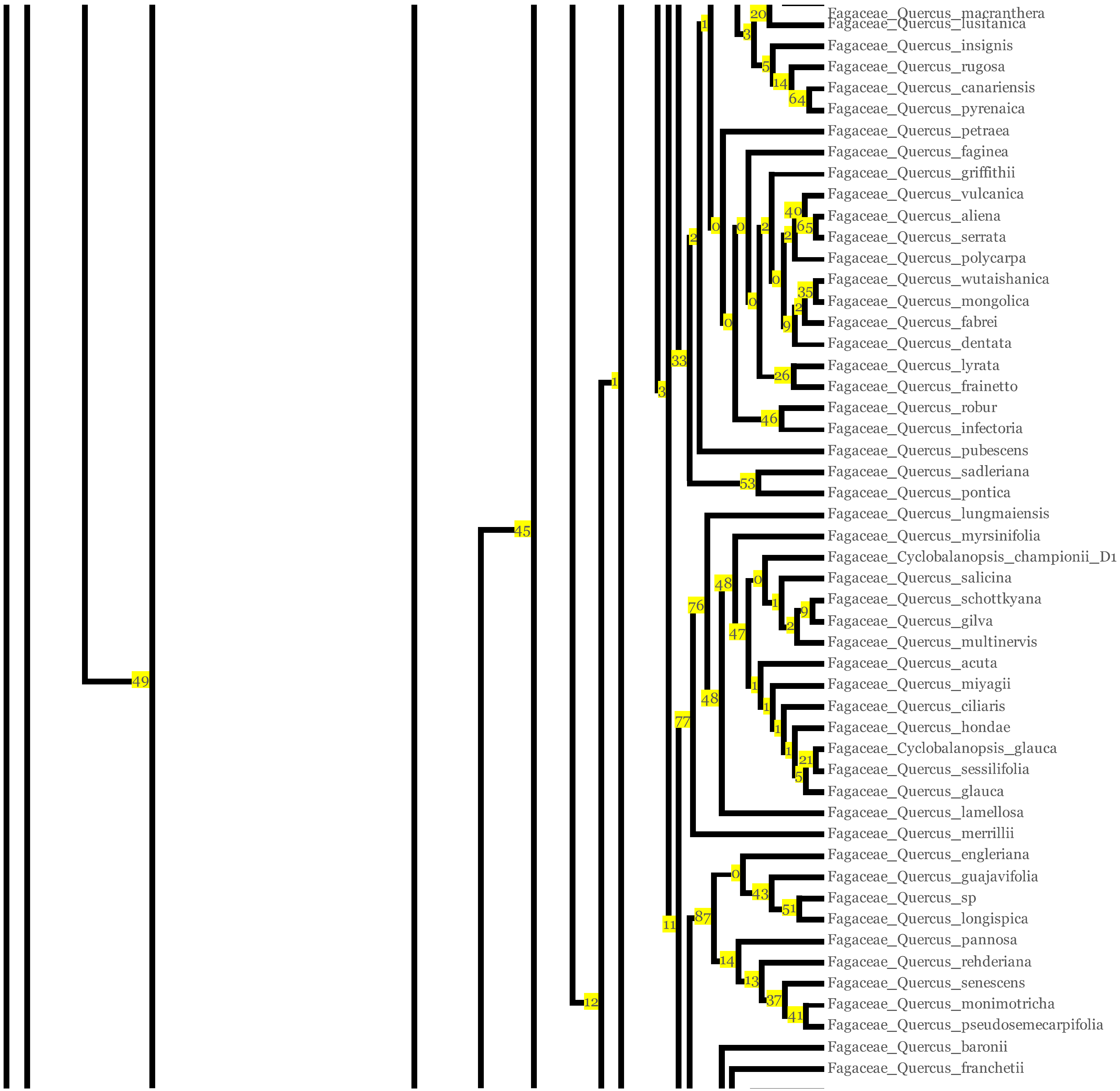
Fagaceae_Quercus_macranthera
Fagaceae_Quercus_lusitanica
Fagaceae_Quercus_insignis
Fagaceae_Quercus_rugosa
Fagaceae_Quercus_canariensis
Fagaceae_Quercus_pyrenaica
Fagaceae_Quercus_petraea
Fagaceae_Quercus_faginea
Fagaceae_Quercus_griffithii
Fagaceae_Quercus_vulcanica
Fagaceae_Quercus_aliena
Fagaceae_Quercus_serrata
Fagaceae_Quercus_polycarpa
Fagaceae_Quercus_wutaishanica
Fagaceae_Quercus_mongolica
Fagaceae_Quercus_fabrei
Fagaceae_Quercus_dentata
Fagaceae_Quercus_lyrata
Fagaceae_Quercus_frainetto
Fagaceae_Quercus_robur
Fagaceae_Quercus_infectoria
Fagaceae_Quercus_pubescens
Fagaceae_Quercus_sadleriana
Fagaceae_Quercus_pontica
Fagaceae_Quercus_lungmaiensis
Fagaceae_Quercus_myrsinifolia
Fagaceae_Cyclobalanopsis_championii_D1
Fagaceae_Quercus_salicina
Fagaceae_Quercus_schottkyana
Fagaceae_Quercus_gilva
Fagaceae_Quercus_multinervis
Fagaceae_Quercus_acuta
Fagaceae_Quercus_miyagii
Fagaceae_Quercus_ciliaris
Fagaceae_Quercus_hondae
Fagaceae_Cyclobalanopsis_glauca
Fagaceae_Quercus_sessilifolia
Fagaceae_Quercus_glauca
Fagaceae_Quercus_lamellosa
Fagaceae_Quercus_merrillii
Fagaceae_Quercus_engleriana
Fagaceae_Quercus_guajavifolia
Fagaceae_Quercus_sp
Fagaceae_Quercus_longispica
Fagaceae_Quercus_pannosa
Fagaceae_Quercus_rehderiana
Fagaceae_Quercus_senescens
Fagaceae_Quercus_monimotricha
Fagaceae_Quercus_pseudosemecarpifolia
Fagaceae_Quercus_baronii
Fagaceae_Quercus_franchetii

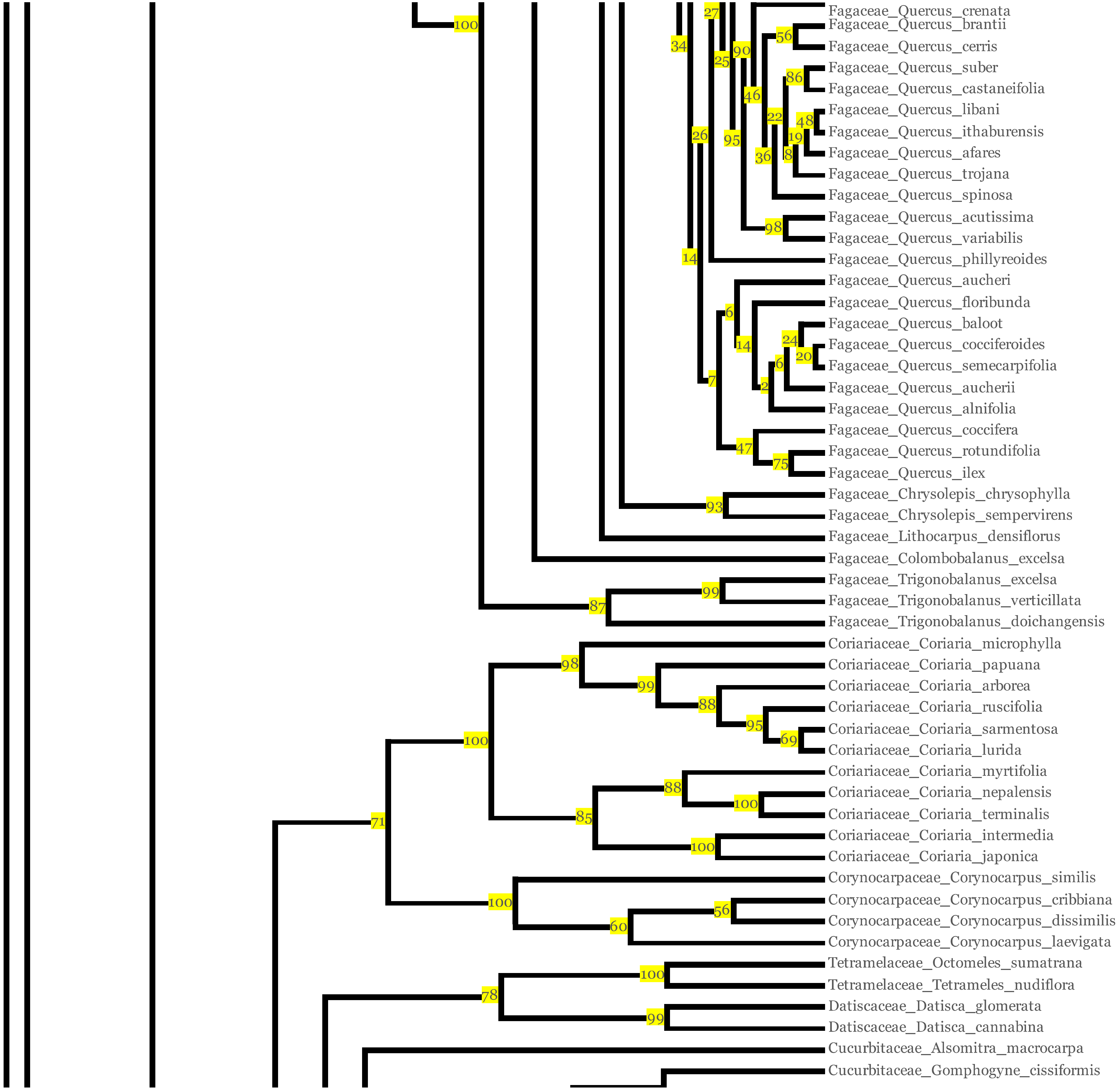
Fagaceae_Quercus_crenata
Fagaceae_Quercus_brantii
Fagaceae_Quercus_cerris
Fagaceae_Quercus_suber
Fagaceae_Quercus_castaneifolia
Fagaceae_Quercus_libani
Fagaceae_Quercus_ithaburensis
Fagaceae_Quercus_afares
Fagaceae_Quercus_trojana
Fagaceae_Quercus_spinosa
Fagaceae_Quercus_acutissima
Fagaceae_Quercus_variabilis
Fagaceae_Quercus_phillyreoides
Fagaceae_Quercus_aucheri
Fagaceae_Quercus_floribunda
Fagaceae_Quercus_baloot
Fagaceae_Quercus_cocciferoides
Fagaceae_Quercus_semecarpifolia
Fagaceae_Quercus_aucherii
Fagaceae_Quercus_alnifolia
Fagaceae_Quercus_coccifera
Fagaceae_Quercus_rotundifolia
Fagaceae_Quercus_ilex
Fagaceae_Chrysolepis_chrysophylla
Fagaceae_Chrysolepis_sempervirens
Fagaceae_Lithocarpus_densiflorus
Fagaceae_Colombobalanus_excelsa
Fagaceae_Trigonobalanus_excelsa
Fagaceae_Trigonobalanus_verticillata
Fagaceae_Trigonobalanus_doichangensis
Coriariaceae_Coriaria_microphylla
Coriariaceae_Coriaria_papuana
Coriariaceae_Coriaria_arborea
Coriariaceae_Coriaria_ruscifolia
Coriariaceae_Coriaria_sarmentosa
Coriariaceae_Coriaria_lurida
Coriariaceae_Coriaria_myrtifolia
Coriariaceae_Coriaria_nepalensis
Coriariaceae_Coriaria_terminalis
Coriariaceae_Coriaria_intermedia
Coriariaceae_Coriaria_japonica
Corynocarpaceae_Corynocarpus_similis
Corynocarpaceae_Corynocarpus_cribbiana
Corynocarpaceae_Corynocarpus_dissimilis
Corynocarpaceae_Corynocarpus_laevigata
Tetramelaceae_Octomeles_sumatrana
Tetramelaceae_Tetrameles_nudiflora
Datiscaceae_Datisca_glomerata
Datiscaceae_Datisca_cannabina
Cucurbitaceae_Alsomitra_macrocarpa
Cucurbitaceae_Gomphogyne_cissiformis

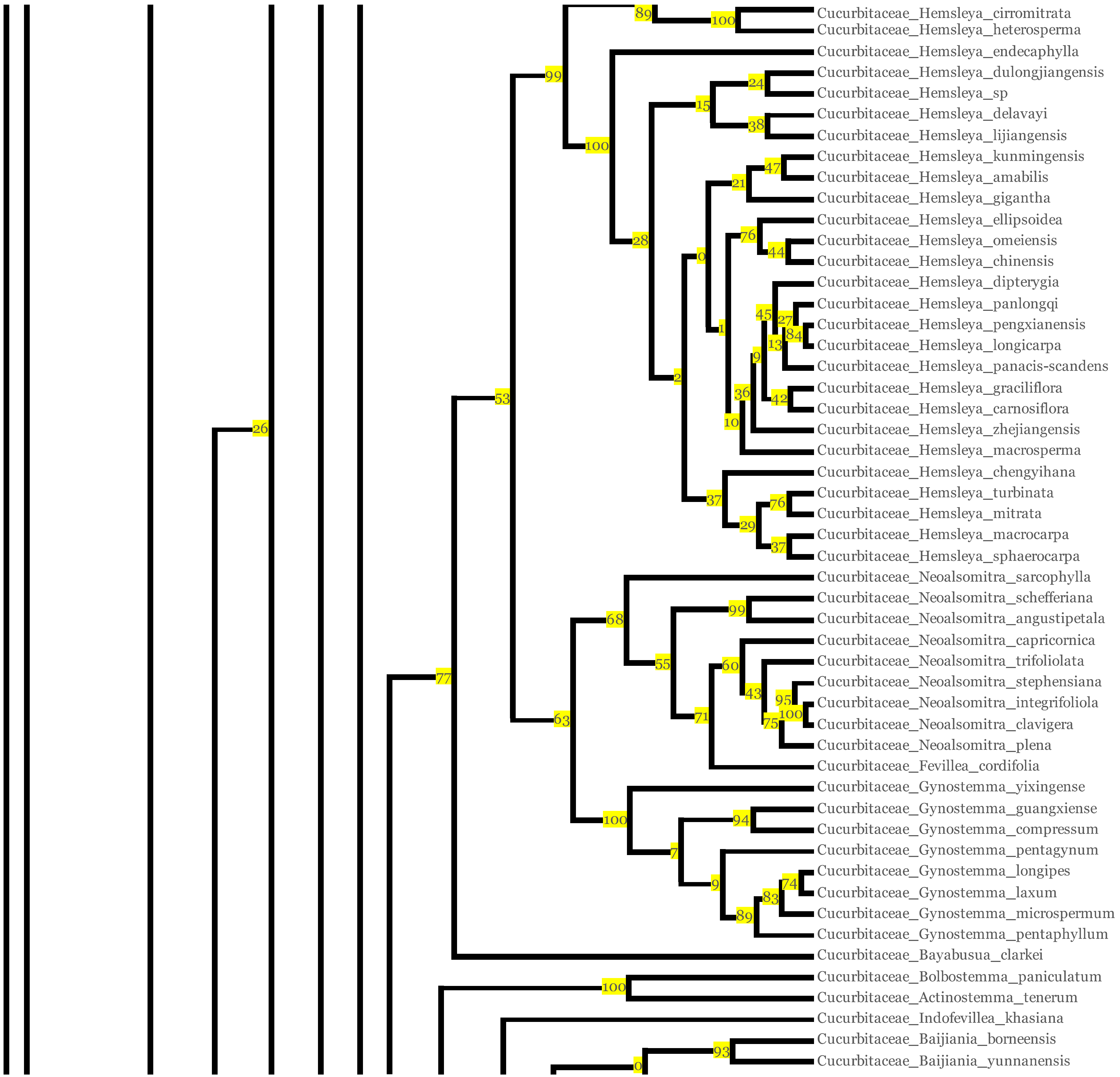
Cucurbitaceae_Hemsleya_cirromitrata
Cucurbitaceae_Hemsleya_heterosperma
Cucurbitaceae_Hemsleya_endecaphylla
Cucurbitaceae_Hemsleya_dulongjiangensis
Cucurbitaceae_Hemsleya_sp
Cucurbitaceae_Hemsleya_delavayi
Cucurbitaceae_Hemsleya_lijiangensis
Cucurbitaceae_Hemsleya_kunmingensis
Cucurbitaceae_Hemsleya_amabilis
Cucurbitaceae_Hemsleya_gigantha
Cucurbitaceae_Hemsleya_ellipsoidea
Cucurbitaceae_Hemsleya_omeiensis
Cucurbitaceae_Hemsleya_chinensis
Cucurbitaceae_Hemsleya_dipterygia
Cucurbitaceae_Hemsleya_panlongqi
Cucurbitaceae_Hemsleya_pengxianensis
Cucurbitaceae_Hemsleya_longicarpa
Cucurbitaceae_Hemsleya_panacis-scandens
Cucurbitaceae_Hemsleya_graciliflora
Cucurbitaceae_Hemsleya_carnosiflora
Cucurbitaceae_Hemsleya_zhejiangensis
Cucurbitaceae_Hemsleya_macrosperma
Cucurbitaceae_Hemsleya_chengyihana
Cucurbitaceae_Hemsleya_turbinata
Cucurbitaceae_Hemsleya_mitrata
Cucurbitaceae_Hemsleya_macrocarpa
Cucurbitaceae_Hemsleya_sphaerocarpa
Cucurbitaceae_Neoalsomitra_sarcophylla
Cucurbitaceae_Neoalsomitra_schefferiana
Cucurbitaceae_Neoalsomitra_angustipetala
Cucurbitaceae_Neoalsomitra_capricornica
Cucurbitaceae_Neoalsomitra_trifoliolata
Cucurbitaceae_Neoalsomitra_stephensiana
Cucurbitaceae_Neoalsomitra_integrifoliola
Cucurbitaceae_Neoalsomitra_clavigera
Cucurbitaceae_Neoalsomitra_plena
Cucurbitaceae_Fevillea_cordifolia
Cucurbitaceae_Gynostemma_yixingense
Cucurbitaceae_Gynostemma_guangxiense
Cucurbitaceae_Gynostemma_compressum
Cucurbitaceae_Gynostemma_pentagynum
Cucurbitaceae_Gynostemma_longipes
Cucurbitaceae_Gynostemma_laxum
Cucurbitaceae_Gynostemma_microspermum
Cucurbitaceae_Gynostemma_pentaphyllum
Cucurbitaceae_Bayabusua_clarkei
Cucurbitaceae_Bolbostemma_paniculatum
Cucurbitaceae_Actinostemma_tenerum
Cucurbitaceae_Indofevillea_khasiana
Cucurbitaceae_Baijiania_borneensis
Cucurbitaceae_Baijiania_yunnanensis

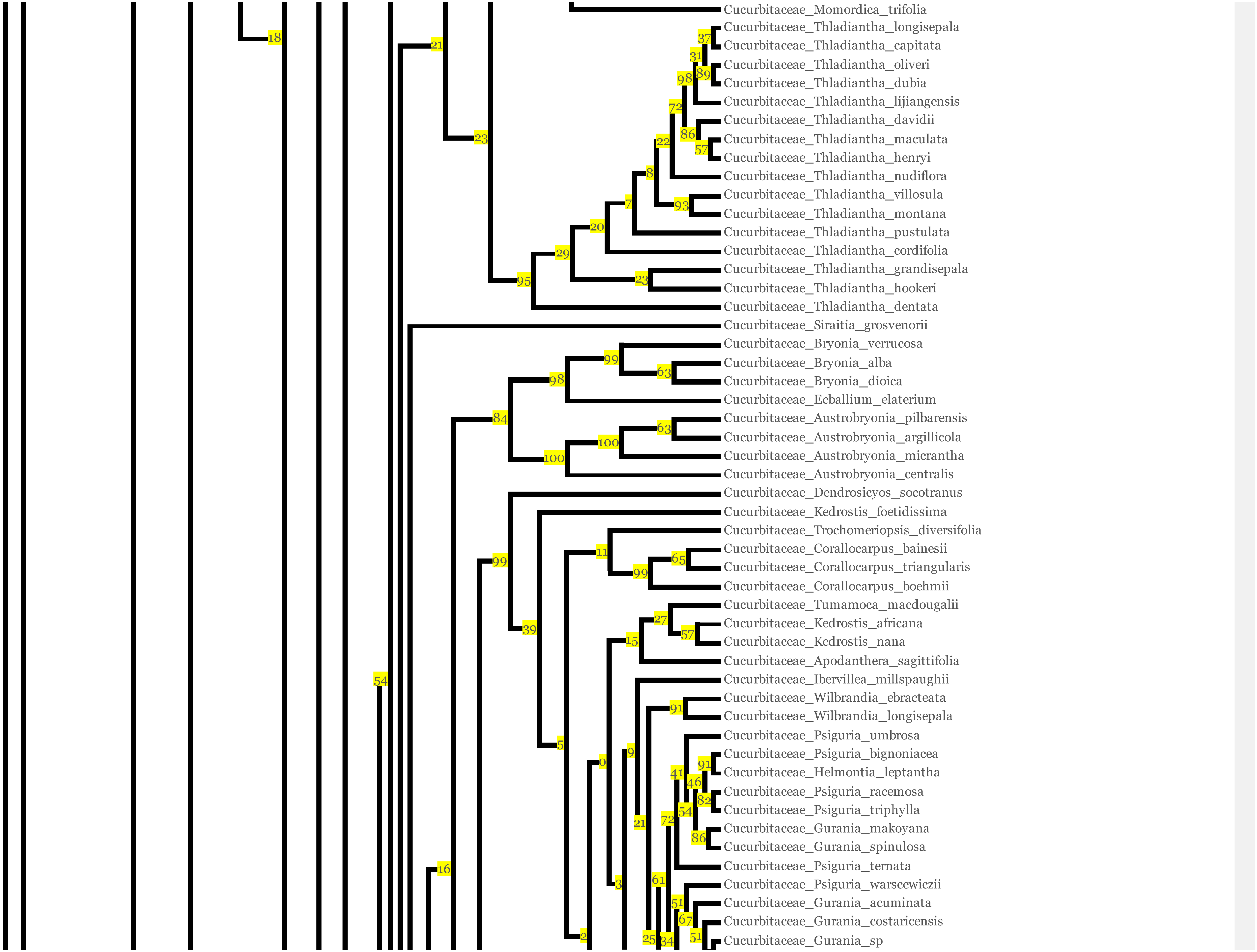
Cucurbitaceae_Momordica_trifolia
Cucurbitaceae_Thladiantha_longisepala
Cucurbitaceae_Thladiantha_capitata
Cucurbitaceae_Thladiantha_oliveri
Cucurbitaceae_Thladiantha_dubia
Cucurbitaceae_Thladiantha_lijiangensis
Cucurbitaceae_Thladiantha_davidii
Cucurbitaceae_Thladiantha_maculata
Cucurbitaceae_Thladiantha_henryi
Cucurbitaceae_Thladiantha_nudiflora
Cucurbitaceae_Thladiantha_villosula
Cucurbitaceae_Thladiantha_montana
Cucurbitaceae_Thladiantha_pustulata
Cucurbitaceae_Thladiantha_cordifolia
Cucurbitaceae_Thladiantha_grandisepala
Cucurbitaceae_Thladiantha_hookeri
Cucurbitaceae_Thladiantha_dentata
Cucurbitaceae_Siraitia_grosvenorii
Cucurbitaceae_Bryonia_verrucosa
Cucurbitaceae_Bryonia_alba
Cucurbitaceae_Bryonia_dioica
Cucurbitaceae_Ecballium_elaterium
Cucurbitaceae_Austrobryonia_pilbarensis
Cucurbitaceae_Austrobryonia_argillicola
Cucurbitaceae_Austrobryonia_micrantha
Cucurbitaceae_Austrobryonia_centralis
Cucurbitaceae_Dendrosicyos_socotranus
Cucurbitaceae_Kedrostis_foetidissima
Cucurbitaceae_Trochomeriopsis_diversifolia
Cucurbitaceae_Corallocarpus_bainesii
Cucurbitaceae_Corallocarpus_triangularis
Cucurbitaceae_Corallocarpus_boehmii
Cucurbitaceae_Tumamoca_macdougalii
Cucurbitaceae_Kedrostis_africana
Cucurbitaceae_Kedrostis_nana
Cucurbitaceae_Apodanthera_sagittifolia
Cucurbitaceae_Ibervillea_millspaughii
Cucurbitaceae_Wilbrandia_ebracteata
Cucurbitaceae_Wilbrandia_longisepala
Cucurbitaceae_Psiguria_umbrosa
Cucurbitaceae_Psiguria_bignoniacea
Cucurbitaceae_Helmontia_leptantha
Cucurbitaceae_Psiguria_racemosa
Cucurbitaceae_Psiguria_triphylla
Cucurbitaceae_Gurania_makoyana
Cucurbitaceae_Gurania_spinulosa
Cucurbitaceae_Psiguria_ternata
Cucurbitaceae_Psiguria_warscewiczii
Cucurbitaceae_Gurania_acuminata
Cucurbitaceae_Gurania_costaricensis
Cucurbitaceae_Gurania_sp

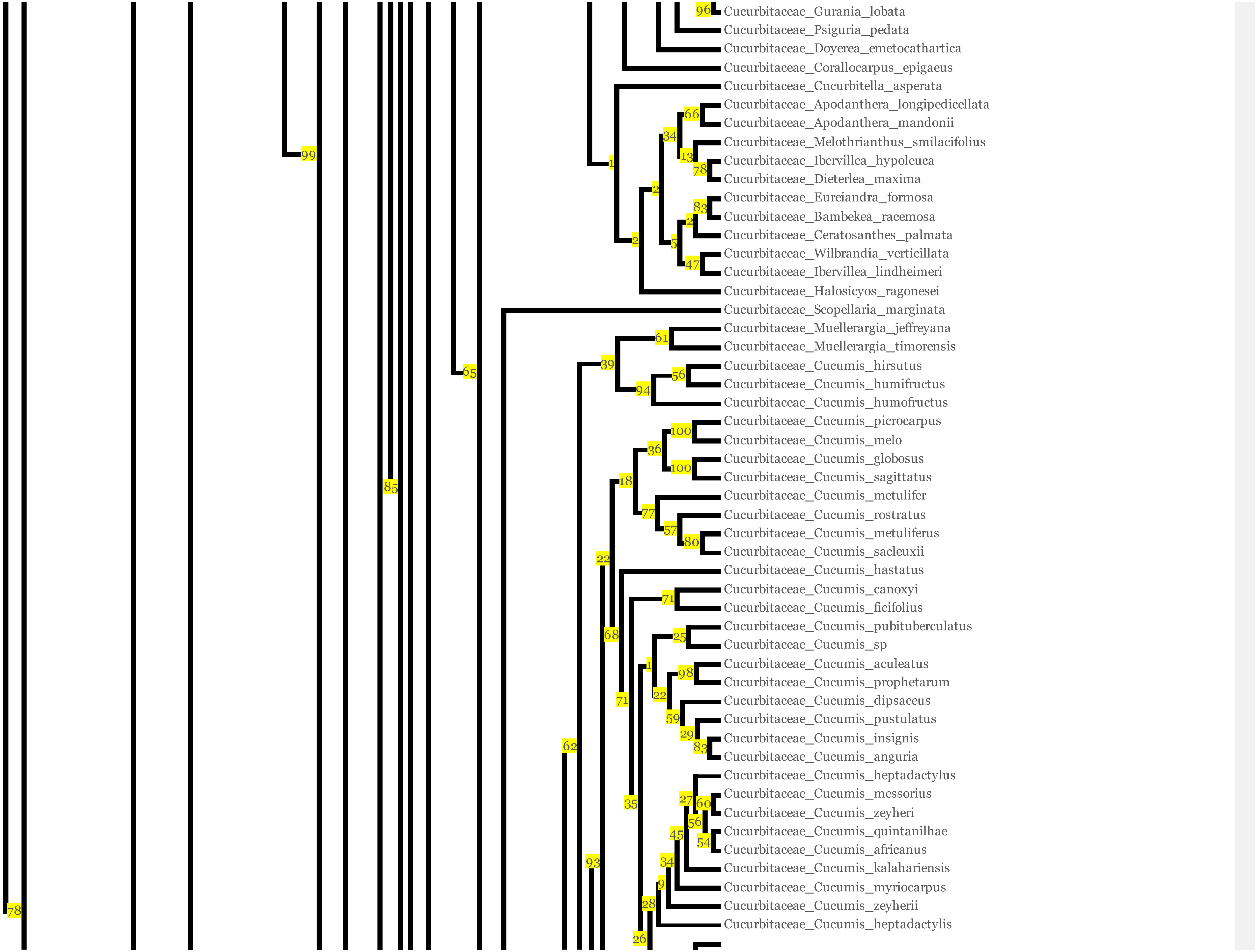
Cucurbitaceae_Gurania_lobata
Cucurbitaceae_Psiguria_pedata
Cucurbitaceae_Doyerea_emetocathartica
Cucurbitaceae_Corallocarpus_epigaeus
Cucurbitaceae_Cucurbitella_asperata
Cucurbitaceae_Apodanthera_longipedicellata
Cucurbitaceae_Apodanthera_mandonii
Cucurbitaceae_Melothrianthus_smilacifolius
Cucurbitaceae_Ibervillea_hypoleuca
Cucurbitaceae_Dieterlea_maxima
Cucurbitaceae_Eureiandra_formosa
Cucurbitaceae_Bambekea_racemosa
Cucurbitaceae_Ceratosanthes_palmata
Cucurbitaceae_Wilbrandia_verticillata
Cucurbitaceae_Ibervillea_lindheimeri
Cucurbitaceae_Halosicyos_ragonesei
Cucurbitaceae_Scopellaria_marginata
Cucurbitaceae_Muellerargia_jeffreyana
Cucurbitaceae_Muellerargia_timorensis
Cucurbitaceae_Cucumis_hirsutus
Cucurbitaceae_Cucumis_humifructus
Cucurbitaceae_Cucumis_humofructus
Cucurbitaceae_Cucumis_picrocarpus
Cucurbitaceae_Cucumis_melo
Cucurbitaceae_Cucumis_globosus
Cucurbitaceae_Cucumis_sagittatus
Cucurbitaceae_Cucumis_metulifer
Cucurbitaceae_Cucumis_rostratus
Cucurbitaceae_Cucumis_metuliferus
Cucurbitaceae_Cucumis_sacleuxii
Cucurbitaceae_Cucumis_hastatus
Cucurbitaceae_Cucumis_canoxyi
Cucurbitaceae_Cucumis_ficifolius
Cucurbitaceae_Cucumis_pubituberculatus
Cucurbitaceae_Cucumis_sp
Cucurbitaceae_Cucumis_aculeatus
Cucurbitaceae_Cucumis_prophetarum
Cucurbitaceae_Cucumis_dipsaceus
Cucurbitaceae_Cucumis_pustulatus
Cucurbitaceae_Cucumis_insignis
Cucurbitaceae_Cucumis_anguria
Cucurbitaceae_Cucumis_heptadactylus
Cucurbitaceae_Cucumis_messorius
Cucurbitaceae_Cucumis_zeyheri
Cucurbitaceae_Cucumis_quintanilhae
Cucurbitaceae_Cucumis_africanus
Cucurbitaceae_Cucumis_kalahariensis
Cucurbitaceae_Cucumis_myriocarpus
Cucurbitaceae_Cucumis_zeyherii
Cucurbitaceae_Cucumis_heptadactylis

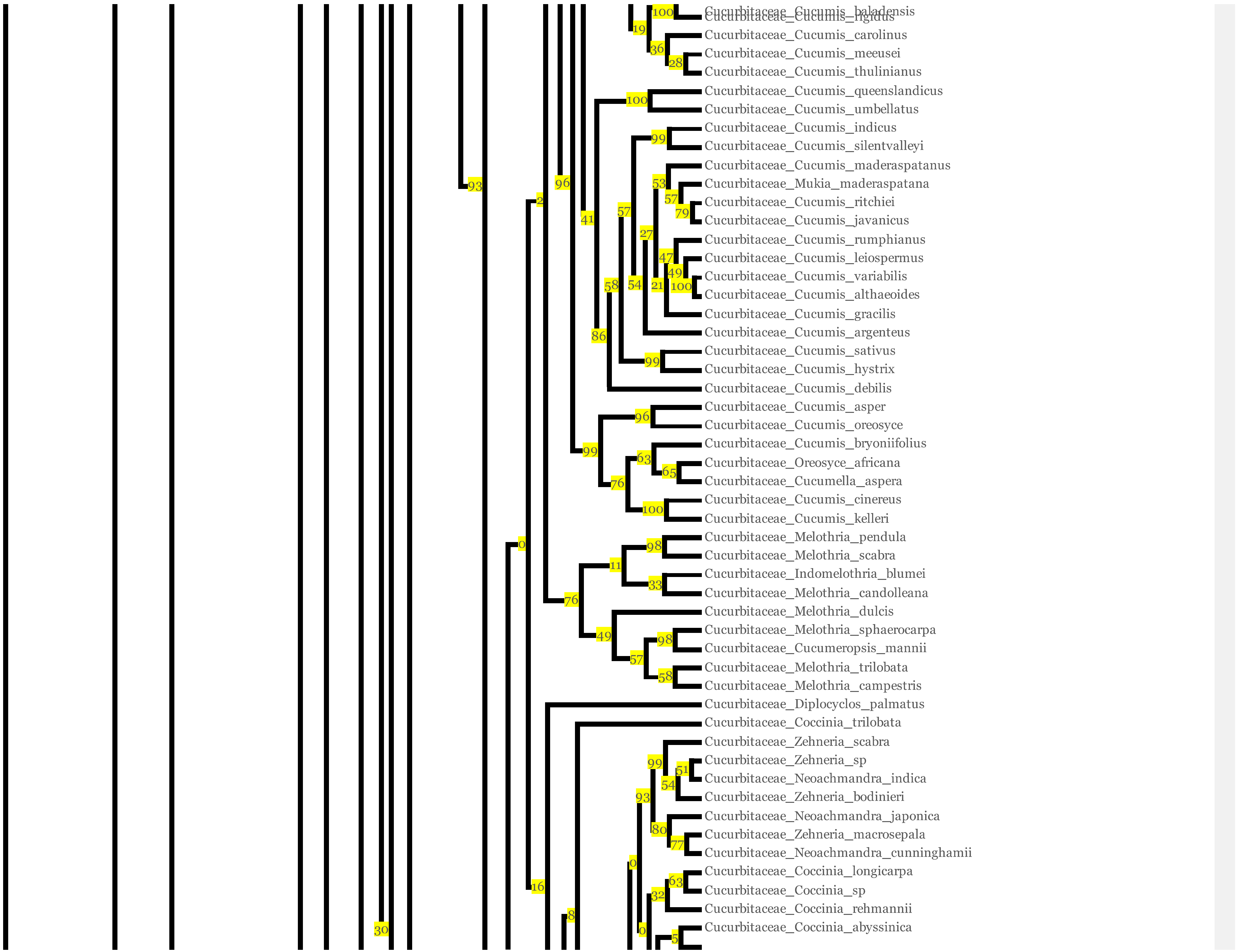

Cucurbitaceae_Cucumis_baladensis
Cucurbitaceae_Cucumis_rigidus
Cucurbitaceae_Cucumis_carolinus
Cucurbitaceae_Cucumis_meeusei
Cucurbitaceae_Cucumis_thulinianus
Cucurbitaceae_Cucumis_queenslandicus
Cucurbitaceae_Cucumis_umbellatus
Cucurbitaceae_Cucumis_indicus
Cucurbitaceae_Cucumis_silentvalleyi
Cucurbitaceae_Cucumis_maderaspatanus
Cucurbitaceae_Mukia_maderaspatana
Cucurbitaceae_Cucumis_ritchiei
Cucurbitaceae_Cucumis_javanicus
Cucurbitaceae_Cucumis_rumphianus
Cucurbitaceae_Cucumis_leiospermus
Cucurbitaceae_Cucumis_variabilis
Cucurbitaceae_Cucumis_althaeoides
Cucurbitaceae_Cucumis_gracilis
Cucurbitaceae_Cucumis_argenteus
Cucurbitaceae_Cucumis_sativus
Cucurbitaceae_Cucumis_hystrix
Cucurbitaceae_Cucumis_debilis
Cucurbitaceae_Cucumis_asper
Cucurbitaceae_Cucumis_oreosyce
Cucurbitaceae_Cucumis_bryoniifolius
Cucurbitaceae_Oreosyce_africana
Cucurbitaceae_Cucumella_aspera
Cucurbitaceae_Cucumis_cinereus
Cucurbitaceae_Cucumis_kelleri
Cucurbitaceae_Melothria_pendula
Cucurbitaceae_Melothria_scabra
Cucurbitaceae_Indomelothria_blumei
Cucurbitaceae_Melothria_candolleana
Cucurbitaceae_Melothria_dulcis
Cucurbitaceae_Melothria_sphaerocarpa
Cucurbitaceae_Cucumeropsis_mannii
Cucurbitaceae_Melothria_trilobata
Cucurbitaceae_Melothria_campestris
Cucurbitaceae_Diplocyclos_palmatus
Cucurbitaceae_Coccinia_trilobata
Cucurbitaceae_Zehneria_scabra
Cucurbitaceae_Zehneria_sp
Cucurbitaceae_Neoachmandra_indica
Cucurbitaceae_Zehneria_bodinieri
Cucurbitaceae_Neoachmandra_japonica
Cucurbitaceae_Zehneria_macrosepala
Cucurbitaceae_Neoachmandra_cunninghamii
Cucurbitaceae_Coccinia_longicarpa
Cucurbitaceae_Coccinia_sp
Cucurbitaceae_Coccinia_rehmannii
Cucurbitaceae_Coccinia_abyssinica

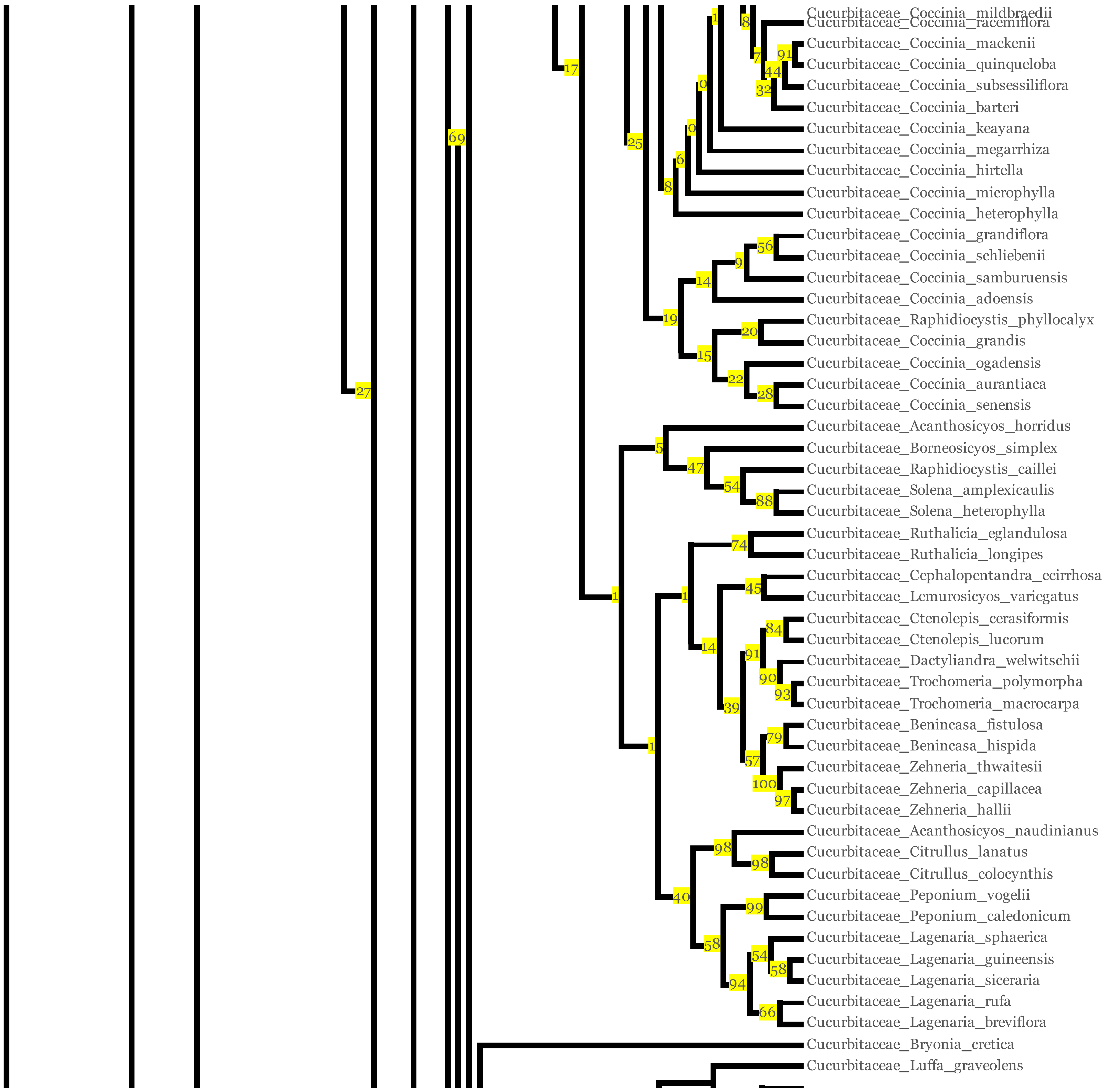
Cucurbitaceae_Coccinia_mildbraedii
Cucurbitaceae_Coccinia_racemiflora
Cucurbitaceae_Coccinia_mackenii
Cucurbitaceae_Coccinia_quinqueloba
Cucurbitaceae_Coccinia_subsessiliflora
Cucurbitaceae_Coccinia_barteri
Cucurbitaceae_Coccinia_keayana
Cucurbitaceae_Coccinia_megarrhiza
Cucurbitaceae_Coccinia_hirtella
Cucurbitaceae_Coccinia_microphylla
Cucurbitaceae_Coccinia_heterophylla
Cucurbitaceae_Coccinia_grandiflora
Cucurbitaceae_Coccinia_schliebenii
Cucurbitaceae_Coccinia_samburuensis
Cucurbitaceae_Coccinia_adoensis
Cucurbitaceae_Raphidiocystis_phyllocalyx
Cucurbitaceae_Coccinia_grandis
Cucurbitaceae_Coccinia_ogadensis
Cucurbitaceae_Coccinia_aurantiaca
Cucurbitaceae_Coccinia_senensis
Cucurbitaceae_Acanthosicyos_horridus
Cucurbitaceae_Borneosicyos_simplex
Cucurbitaceae_Raphidiocystis_caillei
Cucurbitaceae_Solena_amplexicaulis
Cucurbitaceae_Solena_heterophylla
Cucurbitaceae_Ruthalicia_eglandulosa
Cucurbitaceae_Ruthalicia_longipes
Cucurbitaceae_Cephalopentandra_ecirrhosa
Cucurbitaceae_Lemurosicyos_variegatus
Cucurbitaceae_Ctenolepis_cerasiformis
Cucurbitaceae_Ctenolepis_lucorum
Cucurbitaceae_Dactyliandra_welwitschii
Cucurbitaceae_Trochomeria_polymorpha
Cucurbitaceae_Trochomeria_macrocarpa
Cucurbitaceae_Benincasa_fistulosa
Cucurbitaceae_Benincasa_hispida
Cucurbitaceae_Zehneria_thwaitesii
Cucurbitaceae_Zehneria_capillacea
Cucurbitaceae_Zehneria_hallii
Cucurbitaceae_Acanthosicyos_naudinianus
Cucurbitaceae_Citrullus_lanatus
Cucurbitaceae_Citrullus_colocynthis
Cucurbitaceae_Peponium_vogelii
Cucurbitaceae_Peponium_caledonicum
Cucurbitaceae_Lagenaria_sphaerica
Cucurbitaceae_Lagenaria_guineensis
Cucurbitaceae_Lagenaria_siceraria
Cucurbitaceae_Lagenaria_rufa
Cucurbitaceae_Lagenaria_breviflora
Cucurbitaceae_Bryonia_cretica
Cucurbitaceae_Luffa_graveolens

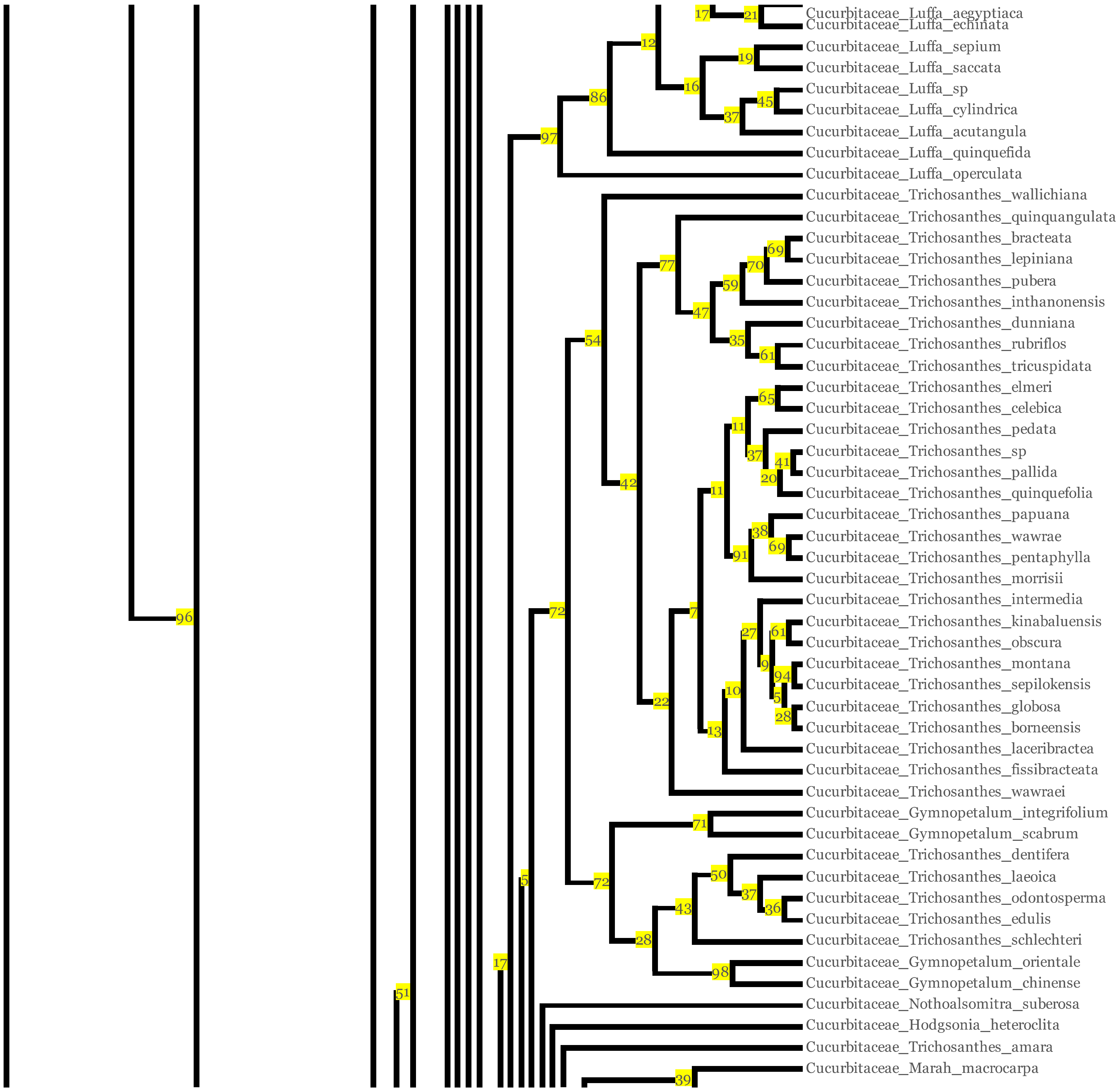
Cucurbitaceae_Luffa_aegyptiaca
Cucurbitaceae_Luffa_echinata
Cucurbitaceae_Luffa_sepium
Cucurbitaceae_Luffa_saccata
Cucurbitaceae_Luffa_sp
Cucurbitaceae_Luffa_cylindrica
Cucurbitaceae_Luffa_acutangula
Cucurbitaceae_Luffa_quinquefida
Cucurbitaceae_Luffa_operculata
Cucurbitaceae_Trichosanthes_wallichiana
Cucurbitaceae_Trichosanthes_quinquangulata
Cucurbitaceae_Trichosanthes_bracteata
Cucurbitaceae_Trichosanthes_lepiniana
Cucurbitaceae_Trichosanthes_pubera
Cucurbitaceae_Trichosanthes_inthanonensis
Cucurbitaceae_Trichosanthes_dunniana
Cucurbitaceae_Trichosanthes_rubriflos
Cucurbitaceae_Trichosanthes_tricuspidata
Cucurbitaceae_Trichosanthes_elmeri
Cucurbitaceae_Trichosanthes_celebica
Cucurbitaceae_Trichosanthes_pedata
Cucurbitaceae_Trichosanthes_sp
Cucurbitaceae_Trichosanthes_pallida
Cucurbitaceae_Trichosanthes_quinquefolia
Cucurbitaceae_Trichosanthes_papuana
Cucurbitaceae_Trichosanthes_wawrae
Cucurbitaceae_Trichosanthes_pentaphylla
Cucurbitaceae_Trichosanthes_morrisii
Cucurbitaceae_Trichosanthes_intermedia
Cucurbitaceae_Trichosanthes_kinabaluensis
Cucurbitaceae_Trichosanthes_obscura
Cucurbitaceae_Trichosanthes_montana
Cucurbitaceae_Trichosanthes_sepilokensis
Cucurbitaceae_Trichosanthes_globosa
Cucurbitaceae_Trichosanthes_borneensis
Cucurbitaceae_Trichosanthes_laceribractea
Cucurbitaceae_Trichosanthes_fissibracteata
Cucurbitaceae_Trichosanthes_wawraei
Cucurbitaceae_Gymnopetalum_integrifolium
Cucurbitaceae_Gymnopetalum_scabrum
Cucurbitaceae_Trichosanthes_dentifera
Cucurbitaceae_Trichosanthes_laeoica
Cucurbitaceae_Trichosanthes_odontosperma
Cucurbitaceae_Trichosanthes_edulis
Cucurbitaceae_Trichosanthes_schlechteri
Cucurbitaceae_Gymnopetalum_orientale
Cucurbitaceae_Gymnopetalum_chinense
Cucurbitaceae_Nothoalsomitra_suberosa
Cucurbitaceae_Hodgsonia_heteroclita
Cucurbitaceae_Trichosanthes_amara
Cucurbitaceae_Marah_macrocarpa

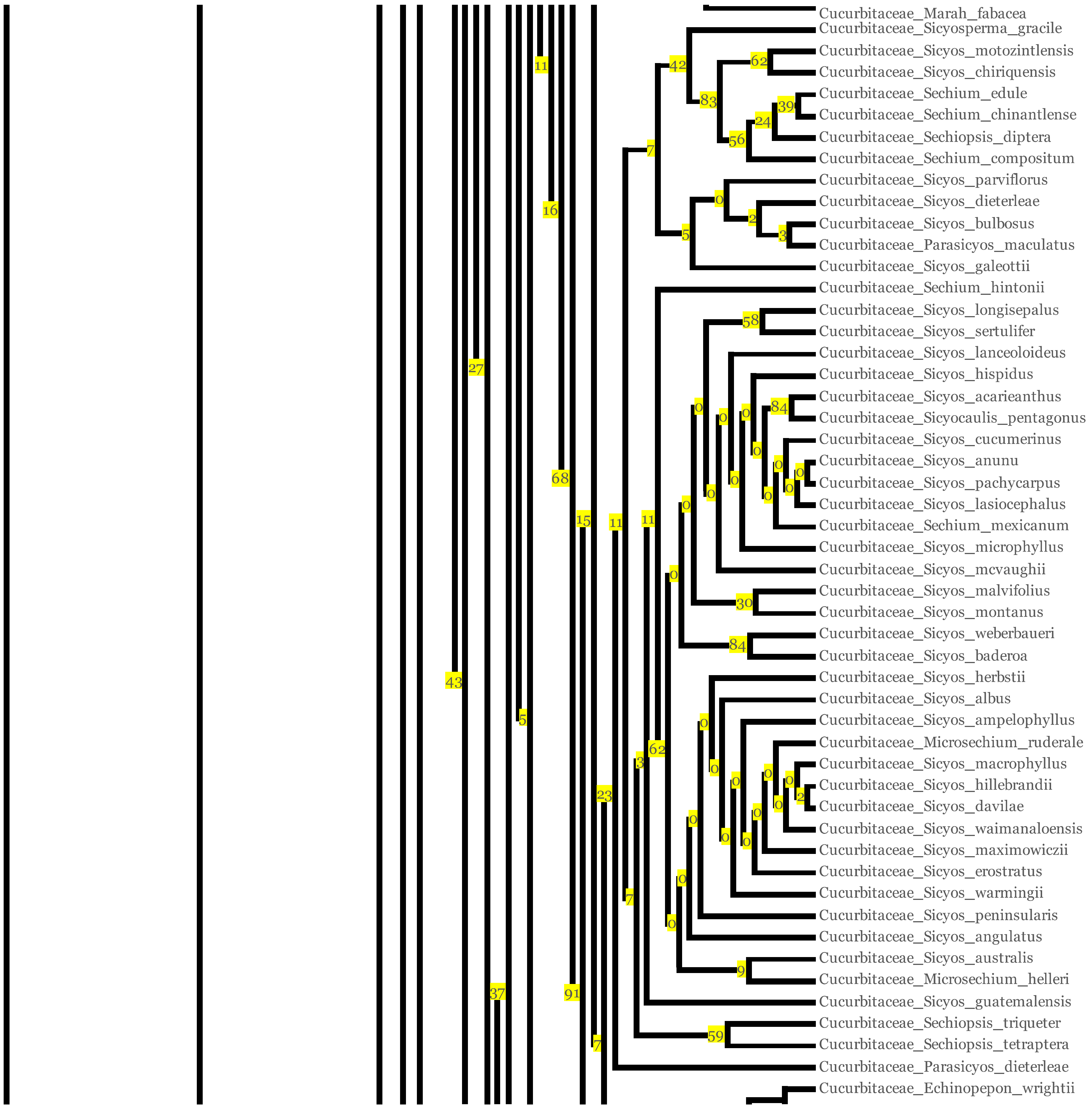
Cucurbitaceae_Marah_fabacea
Cucurbitaceae_Sicyosperma_gracile
Cucurbitaceae_Sicyos_motozintlensis
Cucurbitaceae_Sicyos_chiriquensis
Cucurbitaceae_Sechium_edule
Cucurbitaceae_Sechium_chinantlense
Cucurbitaceae_Sechiopsis_diptera
Cucurbitaceae_Sechium_compositum
Cucurbitaceae_Sicyos_parviflorus
Cucurbitaceae_Sicyos_dieterleae
Cucurbitaceae_Sicyos_bulbosus
Cucurbitaceae_Parasicyos_maculatus
Cucurbitaceae_Sicyos_galeottii
Cucurbitaceae_Sechium_hintonii
Cucurbitaceae_Sicyos_longisepalus
Cucurbitaceae_Sicyos_sertulifer
Cucurbitaceae_Sicyos_lanceoloideus
Cucurbitaceae_Sicyos_hispidus
Cucurbitaceae_Sicyos_acarieanthus
Cucurbitaceae_Sicyocaulis_pentagonus
Cucurbitaceae_Sicyos_cucumerinus
Cucurbitaceae_Sicyos_anunu
Cucurbitaceae_Sicyos_pachycarpus
Cucurbitaceae_Sicyos_lasiocephalus
Cucurbitaceae_Sechium_mexicanum
Cucurbitaceae_Sicyos_microphyllus
Cucurbitaceae_Sicyos_mcvaughii
Cucurbitaceae_Sicyos_malvifolius
Cucurbitaceae_Sicyos_montanus
Cucurbitaceae_Sicyos_weberbaueri
Cucurbitaceae_Sicyos_baderoa
Cucurbitaceae_Sicyos_herbstii
Cucurbitaceae_Sicyos_albus
Cucurbitaceae_Sicyos_ampelophyllus
Cucurbitaceae_Microsechium_ruderale
Cucurbitaceae_Sicyos_macrophyllus
Cucurbitaceae_Sicyos_hillebrandii
Cucurbitaceae_Sicyos_davilae
Cucurbitaceae_Sicyos_waimanaloensis
Cucurbitaceae_Sicyos_maximowiczii
Cucurbitaceae_Sicyos_erostratus
Cucurbitaceae_Sicyos_warmingii
Cucurbitaceae_Sicyos_peninsularis
Cucurbitaceae_Sicyos_angulatus
Cucurbitaceae_Sicyos_australis
Cucurbitaceae_Microsechium_helleri
Cucurbitaceae_Sicyos_guatemalensis
Cucurbitaceae_Sechiopsis_triqueter
Cucurbitaceae_Sechiopsis_tetraptera
Cucurbitaceae_Parasicyos_dieterleae
Cucurbitaceae_Echinopepon_wrightii

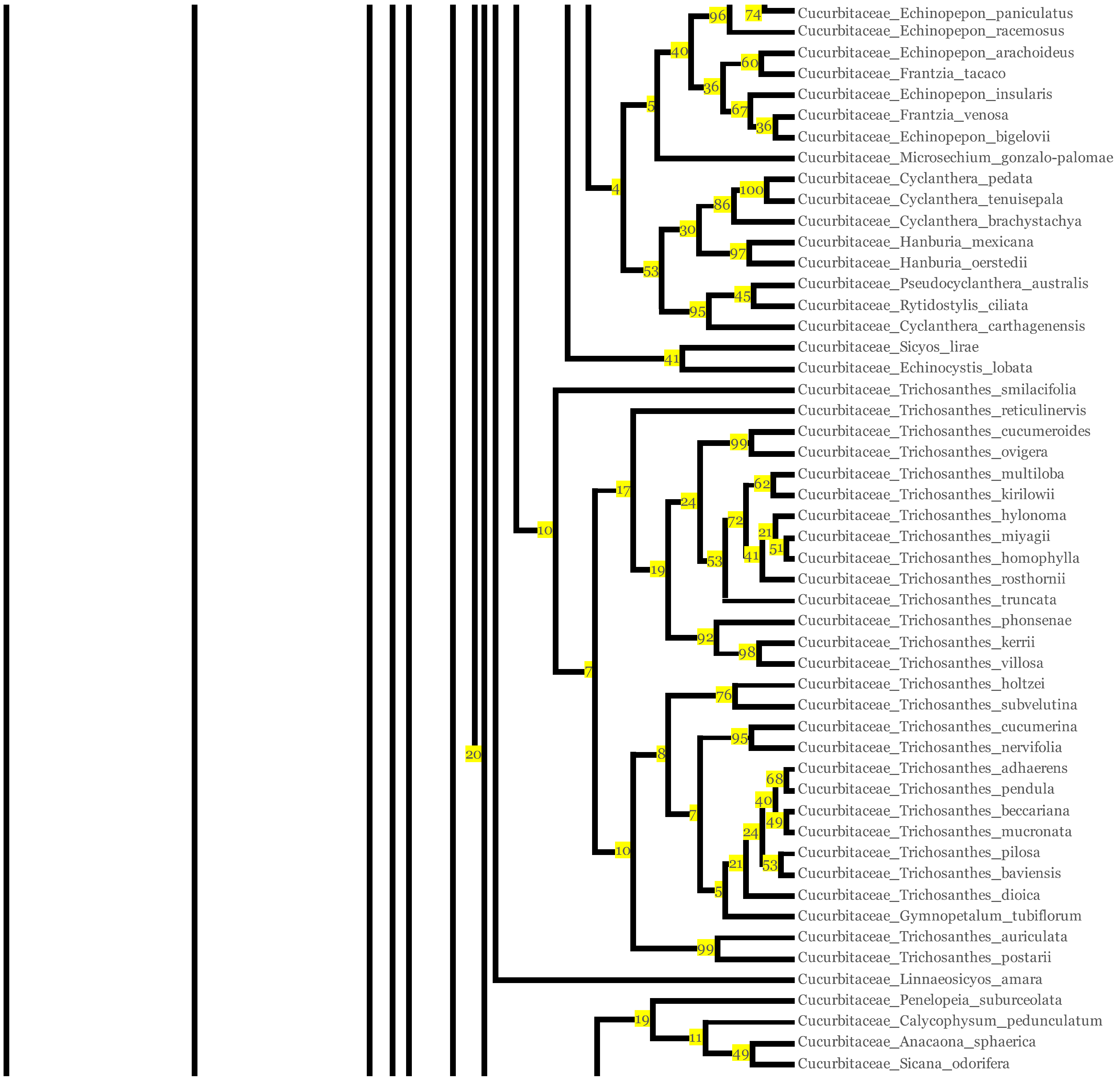
Cucurbitaceae_Echinopepon_paniculatus
Cucurbitaceae_Echinopepon_racemosus
Cucurbitaceae_Echinopepon_arachoideus
Cucurbitaceae_Frantzia_tacaco
Cucurbitaceae_Echinopepon_insularis
Cucurbitaceae_Frantzia_venosa
Cucurbitaceae_Echinopepon_bigelovii
Cucurbitaceae_Microsechium_gonzalo-palomae
Cucurbitaceae_Cyclanthera_pedata
Cucurbitaceae_Cyclanthera_tenuisepala
Cucurbitaceae_Cyclanthera_brachystachya
Cucurbitaceae_Hanburia_mexicana
Cucurbitaceae_Hanburia_oerstedii
Cucurbitaceae_Pseudocyclanthera_australis
Cucurbitaceae_Rytidostylis_ciliata
Cucurbitaceae_Cyclanthera_carthagenensis
Cucurbitaceae_Sicyos_lirae
Cucurbitaceae_Echinocystis_lobata
Cucurbitaceae_Trichosanthes_smilacifolia
Cucurbitaceae_Trichosanthes_reticulinervis
Cucurbitaceae_Trichosanthes_cucumeroides
Cucurbitaceae_Trichosanthes_ovigera
Cucurbitaceae_Trichosanthes_multiloba
Cucurbitaceae_Trichosanthes_kirilowii
Cucurbitaceae_Trichosanthes_hylonoma
Cucurbitaceae_Trichosanthes_miyagii
Cucurbitaceae_Trichosanthes_homophylla
Cucurbitaceae_Trichosanthes_rosthornii
Cucurbitaceae_Trichosanthes_truncata
Cucurbitaceae_Trichosanthes_phonsenae
Cucurbitaceae_Trichosanthes_kerrii
Cucurbitaceae_Trichosanthes_villosa
Cucurbitaceae_Trichosanthes_holtzei
Cucurbitaceae_Trichosanthes_subvelutina
Cucurbitaceae_Trichosanthes_cucumerina
Cucurbitaceae_Trichosanthes_nervifolia
Cucurbitaceae_Trichosanthes_adhaerens
Cucurbitaceae_Trichosanthes_pendula
Cucurbitaceae_Trichosanthes_beccariana
Cucurbitaceae_Trichosanthes_mucronata
Cucurbitaceae_Trichosanthes_pilosa
Cucurbitaceae_Trichosanthes_baviensis
Cucurbitaceae_Trichosanthes_dioica
Cucurbitaceae_Gymnopetalum_tubiflorum
Cucurbitaceae_Trichosanthes_auriculata
Cucurbitaceae_Trichosanthes_postarii
Cucurbitaceae_Linnaeosicyos_amara
Cucurbitaceae_Penelopeia_suburceolata
Cucurbitaceae_Calycophysum_pedunculatum
Cucurbitaceae_Anacaona_sphaerica
Cucurbitaceae_Sicana_odorifera

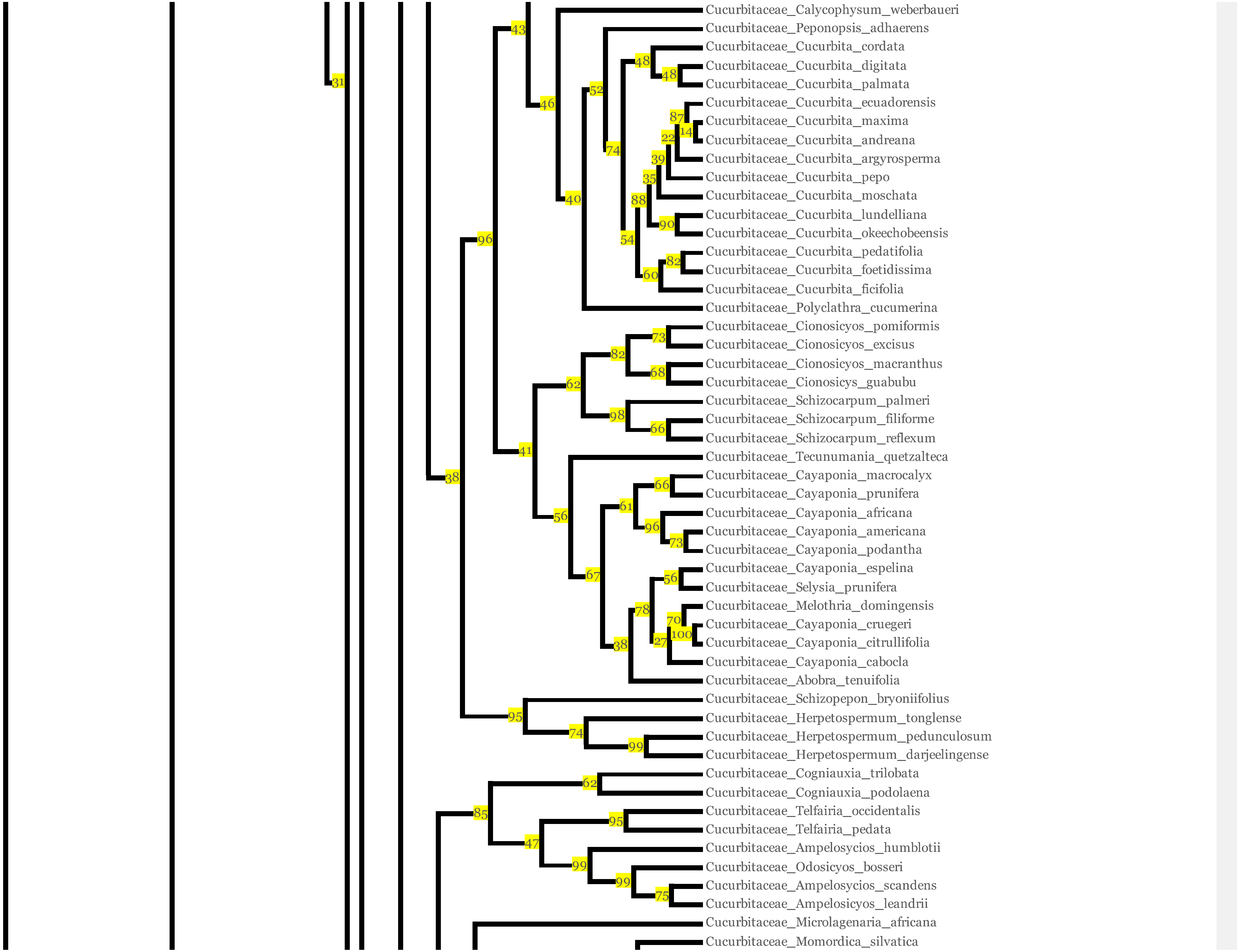

Cucurbitaceae_Calycophysum_weberbaueri
Cucurbitaceae_Peponopsis_adhaerens
Cucurbitaceae_Cucurbita_cordata
Cucurbitaceae_Cucurbita_digitata
Cucurbitaceae_Cucurbita_palmata
Cucurbitaceae_Cucurbita_ecuadorensis
Cucurbitaceae_Cucurbita_maxima
Cucurbitaceae_Cucurbita_andreana
Cucurbitaceae_Cucurbita_argyrosperma
Cucurbitaceae_Cucurbita_pepo
Cucurbitaceae_Cucurbita_moschata
Cucurbitaceae_Cucurbita_lundelliana
Cucurbitaceae_Cucurbita_okeechobeensis
Cucurbitaceae_Cucurbita_pedatifolia
Cucurbitaceae_Cucurbita_foetidissima
Cucurbitaceae_Cucurbita_ficifolia
Cucurbitaceae_Polyclathra_cucumerina
Cucurbitaceae_Cionosicyos_pomiformis
Cucurbitaceae_Cionosicyos_excisus
Cucurbitaceae_Cionosicyos_macranthus
Cucurbitaceae_Cionosicys_guabubu
Cucurbitaceae_Schizocarpum_palmeri
Cucurbitaceae_Schizocarpum_filiforme
Cucurbitaceae_Schizocarpum_reflexum
Cucurbitaceae_Tecunumania_quetzalteca
Cucurbitaceae_Cayaponia_macrocalyx
Cucurbitaceae_Cayaponia_prunifera
Cucurbitaceae_Cayaponia_africana
Cucurbitaceae_Cayaponia_americana
Cucurbitaceae_Cayaponia_podantha
Cucurbitaceae_Cayaponia_espelina
Cucurbitaceae_Selysia_prunifera
Cucurbitaceae_Melothria_domingensis
Cucurbitaceae_Cayaponia_cruegeri
Cucurbitaceae_Cayaponia_citrullifolia
Cucurbitaceae_Cayaponia_cabocla
Cucurbitaceae_Abobra_tenuifolia
Cucurbitaceae_Schizopepon_bryoniifolius
Cucurbitaceae_Herpetospermum_tonglense
Cucurbitaceae_Herpetospermum_pedunculosum
Cucurbitaceae_Herpetospermum_darjeelingense
Cucurbitaceae_Cogniauxia_trilobata
Cucurbitaceae_Cogniauxia_podolaena
Cucurbitaceae_Telfairia_occidentalis
Cucurbitaceae_Telfairia_pedata
Cucurbitaceae_Ampelosycios_humblotii
Cucurbitaceae_Odosicyos_bosseri
Cucurbitaceae_Ampelosycios_scandens
Cucurbitaceae_Ampelosicyos_leandrii
Cucurbitaceae_Microlagenaria_africana
Cucurbitaceae_Momordica_silvatica

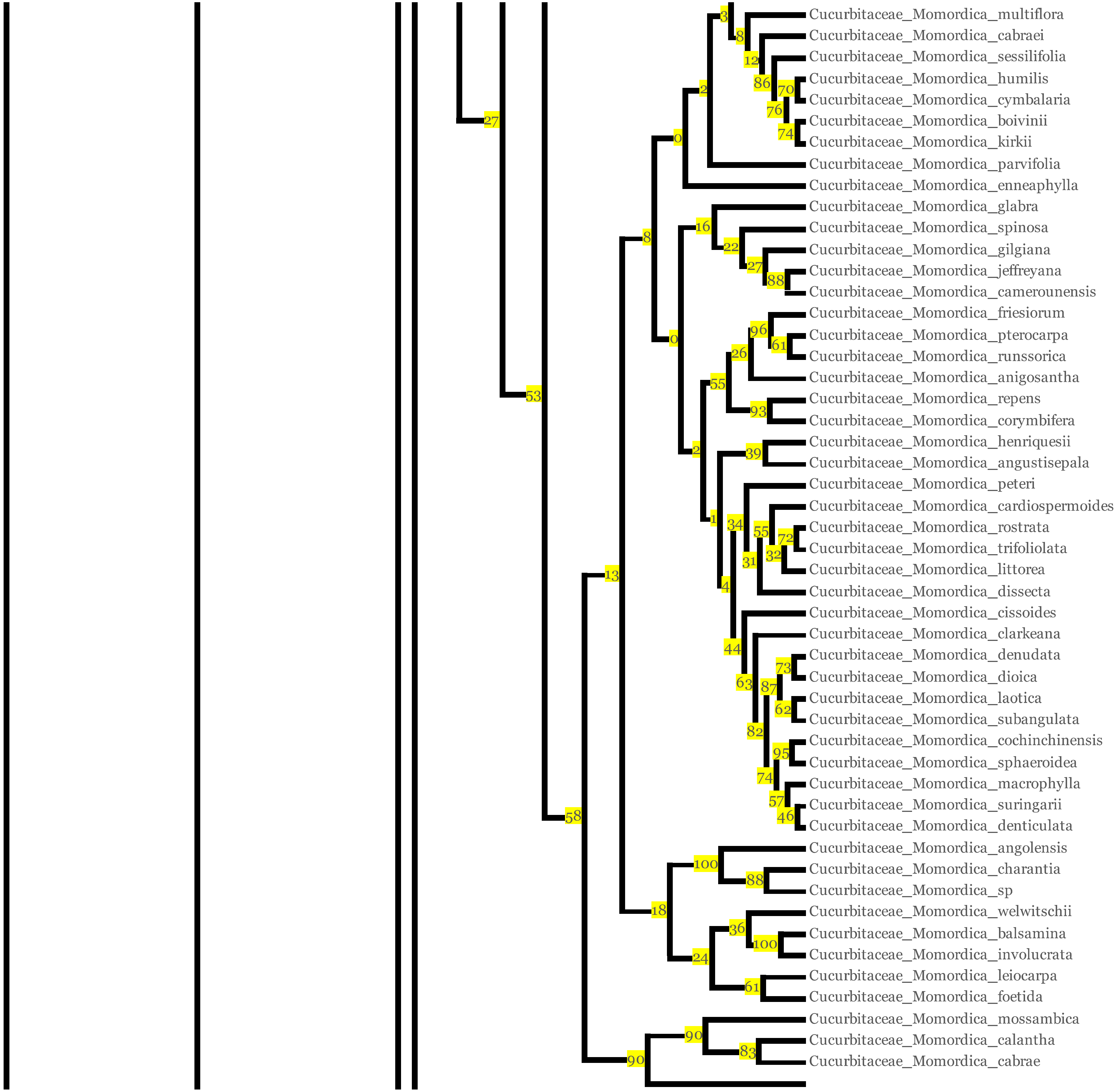
Cucurbitaceae_Momordica_multiflora
Cucurbitaceae_Momordica_cabraei
Cucurbitaceae_Momordica_sessilifolia
Cucurbitaceae_Momordica_humilis
Cucurbitaceae_Momordica_cymbalaria
Cucurbitaceae_Momordica_boivinii
Cucurbitaceae_Momordica_kirkii
Cucurbitaceae_Momordica_parvifolia
Cucurbitaceae_Momordica_enneaphylla
Cucurbitaceae_Momordica_glabra
Cucurbitaceae_Momordica_spinosa
Cucurbitaceae_Momordica_gilgiana
Cucurbitaceae_Momordica_jeffreyana
Cucurbitaceae_Momordica_camerounensis
Cucurbitaceae_Momordica_friesiorum
Cucurbitaceae_Momordica_pterocarpa
Cucurbitaceae_Momordica_runssorica
Cucurbitaceae_Momordica_anigosantha
Cucurbitaceae_Momordica_repens
Cucurbitaceae_Momordica_corymbifera
Cucurbitaceae_Momordica_henriquesii
Cucurbitaceae_Momordica_angustisepala
Cucurbitaceae_Momordica_peteri
Cucurbitaceae_Momordica_cardiospermoides
Cucurbitaceae_Momordica_rostrata
Cucurbitaceae_Momordica_trifoliolata
Cucurbitaceae_Momordica_littorea
Cucurbitaceae_Momordica_dissecta
Cucurbitaceae_Momordica_cissoides
Cucurbitaceae_Momordica_clarkeana
Cucurbitaceae_Momordica_denudata
Cucurbitaceae_Momordica_dioica
Cucurbitaceae_Momordica_laotica
Cucurbitaceae_Momordica_subangulata
Cucurbitaceae_Momordica_cochinchinensis
Cucurbitaceae_Momordica_sphaeroidea
Cucurbitaceae_Momordica_macrophylla
Cucurbitaceae_Momordica_suringarii
Cucurbitaceae_Momordica_denticulata
Cucurbitaceae_Momordica_angolensis
Cucurbitaceae_Momordica_charantia
Cucurbitaceae_Momordica_sp
Cucurbitaceae_Momordica_welwitschii
Cucurbitaceae_Momordica_balsamina
Cucurbitaceae_Momordica_involucrata
Cucurbitaceae_Momordica_leiocarpa
Cucurbitaceae_Momordica_foetida
Cucurbitaceae_Momordica_mossambica
Cucurbitaceae_Momordica_calantha
Cucurbitaceae_Momordica_cabrae

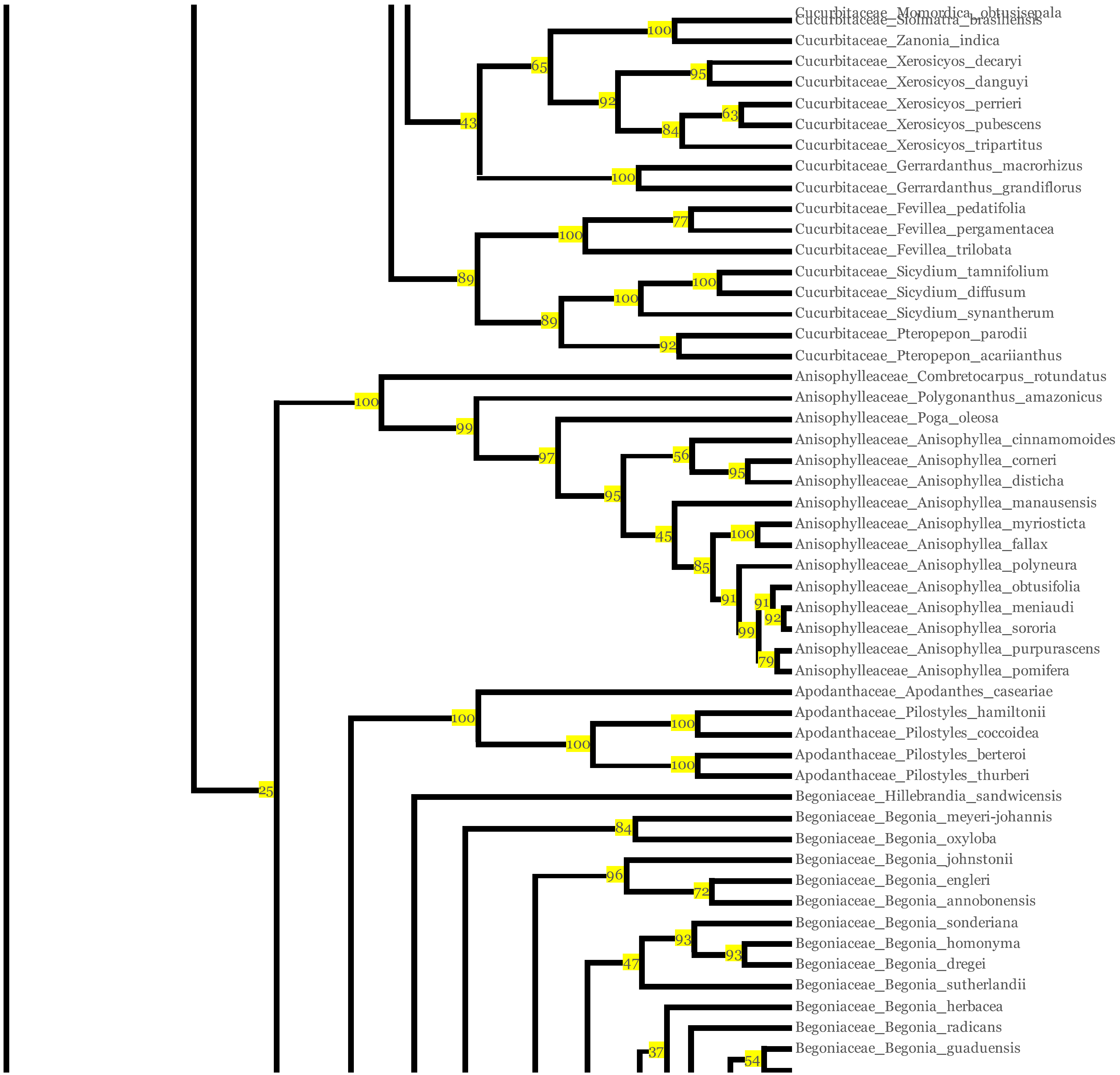
Cucurbitaceae_Momordica_obtusisepala
Cucurbitaceae_Siolmatra_brasiliensis
Cucurbitaceae_Zanonia_indica
Cucurbitaceae_Xerosicyos_decaryi
Cucurbitaceae_Xerosicyos_danguyi
Cucurbitaceae_Xerosicyos_perrieri
Cucurbitaceae_Xerosicyos_pubescens
Cucurbitaceae_Xerosicyos_tripartitus
Cucurbitaceae_Gerrardanthus_macrorhizus
Cucurbitaceae_Gerrardanthus_grandiflorus
Cucurbitaceae_Fevillea_pedatifolia
Cucurbitaceae_Fevillea_pergamentacea
Cucurbitaceae_Fevillea_trilobata
Cucurbitaceae_Sicydium_tamnifolium
Cucurbitaceae_Sicydium_diffusum
Cucurbitaceae_Sicydium_synantherum
Cucurbitaceae_Pteropepon_parodii
Cucurbitaceae_Pteropepon_acariianthus
Anisophylleaceae_Combretocarpus_rotundatus
Anisophylleaceae_Polygonanthus_amazonicus
Anisophylleaceae_Poga_oleosa
Anisophylleaceae_Anisophyllea_cinnamomoides
Anisophylleaceae_Anisophyllea_corneri
Anisophylleaceae_Anisophyllea_disticha
Anisophylleaceae_Anisophyllea_manausensis
Anisophylleaceae_Anisophyllea_myriosticta
Anisophylleaceae_Anisophyllea_fallax
Anisophylleaceae_Anisophyllea_polyneura
Anisophylleaceae_Anisophyllea_obtusifolia
Anisophylleaceae_Anisophyllea_meniaudi
Anisophylleaceae_Anisophyllea_sororia
Anisophylleaceae_Anisophyllea_purpurascens
Anisophylleaceae_Anisophyllea_pomifera
Apodanthaceae_Apodanthes_caseariae
Apodanthaceae_Pilostyles_hamiltonii
Apodanthaceae_Pilostyles_coccoidea
Apodanthaceae_Pilostyles_berteroi
Apodanthaceae_Pilostyles_thurberi
Begoniaceae_Hillebrandia_sandwicensis
Begoniaceae_Begonia_meyeri-johannis
Begoniaceae_Begonia_oxyloba
Begoniaceae_Begonia_johnstonii
Begoniaceae_Begonia_engleri
Begoniaceae_Begonia_annobonensis
Begoniaceae_Begonia_sonderiana
Begoniaceae_Begonia_homonyma
Begoniaceae_Begonia_dregei
Begoniaceae_Begonia_sutherlandii
Begoniaceae_Begonia_herbacea
Begoniaceae_Begonia_radicans
Begoniaceae_Begonia_guaduensis

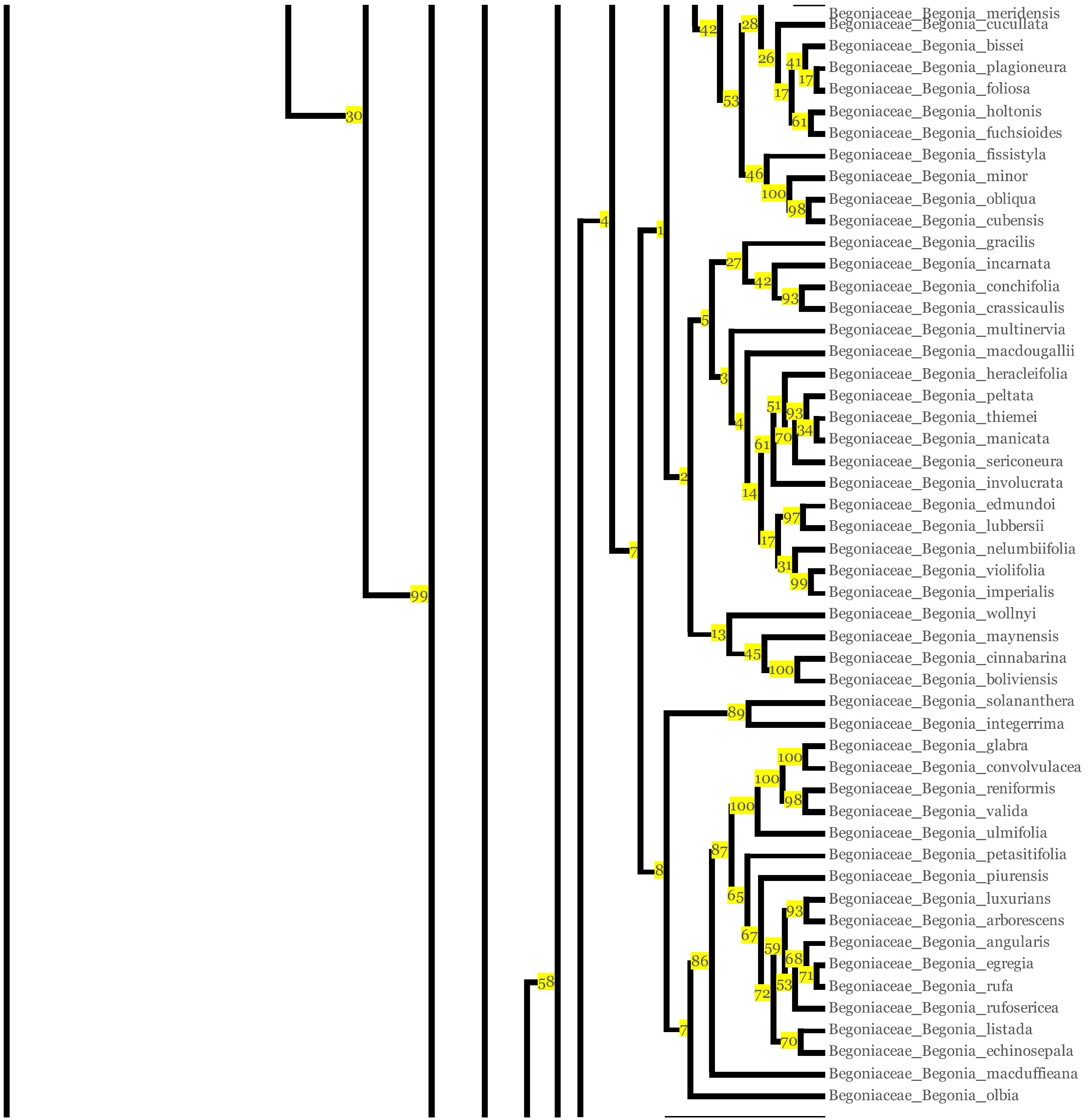
Begoniaceae_Begonia_meridensis
Begoniaceae_Begonia_cucullata
Begoniaceae_Begonia_bissei
Begoniaceae_Begonia_plagioneura
Begoniaceae_Begonia_foliosa
Begoniaceae_Begonia_holtonis
Begoniaceae_Begonia_fuchsioides
Begoniaceae_Begonia_fissistyla
Begoniaceae_Begonia_minor
Begoniaceae_Begonia_obliqua
Begoniaceae_Begonia_cubensis
Begoniaceae_Begonia_gracilis
Begoniaceae_Begonia_incarnata
Begoniaceae_Begonia_conchifolia
Begoniaceae_Begonia_crassicaulis
Begoniaceae_Begonia_multinervia
Begoniaceae_Begonia_macdougallii
Begoniaceae_Begonia_heracleifolia
Begoniaceae_Begonia_peltata
Begoniaceae_Begonia_thiemei
Begoniaceae_Begonia_manicata
Begoniaceae_Begonia_sericoneura
Begoniaceae_Begonia_involucrata
Begoniaceae_Begonia_edmundoi
Begoniaceae_Begonia_lubbersii
Begoniaceae_Begonia_nelumbiifolia
Begoniaceae_Begonia_violifolia
Begoniaceae_Begonia_imperialis
Begoniaceae_Begonia_wollnyi
Begoniaceae_Begonia_maynensis
Begoniaceae_Begonia_cinnabarina
Begoniaceae_Begonia_boliviensis
Begoniaceae_Begonia_solananthera
Begoniaceae_Begonia_integerrima
Begoniaceae_Begonia_glabra
Begoniaceae_Begonia_convolvulacea
Begoniaceae_Begonia_reniformis
Begoniaceae_Begonia_valida
Begoniaceae_Begonia_ulmifolia
Begoniaceae_Begonia_petasitifolia
Begoniaceae_Begonia_piurensis
Begoniaceae_Begonia_luxurians
Begoniaceae_Begonia_arborescens
Begoniaceae_Begonia_angularis
Begoniaceae_Begonia_egregia
Begoniaceae_Begonia_rufa
Begoniaceae_Begonia_rufosericea
Begoniaceae_Begonia_listada
Begoniaceae_Begonia_echinosepala
Begoniaceae_Begonia_macduffieana
Begoniaceae_Begonia_olbia

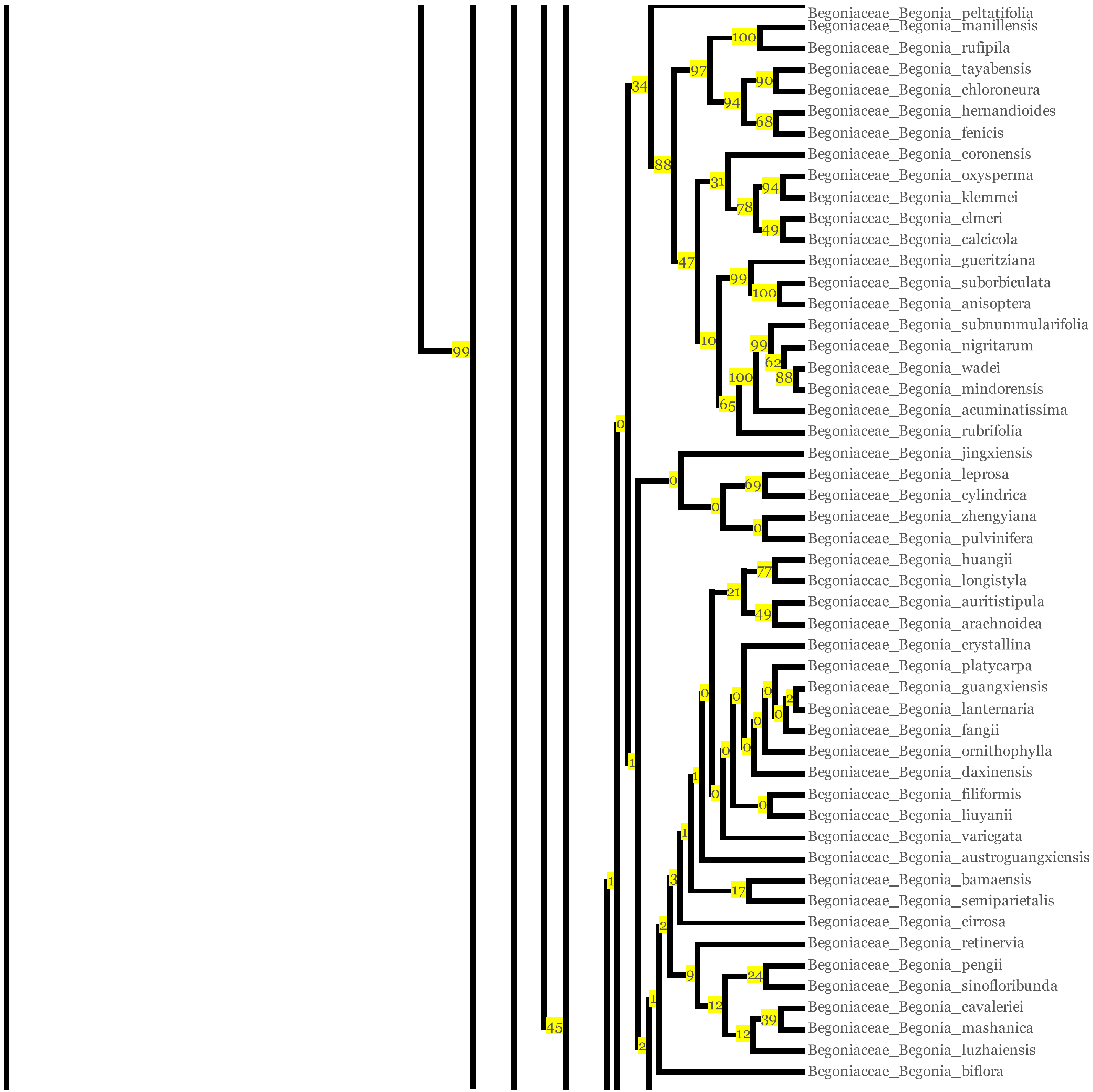
Begoniaceae_Begonia_peltatifolia
Begoniaceae_Begonia_manillensis
Begoniaceae_Begonia_rufipila
Begoniaceae_Begonia_tayabensis
Begoniaceae_Begonia_chloroneura
Begoniaceae_Begonia_hernandioides
Begoniaceae_Begonia_fenicis
Begoniaceae_Begonia_coronensis
Begoniaceae_Begonia_oxysperma
Begoniaceae_Begonia_klemmei
Begoniaceae_Begonia_elmeri
Begoniaceae_Begonia_calcicola
Begoniaceae_Begonia_gueritziana
Begoniaceae_Begonia_suborbiculata
Begoniaceae_Begonia_anisoptera
Begoniaceae_Begonia_subnummularifolia
Begoniaceae_Begonia_nigritarum
Begoniaceae_Begonia_wadei
Begoniaceae_Begonia_mindorensis
Begoniaceae_Begonia_acuminatissima
Begoniaceae_Begonia_rubrifolia
Begoniaceae_Begonia_jingxiensis
Begoniaceae_Begonia_leprosa
Begoniaceae_Begonia_cylindrica
Begoniaceae_Begonia_zhengyiana
Begoniaceae_Begonia_pulvinifera
Begoniaceae_Begonia_huangii
Begoniaceae_Begonia_longistyla
Begoniaceae_Begonia_auritistipula
Begoniaceae_Begonia_arachnoidea
Begoniaceae_Begonia_crystallina
Begoniaceae_Begonia_platycarpa
Begoniaceae_Begonia_guangxiensis
Begoniaceae_Begonia_lanternaria
Begoniaceae_Begonia_fangii
Begoniaceae_Begonia_ornithophylla
Begoniaceae_Begonia_daxinensis
Begoniaceae_Begonia_filiformis
Begoniaceae_Begonia_liuyanii
Begoniaceae_Begonia_variegata
Begoniaceae_Begonia_austroguangxiensis
Begoniaceae_Begonia_bamaensis
Begoniaceae_Begonia_semiparietalis
Begoniaceae_Begonia_cirrosa
Begoniaceae_Begonia_retinervia
Begoniaceae_Begonia_pengii
Begoniaceae_Begonia_sinofloribunda
Begoniaceae_Begonia_cavaleriei
Begoniaceae_Begonia_mashanica
Begoniaceae_Begonia_luzhaiensis
Begoniaceae_Begonia_biflora

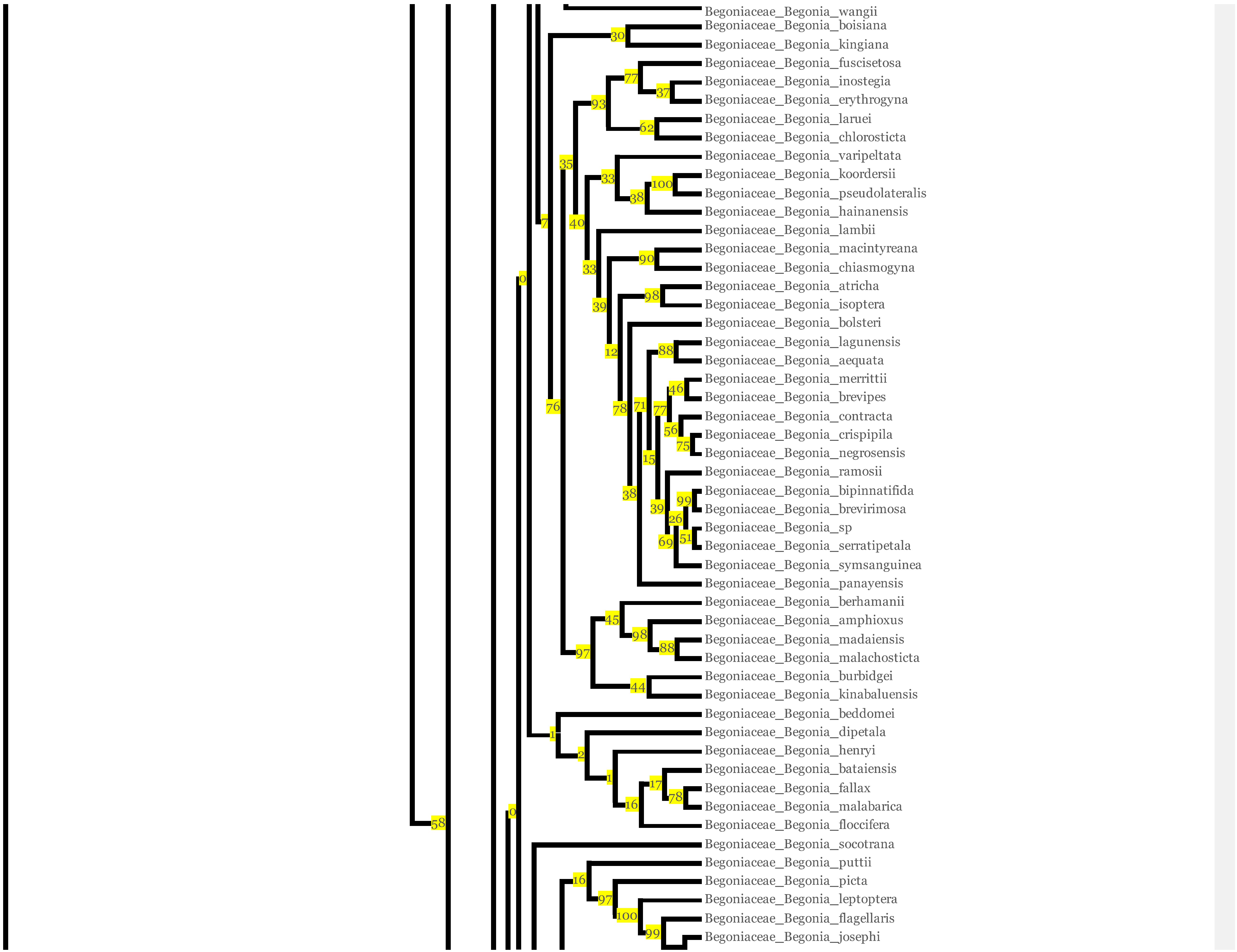
Begoniaceae_Begonia_wangii
Begoniaceae_Begonia_boisiana
Begoniaceae_Begonia_kingiana
Begoniaceae_Begonia_fuscisetosa
Begoniaceae_Begonia_inostegia
Begoniaceae_Begonia_erythrogyna
Begoniaceae_Begonia_laruei
Begoniaceae_Begonia_chlorosticta
Begoniaceae_Begonia_varipeltata
Begoniaceae_Begonia_koordersii
Begoniaceae_Begonia_pseudolateralis
Begoniaceae_Begonia_hainanensis
Begoniaceae_Begonia_lambii
Begoniaceae_Begonia_macintyreana
Begoniaceae_Begonia_chiasmogyna
Begoniaceae_Begonia_atricha
Begoniaceae_Begonia_isoptera
Begoniaceae_Begonia_bolsteri
Begoniaceae_Begonia_lagunensis
Begoniaceae_Begonia_aequata
Begoniaceae_Begonia_merrittii
Begoniaceae_Begonia_brevipes
Begoniaceae_Begonia_contracta
Begoniaceae_Begonia_crispipila
Begoniaceae_Begonia_negrosensis
Begoniaceae_Begonia_ramosii
Begoniaceae_Begonia_bipinnatifida
Begoniaceae_Begonia_brevirimosa
Begoniaceae_Begonia_sp
Begoniaceae_Begonia_serratipetala
Begoniaceae_Begonia_symsanguinea
Begoniaceae_Begonia_panayensis
Begoniaceae_Begonia_berhamanii
Begoniaceae_Begonia_amphioxus
Begoniaceae_Begonia_madaiensis
Begoniaceae_Begonia_malachosticta
Begoniaceae_Begonia_burbidgei
Begoniaceae_Begonia_kinabaluensis
Begoniaceae_Begonia_beddomei
Begoniaceae_Begonia_dipetala
Begoniaceae_Begonia_henryi
Begoniaceae_Begonia_bataiensis
Begoniaceae_Begonia_fallax
Begoniaceae_Begonia_malabarica
Begoniaceae_Begonia_floccifera
Begoniaceae_Begonia_socotrana
Begoniaceae_Begonia_puttii
Begoniaceae_Begonia_picta
Begoniaceae_Begonia_leptoptera
Begoniaceae_Begonia_flagellaris
Begoniaceae_Begonia_josephi

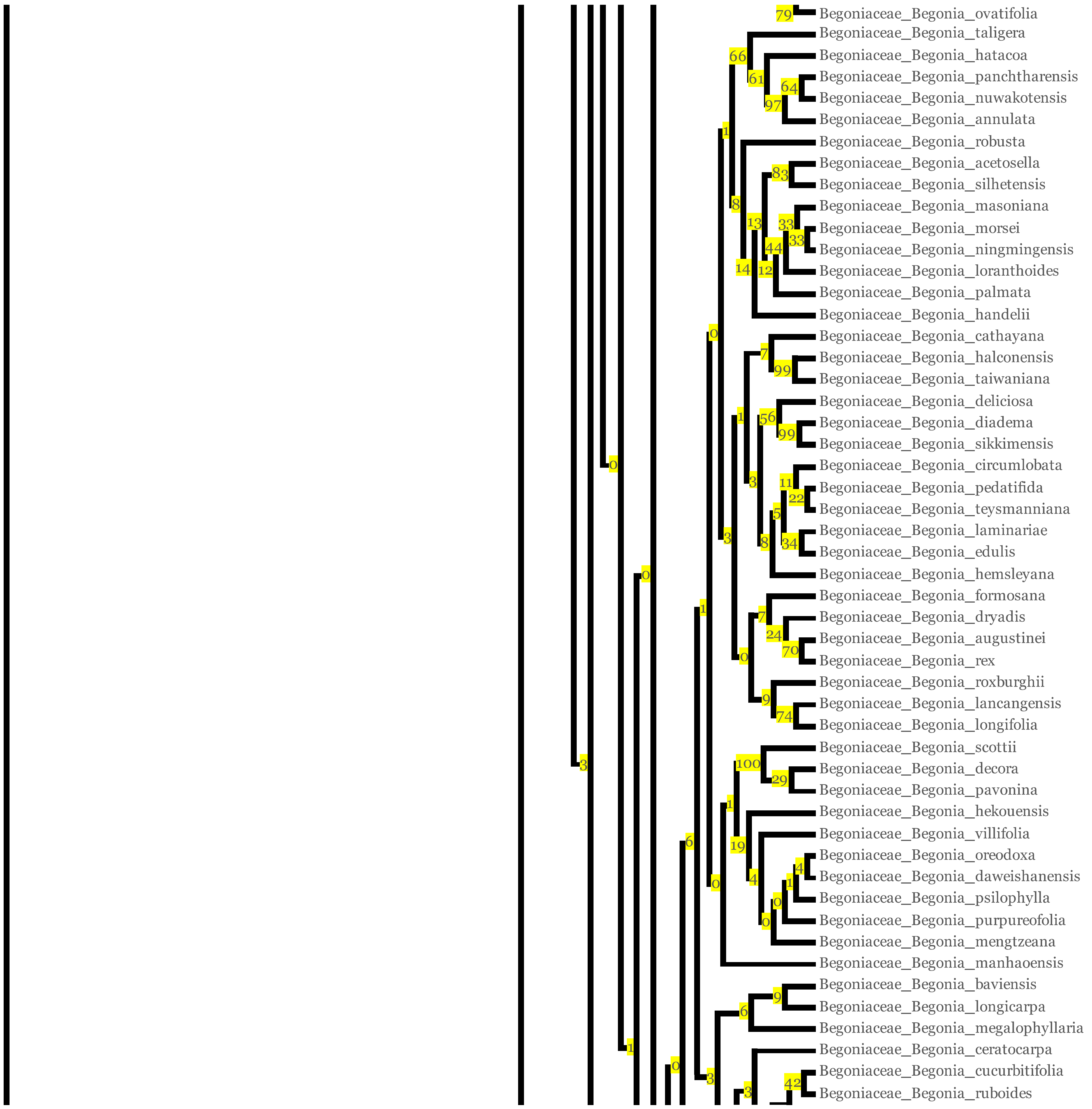
Begoniaceae_Begonia_ovatifolia
Begoniaceae_Begonia_taligera
Begoniaceae_Begonia_hatacoa
Begoniaceae_Begonia_panchtharensis
Begoniaceae_Begonia_nuwakotensis
Begoniaceae_Begonia_annulata
Begoniaceae_Begonia_robusta
Begoniaceae_Begonia_acetosella
Begoniaceae_Begonia_silhetensis
Begoniaceae_Begonia_masoniana
Begoniaceae_Begonia_morsei
Begoniaceae_Begonia_ningmingensis
Begoniaceae_Begonia_loranthoides
Begoniaceae_Begonia_palmata
Begoniaceae_Begonia_handelii
Begoniaceae_Begonia_cathayana
Begoniaceae_Begonia_halconensis
Begoniaceae_Begonia_taiwaniana
Begoniaceae_Begonia_deliciosa
Begoniaceae_Begonia_diadema
Begoniaceae_Begonia_sikkimensis
Begoniaceae_Begonia_circumlobata
Begoniaceae_Begonia_pedatifida
Begoniaceae_Begonia_teysmanniana
Begoniaceae_Begonia_laminariae
Begoniaceae_Begonia_edulis
Begoniaceae_Begonia_hemsleyana
Begoniaceae_Begonia_formosana
Begoniaceae_Begonia_dryadis
Begoniaceae_Begonia_augustinei
Begoniaceae_Begonia_rex
Begoniaceae_Begonia_roxburghii
Begoniaceae_Begonia_lancangensis
Begoniaceae_Begonia_longifolia
Begoniaceae_Begonia_scottii
Begoniaceae_Begonia_decora
Begoniaceae_Begonia_pavonina
Begoniaceae_Begonia_hekouensis
Begoniaceae_Begonia_villifolia
Begoniaceae_Begonia_oreodoxa
Begoniaceae_Begonia_daweishanensis
Begoniaceae_Begonia_psilophylla
Begoniaceae_Begonia_purpureofolia
Begoniaceae_Begonia_mengtzeana
Begoniaceae_Begonia_manhaoensis
Begoniaceae_Begonia_baviensis
Begoniaceae_Begonia_longicarpa
Begoniaceae_Begonia_megalophyllaria
Begoniaceae_Begonia_ceratocarpa
Begoniaceae_Begonia_cucurbitifolia
Begoniaceae_Begonia_ruboides

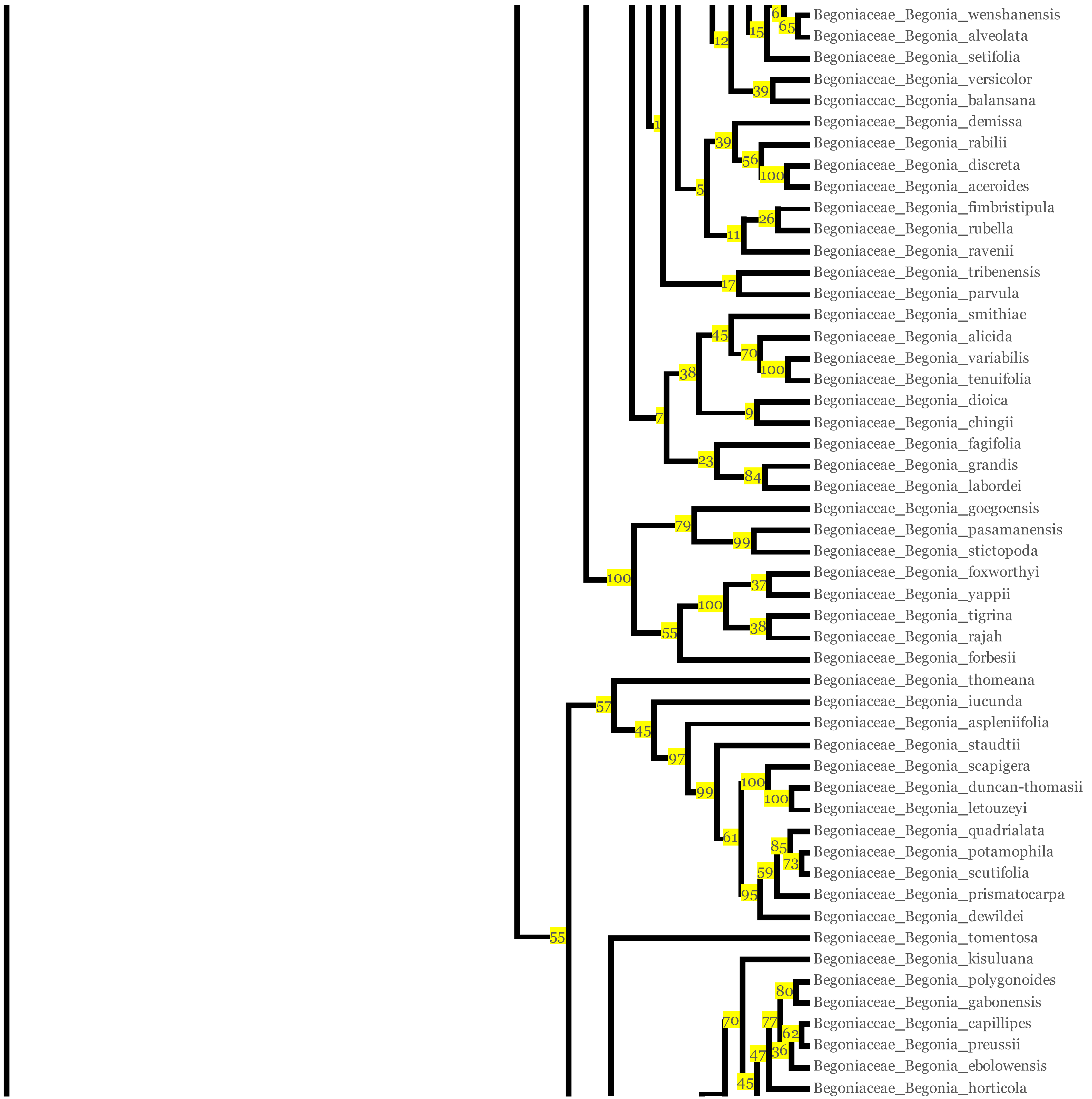
Begoniaceae_Begonia_wenshanensis
Begoniaceae_Begonia_alveolata
Begoniaceae_Begonia_setifolia
Begoniaceae_Begonia_versicolor
Begoniaceae_Begonia_balansana
Begoniaceae_Begonia_demissa
Begoniaceae_Begonia_rabilii
Begoniaceae_Begonia_discreta
Begoniaceae_Begonia_aceroides
Begoniaceae_Begonia_fimbristipula
Begoniaceae_Begonia_rubella
Begoniaceae_Begonia_ravenii
Begoniaceae_Begonia_tribenensis
Begoniaceae_Begonia_parvula
Begoniaceae_Begonia_smithiae
Begoniaceae_Begonia_alicida
Begoniaceae_Begonia_variabilis
Begoniaceae_Begonia_tenuifolia
Begoniaceae_Begonia_dioica
Begoniaceae_Begonia_chingii
Begoniaceae_Begonia_fagifolia
Begoniaceae_Begonia_grandis
Begoniaceae_Begonia_labordei
Begoniaceae_Begonia_goegoensis
Begoniaceae_Begonia_pasamanensis
Begoniaceae_Begonia_stictopoda
Begoniaceae_Begonia_foxworthyi
Begoniaceae_Begonia_yappii
Begoniaceae_Begonia_tigrina
Begoniaceae_Begonia_rajah
Begoniaceae_Begonia_forbesii
Begoniaceae_Begonia_thomeana
Begoniaceae_Begonia_iucunda
Begoniaceae_Begonia_aspleniifolia
Begoniaceae_Begonia_staudtii
Begoniaceae_Begonia_scapigera
Begoniaceae_Begonia_duncan-thomasii
Begoniaceae_Begonia_letouzeyi
Begoniaceae_Begonia_quadrialata
Begoniaceae_Begonia_potamophila
Begoniaceae_Begonia_scutifolia
Begoniaceae_Begonia_prismatocarpa
Begoniaceae_Begonia_dewildei
Begoniaceae_Begonia_tomentosa
Begoniaceae_Begonia_kisuluana
Begoniaceae_Begonia_polygonoides
Begoniaceae_Begonia_gabonensis
Begoniaceae_Begonia_capillipes
Begoniaceae_Begonia_preussii
Begoniaceae_Begonia_ebolowensis
Begoniaceae_Begonia_horticola

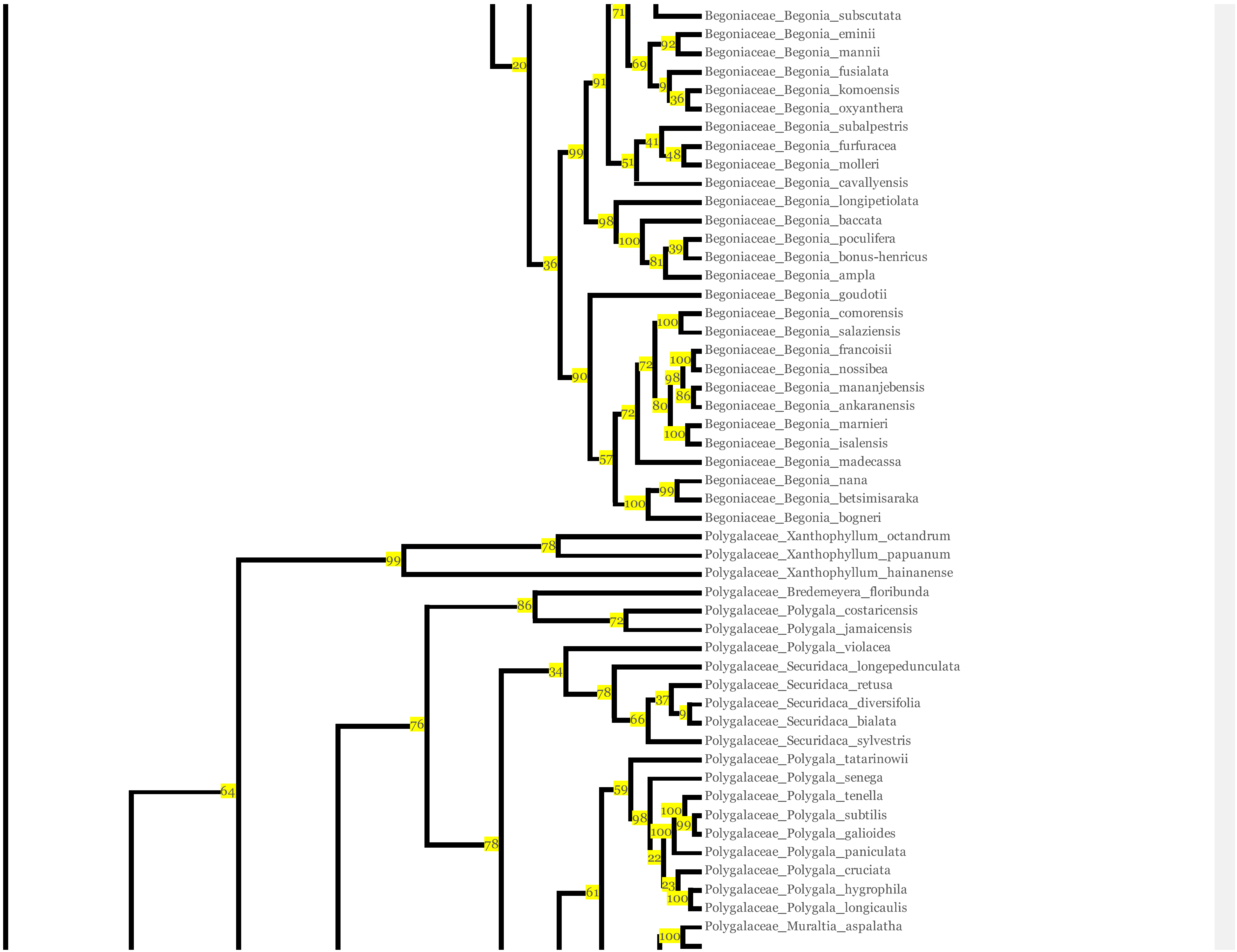

Begoniaceae_Begonia_subscutata
Begoniaceae_Begonia_eminii
Begoniaceae_Begonia_mannii
Begoniaceae_Begonia_fusialata
Begoniaceae_Begonia_komoensis
Begoniaceae_Begonia_oxyanthera
Begoniaceae_Begonia_subalpestris
Begoniaceae_Begonia_furfuracea
Begoniaceae_Begonia_molleri
Begoniaceae_Begonia_cavallyensis
Begoniaceae_Begonia_longipetiolata
Begoniaceae_Begonia_baccata
Begoniaceae_Begonia_poculifera
Begoniaceae_Begonia_bonus-henricus
Begoniaceae_Begonia_ampla
Begoniaceae_Begonia_goudotii
Begoniaceae_Begonia_comorensis
Begoniaceae_Begonia_salaziensis
Begoniaceae_Begonia_francoisii
Begoniaceae_Begonia_nossibea
Begoniaceae_Begonia_mananjebensis
Begoniaceae_Begonia_ankaranensis
Begoniaceae_Begonia_marnieri
Begoniaceae_Begonia_isalensis
Begoniaceae_Begonia_madecassa
Begoniaceae_Begonia_nana
Begoniaceae_Begonia_betsimisaraka
Begoniaceae_Begonia_bogneri
Polygalaceae_Xanthophyllum_octandrum
Polygalaceae_Xanthophyllum_papuanum
Polygalaceae_Xanthophyllum_hainanense
Polygalaceae_Bredemeyera_floribunda
Polygalaceae_Polygala_costaricensis
Polygalaceae_Polygala_jamaicensis
Polygalaceae_Polygala_violacea
Polygalaceae_Securidaca_longepedunculata
Polygalaceae_Securidaca_retusa
Polygalaceae_Securidaca_diversifolia
Polygalaceae_Securidaca_bialata
Polygalaceae_Securidaca_sylvestris
Polygalaceae_Polygala_tatarinowii
Polygalaceae_Polygala_senega
Polygalaceae_Polygala_tenella
Polygalaceae_Polygala_subtilis
Polygalaceae_Polygala_galioides
Polygalaceae_Polygala_paniculata
Polygalaceae_Polygala_cruciata
Polygalaceae_Polygala_hygrophila
Polygalaceae_Polygala_longicaulis
Polygalaceae_Muraltia_aspalatha

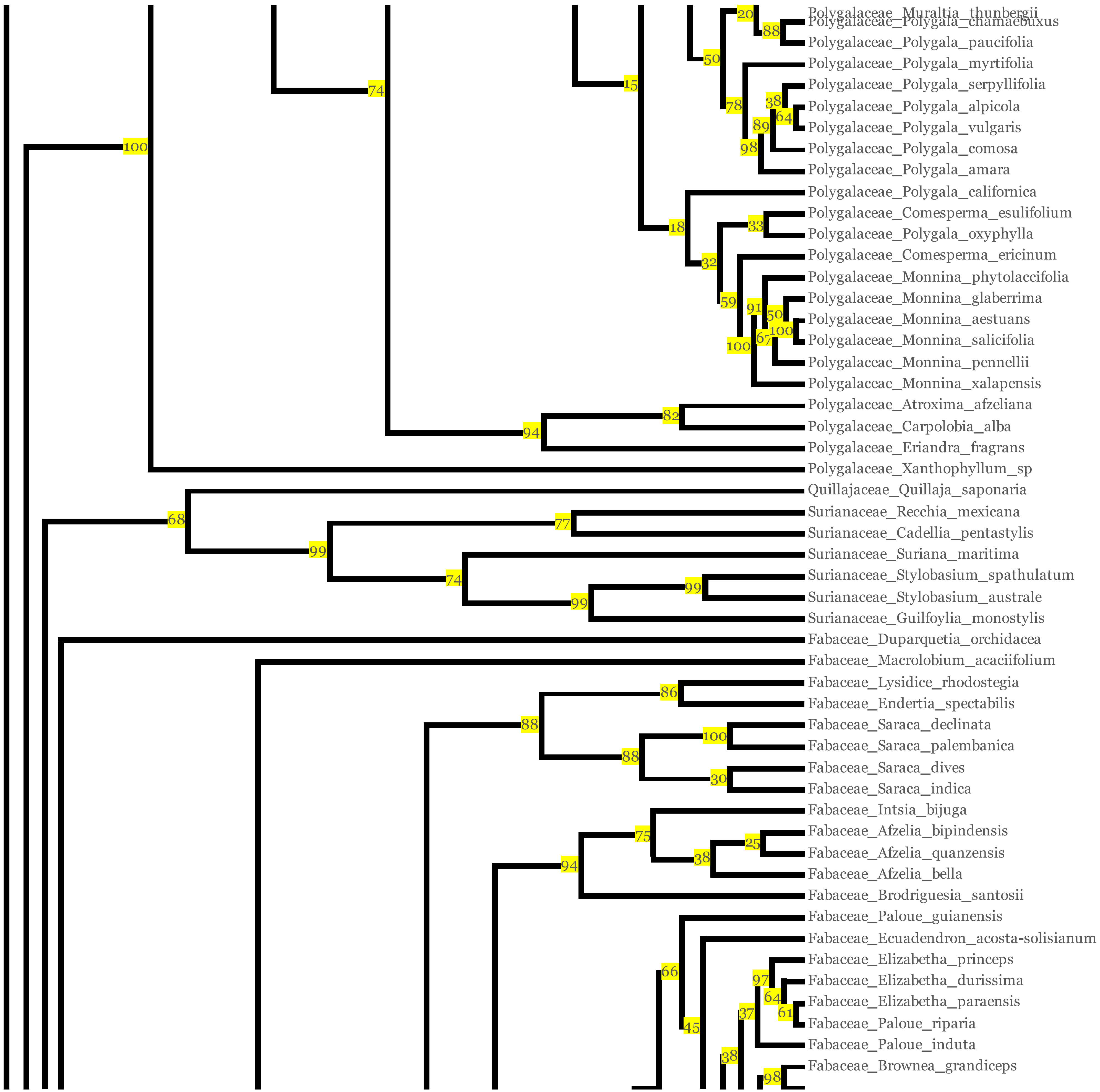
Polygalaceae_Muraltia_thunbergii
Polygalaceae_Polygala_chamaebuxus
Polygalaceae_Polygala_paucifolia
Polygalaceae_Polygala_myrtifolia
Polygalaceae_Polygala_serpyllifolia
Polygalaceae_Polygala_alpicola
Polygalaceae_Polygala_vulgaris
Polygalaceae_Polygala_comosa
Polygalaceae_Polygala_amara
Polygalaceae_Polygala_californica
Polygalaceae_Comesperma_esulifolium
Polygalaceae_Polygala_oxyphylla
Polygalaceae_Comesperma_ericinum
Polygalaceae_Monnina_phytolaccifolia
Polygalaceae_Monnina_glaberrima
Polygalaceae_Monnina_aestuans
Polygalaceae_Monnina_salicifolia
Polygalaceae_Monnina_pennellii
Polygalaceae_Monnina_xalapensis
Polygalaceae_Atroxima_afzeliana
Polygalaceae_Carpolobia_alba
Polygalaceae_Eriandra_fragrans
Polygalaceae_Xanthophyllum_sp
Quillajaceae_Quillaja_saponaria
Surianaceae_Recchia_mexicana
Surianaceae_Cadellia_pentastylis
Surianaceae_Suriana_maritima
Surianaceae_Stylobasium_spathulatum
Surianaceae_Stylobasium_australe
Surianaceae_Guilfoylia_monostylis
Fabaceae_Duparquetia_orchidacea
Fabaceae_Macrolobium_acaciifolium
Fabaceae_Lysidice_rhodostegia
Fabaceae_Endertia_spectabilis
Fabaceae_Saraca_declinata
Fabaceae_Saraca_palembanica
Fabaceae_Saraca_dives
Fabaceae_Saraca_indica
Fabaceae_Intsia_bijuga
Fabaceae_Afzelia_bipindensis
Fabaceae_Afzelia_quanzensis
Fabaceae_Afzelia_bella
Fabaceae_Brodriguesia_santosii
Fabaceae_Paloue_guianensis
Fabaceae_Ecuadendron_acosta-solisianum
Fabaceae_Elizabetha_princeps
Fabaceae_Elizabetha_durissima
Fabaceae_Elizabetha_paraensis
Fabaceae_Paloue_riparia
Fabaceae_Paloue_induta
Fabaceae_Brownea_grandiceps

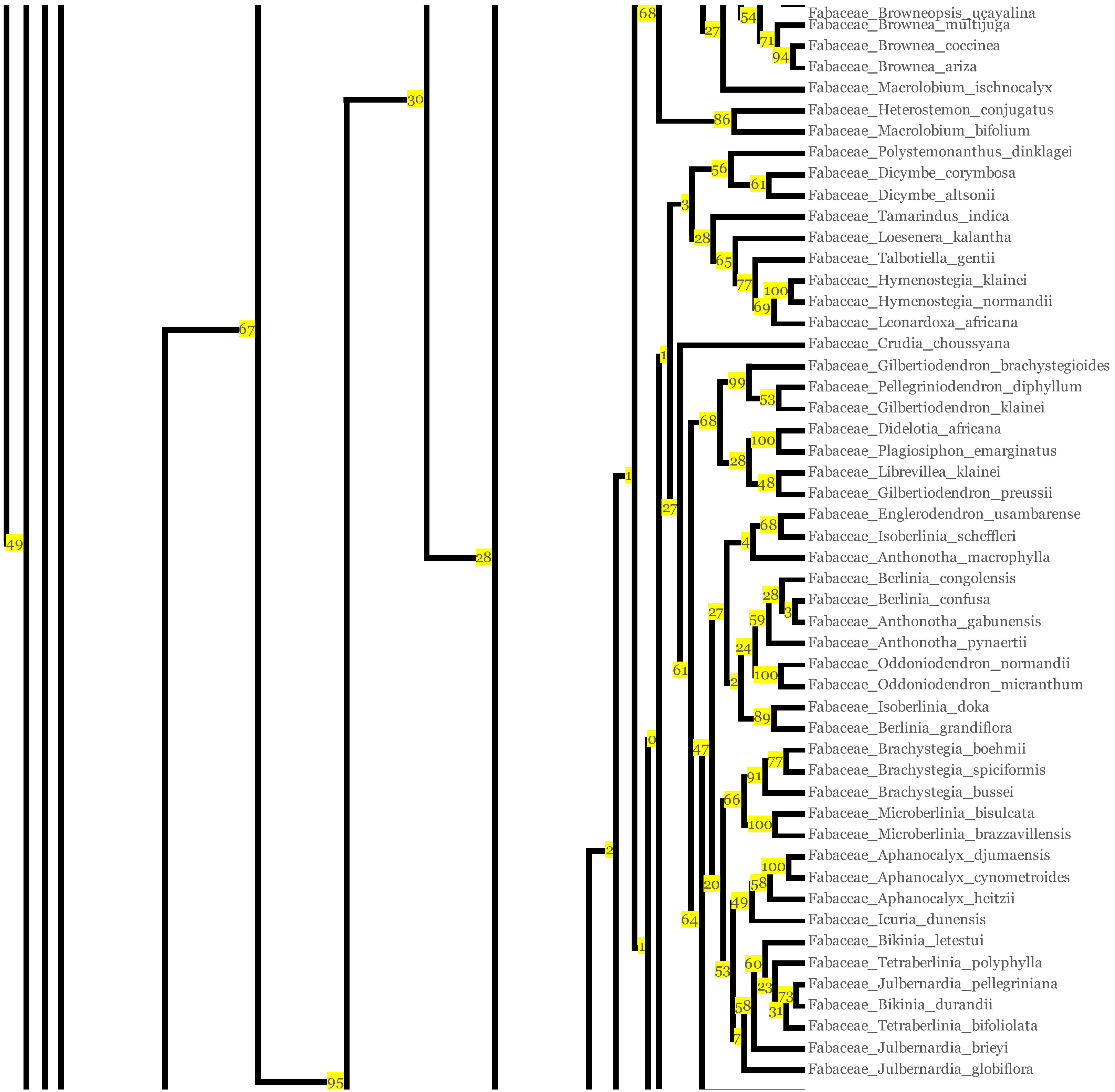

Fabaceae_Browneopsis_ucayalina
Fabaceae_Brownea_multijuga
Fabaceae_Brownea_coccinea
Fabaceae_Brownea_ariza
Fabaceae_Macrolobium_ischnocalyx
Fabaceae_Heterostemon_conjugatus
Fabaceae_Macrolobium_bifolium
Fabaceae_Polystemonanthus_dinklagei
Fabaceae_Dicymbe_corymbosa
Fabaceae_Dicymbe_altsonii
Fabaceae_Tamarindus_indica
Fabaceae_Loesenera_kalantha
Fabaceae_Talbotiella_gentii
Fabaceae_Hymenostegia_klainei
Fabaceae_Hymenostegia_normandii
Fabaceae_Leonardoxa_africana
Fabaceae_Crudia_choussyana
Fabaceae_Gilbertiodendron_brachystegioides
Fabaceae_Pellegriniodendron_diphyllum
Fabaceae_Gilbertiodendron_klainei
Fabaceae_Didelotia_africana
Fabaceae_Plagiosiphon_emarginatus
Fabaceae_Librevillea_klainei
Fabaceae_Gilbertiodendron_preussii
Fabaceae_Englerodendron_usambarense
Fabaceae_Isoberlinia_scheffleri
Fabaceae_Anthonotha_macrophylla
Fabaceae_Berlinia_congolensis
Fabaceae_Berlinia_confusa
Fabaceae_Anthonotha_gabunensis
Fabaceae_Anthonotha_pynaertii
Fabaceae_Oddoniodendron_normandii
Fabaceae_Oddoniodendron_micranthum
Fabaceae_Isoberlinia_doka
Fabaceae_Berlinia_grandiflora
Fabaceae_Brachystegia_boehmii
Fabaceae_Brachystegia_spiciformis
Fabaceae_Brachystegia_bussei
Fabaceae_Microberlinia_bisulcata
Fabaceae_Microberlinia_brazzavillensis
Fabaceae_Aphanocalyx_djumaensis
Fabaceae_Aphanocalyx_cynometroides
Fabaceae_Aphanocalyx_heitzii
Fabaceae_Icuria_dunensis
Fabaceae_Bikinia_letestui
Fabaceae_Tetraberlinia_polyphylla
Fabaceae_Julbernardia_pellegriniana
Fabaceae_Bikinia_durandii
Fabaceae_Tetraberlinia_bifoliolata
Fabaceae_Julbernardia_brieyi
Fabaceae_Julbernardia_globiflora

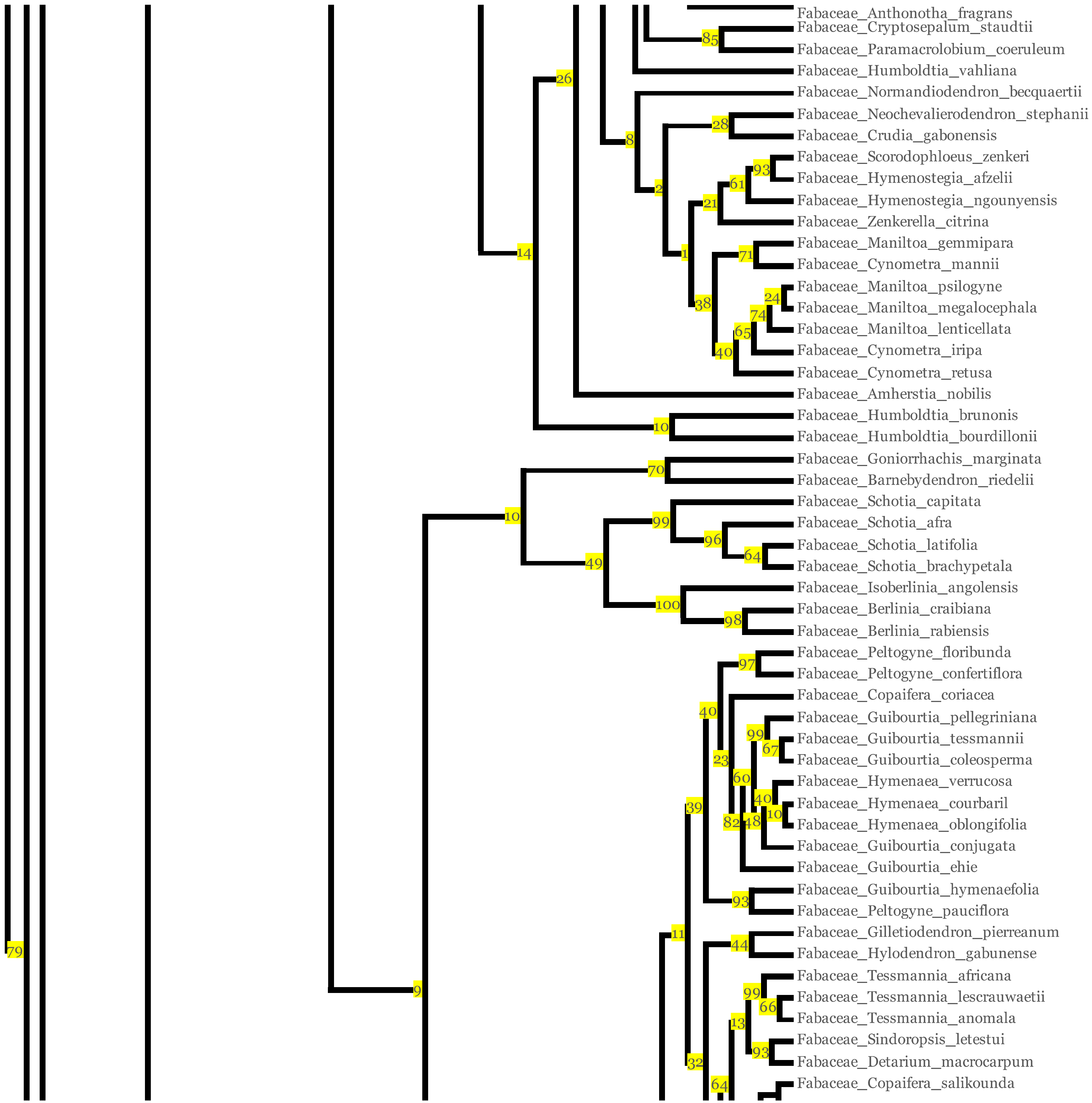
Fabaceae_Anthonotha_fragrans
Fabaceae_Cryptosepalum_staudtii
Fabaceae_Paramacrolobium_coeruleum
Fabaceae_Humboldtia_vahliana
Fabaceae_Normandiodendron_becquaertii
Fabaceae_Neochevalierodendron_stephanii
Fabaceae_Crudia_gabonensis
Fabaceae_Scorodophloeus_zenkeri
Fabaceae_Hymenostegia_afzelii
Fabaceae_Hymenostegia_ngounyensis
Fabaceae_Zenkerella_citrina
Fabaceae_Maniltoa_gemmipara
Fabaceae_Cynometra_mannii
Fabaceae_Maniltoa_psilogyne
Fabaceae_Maniltoa_megalocephala
Fabaceae_Maniltoa_lenticellata
Fabaceae_Cynometra_iripa
Fabaceae_Cynometra_retusa
Fabaceae_Amherstia_nobilis
Fabaceae_Humboldtia_brunonis
Fabaceae_Humboldtia_bourdillonii
Fabaceae_Goniorrhachis_marginata
Fabaceae_Barnebydendron_riedelii
Fabaceae_Schotia_capitata
Fabaceae_Schotia_afra
Fabaceae_Schotia_latifolia
Fabaceae_Schotia_brachypetala
Fabaceae_Isoberlinia_angolensis
Fabaceae_Berlinia_craibiana
Fabaceae_Berlinia_rabiensis
Fabaceae_Peltogyne_floribunda
Fabaceae_Peltogyne_confertiflora
Fabaceae_Copaifera_coriacea
Fabaceae_Guibourtia_pellegriniana
Fabaceae_Guibourtia_tessmannii
Fabaceae_Guibourtia_coleosperma
Fabaceae_Hymenaea_verrucosa
Fabaceae_Hymenaea_courbaril
Fabaceae_Hymenaea_oblongifolia
Fabaceae_Guibourtia_conjugata
Fabaceae_Guibourtia_ehie
Fabaceae_Guibourtia_hymenaefolia
Fabaceae_Peltogyne_pauciflora
Fabaceae_Gilletiodendron_pierreanum
Fabaceae_Hylodendron_gabunense
Fabaceae_Tessmannia_africana
Fabaceae_Tessmannia_lescrauwaetii
Fabaceae_Tessmannia_anomala
Fabaceae_Sindoropsis_letestui
Fabaceae_Detarium_macrocarpum
Fabaceae_Copaifera_salikounda

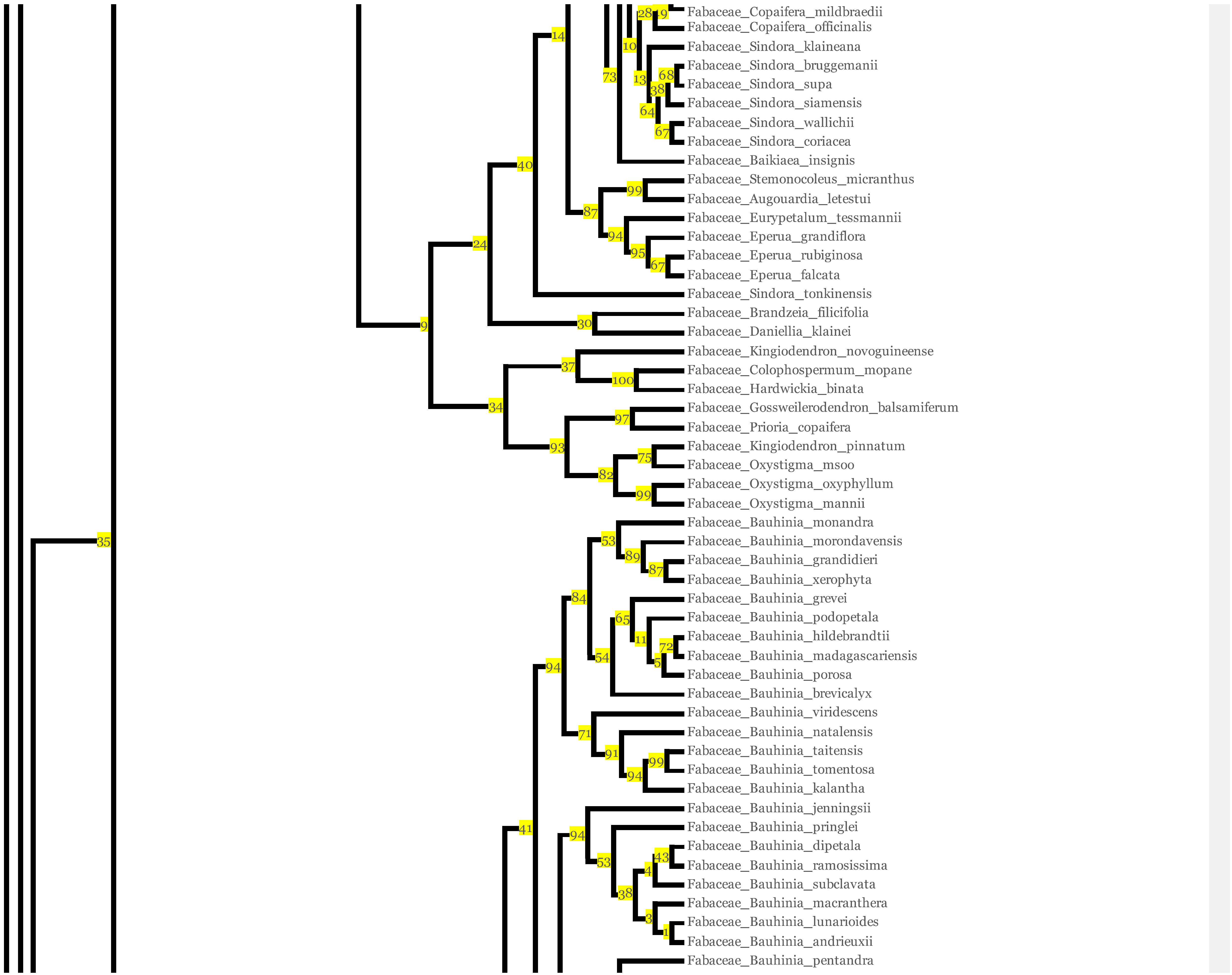
Fabaceae_Copaifera_mildbraedii
Fabaceae_Copaifera_officinalis
Fabaceae_Sindora_klaineana
Fabaceae_Sindora_bruggemanii
Fabaceae_Sindora_supa
Fabaceae_Sindora_siamensis
Fabaceae_Sindora_wallichii
Fabaceae_Sindora_coriacea
Fabaceae_Baikiaea_insignis
Fabaceae_Stemonocoleus_micranthus
Fabaceae_Augouardia_letestui
Fabaceae_Eurypetalum_tessmannii
Fabaceae_Eperua_grandiflora
Fabaceae_Eperua_rubiginosa
Fabaceae_Eperua_falcata
Fabaceae_Sindora_tonkinensis
Fabaceae_Brandzeia_filicifolia
Fabaceae_Daniellia_klainei
Fabaceae_Kingiodendron_novoguineense
Fabaceae_Colophospermum_mopane
Fabaceae_Hardwickia_binata
Fabaceae_Gossweilerodendron_balsamiferum
Fabaceae_Prioria_copaifera
Fabaceae_Kingiodendron_pinnatum
Fabaceae_Oxystigma_msoo
Fabaceae_Oxystigma_oxyphyllum
Fabaceae_Oxystigma_mannii
Fabaceae_Bauhinia_monandra
Fabaceae_Bauhinia_morondavensis
Fabaceae_Bauhinia_grandidieri
Fabaceae_Bauhinia_xerophyta
Fabaceae_Bauhinia_grevei
Fabaceae_Bauhinia_podopetala
Fabaceae_Bauhinia_hildebrandtii
Fabaceae_Bauhinia_madagascariensis
Fabaceae_Bauhinia_porosa
Fabaceae_Bauhinia_brevicalyx
Fabaceae_Bauhinia_viridescens
Fabaceae_Bauhinia_natalensis
Fabaceae_Bauhinia_taitensis
Fabaceae_Bauhinia_tomentosa
Fabaceae_Bauhinia_kalantha
Fabaceae_Bauhinia_jenningsii
Fabaceae_Bauhinia_pringlei
Fabaceae_Bauhinia_dipetala
Fabaceae_Bauhinia_ramosissima
Fabaceae_Bauhinia_subclavata
Fabaceae_Bauhinia_macranthera
Fabaceae_Bauhinia_lunarioides
Fabaceae_Bauhinia_andrieuxii
Fabaceae_Bauhinia_pentandra

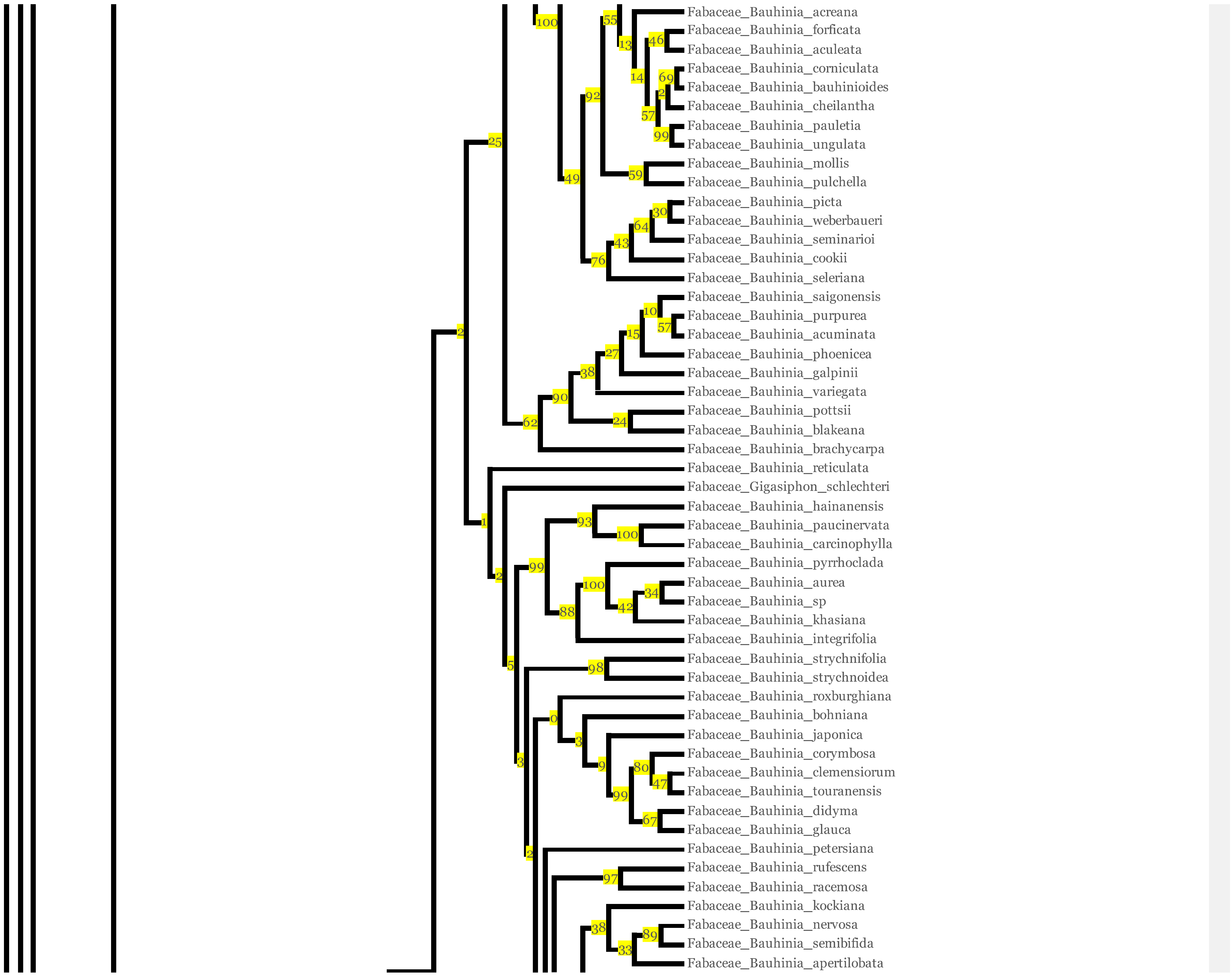
Fabaceae_Bauhinia_acreana
Fabaceae_Bauhinia_forficata
Fabaceae_Bauhinia_aculeata
Fabaceae_Bauhinia_corniculata
Fabaceae_Bauhinia_bauhinioides
Fabaceae_Bauhinia_cheilantha
Fabaceae_Bauhinia_pauletia
Fabaceae_Bauhinia_ungulata
Fabaceae_Bauhinia_mollis
Fabaceae_Bauhinia_pulchella
Fabaceae_Bauhinia_picta
Fabaceae_Bauhinia_weberbaueri
Fabaceae_Bauhinia_seminarioi
Fabaceae_Bauhinia_cookii
Fabaceae_Bauhinia_seleriana
Fabaceae_Bauhinia_saigonensis
Fabaceae_Bauhinia_purpurea
Fabaceae_Bauhinia_acuminata
Fabaceae_Bauhinia_phoenicea
Fabaceae_Bauhinia_galpinii
Fabaceae_Bauhinia_variegata
Fabaceae_Bauhinia_pottsii
Fabaceae_Bauhinia_blakeana
Fabaceae_Bauhinia_brachycarpa
Fabaceae_Bauhinia_reticulata
Fabaceae_Gigasiphon_schlechteri
Fabaceae_Bauhinia_hainanensis
Fabaceae_Bauhinia_paucinervata
Fabaceae_Bauhinia_carcinophylla
Fabaceae_Bauhinia_pyrrhoclada
Fabaceae_Bauhinia_aurea
Fabaceae_Bauhinia_sp
Fabaceae_Bauhinia_khasiana
Fabaceae_Bauhinia_integrifolia
Fabaceae_Bauhinia_strychnifolia
Fabaceae_Bauhinia_strychnoidea
Fabaceae_Bauhinia_roxburghiana
Fabaceae_Bauhinia_bohniana
Fabaceae_Bauhinia_japonica
Fabaceae_Bauhinia_corymbosa
Fabaceae_Bauhinia_clemensiorum
Fabaceae_Bauhinia_touranensis
Fabaceae_Bauhinia_didyma
Fabaceae_Bauhinia_glauca
Fabaceae_Bauhinia_petersiana
Fabaceae_Bauhinia_rufescens
Fabaceae_Bauhinia_racemosa
Fabaceae_Bauhinia_kockiana
Fabaceae_Bauhinia_nervosa
Fabaceae_Bauhinia_semibifida
Fabaceae_Bauhinia_apertilobata

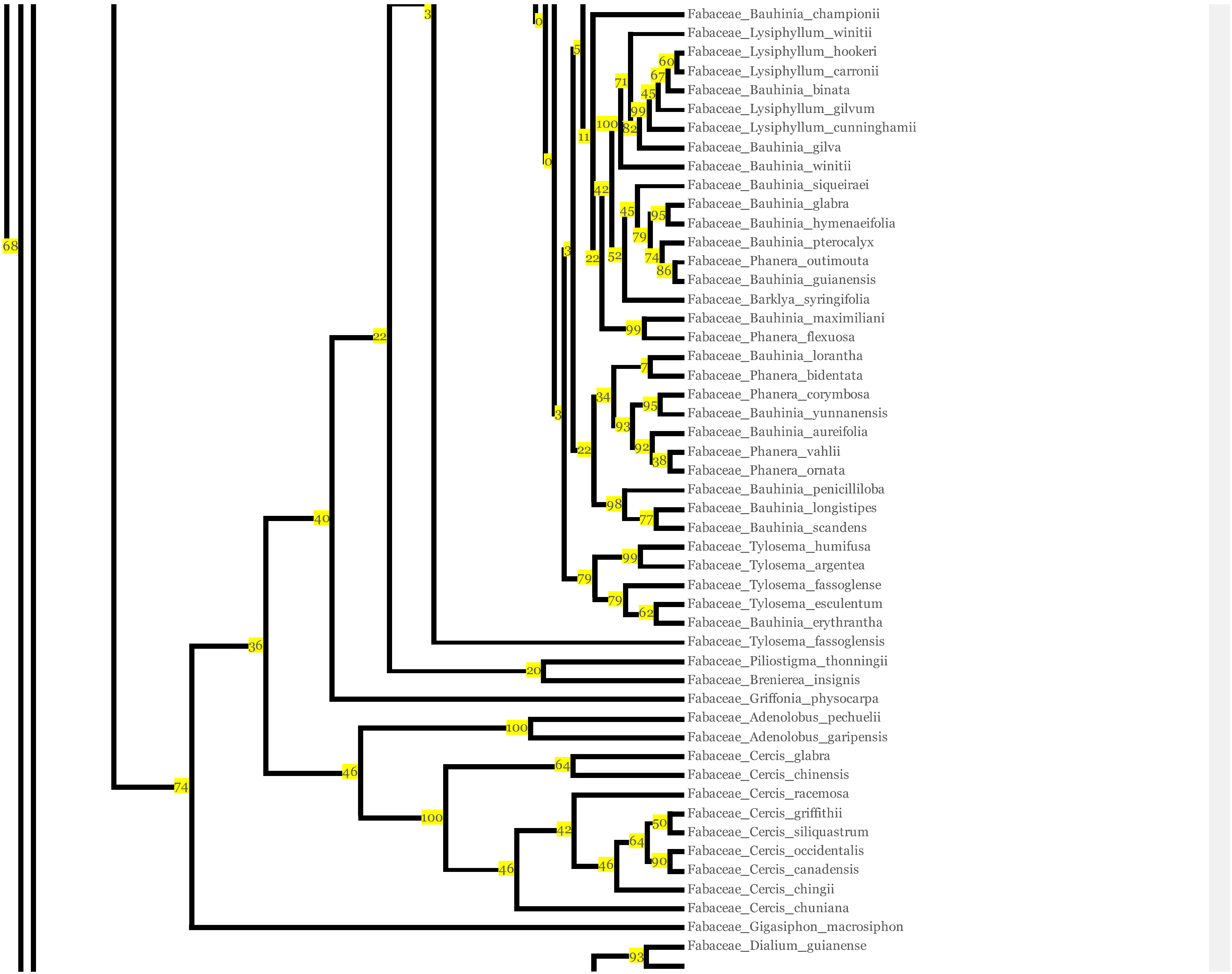

Fabaceae_Bauhinia_championii
Fabaceae_Lysiphyllum_winitii
Fabaceae_Lysiphyllum_hookeri
Fabaceae_Lysiphyllum_carronii
Fabaceae_Bauhinia_binata
Fabaceae_Lysiphyllum_gilvum
Fabaceae_Lysiphyllum_cunninghamii
Fabaceae_Bauhinia_gilva
Fabaceae_Bauhinia_winitii
Fabaceae_Bauhinia_siqueiraei
Fabaceae_Bauhinia_glabra
Fabaceae_Bauhinia_hymenaeifolia
Fabaceae_Bauhinia_pterocalyx
Fabaceae_Phanera_outimouta
Fabaceae_Bauhinia_guianensis
Fabaceae_Barklya_syringifolia
Fabaceae_Bauhinia_maximiliani
Fabaceae_Phanera_flexuosa
Fabaceae_Bauhinia_lorantha
Fabaceae_Phanera_bidentata
Fabaceae_Phanera_corymbosa
Fabaceae_Bauhinia_yunnanensis
Fabaceae_Bauhinia_aureifolia
Fabaceae_Phanera_vahlii
Fabaceae_Phanera_ornata
Fabaceae_Bauhinia_penicilliloba
Fabaceae_Bauhinia_longistipes
Fabaceae_Bauhinia_scandens
Fabaceae_Tylosema_humifusa
Fabaceae_Tylosema_argentea
Fabaceae_Tylosema_fassoglense
Fabaceae_Tylosema_esculentum
Fabaceae_Bauhinia_erythrantha
Fabaceae_Tylosema_fassoglensis
Fabaceae_Piliostigma_thonningii
Fabaceae_Brenierea_insignis
Fabaceae_Griffonia_physocarpa
Fabaceae_Adenolobus_pechuelii
Fabaceae_Adenolobus_garipensis
Fabaceae_Cercis_glabra
Fabaceae_Cercis_chinensis
Fabaceae_Cercis_racemosa
Fabaceae_Cercis_griffithii
Fabaceae_Cercis_siliquastrum
Fabaceae_Cercis_occidentalis
Fabaceae_Cercis_canadensis
Fabaceae_Cercis_chingii
Fabaceae_Cercis_chuniana
Fabaceae_Gigasiphon_macrosiphon
Fabaceae_Dialium_guianense

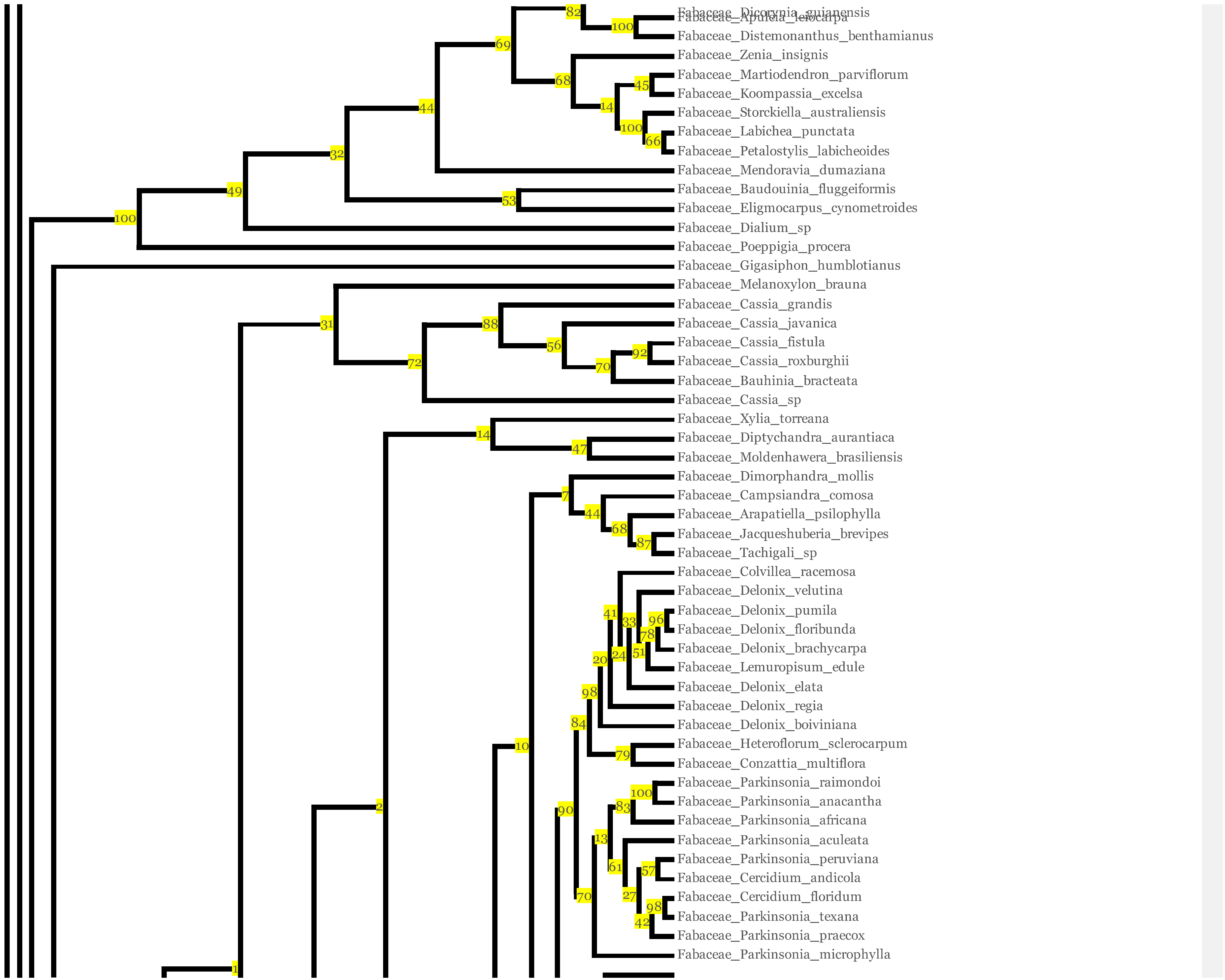
Fabaceae_Dicorynia_guianensis
Fabaceae_Apuleia_leiocarpa
Fabaceae_Distemonanthus_benthamianus
Fabaceae_Zenia_insignis
Fabaceae_Martiodendron_parviflorum
Fabaceae_Koompassia_excelsa
Fabaceae_Storckiella_australiensis
Fabaceae_Labichea_punctata
Fabaceae_Petalostylis_labicheoides
Fabaceae_Mendoravia_dumaziana
Fabaceae_Baudouinia_fluggeiformis
Fabaceae_Eligmocarpus_cynometroides
Fabaceae_Dialium_sp
Fabaceae_Poeppigia_procera
Fabaceae_Gigasiphon_humblotianus
Fabaceae_Melanoxylon_brauna
Fabaceae_Cassia_grandis
Fabaceae_Cassia_javanica
Fabaceae_Cassia_fistula
Fabaceae_Cassia_roxburghii
Fabaceae_Bauhinia_bracteata
Fabaceae_Cassia_sp
Fabaceae_Xylia_torreana
Fabaceae_Diptychandra_aurantiaca
Fabaceae_Moldenhawera_brasiliensis
Fabaceae_Dimorphandra_mollis
Fabaceae_Campsiandra_comosa
Fabaceae_Arapatiella_psilophylla
Fabaceae_Jacqueshuberia_brevipes
Fabaceae_Tachigali_sp
Fabaceae_Colvillea_racemosa
Fabaceae_Delonix_velutina
Fabaceae_Delonix_pumila
Fabaceae_Delonix_floribunda
Fabaceae_Delonix_brachycarpa
Fabaceae_Lemuropisum_edule
Fabaceae_Delonix_elata
Fabaceae_Delonix_regia
Fabaceae_Delonix_boiviniana
Fabaceae_Heteroflorum_sclerocarpum
Fabaceae_Conzattia_multiflora
Fabaceae_Parkinsonia_raimondoi
Fabaceae_Parkinsonia_anacantha
Fabaceae_Parkinsonia_africana
Fabaceae_Parkinsonia_aculeata
Fabaceae_Parkinsonia_peruviana
Fabaceae_Cercidium_andicola
Fabaceae_Cercidium_floridum
Fabaceae_Parkinsonia_texana
Fabaceae_Parkinsonia_praecox
Fabaceae_Parkinsonia_microphylla

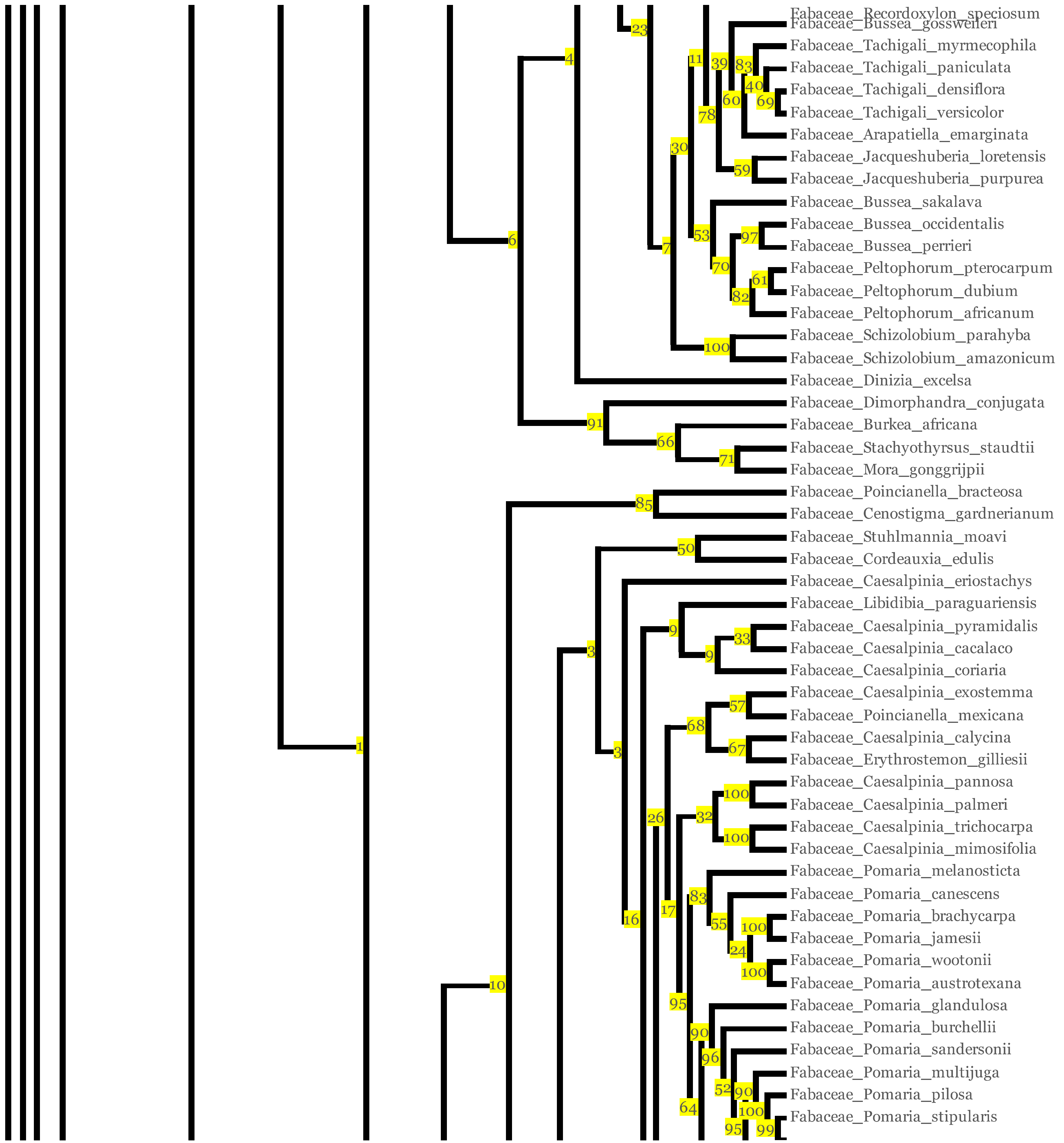
Fabaceae_Recordoxylon_speciosum
Fabaceae_Bussea_gossweileri
Fabaceae_Tachigali_myrmecophila
Fabaceae_Tachigali_paniculata
Fabaceae_Tachigali_densiflora
Fabaceae_Tachigali_versicolor
Fabaceae_Arapatiella_emarginata
Fabaceae_Jacqueshuberia_loretensis
Fabaceae_Jacqueshuberia_purpurea
Fabaceae_Bussea_sakalava
Fabaceae_Bussea_occidentalis
Fabaceae_Bussea_perrieri
Fabaceae_Peltophorum_pterocarpum
Fabaceae_Peltophorum_dubium
Fabaceae_Peltophorum_africanum
Fabaceae_Schizolobium_parahyba
Fabaceae_Schizolobium_amazonicum
Fabaceae_Dinizia_excelsa
Fabaceae_Dimorphandra_conjugata
Fabaceae_Burkea_africana
Fabaceae_Stachyothyrsus_staudtii
Fabaceae_Mora_gonggrijpii
Fabaceae_Poincianella_bracteosa
Fabaceae_Cenostigma_gardnerianum
Fabaceae_Stuhlmannia_moavi
Fabaceae_Cordeauxia_edulis
Fabaceae_Caesalpinia_eriostachys
Fabaceae_Libidibia_paraguariensis
Fabaceae_Caesalpinia_pyramidalis
Fabaceae_Caesalpinia_cacalaco
Fabaceae_Caesalpinia_coriaria
Fabaceae_Caesalpinia_exostemma
Fabaceae_Poincianella_mexicana
Fabaceae_Caesalpinia_calycina
Fabaceae_Erythrostemon_gilliesii
Fabaceae_Caesalpinia_pannosa
Fabaceae_Caesalpinia_palmeri
Fabaceae_Caesalpinia_trichocarpa
Fabaceae_Caesalpinia_mimosifolia
Fabaceae_Pomaria_melanosticta
Fabaceae_Pomaria_canescens
Fabaceae_Pomaria_brachycarpa
Fabaceae_Pomaria_jamesii
Fabaceae_Pomaria_wootonii
Fabaceae_Pomaria_austrotexana
Fabaceae_Pomaria_glandulosa
Fabaceae_Pomaria_burchellii
Fabaceae_Pomaria_sandersonii
Fabaceae_Pomaria_multijuga
Fabaceae_Pomaria_pilosa
Fabaceae_Pomaria_stipularis

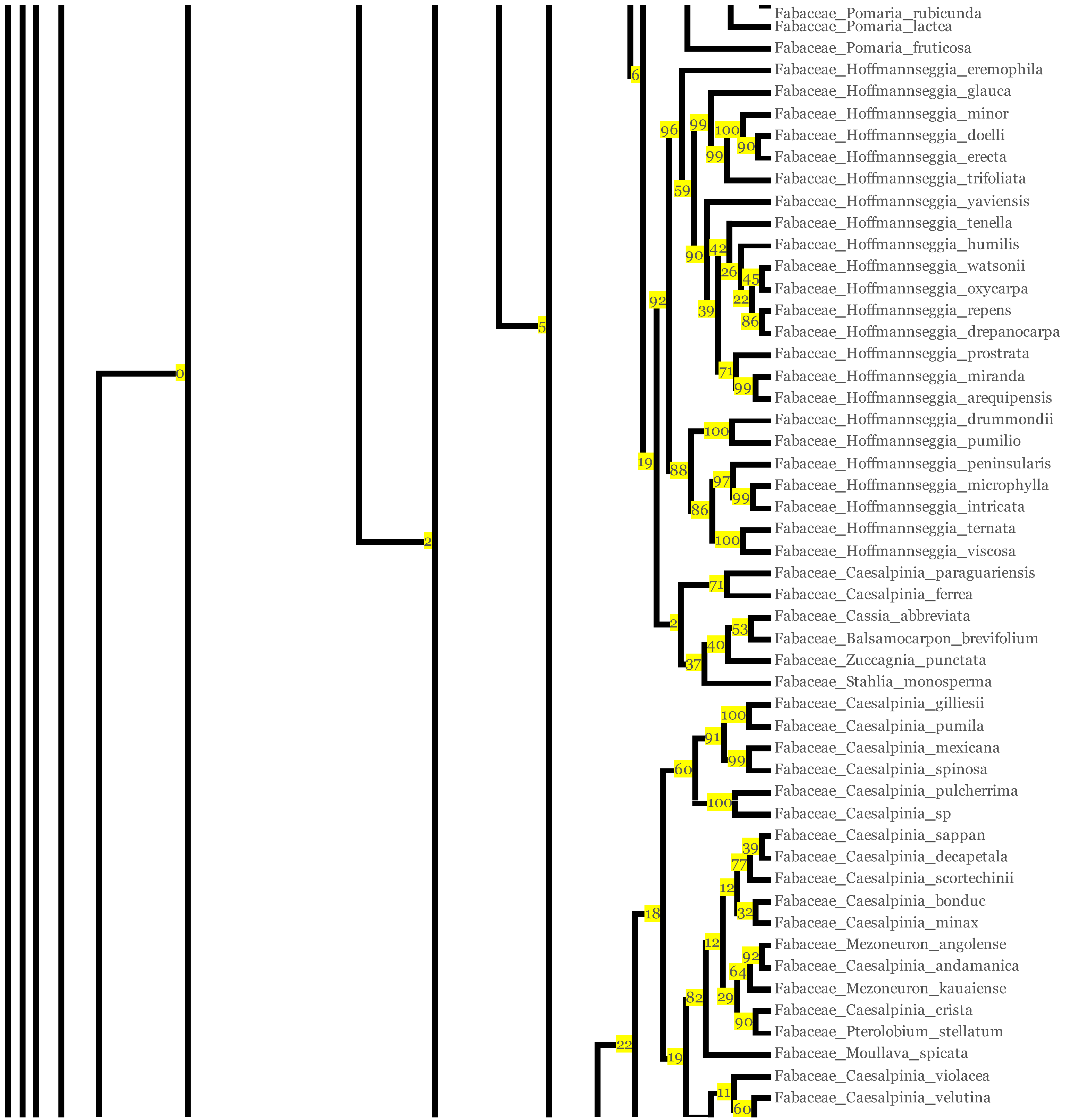
Fabaceae_Pomaria_rubicunda
Fabaceae_Pomaria_lactea
Fabaceae_Pomaria_fruticosa
Fabaceae_Hoffmannseggia_eremophila
Fabaceae_Hoffmannseggia_glauca
Fabaceae_Hoffmannseggia_minor
Fabaceae_Hoffmannseggia_doelli
Fabaceae_Hoffmannseggia_erecta
Fabaceae_Hoffmannseggia_trifoliata
Fabaceae_Hoffmannseggia_yaviensis
Fabaceae_Hoffmannseggia_tenella
Fabaceae_Hoffmannseggia_humilis
Fabaceae_Hoffmannseggia_watsonii
Fabaceae_Hoffmannseggia_oxycarpa
Fabaceae_Hoffmannseggia_repens
Fabaceae_Hoffmannseggia_drepanocarpa
Fabaceae_Hoffmannseggia_prostrata
Fabaceae_Hoffmannseggia_miranda
Fabaceae_Hoffmannseggia_arequipensis
Fabaceae_Hoffmannseggia_drummondii
Fabaceae_Hoffmannseggia_pumilio
Fabaceae_Hoffmannseggia_peninsularis
Fabaceae_Hoffmannseggia_microphylla
Fabaceae_Hoffmannseggia_intricata
Fabaceae_Hoffmannseggia_ternata
Fabaceae_Hoffmannseggia_viscosa
Fabaceae_Caesalpinia_paraguariensis
Fabaceae_Caesalpinia_ferrea
Fabaceae_Cassia_abbreviata
Fabaceae_Balsamocarpon_brevifolium
Fabaceae_Zuccagnia_punctata
Fabaceae_Stahlia_monosperma
Fabaceae_Caesalpinia_gilliesii
Fabaceae_Caesalpinia_pumila
Fabaceae_Caesalpinia_mexicana
Fabaceae_Caesalpinia_spinosa
Fabaceae_Caesalpinia_pulcherrima
Fabaceae_Caesalpinia_sp
Fabaceae_Caesalpinia_sappan
Fabaceae_Caesalpinia_decapetala
Fabaceae_Caesalpinia_scortechinii
Fabaceae_Caesalpinia_bonduc
Fabaceae_Caesalpinia_minax
Fabaceae_Mezoneuron_angolense
Fabaceae_Caesalpinia_andamanica
Fabaceae_Mezoneuron_kauaiense
Fabaceae_Caesalpinia_crista
Fabaceae_Pterolobium_stellatum
Fabaceae_Moullava_spicata
Fabaceae_Caesalpinia_violacea
Fabaceae_Caesalpinia_velutina

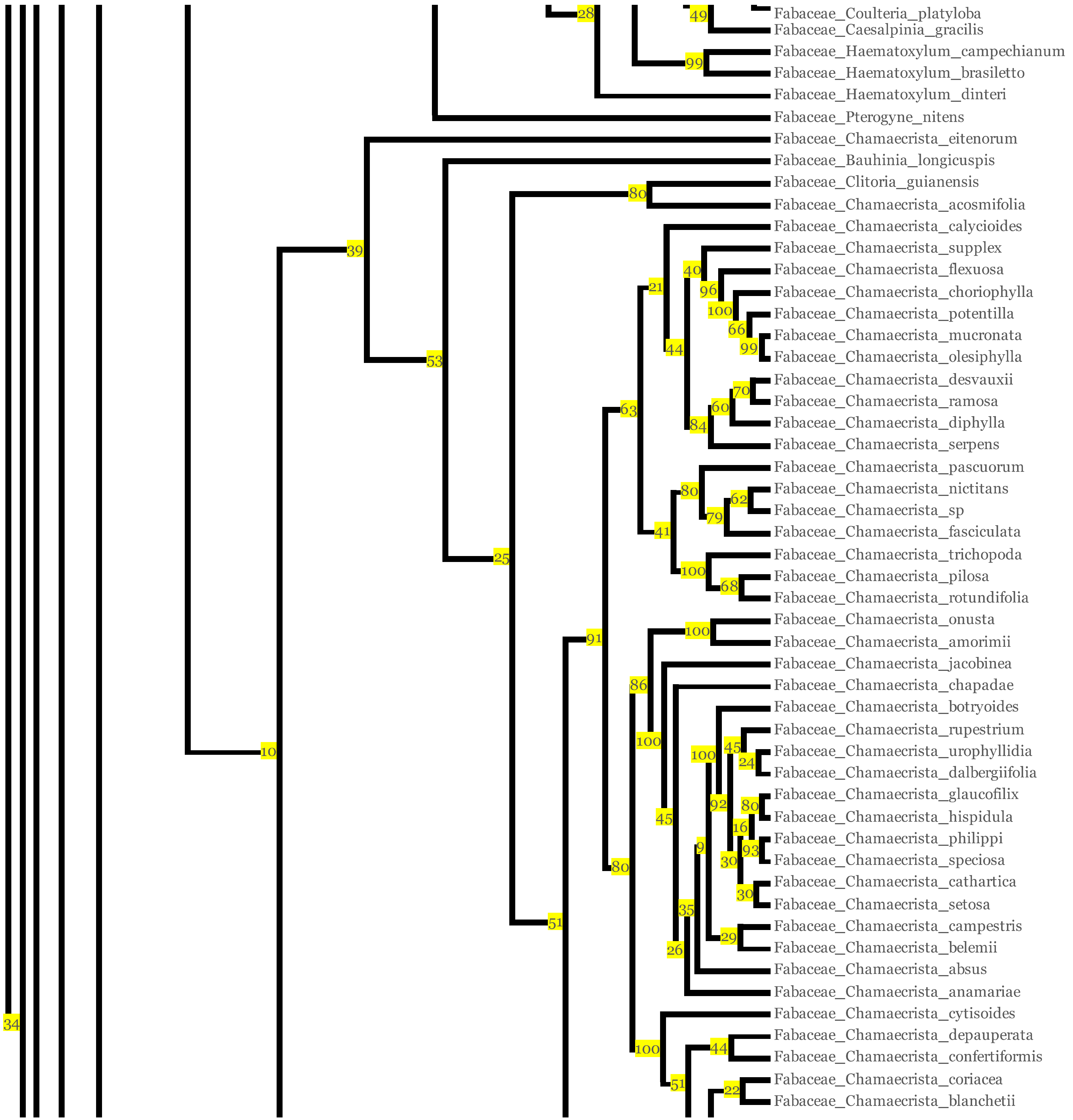
Fabaceae_Coulteria_platyloba
Fabaceae_Caesalpinia_gracilis
Fabaceae_Haematoxylum_campechianum
Fabaceae_Haematoxylum_brasiletto
Fabaceae_Haematoxylum_dinteri
Fabaceae_Pterogyne_nitens
Fabaceae_Chamaecrista_eitenorum
Fabaceae_Bauhinia_longicuspis
Fabaceae_Clitoria_guianensis
Fabaceae_Chamaecrista_acosmifolia
Fabaceae_Chamaecrista_calycioides
Fabaceae_Chamaecrista_supplex
Fabaceae_Chamaecrista_flexuosa
Fabaceae_Chamaecrista_choriophylla
Fabaceae_Chamaecrista_potentilla
Fabaceae_Chamaecrista_mucronata
Fabaceae_Chamaecrista_olesiphylla
Fabaceae_Chamaecrista_desvauxii
Fabaceae_Chamaecrista_ramosa
Fabaceae_Chamaecrista_diphylla
Fabaceae_Chamaecrista_serpens
Fabaceae_Chamaecrista_pascuorum
Fabaceae_Chamaecrista_nictitans
Fabaceae_Chamaecrista_sp
Fabaceae_Chamaecrista_fasciculata
Fabaceae_Chamaecrista_trichopoda
Fabaceae_Chamaecrista_pilosa
Fabaceae_Chamaecrista_rotundifolia
Fabaceae_Chamaecrista_onusta
Fabaceae_Chamaecrista_amorimii
Fabaceae_Chamaecrista_jacobinea
Fabaceae_Chamaecrista_chapadae
Fabaceae_Chamaecrista_botryoides
Fabaceae_Chamaecrista_rupestrium
Fabaceae_Chamaecrista_urophyllidia
Fabaceae_Chamaecrista_dalbergiifolia
Fabaceae_Chamaecrista_glaucofilix
Fabaceae_Chamaecrista_hispidula
Fabaceae_Chamaecrista_philippi
Fabaceae_Chamaecrista_speciosa
Fabaceae_Chamaecrista_cathartica
Fabaceae_Chamaecrista_setosa
Fabaceae_Chamaecrista_campestris
Fabaceae_Chamaecrista_belemii
Fabaceae_Chamaecrista_absus
Fabaceae_Chamaecrista_anamariae
Fabaceae_Chamaecrista_cytisoides
Fabaceae_Chamaecrista_depauperata
Fabaceae_Chamaecrista_confertiformis
Fabaceae_Chamaecrista_coriacea
Fabaceae_Chamaecrista_blanchetii

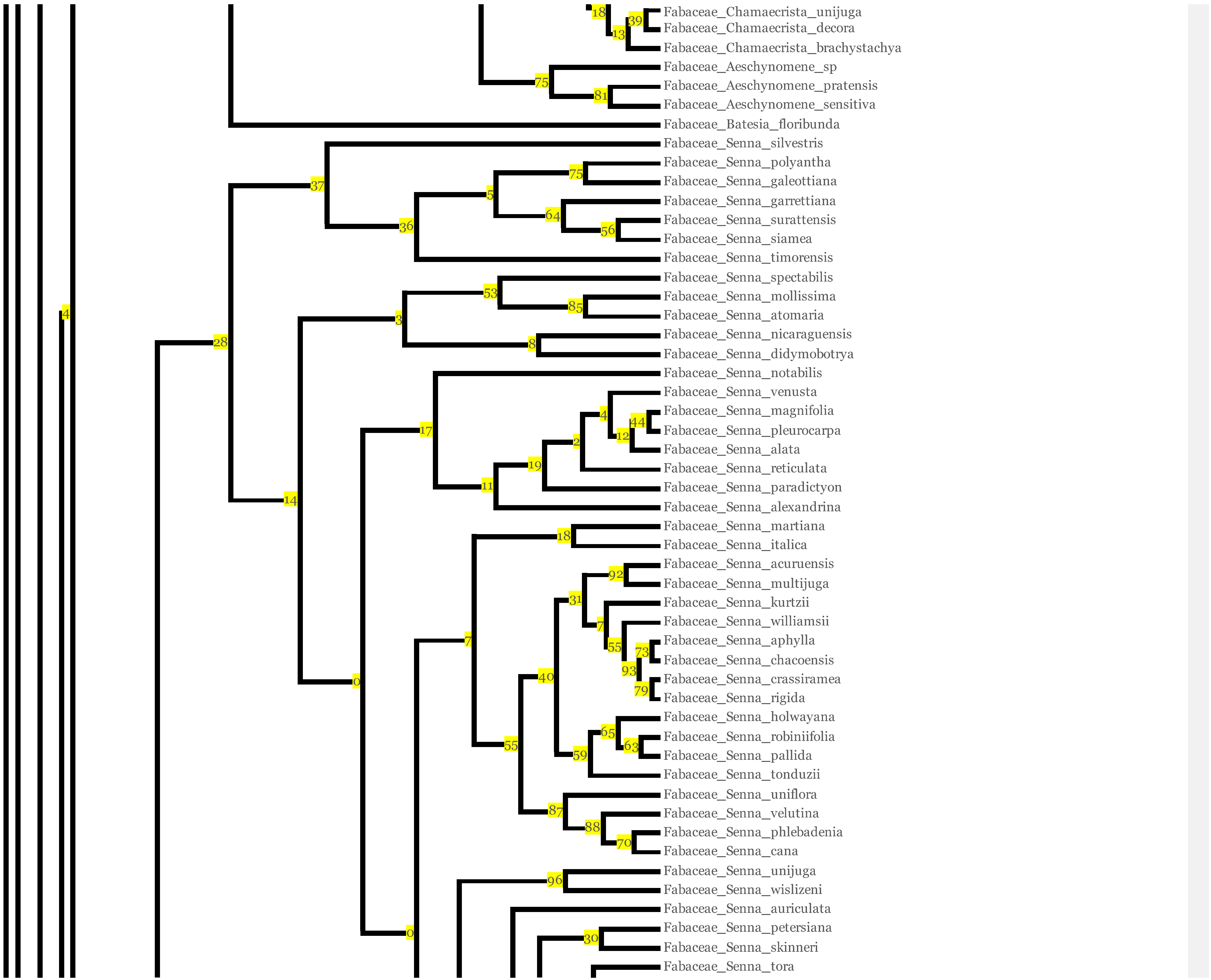
Fabaceae_Chamaecrista_unijuga
Fabaceae_Chamaecrista_decora
Fabaceae_Chamaecrista_brachystachya
Fabaceae_Aeschynomene_sp
Fabaceae_Aeschynomene_pratensis
Fabaceae_Aeschynomene_sensitiva
Fabaceae_Batesia_floribunda
Fabaceae_Senna_silvestris
Fabaceae_Senna_polyantha
Fabaceae_Senna_galeottiana
Fabaceae_Senna_garrettiana
Fabaceae_Senna_surattensis
Fabaceae_Senna_siamea
Fabaceae_Senna_timorensis
Fabaceae_Senna_spectabilis
Fabaceae_Senna_mollissima
Fabaceae_Senna_atomaria
Fabaceae_Senna_nicaraguensis
Fabaceae_Senna_didymobotrya
Fabaceae_Senna_notabilis
Fabaceae_Senna_venusta
Fabaceae_Senna_magnifolia
Fabaceae_Senna_pleurocarpa
Fabaceae_Senna_alata
Fabaceae_Senna_reticulata
Fabaceae_Senna_paradictyon
Fabaceae_Senna_alexandrina
Fabaceae_Senna_martiana
Fabaceae_Senna_italica
Fabaceae_Senna_acuruensis
Fabaceae_Senna_multijuga
Fabaceae_Senna_kurtzii
Fabaceae_Senna_williamsii
Fabaceae_Senna_aphylla
Fabaceae_Senna_chacoensis
Fabaceae_Senna_crassiramea
Fabaceae_Senna_rigida
Fabaceae_Senna_holwayana
Fabaceae_Senna_robiniifolia
Fabaceae_Senna_pallida
Fabaceae_Senna_tonduzii
Fabaceae_Senna_uniflora
Fabaceae_Senna_velutina
Fabaceae_Senna_phlebadenia
Fabaceae_Senna_cana
Fabaceae_Senna_unijuga
Fabaceae_Senna_wislizeni
Fabaceae_Senna_auriculata
Fabaceae_Senna_petersiana
Fabaceae_Senna_skinneri
Fabaceae_Senna_tora

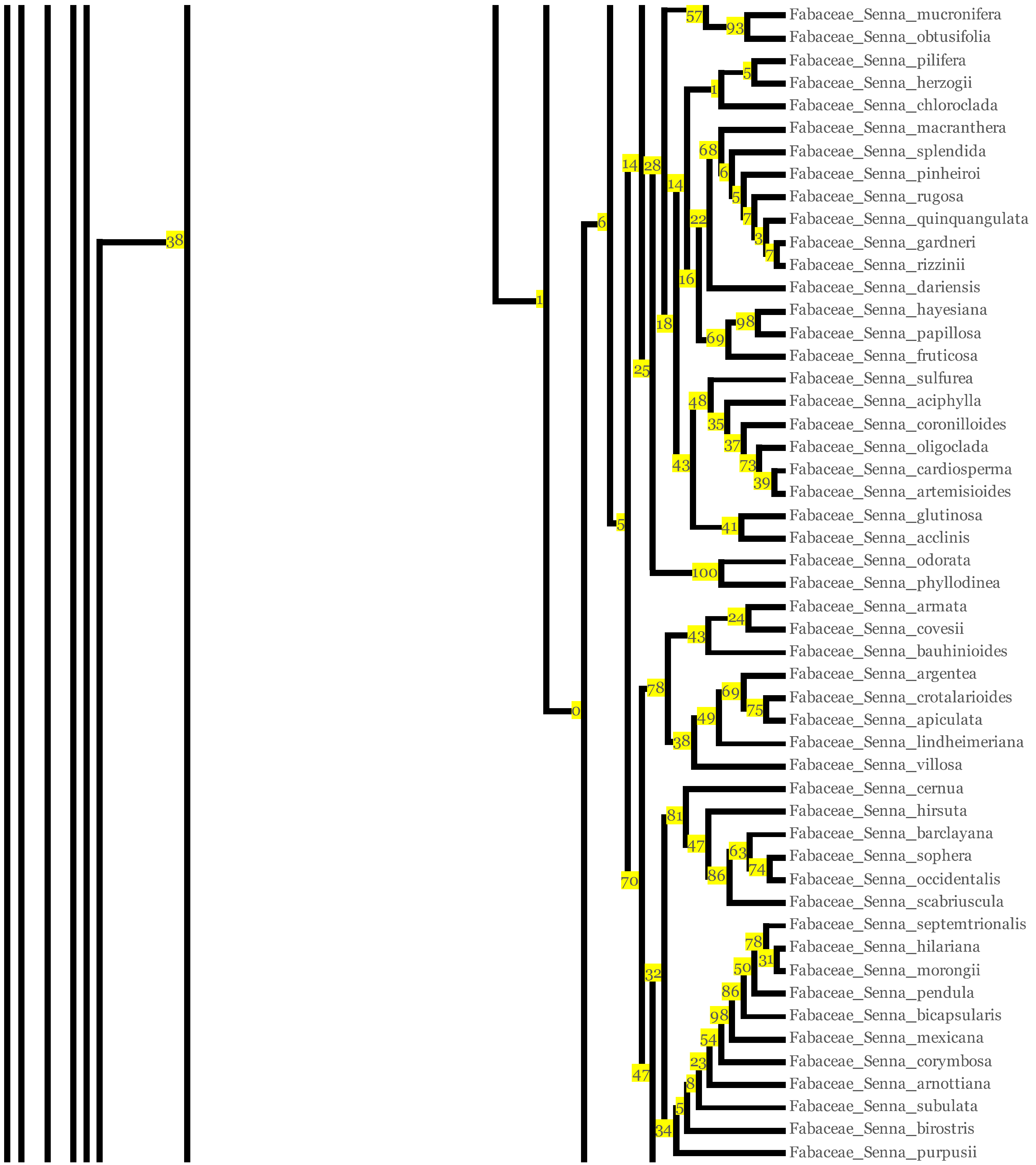
Fabaceae_Senna_mucronifera
Fabaceae_Senna_obtusifolia
Fabaceae_Senna_pilifera
Fabaceae_Senna_herzogii
Fabaceae_Senna_chloroclada
Fabaceae_Senna_macranthera
Fabaceae_Senna_splendida
Fabaceae_Senna_pinheiroi
Fabaceae_Senna_rugosa
Fabaceae_Senna_quinquangulata
Fabaceae_Senna_gardneri
Fabaceae_Senna_rizzinii
Fabaceae_Senna_dariensis
Fabaceae_Senna_hayesiana
Fabaceae_Senna_papillosa
Fabaceae_Senna_fruticosa
Fabaceae_Senna_sulfurea
Fabaceae_Senna_aciphylla
Fabaceae_Senna_coronilloides
Fabaceae_Senna_oligoclada
Fabaceae_Senna_cardiosperma
Fabaceae_Senna_artemisioides
Fabaceae_Senna_glutinosa
Fabaceae_Senna_acclinis
Fabaceae_Senna_odorata
Fabaceae_Senna_phyllodinea
Fabaceae_Senna_armata
Fabaceae_Senna_covesii
Fabaceae_Senna_bauhinioides
Fabaceae_Senna_argentea
Fabaceae_Senna_crotalarioides
Fabaceae_Senna_apiculata
Fabaceae_Senna_lindheimeriana
Fabaceae_Senna_villosa
Fabaceae_Senna_cernua
Fabaceae_Senna_hirsuta
Fabaceae_Senna_barclayana
Fabaceae_Senna_sophera
Fabaceae_Senna_occidentalis
Fabaceae_Senna_scabriuscula
Fabaceae_Senna_septemtrionalis
Fabaceae_Senna_hilariana
Fabaceae_Senna_morongii
Fabaceae_Senna_pendula
Fabaceae_Senna_bicapsularis
Fabaceae_Senna_mexicana
Fabaceae_Senna_corymbosa
Fabaceae_Senna_arnottiana
Fabaceae_Senna_subulata
Fabaceae_Senna_birostris
Fabaceae_Senna_purpusii

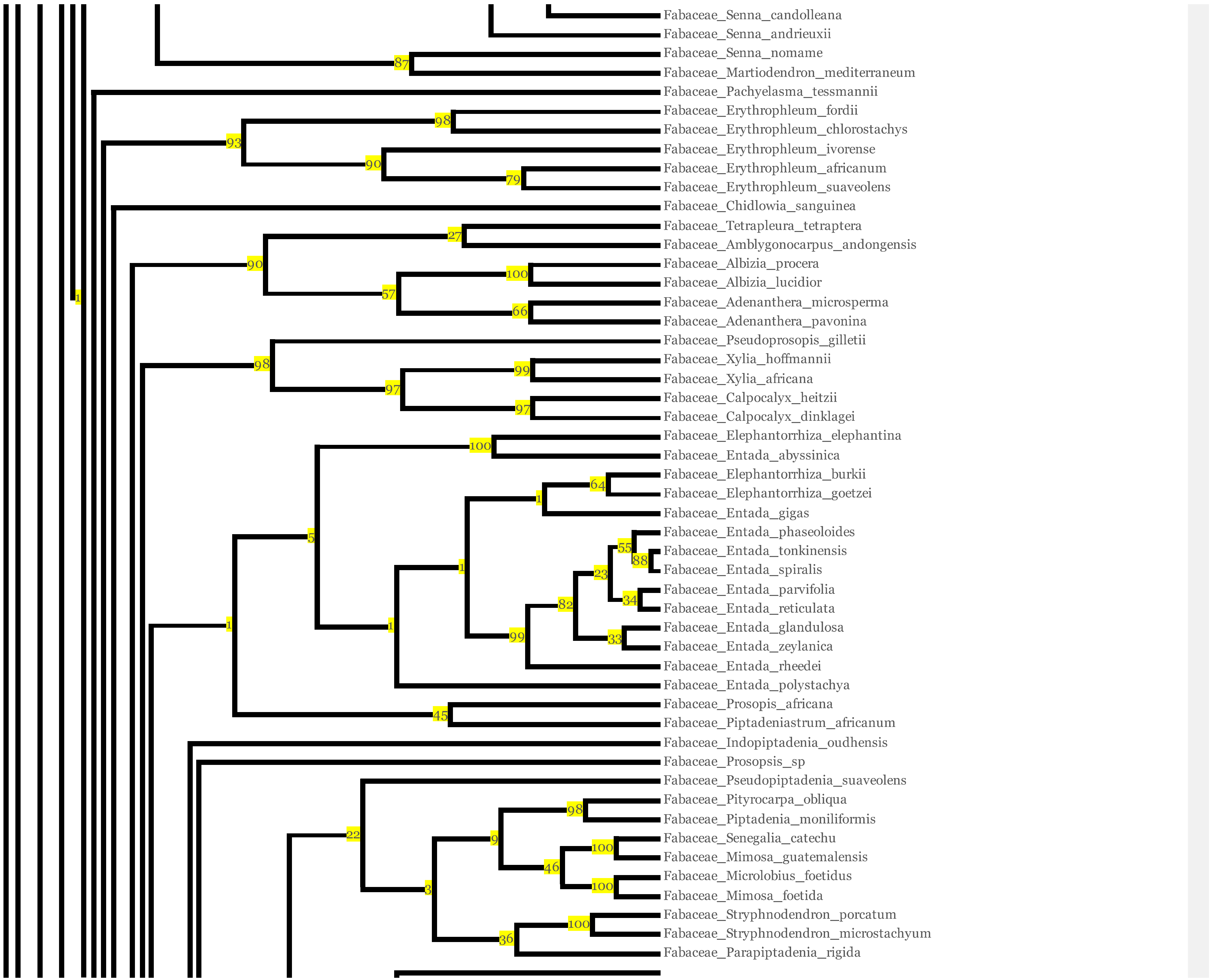
Fabaceae_Senna_candolleana
Fabaceae_Senna_andrieuxii
Fabaceae_Senna_nomame
Fabaceae_Martiodendron_mediterraneum
Fabaceae_Pachyelasma_tessmannii
Fabaceae_Erythrophleum_fordii
Fabaceae_Erythrophleum_chlorostachys
Fabaceae_Erythrophleum_ivorense
Fabaceae_Erythrophleum_africanum
Fabaceae_Erythrophleum_suaveolens
Fabaceae_Chidlowia_sanguinea
Fabaceae_Tetrapleura_tetraptera
Fabaceae_Amblygonocarpus_andongensis
Fabaceae_Albizia_procera
Fabaceae_Albizia_lucidior
Fabaceae_Adenanthera_microsperma
Fabaceae_Adenanthera_pavonina
Fabaceae_Pseudoprosopis_gilletii
Fabaceae_Xylia_hoffmannii
Fabaceae_Xylia_africana
Fabaceae_Calpocalyx_heitzii
Fabaceae_Calpocalyx_dinklagei
Fabaceae_Elephantorrhiza_elephantina
Fabaceae_Entada_abyssinica
Fabaceae_Elephantorrhiza_burkii
Fabaceae_Elephantorrhiza_goetzei
Fabaceae_Entada_gigas
Fabaceae_Entada_phaseoloides
Fabaceae_Entada_tonkinensis
Fabaceae_Entada_spiralis
Fabaceae_Entada_parvifolia
Fabaceae_Entada_reticulata
Fabaceae_Entada_glandulosa
Fabaceae_Entada_zeylanica
Fabaceae_Entada_rheedei
Fabaceae_Entada_polystachya
Fabaceae_Prosopis_africana
Fabaceae_Piptadeniastrum_africanum
Fabaceae_Indopiptadenia_oudhensis
Fabaceae_Prosopsis_sp
Fabaceae_Pseudopiptadenia_suaveolens
Fabaceae_Pityrocarpa_obliqua
Fabaceae_Piptadenia_moniliformis
Fabaceae_Senegalia_catechu
Fabaceae_Mimosa_guatemalensis
Fabaceae_Microlobius_foetidus
Fabaceae_Mimosa_foetida
Fabaceae_Stryphnodendron_porcatum
Fabaceae_Stryphnodendron_microstachyum
Fabaceae_Parapiptadenia_rigida

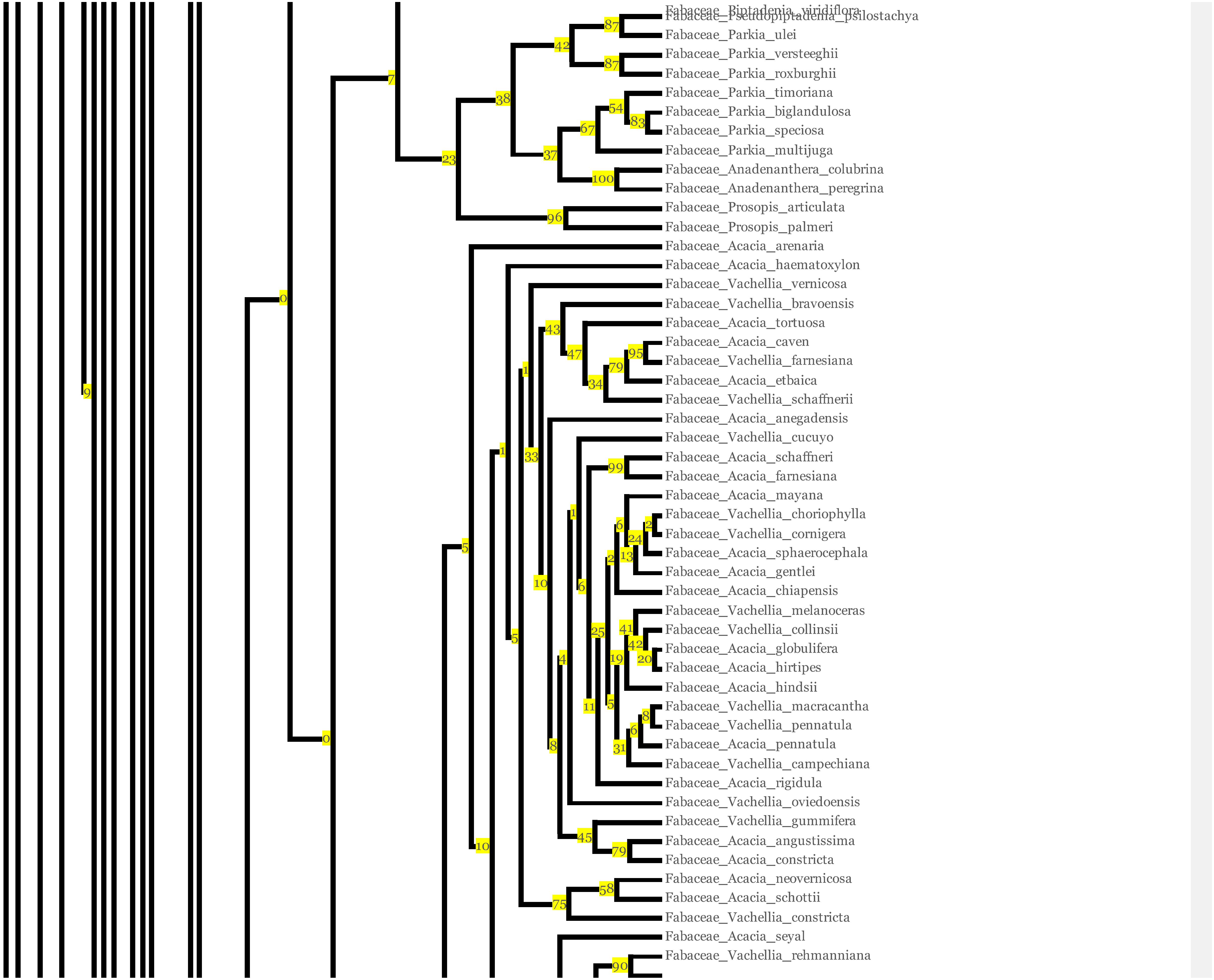
Fabaceae_Piptadenia_viridiflora
Fabaceae_Pseudopiptadenia_psilostachya
Fabaceae_Parkia_ulei
Fabaceae_Parkia_versteeghii
Fabaceae_Parkia_roxburghii
Fabaceae_Parkia_timoriana
Fabaceae_Parkia_biglandulosa
Fabaceae_Parkia_speciosa
Fabaceae_Parkia_multijuga
Fabaceae_Anadenanthera_colubrina
Fabaceae_Anadenanthera_peregrina
Fabaceae_Prosopis_articulata
Fabaceae_Prosopis_palmeri
Fabaceae_Acacia_arenaria
Fabaceae_Acacia_haematoxylon
Fabaceae_Vachellia_vernicosa
Fabaceae_Vachellia_bravoensis
Fabaceae_Acacia_tortuosa
Fabaceae_Acacia_caven
Fabaceae_Vachellia_farnesiana
Fabaceae_Acacia_etbaica
Fabaceae_Vachellia_schaffnerii
Fabaceae_Acacia_anegadensis
Fabaceae_Vachellia_cucuyo
Fabaceae_Acacia_schaffneri
Fabaceae_Acacia_farnesiana
Fabaceae_Acacia_mayana
Fabaceae_Vachellia_choriophylla
Fabaceae_Vachellia_cornigera
Fabaceae_Acacia_sphaerocephala
Fabaceae_Acacia_gentlei
Fabaceae_Acacia_chiapensis
Fabaceae_Vachellia_melanoceras
Fabaceae_Vachellia_collinsii
Fabaceae_Acacia_globulifera
Fabaceae_Acacia_hirtipes
Fabaceae_Acacia_hindsii
Fabaceae_Vachellia_macracantha
Fabaceae_Vachellia_pennatula
Fabaceae_Acacia_pennatula
Fabaceae_Vachellia_campechiana
Fabaceae_Acacia_rigidula
Fabaceae_Vachellia_oviedoensis
Fabaceae_Vachellia_gummifera
Fabaceae_Acacia_angustissima
Fabaceae_Acacia_constricta
Fabaceae_Acacia_neovernicosa
Fabaceae_Acacia_schottii
Fabaceae_Vachellia_constricta
Fabaceae_Acacia_seyal
Fabaceae_Vachellia_rehmanniana

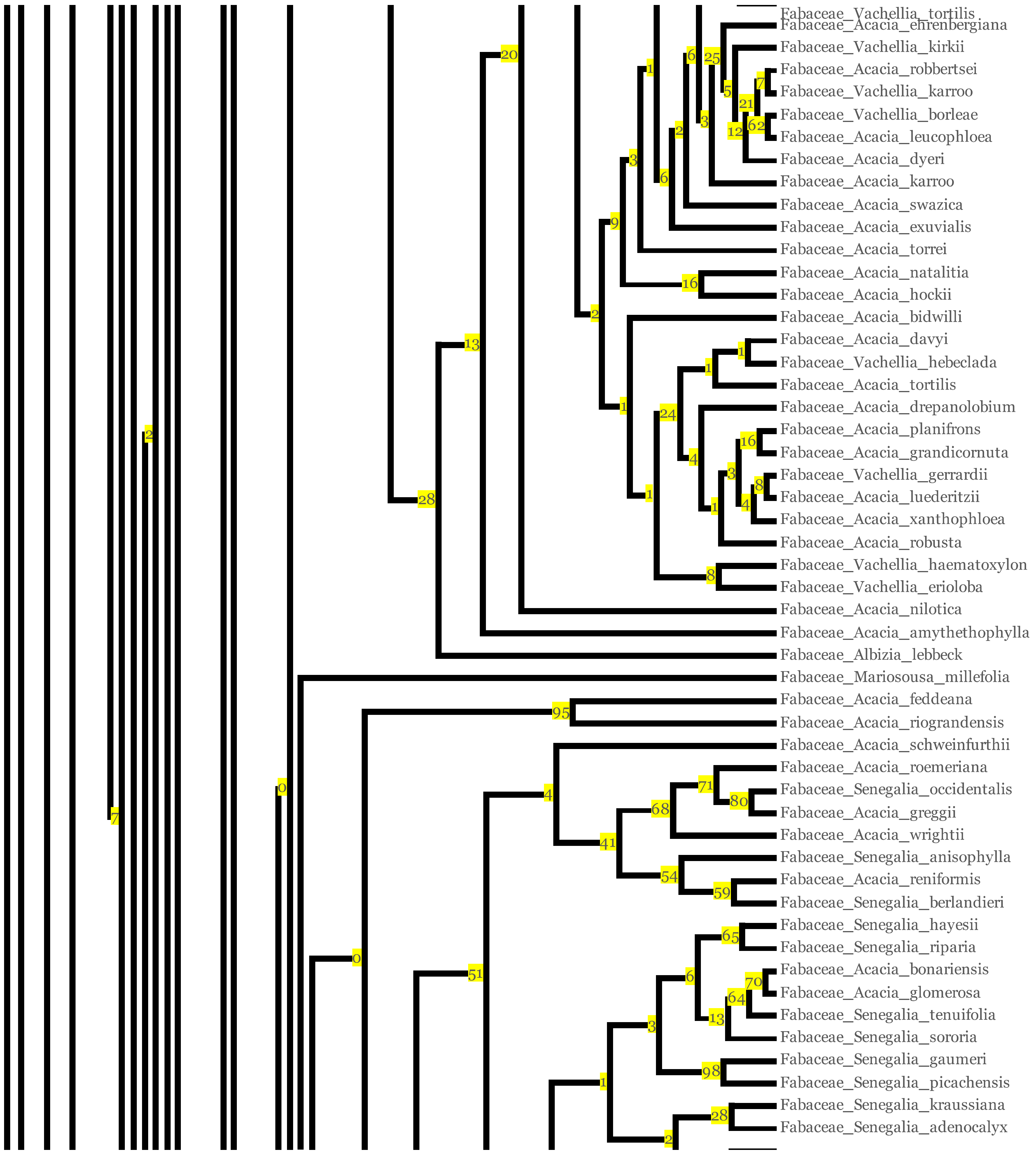
Fabaceae_Vachellia_tortilis
Fabaceae_Acacia_ehrenbergiana
Fabaceae_Vachellia_kirkii
Fabaceae_Acacia_robbertsei
Fabaceae_Vachellia_karroo
Fabaceae_Vachellia_borleae
Fabaceae_Acacia_leucophloea
Fabaceae_Acacia_dyeri
Fabaceae_Acacia_karroo
Fabaceae_Acacia_swazica
Fabaceae_Acacia_exuvialis
Fabaceae_Acacia_torrei
Fabaceae_Acacia_natalitia
Fabaceae_Acacia_hockii
Fabaceae_Acacia_bidwilli
Fabaceae_Acacia_davyi
Fabaceae_Vachellia_hebeclada
Fabaceae_Acacia_tortilis
Fabaceae_Acacia_drepanolobium
Fabaceae_Acacia_planifrons
Fabaceae_Acacia_grandicornuta
Fabaceae_Vachellia_gerrardii
Fabaceae_Acacia_luederitzii
Fabaceae_Acacia_xanthophloea
Fabaceae_Acacia_robusta
Fabaceae_Vachellia_haematoxylon
Fabaceae_Vachellia_erioloba
Fabaceae_Acacia_nilotica
Fabaceae_Acacia_amythethophylla
Fabaceae_Albizia_lebbeck
Fabaceae_Mariosousa_millefolia
Fabaceae_Acacia_feddeana
Fabaceae_Acacia_riograndensis
Fabaceae_Acacia_schweinfurthii
Fabaceae_Acacia_roemeriana
Fabaceae_Senegalia_occidentalis
Fabaceae_Acacia_greggii
Fabaceae_Acacia_wrightii
Fabaceae_Senegalia_anisophylla
Fabaceae_Acacia_reniformis
Fabaceae_Senegalia_berlandieri
Fabaceae_Senegalia_hayesii
Fabaceae_Senegalia_riparia
Fabaceae_Acacia_bonariensis
Fabaceae_Acacia_glomerosa
Fabaceae_Senegalia_tenuifolia
Fabaceae_Senegalia_sororia
Fabaceae_Senegalia_gaumeri
Fabaceae_Senegalia_picachensis
Fabaceae_Senegalia_kraussiana
Fabaceae_Senegalia_adenocalyx

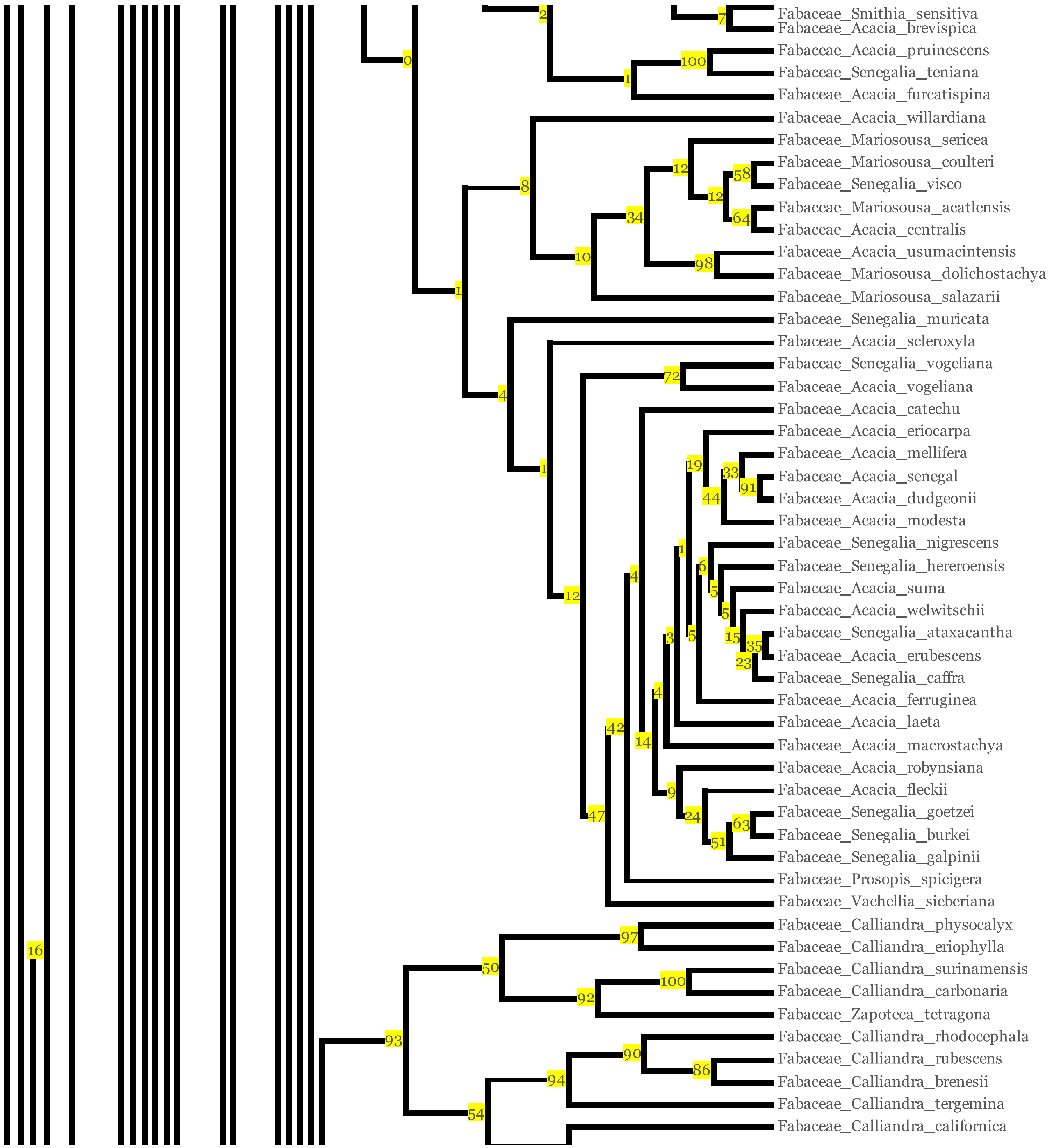
Fabaceae_Smithia_sensitiva
Fabaceae_Acacia_brevispica
Fabaceae_Acacia_pruinescens
Fabaceae_Senegalia_teniana
Fabaceae_Acacia_furcatispina
Fabaceae_Acacia_willardiana
Fabaceae_Mariosousa_sericea
Fabaceae_Mariosousa_coulteri
Fabaceae_Senegalia_visco
Fabaceae_Mariosousa_acatlensis
Fabaceae_Acacia_centralis
Fabaceae_Acacia_usumacintensis
Fabaceae_Mariosousa_dolichostachya
Fabaceae_Mariosousa_salazarii
Fabaceae_Senegalia_muricata
Fabaceae_Acacia_scleroxyla
Fabaceae_Senegalia_vogeliana
Fabaceae_Acacia_vogeliana
Fabaceae_Acacia_catechu
Fabaceae_Acacia_eriocarpa
Fabaceae_Acacia_mellifera
Fabaceae_Acacia_senegal
Fabaceae_Acacia_dudgeonii
Fabaceae_Acacia_modesta
Fabaceae_Senegalia_nigrescens
Fabaceae_Senegalia_hereroensis
Fabaceae_Acacia_suma
Fabaceae_Acacia_welwitschii
Fabaceae_Senegalia_ataxacantha
Fabaceae_Acacia_erubescens
Fabaceae_Senegalia_caffra
Fabaceae_Acacia_ferruginea
Fabaceae_Acacia_laeta
Fabaceae_Acacia_macrostachya
Fabaceae_Acacia_robynsiana
Fabaceae_Acacia_fleckii
Fabaceae_Senegalia_goetzei
Fabaceae_Senegalia_burkei
Fabaceae_Senegalia_galpinii
Fabaceae_Prosopis_spicigera
Fabaceae_Vachellia_sieberiana
Fabaceae_Calliandra_physocalyx
Fabaceae_Calliandra_eriophylla
Fabaceae_Calliandra_surinamensis
Fabaceae_Calliandra_carbonaria
Fabaceae_Zapoteca_tetragona
Fabaceae_Calliandra_rhodocephala
Fabaceae_Calliandra_rubescens
Fabaceae_Calliandra_brenesii
Fabaceae_Calliandra_tergemina
Fabaceae_Calliandra_californica

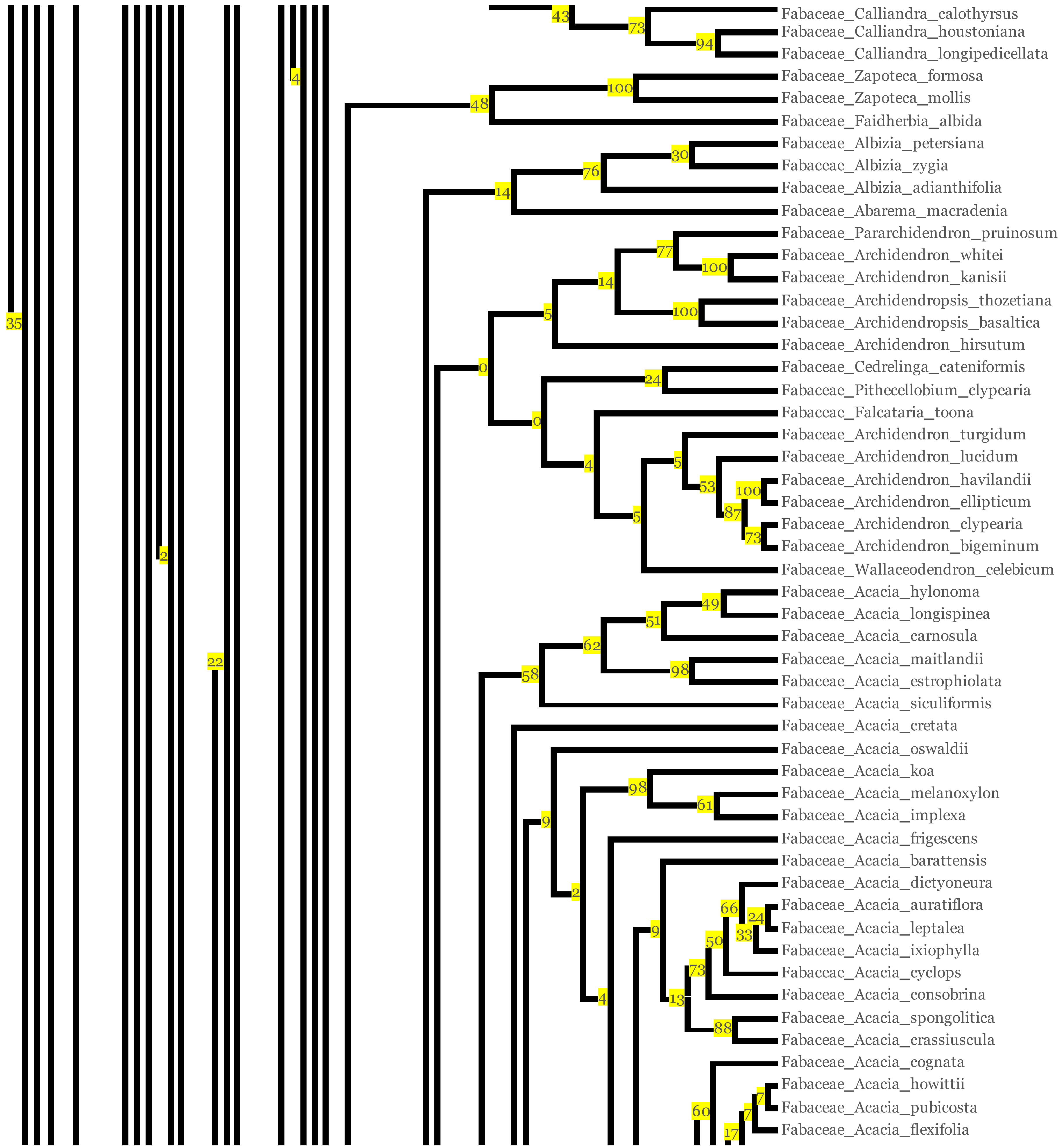
Fabaceae_Calliandra_calothyrsus
Fabaceae_Calliandra_houstoniana
Fabaceae_Calliandra_longipedicellata
Fabaceae_Zapoteca_formosa
Fabaceae_Zapoteca_mollis
Fabaceae_Faidherbia_albida
Fabaceae_Albizia_petersiana
Fabaceae_Albizia_zygia
Fabaceae_Albizia_adianthifolia
Fabaceae_Abarema_macradenia
Fabaceae_Pararchidendron_pruinosum
Fabaceae_Archidendron_whitei
Fabaceae_Archidendron_kanisii
Fabaceae_Archidendropsis_thozetiana
Fabaceae_Archidendropsis_basaltica
Fabaceae_Archidendron_hirsutum
Fabaceae_Cedrelinga_cateniformis
Fabaceae_Pithecellobium_clypearia
Fabaceae_Falcataria_toona
Fabaceae_Archidendron_turgidum
Fabaceae_Archidendron_lucidum
Fabaceae_Archidendron_havilandii
Fabaceae_Archidendron_ellipticum
Fabaceae_Archidendron_clypearia
Fabaceae_Archidendron_bigeminum
Fabaceae_Wallaceodendron_celebicum
Fabaceae_Acacia_hylonoma
Fabaceae_Acacia_longispinea
Fabaceae_Acacia_carnosula
Fabaceae_Acacia_maitlandii
Fabaceae_Acacia_estrophiolata
Fabaceae_Acacia_siculiformis
Fabaceae_Acacia_cretata
Fabaceae_Acacia_oswaldii
Fabaceae_Acacia_koa
Fabaceae_Acacia_melanoxylon
Fabaceae_Acacia_implexa
Fabaceae_Acacia_frigescens
Fabaceae_Acacia_barattensis
Fabaceae_Acacia_dictyoneura
Fabaceae_Acacia_auratiflora
Fabaceae_Acacia_leptalea
Fabaceae_Acacia_ixiophylla
Fabaceae_Acacia_cyclops
Fabaceae_Acacia_consobrina
Fabaceae_Acacia_spongolitica
Fabaceae_Acacia_crassiuscula
Fabaceae_Acacia_cognata
Fabaceae_Acacia_howittii
Fabaceae_Acacia_pubicosta
Fabaceae_Acacia_flexifolia

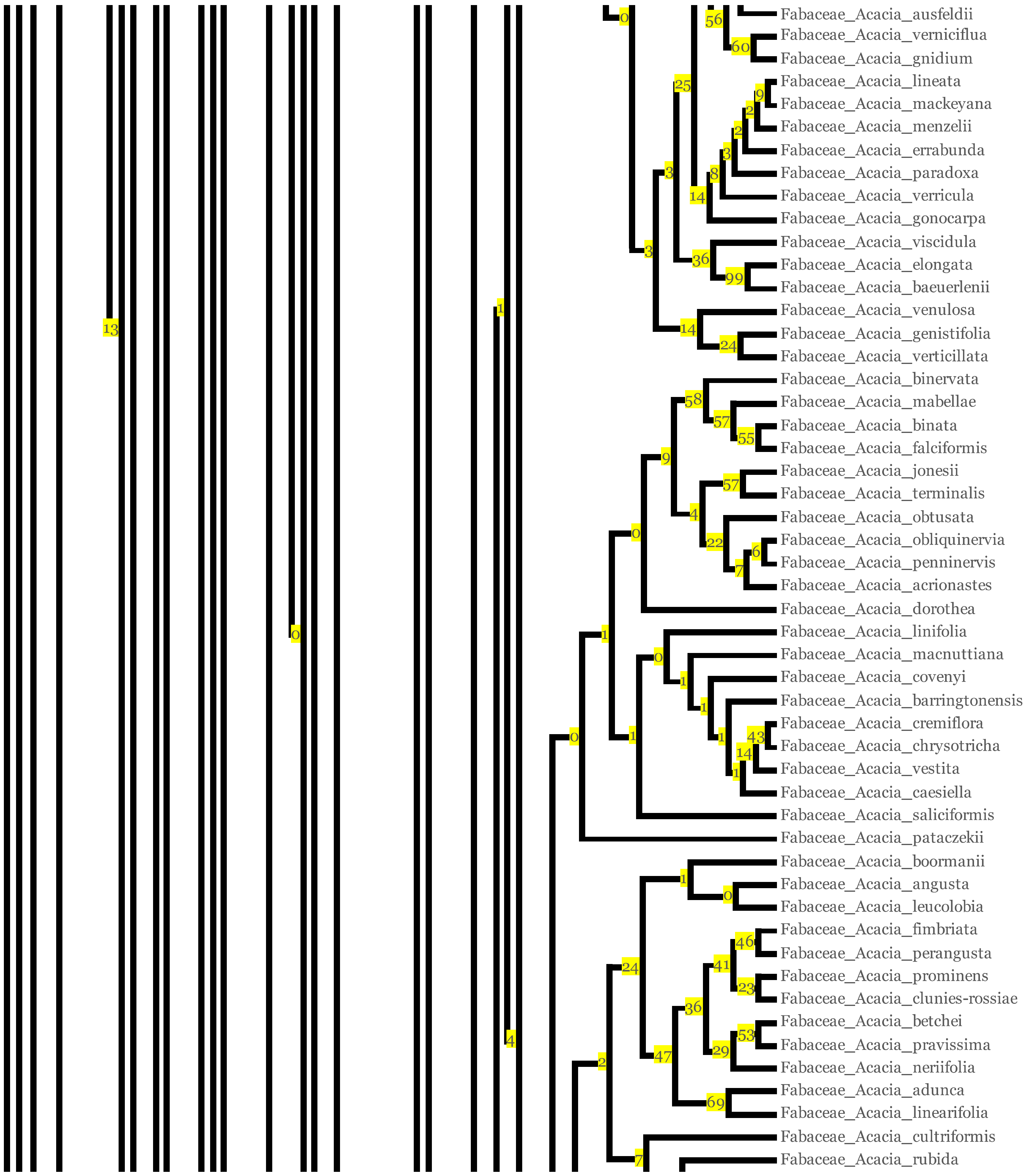
Fabaceae_Acacia_ausfeldii
Fabaceae_Acacia_verniciflua
Fabaceae_Acacia_gnidium
Fabaceae_Acacia_lineata
Fabaceae_Acacia_mackeyana
Fabaceae_Acacia_menzelii
Fabaceae_Acacia_errabunda
Fabaceae_Acacia_paradoxa
Fabaceae_Acacia_verricula
Fabaceae_Acacia_gonocarpa
Fabaceae_Acacia_viscidula
Fabaceae_Acacia_elongata
Fabaceae_Acacia_baeuerlenii
Fabaceae_Acacia_venulosa
Fabaceae_Acacia_genistifolia
Fabaceae_Acacia_verticillata
Fabaceae_Acacia_binervata
Fabaceae_Acacia_mabellae
Fabaceae_Acacia_binata
Fabaceae_Acacia_falciformis
Fabaceae_Acacia_jonesii
Fabaceae_Acacia_terminalis
Fabaceae_Acacia_obtusata
Fabaceae_Acacia_obliquinervia
Fabaceae_Acacia_penninervis
Fabaceae_Acacia_acrionastes
Fabaceae_Acacia_dorothea
Fabaceae_Acacia_linifolia
Fabaceae_Acacia_macnuttiana
Fabaceae_Acacia_covenyi
Fabaceae_Acacia_barringtonensis
Fabaceae_Acacia_cremiflora
Fabaceae_Acacia_chrysotricha
Fabaceae_Acacia_vestita
Fabaceae_Acacia_caesiella
Fabaceae_Acacia_saliciformis
Fabaceae_Acacia_pataczekii
Fabaceae_Acacia_boormanii
Fabaceae_Acacia_angusta
Fabaceae_Acacia_leucolobia
Fabaceae_Acacia_fimbriata
Fabaceae_Acacia_perangusta
Fabaceae_Acacia_prominens
Fabaceae_Acacia_clunies-rossiae
Fabaceae_Acacia_betchei
Fabaceae_Acacia_pravissima
Fabaceae_Acacia_neriifolia
Fabaceae_Acacia_adunca
Fabaceae_Acacia_linearifolia
Fabaceae_Acacia_cultriformis
Fabaceae_Acacia_rubida

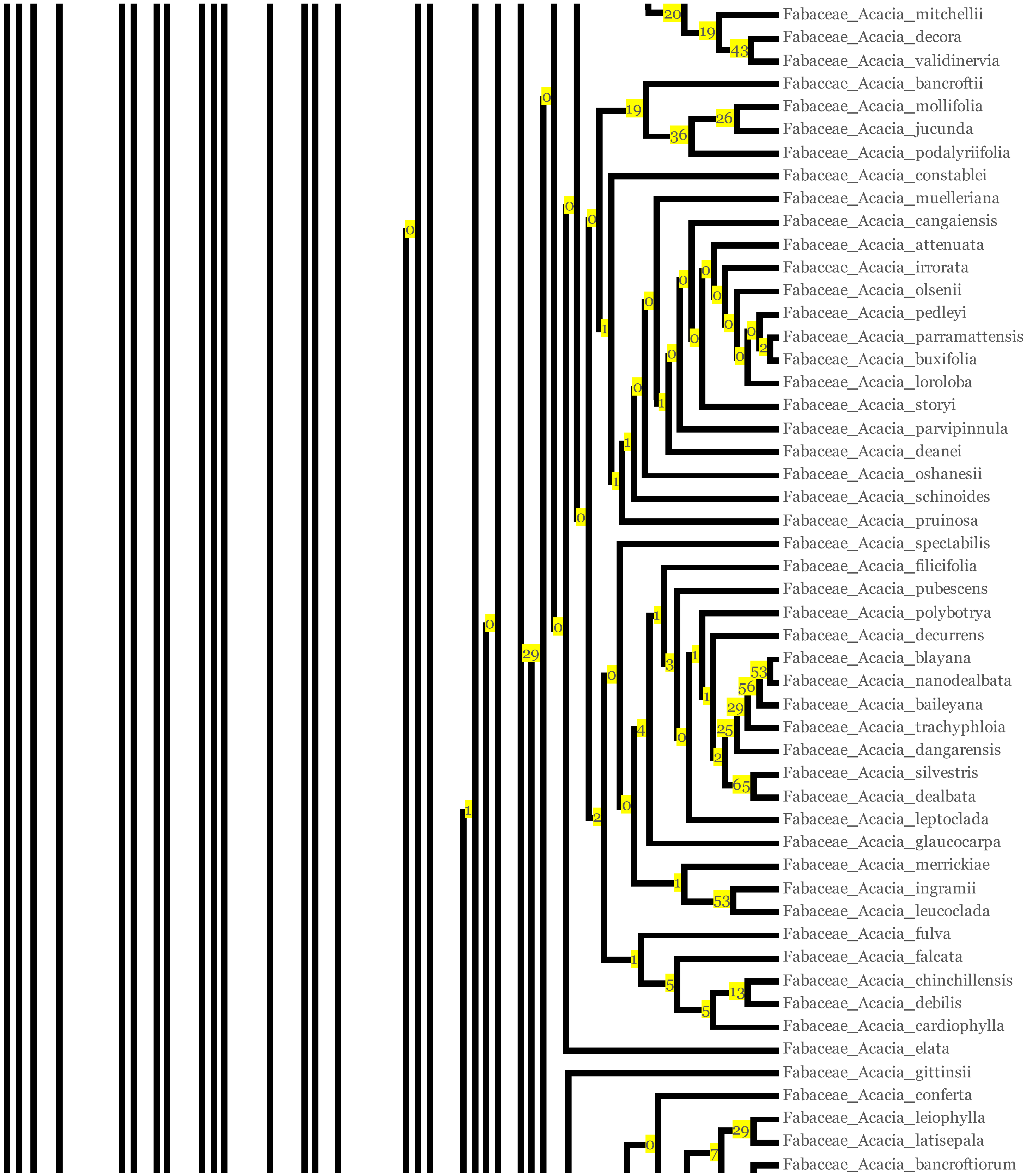
Fabaceae_Acacia_mitchellii
Fabaceae_Acacia_decora
Fabaceae_Acacia_validinervia
Fabaceae_Acacia_bancroftii
Fabaceae_Acacia_mollifolia
Fabaceae_Acacia_jucunda
Fabaceae_Acacia_podalyriifolia
Fabaceae_Acacia_constablei
Fabaceae_Acacia_muelleriana
Fabaceae_Acacia_cangaiensis
Fabaceae_Acacia_attenuata
Fabaceae_Acacia_irrorata
Fabaceae_Acacia_olsenii
Fabaceae_Acacia_pedleyi
Fabaceae_Acacia_parramattensis
Fabaceae_Acacia_buxifolia
Fabaceae_Acacia_loroloba
Fabaceae_Acacia_storyi
Fabaceae_Acacia_parvipinnula
Fabaceae_Acacia_deanei
Fabaceae_Acacia_oshanesii
Fabaceae_Acacia_schinoides
Fabaceae_Acacia_pruinosa
Fabaceae_Acacia_spectabilis
Fabaceae_Acacia_filicifolia
Fabaceae_Acacia_pubescens
Fabaceae_Acacia_polybotrya
Fabaceae_Acacia_decurrens
Fabaceae_Acacia_blayana
Fabaceae_Acacia_nanodealbata
Fabaceae_Acacia_baileyana
Fabaceae_Acacia_trachyphloia
Fabaceae_Acacia_dangarensis
Fabaceae_Acacia_silvestris
Fabaceae_Acacia_dealbata
Fabaceae_Acacia_leptoclada
Fabaceae_Acacia_glaucocarpa
Fabaceae_Acacia_merrickiae
Fabaceae_Acacia_ingramii
Fabaceae_Acacia_leucoclada
Fabaceae_Acacia_fulva
Fabaceae_Acacia_falcata
Fabaceae_Acacia_chinchillensis
Fabaceae_Acacia_debilis
Fabaceae_Acacia_cardiophylla
Fabaceae_Acacia_elata
Fabaceae_Acacia_gittinsii
Fabaceae_Acacia_conferta
Fabaceae_Acacia_leiophylla
Fabaceae_Acacia_latisepala
Fabaceae_Acacia_bancroftiorum

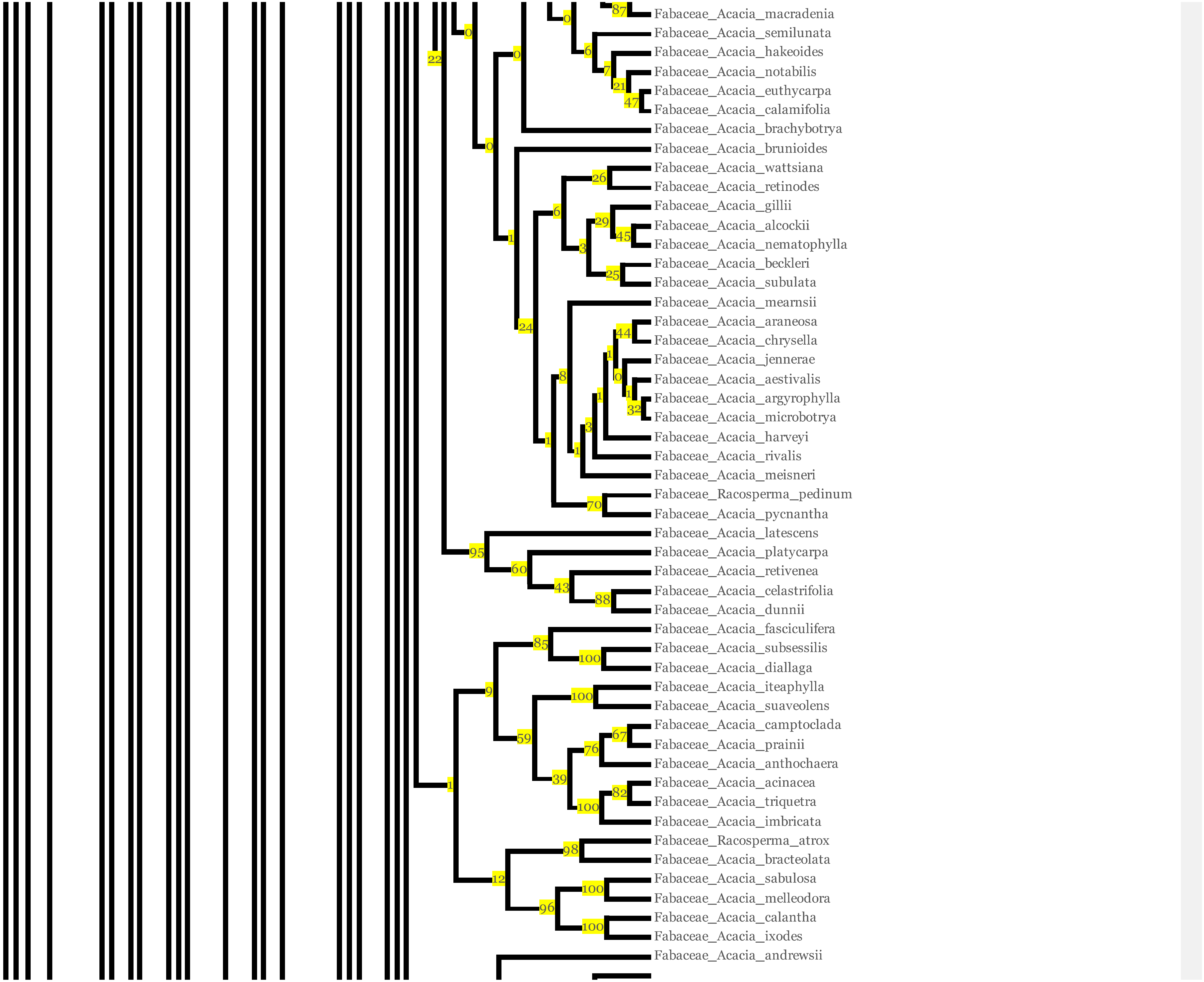
Fabaceae_Acacia_macradenia
Fabaceae_Acacia_semilunata
Fabaceae_Acacia_hakeoides
Fabaceae_Acacia_notabilis
Fabaceae_Acacia_euthycarpa
Fabaceae_Acacia_calamifolia
Fabaceae_Acacia_brachybotrya
Fabaceae_Acacia_brunioides
Fabaceae_Acacia_wattsiana
Fabaceae_Acacia_retinodes
Fabaceae_Acacia_gillii
Fabaceae_Acacia_alcockii
Fabaceae_Acacia_nematophylla
Fabaceae_Acacia_beckleri
Fabaceae_Acacia_subulata
Fabaceae_Acacia_mearnsii
Fabaceae_Acacia_araneosa
Fabaceae_Acacia_chrysella
Fabaceae_Acacia_jennerae
Fabaceae_Acacia_aestivalis
Fabaceae_Acacia_argyrophylla
Fabaceae_Acacia_microbotrya
Fabaceae_Acacia_harveyi
Fabaceae_Acacia_rivalis
Fabaceae_Acacia_meisneri
Fabaceae_Racosperma_pedinum
Fabaceae_Acacia_pycnantha
Fabaceae_Acacia_latescens
Fabaceae_Acacia_platycarpa
Fabaceae_Acacia_retivenea
Fabaceae_Acacia_celastrifolia
Fabaceae_Acacia_dunnii
Fabaceae_Acacia_fasciculifera
Fabaceae_Acacia_subsessilis
Fabaceae_Acacia_diallaga
Fabaceae_Acacia_iteaphylla
Fabaceae_Acacia_suaveolens
Fabaceae_Acacia_camptoclada
Fabaceae_Acacia_prainii
Fabaceae_Acacia_anthochaera
Fabaceae_Acacia_acinacea
Fabaceae_Acacia_triquetra
Fabaceae_Acacia_imbricata
Fabaceae_Racosperma_atrox
Fabaceae_Acacia_bracteolata
Fabaceae_Acacia_sabulosa
Fabaceae_Acacia_melleodora
Fabaceae_Acacia_calantha
Fabaceae_Acacia_ixodes
Fabaceae_Acacia_andrewsii

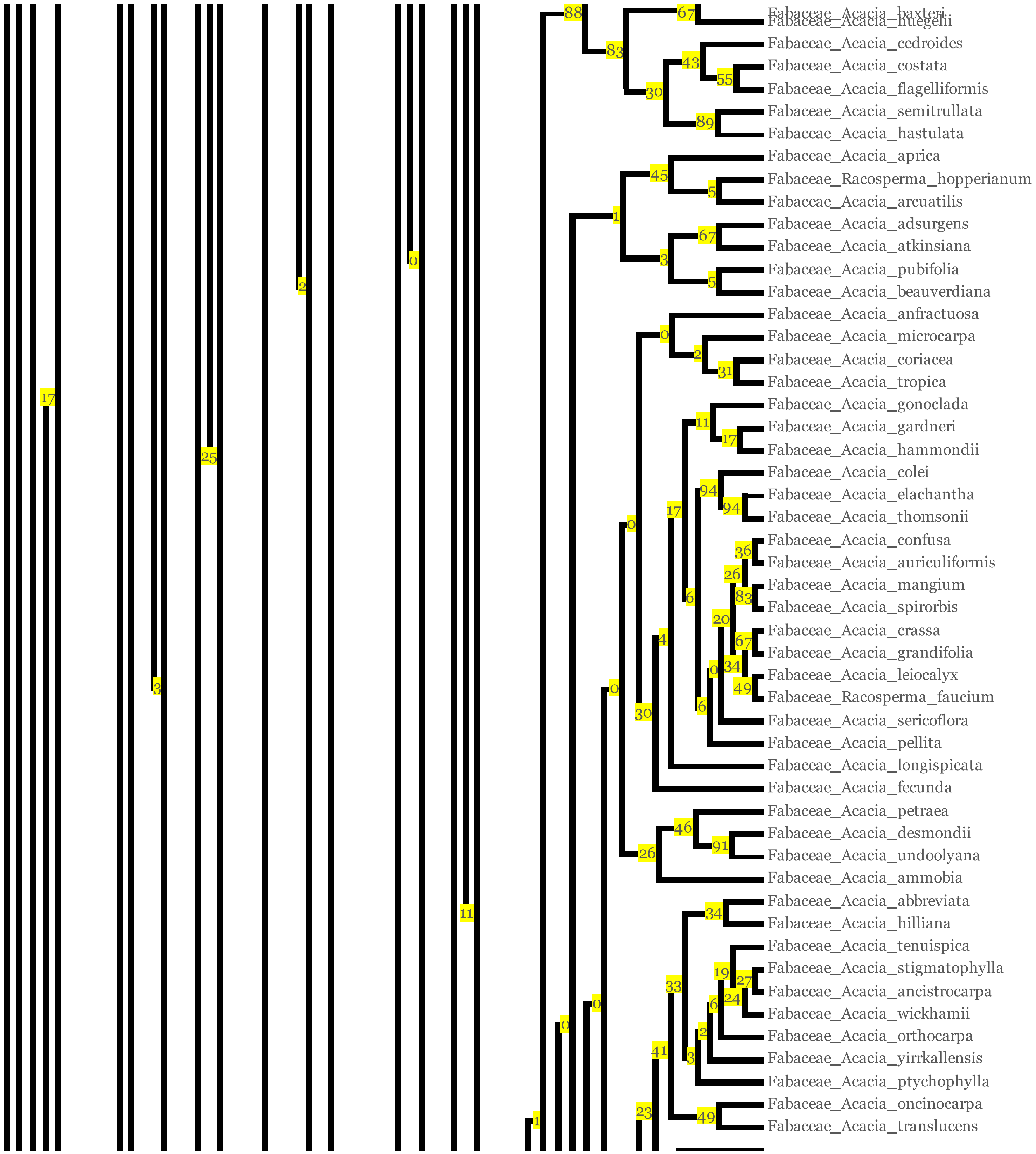
Fabaceae_Acacia_baxteri
Fabaceae_Acacia_huegelii
Fabaceae_Acacia_cedroides
Fabaceae_Acacia_costata
Fabaceae_Acacia_flagelliformis
Fabaceae_Acacia_semitrullata
Fabaceae_Acacia_hastulata
Fabaceae_Acacia_aprica
Fabaceae_Racosperma_hopperianum
Fabaceae_Acacia_arcuatilis
Fabaceae_Acacia_adsurgens
Fabaceae_Acacia_atkinsiana
Fabaceae_Acacia_pubifolia
Fabaceae_Acacia_beauverdiana
Fabaceae_Acacia_anfractuosa
Fabaceae_Acacia_microcarpa
Fabaceae_Acacia_coriacea
Fabaceae_Acacia_tropica
Fabaceae_Acacia_gonoclada
Fabaceae_Acacia_gardneri
Fabaceae_Acacia_hammondii
Fabaceae_Acacia_colei
Fabaceae_Acacia_elachantha
Fabaceae_Acacia_thomsonii
Fabaceae_Acacia_confusa
Fabaceae_Acacia_auriculiformis
Fabaceae_Acacia_mangium
Fabaceae_Acacia_spirorbis
Fabaceae_Acacia_crassa
Fabaceae_Acacia_grandifolia
Fabaceae_Acacia_leiocalyx
Fabaceae_Racosperma_faucium
Fabaceae_Acacia_sericoflora
Fabaceae_Acacia_pellita
Fabaceae_Acacia_longispicata
Fabaceae_Acacia_fecunda
Fabaceae_Acacia_petraea
Fabaceae_Acacia_desmondii
Fabaceae_Acacia_undoolyana
Fabaceae_Acacia_ammobia
Fabaceae_Acacia_abbreviata
Fabaceae_Acacia_hilliana
Fabaceae_Acacia_tenuispica
Fabaceae_Acacia_stigmatophylla
Fabaceae_Acacia_ancistrocarpa
Fabaceae_Acacia_wickhamii
Fabaceae_Acacia_orthocarpa
Fabaceae_Acacia_yirrkallensis
Fabaceae_Acacia_ptychophylla
Fabaceae_Acacia_oncinocarpa
Fabaceae_Acacia_translucens

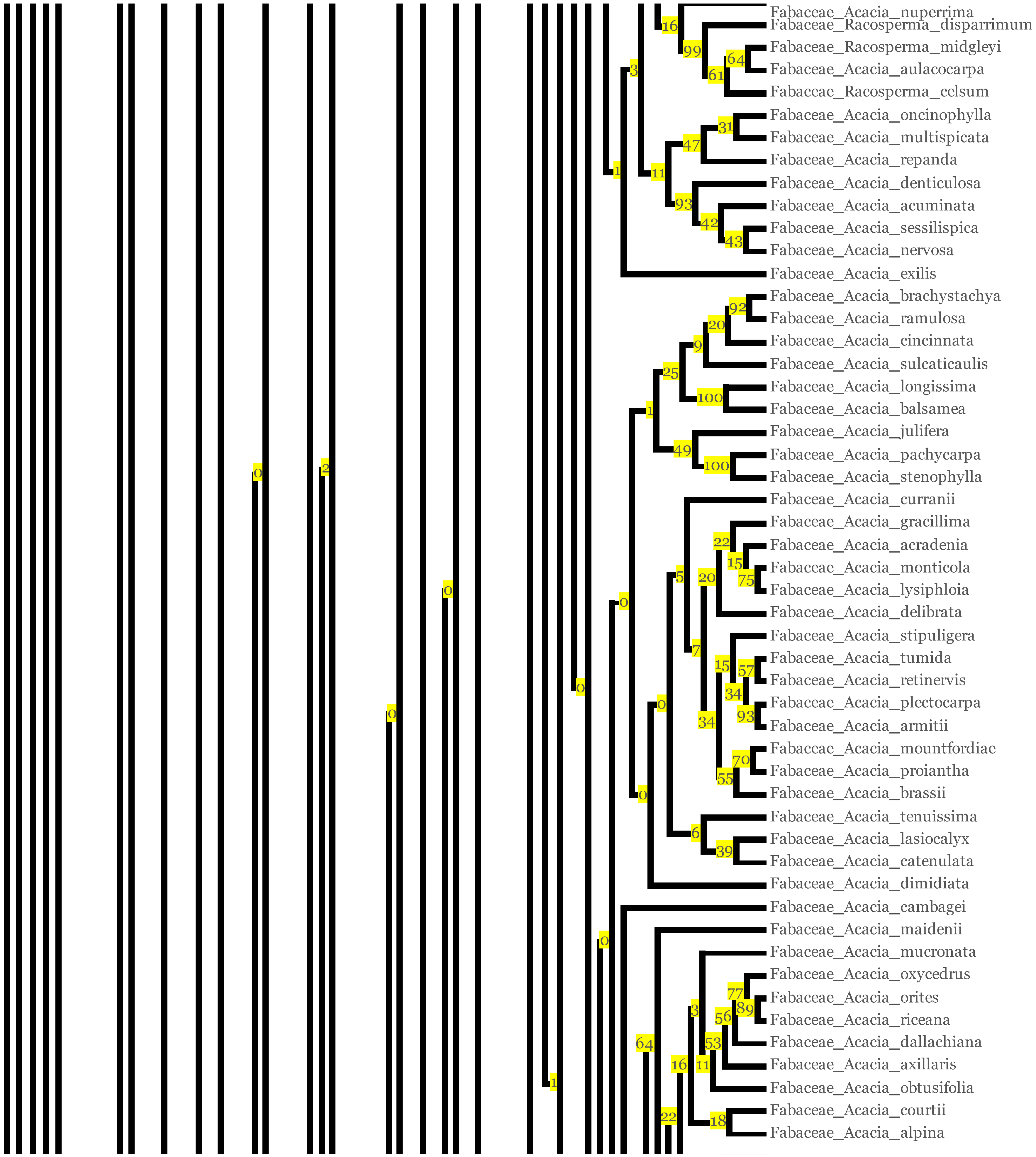
Fabaceae_Acacia_nuperrima
Fabaceae_Racosperma_disparrimum
Fabaceae_Racosperma_midgleyi
Fabaceae_Acacia_aulacocarpa
Fabaceae_Racosperma_celsum
Fabaceae_Acacia_oncinophylla
Fabaceae_Acacia_multispicata
Fabaceae_Acacia_repanda
Fabaceae_Acacia_denticulosa
Fabaceae_Acacia_acuminata
Fabaceae_Acacia_sessilispica
Fabaceae_Acacia_nervosa
Fabaceae_Acacia_exilis
Fabaceae_Acacia_brachystachya
Fabaceae_Acacia_ramulosa
Fabaceae_Acacia_cincinnata
Fabaceae_Acacia_sulcaticaulis
Fabaceae_Acacia_longissima
Fabaceae_Acacia_balsamea
Fabaceae_Acacia_julifera
Fabaceae_Acacia_pachycarpa
Fabaceae_Acacia_stenophylla
Fabaceae_Acacia_curranii
Fabaceae_Acacia_gracillima
Fabaceae_Acacia_acradenia
Fabaceae_Acacia_monticola
Fabaceae_Acacia_lysiphloia
Fabaceae_Acacia_delibrata
Fabaceae_Acacia_stipuligera
Fabaceae_Acacia_tumida
Fabaceae_Acacia_retinervis
Fabaceae_Acacia_plectocarpa
Fabaceae_Acacia_armitii
Fabaceae_Acacia_mountfordiae
Fabaceae_Acacia_proiantha
Fabaceae_Acacia_brassii
Fabaceae_Acacia_tenuissima
Fabaceae_Acacia_lasiocalyx
Fabaceae_Acacia_catenulata
Fabaceae_Acacia_dimidiata
Fabaceae_Acacia_cambagei
Fabaceae_Acacia_maidenii
Fabaceae_Acacia_mucronata
Fabaceae_Acacia_oxycedrus
Fabaceae_Acacia_orites
Fabaceae_Acacia_riceana
Fabaceae_Acacia_dallachiana
Fabaceae_Acacia_axillaris
Fabaceae_Acacia_obtusifolia
Fabaceae_Acacia_courtii
Fabaceae_Acacia_alpina

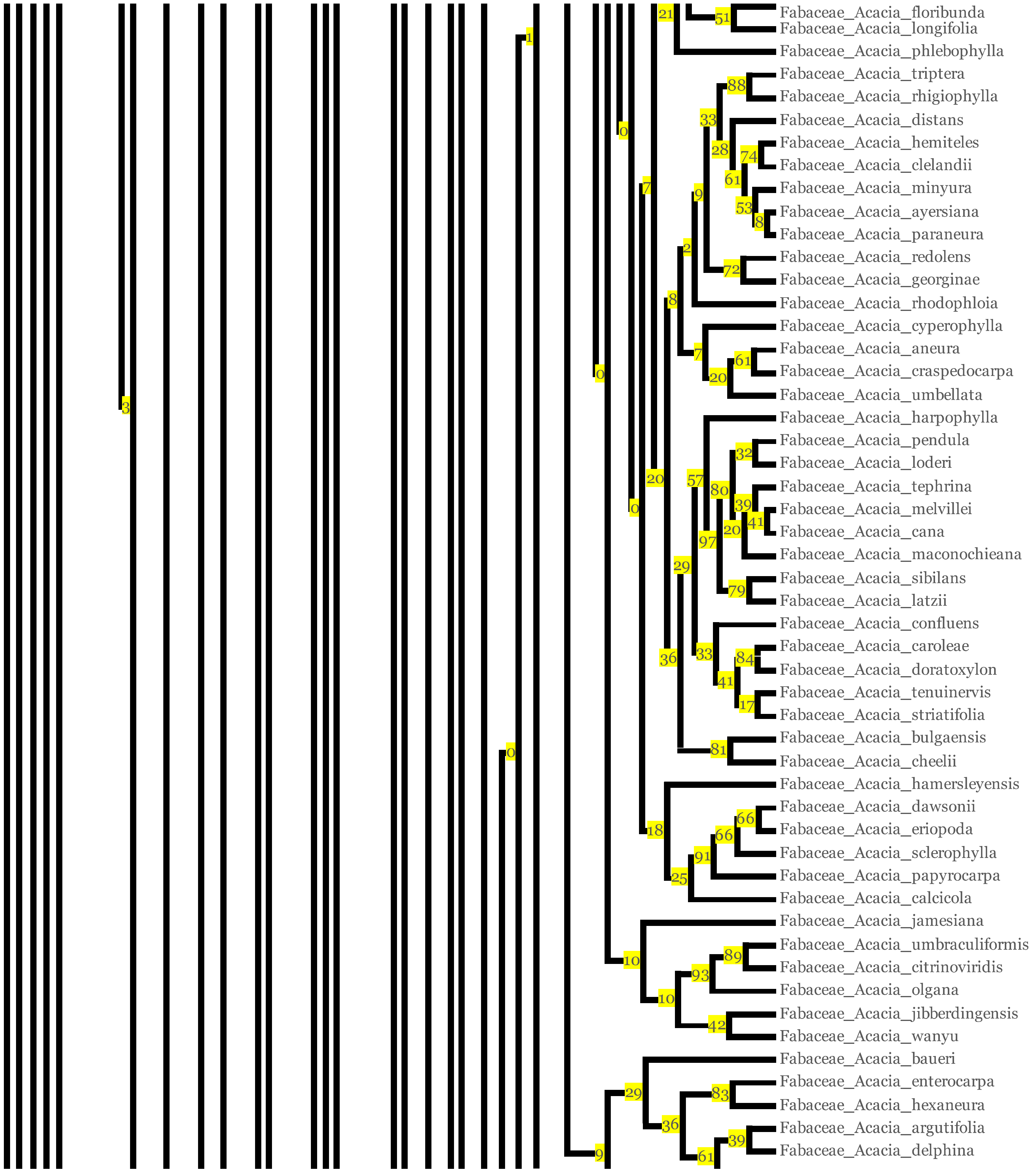
Fabaceae_Acacia_floribunda
Fabaceae_Acacia_longifolia
Fabaceae_Acacia_phlebophylla
Fabaceae_Acacia_triptera
Fabaceae_Acacia_rhigiophylla
Fabaceae_Acacia_distans
Fabaceae_Acacia_hemiteles
Fabaceae_Acacia_clelandii
Fabaceae_Acacia_minyura
Fabaceae_Acacia_ayersiana
Fabaceae_Acacia_paraneura
Fabaceae_Acacia_redolens
Fabaceae_Acacia_georginae
Fabaceae_Acacia_rhodophloia
Fabaceae_Acacia_cyperophylla
Fabaceae_Acacia_aneura
Fabaceae_Acacia_craspedocarpa
Fabaceae_Acacia_umbellata
Fabaceae_Acacia_harpophylla
Fabaceae_Acacia_pendula
Fabaceae_Acacia_loderi
Fabaceae_Acacia_tephrina
Fabaceae_Acacia_melvillei
Fabaceae_Acacia_cana
Fabaceae_Acacia_maconochieana
Fabaceae_Acacia_sibilans
Fabaceae_Acacia_latzii
Fabaceae_Acacia_confluens
Fabaceae_Acacia_caroleae
Fabaceae_Acacia_doratoxylon
Fabaceae_Acacia_tenuinervis
Fabaceae_Acacia_striatifolia
Fabaceae_Acacia_bulgaensis
Fabaceae_Acacia_cheelii
Fabaceae_Acacia_hamersleyensis
Fabaceae_Acacia_dawsonii
Fabaceae_Acacia_eriopoda
Fabaceae_Acacia_sclerophylla
Fabaceae_Acacia_papyrocarpa
Fabaceae_Acacia_calcicola
Fabaceae_Acacia_jamesiana
Fabaceae_Acacia_umbraculiformis
Fabaceae_Acacia_citrinoviridis
Fabaceae_Acacia_olgana
Fabaceae_Acacia_jibberdingensis
Fabaceae_Acacia_wanyu
Fabaceae_Acacia_baueri
Fabaceae_Acacia_enterocarpa
Fabaceae_Acacia_hexaneura
Fabaceae_Acacia_argutifolia
Fabaceae_Acacia_delphina

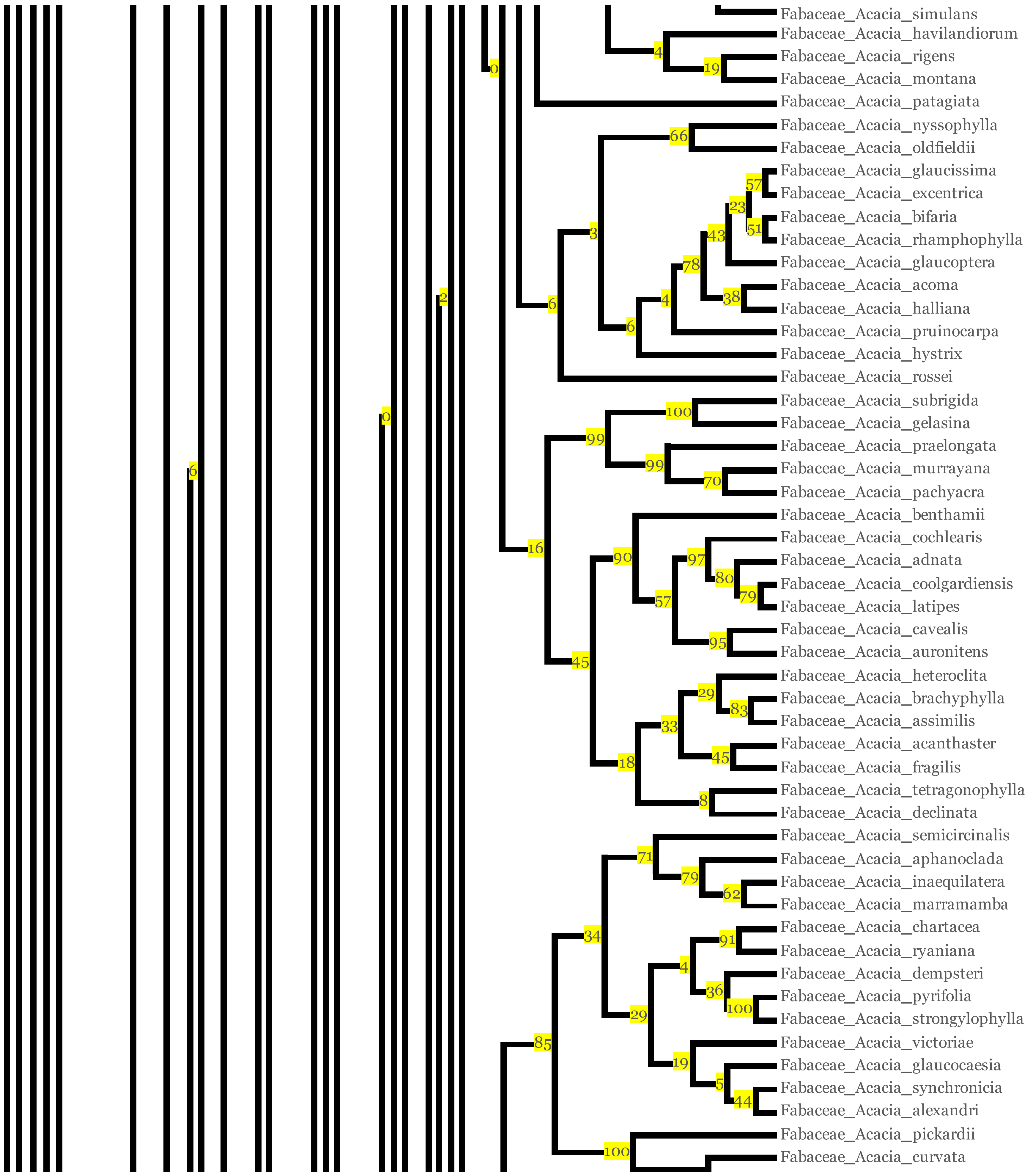
Fabaceae_Acacia_simulans
Fabaceae_Acacia_havilandiorum
Fabaceae_Acacia_rigens
Fabaceae_Acacia_montana
Fabaceae_Acacia_patagiata
Fabaceae_Acacia_nyssophylla
Fabaceae_Acacia_oldfieldii
Fabaceae_Acacia_glaucissima
Fabaceae_Acacia_excentrica
Fabaceae_Acacia_bifaria
Fabaceae_Acacia_rhamphophylla
Fabaceae_Acacia_glaucoptera
Fabaceae_Acacia_acoma
Fabaceae_Acacia_halliana
Fabaceae_Acacia_pruinocarpa
Fabaceae_Acacia_hystrix
Fabaceae_Acacia_rossei
Fabaceae_Acacia_subrigida
Fabaceae_Acacia_gelasina
Fabaceae_Acacia_praelongata
Fabaceae_Acacia_murrayana
Fabaceae_Acacia_pachyacra
Fabaceae_Acacia_benthamii
Fabaceae_Acacia_cochlearis
Fabaceae_Acacia_adnata
Fabaceae_Acacia_coolgardiensis
Fabaceae_Acacia_latipes
Fabaceae_Acacia_cavealis
Fabaceae_Acacia_auronitens
Fabaceae_Acacia_heteroclita
Fabaceae_Acacia_brachyphylla
Fabaceae_Acacia_assimilis
Fabaceae_Acacia_acanthaster
Fabaceae_Acacia_fragilis
Fabaceae_Acacia_tetragonophylla
Fabaceae_Acacia_declinata
Fabaceae_Acacia_semicircinalis
Fabaceae_Acacia_aphanoclada
Fabaceae_Acacia_inaequilatera
Fabaceae_Acacia_marramamba
Fabaceae_Acacia_chartacea
Fabaceae_Acacia_ryaniana
Fabaceae_Acacia_dempsteri
Fabaceae_Acacia_pyrifolia
Fabaceae_Acacia_strongylophylla
Fabaceae_Acacia_victoriae
Fabaceae_Acacia_glaucocaesia
Fabaceae_Acacia_synchronicia
Fabaceae_Acacia_alexandri
Fabaceae_Acacia_pickardii
Fabaceae_Acacia_curvata

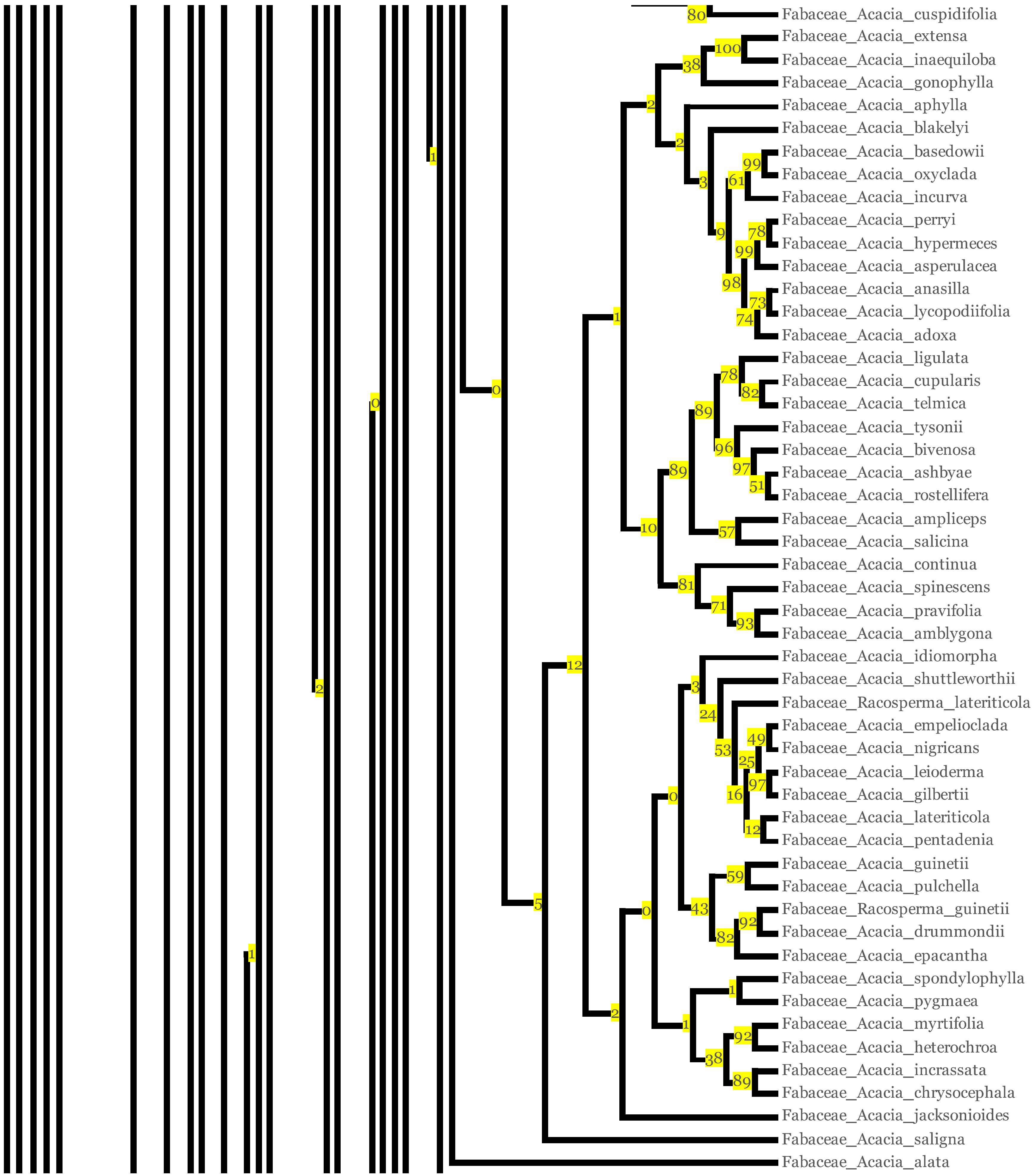
Fabaceae_Acacia_cuspidifolia
Fabaceae_Acacia_extensa
Fabaceae_Acacia_inaequiloba
Fabaceae_Acacia_gonophylla
Fabaceae_Acacia_aphylla
Fabaceae_Acacia_blakelyi
Fabaceae_Acacia_basedowii
Fabaceae_Acacia_oxyclada
Fabaceae_Acacia_incurva
Fabaceae_Acacia_perryi
Fabaceae_Acacia_hypermeces
Fabaceae_Acacia_asperulacea
Fabaceae_Acacia_anasilla
Fabaceae_Acacia_lycopodiifolia
Fabaceae_Acacia_adoxa
Fabaceae_Acacia_ligulata
Fabaceae_Acacia_cupularis
Fabaceae_Acacia_telmica
Fabaceae_Acacia_tysonii
Fabaceae_Acacia_bivenosa
Fabaceae_Acacia_ashbyae
Fabaceae_Acacia_rostellifera
Fabaceae_Acacia_ampliceps
Fabaceae_Acacia_salicina
Fabaceae_Acacia_continua
Fabaceae_Acacia_spinescens
Fabaceae_Acacia_pravifolia
Fabaceae_Acacia_amblygona
Fabaceae_Acacia_idiomorpha
Fabaceae_Acacia_shuttleworthii
Fabaceae_Racosperma_lateriticola
Fabaceae_Acacia_empelioclada
Fabaceae_Acacia_nigricans
Fabaceae_Acacia_leioderma
Fabaceae_Acacia_gilbertii
Fabaceae_Acacia_lateriticola
Fabaceae_Acacia_pentadenia
Fabaceae_Acacia_guinetii
Fabaceae_Acacia_pulchella
Fabaceae_Racosperma_guinetii
Fabaceae_Acacia_drummondii
Fabaceae_Acacia_epacantha
Fabaceae_Acacia_spondylophylla
Fabaceae_Acacia_pygmaea
Fabaceae_Acacia_myrtifolia
Fabaceae_Acacia_heterochroa
Fabaceae_Acacia_incrassata
Fabaceae_Acacia_chrysocephala
Fabaceae_Acacia_jacksonioides
Fabaceae_Acacia_saligna
Fabaceae_Acacia_alata

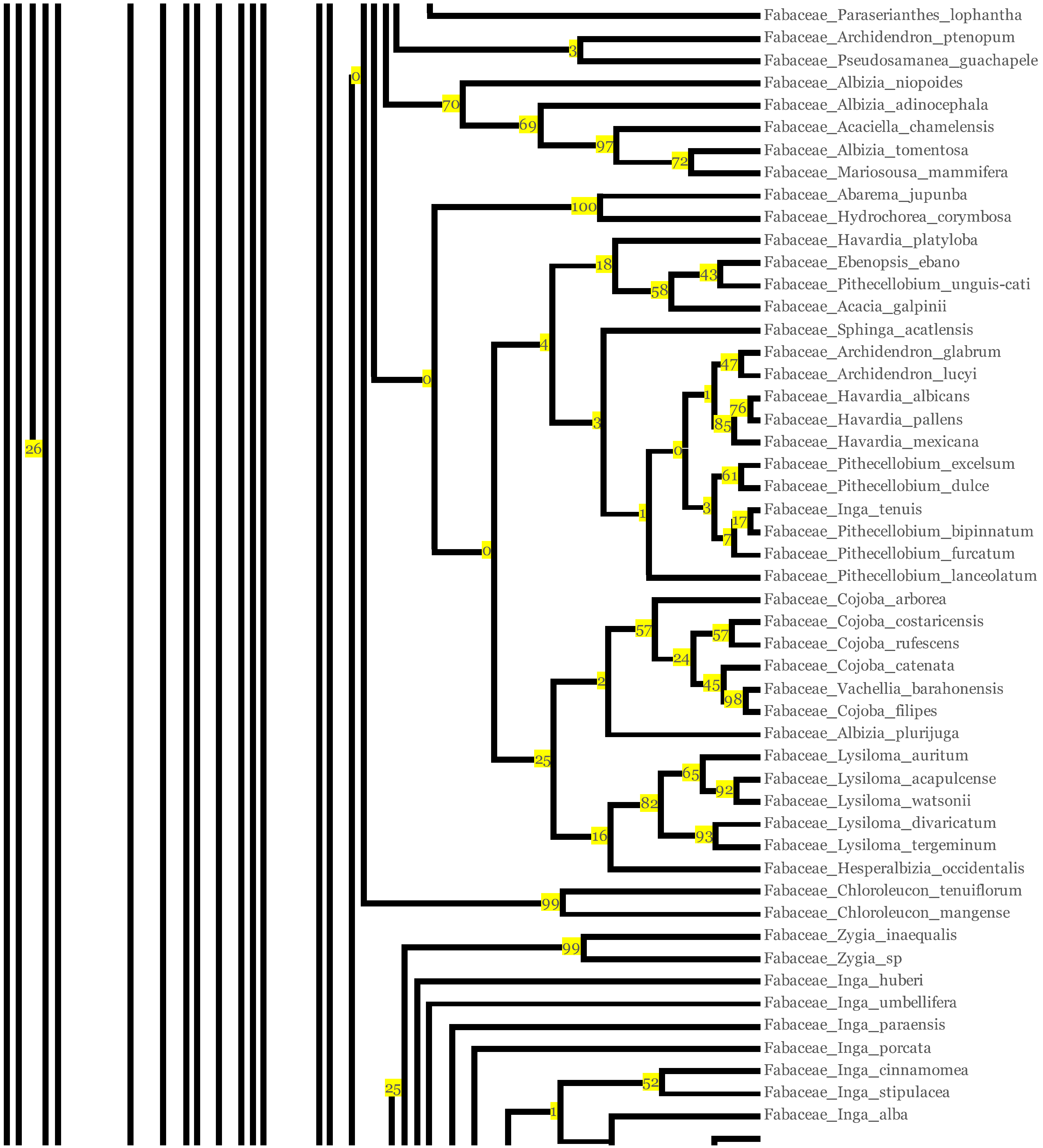
Fabaceae_Paraserianthes_lophantha
Fabaceae_Archidendron_ptenopum
Fabaceae_Pseudosamanea_guachapele
Fabaceae_Albizia_niopoides
Fabaceae_Albizia_adinocephala
Fabaceae_Acaciella_chamelensis
Fabaceae_Albizia_tomentosa
Fabaceae_Mariosousa_mammifera
Fabaceae_Abarema_jupunba
Fabaceae_Hydrochorea_corymbosa
Fabaceae_Havardia_platyloba
Fabaceae_Ebenopsis_ebano
Fabaceae_Pithecellobium_unguis-cati
Fabaceae_Acacia_galpinii
Fabaceae_Sphinga_acatlensis
Fabaceae_Archidendron_glabrum
Fabaceae_Archidendron_lucyi
Fabaceae_Havardia_albicans
Fabaceae_Havardia_pallens
Fabaceae_Havardia_mexicana
Fabaceae_Pithecellobium_excelsum
Fabaceae_Pithecellobium_dulce
Fabaceae_Inga_tenuis
Fabaceae_Pithecellobium_bipinnatum
Fabaceae_Pithecellobium_furcatum
Fabaceae_Pithecellobium_lanceolatum
Fabaceae_Cojoba_arborea
Fabaceae_Cojoba_costaricensis
Fabaceae_Cojoba_rufescens
Fabaceae_Cojoba_catenata
Fabaceae_Vachellia_barahonensis
Fabaceae_Cojoba_filipes
Fabaceae_Albizia_plurijuga
Fabaceae_Lysiloma_auritum
Fabaceae_Lysiloma_acapulcense
Fabaceae_Lysiloma_watsonii
Fabaceae_Lysiloma_divaricatum
Fabaceae_Lysiloma_tergeminum
Fabaceae_Hesperalbizia_occidentalis
Fabaceae_Chloroleucon_tenuiflorum
Fabaceae_Chloroleucon_mangense
Fabaceae_Zygia_inaequalis
Fabaceae_Zygia_sp
Fabaceae_Inga_huberi
Fabaceae_Inga_umbellifera
Fabaceae_Inga_paraensis
Fabaceae_Inga_porcata
Fabaceae_Inga_cinnamomea
Fabaceae_Inga_stipulacea
Fabaceae_Inga_alba

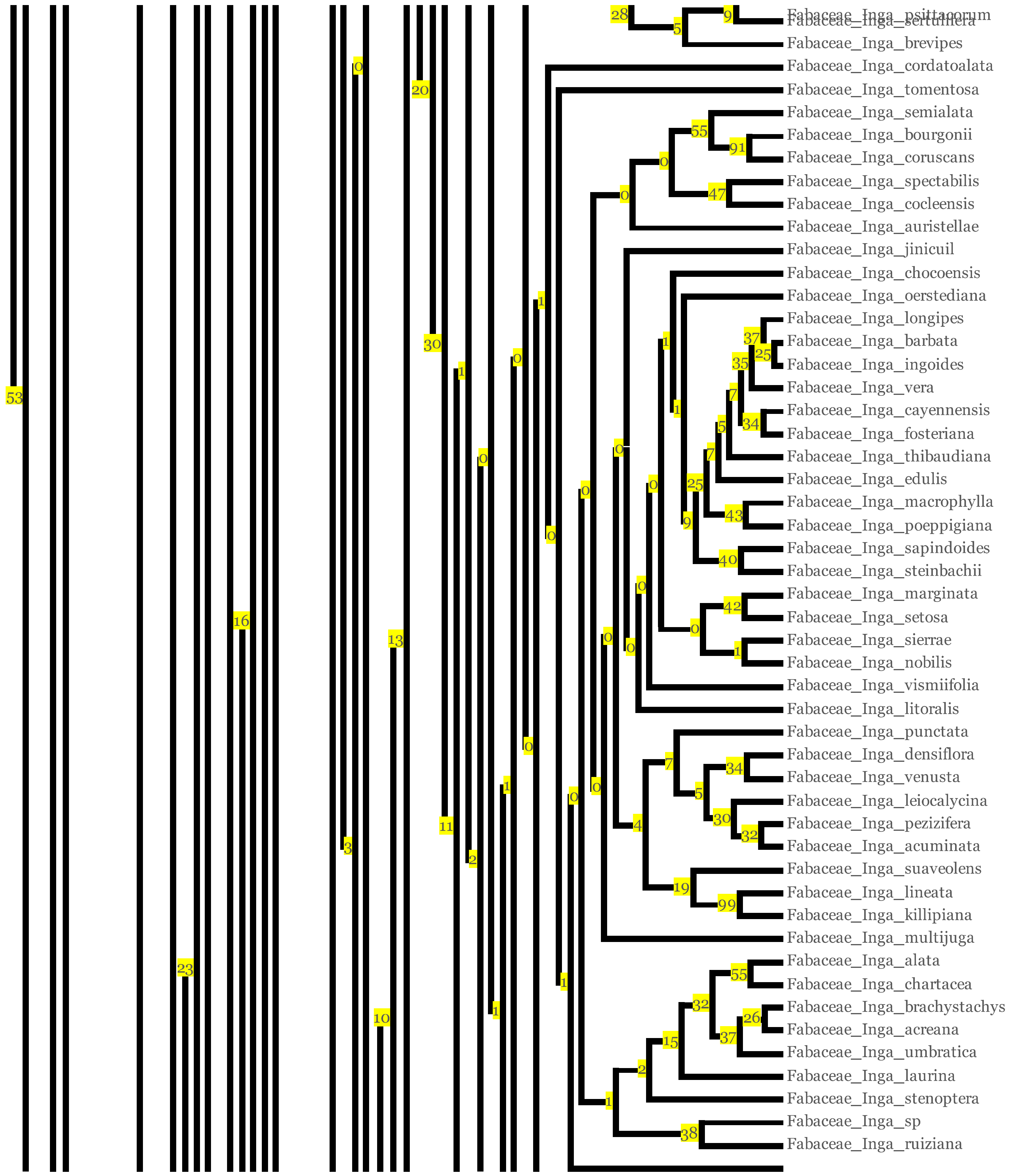
Fabaceae_Inga_psittacorum
Fabaceae_Inga_sertulifera
Fabaceae_Inga_brevipes
Fabaceae_Inga_cordatoalata
Fabaceae_Inga_tomentosa
Fabaceae_Inga_semialata
Fabaceae_Inga_bourgonii
Fabaceae_Inga_coruscans
Fabaceae_Inga_spectabilis
Fabaceae_Inga_cocleensis
Fabaceae_Inga_auristellae
Fabaceae_Inga_jinicuil
Fabaceae_Inga_chocoensis
Fabaceae_Inga_oerstediana
Fabaceae_Inga_longipes
Fabaceae_Inga_barbata
Fabaceae_Inga_ingoides
Fabaceae_Inga_vera
Fabaceae_Inga_cayennensis
Fabaceae_Inga_fosteriana
Fabaceae_Inga_thibaudiana
Fabaceae_Inga_edulis
Fabaceae_Inga_macrophylla
Fabaceae_Inga_poeppigiana
Fabaceae_Inga_sapindoides
Fabaceae_Inga_steinbachii
Fabaceae_Inga_marginata
Fabaceae_Inga_setosa
Fabaceae_Inga_sierrae
Fabaceae_Inga_nobilis
Fabaceae_Inga_vismiifolia
Fabaceae_Inga_litoralis
Fabaceae_Inga_punctata
Fabaceae_Inga_densiflora
Fabaceae_Inga_venusta
Fabaceae_Inga_leiocalycina
Fabaceae_Inga_pezizifera
Fabaceae_Inga_acuminata
Fabaceae_Inga_suaveolens
Fabaceae_Inga_lineata
Fabaceae_Inga_killipiana
Fabaceae_Inga_multijuga
Fabaceae_Inga_alata
Fabaceae_Inga_chartacea
Fabaceae_Inga_brachystachys
Fabaceae_Inga_acreana
Fabaceae_Inga_umbratica
Fabaceae_Inga_laurina
Fabaceae_Inga_stenoptera
Fabaceae_Inga_sp
Fabaceae_Inga_ruiziana

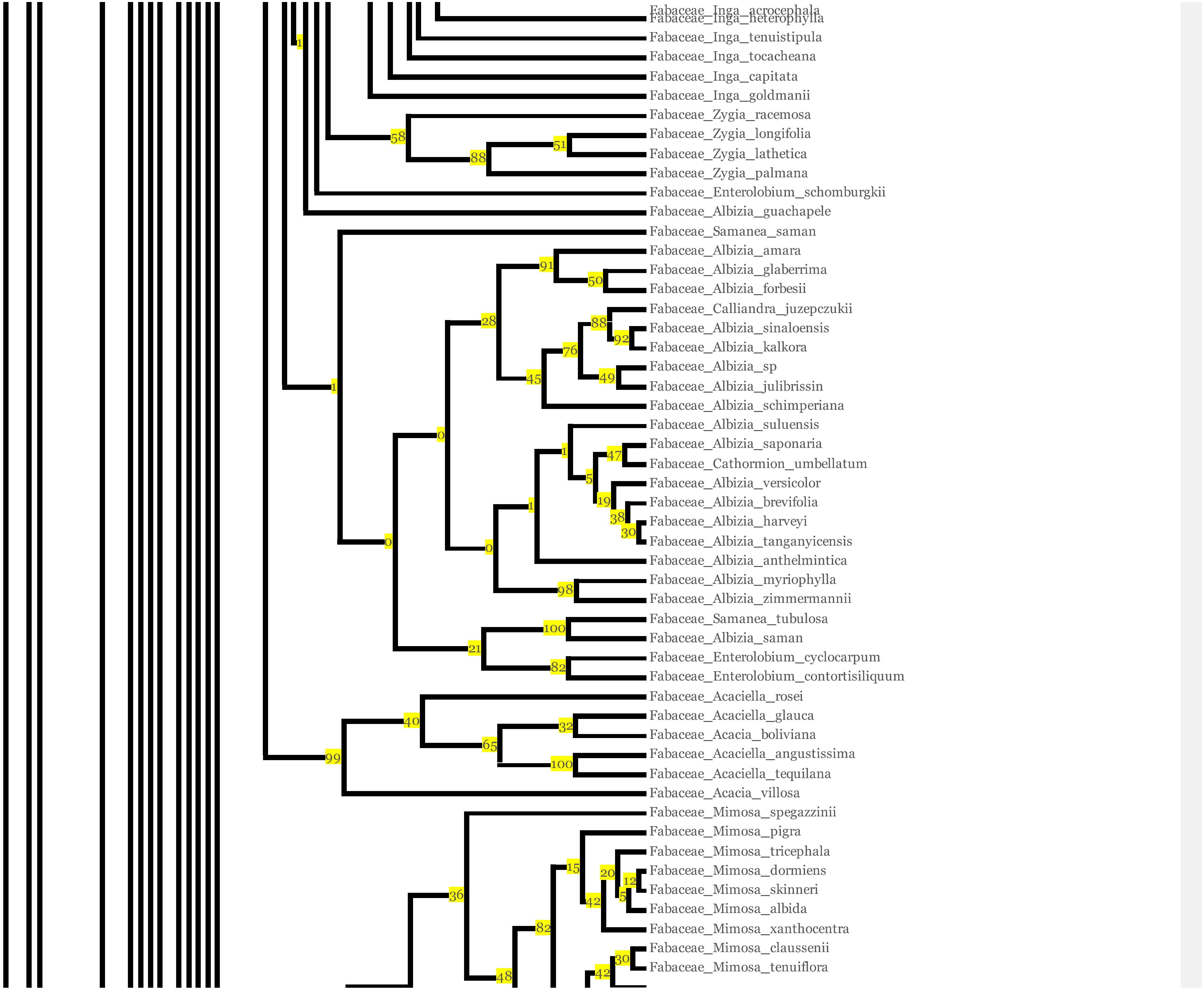
Fabaceae_Inga_acrocephala
Fabaceae_Inga_heterophylla
Fabaceae_Inga_tenuistipula
Fabaceae_Inga_tocacheana
Fabaceae_Inga_capitata
Fabaceae_Inga_goldmanii
Fabaceae_Zygia_racemosa
Fabaceae_Zygia_longifolia
Fabaceae_Zygia_lathetica
Fabaceae_Zygia_palmana
Fabaceae_Enterolobium_schomburgkii
Fabaceae_Albizia_guachapele
Fabaceae_Samanea_saman
Fabaceae_Albizia_amara
Fabaceae_Albizia_glaberrima
Fabaceae_Albizia_forbesii
Fabaceae_Calliandra_juzepczukii
Fabaceae_Albizia_sinaloensis
Fabaceae_Albizia_kalkora
Fabaceae_Albizia_sp
Fabaceae_Albizia_julibrissin
Fabaceae_Albizia_schimperiana
Fabaceae_Albizia_suluensis
Fabaceae_Albizia_saponaria
Fabaceae_Cathormion_umbellatum
Fabaceae_Albizia_versicolor
Fabaceae_Albizia_brevifolia
Fabaceae_Albizia_harveyi
Fabaceae_Albizia_tanganyicensis
Fabaceae_Albizia_anthelmintica
Fabaceae_Albizia_myriophylla
Fabaceae_Albizia_zimmermannii
Fabaceae_Samanea_tubulosa
Fabaceae_Albizia_saman
Fabaceae_Enterolobium_cyclocarpum
Fabaceae_Enterolobium_contortisiliquum
Fabaceae_Acaciella_rosei
Fabaceae_Acaciella_glauca
Fabaceae_Acacia_boliviana
Fabaceae_Acaciella_angustissima
Fabaceae_Acaciella_tequilana
Fabaceae_Acacia_villosa
Fabaceae_Mimosa_spegazzinii
Fabaceae_Mimosa_pigra
Fabaceae_Mimosa_tricephala
Fabaceae_Mimosa_dormiens
Fabaceae_Mimosa_skinneri
Fabaceae_Mimosa_albida
Fabaceae_Mimosa_xanthocentra
Fabaceae_Mimosa_claussenii
Fabaceae_Mimosa_tenuiflora

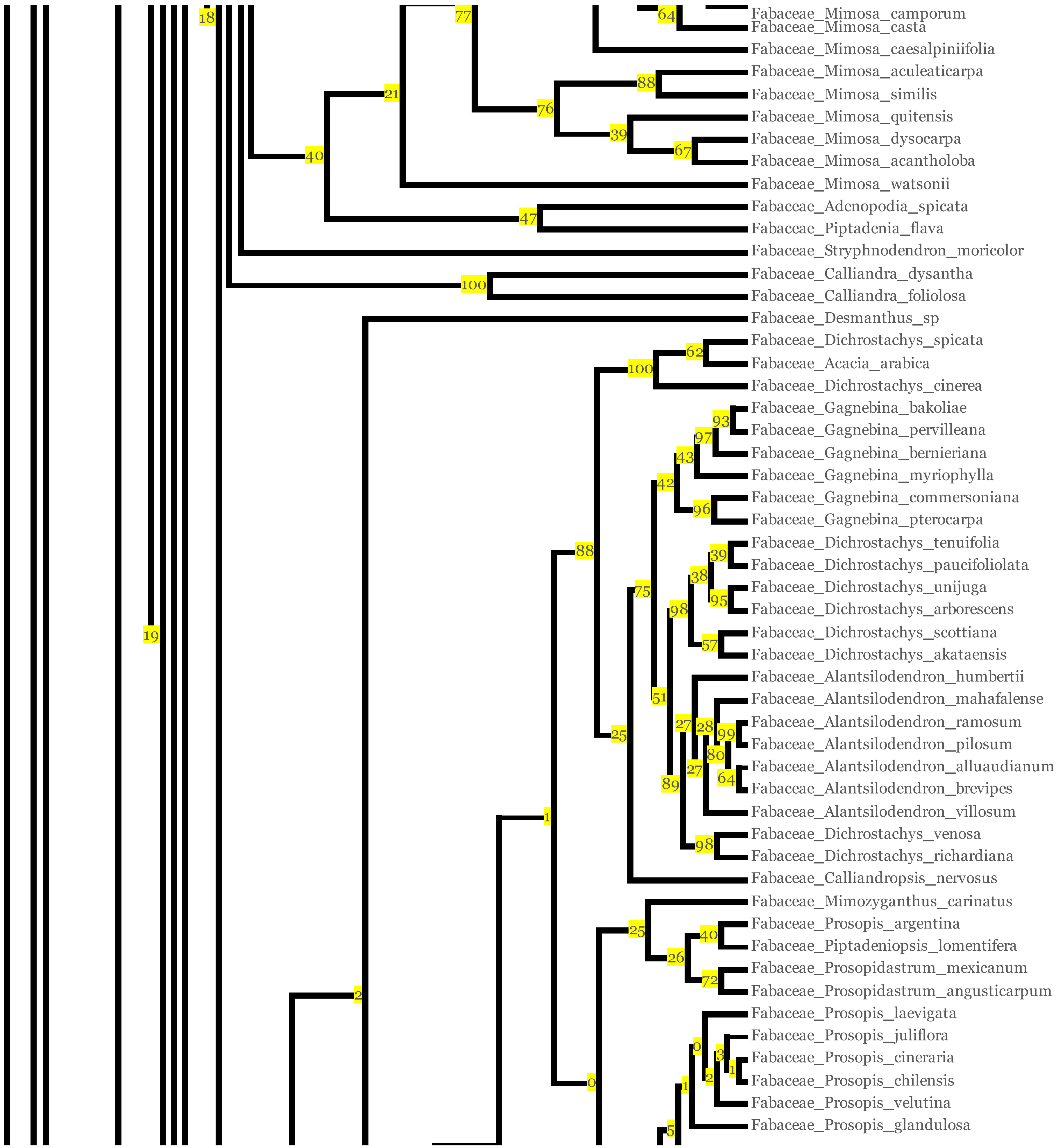

Fabaceae_Mimosa_camporum
Fabaceae_Mimosa_casta
Fabaceae_Mimosa_caesalpiniifolia
Fabaceae_Mimosa_aculeaticarpa
Fabaceae_Mimosa_similis
Fabaceae_Mimosa_quitensis
Fabaceae_Mimosa_dysocarpa
Fabaceae_Mimosa_acantholoba
Fabaceae_Mimosa_watsonii
Fabaceae_Adenopodia_spicata
Fabaceae_Piptadenia_flava
Fabaceae_Stryphnodendron_moricolor
Fabaceae_Calliandra_dysantha
Fabaceae_Calliandra_foliolosa
Fabaceae_Desmanthus_sp
Fabaceae_Dichrostachys_spicata
Fabaceae_Acacia_arabica
Fabaceae_Dichrostachys_cinerea
Fabaceae_Gagnebina_bakoliae
Fabaceae_Gagnebina_pervilleana
Fabaceae_Gagnebina_bernieriana
Fabaceae_Gagnebina_myriophylla
Fabaceae_Gagnebina_commersoniana
Fabaceae_Gagnebina_pterocarpa
Fabaceae_Dichrostachys_tenuifolia
Fabaceae_Dichrostachys_paucifoliolata
Fabaceae_Dichrostachys_unijuga
Fabaceae_Dichrostachys_arborescens
Fabaceae_Dichrostachys_scottiana
Fabaceae_Dichrostachys_akataensis
Fabaceae_Alantsilodendron_humbertii
Fabaceae_Alantsilodendron_mahafalense
Fabaceae_Alantsilodendron_ramosum
Fabaceae_Alantsilodendron_pilosum
Fabaceae_Alantsilodendron_alluaudianum
Fabaceae_Alantsilodendron_brevipes
Fabaceae_Alantsilodendron_villosum
Fabaceae_Dichrostachys_venosa
Fabaceae_Dichrostachys_richardiana
Fabaceae_Calliandropsis_nervosus
Fabaceae_Mimozyganthus_carinatus
Fabaceae_Prosopis_argentina
Fabaceae_Piptadeniopsis_lomentifera
Fabaceae_Prosopidastrum_mexicanum
Fabaceae_Prosopidastrum_angusticarpum
Fabaceae_Prosopis_laevigata
Fabaceae_Prosopis_juliflora
Fabaceae_Prosopis_cineraria
Fabaceae_Prosopis_chilensis
Fabaceae_Prosopis_velutina
Fabaceae_Prosopis_glandulosa

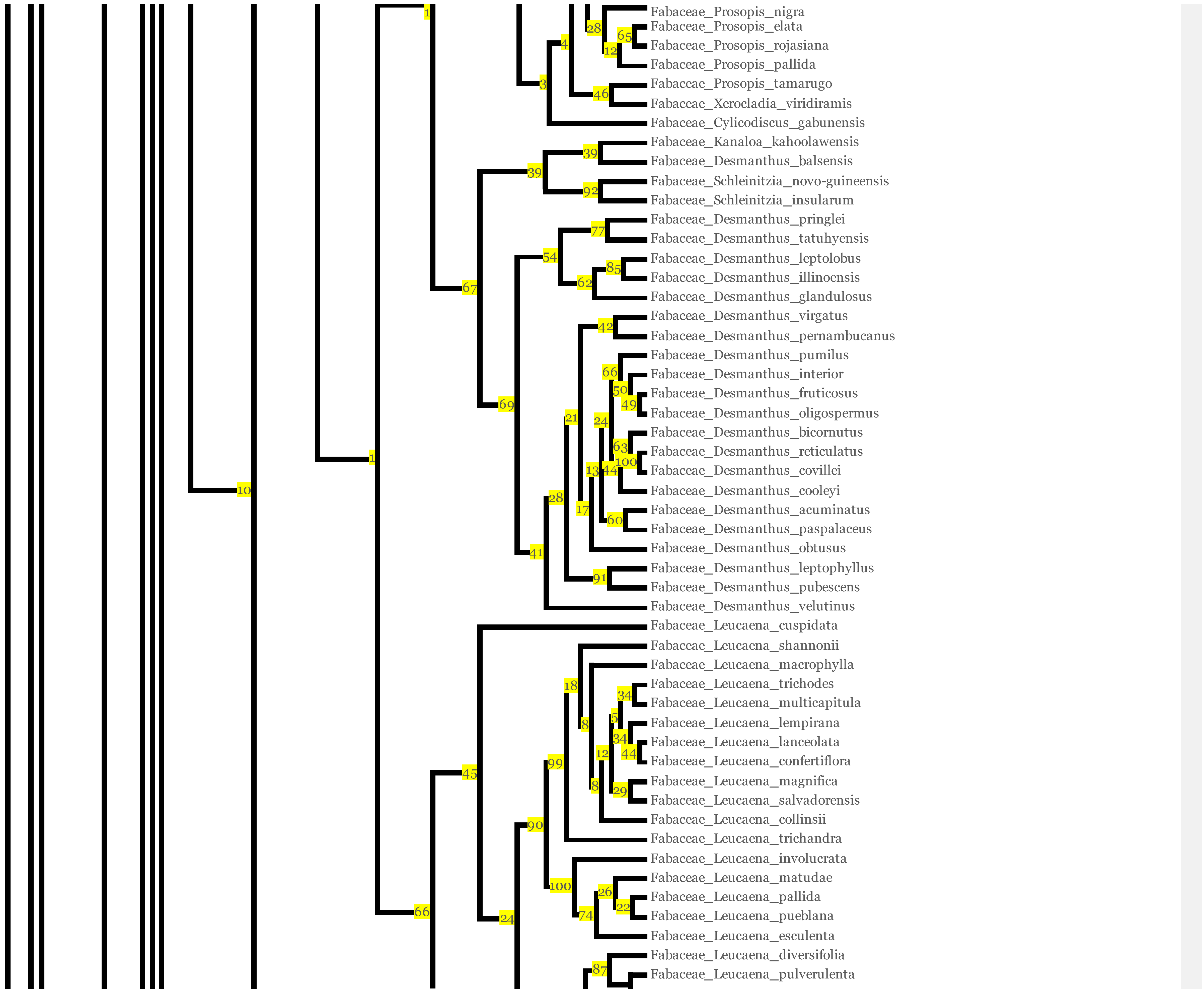
Fabaceae_Prosopis_nigra
Fabaceae_Prosopis_elata
Fabaceae_Prosopis_rojasiana
Fabaceae_Prosopis_pallida
Fabaceae_Prosopis_tamarugo
Fabaceae_Xerocladia_viridiramis
Fabaceae_Cylicodiscus_gabunensis
Fabaceae_Kanaloa_kahoolawensis
Fabaceae_Desmanthus_balsensis
Fabaceae_Schleinitzia_novo-guineensis
Fabaceae_Schleinitzia_insularum
Fabaceae_Desmanthus_pringlei
Fabaceae_Desmanthus_tatuhyensis
Fabaceae_Desmanthus_leptolobus
Fabaceae_Desmanthus_illinoensis
Fabaceae_Desmanthus_glandulosus
Fabaceae_Desmanthus_virgatus
Fabaceae_Desmanthus_pernambucanus
Fabaceae_Desmanthus_pumilus
Fabaceae_Desmanthus_interior
Fabaceae_Desmanthus_fruticosus
Fabaceae_Desmanthus_oligospermus
Fabaceae_Desmanthus_bicornutus
Fabaceae_Desmanthus_reticulatus
Fabaceae_Desmanthus_covillei
Fabaceae_Desmanthus_cooleyi
Fabaceae_Desmanthus_acuminatus
Fabaceae_Desmanthus_paspalaceus
Fabaceae_Desmanthus_obtusus
Fabaceae_Desmanthus_leptophyllus
Fabaceae_Desmanthus_pubescens
Fabaceae_Desmanthus_velutinus
Fabaceae_Leucaena_cuspidata
Fabaceae_Leucaena_shannonii
Fabaceae_Leucaena_macrophylla
Fabaceae_Leucaena_trichodes
Fabaceae_Leucaena_multicapitula
Fabaceae_Leucaena_lempirana
Fabaceae_Leucaena_lanceolata
Fabaceae_Leucaena_confertiflora
Fabaceae_Leucaena_magnifica
Fabaceae_Leucaena_salvadorensis
Fabaceae_Leucaena_collinsii
Fabaceae_Leucaena_trichandra
Fabaceae_Leucaena_involucrata
Fabaceae_Leucaena_matudae
Fabaceae_Leucaena_pallida
Fabaceae_Leucaena_pueblana
Fabaceae_Leucaena_esculenta
Fabaceae_Leucaena_diversifolia
Fabaceae_Leucaena_pulverulenta

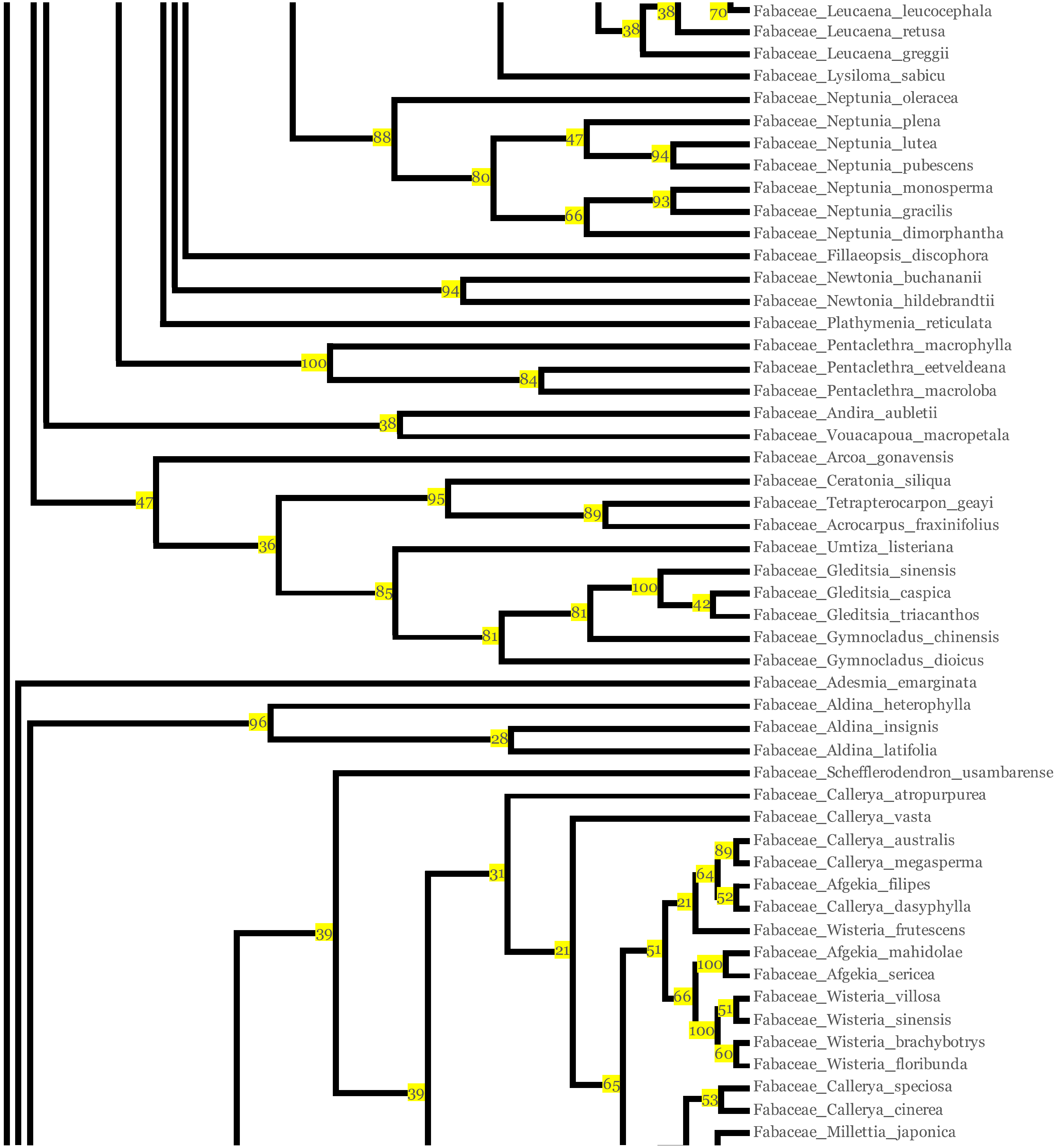
Fabaceae_Leucaena_leucocephala
Fabaceae_Leucaena_retusa
Fabaceae_Leucaena_greggii
Fabaceae_Lysiloma_sabicu
Fabaceae_Neptunia_oleracea
Fabaceae_Neptunia_plena
Fabaceae_Neptunia_lutea
Fabaceae_Neptunia_pubescens
Fabaceae_Neptunia_monosperma
Fabaceae_Neptunia_gracilis
Fabaceae_Neptunia_dimorphantha
Fabaceae_Fillaeopsis_discophora
Fabaceae_Newtonia_buchananii
Fabaceae_Newtonia_hildebrandtii
Fabaceae_Plathymenia_reticulata
Fabaceae_Pentaclethra_macrophylla
Fabaceae_Pentaclethra_eetveldeana
Fabaceae_Pentaclethra_macroloba
Fabaceae_Andira_aubletii
Fabaceae_Vouacapoua_macropetala
Fabaceae_Arcoa_gonavensis
Fabaceae_Ceratonia_siliqua
Fabaceae_Tetrapterocarpon_geayi
Fabaceae_Acrocarpus_fraxinifolius
Fabaceae_Umtiza_listeriana
Fabaceae_Gleditsia_sinensis
Fabaceae_Gleditsia_caspica
Fabaceae_Gleditsia_triacanthos
Fabaceae_Gymnocladus_chinensis
Fabaceae_Gymnocladus_dioicus
Fabaceae_Adesmia_emarginata
Fabaceae_Aldina_heterophylla
Fabaceae_Aldina_insignis
Fabaceae_Aldina_latifolia
Fabaceae_Schefflerodendron_usambarense
Fabaceae_Callerya_atropurpurea
Fabaceae_Callerya_vasta
Fabaceae_Callerya_australis
Fabaceae_Callerya_megasperma
Fabaceae_Afgekia_filipes
Fabaceae_Callerya_dasyphylla
Fabaceae_Wisteria_frutescens
Fabaceae_Afgekia_mahidolae
Fabaceae_Afgekia_sericea
Fabaceae_Wisteria_villosa
Fabaceae_Wisteria_sinensis
Fabaceae_Wisteria_brachybotrys
Fabaceae_Wisteria_floribunda
Fabaceae_Callerya_speciosa
Fabaceae_Callerya_cinerea
Fabaceae_Millettia_japonica

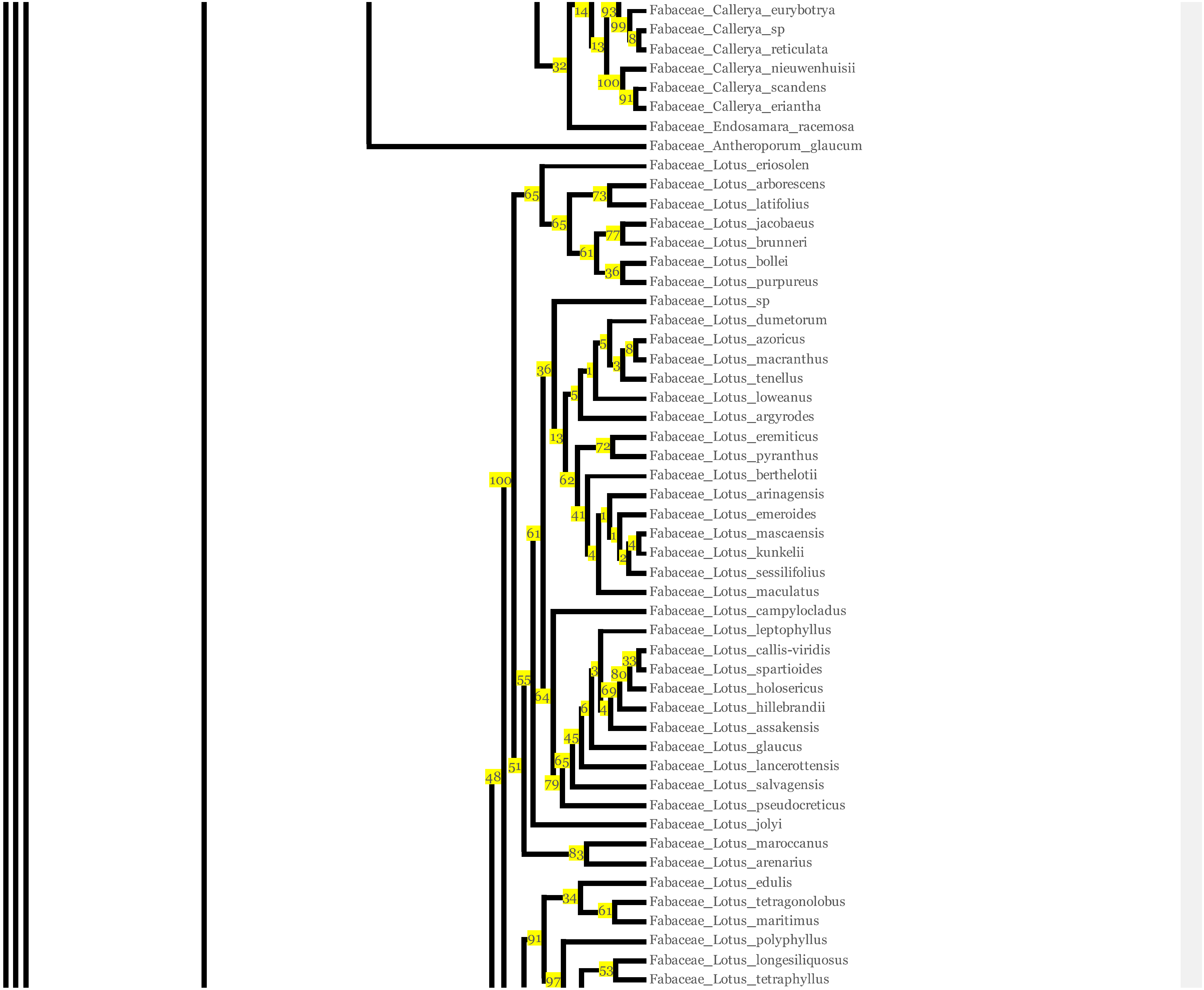
Fabaceae_Callerya_eurybotrya
Fabaceae_Callerya_sp
Fabaceae_Callerya_reticulata
Fabaceae_Callerya_nieuwenhuisii
Fabaceae_Callerya_scandens
Fabaceae_Callerya_eriantha
Fabaceae_Endosamara_racemosa
Fabaceae_Antheroporum_glaucum
Fabaceae_Lotus_eriosolen
Fabaceae_Lotus_arborescens
Fabaceae_Lotus_latifolius
Fabaceae_Lotus_jacobaeus
Fabaceae_Lotus_brunneri
Fabaceae_Lotus_bollei
Fabaceae_Lotus_purpureus
Fabaceae_Lotus_sp
Fabaceae_Lotus_dumetorum
Fabaceae_Lotus_azoricus
Fabaceae_Lotus_macranthus
Fabaceae_Lotus_tenellus
Fabaceae_Lotus_loweanus
Fabaceae_Lotus_argyrodes
Fabaceae_Lotus_eremiticus
Fabaceae_Lotus_pyranthus
Fabaceae_Lotus_berthelotii
Fabaceae_Lotus_arinagensis
Fabaceae_Lotus_emeroides
Fabaceae_Lotus_mascaensis
Fabaceae_Lotus_kunkelii
Fabaceae_Lotus_sessilifolius
Fabaceae_Lotus_maculatus
Fabaceae_Lotus_campylocladus
Fabaceae_Lotus_leptophyllus
Fabaceae_Lotus_callis-viridis
Fabaceae_Lotus_spartioides
Fabaceae_Lotus_holosericus
Fabaceae_Lotus_hillebrandii
Fabaceae_Lotus_assakensis
Fabaceae_Lotus_glaucus
Fabaceae_Lotus_lancerottensis
Fabaceae_Lotus_salvagensis
Fabaceae_Lotus_pseudocreticus
Fabaceae_Lotus_jolyi
Fabaceae_Lotus_maroccanus
Fabaceae_Lotus_arenarius
Fabaceae_Lotus_edulis
Fabaceae_Lotus_tetragonolobus
Fabaceae_Lotus_maritimus
Fabaceae_Lotus_polyphyllus
Fabaceae_Lotus_longesiliquosus
Fabaceae_Lotus_tetraphyllus

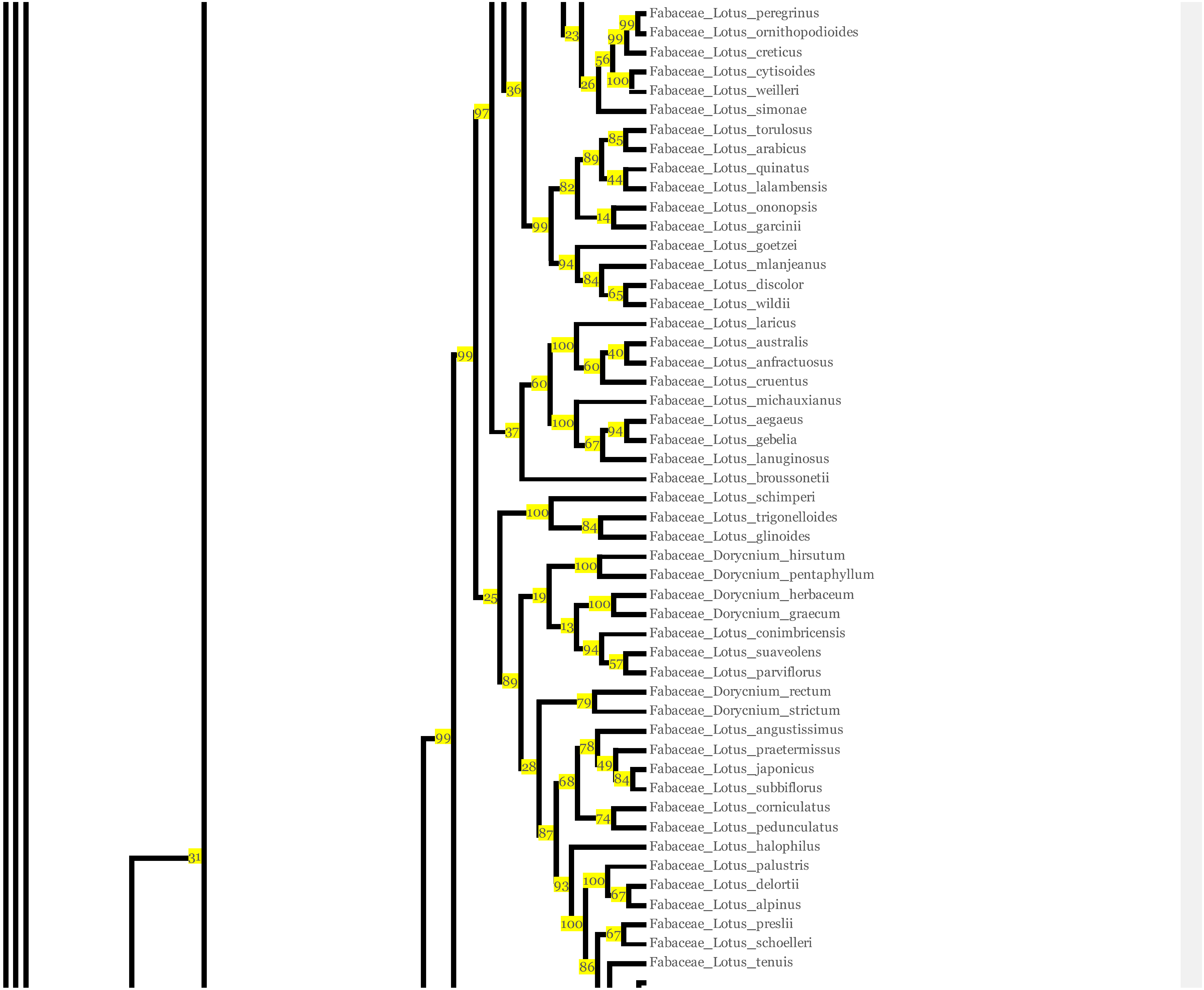
Fabaceae_Lotus_peregrinus
Fabaceae_Lotus_ornithopodioides
Fabaceae_Lotus_creticus
Fabaceae_Lotus_cytisoides
Fabaceae_Lotus_weilleri
Fabaceae_Lotus_simonae
Fabaceae_Lotus_torulosus
Fabaceae_Lotus_arabicus
Fabaceae_Lotus_quinatus
Fabaceae_Lotus_lalambensis
Fabaceae_Lotus_ononopsis
Fabaceae_Lotus_garcinii
Fabaceae_Lotus_goetzei
Fabaceae_Lotus_mlanjeanus
Fabaceae_Lotus_discolor
Fabaceae_Lotus_wildii
Fabaceae_Lotus_laricus
Fabaceae_Lotus_australis
Fabaceae_Lotus_anfractuosus
Fabaceae_Lotus_cruentus
Fabaceae_Lotus_michauxianus
Fabaceae_Lotus_aegaeus
Fabaceae_Lotus_gebelia
Fabaceae_Lotus_lanuginosus
Fabaceae_Lotus_broussonetii
Fabaceae_Lotus_schimperi
Fabaceae_Lotus_trigonelloides
Fabaceae_Lotus_glinoides
Fabaceae_Dorycnium_hirsutum
Fabaceae_Dorycnium_pentaphyllum
Fabaceae_Dorycnium_herbaceum
Fabaceae_Dorycnium_graecum
Fabaceae_Lotus_conimbricensis
Fabaceae_Lotus_suaveolens
Fabaceae_Lotus_parviflorus
Fabaceae_Dorycnium_rectum
Fabaceae_Dorycnium_strictum
Fabaceae_Lotus_angustissimus
Fabaceae_Lotus_praetermissus
Fabaceae_Lotus_japonicus
Fabaceae_Lotus_subbiflorus
Fabaceae_Lotus_corniculatus
Fabaceae_Lotus_pedunculatus
Fabaceae_Lotus_halophilus
Fabaceae_Lotus_palustris
Fabaceae_Lotus_delortii
Fabaceae_Lotus_alpinus
Fabaceae_Lotus_preslii
Fabaceae_Lotus_schoelleri
Fabaceae_Lotus_tenuis

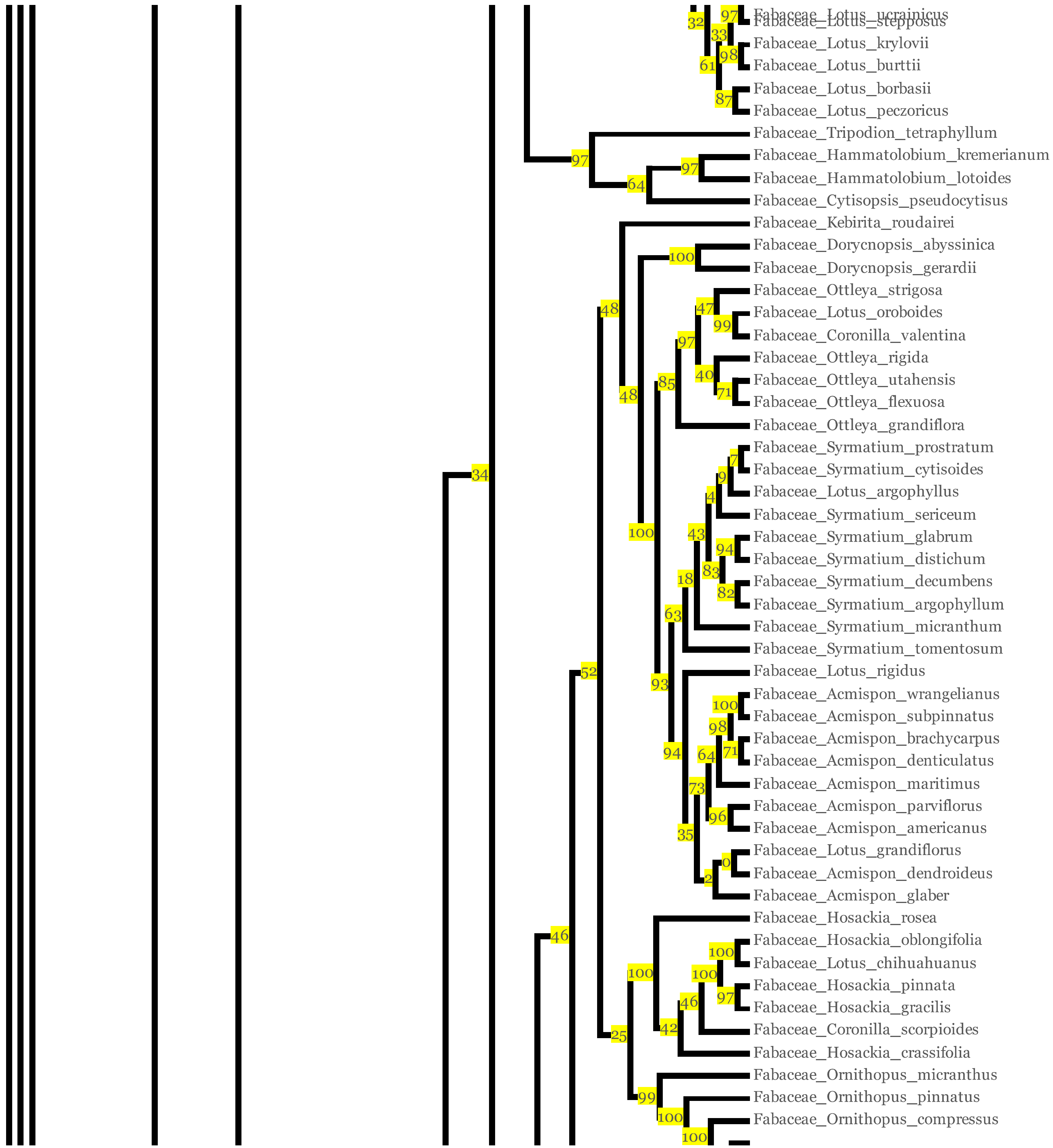
Fabaceae_Lotus_ucrainicus
Fabaceae_Lotus_stepposus
Fabaceae_Lotus_krylovii
Fabaceae_Lotus_burttii
Fabaceae_Lotus_borbasii
Fabaceae_Lotus_peczoricus
Fabaceae_Tripodion_tetraphyllum
Fabaceae_Hammatolobium_kremerianum
Fabaceae_Hammatolobium_lotoides
Fabaceae_Cytisopsis_pseudocytisus
Fabaceae_Kebirita_roudairei
Fabaceae_Dorycnopsis_abyssinica
Fabaceae_Dorycnopsis_gerardii
Fabaceae_Ottleya_strigosa
Fabaceae_Lotus_oroboides
Fabaceae_Coronilla_valentina
Fabaceae_Ottleya_rigida
Fabaceae_Ottleya_utahensis
Fabaceae_Ottleya_flexuosa
Fabaceae_Ottleya_grandiflora
Fabaceae_Syrmatium_prostratum
Fabaceae_Syrmatium_cytisoides
Fabaceae_Lotus_argophyllus
Fabaceae_Syrmatium_sericeum
Fabaceae_Syrmatium_glabrum
Fabaceae_Syrmatium_distichum
Fabaceae_Syrmatium_decumbens
Fabaceae_Syrmatium_argophyllum
Fabaceae_Syrmatium_micranthum
Fabaceae_Syrmatium_tomentosum
Fabaceae_Lotus_rigidus
Fabaceae_Acmispon_wrangelianus
Fabaceae_Acmispon_subpinnatus
Fabaceae_Acmispon_brachycarpus
Fabaceae_Acmispon_denticulatus
Fabaceae_Acmispon_maritimus
Fabaceae_Acmispon_parviflorus
Fabaceae_Acmispon_americanus
Fabaceae_Lotus_grandiflorus
Fabaceae_Acmispon_dendroideus
Fabaceae_Acmispon_glaber
Fabaceae_Hosackia_rosea
Fabaceae_Hosackia_oblongifolia
Fabaceae_Lotus_chihuahuanus
Fabaceae_Hosackia_pinnata
Fabaceae_Hosackia_gracilis
Fabaceae_Coronilla_scorpioides
Fabaceae_Hosackia_crassifolia
Fabaceae_Ornithopus_micranthus
Fabaceae_Ornithopus_pinnatus
Fabaceae_Ornithopus_compressus

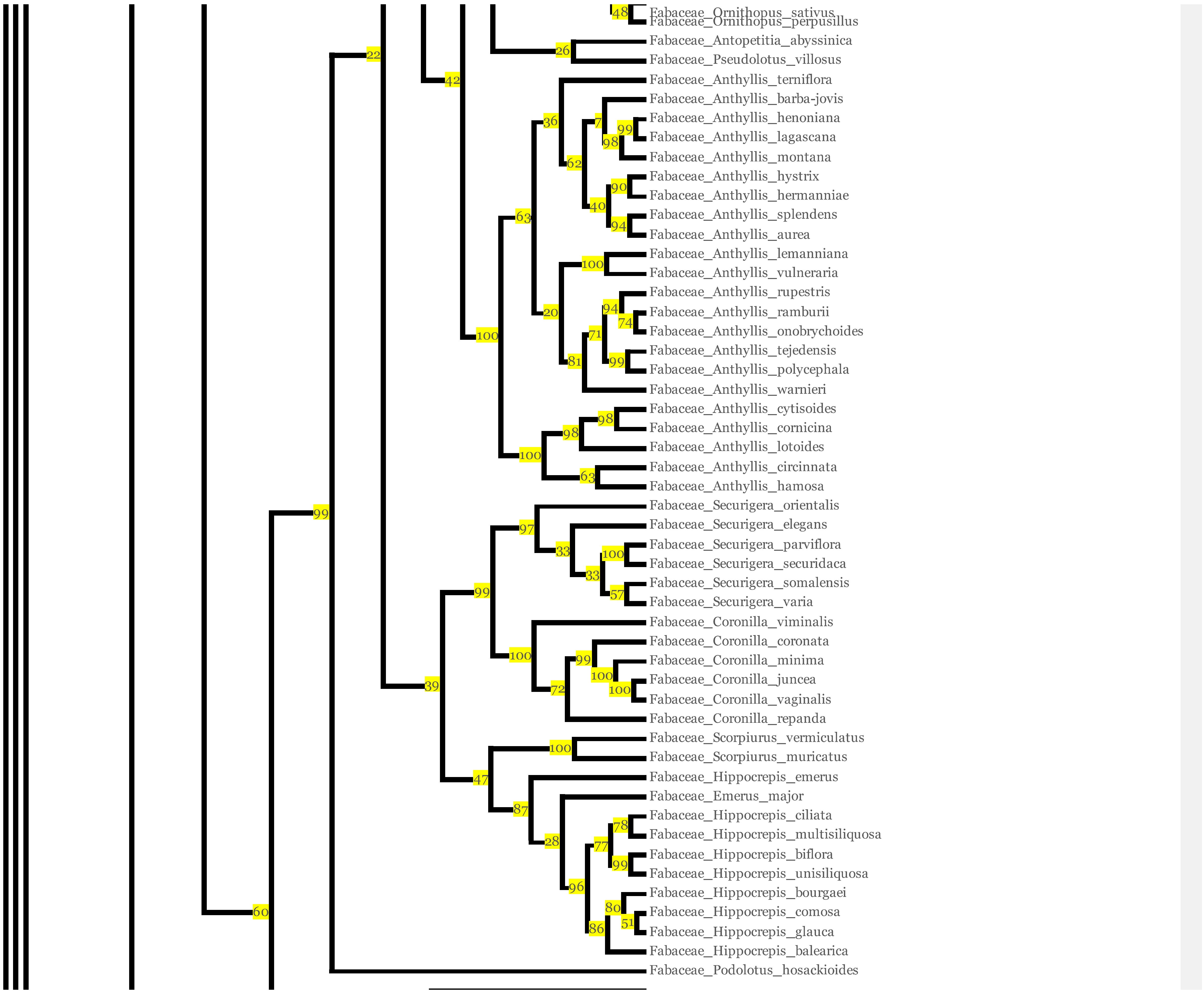
Fabaceae_Ornithopus_sativus
Fabaceae_Ornithopus_perpusillus
Fabaceae_Antopetitia_abyssinica
Fabaceae_Pseudolotus_villosus
Fabaceae_Anthyllis_terniflora
Fabaceae_Anthyllis_barba-jovis
Fabaceae_Anthyllis_henoniana
Fabaceae_Anthyllis_lagascana
Fabaceae_Anthyllis_montana
Fabaceae_Anthyllis_hystrix
Fabaceae_Anthyllis_hermanniae
Fabaceae_Anthyllis_splendens
Fabaceae_Anthyllis_aurea
Fabaceae_Anthyllis_lemanniana
Fabaceae_Anthyllis_vulneraria
Fabaceae_Anthyllis_rupestris
Fabaceae_Anthyllis_ramburii
Fabaceae_Anthyllis_onobrychoides
Fabaceae_Anthyllis_tejedensis
Fabaceae_Anthyllis_polycephala
Fabaceae_Anthyllis_warnieri
Fabaceae_Anthyllis_cytisoides
Fabaceae_Anthyllis_cornicina
Fabaceae_Anthyllis_lotoides
Fabaceae_Anthyllis_circinnata
Fabaceae_Anthyllis_hamosa
Fabaceae_Securigera_orientalis
Fabaceae_Securigera_elegans
Fabaceae_Securigera_parviflora
Fabaceae_Securigera_securidaca
Fabaceae_Securigera_somalensis
Fabaceae_Securigera_varia
Fabaceae_Coronilla_viminalis
Fabaceae_Coronilla_coronata
Fabaceae_Coronilla_minima
Fabaceae_Coronilla_juncea
Fabaceae_Coronilla_vaginalis
Fabaceae_Coronilla_repanda
Fabaceae_Scorpiurus_vermiculatus
Fabaceae_Scorpiurus_muricatus
Fabaceae_Hippocrepis_emerus
Fabaceae_Emerus_major
Fabaceae_Hippocrepis_ciliata
Fabaceae_Hippocrepis_multisiliquosa
Fabaceae_Hippocrepis_biflora
Fabaceae_Hippocrepis_unisiliquosa
Fabaceae_Hippocrepis_bourgaei
Fabaceae_Hippocrepis_comosa
Fabaceae_Hippocrepis_glauca
Fabaceae_Hippocrepis_balearica
Fabaceae_Podolotus_hosackioides

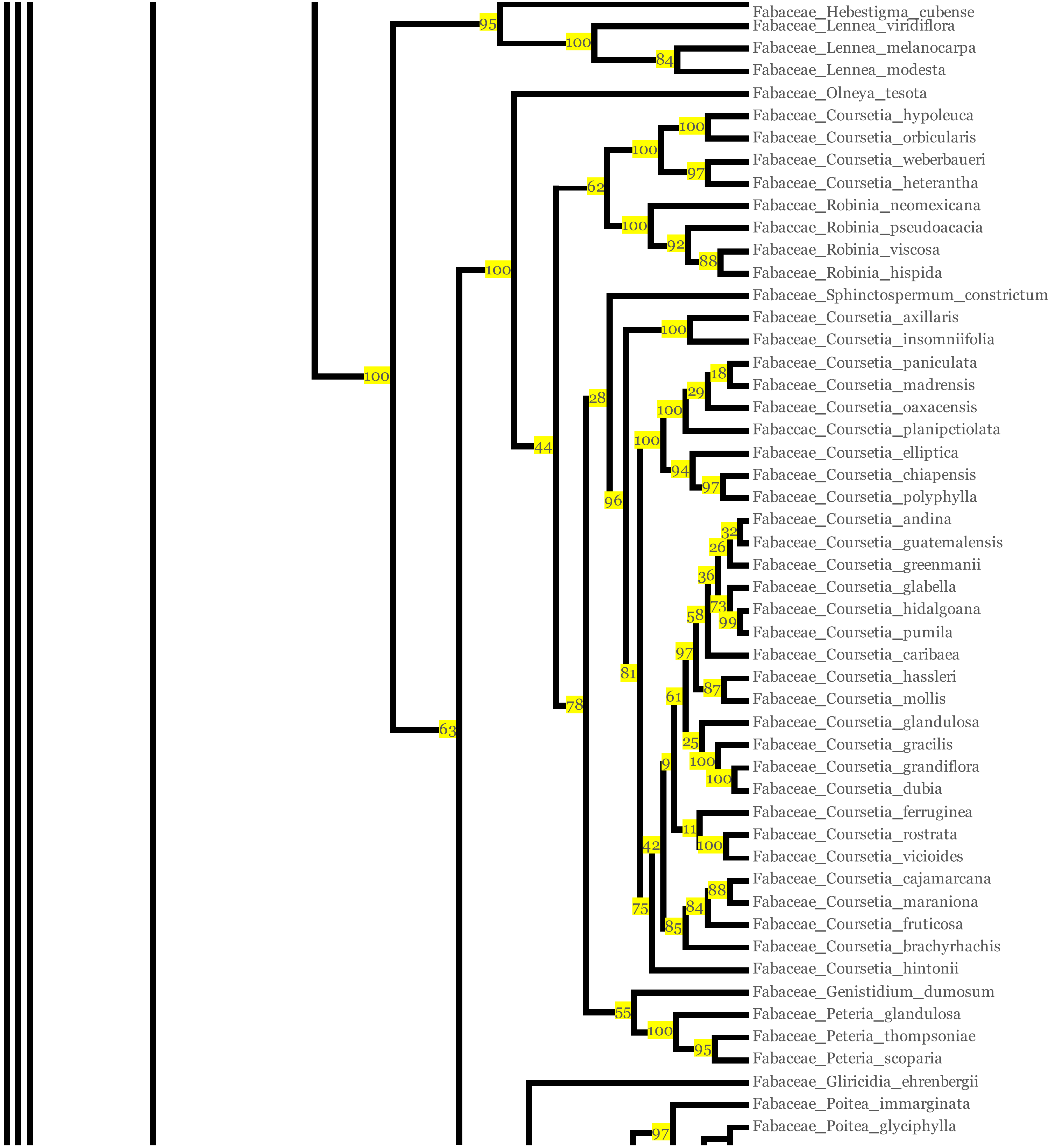
Fabaceae_Hebestigma_cubense
Fabaceae_Lennea_viridiflora
Fabaceae_Lennea_melanocarpa
Fabaceae_Lennea_modesta
Fabaceae_Olneya_tesota
Fabaceae_Coursetia_hypoleuca
Fabaceae_Coursetia_orbicularis
Fabaceae_Coursetia_weberbaueri
Fabaceae_Coursetia_heterantha
Fabaceae_Robinia_neomexicana
Fabaceae_Robinia_pseudoacacia
Fabaceae_Robinia_viscosa
Fabaceae_Robinia_hispida
Fabaceae_Sphinctospermum_constrictum
Fabaceae_Coursetia_axillaris
Fabaceae_Coursetia_insomniifolia
Fabaceae_Coursetia_paniculata
Fabaceae_Coursetia_madrensis
Fabaceae_Coursetia_oaxacensis
Fabaceae_Coursetia_planipetiolata
Fabaceae_Coursetia_elliptica
Fabaceae_Coursetia_chiapensis
Fabaceae_Coursetia_polyphylla
Fabaceae_Coursetia_andina
Fabaceae_Coursetia_guatemalensis
Fabaceae_Coursetia_greenmanii
Fabaceae_Coursetia_glabella
Fabaceae_Coursetia_hidalgoana
Fabaceae_Coursetia_pumila
Fabaceae_Coursetia_caribaea
Fabaceae_Coursetia_hassleri
Fabaceae_Coursetia_mollis
Fabaceae_Coursetia_glandulosa
Fabaceae_Coursetia_gracilis
Fabaceae_Coursetia_grandiflora
Fabaceae_Coursetia_dubia
Fabaceae_Coursetia_ferruginea
Fabaceae_Coursetia_rostrata
Fabaceae_Coursetia_vicioides
Fabaceae_Coursetia_cajamarcana
Fabaceae_Coursetia_maraniona
Fabaceae_Coursetia_fruticosa
Fabaceae_Coursetia_brachyrhachis
Fabaceae_Coursetia_hintonii
Fabaceae_Genistidium_dumosum
Fabaceae_Peteria_glandulosa
Fabaceae_Peteria_thompsoniae
Fabaceae_Peteria_scoparia
Fabaceae_Gliricidia_ehrenbergii
Fabaceae_Poitea_immarginata
Fabaceae_Poitea_glyciphylla

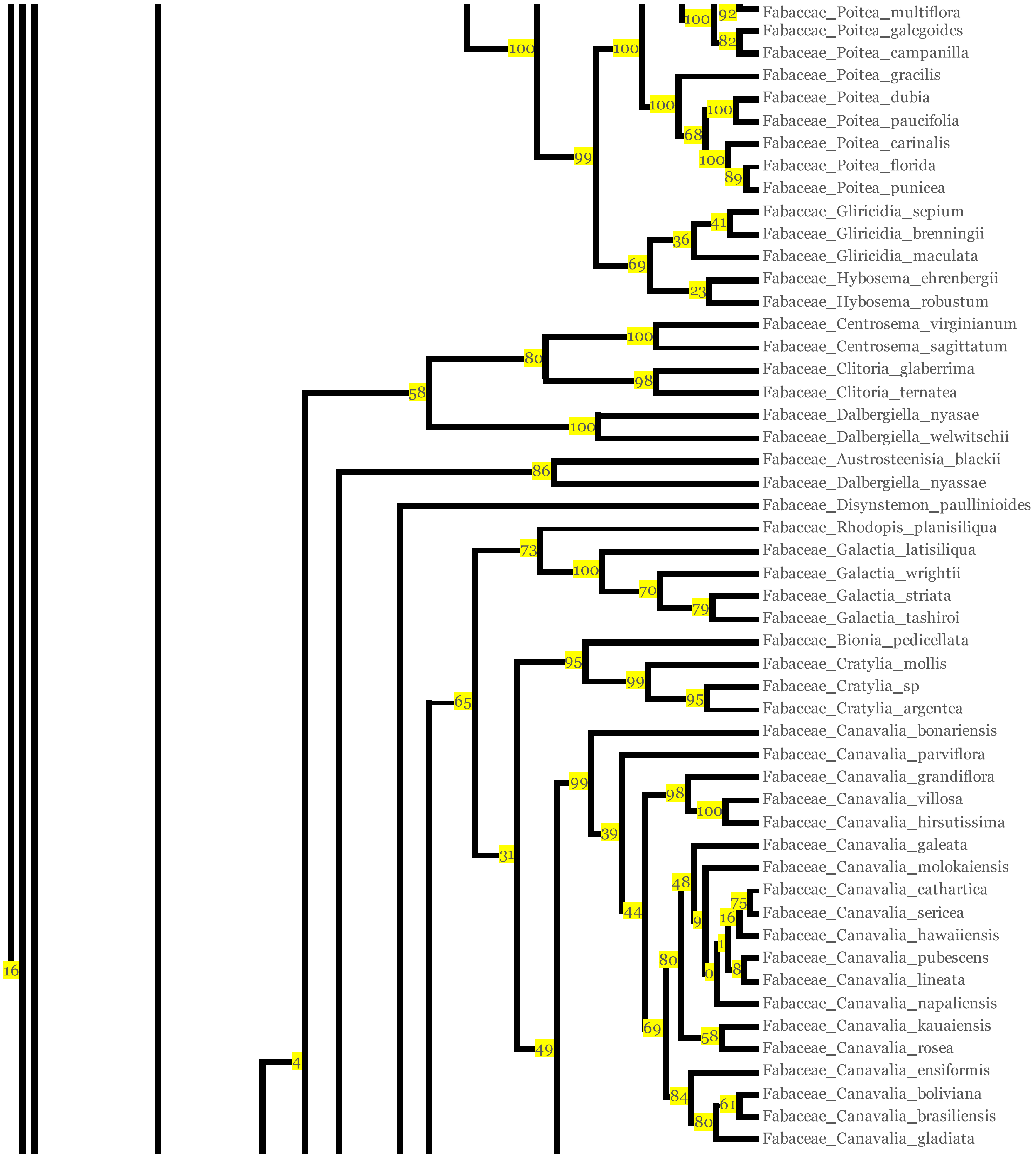
Fabaceae_Poitea_multiflora
Fabaceae_Poitea_galegoides
Fabaceae_Poitea_campanilla
Fabaceae_Poitea_gracilis
Fabaceae_Poitea_dubia
Fabaceae_Poitea_paucifolia
Fabaceae_Poitea_carinalis
Fabaceae_Poitea_florida
Fabaceae_Poitea_punicea
Fabaceae_Gliricidia_sepium
Fabaceae_Gliricidia_brenningii
Fabaceae_Gliricidia_maculata
Fabaceae_Hybosema_ehrenbergii
Fabaceae_Hybosema_robustum
Fabaceae_Centrosema_virginianum
Fabaceae_Centrosema_sagittatum
Fabaceae_Clitoria_glaberrima
Fabaceae_Clitoria_ternatea
Fabaceae_Dalbergiella_nyasae
Fabaceae_Dalbergiella_welwitschii
Fabaceae_Austrosteenisia_blackii
Fabaceae_Dalbergiella_nyassae
Fabaceae_Disynstemon_paullinioides
Fabaceae_Rhodopis_planisiliqua
Fabaceae_Galactia_latisiliqua
Fabaceae_Galactia_wrightii
Fabaceae_Galactia_striata
Fabaceae_Galactia_tashiroi
Fabaceae_Bionia_pedicellata
Fabaceae_Cratylia_mollis
Fabaceae_Cratylia_sp
Fabaceae_Cratylia_argentea
Fabaceae_Canavalia_bonariensis
Fabaceae_Canavalia_parviflora
Fabaceae_Canavalia_grandiflora
Fabaceae_Canavalia_villosa
Fabaceae_Canavalia_hirsutissima
Fabaceae_Canavalia_galeata
Fabaceae_Canavalia_molokaiensis
Fabaceae_Canavalia_cathartica
Fabaceae_Canavalia_sericea
Fabaceae_Canavalia_hawaiiensis
Fabaceae_Canavalia_pubescens
Fabaceae_Canavalia_lineata
Fabaceae_Canavalia_napaliensis
Fabaceae_Canavalia_kauaiensis
Fabaceae_Canavalia_rosea
Fabaceae_Canavalia_ensiformis
Fabaceae_Canavalia_boliviana
Fabaceae_Canavalia_brasiliensis
Fabaceae_Canavalia_gladiata

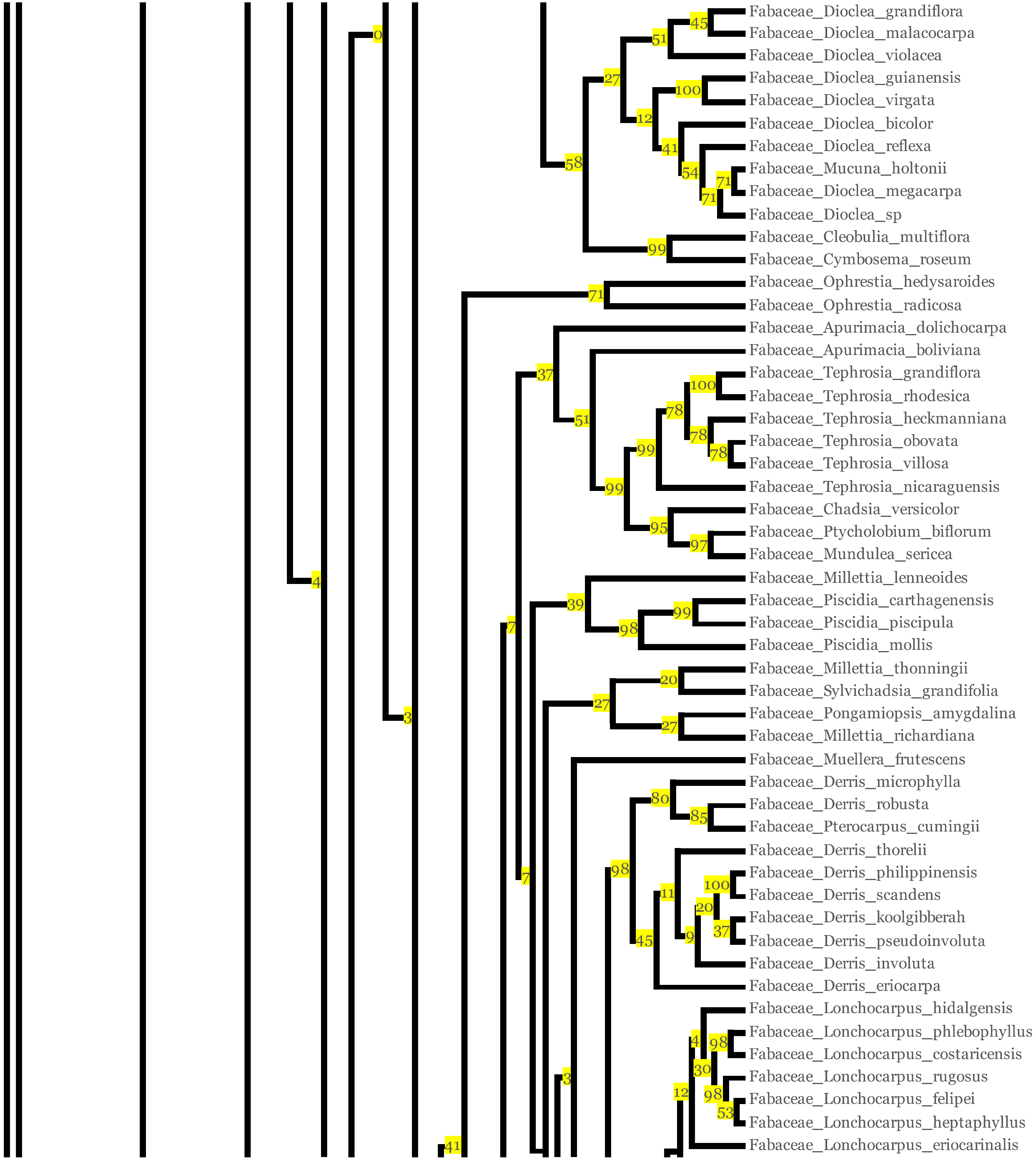
Fabaceae_Dioclea_grandiflora
Fabaceae_Dioclea_malacocarpa
Fabaceae_Dioclea_violacea
Fabaceae_Dioclea_guianensis
Fabaceae_Dioclea_virgata
Fabaceae_Dioclea_bicolor
Fabaceae_Dioclea_reflexa
Fabaceae_Mucuna_holtonii
Fabaceae_Dioclea_megacarpa
Fabaceae_Dioclea_sp
Fabaceae_Cleobulia_multiflora
Fabaceae_Cymbosema_roseum
Fabaceae_Ophrestia_hedysaroides
Fabaceae_Ophrestia_radicosa
Fabaceae_Apurimacia_dolichocarpa
Fabaceae_Apurimacia_boliviana
Fabaceae_Tephrosia_grandiflora
Fabaceae_Tephrosia_rhodesica
Fabaceae_Tephrosia_heckmanniana
Fabaceae_Tephrosia_obovata
Fabaceae_Tephrosia_villosa
Fabaceae_Tephrosia_nicaraguensis
Fabaceae_Chadsia_versicolor
Fabaceae_Ptycholobium_biflorum
Fabaceae_Mundulea_sericea
Fabaceae_Millettia_lenneoides
Fabaceae_Piscidia_carthagenensis
Fabaceae_Piscidia_piscipula
Fabaceae_Piscidia_mollis
Fabaceae_Millettia_thonningii
Fabaceae_Sylvichadsia_grandifolia
Fabaceae_Pongamiopsis_amygdalina
Fabaceae_Millettia_richardiana
Fabaceae_Muellera_frutescens
Fabaceae_Derris_microphylla
Fabaceae_Derris_robusta
Fabaceae_Pterocarpus_cumingii
Fabaceae_Derris_thorelii
Fabaceae_Derris_philippinensis
Fabaceae_Derris_scandens
Fabaceae_Derris_koolgibberah
Fabaceae_Derris_pseudoinvoluta
Fabaceae_Derris_involuta
Fabaceae_Derris_eriocarpa
Fabaceae_Lonchocarpus_hidalgensis
Fabaceae_Lonchocarpus_phlebophyllus
Fabaceae_Lonchocarpus_costaricensis
Fabaceae_Lonchocarpus_rugosus
Fabaceae_Lonchocarpus_felipei
Fabaceae_Lonchocarpus_heptaphyllus
Fabaceae_Lonchocarpus_eriocarinalis

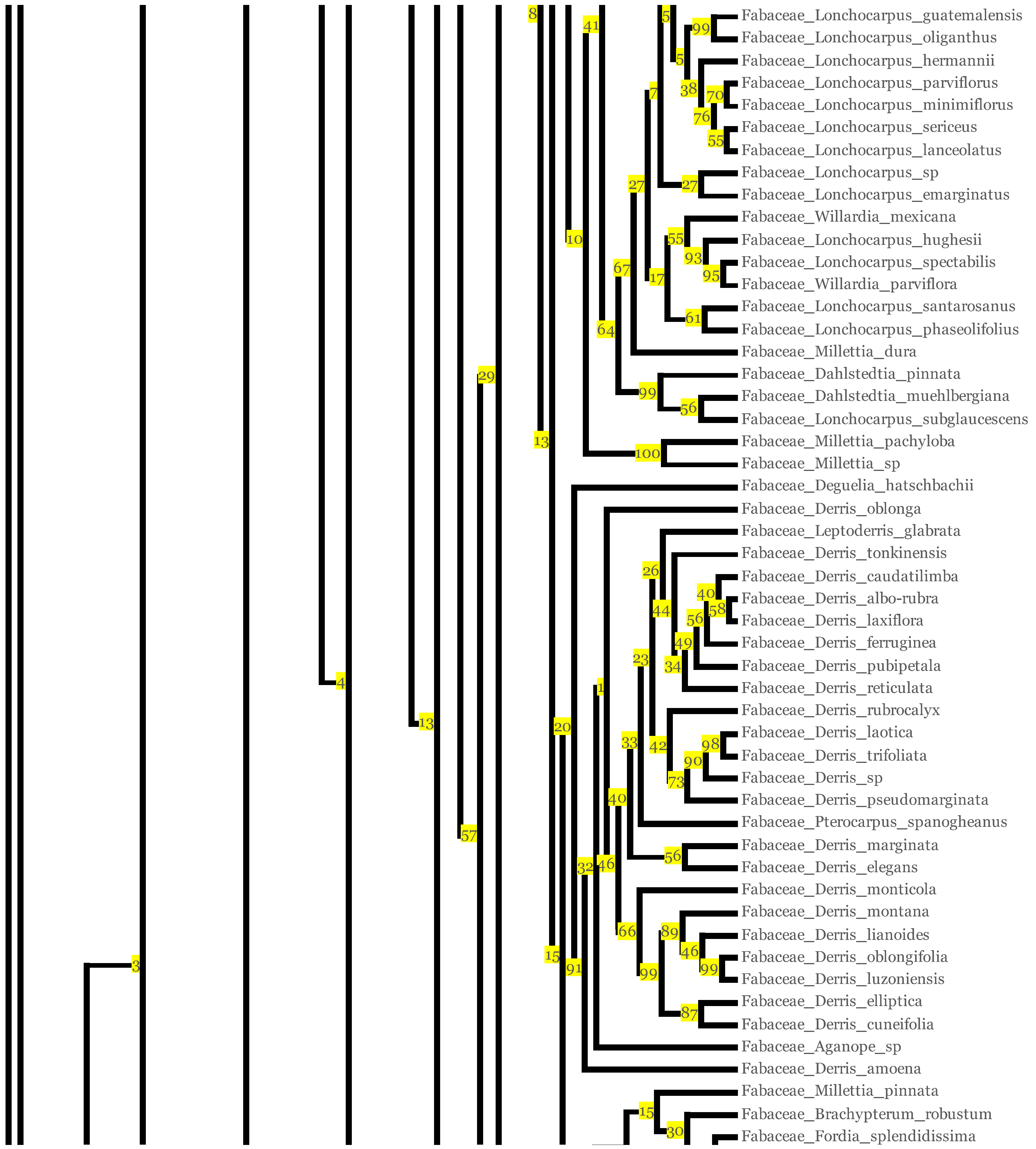
Fabaceae_Lonchocarpus_guatemalensis
Fabaceae_Lonchocarpus_oliganthus
Fabaceae_Lonchocarpus_hermannii
Fabaceae_Lonchocarpus_parviflorus
Fabaceae_Lonchocarpus_minimiflorus
Fabaceae_Lonchocarpus_sericeus
Fabaceae_Lonchocarpus_lanceolatus
Fabaceae_Lonchocarpus_sp
Fabaceae_Lonchocarpus_emarginatus
Fabaceae_Willardia_mexicana
Fabaceae_Lonchocarpus_hughesii
Fabaceae_Lonchocarpus_spectabilis
Fabaceae_Willardia_parviflora
Fabaceae_Lonchocarpus_santarosanus
Fabaceae_Lonchocarpus_phaseolifolius
Fabaceae_Millettia_dura
Fabaceae_Dahlstedtia_pinnata
Fabaceae_Dahlstedtia_muehlbergiana
Fabaceae_Lonchocarpus_subglaucescens
Fabaceae_Millettia_pachyloba
Fabaceae_Millettia_sp
Fabaceae_Deguelia_hatschbachii
Fabaceae_Derris_oblonga
Fabaceae_Leptoderris_glabrata
Fabaceae_Derris_tonkinensis
Fabaceae_Derris_caudatilimba
Fabaceae_Derris_albo-rubra
Fabaceae_Derris_laxiflora
Fabaceae_Derris_ferruginea
Fabaceae_Derris_pubipetala
Fabaceae_Derris_reticulata
Fabaceae_Derris_rubrocalyx
Fabaceae_Derris_laotica
Fabaceae_Derris_trifoliata
Fabaceae_Derris_sp
Fabaceae_Derris_pseudomarginata
Fabaceae_Pterocarpus_spanogheanus
Fabaceae_Derris_marginata
Fabaceae_Derris_elegans
Fabaceae_Derris_monticola
Fabaceae_Derris_montana
Fabaceae_Derris_lianoides
Fabaceae_Derris_oblongifolia
Fabaceae_Derris_luzoniensis
Fabaceae_Derris_elliptica
Fabaceae_Derris_cuneifolia
Fabaceae_Aganope_sp
Fabaceae_Derris_amoena
Fabaceae_Millettia_pinnata
Fabaceae_Brachypterum_robustum
Fabaceae_Fordia_splendidissima

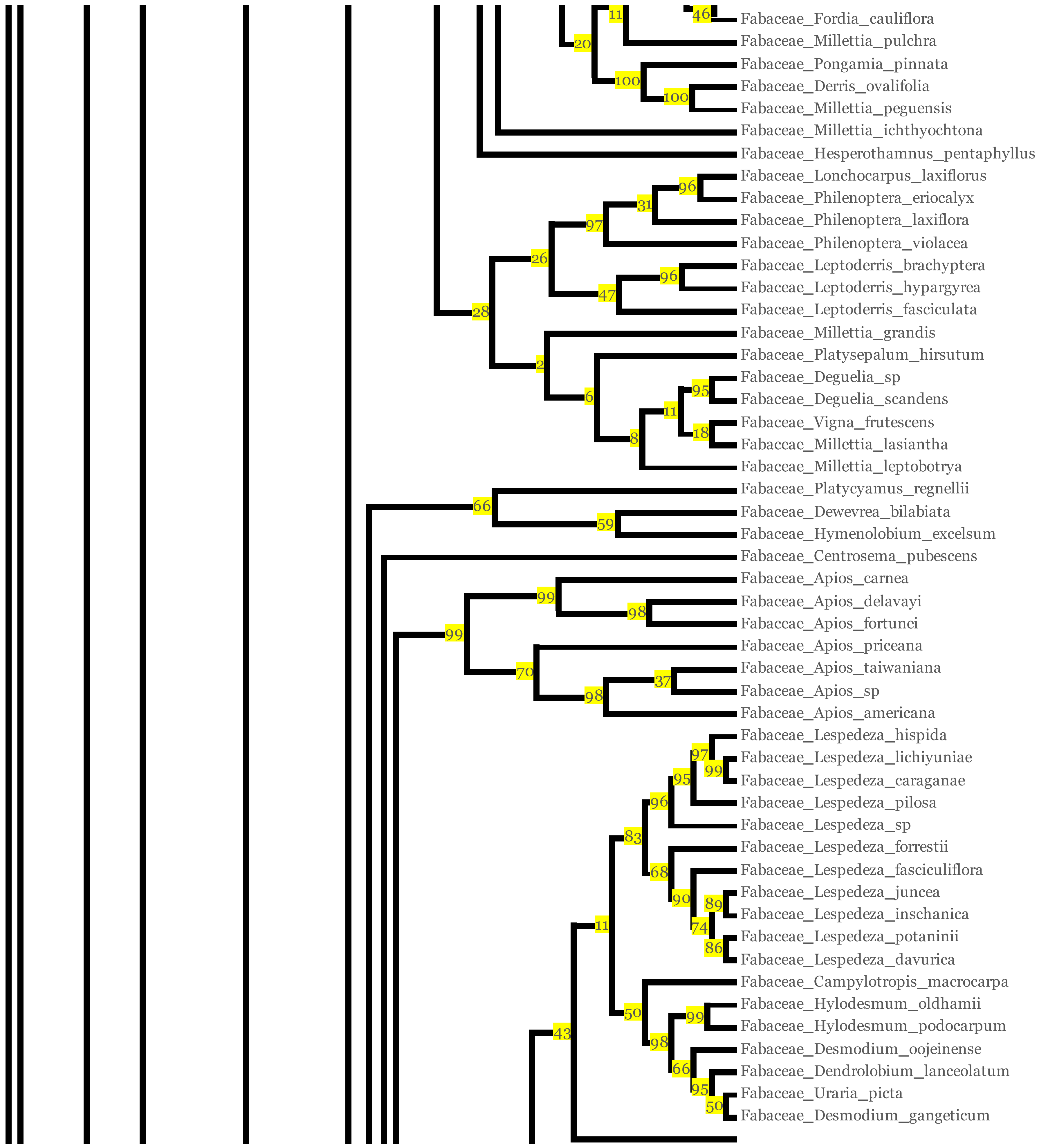
Fabaceae_Fordia_cauliflora
Fabaceae_Millettia_pulchra
Fabaceae_Pongamia_pinnata
Fabaceae_Derris_ovalifolia
Fabaceae_Millettia_peguensis
Fabaceae_Millettia_ichthyochtona
Fabaceae_Hesperothamnus_pentaphyllus
Fabaceae_Lonchocarpus_laxiflorus
Fabaceae_Philenoptera_eriocalyx
Fabaceae_Philenoptera_laxiflora
Fabaceae_Philenoptera_violacea
Fabaceae_Leptoderris_brachyptera
Fabaceae_Leptoderris_hypargyrea
Fabaceae_Leptoderris_fasciculata
Fabaceae_Millettia_grandis
Fabaceae_Platysepalum_hirsutum
Fabaceae_Deguelia_sp
Fabaceae_Deguelia_scandens
Fabaceae_Vigna_frutescens
Fabaceae_Millettia_lasiantha
Fabaceae_Millettia_leptobotrya
Fabaceae_Platycyamus_regnellii
Fabaceae_Dewevrea_bilabiata
Fabaceae_Hymenolobium_excelsum
Fabaceae_Centrosema_pubescens
Fabaceae_Apios_carnea
Fabaceae_Apios_delavayi
Fabaceae_Apios_fortunei
Fabaceae_Apios_priceana
Fabaceae_Apios_taiwaniana
Fabaceae_Apios_sp
Fabaceae_Apios_americana
Fabaceae_Lespedeza_hispida
Fabaceae_Lespedeza_lichiyuniae
Fabaceae_Lespedeza_caraganae
Fabaceae_Lespedeza_pilosa
Fabaceae_Lespedeza_sp
Fabaceae_Lespedeza_forrestii
Fabaceae_Lespedeza_fasciculiflora
Fabaceae_Lespedeza_juncea
Fabaceae_Lespedeza_inschanica
Fabaceae_Lespedeza_potaninii
Fabaceae_Lespedeza_davurica
Fabaceae_Campylotropis_macrocarpa
Fabaceae_Hylodesmum_oldhamii
Fabaceae_Hylodesmum_podocarpum
Fabaceae_Desmodium_oojeinense
Fabaceae_Dendrolobium_lanceolatum
Fabaceae_Uraria_picta
Fabaceae_Desmodium_gangeticum

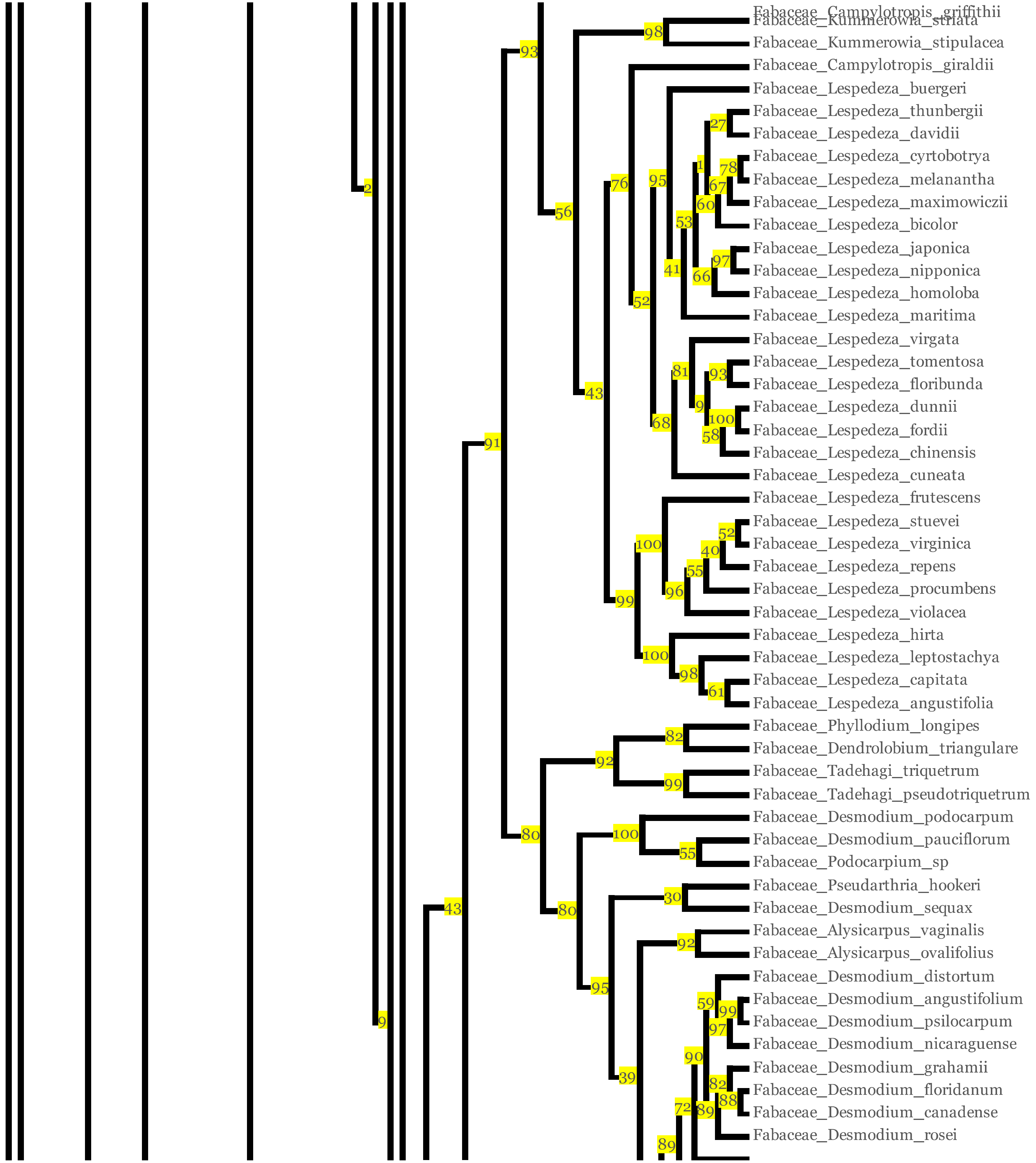
Fabaceae_Campylotropis_griffithii
Fabaceae_Kummerowia_striata
Fabaceae_Kummerowia_stipulacea
Fabaceae_Campylotropis_giraldii
Fabaceae_Lespedeza_buergeri
Fabaceae_Lespedeza_thunbergii
Fabaceae_Lespedeza_davidii
Fabaceae_Lespedeza_cyrtobotrya
Fabaceae_Lespedeza_melanantha
Fabaceae_Lespedeza_maximowiczii
Fabaceae_Lespedeza_bicolor
Fabaceae_Lespedeza_japonica
Fabaceae_Lespedeza_nipponica
Fabaceae_Lespedeza_homoloba
Fabaceae_Lespedeza_maritima
Fabaceae_Lespedeza_virgata
Fabaceae_Lespedeza_tomentosa
Fabaceae_Lespedeza_floribunda
Fabaceae_Lespedeza_dunnii
Fabaceae_Lespedeza_fordii
Fabaceae_Lespedeza_chinensis
Fabaceae_Lespedeza_cuneata
Fabaceae_Lespedeza_frutescens
Fabaceae_Lespedeza_stuevei
Fabaceae_Lespedeza_virginica
Fabaceae_Lespedeza_repens
Fabaceae_Lespedeza_procumbens
Fabaceae_Lespedeza_violacea
Fabaceae_Lespedeza_hirta
Fabaceae_Lespedeza_leptostachya
Fabaceae_Lespedeza_capitata
Fabaceae_Lespedeza_angustifolia
Fabaceae_Phyllodium_longipes
Fabaceae_Dendrolobium_triangulare
Fabaceae_Tadehagi_triquetrum
Fabaceae_Tadehagi_pseudotriquetrum
Fabaceae_Desmodium_podocarpum
Fabaceae_Desmodium_pauciflorum
Fabaceae_Podocarpium_sp
Fabaceae_Pseudarthria_hookeri
Fabaceae_Desmodium_sequax
Fabaceae_Alysicarpus_vaginalis
Fabaceae_Alysicarpus_ovalifolius
Fabaceae_Desmodium_distortum
Fabaceae_Desmodium_angustifolium
Fabaceae_Desmodium_psilocarpum
Fabaceae_Desmodium_nicaraguense
Fabaceae_Desmodium_grahamii
Fabaceae_Desmodium_floridanum
Fabaceae_Desmodium_canadense
Fabaceae_Desmodium_rosei

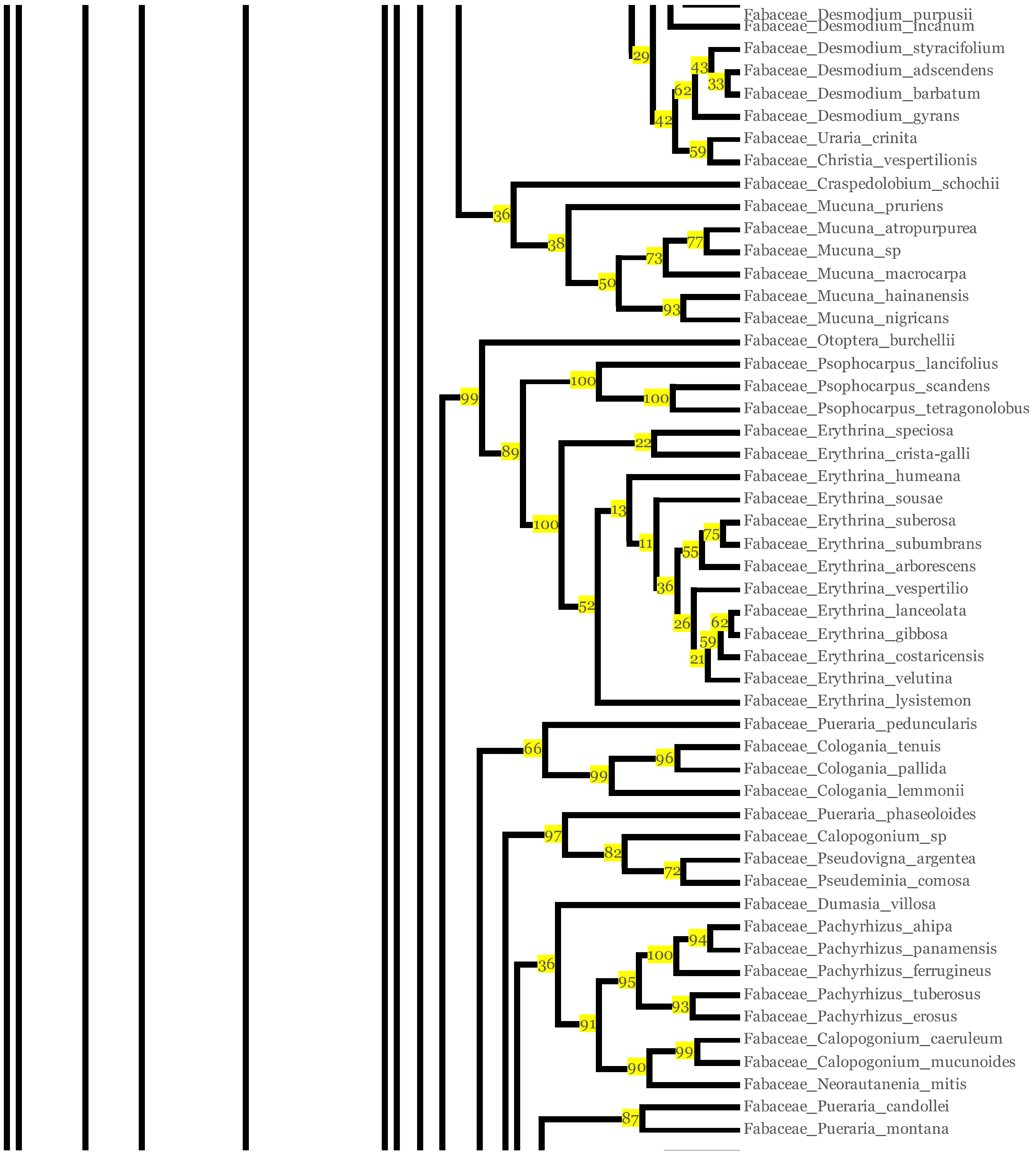
Fabaceae_Desmodium_purpusii
Fabaceae_Desmodium_incanum
Fabaceae_Desmodium_styracifolium
Fabaceae_Desmodium_adscendens
Fabaceae_Desmodium_barbatum
Fabaceae_Desmodium_gyrans
Fabaceae_Uraria_crinita
Fabaceae_Christia_vespertilionis
Fabaceae_Craspedolobium_schochii
Fabaceae_Mucuna_pruriens
Fabaceae_Mucuna_atropurpurea
Fabaceae_Mucuna_sp
Fabaceae_Mucuna_macrocarpa
Fabaceae_Mucuna_hainanensis
Fabaceae_Mucuna_nigricans
Fabaceae_Otoptera_burchellii
Fabaceae_Psophocarpus_lancifolius
Fabaceae_Psophocarpus_scandens
Fabaceae_Psophocarpus_tetragonolobus
Fabaceae_Erythrina_speciosa
Fabaceae_Erythrina_crista-galli
Fabaceae_Erythrina_humeana
Fabaceae_Erythrina_sousae
Fabaceae_Erythrina_suberosa
Fabaceae_Erythrina_subumbrans
Fabaceae_Erythrina_arborescens
Fabaceae_Erythrina_vespertilio
Fabaceae_Erythrina_lanceolata
Fabaceae_Erythrina_gibbosa
Fabaceae_Erythrina_costaricensis
Fabaceae_Erythrina_velutina
Fabaceae_Erythrina_lysistemon
Fabaceae_Pueraria_peduncularis
Fabaceae_Cologania_tenuis
Fabaceae_Cologania_pallida
Fabaceae_Cologania_lemmonii
Fabaceae_Pueraria_phaseoloides
Fabaceae_Calopogonium_sp
Fabaceae_Pseudovigna_argentea
Fabaceae_Pseudeminia_comosa
Fabaceae_Dumasia_villosa
Fabaceae_Pachyrhizus_ahipa
Fabaceae_Pachyrhizus_panamensis
Fabaceae_Pachyrhizus_ferrugineus
Fabaceae_Pachyrhizus_tuberosus
Fabaceae_Pachyrhizus_erosus
Fabaceae_Calopogonium_caeruleum
Fabaceae_Calopogonium_mucunoides
Fabaceae_Neorautanenia_mitis
Fabaceae_Pueraria_candollei
Fabaceae_Pueraria_montana

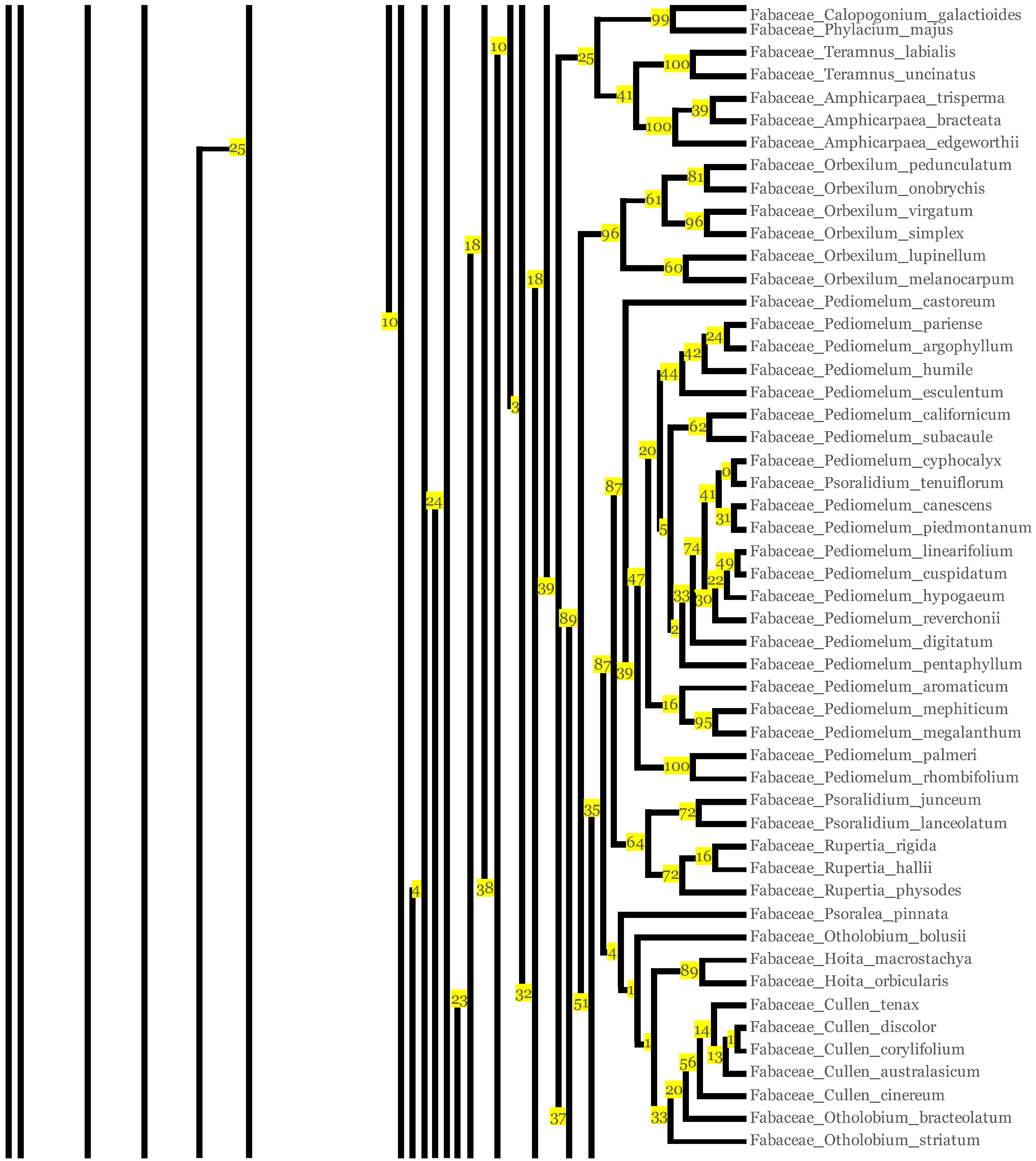
Fabaceae_Calopogonium_galactioides
Fabaceae_Phylacium_majus
Fabaceae_Teramnus_labialis
Fabaceae_Teramnus_uncinatus
Fabaceae_Amphicarpaea_trisperma
Fabaceae_Amphicarpaea_bracteata
Fabaceae_Amphicarpaea_edgeworthii
Fabaceae_Orbexilum_pedunculatum
Fabaceae_Orbexilum_onobrychis
Fabaceae_Orbexilum_virgatum
Fabaceae_Orbexilum_simplex
Fabaceae_Orbexilum_lupinellum
Fabaceae_Orbexilum_melanocarpum
Fabaceae_Pediomelum_castoreum
Fabaceae_Pediomelum_pariense
Fabaceae_Pediomelum_argophyllum
Fabaceae_Pediomelum_humile
Fabaceae_Pediomelum_esculentum
Fabaceae_Pediomelum_californicum
Fabaceae_Pediomelum_subacaule
Fabaceae_Pediomelum_cyphocalyx
Fabaceae_Psoralidium_tenuiflorum
Fabaceae_Pediomelum_canescens
Fabaceae_Pediomelum_piedmontanum
Fabaceae_Pediomelum_linearifolium
Fabaceae_Pediomelum_cuspidatum
Fabaceae_Pediomelum_hypogaeum
Fabaceae_Pediomelum_reverchonii
Fabaceae_Pediomelum_digitatum
Fabaceae_Pediomelum_pentaphyllum
Fabaceae_Pediomelum_aromaticum
Fabaceae_Pediomelum_mephiticum
Fabaceae_Pediomelum_megalanthum
Fabaceae_Pediomelum_palmeri
Fabaceae_Pediomelum_rhombifolium
Fabaceae_Psoralidium_junceum
Fabaceae_Psoralidium_lanceolatum
Fabaceae_Rupertia_rigida
Fabaceae_Rupertia_hallii
Fabaceae_Rupertia_physodes
Fabaceae_Psoralea_pinnata
Fabaceae_Otholobium_bolusii
Fabaceae_Hoita_macrostachya
Fabaceae_Hoita_orbicularis
Fabaceae_Cullen_tenax
Fabaceae_Cullen_discolor
Fabaceae_Cullen_corylifolium
Fabaceae_Cullen_australasicum
Fabaceae_Cullen_cinereum
Fabaceae_Otholobium_bracteolatum
Fabaceae_Otholobium_striatum

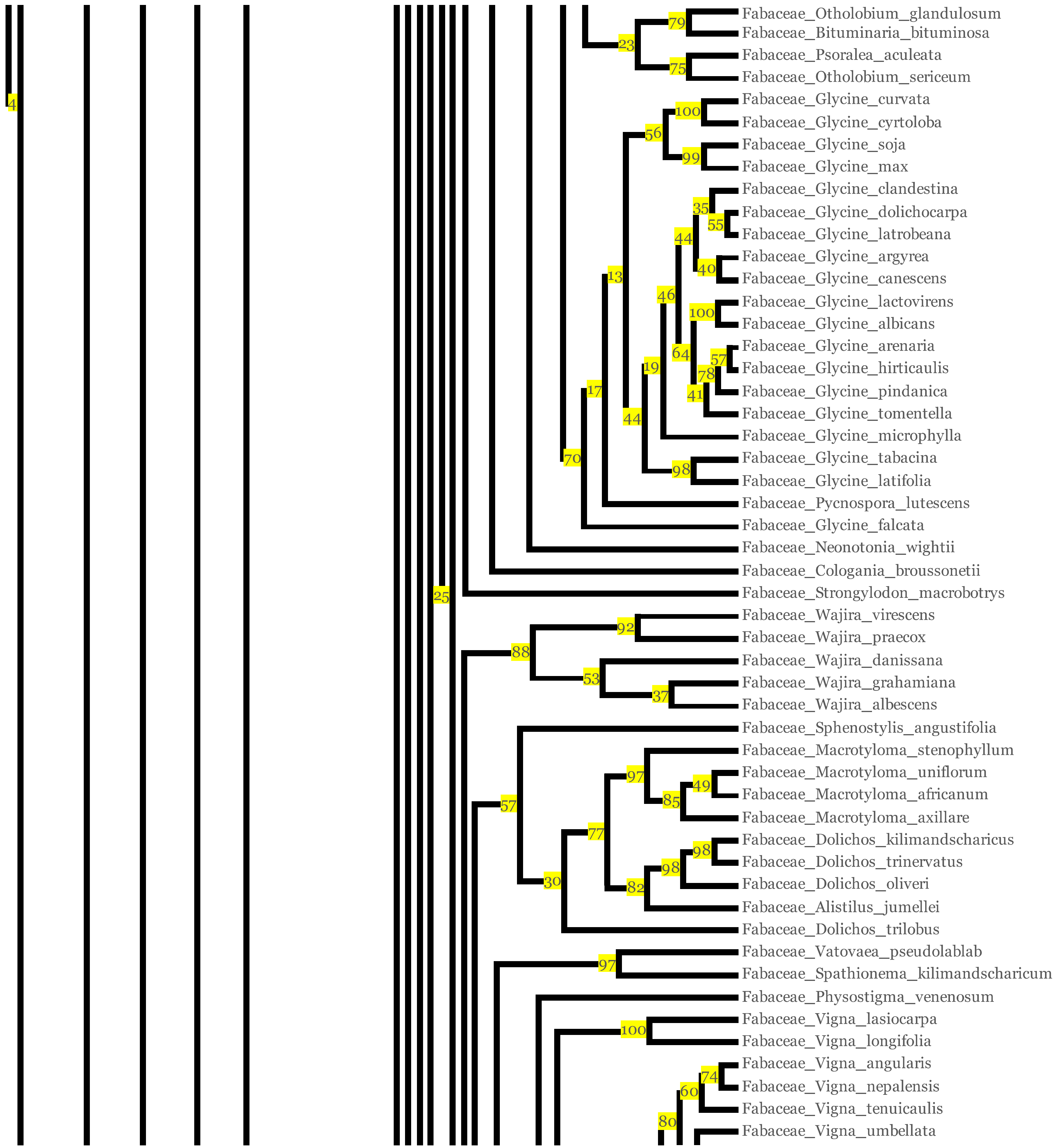
Fabaceae_Otholobium_glandulosum
Fabaceae_Bituminaria_bituminosa
Fabaceae_Psoralea_aculeata
Fabaceae_Otholobium_sericeum
Fabaceae_Glycine_curvata
Fabaceae_Glycine_cyrtoloba
Fabaceae_Glycine_soja
Fabaceae_Glycine_max
Fabaceae_Glycine_clandestina
Fabaceae_Glycine_dolichocarpa
Fabaceae_Glycine_latrobeana
Fabaceae_Glycine_argyrea
Fabaceae_Glycine_canescens
Fabaceae_Glycine_lactovirens
Fabaceae_Glycine_albicans
Fabaceae_Glycine_arenaria
Fabaceae_Glycine_hirticaulis
Fabaceae_Glycine_pindanica
Fabaceae_Glycine_tomentella
Fabaceae_Glycine_microphylla
Fabaceae_Glycine_tabacina
Fabaceae_Glycine_latifolia
Fabaceae_Pycnospora_lutescens
Fabaceae_Glycine_falcata
Fabaceae_Neonotonia_wightii
Fabaceae_Cologania_broussonetii
Fabaceae_Strongylodon_macrobotrys
Fabaceae_Wajira_virescens
Fabaceae_Wajira_praecox
Fabaceae_Wajira_danissana
Fabaceae_Wajira_grahamiana
Fabaceae_Wajira_albescens
Fabaceae_Sphenostylis_angustifolia
Fabaceae_Macrotyloma_stenophyllum
Fabaceae_Macrotyloma_uniflorum
Fabaceae_Macrotyloma_africanum
Fabaceae_Macrotyloma_axillare
Fabaceae_Dolichos_kilimandscharicus
Fabaceae_Dolichos_trinervatus
Fabaceae_Dolichos_oliveri
Fabaceae_Alistilus_jumellei
Fabaceae_Dolichos_trilobus
Fabaceae_Vatovaea_pseudolablab
Fabaceae_Spathionema_kilimandscharicum
Fabaceae_Physostigma_venenosum
Fabaceae_Vigna_lasiocarpa
Fabaceae_Vigna_longifolia
Fabaceae_Vigna_angularis
Fabaceae_Vigna_nepalensis
Fabaceae_Vigna_tenuicaulis
Fabaceae_Vigna_umbellata

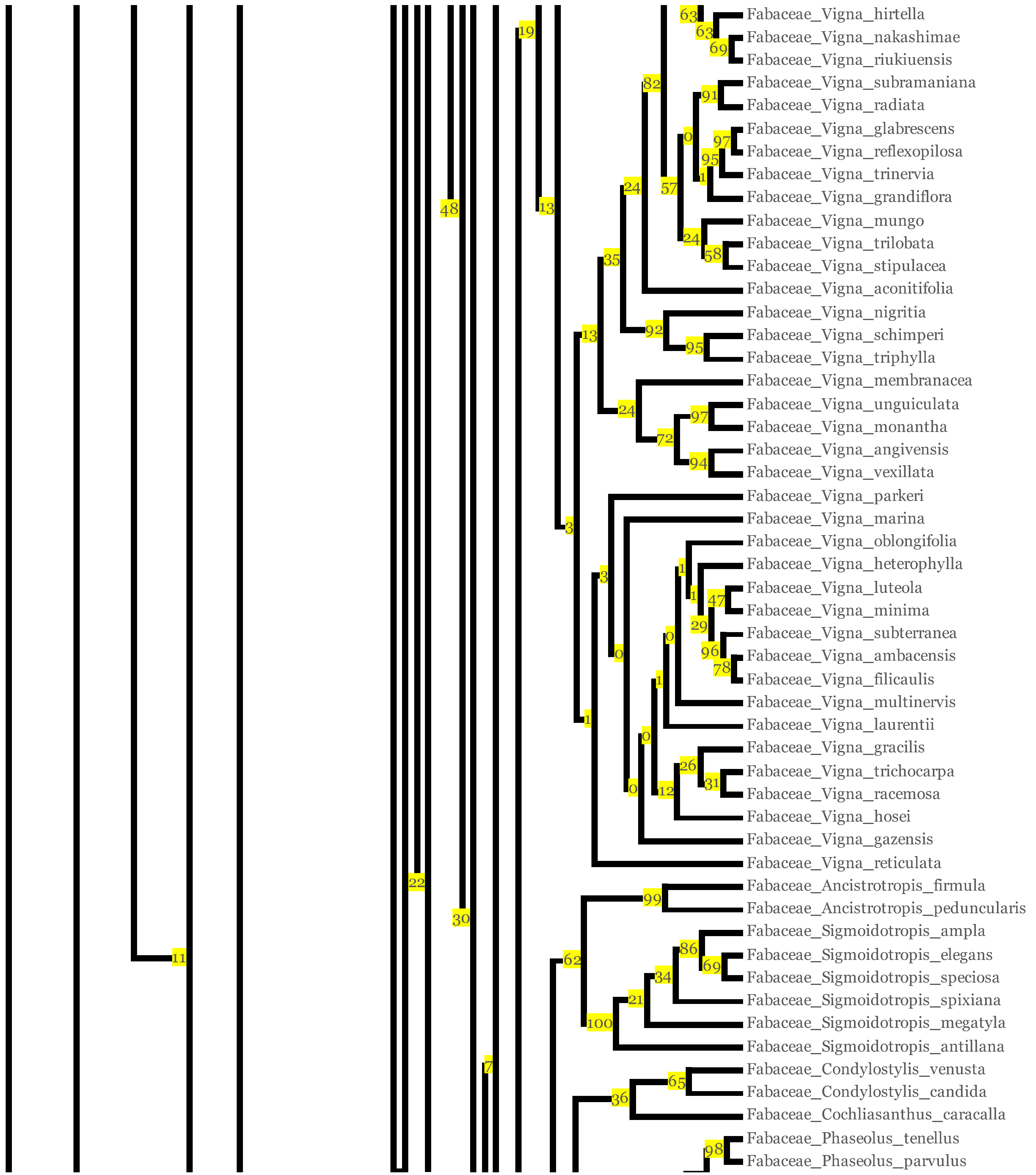

Fabaceae_Vigna_hirtella
Fabaceae_Vigna_nakashimae
Fabaceae_Vigna_riukiuensis
Fabaceae_Vigna_subramaniana
Fabaceae_Vigna_radiata
Fabaceae_Vigna_glabrescens
Fabaceae_Vigna_reflexopilosa
Fabaceae_Vigna_trinervia
Fabaceae_Vigna_grandiflora
Fabaceae_Vigna_mungo
Fabaceae_Vigna_trilobata
Fabaceae_Vigna_stipulacea
Fabaceae_Vigna_aconitifolia
Fabaceae_Vigna_nigritia
Fabaceae_Vigna_schimperi
Fabaceae_Vigna_triphylla
Fabaceae_Vigna_membranacea
Fabaceae_Vigna_unguiculata
Fabaceae_Vigna_monantha
Fabaceae_Vigna_angivensis
Fabaceae_Vigna_vexillata
Fabaceae_Vigna_parkeri
Fabaceae_Vigna_marina
Fabaceae_Vigna_oblongifolia
Fabaceae_Vigna_heterophylla
Fabaceae_Vigna_luteola
Fabaceae_Vigna_minima
Fabaceae_Vigna_subterranea
Fabaceae_Vigna_ambacensis
Fabaceae_Vigna_filicaulis
Fabaceae_Vigna_multinervis
Fabaceae_Vigna_laurentii
Fabaceae_Vigna_gracilis
Fabaceae_Vigna_trichocarpa
Fabaceae_Vigna_racemosa
Fabaceae_Vigna_hosei
Fabaceae_Vigna_gazensis
Fabaceae_Vigna_reticulata
Fabaceae_Ancistrotropis_firmula
Fabaceae_Ancistrotropis_peduncularis
Fabaceae_Sigmoidotropis_ampla
Fabaceae_Sigmoidotropis_elegans
Fabaceae_Sigmoidotropis_speciosa
Fabaceae_Sigmoidotropis_spixiana
Fabaceae_Sigmoidotropis_megatyla
Fabaceae_Sigmoidotropis_antillana
Fabaceae_Condylostylis_venusta
Fabaceae_Condylostylis_candida
Fabaceae_Cochliasanthus_caracalla
Fabaceae_Phaseolus_tenellus
Fabaceae_Phaseolus_parvulus

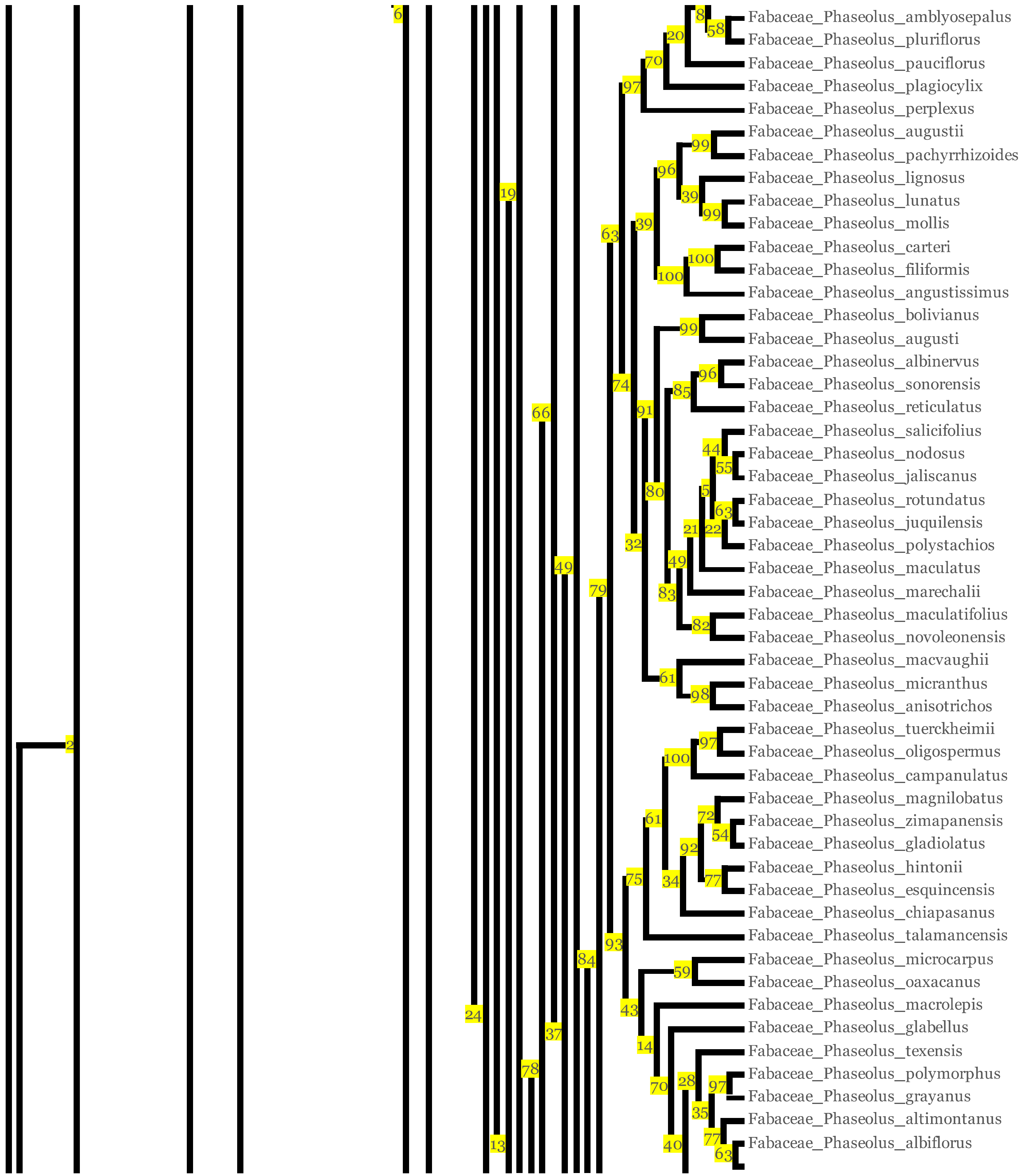

Fabaceae_Phaseolus_amblyosepalus
Fabaceae_Phaseolus_pluriflorus
Fabaceae_Phaseolus_pauciflorus
Fabaceae_Phaseolus_plagiocylix
Fabaceae_Phaseolus_perplexus
Fabaceae_Phaseolus_augustii
Fabaceae_Phaseolus_pachyrrhizoides
Fabaceae_Phaseolus_lignosus
Fabaceae_Phaseolus_lunatus
Fabaceae_Phaseolus_mollis
Fabaceae_Phaseolus_carteri
Fabaceae_Phaseolus_filiformis
Fabaceae_Phaseolus_angustissimus
Fabaceae_Phaseolus_bolivianus
Fabaceae_Phaseolus_augusti
Fabaceae_Phaseolus_albinervus
Fabaceae_Phaseolus_sonorensis
Fabaceae_Phaseolus_reticulatus
Fabaceae_Phaseolus_salicifolius
Fabaceae_Phaseolus_nodosus
Fabaceae_Phaseolus_jaliscanus
Fabaceae_Phaseolus_rotundatus
Fabaceae_Phaseolus_juquilensis
Fabaceae_Phaseolus_polystachios
Fabaceae_Phaseolus_maculatus
Fabaceae_Phaseolus_marechalii
Fabaceae_Phaseolus_maculatifolius
Fabaceae_Phaseolus_novoleonensis
Fabaceae_Phaseolus_macvaughii
Fabaceae_Phaseolus_micranthus
Fabaceae_Phaseolus_anisotrichos
Fabaceae_Phaseolus_tuerckheimii
Fabaceae_Phaseolus_oligospermus
Fabaceae_Phaseolus_campanulatus
Fabaceae_Phaseolus_magnilobatus
Fabaceae_Phaseolus_zimapanensis
Fabaceae_Phaseolus_gladiolatus
Fabaceae_Phaseolus_hintonii
Fabaceae_Phaseolus_esquincensis
Fabaceae_Phaseolus_chiapasanus
Fabaceae_Phaseolus_talamancensis
Fabaceae_Phaseolus_microcarpus
Fabaceae_Phaseolus_oaxacanus
Fabaceae_Phaseolus_macrolepis
Fabaceae_Phaseolus_glabellus
Fabaceae_Phaseolus_texensis
Fabaceae_Phaseolus_polymorphus
Fabaceae_Phaseolus_grayanus
Fabaceae_Phaseolus_altimontanus
Fabaceae_Phaseolus_albiflorus

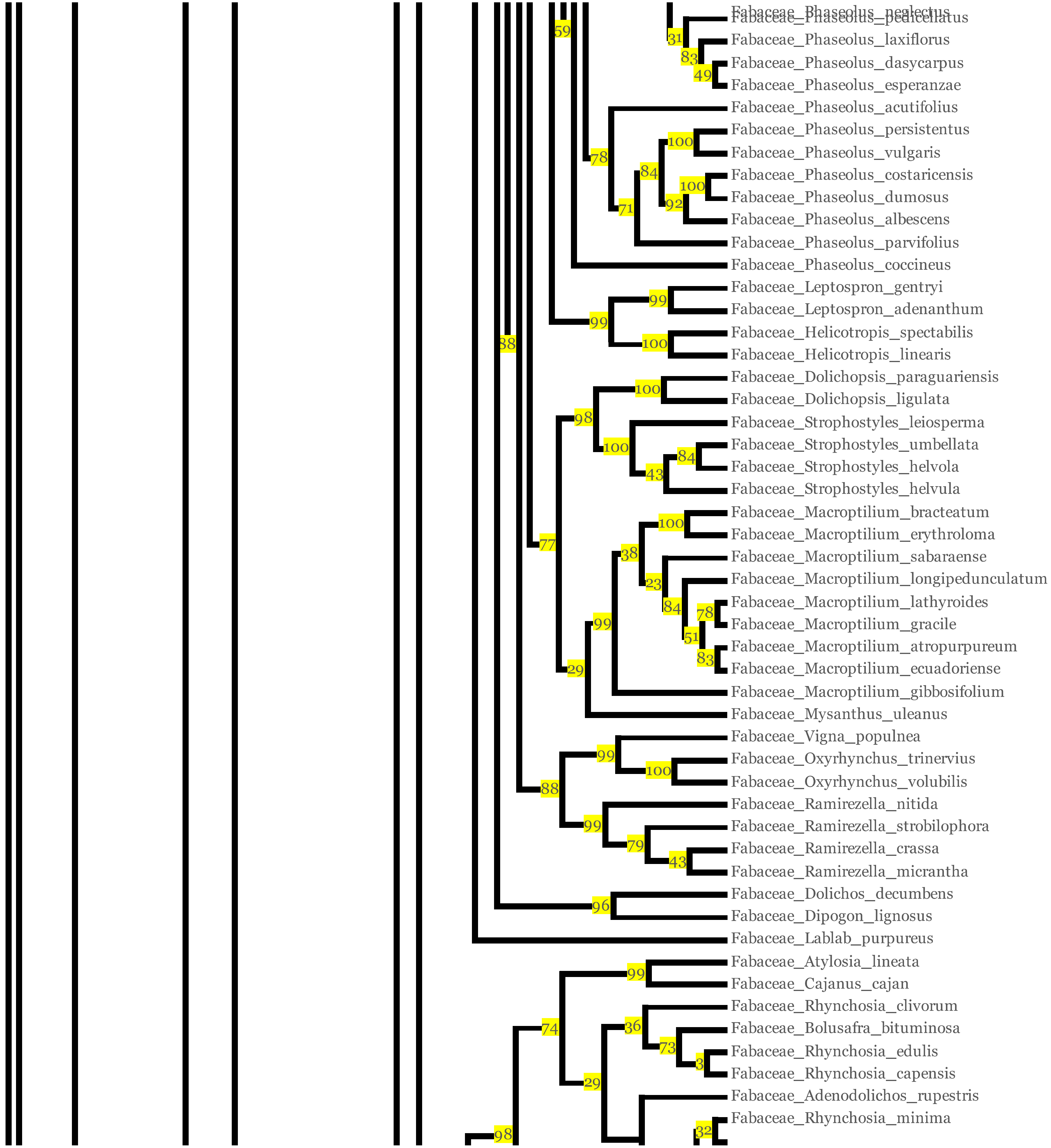

Fabaceae_Phaseolus_neglectus
Fabaceae_Phaseolus_pedicellatus
Fabaceae_Phaseolus_laxiflorus
Fabaceae_Phaseolus_dasycarpus
Fabaceae_Phaseolus_esperanzae
Fabaceae_Phaseolus_acutifolius
Fabaceae_Phaseolus_persistentus
Fabaceae_Phaseolus_vulgaris
Fabaceae_Phaseolus_costaricensis
Fabaceae_Phaseolus_dumosus
Fabaceae_Phaseolus_albescens
Fabaceae_Phaseolus_parvifolius
Fabaceae_Phaseolus_coccineus
Fabaceae_Leptospron_gentryi
Fabaceae_Leptospron_adenanthum
Fabaceae_Helicotropis_spectabilis
Fabaceae_Helicotropis_linearis
Fabaceae_Dolichopsis_paraguariensis
Fabaceae_Dolichopsis_ligulata
Fabaceae_Strophostyles_leiosperma
Fabaceae_Strophostyles_umbellata
Fabaceae_Strophostyles_helvola
Fabaceae_Strophostyles_helvula
Fabaceae_Macroptilium_bracteatum
Fabaceae_Macroptilium_erythroloma
Fabaceae_Macroptilium_sabaraense
Fabaceae_Macroptilium_longipedunculatum
Fabaceae_Macroptilium_lathyroides
Fabaceae_Macroptilium_gracile
Fabaceae_Macroptilium_atropurpureum
Fabaceae_Macroptilium_ecuadoriense
Fabaceae_Macroptilium_gibbosifolium
Fabaceae_Mysanthus_uleanus
Fabaceae_Vigna_populnea
Fabaceae_Oxyrhynchus_trinervius
Fabaceae_Oxyrhynchus_volubilis
Fabaceae_Ramirezella_nitida
Fabaceae_Ramirezella_strobilophora
Fabaceae_Ramirezella_crassa
Fabaceae_Ramirezella_micrantha
Fabaceae_Dolichos_decumbens
Fabaceae_Dipogon_lignosus
Fabaceae_Lablab_purpureus
Fabaceae_Atylosia_lineata
Fabaceae_Cajanus_cajan
Fabaceae_Rhynchosia_clivorum
Fabaceae_Bolusafra_bituminosa
Fabaceae_Rhynchosia_edulis
Fabaceae_Rhynchosia_capensis
Fabaceae_Adenodolichos_rupestris
Fabaceae_Rhynchosia_minima

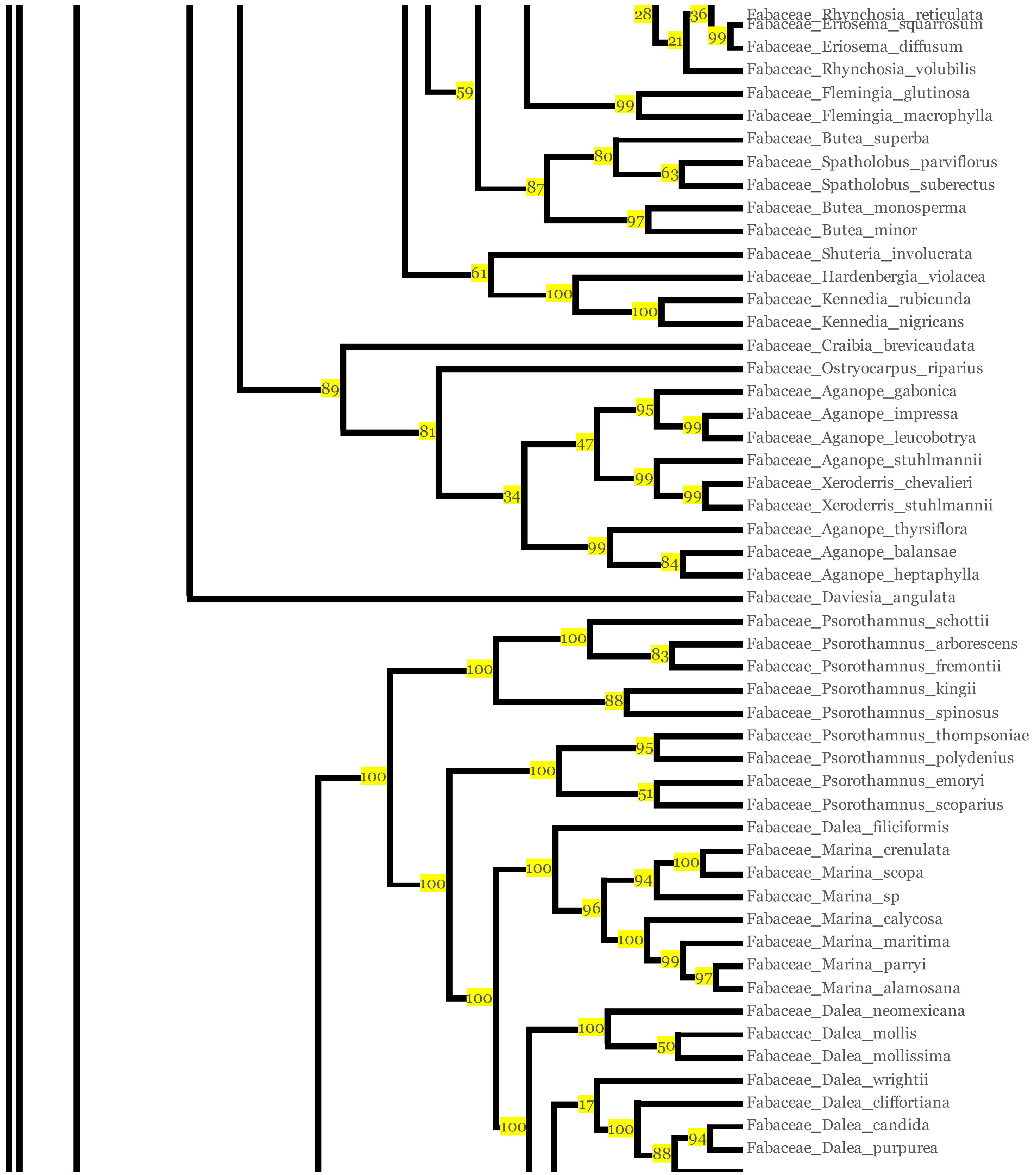
Fabaceae_Rhynchosia_reticulata
Fabaceae_Eriosema_squarrosum
Fabaceae_Eriosema_diffusum
Fabaceae_Rhynchosia_volubilis
Fabaceae_Flemingia_glutinosa
Fabaceae_Flemingia_macrophylla
Fabaceae_Butea_superba
Fabaceae_Spatholobus_parviflorus
Fabaceae_Spatholobus_suberectus
Fabaceae_Butea_monosperma
Fabaceae_Butea_minor
Fabaceae_Shuteria_involucrata
Fabaceae_Hardenbergia_violacea
Fabaceae_Kennedia_rubicunda
Fabaceae_Kennedia_nigricans
Fabaceae_Craibia_brevicaudata
Fabaceae_Ostryocarpus_riparius
Fabaceae_Aganope_gabonica
Fabaceae_Aganope_impressa
Fabaceae_Aganope_leucobotrya
Fabaceae_Aganope_stuhlmannii
Fabaceae_Xeroderris_chevalieri
Fabaceae_Xeroderris_stuhlmannii
Fabaceae_Aganope_thyrsiflora
Fabaceae_Aganope_balansae
Fabaceae_Aganope_heptaphylla
Fabaceae_Daviesia_angulata
Fabaceae_Psorothamnus_schottii
Fabaceae_Psorothamnus_arborescens
Fabaceae_Psorothamnus_fremontii
Fabaceae_Psorothamnus_kingii
Fabaceae_Psorothamnus_spinosus
Fabaceae_Psorothamnus_thompsoniae
Fabaceae_Psorothamnus_polydenius
Fabaceae_Psorothamnus_emoryi
Fabaceae_Psorothamnus_scoparius
Fabaceae_Dalea_filiciformis
Fabaceae_Marina_crenulata
Fabaceae_Marina_scopa
Fabaceae_Marina_sp
Fabaceae_Marina_calycosa
Fabaceae_Marina_maritima
Fabaceae_Marina_parryi
Fabaceae_Marina_alamosana
Fabaceae_Dalea_neomexicana
Fabaceae_Dalea_mollis
Fabaceae_Dalea_mollissima
Fabaceae_Dalea_wrightii
Fabaceae_Dalea_cliffortiana
Fabaceae_Dalea_candida
Fabaceae_Dalea_purpurea

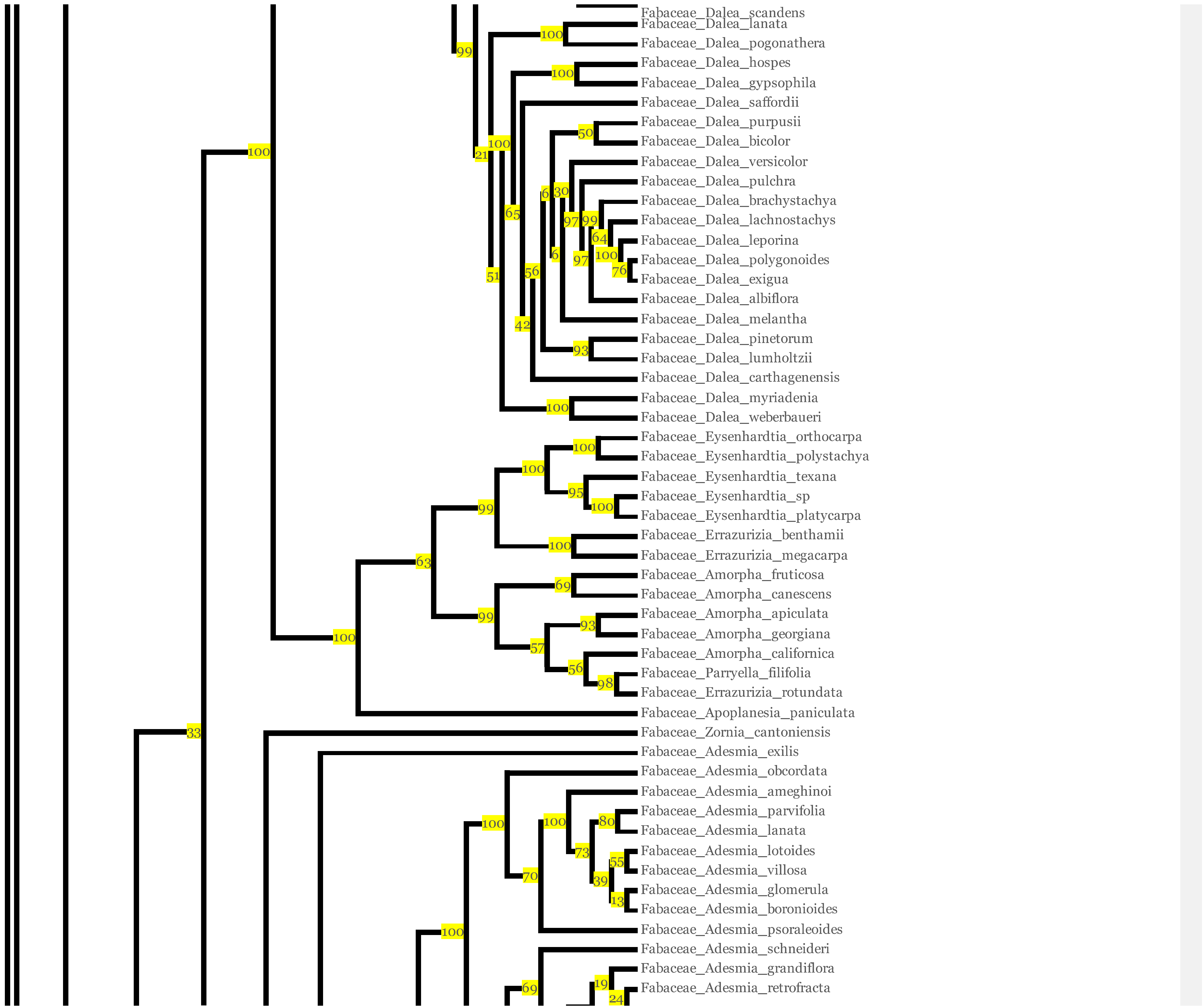
Fabaceae_Dalea_scandens
Fabaceae_Dalea_lanata
Fabaceae_Dalea_pogonathera
Fabaceae_Dalea_hospes
Fabaceae_Dalea_gypsophila
Fabaceae_Dalea_saffordii
Fabaceae_Dalea_purpusii
Fabaceae_Dalea_bicolor
Fabaceae_Dalea_versicolor
Fabaceae_Dalea_pulchra
Fabaceae_Dalea_brachystachya
Fabaceae_Dalea_lachnostachys
Fabaceae_Dalea_leporina
Fabaceae_Dalea_polygonoides
Fabaceae_Dalea_exigua
Fabaceae_Dalea_albiflora
Fabaceae_Dalea_melantha
Fabaceae_Dalea_pinetorum
Fabaceae_Dalea_lumholtzii
Fabaceae_Dalea_carthagenensis
Fabaceae_Dalea_myriadenia
Fabaceae_Dalea_weberbaueri
Fabaceae_Eysenhardtia_orthocarpa
Fabaceae_Eysenhardtia_polystachya
Fabaceae_Eysenhardtia_texana
Fabaceae_Eysenhardtia_sp
Fabaceae_Eysenhardtia_platycarpa
Fabaceae_Errazurizia_benthamii
Fabaceae_Errazurizia_megacarpa
Fabaceae_Amorpha_fruticosa
Fabaceae_Amorpha_canescens
Fabaceae_Amorpha_apiculata
Fabaceae_Amorpha_georgiana
Fabaceae_Amorpha_californica
Fabaceae_Parryella_filifolia
Fabaceae_Errazurizia_rotundata
Fabaceae_Apoplanesia_paniculata
Fabaceae_Zornia_cantoniensis
Fabaceae_Adesmia_exilis
Fabaceae_Adesmia_obcordata
Fabaceae_Adesmia_ameghinoi
Fabaceae_Adesmia_parvifolia
Fabaceae_Adesmia_lanata
Fabaceae_Adesmia_lotoides
Fabaceae_Adesmia_villosa
Fabaceae_Adesmia_glomerula
Fabaceae_Adesmia_boronioides
Fabaceae_Adesmia_psoraleoides
Fabaceae_Adesmia_schneideri
Fabaceae_Adesmia_grandiflora
Fabaceae_Adesmia_retrofracta

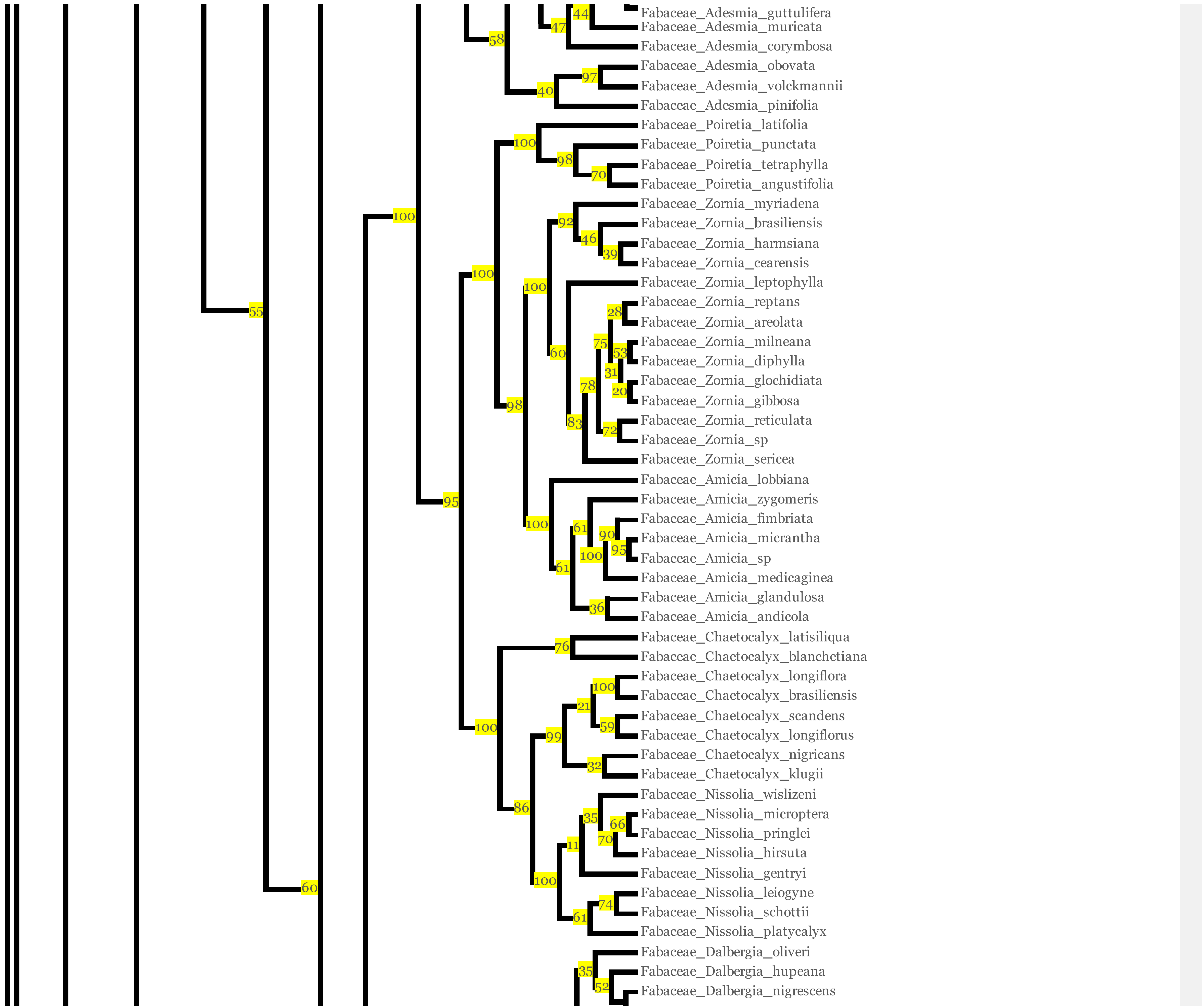
Fabaceae_Adesmia_guttulifera
Fabaceae_Adesmia_muricata
Fabaceae_Adesmia_corymbosa
Fabaceae_Adesmia_obovata
Fabaceae_Adesmia_volckmannii
Fabaceae_Adesmia_pinifolia
Fabaceae_Poiretia_latifolia
Fabaceae_Poiretia_punctata
Fabaceae_Poiretia_tetraphylla
Fabaceae_Poiretia_angustifolia
Fabaceae_Zornia_myriadena
Fabaceae_Zornia_brasiliensis
Fabaceae_Zornia_harmsiana
Fabaceae_Zornia_cearensis
Fabaceae_Zornia_leptophylla
Fabaceae_Zornia_reptans
Fabaceae_Zornia_areolata
Fabaceae_Zornia_milneana
Fabaceae_Zornia_diphylla
Fabaceae_Zornia_glochidiata
Fabaceae_Zornia_gibbosa
Fabaceae_Zornia_reticulata
Fabaceae_Zornia_sp
Fabaceae_Zornia_sericea
Fabaceae_Amicia_lobbiana
Fabaceae_Amicia_zygomeris
Fabaceae_Amicia_fimbriata
Fabaceae_Amicia_micrantha
Fabaceae_Amicia_sp
Fabaceae_Amicia_medicaginea
Fabaceae_Amicia_glandulosa
Fabaceae_Amicia_andicola
Fabaceae_Chaetocalyx_latisiliqua
Fabaceae_Chaetocalyx_blanchetiana
Fabaceae_Chaetocalyx_longiflora
Fabaceae_Chaetocalyx_brasiliensis
Fabaceae_Chaetocalyx_scandens
Fabaceae_Chaetocalyx_longiflorus
Fabaceae_Chaetocalyx_nigricans
Fabaceae_Chaetocalyx_klugii
Fabaceae_Nissolia_wislizeni
Fabaceae_Nissolia_microptera
Fabaceae_Nissolia_pringlei
Fabaceae_Nissolia_hirsuta
Fabaceae_Nissolia_gentryi
Fabaceae_Nissolia_leiogyne
Fabaceae_Nissolia_schottii
Fabaceae_Nissolia_platycalyx
Fabaceae_Dalbergia_oliveri
Fabaceae_Dalbergia_hupeana
Fabaceae_Dalbergia_nigrescens

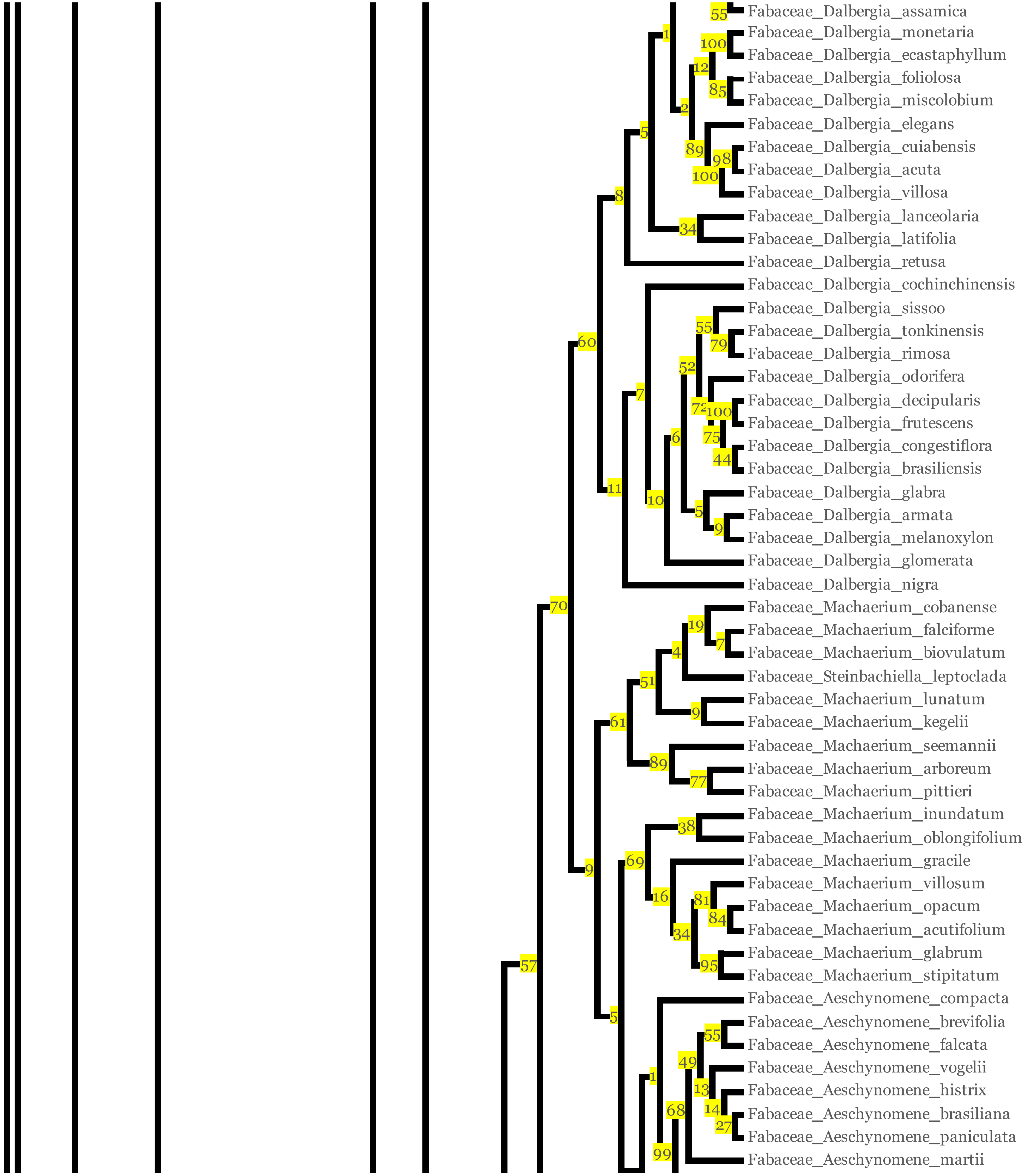
Fabaceae_Dalbergia_assamica
Fabaceae_Dalbergia_monetaria
Fabaceae_Dalbergia_ecastaphyllum
Fabaceae_Dalbergia_foliolosa
Fabaceae_Dalbergia_miscolobium
Fabaceae_Dalbergia_elegans
Fabaceae_Dalbergia_cuiabensis
Fabaceae_Dalbergia_acuta
Fabaceae_Dalbergia_villosa
Fabaceae_Dalbergia_lanceolaria
Fabaceae_Dalbergia_latifolia
Fabaceae_Dalbergia_retusa
Fabaceae_Dalbergia_cochinchinensis
Fabaceae_Dalbergia_sissoo
Fabaceae_Dalbergia_tonkinensis
Fabaceae_Dalbergia_rimosa
Fabaceae_Dalbergia_odorifera
Fabaceae_Dalbergia_decipularis
Fabaceae_Dalbergia_frutescens
Fabaceae_Dalbergia_congestiflora
Fabaceae_Dalbergia_brasiliensis
Fabaceae_Dalbergia_glabra
Fabaceae_Dalbergia_armata
Fabaceae_Dalbergia_melanoxylon
Fabaceae_Dalbergia_glomerata
Fabaceae_Dalbergia_nigra
Fabaceae_Machaerium_cobanense
Fabaceae_Machaerium_falciforme
Fabaceae_Machaerium_biovulatum
Fabaceae_Steinbachiella_leptoclada
Fabaceae_Machaerium_lunatum
Fabaceae_Machaerium_kegelii
Fabaceae_Machaerium_seemannii
Fabaceae_Machaerium_arboreum
Fabaceae_Machaerium_pittieri
Fabaceae_Machaerium_inundatum
Fabaceae_Machaerium_oblongifolium
Fabaceae_Machaerium_gracile
Fabaceae_Machaerium_villosum
Fabaceae_Machaerium_opacum
Fabaceae_Machaerium_acutifolium
Fabaceae_Machaerium_glabrum
Fabaceae_Machaerium_stipitatum
Fabaceae_Aeschynomene_compacta
Fabaceae_Aeschynomene_brevifolia
Fabaceae_Aeschynomene_falcata
Fabaceae_Aeschynomene_vogelii
Fabaceae_Aeschynomene_histrix
Fabaceae_Aeschynomene_brasiliana
Fabaceae_Aeschynomene_paniculata
Fabaceae_Aeschynomene_martii

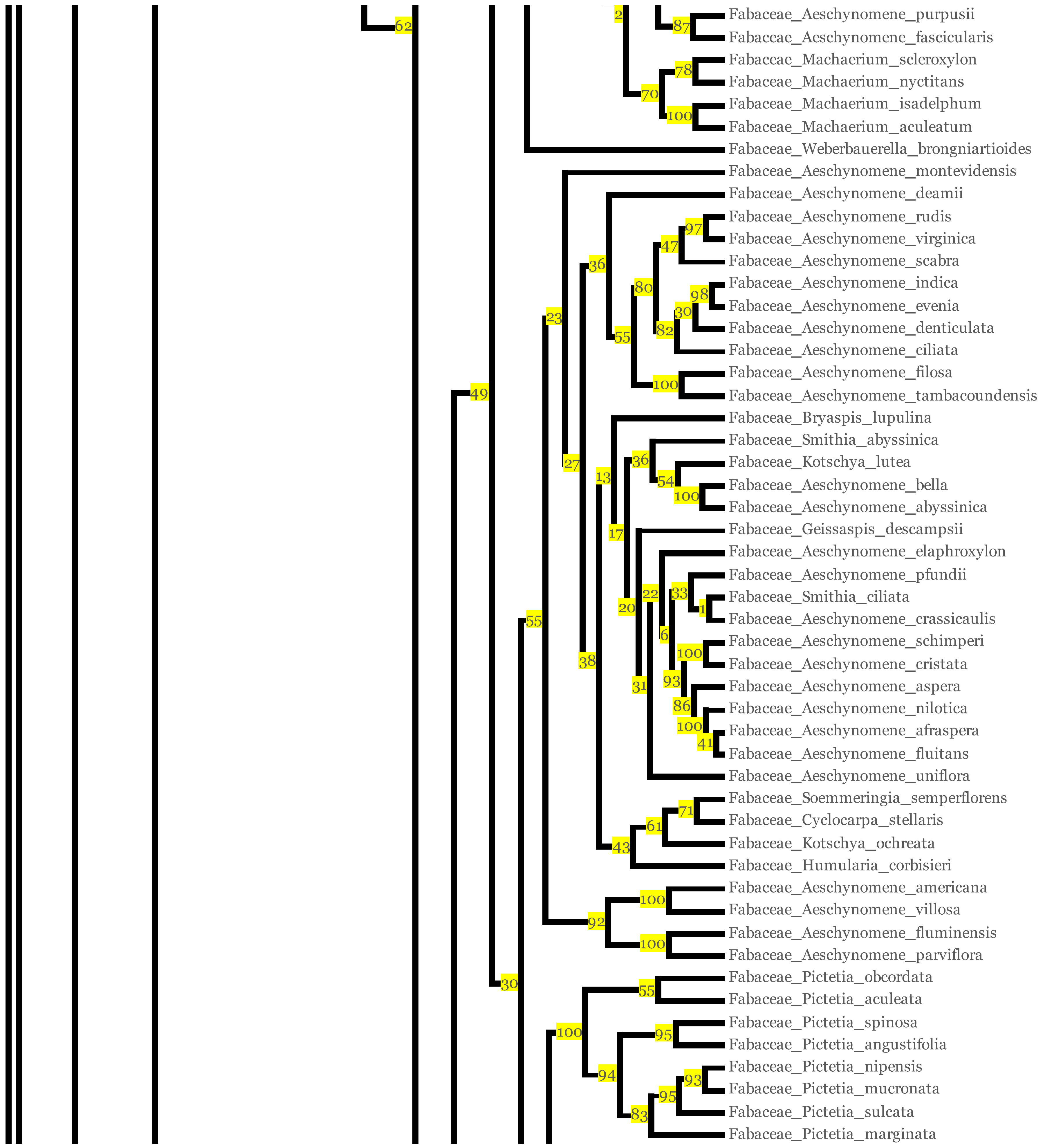
Fabaceae_Aeschynomene_purpusii
Fabaceae_Aeschynomene_fascicularis
Fabaceae_Machaerium_scleroxylon
Fabaceae_Machaerium_nyctitans
Fabaceae_Machaerium_isadelphum
Fabaceae_Machaerium_aculeatum
Fabaceae_Weberbauerella_brongniartioides
Fabaceae_Aeschynomene_montevidensis
Fabaceae_Aeschynomene_deamii
Fabaceae_Aeschynomene_rudis
Fabaceae_Aeschynomene_virginica
Fabaceae_Aeschynomene_scabra
Fabaceae_Aeschynomene_indica
Fabaceae_Aeschynomene_evenia
Fabaceae_Aeschynomene_denticulata
Fabaceae_Aeschynomene_ciliata
Fabaceae_Aeschynomene_filosa
Fabaceae_Aeschynomene_tambacoundensis
Fabaceae_Bryaspis_lupulina
Fabaceae_Smithia_abyssinica
Fabaceae_Kotschya_lutea
Fabaceae_Aeschynomene_bella
Fabaceae_Aeschynomene_abyssinica
Fabaceae_Geissaspis_descampsii
Fabaceae_Aeschynomene_elaphroxylon
Fabaceae_Aeschynomene_pfundii
Fabaceae_Smithia_ciliata
Fabaceae_Aeschynomene_crassicaulis
Fabaceae_Aeschynomene_schimperi
Fabaceae_Aeschynomene_cristata
Fabaceae_Aeschynomene_aspera
Fabaceae_Aeschynomene_nilotica
Fabaceae_Aeschynomene_afraspera
Fabaceae_Aeschynomene_fluitans
Fabaceae_Aeschynomene_uniflora
Fabaceae_Soemmeringia_semperflorens
Fabaceae_Cyclocarpa_stellaris
Fabaceae_Kotschya_ochreata
Fabaceae_Humularia_corbisieri
Fabaceae_Aeschynomene_americana
Fabaceae_Aeschynomene_villosa
Fabaceae_Aeschynomene_fluminensis
Fabaceae_Aeschynomene_parviflora
Fabaceae_Pictetia_obcordata
Fabaceae_Pictetia_aculeata
Fabaceae_Pictetia_spinosa
Fabaceae_Pictetia_angustifolia
Fabaceae_Pictetia_nipensis
Fabaceae_Pictetia_mucronata
Fabaceae_Pictetia_sulcata
Fabaceae_Pictetia_marginata

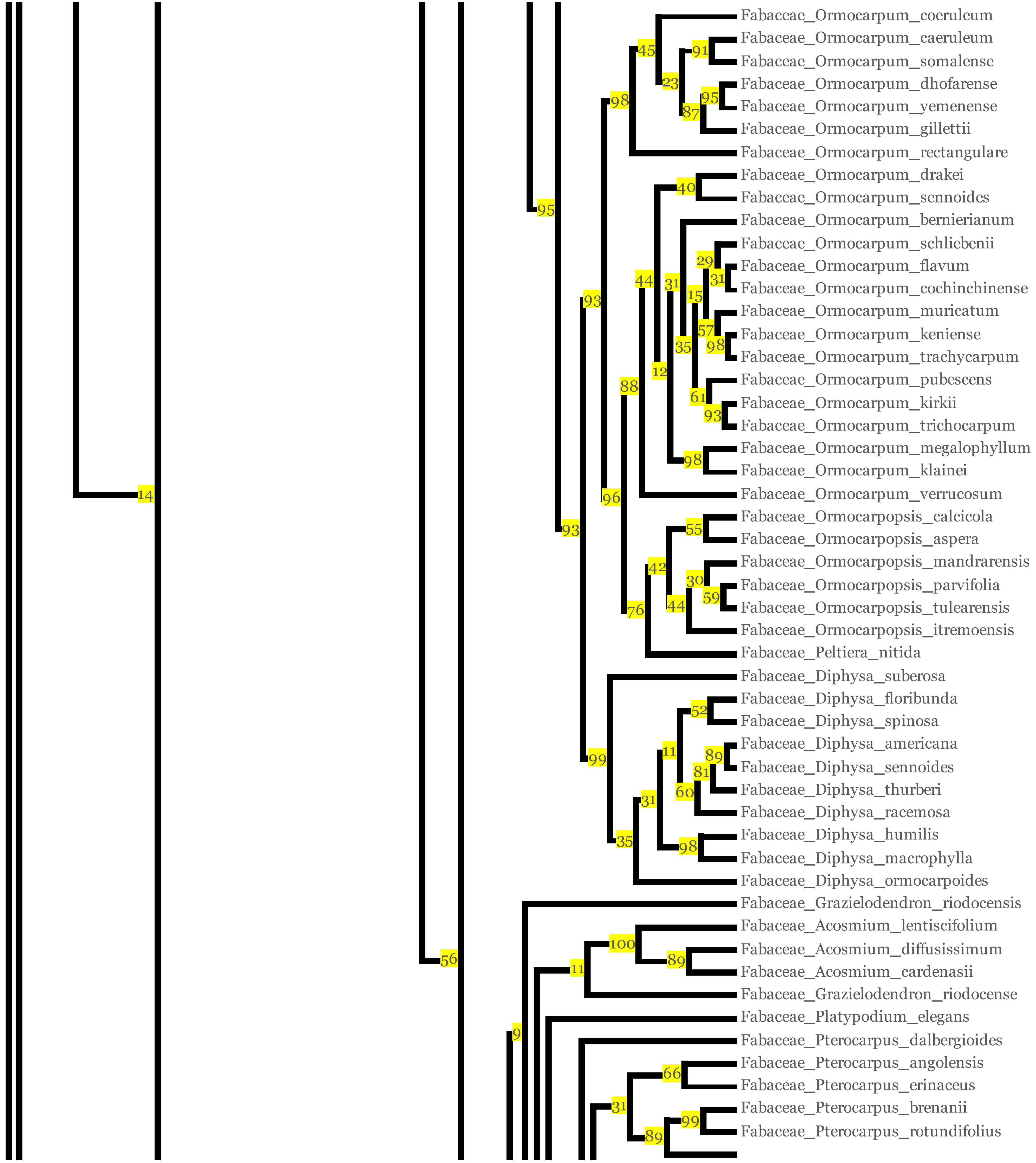
Fabaceae_Ormocarpum_coeruleum
Fabaceae_Ormocarpum_caeruleum
Fabaceae_Ormocarpum_somalense
Fabaceae_Ormocarpum_dhofarense
Fabaceae_Ormocarpum_yemenense
Fabaceae_Ormocarpum_gillettii
Fabaceae_Ormocarpum_rectangulare
Fabaceae_Ormocarpum_drakei
Fabaceae_Ormocarpum_sennoides
Fabaceae_Ormocarpum_bernierianum
Fabaceae_Ormocarpum_schliebenii
Fabaceae_Ormocarpum_flavum
Fabaceae_Ormocarpum_cochinchinense
Fabaceae_Ormocarpum_muricatum
Fabaceae_Ormocarpum_keniense
Fabaceae_Ormocarpum_trachycarpum
Fabaceae_Ormocarpum_pubescens
Fabaceae_Ormocarpum_kirkii
Fabaceae_Ormocarpum_trichocarpum
Fabaceae_Ormocarpum_megalophyllum
Fabaceae_Ormocarpum_klainei
Fabaceae_Ormocarpum_verrucosum
Fabaceae_Ormocarpopsis_calcicola
Fabaceae_Ormocarpopsis_aspera
Fabaceae_Ormocarpopsis_mandrarensis
Fabaceae_Ormocarpopsis_parvifolia
Fabaceae_Ormocarpopsis_tulearensis
Fabaceae_Ormocarpopsis_itremoensis
Fabaceae_Peltiera_nitida
Fabaceae_Diphysa_suberosa
Fabaceae_Diphysa_floribunda
Fabaceae_Diphysa_spinosa
Fabaceae_Diphysa_americana
Fabaceae_Diphysa_sennoides
Fabaceae_Diphysa_thurberi
Fabaceae_Diphysa_racemosa
Fabaceae_Diphysa_humilis
Fabaceae_Diphysa_macrophylla
Fabaceae_Diphysa_ormocarpoides
Fabaceae_Grazielodendron_riodocensis
Fabaceae_Acosmium_lentiscifolium
Fabaceae_Acosmium_diffusissimum
Fabaceae_Acosmium_cardenasii
Fabaceae_Grazielodendron_riodocense
Fabaceae_Platypodium_elegans
Fabaceae_Pterocarpus_dalbergioides
Fabaceae_Pterocarpus_angolensis
Fabaceae_Pterocarpus_erinaceus
Fabaceae_Pterocarpus_brenanii
Fabaceae_Pterocarpus_rotundifolius

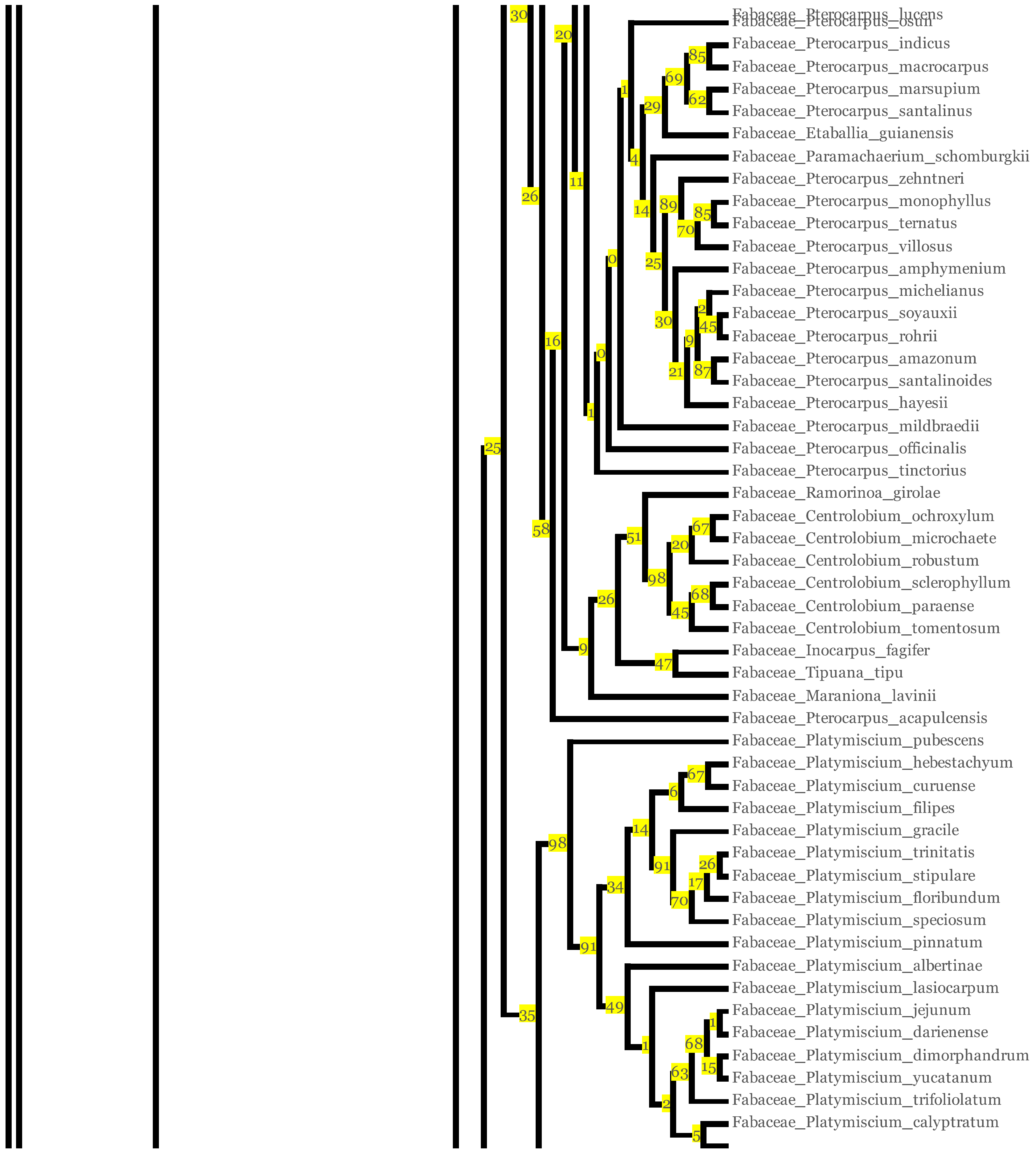
Fabaceae_Pterocarpus_lucens
Fabaceae_Pterocarpus_osun
Fabaceae_Pterocarpus_indicus
Fabaceae_Pterocarpus_macrocarpus
Fabaceae_Pterocarpus_marsupium
Fabaceae_Pterocarpus_santalinus
Fabaceae_Etaballia_guianensis
Fabaceae_Paramachaerium_schomburgkii
Fabaceae_Pterocarpus_zehntneri
Fabaceae_Pterocarpus_monophyllus
Fabaceae_Pterocarpus_ternatus
Fabaceae_Pterocarpus_villosus
Fabaceae_Pterocarpus_amphymenium
Fabaceae_Pterocarpus_michelianus
Fabaceae_Pterocarpus_soyauxii
Fabaceae_Pterocarpus_rohrii
Fabaceae_Pterocarpus_amazonum
Fabaceae_Pterocarpus_santalinoides
Fabaceae_Pterocarpus_hayesii
Fabaceae_Pterocarpus_mildbraedii
Fabaceae_Pterocarpus_officinalis
Fabaceae_Pterocarpus_tinctorius
Fabaceae_Ramorinoa_girolae
Fabaceae_Centrolobium_ochroxylum
Fabaceae_Centrolobium_microchaete
Fabaceae_Centrolobium_robustum
Fabaceae_Centrolobium_sclerophyllum
Fabaceae_Centrolobium_paraense
Fabaceae_Centrolobium_tomentosum
Fabaceae_Inocarpus_fagifer
Fabaceae_Tipuana_tipu
Fabaceae_Maraniona_lavinii
Fabaceae_Pterocarpus_acapulcensis
Fabaceae_Platymiscium_pubescens
Fabaceae_Platymiscium_hebestachyum
Fabaceae_Platymiscium_curuense
Fabaceae_Platymiscium_filipes
Fabaceae_Platymiscium_gracile
Fabaceae_Platymiscium_trinitatis
Fabaceae_Platymiscium_stipulare
Fabaceae_Platymiscium_floribundum
Fabaceae_Platymiscium_speciosum
Fabaceae_Platymiscium_pinnatum
Fabaceae_Platymiscium_albertinae
Fabaceae_Platymiscium_lasiocarpum
Fabaceae_Platymiscium_jejunum
Fabaceae_Platymiscium_darienense
Fabaceae_Platymiscium_dimorphandrum
Fabaceae_Platymiscium_yucatanum
Fabaceae_Platymiscium_trifoliolatum
Fabaceae_Platymiscium_calyptratum

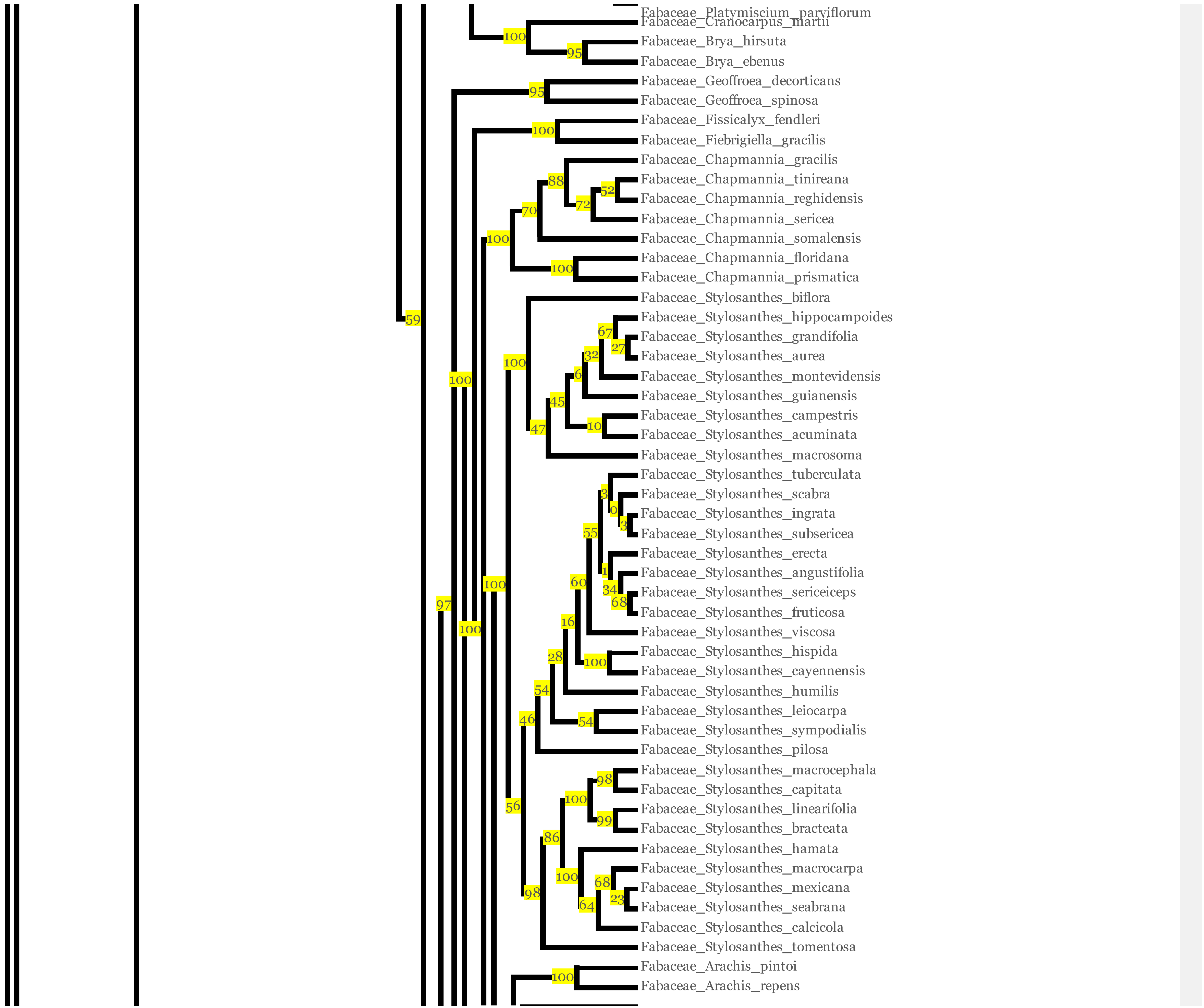
Fabaceae_Platymiscium_parviflorum
Fabaceae_Cranocarpus_martii
Fabaceae_Brya_hirsuta
Fabaceae_Brya_ebenus
Fabaceae_Geoffroea_decorticans
Fabaceae_Geoffroea_spinosa
Fabaceae_Fissicalyx_fendleri
Fabaceae_Fiebrigiella_gracilis
Fabaceae_Chapmannia_gracilis
Fabaceae_Chapmannia_tinireana
Fabaceae_Chapmannia_reghidensis
Fabaceae_Chapmannia_sericea
Fabaceae_Chapmannia_somalensis
Fabaceae_Chapmannia_floridana
Fabaceae_Chapmannia_prismatica
Fabaceae_Stylosanthes_biflora
Fabaceae_Stylosanthes_hippocampoides
Fabaceae_Stylosanthes_grandifolia
Fabaceae_Stylosanthes_aurea
Fabaceae_Stylosanthes_montevidensis
Fabaceae_Stylosanthes_guianensis
Fabaceae_Stylosanthes_campestris
Fabaceae_Stylosanthes_acuminata
Fabaceae_Stylosanthes_macrosoma
Fabaceae_Stylosanthes_tuberculata
Fabaceae_Stylosanthes_scabra
Fabaceae_Stylosanthes_ingrata
Fabaceae_Stylosanthes_subsericea
Fabaceae_Stylosanthes_erecta
Fabaceae_Stylosanthes_angustifolia
Fabaceae_Stylosanthes_sericeiceps
Fabaceae_Stylosanthes_fruticosa
Fabaceae_Stylosanthes_viscosa
Fabaceae_Stylosanthes_hispida
Fabaceae_Stylosanthes_cayennensis
Fabaceae_Stylosanthes_humilis
Fabaceae_Stylosanthes_leiocarpa
Fabaceae_Stylosanthes_sympodialis
Fabaceae_Stylosanthes_pilosa
Fabaceae_Stylosanthes_macrocephala
Fabaceae_Stylosanthes_capitata
Fabaceae_Stylosanthes_linearifolia
Fabaceae_Stylosanthes_bracteata
Fabaceae_Stylosanthes_hamata
Fabaceae_Stylosanthes_macrocarpa
Fabaceae_Stylosanthes_mexicana
Fabaceae_Stylosanthes_seabrana
Fabaceae_Stylosanthes_calcicola
Fabaceae_Stylosanthes_tomentosa
Fabaceae_Arachis_pintoi
Fabaceae_Arachis_repens

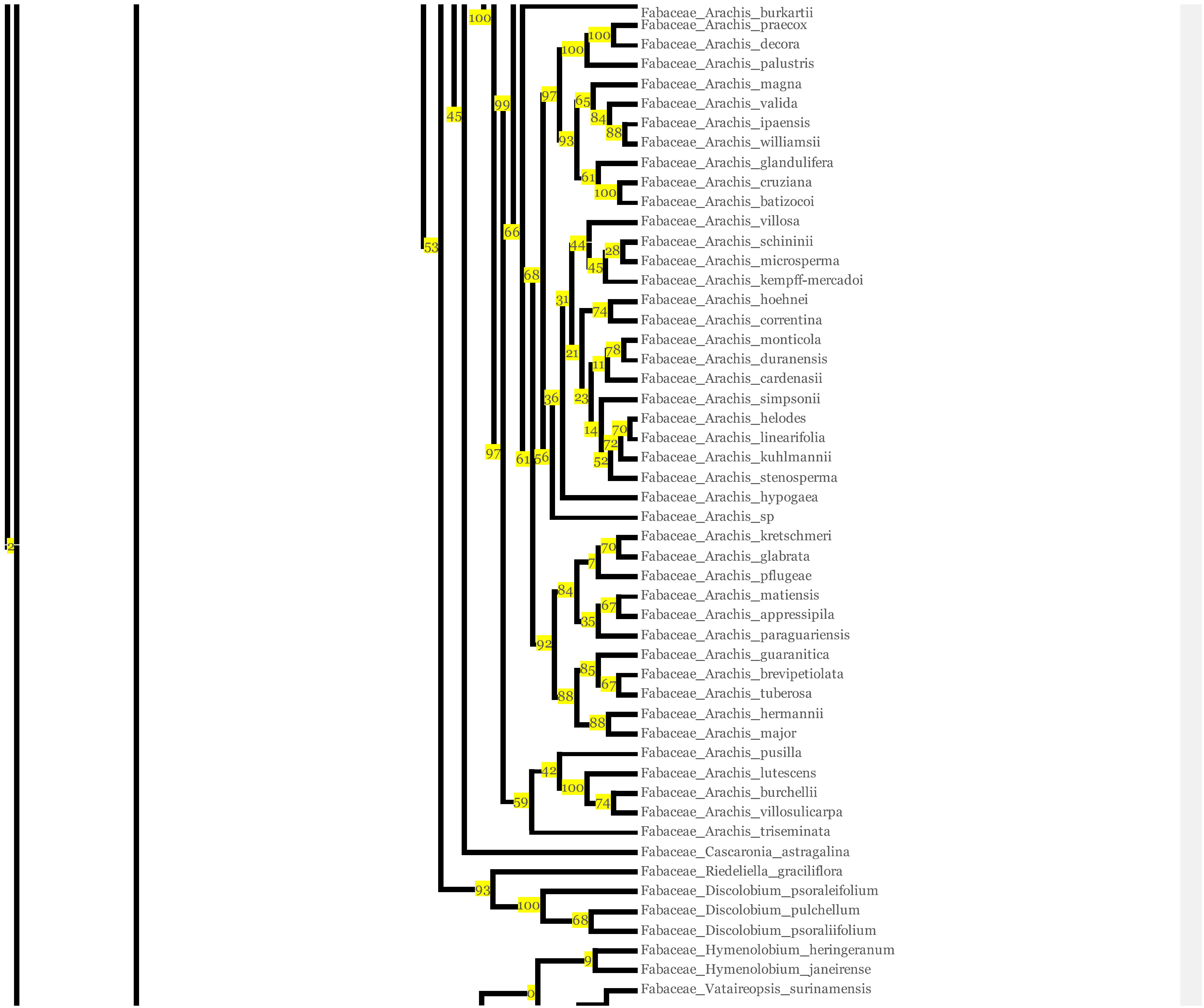
Fabaceae_Arachis_burkartii
Fabaceae_Arachis_praecox
Fabaceae_Arachis_decora
Fabaceae_Arachis_palustris
Fabaceae_Arachis_magna
Fabaceae_Arachis_valida
Fabaceae_Arachis_ipaensis
Fabaceae_Arachis_williamsii
Fabaceae_Arachis_glandulifera
Fabaceae_Arachis_cruziana
Fabaceae_Arachis_batizocoi
Fabaceae_Arachis_villosa
Fabaceae_Arachis_schininii
Fabaceae_Arachis_microsperma
Fabaceae_Arachis_kempff-mercadoi
Fabaceae_Arachis_hoehnei
Fabaceae_Arachis_correntina
Fabaceae_Arachis_monticola
Fabaceae_Arachis_duranensis
Fabaceae_Arachis_cardenasii
Fabaceae_Arachis_simpsonii
Fabaceae_Arachis_helodes
Fabaceae_Arachis_linearifolia
Fabaceae_Arachis_kuhlmannii
Fabaceae_Arachis_stenosperma
Fabaceae_Arachis_hypogaea
Fabaceae_Arachis_sp
Fabaceae_Arachis_kretschmeri
Fabaceae_Arachis_glabrata
Fabaceae_Arachis_pflugeae
Fabaceae_Arachis_matiensis
Fabaceae_Arachis_appressipila
Fabaceae_Arachis_paraguariensis
Fabaceae_Arachis_guaranitica
Fabaceae_Arachis_brevipetiolata
Fabaceae_Arachis_tuberosa
Fabaceae_Arachis_hermannii
Fabaceae_Arachis_major
Fabaceae_Arachis_pusilla
Fabaceae_Arachis_lutescens
Fabaceae_Arachis_burchellii
Fabaceae_Arachis_villosulicarpa
Fabaceae_Arachis_triseminata
Fabaceae_Cascaronia_astragalina
Fabaceae_Riedeliella_graciliflora
Fabaceae_Discolobium_psoraleifolium
Fabaceae_Discolobium_pulchellum
Fabaceae_Discolobium_psoraliifolium
Fabaceae_Hymenolobium_heringeranum
Fabaceae_Hymenolobium_janeirense
Fabaceae_Vataireopsis_surinamensis

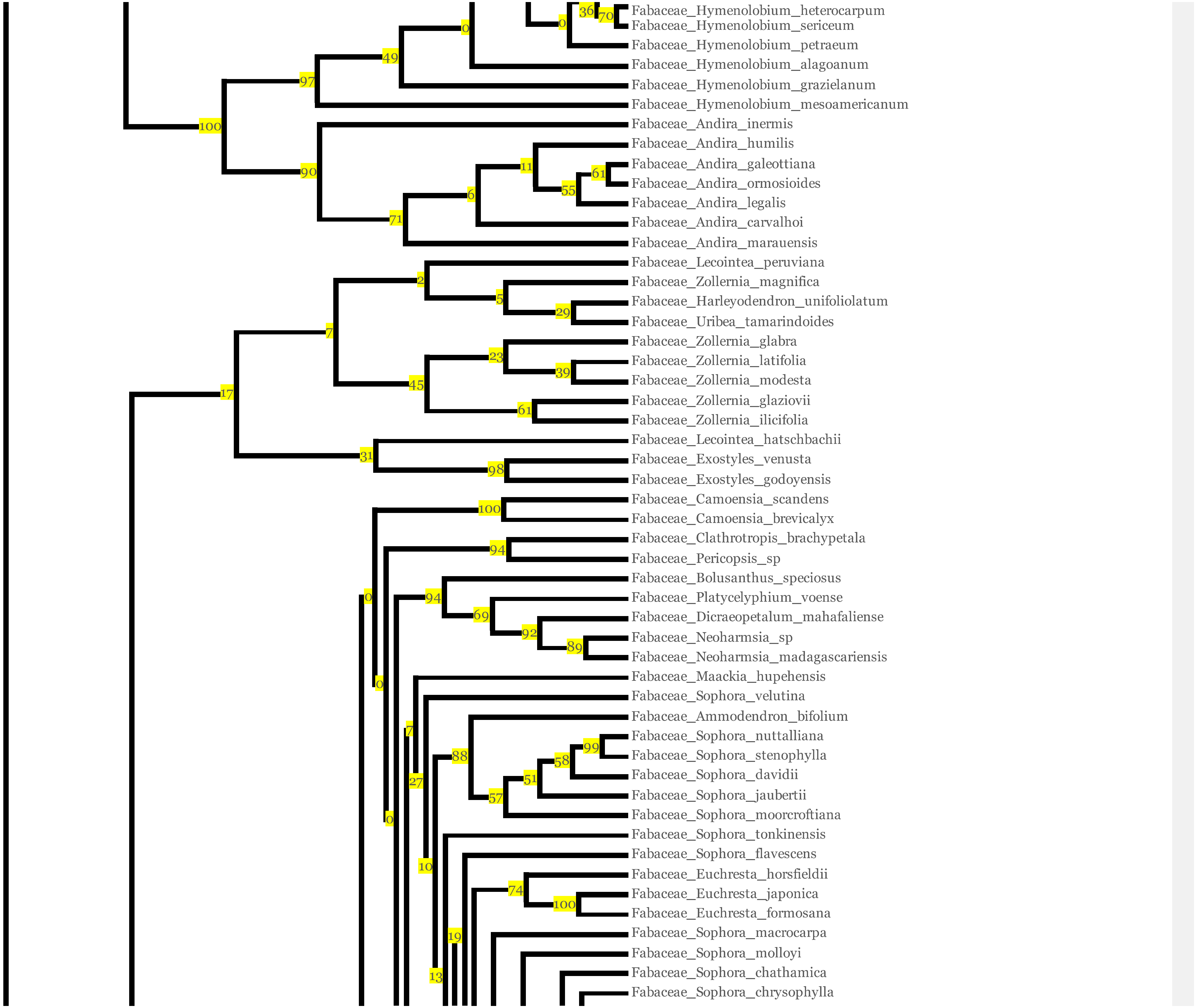
Fabaceae_Hymenolobium_heterocarpum
Fabaceae_Hymenolobium_sericeum
Fabaceae_Hymenolobium_petraeum
Fabaceae_Hymenolobium_alagoanum
Fabaceae_Hymenolobium_grazielanum
Fabaceae_Hymenolobium_mesoamericanum
Fabaceae_Andira_inermis
Fabaceae_Andira_humilis
Fabaceae_Andira_galeottiana
Fabaceae_Andira_ormosioides
Fabaceae_Andira_legalis
Fabaceae_Andira_carvalhoi
Fabaceae_Andira_marauensis
Fabaceae_Lecointea_peruviana
Fabaceae_Zollernia_magnifica
Fabaceae_Harleyodendron_unifoliolatum
Fabaceae_Uribea_tamarindoides
Fabaceae_Zollernia_glabra
Fabaceae_Zollernia_latifolia
Fabaceae_Zollernia_modesta
Fabaceae_Zollernia_glaziovii
Fabaceae_Zollernia_ilicifolia
Fabaceae_Lecointea_hatschbachii
Fabaceae_Exostyles_venusta
Fabaceae_Exostyles_godoyensis
Fabaceae_Camoensia_scandens
Fabaceae_Camoensia_brevicalyx
Fabaceae_Clathrotropis_brachypetala
Fabaceae_Pericopsis_sp
Fabaceae_Bolusanthus_speciosus
Fabaceae_Platycelyphium_voense
Fabaceae_Dicraeopetalum_mahafaliense
Fabaceae_Neoharmsia_sp
Fabaceae_Neoharmsia_madagascariensis
Fabaceae_Maackia_hupehensis
Fabaceae_Sophora_velutina
Fabaceae_Ammodendron_bifolium
Fabaceae_Sophora_nuttalliana
Fabaceae_Sophora_stenophylla
Fabaceae_Sophora_davidii
Fabaceae_Sophora_jaubertii
Fabaceae_Sophora_moorcroftiana
Fabaceae_Sophora_tonkinensis
Fabaceae_Sophora_flavescens
Fabaceae_Euchresta_horsfieldii
Fabaceae_Euchresta_japonica
Fabaceae_Euchresta_formosana
Fabaceae_Sophora_macrocarpa
Fabaceae_Sophora_molloyi
Fabaceae_Sophora_chathamica
Fabaceae_Sophora_chrysophylla

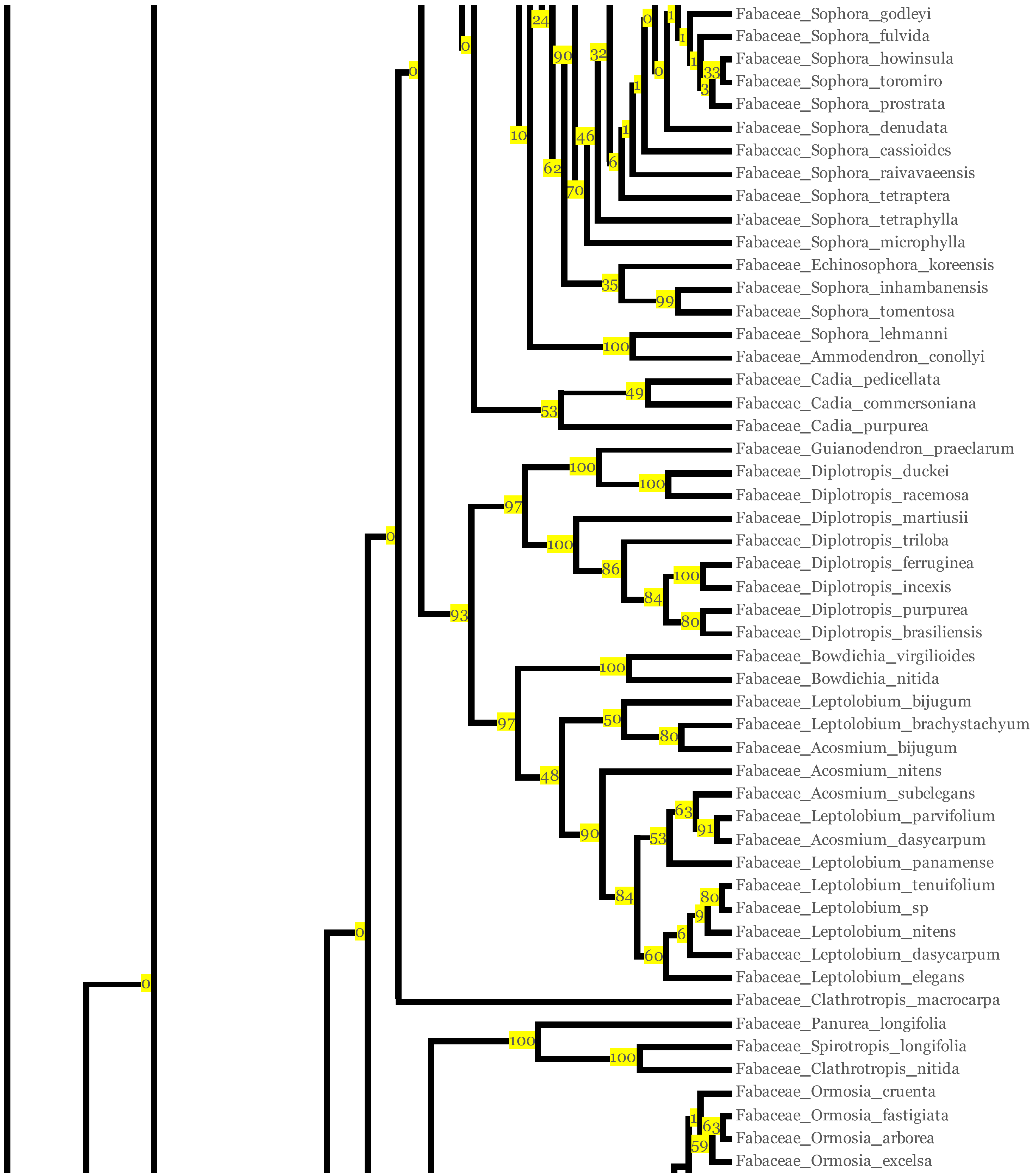
Fabaceae_Sophora_godleyi
Fabaceae_Sophora_fulvida
Fabaceae_Sophora_howinsula
Fabaceae_Sophora_toromiro
Fabaceae_Sophora_prostrata
Fabaceae_Sophora_denudata
Fabaceae_Sophora_cassioides
Fabaceae_Sophora_raivavaeensis
Fabaceae_Sophora_tetraptera
Fabaceae_Sophora_tetraphylla
Fabaceae_Sophora_microphylla
Fabaceae_Echinosophora_koreensis
Fabaceae_Sophora_inhambanensis
Fabaceae_Sophora_tomentosa
Fabaceae_Sophora_lehmanni
Fabaceae_Ammodendron_conollyi
Fabaceae_Cadia_pedicellata
Fabaceae_Cadia_commersoniana
Fabaceae_Cadia_purpurea
Fabaceae_Guianodendron_praeclarum
Fabaceae_Diplotropis_duckei
Fabaceae_Diplotropis_racemosa
Fabaceae_Diplotropis_martiusii
Fabaceae_Diplotropis_triloba
Fabaceae_Diplotropis_ferruginea
Fabaceae_Diplotropis_incexis
Fabaceae_Diplotropis_purpurea
Fabaceae_Diplotropis_brasiliensis
Fabaceae_Bowdichia_virgilioides
Fabaceae_Bowdichia_nitida
Fabaceae_Leptolobium_bijugum
Fabaceae_Leptolobium_brachystachyum
Fabaceae_Acosmium_bijugum
Fabaceae_Acosmium_nitens
Fabaceae_Acosmium_subelegans
Fabaceae_Leptolobium_parvifolium
Fabaceae_Acosmium_dasycarpum
Fabaceae_Leptolobium_panamense
Fabaceae_Leptolobium_tenuifolium
Fabaceae_Leptolobium_sp
Fabaceae_Leptolobium_nitens
Fabaceae_Leptolobium_dasycarpum
Fabaceae_Leptolobium_elegans
Fabaceae_Clathrotropis_macrocarpa
Fabaceae_Panurea_longifolia
Fabaceae_Spirotropis_longifolia
Fabaceae_Clathrotropis_nitida
Fabaceae_Ormosia_cruenta
Fabaceae_Ormosia_fastigiata
Fabaceae_Ormosia_arborea
Fabaceae_Ormosia_excelsa

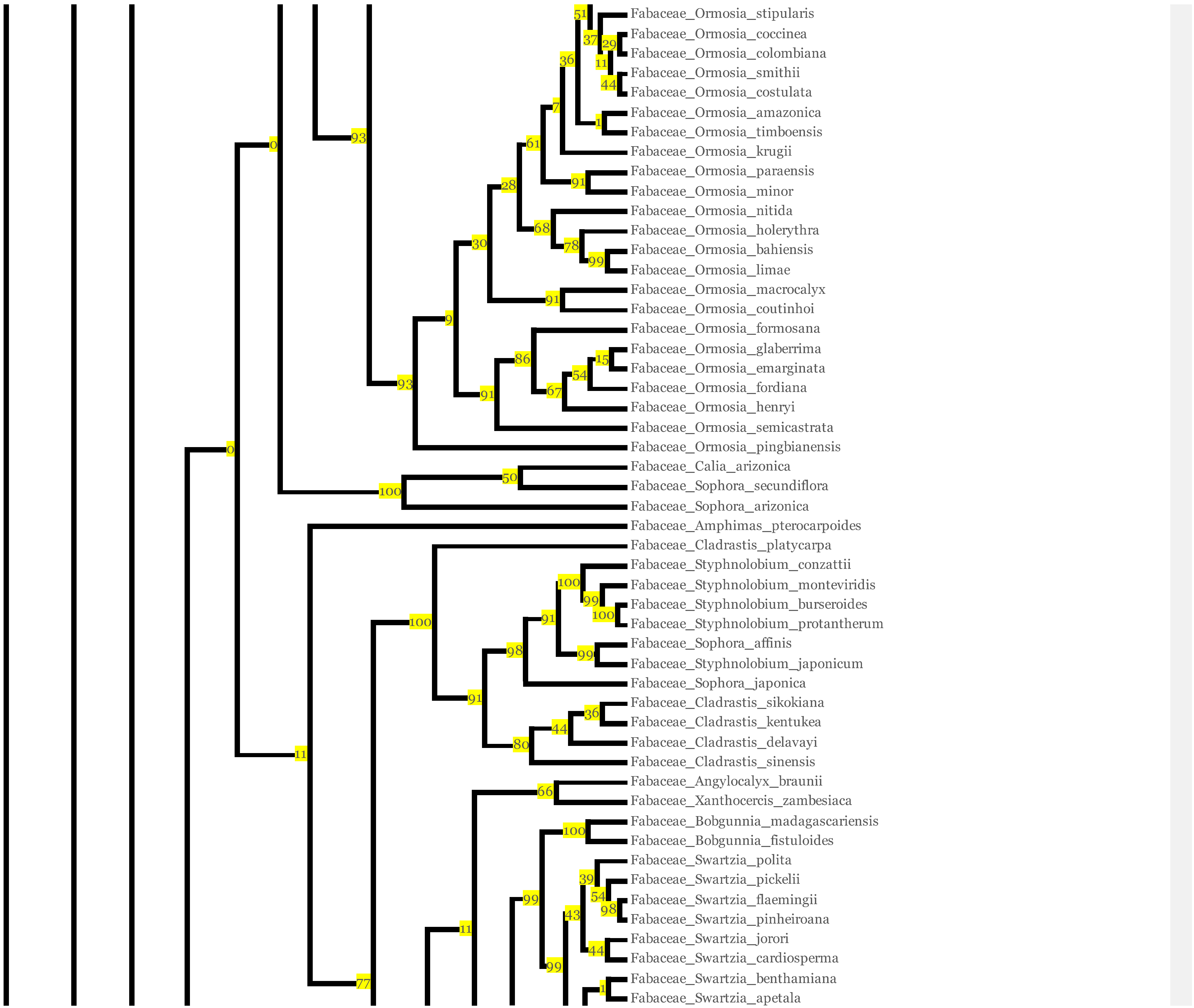

Fabaceae_Ormosia_stipularis
Fabaceae_Ormosia_coccinea
Fabaceae_Ormosia_colombiana
Fabaceae_Ormosia_smithii
Fabaceae_Ormosia_costulata
Fabaceae_Ormosia_amazonica
Fabaceae_Ormosia_timboensis
Fabaceae_Ormosia_krugii
Fabaceae_Ormosia_paraensis
Fabaceae_Ormosia_minor
Fabaceae_Ormosia_nitida
Fabaceae_Ormosia_holerythra
Fabaceae_Ormosia_bahiensis
Fabaceae_Ormosia_limae
Fabaceae_Ormosia_macrocalyx
Fabaceae_Ormosia_coutinhoi
Fabaceae_Ormosia_formosana
Fabaceae_Ormosia_glaberrima
Fabaceae_Ormosia_emarginata
Fabaceae_Ormosia_fordiana
Fabaceae_Ormosia_henryi
Fabaceae_Ormosia_semicastrata
Fabaceae_Ormosia_pingbianensis
Fabaceae_Calia_arizonica
Fabaceae_Sophora_secundiflora
Fabaceae_Sophora_arizonica
Fabaceae_Amphimas_pterocarpoides
Fabaceae_Cladrastis_platycarpa
Fabaceae_Styphnolobium_conzattii
Fabaceae_Styphnolobium_monteviridis
Fabaceae_Styphnolobium_burseroides
Fabaceae_Styphnolobium_protantherum
Fabaceae_Sophora_affinis
Fabaceae_Styphnolobium_japonicum
Fabaceae_Sophora_japonica
Fabaceae_Cladrastis_sikokiana
Fabaceae_Cladrastis_kentukea
Fabaceae_Cladrastis_delavayi
Fabaceae_Cladrastis_sinensis
Fabaceae_Angylocalyx_braunii
Fabaceae_Xanthocercis_zambesiaca
Fabaceae_Bobgunnia_madagascariensis
Fabaceae_Bobgunnia_fistuloides
Fabaceae_Swartzia_polita
Fabaceae_Swartzia_pickelii
Fabaceae_Swartzia_flaemingii
Fabaceae_Swartzia_pinheiroana
Fabaceae_Swartzia_jorori
Fabaceae_Swartzia_cardiosperma
Fabaceae_Swartzia_benthamiana
Fabaceae_Swartzia_apetala

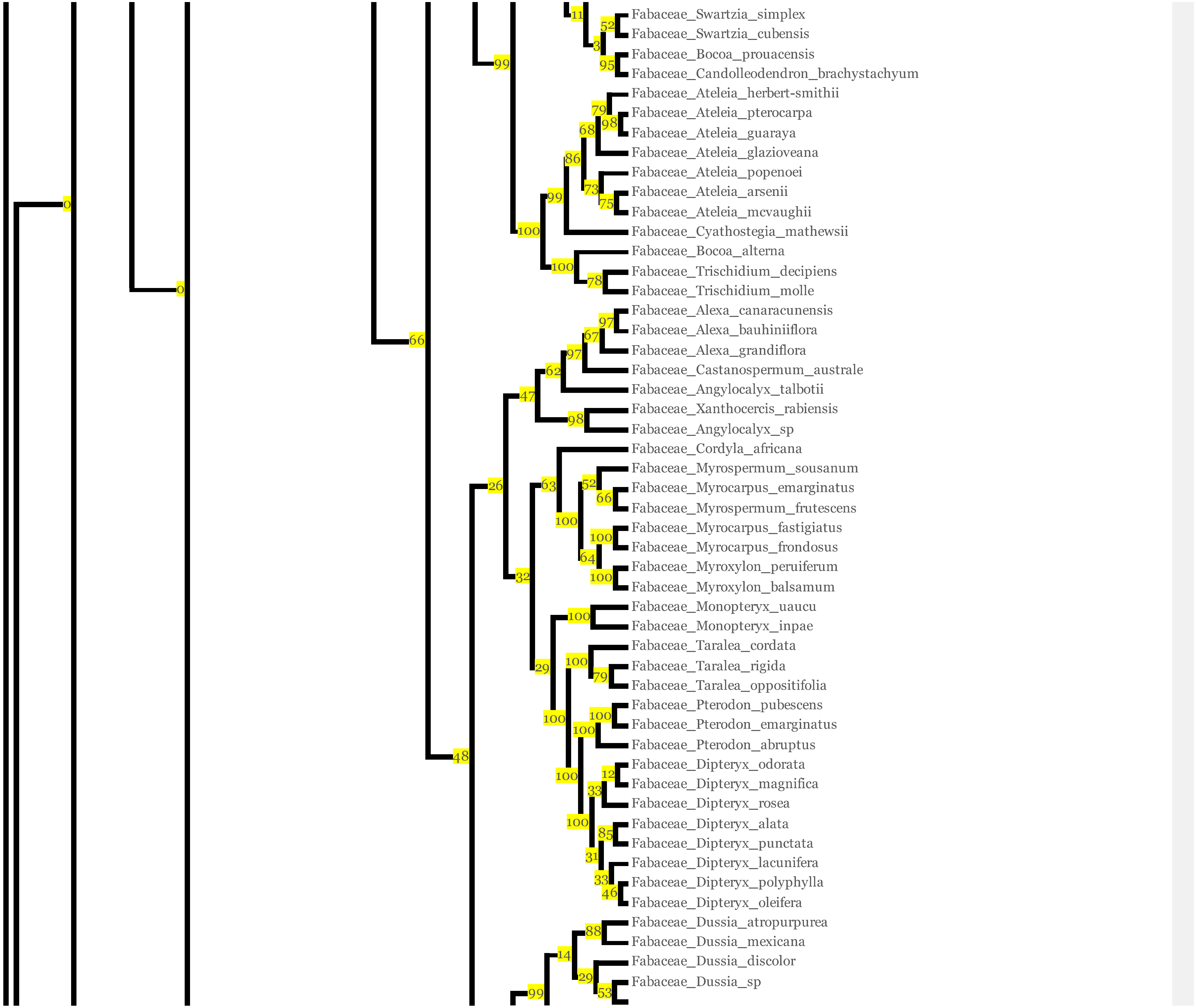
Fabaceae_Swartzia_simplex
Fabaceae_Swartzia_cubensis
Fabaceae_Bocoa_prouacensis
Fabaceae_Candolleodendron_brachystachyum
Fabaceae_Ateleia_herbert-smithii
Fabaceae_Ateleia_pterocarpa
Fabaceae_Ateleia_guaraya
Fabaceae_Ateleia_glazioveana
Fabaceae_Ateleia_popenoei
Fabaceae_Ateleia_arsenii
Fabaceae_Ateleia_mcvaughii
Fabaceae_Cyathostegia_mathewsii
Fabaceae_Bocoa_alterna
Fabaceae_Trischidium_decipiens
Fabaceae_Trischidium_molle
Fabaceae_Alexa_canaracunensis
Fabaceae_Alexa_bauhiniiflora
Fabaceae_Alexa_grandiflora
Fabaceae_Castanospermum_australe
Fabaceae_Angylocalyx_talbotii
Fabaceae_Xanthocercis_rabiensis
Fabaceae_Angylocalyx_sp
Fabaceae_Cordyla_africana
Fabaceae_Myrospermum_sousanum
Fabaceae_Myrocarpus_emarginatus
Fabaceae_Myrospermum_frutescens
Fabaceae_Myrocarpus_fastigiatus
Fabaceae_Myrocarpus_frondosus
Fabaceae_Myroxylon_peruiferum
Fabaceae_Myroxylon_balsamum
Fabaceae_Monopteryx_uaucu
Fabaceae_Monopteryx_inpae
Fabaceae_Taralea_cordata
Fabaceae_Taralea_rigida
Fabaceae_Taralea_oppositifolia
Fabaceae_Pterodon_pubescens
Fabaceae_Pterodon_emarginatus
Fabaceae_Pterodon_abruptus
Fabaceae_Dipteryx_odorata
Fabaceae_Dipteryx_magnifica
Fabaceae_Dipteryx_rosea
Fabaceae_Dipteryx_alata
Fabaceae_Dipteryx_punctata
Fabaceae_Dipteryx_lacunifera
Fabaceae_Dipteryx_polyphylla
Fabaceae_Dipteryx_oleifera
Fabaceae_Dussia_atropurpurea
Fabaceae_Dussia_mexicana
Fabaceae_Dussia_discolor
Fabaceae_Dussia_sp

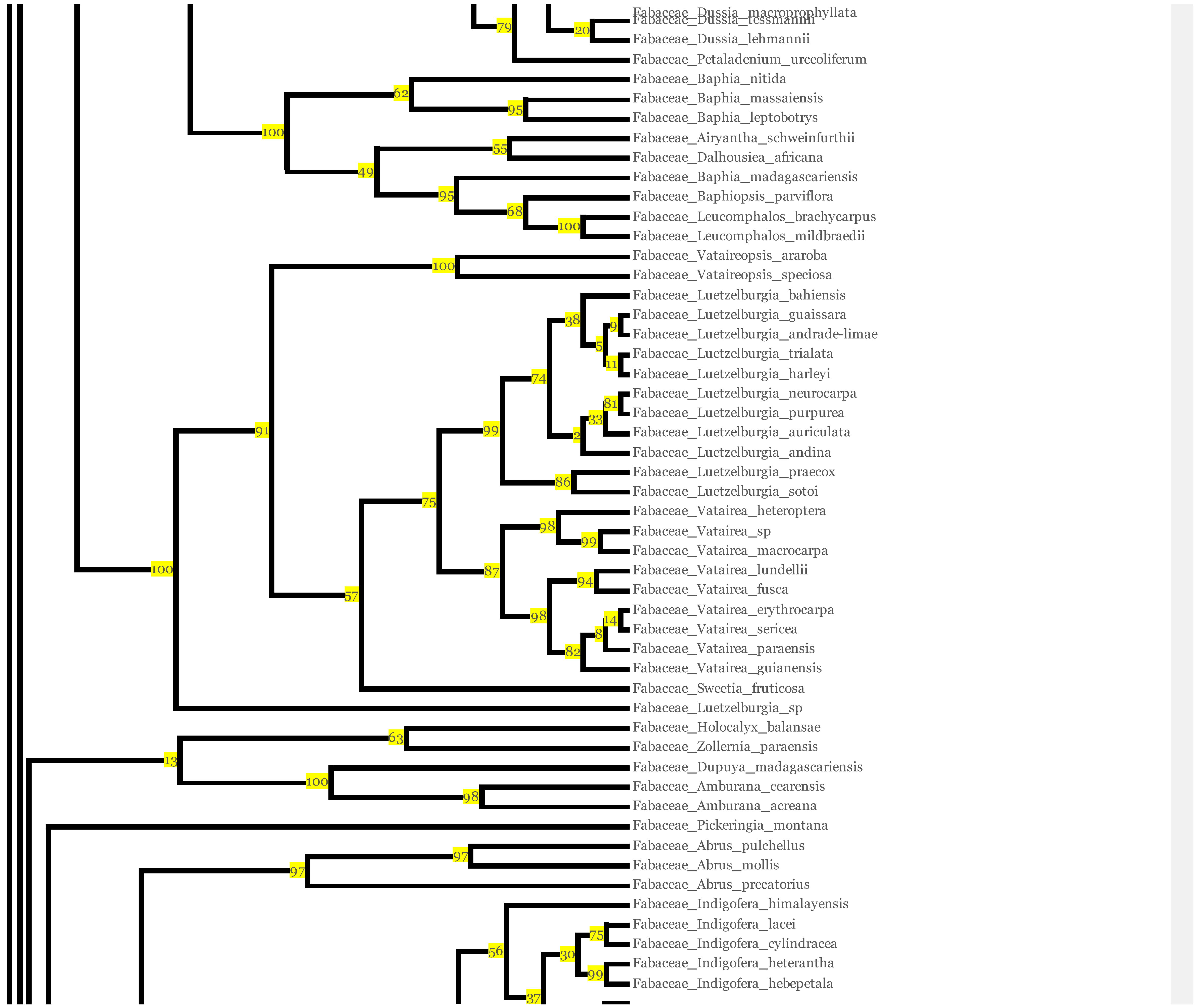

Fabaceae_Dussia_macroprophyllata
Fabaceae_Dussia_tessmannii
Fabaceae_Dussia_lehmannii
Fabaceae_Petaladenium_urceoliferum
Fabaceae_Baphia_nitida
Fabaceae_Baphia_massaiensis
Fabaceae_Baphia_leptobotrys
Fabaceae_Airyantha_schweinfurthii
Fabaceae_Dalhousiea_africana
Fabaceae_Baphia_madagascariensis
Fabaceae_Baphiopsis_parviflora
Fabaceae_Leucomphalos_brachycarpus
Fabaceae_Leucomphalos_mildbraedii
Fabaceae_Vataireopsis_araroba
Fabaceae_Vataireopsis_speciosa
Fabaceae_Luetzelburgia_bahiensis
Fabaceae_Luetzelburgia_guaissara
Fabaceae_Luetzelburgia_andrade-limae
Fabaceae_Luetzelburgia_trialata
Fabaceae_Luetzelburgia_harleyi
Fabaceae_Luetzelburgia_neurocarpa
Fabaceae_Luetzelburgia_purpurea
Fabaceae_Luetzelburgia_auriculata
Fabaceae_Luetzelburgia_andina
Fabaceae_Luetzelburgia_praecox
Fabaceae_Luetzelburgia_sotoi
Fabaceae_Vatairea_heteroptera
Fabaceae_Vatairea_sp
Fabaceae_Vatairea_macrocarpa
Fabaceae_Vatairea_lundellii
Fabaceae_Vatairea_fusca
Fabaceae_Vatairea_erythrocarpa
Fabaceae_Vatairea_sericea
Fabaceae_Vatairea_paraensis
Fabaceae_Vatairea_guianensis
Fabaceae_Sweetia_fruticosa
Fabaceae_Luetzelburgia_sp
Fabaceae_Holocalyx_balansae
Fabaceae_Zollernia_paraensis
Fabaceae_Dupuya_madagascariensis
Fabaceae_Amburana_cearensis
Fabaceae_Amburana_acreana
Fabaceae_Pickeringia_montana
Fabaceae_Abrus_pulchellus
Fabaceae_Abrus_mollis
Fabaceae_Abrus_precatorius
Fabaceae_Indigofera_himalayensis
Fabaceae_Indigofera_lacei
Fabaceae_Indigofera_cylindracea
Fabaceae_Indigofera_heterantha
Fabaceae_Indigofera_hebepetala

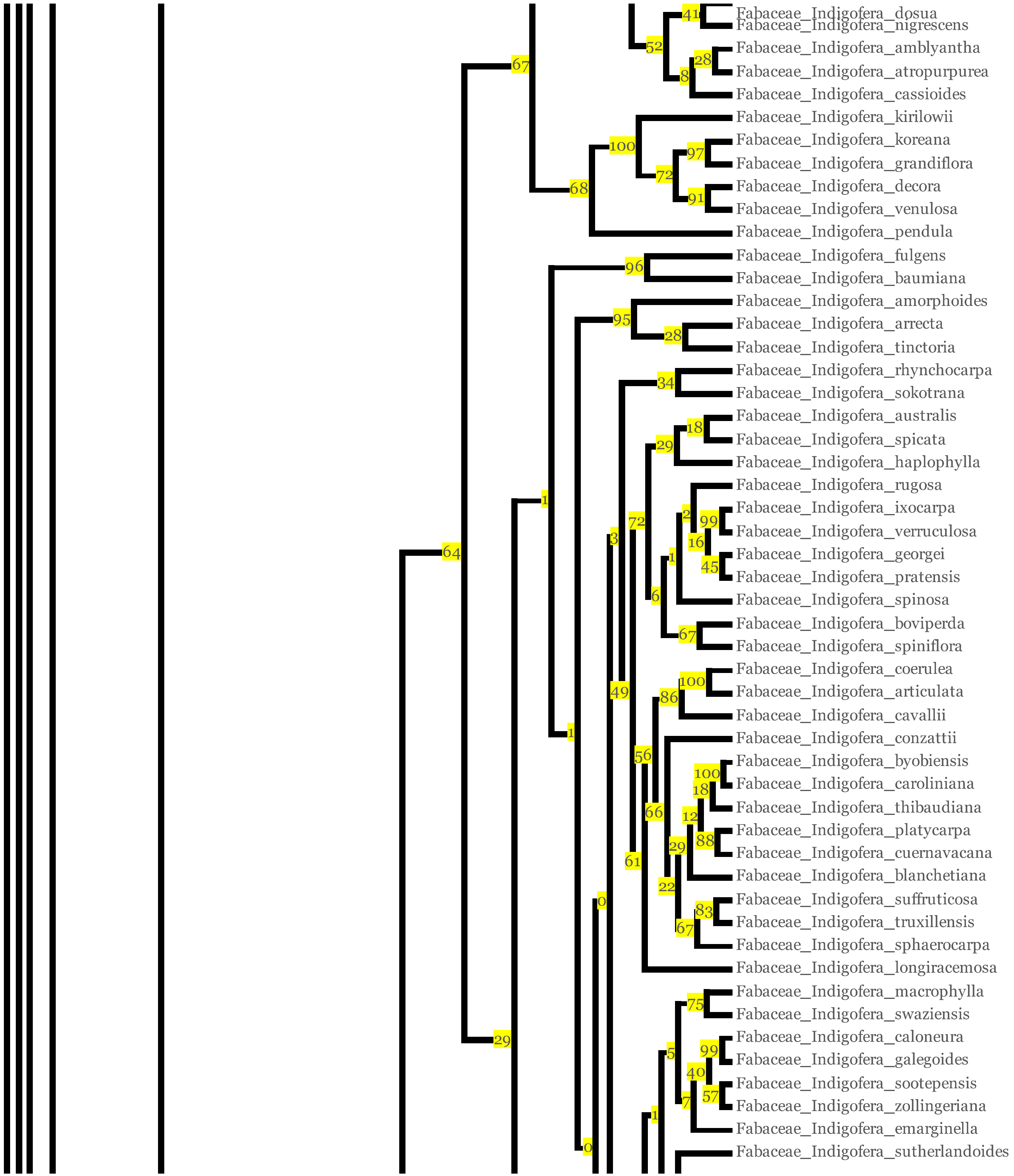
Fabaceae_Indigofera_dosua
Fabaceae_Indigofera_nigrescens
Fabaceae_Indigofera_amblyantha
Fabaceae_Indigofera_atropurpurea
Fabaceae_Indigofera_cassioides
Fabaceae_Indigofera_kirilowii
Fabaceae_Indigofera_koreana
Fabaceae_Indigofera_grandiflora
Fabaceae_Indigofera_decora
Fabaceae_Indigofera_venulosa
Fabaceae_Indigofera_pendula
Fabaceae_Indigofera_fulgens
Fabaceae_Indigofera_baumiana
Fabaceae_Indigofera_amorphoides
Fabaceae_Indigofera_arrecta
Fabaceae_Indigofera_tinctoria
Fabaceae_Indigofera_rhynchocarpa
Fabaceae_Indigofera_sokotrana
Fabaceae_Indigofera_australis
Fabaceae_Indigofera_spicata
Fabaceae_Indigofera_haplophylla
Fabaceae_Indigofera_rugosa
Fabaceae_Indigofera_ixocarpa
Fabaceae_Indigofera_verruculosa
Fabaceae_Indigofera_georgei
Fabaceae_Indigofera_pratensis
Fabaceae_Indigofera_spinosa
Fabaceae_Indigofera_boviperda
Fabaceae_Indigofera_spiniflora
Fabaceae_Indigofera_coerulea
Fabaceae_Indigofera_articulata
Fabaceae_Indigofera_cavallii
Fabaceae_Indigofera_conzattii
Fabaceae_Indigofera_byobiensis
Fabaceae_Indigofera_caroliniana
Fabaceae_Indigofera_thibaudiana
Fabaceae_Indigofera_platycarpa
Fabaceae_Indigofera_cuernavacana
Fabaceae_Indigofera_blanchetiana
Fabaceae_Indigofera_suffruticosa
Fabaceae_Indigofera_truxillensis
Fabaceae_Indigofera_sphaerocarpa
Fabaceae_Indigofera_longiracemosa
Fabaceae_Indigofera_macrophylla
Fabaceae_Indigofera_swaziensis
Fabaceae_Indigofera_caloneura
Fabaceae_Indigofera_galegoides
Fabaceae_Indigofera_sootepensis
Fabaceae_Indigofera_zollingeriana
Fabaceae_Indigofera_emarginella
Fabaceae_Indigofera_sutherlandoides

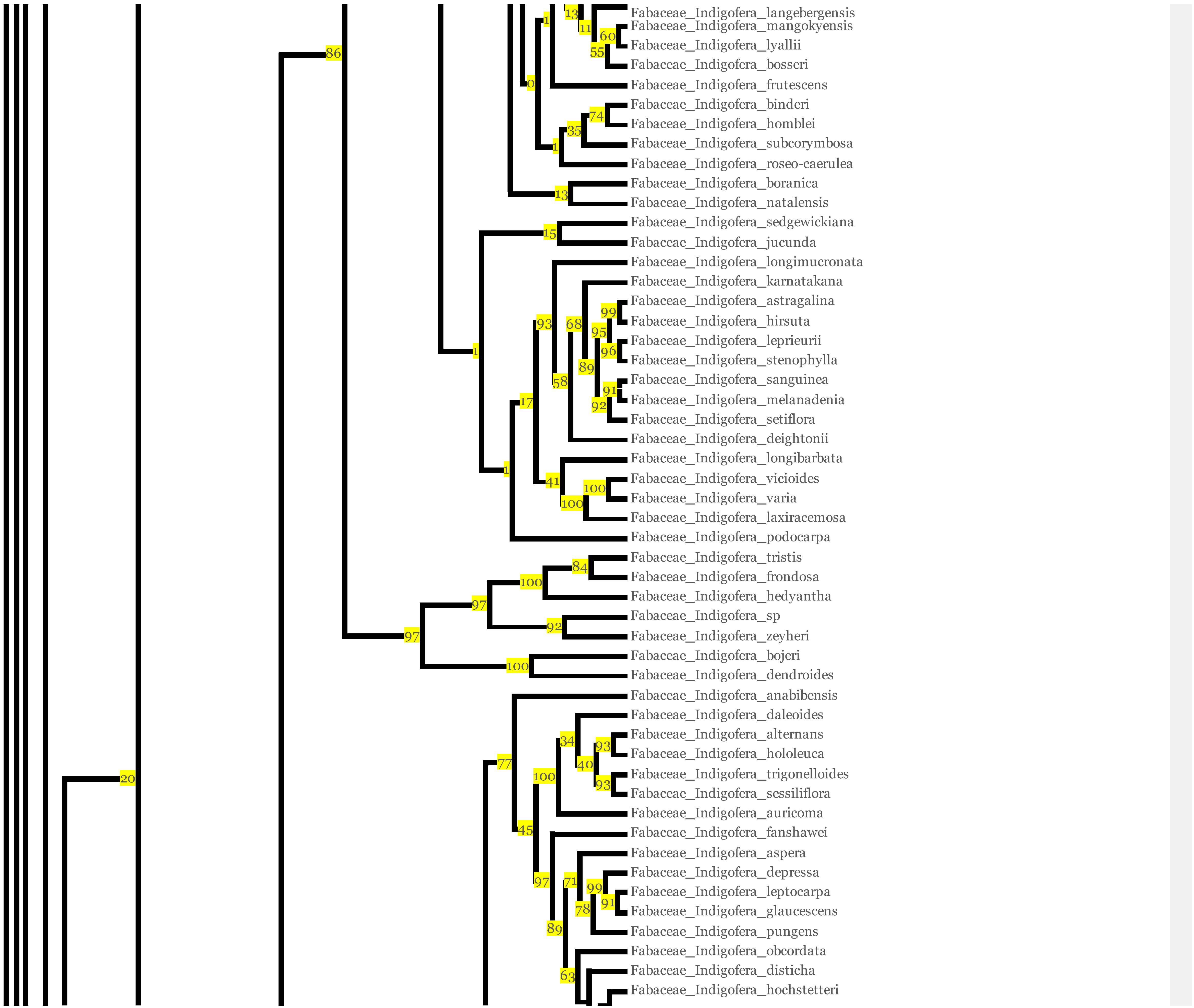
Fabaceae_Indigofera_langebergensis
Fabaceae_Indigofera_mangokyensis
Fabaceae_Indigofera_lyallii
Fabaceae_Indigofera_bosseri
Fabaceae_Indigofera_frutescens
Fabaceae_Indigofera_binderi
Fabaceae_Indigofera_homblei
Fabaceae_Indigofera_subcorymbosa
Fabaceae_Indigofera_roseo-caerulea
Fabaceae_Indigofera_boranica
Fabaceae_Indigofera_natalensis
Fabaceae_Indigofera_sedgewickiana
Fabaceae_Indigofera_jucunda
Fabaceae_Indigofera_longimucronata
Fabaceae_Indigofera_karnatakana
Fabaceae_Indigofera_astragalina
Fabaceae_Indigofera_hirsuta
Fabaceae_Indigofera_leprieurii
Fabaceae_Indigofera_stenophylla
Fabaceae_Indigofera_sanguinea
Fabaceae_Indigofera_melanadenia
Fabaceae_Indigofera_setiflora
Fabaceae_Indigofera_deightonii
Fabaceae_Indigofera_longibarbata
Fabaceae_Indigofera_vicioides
Fabaceae_Indigofera_varia
Fabaceae_Indigofera_laxiracemosa
Fabaceae_Indigofera_podocarpa
Fabaceae_Indigofera_tristis
Fabaceae_Indigofera_frondosa
Fabaceae_Indigofera_hedyantha
Fabaceae_Indigofera_sp
Fabaceae_Indigofera_zeyheri
Fabaceae_Indigofera_bojeri
Fabaceae_Indigofera_dendroides
Fabaceae_Indigofera_anabibensis
Fabaceae_Indigofera_daleoides
Fabaceae_Indigofera_alternans
Fabaceae_Indigofera_hololeuca
Fabaceae_Indigofera_trigonelloides
Fabaceae_Indigofera_sessiliflora
Fabaceae_Indigofera_auricoma
Fabaceae_Indigofera_fanshawei
Fabaceae_Indigofera_aspera
Fabaceae_Indigofera_depressa
Fabaceae_Indigofera_leptocarpa
Fabaceae_Indigofera_glaucescens
Fabaceae_Indigofera_pungens
Fabaceae_Indigofera_obcordata
Fabaceae_Indigofera_disticha
Fabaceae_Indigofera_hochstetteri

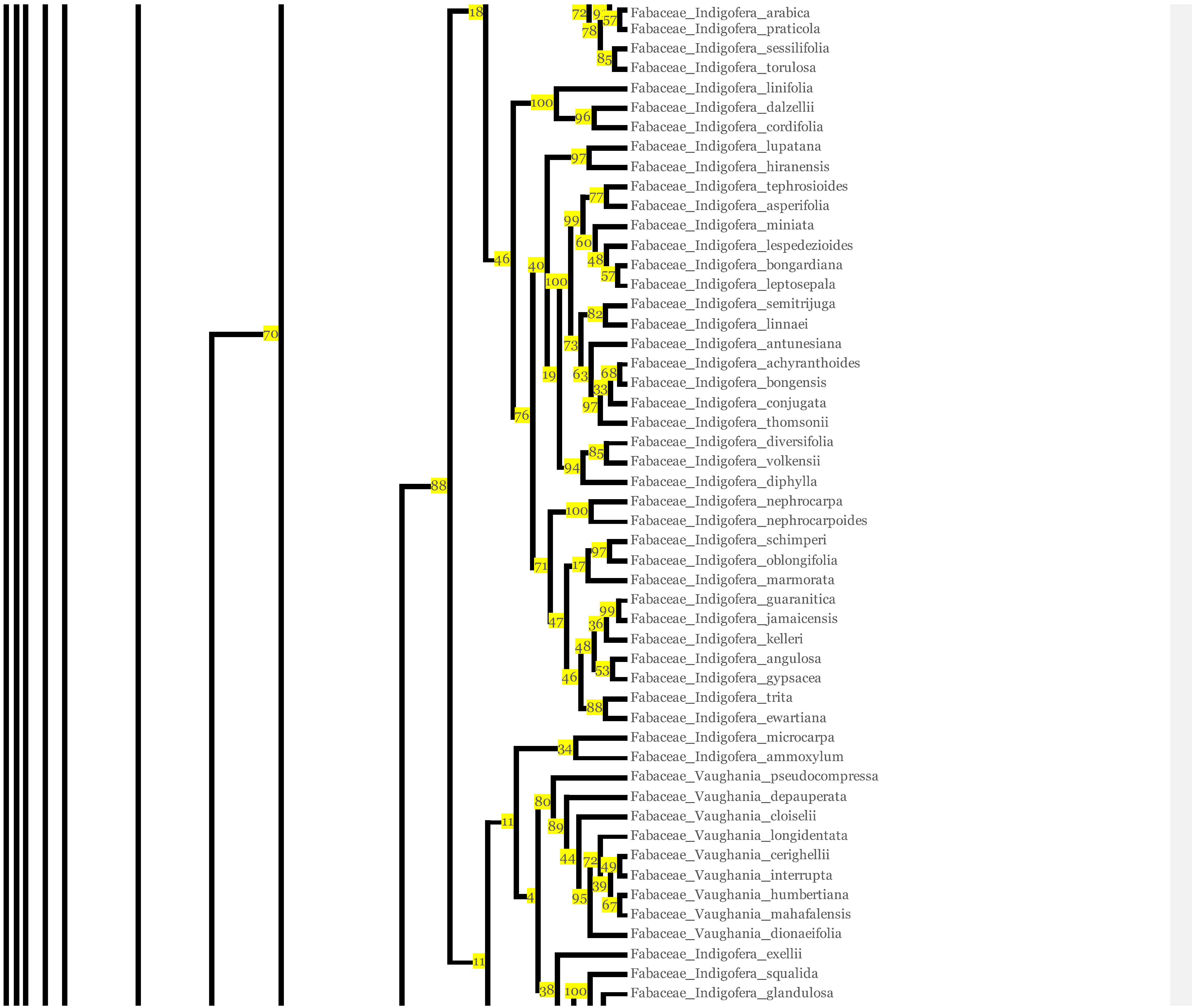
Fabaceae_Indigofera_arabica
Fabaceae_Indigofera_praticola
Fabaceae_Indigofera_sessilifolia
Fabaceae_Indigofera_torulosa
Fabaceae_Indigofera_linifolia
Fabaceae_Indigofera_dalzellii
Fabaceae_Indigofera_cordifolia
Fabaceae_Indigofera_lupatana
Fabaceae_Indigofera_hiranensis
Fabaceae_Indigofera_tephrosioides
Fabaceae_Indigofera_asperifolia
Fabaceae_Indigofera_miniata
Fabaceae_Indigofera_lespedezioides
Fabaceae_Indigofera_bongardiana
Fabaceae_Indigofera_leptosepala
Fabaceae_Indigofera_semitrijuga
Fabaceae_Indigofera_linnaei
Fabaceae_Indigofera_antunesiana
Fabaceae_Indigofera_achyranthoides
Fabaceae_Indigofera_bongensis
Fabaceae_Indigofera_conjugata
Fabaceae_Indigofera_thomsonii
Fabaceae_Indigofera_diversifolia
Fabaceae_Indigofera_volkensii
Fabaceae_Indigofera_diphylla
Fabaceae_Indigofera_nephrocarpa
Fabaceae_Indigofera_nephrocarpoides
Fabaceae_Indigofera_schimperi
Fabaceae_Indigofera_oblongifolia
Fabaceae_Indigofera_marmorata
Fabaceae_Indigofera_guaranitica
Fabaceae_Indigofera_jamaicensis
Fabaceae_Indigofera_kelleri
Fabaceae_Indigofera_angulosa
Fabaceae_Indigofera_gypsacea
Fabaceae_Indigofera_trita
Fabaceae_Indigofera_ewartiana
Fabaceae_Indigofera_microcarpa
Fabaceae_Indigofera_ammoxylum
Fabaceae_Vaughania_pseudocompressa
Fabaceae_Vaughania_depauperata
Fabaceae_Vaughania_cloiselii
Fabaceae_Vaughania_longidentata
Fabaceae_Vaughania_cerighellii
Fabaceae_Vaughania_interrupta
Fabaceae_Vaughania_humbertiana
Fabaceae_Vaughania_mahafalensis
Fabaceae_Vaughania_dionaeifolia
Fabaceae_Indigofera_exellii
Fabaceae_Indigofera_squalida
Fabaceae_Indigofera_glandulosa

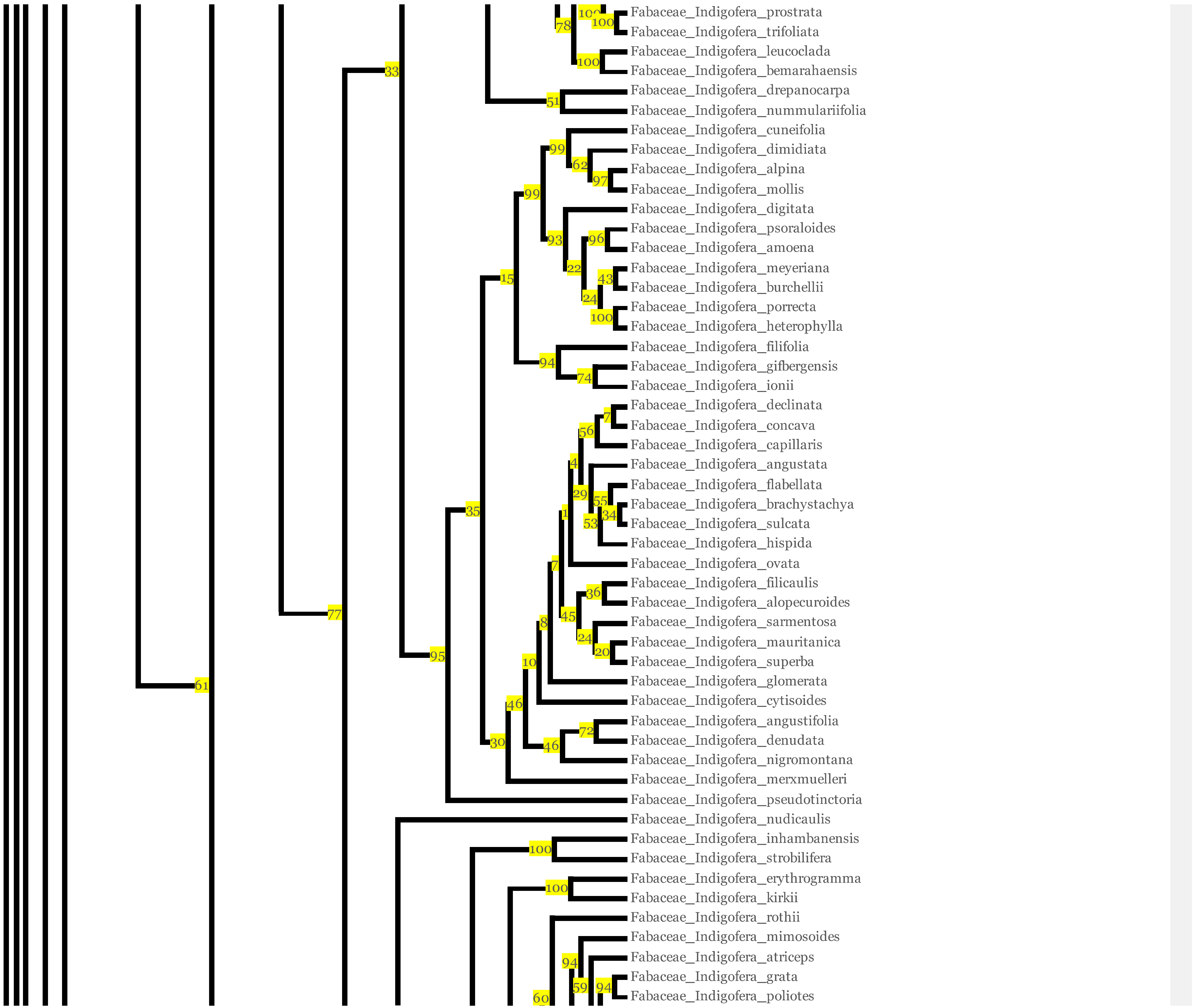

Fabaceae_Indigofera_prostrata
Fabaceae_Indigofera_trifoliata
Fabaceae_Indigofera_leucoclada
Fabaceae_Indigofera_bemarahaensis
Fabaceae_Indigofera_drepanocarpa
Fabaceae_Indigofera_nummulariifolia
Fabaceae_Indigofera_cuneifolia
Fabaceae_Indigofera_dimidiata
Fabaceae_Indigofera_alpina
Fabaceae_Indigofera_mollis
Fabaceae_Indigofera_digitata
Fabaceae_Indigofera_psoraloides
Fabaceae_Indigofera_amoena
Fabaceae_Indigofera_meyeriana
Fabaceae_Indigofera_burchellii
Fabaceae_Indigofera_porrecta
Fabaceae_Indigofera_heterophylla
Fabaceae_Indigofera_filifolia
Fabaceae_Indigofera_gifbergensis
Fabaceae_Indigofera_ionii
Fabaceae_Indigofera_declinata
Fabaceae_Indigofera_concava
Fabaceae_Indigofera_capillaris
Fabaceae_Indigofera_angustata
Fabaceae_Indigofera_flabellata
Fabaceae_Indigofera_brachystachya
Fabaceae_Indigofera_sulcata
Fabaceae_Indigofera_hispida
Fabaceae_Indigofera_ovata
Fabaceae_Indigofera_filicaulis
Fabaceae_Indigofera_alopecuroides
Fabaceae_Indigofera_sarmentosa
Fabaceae_Indigofera_mauritanica
Fabaceae_Indigofera_superba
Fabaceae_Indigofera_glomerata
Fabaceae_Indigofera_cytisoides
Fabaceae_Indigofera_angustifolia
Fabaceae_Indigofera_denudata
Fabaceae_Indigofera_nigromontana
Fabaceae_Indigofera_merxmuelleri
Fabaceae_Indigofera_pseudotinctoria
Fabaceae_Indigofera_nudicaulis
Fabaceae_Indigofera_inhambanensis
Fabaceae_Indigofera_strobilifera
Fabaceae_Indigofera_erythrogramma
Fabaceae_Indigofera_kirkii
Fabaceae_Indigofera_rothii
Fabaceae_Indigofera_mimosoides
Fabaceae_Indigofera_atriceps
Fabaceae_Indigofera_grata
Fabaceae_Indigofera_poliotes

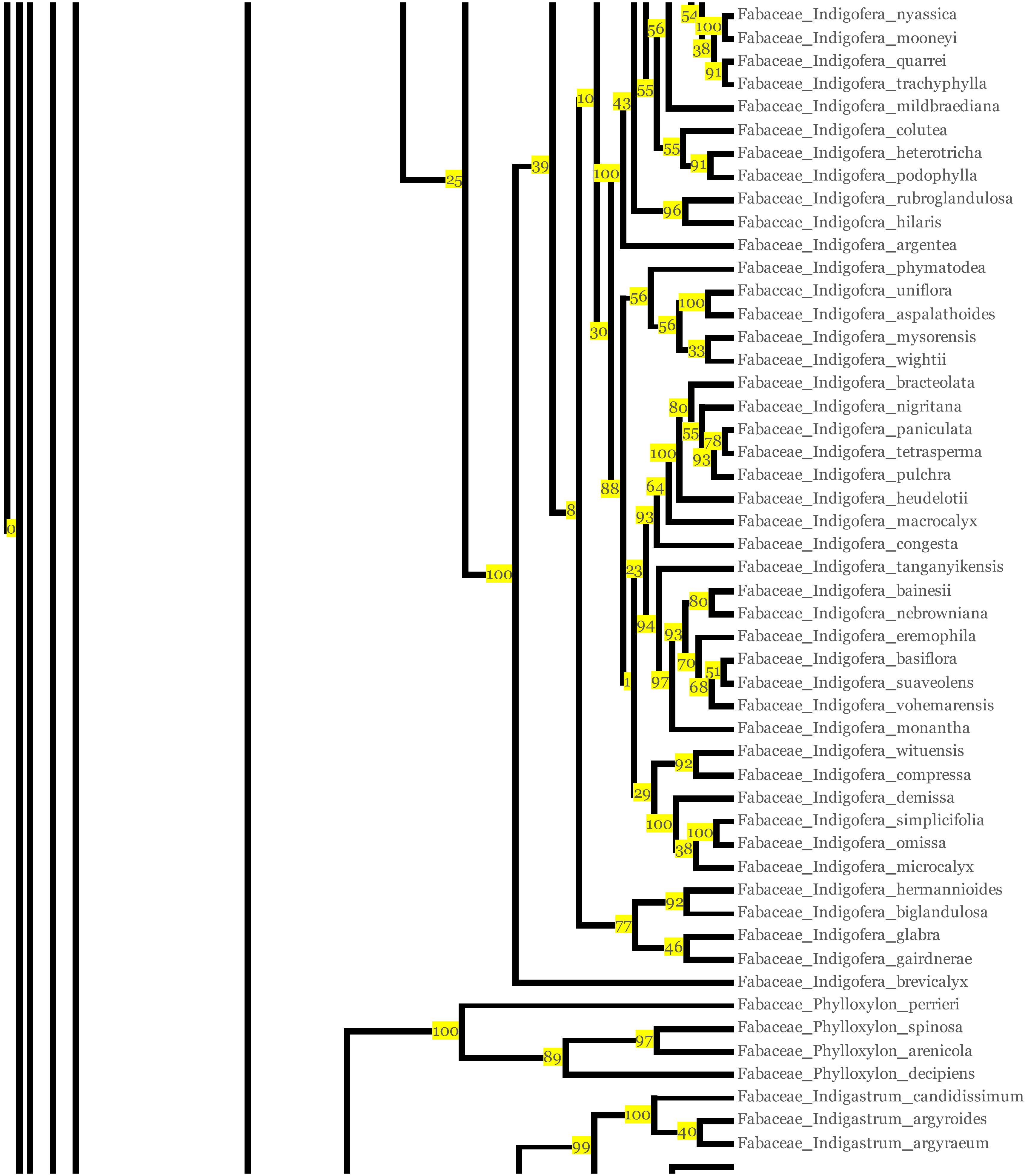
Fabaceae_Indigofera_nyassica
Fabaceae_Indigofera_mooneyi
Fabaceae_Indigofera_quarrei
Fabaceae_Indigofera_trachyphylla
Fabaceae_Indigofera_mildbraediana
Fabaceae_Indigofera_colutea
Fabaceae_Indigofera_heterotricha
Fabaceae_Indigofera_podophylla
Fabaceae_Indigofera_rubroglandulosa
Fabaceae_Indigofera_hilaris
Fabaceae_Indigofera_argentea
Fabaceae_Indigofera_phymatodea
Fabaceae_Indigofera_uniflora
Fabaceae_Indigofera_aspalathoides
Fabaceae_Indigofera_mysorensis
Fabaceae_Indigofera_wightii
Fabaceae_Indigofera_bracteolata
Fabaceae_Indigofera_nigritana
Fabaceae_Indigofera_paniculata
Fabaceae_Indigofera_tetrasperma
Fabaceae_Indigofera_pulchra
Fabaceae_Indigofera_heudelotii
Fabaceae_Indigofera_macrocalyx
Fabaceae_Indigofera_congesta
Fabaceae_Indigofera_tanganyikensis
Fabaceae_Indigofera_bainesii
Fabaceae_Indigofera_nebrowniana
Fabaceae_Indigofera_eremophila
Fabaceae_Indigofera_basiflora
Fabaceae_Indigofera_suaveolens
Fabaceae_Indigofera_vohemarensis
Fabaceae_Indigofera_monantha
Fabaceae_Indigofera_wituensis
Fabaceae_Indigofera_compressa
Fabaceae_Indigofera_demissa
Fabaceae_Indigofera_simplicifolia
Fabaceae_Indigofera_omissa
Fabaceae_Indigofera_microcalyx
Fabaceae_Indigofera_hermannioides
Fabaceae_Indigofera_biglandulosa
Fabaceae_Indigofera_glabra
Fabaceae_Indigofera_gairdnerae
Fabaceae_Indigofera_brevicalyx
Fabaceae_Phylloxylon_perrieri
Fabaceae_Phylloxylon_spinosa
Fabaceae_Phylloxylon_arenicola
Fabaceae_Phylloxylon_decipiens
Fabaceae_Indigastrum_candidissimum
Fabaceae_Indigastrum_argyroides
Fabaceae_Indigastrum_argyraeum

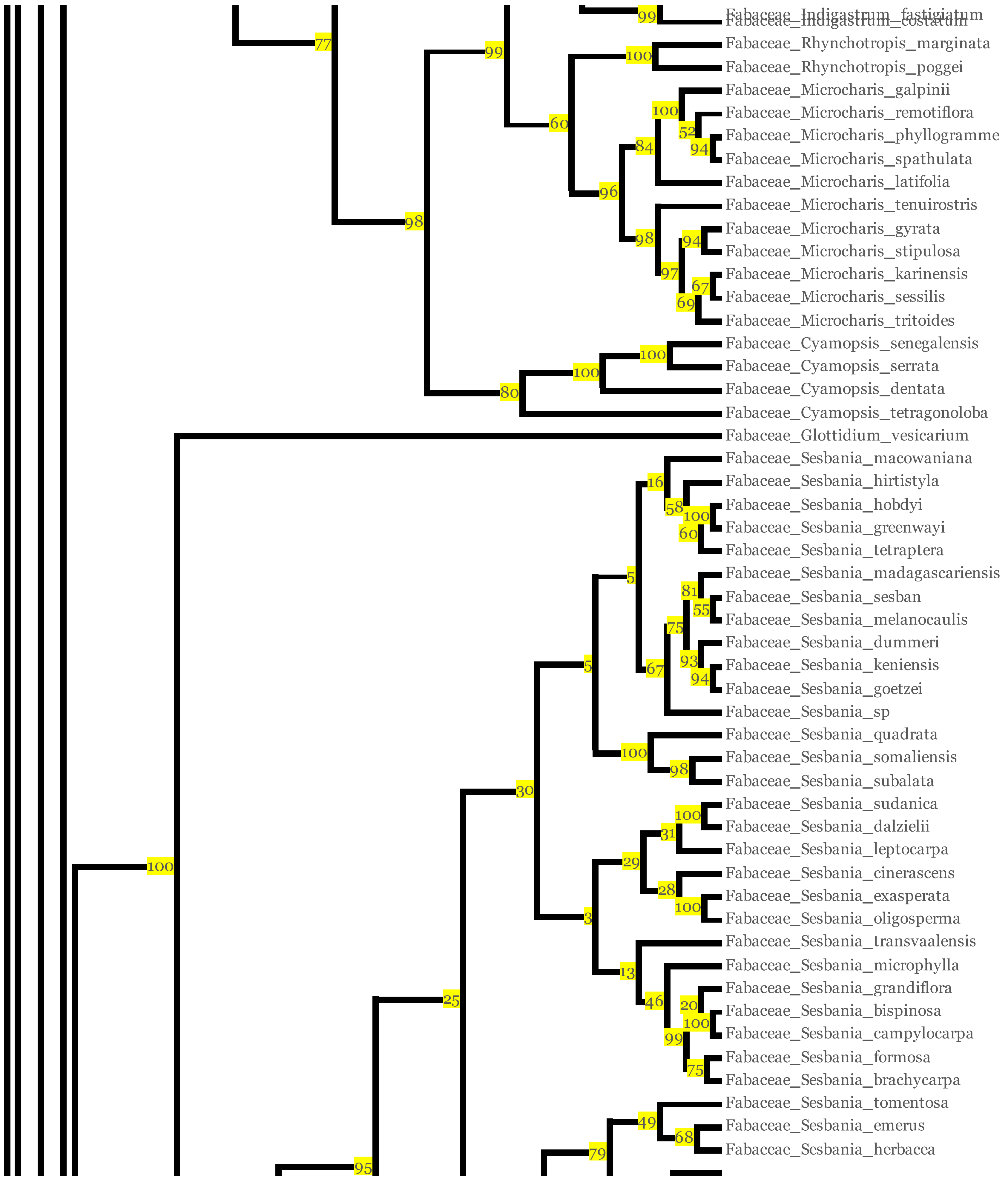
Fabaceae_Indigastrum_fastigiatum
Fabaceae_Indigastrum_costatum
Fabaceae_Rhynchotropis_marginata
Fabaceae_Rhynchotropis_poggei
Fabaceae_Microcharis_galpinii
Fabaceae_Microcharis_remotiflora
Fabaceae_Microcharis_phyllogramme
Fabaceae_Microcharis_spathulata
Fabaceae_Microcharis_latifolia
Fabaceae_Microcharis_tenuirostris
Fabaceae_Microcharis_gyrata
Fabaceae_Microcharis_stipulosa
Fabaceae_Microcharis_karinensis
Fabaceae_Microcharis_sessilis
Fabaceae_Microcharis_tritoides
Fabaceae_Cyamopsis_senegalensis
Fabaceae_Cyamopsis_serrata
Fabaceae_Cyamopsis_dentata
Fabaceae_Cyamopsis_tetragonoloba
Fabaceae_Glottidium_vesicarium
Fabaceae_Sesbania_macowaniana
Fabaceae_Sesbania_hirtistyla
Fabaceae_Sesbania_hobdyi
Fabaceae_Sesbania_greenwayi
Fabaceae_Sesbania_tetraptera
Fabaceae_Sesbania_madagascariensis
Fabaceae_Sesbania_sesban
Fabaceae_Sesbania_melanocaulis
Fabaceae_Sesbania_dummeri
Fabaceae_Sesbania_keniensis
Fabaceae_Sesbania_goetzei
Fabaceae_Sesbania_sp
Fabaceae_Sesbania_quadrata
Fabaceae_Sesbania_somaliensis
Fabaceae_Sesbania_subalata
Fabaceae_Sesbania_sudanica
Fabaceae_Sesbania_dalzielii
Fabaceae_Sesbania_leptocarpa
Fabaceae_Sesbania_cinerascens
Fabaceae_Sesbania_exasperata
Fabaceae_Sesbania_oligosperma
Fabaceae_Sesbania_transvaalensis
Fabaceae_Sesbania_microphylla
Fabaceae_Sesbania_grandiflora
Fabaceae_Sesbania_bispinosa
Fabaceae_Sesbania_campylocarpa
Fabaceae_Sesbania_formosa
Fabaceae_Sesbania_brachycarpa
Fabaceae_Sesbania_tomentosa
Fabaceae_Sesbania_emerus
Fabaceae_Sesbania_herbacea

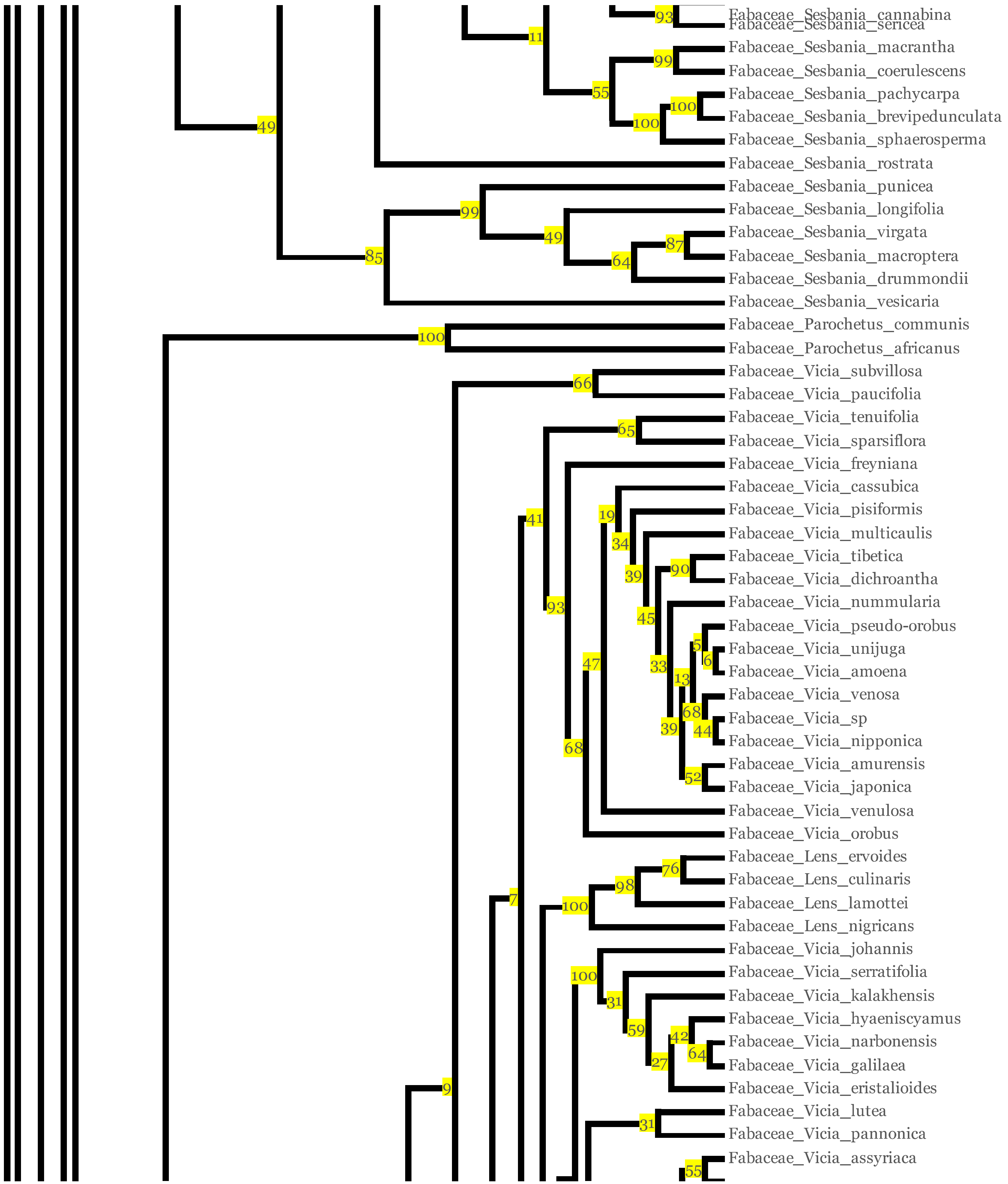
Fabaceae_Sesbania_cannabina
Fabaceae_Sesbania_sericea
Fabaceae_Sesbania_macrantha
Fabaceae_Sesbania_coerulescens
Fabaceae_Sesbania_pachycarpa
Fabaceae_Sesbania_brevipedunculata
Fabaceae_Sesbania_sphaerosperma
Fabaceae_Sesbania_rostrata
Fabaceae_Sesbania_punicea
Fabaceae_Sesbania_longifolia
Fabaceae_Sesbania_virgata
Fabaceae_Sesbania_macroptera
Fabaceae_Sesbania_drummondii
Fabaceae_Sesbania_vesicaria
Fabaceae_Parochetus_communis
Fabaceae_Parochetus_africanus
Fabaceae_Vicia_subvillosa
Fabaceae_Vicia_paucifolia
Fabaceae_Vicia_tenuifolia
Fabaceae_Vicia_sparsiflora
Fabaceae_Vicia_freyniana
Fabaceae_Vicia_cassubica
Fabaceae_Vicia_pisiformis
Fabaceae_Vicia_multicaulis
Fabaceae_Vicia_tibetica
Fabaceae_Vicia_dichroantha
Fabaceae_Vicia_nummularia
Fabaceae_Vicia_pseudo-orobus
Fabaceae_Vicia_unijuga
Fabaceae_Vicia_amoena
Fabaceae_Vicia_venosa
Fabaceae_Vicia_sp
Fabaceae_Vicia_nipponica
Fabaceae_Vicia_amurensis
Fabaceae_Vicia_japonica
Fabaceae_Vicia_venulosa
Fabaceae_Vicia_orobus
Fabaceae_Lens_ervoides
Fabaceae_Lens_culinaris
Fabaceae_Lens_lamottei
Fabaceae_Lens_nigricans
Fabaceae_Vicia_johannis
Fabaceae_Vicia_serratifolia
Fabaceae_Vicia_kalakhensis
Fabaceae_Vicia_hyaeniscyamus
Fabaceae_Vicia_narbonensis
Fabaceae_Vicia_galilaea
Fabaceae_Vicia_eristalioides
Fabaceae_Vicia_lutea
Fabaceae_Vicia_pannonica
Fabaceae_Vicia_assyriaca

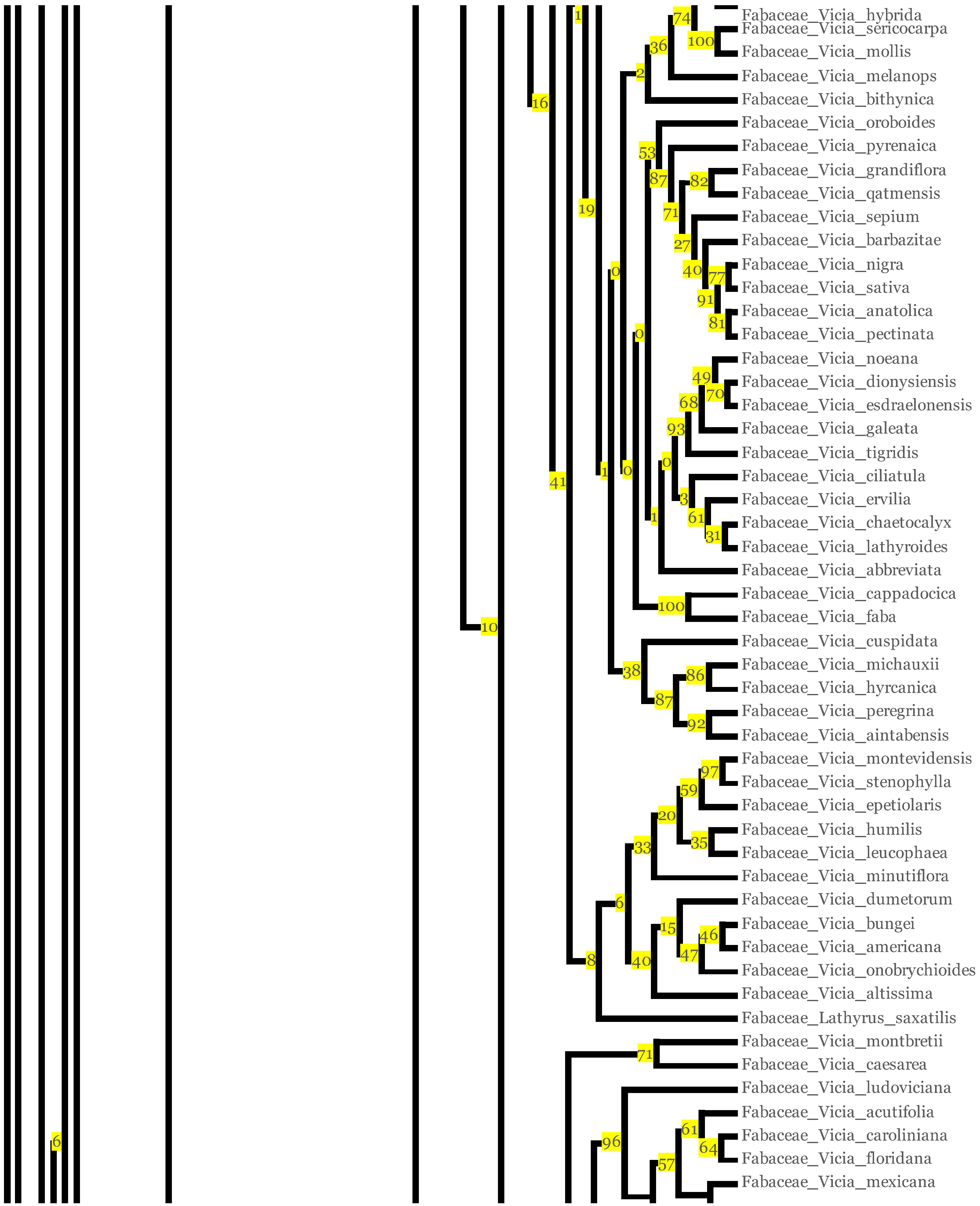
Fabaceae_Vicia_hybrida
Fabaceae_Vicia_sericocarpa
Fabaceae_Vicia_mollis
Fabaceae_Vicia_melanops
Fabaceae_Vicia_bithynica
Fabaceae_Vicia_oroboides
Fabaceae_Vicia_pyrenaica
Fabaceae_Vicia_grandiflora
Fabaceae_Vicia_qatmensis
Fabaceae_Vicia_sepium
Fabaceae_Vicia_barbazitae
Fabaceae_Vicia_nigra
Fabaceae_Vicia_sativa
Fabaceae_Vicia_anatolica
Fabaceae_Vicia_pectinata
Fabaceae_Vicia_noeana
Fabaceae_Vicia_dionysiensis
Fabaceae_Vicia_esdraelonensis
Fabaceae_Vicia_galeata
Fabaceae_Vicia_tigridis
Fabaceae_Vicia_ciliatula
Fabaceae_Vicia_ervilia
Fabaceae_Vicia_chaetocalyx
Fabaceae_Vicia_lathyroides
Fabaceae_Vicia_abbreviata
Fabaceae_Vicia_cappadocica
Fabaceae_Vicia_faba
Fabaceae_Vicia_cuspidata
Fabaceae_Vicia_michauxii
Fabaceae_Vicia_hyrcanica
Fabaceae_Vicia_peregrina
Fabaceae_Vicia_aintabensis
Fabaceae_Vicia_montevidensis
Fabaceae_Vicia_stenophylla
Fabaceae_Vicia_epetiolaris
Fabaceae_Vicia_humilis
Fabaceae_Vicia_leucophaea
Fabaceae_Vicia_minutiflora
Fabaceae_Vicia_dumetorum
Fabaceae_Vicia_bungei
Fabaceae_Vicia_americana
Fabaceae_Vicia_onobrychioides
Fabaceae_Vicia_altissima
Fabaceae_Lathyrus_saxatilis
Fabaceae_Vicia_montbretii
Fabaceae_Vicia_caesarea
Fabaceae_Vicia_ludoviciana
Fabaceae_Vicia_acutifolia
Fabaceae_Vicia_caroliniana
Fabaceae_Vicia_floridana
Fabaceae_Vicia_mexicana

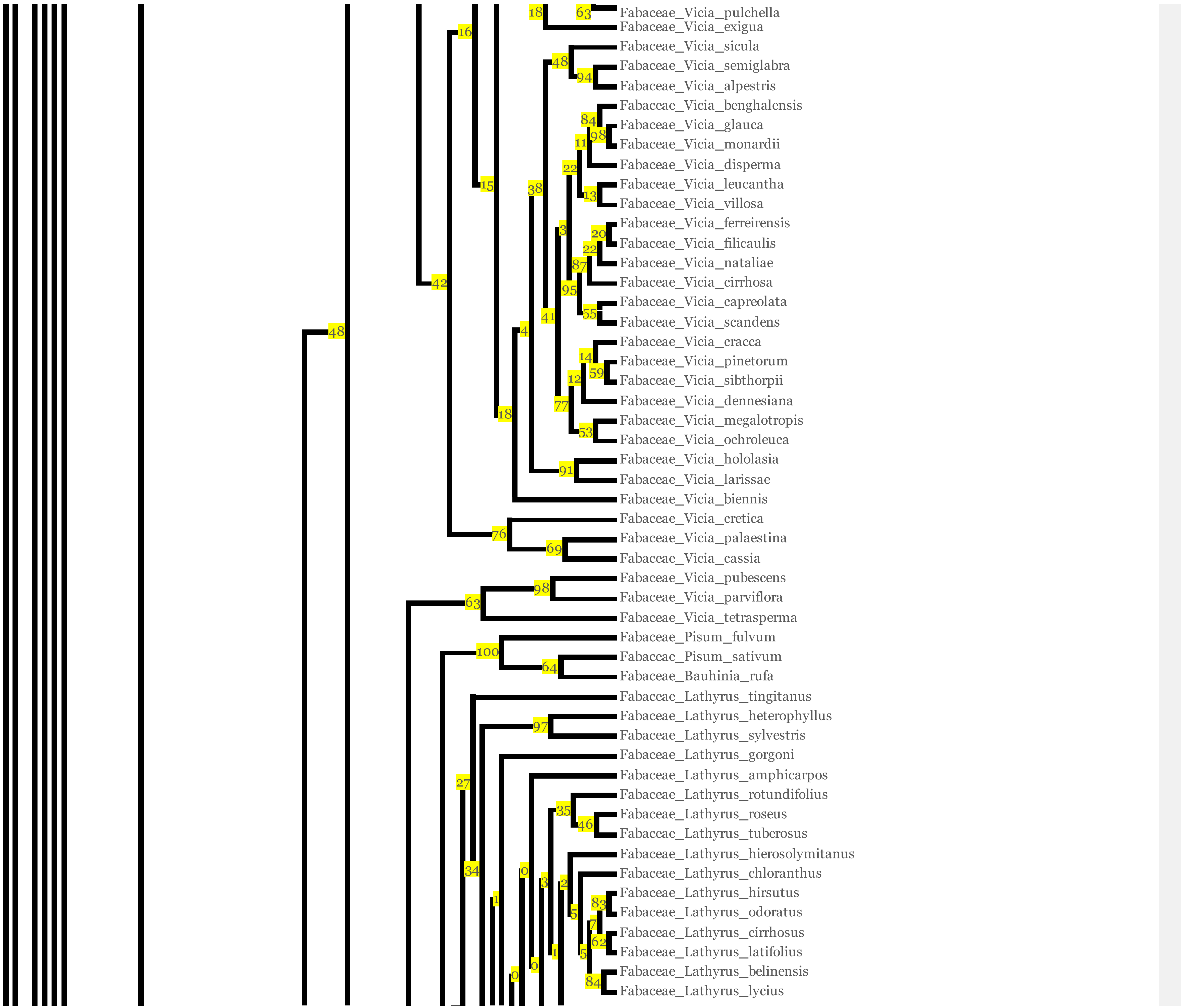
Fabaceae_Vicia_pulchella
Fabaceae_Vicia_exigua
Fabaceae_Vicia_sicula
Fabaceae_Vicia_semiglabra
Fabaceae_Vicia_alpestris
Fabaceae_Vicia_benghalensis
Fabaceae_Vicia_glauca
Fabaceae_Vicia_monardii
Fabaceae_Vicia_disperma
Fabaceae_Vicia_leucantha
Fabaceae_Vicia_villosa
Fabaceae_Vicia_ferreirensis
Fabaceae_Vicia_filicaulis
Fabaceae_Vicia_nataliae
Fabaceae_Vicia_cirrhosa
Fabaceae_Vicia_capreolata
Fabaceae_Vicia_scandens
Fabaceae_Vicia_cracca
Fabaceae_Vicia_pinetorum
Fabaceae_Vicia_sibthorpii
Fabaceae_Vicia_dennesiana
Fabaceae_Vicia_megalotropis
Fabaceae_Vicia_ochroleuca
Fabaceae_Vicia_hololasia
Fabaceae_Vicia_larissae
Fabaceae_Vicia_biennis
Fabaceae_Vicia_cretica
Fabaceae_Vicia_palaestina
Fabaceae_Vicia_cassia
Fabaceae_Vicia_pubescens
Fabaceae_Vicia_parviflora
Fabaceae_Vicia_tetrasperma
Fabaceae_Pisum_fulvum
Fabaceae_Pisum_sativum
Fabaceae_Bauhinia_rufa
Fabaceae_Lathyrus_tingitanus
Fabaceae_Lathyrus_heterophyllus
Fabaceae_Lathyrus_sylvestris
Fabaceae_Lathyrus_gorgoni
Fabaceae_Lathyrus_amphicarpos
Fabaceae_Lathyrus_rotundifolius
Fabaceae_Lathyrus_roseus
Fabaceae_Lathyrus_tuberosus
Fabaceae_Lathyrus_hierosolymitanus
Fabaceae_Lathyrus_chloranthus
Fabaceae_Lathyrus_hirsutus
Fabaceae_Lathyrus_odoratus
Fabaceae_Lathyrus_cirrhosus
Fabaceae_Lathyrus_latifolius
Fabaceae_Lathyrus_belinensis
Fabaceae_Lathyrus_lycius

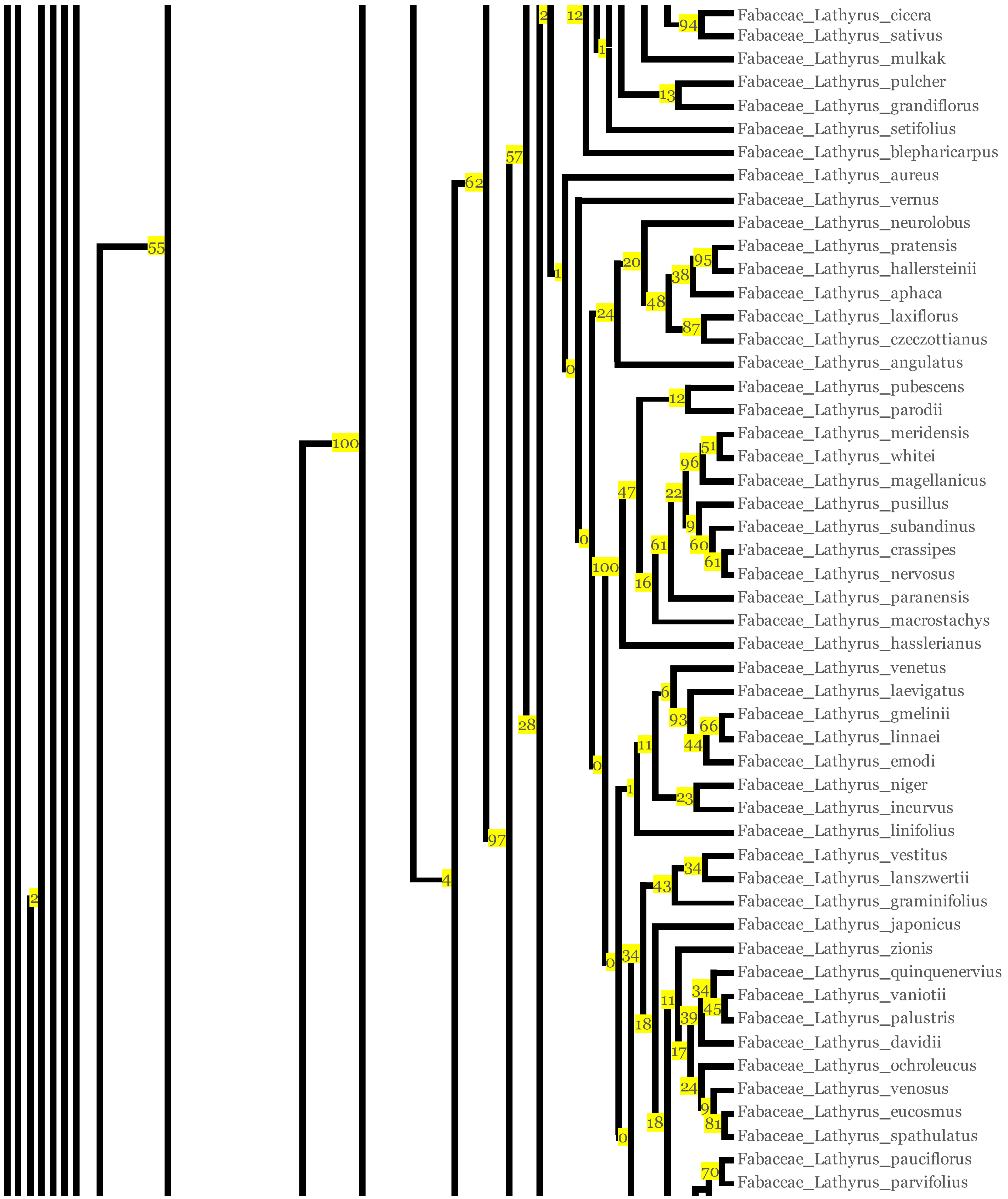

Fabaceae_Lathyrus_cicera
Fabaceae_Lathyrus_sativus
Fabaceae_Lathyrus_mulkak
Fabaceae_Lathyrus_pulcher
Fabaceae_Lathyrus_grandiflorus
Fabaceae_Lathyrus_setifolius
Fabaceae_Lathyrus_blepharicarpus
Fabaceae_Lathyrus_aureus
Fabaceae_Lathyrus_vernus
Fabaceae_Lathyrus_neurolobus
Fabaceae_Lathyrus_pratensis
Fabaceae_Lathyrus_hallersteinii
Fabaceae_Lathyrus_aphaca
Fabaceae_Lathyrus_laxiflorus
Fabaceae_Lathyrus_czeczottianus
Fabaceae_Lathyrus_angulatus
Fabaceae_Lathyrus_pubescens
Fabaceae_Lathyrus_parodii
Fabaceae_Lathyrus_meridensis
Fabaceae_Lathyrus_whitei
Fabaceae_Lathyrus_magellanicus
Fabaceae_Lathyrus_pusillus
Fabaceae_Lathyrus_subandinus
Fabaceae_Lathyrus_crassipes
Fabaceae_Lathyrus_nervosus
Fabaceae_Lathyrus_paranensis
Fabaceae_Lathyrus_macrostachys
Fabaceae_Lathyrus_hasslerianus
Fabaceae_Lathyrus_venetus
Fabaceae_Lathyrus_laevigatus
Fabaceae_Lathyrus_gmelinii
Fabaceae_Lathyrus_linnaei
Fabaceae_Lathyrus_emodi
Fabaceae_Lathyrus_niger
Fabaceae_Lathyrus_incurvus
Fabaceae_Lathyrus_linifolius
Fabaceae_Lathyrus_vestitus
Fabaceae_Lathyrus_lanszwertii
Fabaceae_Lathyrus_graminifolius
Fabaceae_Lathyrus_japonicus
Fabaceae_Lathyrus_zionis
Fabaceae_Lathyrus_quinquenervius
Fabaceae_Lathyrus_vaniotii
Fabaceae_Lathyrus_palustris
Fabaceae_Lathyrus_davidii
Fabaceae_Lathyrus_ochroleucus
Fabaceae_Lathyrus_venosus
Fabaceae_Lathyrus_eucosmus
Fabaceae_Lathyrus_spathulatus
Fabaceae_Lathyrus_pauciflorus
Fabaceae_Lathyrus_parvifolius

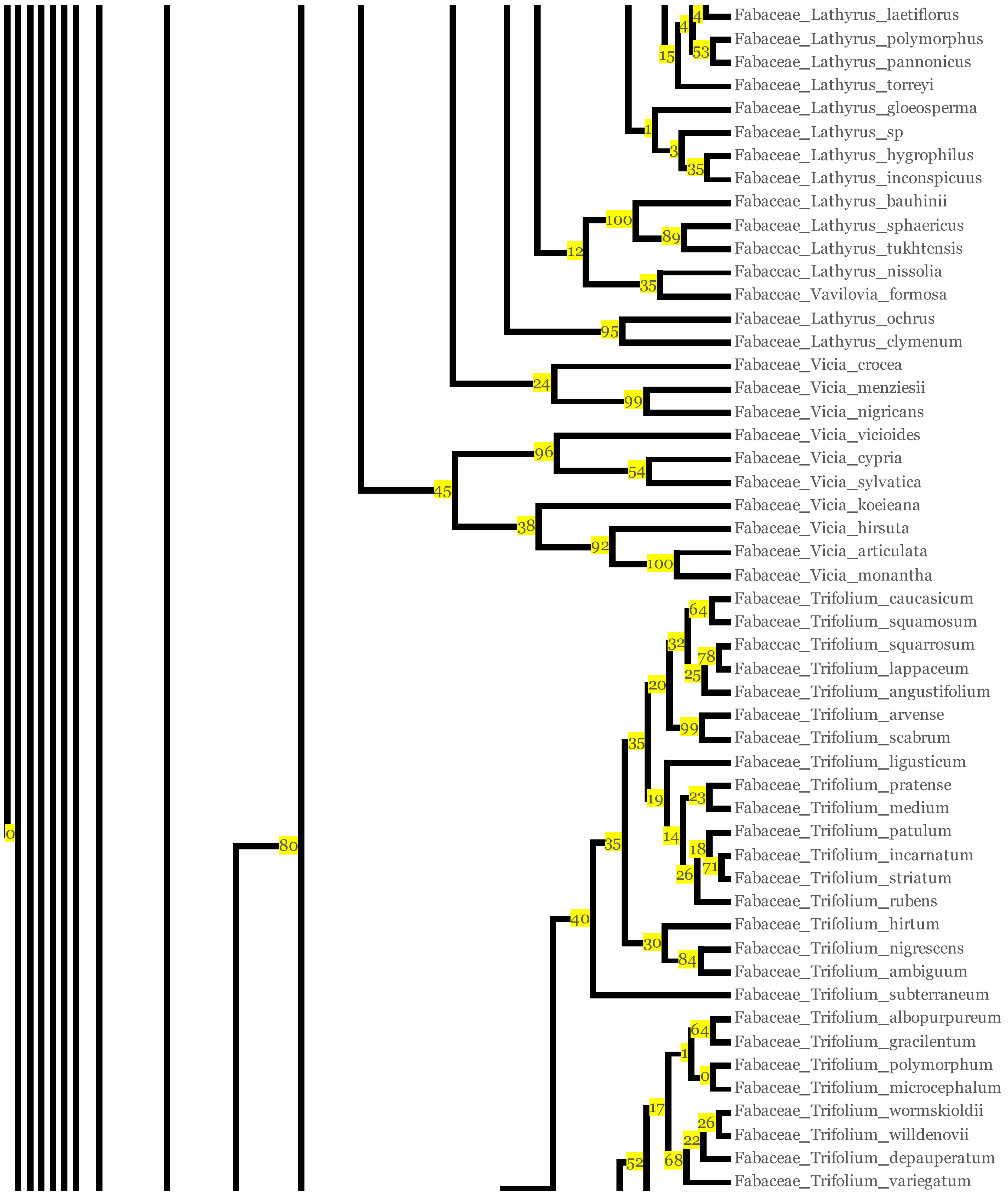
Fabaceae_Lathyrus_laetiflorus
Fabaceae_Lathyrus_polymorphus
Fabaceae_Lathyrus_pannonicus
Fabaceae_Lathyrus_torreyi
Fabaceae_Lathyrus_gloeosperma
Fabaceae_Lathyrus_sp
Fabaceae_Lathyrus_hygrophilus
Fabaceae_Lathyrus_inconspicuus
Fabaceae_Lathyrus_bauhinii
Fabaceae_Lathyrus_sphaericus
Fabaceae_Lathyrus_tukhtensis
Fabaceae_Lathyrus_nissolia
Fabaceae_Vavilovia_formosa
Fabaceae_Lathyrus_ochrus
Fabaceae_Lathyrus_clymenum
Fabaceae_Vicia_crocea
Fabaceae_Vicia_menziesii
Fabaceae_Vicia_nigricans
Fabaceae_Vicia_vicioides
Fabaceae_Vicia_cypria
Fabaceae_Vicia_sylvatica
Fabaceae_Vicia_koeieana
Fabaceae_Vicia_hirsuta
Fabaceae_Vicia_articulata
Fabaceae_Vicia_monantha
Fabaceae_Trifolium_caucasicum
Fabaceae_Trifolium_squamosum
Fabaceae_Trifolium_squarrosum
Fabaceae_Trifolium_lappaceum
Fabaceae_Trifolium_angustifolium
Fabaceae_Trifolium_arvense
Fabaceae_Trifolium_scabrum
Fabaceae_Trifolium_ligusticum
Fabaceae_Trifolium_pratense
Fabaceae_Trifolium_medium
Fabaceae_Trifolium_patulum
Fabaceae_Trifolium_incarnatum
Fabaceae_Trifolium_striatum
Fabaceae_Trifolium_rubens
Fabaceae_Trifolium_hirtum
Fabaceae_Trifolium_nigrescens
Fabaceae_Trifolium_ambiguum
Fabaceae_Trifolium_subterraneum
Fabaceae_Trifolium_albopurpureum
Fabaceae_Trifolium_gracilentum
Fabaceae_Trifolium_polymorphum
Fabaceae_Trifolium_microcephalum
Fabaceae_Trifolium_wormskioldii
Fabaceae_Trifolium_willdenovii
Fabaceae_Trifolium_depauperatum
Fabaceae_Trifolium_variegatum
4
4
53
15
1
3
35
100
89
12
35
95
24
99
96
54
45
38
92
100
64
32
78
25
20
99
35
19
23
14
18
71
26
35
80
0
40
30
84
64
1
0
17
26
22
68
52

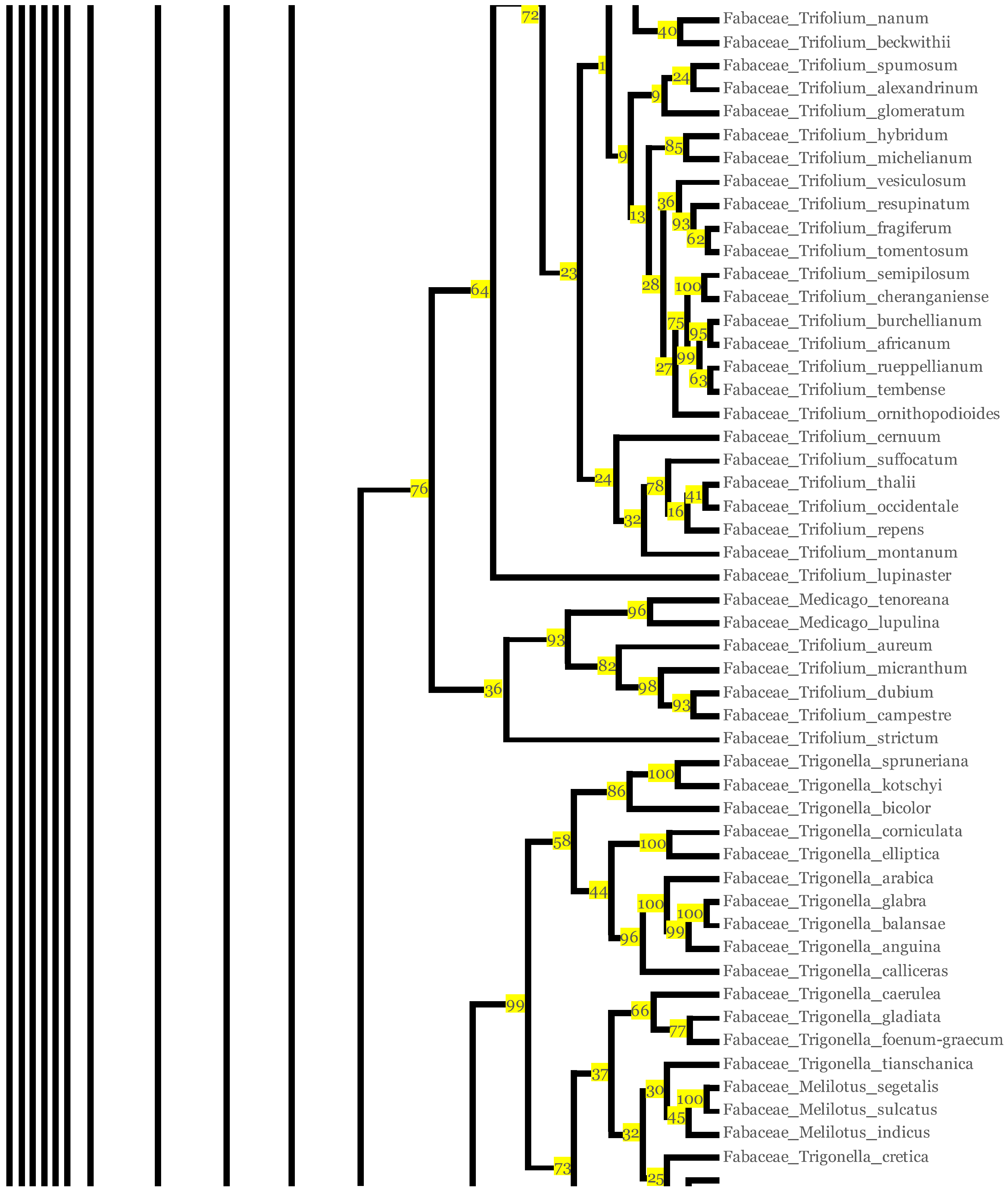
Fabaceae_Trifolium_nanum
Fabaceae_Trifolium_beckwithii
Fabaceae_Trifolium_spumosum
Fabaceae_Trifolium_alexandrinum
Fabaceae_Trifolium_glomeratum
Fabaceae_Trifolium_hybridum
Fabaceae_Trifolium_michelianum
Fabaceae_Trifolium_vesiculosum
Fabaceae_Trifolium_resupinatum
Fabaceae_Trifolium_fragiferum
Fabaceae_Trifolium_tomentosum
Fabaceae_Trifolium_semipilosum
Fabaceae_Trifolium_cheranganiense
Fabaceae_Trifolium_burchellianum
Fabaceae_Trifolium_africanum
Fabaceae_Trifolium_rueppellianum
Fabaceae_Trifolium_tembense
Fabaceae_Trifolium_ornithopodioides
Fabaceae_Trifolium_cernuum
Fabaceae_Trifolium_suffocatum
Fabaceae_Trifolium_thalii
Fabaceae_Trifolium_occidentale
Fabaceae_Trifolium_repens
Fabaceae_Trifolium_montanum
Fabaceae_Trifolium_lupinaster
Fabaceae_Medicago_tenoreana
Fabaceae_Medicago_lupulina
Fabaceae_Trifolium_aureum
Fabaceae_Trifolium_micranthum
Fabaceae_Trifolium_dubium
Fabaceae_Trifolium_campestre
Fabaceae_Trifolium_strictum
Fabaceae_Trigonella_spruneriana
Fabaceae_Trigonella_kotschyi
Fabaceae_Trigonella_bicolor
Fabaceae_Trigonella_corniculata
Fabaceae_Trigonella_elliptica
Fabaceae_Trigonella_arabica
Fabaceae_Trigonella_glabra
Fabaceae_Trigonella_balansae
Fabaceae_Trigonella_anguina
Fabaceae_Trigonella_calliceras
Fabaceae_Trigonella_caerulea
Fabaceae_Trigonella_gladiata
Fabaceae_Trigonella_foenum-graecum
Fabaceae_Trigonella_tianschanica
Fabaceae_Melilotus_segetalis
Fabaceae_Melilotus_sulcatus
Fabaceae_Melilotus_indicus
Fabaceae_Trigonella_cretica

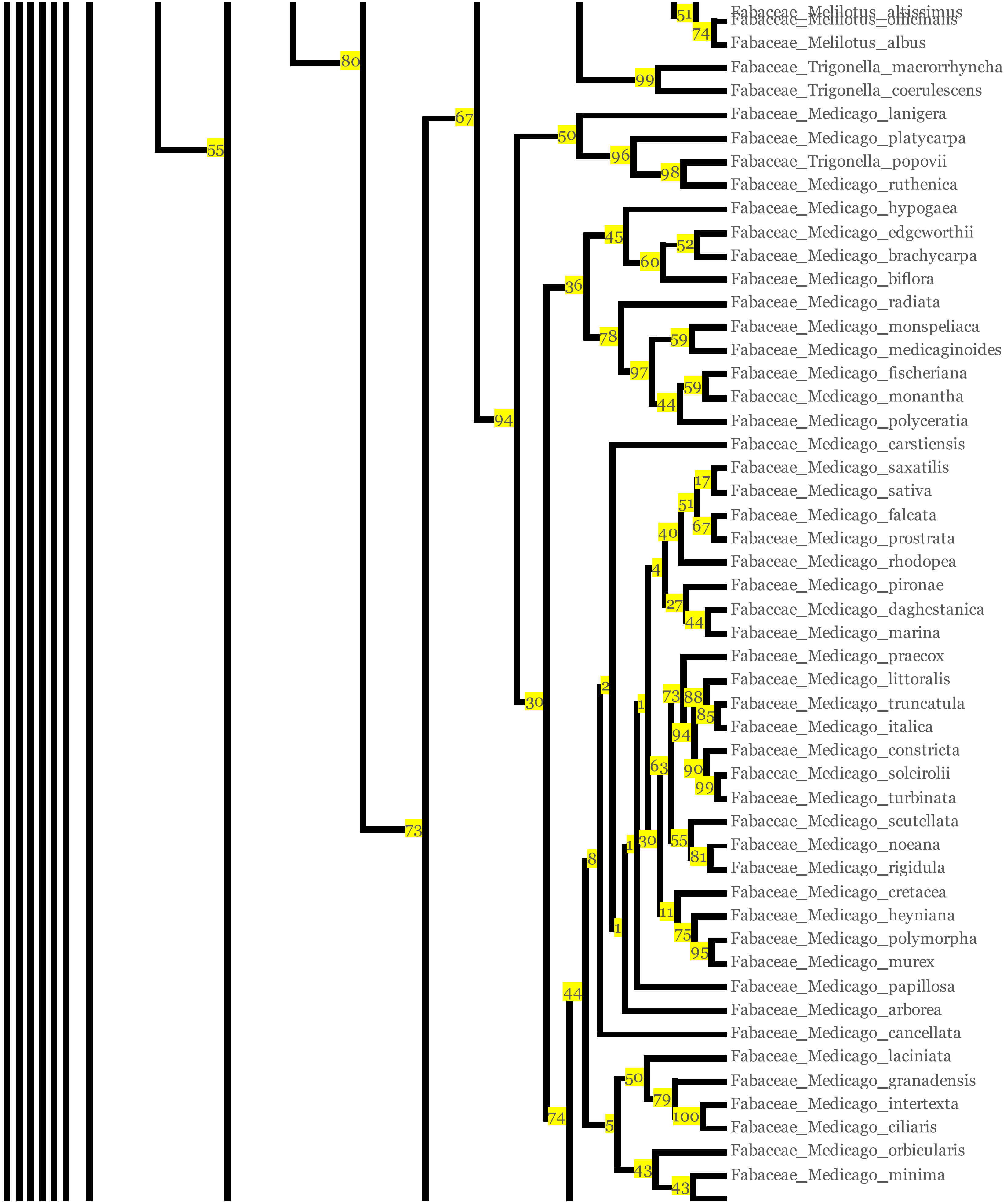
Fabaceae_Melilotus_altissimus
Fabaceae_Melilotus_officinalis
Fabaceae_Melilotus_albus
Fabaceae_Trigonella_macrorrhyncha
Fabaceae_Trigonella_coerulescens
Fabaceae_Medicago_lanigera
Fabaceae_Medicago_platycarpa
Fabaceae_Trigonella_popovii
Fabaceae_Medicago_ruthenica
Fabaceae_Medicago_hypogaea
Fabaceae_Medicago_edgeworthii
Fabaceae_Medicago_brachycarpa
Fabaceae_Medicago_biflora
Fabaceae_Medicago_radiata
Fabaceae_Medicago_monspeliaca
Fabaceae_Medicago_medicaginoides
Fabaceae_Medicago_fischeriana
Fabaceae_Medicago_monantha
Fabaceae_Medicago_polyceratia
Fabaceae_Medicago_carstiensis
Fabaceae_Medicago_saxatilis
Fabaceae_Medicago_sativa
Fabaceae_Medicago_falcata
Fabaceae_Medicago_prostrata
Fabaceae_Medicago_rhodopea
Fabaceae_Medicago_pironae
Fabaceae_Medicago_daghestanica
Fabaceae_Medicago_marina
Fabaceae_Medicago_praecox
Fabaceae_Medicago_littoralis
Fabaceae_Medicago_truncatula
Fabaceae_Medicago_italica
Fabaceae_Medicago_constricta
Fabaceae_Medicago_soleirolii
Fabaceae_Medicago_turbinata
Fabaceae_Medicago_scutellata
Fabaceae_Medicago_noeana
Fabaceae_Medicago_rigidula
Fabaceae_Medicago_cretacea
Fabaceae_Medicago_heyniana
Fabaceae_Medicago_polymorpha
Fabaceae_Medicago_murex
Fabaceae_Medicago_papillosa
Fabaceae_Medicago_arborea
Fabaceae_Medicago_cancellata
Fabaceae_Medicago_laciniata
Fabaceae_Medicago_granadensis
Fabaceae_Medicago_intertexta
Fabaceae_Medicago_ciliaris
Fabaceae_Medicago_orbicularis
Fabaceae_Medicago_minima

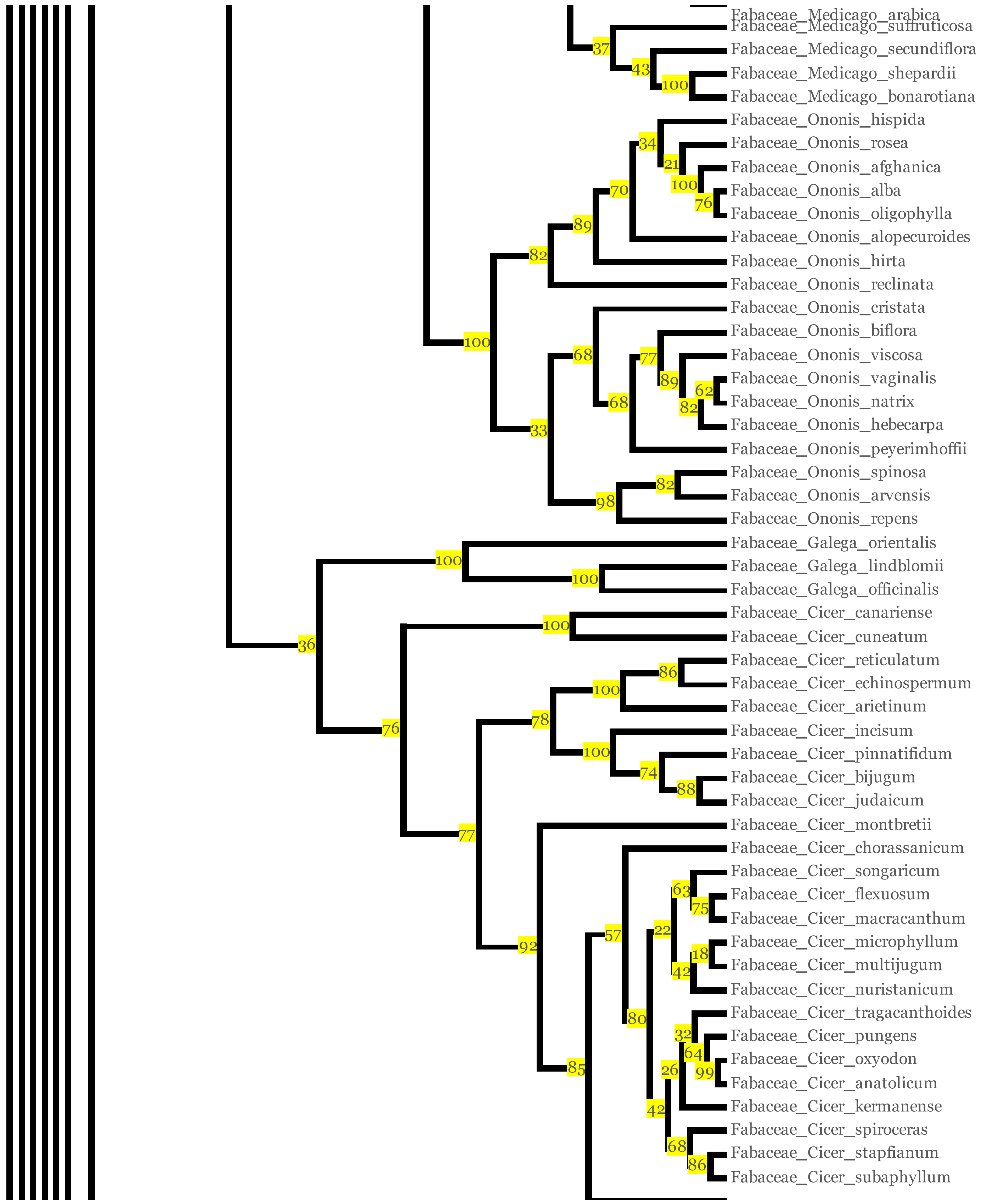
Fabaceae_Medicago_arabica
Fabaceae_Medicago_suffruticosa
Fabaceae_Medicago_secundiflora
Fabaceae_Medicago_shepardii
Fabaceae_Medicago_bonarotiana
Fabaceae_Ononis_hispida
Fabaceae_Ononis_rosea
Fabaceae_Ononis_afghanica
Fabaceae_Ononis_alba
Fabaceae_Ononis_oligophylla
Fabaceae_Ononis_alopecuroides
Fabaceae_Ononis_hirta
Fabaceae_Ononis_reclinata
Fabaceae_Ononis_cristata
Fabaceae_Ononis_biflora
Fabaceae_Ononis_viscosa
Fabaceae_Ononis_vaginalis
Fabaceae_Ononis_natrix
Fabaceae_Ononis_hebecarpa
Fabaceae_Ononis_peyerimhoffii
Fabaceae_Ononis_spinosa
Fabaceae_Ononis_arvensis
Fabaceae_Ononis_repens
Fabaceae_Galega_orientalis
Fabaceae_Galega_lindblomii
Fabaceae_Galega_officinalis
Fabaceae_Cicer_canariense
Fabaceae_Cicer_cuneatum
Fabaceae_Cicer_reticulatum
Fabaceae_Cicer_echinospermum
Fabaceae_Cicer_arietinum
Fabaceae_Cicer_incisum
Fabaceae_Cicer_pinnatifidum
Fabaceae_Cicer_bijugum
Fabaceae_Cicer_judaicum
Fabaceae_Cicer_montbretii
Fabaceae_Cicer_chorassanicum
Fabaceae_Cicer_songaricum
Fabaceae_Cicer_flexuosum
Fabaceae_Cicer_macracanthum
Fabaceae_Cicer_microphyllum
Fabaceae_Cicer_multijugum
Fabaceae_Cicer_nuristanicum
Fabaceae_Cicer_tragacanthoides
Fabaceae_Cicer_pungens
Fabaceae_Cicer_oxyodon
Fabaceae_Cicer_anatolicum
Fabaceae_Cicer_kermanense
Fabaceae_Cicer_spiroceras
Fabaceae_Cicer_stapfianum
Fabaceae_Cicer_subaphyllum

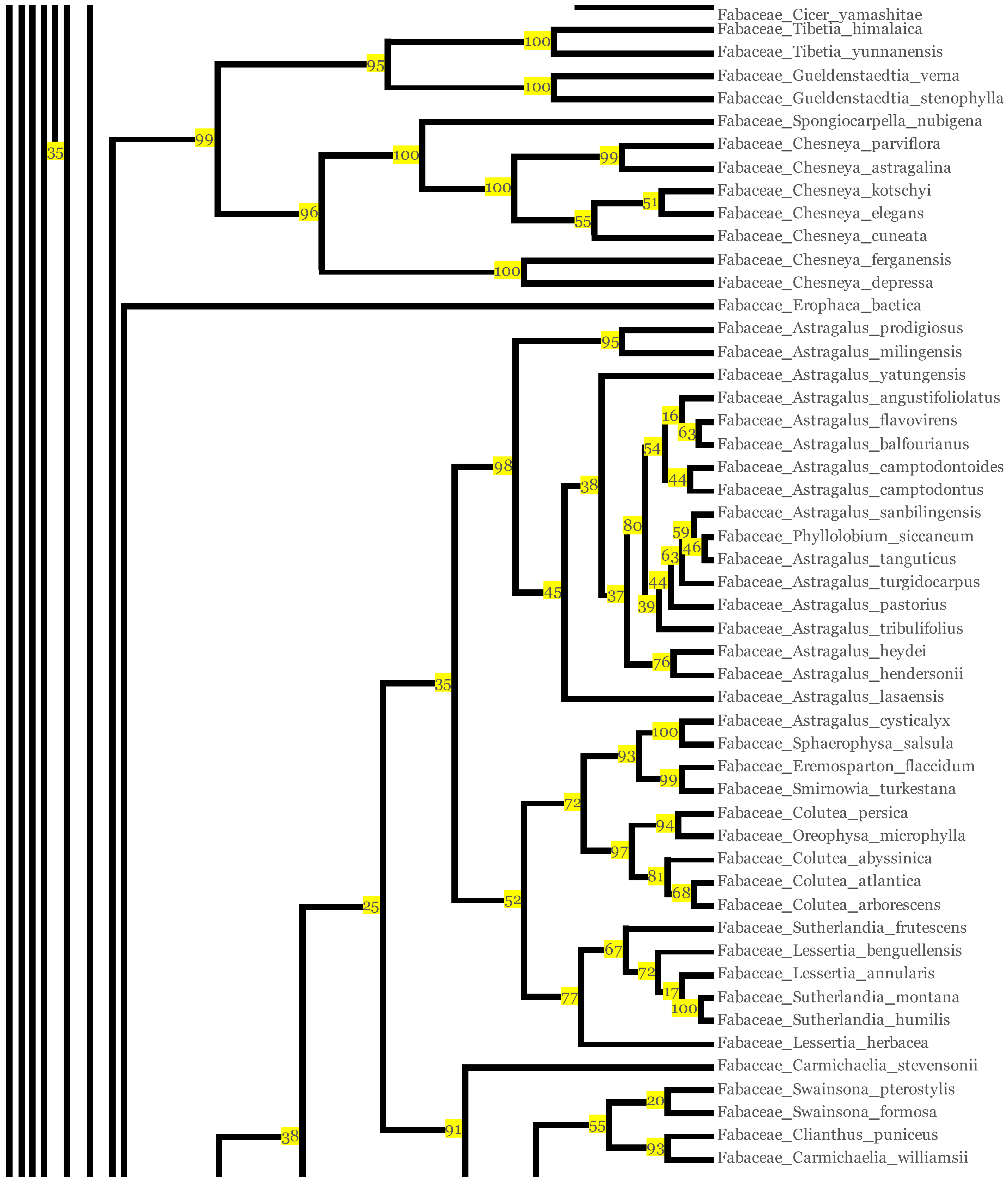
Fabaceae_Cicer_yamashitae
Fabaceae_Tibetia_himalaica
Fabaceae_Tibetia_yunnanensis
Fabaceae_Gueldenstaedtia_verna
Fabaceae_Gueldenstaedtia_stenophylla
Fabaceae_Spongiocarpella_nubigena
Fabaceae_Chesneya_parviflora
Fabaceae_Chesneya_astragalina
Fabaceae_Chesneya_kotschyi
Fabaceae_Chesneya_elegans
Fabaceae_Chesneya_cuneata
Fabaceae_Chesneya_ferganensis
Fabaceae_Chesneya_depressa
Fabaceae_Erophaca_baetica
Fabaceae_Astragalus_prodigiosus
Fabaceae_Astragalus_milingensis
Fabaceae_Astragalus_yatungensis
Fabaceae_Astragalus_angustifoliolatus
Fabaceae_Astragalus_flavovirens
Fabaceae_Astragalus_balfourianus
Fabaceae_Astragalus_camptodontoides
Fabaceae_Astragalus_camptodontus
Fabaceae_Astragalus_sanbilingensis
Fabaceae_Phyllolobium_siccaneum
Fabaceae_Astragalus_tanguticus
Fabaceae_Astragalus_turgidocarpus
Fabaceae_Astragalus_pastorius
Fabaceae_Astragalus_tribulifolius
Fabaceae_Astragalus_heydei
Fabaceae_Astragalus_hendersonii
Fabaceae_Astragalus_lasaensis
Fabaceae_Astragalus_cysticalyx
Fabaceae_Sphaerophysa_salsula
Fabaceae_Eremosparton_flaccidum
Fabaceae_Smirnowia_turkestana
Fabaceae_Colutea_persica
Fabaceae_Oreophysa_microphylla
Fabaceae_Colutea_abyssinica
Fabaceae_Colutea_atlantica
Fabaceae_Colutea_arborescens
Fabaceae_Sutherlandia_frutescens
Fabaceae_Lessertia_benguellensis
Fabaceae_Lessertia_annularis
Fabaceae_Sutherlandia_montana
Fabaceae_Sutherlandia_humilis
Fabaceae_Lessertia_herbacea
Fabaceae_Carmichaelia_stevensonii
Fabaceae_Swainsona_pterostylis
Fabaceae_Swainsona_formosa
Fabaceae_Clianthus_puniceus
Fabaceae_Carmichaelia_williamsii

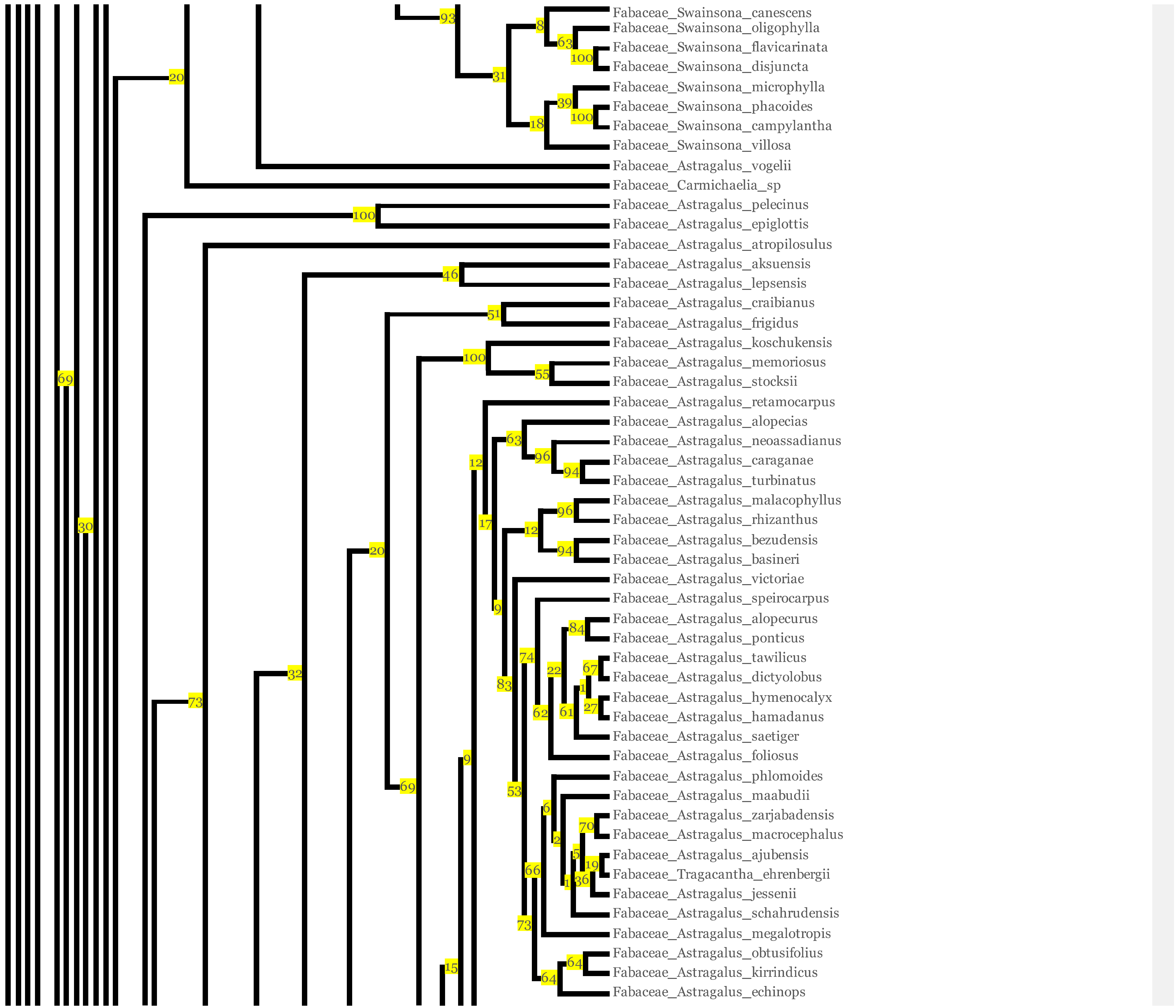

Fabaceae_Swainsona_canescens
Fabaceae_Swainsona_oligophylla
Fabaceae_Swainsona_flavicarinata
Fabaceae_Swainsona_disjuncta
Fabaceae_Swainsona_microphylla
Fabaceae_Swainsona_phacoides
Fabaceae_Swainsona_campylantha
Fabaceae_Swainsona_villosa
Fabaceae_Astragalus_vogelii
Fabaceae_Carmichaelia_sp
Fabaceae_Astragalus_pelecinus
Fabaceae_Astragalus_epiglottis
Fabaceae_Astragalus_atropilosulus
Fabaceae_Astragalus_aksuensis
Fabaceae_Astragalus_lepsensis
Fabaceae_Astragalus_craibianus
Fabaceae_Astragalus_frigidus
Fabaceae_Astragalus_koschukensis
Fabaceae_Astragalus_memoriosus
Fabaceae_Astragalus_stocksii
Fabaceae_Astragalus_retamocarpus
Fabaceae_Astragalus_alopecias
Fabaceae_Astragalus_neoassadianus
Fabaceae_Astragalus_caraganae
Fabaceae_Astragalus_turbinatus
Fabaceae_Astragalus_malacophyllus
Fabaceae_Astragalus_rhizanthus
Fabaceae_Astragalus_bezudensis
Fabaceae_Astragalus_basineri
Fabaceae_Astragalus_victoriae
Fabaceae_Astragalus_speirocarpus
Fabaceae_Astragalus_alopecurus
Fabaceae_Astragalus_ponticus
Fabaceae_Astragalus_tawilicus
Fabaceae_Astragalus_dictyolobus
Fabaceae_Astragalus_hymenocalyx
Fabaceae_Astragalus_hamadanus
Fabaceae_Astragalus_saetiger
Fabaceae_Astragalus_foliosus
Fabaceae_Astragalus_phlomoides
Fabaceae_Astragalus_maabudii
Fabaceae_Astragalus_zarjabadensis
Fabaceae_Astragalus_macrocephalus
Fabaceae_Astragalus_ajubensis
Fabaceae_Tragacantha_ehrenbergii
Fabaceae_Astragalus_jessenii
Fabaceae_Astragalus_schahrudensis
Fabaceae_Astragalus_megalotropis
Fabaceae_Astragalus_obtusifolius
Fabaceae_Astragalus_kirrindicus
Fabaceae_Astragalus_echinops
93
8
63
100
31
39
100
18
20
100
46
51
100
55
69
30
63
96
12
94
17
96
12
94
9
20
83
74
22
67
1
62
61
27
32
73
9
69
53
6
2
70
5
19
1
36
66
73
15
64
64

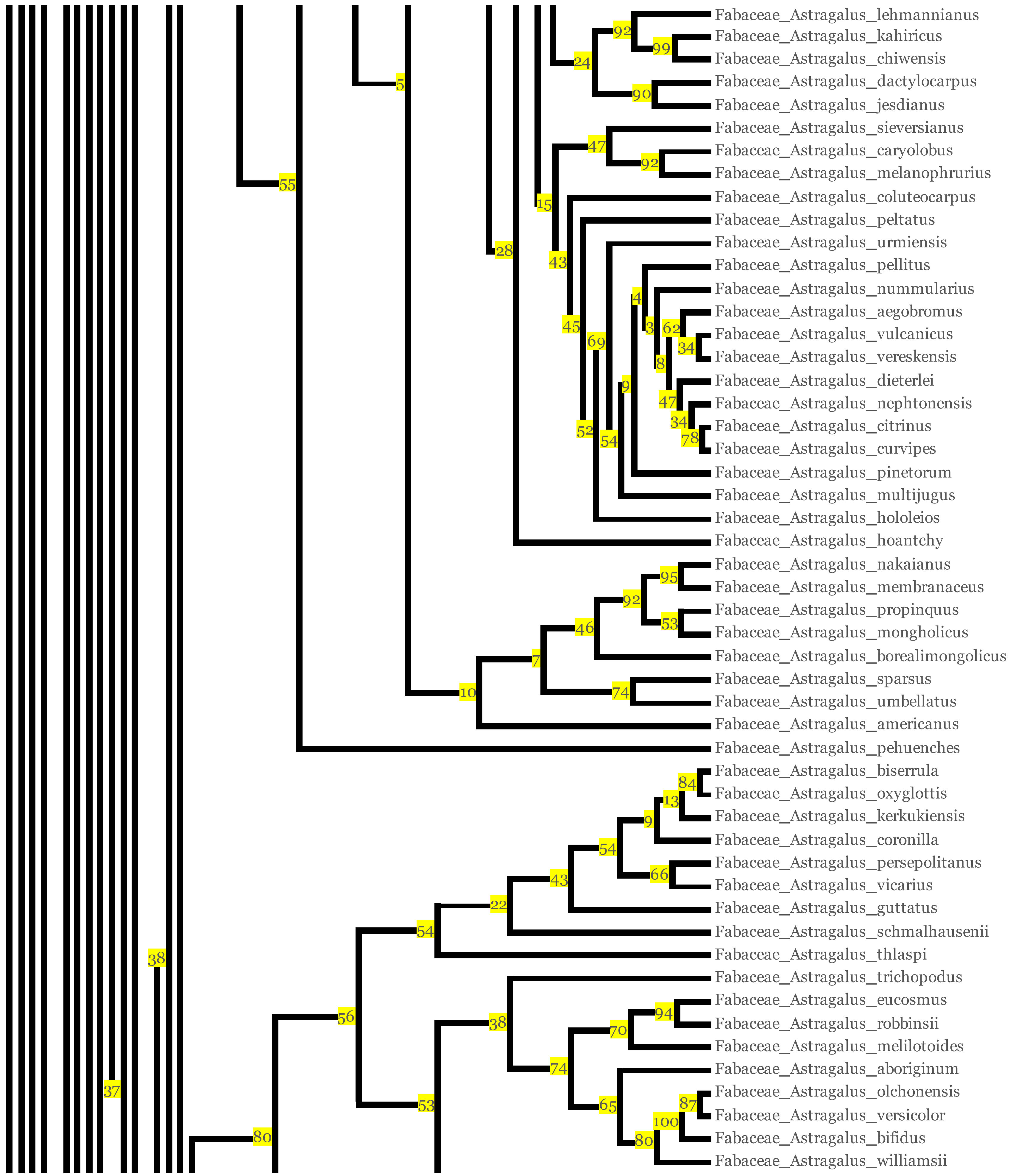
Fabaceae_Astragalus_lehmannianus
Fabaceae_Astragalus_kahiricus
Fabaceae_Astragalus_chiwensis
Fabaceae_Astragalus_dactylocarpus
Fabaceae_Astragalus_jesdianus
Fabaceae_Astragalus_sieversianus
Fabaceae_Astragalus_caryolobus
Fabaceae_Astragalus_melanophrurius
Fabaceae_Astragalus_coluteocarpus
Fabaceae_Astragalus_peltatus
Fabaceae_Astragalus_urmiensis
Fabaceae_Astragalus_pellitus
Fabaceae_Astragalus_nummularius
Fabaceae_Astragalus_aegobromus
Fabaceae_Astragalus_vulcanicus
Fabaceae_Astragalus_vereskensis
Fabaceae_Astragalus_dieterlei
Fabaceae_Astragalus_nephtonensis
Fabaceae_Astragalus_citrinus
Fabaceae_Astragalus_curvipes
Fabaceae_Astragalus_pinetorum
Fabaceae_Astragalus_multijugus
Fabaceae_Astragalus_hololeios
Fabaceae_Astragalus_hoantchy
Fabaceae_Astragalus_nakaianus
Fabaceae_Astragalus_membranaceus
Fabaceae_Astragalus_propinquus
Fabaceae_Astragalus_mongholicus
Fabaceae_Astragalus_borealimongolicus
Fabaceae_Astragalus_sparsus
Fabaceae_Astragalus_umbellatus
Fabaceae_Astragalus_americanus
Fabaceae_Astragalus_pehuenches
Fabaceae_Astragalus_biserrula
Fabaceae_Astragalus_oxyglottis
Fabaceae_Astragalus_kerkukiensis
Fabaceae_Astragalus_coronilla
Fabaceae_Astragalus_persepolitanus
Fabaceae_Astragalus_vicarius
Fabaceae_Astragalus_guttatus
Fabaceae_Astragalus_schmalhausenii
Fabaceae_Astragalus_thlaspi
Fabaceae_Astragalus_trichopodus
Fabaceae_Astragalus_eucosmus
Fabaceae_Astragalus_robbinsii
Fabaceae_Astragalus_melilotoides
Fabaceae_Astragalus_aboriginum
Fabaceae_Astragalus_olchonensis
Fabaceae_Astragalus_versicolor
Fabaceae_Astragalus_bifidus
Fabaceae_Astragalus_williamsii

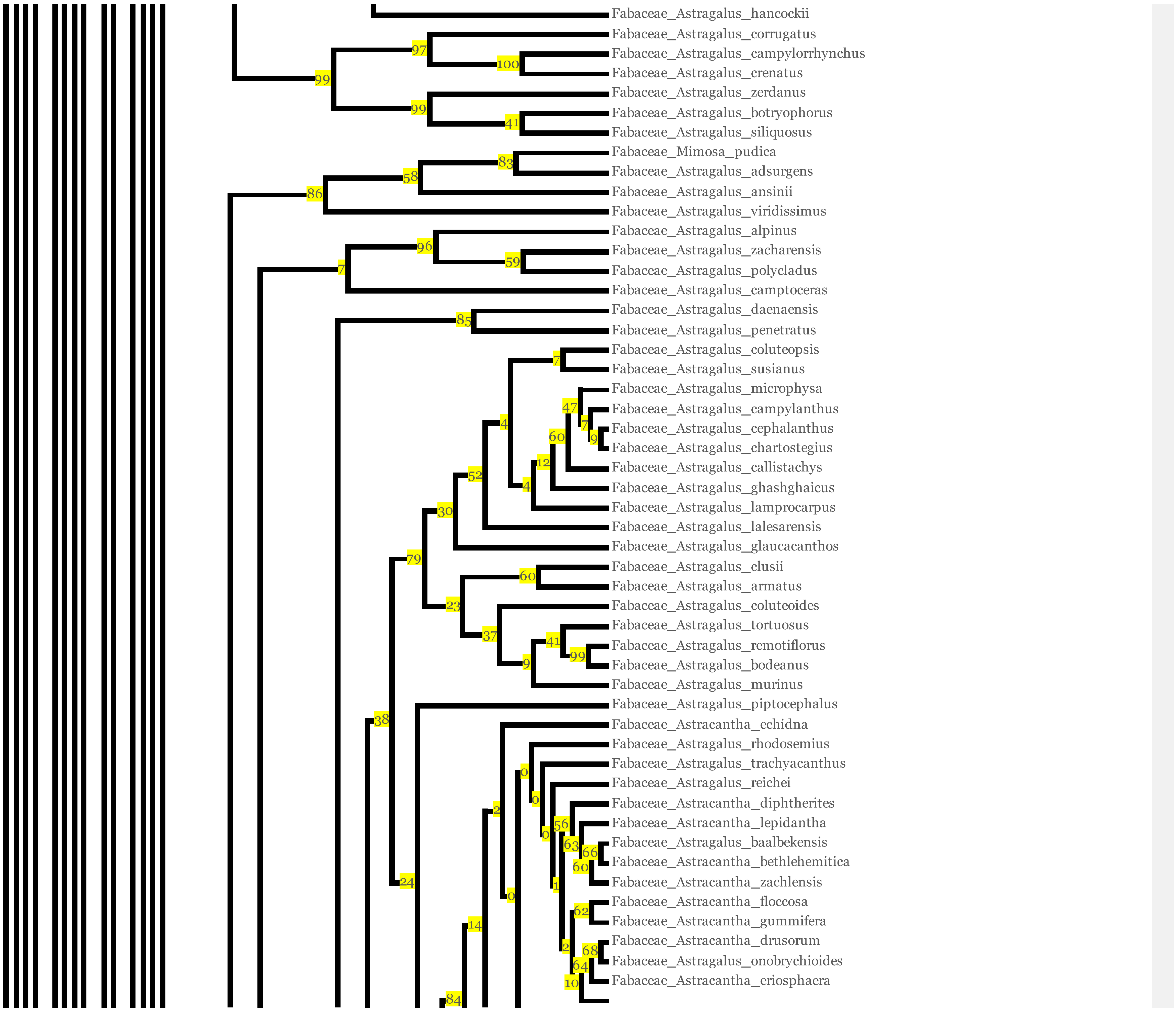
Fabaceae_Astragalus_hancockii
Fabaceae_Astragalus_corrugatus
Fabaceae_Astragalus_campylorrhynchus
Fabaceae_Astragalus_crenatus
Fabaceae_Astragalus_zerdanus
Fabaceae_Astragalus_botryophorus
Fabaceae_Astragalus_siliquosus
Fabaceae_Mimosa_pudica
Fabaceae_Astragalus_adsurgens
Fabaceae_Astragalus_ansinii
Fabaceae_Astragalus_viridissimus
Fabaceae_Astragalus_alpinus
Fabaceae_Astragalus_zacharensis
Fabaceae_Astragalus_polycladus
Fabaceae_Astragalus_camptoceras
Fabaceae_Astragalus_daenaensis
Fabaceae_Astragalus_penetratus
Fabaceae_Astragalus_coluteopsis
Fabaceae_Astragalus_susianus
Fabaceae_Astragalus_microphysa
Fabaceae_Astragalus_campylanthus
Fabaceae_Astragalus_cephalanthus
Fabaceae_Astragalus_chartostegius
Fabaceae_Astragalus_callistachys
Fabaceae_Astragalus_ghashghaicus
Fabaceae_Astragalus_lamprocarpus
Fabaceae_Astragalus_lalesarensis
Fabaceae_Astragalus_glaucacanthos
Fabaceae_Astragalus_clusii
Fabaceae_Astragalus_armatus
Fabaceae_Astragalus_coluteoides
Fabaceae_Astragalus_tortuosus
Fabaceae_Astragalus_remotiflorus
Fabaceae_Astragalus_bodeanus
Fabaceae_Astragalus_murinus
Fabaceae_Astragalus_piptocephalus
Fabaceae_Astracantha_echidna
Fabaceae_Astragalus_rhodosemius
Fabaceae_Astragalus_trachyacanthus
Fabaceae_Astragalus_reichei
Fabaceae_Astracantha_diphtherites
Fabaceae_Astracantha_lepidantha
Fabaceae_Astragalus_baalbekensis
Fabaceae_Astracantha_bethlehemitica
Fabaceae_Astracantha_zachlensis
Fabaceae_Astracantha_floccosa
Fabaceae_Astracantha_gummifera
Fabaceae_Astracantha_drusorum
Fabaceae_Astragalus_onobrychioides
Fabaceae_Astracantha_eriosphaera
97
100
99
99
41
83
58
86
96
59
7
85
7
47
7
9
60
4
12
52
4
30
79
60
23
37
41
99
9
38
0
0
2
0
56
63
66
60
1
0
24
62
14
2
68
64
10
84

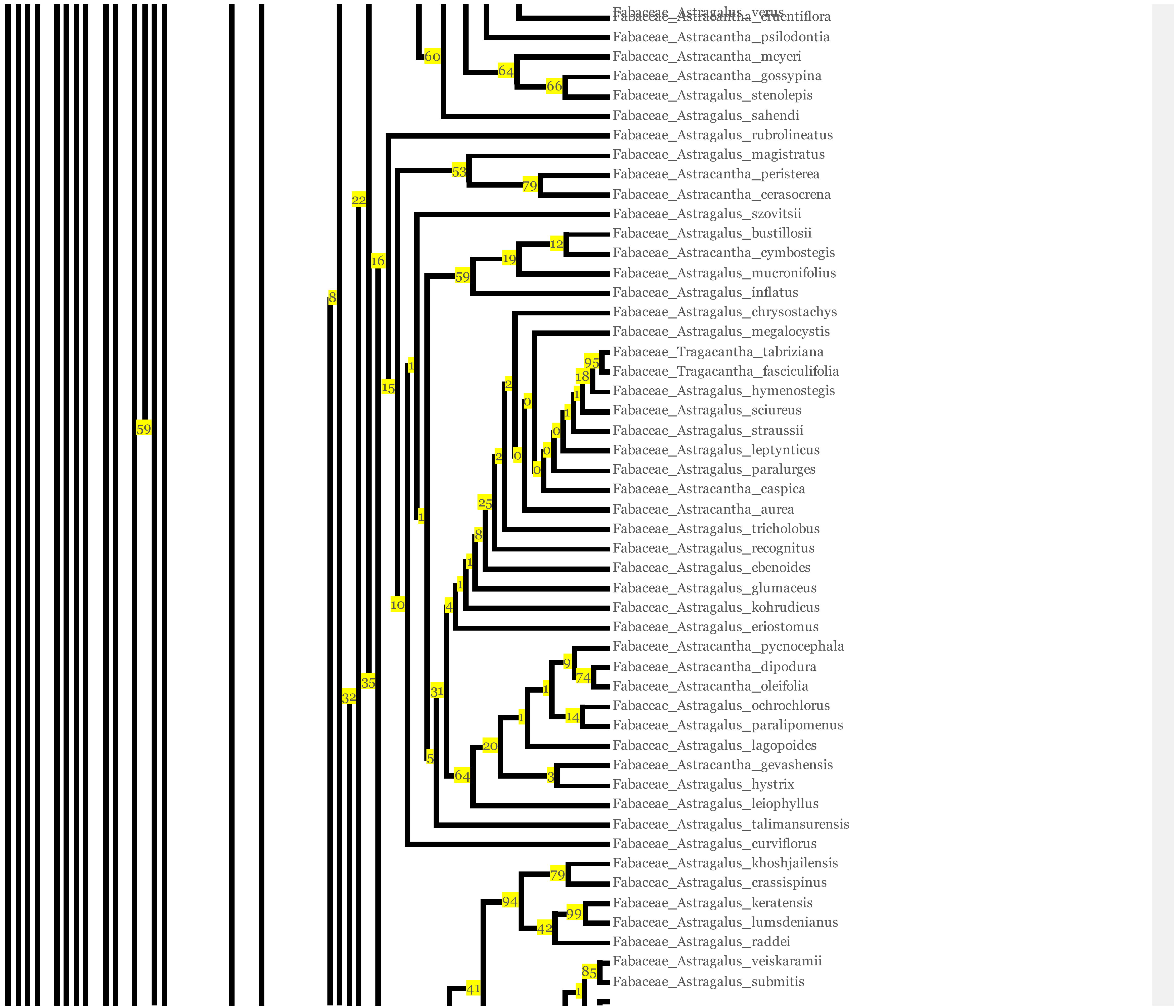

Fabaceae_Astragalus_verus
Fabaceae_Astracantha_cruentiflora
Fabaceae_Astracantha_psilodontia
Fabaceae_Astracantha_meyeri
Fabaceae_Astracantha_gossypina
Fabaceae_Astragalus_stenolepis
Fabaceae_Astragalus_sahendi
Fabaceae_Astragalus_rubrolineatus
Fabaceae_Astragalus_magistratus
Fabaceae_Astracantha_peristerea
Fabaceae_Astracantha_cerasocrena
Fabaceae_Astragalus_szovitsii
Fabaceae_Astragalus_bustillosii
Fabaceae_Astracantha_cymbostegis
Fabaceae_Astragalus_mucronifolius
Fabaceae_Astragalus_inflatus
Fabaceae_Astragalus_chrysostachys
Fabaceae_Astragalus_megalocystis
Fabaceae_Tragacantha_tabriziana
Fabaceae_Tragacantha_fasciculifolia
Fabaceae_Astragalus_hymenostegis
Fabaceae_Astragalus_sciureus
Fabaceae_Astragalus_straussii
Fabaceae_Astragalus_leptynticus
Fabaceae_Astragalus_paralurges
Fabaceae_Astracantha_caspica
Fabaceae_Astracantha_aurea
Fabaceae_Astragalus_tricholobus
Fabaceae_Astragalus_recognitus
Fabaceae_Astragalus_ebenoides
Fabaceae_Astragalus_glumaceus
Fabaceae_Astragalus_kohrudicus
Fabaceae_Astragalus_eriostomus
Fabaceae_Astracantha_pycnocephala
Fabaceae_Astracantha_dipodura
Fabaceae_Astracantha_oleifolia
Fabaceae_Astragalus_ochrochlorus
Fabaceae_Astragalus_paralipomenus
Fabaceae_Astragalus_lagopoides
Fabaceae_Astracantha_gevashensis
Fabaceae_Astragalus_hystrix
Fabaceae_Astragalus_leiophyllus
Fabaceae_Astragalus_talimansurensis
Fabaceae_Astragalus_curviflorus
Fabaceae_Astragalus_khoshjailensis
Fabaceae_Astragalus_crassispinus
Fabaceae_Astragalus_keratensis
Fabaceae_Astragalus_lumsdenianus
Fabaceae_Astragalus_raddei
Fabaceae_Astragalus_veiskaramii
Fabaceae_Astragalus_submitis

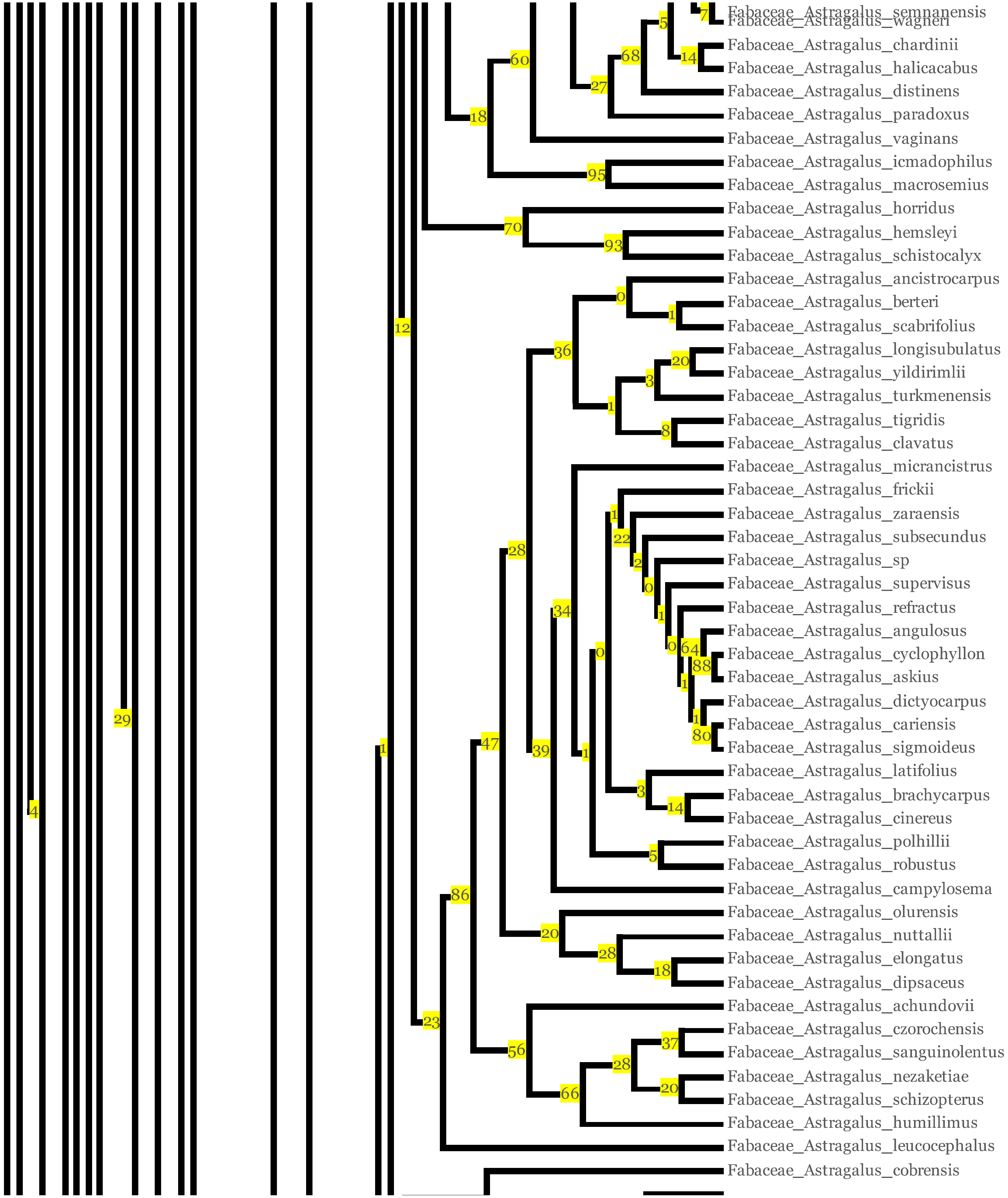
Fabaceae_Astragalus_semnanensis
Fabaceae_Astragalus_wagneri
Fabaceae_Astragalus_chardinii
Fabaceae_Astragalus_halicacabus
Fabaceae_Astragalus_distinens
Fabaceae_Astragalus_paradoxus
Fabaceae_Astragalus_vaginans
Fabaceae_Astragalus_icmadophilus
Fabaceae_Astragalus_macrosemius
Fabaceae_Astragalus_horridus
Fabaceae_Astragalus_hemsleyi
Fabaceae_Astragalus_schistocalyx
Fabaceae_Astragalus_ancistrocarpus
Fabaceae_Astragalus_berteri
Fabaceae_Astragalus_scabrifolius
Fabaceae_Astragalus_longisubulatus
Fabaceae_Astragalus_yildirimlii
Fabaceae_Astragalus_turkmenensis
Fabaceae_Astragalus_tigridis
Fabaceae_Astragalus_clavatus
Fabaceae_Astragalus_micrancistrus
Fabaceae_Astragalus_frickii
Fabaceae_Astragalus_zaraensis
Fabaceae_Astragalus_subsecundus
Fabaceae_Astragalus_sp
Fabaceae_Astragalus_supervisus
Fabaceae_Astragalus_refractus
Fabaceae_Astragalus_angulosus
Fabaceae_Astragalus_cyclophyllon
Fabaceae_Astragalus_askius
Fabaceae_Astragalus_dictyocarpus
Fabaceae_Astragalus_cariensis
Fabaceae_Astragalus_sigmoideus
Fabaceae_Astragalus_latifolius
Fabaceae_Astragalus_brachycarpus
Fabaceae_Astragalus_cinereus
Fabaceae_Astragalus_polhillii
Fabaceae_Astragalus_robustus
Fabaceae_Astragalus_campylosema
Fabaceae_Astragalus_olurensis
Fabaceae_Astragalus_nuttallii
Fabaceae_Astragalus_elongatus
Fabaceae_Astragalus_dipsaceus
Fabaceae_Astragalus_achundovii
Fabaceae_Astragalus_czorochensis
Fabaceae_Astragalus_sanguinolentus
Fabaceae_Astragalus_nezaketiae
Fabaceae_Astragalus_schizopterus
Fabaceae_Astragalus_humillimus
Fabaceae_Astragalus_leucocephalus
Fabaceae_Astragalus_cobrensis

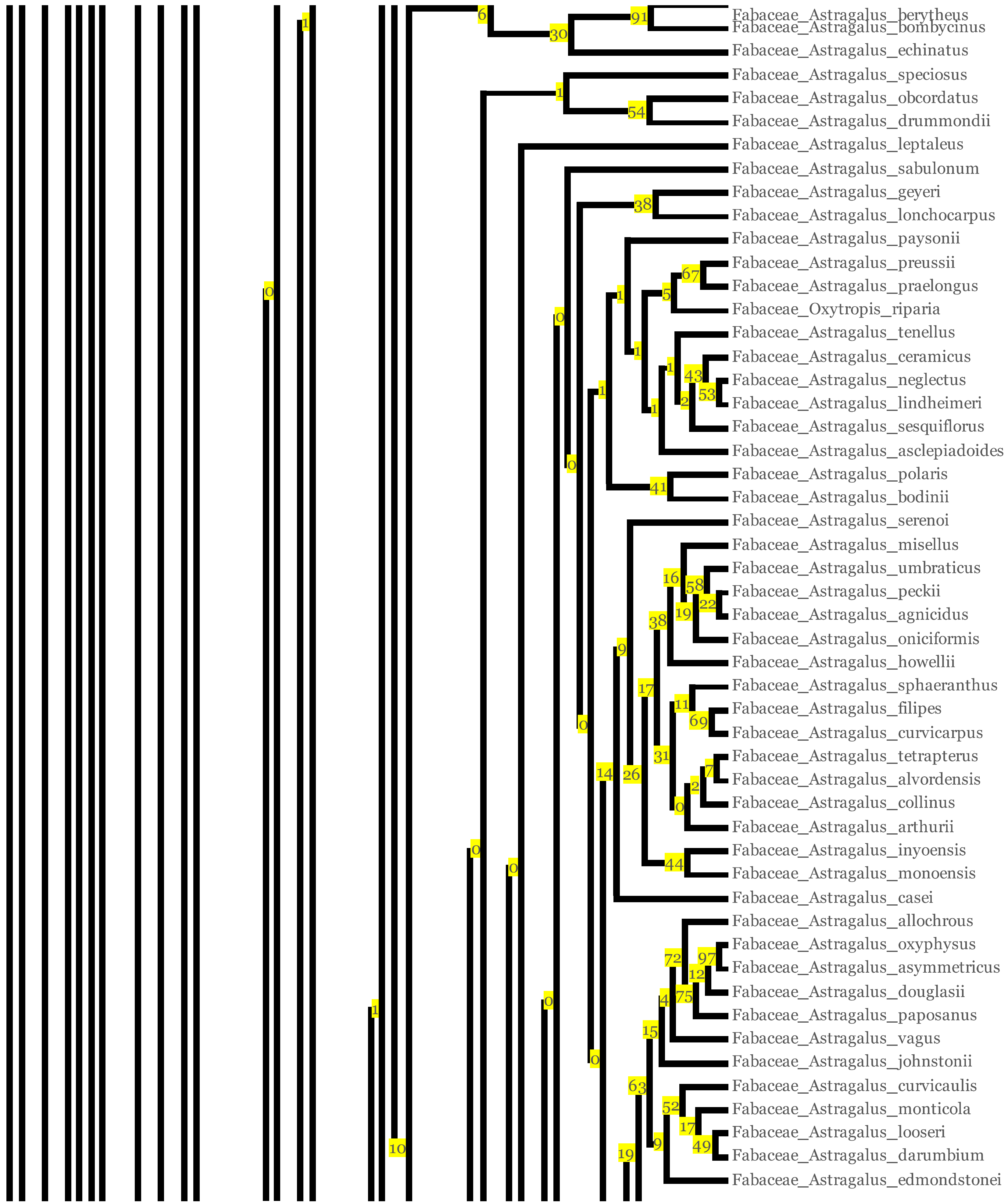
Fabaceae_Astragalus_berytheus
Fabaceae_Astragalus_bombycinus
Fabaceae_Astragalus_echinatus
Fabaceae_Astragalus_speciosus
Fabaceae_Astragalus_obcordatus
Fabaceae_Astragalus_drummondii
Fabaceae_Astragalus_leptaleus
Fabaceae_Astragalus_sabulonum
Fabaceae_Astragalus_geyeri
Fabaceae_Astragalus_lonchocarpus
Fabaceae_Astragalus_paysonii
Fabaceae_Astragalus_preussii
Fabaceae_Astragalus_praelongus
Fabaceae_Oxytropis_riparia
Fabaceae_Astragalus_tenellus
Fabaceae_Astragalus_ceramicus
Fabaceae_Astragalus_neglectus
Fabaceae_Astragalus_lindheimeri
Fabaceae_Astragalus_sesquiflorus
Fabaceae_Astragalus_asclepiadoides
Fabaceae_Astragalus_polaris
Fabaceae_Astragalus_bodinii
Fabaceae_Astragalus_serenoi
Fabaceae_Astragalus_misellus
Fabaceae_Astragalus_umbraticus
Fabaceae_Astragalus_peckii
Fabaceae_Astragalus_agnicidus
Fabaceae_Astragalus_oniciformis
Fabaceae_Astragalus_howellii
Fabaceae_Astragalus_sphaeranthus
Fabaceae_Astragalus_filipes
Fabaceae_Astragalus_curvicarpus
Fabaceae_Astragalus_tetrapterus
Fabaceae_Astragalus_alvordensis
Fabaceae_Astragalus_collinus
Fabaceae_Astragalus_arthurii
Fabaceae_Astragalus_inyoensis
Fabaceae_Astragalus_monoensis
Fabaceae_Astragalus_casei
Fabaceae_Astragalus_allochrous
Fabaceae_Astragalus_oxyphysus
Fabaceae_Astragalus_asymmetricus
Fabaceae_Astragalus_douglasii
Fabaceae_Astragalus_paposanus
Fabaceae_Astragalus_vagus
Fabaceae_Astragalus_johnstonii
Fabaceae_Astragalus_curvicaulis
Fabaceae_Astragalus_monticola
Fabaceae_Astragalus_looseri
Fabaceae_Astragalus_darumbium
Fabaceae_Astragalus_edmondstonei

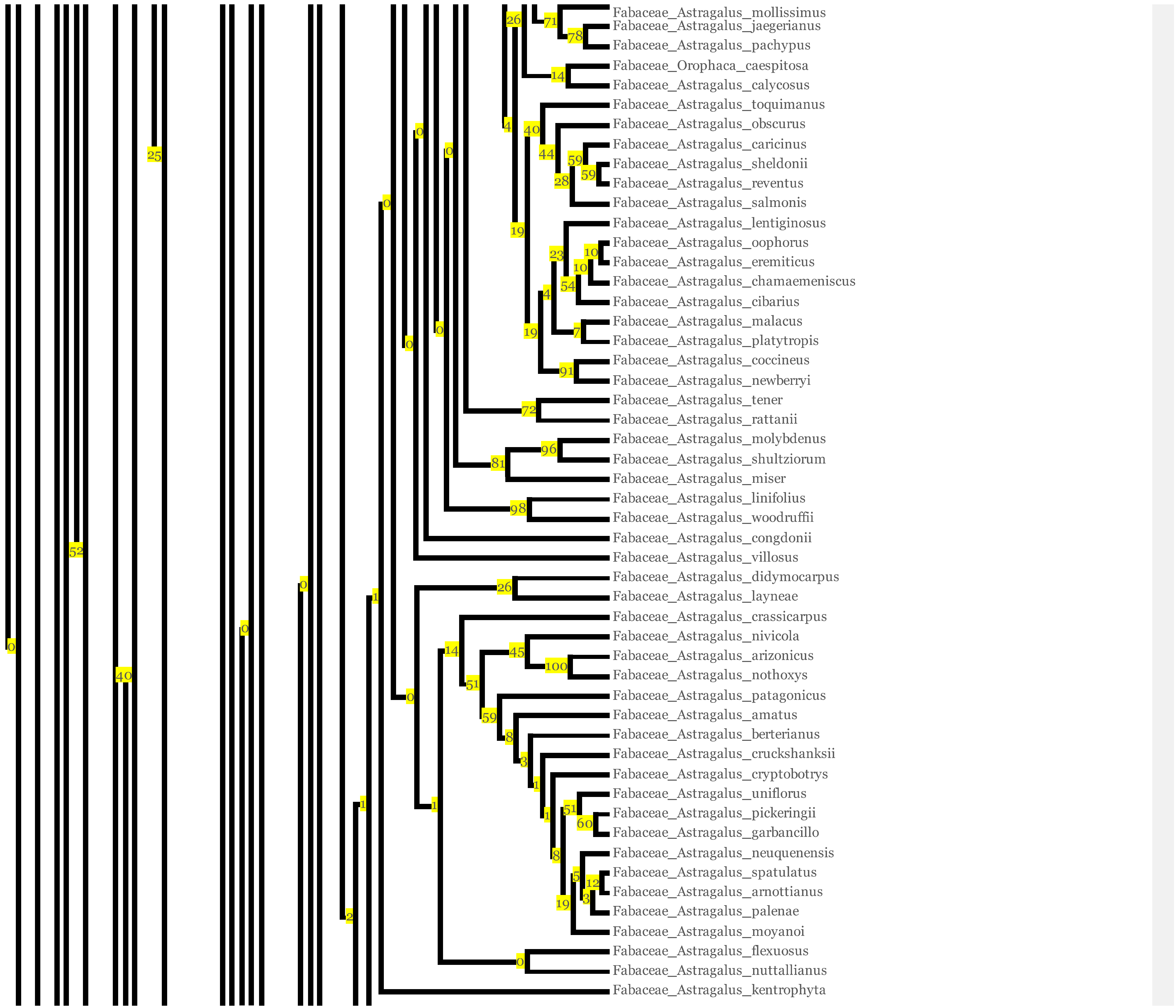
Fabaceae_Astragalus_mollissimus
Fabaceae_Astragalus_jaegerianus
Fabaceae_Astragalus_pachypus
Fabaceae_Orophaca_caespitosa
Fabaceae_Astragalus_calycosus
Fabaceae_Astragalus_toquimanus
Fabaceae_Astragalus_obscurus
Fabaceae_Astragalus_caricinus
Fabaceae_Astragalus_sheldonii
Fabaceae_Astragalus_reventus
Fabaceae_Astragalus_salmonis
Fabaceae_Astragalus_lentiginosus
Fabaceae_Astragalus_oophorus
Fabaceae_Astragalus_eremiticus
Fabaceae_Astragalus_chamaemeniscus
Fabaceae_Astragalus_cibarius
Fabaceae_Astragalus_malacus
Fabaceae_Astragalus_platytropis
Fabaceae_Astragalus_coccineus
Fabaceae_Astragalus_newberryi
Fabaceae_Astragalus_tener
Fabaceae_Astragalus_rattanii
Fabaceae_Astragalus_molybdenus
Fabaceae_Astragalus_shultziorum
Fabaceae_Astragalus_miser
Fabaceae_Astragalus_linifolius
Fabaceae_Astragalus_woodruffii
Fabaceae_Astragalus_congdonii
Fabaceae_Astragalus_villosus
Fabaceae_Astragalus_didymocarpus
Fabaceae_Astragalus_layneae
Fabaceae_Astragalus_crassicarpus
Fabaceae_Astragalus_nivicola
Fabaceae_Astragalus_arizonicus
Fabaceae_Astragalus_nothoxys
Fabaceae_Astragalus_patagonicus
Fabaceae_Astragalus_amatus
Fabaceae_Astragalus_bertierianus
Fabaceae_Astragalus_cruckshanksii
Fabaceae_Astragalus_cryptobotrys
Fabaceae_Astragalus_uniflorus
Fabaceae_Astragalus_pickeringii
Fabaceae_Astragalus_garbancillo
Fabaceae_Astragalus_neuquenensis
Fabaceae_Astragalus_spatulatus
Fabaceae_Astragalus_arnottianus
Fabaceae_Astragalus_palenae
Fabaceae_Astragalus_moyanoi
Fabaceae_Astragalus_flexuosus
Fabaceae_Astragalus_nuttallianus
Fabaceae_Astragalus_kentrophyta

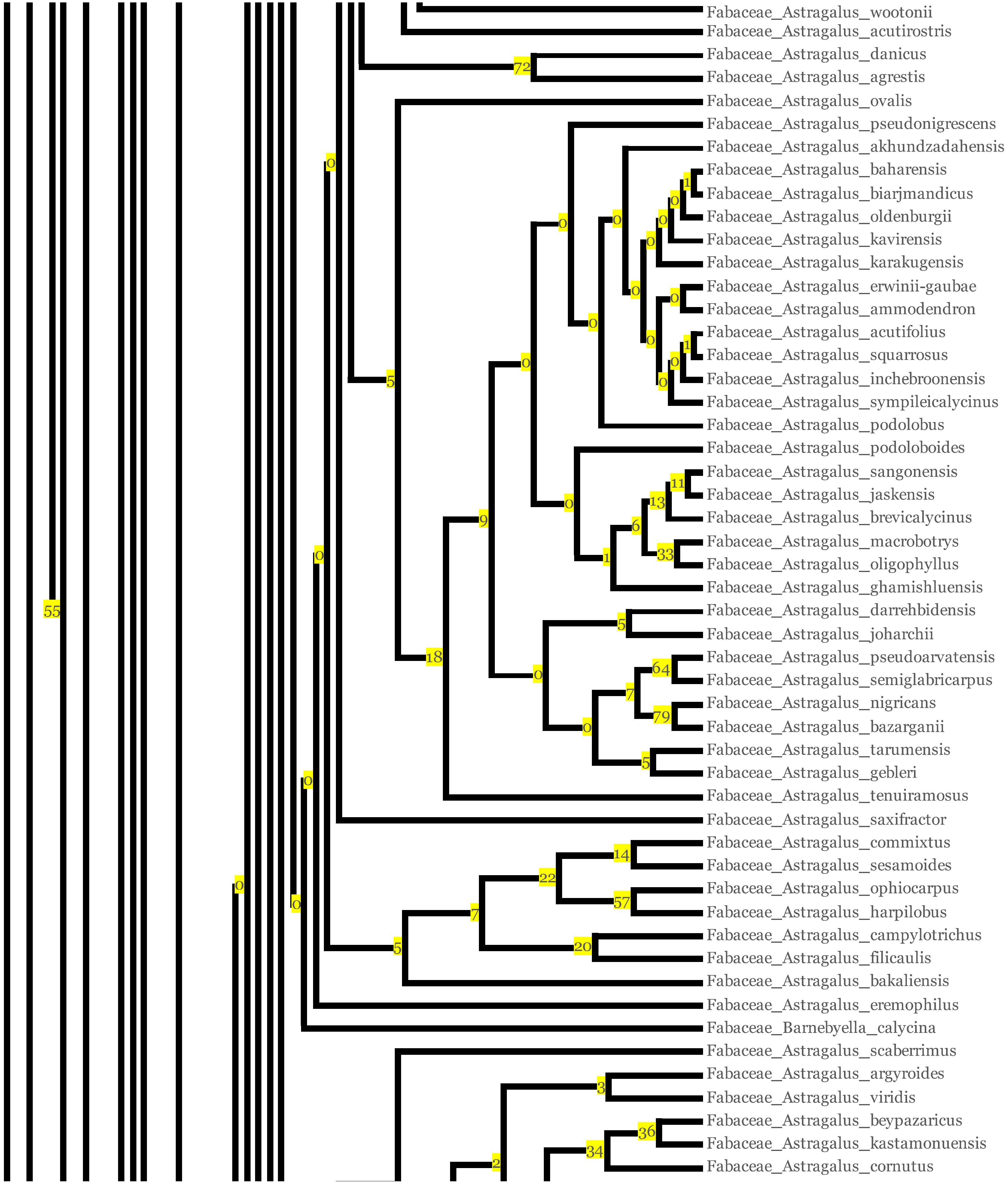
Fabaceae_Astragalus_wootonii
Fabaceae_Astragalus_acutirostris
Fabaceae_Astragalus_danicus
Fabaceae_Astragalus_agrestis
Fabaceae_Astragalus_ovalis
Fabaceae_Astragalus_pseudonigrescens
Fabaceae_Astragalus_akhundzadahensis
Fabaceae_Astragalus_baharensis
Fabaceae_Astragalus_biarjmandicus
Fabaceae_Astragalus_oldenburgii
Fabaceae_Astragalus_kavirensis
Fabaceae_Astragalus_karakugensis
Fabaceae_Astragalus_erwinii-gaubae
Fabaceae_Astragalus_ammodendron
Fabaceae_Astragalus_acutifolius
Fabaceae_Astragalus_squarrosus
Fabaceae_Astragalus_inchebroonensis
Fabaceae_Astragalus_sympileicalycinus
Fabaceae_Astragalus_podolobus
Fabaceae_Astragalus_podoloboides
Fabaceae_Astragalus_sangonensis
Fabaceae_Astragalus_jaskensis
Fabaceae_Astragalus_brevicalycinus
Fabaceae_Astragalus_macrobotrys
Fabaceae_Astragalus_oligophyllus
Fabaceae_Astragalus_ghamishluensis
Fabaceae_Astragalus_darrehbidensis
Fabaceae_Astragalus_joharchii
Fabaceae_Astragalus_pseudoarvatensis
Fabaceae_Astragalus_semiglabricarpus
Fabaceae_Astragalus_nigricans
Fabaceae_Astragalus_bazarganii
Fabaceae_Astragalus_tarumensis
Fabaceae_Astragalus_gebleri
Fabaceae_Astragalus_tenuiramosus
Fabaceae_Astragalus_saxifractor
Fabaceae_Astragalus_commixtus
Fabaceae_Astragalus_sesamoides
Fabaceae_Astragalus_ophiocarpus
Fabaceae_Astragalus_harpilobus
Fabaceae_Astragalus_campylotrichus
Fabaceae_Astragalus_filicaulis
Fabaceae_Astragalus_bakaliensis
Fabaceae_Astragalus_eremophilus
Fabaceae_Barnebyella_calycina
Fabaceae_Astragalus_scaberrimus
Fabaceae_Astragalus_argyroides
Fabaceae_Astragalus_viridis
Fabaceae_Astragalus_beypazaricus
Fabaceae_Astragalus_kastamonuensis
Fabaceae_Astragalus_cornutus

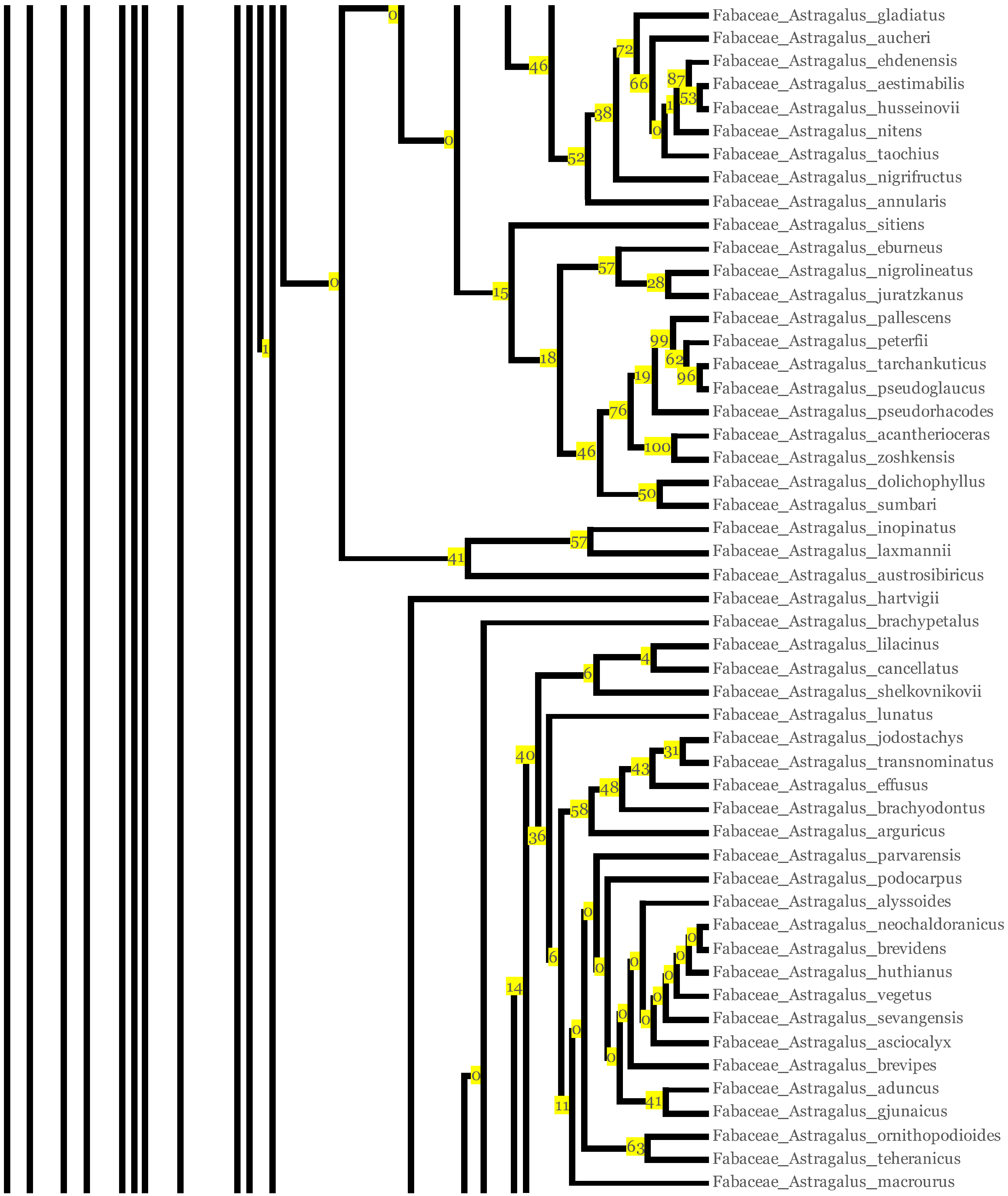
Fabaceae_Astragalus_gladiatus
Fabaceae_Astragalus_aucheri
Fabaceae_Astragalus_ehdenensis
Fabaceae_Astragalus_aestimabilis
Fabaceae_Astragalus_husseinovii
Fabaceae_Astragalus_nitens
Fabaceae_Astragalus_taochius
Fabaceae_Astragalus_nigrifructus
Fabaceae_Astragalus_annularis
Fabaceae_Astragalus_sitiens
Fabaceae_Astragalus_eburneus
Fabaceae_Astragalus_nigrolineatus
Fabaceae_Astragalus_juratzkanus
Fabaceae_Astragalus_pallescens
Fabaceae_Astragalus_peterfii
Fabaceae_Astragalus_tarchankuticus
Fabaceae_Astragalus_pseudoglaucus
Fabaceae_Astragalus_pseudorhacodes
Fabaceae_Astragalus_acantherioceras
Fabaceae_Astragalus_zoshkensis
Fabaceae_Astragalus_dolichophyllus
Fabaceae_Astragalus_sumbari
Fabaceae_Astragalus_inopinatus
Fabaceae_Astragalus_laxmannii
Fabaceae_Astragalus_austrosibiricus
Fabaceae_Astragalus_hartvigii
Fabaceae_Astragalus_brachypetalus
Fabaceae_Astragalus_lilacinus
Fabaceae_Astragalus_cancellatus
Fabaceae_Astragalus_shelkovnikovii
Fabaceae_Astragalus_lunatus
Fabaceae_Astragalus_jodostachys
Fabaceae_Astragalus_transnominatus
Fabaceae_Astragalus_effusus
Fabaceae_Astragalus_brachyodontus
Fabaceae_Astragalus_arguricus
Fabaceae_Astragalus_parvarensis
Fabaceae_Astragalus_podocarpus
Fabaceae_Astragalus_alyssoides
Fabaceae_Astragalus_neochaldoranicus
Fabaceae_Astragalus_brevidens
Fabaceae_Astragalus_huthianus
Fabaceae_Astragalus_vegetus
Fabaceae_Astragalus_sevangensis
Fabaceae_Astragalus_asciocalyx
Fabaceae_Astragalus_brevipes
Fabaceae_Astragalus_aduncus
Fabaceae_Astragalus_gjunaicus
Fabaceae_Astragalus_ornithopodioides
Fabaceae_Astragalus_teheranicus
Fabaceae_Astragalus_macrourus

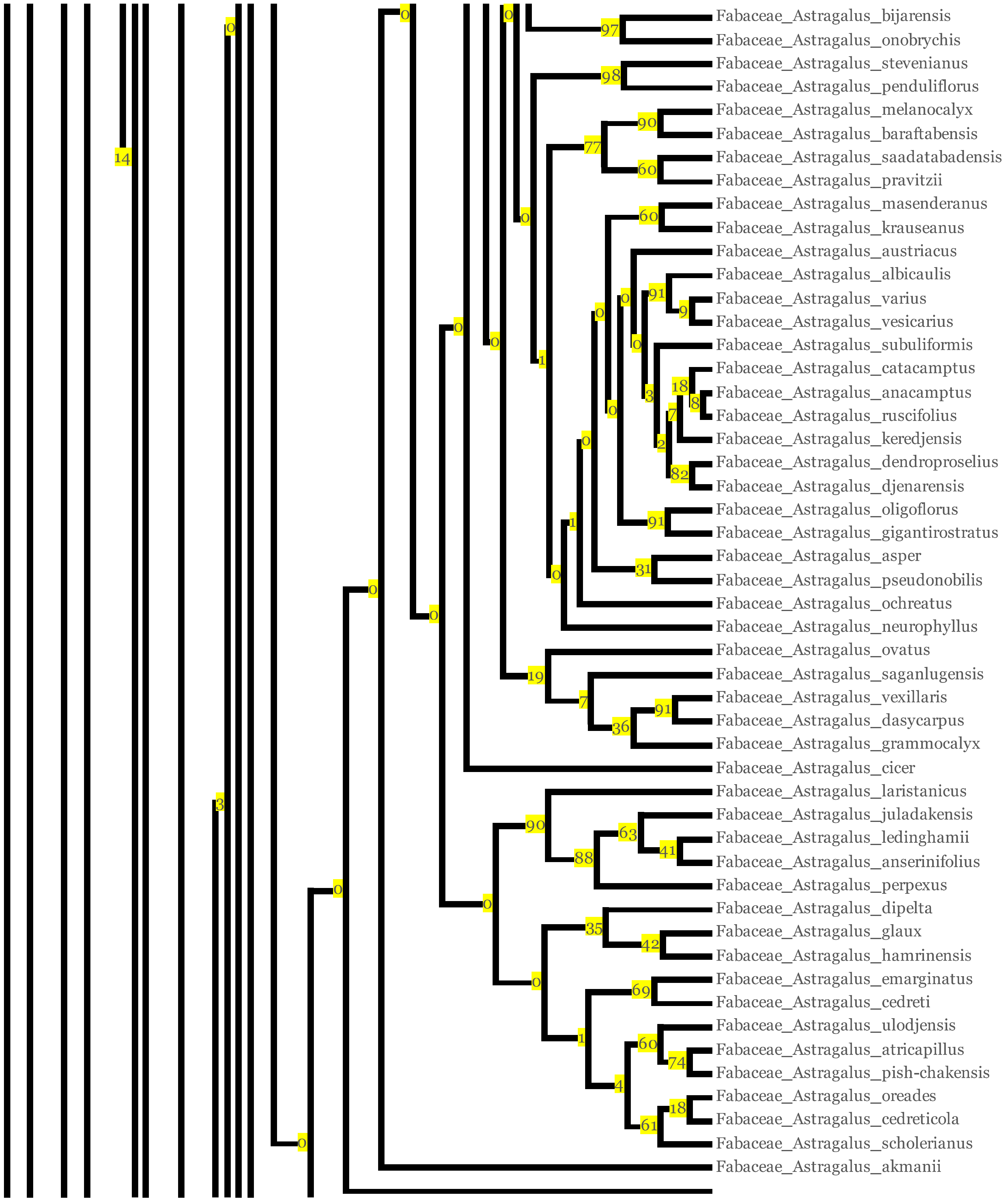
Fabaceae_Astragalus_bijarensis
Fabaceae_Astragalus_onobrychis
Fabaceae_Astragalus_stevenianus
Fabaceae_Astragalus_penduliflorus
Fabaceae_Astragalus_melanocalyx
Fabaceae_Astragalus_baraftabensis
Fabaceae_Astragalus_saadatabadensis
Fabaceae_Astragalus_pravitzii
Fabaceae_Astragalus_masenderanus
Fabaceae_Astragalus_krauseanus
Fabaceae_Astragalus_austriacus
Fabaceae_Astragalus_albicaulis
Fabaceae_Astragalus_varius
Fabaceae_Astragalus_vesicarius
Fabaceae_Astragalus_subuliformis
Fabaceae_Astragalus_catacamptus
Fabaceae_Astragalus_anacamptus
Fabaceae_Astragalus_ruscifolius
Fabaceae_Astragalus_keredjensis
Fabaceae_Astragalus_dendroproselius
Fabaceae_Astragalus_djenarensis
Fabaceae_Astragalus_oligoflorus
Fabaceae_Astragalus_gigantirostratus
Fabaceae_Astragalus_asper
Fabaceae_Astragalus_pseudonobilis
Fabaceae_Astragalus_ochreatus
Fabaceae_Astragalus_neurophyllus
Fabaceae_Astragalus_ovatus
Fabaceae_Astragalus_saganlugensis
Fabaceae_Astragalus_vexillaris
Fabaceae_Astragalus_dasycarpus
Fabaceae_Astragalus_grammocalyx
Fabaceae_Astragalus_cicer
Fabaceae_Astragalus_laristanicus
Fabaceae_Astragalus_juladakensis
Fabaceae_Astragalus_ledinghamii
Fabaceae_Astragalus_anserinifolius
Fabaceae_Astragalus_perpexus
Fabaceae_Astragalus_dipelta
Fabaceae_Astragalus_glaux
Fabaceae_Astragalus_hamrinensis
Fabaceae_Astragalus_emarginatus
Fabaceae_Astragalus_cedreti
Fabaceae_Astragalus_ulodjensis
Fabaceae_Astragalus_atricapillus
Fabaceae_Astragalus_pish-chakensis
Fabaceae_Astragalus_oreades
Fabaceae_Astragalus_cedreticola
Fabaceae_Astragalus_scholerianus
Fabaceae_Astragalus_akmanii

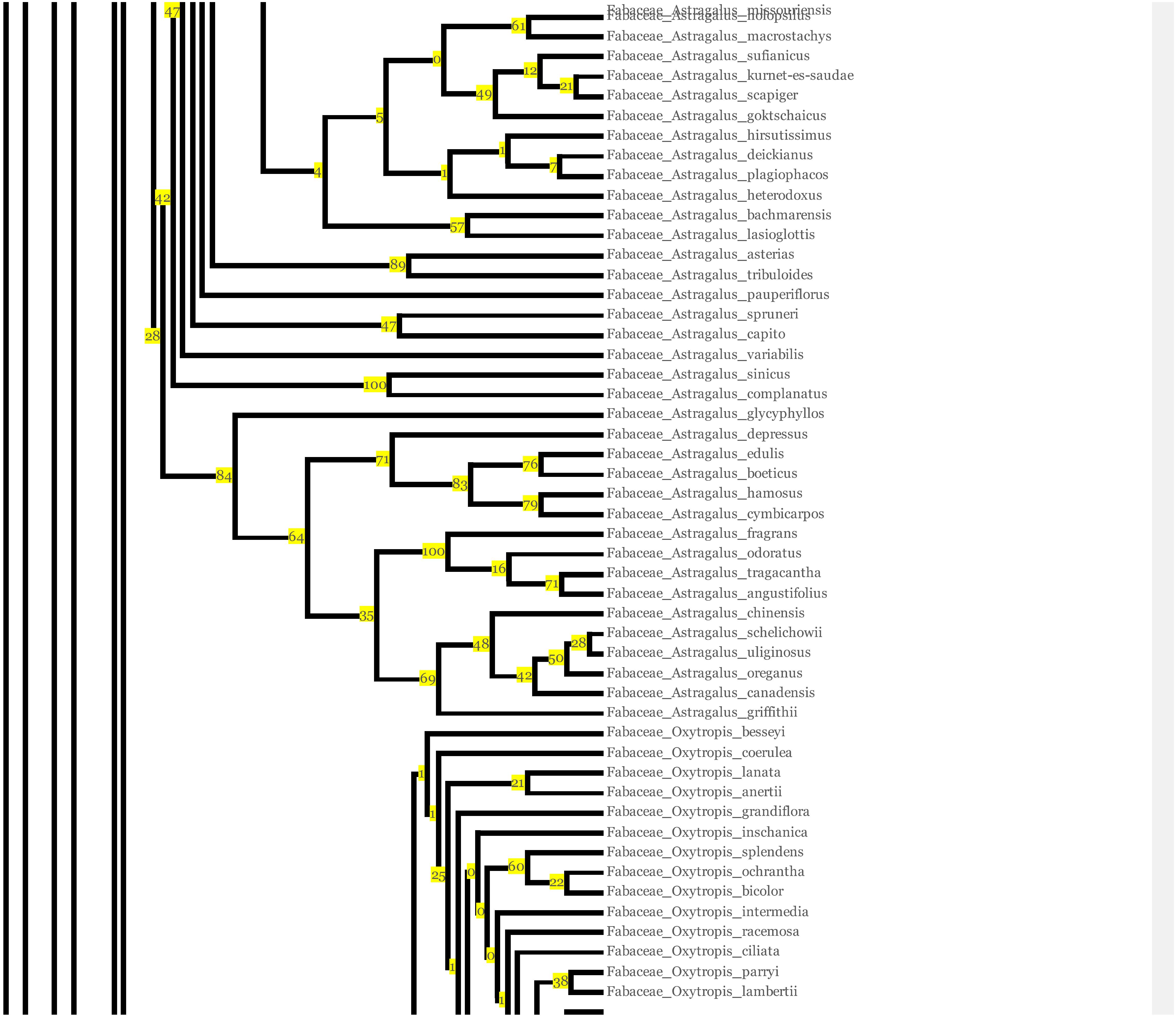
Fabaceae_Astragalus_missouriensis
Fabaceae_Astragalus_holopsilus
Fabaceae_Astragalus_macrostachys
Fabaceae_Astragalus_sufianicus
Fabaceae_Astragalus_kurnet-es-saudae
Fabaceae_Astragalus_scapiger
Fabaceae_Astragalus_goktschaicus
Fabaceae_Astragalus_hirsutissimus
Fabaceae_Astragalus_deickianus
Fabaceae_Astragalus_plagiophacos
Fabaceae_Astragalus_heterodoxus
Fabaceae_Astragalus_bachmarensis
Fabaceae_Astragalus_lasioglottis
Fabaceae_Astragalus_asterias
Fabaceae_Astragalus_tribuloides
Fabaceae_Astragalus_pauperiflorus
Fabaceae_Astragalus_spruneri
Fabaceae_Astragalus_capito
Fabaceae_Astragalus_variabilis
Fabaceae_Astragalus_sinicus
Fabaceae_Astragalus_complanatus
Fabaceae_Astragalus_glycyphyllos
Fabaceae_Astragalus_depressus
Fabaceae_Astragalus_edulis
Fabaceae_Astragalus_boeticus
Fabaceae_Astragalus_hamosus
Fabaceae_Astragalus_cymbicarpos
Fabaceae_Astragalus_fragrans
Fabaceae_Astragalus_odoratus
Fabaceae_Astragalus_tragacantha
Fabaceae_Astragalus_angustifolius
Fabaceae_Astragalus_chinensis
Fabaceae_Astragalus_schelichowii
Fabaceae_Astragalus_uliginosus
Fabaceae_Astragalus_oreganus
Fabaceae_Astragalus_canadensis
Fabaceae_Astragalus_griffithii
Fabaceae_Oxytropis_besseyi
Fabaceae_Oxytropis_coerulea
Fabaceae_Oxytropis_lanata
Fabaceae_Oxytropis_anertii
Fabaceae_Oxytropis_grandiflora
Fabaceae_Oxytropis_inschanica
Fabaceae_Oxytropis_splendens
Fabaceae_Oxytropis_ochrantha
Fabaceae_Oxytropis_bicolor
Fabaceae_Oxytropis_intermedia
Fabaceae_Oxytropis_racemosa
Fabaceae_Oxytropis_ciliata
Fabaceae_Oxytropis_parryi
Fabaceae_Oxytropis_lambertii

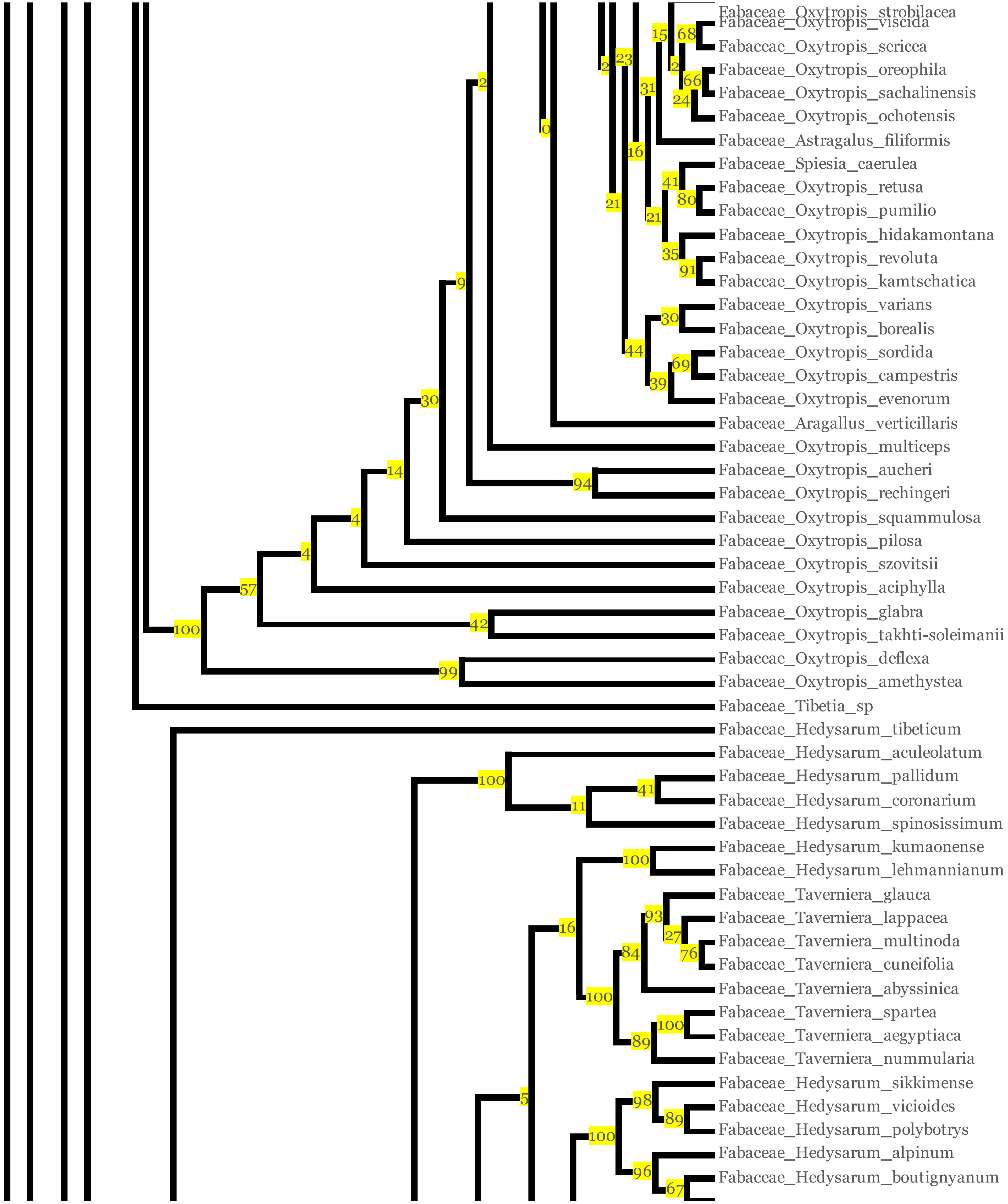
Fabaceae_Oxytropis_strobilacea
Fabaceae_Oxytropis_viscida
Fabaceae_Oxytropis_sericea
Fabaceae_Oxytropis_oreophila
Fabaceae_Oxytropis_sachalinensis
Fabaceae_Oxytropis_ochotensis
Fabaceae_Astragalus_filiformis
Fabaceae_Spiesia_caerulea
Fabaceae_Oxytropis_retusa
Fabaceae_Oxytropis_pumilio
Fabaceae_Oxytropis_hidakamontana
Fabaceae_Oxytropis_revoluta
Fabaceae_Oxytropis_kamtschatica
Fabaceae_Oxytropis_varians
Fabaceae_Oxytropis_borealis
Fabaceae_Oxytropis_sordida
Fabaceae_Oxytropis_campestris
Fabaceae_Oxytropis_evenorum
Fabaceae_Aragallus_verticillaris
Fabaceae_Oxytropis_multiceps
Fabaceae_Oxytropis_aucheri
Fabaceae_Oxytropis_rechingeri
Fabaceae_Oxytropis_squammulosa
Fabaceae_Oxytropis_pilosa
Fabaceae_Oxytropis_szovitsii
Fabaceae_Oxytropis_aciphylla
Fabaceae_Oxytropis_glabra
Fabaceae_Oxytropis_takhti-soleimanii
Fabaceae_Oxytropis_deflexa
Fabaceae_Oxytropis_amethystea
Fabaceae_Tibetia_sp
Fabaceae_Hedysarum_tibeticum
Fabaceae_Hedysarum_aculeolatum
Fabaceae_Hedysarum_pallidum
Fabaceae_Hedysarum_coronarium
Fabaceae_Hedysarum_spinosissimum
Fabaceae_Hedysarum_kumaonense
Fabaceae_Hedysarum_lehmannianum
Fabaceae_Taverniera_glauca
Fabaceae_Taverniera_lappacea
Fabaceae_Taverniera_multinoda
Fabaceae_Taverniera_cuneifolia
Fabaceae_Taverniera_abyssinica
Fabaceae_Taverniera_spartea
Fabaceae_Taverniera_aegyptiaca
Fabaceae_Taverniera_nummularia
Fabaceae_Hedysarum_sikkimense
Fabaceae_Hedysarum_vicioides
Fabaceae_Hedysarum_polybotrys
Fabaceae_Hedysarum_alpinum
Fabaceae_Hedysarum_boutignyanum

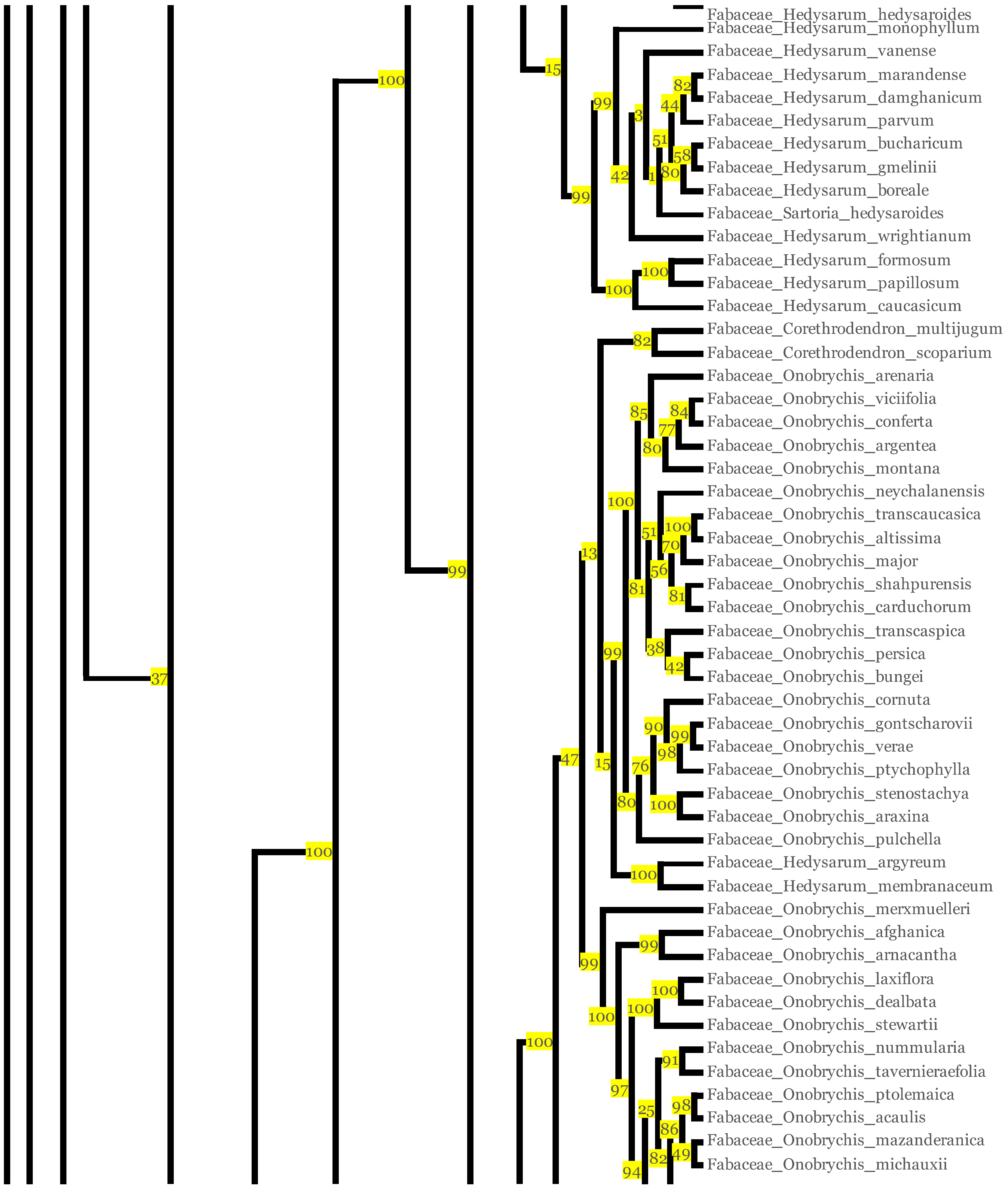
Fabaceae_Hedysarum_hedysaroides
Fabaceae_Hedysarum_monophyllum
Fabaceae_Hedysarum_vanense
Fabaceae_Hedysarum_marandense
Fabaceae_Hedysarum_damghanicum
Fabaceae_Hedysarum_parvum
Fabaceae_Hedysarum_bucharicum
Fabaceae_Hedysarum_gmelinii
Fabaceae_Hedysarum_boreale
Fabaceae_Sartoria_hedysaroides
Fabaceae_Hedysarum_wrightianum
Fabaceae_Hedysarum_formosum
Fabaceae_Hedysarum_papillosum
Fabaceae_Hedysarum_caucasicum
Fabaceae_Corethrodendron_multijugum
Fabaceae_Corethrodendron_scoparium
Fabaceae_Onobrychis_arenaria
Fabaceae_Onobrychis_viciifolia
Fabaceae_Onobrychis_conferta
Fabaceae_Onobrychis_argentea
Fabaceae_Onobrychis_montana
Fabaceae_Onobrychis_neychalanensis
Fabaceae_Onobrychis_transcaucasica
Fabaceae_Onobrychis_altissima
Fabaceae_Onobrychis_major
Fabaceae_Onobrychis_shahpurensis
Fabaceae_Onobrychis_carduchorum
Fabaceae_Onobrychis_transcaspica
Fabaceae_Onobrychis_persica
Fabaceae_Onobrychis_bungei
Fabaceae_Onobrychis_cornuta
Fabaceae_Onobrychis_gontscharovii
Fabaceae_Onobrychis_verae
Fabaceae_Onobrychis_ptychophylla
Fabaceae_Onobrychis_stenostachya
Fabaceae_Onobrychis_araxina
Fabaceae_Onobrychis_pulchella
Fabaceae_Hedysarum_argyreum
Fabaceae_Hedysarum_membranaceum
Fabaceae_Onobrychis_merxmuelleri
Fabaceae_Onobrychis_afghanica
Fabaceae_Onobrychis_arnacantha
Fabaceae_Onobrychis_laxiflora
Fabaceae_Onobrychis_dealbata
Fabaceae_Onobrychis_stewartii
Fabaceae_Onobrychis_nummularia
Fabaceae_Onobrychis_tavernieraefolia
Fabaceae_Onobrychis_ptolemaica
Fabaceae_Onobrychis_acaulis
Fabaceae_Onobrychis_mazanderanica
Fabaceae_Onobrychis_michauxii
100
15
99
42
3
82
44
51
58
1
80
99
100
100
99
37
82
85
84
77
80
100
51
100
70
56
81
81
38
42
13
99
90
99
98
15
76
80
100
47
100
100
99
99
100
100
100
91
97
25
98
86
49
82
94

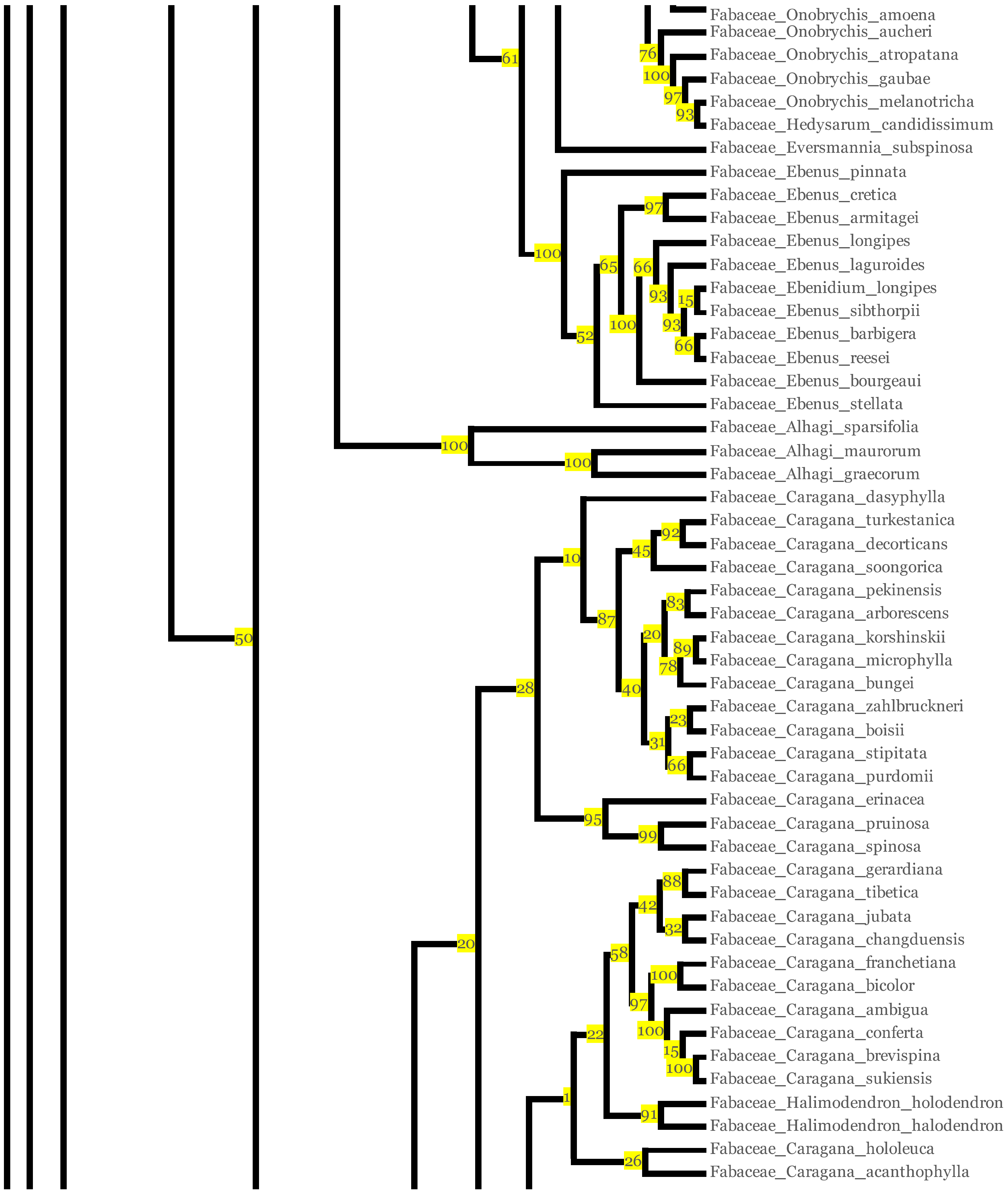
Fabaceae_Onobrychis_amoena
Fabaceae_Onobrychis_aucheri
Fabaceae_Onobrychis_atropatana
Fabaceae_Onobrychis_gaubae
Fabaceae_Onobrychis_melanotricha
Fabaceae_Hedysarum_candidissimum
Fabaceae_Eversmannia_subspinosa
Fabaceae_Ebenus_pinnata
Fabaceae_Ebenus_cretica
Fabaceae_Ebenus_armitagei
Fabaceae_Ebenus_longipes
Fabaceae_Ebenus_laguroides
Fabaceae_Ebenidium_longipes
Fabaceae_Ebenus_sibthorpii
Fabaceae_Ebenus_barbigera
Fabaceae_Ebenus_reesei
Fabaceae_Ebenus_bourgeaui
Fabaceae_Ebenus_stellata
Fabaceae_Alhagi_sparsifolia
Fabaceae_Alhagi_maurorum
Fabaceae_Alhagi_graecorum
Fabaceae_Caragana_dasyphylla
Fabaceae_Caragana_turkestanica
Fabaceae_Caragana_decorticans
Fabaceae_Caragana_soongorica
Fabaceae_Caragana_pekinensis
Fabaceae_Caragana_arborescens
Fabaceae_Caragana_korshinskii
Fabaceae_Caragana_microphylla
Fabaceae_Caragana_bungei
Fabaceae_Caragana_zahlbruckneri
Fabaceae_Caragana_boisii
Fabaceae_Caragana_stipitata
Fabaceae_Caragana_purdomii
Fabaceae_Caragana_erinacea
Fabaceae_Caragana_pruinosa
Fabaceae_Caragana_spinosa
Fabaceae_Caragana_gerardiana
Fabaceae_Caragana_tibetica
Fabaceae_Caragana_jubata
Fabaceae_Caragana_changduensis
Fabaceae_Caragana_franchetiana
Fabaceae_Caragana_bicolor
Fabaceae_Caragana_ambigua
Fabaceae_Caragana_conferta
Fabaceae_Caragana_brevispina
Fabaceae_Caragana_sukiensis
Fabaceae_Halimodendron_holodendron
Fabaceae_Halimodendron_halodendron
Fabaceae_Caragana_hololeuca
Fabaceae_Caragana_acanthophylla

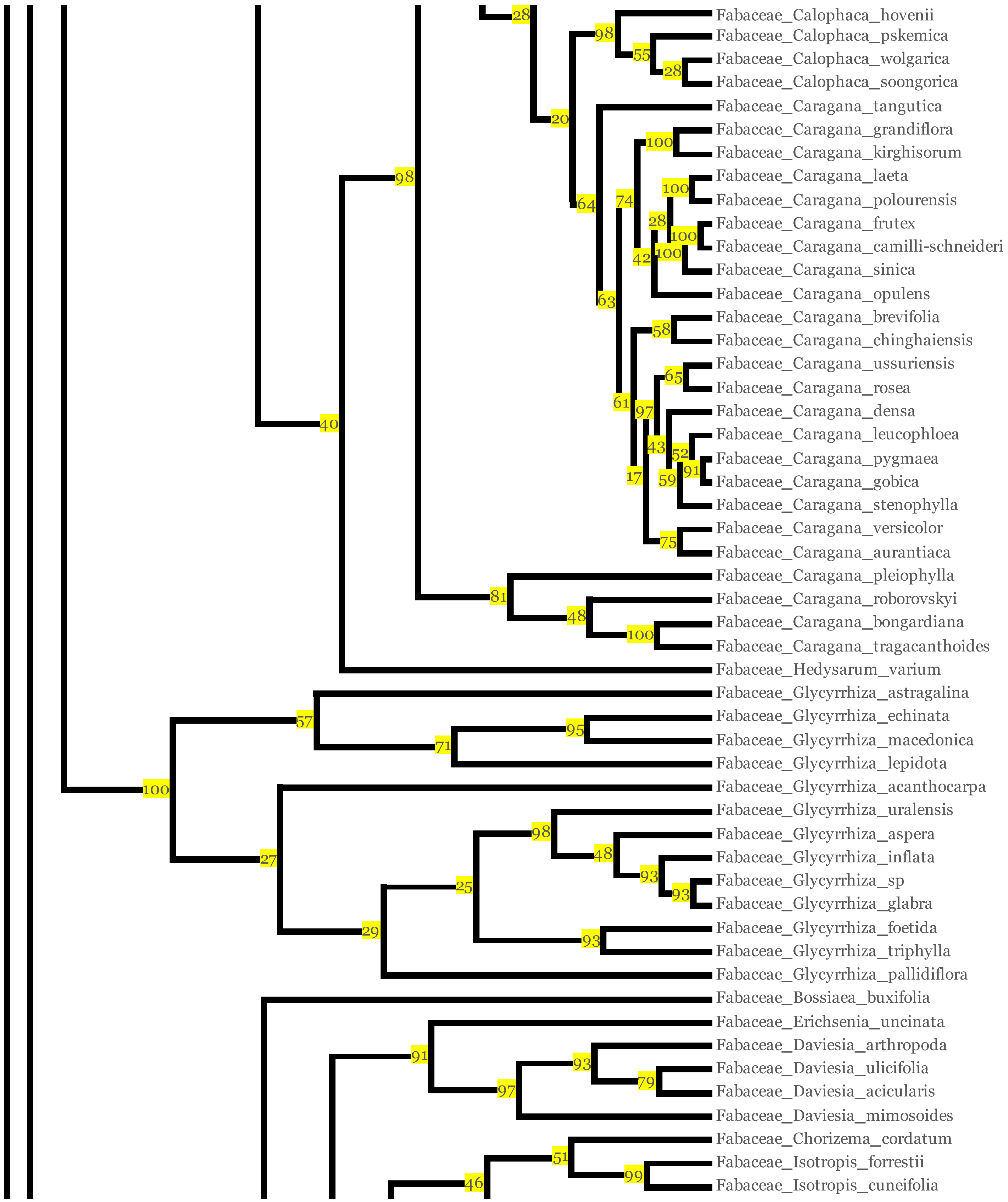
Fabaceae_Calophaca_hovenii
Fabaceae_Calophaca_pskemica
Fabaceae_Calophaca_wolgarica
Fabaceae_Calophaca_soongorica
Fabaceae_Caragana_tangutica
Fabaceae_Caragana_grandiflora
Fabaceae_Caragana_kirghisorum
Fabaceae_Caragana_laeta
Fabaceae_Caragana_polourensis
Fabaceae_Caragana_frutex
Fabaceae_Caragana_camilli-schneideri
Fabaceae_Caragana_sinica
Fabaceae_Caragana_opulens
Fabaceae_Caragana_brevifolia
Fabaceae_Caragana_chinghaiensis
Fabaceae_Caragana_ussuriensis
Fabaceae_Caragana_rosea
Fabaceae_Caragana_densa
Fabaceae_Caragana_leucophloea
Fabaceae_Caragana_pygmaea
Fabaceae_Caragana_gobica
Fabaceae_Caragana_stenophylla
Fabaceae_Caragana_versicolor
Fabaceae_Caragana_aurantiaca
Fabaceae_Caragana_pleiophylla
Fabaceae_Caragana_roborovskyi
Fabaceae_Caragana_bongardiana
Fabaceae_Caragana_tragacanthoides
Fabaceae_Hedysarum_varium
Fabaceae_Glycyrrhiza_astragalina
Fabaceae_Glycyrrhiza_echinata
Fabaceae_Glycyrrhiza_macedonica
Fabaceae_Glycyrrhiza_lepidota
Fabaceae_Glycyrrhiza_acanthocarpa
Fabaceae_Glycyrrhiza_uralensis
Fabaceae_Glycyrrhiza_aspera
Fabaceae_Glycyrrhiza_inflata
Fabaceae_Glycyrrhiza_sp
Fabaceae_Glycyrrhiza_glabra
Fabaceae_Glycyrrhiza_foetida
Fabaceae_Glycyrrhiza_triphylla
Fabaceae_Glycyrrhiza_pallidiflora
Fabaceae_Bossiaea_buxifolia
Fabaceae_Erichsenia_uncinata
Fabaceae_Daviesia_arthropoda
Fabaceae_Daviesia_ulicifolia
Fabaceae_Daviesia_acicularis
Fabaceae_Daviesia_mimosoides
Fabaceae_Chorizema_cordatum
Fabaceae_Isotropis_forrestii
Fabaceae_Isotropis_cuneifolia
28
98
55
28
20
100
100
64
74
28
100
42
100
63
58
65
61
97
43
52
91
17
59
75
98
40
81
48
100
57
95
71
100
98
48
27
93
93
25
29
93
91
93
79
97
51
99
46

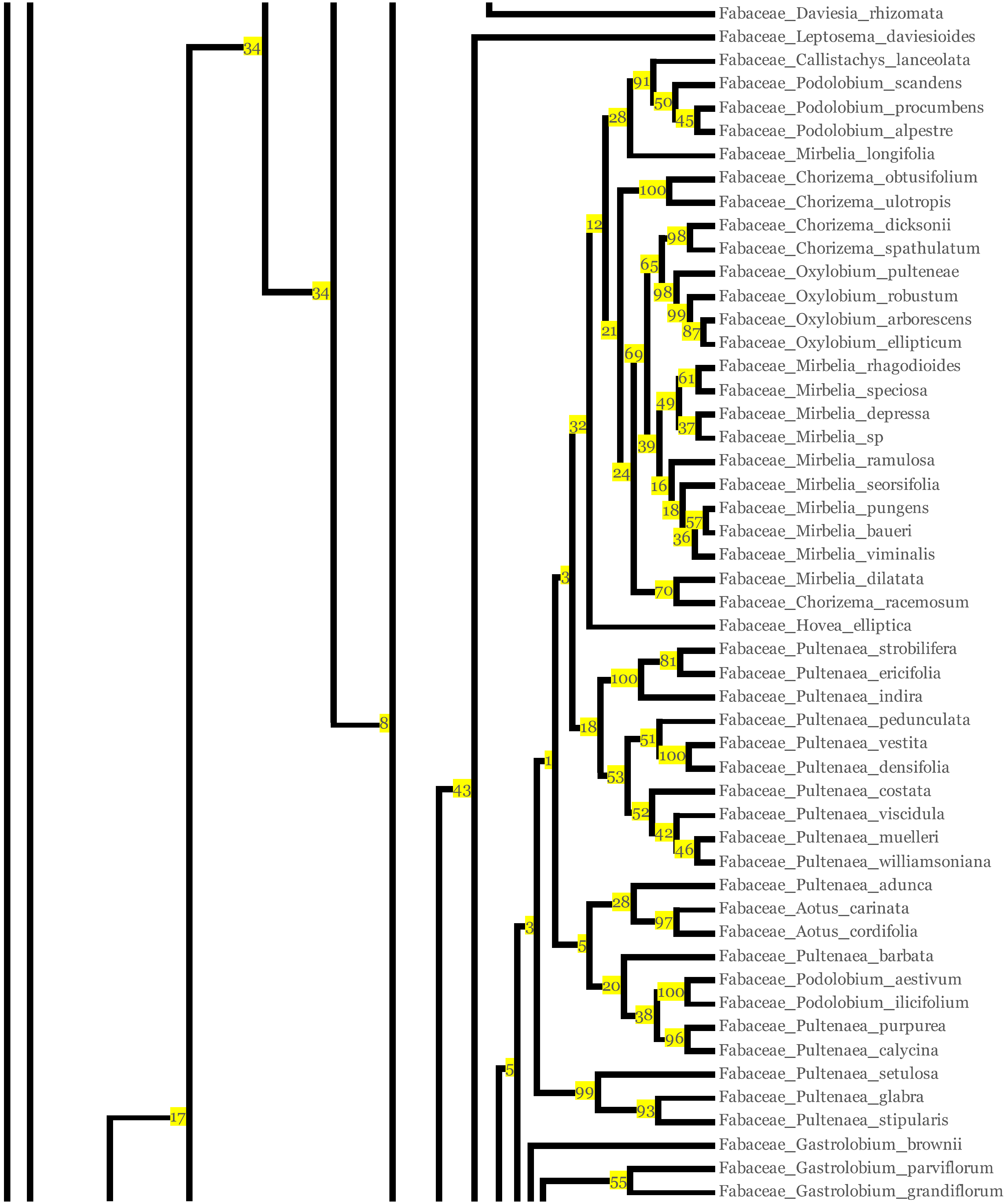
Fabaceae_Daviesia_rhizomata
Fabaceae_Leptosema_daviesioides
Fabaceae_Callistachys_lanceolata
Fabaceae_Podolobium_scandens
Fabaceae_Podolobium_procumbens
Fabaceae_Podolobium_alpestre
Fabaceae_Mirbelia_longifolia
Fabaceae_Chorizema_obtusifolium
Fabaceae_Chorizema_ulotropis
Fabaceae_Chorizema_dicksonii
Fabaceae_Chorizema_spathulatum
Fabaceae_Oxylobium_pulteneae
Fabaceae_Oxylobium_robustum
Fabaceae_Oxylobium_arborescens
Fabaceae_Oxylobium_ellipticum
Fabaceae_Mirbelia_rhagodioides
Fabaceae_Mirbelia_speciosa
Fabaceae_Mirbelia_depressa
Fabaceae_Mirbelia_sp
Fabaceae_Mirbelia_ramulosa
Fabaceae_Mirbelia_seorsifolia
Fabaceae_Mirbelia_pungens
Fabaceae_Mirbelia_baueri
Fabaceae_Mirbelia_viminalis
Fabaceae_Mirbelia_dilatata
Fabaceae_Chorizema_racemosum
Fabaceae_Hovea_elliptica
Fabaceae_Pultenaea_strobilifera
Fabaceae_Pultenaea_ericifolia
Fabaceae_Pultenaea_indira
Fabaceae_Pultenaea_pedunculata
Fabaceae_Pultenaea_vestita
Fabaceae_Pultenaea_densifolia
Fabaceae_Pultenaea_costata
Fabaceae_Pultenaea_viscidula
Fabaceae_Pultenaea_muelleri
Fabaceae_Pultenaea_williamsoniana
Fabaceae_Pultenaea_adunca
Fabaceae_Aotus_carinata
Fabaceae_Aotus_cordifolia
Fabaceae_Pultenaea_barbata
Fabaceae_Podolobium_aestivum
Fabaceae_Podolobium_ilicifolium
Fabaceae_Pultenaea_purpurea
Fabaceae_Pultenaea_calycina
Fabaceae_Pultenaea_setulosa
Fabaceae_Pultenaea_glabra
Fabaceae_Pultenaea_stipularis
Fabaceae_Gastrolobium_brownii
Fabaceae_Gastrolobium_parviflorum
Fabaceae_Gastrolobium_grandiflorum
34
34
91
50
45
28
100
12
98
65
98
99
87
21
69
61
49
37
32
39
24
16
18
57
36
3
70
81
100
8
18
51
100
1
53
43
52
42
46
28
97
5
3
20
100
38
96
5
99
93
17
55

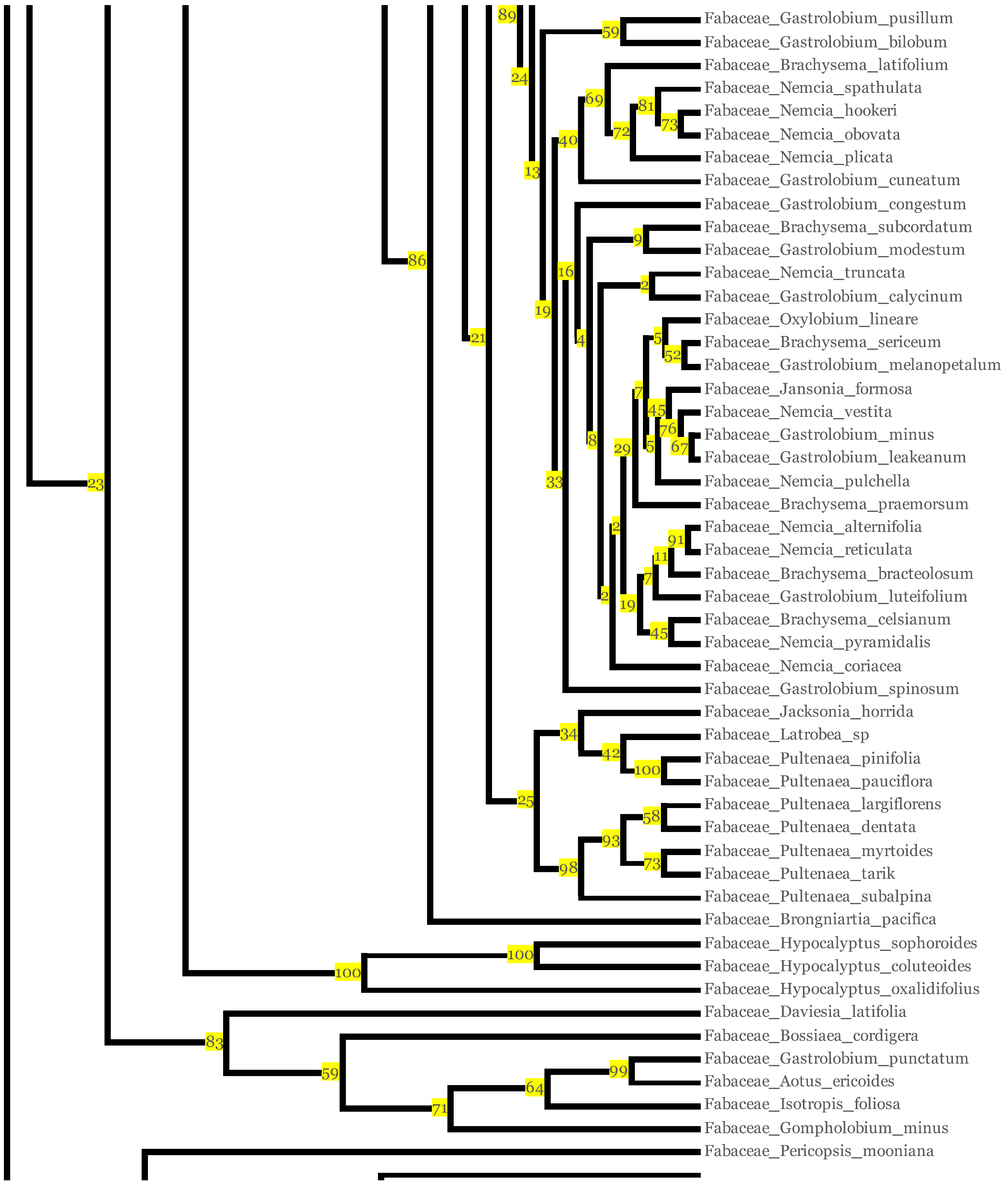

Fabaceae_Gastrolobium_pusillum
Fabaceae_Gastrolobium_bilobum
Fabaceae_Brachysema_latifolium
Fabaceae_Nemcia_spathulata
Fabaceae_Nemcia_hookeri
Fabaceae_Nemcia_obovata
Fabaceae_Nemcia_plicata
Fabaceae_Gastrolobium_cuneatum
Fabaceae_Gastrolobium_congestum
Fabaceae_Brachysema_subcordatum
Fabaceae_Gastrolobium_modestum
Fabaceae_Nemcia_truncata
Fabaceae_Gastrolobium_calycinum
Fabaceae_Oxylobium_lineare
Fabaceae_Brachysema_sericeum
Fabaceae_Gastrolobium_melanopetalum
Fabaceae_Jansonia_formosa
Fabaceae_Nemcia_vestita
Fabaceae_Gastrolobium_minus
Fabaceae_Gastrolobium_leakeanum
Fabaceae_Nemcia_pulchella
Fabaceae_Brachysema_praemorsum
Fabaceae_Nemcia_alternifolia
Fabaceae_Nemcia_reticulata
Fabaceae_Brachysema_bracteolosum
Fabaceae_Gastrolobium_luteifolium
Fabaceae_Brachysema_celsianum
Fabaceae_Nemcia_pyramidalis
Fabaceae_Nemcia_coriacea
Fabaceae_Gastrolobium_spinosum
Fabaceae_Jacksonia_horrida
Fabaceae_Latrobea_sp
Fabaceae_Pultenaea_pinifolia
Fabaceae_Pultenaea_pauciflora
Fabaceae_Pultenaea_largiflorens
Fabaceae_Pultenaea_dentata
Fabaceae_Pultenaea_myrtoides
Fabaceae_Pultenaea_tarik
Fabaceae_Pultenaea_subalpina
Fabaceae_Brongniartia_pacifica
Fabaceae_Hypocalyptus_sophoroides
Fabaceae_Hypocalyptus_coluteoides
Fabaceae_Hypocalyptus_oxalidifolius
Fabaceae_Daviesia_latifolia
Fabaceae_Bossiaea_cordigera
Fabaceae_Gastrolobium_punctatum
Fabaceae_Aotus_ericoides
Fabaceae_Isotropis_foliosa
Fabaceae_Gompholobium_minus
Fabaceae_Pericopsis_mooniana

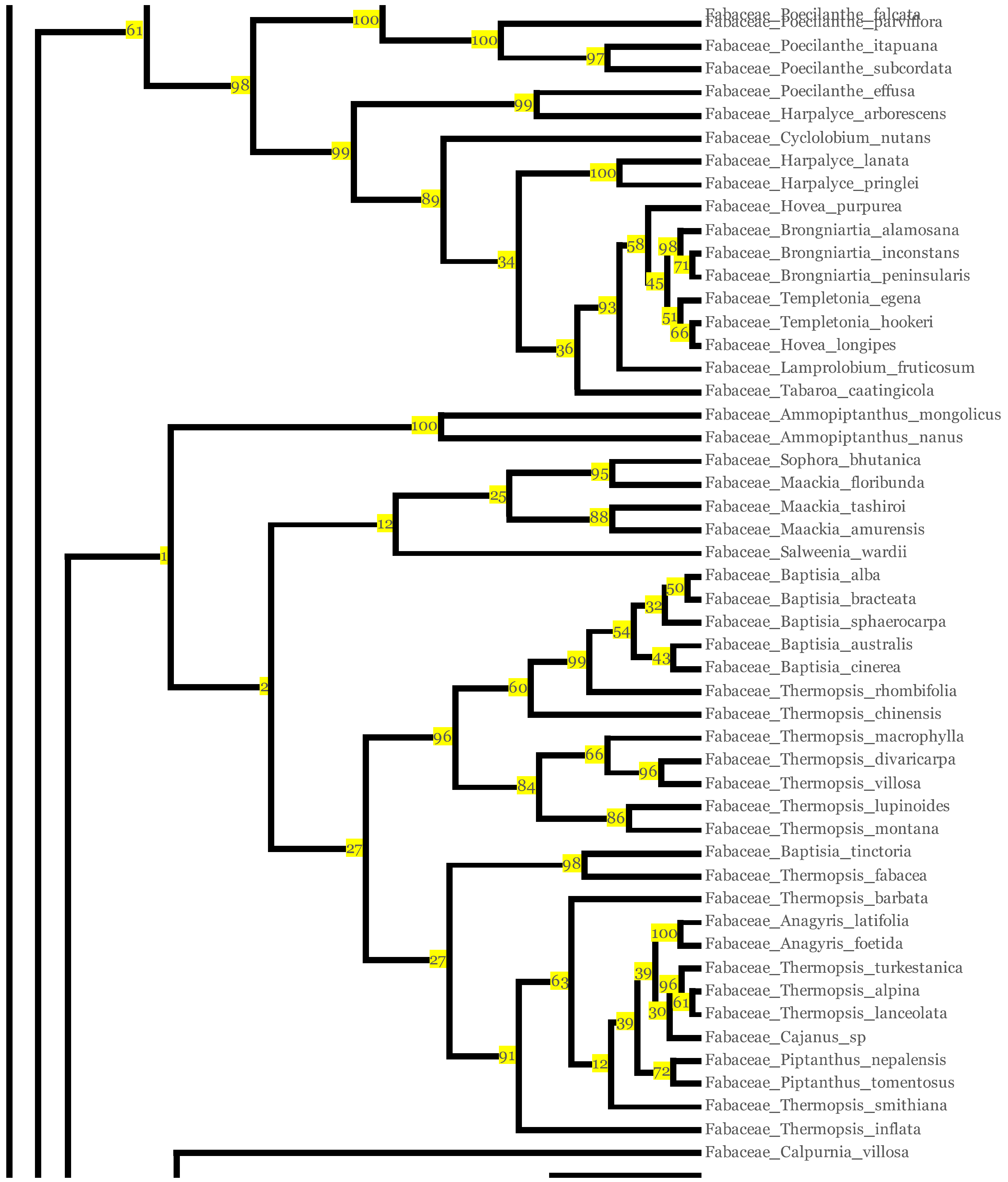
Fabaceae_Poecilanthe_falcata
Fabaceae_Poecilanthe_parviflora
Fabaceae_Poecilanthe_itapuana
Fabaceae_Poecilanthe_subcordata
Fabaceae_Poecilanthe_effusa
Fabaceae_Harpalyce_arborescens
Fabaceae_Cyclolobium_nutans
Fabaceae_Harpalyce_lanata
Fabaceae_Harpalyce_pringlei
Fabaceae_Hovea_purpurea
Fabaceae_Brongniartia_alamosana
Fabaceae_Brongniartia_inconstans
Fabaceae_Brongniartia_peninsularis
Fabaceae_Templetonia_egena
Fabaceae_Templetonia_hookeri
Fabaceae_Hovea_longipes
Fabaceae_Lamprolobium_fruticosum
Fabaceae_Tabaroa_caatingicola
Fabaceae_Ammopiptanthus_mongolicus
Fabaceae_Ammopiptanthus_nanus
Fabaceae_Sophora_bhutanica
Fabaceae_Maackia_floribunda
Fabaceae_Maackia_tashiroi
Fabaceae_Maackia_amurensis
Fabaceae_Salweenia_wardii
Fabaceae_Baptisia_alba
Fabaceae_Baptisia_bracteata
Fabaceae_Baptisia_sphaerocarpa
Fabaceae_Baptisia_australis
Fabaceae_Baptisia_cinerea
Fabaceae_Thermopsis_rhombifolia
Fabaceae_Thermopsis_chinensis
Fabaceae_Thermopsis_macrophylla
Fabaceae_Thermopsis_divaricarpa
Fabaceae_Thermopsis_villosa
Fabaceae_Thermopsis_lupinoides
Fabaceae_Thermopsis_montana
Fabaceae_Baptisia_tinctoria
Fabaceae_Thermopsis_fabacea
Fabaceae_Thermopsis_barbata
Fabaceae_Anagyris_latifolia
Fabaceae_Anagyris_foetida
Fabaceae_Thermopsis_turkestanica
Fabaceae_Thermopsis_alpina
Fabaceae_Thermopsis_lanceolata
Fabaceae_Cajanus_sp
Fabaceae_Piptanthus_nepalensis
Fabaceae_Piptanthus_tomentosus
Fabaceae_Thermopsis_smithiana
Fabaceae_Thermopsis_inflata
Fabaceae_Calpurnia_villosa

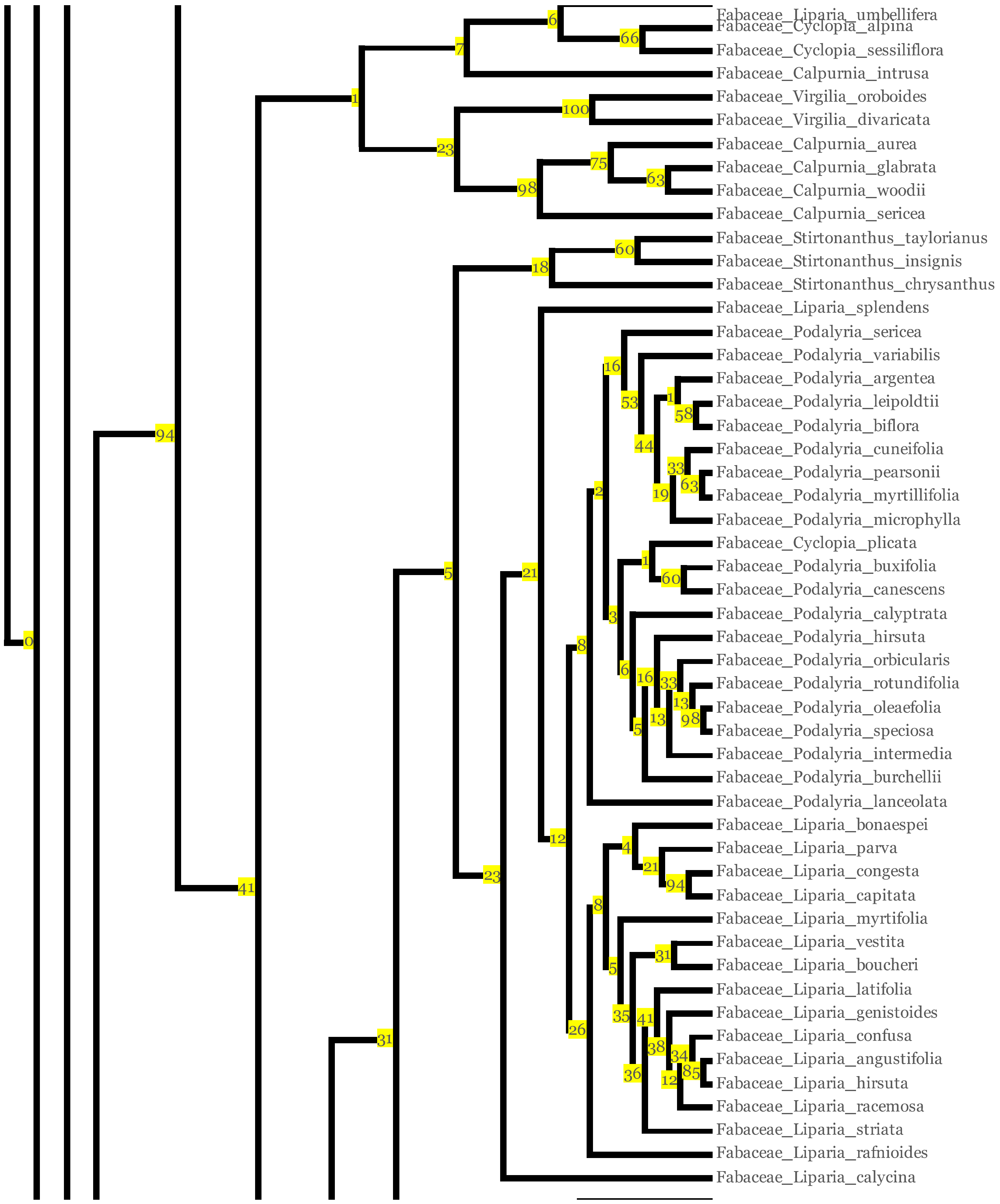
Fabaceae_Liparia_umbellifera
Fabaceae_Cyclopia_alpina
Fabaceae_Cyclopia_sessiliflora
Fabaceae_Calpurnia_intrusa
Fabaceae_Virgilia_oroboides
Fabaceae_Virgilia_divaricata
Fabaceae_Calpurnia_aurea
Fabaceae_Calpurnia_glabrata
Fabaceae_Calpurnia_woodii
Fabaceae_Calpurnia_sericea
Fabaceae_Stirtonanthus_taylorianus
Fabaceae_Stirtonanthus_insignis
Fabaceae_Stirtonanthus_chrysanthus
Fabaceae_Liparia_splendens
Fabaceae_Podalyria_sericea
Fabaceae_Podalyria_variabilis
Fabaceae_Podalyria_argentea
Fabaceae_Podalyria_leipoldtii
Fabaceae_Podalyria_biflora
Fabaceae_Podalyria_cuneifolia
Fabaceae_Podalyria_pearsonii
Fabaceae_Podalyria_myrtillifolia
Fabaceae_Podalyria_microphylla
Fabaceae_Cyclopia_plicata
Fabaceae_Podalyria_buxifolia
Fabaceae_Podalyria_canescens
Fabaceae_Podalyria_calyptrata
Fabaceae_Podalyria_hirsuta
Fabaceae_Podalyria_orbicularis
Fabaceae_Podalyria_rotundifolia
Fabaceae_Podalyria_oleaefolia
Fabaceae_Podalyria_speciosa
Fabaceae_Podalyria_intermedia
Fabaceae_Podalyria_burchellii
Fabaceae_Podalyria_lanceolata
Fabaceae_Liparia_bonaespei
Fabaceae_Liparia_parva
Fabaceae_Liparia_congesta
Fabaceae_Liparia_capitata
Fabaceae_Liparia_myrtifolia
Fabaceae_Liparia_vestita
Fabaceae_Liparia_boucheri
Fabaceae_Liparia_latifolia
Fabaceae_Liparia_genistoides
Fabaceae_Liparia_confusa
Fabaceae_Liparia_angustifolia
Fabaceae_Liparia_hirsuta
Fabaceae_Liparia_racemosa
Fabaceae_Liparia_striata
Fabaceae_Liparia_rafnioides
Fabaceae_Liparia_calycina

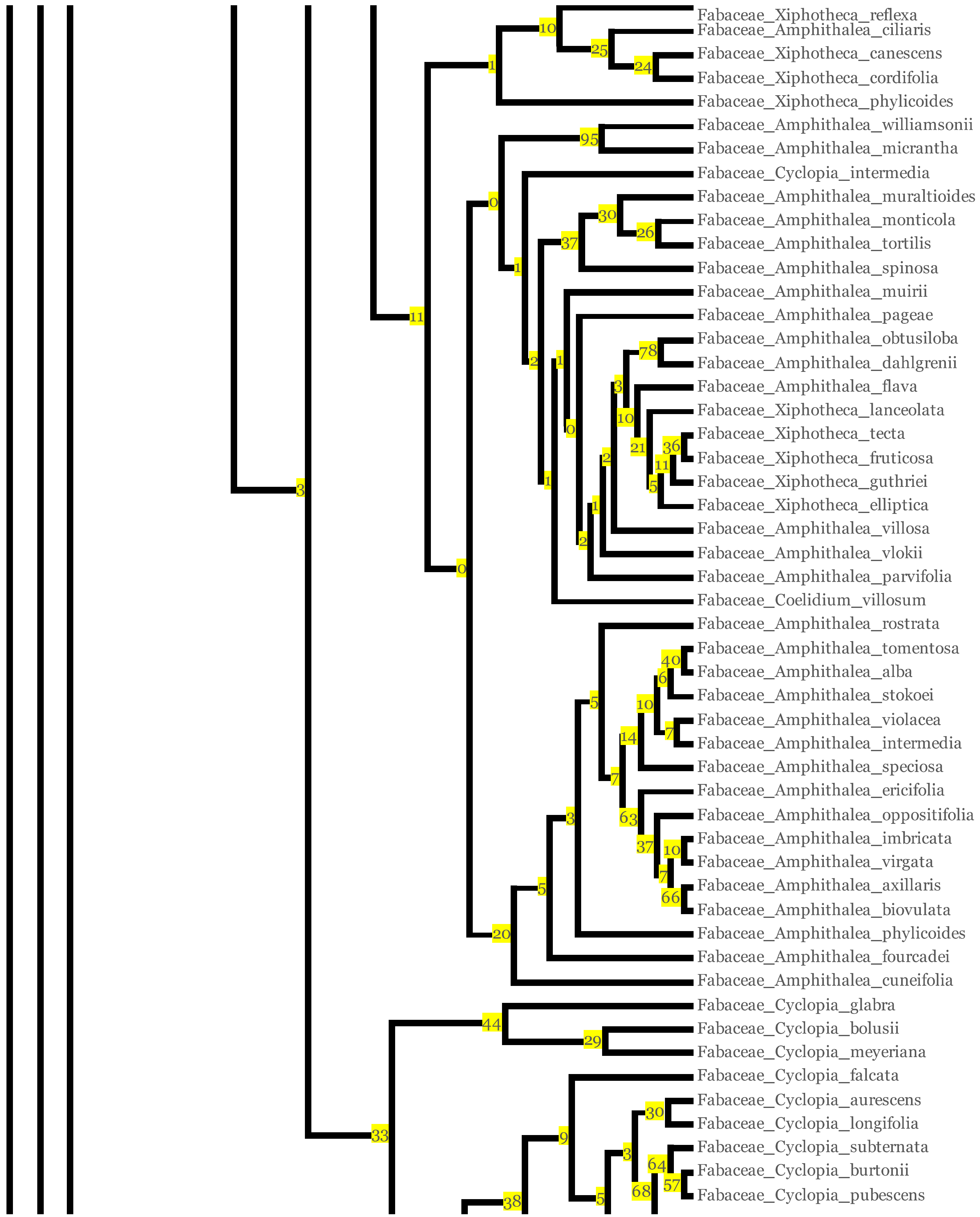
Fabaceae_Xiphotheca_reflexa
Fabaceae_Amphithalea_ciliaris
Fabaceae_Xiphotheca_canescens
Fabaceae_Xiphotheca_cordifolia
Fabaceae_Xiphotheca_phylicoides
Fabaceae_Amphithalea_williamsonii
Fabaceae_Amphithalea_micrantha
Fabaceae_Cyclopia_intermedia
Fabaceae_Amphithalea_muraltioides
Fabaceae_Amphithalea_monticola
Fabaceae_Amphithalea_tortilis
Fabaceae_Amphithalea_spinosa
Fabaceae_Amphithalea_muirii
Fabaceae_Amphithalea_pageae
Fabaceae_Amphithalea_obtusiloba
Fabaceae_Amphithalea_dahlgrenii
Fabaceae_Amphithalea_flava
Fabaceae_Xiphotheca_lanceolata
Fabaceae_Xiphotheca_tecta
Fabaceae_Xiphotheca_fruticosa
Fabaceae_Xiphotheca_guthriei
Fabaceae_Xiphotheca_elliptica
Fabaceae_Amphithalea_villosa
Fabaceae_Amphithalea_vlokii
Fabaceae_Amphithalea_parvifolia
Fabaceae_Coelidium_villosum
Fabaceae_Amphithalea_rostrata
Fabaceae_Amphithalea_tomentosa
Fabaceae_Amphithalea_alba
Fabaceae_Amphithalea_stokoei
Fabaceae_Amphithalea_violacea
Fabaceae_Amphithalea_intermedia
Fabaceae_Amphithalea_speciosa
Fabaceae_Amphithalea_ericifolia
Fabaceae_Amphithalea_oppositifolia
Fabaceae_Amphithalea_imbricata
Fabaceae_Amphithalea_virgata
Fabaceae_Amphithalea_axillaris
Fabaceae_Amphithalea_biovulata
Fabaceae_Amphithalea_phylicoides
Fabaceae_Amphithalea_fourcadei
Fabaceae_Amphithalea_cuneifolia
Fabaceae_Cyclopia_glabra
Fabaceae_Cyclopia_bolusii
Fabaceae_Cyclopia_meyeriana
Fabaceae_Cyclopia_falcata
Fabaceae_Cyclopia_aurescens
Fabaceae_Cyclopia_longifolia
Fabaceae_Cyclopia_subternata
Fabaceae_Cyclopia_burtonii
Fabaceae_Cyclopia_pubescens

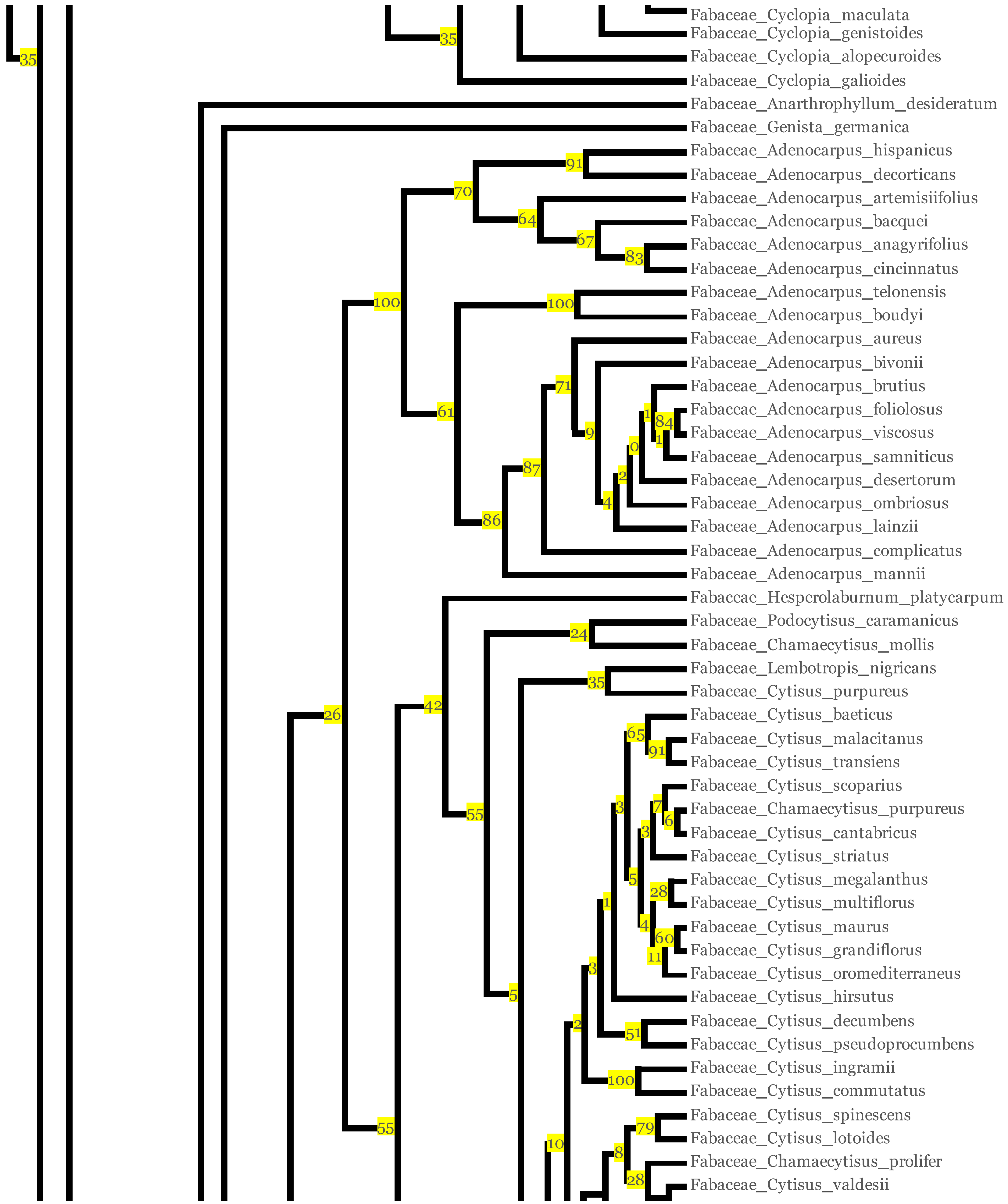
Fabaceae_Cyclopia_maculata
Fabaceae_Cyclopia_genistoides
Fabaceae_Cyclopia_alopecuroides
Fabaceae_Cyclopia_galioides
Fabaceae_Anarthrophyllum_desideratum
Fabaceae_Genista_germanica
Fabaceae_Adenocarpus_hispanicus
Fabaceae_Adenocarpus_decorticans
Fabaceae_Adenocarpus_artemisiifolius
Fabaceae_Adenocarpus_bacquei
Fabaceae_Adenocarpus_anagyrifolius
Fabaceae_Adenocarpus_cincinnatus
Fabaceae_Adenocarpus_telonensis
Fabaceae_Adenocarpus_boudyi
Fabaceae_Adenocarpus_aureus
Fabaceae_Adenocarpus_bivonii
Fabaceae_Adenocarpus_brutius
Fabaceae_Adenocarpus_foliolosus
Fabaceae_Adenocarpus_viscosus
Fabaceae_Adenocarpus_samniticus
Fabaceae_Adenocarpus_desertorum
Fabaceae_Adenocarpus_ombriosus
Fabaceae_Adenocarpus_lainzii
Fabaceae_Adenocarpus_complicatus
Fabaceae_Adenocarpus_mannii
Fabaceae_Hesperolaburnum_platycarpum
Fabaceae_Podocytisus_caramanicus
Fabaceae_Chamaecytisus_mollis
Fabaceae_Lembotropis_nigricans
Fabaceae_Cytisus_purpureus
Fabaceae_Cytisus_baeticus
Fabaceae_Cytisus_malacitanus
Fabaceae_Cytisus_transiens
Fabaceae_Cytisus_scoparius
Fabaceae_Chamaecytisus_purpureus
Fabaceae_Cytisus_cantabricus
Fabaceae_Cytisus_striatus
Fabaceae_Cytisus_megalanthus
Fabaceae_Cytisus_multiflorus
Fabaceae_Cytisus_maurus
Fabaceae_Cytisus_grandiflorus
Fabaceae_Cytisus_oromediterraneus
Fabaceae_Cytisus_hirsutus
Fabaceae_Cytisus_decumbens
Fabaceae_Cytisus_pseudoprocumbens
Fabaceae_Cytisus_ingramii
Fabaceae_Cytisus_commutatus
Fabaceae_Cytisus_spinescens
Fabaceae_Cytisus_lotoides
Fabaceae_Chamaecytisus_prolifer
Fabaceae_Cytisus_valdesii

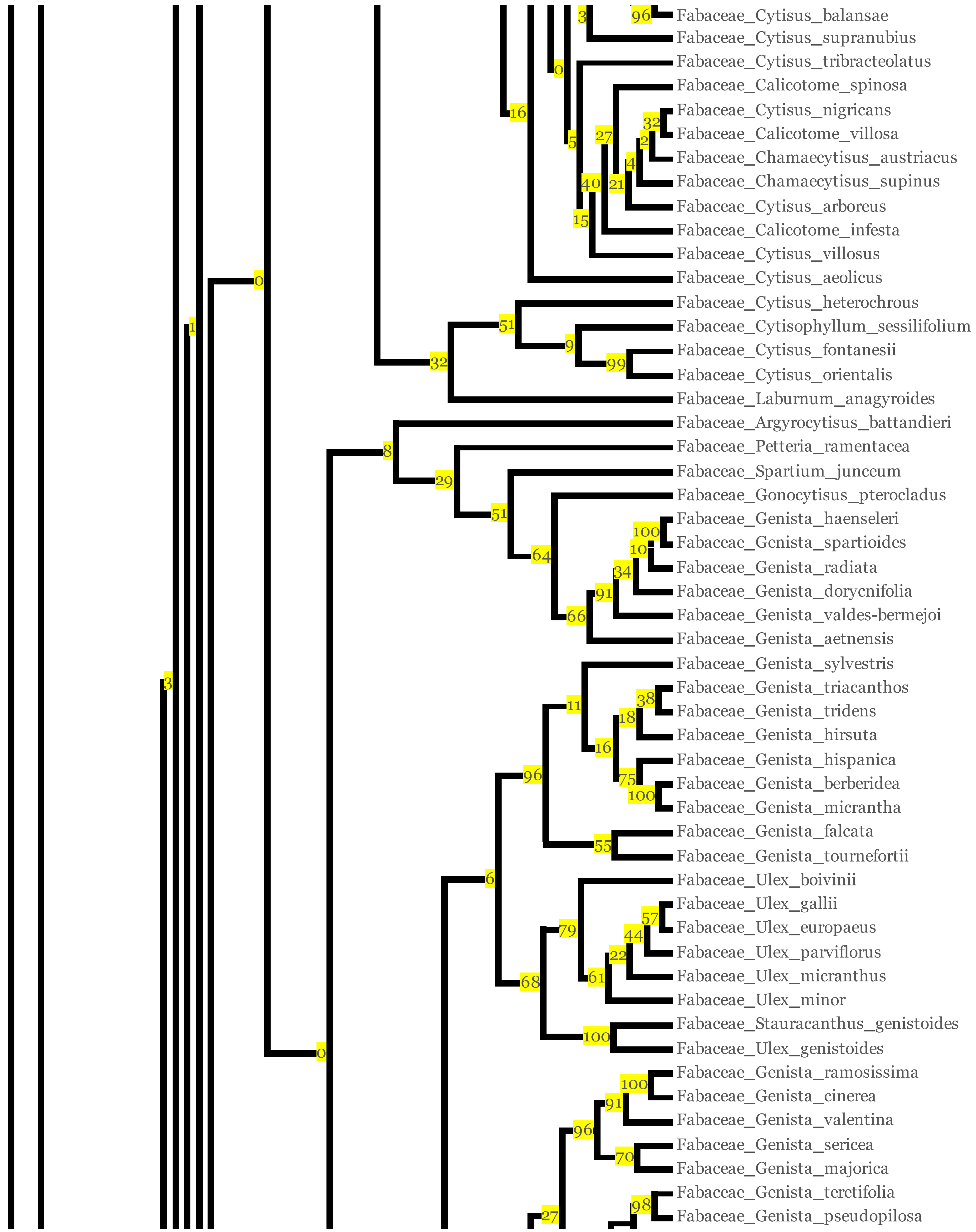
Fabaceae_Cytisus_balansae
Fabaceae_Cytisus_supranubius
Fabaceae_Cytisus_tribracteolatus
Fabaceae_Calicotome_spinosa
Fabaceae_Cytisus_nigricans
Fabaceae_Calicotome_villosa
Fabaceae_Chamaecytisus_austriacus
Fabaceae_Chamaecytisus_supinus
Fabaceae_Cytisus_arboreus
Fabaceae_Calicotome_infesta
Fabaceae_Cytisus_villosus
Fabaceae_Cytisus_aeolicus
Fabaceae_Cytisus_heterochrous
Fabaceae_Cytisophyllum_sessilifolium
Fabaceae_Cytisus_fontanesii
Fabaceae_Cytisus_orientalis
Fabaceae_Laburnum_anagyroides
Fabaceae_Argyrocytisus_battandieri
Fabaceae_Petteria_ramentacea
Fabaceae_Spartium_junceum
Fabaceae_Gonocytisus_pterocladus
Fabaceae_Genista_haenseleri
Fabaceae_Genista_spartioides
Fabaceae_Genista_radiata
Fabaceae_Genista_dorycnifolia
Fabaceae_Genista_valdes-bermejoi
Fabaceae_Genista_aetnensis
Fabaceae_Genista_sylvestris
Fabaceae_Genista_triacanthos
Fabaceae_Genista_tridens
Fabaceae_Genista_hirsuta
Fabaceae_Genista_hispanica
Fabaceae_Genista_berberidea
Fabaceae_Genista_micrantha
Fabaceae_Genista_falcata
Fabaceae_Genista_tournefortii
Fabaceae_Ulex_boivinii
Fabaceae_Ulex_gallii
Fabaceae_Ulex_europaeus
Fabaceae_Ulex_parviflorus
Fabaceae_Ulex_micranthus
Fabaceae_Ulex_minor
Fabaceae_Stauracanthus_genistoides
Fabaceae_Ulex_genistoides
Fabaceae_Genista_ramosissima
Fabaceae_Genista_cinerea
Fabaceae_Genista_valentina
Fabaceae_Genista_sericea
Fabaceae_Genista_majorica
Fabaceae_Genista_teretifolia
Fabaceae_Genista_pseudopilosa

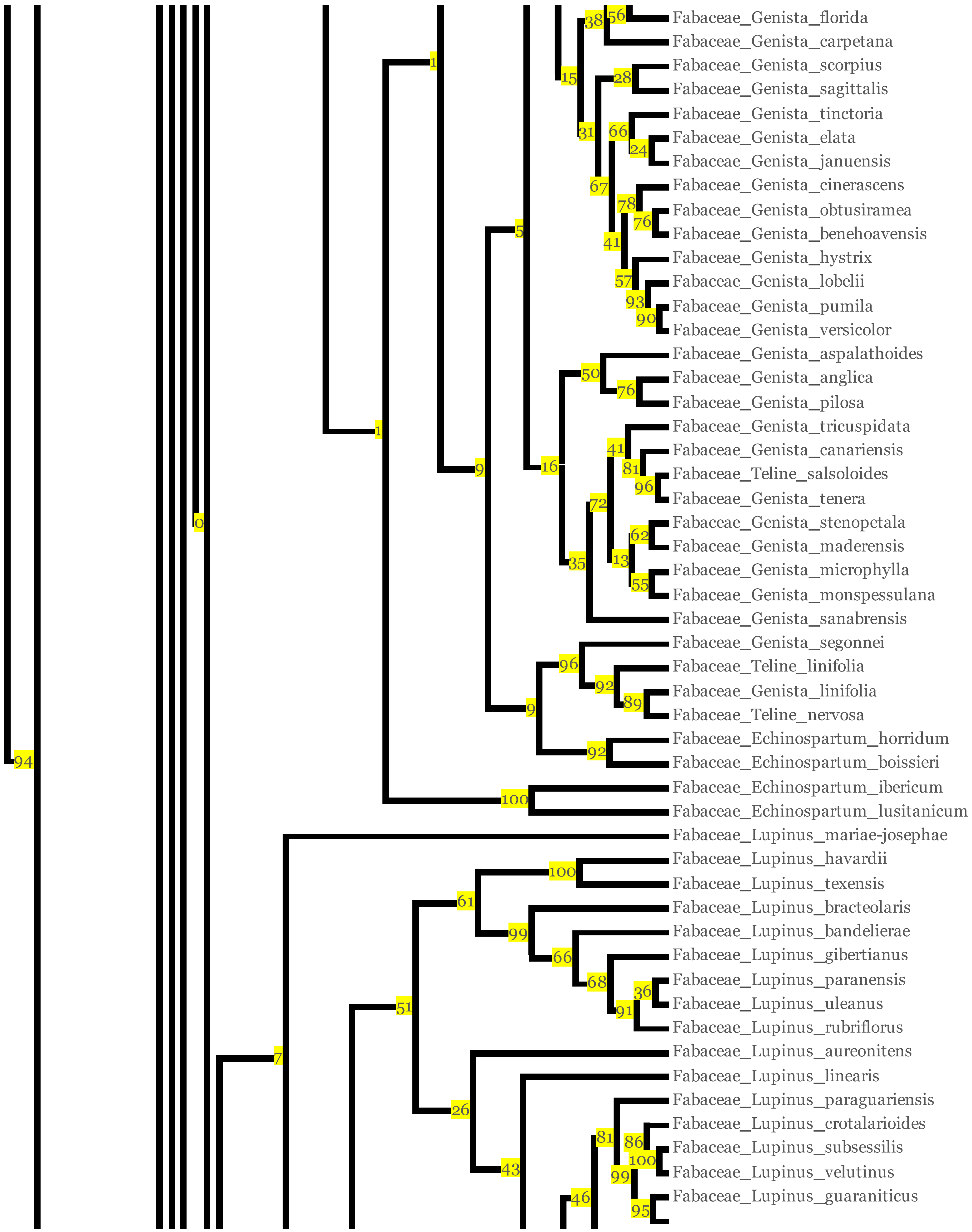
Fabaceae_Genista_florida
Fabaceae_Genista_carpetana
Fabaceae_Genista_scorpius
Fabaceae_Genista_sagittalis
Fabaceae_Genista_tinctoria
Fabaceae_Genista_elata
Fabaceae_Genista_januensis
Fabaceae_Genista_cinerascens
Fabaceae_Genista_obtusiramea
Fabaceae_Genista_benehoavensis
Fabaceae_Genista_hystrix
Fabaceae_Genista_lobelii
Fabaceae_Genista_pumila
Fabaceae_Genista_versicolor
Fabaceae_Genista_aspalathoides
Fabaceae_Genista_anglica
Fabaceae_Genista_pilosa
Fabaceae_Genista_tricuspidata
Fabaceae_Genista_canariensis
Fabaceae_Teline_salsoloides
Fabaceae_Genista_tenera
Fabaceae_Genista_stenopetala
Fabaceae_Genista_maderensis
Fabaceae_Genista_microphylla
Fabaceae_Genista_monspessulana
Fabaceae_Genista_sanabrensis
Fabaceae_Genista_segonnei
Fabaceae_Teline_linifolia
Fabaceae_Genista_linifolia
Fabaceae_Teline_nervosa
Fabaceae_Echinospartum_horridum
Fabaceae_Echinospartum_boissieri
Fabaceae_Echinospartum_ibericum
Fabaceae_Echinospartum_lusitanicum
Fabaceae_Lupinus_mariae-josephae
Fabaceae_Lupinus_havardii
Fabaceae_Lupinus_texensis
Fabaceae_Lupinus_bracteolaris
Fabaceae_Lupinus_bandelierae
Fabaceae_Lupinus_gibertianus
Fabaceae_Lupinus_paranensis
Fabaceae_Lupinus_uleanus
Fabaceae_Lupinus_rubriflorus
Fabaceae_Lupinus_aureonitens
Fabaceae_Lupinus_linearis
Fabaceae_Lupinus_paraguariensis
Fabaceae_Lupinus_crotalarioides
Fabaceae_Lupinus_subsessilis
Fabaceae_Lupinus_velutinus
Fabaceae_Lupinus_guaraniticus

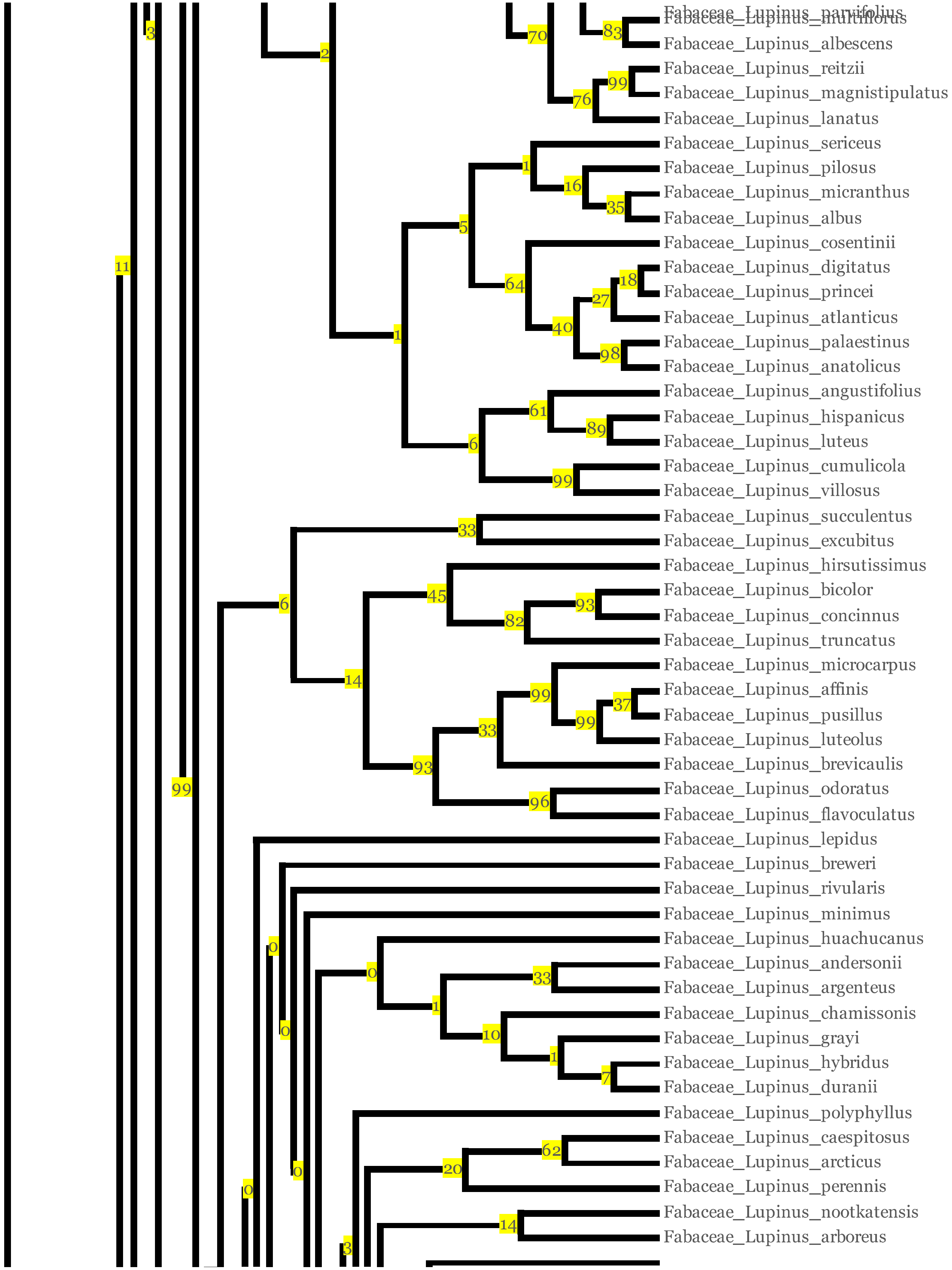
Fabaceae_Lupinus_parvifolius
Fabaceae_Lupinus_multiflorus
Fabaceae_Lupinus_albescens
Fabaceae_Lupinus_reitzii
Fabaceae_Lupinus_magnistipulatus
Fabaceae_Lupinus_lanatus
Fabaceae_Lupinus_sericeus
Fabaceae_Lupinus_pilosus
Fabaceae_Lupinus_micranthus
Fabaceae_Lupinus_albus
Fabaceae_Lupinus_cosentinii
Fabaceae_Lupinus_digitatus
Fabaceae_Lupinus_princei
Fabaceae_Lupinus_atlanticus
Fabaceae_Lupinus_palaestinus
Fabaceae_Lupinus_anatolicus
Fabaceae_Lupinus_angustifolius
Fabaceae_Lupinus_hispanicus
Fabaceae_Lupinus_luteus
Fabaceae_Lupinus_cumulicola
Fabaceae_Lupinus_villosus
Fabaceae_Lupinus_succulentus
Fabaceae_Lupinus_excubitus
Fabaceae_Lupinus_hirsutissimus
Fabaceae_Lupinus_bicolor
Fabaceae_Lupinus_concinnus
Fabaceae_Lupinus_truncatus
Fabaceae_Lupinus_microcarpus
Fabaceae_Lupinus_affinis
Fabaceae_Lupinus_pusillus
Fabaceae_Lupinus_luteolus
Fabaceae_Lupinus_brevicaulis
Fabaceae_Lupinus_odoratus
Fabaceae_Lupinus_flavoculatus
Fabaceae_Lupinus_lepidus
Fabaceae_Lupinus_breweri
Fabaceae_Lupinus_rivularis
Fabaceae_Lupinus_minimus
Fabaceae_Lupinus_huachucanus
Fabaceae_Lupinus_andersonii
Fabaceae_Lupinus_argenteus
Fabaceae_Lupinus_chamissonis
Fabaceae_Lupinus_grayi
Fabaceae_Lupinus_hybridus
Fabaceae_Lupinus_duranii
Fabaceae_Lupinus_polyphyllus
Fabaceae_Lupinus_caespitosus
Fabaceae_Lupinus_arcticus
Fabaceae_Lupinus_perennis
Fabaceae_Lupinus_nootkatensis
Fabaceae_Lupinus_arboreus

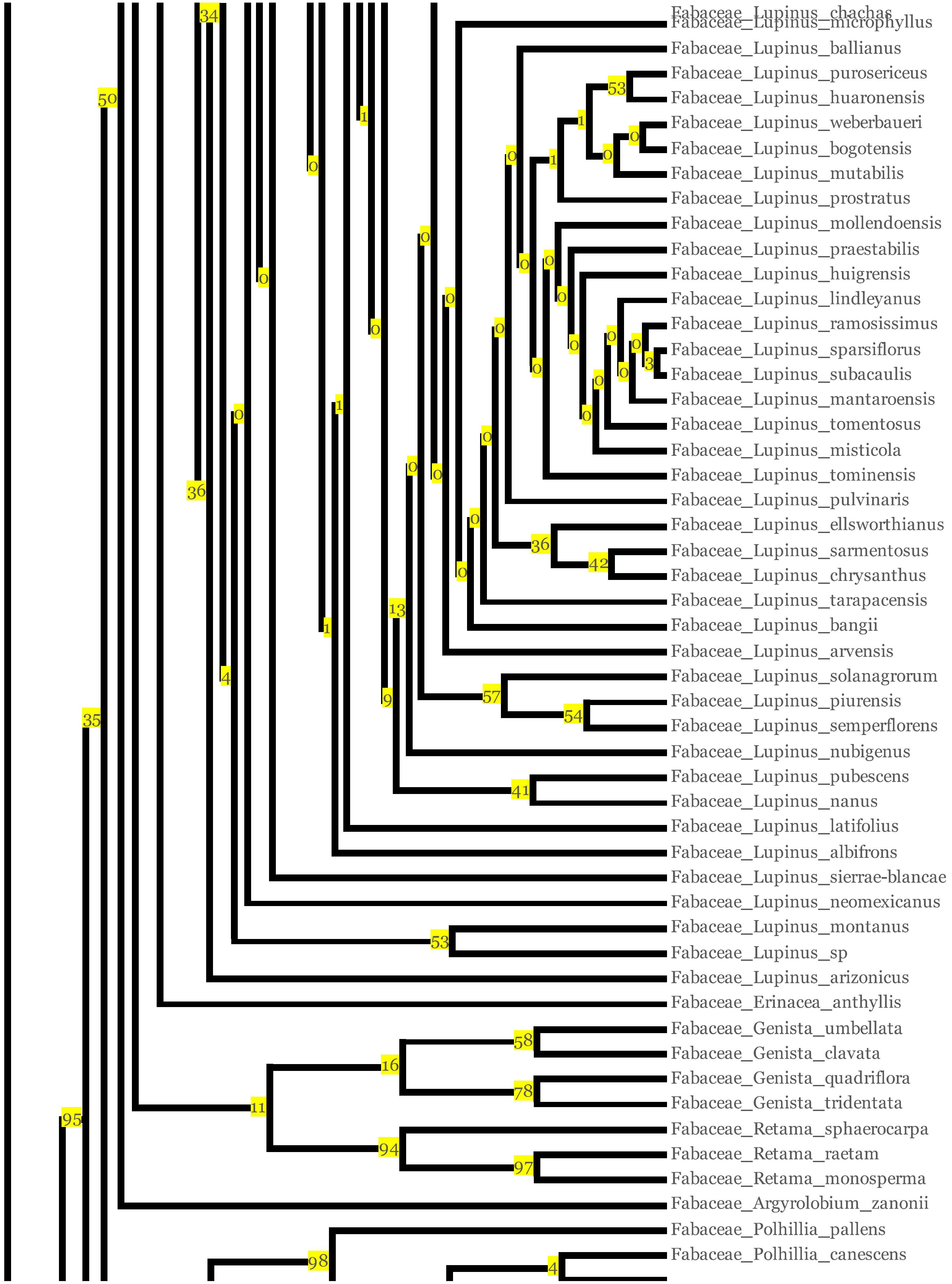
Fabaceae_Lupinus_chachas
Fabaceae_Lupinus_microphyllus
Fabaceae_Lupinus_ballianus
Fabaceae_Lupinus_purosericeus
Fabaceae_Lupinus_huaronensis
Fabaceae_Lupinus_weberbaueri
Fabaceae_Lupinus_bogotensis
Fabaceae_Lupinus_mutabilis
Fabaceae_Lupinus_prostratus
Fabaceae_Lupinus_mollendoensis
Fabaceae_Lupinus_praestabilis
Fabaceae_Lupinus_huigrensis
Fabaceae_Lupinus_lindleyanus
Fabaceae_Lupinus_ramosissimus
Fabaceae_Lupinus_sparsiflorus
Fabaceae_Lupinus_subacaulis
Fabaceae_Lupinus_mantaroensis
Fabaceae_Lupinus_tomentosus
Fabaceae_Lupinus_misticola
Fabaceae_Lupinus_tominensis
Fabaceae_Lupinus_pulvinaris
Fabaceae_Lupinus_ellsworthianus
Fabaceae_Lupinus_sarmentosus
Fabaceae_Lupinus_chrysanthus
Fabaceae_Lupinus_tarapacensis
Fabaceae_Lupinus_bangii
Fabaceae_Lupinus_arvensis
Fabaceae_Lupinus_solanagrorum
Fabaceae_Lupinus_piurensis
Fabaceae_Lupinus_semperflorens
Fabaceae_Lupinus_nubigenus
Fabaceae_Lupinus_pubescens
Fabaceae_Lupinus_nanus
Fabaceae_Lupinus_latifolius
Fabaceae_Lupinus_albifrons
Fabaceae_Lupinus_sierrae-blancae
Fabaceae_Lupinus_neomexicanus
Fabaceae_Lupinus_montanus
Fabaceae_Lupinus_sp
Fabaceae_Lupinus_arizonicus
Fabaceae_Erinacea_anthyllis
Fabaceae_Genista_umbellata
Fabaceae_Genista_clavata
Fabaceae_Genista_quadriflora
Fabaceae_Genista_tridentata
Fabaceae_Retama_sphaerocarpa
Fabaceae_Retama_raetam
Fabaceae_Retama_monosperma
Fabaceae_Argyrolobium_zanonii
Fabaceae_Polhillia_pallens
Fabaceae_Polhillia_canescens

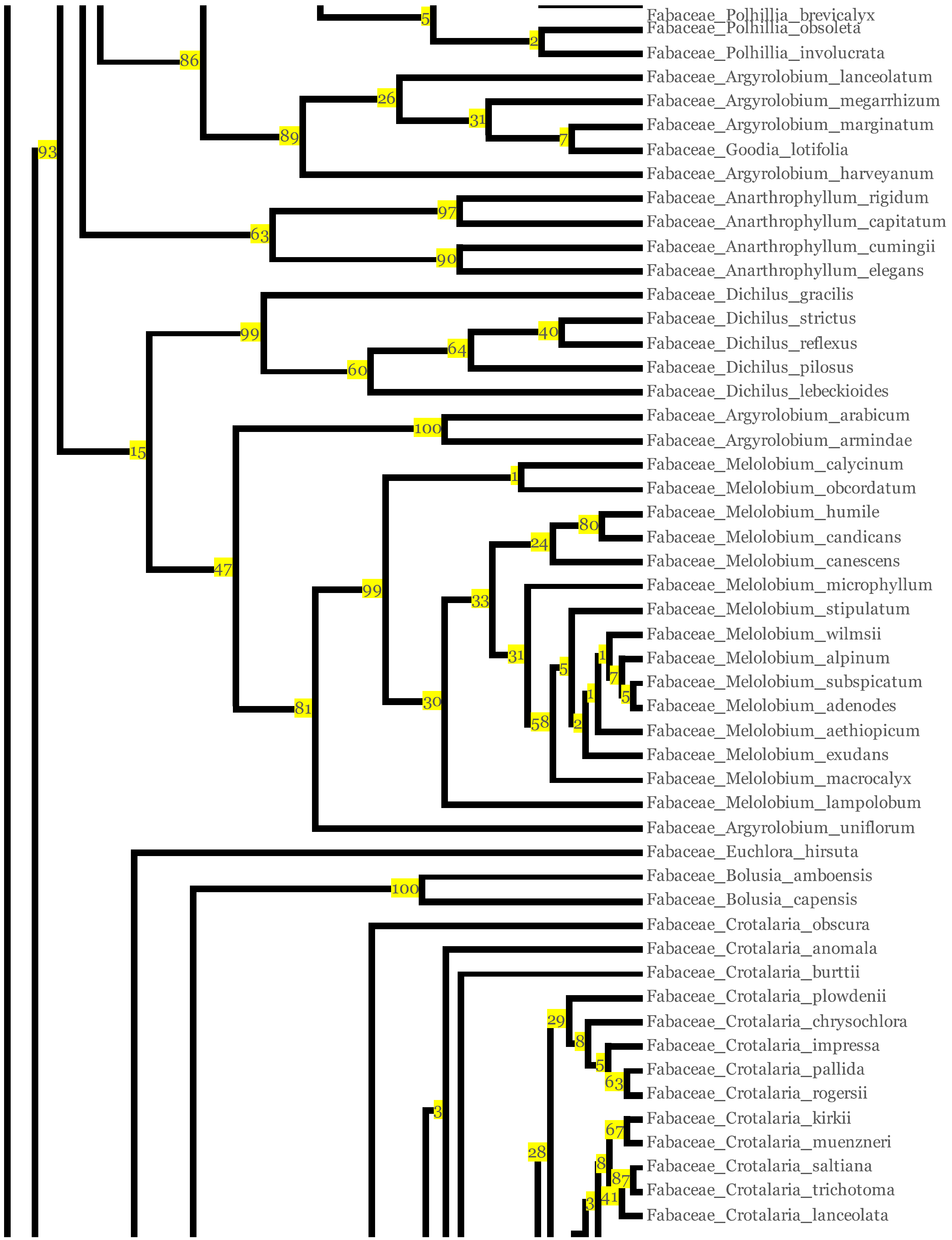
Fabaceae_Polhillia_brevicalyx
Fabaceae_Polhillia_obsoleta
Fabaceae_Polhillia_involucrata
Fabaceae_Argyrolobium_lanceolatum
Fabaceae_Argyrolobium_megarrhizum
Fabaceae_Argyrolobium_marginatum
Fabaceae_Goodia_lotifolia
Fabaceae_Argyrolobium_harveyanum
Fabaceae_Anarthrophyllum_rigidum
Fabaceae_Anarthrophyllum_capitatum
Fabaceae_Anarthrophyllum_cumingii
Fabaceae_Anarthrophyllum_elegans
Fabaceae_Dichilus_gracilis
Fabaceae_Dichilus_strictus
Fabaceae_Dichilus_reflexus
Fabaceae_Dichilus_pilosus
Fabaceae_Dichilus_lebeckioides
Fabaceae_Argyrolobium_arabicum
Fabaceae_Argyrolobium_armindae
Fabaceae_Melolobium_calycinum
Fabaceae_Melolobium_obcordatum
Fabaceae_Melolobium_humile
Fabaceae_Melolobium_candicans
Fabaceae_Melolobium_canescens
Fabaceae_Melolobium_microphyllum
Fabaceae_Melolobium_stipulatum
Fabaceae_Melolobium_wilmsii
Fabaceae_Melolobium_alpinum
Fabaceae_Melolobium_subspicatum
Fabaceae_Melolobium_adenodes
Fabaceae_Melolobium_aethiopicum
Fabaceae_Melolobium_exudans
Fabaceae_Melolobium_macrocalyx
Fabaceae_Melolobium_lampolobum
Fabaceae_Argyrolobium_uniflorum
Fabaceae_Euchlora_hirsuta
Fabaceae_Bolusia_amboensis
Fabaceae_Bolusia_capensis
Fabaceae_Crotalaria_obscura
Fabaceae_Crotalaria_anomala
Fabaceae_Crotalaria_burttii
Fabaceae_Crotalaria_plowdenii
Fabaceae_Crotalaria_chrysochlora
Fabaceae_Crotalaria_impressa
Fabaceae_Crotalaria_pallida
Fabaceae_Crotalaria_rogersii
Fabaceae_Crotalaria_kirkii
Fabaceae_Crotalaria_muenzneri
Fabaceae_Crotalaria_saltiana
Fabaceae_Crotalaria_trichotoma
Fabaceae_Crotalaria_lanceolata

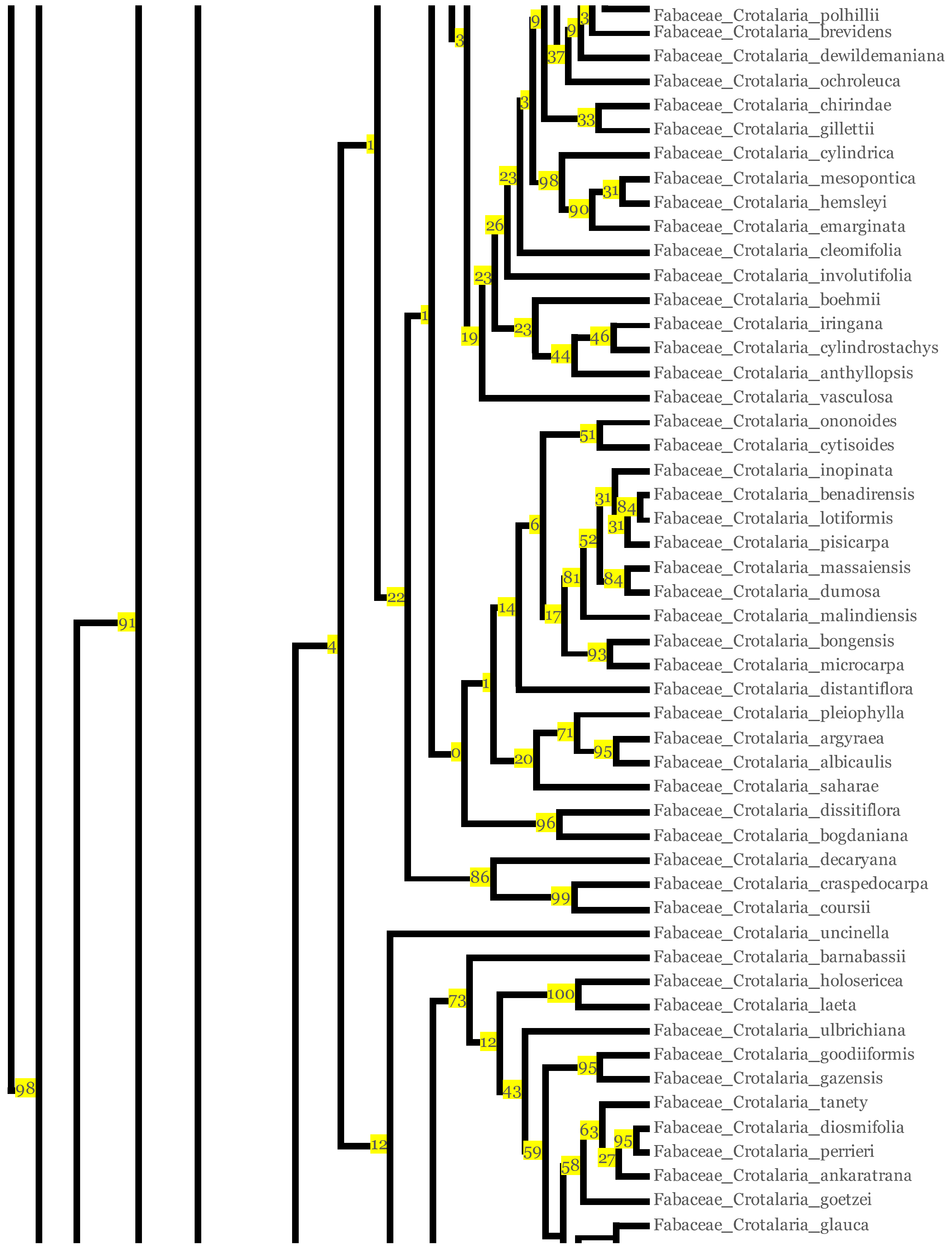
Fabaceae_Crotalaria_polhillii
Fabaceae_Crotalaria_brevidens
Fabaceae_Crotalaria_dewildemaniana
Fabaceae_Crotalaria_ochroleuca
Fabaceae_Crotalaria_chirindae
Fabaceae_Crotalaria_gillettii
Fabaceae_Crotalaria_cylindrica
Fabaceae_Crotalaria_mesopontica
Fabaceae_Crotalaria_hemsleyi
Fabaceae_Crotalaria_emarginata
Fabaceae_Crotalaria_cleomifolia
Fabaceae_Crotalaria_involutifolia
Fabaceae_Crotalaria_boehmii
Fabaceae_Crotalaria_iringana
Fabaceae_Crotalaria_cylindrostachys
Fabaceae_Crotalaria_anthyllopsis
Fabaceae_Crotalaria_vasculosa
Fabaceae_Crotalaria_ononoides
Fabaceae_Crotalaria_cytisoides
Fabaceae_Crotalaria_inopinata
Fabaceae_Crotalaria_benadirensis
Fabaceae_Crotalaria_lotiformis
Fabaceae_Crotalaria_pisicarpa
Fabaceae_Crotalaria_massaiensis
Fabaceae_Crotalaria_dumosa
Fabaceae_Crotalaria_malindiensis
Fabaceae_Crotalaria_bongensis
Fabaceae_Crotalaria_microcarpa
Fabaceae_Crotalaria_distantiflora
Fabaceae_Crotalaria_pleiophylla
Fabaceae_Crotalaria_argyraea
Fabaceae_Crotalaria_albicaulis
Fabaceae_Crotalaria_saharae
Fabaceae_Crotalaria_dissitiflora
Fabaceae_Crotalaria_bogdaniana
Fabaceae_Crotalaria_decaryana
Fabaceae_Crotalaria_craspedocarpa
Fabaceae_Crotalaria_coursii
Fabaceae_Crotalaria_uncinella
Fabaceae_Crotalaria_barnabassii
Fabaceae_Crotalaria_holosericea
Fabaceae_Crotalaria_laeta
Fabaceae_Crotalaria_ulbrichiana
Fabaceae_Crotalaria_goodiiformis
Fabaceae_Crotalaria_gazensis
Fabaceae_Crotalaria_tanety
Fabaceae_Crotalaria_diosmifolia
Fabaceae_Crotalaria_perrieri
Fabaceae_Crotalaria_ankaratrana
Fabaceae_Crotalaria_goetzei
Fabaceae_Crotalaria_glauca

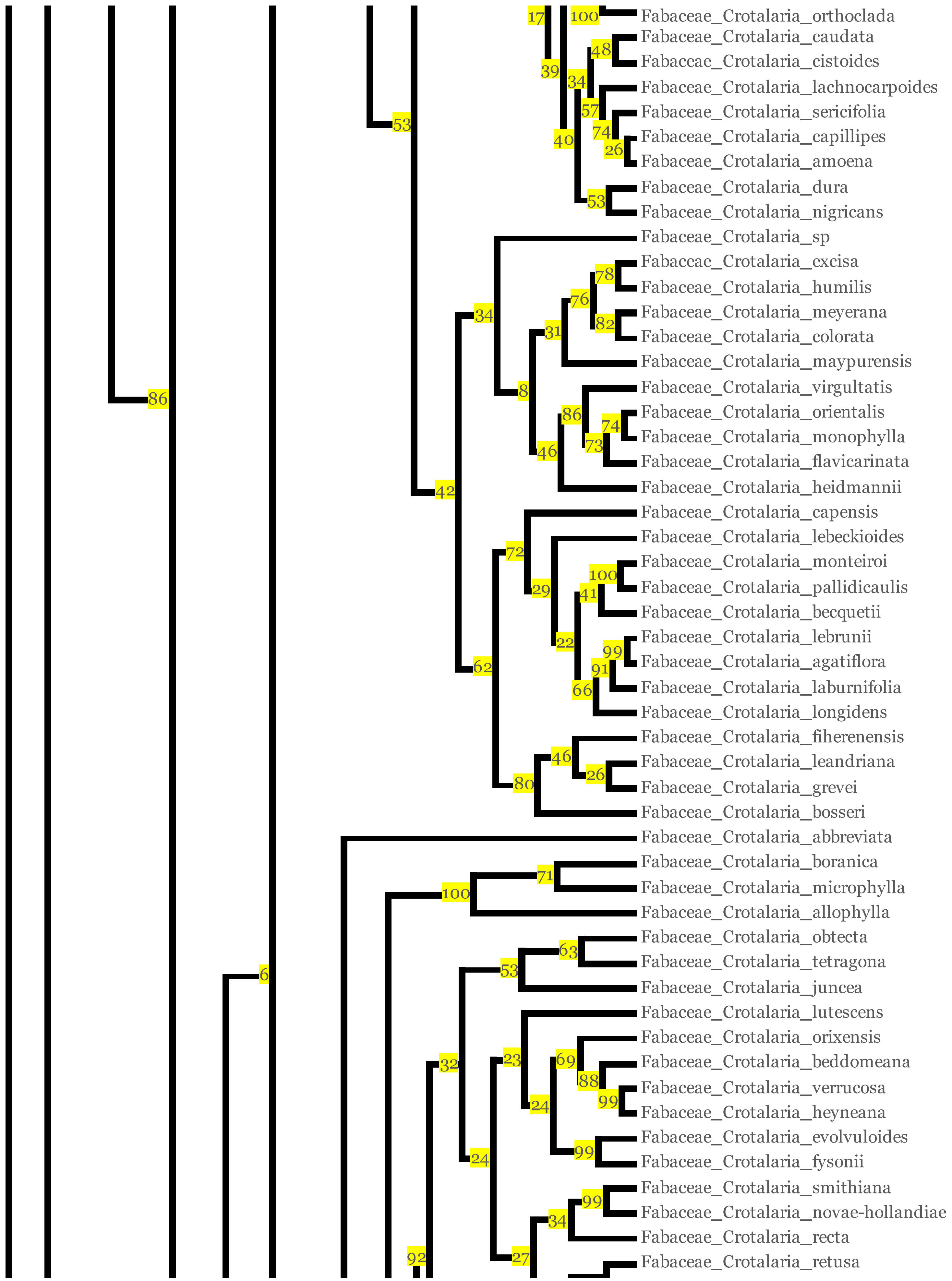
Fabaceae_Crotalaria_orthoclada
Fabaceae_Crotalaria_caudata
Fabaceae_Crotalaria_cistoides
Fabaceae_Crotalaria_lachnocarpoides
Fabaceae_Crotalaria_sericifolia
Fabaceae_Crotalaria_capillipes
Fabaceae_Crotalaria_amoena
Fabaceae_Crotalaria_dura
Fabaceae_Crotalaria_nigricans
Fabaceae_Crotalaria_sp
Fabaceae_Crotalaria_excisa
Fabaceae_Crotalaria_humilis
Fabaceae_Crotalaria_meyerana
Fabaceae_Crotalaria_colorata
Fabaceae_Crotalaria_maypurensis
Fabaceae_Crotalaria_virgultatis
Fabaceae_Crotalaria_orientalis
Fabaceae_Crotalaria_monophylla
Fabaceae_Crotalaria_flavicarinata
Fabaceae_Crotalaria_heidmannii
Fabaceae_Crotalaria_capensis
Fabaceae_Crotalaria_lebeckioides
Fabaceae_Crotalaria_monteiroi
Fabaceae_Crotalaria_pallidicaulis
Fabaceae_Crotalaria_becquetii
Fabaceae_Crotalaria_lebrunii
Fabaceae_Crotalaria_agatiflora
Fabaceae_Crotalaria_laburnifolia
Fabaceae_Crotalaria_longidens
Fabaceae_Crotalaria_fiherenensis
Fabaceae_Crotalaria_leandriana
Fabaceae_Crotalaria_grevei
Fabaceae_Crotalaria_bosseri
Fabaceae_Crotalaria_abbreviata
Fabaceae_Crotalaria_boranica
Fabaceae_Crotalaria_microphylla
Fabaceae_Crotalaria_allophylla
Fabaceae_Crotalaria_obtecta
Fabaceae_Crotalaria_tetragona
Fabaceae_Crotalaria_juncea
Fabaceae_Crotalaria_lutescens
Fabaceae_Crotalaria_orixensis
Fabaceae_Crotalaria_beddomeana
Fabaceae_Crotalaria_verrucosa
Fabaceae_Crotalaria_heyneana
Fabaceae_Crotalaria_evolvuloides
Fabaceae_Crotalaria_fysonii
Fabaceae_Crotalaria_smithiana
Fabaceae_Crotalaria_novae-hollandiae
Fabaceae_Crotalaria_recta
Fabaceae_Crotalaria_retusa

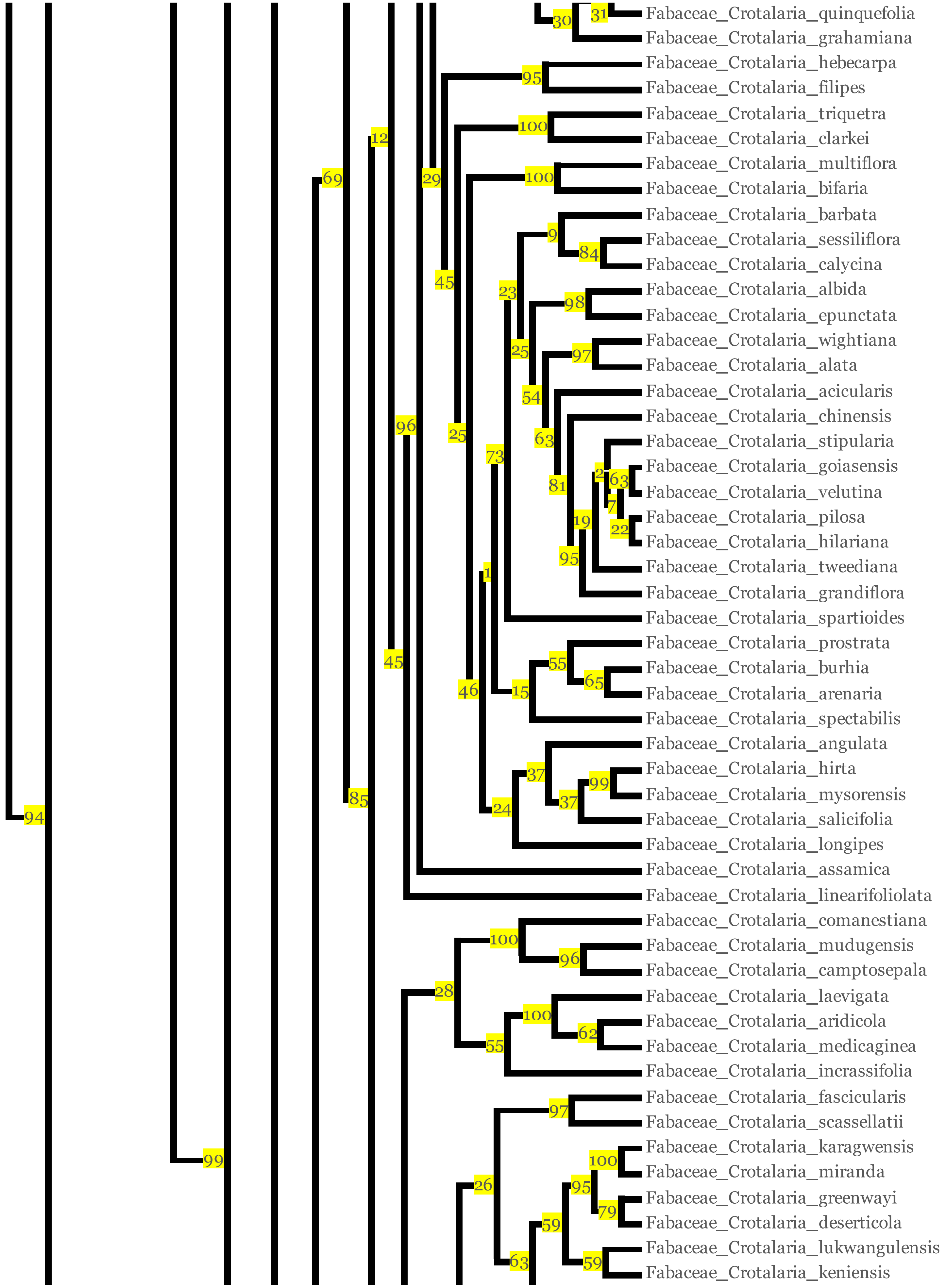
Fabaceae_Crotalaria_quinquefolia
Fabaceae_Crotalaria_grahamiana
Fabaceae_Crotalaria_hebecarpa
Fabaceae_Crotalaria_filipes
Fabaceae_Crotalaria_triquetra
Fabaceae_Crotalaria_clarkei
Fabaceae_Crotalaria_multiflora
Fabaceae_Crotalaria_bifaria
Fabaceae_Crotalaria_barbata
Fabaceae_Crotalaria_sessiliflora
Fabaceae_Crotalaria_calycina
Fabaceae_Crotalaria_albida
Fabaceae_Crotalaria_epunctata
Fabaceae_Crotalaria_wightiana
Fabaceae_Crotalaria_alata
Fabaceae_Crotalaria_acicularis
Fabaceae_Crotalaria_chinensis
Fabaceae_Crotalaria_stipularia
Fabaceae_Crotalaria_goiasensis
Fabaceae_Crotalaria_velutina
Fabaceae_Crotalaria_pilosa
Fabaceae_Crotalaria_hilariana
Fabaceae_Crotalaria_tweediana
Fabaceae_Crotalaria_grandiflora
Fabaceae_Crotalaria_spartioides
Fabaceae_Crotalaria_prostrata
Fabaceae_Crotalaria_burhia
Fabaceae_Crotalaria_arenaria
Fabaceae_Crotalaria_spectabilis
Fabaceae_Crotalaria_angulata
Fabaceae_Crotalaria_hirta
Fabaceae_Crotalaria_mysorensis
Fabaceae_Crotalaria_salicifolia
Fabaceae_Crotalaria_longipes
Fabaceae_Crotalaria_assamica
Fabaceae_Crotalaria_linearifoliolata
Fabaceae_Crotalaria_comanestiana
Fabaceae_Crotalaria_mudugensis
Fabaceae_Crotalaria_camptosepala
Fabaceae_Crotalaria_laevigata
Fabaceae_Crotalaria_aridicola
Fabaceae_Crotalaria_medicaginea
Fabaceae_Crotalaria_incrassifolia
Fabaceae_Crotalaria_fascicularis
Fabaceae_Crotalaria_scassellatii
Fabaceae_Crotalaria_karagwensis
Fabaceae_Crotalaria_miranda
Fabaceae_Crotalaria_greenwayi
Fabaceae_Crotalaria_deserticola
Fabaceae_Crotalaria_lukwangulensis
Fabaceae_Crotalaria_keniensis

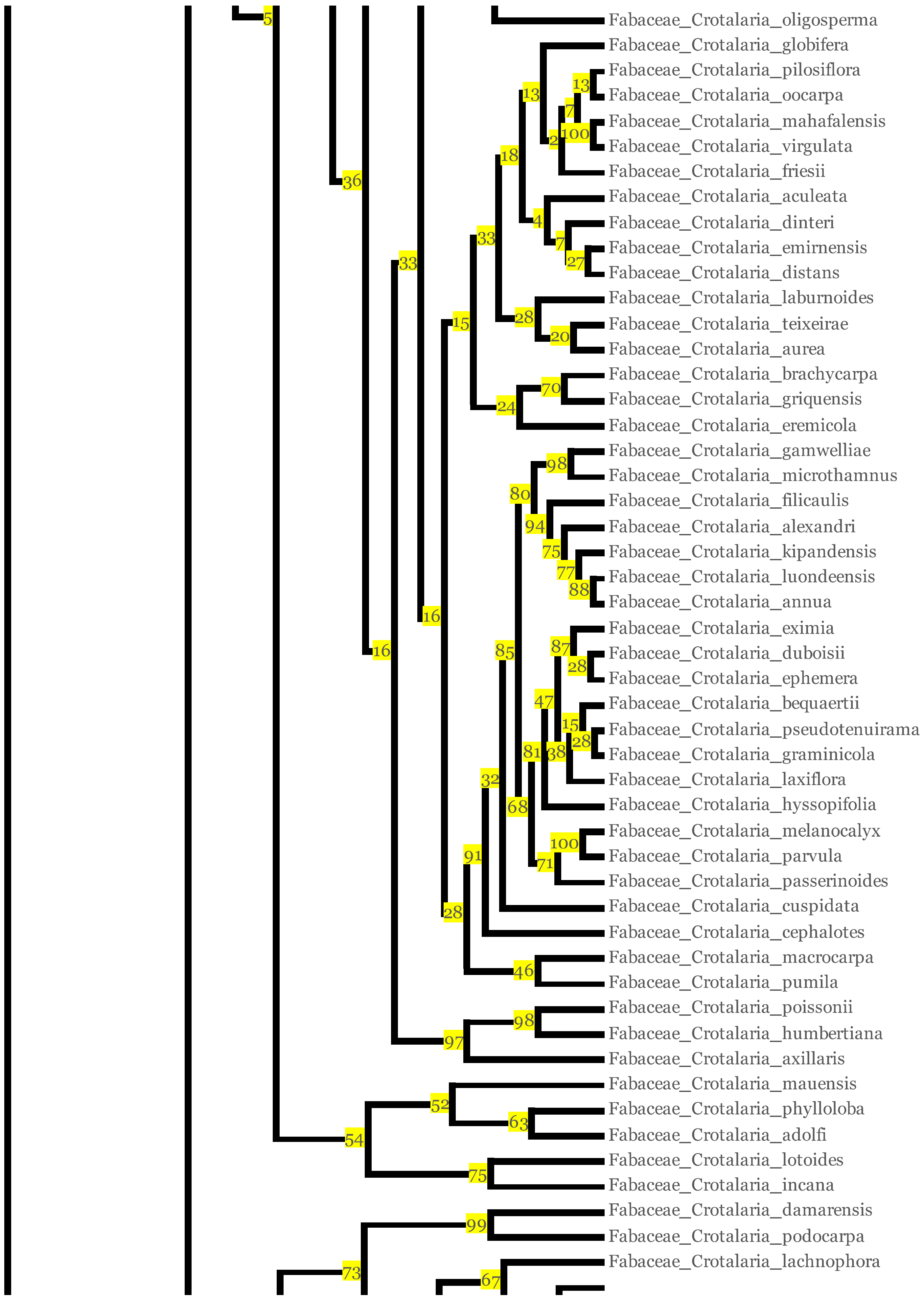
Fabaceae_Crotalaria_oligosperma
Fabaceae_Crotalaria_globifera
Fabaceae_Crotalaria_pilosiflora
Fabaceae_Crotalaria_oocarpa
Fabaceae_Crotalaria_mahafalensis
Fabaceae_Crotalaria_virgulata
Fabaceae_Crotalaria_friesii
Fabaceae_Crotalaria_aculeata
Fabaceae_Crotalaria_dinteri
Fabaceae_Crotalaria_emirnensis
Fabaceae_Crotalaria_distans
Fabaceae_Crotalaria_laburnoides
Fabaceae_Crotalaria_teixeirae
Fabaceae_Crotalaria_aurea
Fabaceae_Crotalaria_brachycarpa
Fabaceae_Crotalaria_griquensis
Fabaceae_Crotalaria_eremicola
Fabaceae_Crotalaria_gamwelliae
Fabaceae_Crotalaria_microthamnus
Fabaceae_Crotalaria_filicaulis
Fabaceae_Crotalaria_alexandri
Fabaceae_Crotalaria_kipandensis
Fabaceae_Crotalaria_luondeensis
Fabaceae_Crotalaria_annua
Fabaceae_Crotalaria_eximia
Fabaceae_Crotalaria_duboisii
Fabaceae_Crotalaria_ephemera
Fabaceae_Crotalaria_bequaertii
Fabaceae_Crotalaria_pseudotenuirama
Fabaceae_Crotalaria_graminicola
Fabaceae_Crotalaria_laxiflora
Fabaceae_Crotalaria_hyssopifolia
Fabaceae_Crotalaria_melanocalyx
Fabaceae_Crotalaria_parvula
Fabaceae_Crotalaria_passerinoides
Fabaceae_Crotalaria_cuspidata
Fabaceae_Crotalaria_cephalotes
Fabaceae_Crotalaria_macrocarpa
Fabaceae_Crotalaria_pumila
Fabaceae_Crotalaria_poissonii
Fabaceae_Crotalaria_humbertiana
Fabaceae_Crotalaria_axillaris
Fabaceae_Crotalaria_mauensis
Fabaceae_Crotalaria_phylloloba
Fabaceae_Crotalaria_adolfi
Fabaceae_Crotalaria_lotoides
Fabaceae_Crotalaria_incana
Fabaceae_Crotalaria_damarensis
Fabaceae_Crotalaria_podocarpa
Fabaceae_Crotalaria_lachnophora

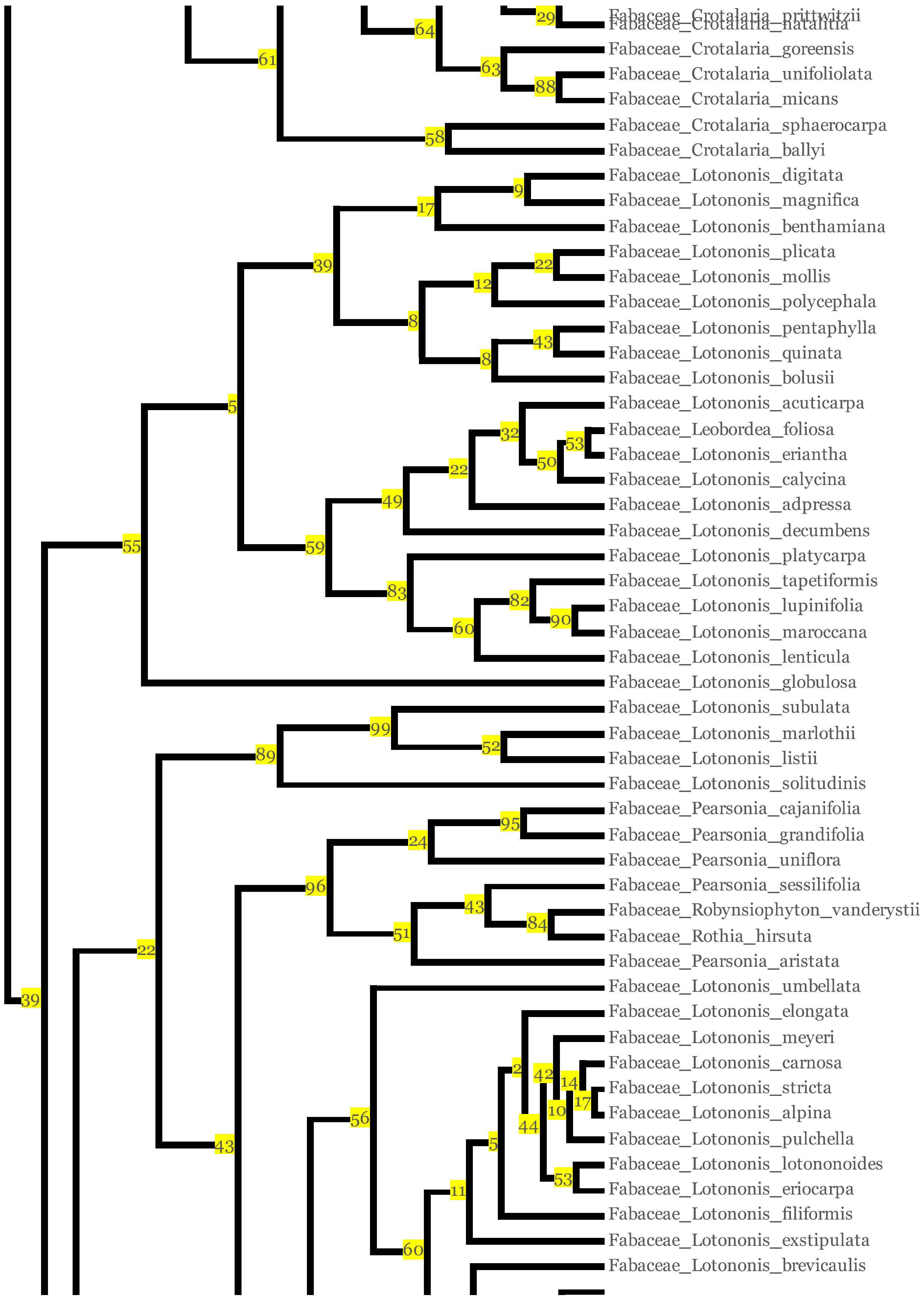
Fabaceae_Crotalaria_prittwitzii
Fabaceae_Crotalaria_natalitia
Fabaceae_Crotalaria_goreensis
Fabaceae_Crotalaria_unifoliolata
Fabaceae_Crotalaria_micans
Fabaceae_Crotalaria_sphaerocarpa
Fabaceae_Crotalaria_ballyi
Fabaceae_Lotononis_digitata
Fabaceae_Lotononis_magnifica
Fabaceae_Lotononis_benthamiana
Fabaceae_Lotononis_plicata
Fabaceae_Lotononis_mollis
Fabaceae_Lotononis_polycephala
Fabaceae_Lotononis_pentaphylla
Fabaceae_Lotononis_quinata
Fabaceae_Lotononis_bolusii
Fabaceae_Lotononis_acuticarpa
Fabaceae_Leobordea_foliosa
Fabaceae_Lotononis_eriantha
Fabaceae_Lotononis_calycina
Fabaceae_Lotononis_adpressa
Fabaceae_Lotononis_decumbens
Fabaceae_Lotononis_platycarpa
Fabaceae_Lotononis_tapetiformis
Fabaceae_Lotononis_lupinifolia
Fabaceae_Lotononis_maroccana
Fabaceae_Lotononis_lenticula
Fabaceae_Lotononis_globulosa
Fabaceae_Lotononis_subulata
Fabaceae_Lotononis_marlothii
Fabaceae_Lotononis_listii
Fabaceae_Lotononis_solitudinis
Fabaceae_Pearsonia_cajanifolia
Fabaceae_Pearsonia_grandifolia
Fabaceae_Pearsonia_uniflora
Fabaceae_Pearsonia_sessilifolia
Fabaceae_Robynsiophyton_vanderystii
Fabaceae_Rothia_hirsuta
Fabaceae_Pearsonia_aristata
Fabaceae_Lotononis_umbellata
Fabaceae_Lotononis_elongata
Fabaceae_Lotononis_meyeri
Fabaceae_Lotononis_carnosa
Fabaceae_Lotononis_stricta
Fabaceae_Lotononis_alpina
Fabaceae_Lotononis_pulchella
Fabaceae_Lotononis_lotononoides
Fabaceae_Lotononis_eriocarpa
Fabaceae_Lotononis_filiformis
Fabaceae_Lotononis_exstipulata
Fabaceae_Lotononis_brevicaulis

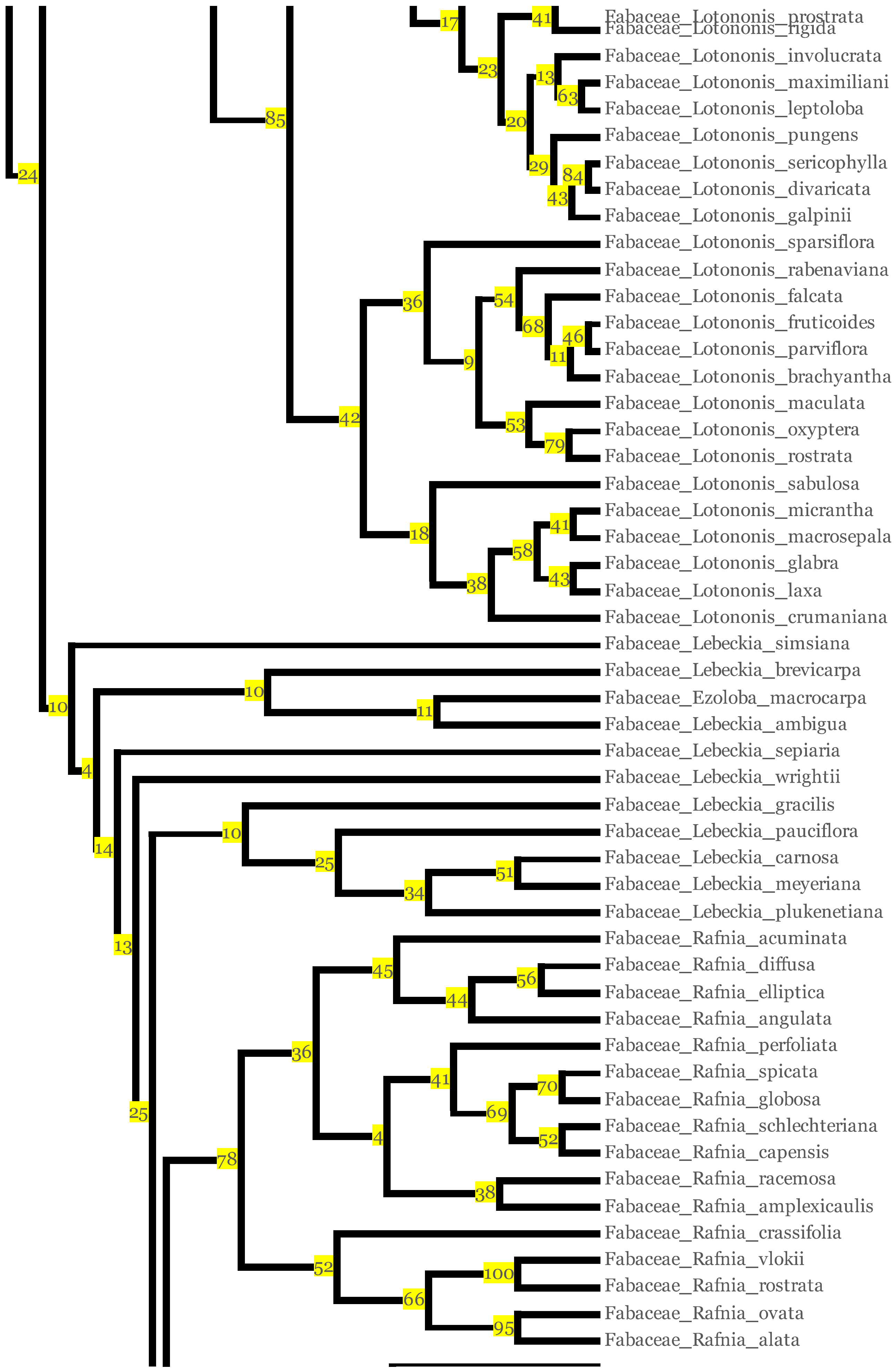
Fabaceae_Lotononis_prostrata
Fabaceae_Lotononis_rigida
Fabaceae_Lotononis_involucrata
Fabaceae_Lotononis_maximiliani
Fabaceae_Lotononis_leptoloba
Fabaceae_Lotononis_pungens
Fabaceae_Lotononis_sericophylla
Fabaceae_Lotononis_divaricata
Fabaceae_Lotononis_galpinii
Fabaceae_Lotononis_sparsiflora
Fabaceae_Lotononis_rabenaviana
Fabaceae_Lotononis_falcata
Fabaceae_Lotononis_fruticoides
Fabaceae_Lotononis_parviflora
Fabaceae_Lotononis_brachyantha
Fabaceae_Lotononis_maculata
Fabaceae_Lotononis_oxyptera
Fabaceae_Lotononis_rostrata
Fabaceae_Lotononis_sabulosa
Fabaceae_Lotononis_micrantha
Fabaceae_Lotononis_macrosepala
Fabaceae_Lotononis_glabra
Fabaceae_Lotononis_laxa
Fabaceae_Lotononis_crumaniana
Fabaceae_Lebeckia_simsiana
Fabaceae_Lebeckia_brevicarpa
Fabaceae_Ezoloba_macrocarpa
Fabaceae_Lebeckia_ambigua
Fabaceae_Lebeckia_sepiaria
Fabaceae_Lebeckia_wrightii
Fabaceae_Lebeckia_gracilis
Fabaceae_Lebeckia_pauciflora
Fabaceae_Lebeckia_carnosa
Fabaceae_Lebeckia_meyeriana
Fabaceae_Lebeckia_plukenetiana
Fabaceae_Rafnia_acuminata
Fabaceae_Rafnia_diffusa
Fabaceae_Rafnia_elliptica
Fabaceae_Rafnia_angulata
Fabaceae_Rafnia_perfoliata
Fabaceae_Rafnia_spicata
Fabaceae_Rafnia_globosa
Fabaceae_Rafnia_schlechteriana
Fabaceae_Rafnia_capensis
Fabaceae_Rafnia_racemosa
Fabaceae_Rafnia_amplexicaulis
Fabaceae_Rafnia_crassifolia
Fabaceae_Rafnia_vlokii
Fabaceae_Rafnia_rostrata
Fabaceae_Rafnia_ovata
Fabaceae_Rafnia_alata

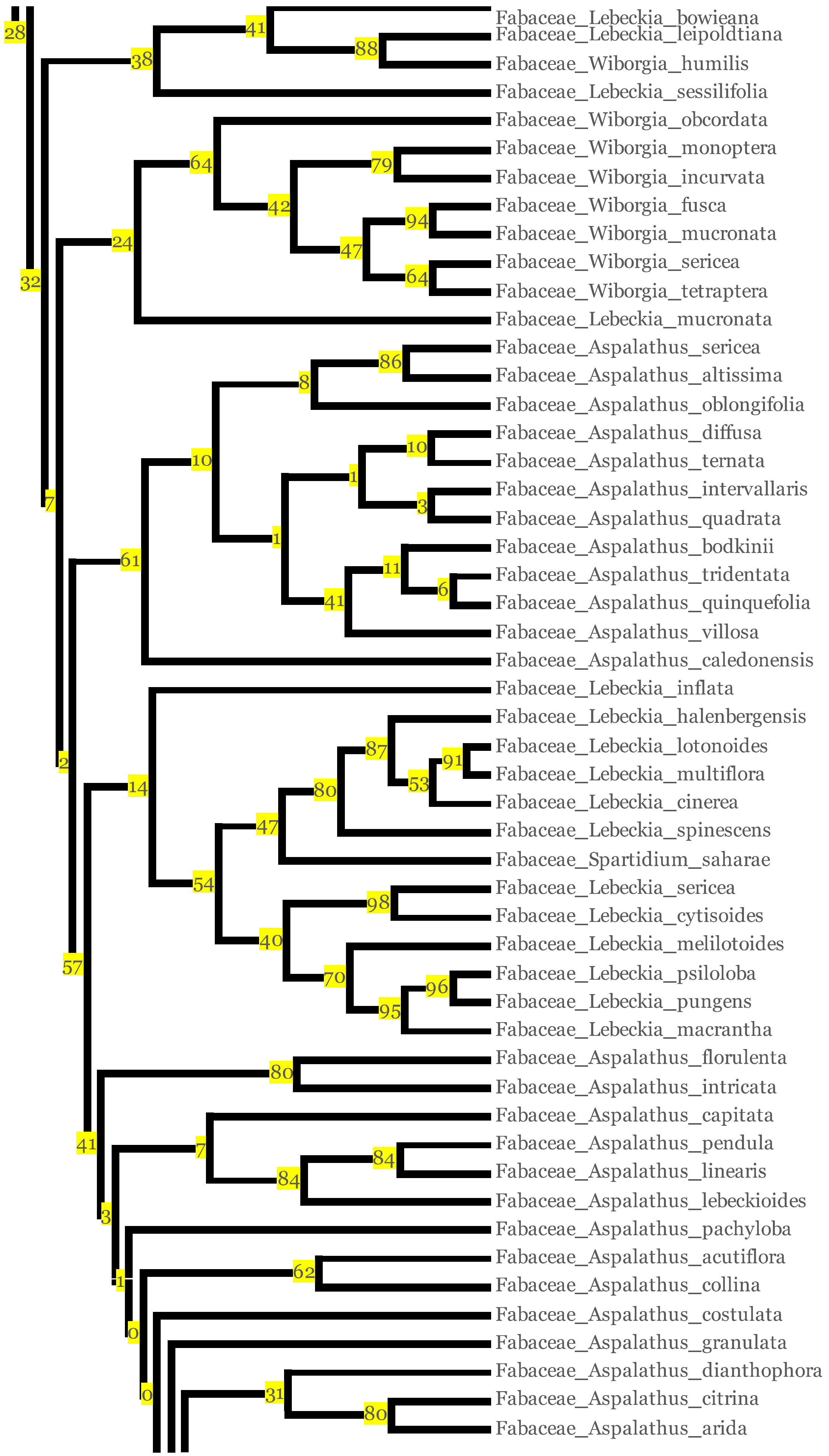
Fabaceae_Lebeckia_bowieana
Fabaceae_Lebeckia_leipoldtiana
Fabaceae_Wiborgia_humilis
Fabaceae_Lebeckia_sessilifolia
Fabaceae_Wiborgia_obcordata
Fabaceae_Wiborgia_monoptera
Fabaceae_Wiborgia_incurvata
Fabaceae_Wiborgia_fusca
Fabaceae_Wiborgia_mucronata
Fabaceae_Wiborgia_sericea
Fabaceae_Wiborgia_tetraptera
Fabaceae_Lebeckia_mucronata
Fabaceae_Aspalathus_sericea
Fabaceae_Aspalathus_altissima
Fabaceae_Aspalathus_oblongifolia
Fabaceae_Aspalathus_diffusa
Fabaceae_Aspalathus_ternata
Fabaceae_Aspalathus_intervallaris
Fabaceae_Aspalathus_quadrata
Fabaceae_Aspalathus_bodkinii
Fabaceae_Aspalathus_tridentata
Fabaceae_Aspalathus_quinquefolia
Fabaceae_Aspalathus_villosa
Fabaceae_Aspalathus_caledonensis
Fabaceae_Lebeckia_inflata
Fabaceae_Lebeckia_halenbergensis
Fabaceae_Lebeckia_lotonoides
Fabaceae_Lebeckia_multiflora
Fabaceae_Lebeckia_cinerea
Fabaceae_Lebeckia_spinescens
Fabaceae_Spartidium_saharae
Fabaceae_Lebeckia_sericea
Fabaceae_Lebeckia_cytisoides
Fabaceae_Lebeckia_melilotoides
Fabaceae_Lebeckia_psiloloba
Fabaceae_Lebeckia_pungens
Fabaceae_Lebeckia_macrantha
Fabaceae_Aspalathus_florulenta
Fabaceae_Aspalathus_intricata
Fabaceae_Aspalathus_capitata
Fabaceae_Aspalathus_pendula
Fabaceae_Aspalathus_linearis
Fabaceae_Aspalathus_lebeckioides
Fabaceae_Aspalathus_pachyloba
Fabaceae_Aspalathus_acutiflora
Fabaceae_Aspalathus_collina
Fabaceae_Aspalathus_costulata
Fabaceae_Aspalathus_granulata
Fabaceae_Aspalathus_dianthophora
Fabaceae_Aspalathus_citrina
Fabaceae_Aspalathus_arida
28
32
41
88
38
64
79
42
94
47
64
24
7
86
8
10
10
1
3
1
11
6
41
61
2
14
87
91
80
53
47
54
98
40
70
96
95
57
80
41
7
84
84
3
1
62
0
0
31
80

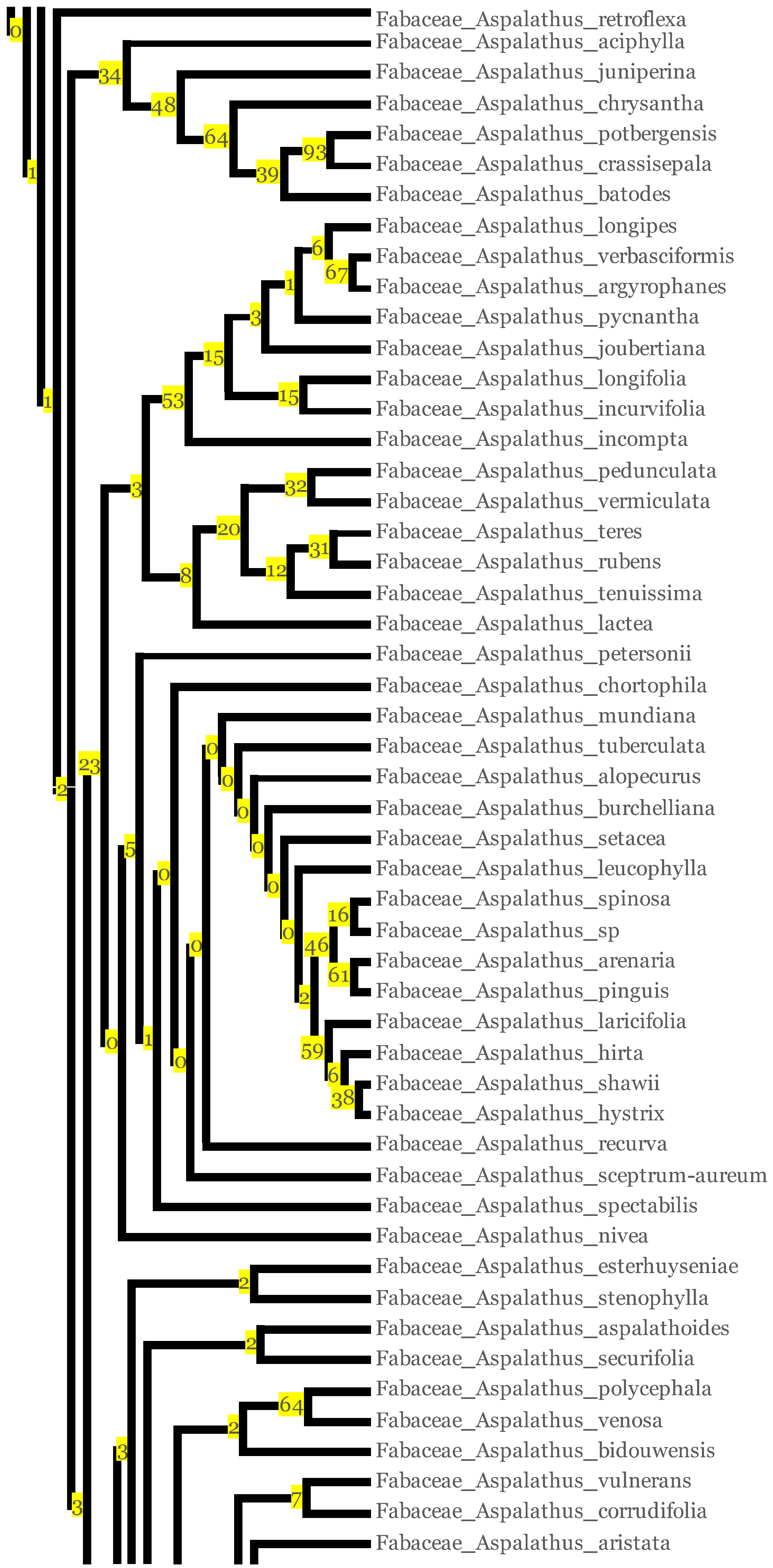
Fabaceae_Aspalathus_retroflexa
Fabaceae_Aspalathus_aciphylla
Fabaceae_Aspalathus_juniperina
Fabaceae_Aspalathus_chrysantha
Fabaceae_Aspalathus_potbergensis
Fabaceae_Aspalathus_crassisepala
Fabaceae_Aspalathus_batodes
Fabaceae_Aspalathus_longipes
Fabaceae_Aspalathus_verbasciformis
Fabaceae_Aspalathus_argyrophanes
Fabaceae_Aspalathus_pycnantha
Fabaceae_Aspalathus_joubertiana
Fabaceae_Aspalathus_longifolia
Fabaceae_Aspalathus_incurvifolia
Fabaceae_Aspalathus_incompta
Fabaceae_Aspalathus_pedunculata
Fabaceae_Aspalathus_vermiculata
Fabaceae_Aspalathus_teres
Fabaceae_Aspalathus_rubens
Fabaceae_Aspalathus_tenuissima
Fabaceae_Aspalathus_lactea
Fabaceae_Aspalathus_petersonii
Fabaceae_Aspalathus_chortophila
Fabaceae_Aspalathus_mundiana
Fabaceae_Aspalathus_tuberculata
Fabaceae_Aspalathus_alopecurus
Fabaceae_Aspalathus_burchelliana
Fabaceae_Aspalathus_setacea
Fabaceae_Aspalathus_leucophylla
Fabaceae_Aspalathus_spinosa
Fabaceae_Aspalathus_sp
Fabaceae_Aspalathus_arenaria
Fabaceae_Aspalathus_pinguis
Fabaceae_Aspalathus_laricifolia
Fabaceae_Aspalathus_hirta
Fabaceae_Aspalathus_shawii
Fabaceae_Aspalathus_hystrix
Fabaceae_Aspalathus_recurva
Fabaceae_Aspalathus_sceptrum-aureum
Fabaceae_Aspalathus_spectabilis
Fabaceae_Aspalathus_nivea
Fabaceae_Aspalathus_esterhuyseniae
Fabaceae_Aspalathus_stenophylla
Fabaceae_Aspalathus_aspalathoides
Fabaceae_Aspalathus_securifolia
Fabaceae_Aspalathus_polycephala
Fabaceae_Aspalathus_venosa
Fabaceae_Aspalathus_bidouwensis
Fabaceae_Aspalathus_vulnerans
Fabaceae_Aspalathus_corrudifolia
Fabaceae_Aspalathus_aristata

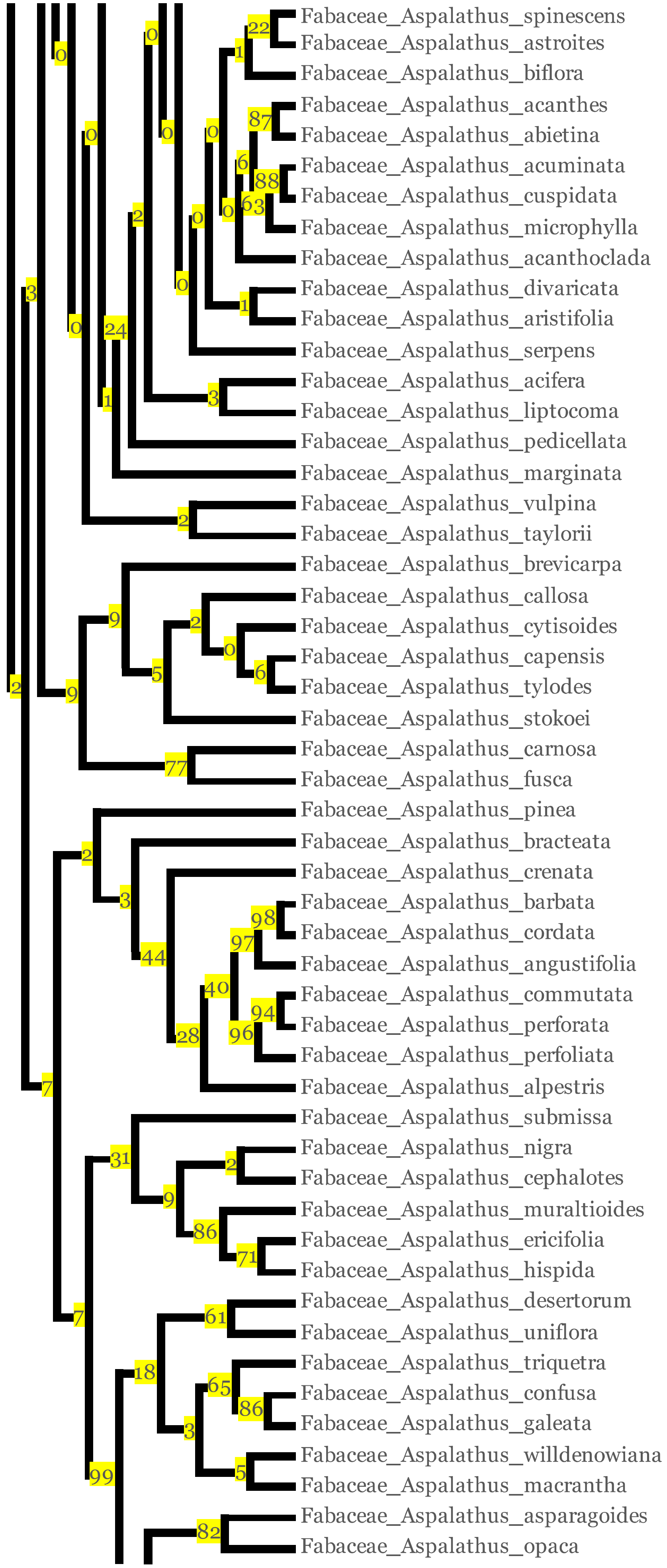
Fabaceae_Aspalathus_spinescens
Fabaceae_Aspalathus_astroites
Fabaceae_Aspalathus_biflora
Fabaceae_Aspalathus_acanthes
Fabaceae_Aspalathus_abietina
Fabaceae_Aspalathus_acuminata
Fabaceae_Aspalathus_cuspidata
Fabaceae_Aspalathus_microphylla
Fabaceae_Aspalathus_acanthoclada
Fabaceae_Aspalathus_divaricata
Fabaceae_Aspalathus_aristifolia
Fabaceae_Aspalathus_serpens
Fabaceae_Aspalathus_acifera
Fabaceae_Aspalathus_liptocoma
Fabaceae_Aspalathus_pedicellata
Fabaceae_Aspalathus_marginata
Fabaceae_Aspalathus_vulpina
Fabaceae_Aspalathus_taylorii
Fabaceae_Aspalathus_brevicarpa
Fabaceae_Aspalathus_callosa
Fabaceae_Aspalathus_cytisoides
Fabaceae_Aspalathus_capensis
Fabaceae_Aspalathus_tylodes
Fabaceae_Aspalathus_stokoei
Fabaceae_Aspalathus_carnosa
Fabaceae_Aspalathus_fusca
Fabaceae_Aspalathus_pinea
Fabaceae_Aspalathus_bracteata
Fabaceae_Aspalathus_crenata
Fabaceae_Aspalathus_barbata
Fabaceae_Aspalathus_cordata
Fabaceae_Aspalathus_angustifolia
Fabaceae_Aspalathus_commutata
Fabaceae_Aspalathus_perforata
Fabaceae_Aspalathus_perfoliata
Fabaceae_Aspalathus_alpestris
Fabaceae_Aspalathus_submissa
Fabaceae_Aspalathus_nigra
Fabaceae_Aspalathus_cephalotes
Fabaceae_Aspalathus_muraltioides
Fabaceae_Aspalathus_ericifolia
Fabaceae_Aspalathus_hispida
Fabaceae_Aspalathus_desertorum
Fabaceae_Aspalathus_uniflora
Fabaceae_Aspalathus_triquetra
Fabaceae_Aspalathus_confusa
Fabaceae_Aspalathus_galeata
Fabaceae_Aspalathus_willdenowiana
Fabaceae_Aspalathus_macrantha
Fabaceae_Aspalathus_asparagoides
Fabaceae_Aspalathus_opaca

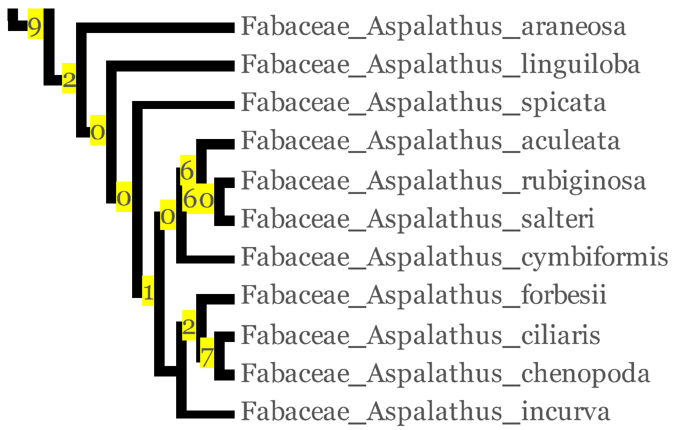

Fabaceae_Aspalathus_araneosa
Fabaceae_Aspalathus_linguiloba
Fabaceae_Aspalathus_spicata
Fabaceae_Aspalathus_aculeata
Fabaceae_Aspalathus_rubiginosa
Fabaceae_Aspalathus_salteri
Fabaceae_Aspalathus_cymbiformis
Fabaceae_Aspalathus_forbesii
Fabaceae_Aspalathus_ciliaris
Fabaceae_Aspalathus_chenopoda
Fabaceae_Aspalathus_incurva
