## Supplementary material for "Exploring the phylogeny of rosids with a five-locus supermatrix from GenBank": Tables S1, S2, S3

**Supporting Information Tables S1–S3**

### **Table S1.** Information of the 59 rosid calibration constraints used in this study. “CP#” is Calibration Point sequential number. “Tip1” and “Tip2” are tips in the both trees to define the node where the calibration point anchored; name formatting is Family_Genus_species. The age unit is million years ago (Myr).

| **CP#** | **Order** | **MinAge** | **MaxAge** | **Tip1** | **Tip2** | **Reference** |
| --- | --- | --- | --- | --- | --- | --- |
| 1 | Proteales | 125 | 125 | Platanaceae_Platanus_occidentalis | Nothofagaceae_Nothofagus_antarctica | Wang et al. (2009) |
| 2 | Proteales | 104 |  | Proteaceae_Lambertia_inermis | Platanaceae_Platanus_occidentalis | Crane et al. (1993); Doyle & Endress (2010);  Magallón et al. (2015) |
| 3 | Saxifragales | 89.3 |  | Altingiaceae_Liquidambar_orientalis | Altingiaceae_Altingia_obovata | Zhou et al. (2001); Magallón et al. (2015) |
| 4 | Saxifragales | 83.5 |  | Hamamelidaceae_Hamamelis_mollis | Hamamelidaceae_Corylopsis_stenopetala | Magallón-Puebla et al. (1996); Magallón et al. (2001); Magallón (2007); Magallón et al. (2015) |
| 5 | Saxifragales | 89.3 |  | Iteaceae_Pterostemon_rotundifolius | Iteaceae_Choristylis_rhamnoides | Hermsen et al. (2003); Magallón et al. (2015) |
| 6 | Saxifragales |  | 94.6 | Iteaceae_Pterostemon_rotundifolius | Grossulariaceae_Ribes_maximowiczianum | Hermsen et al. (2003); Bell et al. (2010) |
| 7 | Saxifragales | 70.6 |  | Haloragaceae_Haloragodendron_baeuerlenii | Penthoraceae_Penthorum_sedoides | Hernández-Castillo & Cevallos-Ferriz (1999);  Magallón et al. (2015) |
| 8 | Vitales | 56.8 |  | Vitaceae_Leea_guineensis | Vitaceae_Vitis_vinifera | Collinson et al. (1993); Bell et al. (2010) |
| 9 | Vitales | 66 |  | Vitaceae_Ampelopsis_grossedentata | Vitaceae_Cissus_repens | Manchester et al. (2013) |
| 10 | Myrtales | 87.5 | 95 | Combretaceae_Terminalia_sericea | Onagraceae_Clarkia_xantiana | Magallón et al. (2015) |
| 11 | Myrtales | 70.6 |  | Lythraceae_Decodon_verticillatus | Lythraceae_Lythrum_ovalifolium | Estrada-Ruíz et al. (2009) |
| 12 | Myrtales |  | 77.5 | Lythraceae_Nesaea_myrtifolia | Onagraceae_Circaea_alpina | Magallón et al., (2015) |
| 13 | Myrtales | 83.5 |  | Myrtaceae_Lophostemon_confertus | Crypteroniaceae_Crypteronia_paniculata | Eklund, (2003); Magallón et al. (2015) |
| 14 | Myrtales | 88.2 |  | Melastomataceae_Memecylon_bakerianum | Lythraceae_Lythrum_salicaria | Takahashi et al. (1999); Bell et al. (2010) |
| 15 | Myrtales | 55.8 |  | Myrtaceae_Heteropyxis_natalensis | Myrtaceae_Callistemon_lanceolatus | Crane et al. (1990); Pigg et al. (1993);  Magallón et al. (2015) |
| 16 | Crossosomatales | 28.4 |  | Staphyleaceae_Euscaphis_japonica | Staphyleaceae_Turpinia_paniculata | Tiffney (1979); Magallón et al. (2015) |
| 17 | Sapindales | 48.6 |  | Burseraceae_Bursera_schlechtendalii | Anacardiaceae_Choerospondias_axillaris | Reid & Chandler (1933); Manchester et al. (2009);  Magallón et al. (2015) |
| 18 | Sapindales | 55.8 |  | Sapindaceae_Acer_rubrum | Sapindaceae_Guindilia_trinervis | Knobloch & Mai (1986); Bell et al. (2010) |
| 19 | Sapindales | 65.5 |  | Rutaceae_Citrus_unshiu | Rutaceae_Spathelia_brittonii | Knoboch & Mai (1986) |
| 20 | Sapindales | 65 |  | Nitrariaceae_Nitraria_roborowskii | Sapindaceae_Guindilia_trinervis | Knobloch & Mai (1986); Bell et al. (2010) |
| 21 | Sapindales | 48.6 |  | Meliaceae_Chisocheton_cumingianus | Meliaceae_Khaya_nyasica | Reid & Chandler (1933); Magallón et al. (2015) |
| 22 | Huerteales | 37.2 |  | Tapisciaceae_Huertea_glandulosa | Tapisciaceae_Tapiscia_sinensis | Manchester (1988) |
| 23 | Malvales | 65.5 |  | Malvaceae_Bombax_ceiba | Thymelaeaceae_Thymelaea_hirsuta | Wheeler et al. (1987, 1994) |
| 24 | Malvales | 55.8 |  | Malvaceae_Microcos_argentata | Bixaceae_Bixa_urucurana | Carvalho et al. (2011); Magallón et al. (2015) |
| 25 | Brassicales | 89.3 |  | Neuradaceae_Neurada_procumbens | Brassicaceae_Brassica_nigra | Becker (1961); Beilstein et al. (2010);  Manchester & O’Leary (2010); Magallón et al. (2015)5 |
| 26 | Brassicales | 23.03 |  | Brassicaceae_Brassica_nigra | Brassicaceae_Arabidopsis_halleri | Becker (1961); Beilstein et al. (2010);  Manchester & O’Leary (2010); Magallón et al. (2015) |
| 27 | Zygophyllales | 23.03 |  | Krameriaceae_Krameria_ixine | Zygophyllaceae_Larrea_tridentata | Manchester & O’Leary (2010); Magallón et al. (2015) |
| 28 | Fabales | 55.8 |  | Polygalaceae_Xanthophyllum_hainanense | Polygalaceae_Polygala_tatarinowii | Pigg et al. (2008); Magallón et al. (2015) |
| 29 | Fabales | 55.8 |  | Surianaceae_Stylobasium_spathulatum | Fabaceae_Astragalus_membranaceus | Herendeen & Crane (1992); Magallón et al. (2015) |
| 30 | Fabales | 55.8 |  | Fabaceae_Cladrastis_delavayi | Fabaceae_Ammodendron_bifolium | Crepet & Taylor (1986); Magallón et al. (2015) |
| 31 | Fabales | 48.6 |  | Fabaceae_Erythrophleum_fordii | Fabaceae_Desmanthus_cooleyi | Crepet & Taylor (1986); Magallón et al. (2015) |
| 32 | Fabales | 59.9 |  | Fabaceae_Cercis_chinensis | Polygalaceae_Polygala_tenella | Herendeen & Crane (1992); Bell et al. (2010) |
| 33 | Rosales | 48.6 |  | Rosaceae_Spiraea_chinensis | Rosaceae_Photinia_prunifolia | Li et al. (2011) |
| 34 | Rosales | 89.8 |  | Rosaceae_Rosa_transmorrisonensis | Rosaceae_Sorbus_discolor | Crepet & Nixon (1996) |
| 35 | Rosales | 70.6 |  | Elaeagnaceae_Elaeagnus_umbellata | Rhamnaceae_Ceanothus_americanus | Calvillo-Canadell & Cevallos-Ferriz (2007) |
| 36 | Rosales | 55.8 |  | Ulmaceae_Zelkova_serrata | Urticaceae_Boehmeria_macrophylla | Manchester (1999) |
| 37 | Rosales | 65.5 |  | Cannabaceae_Aphananthe_monoica | Cannabaceae_Gironniera_subaequalis | Knobloch & Mai (1986); Magallón et al. (2015) |
| 38 | Rosales | 48.6 |  | Rhamnaceae_Ziziphus_angustifolia | Rhamnaceae_Alphitonia_incana | Manchester (1999); Magallón et al. (2015) |
| 39 | Rosales |  | 81.2 | Cannabaceae_Cannabis_sativa | Moraceae_Ficus_microcarpa | Magallón et al. (2015) |
| 40 | Fagales | 87.5 |  | Nothofagaceae_Nothofagus_antarctica | Cucurbitaceae_Coccinia_subsessiliflora | Chandler (1961); Collinson (1983);  Collinson et al. (1993) |
| 41 | Fagales | 64.4 |  | Juglandaceae_Pterocarya_hupehensis | Juglandaceae_Engelhardia_fenzelii | Manchester & Dilcher (1982) |
| 42 | Fagales | 64 |  | Juglandaceae_Carya_cathayensis | Juglandaceae_Annamocarya_sinensis | Manchester & Dilcher (1997) |
| 43 | Fagales | 59.8 | 65.5 | Betulaceae_Carpinus_caroliniana | Betulaceae_Betula_papyrifera | Pigg et al. (2003); Manchester et al. (2004) |
| 44 | Fagales | 83.5 | 85.5 | Fagaceae_Fagus_sylvatica | Betulaceae_Alnus_nitida | Sims et al. (1998) |
| 45 | Fagales | 37.2 |  | Fagaceae_Castanopsis_tibetana | Fagaceae_Castanea_seguinii | Nixon & Crepet (1989) |
| 46 | Fagales | 37.2 |  | Fagaceae_Lithocarpus_henryi | Fagaceae_Fagus_engleriana | Crepet & Daghlian (1980); Magallón et al. (2015) |
| 47 | Cucurbitales | 48.6 |  | Coriariaceae_Coriaria_nepalensis | Cucurbitaceae_Coccinia_subsessiliflora | Chandler (1961); Magallón et al. (2015) |
| 48 | Celastrales | 2.6 |  | Celastraceae_Tripterygium_wilfordii | Celastraceae_Celastrus_orbiculatus | Ozaki (1991); Magallón et al. (2015) |
| 49 | Celastrales | 37.2 |  | Celastraceae_Microtropis_fokienensis | Celastraceae_Euonymus_alatus | MacGinitie (1969); Wolfe (1977);  Magallón et al. (2015) |
| 50 | Oxalidales | 79.2 |  | Cephalotaceae_Cephalotus_follicularis | Cunoniaceae_Davidsonia_pruriens | Schönenberger et al. (2001) |
| 51 | Oxalidales | 61.7 |  | Elaeocarpaceae_Sloanea_latifolia | Elaeocarpaceae_Crinodendron_patagua | Manchester & Kvaček (2009); |
| 52 | Malpighiales | 33.9 | 37.2 | Humiriaceae_Humiria_balsamifera | Humiriaceae_Sacoglottis_gabonensis | Herrera et al. (2014) |
| 53 | Malpighiales | 89.3 |  | Clusiaceae_Symphonia_tanalensis | Hypericaceae_Cratoxylum_cochinchinense | Crepet & Nixon (1998); Ruhfel et al. (2013) |
| 54 | Malpighiales | 48 |  | Salicaceae_Idesia_polycarpa | Salicaceae_Populus_deltoides | Manchester et al. (2006) |
| 55 | Malpighiales | 37.2 |  | Euphorbiaceae_Pimelodendron_zoanthogyne | Euphorbiaceae_Acalypha_californica | Dilcher & Manchester (1988) |
| 56 | Malpighiales | 37.2 |  | Elatinaceae_Elatine_hexandra | Malpighiaceae_Malpighia_romeroana | Taylor & Crepet (1987) |
| 57 | Malpighiales | 33 |  | Malpighiaceae_Tetrapterys_glabrifolia | Malpighiaceae_Heteropterys_bahiensis | Hably & Manchester (2000) |
| 58 | Malpighiales | 37.2 |  | Salicaceae_Salix_lucida | Salicaceae_Populus_deltoides | Manchester et al. (2006) |
| 59 | Malpighiales | 33.9 |  | Rhizophoraceae_Rhizophora_apiculata | Rhizophoraceae_Bruguiera_gymnorhiza | Germeraad et al. (1968); Magallón et al. (2015) |

### **Table S2.** Family level summary sampling table for the 5-locus supermatrix (“Matrix”) compared to the rosid clade of the Open Tree Taxonomy (“OTT”) database v.3.0 (https://devtree.opentreeoflife.org/about/taxonomy-version/ott3.0; Hinchliff et al., 2015) and matching taxon names between these data sets. Families follow APG IV (2016); the circumscriptions of some families indicated as asterisk symbol in OTT were adjusted to comply with APG IV (2016), see "Note" column for details.

| **Order** | **Family*** | **Match(Matrix genera)**  **/OTT genera** | **Matched genus %** | **Match(Matrix species)**  **/OTT species** | **Matched species %** | **Note** |
| --- | --- | --- | --- | --- | --- | --- |
| Brassicales | Akaniaceae | 2(2)/2 | 100.00% | 2(2)/5 | 40.00% |  |
|  | Bataceae | 1(1)/1 | 100.00% | 1(1)/2 | 50.00% |  |
|  | Brassicaceae | 299(307)/407 | 73.46% | 1397(1502)/4862 | 28.73% |  |
|  | Capparaceae* | 17(17)/39 | 43.59% | 48(62)/459 | 10.46% | Excluding *Stixis* and *Tirania* (APG IV, 2016) |
|  | Caricaceae | 6(6)/7 | 85.71% | 33(34)/44 | 75.00% |  |
|  | Cleomaceae | 6(10)/14 | 42.86% | 66(90)/260 | 25.38% |  |
|  | Emblingiaceae | 1(1)/1 | 100.00% | 1(1)/1 | 100.00% |  |
|  | Gyrostemonaceae | 3(3)/5 | 60.00% | 5(5)/19 | 26.32% |  |
|  | Koeberliniaceae | 1(1)/1 | 100.00% | 1(1)/2 | 50.00% |  |
|  | Limnanthaceae | 2(2)/2 | 100.00% | 8(9)/12 | 66.67% |  |
|  | Moringaceae | 1(1)/2 | 50.00% | 7(7)/15 | 46.67% |  |
|  | Pentadiplandraceae | 1(1)/1 | 100.00% | 1(1)/1 | 100.00% |  |
|  | Resedaceae* | 10(10)/11 | 90.91% | 74(76)/132 | 56.06% | Including *Borthwickia*, (*Neothorelia*),  *Stixis* and *Tirania* (APG IV, 2016) |
|  | Salvadoraceae | 2(2)/3 | 66.67% | 4(4)/9 | 44.44% |  |
|  | Setchellanthaceae | 1(1)/1 | 100.00% | 1(1)/2 | 50.00% |  |
|  | Tovariaceae | 1(1)/1 | 100.00% | 1(1)/2 | 50.00% |  |
|  | Tropaeolaceae | 3(3)/4 | 75.00% | 43(46)/113 | 38.05% |  |
| Celastrales | Celastraceae | 66(67)/109 | 60.55% | 270(279)/1497 | 18.04% |  |
|  | Lepidobotryaceae | 2(2)/2 | 100.00% | 2(2)/2 | 100.00% |  |
| Crossosomatales | Aphloiaceae | 1(1)/1 | 100.00% | 1(1)/1 | 100.00% |  |
|  | Crossosomataceae | 4(4)/4 | 100.00% | 5(5)/10 | 50.00% |  |
|  | Geissolomataceae* | 1(1)/1 | 100.00% | 1(1)/1 | 100.00% | *Geissoloma marginata* is the accepted name in The Plant List,  while the name in OTL is *Geissoloma marginatum* |
|  | Guamatelaceae | 1(1)/1 | 100.00% | 1(1)/1 | 100.00% |  |
|  | Stachyuraceae | 1(1)/1 | 100.00% | 6(6)/15 | 40.00% |  |
|  | Staphyleaceae | 3(3)/4 | 75.00% | 8(8)/52 | 15.38% |  |
|  | Strasburgeriaceae | 2(2)/2 | 100.00% | 2(2)/2 | 100.00% |  |
| Cucurbitales | Anisophylleaceae | 4(4)/4 | 100.00% | 15(15)/60 | 25.00% |  |
|  | Apodanthaceae | 2(2)/3 | 66.67% | 5(5)/27 | 18.52% |  |
|  | Begoniaceae | 2(2)/4 | 50.00% | 291(296)/1755 | 16.58% |  |
|  | Coriariaceae | 1(1)/1 | 100.00% | 10(11)/19 | 52.63% |  |
|  | Corynocarpaceae | 1(1)/1 | 100.00% | 2(4)/5 | 40.00% |  |
|  | Cucurbitaceae | 104(112)/117 | 88.89% | 496(528)/1223 | 40.56% |  |
|  | Datiscaceae | 1(1)/1 | 100.00% | 2(2)/2 | 100.00% |  |
|  | Tetramelaceae | 2(2)/2 | 100.00% | 2(2)/3 | 66.67% |  |
| Fabales | Fabaceae | 641(652)/827 | 77.51% | 5260(5626)/22780 | 23.09% |  |
|  | Polygalaceae | 10(10)/31 | 32.26% | 44(45)/1400 | 3.14% |  |
|  | Quillajaceae | 1(1)/1 | 100.00% | 1(1)/2 | 50.00% |  |
|  | Surianaceae | 5(5)/5 | 100.00% | 6(6)/9 | 66.67% |  |
| Fagales | Betulaceae | 6(6)/9 | 66.67% | 105(110)/359 | 29.25% |  |
|  | Casuarinaceae | 4(4)/4 | 100.00% | 75(77)/104 | 72.12% |  |
|  | Fagaceae | 9(10)/29 | 31.03% | 221(235)/1478 | 14.95% |  |
|  | Juglandaceae | 10(11)/21 | 47.62% | 55(59)/137 | 40.15% |  |
|  | Myricaceae | 4(4)/5 | 80.00% | 30(34)/66 | 45.45% |  |
|  | Nothofagaceae | 2(2)/5 | 40.00% | 9(24)/118 | 7.63% |  |
|  | Ticodendraceae | 1(1)/1 | 100.00% | 1(1)/1 | 100.00% |  |
| Geraniales | Francoaceae* | 9(11)/9 | 100.00% | 26(28)/49 | 53.06% | New family name propoased in APG IV (2016) including  Melianthaceae, Vivianiaceae |
|  | Geraniaceae | 6(7)/11 | 54.55% | 269(277)/913 | 29.46% |  |
| Huerteales | Dipentodontaceae | 2(2)/2 | 100.00% | 3(3)/20 | 15.00% |  |
|  | Gerrardinaceae | 1(1)/1 | 100.00% | 1(1)/2 | 50.00% |  |
|  | Petenaeaceae | 1(1)/1 | 100.00% | 1(1)/1 | 100.00% |  |
|  | Tapisciaceae | 2(2)/2 | 100.00% | 2(2)/7 | 28.57% |  |
| Malpighiales | Achariaceae | 18(18)/31 | 58.06% | 20(22)/168 | 11.90% |  |
|  | Balanopaceae | 1(1)/1 | 100.00% | 4(4)/9 | 44.44% |  |
|  | Bonnetiaceae | 3(3)/3 | 100.00% | 10(10)/38 | 26.32% |  |
|  | Calophyllaceae | 12(12)/17 | 70.59% | 48(51)/432 | 11.11% |  |
|  | Caryocaraceae | 2(2)/2 | 100.00% | 3(3)/29 | 10.34% |  |
|  | Centroplacaceae | 2(2)/2 | 100.00% | 1(2)/4 | 25.00% |  |
|  | Chrysobalanaceae | 16(16)/21 | 76.19% | 39(40)/573 | 6.81% |  |
|  | Clusiaceae | 15(15)/28 | 53.57% | 77(80)/898 | 8.57% |  |
|  | Ctenolophonaceae | 1(1)/1 | 100.00% | 1(1)/2 | 50.00% |  |
|  | Dichapetalaceae | 2(2)/3 | 66.67% | 8(8)/214 | 3.74% |  |
|  | Elatinaceae | 2(2)/5 | 40.00% | 7(7)/58 | 12.07% |  |
|  | Erythroxylaceae | 3(3)/5 | 60.00% | 10(11)/286 | 3.50% |  |
|  | Euphorbiaceae | 176(177)/282 | 62.41% | 1645(1693)/7493 | 21.95% |  |
|  | Euphroniaceae | 1(1)/1 | 100.00% | 1(1)/3 | 33.33% |  |
|  | Goupiaceae | 1(1)/1 | 100.00% | 1(1)/1 | 100.00% |  |
|  | Humiriaceae | 4(4)/8 | 50.00% | 4(5)/75 | 5.33% |  |
|  | Hypericaceae | 7(7)/10 | 70.00% | 256(263)/685 | 37.37% |  |
|  | Irvingiaceae | 3(3)/3 | 100.00% | 3(3)/11 | 27.27% |  |
|  | Ixonanthaceae | 3(3)/5 | 60.00% | 4(4)/20 | 20.00% |  |
|  | Lacistemataceae | 2(2)/2 | 100.00% | 3(3)/15 | 20.00% |  |
|  | Linaceae | 16(16)/17 | 94.12% | 83(85)/304 | 27.30% |  |
|  | Lophopyxidaceae | 1(1)/1 | 100.00% | 1(1)/1 | 100.00% |  |
|  | Malpighiaceae | 70(71)/84 | 83.33% | 295(307)/1614 | 18.28% |  |
|  | Ochnaceae | 17(18)/40 | 42.50% | 28(33)/686 | 4.08% |  |
|  | Pandaceae | 3(3)/5 | 60.00% | 6(6)/24 | 25.00% |  |
|  | Passifloraceae | 27(28)/34 | 79.41% | 285(293)/1090 | 26.15% |  |
|  | Peraceae | 4(5)/4 | 100.00% | 12(13)/116 | 10.34% |  |
|  | Phyllanthaceae | 50(54)/61 | 81.97% | 269(302)/2391 | 11.25% |  |
|  | Picrodendraceae | 13(13)/27 | 48.15% | 14(14)/122 | 11.48% |  |
|  | Podostemaceae | 45(46)/56 | 80.36% | 135(148)/368 | 36.68% |  |
|  | Putranjivaceae | 3(3)/3 | 100.00% | 22(23)/223 | 9.87% |  |
|  | Rafflesiaceae | 3(3)/3 | 100.00% | 21(23)/39 | 53.85% |  |
|  | Rhizophoraceae | 14(14)/19 | 73.68% | 38(39)/164 | 23.17% |  |
|  | Salicaceae | 23(23)/66 | 34.85% | 192(199)/1723 | 11.14% |  |
|  | Trigoniaceae | 2(2)/5 | 40.00% | 5(5)/36 | 13.89% |  |
|  | Violaceae | 21(21)/35 | 60.00% | 152(165)/1401 | 10.85% |  |
| Malvales | Bixaceae* | 4(4)/8 | 50.00% | 6(6)/31 | 19.35% | Including Cochlospermaceae (*Maximilianea*) |
|  | Cistaceae | 8(8)/9 | 88.89% | 60(62)/356 | 16.85% |  |
|  | Cytinaceae | 2(2)/2 | 100.00% | 2(2)/11 | 18.18% |  |
|  | Dipterocarpaceae | 14(14)/24 | 58.33% | 135(135)/575 | 23.48% |  |
|  | Malvaceae | 185(186)/291 | 63.57% | 802(850)/5740 | 13.97% |  |
|  | Muntingiaceae | 2(2)/2 | 100.00% | 2(2)/2 | 100.00% |  |
|  | Neuradaceae | 2(2)/3 | 66.67% | 2(2)/7 | 28.57% |  |
|  | Sarcolaenaceae | 2(2)/10 | 20.00% | 3(3)/73 | 4.11% |  |
|  | Sphaerosepalaceae | 2(2)/3 | 66.67% | 3(3)/21 | 14.29% |  |
|  | Thymelaeaceae | 34(35)/53 | 64.15% | 270(284)/955 | 28.27% |  |
| Myrtales | Alzateaceae | 1(1)/1 | 100.00% | 1(1)/1 | 100.00% |  |
|  | Combretaceae | 16(16)/23 | 69.57% | 99(100)/602 | 16.45% |  |
|  | Crypteroniaceae | 3(3)/3 | 100.00% | 7(7)/13 | 53.85% |  |
|  | Lythraceae | 27(28)/35 | 77.14% | 123(132)/724 | 16.99% |  |
|  | Melastomataceae | 79(80)/192 | 41.15% | 278(297)/6010 | 4.63% |  |
|  | Myrtaceae | 93(95)/167 | 55.69% | 524(548)/6909 | 7.58% |  |
|  | Onagraceae | 22(24)/37 | 59.46% | 214(247)/990 | 21.62% |  |
|  | Penaeaceae | 9(9)/9 | 100.00% | 26(26)/32 | 81.25% |  |
|  | Vochysiaceae | 7(7)/8 | 87.50% | 14(15)/248 | 5.65% |  |
| Oxalidales | Brunelliaceae | 1(1)/1 | 100.00% | 3(3)/58 | 5.17% |  |
|  | Cephalotaceae | 1(1)/1 | 100.00% | 1(1)/1 | 100.00% |  |
|  | Connaraceae | 4(4)/15 | 26.67% | 7(7)/251 | 2.79% |  |
|  | Cunoniaceae | 23(25)/30 | 76.67% | 56(64)/355 | 15.77% |  |
|  | Elaeocarpaceae* | 8(8)/14 | 57.14% | 41(42)/806 | 5.09% |  |
|  | Huaceae | 2(2)/2 | 100.00% | 1(2)/5 | 20.00% |  |
|  | Oxalidaceae | 4(4)/6 | 66.67% | 73(74)/731 | 9.99% |  |
| Picramniales | Picramniaceae | 2(2)/3 | 66.67% | 5(5)/57 | 8.77% |  |
| Rosales | Barbeyaceae | 1(1)/1 | 100.00% | 1(1)/1 | 100.00% |  |
|  | Cannabaceae* | 10(10)/10 | 100.00% | 31(31)/126 | 24.60% | Including *Chaetachme* |
|  | Dirachmaceae | 1(1)/1 | 100.00% | 1(1)/3 | 33.33% |  |
|  | Elaeagnaceae | 3(3)/3 | 100.00% | 14(15)/115 | 12.17% |  |
|  | Moraceae | 24(24)/46 | 52.17% | 197(199)/1554 | 12.68% |  |
|  | Rhamnaceae | 46(46)/65 | 70.77% | 162(172)/1332 | 12.16% |  |
|  | Rosaceae | 97(107)/161 | 60.25% | 1179(1253)/15235 | 7.74% |  |
|  | Ulmaceae* | 6(6)/11 | 54.55% | 22(22)/103 | 21.36% | Excluding *Chaetachme* |
|  | Urticaceae | 29(29)/61 | 47.54% | 87(93)/2151 | 4.04% |  |
| Sapindales | Anacardiaceae | 50(50)/92 | 54.35% | 121(128)/1034 | 11.70% |  |
|  | Biebersteiniaceae | 1(1)/1 | 100.00% | 4(4)/4 | 100.00% |  |
|  | Burseraceae | 15(15)/24 | 62.50% | 225(227)/703 | 32.01% |  |
|  | Kirkiaceae* | 1(1)/1 | 100.00% | 2(2)/6 | 33.33% | This family is not presented in OTL V.9.0 |
|  | Meliaceae | 47(47)/59 | 79.66% | 126(133)/835 | 15.09% |  |
|  | Nitrariaceae | 4(4)/4 | 100.00% | 9(9)/17 | 52.94% |  |
|  | Rutaceae* | 97(100)/175 | 55.43% | 391(410)/2561 | 15.27% | This family is not presented in OTL V.9.0 |
|  | Sapindaceae | 107(109)/171 | 62.57% | 422(460)/2127 | 19.84% |  |
|  | Simaroubaceae | 20(20)/24 | 83.33% | 61(61)/134 | 45.52% |  |
| Vitales | Vitaceae | 9(9)/15 | 60.00% | 110(119)/1155 | 9.52% |  |
| Zygophyllales | Krameriaceae | 1(1)/1 | 100.00% | 12(12)/20 | 60.00% |  |
|  | Zygophyllaceae | 17(17)/26 | 65.38% | 48(59)/320 | 15.00% |  |

### **Table S3.** List of non-monophyletic families in the *matR* and ITS locus trees.

**#1. 12 Non-monophyletic families in the *matR* gene tree**

Achariaceae

Betulaceae

Cannabaceae

Capparaceae

Elaeocarpaceae

Euphorbiaceae

Fagaceae

Hypericaceae

Juglandaceae

Meliaceae

Rutaceae

Simaroubaceae

**#2. 18 Non-monophyletic families in the ITS locus tree**

Combretaceae

Euphorbiaceae

Fabaceae

Francoaceae

Hypericaceae

Malvaceae

Meliaceae

Nitrariaceae

Phyllanthaceae

Podostemaceae

Rafflesiaceae

Resedaceae

Rhizophoraceae

Rutaceae

Sapindaceae

Simaroubaceae

Ulmaceae

Zygophyllaceae
